# Hypothesis: A modern human range expansion ∼300,000 years ago had a critical role in Neandertal origins

**DOI:** 10.64898/2026.03.11.711219

**Authors:** David Reich

## Abstract

This paper proposes that a critical event in Neandertal evolution was a range expansion of early modern humans into Europe 300 thousand years ago (ka). As bands of early modern humans spread, they incorporated resident archaic people leading ultimately to so much admixture that the resulting population carried ∼3-7% modern ancestry. I identify multiple ways in which this would lead to a high probability of these direct ancestors of Neandertals retaining early modern human lineages at both mitochondrial DNA and Y chromosomes while clustering with Denisovans genome-wide. A previous explanation for these patterns without the unique population dynamics associated with a range expansion has less than a 3% chance of matching the data. In Africa, the hypothesis also has explanatory power: if the expanding population also mixed African archaic humans, it could account for the evidence that all humans living today are admixed between lineages that separated more than a million years ago and remixed ∼300 ka. This hypothesis of a bicontinental range expansion correlates with shared archaeological traits between West Eurasia and Africa that were far less present in East Asia, including Levallois point stone tools, improved hafting, use of fire and ochre, and larger and more organized campsites. Motivated by the evidence that an early modern human expansion had a role in Neandertal and modern human evolution, I identify 66 loci spanning <0.1% of the genome where people today share ancestry with Neandertals more recently than separation from Denisovans. This includes a haplotype at *FADS1/FADS2* that codes for an enzyme modifying polyunsaturated plant-derived fatty acids and may reflect adaptation to a more diversified diet, and an amino acid substitution at *ARHGAP32* modulating neural connections that may be important in modern human cognitive evolution.

## A genetic puzzle

Neandertals and Denisovans were extinct populations of humans who lived across Eurasia until ∼40 ka. Analysis of high-coverage genome sequences from them has led to the inference that they separated from a common archaic ancestral population 585-504 thousand years ago (ka), which in turn separated from a common ancestral population with modern humans 825-694 ka (Figure 1; ranges of dates from ref. 1). This reconstruction is consistent with the estimated genetic divergence of Denisovans and modern humans at two loci with enough informative sites to allow accurate estimates: at mitochondrial DNA (mtDNA) 967-800 ka (95% confidence intervals from ref. 1 here and in what follows), and on the Y chromosome 876-697 ka. But the estimated genetic splits of Neandertals and modern humans living today are hundreds of thousands of years younger than the inferred population separation—for mtDNA 495-387 ka^1,2^, and for the Y chromosome 433-346 ka^1,3^—which can only be explained by gene flow between the lineages after the inferred population split time. Multiple studies of the rest of the genome (chromosomes 1-22 and X) have found confirmatory evidence for this gene flow, inferring that Neandertals harbor 3-7% admixture^4,5,6^ from an early modern human lineage ∼300-200 ka^5,6,7^, clearly the vector for the shared mtDNA and Y lineages (genetic divergence times must be at least as old as the time since populations share ancestors, but can be hundreds of thousands of years older). A final line of evidence consistent with a ∼300-200 ka gene flow from early modern humans into Neandertals is ancient DNA from an individual from Sima de los Huesos in Spain who lived 430-300 ka^8^, and who carried archaic lineage mtDNA related to Denisovans^9^ but a nuclear genome that shared drift with Neandertals^10^, as expected for an archaic European who lived prior to admixture with modern humans (Figure 1).

**Figure 1:**
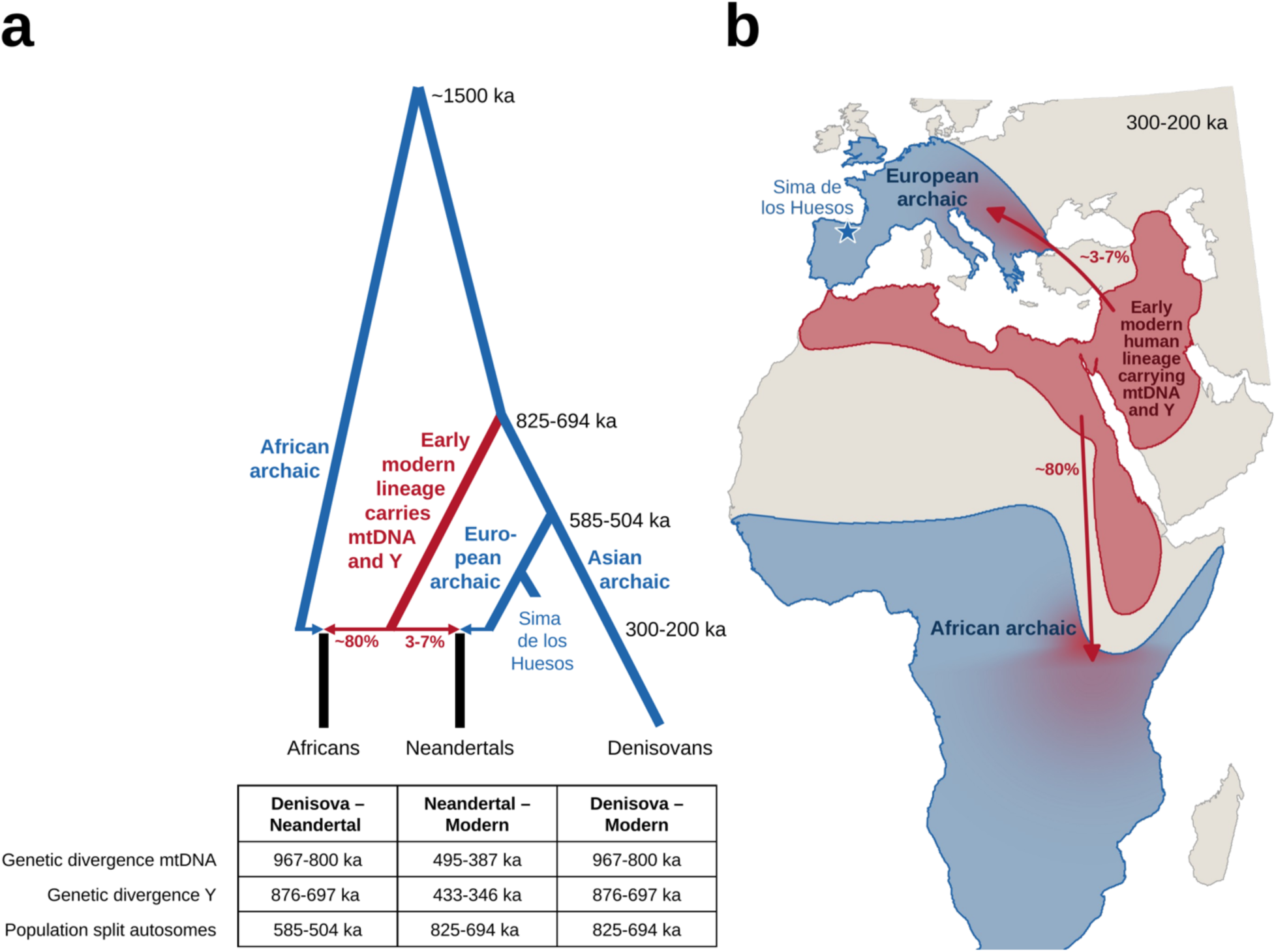
A bicontinental range expansion ∼300 ka. (a) I start with a previously proposed model for the relationship of Neandertals, Denisovans, and modern Africans in which an early modern lineage contributed mtDNA and Y chromosomes to Neandertals and ∼3-7% of the rest of the genome^3^. I graft onto it the deep substructure detected in people today, inferred in one study to contribute ∼80% of the ancestry of present-day Africans^11^, and inferred in several studies to have ended around ∼300 ka^7,11,12^. At bottom are estimates of divergences and population splits: range of estimated dates for the autosomes, and 95% confidence intervals for mtDNA and Y^1^. (Not shown: <50 ka gene flow from Neandertals and Denisovans into non-Africans, and superarchaic gene flow to Denisovans.) (b) Schematic of what the range expansion might have looked like after ∼300 ka, with an early modern human lineage in northeast Africa or the Near East expanding both into Europe and Africa and mixing with local archaic populations; ancestry within these regions would then have homogenized due to admixture among neighbors; the red-to-blue shading indicates mixing. The European archaic range is drawn based on an estimate of the later Neandertal range^13^. Coastlines are from the GLOBE 1-km digital elevation model^14^ via the global-land-mask package^15^, in an Albers equal-area projection.

## Models without range expansion have a less than 3% chance of explaining these patterns

The conundrum posed by the fact that Neandertals harbor modern human-associated mtDNA and Y chromosomes but are more closely related to Denisovans on chromosomes 1-22 and X was recognized in the paper that reported the Denisovan Y chromosome sequences^3^. To explain it, the authors hypothesized that following admixture ∼300-200 ka—which they modeled as around 5% early modern human admixture into an archaic human population in a non-geographically-explicit model—there was selection for the early modern human lineage on the entirely male line (tracked by the Y chromosome), or the female line (tracked by mtDNA), or both^3^. Using simulations of a non-recombining, uniparentally transmitted locus and a homogeneous population, they demonstrated that a selection coefficient of 1-2% for the genetic material from the early modern human lineage over the archaic one for either mtDNA or the Y chromosome would be sufficient to lead to probable fixation^3^. They proposed that such a fitness gap could have arisen through a greater accumulation of slightly deleterious mutations expected in populations of relatively smaller size (the archaic ancestors of Neandertals) than in ones of relatively larger size (the early modern ancestors of Neandertals)^16^. However, they did not test if the development of a fitness gap of this magnitude was consistent with biological constraints on mtDNA and the Y chromosome, or the probability of both the mtDNA and Y chromosome patterns being observed together.

To determine whether a fitness gap this large for mtDNA was likely given the biological constraints, I started with a previous framework for estimating the distribution of fitness effects of new mutations^17^, expanding it to include not just the approximately two-thirds of mtDNA in protein coding sequence but also the non-coding parts of mtDNA which are largely RNA genes (Supplementary Information section 1). I took into account all the sources of uncertainty I could identify, including in mutation rate, the fitness effects of mutations, and demography. I constrained the mutation rate using the synonymous substitution counts on the modern and Denisovan mtDNA lineages, the fitness effects based on patterns of polymorphism within humans and divergence from archaic humans, and the fitness gap by the observed ratio of nonsynonymous substitutions on the two lineages (which is slightly higher on the modern than on the Denisovan side, opposite to what an excess archaic genetic load would predict). I inferred a posterior distribution on the mtDNA fitness gap with a median of 0.32% (95% of the density between [-0.2%, +1.3%]), leading to a probability of 23% of fixation of modern human mtDNA when I carried out simulations integrated over the uncertainty in the fitness gap (Figure 2a).

**Figure 2:**
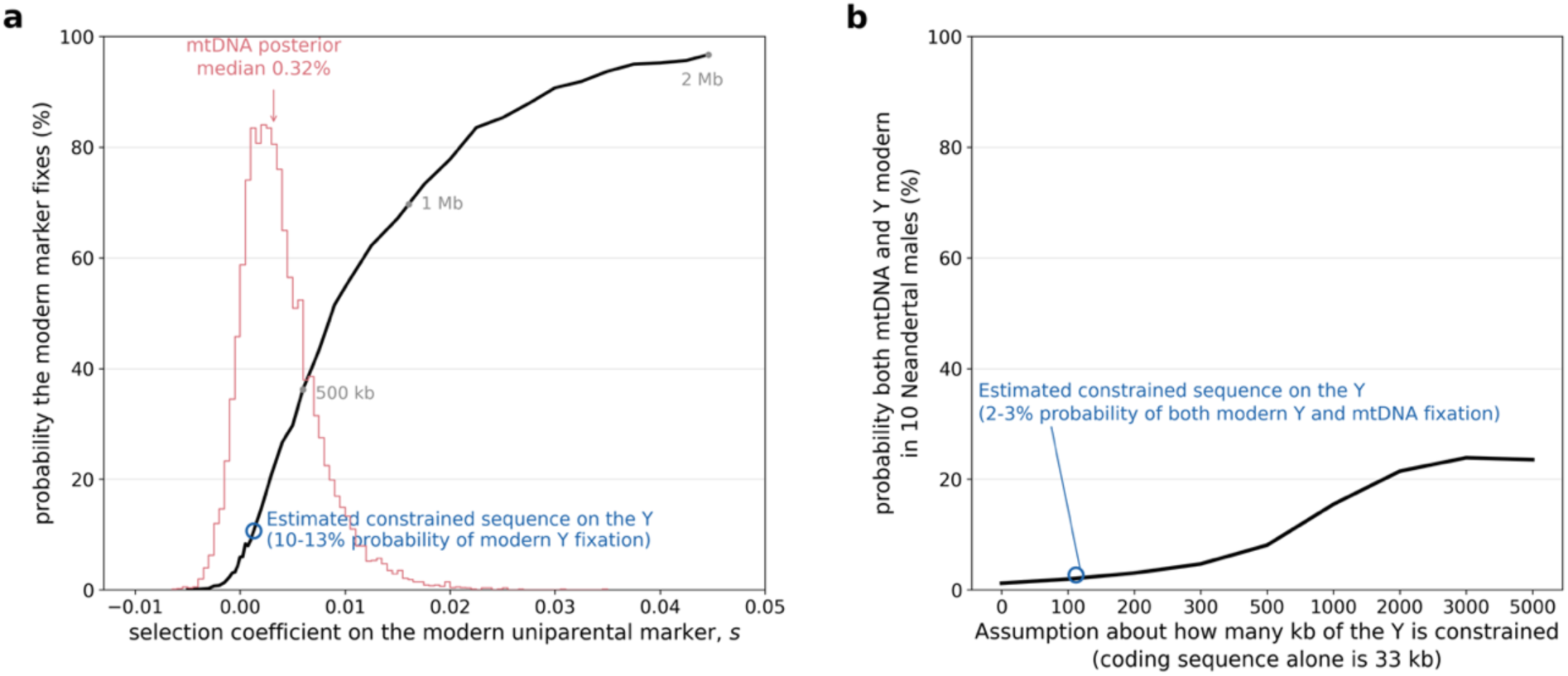
Archaic-modern fitness gap is highly unlikely to have been sufficient to retain modern DNA at both uniparentally inherited markers. (a) To estimate the fitness gap between the archaic and the early modern ancestors of Neandertals, I conditioned on the observed patterns of polymorphism and divergence in mtDNA and estimates of the demographic parameters. I infer a 23% probability of retaining modern human lineage mtDNA integrating over the uncertainty (red). For the Y chromosome, I show the point estimate of the fitness gap. (b) The low probability of the data is evident from considering both loci together. I show the probability that 10 sampled Neandertals 50 ka all harbor early modern human mtDNA and Y chromosomes, similar to real data. This is 2-3% at the estimated 112 kb of constrained sequence in the non-palindromic Y.

To determine whether a fitness gap this large for the Y chromosome was likely, I could not use the same direct approach as for mtDNA, because the fraction of the Y chromosome that is coding is on the order of 0.1%, providing too little data to constrain the inference (a previous estimate of Y chromosome load was not usable for my purposes, as its authors attribute it largely to palindromic repeats, which gene conversion between near-identical copies protects from irreversible accumulation of deleterious mutations^18^). As an alternative, I estimated the amount of constrained sequence in the single-copy part of the Y chromosome by analyzing conservation scores in an alignment of mammalian genomes^19^. I identified ∼112 kb of conserved sequences, and by assuming that the distribution of selection coefficients for newly arising mutations is the same as for coding bases elsewhere in the genome (or weaker, translating to a faster rate of accumulation of slightly deleterious mutations), I inferred a median 0.09-0.17% fitness gap, leading to 10-13% probability of fixation of modern human Y chromosomes (Supplementary Information section 1).

Combining mtDNA and the Y chromosome, my simulations infer a 2-3% probability of modern human lineage fixation at both loci (Figure 2b). This is in fact an overestimate, because hybridization is expected to pull down the frequencies of functionally important alleles from the minority admixing population due to “Dobzhansky-Muller Incompatibilities” (DMIs): co-evolved variants that only work when they are on the same genetic background causing compromised reproduction in hybrids that do not carry both^20^. DMIs appear to have affected many loci in the ∼300-200 ka mixture^4,6^, and since they often involve mtDNA^21^, it is reasonable to expect them to have pushed down the probability of modern mtDNA fixation. I was not able to identify substitutions on both sides of an interface between mtDNA and nuclear interacting proteins that are often involved in a mtDNA DMI^22^ (Supplementary Information section 2), but since mito-nuclear DMIs are not always mediated by such interactions^20^, 2-3% is conservative.

## Geographically explicit scenarios predict the data with high probability

Studies of diverse species have shown that when a population moves into territory occupied by a previously established group, large-scale incorporation of local genes occurs if there is even a small rate of interbreeding at the wavefront of the expansion. By the time the expansion finishes, the genetic composition can be largely local^23^, consistent with the observation that Neandertals carried only a small proportion (∼3-7%) of early modern human lineage ancestry genome-wide. A previous study considered how range expansion might have affected patterns of introgression in admixtures between modern humans and Neandertals, focusing not on the ∼300-200 ka event, but instead on the ∼50 ka admixture in which modern humans bearing Upper Paleolithic technology expanded into Europe^26^. It inferred a <0.1% per generation rate of gene flow from Neandertals into expanding modern humans at locations where they overlapped, based on the absence of Neandertal ancestry in later European mtDNAs. The larger introgression proportion in the earlier ∼300-200 ka event would imply a higher interbreeding rate between groups or a slower expansion, likely reflecting more modest biological or cultural gaps between the interacting groups.

The geographic explicitness of a range expansion results in population genetic dynamics that do not occur in non-geographically-explicit models^23^. I hypothesized that this could help explain retention of modern ancestry not only in the autosomal genome, but also at the uniparentally inherited markers. In human foragers and great apes, dispersal events to join neighboring groups are nearly always mediated by one sex^24,25^. In a range expansion, this would be expected to lead to the marker associated with the non-dispersing sex being retained by the expanding population, shielded from the archaic ancestry flooding the rest of the genome. In contrast, sex-bias does not help to explain the data for the previously proposed non-geographically-explicit model of a ∼5% pulse of modern human flow into Neandertal ancestors^3^, since the lineage associated with the sex that mates across populations would still have a small probability of fixing due to drift (Extended Data Figure 1.) Retention of modern human ancestry at the other uniparentally inherited marker is also more plausible in a range expansion, since at the beginning modern human lineages at both mtDNA and Y chromosomes are in the majority, so what is necessary to explain the data is to block archaic lineages from coming in; I show four plausible scenarios that do this in what follows.

To identify if there were plausible scenarios whereby both uniparentally transmitted markers could be retained with high probability during a range expansion, I developed two computer simulation frameworks capturing different features of the geography, timing, and population sizes in Neandertal ancestors before and after the ∼300-200 ka mixture. In the “deme model” (inspired by previous modeling of range expansions^23,26^), individuals live in a grid of usually 12x12 squares each of which can include individuals of both groups (archaic, or early modern human). Modern individuals have a reproductive advantage over archaic ones that live in the same deme, and occasionally interbreed across groups within a deme with the group identity of the offspring determined by either mothers or fathers. In the “roving band model”, there are on the order of 70 bands of typical size 25 scattered across a landscape that I match to the geography of the area of Europe where Neandertals lived^13^. Bands are 100% archaic or 100% early modern human, diffuse across the landscape each generation, and modern human bands have reproductive advantages over archaic ones when near enough to compete for the same resources. Individuals occasionally switch groups (males join their partner’s band in some sets of simulations, females join their partner’s band in other sets of simulations). Supplementary Information section 3 gives full details.

The results replicate the qualitative behavior of previously reported simulations of range expansions^23,26^. The reproductive advantage of early modern humans at the wavefront of expansion amplifies the archaic DNA they take on through admixture, so even after the first hundred generations more than half of the expanding population’s ancestry is archaic. By the time the wavefront sweeps across the landscape, there is typically only a small fraction of early modern human lineage genes left (Figure 3a, animated in the Supplementary Video, Extended Data Figure 2). These patterns are robust to a wide range of assumptions (Extended Data Figure 3).

**Figure 3:**
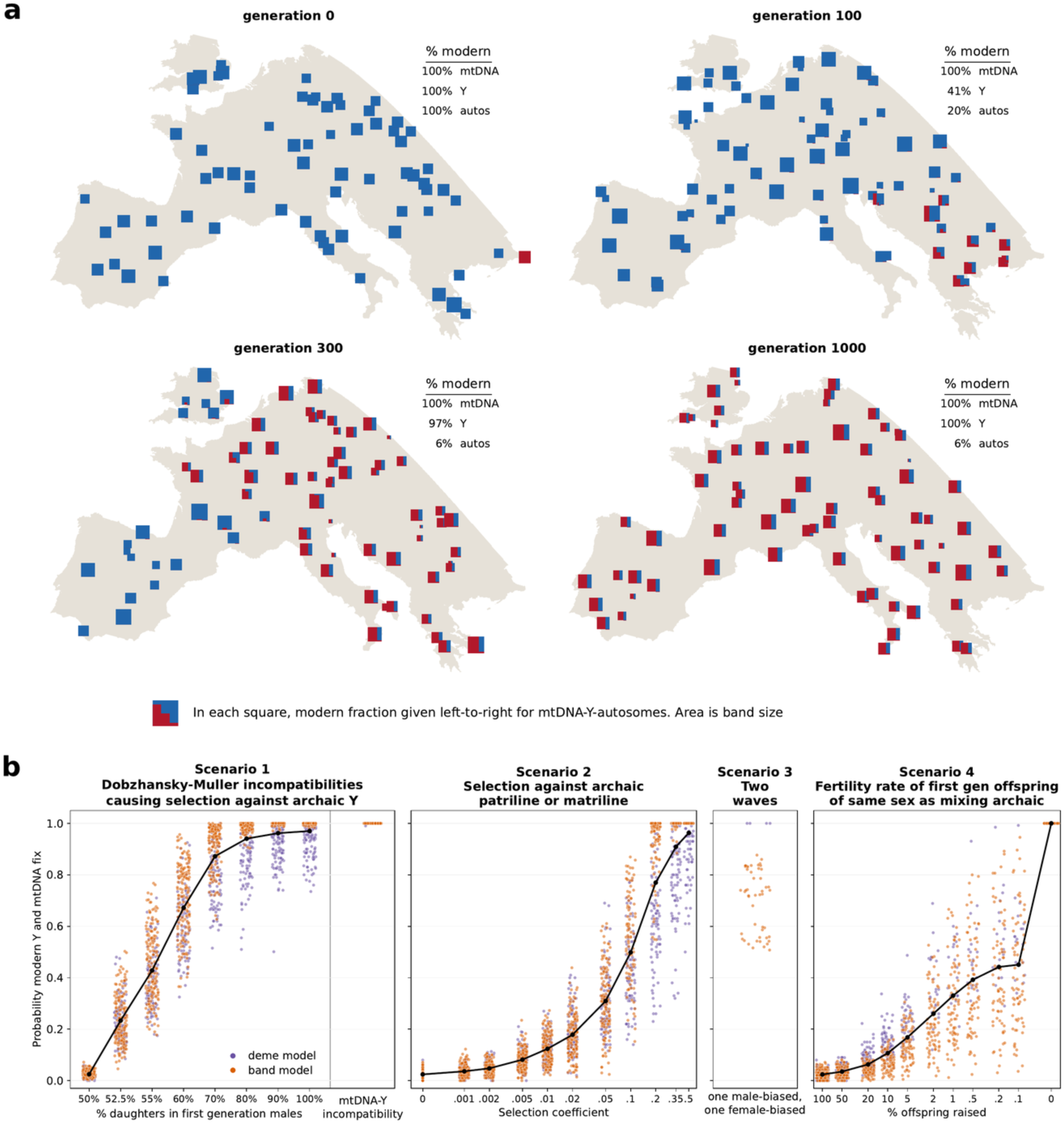
Range expansion scenarios that retain early modern mtDNA and Y with high probability. (a) Example of an early modern human expansion into Europe in a “roving band” model with baseline demographic parameters and the Y retained due to a 16% higher reproductive rate for prestigious male patrilines. The landscape is seeded by a single early modern human band of size 25 (the remaining 69 are archaic), and bands diffuse every generation and exchange male migrants mixing modern (red) and archaic (blue) lineages. Each square has three vertical bars showing the modern proportion for mtDNA (left), Y (middle), and autosomes (right). Next to each plot is a count of the percent modern genetic ancestry in bands with a modern affiliation at that generation; because only males switch bands, a band’s mtDNA bar is either fully modern or fully archaic. Modern ancestry at the Y and autosomes decreases because of archaic gene flow (but recovers at the Y due to advantage of prestigious male patrilines). (b) Simulations of a wide range of parameter combinations for both the deme model (purple) and roving band model (orange) show four scenarios through which modern ancestry can be retained at both mtDNA and Y with high probability.

For each mechanism I hypothesized could retain the other uniparentally inherited marker, I tested a range of parameters (744 for the deme model, 567 for the band model) (Extended Data Figure 4). For parameter combinations matching the chromosome 1-22 ancestry (3-7% early modern human), and a plausible time for range expansion completion (100-1000 generations), I tabulated the rate of fixation of early modern human lineage mtDNA and Y chromosomes in 10 simulated Neandertal males (Supplementary Table 1). Four scenarios match the data with high probability (Figure 3b, Supplementary Information section 3).

The first scenario I found only works when males are the sex that switches groups. “Haldane’s Rule”^27^ states that when two populations have been isolated for long enough that first generation hybrids have fragile reproductive phenotypes (due to DMIs: pairs of co-evolved alleles that do not work if they aren’t inherited together), it is almost always the heterogametic sex that is affected (males in mammals), blocking introgression of archaic Y chromosomes. If the fragile male reproductive phenotype is reduced fertility, this is not an effective mechanism for retaining early modern human Y (Extended Data Figure 4); This is because selection in hybrids against both partner alleles in the DMI rapidly ends the block to Y introgression. However, the other frequent manifestation of DMIs—hybrids of the heterogametic sex (males in mammals) having a skew toward mostly having children of the homogametic sex (daughters in mammals), a pattern seen in mice and fruit flies^27,28,29,30,31^—effectively prevents archaic Y introgression (Figure 3b, Extended Data Figure 4). Daughter-skew arises due to selfish mutations on the X that cause men to overproduce sperm carrying X chromosomes, which are normally masked by suppressor alleles on the Y. But in hybrids, archaic Y lineages not carrying the suppressor are outcompeted by early modern ones carrying it. I show that a daughter skew as little as 60% is enough to make it likely to retain the modern Y chromosome (Figure 3b, Supplementary Information section 3).

The second scenario that provides a high probability match to the mtDNA and Y chromosome patterns works for both directions of sex-bias. Human cultures often track clan membership generations into the past^32^, and in such a context the prestige associated with modern patrilines or matrilines can translate to increased fertility. In an expanding early human population impacted by hybridization by local archaic humans, it would not be surprising if individuals who traced their ancestry to early modern humans on the entirely male line (prestigious patrilines) or female line (prestigious matrilines) would have been more successful at competing for mates and therefore have a higher reproductive rate. My simulations show that if the fertility of modern patrilines or matrilines within modern groups was higher by roughly 9-14%, it would be sufficient to drive the observations (Figure 3b, Supplementary Information section 3).

The third scenario that can explain the data is two successful range expansions. In one, males are the sex that interbreeds across populations leading to retention of early modern human mtDNA. In the other that precedes or succeeds it, females interbreed across populations leading to retention of early modern human Y chromosomes (Figure 3b). Cultural adaptation in humans can be rapid, and two waves with different social structures, on the order of 10,000 years apart, seem plausible.

A fourth scenario that could explain the data is if first-generation offspring of the same sex as their archaic parent were not raised. There are ethnographic examples in forager societies today of offspring of different sexes being incorporated into the two parental populations differently depending on their sex^33,34^, but I view this as the least anthropologically plausible of the four scenarios, as my simulations in Figure 3b show that the differential treatment of males and females would have to be more than a hundred-fold for this process to explain the data.

## Genetic substructure in people today suggests that the range expansion also impacted Africa

I now highlight an independent genetic inference that has not previously been co-considered with the Neandertal evidence, and also points to a major range expansion ∼300 ka. Multiple studies have inferred that the ancestors of all living humans including sub-Saharan Africans were deeply substructured into lineages that diverged more than a million years ago^7,11,12,35,36,37^. Several studies using independent methods have now dated the mixture of these divergent lineages to form the ancestors of modern humans to approximately 300 ka^7,11,12^, although a caveat is that the inference is not stable across populations plausibly reflecting multiple layers of mixture in Africa^11,36^, and some studies have preferred models of persistent substructure^37^ consistent with a view of human ancestors as a deeply structured metapopulation^38^. Nevertheless, the coincidence of the ∼300 ka inference over multiple studies is significant, as it increases the weight of evidence for the theory that an expanding early modern human lineage impacted both Europe and Africa around this time.

Taken together, these results—now corroborated by an independent study that analyzed DNA sequence motifs that are bound by the *PRDM9* gene determining where in the genome recombinations occur^7^—imply a view in which before ∼300 ka, there were at least three major lineages that contributed to high coverage sequenced human genomes (Figure 1). The first contributed to modern humans living today but not to Eurasian archaic humans (Denisovans and Neandertals), likely reflecting African archaic humans. The second contributed to Denisovans and Neandertals but not to modern Africans, likely representing Eurasian archaic humans. The third— showing evidence of activity of the “*PRDM9*-A” allele, which Denisovans lack—is the expanding early modern human lineage represented in admixed form in Neandertals and modern humans living today, but not in Denisovans: a successful early modern human population. In Europe, the expansion of this population led to large-scale incorporation of local archaic genes because barriers to producing viable offspring were few, giving rise to Neandertals who ultimately harbored 3-7% ancestry from the expanding early modern human lineage. In Africa, the mixture was with a more divergent lineage, which meant that there was less permeability of the expanding population to admixture with the local archaic group. Thus, the final population which became our own ancestors had a much higher proportion of ancestry from the expanding early modern human lineage^11^.

## A range expansion affecting Africa and West Eurasia correlates to archaeological evidence

Is the genetic evidence for major movements of people linking Neandertals in Europe and early modern humans in Africa ∼300 ka, to the exclusion of Denisovan ancestors in East Asia, mirrored by archaeological data? Indeed it is. The genetically inferred expansion follows the time when a tool production technique (Levallois points) based on prepared stone cores spread across a wide area of Africa where it was used by early modern humans (known there as the Middle Stone Age and starting by at least 400 ka^39,40^), and to the Middle East and Europe where it was used by Neandertals (known there as the Middle Paleolithic, starting by at least 480-300 ka^41^). There is evidence that these technologies were used much less intensively or not at all in East Asia where the closest cousins of early modern humans and Neandertals were Denisovans. In China, the sole strong claim of Levallois in the Middle Paleolithic is at the site of Guanyindong Cave dating to ∼170-80 ka^42^, but this is not a confident inference as another study refutes this interpretation^43^. Levallois has been documented in the Altai Mountains^44^, but this region was proximate to West Eurasia and the technology could have spread from there through contact with Neandertal populations which are well-documented in the Altai. Similarly, evidence of Levallois in India^45^ ∼250-200 ka could have been spread by modern human populations from the Near East.

The theory that the spread of Levallois stone tools may be a material marker for early human migrations was suggested decades ago as the “Mode 3 Hypothesis,” with the authors writing: “Neanderthal and modern lineages share a common ancestor… 350,000 and 250,000 years ago rather than in the earlier Middle Pleistocene; this ancestral population, which developed mode 3 technology (Levallois/Middle Stone Age), dispersed across Africa and western Eurasia in a warmer period prior to independent evolution towards Neanderthals and modern humans.”^46^ This proposal has been questioned based on evidence that prepared core techniques have been invented on multiple occasions including on the order of one million years ago in Africa^47^, and reconstructions suggesting that Levallois itself may have been invented independently^48^. Whether or not Levallois points were independently invented, the period preceding the ∼300 ka range expansion also corresponds to at least four other shared material culture features that increase the plausibility of a spread of an early modern human lineage from a core region to broader Europe and Africa around this time: improved hafting, expansion in the use of controlled fire^49^, use of ochre, and larger and more organized campsites^50^. All of these developments became widespread both in West Eurasia and in Africa and were much less present or entirely absent in East Asia, providing independent archaeological evidence for connections. The archaeological record from this period is thin and the accuracy of date estimates too poor to determine if the timings of appearance of each these cultural traits were coincidental enough to point to a range expansion.

But taken together, these results increase the weight of evidence for the hypothesis^10^ that some of these shared material culture features may have spread through the demographic expansion of a successful early modern human group that has been inferred here entirely through genetic analysis.

## Catalog of mutations that may contribute to distinctively early modern human biology

Previous studies have compiled lists of genetic variants present in nearly all living people, but absent in Neandertals and Denisovans, under the hypothesis that these are likely to be enriched in alleles that contributed to unique modern human innovations^51^. But the evidence presented here indicates that these catalogs are likely to screen out an important class of mutations: ones that evolved in the early modern human population prior to its bi-continental range expansion, and that may have supported behaviors shared between Neandertals and African modern humans like the intensive use of Levallois point tools, ochre, fire, and hafting; and the propensity to use larger and more organized campsites.

To search for candidate mutations that are recently shared between modern humans and Neandertals and absent in Denisovans, I analyzed the output of an Ancestral Recombination Graph inferred by the ARGweaver-D software^4^. This process identified 0.06% of chromosomes 1-22 where 16 African haplotypes and 4 Neandertal haplotypes are estimated to share a common ancestor more recently than 625 ka, while the 2 Denisovan haplotypes share an ancestor with them only more than 700 ka ago (Supplementary Information section 4). Altogether I identified 1,576 one kilobase windows that merge into 141 loci and in turn fall into 66 clusters (Extended Data Table 1). I estimate that 77% of these windows are genuine instances of shared ancestry of the sampled African and Neandertal haplotypes after their separation from the population ancestral to Denisovans, based on the excess over what I obtain when I repeat the experiment with the Neandertal and Denisovan labels exchanged. Ten of these clusters are visualized in Extended Data Figure 5. One includes the genes *FADS1* and *FADS2*, where modern humans and Neandertals share a common ancestor around 544 ka. This locus has been under repeated selection in the last hundreds of thousands of years, allowing more efficient conversion of polyunsaturated fatty acids that typically come from plants^52^, and raising the possibility of selection for a more diversified diet during the early modern human expansion. A second overlaps *ARHGAP32* which modulates the structure of neural connections, and contains Glu1489Asp, one of six amino-acid-changing substitutions where nearly all sampled modern humans and Neandertals share an ancestor <700 ka while Denisovans are much more distantly related (Table 1 and Extended Data Table 2).

**Table 1.**
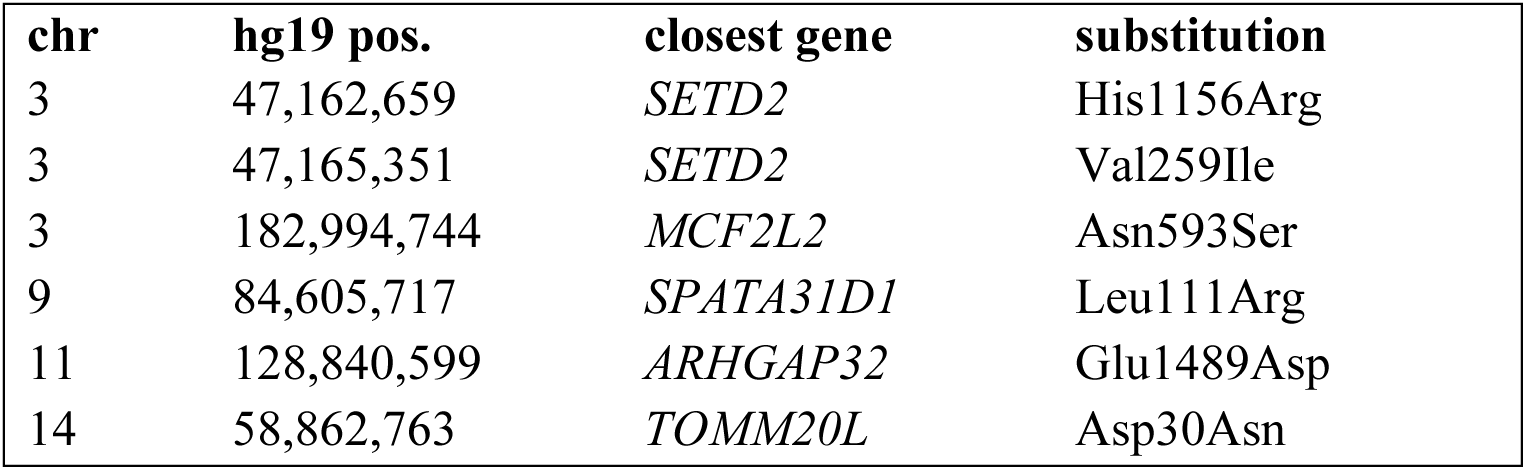
Nonsynonymous substitutions where modern humans and Neandertals share a common ancestor <625 ka and Denisovans are more distantly related.

I tested if the catalog is enriched for particular categories of gene function^53^, thereby identifying nine groupings of genes (of 536 tested) largely associated with protein catabolism that showed significant enrichment even after correction for multiple hypothesis testing (False Discovery Rate q-values from 0.001 to 0.006) (Extended Data Table 3). A notable enrichment is seen for genes involved in proteasome-mediated ubiquitin-dependent protein catabolism, suggestive of viral interactions^54^. Virus-interacting proteins account for an estimated 30% of adaptive amino acid substitutions in the mammalian-conserved part of the human proteome^55^, and so are an intriguing candidate for being involved in early-modern-human-specific evolution in this period.

## Discussion

The hypothesis of an early modern human range expansion ∼300 ka that impacted Neandertal ancestors is not new to this study: multiple studies have provided evidence of early modern human gene flow into Neandertals^2,3,4,5,6^, and since the gene flow occurred in a geographic context, it was almost certainly a range expansion. What is new to this study is the demonstration that the only plausible way to explain the data is the unique population genetic dynamics associated with a geographically explicit range expansion—large-scale incorporation of local genes into a population that maintained coherence from generation to generation as it expanded into a new territory—which means that the range expansion would have been a plausible vector for spreading culture even if the ultimate proportion of ancestry from the early modern human ancestors was small (∼3-7%). In Europe, Neandertal ancestors formed as this expanding early modern human population incorporated a stream of local archaic people, retaining early modern human mtDNA and Y chromosomes while having most of the rest of their genome replaced by archaic genes. In Africa, the ancestors of living humans formed as this expanding population also incorporated local archaic genes. By establishing a vector of major movements connecting West Eurasia and Africa, these results increase the weight of evidence that at least some of the distinctive shared archaeological patterns between West Eurasia and Africa—including use of Levallois points, controlled use of fire, use of ochre, improved hafting, and larger and more organized campsites— spread by the bi-continental, large-scale early modern human range expansion hypothesized here.

## Supporting information

Supplementary Information section 1

Supplementary Information section 1 code base

Supplementary Information section 2

Supplementary Information section 2 code base

Supplementary Information section 3Supplementary Information section 1

Supplementary Information section 3 code base

Supplementary Information section 4

Supplementary Information section 4 code base

## Materials and Methods

Details of the methods used in this paper are presented in the Supplementary Information.

## Acknowledgments

This research was inspired by a presentation Janet Kelso gave at the Boston Evolutionary Genomics Supergroup in September 2022, in which she proposed the puzzle addressed here. The work benefited from comments either in-person or through public forums by Bridget Alex, Mira Bernstein, Amy Clark, Trevor Cousins, Rob Foley, David Gokhman, John Hawks, Alexander Kim, Hrushikesh Loya, Dan Lieberman, Iain Mathieson, Simon Myers, Svante Pääbo, Nick Patterson, Molly Schumer, Ashwin Sivakumar, Pontus Skoglund, Daniel Tabin, Shamam Waldman, Tianyi Wang, and Richard Wrangham. I am grateful to Melissa Hubisz and Adam Siepel for sharing the output of their ancestral recombination graph reconstruction of relationships among 8 Africans, 2 Neandertals, and a Denisovan, and to Leo Speidel for sharing his reconstruction for individuals from the 1000 Genomes project. I acknowledge support from the National Institutes of Health (grants HG012287 and HG013987), the Howard Hughes Medical Institute.

## Use of artificial intelligence tools

Many of the analyses in this study are the product of a dialogue with artificial intelligence tools (Claude Sonnet, Opus, and Fable models, sometimes implemented within Cursor Integrated Development Environment). The AI tools wrote the code and generated figures using Python, and I checked the robustness of the results. For the supplementary notes but not for the main manuscript, some of the text was composed by the AI and I edited it.

## Conflict of Interest Statement

I declare no competing interests.

## Data Availability Statement

The quantitative simulation results are available as Supplementary Table 1, and the results of the scan for loci and variants that may have provided advantages to early modern humans are available as Supplementary Tables 2 and 3.

## Code Availability Statement

The scripts used to generate the analyses and figures are available as four zipped files, one for each Supplementary Information section.

## Supplementary Information Guide

### Supplementary Information sections

(SI 1) Biological constraints mean that in the absence of a range expansion, fixation of early modern human lineage mtDNA and Y chromosomes in Neandertals is unlikely

(SI 2) No evidence for a Dobzhansky-Muller Incompatibility between Denisovan mtDNA and modern human nuclear DNA

(SI 3) Modern human range expansions predict fixation of modern mtDNA and Y in Neandertals (SI 4) A catalog of mutations that may have been important in uniquely modern human biology

### Analysis Scripts (separate zipped folder with a README for each SI)

1. SI1codebase.zip
2. SI2codebase.zip
3. SI3codebase.zip
4. SI4codebase.zip

### Supplementary Video

(Supplementary Video) Animated version of Figure 3a

### Supplementary Tables

(Supplementary Table 1) Simulation results: Excel file with 15 tabs providing all the simulation results and details on the simulation setup

(Supplementary Table 2) gzipped tsv consisting of 2.44 million rows and the results of the scan for loci that may have provided advantages to early modern humans

(Supplementary Table 3) tsv with 1,223 rows: sites in candidate introgressed regions that are high-frequency-derived in Neandertals and modern humans and absent in Denisovans

---

**Extended Data Figure 1:**
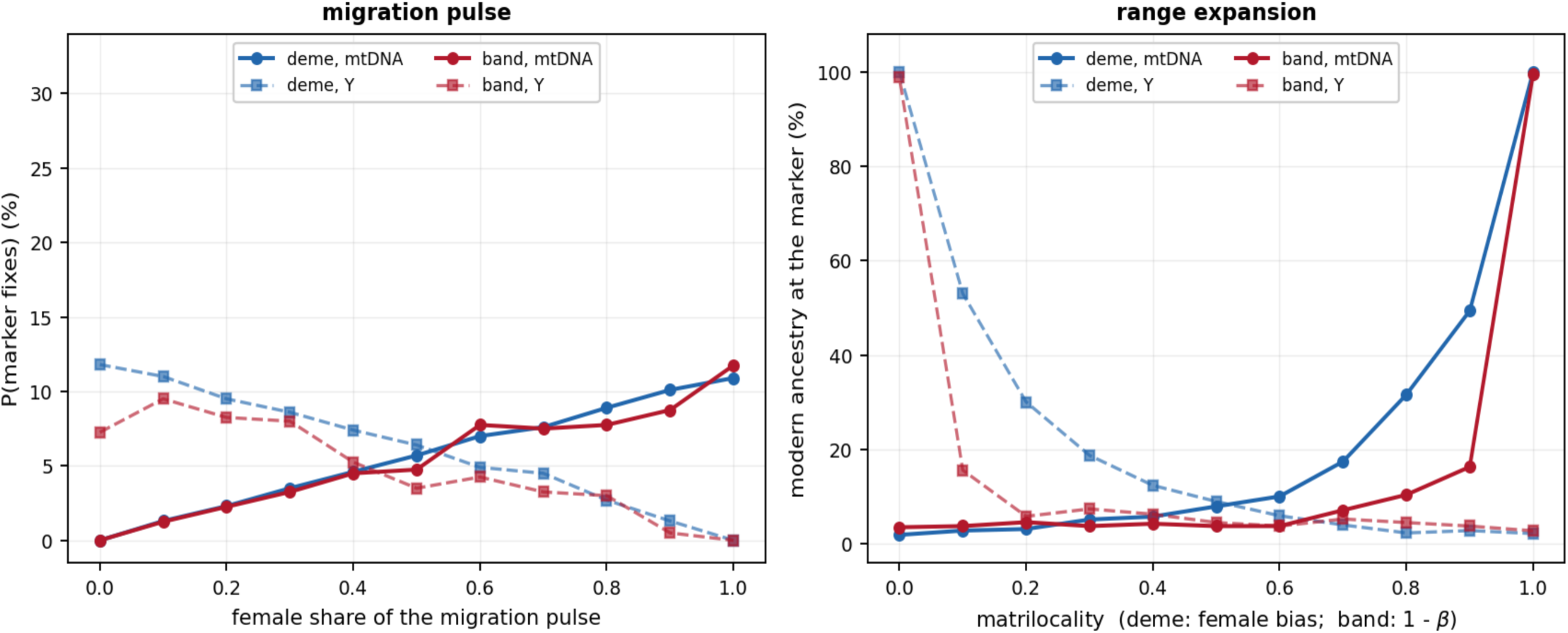
Sex-bias drives one uniparentally inherited marker to fixation in a range expansion, but not in a non-geographically-explicit model. 1,000 replicates per point. *Left: Non-geographically-explicit.* 5% gene flow from early modern humans into archaic humans is followed by drift. Even under complete sex bias, fixation is unlikely (∼10%). *Right: Range expansion.* As the probability that the child of a cross-lineage pairing joins its mother’s group nears 1, fixation of the corresponding marker becomes likely. This does not explain the pattern at the other uniparentally transmitted marker, where biological or cultural selection is needed.

**Extended Data Figure 2.**
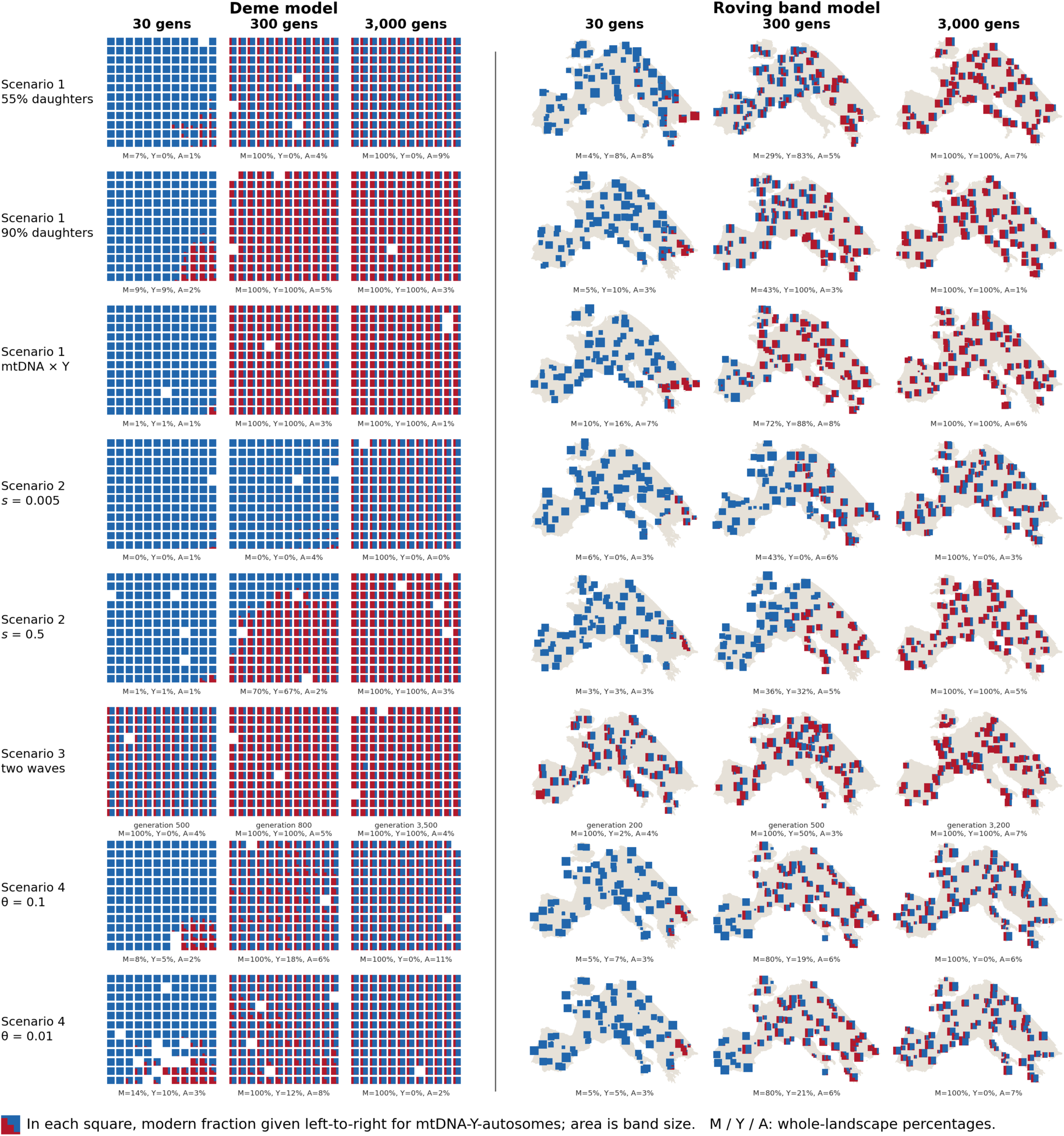
Robustness across both simulation frameworks. Each row is an example of one mechanism at one strength, drawn on the left in the deme framework and on the right in the roving band framework, at 30, 300 and 3,000 generations after contact. Each square is one deme or one band, with three contiguous bars giving the modern fraction at mtDNA, the Y and the autosomes from left to right. The percentages below each panel are the whole-landscape values at that generation for the modern human subset of individuals.

**Extended Data Figure 3.**
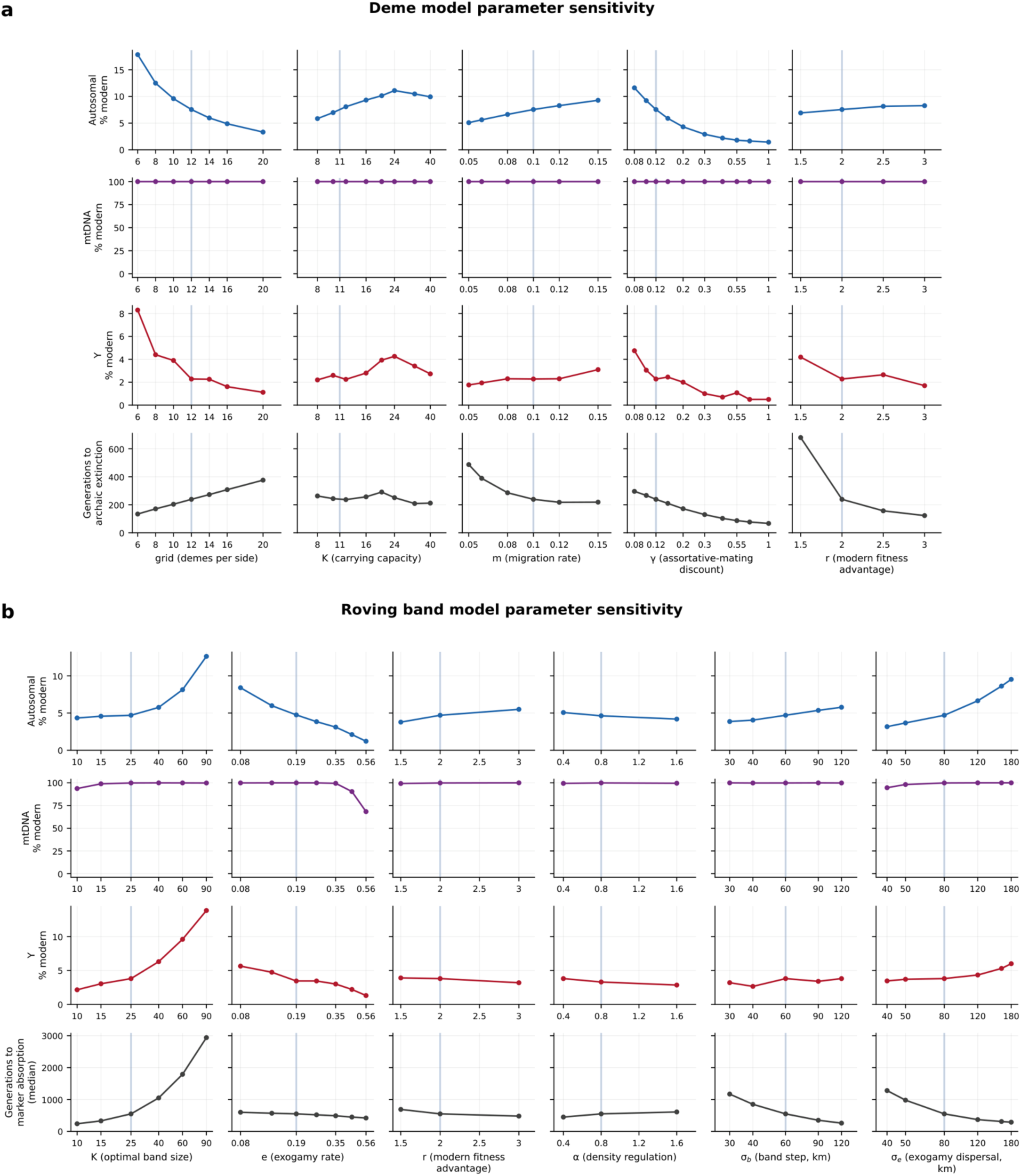
Sensitivity of deme and band frameworks to each parameter. 2,000 replicates per point in both. Light vertical line indicates the baseline value from which each axis varies the parameter.

**Extended Data Figure 4:**
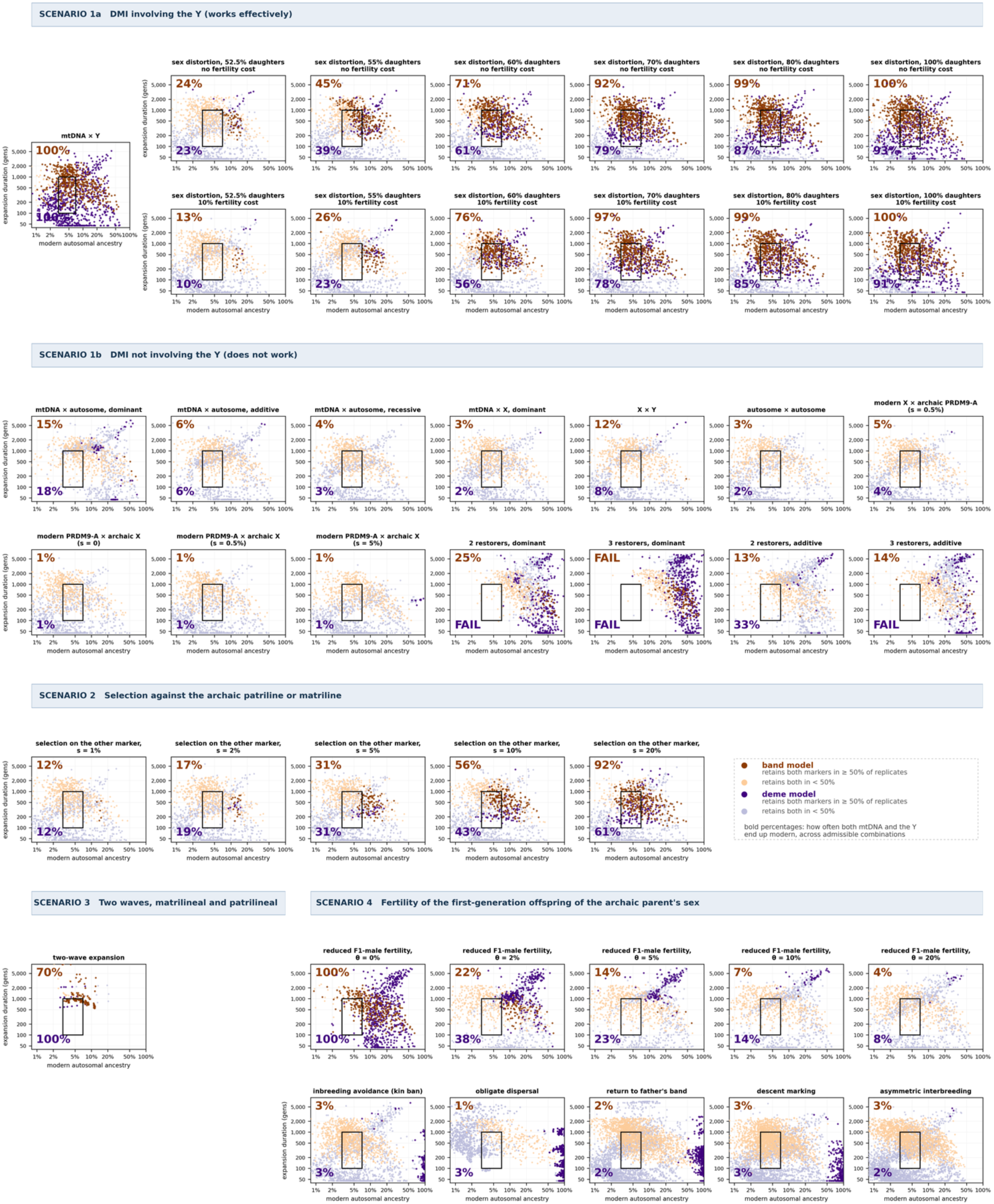
Systematic survey of biological and cultural scenarios matching the data. Orange band model, purple deme model, points full grid of parameters. Black rectangle is acceptance region modern autosomal ancestry 3-7%, expansion in 100-1,000 generations). Mean probability that all ten sampled males are modern at both mtDNA and the Y is shown for band top left, deme bottom left, averaged over admissible combinations (FAIL marks no admissible combination). Dark points are where a sample of ten males is modern at both markers with probability ≥50%. Every point inside the box is plotted in the main Figure 3b associated with the same scenario. Each cell is judged on 100 or 200 replicates per point.

**Extended Data Figure 5:**
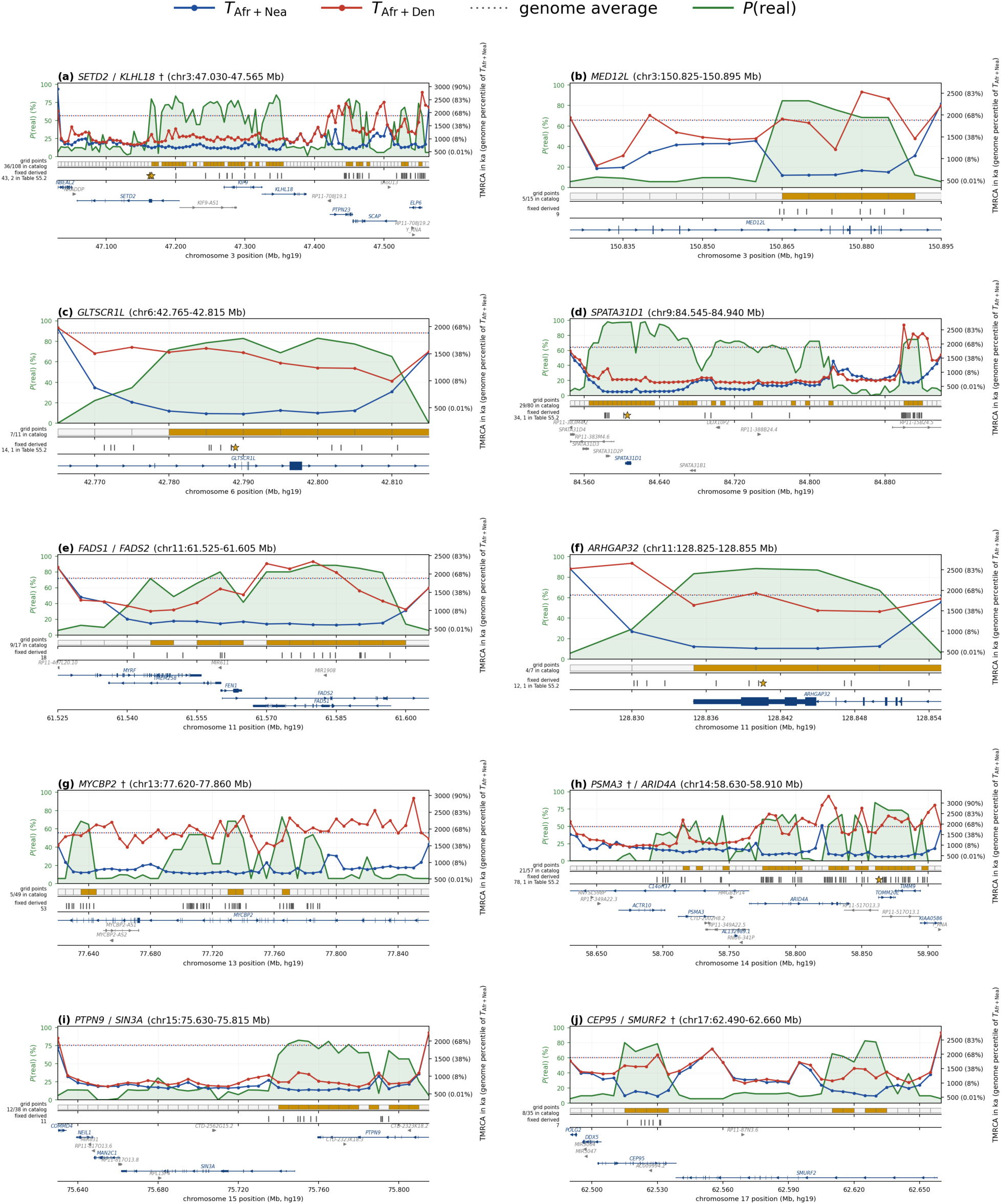
Ten notable genomic regions where present-day humans and Neandertals hare an extraordinarily recent ancestor making them candidates for harboring evolutionary adaptations important to early modern humans. The estimated probability of modern humans and Neandertals sharing a common ancestor more recently than the population split from Denisovans is shown on the left axis as *P(real)*. The right axis shows the mean time to the most recent common ancestor of both Africans and Neandertals TAfr+Nea, and Africans and Denisovans TAfr+Den (parentheses give percentile in the genome-wide distribution of TAfr+Nea; dotted lines genome-wide averages). Below the graphs, I show the grid points that qualify for inclusion in the catalog, fixed derived sites (starred if predicted to be highly functional), and GENCODE v19 genes with coding sequence drawn thickly.

**Extended Data Table 1:**
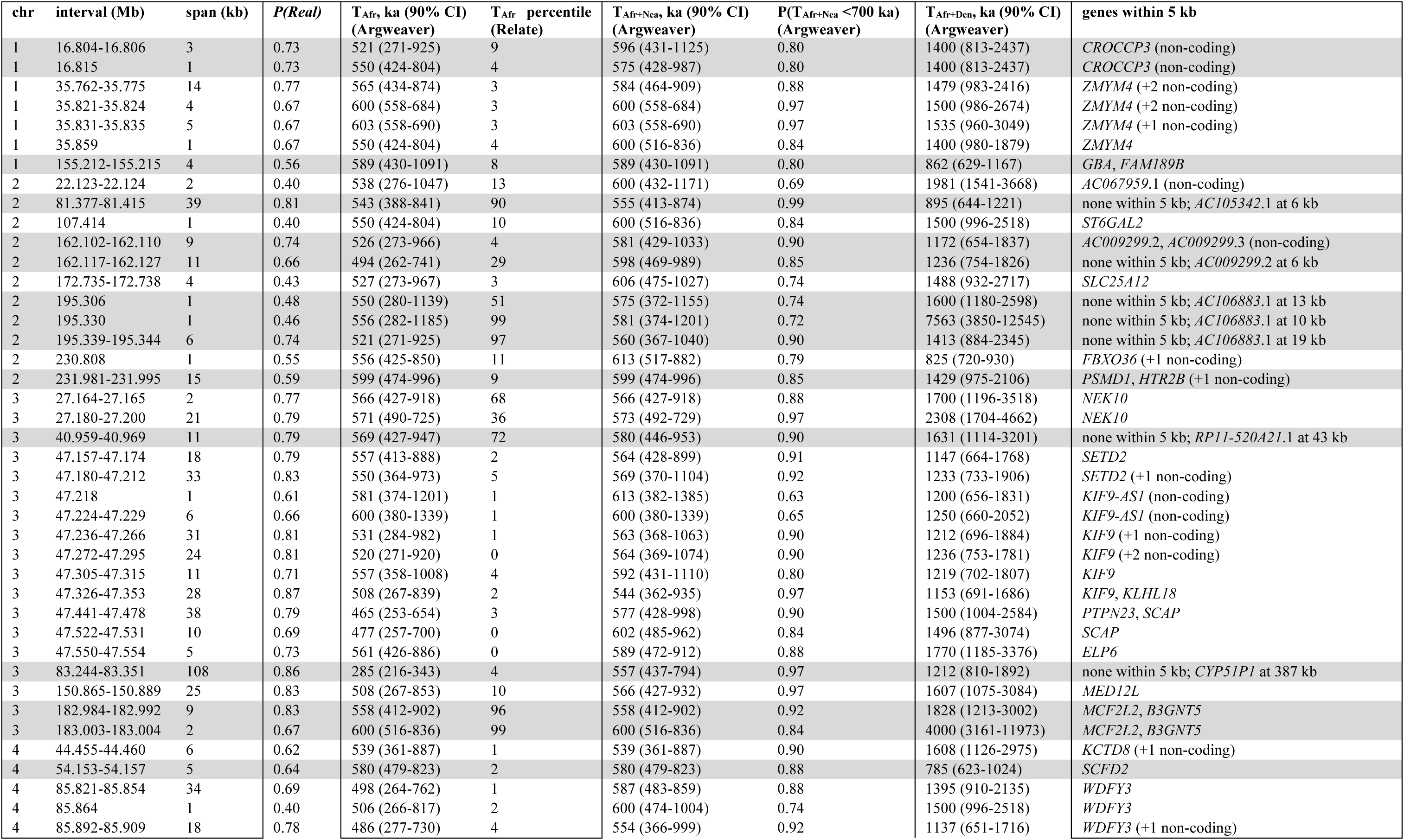
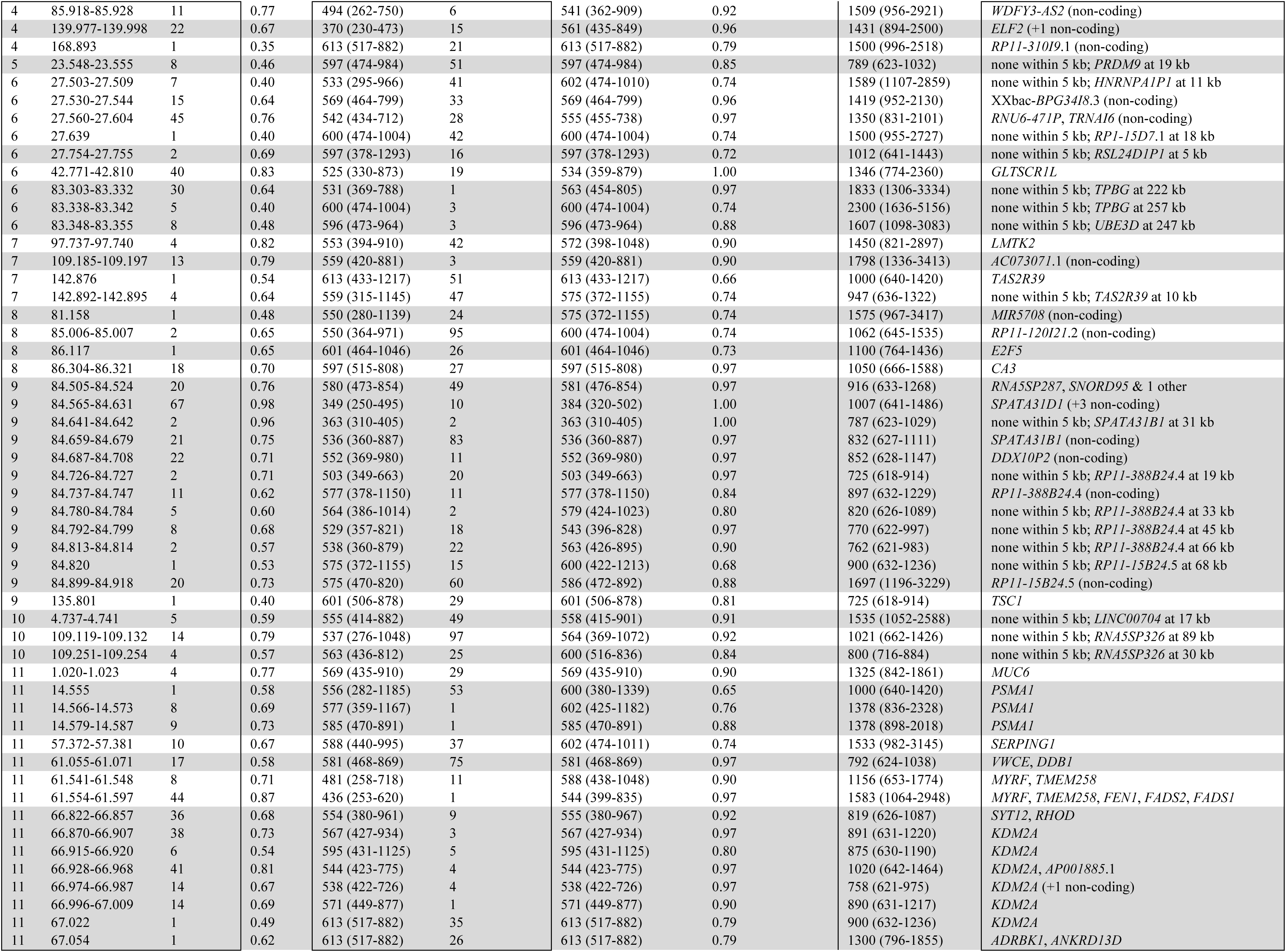
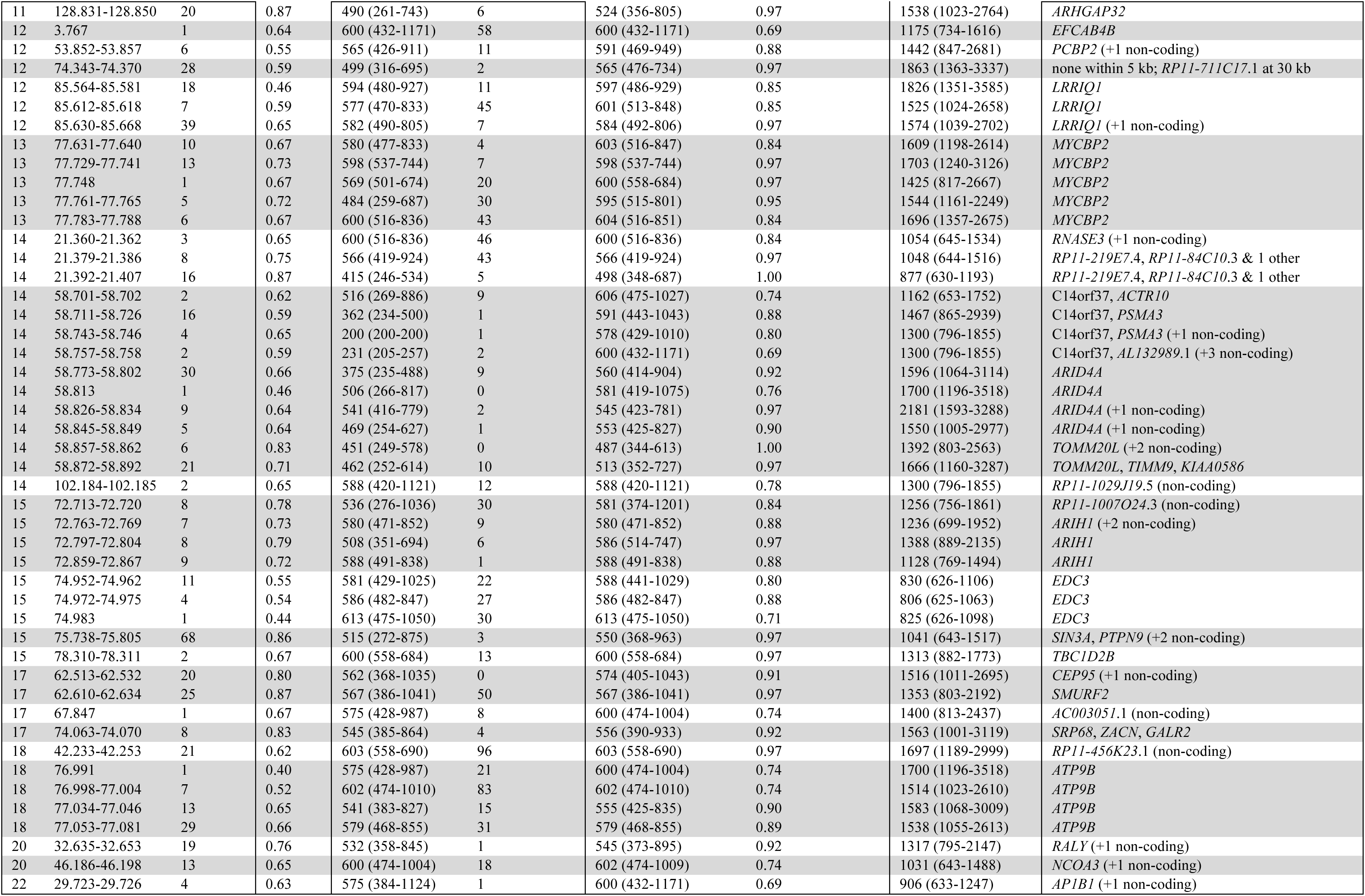
Catalog of candidate loci for early modern human adaptation. 141 runs of 1 kb autosomal grid points (points within 5 kb ne another merged into a single locus) where 16 sampled modern human and 4 Neandertal haplotypes derive from a common ancestor ≤625 ka le 2 Denisovan haplotypes diverge ≥700 ka based on parsing of an ARGweaver-D ancestral recombination graph (ARG). I pool loci within 100 kb orm 66 clusters (gray striping). Each TMRCA is the posterior median with its 90% central interval. “P(<700 ka)” is the posterior probability that modern-plus-Neandertal coalescence is below the 700 ka threshold at the gridpoint within the locus where this quantity is highest. “Relate centile” is the inferred time to the most recent common ancestor compared to the autosome-wide distribution in a larger panel of 216 West African uba haplotypes. “P(real)” from the 1 kb grid point in each locus showing the strongest posterior signal of enrichment.

**Extended Data Table 2.**
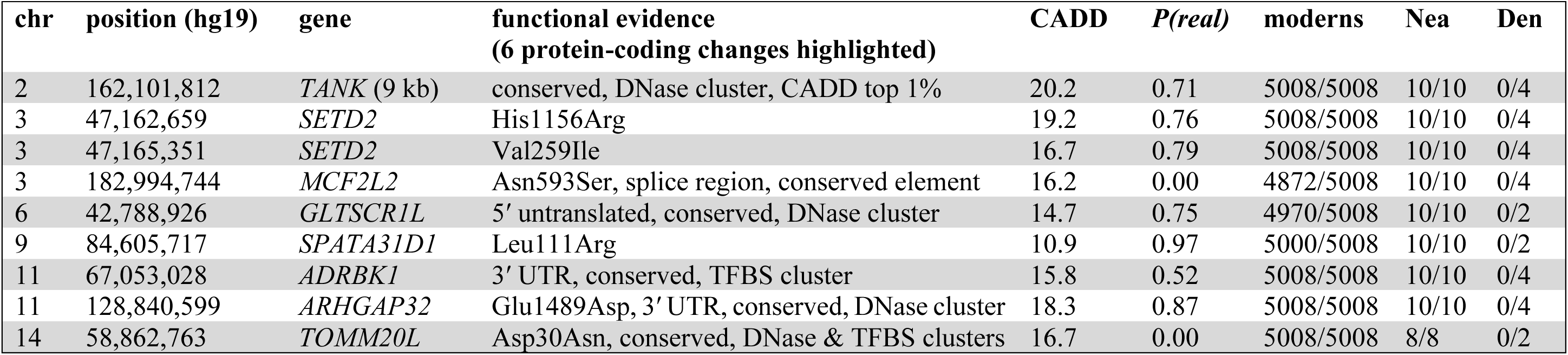
Candidate variants of high predicted functional significance. The 6 amino-acid-changing substitutions in Table 1 of the main text are a subset of the 9 variants here. CADD measures predicted deleteriousness by combining functional annotations into one score on the PHRED scale of -10 × log10(percentile), for this table running from 10.9 (top 10%) through 20.2 (top 1%). Haplotype columns give derived over total count for 5,008 modern human haplotypes, up to 10 haplotypes from high-coverage Neandertals, and up to 4 haplotypes from high-coverage Denisovans. Gene column gives the (nearest) gene body. P(real) is evaluated at the substitution itself rather than across the locus around it, and is 0.00 at MCF2L2 and TOMM20L because the modern-plus-Neandertal coalescence inferred at those exact positions is older than the 625 ka catalog ceiling even though the surrounding locus qualifies; both loci are at P(real) = 0.83 in Extended Data Table 1

**Extended Data Table 3.**
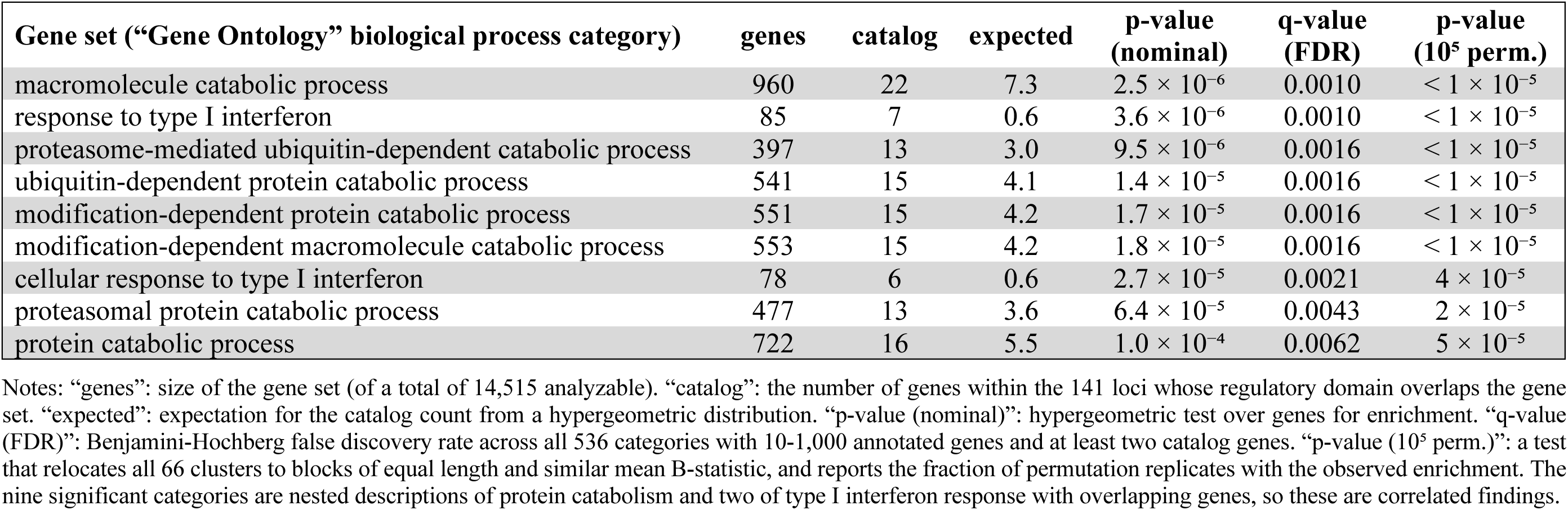
Gene set most strongly enriched for recent modern human - Neandertal ancestors.

## Notes

### Competing Interest Statement

The authors have declared no competing interest.

### Summary of Updates

This new submission reflects extensive new analysis, detailed in four supplementary notes and three supplementary tables. It is changed relative to the original manuscript in three main ways: (1) It is reorganized to be *genetics first*. The original manuscript was presented as a synthesis of genetic and archaeological observations. Rather than starting from a premise that depended on an interpretation of archaeology, I now use entirely genetic arguments to show that models that have been previously used in simulations of the ∼300-200 ka interbreeding of early modern humans and Neandertal ancestors are highly unlikely to fit. But the proposed new model explains disparate observations. I begin by showing that a non-geographically-explicit model that was simulated by Petr et al. Science 2020 is unlikely to explain the observed joint patterns for mtDNA, the Y chromosome, and the autosomes. Given what we know about mtDNA biology, Y chromosome conservation patterns across mammals, and selective constraints, I show that this model has less than a 2-3% probability of being consistent with the data. I then go on to show that geographically explicit simulations of a range expansion accommodate multiple scenarios that predict the data pattern with high probability. Only at the end do I link to the archaeological evidence. (2) Through my new simulation studies, I identify multiple plausible scenarios that would explain the Y chromosome and mtDNA findings with high probability. One is a sex-biased modern human range expansion combined with individuals from matrilines or patrilines tracing their lineages back to modern humans. A second is a modern human range expansion where females stay within the group and where hybrid males have a skew toward at least 60% daughters. A third example is two successive range expansion: one where females stay within the group (spreading modern human lineage mtDNA) and one where males stay within the group (spreading modern human lineage Y). (3) I present a catalog of locations where all living humans and Neandertals share ancestors <625,000 years ago with Denisovans far more distantly related; these previously overlooked loci are candidates for key early modern human innovations. One example is a haplotype at the FADS1 and FADS2 genes which is involved in more efficient conversion of plant-derived fatty acids raising the possibility that a diversified diet could have been adaptive during the early modern human population expansion. Another is at the gene ARHGAP32 which modulates the structure of neural connections.

