## Supplementary Information section 1 for "Hypothesis: A modern human range expansion ∼300,000 years ago had a critical role in Neandertal origins"

Biological constraints mean that in the absence of a range expansion, fixation of early modern human lineage mtDNA and Y chromosomes in Neandertals is unlikely

**Findings**

- Muller's Ratchet is unlikely to have driven modern mtDNA to fixation in Neandertals: I estimate its fixation probability to have been less than 23%.
- Muller's Ratchet is unlikely to have driven the modern Y chromosome to fixation in Neandertals: I estimate its fixation probability to have been less than 10-13%.
- Muller's Ratchet is unlikely to have driven both modern mtDNA and Y chromosome to fixation in Neandertals: I estimate their joint fixation probability to have been less than 2-3%.

**Motivation**

Mixture of archaic and early modern human lineages in Europe 300-200 thousand years ago (ka) has been inferred to have produced a population with ~3-7% early modern human ancestry and the remainder archaic ancestry: the ancestors of Neandertals[1,2,3]. Natural selection had differential impacts on the DNA inherited from these two mixing populations. On average across the autosomes, early modern human ancestry was selected against, as can be seen from the fact that close to genes, DNA derived from early modern humans occurs less often than the genomic average[1,3]. This is the pattern expected if large numbers of Dobzhansky-Muller Incompatibilities (DMIs) existed between early modern human and archaic human genomes, leading to selection against minority ancestry in hybrids[4].

This note is motivated by the hypothesis that mitochondrial DNA (mtDNA) and the Y chromosome could be an exception to the pattern of selection against early modern human ancestry. These loci are unusual in that they contain a large number of fully linked genes that are not decoupled by recombination, making them candidates for being affected by "Muller's Ratchet", whereby an accumulation of slightly deleterious mutations at a large non-recombining locus is expected to reduce fitness faster in a population of smaller size[5]. From ~750 to ~250 ka, between the separation of the archaic and early modern human populations and their putative re-mixture, my digitization of the curves in Figure 4A of ref. 6 suggests that the effective number of diploid individuals in the early modern lineage was $N_{M\text{-diploid}}$ = 11000-21900, with a harmonic mean of around 14000 (the harmonic mean is the quantity that matters most for the effectiveness of selection). For the archaic lineage, the comparable estimates are $N_{A\text{-diploid}}$ = 1300-10300, with a harmonic mean of around 4500. For the Neandertal ancestral population following mixture between ~250 and ~50 ka, the estimates are $N_{N\text{-diploid}}$ = 400-2300, with a harmonic mean of around 1450. Because mtDNA is transmitted only through females and the Y only through males, the effective number of copies of each is half of the diploid value, so $N_{M\text{-uniparental}}$ = 7000, $N_{A\text{-uniparental}}$ = 2250 and $N_{N\text{-uniparental}}$ = 725. These are the primary values I use in my simulations.

Previously published simulations of Y chromosome dynamics by Petr and colleagues[7] used central values of $N_{M\text{-uniparental}}$ = 5000 and $N_{A\text{-uniparental}}$ = 500, with divergence at 600 ka and gene flow between 450 and 150 ka (midpoint 300 ka, which I use as a fixed time of mixture in my simulations to replicate their setting). They held the archaic population constant from divergence onward, so their post-mixture size is also $N_{N\text{-uniparental}} = 500$. The lack of modeling of a larger population size prior to mixture, followed by a contraction after mixture, is a limitation of the Petr et al. simulations[7] that my new simulations sought to address. This fuller modeling is potentially important, since deleterious mutations accumulate at one population size but the fate of the introgressed haplotype is decided at another. In what follows, the main numbers are therefore from my simulations, but I compare to results I obtain using parameters similar to those of Petr et al.[7]

In the preprint version of the present study[8], I also carried out simulations of Muller's Ratchet and inferred that it is unlikely to have driven the modern human mtDNA to fixation. Here, I greatly expand these simulations by simulating a semi-realistic distribution of fitness effects (DFE), by considering the impact of the non-protein coding part of the mitochondrial genome, by conditioning on observed patterns of non-synonymous substitutions, and by also extending the analysis to the Y chromosome.

The text and analyses in this note are the product of an interaction between the author and AI coding agents of the Anthropic Claude family (Sonnet, Opus, and Fable models), for the most part running inside the Cursor Integrated Development Environment. I used the agents to write the simulations and ABC code in Python, to run diagnostics, and to help compose the text (any suggestions from the AI agents were reviewed and edited by me). All quantitative results are produced by my execution of the cited scripts on the O2 computing cluster at Harvard Medical School, or on my personal computer. I checked the robustness of the output, and reviewed the scientific interpretations, modeling choices, and final claims.

### Methods

I simulate two isolated, randomly mating populations, which split from a common ancestral population at time $T_{sep}$, remixed at time $T_{mix}$, and were sampled at time $T_{samp}$. I assume the same effective sizes for females (mtDNA) and males (Y), which ignores effects due to differences in variability of reproductive success (expected to have a larger impact on Y). I held the modern lineage size $N_{M\text{-uniparental}}$ constant throughout. I held the archaic lineage size $N_{A\text{-uniparental}}$ constant during the period of isolation and then contracted it at the time of mixture to $N_{N\text{-uniparental}}$, in an attempt to capture the decline evident in Figure 4A of ref. 6:

- Modern-lineage constant effective size $N_{M\text{-uniparental}} = 7000$
- Archaic-lineage constant effective size during isolation $N_{A\text{-uniparental}} = 2250$ (for comparison I also simulate $N_{A\text{-uniparental}} = 1500$ and $3000$, bracketing this central value)
- Post-mixture Neandertal constant size $N_{N\text{-uniparental}} = 725$ (the fixation-probability analysis which simulates the post-mixture phase rounds this to 750 and brackets it with $N_{N\text{-uniparental}} = 500$ and $1000$, paired with $N_{A\text{-uniparental}} = 1500$ and $3000$ respectively)
- Separation time $T_{sep} = 26224$ generations ago (750 ka at 28.6 years per generation)
- Mixture time $T_{mix} = 8741$ generations ago (250 ka at 28.6 years per generation), giving 17483 generations of isolation
- Sampling time $T_{samp} = 1748$ generations ago (50 ka at 28.6 years per generation), giving 6993 generations between mixture and sampling
- For comparison, I also simulate a parameter set modeled on Petr et al.[7]: $N_{M\text{-uniparental}} = 5000$, $N_{A\text{-uniparental}} = 500$, $N_{N\text{-uniparental}} = 500$ (no contraction after mixture), $T_{sep} = 20979$ generations

ago (600 ka), $T_{mix}$ = 10490 generations ago (300 ka, giving 10489 generations of isolation), and $T_{samp}$ = 1748 generations ago (50 ka).

Both populations start at mean fitness 1.0, so the modern-archaic difference at the time of mixture $T_{mix}$ estimates the advantage the modern copy would have had when the lineages meet. The target is a non-recombining locus with no back-mutation (infinite-sites model). Fitness is multiplicative and tracked in log space (combining mutations via addition). I use a haploid Wright-Fisher process with viability selection and a per-mutation DFE:

1. Selection + reproduction: parents for the next generation are resampled with probability proportional to multiplicative fitness *w*.
2. Mutation: For analyses of protein coding sequence, each offspring receives a Poisson(U_missense) number of amino-acid-altering mutations, each multiplying its fitness by (1-s) with s drawn from the locus's gamma-distributed DFE (capped at 1); plus a Poisson(U_nonsense) number of nonsense mutations, each lethal (s = 1). For analyses that include non-coding sequence, I model a second deleterious class in the same way, with its own rate and its own gamma. I do this in most detail for the RNA gene class, but also use the same framework for non-coding sequences on the Y chromosome. The mtDNA and Y chromosome cases differ in the constraints that the data support. For the mtDNA RNA genes, a separate fit gives a DFE indistinguishable from the coding one in the nearly neutral range, which I therefore use, and divergence shows about 0.77 of their mutations to be deleterious. For Y chromosome non-coding sequence, there is not enough data to meaningfully estimate the DFE, so I count every mutation at a constrained base and scan the mean from that of the coding DFE down to 1/30 of it (this is conservative: weaker mean selection coefficients can translate to a large fitness gap, since what contributes to a fitness gap is coefficients that are more efficiently removed in the larger modern than archaic population).

**Table S1.1: Parameters used for mitochondrial DNA and Y chromosome analyses**

| **Property** | **Whole mitochondrial DNA** | **Y chromosome** |
|---|---|---|
| Inheritance | Maternal; haploid, clonal | Paternal; haploid, clonal |
| Recombination | None | None in single-copy region |
| Modern lineage size $N_{M\text{-uniparental}}$ | 7000 (female) | 7000 (male) |
| Archaic lineage size $N_{A\text{-uniparental}}$ | 2250 (female) | 2250 (male) |
| Neandertal post-mixture $N_{N\text{-uniparental}}$ | 725 (female) | 725 (male) |
| Neutral substitution rate (per bp/gen) | $9.35\times10^{-7}$ (ref. 9) | $3.0\times10^{-8}$ (range 2.5-3.5)[10,11] |
| Non-recombining constrained target | 11,388 bp (13 genes) plus 4,013 bp RNA genes, of 16,569 bp | 33,156 bp codons plus 79 kb measured constrained non-coding, of 16.5 Mb single-copy euchromatic |
| Fraction of mutations drawn from the DFE | 0.60 coding; 0.77 RNA genes | 0.72 codons; 1.0 constrained non-coding |
| Deleterious input U (per gen) | 0.0093 including RNA genes (range 0.0055-0.0133); coding only 0.0064 | 0.0031 at the measured target (0.00072 codons alone) |
| Distribution of Fitness Effects (DFE) | Appendix A, extending ref. 12; same for RNA genes | Ref. 13, rescaled; non-coding same shape, mean scaled by 1 to 1/30 |
| Gamma shape $\beta$ | 0.245 (95% CI 0.13-0.35) | 0.23 (95% CI 0.19-0.27) |
| Mean unscaled *s* | 0.069 | 0.043 |

*Mitochondrial DNA*

I simulate new mutations in mtDNA by drawing selection coefficients from a gamma-distributed DFE, following the approach of an earlier study (Table S1.1)[12]. That study, focusing on non-synonymous sites at protein-coding bases, analyzed the joint patterns of divergence across species and polymorphism within species to infer the mtDNA DFE. In humans, using data from 53 human mtDNAs, the study inferred a shape parameter $\beta = 0.39$ (95% CI [0.36, 0.50]) and a mean scaled coefficient $\bar{S} = -1400$ (95% CI [-2400, -420]). I re-implemented the estimator of that study and found, running at the same sample size, that I could closely reproduce these results ($\beta = 0.39$, $\bar{S} = -1553$; Appendix A, Table S1.A1). I then extended the estimator beyond the 13 protein genes to the tRNA and rRNA genes, so that one DFE is fitted to the whole non-recombining locus. This whole-mtDNA fit gives $\beta = 0.245$ (95% CI [0.13, 0.35]) and $\bar{S} = -968$, with a near-neutral mass $P(1 < |S| < 10) = 0.110$ (95% CI [0.063, 0.146]). The near-neutral mass, the only feature of the DFE the ratchet is sensitive to, is well-determined even if the absolute $|\bar{S}|$ is not (Appendix A).

The scaled coefficient $\bar{S} = 2N_e s$ must be converted to an unscaled s. Here $N_e$ is the effective number of haploid mtDNA copies (the female effective size), not a diploid autosomal size (the factor of 2 is the standard convention for haploid loci, and the same convention is used in the DFE fit and in the simulations). Because I estimated the DFE from present-day human mtDNA, I anchor it at the modern female size, $N_{M\text{-uniparental}} = 7000$, giving $\bar{s} = \bar{S} / (2 \cdot N_{M\text{-uniparental}}) \approx 0.069$, with a median of 0.012 and a heavy lethal tail.

I leveraged the ancient-DNA-calibrated mitochondrial rate of ref. 9, whose Table 2 third-codon-position rate is $3.27 \times 10^{-8}$ substitutions/site/yr (95% CI $[1.94, 4.62] \times 10^{-8}$). Third-codon-position sites are approximately neutral, so this is effectively the neutral per-site rate, which at 28.6 yr/generation is $9.35 \times 10^{-7}$/bp/gen (CI $[5.55\text{-}13.2] \times 10^{-7}$). I do not use the higher deCODE Icelandic pedigree rate[14], which at synonymous sites is $6.36 \times 10^{-6}$/bp/gen, about sevenfold the ancient-DNA-calibrated rate. Pedigree studies count variants at the moment they arise, as low-frequency heteroplasmies, the great majority of which are lost by drift through the germline bottleneck before they ever fix in a maternal lineage, and recurrent mutation at hypervariable sites further separates one-generation from long-term estimates. What the ratchet acts on is the rate at which mutations accumulate over tens of thousands of years, which the ancient-DNA calibration measures directly.

To apportion mtDNA mutations, I used the vertebrate mitochondrial code for the revised Cambridge Reference Sequence (NC_012920[15]), weighting each possible change by the extreme transition/transversion bias characteristic of mtDNA of 24.3.[14] The missense fraction is $f \approx 0.60$, nonsense $\approx 0.04$, and synonymous $\approx 0.36$. Multiplying $f$ by the protein-coding length (11,388 bp) and the per-bp rate gives a missense input U_missense ≈ 0.0064 (range [0.0038, 0.0091] across the 95% CI of the ancient-DNA-calibrated mutation rate). I treat premature stops as a separate lethal class, and synonymous mutations as fully neutral; neither matters to Muller's Ratchet, which is driven by nearly neutral mutations.

The mutational input above is restricted to the 13 protein-coding genes. mtDNA also carries 2 rRNA (12S + 16S, ~2513 bp) and 22 tRNA (~1500 bp) genes. Since previous work did not fit a DFE including the RNA genes, but doing so is essential in order to understand the rate of accumulation of deleterious mutations at mtDNA, I fit one with the same estimator used for the coding genes (Appendix A, section A4). The fit shows the RNA DFE to be statistically indistinguishable from the coding DFE in the near-neutral band that drives the ratchet. Not every RNA-gene mutation is deleterious: comparing the rCRS against high-coverage Denisovan,

Neandertal and chimpanzee mtDNA, with each site class normalized by its spectrum-weighted mutational opportunity, I estimate that purifying selection removes 0.76-0.80 of new RNA-gene mutations (0.80-0.84 in tRNA, 0.73-0.81 in rRNA). The spectrum-weighted deleterious RNA target is thus ~0.77 × 4,013 bp ≈ 3,100 bp, which at $9.35\times10^{-7}$/bp/gen adds U_RNA ≈ 0.0029 to the missense input of ≈ 0.0064. The Results therefore present the fitness gap with an RNA-inclusive input U ≈ 0.0093 (band 0.0055-0.0133).

To constrain the parameters relevant to the simulations, I also conditioned inferences on the observed numbers of synonymous and nonsynonymous substitutions on each lineage. For this purpose I use Approximate Bayesian Computation (ABC, rejection sampling from simulations)[16], simulating lineage pairs forward under the demographic model, accepting those whose simulated nonsynonymous counts are close to the observed values, and reading off the posterior distribution of the modern-archaic fitness gap at the time the lineages met. The synonymous count is neutral and so calibrates the molecular rate internally (the mutation rate is not an external free parameter, which means that uncertainty in the ancient-DNA-calibrated mutation rate is not a major source of variation in the inference). Across the 150,000 RNA-inclusive prior draws (50,000 per archaic size $N_{A\text{-uniparental}}$ = 1500 / 2250 / 3000), 11,698 (7.8%) met the acceptance criterion. The coding-only companion to this analysis accepted 11,224 of 150,000 (7.5%), and the comparison to the parameters used in the previously published study[7] accepted 1,270 of 50,000 (2.5%).

The simulation engine for the ABC analysis does the following. (i) It starts a Wright-Fisher simulation for a homogeneous population at 1000 ka; (ii) it draws a stochastic coalescence time per pair (an exponential tail older than the fixed 750 ka split, scaled by $N_{A\text{-uniparental}}$), giving a time to the most recent common ancestor (TMRCA) distribution centered near ~815 ka for $N_{A\text{-uniparental}}$ = 2250, consistent with the empirical mtDNA coalescence CI (800-967 ka); (iii) it tracks one focal modern and one focal archaic lineage; (iv) it defines the fitness gap on those two lineages at 250 ka (the introgressing modern haplotype vs. the archaic); and (v) it reads the synonymous/nonsynonymous counts at the actual sampling times (present-day modern; ~65 ka Denisovan). I simulate three ancestral sizes ($N_{A\text{-uniparental}}$ = 1500, 2250, and 3000, with an equal prior on all three), and integrate over uncertainty in the DFE and the molecular rate by drawing the pair (β, $|\bar{S}|$) jointly from the bootstrap distribution of the DFE fit and drawing the rate from its synonymous-count calibration.

The constraints used for the ABC analysis are shown in Table S1.2, obtained in ref. 17 from a maximum-likelihood estimate of synonymous and nonsynonymous substitution rates on each branch of the ((modern, Neandertal), Denisova) mtDNA tree. For the modern-vs-Denisova comparison since their common ancestor (the tree root), the modern lineage's path is the modern-human branch plus the modern-Neandertal ancestral branch: synonymous 38.3 + 79.5 = 117.8 and nonsynonymous 11.8 + 9.8 = 21.6. The Denisovan path is 108 synonymous and 17.9 nonsynonymous. The modern:Denisovan nonsynonymous ratio is 21.6 / 17.9 = 1.21, so there are actually more nonsynonymous substitutions on the modern lineage than the Denisovan one, even after controlling for the antiquity of the Denisovan individual, providing no evidence of an excess accumulation of deleterious mutations on the archaic lineage that could lead to fixation of the modern human lineage mtDNA. The observed value is within sampling error of the model, so it does not refute a ratcheting in the direction of retaining modern human lineage mtDNA, but it provides no support for it and helps to tighten the upper bound on the fitness gap. My ABC conditions on this ratio (acceptance window 1.16-1.26) and constrains the molecular rate using the synonymous count (117.8 + 108 = 225.8, neutral). I draw DFE parameters (β, $|\bar{S}|$) jointly from the bootstrap distribution of the whole-mtDNA fit (Appendix A) in each replicate. The ABC analysis conditions on protein-coding substitutions only; Appendix B sets out how the RNA load is carried through it without contaminating the acceptance statistic.

**Table S1.2: Empirically estimated branch mutation counts in mtDNA[17]**

| mtDNA lineage / branch | Synonymous | Nonsynonymous | dN/dS |
|---|---|---|---|
| Modern human | 38.3 | 11.8 | 0.105 |
| Neandertal | 39.2 | 13.6 | 0.118 |
| Modern-Neandertal ancestor | 79.5 | 9.8 | 0.042 |
| Denisova | 108 | 17.9 | 0.056 |

Given this setup, how large a modern-archaic mean-fitness gap does the ratchet produce? I obtain a single estimate for each parameter combination by propagating the uncertainty. In each of 600 replicates, I randomly draw the DFE and the deleterious input U from their distributions, simulate one modern ($N_{M\text{-uniparental}} = 7000$) and one archaic (equiprobable $N_{A\text{-uniparental}} = 1500, 2250, 3000$) lineage under the drawn values, and record the modern-archaic mean-fitness gap at $T_{mix}$. I draw ($\beta$, $|\bar{S}|$) jointly from the bootstrap distribution of the DFE fit (the two are strongly anticorrelated, $r \approx -0.9$, so independent draws from the marginal intervals would produce a very large number of parameter combinations the data exclude), and U from a triangular prior spanning its 95% CI (mode at the central estimate).

*Y chromosome-specific parameters*

To infer a Y chromosome-relevant DFE, I took inspiration from the approach of ref.18, who rescaled the autosomal DFE to the Y chromosome. Specifically, I use the inference of the autosomal DFE from ref. 13 with shape $\beta = 0.23$ (95% CI [0.19,0.27]) and $\bar{S} = 425$ (95% CI [225,766]), anchored at the autosomal reference effective size $N_{ref} = 10000$ used in that study, giving an unscaled mean $\bar{s} \approx 0.043$ (ref. 13 defines its scaled mean as $N_e s$, not $2N_e s$). Because a single-copy Y gene is hemizygous and unmasked, it could be argued that the same alleles will have stronger effects on the Y chromosome, and hence the comparable quantity may be the homozygous effect $2 \cdot \bar{s} \approx 0.085$. Stronger selection removes deleterious mutations before drift can fix them, so this larger value might be expected to produce smaller fitness gaps. A rerun under the hemizygous conversion confirms this: pooled over the archaic sizes, the expected gap for 500 kb of coding-equivalent sequence falls from 0.63% to 0.47%, so the conversion I use is more conservative from the point of view of inferring the effect of Muller's Ratchet on the Y chromosome.

The Y chromosome point-mutation rate has been estimated at $\sim 3.0\times10^{-8}$/bp/gen (range 2.5-$3.5\times10^{-8}$)[10,11]. The non-recombining, single-copy target that I use for this analysis is the X-degenerate coding sequence[19]. (I exclude the multi-copy 'ampliconic' palindromes from the computation relevant to the ratchet, as intra-chromosomal gene conversion acts like recombination, removing deleterious alleles by copying the more functional one from the other copy.) I use 33,156 bp of codons as the coding class, and treat constrained non-coding sequence as a second class whose target I measure directly and in which every mutation counts. Simulating the 33,156 bp of codons alone gives a mean gap of 0.021%, pooled over the three archaic sizes, and a fixation probability of 6%. Of all possible point mutations in those codons, 0.72 alter an amino acid under the standard genetic code and a 2:1 transition:transversion ratio, so the coding input is $U\_coding \approx 3.0\times10^{-8} \times 33{,}156 \times 0.72 \approx 0.00072$ per chromosome per generation, to which the measured non-coding class adds 0.0024. The corresponding mtDNA fraction in Table S1.1 counts missense changes only, because nonsense mutations there are a separate lethal class.

*Measuring the Y chromosome's constrained target*

In contrast to mtDNA, I was not able to use empirical data to obtain a direct constraint on the Y chromosome fitness gap. For mtDNA, protein sequence is almost 3/4 of the locus, and here I leverage the substantial counts of nonsynonymous to synonymous sites to estimate the DFE. But for the Y chromosome, the coding sequence is on the order of 1% of the total, and at its mutation rate the 33,156 bp of codons accumulate only ~24 substitutions per lineage since the modern-Denisovan Y divergence, of which ~2-4 are nonsynonymous, many fewer than for the mtDNA case so a modern:archaic nonsynonymous ratio has essentially no power.

As an alternative approach to estimating the size of the non-coding class of constrained mutations on the Y chromosome, I performed a direct measurement, studying Zoonomia alignment of large numbers of mammals, which includes chromosome Y sequences. I take the single-copy target to be the 23.8 Mb euchromatic male-specific region less the 7.3 Mb of intra-chromosomal segmental duplications at 95% identity or better, leaving 16.5 Mb. At the phyloP threshold of 2.27 that the Zoonomia analysis calibrates to a 5% false discovery rate, I call 139,380 bases of that region as constrained: 19,073 in codons and 120,307 outside them.

To obtain a sense of the reliability of these estimates, I first asked whether phyloP has power on the Y chromosome, since Y chromosomes are poorly assembled in most of the other mammals and a shallower alignment would leave constrained bases uncalled. Encouragingly Y chromosome codons, which align to a median of 206 other species, are constrained at 58.3%, compared to 58.7% for a matched sample of 400 autosomal genes put through the same pipeline. Less encouragingly, away from the genes, half of single-copy bases align to at most one other species, and 85% of distal bases (more than 10 kb from a gene) are aligned across too little of the mammalian tree for phyloP to call any base constrained. This also lowers the neutral baseline: the Y chromosome's own ancient L2 and MIR repeats are called constrained at 1.06%, but they align more deeply than distal sequence, and at its alignment depth they are called at only 0.23%. Motivated by this, I subtracted a background matched on alignment depth, compartment by compartment, so that my point estimate is that 41,458 of the 120,307 bases in the raw count are due to background noise. The non-coding total is 78,849 bases: 7,898 in untranslated exonic sequence, 9,095 in introns, 3,130 in the 10 kb flanking each gene and 58,726 in distal sequence (95% interval 68-88 kb). A concern with this analysis is that the distal elements may be conserved mainly through the X-linked copy of the sequence: at 55% of their bases at least one species aligns through a contig named as its X chromosome, and at only 1% through one named as its Y (100% and 66% for codons). On the Y itself they show at most weak constraint, with human-chimpanzee divergence 0.91 (95% interval 0.79-1.05) times that of neighboring sequence, against 0.75 for untranslated exonic elements and 0.55 for codons. Since weakly deleterious mutations are the ones that drive the ratchet, I conservatively count them, and I thus simulate 79 kb of constrained non-coding sequence. Constraint in the 85% of distal sequence that phyloP cannot test remains unmeasured; the larger targets in Table S1.3 show what more constrained sequence would imply.

*Checking of Python simulations against a Perl-based implementation*

I confirmed that the DFE framework reproduces the results from the first version of Extended Data Figure 1 in my preprint (there "ratchet.pl")[8]. The original model assumed one selection coefficient *s* for all mutation and no separate lethal class. Collapsing the gamma DFE to a near-delta function at a single s (very large shape parameter), with the nonsense class switched off, I compared results across the complete published grid (6 mutation rates × 9 selection coefficients × 2 archaic sizes = 108 parameter combinations), with 30 replicates per combination. Figure S1.1 shows a scatterplot of the new framework output against the original values for modern lineage fitness, archaic lineage fitness, and their difference. All three panels lie on the y = x line with $R^2 \geq 0.995$.

**Figure S1.1: Validation against previous simulations.** New Python framework vs. original Perl simulation for all 108 grid points in the original Extended Data Figure 1. Blue: $N_{A\text{-uniparental}} = 1500$; orange: $N_{A\text{-uniparental}} = 3000$. Left: modern lineage fitness. Center: archaic lineage fitness. Right: modern - archaic fitness difference. Dashed line: y = x.

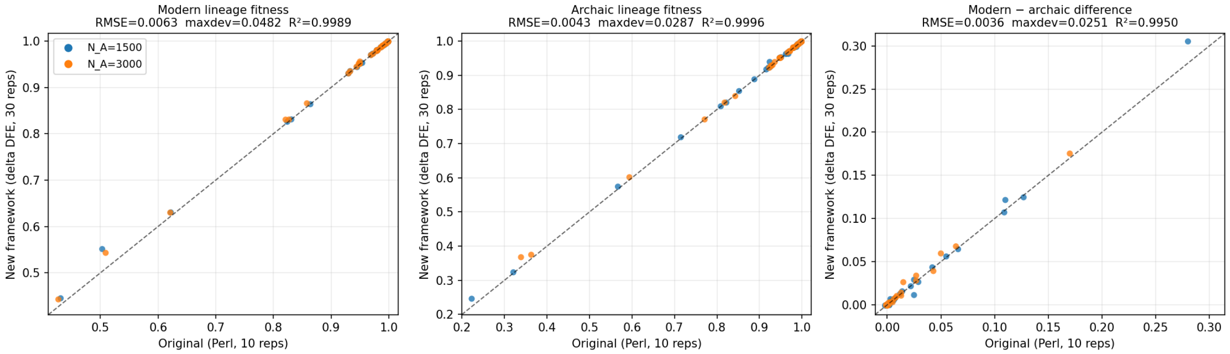


### Results

*Muller's Ratchet is unlikely to have driven modern mtDNA to fixation in Neandertals: I estimate its fixation probability to have been less than 23%.*

Table S1.3 and Figure S1.2 present the fitness gaps that emerges for different simulation settings, and their implication for retention of early modern human lineages during a not-geographically-explicit, 5% admixture from early modern humans into European archaic humans (Appendix C).

**Table S1.3: Mean fitness gap, 95% one-sided upper bound, and fixation probability**

| Genome / analysis | Mean fitness gap | 95% upper bound on fitness gap | Probability of fixation |
|---|---|---|---|
| mtDNA coding only | 0.34% | 0.93% | 20% |
| mtDNA coding only (ABC) | 0.20% | 0.63% | 15% |
| mtDNA with RNA genes | 0.57% | 1.5% | 30% |
| mtDNA with RNA genes (ABC) | 0.39% | 1.1% | 23% |
| Y coding only, 100 kb | 0.07% | 0.29% | 8% |
| Y, codons only (33 kb) | 0.021% | 0.13% | 6% |
| Y, measured target (33 kb + 79 kb) | 0.11-0.18% | 0.50% | 10-13% |
| Y, 500 kb of coding equivalent | 0.63% | 1.5% | 33% |
| Y, 1000 kb of coding equivalent | 1.7% | 3.4% | 64% |

Notes: All rows for both mtDNA and the Y are from the $N_{M\text{-uniparental}} = 7000$ runs, integrating over uncertainty in the parameters and pooling with equal weight over archaic sizes $N_A$-uniparental = 1500/2250/3000; in ABC rows accepted draws are pooled. Probabilities average, over the pooled draws, the probability read from the fixation curve of each draw's post-mixture size ($N_{N\text{-uniparental}}$ = 500/750/1000 for $N_{A\text{-uniparental}}$ = 1500/2250/3000; Petr et al. comparison, 500).

**Figure S1.2: Fitness gap for mtDNA.** The fitness gap is computed at the time of mixture $T_{mix}$. Histograms (gray for primary inference) show the results from simulations. Panels: (a) posterior as a function of deleterious input U; (b) DFE shape β; (c) DFE mean $|\bar{S}|$; (d) archaic size $N_{A\text{-}uniparental}$; (e) model variants (coding-only and RNA-inclusive, with and without ABC conditioning), and the Petr et al. parameters; (f) the gap translated into fixation probability. Percentages in the panel legends give the probability of fixation of modern mtDNA for that scenario, integrating each draw against the fixation curve of its post-mixture size (Petr et al. against 500). Panel f translates the gap posterior into the probability of fixation of modern human mtDNA using the fixation curves derived in Supplementary Information section 3.

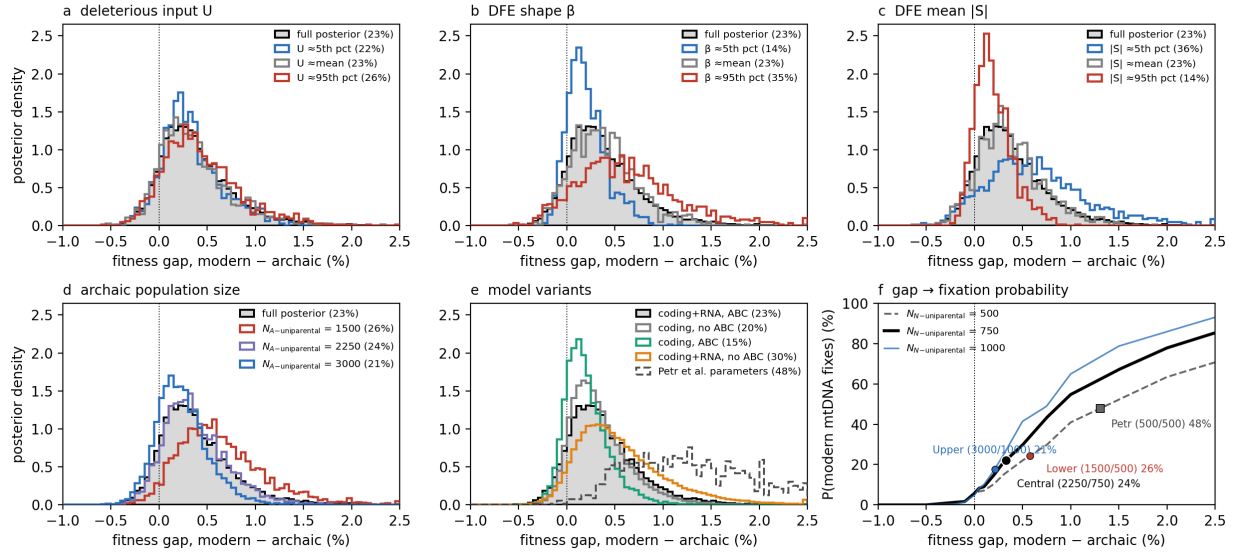


Key observations are:

- For the full posterior distribution, the median is 0.32% (95% of density [-0.2%, +1.3%]). Integrating each posterior draw against the fixation curve of its paired post-mixture size ($N_{N\text{-}uniparental}$ = 500/750/1000 for $N_{A\text{-}uniparental}$ = 1500/2250/3000; the companion analysis rounds the central 725 to 750) gives a ~23% probability that the modern mtDNA fixes, against a ~6% neutral baseline. For all the variant models I considered in panels a-e (excluding the Petr et al.[7] parameters), fixation of modern mtDNA remains unlikely, with probabilities of 15-30%.
- The assumed population size matters little: pairing the $N_{A\text{-}uniparental}$ = 1500 and 3000 bracket posteriors with their correspondingly contracted post-mixture curves of $N_{N\text{-}uniparental}$ = 500 and 1000 gives 26% and 21% probabilities of fixation of modern mtDNA, so fixation probability is essentially independent of the archaic size.
- Fixation probability is larger for the Petr et al. demographic scenario[7], reflecting the much larger (10-fold) population size gap that it assumes for the isolation period during which deleterious mutations are expected to build up[7], compared to the approximately 3-fold harmonic mean supported by demographic inference[6] (Figure S1.2f). Even under the Petr et al. parameters, modern mtDNA fixation remains less likely than not (48%).
- The ABC tightens the inference by conditioning on observed nonsynonymous counts, pulling the 95% upper bound of the RNA-inclusive gap from 1.5% to 1.1%.

In fact, the 23% estimate for the probability of modern mtDNA fixation is conservatively too high.

First, mtDNA frequently participates in DMIs[20], and DMIs have the effect of decreasing minority ancestry[4]. There is evidence of DMIs reducing early modern human ancestry around genes during the interbreeding of early modern humans and archaic humans to form the ancestors of Neandertals[13,3], so there is a reasonable chance that it was involved in some of these DMIs (I carry out a scan for signatures of such DMIs in Supplementary Information section 2 and find no evidence for them, but it would have picked up only a fraction of DMIs known to involve mtDNA).

Second, the literature finds no molecular signal of ratchet-like accumulation of deleterious mtDNA substitutions across divergent primates, birds and fruit flies[21]. Although the population sizes and divergence times in these lineages are different from the archaic/modern human case, the comparisons further suggest that mtDNA is unlikely to have developed a fitness gap sufficient to drive one lineage to fixation.

*Muller's Ratchet is unlikely to have driven the modern Y chromosome to fixation in Neandertals: I estimate its fixation probability to have been less than 10-13%.*

For the Y chromosome, Figure S1.3 and Table S1.3 show how the distribution of fitness gaps varies with the magnitude of the nearly neutral deleterious mutational input to the Y chromosome: small at the measured target, whose deleterious input matches that of ~140 kb of coding-equivalent sequence, and increasing for the larger coding-equivalent targets considered for comparison. Each of 300 replicates per target and archaic size draws ($\beta$, $\bar{S}$, U) from their prior distributions before simulating (Methods), and the three archaic sizes $N_{A\text{-uniparental}}$ = 1500/2250/3000 are pooled with equal weight, exactly as for the mtDNA rows of Table S1.3, so the distributions integrate over demographic and parameter uncertainty as well as evolutionary stochasticity. I also simulated under the Petr et al. parameters[7], which for their 500 kb reference target gives a mean gap of 1.8%, similar to their reported ~2%. This gives a fixation probability of ~55% at its 500 kb reference target, consistent with the ~50% replacement probability they report.

However, Petr et al.[7] made no attempt to biologically constrain the parameters. The measurement of the amount of constrained sequence on the Y chromosome that I present—which suggests that there are only ~112 kb of constrained sequence which is consistent with the fact that evolution disfavors the Y chromosome as a place to maintain functionally important sequence[22]—implies that the fitness gap on the Y chromosome is likely to be far lower than ~1.8%. Assuming 33 kb of constrained coding sequence and 79 kb of constrained non-coding sequence, I estimate that the mean fitness gap is 0.11-0.18% (median 0.09-0.17%) and the fixation probability 10-13% (higher for the weaker DFEs), against the codons-only floor of 6%.

*Muller's Ratchet is unlikely to have driven both modern mtDNA and Y chromosome to fixation in Neandertals: I estimate their joint fixation probability to have been less than 2-3%.*

I finally carried out simulations jointly combining the constraints on mtDNA (23% probability of modern human fixation) and the Y chromosome (10-13% probability of modern human fixation). Together, the probability of fixation of modern human lineages at both loci, in a random sample of 10 Neandertals males, is 2-3%. This is in fact a conservative estimate, because DMIs are expected to pull down the average frequency of modern human lineages at a typical locus

**Figure S1.3: Fitness gap for Y measured at the mixture time $T_{mix}$.** Violin plots show the distribution across 900 replicate simulations (300 per archaic size), each drawing ($\beta$, $\bar{S}$, U) from their priors, of the modern-archaic gap for the measured target (the leftmost violin, the primary inference, pooling the scan over the strength r of the non-coding DFE as well as the archaic sizes) and for coding-equivalent constrained targets from 100 kb to 3 Mb, pooled with equal weight over the archaic sizes $N_{A\text{-uniparental}}$ = 1500/2250/3000; horizontal bars mark medians, dots means. The dashed outline at far right is my simulation under Petr et al. parameters at their 500 kb target, and the dotted line marks their reported ~2% deficit. The annotation above each violin gives the mean gap and the probability of fixation of modern human Y chromosomes, computed from the fixation curves, each drawn at the post-mixture size paired with its archaic size ($N_{N\text{-uniparental}}$ = 500/750/1000; 500 for the Petr et al. set); ≥ marks rungs whose gaps extend beyond the tabulated curves. Priors centered on $\beta = 0.23$ and $\bar{S} = 425$; $N_{M\text{-uniparental}} = 7000$.

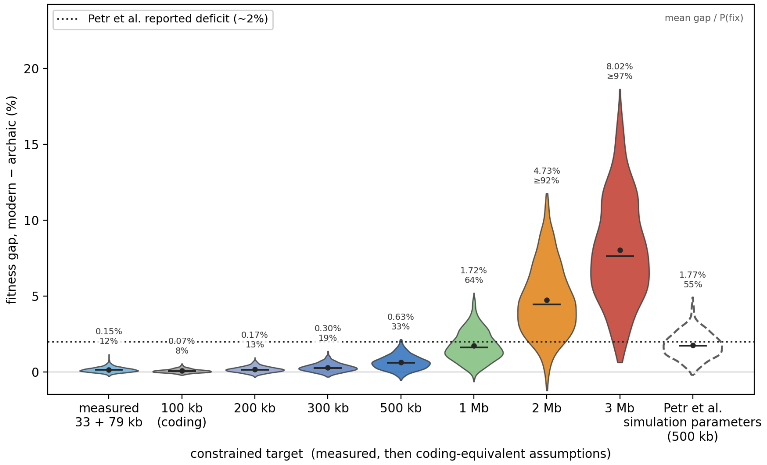


### Appendix A: Inferring mtDNA DFE from polymorphism and divergence

The simulations described in this note draw missense selection coefficients from a gamma DFE fitted to the entire mitochondrial coding-plus-RNA sequence, with shape $\beta = 0.245$ and mean scaled coefficient $\bar{S} = -968$. The present appendix, written largely by AI with editing by the author, sets out the estimator in full: it rederives the Piganeau-Eyre-Walker estimator (ref. 12), verifies that an independent reimplementation reproduces their published human protein-coding values ($\beta = 0.39$, $\bar{S} = -1400$), then applies the identical estimator by class (nonsynonymous, tRNA, rRNA) so that the assumption that the RNA-gene mutations follow the same DFE as the coding genes can be tested. It applies the estimator to the pooled whole-mtDNA class for the main-text simulations.

#### A1. The Poisson Random Field model

Consider a non-recombining, effectively haploid locus (mtDNA) under the infinite-sites assumption, so that each new mutation strikes a previously monomorphic site. Let u be the per-site mutation rate per generation, $N_e$ the effective population size, and s the fitness reduction of a derived allele ($s > 0$ deleterious). In the diffusion limit, the relevant compound parameters are the scaled mutation rate $\theta = 2N_e u$ and scaled selection coefficient

$$S = 2N_e s. \qquad \text{(A1)}$$

Under the Poisson Random Field formulation[23,24], the derived-allele frequencies of the segregating sites in one class form a Poisson process whose intensity (the expected number of sites at which the derived allele segregates at population frequency in $(x, x + dx)$) is

$$f(x; S)\,dx = \theta \cdot g(x; S)\,dx$$
$$g(x; S) = (1 - e^{-S(1-x)}) / [(1 - e^{-S}) \cdot x(1-x)] \tag{A2}$$

Here $g(x; S)$ is proportional to the expected sojourn time that a new semidominant mutation of scaled effect S spends near frequency x before loss or fixation (the Green's function of the Wright-Fisher diffusion), and a deleterious mutation has $S < 0$. In the neutral limit $S \to 0$, $g(x; 0) = 1/x$ (the expected number of neutral sites segregating at frequency x is $\theta/x$).

In a sample of n chromosomes, a site at population frequency x is seen as segregating unless all n draws are ancestral (probability $(1-x)^n$) or all derived (probability $x^n$). Hence a class whose mutational target is L sites contributes

$$E[P] = L\,\theta \int_0^1 g(x; S)\,[1 - x^n - (1-x)^n]\,dx. \tag{A3}$$

For a neutral class this reduces, using $g(x; 0) = 1/x$ and $\int_0^1 (1/x)[1 - x^n - (1-x)^n]\,dx = a_n$, to Watterson's expectation[25]

$$E[P_s] = L_s\,\theta\,a_n$$
$$a_n = \Sigma_{i=1}^{n-1} 1/i. \tag{A4}$$

When a class carries a mixture of effects distributed as Z(S), average (A3) over Z:

$$E[P_n] = L_n\,\theta \int Z(S)\,Q(S, n)\,dS,$$
$$Q(S, n) = \int_0^1 g(x; S)[1 - x^n - (1-x)^n]\,dx \tag{A5}$$

Over a divergence time separating the sample from an outgroup, the fixation probability of a semidominant mutation of scaled effect S, relative to a neutral mutation[26] is

$$F(S) = S / (1 - e^{-S})$$
$$F(0) = 1 \tag{A6}$$

so the expected substitution counts are

$$E[D_s] = L_s\,\lambda,$$
$$E[D_n] = L_n\,\lambda \int Z(S)\,F(S)\,dS$$
$$\lambda = 2u\tau \tag{A7}$$

Z is a gamma distribution on the magnitude of the (deleterious) scaled coefficient[12]

$$Z(S) = \alpha^\beta\,|S|^{\beta-1}\,e^{-\alpha|S|} / \Gamma(\beta)$$
$$\bar{S} = \beta/\alpha \tag{A8}$$

with shape $\beta$ and rate $\alpha$. $\beta < 1$ gives a leptokurtic DFE with mass concentrated near neutrality and in a strongly deleterious tail. $\beta = 1$ is exponential. $\beta > 1$ concentrates mass at intermediate effects. Treating the four counts ($P_n$, $P_s$, $D_n$, $D_s$) as independent Poisson variables with the means above gives the log-likelihood

$$\ln \mathscr{L} = \Sigma_k\,[\,k \ln \hat{E}_k - \hat{E}_k - \ln k!\,]$$
$$k \in \{P_n, P_s, D_n, D_s\} \tag{A9}$$

The four observations determine the four parameters (θ, λ, α, β). For the gamma model there is generically an exact fit, as ref. 12 notes.

**A2. Reduction to two moment equations**

The neutral class fixes the two nuisance parameters, $\hat{\theta} = P_s/(L_s\, a_n)$ and $\hat{\lambda} = D_s/L_s$. Substituting these into (A5) and (A7), and writing the site-count ratio as $\rho_n/\rho_s = L_n/L_s$, leaves two equations in (α, β):

$$I_P(\alpha, \beta) \equiv \int Z(S)\, Q(S, n)\, dS = (P_n/P_s)\cdot(\rho_s/\rho_n)\cdot a_n, \qquad \text{(A10)}$$
$$I_D(\alpha, \beta) \equiv \int Z(S)\, F(S)\, dS = (D_n/D_s)\cdot(\rho_s/\rho_n). \qquad \text{(A11)}$$

The right side of (A11) is the dN/dS ratio. The right side of (A10) is the reduction in nonsynonymous polymorphism relative to the neutral expectation $a_n$. (A10)-(A11) is solved for (α, β) by minimizing the sum of squared log-residuals on a fixed logarithmic grid of S, integrating the gamma against precomputed Q(S, n) and F(S) (pew_replicate.py).

**A3. Validation against published estimates**

Run on the counts from ref. 12 (their Table 1), this estimator qualitatively reproduces the model 2 estimates of ref. 12 across all sixteen usable data sets (Table S1.A1). For the complete human mtDNA data set[27], it returns $\beta = 0.39$ and $\bar{S} = -1.55\times10^3$, which is very similar to their $\beta = 0.39$ and $\bar{S} = -1.4\times10^3$. The shape parameter agrees to within ±0.02 for every data set. The few data sets with very large $|\bar{S}|$ are those in which $\alpha \to 0$ leaves $\bar{S}$ nearly unidentified, as ref. 12 reports. The shape β, which is what matters here, is stably inferred.

**Table S1.A1: Reimplementing ref. 12 model 2 (representative data sets; all 16 agree)**

| Data set | β (this study) | β (ref. 12) | S̄ (this study) | S̄ (ref. 12) |
|---|---|---|---|---|
| Ambystoma (AL) | 1.06 | 1.10 | -99 | -100 |
| Brachyramphus (BM) | 0.90 | 0.91 | -1027 | -990 |
| D. melanogaster (DM) | 0.58 | 0.58 | -1599 | -1700 |
| Human (HS) | 0.39 | 0.39 | -1553 | -1400 |
| Mesomys (MH) | 0.62 | 0.62 | -379 | -390 |
| Phyllobates (PL) | 0.75 | 0.75 | -253 | -250 |
| Sciurus (SA) | 0.85 | 0.85 | -108 | -110 |
| Mus (MD) | 2.23 | 2.20 | -15 | -15 |

**A4. Extension to the mitochondrial RNA genes**

I then applied the identical estimator to a modern human data set, taking synonymous sites as the neutral reference and, in turn, nonsynonymous, tRNA, and rRNA sites as the selected class. The nonsynonymous class is a control that must reproduce ref. 12 for the RNA estimates to be believed. I started by assembling inputs as follows.

- Polymorphism: the number of segregating sites per class in gnomAD v3.1[28], restricted to the African sample with 14,347 called chromosomes. This is the same quantity as $P_n$, $P_s$ above, with synonymous 2776, nonsynonymous 1360, tRNA 342, and rRNA 500.
- Mutational target: because mtDNA has an extreme transition bias (Ti:Tv ≈ 24), I count sites as spectrum-weighted mutational opportunities (each change weighted 2·(Ti:Tv) for a transition, 1 for a transversion), the same convention used for U in the Methods; only the ratio $\rho_s/\rho_n$ enters.

- Divergence: substitutions per class against a high-coverage archaic outgroup (Denisovan mtDNA) from a global alignment to the rCRS, multiple-hit corrected per class by the Jukes-Cantor transform $d^* = -(3/4) \ln(1 - 4d/3)$ (ref. 29); only the ratio $d_n/d_s = I_D$ enters.

At the full African sample size, the control does not reproduce ref. 12: it returns $\beta \approx 0.16$ rather than 0.39. This is likely because the sample is large and, even within one population, far from the mutation-drift-selection equilibrium the model assumes. Recent explosive growth loads it with an excess of rare variants, which the equilibrium model reads as informative about DFE. Because the estimator uses only the total number of segregating sites per class, a class-independent inflation is absorbed into θ and cancels from the ratios in (A10)-(A11). What does not cancel is the interaction between selection and demography, which shifts β. Its size can be read directly by projecting the observed counts down to a smaller sample of m chromosomes: a site carrying i minor alleles in the full sample of n is segregating in a random subsample of m with probability

$$P_{seg}(i; n, m) = 1 - [C(n-i, m) + C(i, m)] / C(n, m). \tag{A12}$$

The expected segregating-site count at depth m is the sum of $P_{seg}$ over sites. Refitting at each m makes β rise monotonically as the sample shrinks, passing through ≈ 0.27 (coding control) at m = 53 (the sample size of ref. 12) and falling to ≈ 0.16 only at the full African sample. β declines with depth for all four classes, so comparisons among classes at a common depth are stable even though the absolute value is depth-dependent.

Table S1.A2 reports the fits at m = 53 (the human sample size of ref. 12), with 95% confidence intervals from a bootstrap that resamples segregating sites within each class and applies Poisson noise to the divergence counts. The nonsynonymous control returns β = 0.27 (95% CI [0.15, 0.40]), broadly consistent with ref. 12's 0.39. The RNA genes return β = 0.18 [0.07, 0.48] (tRNA) and 0.24 [0.08, 0.47] (rRNA), close to the coding value. Most important for the ratchet, the near-neutral mass, the probability that $1 < |S| < 10$, where mutations are simultaneously drift-permitted and costly, is statistically indistinguishable from coding, motivating the decision to treat the DFEs as the same for the main analyses.

Two caveats apply, both shared with ref. 12. First, mtDNA does not recombine, so the sites are not independent and linked selection biases β downward; this is why the focus is on class comparisons at a matched depth rather than on absolute values. Second, the mean coefficient $\bar{S}$ is only imprecisely inferred: calibrating divergence against the unsaturated Denisovan outgroup brings the coding $\bar{S}$ to about -968, close to ref. 12's -1400, but its upper magnitude remains poorly determined. The estimator and its extension are implemented in pew_replicate.py and pew_rna.py.

**Table S1.A2: DFE fits for the African sample at matched downsampled depth m = 53 (divergence vs. the high-coverage Denisovan mtDNA, Jukes-Cantor corrected).**

| Class | β (95% CI) | Near-neutral mass P(1<\|S\|<10) | Near-neutral mass relative to nonsyn |
|---|---|---|---|
| Nonsynonymous only | 0.27 (0.15-0.40) | 0.10 (0.06-0.14) | 1.00 (reference) |
| tRNA | 0.18 (0.07-0.48) | 0.11 (0.04-0.25) | 1.16 (0.35-2.44) |
| rRNA | 0.24 (0.08-0.47) | 0.12 (0.04-0.21) | 1.21 (0.42-2.30) |
| tRNA + rRNA | 0.22 (0.08-0.39) | 0.12 (0.04-0.19) | 1.19 (0.39-2.08) |
| Nonsyn + tRNA + rRNA | 0.24 (0.13-0.35) | 0.11 (0.06-0.15) | 1.06 (0.65-1.78) |

### Appendix B: Combining the coding and RNA contributions in the ABC

The introgression ABC presented in the Methods (Table S1.2) conditions on protein-coding data alone, yet the deleterious mutational input includes the mitochondrial RNA genes. This appendix describes how I combine the two contributions.

**B1. The ABC provides a direct constraint on protein coding, but not RNA genes**
The ABC accepts (modern, Denisovan) lineage pairs whose ratio of carried nonsynonymous substitutions is close to the observed value (modern/Denisova ≈ 1.21; acceptance window [1.16, 1.26]). This is a protein-coding summary statistic, so what it constrains is the coding deleterious load: the combination of missense input rate and coding DFE that reproduces the observed nonsynonymous branch counts. No RNA substitution data enter the acceptance, so the ABC places no direct constraint on the strength of selection acting on the RNA genes. That is supplied separately by the DFE in Appendix A.

**B2. The mutation rate is shared, so the ABC indirectly constrains RNA gene evolution**
The per-generation mutation rate is not a free parameter. It is constrained from each lineage's synonymous substitution count. Because the RNA genes are replicated by the same machinery, they are expected to experience the same mutation rate. The RNA deleterious input enters as U_RNA = U_coding × (U_incl / U_coding - 1), with the ratio U_incl / U_coding = 0.0093 / 0.0064 ≈ 1.45 fixed by the relative mutational targets (main text). Because each accepted draw's U_coding carries the synonymous-calibrated rate, the RNA input scales with it (abc_joint_rna.py). The ratio is derived in the Methods paragraph on the RNA genes (RNA-inclusive U ≈ 0.0093 versus coding-only U_missense ≈ 0.0064). Both inputs are the same per-base rate multiplied by their spectrum-weighted deleterious target sizes, so the mutation rate cancels and the ratio is fixed by the relative target sizes alone. What it holds fixed is the assumption that the RNA genes are constrained according to the whole-mtDNA DFE, the modeling choice tested in Appendix A4.

**B3. Adding the RNA load without contaminating the acceptance**
In each simulated lineage, the coding missense mutations both reduce fitness and are tallied into the nonsynonymous count. The RNA mutations are drawn at rate U_RNA from the same whole-mtDNA gamma DFE (the 'all' fit, $\beta \approx 0.245$, $\bar{S} \approx -968$) and likewise reduce fitness, but they are held in a separate counter and are excluded from the nonsynonymous tally. The recorded modern-Denisova fitness gap at introgression then reflects coding load (ABC-constrained) plus RNA load (rate-constrained, with its selection taken from the DFE), while the acceptance remains a protein-coding comparison, matching exactly the data that inform it. The corresponding coding-only and RNA-inclusive gap posteriors, with their one-sided 95% upper bounds, are reported in Table S1.3.

### Appendix C: Simulations of non-geographically-explicit mixture event

The fixation probabilities quoted in this note are derived from simulations of a single pulse of gene flow into a resident population; Supplementary Information section 3 presents simulations of range expansions. This appendix describes the non-geographically-explicit model of admixture.

**The panmictic model.** The population is simulated as a panmictic population of N diploid individuals in which, at generation zero, a fraction f = 5% of individuals is replaced by modern individuals, with one parameter setting the sex composition of the pulse. Replacing 5% of all individuals with women puts modern mtDNA in 5% of the population but in 10% of females, so the neutral fixation probability for modern mtDNA is 10% in this case.

**The pulse within the roving-band landscape.** The same pulse is also simulated inside the geographically explicit band framework of Supplementary Information section 3, as the comparator for the range expansions reported there. The landscape begins entirely archaic, and at generation zero, 5% of individuals at random are replaced by modern lineage individuals, regardless of their band's position.

**Implementation.** panmictic_sim.py is the migration-pulse engine, with no geographical specification and a single 5% pulse into a constant-size population reproducing under a Wright-Fisher model with separate sexes. It ships in fixation_curves_code/, together with panm3_run.py and the cluster and single-machine entry points that generate fixation_curves_mtdna.csv, the table of fixation probabilities that every probability quoted in this note interpolates.

### Software and Reproducibility

A zipped folder associated with this supplementary note (SI1codebase-v7.zip) includes all the code and the collected simulation outputs necessary to replicate the results, using Python 3.10 (see also requirements.txt and the README file in that folder):

**Inputs.** compute_fn.py: missense fraction from NC_012920.

**The ratchet engine.** ratchet_dfe.py: core forward Wright-Fisher simulation of Muller's Ratchet under a gamma DFE (one combination per call; --anchor-ne sets the DFE reference Ne). ratchet_dfe_annotated.py is a line-by-line commented reading copy.

**DFE and mutation-rate estimates.** save_dfe_draws.py: exports the joint ($\beta$, $|\bar{S}|$) bootstrap draws of the Denisova-calibrated DFE fit (dfe_draws.npz), the prior sampled by the ABC and by the parameter-integrated simulations. pew_replicate.py: reimplementation of the Piganeau & Eyre-Walker (2003) model-2 estimator, validated against their published results (Appendix A, Table S1.A1). pew_rna.py: the same estimator extended to the mitochondrial tRNA and rRNA genes (gnomAD African polymorphism, outgroup divergence), with sample-depth projection and bootstrap (Appendix A, Table S1.A2). mt_dfe_sfs.py: gnomAD chrM VCF reader and mutational-opportunity helpers imported by pew_rna.py and save_dfe_draws.py. rna_constraint.py: divergence-based constraint of the mitochondrial tRNA and rRNA genes against the synonymous neutral clock (rCRS versus Denisovan, Neandertal and chimpanzee mtDNA, spectrum-weighted mutational opportunities; NCBI accessions in data/mtseq/ (included, with NC_012920.gb and the gnomAD chrM VCF)): the source of the measured RNA deleterious fraction of 0.76-0.80 behind the RNA-inclusive $U \approx 0.0093$.

**Validation.** validate_delta.py / make_grid_replicate.py / run_slurm_replicate.sbatch / collect_replicate.py / plot_replicate.py: validation against the original Perl simulation of ref. 8 (Figure S1.1); collected output new_replicate.csv.

**The mtDNA grid and ABC.** make_grid_v22.py / run_task.py / run_slurm.sbatch / merge_chunks.py: the current-demography mtDNA forward grid ($N_M$-uniparental = 7000, $N_A$-

uniparental = 1500/2250/3000, Petr et al. comparison). run_part1_bound.py gives unconstrained (prior-predictive) 95% upper bounds in Table S1.3, integrating over rate and DFE uncertainty. abc_joint_rna.py / run_slurm_joint_rna.sbatch / run_slurm_joint_cod.sbatch / collect_joint_rna.py: the RNA-inclusive ABC of Appendix B and the coding-only companion (collected output: joint_rna_{1500,2250,3000,petr500}.npz and joint_cod_{...}.npz). Coding parameters are constrained by the nonsynonymous acceptance statistic; the synonymous clock constrains the shared mutation rate; RNA deleterious input contributes to fitness but not to acceptance. plot_abc_fig2.py renders Figure S1.2 from those collected posteriors and fixation_curves_mtdna.csv.

**The Y ladder.** make_grid_y_draw.py / run_task_ydraw.py / run_slurm_ydraw.sbatch / collect_y_draw.py: the parameter-integrated Y constrained-target ladder used in Figure S1.3 and Table S1.3 (each of 300 replicates per target and archaic size draws β, $\bar{S}$ and U from their priors). plot_y_fig3_violin.py renders Figure S1.3, pooling the three archaic sizes with equal weight. y_numbers_stated.py derives the Y numbers quoted in the Results and Table S1.3, and the collected outputs are y_stated_summary.csv, y_hemi_summary.csv and y_draw_{stated,hemi}_{reps.npz,summary.csv}. The grid script takes --conv to select the $\bar{S} \rightarrow s$ conversion (stated $\bar{S}$/Nref = 0.0425, the main analysis; hemizygous 2·$\bar{S}$/Nref, the sensitivity). build_ychr_cds_map.py: builds the hg19 X-degenerate codon map (ychr_cds_map.json) that gives the 33,156 bp of complete codons in Table S1.1. sweep_y_U.py: sweeps the whole-Y deleterious input to find the coding-equivalent constrained sequence at which the gap reaches 1%. ratchet_dfe2.py / make_grid_y_twoclass.py / run_task_y2.py / run_slurm_y2.sbatch / collect_y_twoclass.py: the two-class Y analysis, simulating the codons and the constrained non-coding sequence as separate mutation classes with separate DFEs (collected output y_twoclass_summary.csv and y_twoclass_reps.npz). With the non-coding class switched off the engine reproduces simulate_one bit for bit, so those rows sit on the same footing as the ladder above. measure_y_constraint.py / measure_y_singlecopy.py: count constrained bases on the Y from the Zoonomia phyloP scores, by compartment and over the single-copy euchromatin left after removing intra-chromosomal segmental duplications. measure_autosomal_control.py: the same compartments on 400 autosomal genes, the power check. measure_y_neutral_fdr.py: the call rate in ancient L2/MIR repeats on chrY and on the autosomes, the neutral calibration. measure_y_alignment_depth.py / measure_y_depth_followup.py / analyze_y_depth_followup.py: read the Zoonomia alignment itself to measure how much of the mammalian tree is aligned at each base, the neutral call rate at each alignment depth, and human-chimpanzee divergence inside and outside called elements. reconcile_y_target_depth.py combines these into y_target_depth_matched.json, the measured target. make_grid_y_depth.py builds the measured-target grid, run through run_task_y2.py and collected by collect_y_measured.py (y_depth_*); collect_measured_numbers.py assembles every quantity quoted here into y_final_numbers.json.

**Tables.** make_krause_table.py: formats the per-lineage synonymous and nonsynonymous mtDNA substitution counts of Table S1.2 from Krause et al. (2010).

**Fixation curves.** fixation_curves_code/ generates fixation_curves_mtdna.csv, the table of fixation probabilities (as a function of selective advantage and post-mixture size) that all quoted fixation probabilities interpolate. It contains forward Wright-Fisher simulations of a single 5% pulse into a constant-size population, with its own README, the supporting modules panmictic_sim.py and panm3_run.py, and cluster (panm3_submit.py / panm3_collect.py) and single-machine (run_local.py) entry points.
