## Supplementary Information section 1 code base for "Hypothesis: A modern human range expansion ∼300,000 years ago had a critical role in Neandertal origins": abc_joint_rna.py

```
"""
ABC introgression simulation, v4: RNA-inclusive.

This extends the joint single-pair engine (abc_joint.py) to the "weird
combination" the note describes for combining the coding and RNA contributions:

  * The molecular mutation rate is calibrated, exactly as before, from the
    SYNONYMOUS substitution counts on the 13 protein-coding genes (Krause 2010).
    Synonymous sites are neutral, so this pins the per-nucleotide neutral clock of
    the whole mtDNA molecule. Because the RNA genes ride on the same molecule,
    that same rate governs the RNA mutational input -- i.e. the ABC's rate
    constraint DOES propagate to the RNA (Appendix B).

  * The ACCEPTANCE statistic is unchanged: the modern/Denisova ratio of
    nonsynonymous substitutions carried on the two sampled genomes, required to be
    in [1.16, 1.26] (observed ~1.21). This is a protein-coding summary, so the ABC
    constrains only the CODING deleterious load. It sees no RNA data.

  * The RNA deleterious input is added on top of the coding missense input, using
    the SAME whole-mtDNA ('all') DFE and scaling the target by the U ratio
    U_incl / U_coding = 0.0093 / 0.0064 (RNA_FACTOR). RNA mutations reduce fitness
    (they enter L, and hence the recorded gap) but are counted separately (Cr) and
    are DELIBERATELY EXCLUDED from the nonsynonymous count Cm that feeds
    acceptance. The RNA selection strength is NOT fit by the ABC; it is taken from
    the polymorphism+divergence DFE (Appendix A).

DFE: (beta, |Sbar|) drawn JOINTLY from the Denisova matched-depth (m=53) 'all'
bootstrap (dfe_draws.npz), the same joint prior used for the forward upper bound
(denis_bound.py). --fixed uses the central estimate beta=0.245, |Sbar|=968.

Everything else (timeline, size schedule, stochastic coalescence, single-lineage
fitness snapshot at the mixture time) matches abc_joint.py. Acceptance is applied
by combine_joint.py on the resulting joint_{Nanc}.npz.

v22 demography (8_20_2026 note): two scenarios, all sizes haploid locus copies.
  main: NM = 7000, Tsep = 26,224 and Tmix = 8,741 generations ago
        (750 / 250 ka at 28.6 yr/gen; 17,483 generations of isolation);
        Nanc in {1500, 2250, 3000}. DFE anchored at NM: s = Sbar/(2*7000) = 0.069.
  petr: NM = 5000, Nanc = 500, Tsep = 20,979 and Tmix = 10,490 generations ago
        (600 / 300 ka; 10,489 generations of isolation); anchor 5000.
The post-mixture Neandertal phase (NN = 725, sampled at 50 ka) is simulated in
the companion fixation analysis, not here; this engine's job ends at Tmix.
Sampling for the Krause acceptance data is unchanged (modern 0 ka, Denisova 65 ka).
"""
import argparse
import time

import numpy as np

GEN_YEARS = 28.6
G_MODERN_END = round(1_000_000 / GEN_YEARS)  # 34965 ->   0 ka (modern sampled)
G_DENI_END = round(935_000 / GEN_YEARS)      # 32692 ->  65 ka (Denisova sampled)

# gen 0 of the simulation axis = 1000 ka; convert "generations ago" to axis gens
SCENARIOS = {
    #        NM (= DFE anchor)   Tsep gens ago   Tmix gens ago
    "main": dict(NM=7000, tsep=26_224, tmix=8_741),   # 750 -> 250 ka
    "petr": dict(NM=5000, tsep=20_979, tmix=10_490),  # 600 -> 300 ka (Petr et al.)
}

# set from --scenario in main(); module-level so the helpers below see them
N_MODERN = 7000
ANCHOR_NE = 7000.0
G_SPLIT = G_MODERN_END - SCENARIOS["main"]["tsep"]   # 8741  (750 ka)
G_INTRO = G_MODERN_END - SCENARIOS["main"]["tmix"]   # 26224 (250 ka)

# --- observed synonymous counts on each terminal branch since the MRCA (Krause) -
SYN_MODERN = 79.5 + 38.3      # 117.8
SYN_DENI = 108.0
SYN_TOTAL = SYN_MODERN + SYN_DENI    # 225.8 (pooled estimator for one molecular rate)
# opportunity ratios (compute_fn.py, deCODE Ti:Tv = 24.3)
F_SYN, F_MIS, F_NON = 0.362, 0.602, 0.0350
OPP_MIS = F_MIS / F_SYN       # 1.663
OPP_NON = F_NON / F_SYN       # 0.0967

# --- DFE: whole-mtDNA ('all') Denisova matched-depth fit -----------------------
BETA_CENTRAL = 0.245          # 'all' central (v18 main text)
SBAR_CENTRAL = 968.0

# --- coding <-> RNA-inclusive deleterious input -------------------------------
U_CODING, U_INCL = 0.0064, 0.0093
RNA_FACTOR = U_INCL / U_CODING          # 1.453; RNA adds (RNA_FACTOR-1) x coding

BRANCH_END = {"modern": G_MODERN_END, "deni": G_DENI_END}


def load_dfe_draws(path="dfe_draws.npz", cls="all"):
    d = np.load(path)
    if cls not in d:
        raise SystemExit(f"{path} has no '{cls}' draws")
    return np.asarray(d[cls], float)     # (N, 2) columns (beta, |Sbar|)


def schedule_size(arm, N_anc, gen):
    if arm == "modern":
        return N_anc if gen < G_SPLIT else N_MODERN
    return N_anc


def draw_coalescence(N_anc, rng):
    t_extra = rng.exponential(N_anc)
    g_c = G_SPLIT - t_extra
    return int(max(0, round(g_c)))


def draw_dfe(rng, draws, uncertainty):
    if uncertainty:
        beta, Sbar = draws[rng.integers(len(draws))]
        beta, Sbar = float(beta), float(Sbar)
    else:
        beta, Sbar = BETA_CENTRAL, SBAR_CENTRAL
    scale = (Sbar / (2.0 * ANCHOR_NE)) / beta
    return beta, Sbar, scale


def draw_rate(g_c, rng, uncertainty):
    lam = float(rng.gamma(SYN_TOTAL, 1.0)) if uncertainty else SYN_TOTAL
    branch_gens = (G_MODERN_END - g_c) + (G_DENI_END - g_c)
    return lam / branch_gens


def _gamma_load(u, N_cur, beta, scale, rng):
    """Poisson(u) new gamma-DFE hits per individual; return (delta_logfit, counts).
    Selection coefficients are gamma(beta, scale), capped at 1 (lethal)."""
    k = rng.poisson(u, size=N_cur)
    m = int(k.sum())
    if not m:
        return None, k
    s = rng.gamma(beta, scale, size=m)
    np.minimum(s, 1.0, out=s)
    off = np.repeat(np.arange(N_cur), k)
    with np.errstate(divide="ignore"):
        dL = np.bincount(off, weights=np.log1p(-s), minlength=N_cur)
    return dL, k


def simulate_lineage(arm, N_anc, g_c, rng, beta, scale, u_syn, rna):
    """Evolve one lineage from the MRCA (clean) to its sampling time.

    Returns (ns, syn, ns_rna, w_intro) for ONE sampled genome:
      ns      : missense (protein-coding) mutations it carries  -> ACCEPTANCE stat
      syn     : synonymous (neutral) mutations it carries
      ns_rna  : RNA deleterious mutations it carries (diagnostic; NOT in acceptance)
      w_intro : relative fitness at 250 ka of that genome's ancestral lineage
    """
    T = BRANCH_END[arm]
    u_mis = u_syn * OPP_MIS                    # coding missense (deleterious)
    u_non = u_syn * OPP_NON                    # coding nonsense (lethal)
    u_rna = u_mis * (RNA_FACTOR - 1.0) if rna else 0.0   # extra RNA input
    N0 = schedule_size(arm, N_anc, g_c)
    L = np.zeros(N0)
    Cm = np.zeros(N0, dtype=np.int64)         # missense carried (coding)
    Cs = np.zeros(N0, dtype=np.int64)         # synonymous carried (neutral)
    Cr = np.zeros(N0, dtype=np.int64)         # RNA deleterious carried
    L_intro = None
    anc = None

    for gen in range(g_c, T):
        N_cur = schedule_size(arm, N_anc, gen)
        w = np.exp(L - L.max())
        p = w / w.sum()
        parents = rng.choice(L.size, size=N_cur, replace=True, p=p)
        L = L[parents]; Cm = Cm[parents]; Cs = Cs[parents]; Cr = Cr[parents]
        if anc is not None:
            anc = anc[parents]

        # coding missense: reduces fitness AND counts toward the acceptance stat
        dL, k = _gamma_load(u_mis, N_cur, beta, scale, rng)
        if dL is not None:
            L = L + dL; Cm = Cm + k

        # RNA deleterious: SAME DFE, reduces fitness, but NOT counted in Cm
        if u_rna > 0:
            dLr, kr = _gamma_load(u_rna, N_cur, beta, scale, rng)
            if dLr is not None:
                L = L + dLr; Cr = Cr + kr

        ks = rng.poisson(u_syn, size=N_cur)   # neutral synonymous
        if ks.sum():
            Cs = Cs + ks

        if u_non > 0:                          # coding nonsense = lethal
            lethal = rng.poisson(u_non, size=N_cur) > 0
            if lethal.any():
                L[lethal] = -np.inf

        if gen == G_INTRO:
            L_intro = L.copy()
            anc = np.arange(N_cur)

    finite = np.isfinite(L)
    living = np.flatnonzero(finite)
    pick = living[rng.integers(living.size)]
    w_intro = float(np.exp(L_intro[anc[pick]]))
    return int(Cm[pick]), int(Cs[pick]), int(Cr[pick]), w_intro


def run_pair(N_anc, rng, draws, uncertainty, rna):
    g_c = draw_coalescence(N_anc, rng)
    beta, Sbar, scale = draw_dfe(rng, draws, uncertainty)
    u_syn = draw_rate(g_c, rng, uncertainty)
    m_ns, m_syn, m_rna, m_w = simulate_lineage(
        "modern", N_anc, g_c, rng, beta, scale, u_syn, rna)
    d_ns, d_syn, d_rna, d_w = simulate_lineage(
        "deni", N_anc, g_c, rng, beta, scale, u_syn, rna)
    return dict(g_c=g_c, beta=beta, Sbar=Sbar,
                m_ns=m_ns, m_syn=m_syn, m_rna=m_rna, m_w=m_w,
                d_ns=d_ns, d_syn=d_syn, d_rna=d_rna, d_w=d_w)


KEYS = ("g_c", "beta", "Sbar", "m_ns", "m_syn", "m_rna", "m_w",
        "d_ns", "d_syn", "d_rna", "d_w")


def run_task(N_anc, K, seed, draws, uncertainty, rna):
    rng = np.random.default_rng(seed)
    out = {k: np.empty(K) for k in KEYS}
    for i in range(K):
        r = run_pair(N_anc, rng, draws, uncertainty, rna)
        for k in KEYS:
            out[k][i] = r[k]
    return out


def main():
    ap = argparse.ArgumentParser(
        description=__doc__,
        formatter_class=argparse.RawDescriptionHelpFormatter)
    ap.add_argument("--Nanc", type=int, required=True,
                    help="ancestral = archaic effective size "
                         "(main: 1500/2250/3000; petr: 500)")
    ap.add_argument("--scenario", choices=sorted(SCENARIOS), default="main",
                    help="demographic scenario (sets NM, DFE anchor, and the "
                         "split/mixture times); default 'main' = v22 central")
    ap.add_argument("--K", type=int, default=200, help="pairs per task")
    ap.add_argument("--seed", type=int, required=True)
    ap.add_argument("--out", type=str, required=True)
    ap.add_argument("--fixed", action="store_true",
                    help="central DFE + rate instead of drawing from priors")
    ap.add_argument("--coding-only", action="store_true",
                    help="disable the RNA input (reproduces the coding-only ABC "
                         "with the new 'all' DFE)")
    ap.add_argument("--draws", type=str, default="dfe_draws.npz")
    args = ap.parse_args()

    uncertainty = not args.fixed
    rna = not args.coding_only
    draws = load_dfe_draws(args.draws, "all") if uncertainty else None

    scen = SCENARIOS[args.scenario]
    global N_MODERN, ANCHOR_NE, G_SPLIT, G_INTRO
    N_MODERN = scen["NM"]
    ANCHOR_NE = float(scen["NM"])            # DFE anchored at the modern size
    G_SPLIT = G_MODERN_END - scen["tsep"]
    G_INTRO = G_MODERN_END - scen["tmix"]

    t0 = time.time()
    out = run_task(args.Nanc, args.K, args.seed, draws, uncertainty, rna)
    dt = time.time() - t0

    np.savez(args.out, Nanc=args.Nanc, seed=args.seed, K=args.K,
             uncertainty=uncertainty, rna=rna, scen=args.scenario,
             NM=N_MODERN, **out)

    tmrca = 1000.0 - out["g_c"] * GEN_YEARS / 1000.0   # ka
    ratio = out["m_ns"] / np.where(out["d_ns"] == 0, np.nan, out["d_ns"])
    print(f"[{args.scenario} Nanc={args.Nanc} NM={N_MODERN} seed={args.seed}] "
          f"K={args.K} {'priors' if uncertainty else 'fixed'} "
          f"{'RNA-incl' if rna else 'coding-only'} in {dt:.1f}s "
          f"({dt/(2*args.K):.2f}s/lineage)")
    print(f"  TMRCA mean={np.nanmean(tmrca):.0f} ka "
          f"[{np.nanpercentile(tmrca, 2.5):.0f}, {np.nanpercentile(tmrca, 97.5):.0f}]")
    print(f"  modern ns~{out['m_ns'].mean():.1f} syn~{out['m_syn'].mean():.1f} "
          f"rna~{out['m_rna'].mean():.1f} | deni ns~{out['d_ns'].mean():.1f} "
          f"rna~{out['d_rna'].mean():.1f} | ns ratio~{np.nanmean(ratio):.3f}")
    print(f"  wrote {args.out}")


if __name__ == "__main__":
    main()
```
