## Supplementary Information section 1 code base for "Hypothesis: A modern human range expansion ∼300,000 years ago had a critical role in Neandertal origins": build_ychr_cds_map.py

```
"""
Build a hg19 chrY coding-site map for the single-copy X-degenerate Y genes and
validate it by translating each gene's reference CDS.

Output: ychr_cds_map.json  ->  {gene: {"strand": "+/-", "codons": [[[pos,ref],
[pos,ref],[pos,ref]], ...]}}  with 1-based hg19 positions and the hg19 reference
base at every coding site (used downstream to auto-detect the pileup coordinate
convention by reference-base concordance).

Run locally (needs internet to UCSC):
    .venv/bin/python build_ychr_cds_map.py
"""
import gzip
import json
import os
import sys
import urllib.request

REFGENE = "/tmp/refGene_hg19.txt.gz"
REFGENE_URL = "http://hgdownload.soe.ucsc.edu/goldenPath/hg19/database/refGene.txt.gz"

# Single-copy X-degenerate MSY protein-coding genes (Skaletsky et al. 2003).
# Excludes ampliconic (TSPY/RBMY/DAZ/CDY/HSFY/PRY/BPY2/VCY/XKRY...) and
# pseudoautosomal genes (SHOX/CSF2RA/CD99/...), which gene-convert/recombine.
GENES = [
    "SRY", "RPS4Y1", "RPS4Y2", "ZFY", "TBL1Y", "AMELY", "TGIF2LY", "PCDH11Y",
    "TMSB4Y", "NLGN4Y", "USP9Y", "DDX3Y", "UTY", "TXLNGY", "EIF1AY", "KDM5D",
]

CODON = {
    "TTT": "F", "TTC": "F", "TTA": "L", "TTG": "L", "CTT": "L", "CTC": "L",
    "CTA": "L", "CTG": "L", "ATT": "I", "ATC": "I", "ATA": "I", "ATG": "M",
    "GTT": "V", "GTC": "V", "GTA": "V", "GTG": "V", "TCT": "S", "TCC": "S",
    "TCA": "S", "TCG": "S", "CCT": "P", "CCC": "P", "CCA": "P", "CCG": "P",
    "ACT": "T", "ACC": "T", "ACA": "T", "ACG": "T", "GCT": "A", "GCC": "A",
    "GCA": "A", "GCG": "A", "TAT": "Y", "TAC": "Y", "TAA": "*", "TAG": "*",
    "CAT": "H", "CAC": "H", "CAA": "Q", "CAG": "Q", "AAT": "N", "AAC": "N",
    "AAA": "K", "AAG": "K", "GAT": "D", "GAC": "D", "GAA": "E", "GAG": "E",
    "TGT": "C", "TGC": "C", "TGA": "*", "TGG": "W", "CGT": "R", "CGC": "R",
    "CGA": "R", "CGG": "R", "AGT": "S", "AGC": "S", "AGA": "R", "AGG": "R",
    "GGT": "G", "GGC": "G", "GGA": "G", "GGG": "G",
}
COMP = {"A": "T", "C": "G", "G": "C", "T": "A", "N": "N"}


def rc(s):
    return "".join(COMP[b] for b in reversed(s))


def fetch_seq(start1, end1):
    """hg19 chrY [start1,end1] inclusive, 1-based -> uppercase sequence."""
    url = ("https://api.genome.ucsc.edu/getData/sequence?genome=hg19;"
           f"chrom=chrY;start={start1-1};end={end1}")
    with urllib.request.urlopen(url) as r:
        return json.load(r)["dna"].upper()


def main():
    if not os.path.exists(REFGENE):
        urllib.request.urlretrieve(REFGENE_URL, REFGENE)

    # best (longest-CDS) NM_ transcript per gene
    best = {}
    with gzip.open(REFGENE, "rt") as f:
        for line in f:
            c = line.rstrip("\n").split("\t")
            chrom, strand, name, gene = c[2], c[3], c[1], c[12]
            if chrom != "chrY" or gene not in GENES or not name.startswith("NM_"):
                continue
            cds_s, cds_e = int(c[6]), int(c[7])
            ex_s = [int(x) for x in c[9].rstrip(",").split(",")]
            ex_e = [int(x) for x in c[10].rstrip(",").split(",")]
            cds_len = 0
            for s, e in zip(ex_s, ex_e):
                a, b = max(s, cds_s), min(e, cds_e)
                if b > a:
                    cds_len += b - a
            if gene not in best or cds_len > best[gene][0]:
                best[gene] = (cds_len, strand, cds_s, cds_e, ex_s, ex_e)

    out = {}
    summary = []
    for gene in GENES:
        if gene not in best:
            summary.append((gene, "NO NM_ TRANSCRIPT", 0, 0))
            continue
        _, strand, cds_s, cds_e, ex_s, ex_e = best[gene]
        # coding intervals (0-based half-open), genomic order
        iv = []
        for s, e in zip(ex_s, ex_e):
            a, b = max(s, cds_s), min(e, cds_e)
            if b > a:
                iv.append((a, b))
        iv.sort()
        # genomic coding positions, 1-based, with reference base
        pos_ref = []
        for a, b in iv:
            seq = fetch_seq(a + 1, b)          # a+1..b inclusive
            for k, p in enumerate(range(a + 1, b + 1)):
                pos_ref.append((p, seq[k]))
        if strand == "-":
            pos_ref = [(p, COMP[r]) for p, r in reversed(pos_ref)]
        # group into codons in transcription order
        codons = [pos_ref[i:i + 3] for i in range(0, len(pos_ref) - len(pos_ref) % 3, 3)]
        prot = "".join(CODON.get("".join(r for _, r in cod), "X") for cod in codons)
        internal_stops = prot[:-1].count("*")
        ok = prot.startswith("M") and prot.endswith("*") and internal_stops == 0
        summary.append((gene, "OK" if ok else f"check(M={prot[0]},end={prot[-1]},"
                        f"internal*={internal_stops})", len(codons), len(prot)))
        out[gene] = {"strand": strand,
                     "codons": [[[p, r] for p, r in cod] for cod in codons]}

    with open("ychr_cds_map.json", "w") as fh:
        json.dump(out, fh)

    print(f"{'gene':10s} {'status':40s} {'codons':>7s}")
    tot = 0
    for gene, status, ncod, _ in summary:
        print(f"{gene:10s} {status:40s} {ncod:7d}")
        tot += ncod
    print(f"\nTotal codons: {tot}  ({tot*3} coding sites)")
    print("Wrote ychr_cds_map.json")


if __name__ == "__main__":
    sys.exit(main())
```
