## Supplementary Information section 1 code base for "Hypothesis: A modern human range expansion ∼300,000 years ago had a critical role in Neandertal origins": collect_joint_rna.py

```
"""
Merge per-task joint npz from an abc_joint_rna.py array into one file per Nanc.
Defaults merge the RNA-inclusive run (results_joint_rna -> joint_rna_{Nanc}.npz);
use --indir results_joint_cod --out joint_cod for the coding-only companion.
Pairs are independent, so this is a plain concatenation. Run on the cluster after
the array finishes; copy the merged files back and run:
  python combine_joint.py --prefix joint_rna    (or --prefix joint_cod)
"""
import argparse
import glob

import numpy as np

ARR = ("g_c", "beta", "Sbar", "m_ns", "m_syn", "m_rna", "m_w",
       "d_ns", "d_syn", "d_rna", "d_w")
GEN_YEARS = 28.6


def main():
    ap = argparse.ArgumentParser(description=__doc__)
    ap.add_argument("--indir", default="results_joint_rna")
    ap.add_argument("--out", default="joint_rna",
                    help="output prefix: <out>_{Nanc}.npz")
    args = ap.parse_args()

    files = sorted(glob.glob(f"{args.indir}/joint_*.npz"))
    if not files:
        raise SystemExit(f"No {args.indir}/joint_*.npz files found.")
    groups = {}
    for fn in files:
        d = np.load(fn)
        Nanc = int(d["Nanc"])
        scen = str(d["scen"]) if "scen" in d else "main"
        g = groups.setdefault((scen, Nanc), {a: [] for a in ARR})
        for a in ARR:
            g[a].append(np.asarray(d[a], dtype=float))
    for (scen, Nanc), g in sorted(groups.items()):
        merged = {a: np.concatenate(g[a]) for a in ARR}
        stem = f"{Nanc}" if scen == "main" else f"{scen}{Nanc}"
        out = f"{args.out}_{stem}.npz"
        np.savez(out, Nanc=Nanc, scen=scen, **merged)
        tmrca = 1000.0 - merged["g_c"] * GEN_YEARS / 1000.0
        n = merged["m_ns"].size
        print(f"{out}: {n:,} pairs  "
              f"TMRCA {np.mean(tmrca):.0f} ka "
              f"[{np.percentile(tmrca, 2.5):.0f}, {np.percentile(tmrca, 97.5):.0f}]  "
              f"beta {merged['beta'].min():.2f}-{merged['beta'].max():.2f}  "
              f"modern ns~{merged['m_ns'].mean():.1f} deni ns~{merged['d_ns'].mean():.1f} "
              f"rna(m)~{merged['m_rna'].mean():.1f}")


if __name__ == "__main__":
    main()
```
