## Supplementary Information section 1 code base for "Hypothesis: A modern human range expansion ∼300,000 years ago had a critical role in Neandertal origins": collect_replicate.py

```
"""
Aggregate results_replicate/rep_*.tsv (written on the cluster) into one CSV:
new_replicate.csv with columns idx, NA, mu, s, new_modern, new_archaic.

Run on the cluster after the array finishes, then copy new_replicate.csv to the
laptop and run plot_replicate.py there (which compares against the Excel).
"""
import csv
import glob
import os

rows = []
for fn in sorted(glob.glob("results_replicate/rep_*.tsv")):
    with open(fn) as f:
        line = f.readline().strip()
    if not line:
        continue
    idx, NA, mu, s, wm, wa = line.split("\t")
    rows.append((int(idx), int(NA), float(mu), float(s), float(wm), float(wa)))

rows.sort()
with open("new_replicate.csv", "w", newline="") as f:
    w = csv.writer(f)
    w.writerow(["idx", "NA", "mu", "s", "new_modern", "new_archaic"])
    w.writerows(rows)

print(f"Wrote new_replicate.csv ({len(rows)} of 108 grid points).")
missing = sorted(set(range(108)) - {r[0] for r in rows})
if missing:
    print(f"WARNING: missing idx: {missing}")
```
