## Supplementary Information section 1 code base for "Hypothesis: A modern human range expansion ∼300,000 years ago had a critical role in Neandertal origins": collect_y_draw.py

```
"""
Pool the parameter-integrated Y ladder (results_y_draw/task_*.jsonl, from
params_y_draw.tsv) into y_draw_summary.csv and y_draw_reps.npz, keyed the
same way as the fixed-parameter run ('y500_2250', 'y500_petr_500', ...) so
plot_y_fig3_violin.py can render either with --npz.

    python collect_y_draw.py [--outdir results_y_draw] [--params params_y_draw.tsv]
"""
import argparse
import csv
import glob
import json
import os

import numpy as np


def main():
    ap = argparse.ArgumentParser()
    ap.add_argument("--outdir", default="results_y_draw")
    ap.add_argument("--params", default="params_y_draw.tsv")
    ap.add_argument("--out", default="y_draw",
                    help="output prefix: <out>_reps.npz + <out>_summary.csv")
    args = ap.parse_args()

    with open(args.params) as f:
        grid = list(csv.DictReader(f, delimiter="\t"))
    meta = {}
    for r in grid:
        meta.setdefault(int(r["combo"]), r)
    expected = {ci: sum(int(r["reps"]) for r in grid if int(r["combo"]) == ci)
                for ci in meta}

    diffs = {ci: [] for ci in meta}
    n_files = 0
    for path in sorted(glob.glob(os.path.join(args.outdir, "task_*.jsonl"))):
        with open(path) as f:
            data = json.loads(f.readline())
        ci = int(data["summary"]["combo"])
        diffs[ci].extend(r["diff"] for r in data["reps"])
        n_files += 1
    print(f"read {n_files} task files from {args.outdir}")

    npz, rows = {}, []
    for ci in sorted(meta):
        m = meta[ci]
        d = np.array(diffs[ci], dtype=float)
        key = f"{m['tag']}_{m['N_archaic']}"
        npz[key] = d
        status = ("OK" if d.size == expected[ci]
                  else f"INCOMPLETE {d.size}/{expected[ci]}")
        rows.append(dict(
            tag=m["tag"], target_kb=m["target_kb"],
            scen="petr" if m["tag"].endswith("_petr") else "main",
            N_modern=m["N_modern"], N_archaic=m["N_archaic"], T=m["T"],
            n_reps=d.size,
            gap_mean=d.mean() if d.size else float("nan"),
            gap_median=float(np.median(d)) if d.size else float("nan"),
            gap_p95=float(np.percentile(d, 95)) if d.size else float("nan"),
            frac_over_1pct=float(np.mean(d > 0.01)) if d.size else float("nan"),
            status=status))
        if status != "OK":
            print(f"  combo {ci} ({key}): {status}")

    np.savez(f"{args.out}_reps.npz", **npz)
    with open(f"{args.out}_summary.csv", "w", newline="") as f:
        w = csv.DictWriter(f, fieldnames=list(rows[0]))
        w.writeheader(); w.writerows(rows)
    done = sum(1 for r in rows if r["status"] == "OK")
    print(f"wrote {args.out}_reps.npz + {args.out}_summary.csv "
          f"({done}/{len(rows)} combinations complete)")


if __name__ == "__main__":
    main()
```
