## Supplementary Information section 1 code base for "Hypothesis: A modern human range expansion ∼300,000 years ago had a critical role in Neandertal origins": compute_fn.py

```
"""
Compute the de novo nonsynonymous (missense) fraction f_n of single-nucleotide
mutations in the human mtDNA protein-coding genes, under the vertebrate
mitochondrial genetic code (transl_table=2) and a transition-biased mutational
spectrum (deCODE pedigree coding Ti:Tv = 24.3; Sun et al., Cell 2024).

For every position of every complete codon in the 13 protein-coding genes we
enumerate the 3 possible single-nucleotide substitutions, classify each as
synonymous / missense / nonsense (and stop-loss), and weight it by the
mutational spectrum. f_n is then the spectrum-weighted fraction of mutations
that change the amino acid.

We report f under several Ti:Tv ratios so the sensitivity of f_n to the
spectrum is explicit, and we derive a de novo missense input rate U_n.
"""
from collections import defaultdict
from Bio import SeqIO
from Bio.Seq import Seq

GB = "data/NC_012920.gb"
TABLE = 2  # vertebrate mitochondrial code

TRANSITION = {"A": "G", "G": "A", "C": "T", "T": "C"}  # purine<->purine, pyr<->pyr
BASES = ["A", "C", "G", "T"]


def classify(ref_codon, pos, alt):
    """Classify a single-nt change at codon position `pos` (0,1,2) -> alt base."""
    alt_codon = ref_codon[:pos] + alt + ref_codon[pos + 1:]
    ref_aa = str(Seq(ref_codon).translate(table=TABLE))
    alt_aa = str(Seq(alt_codon).translate(table=TABLE))
    if ref_aa == alt_aa:
        return "synonymous"
    if alt_aa == "*":
        return "nonsense"          # gain of stop
    if ref_aa == "*":
        return "stoploss"          # loss of stop (count with nonsynonymous)
    return "missense"


def spectrum_weight(ref_base, alt, titv):
    """Relative per-event weight. Ti:Tv ratio (counts) = titv.
    Each site has 1 transition + 2 transversion opportunities, so per-opportunity
    rates satisfy r_ti / r_tv = 2 * titv."""
    if alt == TRANSITION[ref_base]:
        return 2.0 * titv
    return 1.0


def main():
    rec = SeqIO.read(GB, "genbank")
    cds = [f for f in rec.features if f.type == "CDS"]
    print(f"Parsed {len(cds)} CDS features from {rec.id}\n")

    titv_grid = [("uniform", 0.5), ("Ti:Tv=12", 12.0),
                 ("Ti:Tv=24.3 (deCODE)", 24.3), ("Ti:Tv=40", 40.0)]

    # accumulate spectrum-weighted opportunity by class, for each titv
    results = {label: defaultdict(float) for label, _ in titv_grid}
    pc_bp = 0
    per_gene_bp = {}

    for f in cds:
        gene = f.qualifiers.get("gene", ["?"])[0]
        nt = str(f.extract(rec.seq)).upper()
        ncomplete = len(nt) // 3
        per_gene_bp[gene] = ncomplete * 3
        pc_bp += ncomplete * 3
        for ci in range(ncomplete):
            codon = nt[ci * 3:ci * 3 + 3]
            if any(b not in BASES for b in codon):
                continue
            for pos in range(3):
                ref_base = codon[pos]
                for alt in BASES:
                    if alt == ref_base:
                        continue
                    cls = classify(codon, pos, alt)
                    for label, titv in titv_grid:
                        w = spectrum_weight(ref_base, alt, titv)
                        results[label][cls] += w

    print(f"Protein-coding sequence analysed (complete codons): {pc_bp:,} bp "
          f"across {len(cds)} genes\n")

    print(f"{'spectrum':<22}{'f_syn':>9}{'f_missense':>12}{'f_nonsense':>12}"
          f"{'f_nonsyn*':>11}")
    central_fn = None
    for label, _ in titv_grid:
        d = results[label]
        tot = sum(d.values())
        f_syn = d["synonymous"] / tot
        f_mis = d["missense"] / tot
        f_non = d["nonsense"] / tot
        f_stoploss = d["stoploss"] / tot
        f_nonsyn = f_mis + f_non + f_stoploss   # all amino-acid-altering
        print(f"{label:<22}{f_syn:>9.3f}{f_mis:>12.3f}{f_non:>12.4f}"
              f"{f_nonsyn:>11.3f}")
        if "deCODE" in label:
            central_fn = (f_mis, f_nonsyn)

    f_mis, f_nonsyn = central_fn
    print("\n* f_nonsyn = missense + nonsense + stop-loss (all amino-acid-altering)\n")

    # ---- derive de novo missense input rate U_n ------------------------------
    L_pc = pc_bp
    rate_all_coding = 2.87e-6   # deCODE whole non-control region, /bp/gen
    rate_syn = 6.36e-6          # deCODE synonymous (near-neutral) /bp/gen
    print("De novo missense input rate U_n = f_missense x mu_neutral x L_pc")
    print(f"  L_pc = {L_pc:,} bp; f_missense(deCODE Ti:Tv) = {f_mis:.3f}")
    for name, mu in [("all-coding rate 2.87e-6 (selection-depressed -> lower)",
                      rate_all_coding),
                     ("synonymous rate 6.36e-6 (neutral -> upper)", rate_syn)]:
        print(f"  mu = {mu:.2e} [{name}]:  U_n = {f_mis * mu * L_pc:.4f} /genome/gen")


if __name__ == "__main__":
    main()
```
