## Supplementary Information section 1 code base for "Hypothesis: A modern human range expansion ∼300,000 years ago had a critical role in Neandertal origins": panm3_collect.py

```
#!/usr/bin/env python3
"""
panm3_collect.py -- write fixation_curves_mtdna.csv, a machine-readable mtDNA fixation curve
for the stable-boundary (panmictic) model.

Statistic: the probability that ALL `NSAMPLE` (default 10) individuals sampled from the final
population carry the modern mtDNA. Estimated as the mean of p**NSAMPLE over replicates, where
p is the modern mtDNA frequency at the sampling generation. Because trajectories are
essentially fixed or lost by then, this is numerically close to the fixation probability and
is also the expected final frequency.

    python panm3_collect.py            # -> fixation_curves_mtdna.csv
    NSAMPLE=3 python panm3_collect.py
"""
import os
import glob
import datetime
import numpy as np
import panmictic_sim as PS
from panm3_run import combos, PULSE

NSAMPLE = int(os.environ.get("NSAMPLE", "10"))
COL = dict(seed=0, mt=1, Y=2, auto=3, X=4)
GEN_YEARS = 28.6                      # as used in the Muller's Ratchet note
SAMPLE_GEN = PS.Config().gens         # generation at which the population is scored


def pool(s_, N):
    rows = []
    for f in glob.glob(f"out_panm3/out_{s_:g}_{N}_s*.txt"):
        for line in open(f).read().strip().split("\n")[1:]:
            p = line.split("\t")
            if len(p) >= 5:
                rows.append([float(x) for x in p])
    return np.array(rows) if rows else None


def main():
    out = "fixation_curves_mtdna.csv"
    with open(out, "w") as fh:
        fh.write(
f"""# fixation_curves_mtdna.csv
# Generated by panm3_collect.py (from panm3_run.py) on {datetime.date.today().isoformat()}.
#
# MODEL: stable-boundary / panmictic. One instantaneous, sex-unbiased pulse in which a
#   fraction {PULSE:g} of the population is of modern origin, into a constant-size randomly
#   mating archaic population; then {SAMPLE_GEN:,} generations of Wright-Fisher reproduction
#   with separate sexes. At {GEN_YEARS} years per generation that is
#   ~{SAMPLE_GEN*GEN_YEARS/1000:.0f} thousand years.
# LOCUS: mtDNA (maternally transmitted, haploid, non-recombining). Because the pulse is
#   sex-unbiased the modern mtDNA starts at frequency {PULSE:g}.
# SELECTION: additive, acting only on mtDNA. s > 0 favours the modern mtDNA; s < 0 favours the
#   archaic mtDNA with strength |s| (NOT clipped to neutrality).
# SCORED: at generation {SAMPLE_GEN:,} (no early stopping).
# STATISTIC pfix_pct: 100 x mean over replicates of p**{NSAMPLE}, where p is the modern mtDNA
#   frequency at that generation -- i.e. the probability that all {NSAMPLE} sampled individuals
#   carry the modern type. se_pct is the standard error of that mean across replicates.
# SIZES: census_N is the number of diploid individuals; locus_Ne = census_N / 2 is the number
#   of mtDNA copies, since only females transmit. locus_Ne is the quantity denoted N_A in the
#   Muller's Ratchet note.
#
census_N,locus_Ne,s,pfix_pct,n_replicates,se_pct
""")
        print(f"{'census_N':>9} {'locus_Ne':>9} {'s':>8} {'pfix_pct':>9} {'n':>6} {'se_pct':>7}")
        for N, s_ in sorted(((N, s_) for s_, N in combos()), key=lambda t: (t[0], t[1])):
                a = pool(s_, N)
                if a is None:
                    continue
                p = a[:, COL["mt"]] ** NSAMPLE
                m, se, n = 100 * p.mean(), 100 * p.std(ddof=1) / np.sqrt(len(p)), len(p)
                fh.write(f"{N},{N//2},{s_:g},{m:.3f},{n},{se:.3f}\n")
                print(f"{N:>9} {N//2:>9} {s_:>8g} {m:>9.2f} {n:>6} {se:>7.2f}")
    print(f"\nwrote {out}")


if __name__ == "__main__":
    main()
```
