## Supplementary Information section 1 code base for "Hypothesis: A modern human range expansion ∼300,000 years ago had a critical role in Neandertal origins": panm3_run.py

```
#!/usr/bin/env python3
"""
panm3_run.py -- dense mtDNA fixation curve for the stable-boundary (panmictic) model, on the
s grid where an ABC posterior on the modern-archaic fitness gap actually has mass.

One task = one (s, census N) x `reps` replicates.  Row: seed  mt  Y  auto  X
Usage: python panm3_run.py <s> <N> <reps> <seed0> <outfile>
"""
import os
import sys
from dataclasses import replace
import panmictic_sim as PS

# s < 0 means the ARCHAIC mtDNA is the fitter type; the engine handles this explicitly rather
# than clipping to neutrality, because the posterior being fed in puts mass there.
SVALS = [-0.005, -0.002, -0.001, 0.0, 0.0005, 0.001, 0.002, 0.003,
         0.005, 0.0075, 0.01, 0.015, 0.02, 0.025, 0.03]
# Extension requested for the Y-chromosome ladder: its 1-3 Mb rungs have mean fitness gaps of
# 1.9-8.4%, past the top of SVALS, so fixation probabilities there could only be quoted as
# lower bounds. These are run only at the two sizes where the sweep actually happens.
HIGH_S = [0.04, 0.05, 0.075, 0.10]
HIGH_S_NVALS = [1000, 1500]
# census diploid individuals; a uniparental locus has N/2 copies, so these give the
# mtDNA population sizes N_A = 1500 and 3000 used in the Muller's Ratchet note.
# census diploid individuals; the mtDNA population is half this.
#   1000 -> haploid  500 : Petr et al.'s Neandertal Ne (1000 diploid), for comparison
#   2000 -> haploid 1000 : the sweep size for the NA = 3000 bracket, which contracts
#                          by the same ~1/3 ratio as the central case
#   1500 -> haploid 750  : the SWEEP size. Peyregne Fig. 4A gives a time-weighted harmonic
#                          mean of 1,447 diploid (723 haploid) for the Neandertal lineage over
#                          250-50 ka, i.e. after admixture. This is the primary case.
#   3000 -> haploid 1500 : the lower ACCUMULATION-phase size (750-250 ka), kept as a bound.
#   6000 -> haploid 3000 : the upper accumulation-phase size, kept as a bound.
NVALS = [1000, 1500, 2000, 3000, 6000]
PULSE = 0.05          # single instantaneous pulse of modern ancestry, unbiased by sex


def combos():
    """(s, census_N) pairs to simulate: a full grid over SVALS, plus the high-s extension
    at the two sizes where the post-mixture sweep occurs. Deliberately not a cross-product."""
    out = [(s_, N) for s_ in SVALS for N in NVALS]
    out += [(s_, N) for s_ in HIGH_S for N in HIGH_S_NVALS]
    return out


def main():
    s_ = float(sys.argv[1]); N = int(sys.argv[2])
    reps, seed0, outfile = int(sys.argv[3]), int(sys.argv[4]), sys.argv[5]
    os.makedirs(os.path.dirname(outfile) or ".", exist_ok=True)
    cfg0 = PS.Config(N=N, f=PULSE, male_frac=0.5,
                     s=s_, sel_locus=("none" if s_ == 0 else "mt"))
    with open(outfile, "w") as f:
        f.write("seed\tmt\tY\tauto\tX\n")
        for seed in range(seed0, seed0 + reps):
            r = PS.run(replace(cfg0, seed=seed))
            f.write(f"{seed}\t{r['mt_mod']:.6g}\t{r['Y_mod']:.6g}\t"
                    f"{r['auto_mod']:.6g}\t{r['X_mod']:.6g}\n")
    print("wrote", outfile)


if __name__ == "__main__":
    main()
```
