## Supplementary Information section 1 code base for "Hypothesis: A modern human range expansion ∼300,000 years ago had a critical role in Neandertal origins": panmictic_sim.py

```
"""
panmictic_sim.py -- the NON-SPATIAL alternative to spatial_sim2.py.

Here the gene exchange between early modern humans and Neandertal ancestors happens at a
stable geographic boundary rather than through a range expansion, so there is no need to model
geography. The exchange is a single INSTANTANEOUS PULSE into one randomly mating (panmictic)
Neandertal population, which then evolves for `gens` generations.

Model
  * One population of N diploid individuals, N/2 males and N/2 females, constant through time.
  * At generation 0 a fraction `f` of the individuals are early modern humans and the rest are
    archaic. `male_frac` sets what fraction of the modern contribution is male, so the pulse
    can be unbiased (0.5), entirely male (1.0), or entirely female (0.0). Note the resulting
    starting frequencies:
        autosomes        f            (regardless of the sex bias)
        Y (among males)  2*f*male_frac
        mtDNA            2*f*(1-male_frac)
    so an all-male pulse starts the modern Y at 2f and the modern mtDNA at 0, and vice versa.
  * Each generation is Wright-Fisher with separate sexes: every offspring draws a father
    uniformly from the males and a mother uniformly from the females.
  * Loci: mtDNA (maternal), Y (paternal, males only), 8 unlinked autosomes (diploid), 2 X loci
    (mother transmits one of her two; father transmits his X to daughters only) -- the same
    complement as the spatial engine, so results are directly comparable.
  * Selection is viability selection with population regulation back to N (candidates are
    oversampled and non-survivors discarded). This differs from the spatial engine, where a
    removed offspring simply lowers local density; in a single panmictic population of fixed
    size the standard treatment is to regulate back to N.

Contrast with the range-expansion model
  Under a pulse the autosomal modern proportion starts at f and stays there in expectation --
  it is set by the pulse, not built up by repeated local introgression. The fixation
  probability of a neutral uniparental marker equals its starting frequency, so with an
  unbiased 5% pulse each of the Y and mtDNA fixes about 5% of the time and both fix about
  0.25% of the time. That is the baseline this model exists to quantify.
"""

import numpy as np
from dataclasses import dataclass, replace

AUTOS = 8
XLOCI = 2


@dataclass
class Config:
    N: int = 1000              # diploid individuals (Petr et al. use Ne ~1000 for Neandertals)
    f: float = 0.05            # modern human fraction in the instantaneous pulse
    male_frac: float = 0.5     # fraction of the modern contribution that is male
    # --- additive selection against the archaic allele ---
    s: float = 0.0         # >0 favours the modern allele, <0 favours the archaic one
    sel_locus: str = "none"    # "none", "Y", "mt", "auto", "X"
    auto_locus: int = 7
    x_locus: int = 1
    # --- epistatic (Dobzhansky-Muller) selection, same semantics as spatial_sim2 ---
    dm: float = 0.0
    dm_focal: str = "mt"
    dm_focal_locus: int = 7
    dm_partner: str = "Y"
    dm_partner_locus: int = 7
    dm_dom: str = "dominant"
    dm_male_limited: bool = True
    # F1 male sterility: sons produced in the FIRST generation after the pulse by a modern
    # mother and an archaic father leave no offspring. With an all-female pulse every son of
    # a modern mother is such an F1, and is identified by his modern mtDNA.
    f1_male_sterile: bool = False
    gens: int = 9091
    oversample: float = 3.0    # candidate offspring per slot when selection is acting
    seed: int = 0


# ---------------------------------------------------------------- initialization
def _init(cfg, rng):
    N = cfg.N
    nm = N // 2
    nf = N - nm
    n_mod_m = int(round(cfg.f * N * cfg.male_frac))
    n_mod_f = int(round(cfg.f * N * (1.0 - cfg.male_frac)))
    if n_mod_m > nm or n_mod_f > nf:
        raise ValueError("modern pulse exceeds the number of individuals of that sex")
    sex = np.concatenate([np.ones(nm, np.int8), np.zeros(nf, np.int8)])
    mod = np.zeros(N, bool)
    mod[:n_mod_m] = True                       # modern males occupy the first slots
    mod[nm:nm + n_mod_f] = True                # modern females the first female slots
    v = mod.astype(np.int8)
    P = dict(
        sex=sex,
        mt=v.copy(),
        Y=np.where(sex == 1, v, -1).astype(np.int8),
        auto=np.repeat((2 * v)[:, None], AUTOS, axis=1).astype(np.int8),
        X=np.repeat((np.where(sex == 0, 2, 1) * v)[:, None], XLOCI, axis=1).astype(np.int8),
    )
    return P


# ---------------------------------------------------------------- selection
def _dm_penetrance(cfg, sx, mt, Yc, auto, X):
    n = sx.shape[0]

    def focal_modern():
        f = cfg.dm_focal
        if f == "mt":   return (mt == 1)
        if f == "Y":    return (sx == 1) & (Yc == 1)
        if f == "auto": return auto[:, cfg.dm_focal_locus] >= 1
        if f == "X":    return X[:, cfg.dm_focal_locus] >= 1
        raise ValueError(cfg.dm_focal)

    def partner_archaic():
        p, dom = cfg.dm_partner, cfg.dm_dom
        if p == "mt":   return (mt == 0).astype(float)
        if p == "Y":    return ((sx == 1) & (Yc == 0)).astype(float)
        if p == "auto":
            k = 2 - auto[:, cfg.dm_partner_locus]; maxd = np.full(n, 2)
        elif p == "X":
            maxd = np.where(sx == 0, 2, 1); k = maxd - X[:, cfg.dm_partner_locus]
        else:
            raise ValueError(p)
        if dom == "dominant":  return (k >= 1).astype(float)
        if dom == "recessive": return (k == maxd).astype(float)
        return k / maxd

    p = cfg.dm * focal_modern().astype(float) * partner_archaic()
    if cfg.dm_male_limited:
        p = p * (sx == 1)
    return np.clip(p, 0.0, 1.0)


def _disfavoured_count(cfg, sx, mt, Yc, auto, X):
    """Copies of the DISFAVOURED allele carried by each candidate.

    cfg.s > 0 selects against the ARCHAIC allele (the modern one is favoured); cfg.s < 0
    reverses this, selecting against the MODERN allele with strength |s|. Allowing negative s
    matters when a posterior on the fitness gap puts mass on the archaic type being the fitter
    of the two, which must not be silently treated as neutral.
    """
    n = _archaic_count(cfg, sx, mt, Yc, auto, X)
    if cfg.s >= 0:
        return n
    if cfg.sel_locus == "Y":    return np.where(sx == 1, 1 - n, 0)   # males only
    if cfg.sel_locus == "mt":   return 1 - n
    if cfg.sel_locus == "auto": return 2 - n
    if cfg.sel_locus == "X":    return np.where(sx == 0, 2, 1) - n
    return n


def _archaic_count(cfg, sx, mt, Yc, auto, X):
    if cfg.sel_locus == "Y":
        return np.where((sx == 1) & (Yc == 0), 1, 0)
    if cfg.sel_locus == "mt":
        return (mt == 0).astype(np.int64)
    if cfg.sel_locus == "auto":
        return 2 - auto[:, cfg.auto_locus]
    if cfg.sel_locus == "X":
        return np.where(sx == 0, 2, 1) - X[:, cfg.x_locus]
    return np.zeros(sx.shape, np.int64)


# ---------------------------------------------------------------- one generation
def _breed(P, cfg, rng, n_needed, sex_val, gen=1):
    """Draw `n_needed` surviving offspring of sex `sex_val`."""
    males = np.nonzero(P["sex"] == 1)[0]
    females = np.nonzero(P["sex"] == 0)[0]
    if males.size == 0 or females.size == 0:
        return None
    selecting = ((cfg.s != 0 and cfg.sel_locus != "none") or cfg.dm > 0
                 or (cfg.f1_male_sterile and gen == 0 and sex_val == 1))
    out = {k: [] for k in ("mt", "Y", "auto", "X")}
    got = 0
    tries = 0
    drawn = 0          # adaptive oversampling: track the realised acceptance rate so that
    kept_tot = 0       # strong selection (e.g. s=1, where nearly every male is culled) does
    while got < n_needed and tries < 200:   # not require hundreds of undersized batches
        tries += 1
        if selecting:
            if drawn > 0 and kept_tot > 0:      # use the realised acceptance rate once known
                factor = min(max(1.1 / max(kept_tot / drawn, 1e-3), 1.05), 500.0)
            else:                                # first batch: fall back to the configured guess
                factor = cfg.oversample
        else:
            factor = 1.0
        B = int(np.ceil((n_needed - got) * factor))
        fa = males[rng.integers(0, males.size, B)]
        mo = females[rng.integers(0, females.size, B)]
        mt = P["mt"][mo].astype(np.int8)
        Yc = (P["Y"][fa] if sex_val == 1 else np.full(B, -1)).astype(np.int8)
        auto = ((rng.random((B, AUTOS)) < P["auto"][fa] / 2.0).astype(np.int8)
                + (rng.random((B, AUTOS)) < P["auto"][mo] / 2.0).astype(np.int8))
        mo_tx = (rng.random((B, XLOCI)) < P["X"][mo] / 2.0).astype(np.int8)
        X = (mo_tx + P["X"][fa]).astype(np.int8) if sex_val == 0 else mo_tx.astype(np.int8)
        sx = np.full(B, sex_val, np.int8)
        if selecting:
            w = np.ones(B)
            if cfg.s != 0 and cfg.sel_locus != "none":
                w *= np.clip(1.0 - abs(cfg.s)
                             * _disfavoured_count(cfg, sx, mt, Yc, auto, X), 0.0, 1.0)
            if cfg.dm > 0:
                w *= (1.0 - _dm_penetrance(cfg, sx, mt, Yc, auto, X))
            if cfg.f1_male_sterile and gen == 0 and sex_val == 1:
                w = w * (mt != 1)      # F1 sons of modern mothers do not reproduce
            keep = np.nonzero(rng.random(B) < w)[0]
        else:
            keep = np.arange(B)
        drawn += B; kept_tot += keep.size
        if keep.size:
            take = keep[:n_needed - got]
            out["mt"].append(mt[take]); out["Y"].append(Yc[take])
            out["auto"].append(auto[take]); out["X"].append(X[take])
            got += take.size
    if got < n_needed:
        return None
    return {k: (np.concatenate(v, 0) if v[0].ndim > 1 else np.concatenate(v))
            for k, v in out.items()}


def _step(P, cfg, rng, gen=1):
    nm = cfg.N // 2
    nf = cfg.N - nm
    sons = _breed(P, cfg, rng, nm, 1, gen)
    if sons is None: return None
    daughters = _breed(P, cfg, rng, nf, 0, gen)
    if daughters is None: return None
    return dict(
        sex=np.concatenate([np.ones(nm, np.int8), np.zeros(nf, np.int8)]),
        mt=np.concatenate([sons["mt"], daughters["mt"]]),
        Y=np.concatenate([sons["Y"], daughters["Y"]]),
        auto=np.concatenate([sons["auto"], daughters["auto"]], 0),
        X=np.concatenate([sons["X"], daughters["X"]], 0),
    )


# ---------------------------------------------------------------- stats & driver
def _stats(P, exclude=(), exclude_x=()):
    male = P["sex"] == 1
    xcols = [c for c in range(XLOCI) if c not in exclude_x]
    acols = [c for c in range(AUTOS) if c not in exclude]

    def xfrac(mask, diploid):
        if not mask.any() or not xcols:
            return np.nan
        cap = 2.0 if diploid else 1.0
        return float((P["X"][mask][:, xcols].sum(1) / (len(xcols) * cap)).mean())

    return dict(
        mt_mod=float((P["mt"] == 1).mean()),
        Y_mod=float((P["Y"][male] == 1).mean()) if male.any() else np.nan,
        auto_mod=float(P["auto"][:, acols].mean() / 2.0),
        X_mod=float(np.nanmean([v for v in (xfrac(~male, True), xfrac(male, False)) if v == v])),
        pop=int(P["sex"].size))


def run(cfg):
    rng = np.random.default_rng(cfg.seed)
    excl = set(); excl_x = set()
    if cfg.sel_locus == "auto": excl.add(cfg.auto_locus)
    if cfg.sel_locus == "X": excl_x.add(cfg.x_locus)
    if cfg.dm > 0 and cfg.dm_focal == "auto": excl.add(cfg.dm_focal_locus)
    if cfg.dm > 0 and cfg.dm_partner == "auto": excl.add(cfg.dm_partner_locus)
    if cfg.dm > 0 and cfg.dm_focal == "X": excl_x.add(cfg.dm_focal_locus)
    if cfg.dm > 0 and cfg.dm_partner == "X": excl_x.add(cfg.dm_partner_locus)
    P = _init(cfg, rng)
    for g in range(cfg.gens):
        nxt = _step(P, cfg, rng, g)
        if nxt is None:
            break
        P = nxt
    return _stats(P, tuple(excl), tuple(excl_x))


if __name__ == "__main__":
    import sys
    n = int(sys.argv[1]) if len(sys.argv) > 1 else 20
    for tag, kw in [("unbiased 5% pulse", {}),
                    ("all-male 5% pulse", dict(male_frac=1.0)),
                    ("all-female 5% pulse", dict(male_frac=0.0))]:
        res = [run(replace(Config(**kw), seed=s)) for s in range(n)]
        mt = np.array([r["mt_mod"] for r in res]); Y = np.array([r["Y_mod"] for r in res])
        au = np.array([r["auto_mod"] for r in res])
        print(f"{tag:22s} n={n}  auto={100*au.mean():5.2f}%  "
              f"P(Y fixed)={100*(Y >= 1).mean():5.1f}%  P(mt fixed)={100*(mt >= 1).mean():5.1f}%")
```
