## Supplementary Information section 1 code base for "Hypothesis: A modern human range expansion ∼300,000 years ago had a critical role in Neandertal origins": README.md

# Fixation probability of introgressed mtDNA at a stable geographic boundary

Code to reproduce `fixation_curves_mtdna.csv`, the table giving the probability that an
introgressed modern human mtDNA replaces the resident archaic mtDNA, as a function of its
selective advantage *s* and of the mtDNA effective population size.

This table is the second half of the argument in the Muller's Ratchet supplementary note. That
note estimates the *fitness gap* the ratchet can generate between the modern and archaic mtDNA
lineages; this code converts a gap into the quantity actually observed — whether the modern
type took over. The two are combined by pushing the ABC posterior on the gap through these
curves.

## The model

A single instantaneous pulse of early modern human ancestry enters an archaic population at a
stable geographic boundary, and the population then evolves forward to the time the Neandertals
in the fossil record were sampled. There is no spatial structure: this is the deliberate
contrast to a range expansion, isolating what selection alone can do in a well-mixed population.

- One population of `census_N` diploid individuals, half male and half female, constant in size.
- At generation 0 a fraction 0.05 of individuals are of modern origin. The pulse is **unbiased
  by sex**, so the modern mtDNA starts at frequency 0.05 among the transmitting females and
  autosomal ancestry is 5% by construction — no calibration is needed.
- Wright-Fisher reproduction with separate sexes for 9,091 generations (~260 kyr at 28.6
  years/generation). Loci tracked: mtDNA, Y, 8 unlinked autosomes and 2 X loci.
- Additive viability selection on mtDNA only. **`s > 0` favours the modern mtDNA; `s < 0`
  favours the archaic one with strength |s|.** Negative values are simulated explicitly rather
  than clipped to neutrality, because the posterior being fed through these curves places real
  mass on the archaic type being the fitter of the two.

**Population size convention.** `census_N` is the number of diploid individuals. Because only
females transmit mtDNA, the effective number of mtDNA copies is `locus_Ne = census_N / 2`.
`locus_Ne` is the quantity denoted N_A / N_N in the note, and **is the column to select on**.

| `locus_Ne` | `census_N` | meaning |
|---|---|---|
| 500 | 1,000 | post-mixture size for the N_A = 1500 bracket; also matches Petr et al.'s Neandertal N_e |
| 750 | 1,500 | post-mixture size for the central N_A = 2250 case |
| 1,000 | 2,000 | post-mixture size for the N_A = 3000 bracket |
| 1,500 | 3,000 | N_A = 1500 with *no* post-mixture contraction (sensitivity only) |
| 3,000 | 6,000 | N_A = 3000 with *no* post-mixture contraction (sensitivity only) |

## The statistic

`pfix_pct` is 100 x the mean over replicates of *p*^10, where *p* is the modern mtDNA frequency
at the scoring generation — the probability that all 10 sampled Neandertals carry the modern
type. Ten is roughly the number of archaic individuals with usable mtDNA. This is reported
rather than literal fixation because it is what the data can speak to; in practice the two are
numerically identical here (trajectories are at 0 or 1 by the scoring generation, verified to
within 0.0 percentage points on every row).

## Files

- `panmictic_sim.py` — the simulation engine. `Config` holds every parameter; `run()` returns
  the modern fraction at each locus at the sampling generation.
- `panm3_run.py` — worker: one (s, census_N) combination x `reps` replicates. Defines the
  s grid, the population sizes, and `combos()`, the list of (s, N) pairs simulated. Note this
  is deliberately **not** a full cross-product: the high-s extension (0.04-0.10) is run only at
  the two sizes where the post-mixture sweep occurs.
- `panm3_collect.py` — pools the per-task output and writes `fixation_curves_mtdna.csv`,
  including the header comments recording model settings and provenance.
- `panm3_submit.py` — builds a SLURM job array (`panm3_params.txt` + `panm3_array.sh`).
- `run_local.py` — reproduces the same table on a single machine, without SLURM.
- `fixation_curves_mtdna.csv` — the output table, included so the analysis can be rerun
  without repeating the simulations.

## Reproducing the table

On a single machine (slow; the full table is ~130 CPU-hours):

```bash
pip install -r requirements.txt
python run_local.py            # or: python run_local.py --reps 100   (10x faster)
python panm3_collect.py
```

A smoke test that finishes in minutes and reproduces the shape of every curve:

```bash
python run_local.py --quick && python panm3_collect.py
```

On a SLURM cluster:

```bash
python panm3_submit.py 1000 100 500 short   # <reps> <batch> <concurrency> <partition>
sbatch panm3_array.sh
python panm3_collect.py
```

Both paths are resumable: re-running skips any task whose output file is already complete, so
an interrupted run can be restarted with the same command.

## Output schema

`census_N, locus_Ne, s, pfix_pct, n_replicates, se_pct`

83 rows: 15 s-values at each of the five sizes, plus four additional high-s points
(0.04, 0.05, 0.075, 0.10) at `locus_Ne` 500 and 750. Every row uses 1,000 replicates; the
largest standard error is 1.6 percentage points.

## Validation

Every row was checked against the diffusion prediction for a haploid locus starting at
frequency *p* = 0.05,

```
P(fix) = (1 - exp(-2 * Ne * s * p)) / (1 - exp(-2 * Ne * s))
```

The largest deviation across all 83 rows is 3.9 percentage points (at `locus_Ne` = 1000,
s = 0.0075), i.e. 2.4 standard errors, with neighbouring points deviating in the opposite
direction — consistent with sampling noise rather than bias. At `locus_Ne` = 500 the curve also
reproduces the independent SLiM result of Petr et al. (their Fig. S28) to within a few points.

## Dependencies

Python 3.10 and NumPy (`requirements.txt`). No other packages are required to generate the
table; `panm3_collect.py` uses only the standard library and NumPy.
