## Supplementary Information section 1 code base for "Hypothesis: A modern human range expansion ∼300,000 years ago had a critical role in Neandertal origins": run_local.py

```
#!/usr/bin/env python3
"""
run_local.py -- reproduce fixation_curves_mtdna.csv on one machine, without SLURM.

The published table was produced on a cluster (panm3_submit.py + panm3_array.sh); this script
runs the identical worker over the same (s, N) combinations using local processes, so the
result can be reproduced anywhere. It is slow: the full table is ~130 CPU-hours, so with 8
cores expect roughly 16 hours. Use --quick for a reduced version that finishes in minutes and
reproduces the shape of every curve.

    python run_local.py                 # full table, all cores
    python run_local.py --reps 100      # 10x faster, SE ~4.5% instead of ~1.4%
    python run_local.py --quick         # smoke test: 20 reps, sweep sizes only
"""
import argparse
import os
import sys
from multiprocessing import Pool, cpu_count
import panm3_run as W


def one(job):
    s_, N, reps, seed0, out = job
    if os.path.exists(out) and sum(1 for _ in open(out)) >= reps + 1:
        return out                                   # resumable
    sys.argv = ["panm3_run.py", str(s_), str(N), str(reps), str(seed0), out]
    W.main()
    return out


def main():
    ap = argparse.ArgumentParser()
    ap.add_argument("--reps", type=int, default=1000)
    ap.add_argument("--batch", type=int, default=100)
    ap.add_argument("--procs", type=int, default=cpu_count())
    ap.add_argument("--quick", action="store_true")
    a = ap.parse_args()
    combos = W.combos()
    if a.quick:
        a.reps, a.batch = 20, 20
        combos = [(s_, N) for s_, N in combos if N in (1000, 1500, 2000)]
    os.makedirs("out_panm3", exist_ok=True)
    jobs = [(s_, N, min(a.batch, a.reps - k), k,
             f"out_panm3/out_{s_:g}_{N}_s{k}.txt")
            for s_, N in combos for k in range(0, a.reps, a.batch)]
    print(f"{len(jobs)} tasks over {len(combos)} (s, N) combinations, "
          f"{a.reps} replicates each, {a.procs} processes")
    with Pool(a.procs) as p:
        for i, _ in enumerate(p.imap_unordered(one, jobs), 1):
            if i % 20 == 0 or i == len(jobs):
                print(f"  {i}/{len(jobs)}", flush=True)
    print("\nnow run:  python panm3_collect.py")


if __name__ == "__main__":
    main()
```
