## Supplementary Information section 1 code base for "Hypothesis: A modern human range expansion ∼300,000 years ago had a critical role in Neandertal origins": make_grid_replicate.py

```
"""
Write params_replicate.tsv: the full Extended Data Figure 1 grid, so we can
re-run BOTH the original Perl simulation (ratchet.pl) and the new DFE framework
(delta DFE) at every point and scatter new-vs-original.

Grid (from the Extended Data Figure 1 caption):
  NM = 7500, NA in {1500, 3000}, T = 17500, reps = 10,
  mu in {0.001,0.002,0.005,0.01,0.02,0.05}  (whole-locus mutation rate / gen),
  s  in {0.0001,0.0003,0.001,0.003,0.01,0.03,0.1,0.3,1}  (single coefficient).
= 6 x 9 x 2 = 108 grid points.
"""
import csv

NM = 7500
T = 17500
REPS = 30   # more than the original 10 so the new-sim estimate is precise;
            # off-diagonal scatter then mostly reflects the original 10-rep noise
MUS = [0.001, 0.002, 0.005, 0.01, 0.02, 0.05]
SS = [0.0001, 0.0003, 0.001, 0.003, 0.01, 0.03, 0.1, 0.3, 1]
NAS = [1500, 3000]


def main():
    rows, idx, seed = [], 0, 4000
    for Na in NAS:
        for mu in MUS:
            for s in SS:
                rows.append([idx, NM, Na, T, mu, s, REPS, seed])
                idx += 1
                seed += 1
    with open("params_replicate.tsv", "w", newline="") as f:
        w = csv.writer(f, delimiter="\t")
        w.writerow(["idx", "NM", "NA", "T", "mu", "s", "reps", "seed"])
        w.writerows(rows)
    print(f"Wrote params_replicate.tsv: {len(rows)} grid points "
          f"({len(MUS)}x{len(SS)} x {len(NAS)} archaic sizes, {REPS} reps).")
    print(f"Set the sbatch array to 0-{len(rows)-1}.")


if __name__ == "__main__":
    main()
```
