## Supplementary Information section 1 code base for "Hypothesis: A modern human range expansion ∼300,000 years ago had a critical role in Neandertal origins": make_grid_v22.py

```
"""
Write params_v22.tsv: the full Part 1 mtDNA grid rerun under the v22 demography
(8_20_2026 note), replacing the retired params_denis300 grid.

What changed from the denis300 grid and why everything reruns:
  * NM = 7000 (was 7500). The DFE is anchored at the modern size, so
    s_bar = Sbar/(2*NM) moves from 0.065 to 0.069 at Sbar=968 -- every grid
    point uses that coefficient, hence the full rerun.
  * T = 17,483 generations of isolation (Tsep 26,224 - Tmix 8,741 at
    28.6 yr/gen; was 17,500).
  * N_archaic gains the new central value 2250 (brackets 1500 / 3000 kept).
  * A "Petr et al." comparison block: NM=5000, N_A=500, T=10,489 generations
    (Tsep 20,979 - Tmix 10,490), DFE anchored at 5000, central DFE, all three
    U values, tagged 'petr'.

Central DFE (this study):  beta = 0.245,  |Sbar| = 968
Deleterious input (RNA-inclusive):  U = 0.0093  (band 0.0055-0.0133)

Same layout as make_grid_denis300, so run_slurm.sbatch, run_task.py and
merge_chunks.py work unchanged (anchor_ne and tag ride along as columns):

    python make_grid_v22.py
    mkdir -p results_v22
    sbatch --export=ALL,PARAMS=params_v22.tsv,OUTDIR=results_v22 \
           --array=0-<N-1> run_slurm.sbatch
    python merge_chunks.py --outdir results_v22 --params params_v22.tsv
"""
import csv
import itertools

N_MODERN = 7000
T = 17483                     # 26,224 - 8,741 generations (750 -> 250 ka)
ANCHOR = 7000                 # DFE anchor = modern size (s_bar = 968/14000 = 0.069)

SBAR_GRID = [200, 300, 420, 600, 850, 968, 1200, 1400, 1700, 2100, 2400, 3000]
NARCH_GRID = [1500, 2250, 3000]             # 2250 central, 1500/3000 brackets
UMIS_GRID = [0.0055, 0.0093, 0.0133]        # RNA-inclusive low / central / high
BETA_CENTRAL = 0.245

# beta sweep across the (wide) matched-depth CI, at central Sbar/U
BETA_SWEEP = [0.15, 0.20, 0.30, 0.36]
SBAR_CENTRAL = 968
UMIS_CENTRAL = 0.0093

# Petr et al. comparison: smaller archaic size but shorter isolation
PETR = dict(N_modern=5000, N_archaic=500, T=10489, anchor=5000)

REPS_TOTAL = 300
CHUNKS = 10
REPS_PER_CHUNK = REPS_TOTAL // CHUNKS
SEED_BASE = 500000            # distinct from the retired denis300 grid (300000)


def combos():
    """(Sbar, beta, U, N_arch, N_mod, T, anchor, tag) tuples, deduplicated."""
    out, seen = [], set()

    def add(c):
        if c not in seen:
            seen.add(c); out.append(c)

    for Sbar, Na, U in itertools.product(SBAR_GRID, NARCH_GRID, UMIS_GRID):
        add((Sbar, BETA_CENTRAL, U, Na, N_MODERN, T, ANCHOR, ""))
    for beta, Na in itertools.product(BETA_SWEEP, NARCH_GRID):
        add((SBAR_CENTRAL, beta, UMIS_CENTRAL, Na, N_MODERN, T, ANCHOR, ""))
    for U in UMIS_GRID:
        add((SBAR_CENTRAL, BETA_CENTRAL, U, PETR["N_archaic"],
             PETR["N_modern"], PETR["T"], PETR["anchor"], "petr"))
    return out


def main():
    uniq = combos()
    rows = []
    for ci, (Sbar, beta, U, Na, Nm, Tgen, anchor, tag) in enumerate(uniq):
        for chunk in range(CHUNKS):
            rows.append(dict(idx=len(rows), combo=ci, chunk=chunk,
                             Sbar=Sbar, beta=beta, U_mis=U, N_archaic=Na,
                             N_modern=Nm, T=Tgen, reps=REPS_PER_CHUNK,
                             seed=SEED_BASE + 100 * ci + chunk,
                             anchor_ne=anchor, tag=tag))
    cols = ["idx", "combo", "chunk", "Sbar", "beta", "U_mis", "N_archaic",
            "N_modern", "T", "reps", "seed", "anchor_ne", "tag"]
    with open("params_v22.tsv", "w", newline="") as f:
        w = csv.DictWriter(f, fieldnames=cols, delimiter="\t",
                           lineterminator="\n")
        w.writeheader(); w.writerows(rows)
    if len({r["seed"] for r in rows}) != len(rows):
        raise SystemExit("seed collision")
    n_petr = sum(1 for c in uniq if c[7] == "petr")
    print(f"wrote params_v22.tsv: {len(uniq)} combinations "
          f"({len(uniq) - n_petr} main + {n_petr} petr) x {CHUNKS} chunks "
          f"= {len(rows)} tasks")
    print(f"submit with:\n  sbatch --export=ALL,PARAMS=params_v22.tsv,"
          f"OUTDIR=results_v22 --array=0-{len(rows)-1} run_slurm.sbatch")


if __name__ == "__main__":
    main()
```
