## Supplementary Information section 1 code base for "Hypothesis: A modern human range expansion ∼300,000 years ago had a critical role in Neandertal origins": make_grid_y_draw.py

```
"""
Write params_y_draw.tsv: the PARAMETER-INTEGRATED Y target-size ladder.
Same 28 combinations x 10 chunks = 280 tasks as params_y_v22.tsv (7 targets
x {N_A = 1500, 2250, 3000, plus the Petr scenario}), but run by
run_task_ydraw.py, which draws (beta, Sbar, U) per replicate from the
triangular priors of the Methods' bound calculation instead of fixing them
at the central values. Seeds 800000+ (disjoint from every earlier grid).

    python make_grid_y_draw.py --conv stated
    mkdir -p logs results_y_draw_stated
    sbatch --export=ALL,PARAMS=params_y_draw_stated.tsv,OUTDIR=results_y_draw_stated \
           --job-name=y_draw_st --array=0-319 run_slurm_ydraw.sbatch
    python collect_y_draw.py --outdir results_y_draw_stated \
           --params params_y_draw_stated.tsv --out y_draw_stated
"""
import argparse
import csv

# 3000 kb appended at the end: combo numbering and seeds of the original
# 280-task submission are unchanged; tasks 280-319 are the 3-Mb extension.
TARGETS_KB = [100, 200, 300, 500, 1000, 2000, 5000, 3000]
N_MODERN = 7000
T = 17483
NARCH_GRID = [1500, 2250, 3000]
PETR = dict(N_modern=5000, N_archaic=500, T=10489)

# run_task_ydraw.py draws Sbar ~ tri(225, 425, 766) (ref. 12's 95% CI in its
# own Ne*s convention) and computes s = Sbar/(2*anchor_ne); the conversion is
# therefore encoded through an effective anchor, exactly as in
# make_grid_y_v22.py: stated s = Sbar/Nref -> anchor 5000; hemizygous
# s = 2*Sbar/Nref -> anchor 2500; old (superseded) -> anchor 10000.
CONVERSIONS = {
    "stated": dict(anchor=5000, seed_base=890000, stem="y_draw_stated"),
    "hemi": dict(anchor=2500, seed_base=920000, stem="y_draw_hemi"),
    "old": dict(anchor=10000, seed_base=800000, stem="y_draw"),
}

REPS_TOTAL = 300
CHUNKS = 10


def main():
    ap = argparse.ArgumentParser(description=__doc__)
    ap.add_argument("--conv", choices=list(CONVERSIONS), default="stated")
    args = ap.parse_args()
    cfg = CONVERSIONS[args.conv]
    params = f"params_{cfg['stem']}.tsv"

    combos = []
    for kb in TARGETS_KB:
        for Na in NARCH_GRID:
            combos.append((f"y{kb}", kb, N_MODERN, Na, T))
        combos.append((f"y{kb}_petr", kb, PETR["N_modern"],
                       PETR["N_archaic"], PETR["T"]))
    rows = []
    for ci, (tag, kb, Nm, Na, Tgen) in enumerate(combos):
        for chunk in range(CHUNKS):
            rows.append(dict(idx=len(rows), combo=ci, chunk=chunk, tag=tag,
                             target_kb=kb, N_archaic=Na, N_modern=Nm, T=Tgen,
                             reps=REPS_TOTAL // CHUNKS,
                             seed=cfg["seed_base"] + 100 * ci + chunk,
                             anchor_ne=cfg["anchor"]))
    cols = ["idx", "combo", "chunk", "tag", "target_kb", "N_archaic",
            "N_modern", "T", "reps", "seed", "anchor_ne"]
    with open(params, "w", newline="") as f:
        w = csv.DictWriter(f, fieldnames=cols, delimiter="\t",
                           lineterminator="\n")
        w.writeheader(); w.writerows(rows)
    if len({r["seed"] for r in rows}) != len(rows):
        raise SystemExit("seed collision")
    print(f"wrote {params} ({args.conv}: central s = "
          f"{425.0 / (2 * cfg['anchor']):.4f}): {len(combos)} combinations "
          f"x {CHUNKS} chunks = {len(rows)} tasks")
    print(f"submit with:\n  mkdir -p logs results_{cfg['stem']}\n"
          f"  sbatch --export=ALL,PARAMS={params},"
          f"OUTDIR=results_{cfg['stem']} --job-name={cfg['stem']} "
          f"--array=0-{len(rows)-1} run_slurm_ydraw.sbatch")


if __name__ == "__main__":
    main()
```
