## Supplementary Information section 1 code base for "Hypothesis: A modern human range expansion ∼300,000 years ago had a critical role in Neandertal origins": make_krause_table.py

```
"""
Generate Krause_substitution_table.docx: per-lineage synonymous vs
nonsynonymous mtDNA substitution counts (Green et al. 2008-style table),
extracted from Krause et al. 2010 (Nature 464:894), Supplementary Figure 5.

Run:  ./mtdna_dfe_sim/.venv/bin/python make_krause_table.py
"""
from docx import Document
from docx.shared import Pt, RGBColor

OUT = "Krause_substitution_table.docx"

doc = Document()
doc.styles["Normal"].font.name = "Calibri"
doc.styles["Normal"].font.size = Pt(11)


def p(text="", italic=False, bold=False, size=11):
    par = doc.add_paragraph()
    r = par.add_run(text)
    r.italic = italic
    r.bold = bold
    r.font.size = Pt(size)
    return par


def table(data):
    t = doc.add_table(rows=len(data), cols=len(data[0]))
    t.style = "Table Grid"
    for i, row in enumerate(data):
        for j, val in enumerate(row):
            cell = t.cell(i, j)
            cell.text = ""
            run = cell.paragraphs[0].add_run(val)
            run.font.size = Pt(10)
            if i == 0:
                run.bold = True
    return t


doc.add_heading("Synonymous vs. nonsynonymous mtDNA substitutions per hominin "
                "lineage", level=0)
p("Source: Krause et al. (2010), Nature 464:894\u2013897, Supplementary "
  "Figure 5 ('Rates of hominin mtDNA protein evolution estimated without or "
  "with the Denisova mtDNA'). For each branch the figure gives dN/dS and, in "
  "parentheses, the estimated numbers of (non-synonymous, synonymous) "
  "substitutions. Format follows Green et al. (2008), Cell 134:416.",
  italic=True, size=9)

doc.add_heading("Table 1. Tree including Denisova (preferred reconstruction)", 2)
table([
    ["Lineage / branch", "Synonymous", "Nonsynonymous", "dN/dS"],
    ["Modern human", "38.3", "11.8", "0.105"],
    ["Neandertal", "39.2", "13.6", "0.118"],
    ["Modern\u2013Neandertal shared ancestor", "79.5", "9.8", "0.042"],
    ["Denisova", "108.0", "17.9", "0.056"],
])
p("")

doc.add_heading("Table 2. Tree without Denisova (for comparison)", 2)
table([
    ["Lineage / branch", "Synonymous", "Nonsynonymous", "dN/dS"],
    ["Modern human", "49.1", "11.9", "0.082"],
    ["Neandertal", "27.5", "13.7", "0.168"],
    ["Modern\u2013Neandertal ancestor (deep internal edge)*", "1,217.2",
     "122.4", "0.034"],
])
p("*In the four-taxon tree without Denisova there is a single internal edge "
  "(shared between the chimpanzee/bonobo side and the hominin side), so this "
  "branch lumps the deep divergence and its large counts are not directly "
  "comparable to the cleanly resolved ancestral branch in Table 1.", size=9,
  italic=True)

doc.add_heading("Notes", 2)
p("\u2022 Counts are fractional because they are maximum-likelihood expected "
  "substitution numbers from a model-based reconstruction (HKY/TrN+I+\u0393), "
  "not direct integer tallies: ancestral states at internal nodes are inferred "
  "with uncertainty (and apportioned fractionally between adjoining branches), "
  "multiple/back substitutions are corrected for, and the synonymous vs. "
  "nonsynonymous classification itself splits ambiguous sites and multi-step "
  "codon changes fractionally.")
p("\u2022 Adding Denisova as a third, deeply diverged lineage reassigns "
  "synonymous substitutions on the Neandertal terminal branch (27.5 \u2192 "
  "39.2) and lowers its dN/dS from 0.168 to 0.118, close to modern humans "
  "(0.105). The apparent Neandertal 'excess' of nonsynonymous change in "
  "earlier analyses is thus largely a long-branch reconstruction artifact.")
p("\u2022 The shared modern\u2013Neandertal ancestral branch has a low dN/dS "
  "(0.042), lower than either terminal lineage; the elevated dN/dS on the "
  "terminal branches reflects recent, slightly deleterious nonsynonymous "
  "variants not yet removed by purifying selection.")
p("\u2022 For reference, Green et al. (2008) reported (on a human+Neanderthal "
  "tree, different method): Neandertal 44 synonymous / 20 nonsynonymous; "
  "modern human 57 synonymous / 18 nonsynonymous.")

doc.save(OUT)
print(f"Wrote {OUT}")
```
