## Supplementary Information section 1 code base for "Hypothesis: A modern human range expansion ∼300,000 years ago had a critical role in Neandertal origins": merge_chunks.py

```
"""
Pool the chunked cluster output into one result per parameter combination.

Reads <outdir>/task_*.jsonl, groups by combination, concatenates the
per-replicate rows and recomputes the summary from the pooled 300 replicates.
Standard errors come from the pooled replicates rather than from averaging the
chunks' standard errors, which would be wrong.

Two output formats, because the two grids feed different downstream scripts:

    # mtDNA grid -> combo_<n>.jsonl, what collect_and_plot.py globs
    python merge_chunks.py --outdir results300 --params params_300.tsv
    mv results results_30reps
    mkdir results && cp results300/combo_*.jsonl results/
    python collect_and_plot.py

    # Y and RNA grid -> extra_results.csv, what plot_y.py and plot_extra.py read
    python merge_chunks.py --outdir results_extra300 \
        --params params_extra_300.tsv --format csv
    cp extra_results.csv extra_results_50reps.csv     # keep the old one first
    cp results_extra300/extra_results.csv .
    python plot_y.py && python plot_extra.py

The run refuses to write anything if any combination is missing chunks, and
prints the array indices to resubmit, so a partial array cannot quietly produce
an underpowered figure that still looks finished.
"""
import argparse
import csv
import glob
import json
import os
from collections import defaultdict

from ratchet_dfe import summarize

CSV_COLS = ["tag", "N_modern", "N_archaic", "anchor_ne", "Sbar", "s_bar",
            "beta", "U_mis", "U_non", "T", "reps", "w_modern_mean",
            "w_archaic_mean", "diff_mean", "diff_sd", "diff_se",
            "frac_reps_over_5pct"]


def main():
    ap = argparse.ArgumentParser()
    ap.add_argument("--outdir", default="results300")
    ap.add_argument("--params", default="params_300.tsv")
    ap.add_argument("--allow-partial", action="store_true",
                    help="write combinations that are missing chunks")
    ap.add_argument("--format", choices=("jsonl", "csv"), default="jsonl",
                    help="jsonl for the mtDNA grid (collect_and_plot.py), "
                         "csv for the Y/RNA grid (plot_y.py, plot_extra.py)")
    args = ap.parse_args()

    with open(args.params) as f:
        grid = list(csv.DictReader(f, delimiter="\t"))
    expected = defaultdict(set)
    for r in grid:
        expected[int(r["combo"])].add(int(r["chunk"]))

    got = defaultdict(dict)
    for path in glob.glob(os.path.join(args.outdir, "task_*.jsonl")):
        with open(path) as f:
            d = json.loads(f.readline())
        s = d["summary"]
        got[s["combo"]][s["chunk"]] = d

    missing = {c: sorted(expected[c] - set(got.get(c, {})))
               for c in expected if expected[c] - set(got.get(c, {}))}
    if missing:
        print(f"incomplete: {len(missing)} of {len(expected)} combinations")
        for c in sorted(missing)[:10]:
            print(f"   combo {c}: missing chunks {missing[c]}")
        if not args.allow_partial:
            idx = sorted(int(r["idx"]) for r in grid
                         if int(r["combo"]) in missing
                         and int(r["chunk"]) in missing[int(r["combo"])])
            print(f"\nresubmit these {len(idx)} tasks with:")
            print("   sbatch --array=" + ",".join(map(str, idx[:50]))
                  + ("..." if len(idx) > 50 else "") + " run_slurm.sbatch")
            raise SystemExit("refusing to merge; pass --allow-partial to override")

    written, csv_rows = 0, []
    for combo, chunks in sorted(got.items()):
        rows, seeds = [], []
        for ch in sorted(chunks):
            d = chunks[ch]
            seeds.append(d["summary"]["seed"])
            for r in d["reps"]:
                rows.append(dict(r, rep=len(rows)))
        s0 = chunks[sorted(chunks)[0]]["summary"]
        summary = summarize(
            rows, N_modern=s0["N_modern"], N_archaic=s0["N_archaic"],
            T=s0["T"], U_mis=s0["U_mis"], U_non=s0["U_non"], beta=s0["beta"],
            Sbar=s0["Sbar"], s_bar=s0["s_bar"], reps=len(rows),
            seed=seeds[0],
            elapsed=sum(chunks[c]["summary"].get("elapsed_sec") or 0
                        for c in chunks))
        summary["combo"] = combo
        summary["seeds"] = seeds
        summary["chunks"] = len(chunks)
        summary["anchor_ne"] = s0.get("anchor_ne")
        if s0.get("tag"):
            summary["tag"] = s0["tag"]

        if args.format == "jsonl":
            out = os.path.join(args.outdir, f"combo_{combo}.jsonl")
            with open(out, "w") as f:
                f.write(json.dumps({"summary": summary, "reps": rows}) + "\n")
            written += 1
        else:
            csv_rows.append(summary)

        print(f"combo {combo:>3}: {s0.get('tag', ''):>9} "
              f"Sbar={s0['Sbar']:>6} beta={s0['beta']} "
              f"U={s0['U_mis']} N_A={s0['N_archaic']:>5} | "
              f"{len(rows):>4} reps  gap={100 * summary['diff_mean']:.4f}% "
              f"+-{100 * summary['diff_se']:.4f}")

    if args.format == "csv":
        out = os.path.join(args.outdir, "extra_results.csv")
        with open(out, "w", newline="") as f:
            w = csv.DictWriter(f, fieldnames=CSV_COLS, extrasaction="ignore")
            w.writeheader()
            w.writerows(csv_rows)
        print(f"\nwrote {len(csv_rows)} rows to {out}")
    else:
        print(f"\nwrote {written} combo files to {args.outdir}/")


if __name__ == "__main__":
    main()
```
