## Supplementary Information section 1 code base for "Hypothesis: A modern human range expansion ∼300,000 years ago had a critical role in Neandertal origins": mt_dfe_sfs.py

```
"""
Extend Piganeau & Eyre-Walker (2003) to the whole mitochondrial genome: fit a
gamma distribution of fitness effects separately to protein-coding, tRNA and
rRNA sites from the site frequency spectrum of 56,434 human mtDNA genomes
(gnomAD v3.1 chrM).

P&EW fitted a gamma DFE to human mtDNA *protein-coding* polymorphism and
divergence and obtained shape beta ~ 0.4 with a large mean effect. The ratchet
analysis in this project assumes mutations in the ~4 kb of tRNA and rRNA can be
treated as if drawn from that same protein DFE. That assumption has never been
tested, because no DFE has been published for mitochondrial RNA genes. This
script tests it, and refits the coding DFE by the identical procedure as a
control so the RNA numbers can be read against a benchmark we already trust.

Model
-----
Poisson Random Field (Sawyer & Hartl 1992). mtDNA is haploid, so the scaled
selection coefficient is S = 2*Ne*s (the convention already used in
krause_load.py). The stationary density of a derived allele at frequency x for
a deleterious variant of strength S is

    f_S(x) = theta * phi(x,S) / (x (1-x)),
    phi(x,S) = (e^{-Sx} - e^{-S}) / (1 - e^{-S}),      phi(x,0) = 1-x

and the expected number of sites where the derived allele is seen i times in a
sample of n is the binomial sampling integral of f_S. Substituting the logit
x = 1/(1+e^-u) gives dx = x(1-x) du, which cancels the singular denominator
exactly and leaves a smooth bounded integrand -- so the integral is computed on
a plain uniform grid in u with no special handling of the endpoints.

Because the S-grid is fixed we precompute H[k,i] = expected SFS at S_k once.
Averaging over a gamma DFE is then just a weight vector times H, which makes
the likelihood cheap enough to optimise directly.

Folding
-------
The spectrum is FOLDED (binned by minor allele count). Polarising human mtDNA
against chimp does not work: at 6-7 Myr with mtDNA's transition bias the
synonymous sites are saturated (we measure d_syn = 0.32 with 62 multiply-hit
codons), so chimp matches the rCRS by chance at a large fraction of variable
sites. Polarising this way put 22.9% of synonymous variants in the top
frequency bin, which is ancestral misidentification, not biology. Folding
removes the problem entirely and costs almost nothing here, because the DFE
signal lives in rare variants where minor = derived essentially always.

Sample size
-----------
No projection. Subsampling 56,433 chromosomes down to n=1000 discarded three
quarters of the variants (1,006 of 4,050 synonymous survived), and it discards
exactly the rare end that carries the selection signal. Instead the binomial
sampling probability for a whole bin is evaluated in closed form with the
regularised incomplete beta function,
    P(X >= k) = I_x(k, n-k+1),   X ~ Binom(n, x)
so we work at the full n with no large matrices and no information loss.

Nuisance parameters
-------------------
demography  Human mtDNA is far from equilibrium (massive recent expansion), so
            the raw PRF shape is wrong. We take the *observed synonymous* SFS as
            the empirical neutral shape and correct every selected class by the
            per-bin ratio r_b = obs_syn_b / expected_neutral_b. This is the
            standard DFE-alpha / polyDFE style correction: it absorbs demography
            and sampling structure to the extent that they act multiplicatively
            across classes.

Caveats that no amount of fitting removes: mtDNA does not recombine, so the PRF
independence assumption is violated and linked selection will bias beta
downward; the multiplicative demographic correction is only an approximation,
because strongly deleterious variants equilibrate faster than neutral ones and
so carry less of the expansion signature than the correction assumes (this
pushes inferred selection to be too weak); and gnomAD is a mixture of
populations, which inflates rare variants in all classes alike.
"""
import argparse
import gzip
import math
import os
from collections import defaultdict

import numpy as np
from scipy.optimize import minimize
from scipy.special import gammainc, betainc
from Bio import SeqIO

from rna_constraint import (build_site_table, align_to_ref, SEQDIR, TITV,
                            analyse as rc_analyse)

VCF = "data/gnomad.chrM.vcf.bgz"
GB = "data/NC_012920.gb"
CHIMP = "NC_001643.1"

N_COMMON = 56433       # gnomAD chrM sample size; counts are rescaled to this
N_BINS = 18            # log-spaced minor-allele-count bins used in the fit
NE_ANCHOR = 7500.0     # for reporting s alongside S (same anchor as krause_load)


# ---------------------------------------------------------------- PRF spectrum
def _phi(x, S):
    """Stationary PRF density factor, with the x(1-x) denominator removed."""
    if S < 1e-8:
        return 1.0 - x
    return (np.exp(-S * x) - math.exp(-S)) / (1.0 - math.exp(-S))


def prf_folded_bins(n, S_grid, bins, n_u=8000, u_max=30.0):
    """H[k,b] = expected number of sites falling in minor-allele-count bin b,
    per unit theta, at selection strength S_grid[k].

    The logit substitution x = 1/(1+e^-u) gives dx = x(1-x) du, cancelling the
    PRF denominator exactly, so the integrand is smooth and bounded. Binomial
    bin probabilities come from the regularised incomplete beta, which lets us
    stay at the full sample size."""
    u = np.linspace(-u_max, u_max, n_u)
    x = 1.0 / (1.0 + np.exp(-u))

    def p_ge(k):
        """P(Binom(n,x) >= k) for the whole x grid."""
        if k <= 0:
            return np.ones_like(x)
        if k > n:
            return np.zeros_like(x)
        return betainc(k, n - k + 1, x)

    # P(count in [a,b)) for the minor allele, plus the mirrored high-frequency
    # copy, since folding maps count i and n-i onto the same bin
    Pb = []
    for a, b in bins:
        low = p_ge(a) - p_ge(b)
        high = p_ge(n - b + 1) - p_ge(n - a + 1)
        Pb.append(low + high)
    Pb = np.array(Pb)                                   # (nbins, n_u)

    H = np.empty((len(S_grid), len(bins)))
    for k, S in enumerate(S_grid):
        H[k] = np.trapezoid(Pb * _phi(x, S), u, axis=1)
    return H


def gamma_weights(S_grid_edges, beta, Sbar):
    """Probability mass of Gamma(shape=beta, mean=Sbar) in each grid cell."""
    scale = Sbar / beta
    cdf = gammainc(beta, S_grid_edges / scale)      # regularised lower incomplete
    w = np.diff(cdf)
    tot = w.sum()
    return w / tot if tot > 0 else w


# ------------------------------------------------------------------ input data
def load_sites():
    rec = SeqIO.read(GB, "genbank")
    site, refseq = build_site_table(rec)
    chimp = str(SeqIO.read(os.path.join(SEQDIR, f"{CHIMP}.fa"), "fasta").seq).upper()
    return site, refseq, align_to_ref(refseq, chimp)


def class_of(site, p, ref, alt):
    """Functional class of a specific substitution at rCRS position p."""
    s = site.get(p)
    if s is None:
        return None
    if s["cls"] != "CDS":
        return s["cls"] if s["cls"] in ("tRNA", "rRNA") else None
    cl = s["alt_class"].get(alt)
    if cl is None:
        return None
    return "synonymous" if cl == "synonymous" else "nonsynonymous"


POPS = ['afr', 'ami', 'amr', 'asj', 'eas', 'fin', 'nfe', 'oth', 'sas', 'mid']


def read_variants(site, refseq, use_het=False, an_min=50000, n_common=56433,
                  pop=None):
    """Return {class: array of minor allele counts} rescaled to a common sample
    size. No polarisation: the spectrum is folded.

    If `pop` is given, restrict to that gnomAD population. mtDNA does not
    recombine, so pooling ten populations superimposes ten haplogroup
    structures and drives an excess of common variants that the Poisson Random
    Field cannot represent; a single population is closer to the model."""
    out = defaultdict(list)
    stats = defaultdict(int)
    ip = POPS.index(pop) if pop else None
    with gzip.open(VCF, "rt") as f:
        for line in f:
            if line.startswith("#"):
                continue
            c = line.rstrip("\n").split("\t")
            if c[6] != "PASS":
                continue
            ref, alt = c[3], c[4]
            if len(ref) != 1 or len(alt) != 1:
                continue
            p = int(c[1])
            info = dict(kv.split("=", 1) for kv in c[7].split(";") if "=" in kv)
            if ip is None:
                an = int(info["AN"])
                ac = int(info["AC_hom"]) + (int(info["AC_het"]) if use_het else 0)
            else:
                an = int(info["pop_AN"].split("|")[ip])
                ac = int(info["pop_AC_hom"].split("|")[ip])
                if use_het:
                    ac += int(info["pop_AC_het"].split("|")[ip])

            if refseq[p - 1] != ref:            # VCF must match rCRS
                stats["ref_mismatch"] += 1
                continue
            if an < an_min:
                stats["low_AN"] += 1
                continue
            cls = class_of(site, p, ref, alt)
            if cls is None:
                stats["unclassified"] += 1
                continue
            minor = min(ac, an - ac)
            if minor <= 0:
                stats["monomorphic"] += 1
                continue
            out[cls].append(max(1, int(round(minor * n_common / an))))
            stats[cls] += 1
    return {k: np.array(v) for k, v in out.items()}, stats


def pop_sample_size(pop):
    """Median allele number for one gnomAD population."""
    ip = POPS.index(pop)
    ans = []
    with gzip.open(VCF, "rt") as f:
        for line in f:
            if line.startswith("#"):
                continue
            c = line.rstrip("\n").split("\t")
            if c[6] != "PASS":
                continue
            info = dict(kv.split("=", 1) for kv in c[7].split(";") if "=" in kv)
            ans.append(int(info["pop_AN"].split("|")[ip]))
    return int(np.median(ans))


def make_bins(n, nbins):
    """Log-spaced bins of minor allele count, from 1 up to n/2."""
    edges = np.unique(np.round(np.logspace(0, math.log10(n // 2), nbins + 1))
                      ).astype(int)
    edges[-1] = n // 2 + 1
    return [(int(edges[k]), int(edges[k + 1])) for k in range(len(edges) - 1)]


def binned(counts, bins):
    return np.array([np.sum((counts >= a) & (counts < b)) for a, b in bins],
                    dtype=float)


# --------------------------------------------------------------------- fitting
def opportunity(site, refseq, titv=TITV):
    """Spectrum-weighted mutational opportunity per class, in site-equivalents.
    Identical convention to rna_constraint.py, but over every classified site
    rather than only those callable in an alignment."""
    from rna_constraint import weight, per_site_weight
    psw = per_site_weight(titv)
    opp = defaultdict(float)
    for p, s in site.items():
        ref = refseq[p - 1]
        if s["cls"] == "CDS":
            for alt, cl in s["alt_class"].items():
                key = "synonymous" if cl == "synonymous" else "nonsynonymous"
                opp[key] += weight(ref, alt, titv) / psw
        elif s["cls"] in ("tRNA", "rRNA"):
            opp[s["cls"]] += 1.0
    return opp


def fit_gamma(obs_b, r_b, H, S_edges, theta, div=None, S_mid=None, warm=None):
    """Gamma DFE for one class. theta is FIXED from the synonymous rate scaled
    by mutational opportunity, so the deficit of selected variants -- not just
    the shape of the spectrum -- informs the deleterious mass. Optionally adds
    a divergence term, div = (k_obs, opportunity, d_syn), which is what
    Piganeau & Eyre-Walker used to pin the strongly deleterious tail."""
    fr = fixrel(S_mid) if div is not None else None

    def nll(par):
        beta, Sbar = math.exp(par[0]), math.exp(par[1])
        if not (0.02 < beta < 20 and 1e-4 < Sbar < 1e8):
            return 1e12
        w = gamma_weights(S_edges, beta, Sbar)
        if not np.isfinite(w).all():
            return 1e12
        e_b = np.maximum((w @ H) * r_b * theta, 1e-12)
        ll = float(np.sum(e_b - obs_b * np.log(e_b)))
        if div is not None:
            k_obs, opp, d_syn = div
            lam = max(opp * d_syn * float(w @ fr), 1e-12)
            ll += lam - k_obs * math.log(lam)
        return ll if np.isfinite(ll) else 1e12

    if warm is not None:
        starts = [(math.log(warm[0]), math.log(warm[1]))]
    else:
        starts = [(math.log(b0), math.log(S0))
                  for b0 in (0.08, 0.15, 0.3, 0.5, 0.9)
                  for S0 in (1.0, 10.0, 100.0, 1e3, 1e4, 1e5)]
    best, best_x = np.inf, None
    for x0 in starts:
        res = minimize(nll, list(x0), method="Nelder-Mead",
                       options={"maxiter": 20000, "maxfev": 20000,
                                "xatol": 1e-7, "fatol": 1e-7})
        if res.fun < best:
            best, best_x = res.fun, res.x
    return {"beta": math.exp(best_x[0]), "Sbar": math.exp(best_x[1]),
            "nll": best}


def fixrel(S):
    """Fixation probability of a deleterious mutation relative to neutral, in a
    haploid population: S/(e^S - 1). Same expression as krause_load.w_fix."""
    S = np.asarray(S, dtype=float)
    out = np.ones_like(S)
    mid = (S > 1e-9) & (S <= 700)
    big = S > 700
    out[mid] = S[mid] / np.expm1(S[mid])
    out[big] = S[big] * np.exp(-S[big])
    return out


def fit_discrete(obs_b, r_b, Hg, theta):
    """DFE as free mass in coarse selection intervals, with theta fixed by
    opportunity. This is the model-free version: it makes no assumption about
    the shape of the tail, and because theta is pinned, the mass in the
    invisible S>100 class is identified by how many variants are MISSING."""
    K = Hg.shape[0]

    def nll(par):
        p = np.exp(par - par.max())
        p = p / p.sum()
        e_b = np.maximum((p @ Hg) * r_b * theta, 1e-12)
        v = float(np.sum(e_b - obs_b * np.log(e_b)))
        return v if np.isfinite(v) else 1e12

    best, best_x = np.inf, None
    for seed in range(12):
        x0 = np.random.default_rng(seed).normal(0, 1.0, K)
        res = minimize(nll, x0, method="Nelder-Mead",
                       options={"maxiter": 60000, "maxfev": 60000,
                                "xatol": 1e-7, "fatol": 1e-7})
        if res.fun < best:
            best, best_x = res.fun, res.x
    p = np.exp(best_x - best_x.max())
    return p / p.sum(), best


def near_neutral_mass(beta, Sbar, lo=1.0, hi=10.0):
    """Fraction of the DFE with S in [lo,hi] -- the window that drives the
    ratchet, where drift and selection are comparable."""
    scale = Sbar / beta
    return float(gammainc(beta, hi / scale) - gammainc(beta, lo / scale))


def main():
    ap = argparse.ArgumentParser()
    ap.add_argument("--het", action="store_true",
                    help="include heteroplasmic carriers in the allele counts")
    ap.add_argument("--boot", type=int, default=200,
                    help="bootstrap replicates for confidence intervals")
    ap.add_argument("--pop", default=None, choices=POPS,
                    help="restrict to one gnomAD population")
    args = ap.parse_args()

    rec = SeqIO.read(GB, "genbank")
    site, refseq = build_site_table(rec)
    if args.pop:
        an_med = pop_sample_size(args.pop)
        variants, stats = read_variants(site, refseq, use_het=args.het,
                                        an_min=int(0.9 * an_med),
                                        n_common=an_med, pop=args.pop)
        N = an_med
    else:
        variants, stats = read_variants(site, refseq, use_het=args.het)
        N = N_COMMON

    n = N
    bins = make_bins(n, N_BINS)

    print(f"gnomAD v3.1 chrM, n = {n:,} chromosomes, homoplasmic calls"
          + (" + heteroplasmic" if args.het else ""))
    print("folded spectrum (minor allele count); no outgroup polarisation")
    print("  variants by class : " + ", ".join(
        f"{k}={stats[k]}" for k in
        ("synonymous", "nonsynonymous", "tRNA", "rRNA")))
    print("  dropped           : " + ", ".join(
        f"{k}={stats[k]}" for k in
        ("unclassified", "monomorphic", "low_AN", "ref_mismatch")))

    S_edges = np.concatenate([[0.0], np.logspace(-3, 7, 200)])
    S_mid = np.sqrt(np.maximum(S_edges[:-1], 1e-4) * S_edges[1:])
    S_mid[0] = 0.0
    print(f"\nprecomputing folded PRF spectra on {len(S_mid)} selection values"
          f" ({len(bins)} bins)...")
    H = prf_folded_bins(n, S_mid, bins)

    # sanity: folded neutral expectation must be theta*(1/i + 1/(n-i))
    i = np.arange(1, n // 2 + 1)
    exact = np.array([np.sum((1.0 / i + 1.0 / (n - i))[(i >= a) & (i < b)])
                      for a, b in bins])
    err = np.max(np.abs(H[0] - exact) / exact)
    print(f"  neutral check vs theta*(1/i + 1/(n-i)): max rel. error "
          f"{err:.2e}")

    # empirical neutral shape from synonymous sites, split in half so the
    # control fit is genuinely held out rather than tautological
    syn = variants["synonymous"]
    rng = np.random.default_rng(0)
    perm = rng.permutation(len(syn))
    syn_a, syn_b_ = syn[perm[::2]], syn[perm[1::2]]

    def distortion(counts):
        obs = binned(counts, bins)
        exp0 = H[0] * obs.sum() / H[0].sum()
        return np.maximum(obs / np.maximum(exp0, 1e-12), 1e-4), obs

    r_full, syn_obs = distortion(syn)
    r_half, _ = distortion(syn_a)
    print("demographic distortion r_b (observed synonymous / equilibrium):")
    print("  " + " ".join(f"{v:.2f}" for v in r_full))

    # theta is pinned, not fitted: the synonymous rate scaled by each class's
    # mutational opportunity. This is what makes the DEFICIT of selected
    # variants informative about the strongly deleterious mass.
    opp_p = opportunity(site, refseq)
    theta_syn = syn_obs.sum() / H[0].sum()
    print("\nmutational opportunity (site-equivalents): " + ", ".join(
        f"{k}={opp_p[k]:.0f}" for k in
        ("synonymous", "nonsynonymous", "tRNA", "rRNA")))
    print(f"theta_syn = {theta_syn:.2f}; theta for each class is fixed at "
          f"theta_syn x opp_class/opp_syn")

    rna = np.concatenate([variants["tRNA"], variants["rRNA"]])
    classes = [("nonsynonymous", variants["nonsynonymous"], ("nonsynonymous",)),
               ("tRNA", variants["tRNA"], ("tRNA",)),
               ("rRNA", variants["rRNA"], ("rRNA",)),
               ("RNA (tRNA+rRNA)", rna, ("tRNA", "rRNA"))]

    def theta_of(keys):
        return theta_syn * sum(opp_p[k] for k in keys) / opp_p["synonymous"]

    # observed vs neutral-expected variant counts: the raw deficit signal
    print(f"\n{'class':<22}{'observed':>10}{'exp. if neutral':>18}{'deficit':>10}")
    for cls, counts, keys in classes:
        exp_n = theta_of(keys) * float(np.sum(H[0] * r_full))
        print(f"{cls:<22}{len(counts):>10}{exp_n:>18.0f}"
              f"{1 - len(counts) / exp_n:>10.2f}")

    print(f"\n{'class':<22}{'n var':>7}{'beta':>9}{'Sbar':>11}{'sbar':>11}"
          f"{'P(S<1)':>9}{'P(1<S<10)':>11}{'P(S>10)':>9}")

    def show(tag, n_var, f):
        b, S = f["beta"], f["Sbar"]
        print(f"{tag:<22}{n_var:>7}{b:>9.3f}{S:>11.1f}{S / (2 * NE_ANCHOR):>11.2e}"
              f"{near_neutral_mass(b, S, 1e-6, 1.0):>9.3f}"
              f"{near_neutral_mass(b, S, 1.0, 10.0):>11.3f}"
              f"{near_neutral_mass(b, S, 10.0, 1e9):>9.3f}")

    # control: held-out synonymous. A correct implementation must return
    # Sbar ~ 0 on genuinely neutral sites.
    ctrl = fit_gamma(binned(syn_b_, bins), r_half, H, S_edges,
                     len(syn_b_) / float(np.sum(H[0] * r_half)))
    show("synonymous (held out)", len(syn_b_), ctrl)

    print("\nPOLYMORPHISM ONLY (theta pinned by opportunity):")
    for cls, counts, keys in classes:
        f = fit_gamma(binned(counts, bins), r_full, H, S_edges, theta_of(keys))
        show(cls, len(counts), f)

    # ---- P&EW-style joint fit: polymorphism + divergence ---------------------
    deni = str(SeqIO.read(os.path.join(SEQDIR, "FN673705.1.fa"), "fasta").seq).upper()
    obs_d, opp_d, _ = rc_analyse(site, refseq, align_to_ref(refseq, deni), TITV)
    d_syn = obs_d["synonymous"] / opp_d["synonymous"]
    print("\nPOLYMORPHISM + DIVERGENCE (the P&EW approach). Divergence is the")
    print("rCRS-vs-Denisova contrast, unsaturated (0 codons with >1 change);")
    print(f"d_syn = {d_syn:.5f} per site-equivalent from "
          f"{obs_d['synonymous']:.0f} synonymous substitutions.")
    joint = {}
    for cls, counts, keys in classes:
        k = sum(obs_d[x] for x in keys)
        opp = sum(opp_d[x] for x in keys)
        f = fit_gamma(binned(counts, bins), r_full, H, S_edges, theta_of(keys),
                      div=(k, opp, d_syn), S_mid=S_mid)
        joint[cls] = f
        show(cls, len(counts), f)

    # ---- model-free discrete DFE --------------------------------------------
    # A sample of 56,433 resolves selection up to S ~ n: even S=100 variants
    # still segregate detectably as singletons, so the intervals must extend
    # well beyond 100 before the DFE becomes genuinely invisible.
    edges_g = [0.0, 1.0, 10.0, 1e2, 1e3, 1e4, 1e5, 1e9]
    groups = list(zip(edges_g[:-1], edges_g[1:]))
    Hg = np.array([H[(S_mid >= a) & (S_mid < b)].mean(axis=0) for a, b in groups])
    labels = ["S<1", "1-10", "10-1e2", "1e2-1e3", "1e3-1e4", "1e4-1e5", ">1e5"]
    print("\nDISCRETE DFE (free mass per interval, no gamma assumption):")
    print(f"{'class':<22}" + "".join(f"{l:>10}" for l in labels))
    for cls, counts, keys in classes:
        p, _ = fit_discrete(binned(counts, bins), r_full, Hg, theta_of(keys))
        print(f"{cls:<22}" + "".join(f"{v:>10.3f}" for v in p))

    # ---- goodness of fit ----------------------------------------------------
    print("\nGOODNESS OF FIT, nonsynonymous (observed vs gamma+divergence fit):")
    k = obs_d["nonsynonymous"]
    f = fit_gamma(binned(variants["nonsynonymous"], bins), r_full, H, S_edges,
                  theta_of(("nonsynonymous",)),
                  div=(k, opp_d["nonsynonymous"], d_syn), S_mid=S_mid)
    w = gamma_weights(S_edges, f["beta"], f["Sbar"])
    e_b = (w @ H) * r_full * theta_of(("nonsynonymous",))
    o_b = binned(variants["nonsynonymous"], bins)
    print("  MAC bin  " + " ".join(f"{a:>6d}" for a, _ in bins))
    print("  observed " + " ".join(f"{v:>6.0f}" for v in o_b))
    print("  fitted   " + " ".join(f"{v:>6.0f}" for v in e_b))
    chi2 = float(np.sum((o_b - e_b) ** 2 / np.maximum(e_b, 1e-9)))
    print(f"  chi-square = {chi2:.1f} on ~{len(bins) - 2} df")

    # ---- bootstrap on the joint fit -----------------------------------------
    if args.boot:
        print(f"\nbootstrapping the joint fit ({args.boot} replicates, warm "
              f"started at the point estimate)...")
        print(f"{'class':<22}{'beta 95% CI':>20}{'P(1<S<10) 95% CI':>24}"
              f"{'nn mass / nonsyn':>19}")
        draws = {}
        for cls, counts, keys in classes:
            k = sum(obs_d[x] for x in keys)
            opp = sum(opp_d[x] for x in keys)
            th = theta_of(keys)
            warm = (joint[cls]["beta"], joint[cls]["Sbar"])
            bs = []
            for _ in range(args.boot):
                samp = rng.choice(counts, size=len(counts), replace=True)
                f = fit_gamma(binned(samp, bins), r_full, H, S_edges, th,
                              div=(rng.poisson(k), opp, d_syn), S_mid=S_mid,
                              warm=warm)
                bs.append((f["beta"],
                           near_neutral_mass(f["beta"], f["Sbar"], 1.0, 10.0)))
            draws[cls] = np.array(bs)

        for cls, _, _ in classes:
            bs = draws[cls]
            bq = np.percentile(bs[:, 0], [2.5, 97.5])
            nq = np.percentile(bs[:, 1], [2.5, 97.5])
            ratio = bs[:, 1] / np.maximum(draws["nonsynonymous"][:, 1], 1e-12)
            rq = np.percentile(ratio, [2.5, 50, 97.5])
            print(f"{cls:<22}{f'[{bq[0]:.2f}, {bq[1]:.2f}]':>20}"
                  f"{f'[{nq[0]:.3f}, {nq[1]:.3f}]':>24}"
                  f"{f'{rq[1]:.2f} [{rq[0]:.2f},{rq[2]:.2f}]':>19}")
        print("\nThe last column is the quantity the write-up actually needs:")
        print("near-neutral mass in each class RELATIVE to nonsynonymous "
              "coding.")
        print("It is far better determined than the absolute DFE, because all")
        print("classes share the same demography, haplogroup structure and")
        print("linkage, so those errors largely cancel in the ratio.")


    print("\nsbar = Sbar / (2*Ne) at Ne = 7,500 (mtDNA female Ne anchor).")
    print("P(1<S<10) is the near-neutral mass that drives Muller's ratchet:")
    print("mutations drift-dominated enough to fix, yet costly enough to matter.")
    print("\nPublished control: Piganeau & Eyre-Walker (2003) fitted the mtDNA")
    print("protein-coding DFE and obtained beta ~ 0.4; this project has been")
    print("using beta = 0.39, Sbar = 1400 anchored at Ne = 7,500. The")
    print("nonsynonymous row above is the like-for-like comparison.")


if __name__ == "__main__":
    main()
```
