## Supplementary Information section 1 code base for "Hypothesis: A modern human range expansion ∼300,000 years ago had a critical role in Neandertal origins": pew_replicate.py

```
"""
Faithful replication of Piganeau & Eyre-Walker (2003) PNAS 100:10335, model 2
(gamma distribution of fitness effects, all nonsynonymous mutations deleterious).

The method is a Poisson / MK-style estimator on four counts per data set:
    Pn, Ps   nonsynonymous / synonymous POLYMORPHISMS (segregating sites)
    Dn, Ds   nonsynonymous / synonymous SUBSTITUTIONS (multiple-hit corrected)
plus the sample size n and the site proportions rho_n, rho_s.

Expected counts (P&EW eqs 1-5), with S the scaled selection coefficient
(S = 2*Ne*s for a haploid such as mtDNA; deleterious mutations have S<0 and the
gamma is placed on the magnitude s = |S| >= 0):

    Ps_hat = L rho_s theta * a_n,                 a_n = sum_{i=1}^{n-1} 1/i
    Pn_hat = L rho_n theta * INT g(s) Q(s,n) ds
    Ds_hat = L rho_s lambda
    Dn_hat = L rho_n lambda * INT g(s) F(s) ds

    Q(s,n) = INT_0^1 H(s,x) (1 - x^n - (1-x)^n) dx,   sojourn time of a
             deleterious mutation weighted by the chance it segregates in n draws
    H(s,x) = (e^{s(1-x)} - 1) / [ (e^s - 1) x (1-x) ]   (Wright 1938; note H->1/x
             as s->0, recovering Watterson's a_n for the neutral class)
    F(s)   = s / (e^s - 1)   = 2Ne * (relative fixation prob of a deleterious S)
    g(s)   = alpha^beta s^{beta-1} e^{-alpha s} / Gamma(beta),  mean s = beta/alpha

Because there are four observations and four parameters (theta, lambda, alpha,
beta) the model fits exactly (their "perfect fit"). Fixing theta and lambda from
the synonymous class reduces this to two equations in (alpha, beta):

    I_P(alpha,beta) = Pn * rho_s * a_n / (rho_n * Ps)
    I_D(alpha,beta) = Dn * rho_s        / (rho_n * Ds)      ( = dN/dS )

which we solve by least squares on the logs. Running this on P&EW's own human
counts should return their published beta = 0.39, alpha = 2.7e-4, Sbar = -1400.
"""
import math

import numpy as np
from scipy.optimize import minimize
from scipy.special import gammainc


# ---------------------------------------------------------------- PRF integrals
def _Q_of_s(s, n, x):
    """Q(s,n) integrand-integrated on grid x: sojourn time of a deleterious
    mutation of scaled strength s (>=0) times the probability it is polymorphic
    in a sample of n. H(s,x) = (e^{s(1-x)}-1)/((e^s-1) x (1-x))."""
    poly = 1.0 - x**n - (1.0 - x)**n           # P(segregating | freq x, n draws)
    if s < 1e-9:
        H = 1.0 / x                            # neutral limit -> Watterson
    else:
        # (e^{s(1-x)}-1)/(e^s-1) = (e^{-sx}-e^{-s})/(1-e^{-s}); stable for all s>0
        H = (np.exp(-s * x) - math.exp(-s)) / (-math.expm1(-s) * x * (1.0 - x))
    return np.trapezoid(H * poly, x)


def build_grids(n, S_grid, n_x=20000):
    """Q[k] = Q(S_grid[k], n) and Fdiv[k] = F(S_grid[k]); precomputed once per n.
    x avoids the exact endpoints where 1/(x(1-x)) is singular (the numerator
    vanishes there, so the product is finite, but the grid must not divide by 0)."""
    x = np.linspace(0.5 / n_x, 1.0 - 0.5 / n_x, n_x)
    Q = np.array([_Q_of_s(s, n, x) for s in S_grid])
    # F(s) = s/(e^s-1); for large s this is s*e^{-s} -> 0 (stable, no overflow)
    F = np.where(S_grid < 1e-9, 1.0,
                 np.where(S_grid < 700.0,
                          S_grid / np.expm1(np.minimum(S_grid, 700.0)),
                          S_grid * np.exp(-np.minimum(S_grid, 700.0))))
    return Q, F


def gamma_weights(S_edges, beta, Sbar):
    """Mass of Gamma(shape=beta, mean=Sbar) in each grid cell defined by edges."""
    scale = Sbar / beta
    cdf = gammainc(beta, S_edges / scale)      # regularised lower incomplete
    w = np.diff(cdf)
    tot = w.sum()
    return w / tot if tot > 0 else w


# --------------------------------------------------------------------- estimator
def a_n(n):
    return float(np.sum(1.0 / np.arange(1, n)))


def fit_model2(n, rho_n, Dn, Ds, Pn, Ps, S_edges, S_mid, Q, F, verbose=False):
    """Return (alpha, beta, Sbar) solving the two P&EW moment equations."""
    rho_s = 1.0 - rho_n
    an = a_n(n)
    IP_target = Pn * rho_s * an / (rho_n * Ps)     # nonsyn polymorphism / neutral
    ID_target = Dn * rho_s / (rho_n * Ds)          # = dN/dS

    def resid(par):
        logbeta, logSbar = par
        beta, Sbar = np.exp(logbeta), np.exp(logSbar)
        if not (1e-3 < beta < 50 and 1e-3 < Sbar < 1e8):
            return 1e6
        w = gamma_weights(S_edges, beta, Sbar)
        if not np.isfinite(w).all() or w.sum() <= 0:
            return 1e6
        IP = float(w @ Q)
        ID = float(w @ F)
        if IP <= 0 or ID <= 0:
            return 1e6
        return (np.log(IP) - np.log(IP_target))**2 + \
               (np.log(ID) - np.log(ID_target))**2

    best, bx = np.inf, None
    for b0 in (0.2, 0.4, 0.6, 1.0, 2.0):
        for S0 in (1e2, 1e3, 1e4, 1e5):
            r = minimize(resid, [np.log(b0), np.log(S0)], method="Nelder-Mead",
                         options={"xatol": 1e-8, "fatol": 1e-12, "maxiter": 20000})
            if r.fun < best:
                best, bx = r.fun, r.x
    beta, Sbar = np.exp(bx[0]), np.exp(bx[1])
    alpha = beta / Sbar
    if verbose:
        w = gamma_weights(S_edges, beta, Sbar)
        print(f"    target I_P={IP_target:.4f} I_D={ID_target:.5f}; "
              f"fit I_P={float(w@Q):.4f} I_D={float(w@F):.5f}; resid={best:.2e}")
    return alpha, beta, Sbar


# P&EW Table 1 counts and Table 3 model-2 published estimates -------------------
# columns: n, L, rho_n, Dn, Ds, Pn, Ps ; then published alpha, beta, Sbar
PEW = {
    "AL": (10, 306, 0.71, 3, 90, 1, 4, 1.1e-2, 1.1, -100),
    "BM": (14, 1041, 0.75, 1, 118, 1, 13, 9.2e-4, 0.91, -990),
    "DM": (59, 1515, 0.90, 19, 157, 11, 17, 3.5e-4, 0.58, -1700),
    "DP": (22, 1399, 0.87, 9, 174, 2, 19, 3.7e-7, 0.34, -920000),
    "FC": (15, 1283, 0.83, 11, 96, 13, 63, 3.7e-6, 0.31, -84000),
    "GA": (9, 1143, 0.78, 3, 41, 7, 10, 6.3e-2, 1.6, -25),
    "GM": (41, 300, 0.66, 0, 20, 3, 22, 3.3e-2, 4.7, -140),
    "HS": (53, 11259, 0.68, 186, 1765, 91, 320, 2.7e-4, 0.39, -1400),
    "IB": (10, 798, 0.71, 15, 150, 15, 103, 4.5e-7, 0.23, -510000),
    "MA": (10, 1143, 0.72, 12, 80, 13, 20, 3.7e-2, 1.0, -27),
    "MD": (56, 342, 0.67, 3, 57, 11, 13, 1.5e-1, 2.2, -15),
    "MH": (29, 798, 0.69, 7, 120, 30, 118, 1.6e-3, 0.62, -390),
    "PI": (19, 431, 0.78, 12, 186, 5, 10, 1.1e-2, 0.99, -90),
    "PL": (8, 292, 0.71, 7, 156, 11, 59, 3.0e-3, 0.75, -250),
    "SA": (20, 1140, 0.68, 26, 487, 12, 38, 7.8e-3, 0.85, -110),
    "UA": (166, 1140, 0.64, 23, 167, 11, 44, 2.8e-6, 0.21, -75000),
}


def main():
    S_edges = np.concatenate([[0.0], np.logspace(-4, 7.5, 600)])
    S_mid = np.sqrt(np.maximum(S_edges[:-1], 1e-6) * np.maximum(S_edges[1:], 1e-6))
    S_mid[0] = 0.0

    print("Replication of Piganeau & Eyre-Walker (2003), Table 3 model 2")
    print("(gamma DFE on nonsynonymous mutations, all deleterious)\n")
    print(f"{'set':<5}{'beta_fit':>9}{'beta_pub':>9}"
          f"{'Sbar_fit':>12}{'Sbar_pub':>12}{'alpha_fit':>11}{'alpha_pub':>11}")

    Qcache = {}
    for name, v in PEW.items():
        n, L, rho_n, Dn, Ds, Pn, Ps, a_pub, b_pub, S_pub = v
        if Pn == 0 or Dn == 0:                 # GM/PT-type: model degenerate
            print(f"{name:<5}{'--':>9}{b_pub:>9}   (excess-substitution / "
                  f"zero-count set, model 2 degenerate)")
            continue
        if n not in Qcache:
            Qcache[n] = build_grids(n, S_mid)
        Q, F = Qcache[n]
        alpha, beta, Sbar = fit_model2(n, rho_n, Dn, Ds, Pn, Ps,
                                       S_edges, S_mid, Q, F)
        print(f"{name:<5}{beta:>9.2f}{b_pub:>9.2f}{-Sbar:>12.0f}{S_pub:>12.0f}"
              f"{alpha:>11.1e}{a_pub:>11.1e}")


if __name__ == "__main__":
    main()
```
