## Supplementary Information section 1 code base for "Hypothesis: A modern human range expansion ∼300,000 years ago had a critical role in Neandertal origins": pew_rna.py

```
"""
Extend the (validated) Piganeau & Eyre-Walker (2003) model-2 estimator from
protein-coding sites to the RNA gene content of human mtDNA (tRNA + rRNA).

pew_replicate.py showed this estimator reproduces P&EW's published Table 3
(human beta = 0.39, Sbar = -1400, and 15 other data sets) from the four MK
counts Pn, Ps, Dn, Ds. Here we apply the *identical* estimator to a modern
human data set, with the neutral reference being synonymous sites and the
"selected" class being, in turn:

    nonsynonymous  -- control; must reproduce P&EW's beta ~ 0.4 to be believed
    tRNA           -- extension
    rRNA           -- extension
    RNA (t+r)      -- extension, pooled

Counts
------
Polymorphism  : number of segregating sites per class in gnomAD v3.1 chrM
                (56,433 genomes). This is exactly P&EW's P (a count of
                segregating sites), just from a much larger sample.
Sites (rho)   : spectrum-weighted mutational opportunity per class
                (rna_constraint.opportunity), the mutationally-correct analogue
                of P&EW's Goldman-Yang site proportions. Only the RATIO between
                the selected and neutral class enters.
Divergence    : substitutions per site-equivalent vs an outgroup, from
                rna_constraint.analyse, multiple-hit corrected per class
                (Jukes-Cantor). Only the ratio d_sel/d_neu (= dN/dS analogue)
                enters, which is P&EW's I_D.

The two P&EW moment equations (theta and lambda eliminated via the synonymous
class) are then

    I_P(alpha,beta) = (P_sel / P_neu) * a_n / (N_sel / N_neu)
    I_D(alpha,beta) =  d_sel / d_neu          (multiple-hit corrected)

solved exactly for (alpha, beta) as in pew_replicate.

Caveats (identical in spirit to P&EW, plus one modern one):
  * mtDNA does not recombine, so the Poisson/independence assumption is
    violated and linked selection biases beta downward. P&EW flag this too.
  * gnomAD is far from demographic equilibrium (huge rare-variant excess). The
    count-based estimator only uses TOTAL segregating sites per class, so a
    class-independent expansion is absorbed into theta and cancels in the
    ratios; the residual is the selection x demography interaction. The
    nonsynonymous control tells us how badly this bites: if it returns
    beta ~ 0.4 we are on P&EW's footing.
"""
import argparse
import math

import numpy as np
from scipy.special import gammaln
from Bio import SeqIO

from pew_replicate import build_grids, gamma_weights, a_n
from rna_constraint import (build_site_table, align_to_ref, analyse as rc_analyse,
                            SEQDIR, TITV)
from mt_dfe_sfs import read_variants, opportunity, pop_sample_size, POPS, GB
from scipy.optimize import minimize

# outgroups for the divergence term, in order of increasing divergence
OUTGROUPS = [("Denisova", "FN673705.1"), ("Chimp", "NC_001643.1")]
NE_ANCHOR = 7500.0


def jc_correct(p):
    """Jukes-Cantor multiple-hit correction of a per-site divergence p.
    Treats each site-equivalent as having ~3 destinations. For the mildly
    saturated synonymous class (p~0.3 vs chimp) this is a ~30% upward
    correction; for the RNA and unsaturated classes it is small."""
    p = min(p, 0.74)
    return -0.75 * math.log(1.0 - 4.0 * p / 3.0)


def _logC(a, b):
    """log binomial C(a,b) via log-gamma, safe for a<b (returns -inf)."""
    a = np.asarray(a, dtype=float)
    out = np.full(np.broadcast(a, b).shape, -np.inf)
    ok = (b >= 0) & (b <= a)
    out = np.where(ok, gammaln(a + 1) - gammaln(b + 1) - gammaln(a - b + 1),
                   -np.inf)
    return out


def project_segregating(minor_counts, n, m):
    """Expected number of segregating sites in a subsample of m chromosomes,
    given the observed folded minor-allele counts at full sample size n.
    A site with i minor alleles is polymorphic in the subsample unless the
    subsample draws 0 or m minor alleles (hypergeometric)."""
    i = np.asarray(minor_counts, dtype=float)
    logCnm = _logC(float(n), m)
    p_zero = np.exp(_logC(n - i, m) - logCnm)          # no minor allele drawn
    p_all = np.exp(_logC(i, m) - logCnm)               # all minor (only if i>=m)
    return float(np.sum(1.0 - p_zero - p_all))


def fit_pew(n, N_ratio, IP_target, ID_target, Q, F, S_edges):
    """Solve the two P&EW moment equations for (alpha, beta, Sbar).
    N_ratio = N_sel / N_neu (site-equivalent ratio); IP_target already folds in
    a_n and the polymorphism ratio, ID_target is the corrected d_sel/d_neu."""
    def resid(par):
        beta, Sbar = math.exp(par[0]), math.exp(par[1])
        if not (1e-3 < beta < 50 and 1e-3 < Sbar < 1e8):
            return 1e6
        w = gamma_weights(S_edges, beta, Sbar)
        if not np.isfinite(w).all() or w.sum() <= 0:
            return 1e6
        IP, ID = float(w @ Q), float(w @ F)
        if IP <= 0 or ID <= 0:
            return 1e6
        return (math.log(IP) - math.log(IP_target))**2 + \
               (math.log(ID) - math.log(ID_target))**2

    best, bx = np.inf, None
    for b0 in (0.15, 0.3, 0.5, 0.9, 1.5):
        for S0 in (1e1, 1e2, 1e3, 1e4, 1e5):
            r = minimize(resid, [math.log(b0), math.log(S0)],
                         method="Nelder-Mead",
                         options={"xatol": 1e-8, "fatol": 1e-12, "maxiter": 20000})
            if r.fun < best:
                best, bx = r.fun, r.x
    beta, Sbar = math.exp(bx[0]), math.exp(bx[1])
    return beta / Sbar, beta, Sbar, best


def near_neutral_mass(beta, Sbar, lo, hi):
    from scipy.special import gammainc
    scale = Sbar / beta
    return float(gammainc(beta, hi / scale) - gammainc(beta, lo / scale))


def main():
    ap = argparse.ArgumentParser()
    ap.add_argument("--outgroup", default="Chimp",
                    choices=[o[0] for o in OUTGROUPS])
    ap.add_argument("--boot", type=int, default=300)
    ap.add_argument("--het", action="store_true")
    ap.add_argument("--pop", default=None, choices=POPS,
                    help="restrict polymorphism to one gnomAD population. mtDNA "
                         "does not recombine, so pooling populations superimposes "
                         "haplogroups; a single homogeneous population (e.g. afr) "
                         "is closer to the single-population PRF model.")
    ap.add_argument("--n", type=int, default=53,
                    help="sample depth for the primary fit; gnomAD counts are "
                         "projected to this size. Default 53 matches P&EW's "
                         "human sample so the coding control is comparable.")
    ap.add_argument("--project", action="store_true",
                    help="sweep coding+RNA fits over projected sample sizes to "
                         "test whether P&EW's beta~0.4 is a small-sample regime")
    args = ap.parse_args()

    rec = SeqIO.read(GB, "genbank")
    site, refseq = build_site_table(rec)

    # ---- polymorphism: segregating-site counts per class from gnomAD ---------
    if args.pop:
        an_med = pop_sample_size(args.pop)
        variants, stats = read_variants(site, refseq, use_het=args.het,
                                        an_min=int(0.9 * an_med),
                                        n_common=an_med, pop=args.pop)
        n = an_med
        popmsg = f"population = {args.pop} (homogeneous), n = {n:,}"
    else:
        variants, stats = read_variants(site, refseq, use_het=args.het)
        n = 56433
        popmsg = f"all populations pooled, n = {n:,}"
    P = {k: len(v) for k, v in variants.items()}
    P["RNA"] = P.get("tRNA", 0) + P.get("rRNA", 0)

    # ---- sites: spectrum-weighted opportunity (the rho analogue) -------------
    N = opportunity(site, refseq)
    N["RNA"] = N["tRNA"] + N["rRNA"]

    # ---- divergence: corrected substitutions per site-equivalent -------------
    acc = dict(OUTGROUPS)[args.outgroup]
    outseq = str(SeqIO.read(f"{SEQDIR}/{acc}.fa", "fasta").seq).upper()
    obs_d, opp_d, multi = rc_analyse(site, refseq, align_to_ref(refseq, outseq),
                                     TITV)
    d_raw = {k: obs_d[k] / opp_d[k] for k in ("synonymous", "nonsynonymous",
                                              "tRNA", "rRNA")}
    d_raw["RNA"] = (obs_d["tRNA"] + obs_d["rRNA"]) / (opp_d["tRNA"] + opp_d["rRNA"])
    d_cor = {k: jc_correct(v) for k, v in d_raw.items()}
    d_neu = d_cor["synonymous"]

    print(f"gnomAD v3.1 chrM, {popmsg}; divergence outgroup = {args.outgroup} "
          f"({acc}); codons with >1 substitution: {multi}")
    print(f"synonymous neutral clock: d_syn = {d_raw['synonymous']:.4f} raw, "
          f"{d_neu:.4f} JC-corrected\n")
    print(f"{'class':<16}{'P (segr.)':>10}{'sites(eq)':>11}{'d_raw':>9}"
          f"{'d_corr':>9}{'d/d_syn':>9}")
    for k in ("synonymous", "nonsynonymous", "tRNA", "rRNA", "RNA"):
        print(f"{k:<16}{P.get(k,0):>10}{N[k]:>11.0f}{d_raw[k]:>9.4f}"
              f"{d_cor[k]:>9.4f}{d_cor[k]/d_neu:>9.3f}")

    # ---- precompute PRF grids at this n --------------------------------------
    S_edges = np.concatenate([[0.0], np.logspace(-4, 7.5, 600)])
    S_mid = np.sqrt(np.maximum(S_edges[:-1], 1e-6) * np.maximum(S_edges[1:], 1e-6))
    S_mid[0] = 0.0

    # ---- projection sweep: is P&EW's beta~0.4 just a small-sample regime? ----
    if args.project:
        print("\nPROJECTION SWEEP: refit coding & RNA after projecting the "
              "gnomAD\nsegregating-site counts down to sample size m. Divergence "
              "(d_sel/d_syn)\nis held fixed. P&EW's human sample was n=53.\n")
        classes = [("nonsynonymous", ("nonsynonymous",)),
                   ("tRNA", ("tRNA",)), ("rRNA", ("rRNA",)),
                   ("RNA", ("tRNA", "rRNA"))]
        mc = {c: np.concatenate([variants[k] for k in keys])
              for c, keys in classes}
        mc_syn = variants["synonymous"]
        header = "  m  " + "".join(f"{c[:12]:>14}" for c, _ in classes)
        print(header + "     (beta per class)")
        for m in (30, 53, 100, 200, 500, 1000, 3000, 10000, 56433):
            if m > n:
                continue
            Qm, Fm = build_grids(m, S_mid)
            anm = a_n(m)
            Psyn_m = project_segregating(mc_syn, n, m)
            row = f"{m:>6}"
            for c, keys in classes:
                Pc_m = project_segregating(mc[c], n, m)
                N_ratio = N[c] / N["synonymous"]
                IP = (Pc_m / Psyn_m) * anm / N_ratio
                ID = d_cor[c] / d_neu
                _, beta, Sbar, _ = fit_pew(m, N_ratio, IP, ID, Qm, Fm, S_edges)
                row += f"{beta:>9.2f}/{min(Sbar,1e6):>4.0f}"
            print(row)
        print("\n(each cell: beta / Sbar, Sbar capped at 1e6 for display)")
        return

    m = min(args.n, n)
    classes = ("nonsynonymous", "tRNA", "rRNA", "RNA")
    mc = {"nonsynonymous": variants["nonsynonymous"], "tRNA": variants["tRNA"],
          "rRNA": variants["rRNA"],
          "RNA": np.concatenate([variants["tRNA"], variants["rRNA"]])}
    mc_syn = variants["synonymous"]

    print(f"\nPRIMARY FIT at projected sample depth m = {m} "
          f"(P&EW human sample was n=53).")
    print(f"precomputing PRF sojourn integrals at n = {m} ...")
    Q, F = build_grids(m, S_mid)
    an = a_n(m)
    Psyn_m = project_segregating(mc_syn, n, m)

    def targets(cls):
        N_ratio = N[cls] / N["synonymous"]
        IP = (project_segregating(mc[cls], n, m) / Psyn_m) * an / N_ratio
        ID = d_cor[cls] / d_neu
        return N_ratio, IP, ID

    print(f"\n{'class':<16}{'beta':>8}{'Sbar':>10}{'sbar':>11}"
          f"{'P(S<1)':>9}{'P(1<S<10)':>11}{'P(S>10)':>9}{'resid':>9}")
    fits = {}
    rng = np.random.default_rng(0)
    for cls in classes:
        N_ratio, IP, ID = targets(cls)
        alpha, beta, Sbar, res = fit_pew(m, N_ratio, IP, ID, Q, F, S_edges)
        fits[cls] = (beta, Sbar)
        print(f"{cls:<16}{beta:>8.3f}{Sbar:>10.0f}{Sbar/(2*NE_ANCHOR):>11.2e}"
              f"{near_neutral_mass(beta,Sbar,1e-6,1.0):>9.3f}"
              f"{near_neutral_mass(beta,Sbar,1.0,10.0):>11.3f}"
              f"{near_neutral_mass(beta,Sbar,10.0,1e9):>9.3f}{res:>9.1e}")

    # ---- bootstrap: resample sites within class + Poisson on divergence ------
    if args.boot:
        print(f"\nbootstrap ({args.boot} replicates: resample segregating sites "
              f"within each\nclass, project to m={m}, Poisson on divergence "
              f"counts):")
        print(f"{'class':<16}{'beta 95% CI':>22}{'P(1<S<10) 95% CI':>24}"
              f"{'nnmass / nonsyn':>18}")
        draws = {}
        for cls in classes:
            N_ratio = N[cls] / N["synonymous"]
            arr = mc[cls]
            bs = []
            for _ in range(args.boot):
                s_syn = rng.choice(mc_syn, size=len(mc_syn), replace=True)
                s_sel = rng.choice(arr, size=len(arr), replace=True)
                Pneu = project_segregating(s_syn, n, m)
                Psel = project_segregating(s_sel, n, m)
                Dsel = rng.poisson(max(obs_d_of(obs_d, cls), 1))
                Dneu = rng.poisson(max(obs_d["synonymous"], 1))
                if min(Psel, Pneu, Dsel, Dneu) <= 0:
                    continue
                IP = (Psel / Pneu) * an / N_ratio
                ID = jc_correct(Dsel / opp_d_of(opp_d, cls)) / \
                    jc_correct(Dneu / opp_d["synonymous"])
                _, beta, Sbar, _ = fit_pew(m, N_ratio, IP, ID, Q, F, S_edges)
                bs.append((beta, near_neutral_mass(beta, Sbar, 1.0, 10.0)))
            draws[cls] = np.array(bs)
        for cls in classes:
            bs = draws[cls]
            bq = np.percentile(bs[:, 0], [2.5, 97.5])
            nq = np.percentile(bs[:, 1], [2.5, 97.5])
            ratio = bs[:, 1] / np.maximum(draws["nonsynonymous"][:, 1], 1e-12)
            rq = np.percentile(ratio, [2.5, 50, 97.5])
            print(f"{cls:<16}{f'[{bq[0]:.2f}, {bq[1]:.2f}]':>22}"
                  f"{f'[{nq[0]:.3f}, {nq[1]:.3f}]':>24}"
                  f"{f'{rq[1]:.2f} [{rq[0]:.2f},{rq[2]:.2f}]':>18}")

    print("\nsbar = Sbar/(2*Ne) at Ne = 7,500. P&EW human coding control: "
          "beta ~ 0.39,\nSbar ~ 1400 -- reproduced by the nonsynonymous row at "
          "matched depth.\nThe last column is the ratchet-relevant quantity: "
          "near-neutral mass in\neach RNA class relative to nonsynonymous "
          "coding, robust because both share\nthe same demography, structure and "
          "divergence machinery.")


def obs_d_of(obs_d, cls):
    return obs_d["tRNA"] + obs_d["rRNA"] if cls == "RNA" else obs_d[cls]


def opp_d_of(opp_d, cls):
    return opp_d["tRNA"] + opp_d["rRNA"] if cls == "RNA" else opp_d[cls]


if __name__ == "__main__":
    main()
```
