## Supplementary Information section 1 code base for "Hypothesis: A modern human range expansion ∼300,000 years ago had a critical role in Neandertal origins": plot_abc_fig2.py

```
"""
Figure 2 top (mtDNA), redesigned: posterior histograms of the modern-archaic
fitness gap from the introgression ABC, with all sources of uncertainty
integrated (U, beta, |Sbar|, archaic size with equal prior on 1500/3000).

Layout: five panels. Every panel repeats the FULL RNA-inclusive ABC-accepted
histogram in black; colors show the conditional histograms obtained by slicing
one parameter while integrating over everything else:
  (a) U     at ~5th pct / mean / ~95th pct   (slices = 0-10 / 45-55 / 90-100 pct)
  (b) beta  at ~5th pct / mean / ~95th pct
  (c) |Sbar| at ~5th pct / mean / ~95th pct
  (d) N_archaic = 1500 vs 3000
  (e) model variants: coding-only (no ABC), coding-only (ABC-accepted),
      coding+RNA (no ABC); the black histogram is coding+RNA (ABC-accepted).

Real data (--real, default once files exist):
  joint_rna_{1500,3000}.npz  : RNA-inclusive pairs (abc_joint_rna.py)
  joint_cod_{1500,3000}.npz  : coding-only pairs   (abc_joint_rna.py --coding-only)
  Acceptance: nonsynonymous ratio in [1.16, 1.26] (combine_joint.py).
  Per-pair U slice uses the realized synonymous branch counts (m_syn + d_syn),
  the observable that calibrates the rate (the drawn u_syn itself is not stored).

Mock mode (--mock): synthetic draws with the response surface calibrated to the
O2 grid (central gaps 0.67% / 0.29%, Sbar and U scaling from results_denis300)
and (beta, |Sbar|) resampled from the real bootstrap (dfe_draws.npz), purely to
preview the layout.
"""
import argparse
import os

import numpy as np

os.environ.setdefault("MPLCONFIGDIR", os.path.abspath(".mplcache"))
os.makedirs(os.environ["MPLCONFIGDIR"], exist_ok=True)

import matplotlib
matplotlib.use("Agg")
import matplotlib.pyplot as plt

RATIO_LO, RATIO_HI = 1.16, 1.26
SWEEP_BAND = (0.5, 5.0)               # % gap range where sweep probability rises
COL_MAIN = "black"
# figure-label notation follows the note: accumulation-phase archaic size
# NA-uniparental, post-mixture Neandertal size NN-uniparental
NA_LAB = "$N_{A\\mathrm{-uniparental}}$"
NN_LAB = "$N_{N\\mathrm{-uniparental}}$"
COL_LMH = ["#2c6fbb", "#7f7f7f", "#c0392b"]      # low / mean / high slices
COL_NA = {1500: "#c0392b", 2250: "#7570b3", 3000: "#2c6fbb"}
COL_E = ["#7f7f7f", "#1b9e77", "#e08214"]        # cod, cod+ABC, rna no-ABC
XLIM = (-1.0, 2.5)
BINS = np.linspace(XLIM[0], XLIM[1], 71)

# ---------------------------------------------------------------------------
# Fixation-probability curves from the companion analysis
# (fixation_curves_mtdna.csv): percent of replicates in which all 10 sampled
# individuals carry the modern type ~260 ka after a 5% sex-unbiased pulse, vs
# the additive coefficient s of the modern mtDNA, by locus_Ne (number of mtDNA
# copies = census/2). Selection: locus_Ne = 750 is the post-mixture Neandertal
# size and is paired with the NA = 2250 accumulation posterior (the sweep
# happens after the contraction); 500 is the Petr comparison; 1500/3000 are a
# no-contraction sensitivity matching the accumulation-phase brackets.
FIXCURVE = "fixation_curves_mtdna.csv"


def load_fixcurves(path=FIXCURVE):
    if not os.path.exists(path):
        return None
    import csv
    curves = {}
    with open(path) as f:
        rows = csv.DictReader(r for r in f if not r.startswith("#"))
        for r in rows:
            curves.setdefault(int(float(r["locus_Ne"])), []).append(
                (float(r["s"]), float(r["pfix_pct"])))
    return {ne: np.array(sorted(pts)) for ne, pts in curves.items()}


CURVES = load_fixcurves()


def pfix(gap_pct, locus_ne):
    """P(fixation) in % for a fitness gap (in %), interpolating the
    companion-analysis curve at the given locus_Ne. The curves are simulated
    down to s = -0.005 (archaic favored) and up to s = 0.03; values outside
    clamp to the endpoints (the s = 0.03 end is therefore a lower bound)."""
    s = np.asarray(gap_pct, float) / 100.0
    c = CURVES[locus_ne]
    return np.interp(s, c[:, 0], c[:, 1])


# each accumulation-phase size NA is paired with its contracted post-mixture
# size NN (same ~1/3 ratio as the central 2250 -> 750)
PAIR_NN = {1500: 500, 2250: 750, 3000: 1000}


def pfix_mix(gaps, nas):
    """Integrated P(fix) in % for a set of draws, each evaluated on the
    fixation curve of its own NA-paired post-mixture size."""
    gaps, nas = np.asarray(gaps, float), np.asarray(nas)
    tot = 0.0
    for na, nn in PAIR_NN.items():
        m = nas == na
        if np.any(m):
            tot += pfix(gaps[m], nn).sum()
    return tot / gaps.size


# --------------------------------------------------------------------------
# data ingestion: returns a dict with per-pair arrays for the RNA-inclusive run
# (gap %, U-proxy, beta, Sbar, NA, accepted) and gaps for the panel-e variants.
# --------------------------------------------------------------------------
def load_real():
    def accept(d):
        R = d["m_ns"] / np.where(d["d_ns"] == 0, np.nan, d["d_ns"])
        return np.isfinite(R) & (R >= RATIO_LO) & (R <= RATIO_HI)

    def one(prefix, nas=(1500, 2250, 3000)):
        gs, us, bs, ss, nass, acc = [], [], [], [], [], []
        for na in nas:
            fn = f"{prefix}_{na}.npz"
            if not os.path.exists(fn):
                raise SystemExit(f"missing {fn} (pull from O2 first)")
            d = np.load(fn)
            gap = (d["m_w"] - d["d_w"]) * 100.0
            gs.append(gap)
            us.append(d["m_syn"] + d["d_syn"])    # realized syn counts ~ rate
            bs.append(d["beta"]); ss.append(d["Sbar"])
            nass.append(np.full(gap.size, na)); acc.append(accept(d))
        a = np.concatenate(acc)
        print(f"{prefix}: accepted {a.sum()}/{a.size} "
              f"({100 * a.mean():.1f}%)  [Methods acceptance fraction]")
        return {k: np.concatenate(v) for k, v in
                zip(("gap", "U", "beta", "Sbar", "NA", "acc"),
                    (gs, us, bs, ss, nass, acc))}

    rna = one("joint_rna")
    out = dict(
        main=rna["gap"][rna["acc"]],
        U=rna["U"][rna["acc"]], beta=rna["beta"][rna["acc"]],
        Sbar=rna["Sbar"][rna["acc"]], NA=rna["NA"][rna["acc"]],
        rna_noabc=rna["gap"], NA_rna_all=rna["NA"], mock=False)
    try:
        cod = one("joint_cod")
        out["cod_noabc"] = cod["gap"]
        out["NA_cod_all"] = cod["NA"]
        out["cod_abc"] = cod["gap"][cod["acc"]]
        out["NA_cod_acc"] = cod["NA"][cod["acc"]]
    except SystemExit:
        print("joint_cod_*.npz not found; panel (e) coding variants omitted")
    if os.path.exists("joint_rna_petr500.npz"):
        d = np.load("joint_rna_petr500.npz")
        gap = (d["m_w"] - d["d_w"]) * 100.0
        a = accept(d)
        print(f"joint_rna_petr500: accepted {a.sum()}/{a.size} "
              f"({100 * a.mean():.1f}%)  [Methods acceptance fraction]")
        out["petr"] = gap[a]
    else:
        print("joint_rna_petr500.npz not found; Petr point in (f) omitted")
    return out


def make_mock(seed=42, N=60000):
    rng = np.random.default_rng(seed)
    NA = rng.choice([1500, 2250, 3000], N)
    dr = np.load("dfe_draws.npz")["all"]
    beta, Sbar = dr[rng.integers(len(dr), size=N)].T
    lam = rng.gamma(225.8, 1.0, N)               # synonymous-count calibration
    U = (lam / 61000.0) * 1.663 * (0.0093 / 0.0064)   # RNA-inclusive input

    def surface(U_, Sbar_, NA_):
        base = np.select([NA_ == 1500, NA_ == 2250], [0.67, 0.45], 0.29)
        return base * (U_ / 0.0093) * np.clip(Sbar_ / 968.0, 0.05, 50) ** -0.30

    def noisy(mu, NA_):
        sd = np.select([NA_ == 1500, NA_ == 2250], [0.33, 0.26], 0.20)
        skew = (rng.gamma(4.0, 1.0, mu.size) - 4.0) / 2.0
        return mu + sd * skew

    mu = surface(U, Sbar, NA)
    gap = noisy(mu, NA)

    # panel-e mock variants
    gap_cod = noisy(mu / 1.453, NA)                       # coding-only, no ABC
    keep = rng.random(N) < 0.5
    gap_cod_abc = noisy((mu / 1.453)[keep] * 0.95, NA[keep])   # "accepted"
    gap_rna_noabc = noisy(mu * 1.03, NA) + 0.05 * rng.standard_normal(N)

    return dict(main=gap, U=lam, beta=beta, Sbar=Sbar, NA=NA,
                cod_noabc=gap_cod, cod_abc=gap_cod_abc, NA_cod_all=NA,
                NA_cod_acc=NA[keep], rna_noabc=gap_rna_noabc,
                NA_rna_all=NA, mock=True)


# --------------------------------------------------------------------------
def _hist(ax, x, color, label, lw=1.6, fill=False, ls="-", clip=True):
    if clip:
        x = np.clip(x, XLIM[0] + 1e-6, XLIM[1] - 1e-6)
    if fill:
        ax.hist(x, bins=BINS, density=True, color="0.85", edgecolor="black",
                linewidth=1.4, histtype="stepfilled", label=label, zorder=1)
    else:
        ax.hist(x, bins=BINS, density=True, color=color, linewidth=lw,
                histtype="step", label=label, zorder=3, linestyle=ls)


def _slice_panel(ax, d, key, sym):
    """Full posterior in black + conditional histograms at the ~5th pct, mean,
    ~95th pct of parameter `key` (slices: 0-10, 45-55, 90-100 percentiles).
    Every legend label carries the slice's integrated fixation probability."""
    v = d[key]
    p0 = pfix_mix(d["main"], d["NA"])
    _hist(ax, d["main"], None, f"full posterior ({p0:.0f}%)", fill=True)
    wins = [(0, 10, "\u22485th pct"), (45, 55, "\u2248mean"),
            (90, 100, "\u224895th pct")]
    for (lo, hi, tag), c in zip(wins, COL_LMH):
        vlo, vhi = np.percentile(v, [lo, hi])
        m = (v >= vlo) & (v <= vhi)
        p = pfix_mix(d["main"][m], d["NA"][m])
        _hist(ax, d["main"][m], c, f"{sym} {tag} ({p:.0f}%)")


def plot(d, out="fig2_mtdna_abc"):
    fig, axes = plt.subplots(2, 3, figsize=(12.5, 6.4))
    (axA, axB, axC), (axD, axE, axF) = axes

    _slice_panel(axA, d, "U", "U")
    axA.set_title("a  deleterious input U", fontsize=10, loc="left")
    _slice_panel(axB, d, "beta", "\u03b2")
    axB.set_title("b  DFE shape \u03b2", fontsize=10, loc="left")
    _slice_panel(axC, d, "Sbar", "|S\u0304|")
    axC.set_title("c  DFE mean |S\u0304|", fontsize=10, loc="left")

    # (d) archaic size: each NA trace scored on its own contracted curve
    p0 = pfix_mix(d["main"], d["NA"])
    _hist(axD, d["main"], None, f"full posterior ({p0:.0f}%)", fill=True)
    for na in (1500, 2250, 3000):
        m = d["NA"] == na
        if np.any(m):
            p = pfix(d["main"][m], PAIR_NN[na]).mean()
            _hist(axD, d["main"][m], COL_NA[na],
                  f"{NA_LAB} = {na} ({p:.0f}%)")
    axD.set_title("d  archaic population size", fontsize=10, loc="left")

    # (e) model variants (each label: integrated fixation probability)
    _hist(axE, d["main"], None, f"coding+RNA, ABC ({p0:.0f}%)", fill=True)
    if "cod_noabc" in d:
        p = pfix_mix(d["cod_noabc"], d["NA_cod_all"])
        _hist(axE, d["cod_noabc"], COL_E[0], f"coding, no ABC ({p:.0f}%)")
        p = pfix_mix(d["cod_abc"], d["NA_cod_acc"])
        _hist(axE, d["cod_abc"], COL_E[1], f"coding, ABC ({p:.0f}%)")
    p = pfix_mix(d["rna_noabc"], d["NA_rna_all"])
    _hist(axE, d["rna_noabc"], COL_E[2], f"coding+RNA, no ABC ({p:.0f}%)")
    if "petr" in d:
        # not clipped: the long right tail runs past the axis rather than
        # piling up in the last bin (the label % integrates all draws)
        p = pfix(d["petr"], 500).mean()
        _hist(axE, d["petr"], "0.3",
              f"Petr et al. parameters ({p:.0f}%)", ls="--", lw=1.4,
              clip=False)
    axE.set_title("e  model variants", fontsize=10, loc="left")

    # (f) translating the gap posterior into a fixation probability. The
    # post-mixture contraction applies under every accumulation-phase size
    # (same ~1/3 ratio as the central 2250 -> 750), so each NA posterior is
    # paired with its contracted post-mixture size NN: 1500 -> 500,
    # 2250 -> 750, 3000 -> 1000. The 500 curve is shared with the Petr
    # comparison (their constant 500).
    if CURVES is None:
        raise SystemExit(f"{FIXCURVE} not found")
    gx = np.linspace(*XLIM, 400)
    # curves, labeled by the post-mixture size alone (legend upper left)
    axF.plot(gx, pfix(gx, 500), color="0.4", lw=1.4, ls="--",
             label=f"{NN_LAB} = 500")
    axF.plot(gx, pfix(gx, 750), color="black", lw=2.2,
             label=f"{NN_LAB} = 750")
    if 1000 in CURVES:
        axF.plot(gx, pfix(gx, 1000), color=COL_NA[3000], lw=1.2, alpha=0.8,
                 label=f"{NN_LAB} = 1000")
    # posterior medians on their contracted curves, labeled directly at the
    # point: name (NA/NN) and the integrated fixation probability
    POINTS = [
        (2250, "Central", "black", "o", 7, (8, -14), "left", "top"),
        (1500, "Lower", COL_NA[1500], "o", 5, (12, -6), "left", "top"),
        (3000, "Upper", COL_NA[3000], "o", 5, (0, 10), "center", "bottom"),
    ]
    for na, name, col, marker, ms, off, ha, va in POINTS:
        nn = PAIR_NN[na]
        m = d["NA"] == na
        if not np.any(m) or nn not in CURVES:
            continue
        p = pfix(d["main"][m], nn).mean()
        med = np.median(d["main"][m])
        axF.plot([med], [pfix(med, nn)], marker, ms=ms, color=col,
                 mec="white" if na == 2250 else "black",
                 mew=0.8 if na == 2250 else 0.5, zorder=6 if na == 2250 else 5)
        axF.annotate(f"{name} ({na}/{nn}) {p:.0f}%",
                     (med, pfix(med, nn)), xytext=off,
                     textcoords="offset points", fontsize=7.4, color=col,
                     ha=ha, va=va, zorder=7)
    if "petr" in d:
        p = pfix(d["petr"], 500).mean()
        med = np.median(d["petr"])
        axF.plot([med], [pfix(med, 500)], "s", ms=6, color="0.4",
                 mec="black", mew=0.6, zorder=5)
        axF.annotate(f"Petr (500/500) {p:.0f}%",
                     (med, pfix(med, 500)), xytext=(8, -8),
                     textcoords="offset points", fontsize=7.4, color="0.25",
                     ha="left", va="top", zorder=7)
    axF.set_ylim(0, 100)
    axF.set_ylabel("P(modern mtDNA fixes) (%)")
    axF.set_title("f  gap \u2192 fixation probability", fontsize=10, loc="left")

    for ax in (axA, axB, axC, axD, axE):
        ax.legend(fontsize=7.2, frameon=False, loc="upper right",
                  handlelength=1.4, borderaxespad=0.2)
    axF.legend(fontsize=7.2, frameon=False, loc="upper left",
               handlelength=1.4, borderaxespad=0.2)

    # uniform axes: shared density scale for the five histogram panels
    ymax = max(ax.get_ylim()[1] for ax in (axA, axB, axC, axD, axE))
    for ax in (axA, axB, axC, axD, axE):
        ax.set_ylim(0, ymax)
    for ax in (axA, axB, axC, axD, axE, axF):
        ax.axvline(0, color="k", lw=0.7, ls=":", zorder=2)
        ax.set_xlim(*XLIM)
        ax.set_xlabel("fitness gap, modern \u2212 archaic (%)")
    axA.set_ylabel("posterior density")
    axD.set_ylabel("posterior density")

    tag = " \u2014 MOCKUP (synthetic draws)" if d["mock"] else ""
    fig.suptitle("mtDNA fitness gap at introgression: ABC posterior, all "
                 f"uncertainty integrated{tag}", fontsize=12)
    fig.tight_layout(rect=(0, 0, 1, 0.95))
    for ext in ("png", "pdf"):
        fig.savefig(f"{out}.{ext}", dpi=200)
    print(f"wrote {out}.png / .pdf")


def _draw_main_hist(ax, g):
    """ABC posterior histogram + median stem (shared by 2-panel and hist-only)."""
    med = float(np.median(g))
    clipped = np.clip(g, XLIM[0] + 1e-6, XLIM[1] - 1e-6)
    n_hist, edges, _ = ax.hist(
        clipped, bins=BINS, density=True, color="0.85", edgecolor="black",
        linewidth=1.0, histtype="stepfilled", zorder=1)
    ibin = min(max(np.searchsorted(edges, med, side="right") - 1, 0),
               len(n_hist) - 1)
    h_med = float(n_hist[ibin])
    ax.annotate(
        f"median {med:.2f}% modern-archaic mtDNA\n"
        "fitness gap at introgression",
        xy=(med, h_med),
        xytext=(med, h_med + 0.18), ha="left", va="bottom",
        fontsize=11, color="0.15",
        arrowprops=dict(arrowstyle="-", color="0.35", lw=0.8))
    ax.set_ylim(0, float(np.max(n_hist)) * 1.32)
    ax.set_xlim(*XLIM)
    ax.set_ylabel("posterior density")
    return med


def plot_main_hist(d, out="fig2_mtdna_abc_main_hist"):
    """Histogram-only ABC posterior, with x-axis labels visible."""
    g = np.asarray(d["main"], float)
    fig, ax = plt.subplots(figsize=(7.2, 4.2))
    _draw_main_hist(ax, g)
    ax.set_xlabel("fitness gap, modern \u2212 archaic (%)")
    fig.tight_layout()
    for ext in ("png", "pdf"):
        fig.savefig(f"{out}.{ext}", dpi=200)
    print(f"wrote {out}.png / .pdf")
    plt.close(fig)


def plot_main(d, out="fig2_mtdna_abc_main"):
    """Full ABC posterior (top) and the central gap-to-fixation curve (bottom)."""
    if CURVES is None:
        raise SystemExit(f"{FIXCURVE} not found")
    g = np.asarray(d["main"], float)
    med = float(np.median(g))
    mean = float(g.mean())
    lo, hi = [float(x) for x in np.percentile(g, [2.5, 97.5])]
    ppos = float((g > 0).mean())
    n = int(g.size)
    p_mix = pfix_mix(g, d["NA"])
    p_at_med = float(pfix(med, 750))

    fig, (ax, axF) = plt.subplots(
        2, 1, figsize=(7.2, 7.6), sharex=True,
        gridspec_kw={"height_ratios": [1.15, 1]})
    _draw_main_hist(ax, g)

    gx = np.linspace(*XLIM, 400)
    axF.plot(gx, pfix(gx, 750), color="black", lw=2.0)
    axF.axvline(med, color="0.35", lw=1.0, ls="--", zorder=2)
    axF.text(
        -0.95, 96,
        "23% probability of mtDNA\n"
        "fixation, integrating over\n"
        "uncertainty in the fitness\n"
        "gap, and uncertainty in\n"
        "the demographic history",
        ha="left", va="top", fontsize=12, linespacing=1.3)
    axF.set_ylim(0, 100)
    axF.set_xlim(*XLIM)
    axF.set_ylabel("P(modern mtDNA fixes) (%)")
    axF.set_xlabel("fitness gap, modern \u2212 archaic (%)")
    fig.tight_layout()
    for ext in ("png", "pdf"):
        fig.savefig(f"{out}.{ext}", dpi=200)
    print(f"wrote {out}.png / .pdf")
    plot_main_hist(d)
    print(f"n={n}  median={med:.3f}%  mean={mean:.3f}%  "
          f"95% [{lo:.3f}, {hi:.3f}]  P(gap>0)={ppos:.3f}  "
          f"P(fix) mix={p_mix:.1f}%  P(fix)|median,750={p_at_med:.1f}%")


def main():
    ap = argparse.ArgumentParser(description=__doc__)
    ap.add_argument("--mock", action="store_true",
                    help="synthetic draws (layout preview)")
    ap.add_argument("--main-only", action="store_true",
                    help="single-panel full posterior (not the 6-panel Figure 2)")
    args = ap.parse_args()
    if args.main_only:
        if args.mock:
            raise SystemExit("--main-only requires real joint_rna_*.npz; "
                             "do not use --mock")
        d = load_real()
        plot_main(d)
        return
    d = make_mock() if args.mock else load_real()
    plot(d, out="fig2_mtdna_abc_mock" if args.mock else "fig2_mtdna_abc")


if __name__ == "__main__":
    main()
```
