## Supplementary Information section 1 code base for "Hypothesis: A modern human range expansion ∼300,000 years ago had a critical role in Neandertal origins": plot_replicate.py

```
"""
LOCAL plotting: compare the NEW framework (delta DFE, new_replicate.csv from the
cluster) against the ORIGINAL published values in the Excel workbook, for every
Extended Data Figure 1 grid point (NA = 1500 and 3000).

Outputs:
  replicate_compare.csv  - per-point original vs new (modern, archaic, diff)
  replicate_scatter.png/.pdf - 3-panel scatter (modern, archaic, difference)

Run:  ./.venv/bin/python plot_replicate.py
"""
import csv
import os

os.environ.setdefault("MPLCONFIGDIR", os.path.abspath(".mplcache"))
import numpy as np
import openpyxl
import matplotlib
matplotlib.use("Agg")
import matplotlib.pyplot as plt

XLSX_NAME = "2_26_2026 Muller's Ratchet simulations.xlsx"
NEW = "new_replicate.csv"


def find_xlsx():
    here = os.path.dirname(os.path.abspath(__file__))
    for p in (XLSX_NAME,
              os.path.join("data", XLSX_NAME),
              os.path.join(here, XLSX_NAME),
              os.path.join(here, "..", XLSX_NAME)):
        if os.path.isfile(p):
            return p
    raise SystemExit(f"missing original Perl-sim workbook {XLSX_NAME}")

MUS = [0.001, 0.002, 0.005, 0.01, 0.02, 0.05]
SS = [0.0001, 0.0003, 0.001, 0.003, 0.01, 0.03, 0.1, 0.3, 1.0]
# Excel column blocks (1-based) for the two archaic sizes we keep.
NA_COL0 = {1500: 14, 3000: 25}      # first s-column of each block
MODERN_ROW0 = 6                     # mu=0.001 row of the Modern table
ARCHAIC_ROW0 = 15                   # mu=0.001 row of the Archaic table


def load_original():
    wb = openpyxl.load_workbook(find_xlsx(), data_only=True)
    ws = wb["Sheet1"]
    orig = {}
    for NA, c0 in NA_COL0.items():
        for i, mu in enumerate(MUS):
            for j, s in enumerate(SS):
                m = ws.cell(MODERN_ROW0 + i, c0 + j).value
                a = ws.cell(ARCHAIC_ROW0 + i, c0 + j).value
                orig[(NA, round(mu, 6), round(s, 6))] = (float(m), float(a))
    return orig


def load_new():
    new = {}
    with open(NEW) as f:
        for row in csv.DictReader(f):
            key = (int(row["NA"]), round(float(row["mu"]), 6),
                   round(float(row["s"]), 6))
            new[key] = (float(row["new_modern"]), float(row["new_archaic"]))
    return new


def main():
    orig, new = load_original(), load_new()
    rows = []
    for key in sorted(orig):
        if key not in new:
            continue
        NA, mu, s = key
        om, oa = orig[key]
        nm, na = new[key]
        rows.append(dict(NA=NA, mu=mu, s=s,
                         orig_modern=om, orig_archaic=oa, orig_diff=om - oa,
                         new_modern=nm, new_archaic=na, new_diff=nm - na))

    with open("replicate_compare.csv", "w", newline="") as f:
        w = csv.DictWriter(f, fieldnames=list(rows[0].keys()))
        w.writeheader()
        w.writerows(rows)
    print(f"Wrote replicate_compare.csv ({len(rows)} points).")

    panels = [("modern", "Modern lineage fitness"),
              ("archaic", "Archaic lineage fitness"),
              ("diff", "Modern \u2212 archaic difference")]
    fig, axes = plt.subplots(1, 3, figsize=(15, 5))
    colors = {1500: "#1f77b4", 3000: "#ff7f0e"}
    for ax, (key, title) in zip(axes, panels):
        xo = np.array([r[f"orig_{key}"] for r in rows])
        yn = np.array([r[f"new_{key}"] for r in rows])
        cols = [colors[r["NA"]] for r in rows]
        ax.scatter(xo, yn, c=cols, s=28, alpha=0.8, edgecolors="none")
        lo = min(xo.min(), yn.min())
        hi = max(xo.max(), yn.max())
        pad = 0.03 * (hi - lo + 1e-9)
        ax.plot([lo - pad, hi + pad], [lo - pad, hi + pad], "k--", lw=1,
                alpha=0.6)
        ax.set_xlim(lo - pad, hi + pad)
        ax.set_ylim(lo - pad, hi + pad)
        rmse = float(np.sqrt(np.mean((yn - xo) ** 2)))
        mad = float(np.max(np.abs(yn - xo)))
        r2 = float(np.corrcoef(xo, yn)[0, 1] ** 2)
        ax.set_title(f"{title}\nRMSE={rmse:.4f}  maxdev={mad:.4f}  "
                     f"R\u00b2={r2:.4f}", fontsize=10)
        ax.set_xlabel("Original (Perl, 10 reps)")
        ax.set_ylabel("New framework (delta DFE, 30 reps)")
        ax.grid(alpha=0.25)
    handles = [plt.Line2D([], [], marker="o", ls="", color=colors[k],
                          label=f"N_A={k}") for k in (1500, 3000)]
    axes[0].legend(handles=handles, loc="upper left", fontsize=9)
    fig.suptitle("Validation: new framework reproduces every Extended Data "
                 "Figure 1 grid point", fontsize=12)
    fig.tight_layout(rect=[0, 0, 1, 0.96])
    fig.savefig("replicate_scatter.png", dpi=150)
    fig.savefig("replicate_scatter.pdf")
    print("Wrote replicate_scatter.png / .pdf")

    # headline numbers across all panels
    allo = np.array([r[f"orig_{k}"] for r in rows for k in ("modern", "archaic")])
    alln = np.array([r[f"new_{k}"] for r in rows for k in ("modern", "archaic")])
    print(f"Overall (modern+archaic fitness): RMSE="
          f"{np.sqrt(np.mean((alln-allo)**2)):.4f}, "
          f"max|dev|={np.max(np.abs(alln-allo)):.4f}")


if __name__ == "__main__":
    main()
```
