## Supplementary Information section 1 code base for "Hypothesis: A modern human range expansion ∼300,000 years ago had a critical role in Neandertal origins": plot_y_fig3_violin.py

```
"""
Alternative Figure 3: violin plot of the Y-chromosome fitness-gap
distributions (y-axis, linear) against the coding-equivalent constrained
target (x-axis categories: 100 kb ... 500 kb, Petr, 1 Mb, 2 Mb, 5 Mb).
Same data as plot_y_fig3.py (y_v22_reps.npz; 300 replicates per violin,
main scenario at the central N_A = 2250; the Petr violin is their
demography at their 500-kb reference target).

    python plot_y_fig3_violin.py     # -> fig3_y_gap_violin.png / .pdf
    python plot_y_fig3_violin.py --npz y_draw_reps.npz --out fig3_y_gap_violin_integrated
                                     # parameter-integrated version
"""
import argparse
import os

import numpy as np

os.environ.setdefault("MPLCONFIGDIR", os.path.abspath(".mplcache"))
os.makedirs(os.environ["MPLCONFIGDIR"], exist_ok=True)
import matplotlib
matplotlib.use("Agg")
import matplotlib.pyplot as plt


PETR_REPORTED = 2.0              # percent; deficit reported by Petr et al.
FIXCURVE = "fixation_curves_mtdna.csv"

# main rungs pool the three archaic sizes with equal weight (like the
# mtDNA rows of Table 3); each draw's P(fix) uses the fixation curve of
# its NA-paired post-mixture size
NAS = (1500, 2250, 3000)
PAIR_NN = {1500: 500, 2250: 750, 3000: 1000}

# (kb tag, x label, fill color); Petr comparison at the far right
ORDER = [
    ("y100",  "100 kb\n(coding)", "0.60"),
    ("y200",  "200 kb",           "#8c9fca"),
    ("y300",  "300 kb",           "#5f81c0"),
    ("y500",  "500 kb",           "#2c6fbb"),
    ("y1000", "1 Mb",             "#7fbf7b"),
    ("y2000", "2 Mb",             "#e08214"),
    ("y3000", "3 Mb",             "#c0392b"),
    ("y500_petr_500", "Petr et al.\nsimulation parameters\n(500 kb)",
     "0.15"),
]


def load_fixcurves(path=FIXCURVE):
    """Curves from the companion introgression analysis: per locus_Ne, the
    probability (fraction) that the modern mtDNA/Y fixes after a 5% pulse,
    as a function of its selective advantage s. locus_Ne = 750 corresponds
    to the post-mixture Neandertal size; 500 to the Petr comparison."""
    if not os.path.exists(path):
        return None
    import csv
    curves = {}
    with open(path) as f:
        rows = csv.DictReader(r for r in f if not r.startswith("#"))
        for r in rows:
            curves.setdefault(int(float(r["locus_Ne"])), []).append(
                (float(r["s"]), float(r["pfix_pct"]) / 100.0))
    return {ne: np.array(sorted(pts)) for ne, pts in curves.items()}


def pfix_label(segments, curves):
    """segments: list of (gaps_frac, locus_ne); the label averages P(fix)
    over all draws, each scored on its own curve."""
    if curves is None or any(ne not in curves for _, ne in segments):
        return "P(fix)\npending"
    vals, clipped = [], []
    for g, ne in segments:
        s, p = curves[ne][:, 0], curves[ne][:, 1]
        vals.append(100 * np.interp(g, s, p))
        clipped.append(np.asarray(g) > s.max())
    vals = np.concatenate(vals)
    # gaps beyond the tabulated s-range clamp to the boundary: a lower bound
    geq = "\u2265" if np.concatenate(clipped).mean() > 0.05 else ""
    return f"P(fix)\n{geq}{vals.mean():.0f}%"


def main():
    ap = argparse.ArgumentParser()
    ap.add_argument("--npz", default="y_draw_stated_reps.npz")
    ap.add_argument("--out", default="fig3_y_gap_violin")
    args = ap.parse_args()
    data = np.load(args.npz)
    curves = load_fixcurves()
    if curves is None:
        print(f"{FIXCURVE} not found; annotations will say 'P(fix) pending'")
    # series: (label, pooled gaps in %, color, [(gaps_frac, locus_ne), ...])
    series = []
    for tag, lab, col in ORDER:
        if tag in data:                      # single-scenario key (Petr)
            segs = [(data[tag], 500)]
        else:                                # pool the three archaic sizes
            keys = [f"{tag}_{na}" for na in NAS]
            if not all(k in data and data[k].size for k in keys):
                print(f"note: {tag} incomplete in {args.npz}; skipped")
                continue
            segs = [(data[k], PAIR_NN[na]) for k, na in zip(keys, NAS)]
        pooled = np.concatenate([g for g, _ in segs]) * 100.0
        series.append((lab, pooled, col, segs))

    fig, ax = plt.subplots(figsize=(9, 5.5))
    ax.axhline(0, color="0.8", lw=0.8, zorder=1)
    ax.axhline(PETR_REPORTED, color="0.15", lw=1.4, ls=":", zorder=2,
               label="Petr et al. reported deficit (~2%)")

    pos = np.arange(1, len(series) + 1)
    parts = ax.violinplot([s[1] for s in series], positions=pos,
                          widths=0.8, showextrema=False)
    for body, (lab, pct, col, segs) in zip(parts["bodies"], series):
        body.set_facecolor(col)
        body.set_edgecolor("0.25")
        body.set_alpha(0.85)
        if lab.startswith("Petr"):
            body.set_linestyle("--")
            body.set_linewidth(1.6)
            body.set_facecolor("none")
    # median/mean markers and per-violin annotation (mean gap, P(fix))
    ymax = max(p.max() for _, p, _, _ in series)
    for x, (lab, pct, col, segs) in zip(pos, series):
        ax.plot([x - 0.18, x + 0.18], [np.median(pct)] * 2,
                color="0.15", lw=1.4, zorder=4)
        ax.plot(x, pct.mean(), "o", color="0.15", ms=3.5, zorder=4)
        ax.text(x, pct.max() + 0.035 * ymax,
                f"mean\n{pct.mean():.2f}%\n{pfix_label(segs, curves)}",
                ha="center", va="bottom", fontsize=7.5, color="0.2",
                linespacing=1.15)

    ymin = min(p.min() for _, p, _, _ in series)
    ax.set_ylim(min(ymin * 1.2, -0.3), ymax * 1.30)
    ax.set_xticks(pos)
    ax.set_xticklabels([s[0] for s in series])
    ax.set_ylabel("fitness gap, modern \u2212 archaic (%)")
    ax.set_xlabel("coding-equivalent constrained target")
    ax.legend(fontsize=9, loc="upper left", framealpha=0.9)
    fig.tight_layout()
    for ext in ("png", "pdf"):
        fig.savefig(f"{args.out}.{ext}", dpi=200)
    print(f"wrote {args.out}.png / .pdf")


if __name__ == "__main__":
    main()
```
