## Supplementary Information section 1 code base for "Hypothesis: A modern human range expansion ∼300,000 years ago had a critical role in Neandertal origins": ratchet_dfe.py

```
"""
Muller's-ratchet simulation for mtDNA fitness decline under a realistic
distribution of fitness effects (DFE), comparing a large "modern" lineage with
a smaller "archaic" lineage.

This is a DFE-based replacement for the single-`s` Perl simulation behind
Extended Data Figure 1. Instead of assuming one selection coefficient for all
mutations, each new amino-acid (missense) mutation draws its (unscaled)
selection coefficient from a gamma DFE estimated for human mtDNA by
Piganeau & Eyre-Walker (2003, PNAS): shape beta = 0.39, mean scaled
coefficient S-bar = -1400.

Scaling / anchoring
-------------------
P&E-W report the *scaled* coefficient S = 2*Ne*s. The DFE was estimated from
the human (= modern) lineage, so we anchor it there: the unscaled mean is
    s_bar = S_bar / (2 * anchor_Ne),   anchor_Ne = N_modern = 7500.
The SAME intrinsic (unscaled) DFE is used in both populations; the smaller
archaic population simply selects less efficiently (efficacy ~ N*s), so it
accumulates more mildly deleterious mutations. The proportionality constant
cancels because the conversion and the simulation use the same N-scaling.

Model
-----
Haploid, clonal, fully linked mtDNA; one haplotype per individual; no
recombination; no back-mutation (infinite-sites). Each generation:
  (1) Selection + reproduction: N parents resampled proportional to fitness
      (Wright-Fisher with viability selection).
  (2) Mutation: each offspring gets Poisson(U_mis) missense mutations, each
      multiplying fitness by (1 - s), s ~ Gamma(beta, s_bar/beta), capped at 1;
      plus Poisson(U_non) nonsense mutations, treated as lethal (s = 1).

Fitness is multiplicative and tracked in log space. Mean fitness is absolute
(both lineages start at 1.0); the readout is the modern - archaic difference,
compared against the ~5% threshold at which the range-expansion simulations
show mtDNA sweeps to fixation.
"""
import argparse
import json
import sys
import time

import numpy as np

ANCHOR_NE = 7500          # modern female Ne; also the DFE anchor (P&E-W reference)
# nonsense:missense opportunity ratio under the deCODE spectrum (compute_fn.py):
# f_nonsense / f_missense = 0.0350 / 0.602
NONSENSE_FRACTION = 0.0350 / 0.602


def simulate_one(N, T, U_mis, U_non, beta, scale, rng, record_every=0):
    """Run one lineage; return final mean (absolute) fitness.

    If record_every > 0, also return the mean-fitness trajectory.
    """
    L = np.zeros(N)                      # log absolute fitness, ln(w)
    idx_all = np.arange(N)
    traj = []
    for gen in range(T):
        # (1) viability selection: sample parents proportional to fitness.
        # Subtract max for numerical stability; this does not change relative p.
        w = np.exp(L - L.max())
        p = w / w.sum()
        parents = rng.choice(N, size=N, replace=True, p=p)
        L = L[parents]

        # (2a) missense mutations (drawn from the gamma DFE)
        k = rng.poisson(U_mis, size=N)
        m = int(k.sum())
        if m:
            s = rng.gamma(beta, scale, size=m)
            np.minimum(s, 1.0, out=s)
            offspring = np.repeat(idx_all, k)
            with np.errstate(divide="ignore"):   # s==1 -> log1p(-1)=-inf (lethal)
                dL = np.bincount(offspring, weights=np.log1p(-s), minlength=N)
            L = L + dL                    # ln(1-s) summed per offspring

        # (2b) nonsense mutations (lethal)
        if U_non > 0:
            lethal = rng.poisson(U_non, size=N) > 0
            if lethal.any():
                L[lethal] = -np.inf

        if record_every and (gen % record_every == 0 or gen == T - 1):
            traj.append((gen, float(np.exp(L).mean())))

    mean_w = float(np.exp(L).mean())
    return (mean_w, traj) if record_every else (mean_w, None)


def run(N_modern, N_archaic, T, U_mis, U_non, beta, Sbar, reps, seed,
        record_every=0, anchor_ne=ANCHOR_NE):
    # The DFE is reported as a *scaled* coefficient S = 2*Ne*s estimated at some
    # reference Ne (anchor_ne). For mtDNA that is the modern female Ne (7500);
    # for the Y we use the autosomal DFE estimated at Ne=10000.
    s_bar = Sbar / (2.0 * anchor_ne)     # anchor the DFE to its reference Ne
    scale = s_bar / beta                 # gamma scale (mean = beta*scale = s_bar)
    rng = np.random.default_rng(seed)

    rows = []
    for r in range(reps):
        wm, tm = simulate_one(N_modern, T, U_mis, U_non, beta, scale, rng,
                              record_every)
        wa, ta = simulate_one(N_archaic, T, U_mis, U_non, beta, scale, rng,
                              record_every)
        rows.append(dict(rep=r, w_modern=wm, w_archaic=wa, diff=wm - wa,
                         traj_modern=tm, traj_archaic=ta))
    return rows, s_bar


def summarize(rows, *, N_modern, N_archaic, T, U_mis, U_non, beta, Sbar,
              s_bar, reps, seed, elapsed=None):
    """Collapse per-replicate rows into a summary dict."""
    diffs = np.array([r["diff"] for r in rows])
    wm = np.array([r["w_modern"] for r in rows])
    wa = np.array([r["w_archaic"] for r in rows])
    n = len(rows)
    sd = float(diffs.std(ddof=1)) if n > 1 else 0.0
    return dict(
        N_modern=N_modern, N_archaic=N_archaic, T=T, U_mis=U_mis, U_non=U_non,
        beta=beta, Sbar=Sbar, s_bar=s_bar, reps=reps, seed=seed,
        w_modern_mean=float(wm.mean()), w_archaic_mean=float(wa.mean()),
        diff_mean=float(diffs.mean()), diff_sd=sd,
        diff_se=sd / np.sqrt(n) if n > 1 else 0.0,
        frac_reps_over_5pct=float(np.mean(diffs > 0.05)),
        elapsed_sec=elapsed,
    )


def main():
    ap = argparse.ArgumentParser(description=__doc__)
    ap.add_argument("--N-modern", type=int, default=7500)
    ap.add_argument("--N-archaic", type=int, default=1500)
    ap.add_argument("--T", type=int, default=17500, help="generations of separation")
    ap.add_argument("--U-mis", type=float, default=0.03,
                    help="de novo missense mutations per genome per generation")
    ap.add_argument("--U-non", type=float, default=None,
                    help="nonsense (lethal) rate; default = U_mis * 0.0581")
    ap.add_argument("--beta", type=float, default=0.39, help="gamma shape")
    ap.add_argument("--Sbar", type=float, default=1400.0,
                    help="mean scaled selection coefficient |S| (P&E-W)")
    ap.add_argument("--reps", type=int, default=10)
    ap.add_argument("--seed", type=int, default=0)
    ap.add_argument("--anchor-ne", type=float, default=ANCHOR_NE,
                    help="reference Ne at which the scaled DFE was estimated "
                         "(7500 for mtDNA P&E-W; 10000 for the autosomal/Y DFE)")
    ap.add_argument("--record-every", type=int, default=0,
                    help="if >0, record mean-fitness trajectory every N gens")
    ap.add_argument("--out", type=str, default=None,
                    help="output .jsonl path (default stdout)")
    args = ap.parse_args()

    U_non = args.U_non if args.U_non is not None else args.U_mis * NONSENSE_FRACTION

    t0 = time.time()
    rows, s_bar = run(args.N_modern, args.N_archaic, args.T, args.U_mis, U_non,
                      args.beta, args.Sbar, args.reps, args.seed,
                      args.record_every, anchor_ne=args.anchor_ne)
    elapsed = time.time() - t0

    summary = summarize(rows, N_modern=args.N_modern, N_archaic=args.N_archaic,
                        T=args.T, U_mis=args.U_mis, U_non=U_non, beta=args.beta,
                        Sbar=args.Sbar, s_bar=s_bar, reps=args.reps,
                        seed=args.seed, elapsed=elapsed)

    out = open(args.out, "a") if args.out else sys.stdout
    out.write(json.dumps({"summary": summary,
                          "reps": [{k: v for k, v in r.items()
                                    if not k.startswith("traj") or args.record_every}
                                   for r in rows]}) + "\n")
    if args.out:
        out.close()
    # human-readable line to stderr for monitoring
    sys.stderr.write(
        f"Sbar={args.Sbar:>6} beta={args.beta} U_mis={args.U_mis} "
        f"N_A={args.N_archaic}: diff={summary['diff_mean']:.4f}"
        f"+-{summary['diff_sd']:.4f} (w_M={summary['w_modern_mean']:.4f} "
        f"w_A={summary['w_archaic_mean']:.4f}) [{elapsed:.1f}s]\n")


if __name__ == "__main__":
    main()
```
