## Supplementary Information section 1 code base for "Hypothesis: A modern human range expansion ∼300,000 years ago had a critical role in Neandertal origins": ratchet_dfe_annotated.py

```
"""
============================================================================
 ANNOTATED READING COPY of the core simulation (ratchet_dfe.py).
============================================================================
This file is functionally identical to `ratchet_dfe.py` but is commented for
someone who knows Perl and C but not Python/NumPy. It is meant to be READ.
(The version actually run on the cluster is ratchet_dfe.py.)

Orientation for a Perl/C programmer
-----------------------------------
* NumPy ("import numpy as np") gives Python C-style numeric arrays plus
  "vectorized" operations: instead of writing a C for-loop over an array,
  you write one call that loops internally in compiled C. So where the old
  Perl `ratchet.pl` had an inner `for (my $i=0;$i<$M;$i++)` over every
  individual, here that loop disappears into single array operations. This is
  the main reason the Python version is fast despite Python being slow.

* An "ndarray" (e.g. `L` below) is like a C array `double L[N]`. Writing
  `L = L[parents]` builds a NEW array by gathering elements at the index
  positions in `parents` -- think of it as a gather/permute, not a copy loop.

* `rng` is a random-number generator object (like a seeded drand48 state).
  `rng.choice`, `rng.poisson`, `rng.gamma` draw whole arrays of random
  numbers at once.

Biology recap (what each lineage is)
------------------------------------
Each individual carries ONE haploid mtDNA with a fitness w in (0,1].
w starts at 1.0 and is multiplied by (1-s) for every deleterious mutation it
accumulates (Muller's ratchet: no recombination, no back-mutation). We store
ln(w) instead of w (called L) so that multiplying fitnesses becomes ADDING
logs -- this avoids underflow and lets us sum many mutations cheaply.
============================================================================
"""
import numpy as np

# The DFE was estimated from the human (= modern) lineage, so we anchor the
# unscaled mean selection coefficient to the modern Ne:  s_bar = S_bar/(2*Ne).
ANCHOR_NE = 7500

# Of protein-coding single-nucleotide mutations, ~3.5% create a premature stop
# (nonsense) and ~60.2% are missense (compute_fn.py). We feed missense through
# the DFE and treat nonsense as a separate lethal class; this is their ratio.
NONSENSE_FRACTION = 0.0350 / 0.602


def simulate_one(N, T, U_mis, U_non, beta, scale, rng):
    """
    Evolve ONE lineage of N haploid mtDNAs for T generations and return the
    mean (absolute) fitness at the end.

    Parameters (all scalars unless noted):
      N      population size (number of individuals / mtDNAs)
      T      number of generations
      U_mis  expected # missense mutations per individual per generation
      U_non  expected # nonsense (lethal) mutations per individual per gen
      beta   gamma SHAPE parameter of the DFE (0.39)
      scale  gamma SCALE parameter (= s_bar / beta); mean of draws = beta*scale
      rng    random-number generator
    """
    # L[i] = natural log of individual i's fitness. "np.zeros(N)" is like
    # C's `double L[N] = {0}`; ln(1)=0, so everyone starts at fitness 1.
    L = np.zeros(N)

    # A constant 0..N-1 index vector, reused below (like `int idx[N]={0,1,...}`).
    idx_all = np.arange(N)

    for gen in range(T):                       # main generation loop (like Perl's $gen loop)

        # ---- (1) SELECTION + REPRODUCTION ---------------------------------
        # Sample N parents for the next generation, each chosen with
        # probability proportional to its fitness w = exp(L). This is
        # Wright-Fisher viability selection -- the same intent as the Perl
        # "pick a parent, reject with prob (1-fitness)" loop, but done in one
        # shot. We subtract max(L) before exponentiating purely for numerical
        # safety; it cancels out when we normalize, so probabilities are
        # unchanged.
        w = np.exp(L - L.max())                # relative fitness weights, all <= 1
        p = w / w.sum()                        # normalize to a probability vector
        parents = rng.choice(N, size=N, replace=True, p=p)   # gather N parent indices
        L = L[parents]                         # offspring inherit parents' fitness

        # ---- (2a) MISSENSE MUTATIONS (drawn from the DFE) -----------------
        # Each offspring gets a Poisson-distributed number of new missense
        # mutations (mean U_mis). k is an array: k[i] = how many for offspring i.
        k = rng.poisson(U_mis, size=N)
        m = int(k.sum())                       # total new missense mutations this gen
        if m:
            # Draw m selection coefficients from the gamma DFE, then cap at 1
            # (a mutation can't reduce fitness below 0). s is an array of m values.
            s = rng.gamma(beta, scale, size=m)
            np.minimum(s, 1.0, out=s)          # in-place: s = min(s, 1)

            # Expand k into a list of offspring indices, one entry per mutation.
            # e.g. if k = [2,0,1] then offspring = [0,0,2]. (np.repeat)
            offspring = np.repeat(idx_all, k)

            # For each mutation, the log-fitness change is ln(1-s). We sum these
            # per offspring with bincount (a fast grouped-sum: like a histogram
            # weighted by ln(1-s)). s==1 gives ln(0) = -inf (lethal); the
            # errstate just silences the expected divide-by-zero warning.
            with np.errstate(divide="ignore"):
                dL = np.bincount(offspring, weights=np.log1p(-s), minlength=N)
            L = L + dL                         # apply: multiply fitnesses by prod(1-s)

        # ---- (2b) NONSENSE MUTATIONS (lethal) -----------------------------
        # Premature stop codons are treated as lethal (s = 1 -> fitness 0).
        if U_non > 0:
            lethal = rng.poisson(U_non, size=N) > 0   # boolean array, True where >=1 hit
            if lethal.any():
                L[lethal] = -np.inf            # exp(-inf) = 0 -> purged next generation

    # Mean ABSOLUTE fitness across the population (both lineages started at 1.0,
    # so comparing this between lineages is meaningful). exp(L) turns logs back
    # into fitnesses; freshly-lethal individuals contribute 0.
    return float(np.exp(L).mean())


def run(N_modern, N_archaic, T, U_mis, U_non, beta, Sbar, reps, seed):
    """
    Run BOTH lineages for `reps` independent replicates and return the list of
    per-replicate results plus the unscaled mean coefficient s_bar.

    The two lineages use the SAME intrinsic DFE (same beta, same scale); only
    N differs, so the smaller archaic population selects less efficiently and
    accumulates more mildly-deleterious mutations -- that asymmetry is the
    whole point.
    """
    s_bar = Sbar / (2.0 * ANCHOR_NE)           # convert scaled S_bar to unscaled s_bar
    scale = s_bar / beta                       # gamma scale so that mean draw = s_bar
    rng = np.random.default_rng(seed)          # seeded RNG (reproducible)

    rows = []
    for r in range(reps):                      # like Perl's replicate loop
        wm = simulate_one(N_modern, T, U_mis, U_non, beta, scale, rng)
        wa = simulate_one(N_archaic, T, U_mis, U_non, beta, scale, rng)
        # diff = how much fitter modern mtDNA is than archaic mtDNA at the end;
        # compared against the ~5% threshold needed for it to sweep.
        rows.append(dict(rep=r, w_modern=wm, w_archaic=wa, diff=wm - wa))
    return rows, s_bar


# This file is for reading; run ratchet_dfe.py to actually produce results.
if __name__ == "__main__":
    # Tiny demonstration so you can SEE it work (short, not the real settings).
    rows, s_bar = run(N_modern=7500, N_archaic=1500, T=500,
                      U_mis=0.03, U_non=0.03 * NONSENSE_FRACTION,
                      beta=0.39, Sbar=1400, reps=2, seed=0)
    for row in rows:
        print(f"rep {row['rep']}: w_modern={row['w_modern']:.4f} "
              f"w_archaic={row['w_archaic']:.4f} diff={row['diff']:.4f}")
    print(f"(unscaled mean coefficient s_bar = {s_bar:.4f})")
```
