## Supplementary Information section 1 code base for "Hypothesis: A modern human range expansion ∼300,000 years ago had a critical role in Neandertal origins": README.md

# Muller's Ratchet in Archaic mtDNA and Y Chromosomes — Code Base

Code and collected simulation output accompanying Supplementary
Information section 1. This folder is the archive named in the note
(`SI1codebase-v7.zip`). Every program listed under Software and
Reproducibility in the note is included here; programs from earlier
drafts that are not used for the presented results are not included.

Everything runs under Python 3.10 with the packages in
`requirements.txt` (numpy, scipy, matplotlib, biopython, openpyxl;
python-docx is needed by `make_krause_table.py`). Grid and ABC runs
were executed on a SLURM cluster with the `.sbatch` scripts included
here; every script can also be run on a laptop for single combinations.
Collected outputs (`.npz` / `.csv`) are included, so all figures and
quoted numbers can be regenerated locally without rerunning the cluster
arrays. Sequence inputs for the DFE/RNA scripts are in `data/`.

## Demography and conventions used throughout

All population sizes are haploid locus copy numbers ("uniparental"
sizes). Main demography: modern lineage NM-uniparental = 7000
throughout; archaic lineage at NA-uniparental = 2250 (central) with
1500 and 3000 brackets during 17,483 generations of isolation
(separation 26,224 generations ago, mixture 8,741, sampling 1,748, at
28.6 yr/generation). Petr et al. comparison: NM = 5000, NA = 500,
10,489 generations of isolation. Unless stated otherwise, results
integrate over the DFE/mutation-rate priors and pool the three archaic
sizes with equal weight. Fixation probabilities score each draw on the
fixation curve of its NA-paired post-mixture size
(NN-uniparental = 500/750/1000 for NA = 1500/2250/3000; 500 for the
Petr et al. comparison). The companion fixation analysis rounds the
central post-mixture size 725 to 750.

## Core simulation

- `compute_fn.py` — missense fraction from NC_012920 (uses biopython;
  reads `data/NC_012920.gb`).
- `ratchet_dfe.py` — core forward Wright-Fisher simulation of Muller's
  Ratchet under a gamma DFE (one combination per call; `--anchor-ne`
  sets the DFE reference Ne). `ratchet_dfe_annotated.py` is a
  line-by-line commented reading copy.

## DFE inference (Appendix A)

- `save_dfe_draws.py` — exports the joint (beta, |S-bar|) bootstrap
  draws of the Denisova-calibrated DFE fit to `dfe_draws.npz`, the
  prior sampled by the ABC and by the parameter-integrated
  simulations. (`dfe_draws.npz` is included.)
- `pew_replicate.py` — reimplementation of the Piganeau & Eyre-Walker
  (2003) model-2 estimator, validated against their published Table 3
  (Table S1.A1 of the note).
- `pew_rna.py` — the same estimator extended to the mitochondrial tRNA
  and rRNA genes (gnomAD African polymorphism, outgroup divergence),
  with sample-depth projection and bootstrap (Table S1.A2).
- `mt_dfe_sfs.py` — gnomAD chrM VCF reader and mutational-opportunity
  helpers imported by `pew_rna.py` and `save_dfe_draws.py`.
- `rna_constraint.py` — divergence-based constraint of the tRNA and
  rRNA genes against the synonymous neutral clock (rCRS vs Denisovan,
  Neandertal and chimpanzee mtDNA, spectrum-weighted opportunities;
  NCBI accessions in `data/mtseq/`, included, with NC_012920.gb and
  the gnomAD chrM VCF): the source of the measured RNA deleterious
  fraction (0.76-0.80) behind the RNA-inclusive U = 0.0093.

## Figure S1.1 (validation against the original Perl simulation)

- `validate_delta.py` / `make_grid_replicate.py` /
  `run_slurm_replicate.sbatch` / `collect_replicate.py` /
  `plot_replicate.py` — validation against the original Perl
  simulation of ref. 7 (Figure S1.1). Collected output:
  `new_replicate.csv`. The original grid values are in
  `2_26_2026 Muller's Ratchet simulations.xlsx`.

## Unconstrained mtDNA bounds (Table S1.3, prior-predictive rows)

- `make_grid_v22.py` / `run_task.py` / `run_slurm.sbatch` /
  `merge_chunks.py` — the current-demography mtDNA forward grid
  (NM = 7000, NA = 1500/2250/3000, plus the Petr et al. comparison).
- `run_part1_bound.py` — the unconstrained 95% upper bounds in Table
  S1.3, integrating over rate and DFE uncertainty.

## ABC and Figure S1.2 (Appendix B)

- `abc_joint_rna.py` / `run_slurm_joint_rna.sbatch` /
  `run_slurm_joint_cod.sbatch` / `collect_joint_rna.py` — the
  RNA-inclusive ABC of Appendix B and the coding-only companion.
  Coding parameters are constrained by the nonsynonymous acceptance
  statistic, the synonymous clock constrains the shared mutation rate,
  and the RNA deleterious input contributes to fitness but not to
  acceptance. Collected output: `joint_rna_{1500,2250,3000,petr500}.npz`
  and `joint_cod_{...}.npz` (included).
- `plot_abc_fig2.py` — renders Figure S1.2 (`fig2_mtdna_abc.png`) from
  those collected posteriors and `fixation_curves_mtdna.csv`.

## Y chromosome ladders (Figure S1.3, Table S1.3)

- `make_grid_y_draw.py` / `run_task_ydraw.py` /
  `run_slurm_ydraw.sbatch` / `collect_y_draw.py` — the
  parameter-integrated ladder used in the note: each of 300 replicates
  per target and archaic size draws (beta, S-bar, U) from the priors
  before simulating. `--conv stated` is the main analysis
  (s = S-bar/Nref = 0.0425); `--conv hemi` is the hemizygous
  sensitivity.
- `plot_y_fig3_violin.py` — renders Figure S1.3 (`fig3_y_gap_violin.png`)
  from the parameter-integrated collection, pooling the three archaic
  sizes with equal weight.
- `y_numbers_stated.py` — derives the Y numbers quoted in the Results
  and Table S1.3 from the collected ladders.
- `build_ychr_cds_map.py` — builds the hg19 X-degenerate codon map
  (`ychr_cds_map.json`) that gives the 33,156 bp of complete codons in
  Table S1.1. The collected map is included.
- `sweep_y_U.py` — sweeps the whole-Y deleterious input to find the
  coding-equivalent constrained sequence at which the gap reaches 1%.
- Collected outputs included: `y_draw_stated_reps.npz`,
  `y_draw_hemi_reps.npz` and the `y_*_summary.csv` files.

## Tables

- `make_krause_table.py` — formats the per-lineage synonymous and
  nonsynonymous mtDNA substitution counts extracted from Krause et
  al. (2010), Supplementary Figure 5, used as the observed statistics
  of the ABC (writes a standalone .docx table; requires python-docx).

## Fixation curves (companion analysis)

- `fixation_curves_code/` — generates `fixation_curves_mtdna.csv`, the
  probability that introgressed modern mtDNA replaces the archaic type
  as a function of selective advantage s and mtDNA effective size,
  from forward Wright-Fisher simulations of a single 5% pulse into a
  constant-size population. Supporting modules `panmictic_sim.py` and
  `panm3_run.py`; cluster entry `panm3_submit.py` / `panm3_collect.py`;
  single-machine entry `run_local.py`. See the README in that folder.
- `fixation_curves_mtdna.csv` — kept at the top level because
  `plot_abc_fig2.py`, `plot_y_fig3_violin.py` and `y_numbers_stated.py`
  read it from the working directory. All fixation probabilities in the
  note interpolate this table.

## Reproducing the main results

```
# Figure S1.2 (mtDNA ABC posteriors + fixation panel)
python plot_abc_fig2.py            # reads joint_rna_*.npz, joint_cod_*.npz,
                                   # fixation_curves_mtdna.csv

# Figure S1.3 (Y violin plot, parameter-integrated, pooled archaic sizes)
python plot_y_fig3_violin.py       # reads y_draw_stated_reps.npz,
                                   # fixation_curves_mtdna.csv

# Numbers quoted in Results / Table S1.3 for the Y
python y_numbers_stated.py

# Figure S1.1 (Python vs original Perl grid)
python plot_replicate.py           # reads new_replicate.csv and the xlsx

# Full rerun on a SLURM cluster (example: parameter-integrated Y ladder)
python make_grid_y_draw.py --conv stated
sbatch --array=0-319 run_slurm_ydraw.sbatch
python collect_y_draw.py --outdir results_y_draw_stated \
    --params params_y_draw_stated.tsv --out y_draw_stated

# Fixation curves (companion analysis; ~130 CPU-hours, or --quick smoke test)
cd fixation_curves_code && python run_local.py --quick && python panm3_collect.py
```
