## Supplementary Information section 1 code base for "Hypothesis: A modern human range expansion ∼300,000 years ago had a critical role in Neandertal origins": rna_constraint.py

```
"""
Empirical constraint of mtDNA RNA genes (tRNA, rRNA) relative to protein-coding
sites, measured on the same lineages and timescale as the ratchet analysis.

Motivation
----------
The write-up currently assumes that a fraction 0.4-0.8 of mutations in the ~4 kb
of mitochondrial RNA genes are deleterious, and treats them as if drawn from the
protein missense DFE. That fraction was asserted, not measured. This script
measures it.

Method
------
Divergence-based constraint. Synonymous sites in mtDNA are ~neutral, so they
provide the local neutral clock. For each site class,

    f_del(class) = 1 - d_class / d_syn

where d is substitutions per *mutational-opportunity-equivalent site*.

The opportunity normalisation matters: mtDNA has an extreme transition bias
(Ti:Tv ~ 24), and synonymous changes are concentrated at 3rd codon positions
whose transitions are disproportionately synonymous. Comparing raw per-bp rates
would therefore confound constraint with codon composition. We instead weight
every possible substitution by the spectrum (as compute_fn.py does):

    weight(transition)   = 2 * TiTv
    weight(transversion) = 1

Every site has exactly 1 transition + 2 transversion opportunities, so the total
weight per site is (2*TiTv + 2) regardless of its base. For classes where any
change is potentially deleterious (tRNA, rRNA, non-coding) the number of
"site-equivalents" is therefore just the number of callable sites. For
synonymous / nonsynonymous we sum only the weights of changes of that class and
divide by (2*TiTv + 2). Rates per site-equivalent are then directly comparable.

Alignment is reference-based: each sequence is globally aligned to the rCRS and
projected onto rCRS coordinates, which is what we need since all annotation is
in rCRS coordinates.

Data: NCBI accessions in data/mtseq/ (fetched separately).
"""
import argparse
import math
import os
from collections import defaultdict

from Bio import Align, SeqIO
from Bio.Seq import Seq

GB = "data/NC_012920.gb"
SEQDIR = "data/mtseq"
TABLE = 2                 # vertebrate mitochondrial code
TITV = 24.3               # deCODE coding Ti:Tv (same value used in compute_fn.py)
BASES = ("A", "C", "G", "T")
TRANSITION = {"A": "G", "G": "A", "C": "T", "T": "C"}

SAMPLES = [
    ("Denisova",        "FN673705.1"),   # Krause et al. 2010 Denisova 3
    ("Neanderthal_Mez", "FM865411.1"),
    ("Neanderthal_Alt", "KC879692.1"),
    ("Chimp",           "NC_001643.1"),
]

_codon_cache = {}


def translate(codon):
    aa = _codon_cache.get(codon)
    if aa is None:
        aa = str(Seq(codon).translate(table=TABLE))
        _codon_cache[codon] = aa
    return aa


def weight(ref, alt, titv):
    """Relative rate of one specific substitution. Each site has 1 transition
    and 2 transversion opportunities, so per-opportunity rates satisfy
    r_ti/r_tv = 2*titv. titv=0.5 recovers a uniform spectrum, which is the
    implicit assumption of Nei-Gojobori site counting."""
    return 2.0 * titv if alt == TRANSITION[ref] else 1.0


def per_site_weight(titv):
    return 2.0 * titv + 2.0


def revcomp(b):
    return {"A": "T", "T": "A", "C": "G", "G": "C"}[b]


def build_site_table(rec):
    """Map each 1-based rCRS position to a functional class, and for coding
    positions to the class of each possible genomic-strand substitution.

    Positions covered by more than one feature type (or two CDS) are dropped, so
    that every counted site belongs unambiguously to one class."""
    seq = str(rec.seq).upper()
    L = len(seq)

    owners = defaultdict(list)          # pos -> list of feature types
    cds_owner = defaultdict(list)       # pos -> list of gene names

    for f in rec.features:
        if f.type not in ("CDS", "tRNA", "rRNA", "D-loop"):
            continue
        for p in f.location:            # 0-based
            owners[p + 1].append(f.type)
            if f.type == "CDS":
                cds_owner[p + 1].append(f.qualifiers.get("gene", ["?"])[0])

    site = {}
    for p in range(1, L + 1):
        types = set(owners.get(p, []))
        if len(types) != 1:
            continue                    # unannotated or ambiguous overlap
        t = types.pop()
        if t == "CDS" and len(cds_owner[p]) != 1:
            continue                    # ATP8/ATP6, ND4L/ND4 overlaps
        if seq[p - 1] not in BASES:
            continue                    # rCRS 'N' placeholder at 3107
        site[p] = {"cls": t}

    # ---- annotate coding positions with codon context ------------------------
    for f in rec.features:
        if f.type != "CDS":
            continue
        minus = f.location.strand == -1
        pos = [p + 1 for p in f.location]        # genomic, transcription order
        if minus:
            pos = pos[::-1]
        nt = str(f.extract(rec.seq)).upper()
        ncomplete = len(nt) // 3
        for ci in range(ncomplete):
            codon = nt[ci * 3:ci * 3 + 3]
            if any(b not in BASES for b in codon):
                continue
            gpos = pos[ci * 3:ci * 3 + 3]
            for k in range(3):
                p = gpos[k]
                s = site.get(p)
                if s is None or s["cls"] != "CDS":
                    continue
                ref_genomic = seq[p - 1]
                alt_class = {}
                for alt_g in BASES:
                    if alt_g == ref_genomic:
                        continue
                    alt_t = revcomp(alt_g) if minus else alt_g
                    mutated = codon[:k] + alt_t + codon[k + 1:]
                    ra, aa = translate(codon), translate(mutated)
                    if ra == aa:
                        cl = "synonymous"
                    elif aa == "*":
                        cl = "nonsense"
                    elif ra == "*":
                        cl = "stoploss"
                    else:
                        cl = "missense"
                    alt_class[alt_g] = cl
                s["alt_class"] = alt_class
                s["gene"] = f.qualifiers.get("gene", ["?"])[0]
                s["codon_id"] = (s["gene"], ci)

    # coding sites that never got codon context (incomplete trailing codon)
    for p, s in list(site.items()):
        if s["cls"] == "CDS" and "alt_class" not in s:
            del site[p]
    return site, seq


def align_to_ref(ref, other):
    """Global align `other` to `ref`; return dict rCRS 1-based pos -> base.
    Positions deleted in `other` are simply absent."""
    aligner = Align.PairwiseAligner()
    aligner.mode = "global"
    aligner.match_score = 2
    aligner.mismatch_score = -3
    aligner.open_gap_score = -7
    aligner.extend_gap_score = -1
    # do not penalise the ends: mtDNA entries differ in start offset conventions
    aligner.target_end_gap_score = 0
    aligner.query_end_gap_score = 0
    aln = aligner.align(ref, other)[0]

    mapping = {}
    for (rs, re_), (qs, qe) in zip(aln.aligned[0], aln.aligned[1]):
        for off in range(re_ - rs):
            mapping[rs + off + 1] = other[qs + off]
    return mapping


def analyse(site, refseq, mapping, titv):
    """Count substitutions and opportunity per class over callable sites."""
    obs = defaultdict(float)
    opp = defaultdict(float)
    hits_per_codon = defaultdict(int)
    psw = per_site_weight(titv)

    for p, s in site.items():
        alt = mapping.get(p)
        if alt is None or alt not in BASES:
            continue                      # gap / ambiguous in this sample
        ref = refseq[p - 1]
        cls = s["cls"]

        if cls == "CDS":
            ac = s["alt_class"]
            # opportunity: spectrum weight of each possible change, by class
            for alt_g, cl in ac.items():
                key = "synonymous" if cl == "synonymous" else "nonsynonymous"
                opp[key] += weight(ref, alt_g, titv) / psw
            if alt != ref:
                cl = ac.get(alt)
                if cl is None:
                    continue
                key = "synonymous" if cl == "synonymous" else "nonsynonymous"
                obs[key] += 1
                hits_per_codon[s["codon_id"]] += 1
        else:
            key = {"tRNA": "tRNA", "rRNA": "rRNA", "D-loop": "D-loop"}[cls]
            opp[key] += 1.0               # any change counts; 1 site-equivalent
            if alt != ref:
                obs[key] += 1
    multi = sum(1 for v in hits_per_codon.values() if v > 1)
    return obs, opp, multi


def rna_multiplier(r):
    """Deleterious mutational input in tRNA+rRNA relative to nonsynonymous
    coding. For each class the input removed by selection per unit time is
    (opportunity in site-equivalents) x f_del, since every class is measured
    against the same synonymous clock."""
    d, opp = r["d"], r["opp"]
    dsyn = r["dsyn"]

    def contrib(k):
        return opp[k] * (1 - d[k] / dsyn)

    return (contrib("tRNA") + contrib("rRNA")) / contrib("nonsynonymous")


def poisson_ci(k):
    """Approximate 95% CI on a Poisson count (Byar)."""
    if k == 0:
        return 0.0, 3.0
    lo = k * (1 - 1 / (9 * k) - 1.96 / (3 * math.sqrt(k))) ** 3
    hi = (k + 1) * (1 - 1 / (9 * (k + 1)) + 1.96 / (3 * math.sqrt(k + 1))) ** 3
    return lo, hi


def report(name, obs, opp, multi, titv, label):
    order = ["synonymous", "nonsynonymous", "tRNA", "rRNA", "D-loop"]
    d = {}
    print(f"\n--- rCRS vs {name}   [{label}]   "
          f"(codons with >1 substitution: {multi})")
    print(f"{'class':<15}{'subs':>7}{'site-equiv':>12}{'d (per site)':>14}"
          f"{'d/d_syn':>10}{'f_del':>9}{'95% CI on f_del':>22}")
    dsyn = obs["synonymous"] / opp["synonymous"]
    for k in order:
        if opp[k] == 0:
            continue
        dk = obs[k] / opp[k]
        d[k] = dk
        ratio = dk / dsyn
        lo, hi = poisson_ci(obs[k])
        rlo, rhi = (lo / opp[k]) / dsyn, (hi / opp[k]) / dsyn
        ci = (f"[{max(0.0, 1 - rhi):.2f}, {1 - rlo:.2f}]"
              if k != "synonymous" else "(reference)")
        print(f"{k:<15}{obs[k]:>7.0f}{opp[k]:>12.1f}{dk:>14.5f}"
              f"{ratio:>10.3f}{1 - ratio:>9.2f}{ci:>22}")
    return {"d": d, "dNdS": d["nonsynonymous"] / dsyn,
            "obs": dict(obs), "opp": dict(opp), "dsyn": dsyn}


def main():
    ap = argparse.ArgumentParser(description=__doc__,
                                 formatter_class=argparse.RawDescriptionHelpFormatter)
    ap.add_argument("--samples", nargs="*", default=None,
                    help="subset of sample names to analyse")
    args = ap.parse_args()

    rec = SeqIO.read(GB, "genbank")
    site, refseq = build_site_table(rec)

    counts = defaultdict(int)
    for s in site.values():
        counts[s["cls"]] += 1
    print(f"rCRS {rec.id}: {len(refseq):,} bp")
    print("Unambiguously classified sites: " +
          ", ".join(f"{k}={v:,}" for k, v in sorted(counts.items())))

    # Two opportunity conventions:
    #  - uniform (titv=0.5) reproduces Nei-Gojobori counting, which is what most
    #    published mtDNA dN/dS values use, so it is the literature-comparable one
    #  - deCODE (titv=24.3) is mutationally correct for mtDNA and is the
    #    convention used everywhere else in this project (compute_fn.py)
    conventions = [(0.5, "uniform spectrum ~ Nei-Gojobori"),
                   (TITV, f"Ti:Tv={TITV} (deCODE)")]

    results = defaultdict(dict)
    for name, acc in SAMPLES:
        if args.samples and name not in args.samples:
            continue
        path = os.path.join(SEQDIR, f"{acc}.fa")
        other = str(SeqIO.read(path, "fasta").seq).upper()
        mapping = align_to_ref(refseq, other)
        print(f"\n{'=' * 78}\n=== {name} ({acc})")
        for titv, label in conventions:
            obs, opp, multi = analyse(site, refseq, mapping, titv)
            results[name][label] = report(name, obs, opp, multi, titv, label)

    # ---- literature cross-check on the coding sequence -----------------------
    print("\n" + "=" * 78)
    print("CODING-SEQUENCE VALIDATION (dN/dS), for comparison with published "
          "values")
    print(f"{'comparison':<20}" +
          "".join(f"{lab:>36}" for _, lab in conventions))
    for name in results:
        row = "".join(f"{results[name][lab]['dNdS']:>36.3f}"
                      for _, lab in conventions)
        print(f"{name:<20}{row}")
    print("\nPublished anchors: human-chimp mtDNA protein-coding dN/dS is "
          "generally\nreported near 0.05-0.10 by Nei-Gojobori-style counting; "
          "within-human and\nvery recent lineages run higher (~0.2-0.4) because "
          "segregating slightly\ndeleterious variants have not yet been purged.")

    # ---- deleterious input multiplier contributed by the RNA genes -----------
    print("\n" + "=" * 78)
    print("DELETERIOUS-INPUT MULTIPLIER from adding tRNA+rRNA to the coding "
          "genes")
    print("(mutations removed by selection per unit time, class opportunity x "
          "f_del,\nrelative to nonsynonymous coding alone)")
    for _, lab in conventions:
        print(f"\n  [{lab}]")
        for name in results:
            r = results[name][lab]
            print(f"    {name:<18} RNA/nonsyn = {rna_multiplier(r):.2f}"
                  f"   -> U multiplier {1 + rna_multiplier(r):.2f}x")


if __name__ == "__main__":
    main()
```
