## Supplementary Information section 1 code base for "Hypothesis: A modern human range expansion ∼300,000 years ago had a critical role in Neandertal origins": run_part1_bound.py

```
"""
Part 1 (unconstrained) 95% confident UPPER BOUND on the modern-archaic fitness
gap, integrating over uncertainty in the mutation rate and the DFE.

Motivation
----------
Part 1 (ratchet_dfe.py + the grids) reports the gap at CENTRAL parameter values
and sweeps one axis at a time. This script instead produces a single summary
number per (genome, archaic size): the prior-predictive distribution of the gap
when the uncertain inputs are drawn from their published 95% confidence
intervals, and its 95th percentile.

This is the unconstrained analogue of the Part 2 ABC (abc_joint.py), which draws
the same parameters from the same triangular priors but then CONDITIONS on the
observed substitution counts. Here there is no conditioning: we simply propagate
input uncertainty through the forward simulation.

What is (and is not) sampled
----------------------------
- DFE shape beta and DFE mean |Sbar|: drawn per replicate from triangular priors
  spanning each parameter's 95% CI (mode = central estimate). This is PARAMETER
  uncertainty. (Drawing individual selection coefficients s ~ Gamma(beta, .) is
  the modelled biology and already happens inside every simulation; it is not
  "uncertainty" and is not what the percentile integrates over.)
- Mutation rate: the deleterious input U is drawn per replicate from a triangular
  prior over its published rate CI.
- Each draw runs ONE modern population (N=7500) and ONE archaic population (N_A),
  so the recorded per-draw gap = mean(w_modern) - mean(w_archaic) also carries a
  small amount of evolutionary (drift) noise on top of the parameter spread. The
  modern realization is shared across the two archaic sizes within a draw, so the
  1500 vs 3000 columns are paired.

Estimand: the population mean-fitness gap (same quantity Part 1 Figures report),
whose 95th percentile across draws is the reported "95% confident upper bound".

Run (laptop, multiprocessing):
    ./.venv/bin/python run_part1_bound.py --draws 500
    ./.venv/bin/python run_part1_bound.py --draws 500 --genome mtDNA
"""
import argparse
import csv
import time
from multiprocessing import Pool, cpu_count

import numpy as np

from ratchet_dfe import simulate_one, NONSENSE_FRACTION

T = 17500
N_MODERN = 7500
NARCH = (1500, 3000)

# --- priors: (low, mode, high) triangular over each parameter's 95% CI ----------
# mtDNA: Piganeau & Eyre-Walker (2003) DFE; Fu et al. (2013) rate. Anchor Ne=7500.
MT = dict(
    anchor_ne=7500.0,
    beta_tri=(0.36, 0.39, 0.50),
    sbar_tri=(420.0, 1400.0, 2400.0),
    u_tri=(0.0038, 0.0064, 0.0091),
)
# Y: autosomal amino-acid DFE (Eyre-Walker et al. 2006) rescaled to the Y; Y point
# rate ~3.0e-8 (2.5-3.5) x 100 kb x 0.72 nonsynonymous. Anchor Ne=10000.
Y = dict(
    anchor_ne=10000.0,
    beta_tri=(0.19, 0.23, 0.27),
    sbar_tri=(225.0, 425.0, 766.0),
    u_tri=(0.0018, 0.00216, 0.0025),
)
GENOMES = {"mtDNA": MT, "Y": Y}


def one_draw(job):
    """One prior draw: sample (beta, Sbar, U), run modern once + both archaic.

    Returns a dict with the drawn parameters and the gap for each archaic size.
    """
    genome, seed = job
    cfg = GENOMES[genome]
    rng = np.random.default_rng(seed)

    beta = float(rng.triangular(*cfg["beta_tri"]))
    Sbar = float(rng.triangular(*cfg["sbar_tri"]))
    U_mis = float(rng.triangular(*cfg["u_tri"]))
    U_non = U_mis * NONSENSE_FRACTION
    scale = (Sbar / (2.0 * cfg["anchor_ne"])) / beta

    w_mod, _ = simulate_one(N_MODERN, T, U_mis, U_non, beta, scale, rng)
    out = dict(genome=genome, seed=seed, beta=beta, Sbar=Sbar, U_mis=U_mis,
               w_modern=w_mod)
    for Na in NARCH:
        w_arch, _ = simulate_one(Na, T, U_mis, U_non, beta, scale, rng)
        out[f"gap_{Na}"] = w_mod - w_arch
    return out


def summarize(draws, genome, Na):
    g = np.array([d[f"gap_{Na}"] for d in draws if d["genome"] == genome]) * 100.0
    return dict(
        genome=genome, N_archaic=Na, n_draws=g.size,
        median=float(np.median(g)), mean=float(g.mean()),
        p95=float(np.percentile(g, 95)), p975=float(np.percentile(g, 97.5)),
        frac_over_1pct=float(np.mean(g > 1.0)),
    )


def main():
    ap = argparse.ArgumentParser(description=__doc__,
                                 formatter_class=argparse.RawDescriptionHelpFormatter)
    ap.add_argument("--draws", type=int, default=500,
                    help="prior draws per genome (each = 1 modern + 2 archaic runs)")
    ap.add_argument("--genome", choices=list(GENOMES) + ["both"], default="both")
    ap.add_argument("--seed0", type=int, default=90000)
    ap.add_argument("--procs", type=int, default=0, help="0 = all cores")
    ap.add_argument("--out", default="part1_bound.csv")
    args = ap.parse_args()

    genomes = list(GENOMES) if args.genome == "both" else [args.genome]
    jobs = [(gname, args.seed0 + i)
            for gi, gname in enumerate(genomes)
            for i in range(gi * 10_000_000, gi * 10_000_000 + args.draws)]

    nproc = args.procs or min(cpu_count(), len(jobs))
    print(f"Part 1 upper-bound: {args.draws} draws x {len(genomes)} genome(s) "
          f"= {len(jobs)} draws on {nproc} procs (T={T})...\n")
    t0 = time.time()
    with Pool(nproc) as pool:
        draws = pool.map(one_draw, jobs)
    dt = time.time() - t0

    rows = [summarize(draws, g, Na) for g in genomes for Na in NARCH]
    with open(args.out, "w", newline="") as f:
        w = csv.DictWriter(f, fieldnames=list(rows[0].keys()))
        w.writeheader()
        w.writerows(rows)
    # also keep the raw per-draw gaps for reproducible percentiles/plots
    raw = args.out.replace(".csv", "_draws.csv")
    with open(raw, "w", newline="") as f:
        cols = ["genome", "seed", "beta", "Sbar", "U_mis", "w_modern",
                "gap_1500", "gap_3000"]
        w = csv.DictWriter(f, fieldnames=cols, extrasaction="ignore")
        w.writeheader()
        w.writerows(draws)

    print(f"{'genome':7s} {'N_A':>5s} {'median%':>8s} {'mean%':>7s} "
          f"{'95%UB':>7s} {'97.5%':>7s} {'P(gap>1%)':>10s}")
    for r in rows:
        print(f"{r['genome']:7s} {r['N_archaic']:>5d} {r['median']:>8.3f} "
              f"{r['mean']:>7.3f} {r['p95']:>7.3f} {r['p975']:>7.3f} "
              f"{r['frac_over_1pct']:>10.3f}")
    print(f"\nWrote {args.out} and {raw}  [{dt:.0f}s]")


if __name__ == "__main__":
    main()
```
