## Supplementary Information section 1 code base for "Hypothesis: A modern human range expansion ∼300,000 years ago had a critical role in Neandertal origins": run_slurm.sbatch

```
#!/bin/bash
# SLURM array job for the 300-replicate ratchet grids on O2.
#
#   cd /n/groups/reich/DAVID/RATCHET
#   python make_grid_300.py            # mtDNA: params_300.tsv, 700 tasks
#   python make_grid_extra_300.py      # Y+RNA: params_extra_300.tsv, 200 tasks
#   mkdir -p logs results300 results_extra300
#
#   sbatch --array=0-699%200 run_slurm.sbatch
#   sbatch --export=ALL,PARAMS=params_extra_300.tsv,OUTDIR=results_extra300 \
#          --array=0-199%200 run_slurm.sbatch
#
# One array task = one row of the params file = 30 replicates of one parameter
# combination, about 12-15 minutes on one core. Ten chunks make up each
# combination; merge_chunks.py pools them afterwards.
#
# Which grid to run is taken from the environment rather than hard-coded, so one
# script serves both, and the array index likewise, so the same script serves a
# full submission and a resubmission of stragglers (sbatch --array=3,17,412 ...).
#
# --export=ALL,... rather than --export=PARAMS,...: the bare form would replace
# the whole environment rather than adding to it, which strips PATH and the
# conda machinery and leaves the task unable to find python.
#SBATCH --job-name=ratchet300
#SBATCH --partition=short
#SBATCH --time=0-01:00:00
#SBATCH --cpus-per-task=1
#SBATCH --mem=2G
#SBATCH --output=logs/ratchet_%A_%a.out
#SBATCH --error=logs/ratchet_%A_%a.err

set -euo pipefail

# numpy must not spawn threads: each task owns exactly one core, and a threaded
# BLAS here would oversubscribe the node without speeding anything up, since the
# arrays are a few thousand elements.
export OMP_NUM_THREADS=1
export OPENBLAS_NUM_THREADS=1
export MKL_NUM_THREADS=1

if command -v conda >/dev/null 2>&1; then
    eval "$(conda shell.bash hook)"
    conda activate ratchet
elif [ -f .venv/bin/activate ]; then
    source .venv/bin/activate
else
    echo "no conda env 'ratchet' and no .venv; run setup_cluster.sh first" >&2
    exit 1
fi

PARAMS="${PARAMS:-params_300.tsv}"
OUTDIR="${OUTDIR:-results300}"

echo "task ${SLURM_ARRAY_TASK_ID} on $(hostname) at $(date)"
echo "params=${PARAMS} outdir=${OUTDIR}"
python run_task.py --idx "${SLURM_ARRAY_TASK_ID}" \
                   --params "${PARAMS}" --outdir "${OUTDIR}"
echo "done at $(date)"
```
