## Supplementary Information section 1 code base for "Hypothesis: A modern human range expansion ∼300,000 years ago had a critical role in Neandertal origins": run_slurm_joint_cod.sbatch

```
#!/bin/bash
#SBATCH --job-name=abc_joint_cod
#SBATCH --partition=short
#SBATCH --array=0-799             # 800 tasks (3 main Nanc + petr) x 200; unthrottled
#SBATCH --time=12:00:00           # Nanc=3000 tasks can exceed 3 h on O2 nodes
#SBATCH --mem=2G
#SBATCH --cpus-per-task=1
#SBATCH --output=logs/joint_cod_%A_%a.out

# Coding-only companion to run_slurm_joint_rna.sbatch: same engine, same v22
# demography (main + petr scenarios), same Denisova 'all' DFE bootstrap prior,
# same acceptance data, but --coding-only (no RNA input). Needed for Figure 2
# panel (e): "coding only, no ABC" and "coding only, ABC-accepted".
#
#   1. python make_grid_joint.py              # params_joint.tsv (already there)
#   2. sbatch run_slurm_joint_cod.sbatch      # 800-task array on 'short'
#   3. python collect_joint_rna.py --indir results_joint_cod --out joint_cod
# Then copy joint_cod_{1500,2250,3000}.npz + joint_cod_petr500.npz back.

set -euo pipefail
mkdir -p logs results_joint_cod

source activate ratchet 2>/dev/null \
  || conda activate ratchet 2>/dev/null \
  || source .venv/bin/activate 2>/dev/null \
  || true

# tr strips the CR that python's csv module puts at line ends; bash arithmetic
# on SEED below fails otherwise
ROW=$(awk -F'\t' -v i="${SLURM_ARRAY_TASK_ID}" '$1==i {print; exit}' params_joint.tsv | tr -d '\r')
if [[ -z "${ROW}" ]]; then
  echo "No params_joint row with idx=${SLURM_ARRAY_TASK_ID}"; exit 1
fi
read -r IDX SCEN NANC K SEED <<< "${ROW}"

echo "Task ${IDX}: scen=${SCEN} Nanc=${NANC} K=${K} seed=$((SEED + 40000)) (coding-only)"
python abc_joint_rna.py --scenario "${SCEN}" --Nanc "${NANC}" --K "${K}" \
  --seed "$((SEED + 40000))" --coding-only --draws dfe_draws.npz \
  --out "results_joint_cod/joint_${IDX}.npz"
```
