## Supplementary Information section 1 code base for "Hypothesis: A modern human range expansion ∼300,000 years ago had a critical role in Neandertal origins": run_slurm_joint_rna.sbatch

```
#!/bin/bash
#SBATCH --job-name=abc_joint_rna
#SBATCH --partition=short
#SBATCH --array=0-799             # 800 tasks (3 main Nanc + petr) x 200; unthrottled
#SBATCH --time=12:00:00           # Nanc=3000 tasks can exceed 3 h on O2 nodes
#SBATCH --mem=2G
#SBATCH --cpus-per-task=1
#SBATCH --output=logs/joint_rna_%A_%a.out

# RNA-inclusive joint single-pair ABC (abc_joint_rna.py), v22 demography:
#   main: NM=7000 (DFE anchor 7000), Nanc in {1500, 2250, 3000},
#         Tsep/Tmix = 26,224/8,741 generations ago (17,483 gens isolation)
#   petr: NM=5000, Nanc=500, Tsep/Tmix = 20,979/10,490 (10,489 gens isolation)
# Coding DFE prior: Denisova matched-depth 'all' bootstrap (dfe_draws.npz);
# RNA deleterious input added on top (same DFE, target scaled by
# U_incl/U_coding = 0.0093/0.0064), fitness-reducing but EXCLUDED from the
# nonsynonymous acceptance statistic.
#
#   1. python make_grid_joint.py             # writes params_joint.tsv (800 rows)
#   2. sbatch run_slurm_joint_rna.sbatch     # 800-task array on 'short'
#   3. python collect_joint_rna.py           # -> joint_rna_{1500,2250,3000}.npz
#                                            #    + joint_rna_petr500.npz
# Then copy the merged files back and run:
#   python combine_joint.py --prefix joint_rna

set -euo pipefail
mkdir -p logs results_joint_rna

source activate ratchet 2>/dev/null \
  || conda activate ratchet 2>/dev/null \
  || source .venv/bin/activate 2>/dev/null \
  || true

# tr guards against CR at line ends (bash read would keep it in SEED)
ROW=$(awk -F'\t' -v i="${SLURM_ARRAY_TASK_ID}" '$1==i {print; exit}' params_joint.tsv | tr -d '\r')
if [[ -z "${ROW}" ]]; then
  echo "No params_joint row with idx=${SLURM_ARRAY_TASK_ID}"; exit 1
fi
read -r IDX SCEN NANC K SEED <<< "${ROW}"

echo "Task ${IDX}: scen=${SCEN} Nanc=${NANC} K=${K} seed=${SEED} (RNA-inclusive)"
python abc_joint_rna.py --scenario "${SCEN}" --Nanc "${NANC}" --K "${K}" \
  --seed "${SEED}" --draws dfe_draws.npz \
  --out "results_joint_rna/joint_${IDX}.npz"
```
