## Supplementary Information section 1 code base for "Hypothesis: A modern human range expansion ∼300,000 years ago had a critical role in Neandertal origins": run_slurm_replicate.sbatch

```
#!/bin/bash
#SBATCH --job-name=ratchet_rep
#SBATCH --partition=short
#SBATCH --array=0-107%64          # 108 grid points (6 mu x 9 s x 2 archaic sizes)
#SBATCH --time=00:30:00           # 30 reps/task at T=17500; 30 min is safe
#SBATCH --mem=1G
#SBATCH --cpus-per-task=1
#SBATCH --output=logs/rep_%A_%a.out

# Re-run every Extended Data Figure 1 grid point through the NEW framework
# (delta DFE) so we can scatter new-vs-original.
#   1. python make_grid_replicate.py     # writes params_replicate.tsv (108 rows)
#   2. sbatch run_slurm_replicate.sbatch # 108-task array on 'short'
#   3. python collect_replicate.py       # writes new_replicate.csv
# Then copy new_replicate.csv back to the laptop for plotting against the Excel.

set -euo pipefail
mkdir -p logs results_replicate

source activate ratchet 2>/dev/null \
  || conda activate ratchet 2>/dev/null \
  || source .venv/bin/activate 2>/dev/null \
  || true

ROW=$(awk -F'\t' -v i="${SLURM_ARRAY_TASK_ID}" '$1==i {print; exit}' params_replicate.tsv)
if [[ -z "${ROW}" ]]; then
  echo "No params_replicate row with idx=${SLURM_ARRAY_TASK_ID}"; exit 1
fi
read -r IDX NM NA T MU S REPS SEED <<< "${ROW}"

echo "Task ${IDX}: N_A=${NA} mu=${MU} s=${S} reps=${REPS}"

OUT=$(python run_replicate_py.py "${NM}" "${NA}" "${T}" "${MU}" "${S}" "${REPS}" "${SEED}")
read -r WM WA <<< "${OUT}"

printf '%s\t%s\t%s\t%s\t%s\t%s\n' "${IDX}" "${NA}" "${MU}" "${S}" "${WM}" "${WA}" \
  > "results_replicate/rep_${IDX}.tsv"
echo "  -> modern=${WM} archaic=${WA}"
```
