## Supplementary Information section 1 code base for "Hypothesis: A modern human range expansion ∼300,000 years ago had a critical role in Neandertal origins": run_slurm_ydraw.sbatch

```
#!/bin/bash
# SLURM array for the parameter-integrated Y target-size ladder.
#
#   cd /n/groups/reich/DAVID/RATCHET
#   python make_grid_y_draw.py
#   mkdir -p logs results_y_draw
#   sbatch --export=ALL,PARAMS=params_y_draw.tsv,OUTDIR=results_y_draw \
#          --job-name=y_draw --array=0-279 run_slurm_ydraw.sbatch
#
# Identical to run_slurm.sbatch except that it calls run_task_ydraw.py,
# which draws (beta, Sbar, U) per replicate from triangular priors.
#SBATCH --job-name=y_draw
#SBATCH --partition=short
#SBATCH --time=0-01:00:00
#SBATCH --cpus-per-task=1
#SBATCH --mem=2G
#SBATCH --output=logs/ydraw_%A_%a.out
#SBATCH --error=logs/ydraw_%A_%a.err

set -euo pipefail

export OMP_NUM_THREADS=1
export OPENBLAS_NUM_THREADS=1
export MKL_NUM_THREADS=1

if command -v conda >/dev/null 2>&1; then
    eval "$(conda shell.bash hook)"
    conda activate ratchet
elif [ -f .venv/bin/activate ]; then
    source .venv/bin/activate
else
    echo "no conda env 'ratchet' and no .venv; run setup_cluster.sh first" >&2
    exit 1
fi

PARAMS="${PARAMS:-params_y_draw.tsv}"
OUTDIR="${OUTDIR:-results_y_draw}"

echo "task ${SLURM_ARRAY_TASK_ID} on $(hostname) at $(date)"
python run_task_ydraw.py --idx "${SLURM_ARRAY_TASK_ID}" \
                         --params "${PARAMS}" --outdir "${OUTDIR}"
echo "done at $(date)"
```
