## Supplementary Information section 1 code base for "Hypothesis: A modern human range expansion ∼300,000 years ago had a critical role in Neandertal origins": run_task.py

```
"""
Run one chunk of a 300-replicate grid: one SLURM array task, one row of a
params file.

    python run_task.py                                    # $SLURM_ARRAY_TASK_ID
    python run_task.py --idx 17                           # mtDNA grid
    python run_task.py --idx 17 --params params_extra_300.tsv \
                       --outdir results_extra300          # Y and RNA grid

Serves both grids. The mtDNA params file has no tag or anchor_ne column, so both
default: anchor_ne to the mtDNA reference Ne of 7500 that ratchet_dfe assumes,
and tag to the empty string. The Y grid sets anchor_ne = 10000, because its DFE
is the autosomal one estimated at that reference size, and getting this wrong
would rescale every Y selection coefficient.

Writes <outdir>/task_<idx>.jsonl in the same shape as run_laptop.py's output (a
summary plus the per-replicate rows). Tasks that already have a non-empty output
file are skipped, so a partially failed array can be resubmitted whole rather
than needing the survivors picked out by hand. Output is written to a .partial
file and renamed, so a task killed mid-write cannot leave a truncated file that
the skip check would later mistake for a finished one.

The per-replicate rows are what matter here; the per-chunk summary is written
only for inspection, since merge_chunks.py recomputes the statistics from the
pooled replicates of all chunks of a combination.
"""
import argparse
import csv
import json
import os
import time

from ratchet_dfe import ANCHOR_NE, NONSENSE_FRACTION, run, summarize

OUTDIR = "results300"


def main():
    ap = argparse.ArgumentParser()
    ap.add_argument("--idx", type=int,
                    default=int(os.environ.get("SLURM_ARRAY_TASK_ID", -1)))
    ap.add_argument("--params", default="params_300.tsv")
    ap.add_argument("--outdir", default=OUTDIR)
    args = ap.parse_args()
    if args.idx < 0:
        raise SystemExit("no --idx and no SLURM_ARRAY_TASK_ID")

    with open(args.params) as f:
        grid = {int(r["idx"]): r for r in csv.DictReader(f, delimiter="\t")}
    if args.idx not in grid:
        raise SystemExit(f"idx {args.idx} not in {args.params}")
    row = grid[args.idx]

    os.makedirs(args.outdir, exist_ok=True)
    out = os.path.join(args.outdir, f"task_{args.idx}.jsonl")
    if os.path.exists(out) and os.path.getsize(out) > 0:
        print(f"task {args.idx}: already done, skipping")
        return

    Sbar, beta = float(row["Sbar"]), float(row["beta"])
    U_mis = float(row["U_mis"])
    U_non = U_mis * NONSENSE_FRACTION
    N_arch, N_mod = int(row["N_archaic"]), int(row["N_modern"])
    T, reps, seed = int(row["T"]), int(row["reps"]), int(row["seed"])
    anchor_ne = float(row.get("anchor_ne") or ANCHOR_NE)
    tag = row.get("tag") or ""

    print(f"task {args.idx}: combo {row['combo']} chunk {row['chunk']} "
          f"{('[' + tag + '] ') if tag else ''}| "
          f"Sbar={Sbar} beta={beta} U={U_mis} N_A={N_arch} "
          f"anchor_Ne={anchor_ne:.0f} reps={reps} seed={seed}", flush=True)

    t0 = time.time()
    rows, s_bar = run(N_mod, N_arch, T, U_mis, U_non, beta, Sbar, reps, seed,
                      anchor_ne=anchor_ne)
    elapsed = time.time() - t0
    summary = summarize(rows, N_modern=N_mod, N_archaic=N_arch, T=T,
                        U_mis=U_mis, U_non=U_non, beta=beta, Sbar=Sbar,
                        s_bar=s_bar, reps=reps, seed=seed, elapsed=elapsed)
    summary["combo"] = int(row["combo"])
    summary["chunk"] = int(row["chunk"])
    summary["anchor_ne"] = anchor_ne
    if tag:
        summary["tag"] = tag

    tmp = out + ".partial"
    with open(tmp, "w") as f:
        f.write(json.dumps({
            "summary": summary,
            "reps": [{k: v for k, v in r.items() if not k.startswith("traj")}
                     for r in rows]}) + "\n")
    os.replace(tmp, out)

    print(f"task {args.idx}: diff={summary['diff_mean']:.5f} "
          f"+-{summary['diff_se']:.5f} [{elapsed:.0f}s]")


if __name__ == "__main__":
    main()
```
