## Supplementary Information section 1 code base for "Hypothesis: A modern human range expansion ∼300,000 years ago had a critical role in Neandertal origins": run_task_ydraw.py

```
"""
One SLURM array task of the PARAMETER-INTEGRATED Y target-size ladder: one row
of params_y_draw.tsv = one chunk of 30 replicates, where EACH replicate first
draws (beta, Sbar, U) from triangular priors and then simulates one modern and
one archaic lineage under the drawn values.

The priors follow the Methods' Part-1 bound calculation (run_part1_bound.py)
exactly: beta ~ tri(0.19, 0.23, 0.27), |Sbar| ~ tri(225, 425, 766) anchored at
Ne = 10000, and the deleterious input U ~ tri over the Y rate CI
(2.5-3.5e-8 /bp/gen, mode 3.0e-8) scaled to the rung's target size L:
U = mu * L * 0.72. As there, the modern and archaic runs of a replicate share
one RNG stream, and scale = (Sbar / (2 * anchor_ne)) / beta.

The resulting per-rung gap distributions mix parameter uncertainty with the
ratchet's own stochasticity -- the parameter-integrated counterpart of the
fixed-central-DFE distributions from params_y_v22.tsv.

Output: <outdir>/task_<idx>.jsonl in the same shape as run_task.py (summary +
per-replicate rows, here with each replicate's drawn parameters), with the
same skip-if-done and atomic-rename behavior.
"""
import argparse
import csv
import json
import os
import time

import numpy as np

from ratchet_dfe import NONSENSE_FRACTION, simulate_one

MU_TRI = (2.5e-8, 3.0e-8, 3.5e-8)     # Y point rate /bp/gen (low, mode, high)
F_NONSYN = 0.72
BETA_TRI = (0.19, 0.23, 0.27)
SBAR_TRI = (225.0, 425.0, 766.0)


def main():
    ap = argparse.ArgumentParser()
    ap.add_argument("--idx", type=int,
                    default=int(os.environ.get("SLURM_ARRAY_TASK_ID", -1)))
    ap.add_argument("--params", default="params_y_draw.tsv")
    ap.add_argument("--outdir", default="results_y_draw")
    args = ap.parse_args()
    if args.idx < 0:
        raise SystemExit("no --idx and no SLURM_ARRAY_TASK_ID")

    with open(args.params) as f:
        grid = {int(r["idx"]): r for r in csv.DictReader(f, delimiter="\t")}
    row = grid[args.idx]

    os.makedirs(args.outdir, exist_ok=True)
    out = os.path.join(args.outdir, f"task_{args.idx}.jsonl")
    if os.path.exists(out) and os.path.getsize(out) > 0:
        print(f"task {args.idx}: already done, skipping")
        return

    kb = int(row["target_kb"])
    N_mod, N_arch = int(row["N_modern"]), int(row["N_archaic"])
    T, reps, seed = int(row["T"]), int(row["reps"]), int(row["seed"])
    anchor = float(row["anchor_ne"])
    u_tri = tuple(mu * kb * 1000 * F_NONSYN for mu in MU_TRI)

    print(f"task {args.idx}: combo {row['combo']} chunk {row['chunk']} "
          f"[{row['tag']}] | target={kb}kb N_M={N_mod} N_A={N_arch} T={T} "
          f"reps={reps} seed={seed} (per-replicate prior draws)", flush=True)

    t0 = time.time()
    rows = []
    for rep in range(reps):
        rng = np.random.default_rng([seed, rep])
        beta = float(rng.triangular(*BETA_TRI))
        Sbar = float(rng.triangular(*SBAR_TRI))
        U_mis = float(rng.triangular(*u_tri))
        U_non = U_mis * NONSENSE_FRACTION
        scale = (Sbar / (2.0 * anchor)) / beta
        wm, _ = simulate_one(N_mod, T, U_mis, U_non, beta, scale, rng)
        wa, _ = simulate_one(N_arch, T, U_mis, U_non, beta, scale, rng)
        rows.append(dict(rep=rep, beta=beta, Sbar=Sbar, U_mis=U_mis,
                         w_modern=wm, w_archaic=wa, diff=wm - wa))
    elapsed = time.time() - t0

    diffs = np.array([r["diff"] for r in rows])
    summary = dict(
        combo=int(row["combo"]), chunk=int(row["chunk"]), tag=row["tag"],
        target_kb=kb, N_modern=N_mod, N_archaic=N_arch, T=T, reps=reps,
        seed=seed, anchor_ne=anchor,
        diff_mean=float(diffs.mean()),
        diff_se=float(diffs.std(ddof=1) / np.sqrt(len(diffs)))
        if len(diffs) > 1 else 0.0,
        elapsed_sec=elapsed)

    tmp = out + ".partial"
    with open(tmp, "w") as f:
        f.write(json.dumps({"summary": summary, "reps": rows}) + "\n")
    os.replace(tmp, out)
    print(f"task {args.idx}: diff={summary['diff_mean']:.5f} "
          f"+-{summary['diff_se']:.5f} [{elapsed:.0f}s]")


if __name__ == "__main__":
    main()
```
