## Supplementary Information section 1 code base for "Hypothesis: A modern human range expansion ∼300,000 years ago had a critical role in Neandertal origins": save_dfe_draws.py

```
"""
Save raw bootstrap draws of (beta, |Sbar|) for the Denisova-calibrated m=53 fit,
for the 'nonsynonymous' (coding-only) and 'all' (whole-mtDNA) classes. These are
consumed by denis_bound.py, which propagates DFE uncertainty into the fitness-gap
upper bound by drawing (beta, Sbar) jointly (the two are strongly correlated, so
triangular marginals over the CIs would misstate the prior).
Same resampling procedure as compute_denis_table.py.
"""
import numpy as np
from Bio import SeqIO

from pew_replicate import build_grids, a_n
from pew_rna import fit_pew, project_segregating, jc_correct, OUTGROUPS
from rna_constraint import build_site_table, align_to_ref, analyse as rc_analyse, \
    SEQDIR, TITV
from mt_dfe_sfs import read_variants, opportunity, pop_sample_size, GB

POP, OUTGROUP, M, BOOT = "afr", "Denisova", 53, 400
FUNC = ("nonsynonymous", "tRNA", "rRNA")
CLASSES = ("nonsynonymous", "all")


def main():
    import argparse
    ap = argparse.ArgumentParser()
    ap.add_argument("--boot", type=int, default=BOOT,
                    help="bootstrap replicates (default 400)")
    args = ap.parse_args()
    rec = SeqIO.read(GB, "genbank")
    site, refseq = build_site_table(rec)
    n = pop_sample_size(POP)
    variants, _ = read_variants(site, refseq, an_min=int(0.9 * n),
                                n_common=n, pop=POP)
    mc = {"nonsynonymous": variants["nonsynonymous"],
          "all": np.concatenate([variants[k] for k in FUNC])}
    mc_syn = variants["synonymous"]

    N = opportunity(site, refseq)
    N["all"] = sum(N[k] for k in FUNC)
    Nr = {c: N[c] / N["synonymous"] for c in CLASSES}

    acc = dict(OUTGROUPS)[OUTGROUP]
    outseq = str(SeqIO.read(f"{SEQDIR}/{acc}.fa", "fasta").seq).upper()
    obs_d, opp_d, _ = rc_analyse(site, refseq, align_to_ref(refseq, outseq), TITV)
    obs_d["all"] = sum(obs_d[k] for k in FUNC)
    opp_d["all"] = sum(opp_d[k] for k in FUNC)

    S_edges = np.concatenate([[0.0], np.logspace(-4, 7.5, 600)])
    S_mid = np.sqrt(np.maximum(S_edges[:-1], 1e-6) * np.maximum(S_edges[1:], 1e-6))
    S_mid[0] = 0.0
    Q, F = build_grids(M, S_mid)
    an = a_n(M)

    rng = np.random.default_rng(2025)
    draws = {c: [] for c in CLASSES}
    for b in range(args.boot):
        s_syn = rng.choice(mc_syn, size=len(mc_syn), replace=True)
        Pneu = project_segregating(s_syn, n, M)
        Dneu = max(rng.poisson(obs_d["synonymous"]), 1)
        dneu = jc_correct(Dneu / opp_d["synonymous"])
        for c in CLASSES:
            s_sel = rng.choice(mc[c], size=len(mc[c]), replace=True)
            Psel = project_segregating(s_sel, n, M)
            Dsel = max(rng.poisson(obs_d[c]), 1)
            if min(Pneu, Psel) <= 0:
                continue
            IP = (Psel / Pneu) * an / Nr[c]
            ID = jc_correct(Dsel / opp_d[c]) / dneu
            _, beta, Sbar, _ = fit_pew(M, Nr[c], IP, ID, Q, F, S_edges)
            draws[c].append((beta, abs(Sbar)))
        if (b + 1) % 50 == 0 or (b + 1) == args.boot:
            print(f"  {b + 1}/{args.boot}", flush=True)
    out = {c: np.array(v) for c, v in draws.items()}
    np.savez("dfe_draws.npz", **{f"{c}": out[c] for c in CLASSES})
    for c in CLASSES:
        a = out[c]
        print(f"{c}: {len(a)} draws, beta [{np.percentile(a[:,0],2.5):.2f},"
              f"{np.percentile(a[:,0],97.5):.2f}], corr(beta,log|S|) = "
              f"{np.corrcoef(a[:,0], np.log(a[:,1]))[0,1]:.2f}", flush=True)
    print("saved dfe_draws.npz", flush=True)


if __name__ == "__main__":
    main()
```
