## Supplementary Information section 1 code base for "Hypothesis: A modern human range expansion ∼300,000 years ago had a critical role in Neandertal origins": sweep_y_U.py

```
"""
How large a deleterious input U (whole-Y, under the Y DFE) is needed before the
modern-archaic fitness gap crosses 1%? Sweeps U at the central Y DFE and reports
the gap for N_archaic = 1500 and 3000, plus the implied constrained target size.

This addresses: over the ENTIRE Y (coding + non-coding, agnostic), could a rain
of slightly deleterious mutations plausibly build a >1% gap over 17,500 gen?
"""
import sys
from multiprocessing import Pool, cpu_count
import numpy as np
from ratchet_dfe import simulate_one, NONSENSE_FRACTION

T = 17500
N_MODERN = 7500
NARCH = (1500, 3000)
# Y DFE (central): autosomal aa DFE rescaled to Y, anchored at Ne=10000.
BETA = 0.23
SBAR = 425.0
ANCHOR = 10000.0
SCALE = (SBAR / (2.0 * ANCHOR)) / BETA
MU_Y = 3.0e-8          # per bp per gen
MISSENSE_FRAC = 0.72   # fraction of point mutations that are amino-acid-altering-equivalent

U_GRID = [0.0022, 0.005, 0.010, 0.020, 0.030, 0.050, 0.080]
REPS = 24


def job(args):
    U_mis, N, rep, T_run = args
    rng = np.random.default_rng(10_000 + rep * 131 + int(U_mis * 1e6))
    w, _ = simulate_one(N, T_run, U_mis, U_mis * NONSENSE_FRACTION, BETA, SCALE, rng)
    return (U_mis, N, rep, w)


def main():
    import argparse
    ap = argparse.ArgumentParser()
    ap.add_argument("--reps", type=int, default=REPS)
    ap.add_argument("--T", type=int, default=T)
    args = ap.parse_args()
    T_run = args.T
    jobs = []
    # modern once per (U, rep); archaic per (U, N, rep)
    for U in U_GRID:
        for rep in range(args.reps):
            jobs.append((U, N_MODERN, rep, T_run))
            for N in NARCH:
                jobs.append((U, N, rep, T_run))
    nproc = min(cpu_count(), 10)
    print(f"{len(jobs)} sims on {nproc} procs (T={T_run})...", file=sys.stderr)
    with Pool(nproc) as pool:
        res = pool.map(job, jobs)

    # index results
    from collections import defaultdict
    wmod = defaultdict(dict)   # U -> rep -> w
    warch = defaultdict(lambda: defaultdict(dict))  # N -> U -> rep -> w
    for U, N, rep, w in res:
        if N == N_MODERN:
            wmod[U][rep] = w
        else:
            warch[N][U][rep] = w

    print(f"\nY DFE central (beta={BETA}, Sbar={SBAR}); T={T_run}; N_mod={N_MODERN}")
    print(f"{'U':>7s} {'~bp_constrained':>15s} {'gap_1500%':>10s} {'gap_3000%':>10s}")
    for U in U_GRID:
        bp = U / (MU_Y * MISSENSE_FRAC)
        gaps = {}
        for N in NARCH:
            g = np.array([wmod[U][r] - warch[N][U][r] for r in range(args.reps)]) * 100.0
            se = 0.0 if len(g) < 2 else g.std(ddof=1) / np.sqrt(len(g))
            gaps[N] = (g.mean(), se)
        print(f"{U:>7.4f} {bp:>15,.0f} "
              f"{gaps[1500][0]:>7.3f}±{gaps[1500][1]:.3f} "
              f"{gaps[3000][0]:>7.3f}±{gaps[3000][1]:.3f}")


if __name__ == "__main__":
    main()
```
