## Supplementary Information section 1 code base for "Hypothesis: A modern human range expansion ∼300,000 years ago had a critical role in Neandertal origins": validate_delta.py

```
"""
Validation: does the new DFE framework reproduce the ORIGINAL single-s Perl
simulation (Extended Data Figure 1) at one grid point?

Original grid point (ratchet.pl / wrapper.pl):
    M = 7500, A = 1500, T = 17500, mu = 0.02 (per-generation, whole locus),
    s = 0.01 (single coefficient, all mutations), no lethal class.
    Reported result: modern w = 0.98, archaic w = 0.85.

We reproduce it by collapsing the gamma DFE to a near-delta at s = 0.01:
    beta -> very large; s_bar = Sbar / (2 * anchor_Ne) = 0.01  (Sbar=150, anchor=7500)
    U_mis = 0.02, U_non = 0  (no separate nonsense/lethal class).

The only model difference from the Perl code is Poisson(U) vs Bernoulli(U) new
mutations per generation; for U = 0.02 these are essentially identical
(P(>=2 mutations) ~ 2e-4).

Run:  ./.venv/bin/python validate_delta.py
"""
import math
from multiprocessing import Pool

from ratchet_dfe import run

N_MODERN = 7500
N_ARCHAIC = 1500
T = 17500
U_MIS = 0.02
U_NON = 0.0
S_TARGET = 0.01
ANCHOR = 7500
SBAR = 2 * ANCHOR * S_TARGET     # = 150 -> s_bar = 0.01
BETA = 1e7                       # huge shape -> gamma ~ delta at the mean
PROCS = 10
REPS_PER = 6                     # total = PROCS * REPS_PER replicates

OLD_MODERN, OLD_ARCHAIC = 0.98, 0.85


def worker(args):
    seed, reps_per = args
    rows, s_bar = run(N_MODERN, N_ARCHAIC, T, U_MIS, U_NON, BETA, SBAR,
                      reps_per, seed, anchor_ne=ANCHOR)
    return rows, s_bar


def main():
    import argparse
    ap = argparse.ArgumentParser()
    ap.add_argument("--procs", type=int, default=PROCS)
    ap.add_argument("--reps-per", type=int, default=REPS_PER)
    args = ap.parse_args()
    print(f"Replicating original grid point: M={N_MODERN} A={N_ARCHAIC} "
          f"T={T} mu={U_MIS} s={S_TARGET}")
    print(f"(delta DFE: beta={BETA:g}, Sbar={SBAR} @ anchor {ANCHOR} "
          f"-> s_bar={SBAR/(2*ANCHOR)}, U_non={U_NON})\n")

    seeds = [(9000 + i, args.reps_per) for i in range(args.procs)]
    with Pool(args.procs) as pool:
        out = pool.map(worker, seeds)

    wm = [r["w_modern"] for rows, _ in out for r in rows]
    wa = [r["w_archaic"] for rows, _ in out for r in rows]
    s_bar = out[0][1]
    n = len(wm)

    def stat(x):
        m = sum(x) / len(x)
        sd = math.sqrt(sum((v - m) ** 2 for v in x) / (len(x) - 1))
        return m, sd / math.sqrt(len(x))

    mm, mse = stat(wm)
    am, ase = stat(wa)
    print(f"replicates: {n};  realized s_bar = {s_bar:.4f}\n")
    print(f"{'lineage':<10}{'new sim (mean +/- SE)':<28}{'original':<12}")
    print(f"{'modern':<10}{mm:.3f} +/- {mse:.3f}{'':<14}{OLD_MODERN}")
    print(f"{'archaic':<10}{am:.3f} +/- {ase:.3f}{'':<14}{OLD_ARCHAIC}")
    print(f"\ngap (modern - archaic): {mm-am:.3f}   (original {OLD_MODERN-OLD_ARCHAIC:.3f})")
    print(f"Haldane-Muller expectation for modern (e^-U): {math.exp(-U_MIS):.3f}")


if __name__ == "__main__":
    main()
```
