## Supplementary Information section 1 code base for "Hypothesis: A modern human range expansion ∼300,000 years ago had a critical role in Neandertal origins": y_numbers_stated.py

```
"""
All document numbers for the Y ladder under the stated conversion
(s-bar = S-bar/Nref = 0.0425): mean gaps, 1% crossing, fixation
probabilities, Table 3 rows, hemi sensitivity, integrated-vs-fixed check.
"""
import csv

import numpy as np

KBS = [100, 200, 300, 500, 1000, 2000, 3000, 5000]
EUCHROMATIC_MSY_KB = 23000.0


def load_curves(path="fixation_curves_mtdna.csv"):
    curves = {}
    with open(path) as f:
        for r in csv.DictReader(x for x in f if not x.startswith("#")):
            curves.setdefault(int(float(r["locus_Ne"])), []).append(
                (float(r["s"]), float(r["pfix_pct"])))
    return {ne: np.array(sorted(p)) for ne, p in curves.items()}


CURVES = load_curves()


def pfix(gap_pct, ne):
    c = CURVES[ne]
    s = np.asarray(gap_pct, float) / 100.0
    return np.interp(s, c[:, 0], c[:, 1])


def crossing(kbs, means, level):
    """kb at which mean gap (in %) crosses `level`, log-log interpolation."""
    kbs, means = np.array(kbs, float), np.array(means, float)
    i = np.searchsorted(means, level)
    if i == 0 or i >= len(means):
        return np.nan
    lk = np.interp(np.log(level), np.log(means[i - 1:i + 1]),
                   np.log(kbs[i - 1:i + 1]))
    return np.exp(lk)


def ladder(npz, na="2250"):
    d = np.load(npz)
    return {kb: d[f"y{kb}_{na}"] * 100.0 for kb in KBS}


def petr_ladder(npz):
    d = np.load(npz)
    return {kb: d[f"y{kb}_petr_500"] * 100.0 for kb in KBS}


def report(name, npz):
    print(f"===== {name} =====")
    for na in ("1500", "2250", "3000"):
        lad = ladder(npz, na)
        means = [lad[kb].mean() for kb in KBS]
        print(f"NA={na}: means " +
              ", ".join(f"{kb}kb {m:.3g}%" for kb, m in zip(KBS, means)))
        print(f"  1% crossing: {crossing(KBS, means, 1.0):.0f} kb")
        if na == "2250":
            pf = {kb: pfix(lad[kb], 750).mean() for kb in KBS}
            print("  P(fix, 750): " +
                  ", ".join(f"{kb}kb {pf[kb]:.0f}%" for kb in KBS))
            pfs = [pf[kb] for kb in KBS]
            print(f"  50% P(fix) crossing: "
                  f"{crossing(KBS, pfs, 50.0):.0f} kb")
            g100 = lad[100]
            print(f"  100kb: mean {g100.mean():.3g}%, "
                  f"95th pct {np.percentile(g100, 95):.2g}%, "
                  f"P(fix) int {pfix(g100, 750).mean():.0f}%, "
                  f"at bound {pfix(np.percentile(g100, 95), 750):.0f}%")
        else:
            g100 = lad[100]
            print(f"  100kb 95th pct: {np.percentile(g100, 95):.2g}%")
    pl = petr_ladder(npz)
    print(f"Petr 500kb: mean {pl[500].mean():.3g}%, "
          f"P(fix, 500-curve) {pfix(pl[500], 500).mean():.0f}%")
    x = crossing(KBS, [ladder(npz, '2250')[kb].mean() for kb in KBS], 1.0)
    print(f"share of euchromatic MSY at 1% crossing: "
          f"{100 * x / EUCHROMATIC_MSY_KB:.1f}%")


if __name__ == "__main__":
    report("stated, parameter-integrated", "y_draw_stated_reps.npz")
    report("hemizygous, parameter-integrated", "y_draw_hemi_reps.npz")

    s = ladder("y_draw_stated_reps.npz")
    h = ladder("y_draw_hemi_reps.npz")

    sp500 = np.concatenate([ladder("y_draw_stated_reps.npz", na)[500]
                            for na in ("1500", "2250", "3000")])
    hp500 = np.concatenate([ladder("y_draw_hemi_reps.npz", na)[500]
                            for na in ("1500", "2250", "3000")])
    print(f"pooled 500kb: stated {sp500.mean():.3g}% vs hemi {hp500.mean():.3g}%")

    x = crossing(KBS, [s[kb].mean() for kb in KBS], 1.0)
    print(f"share of euchromatic MSY at 1% crossing (NA=2250 stated): "
          f"{100 * x / EUCHROMATIC_MSY_KB:.1f}%")
    print(f"non-coding must contribute ~{(x - 100) / 100:.1f}x the coding "
          f"input (crossing {x:.0f} kb)")
```
