## Supplementary Information section 2 for "Hypothesis: A modern human range expansion ∼300,000 years ago had a critical role in Neandertal origins"

No evidence for a Dobzhansky-Muller Incompatibility between Denisovan mtDNA and modern human nuclear DNA

#### Findings

- Eight nonsynonymous mtDNA changes are fixed across the four sequenced Denisovans, and absent from Neandertals and modern humans. Three of the eight are situated at interfaces with nuclear-encoded OXPHOS subunits. However, none of the contacting nuclear residues carries an archaic-derived substitution of the kind a Dobzhansky-Muller Incompatibility would predict.
- These results reduce the probability that the rise in frequency of early-modern-human-derived mtDNA following admixture into Neandertal ancestors ~300-200 thousand years ago (ka) is due to a Dobzhansky-Muller Incompatibility (DMI) involving mtDNA.

#### Motivation

Neandertal mitochondrial genomes are far more similar to those of modern humans than to deeply divergent Denisovan mtDNA, despite the fact that Neandertals are closer to Denisovans on average in the nuclear genome. This discordance has been shown to be due to a history of introgression of a modern-human-like mtDNA into the ancestors of the Neandertals, and replacement of an originally Denisovan-like mtDNA[1]. Analyses of whole genome data infer that this admixture event occurred ~300-200 ka, and resulted in ~3-7% of Neandertal DNA deriving from lineages related to modern humans on the autosomes, even though modern human mtDNA rose to fixation[2,3,4].

In what follows, I search for evidence that a mitochondrial-nuclear Dobzhansky-Muller Incompatibility (DMI) caused the Denisovan-like mtDNA to be incompatible with the incoming (modern-human-like) nuclear background, under the hypothesis that such a process would have led to selection for the replacement of this allele. Mitochondrial-nuclear incompatibilities are well described across diverse species, and have in multiple instances been implicated in lowered hybrid fitness, including in reduced male fertility[5]. Previous work has shown that these incompatibilities often arise due to interactions between nonsynonymous mtDNA changes at interfaces with nuclear-encoded OXPHOS subunits whose partner residues carry a coevolved variant[6,7]. Thus, a prediction of this hypothesis is that Denisovan-specific, derived, nonsynonymous mtDNA changes should fall at interfaces with nuclear-encoded OXPHOS subunits carrying an archaic-versus-modern-distinct allele. I test this prediction using computational approaches, mining datasets of protein structure and of archaic and modern human genetic variation.

The text and analyses in this note are the product of an interaction between the author and AI coding agents of the Anthropic Claude family (Opus and Fable models). I used the agents to write the analysis code in Python, to run diagnostics, and to help compose the text (any suggestions from the AI agents were reviewed and edited by me). All quantitative results are produced by my execution of the cited scripts on the O2 computing cluster at Harvard Medical School, or on my personal computer. I checked the robustness of the output, and reviewed the scientific interpretations, modeling choices, and final claims.

### Data and methods

I called coding changes per gene from NCBI CDS translations for the revised Cambridge Reference Sequence (rCRS) for human mtDNA (NC_012920), Denisova 3 (FN673705), Vindija 33.19 (KJ533545), and great-ape outgroups (chimpanzee NC_001643, bonobo NC_001644, gorilla NC_011120), aligned to rCRS and polarized by great-ape majority under the vertebrate-mitochondrial code (transl_table 2). For the fixation filter, I used four Denisovan and eleven Neandertal mtDNAs.

I identified nuclear partners in human cryo-EM structures of the OXPHOS complexes: Complex I (5XTD)[8], Complex III (9HZL), Complex IV (5Z62) and ATP synthase (8H9S)[9,10,11]. I inferred inter-chain contacts with Biopython (heavy-atom ≤4 Å = interface, ≤8 Å = shell). I reconciled residue numbering to UniProt via SIFTS and verified it by residue identity.

I carried out allele-level tests drawing on the archaic-versus-modern coding catalogs of Prüfer et al. 2014 (Altai; 789-gene coverage)[12], Meyer et al. 2012 (Denisova-derived; 2,002-genes)[13], and Kuhlwilm and Boeckx 2019 (647 high-frequency modern-derived missense changes in 571 genes, and 1,542 genes carrying a high-frequency archaic-derived missense change)[14]. I used modern-human variation information from gnomAD[15] v4; archaic genotypes from the Prüfer et al. 2017 VCF release[16]; and intrinsic mtDNA variant impact from MitImpact 3.1.3 (APOGEE2)[17,18].

### Eight nonsynonymous mtDNA changes that are fixed in Denisovans

I identified eight nonsynonymous sites that were fixed derived in the four-genome fixation panel (Denisova 2, 3, 4 and 8), and absent from all eleven sampled Neandertals and from rCRS. I corroborated all eight by MitImpact's position-to-amino-acid annotation. Counting every change derived in any single Denisovan raises the apparent count to fifteen. The additional seven are non-fixed, seen in only a subset of the four Denisovans (Table S2.1).

I carried out a codon-aware partition of all coding substitutions among Denisova 3, Vindija 33.19, and rCRS (ape-polarized). This reveals the expected time-dependence of dN/dS: low nonsynonymous-to-synonymous ratios on the deep Denisova-only and shared modern+Neandertal branches, and elevated ratios on the short terminal branches. This is consistent with ref. 19, who report a maximum-likelihood estimate of ~18 nonsynonymous substitutions on the Denisovan terminal branch (dN/dS = 0.056; their Supplementary Figure 5). The eight high-confidence sites are a fixed-and-derived subset of that total.

The eight fixed, derived mtDNA changes are called benign or likely-benign by APOGEE2, are physicochemically conservative, and in five of eight cases substitute a residue already present in other mammals.

**Table S2.1. Denisovan-specific mtDNA changes and their nuclear OXPHOS partners.** Rows list every nonsynonymous change derived in any of the four sequenced Denisovans (Denisova 2, 3, 4, 8): eight are fixed across all four (or across all three with coverage), plus seven non-fixed (carried by a subset: two further private premature-stop calls in Denisova 2 are excluded as probable artifacts); sites with no nuclear partner within 8 Å are shown as "none ≤8 Å".

| **mtDNA change (ancestral→derived)** | **Denisovan mtDNA genotype (Den2 / Den3 / Den4 / Den8)** | **Fixed or polymorphic** | **Nuclear partner residue (closest heavy-atom distance)** | **Derived allele at partner? (Den3 / Den25 / Nea)** | **APOGEE2 (mt variant)** |
|---|---|---|---|---|---|
| ***Fixed across all four Denisovans — derived on the Denisovan lineage (the eight high-confidence sites)*** | | | | | |
| *ND2* Ala64→Thr (A64T) | Thr / Thr / Thr / Thr | Fixed (4/4) | none ≤8 Å | n/a | Likely-benign |
| *ND2* Ala164→Thr (A164T) | Thr / Thr / Thr / Thr | Fixed (4/4) | none ≤8 Å | n/a | Benign |
| *ND5* Thr398→Ala (T398A) | Ala / Ala / Ala / Ala | Fixed (4/4) | *NDUFB8* Gln154 (7.8 Å, shell) | anc / anc / anc | Benign |
| *CYTB* Thr70→Ala (T70A) | Ala / Ala / Ala / Ala | Fixed (4/4) | *CYC1* Tyr199 (3.0 Å, interface) | anc / anc / anc | Likely-benign |
| *COX1* Val456→Ala (V456A) | Ala / Ala / Ala / Ala | Fixed (4/4) | *COX7B* Trp53 (3.7 Å); COX4I1 Ile118 (3.8 Å) | anc / anc / anc | Benign |
| *COX2* Ala27→Thr (A27T) | Thr / Thr / Thr / Thr | Fixed (4/4) | *COX6C* Tyr37 (6.4 Å, shell) | anc / anc / anc | Benign |
| *ATP6* Ala7→Thr (A7T) ‡‡ | Thr / Thr / ? / Thr | Fixed (3/3 cov.) | *ATP5MF* His84 (3.8 Å, interface) | anc / n.c. / anc | Benign |
| *ATP6* Asn185→Ser (N185S) ‡ | Ser / Ser / ? / Ser | Fixed (3/3 cov.) | none ≤8 Å | n/a | Benign |
| ***Non-fixed — derived in only a subset of the four Denisovans (carriers in parentheses); none sits at a nuclear interface*** | | | | | |
| *ND1* Thr153→Met (T153M) | Thr / Met / Met / Thr | Polymorphic (Den3, Den4) | none ≤8 Å | n/a | not assessed |
| *ND2* Met163→Val (M163V) | Met / Val / Val / Met | Polymorphic (Den3, Den4) | none ≤8 Å | n/a | not assessed |
| *COX1* Val83→Ile (V83I) | Val / Ile / Ile / Val | Polymorphic (Den3, Den4) | none ≤8 Å | n/a | not assessed |
| *COX3* Ala32→Thr (A32T) | Ala / Thr / Thr / Ala | Polymorphic (Den3, Den4) | none ≤8 Å | n/a | not assessed |
| *ND1* Ala112→Thr (A112T) | Thr / Ala / Ala / Ala | Polymorphic (Den2) | none ≤8 Å | n/a | not assessed |
| *ATP6* Thr45→Ala (T45A) | Ala / Thr / ? / Ala | Polymorphic (Den2, Den8) | none ≤8 Å | n/a | not assessed |
| *CYTB* Leu149→Met (L149M) | Leu / Leu / Leu / Met | Polymorphic (Den8) | none ≤8 Å | n/a | not assessed |

Notes: mtDNA change: ancestral→derived amino acid (three-letter), great-ape-polarized, with the one-letter shorthand in parentheses; positions in rCRS coordinates. Denisovan mtDNA genotype: residue in each of the four Denisovan mtDNAs: Den2 (KX663333), Den3 (FN673705), Den4 (FR695060), Den8 (KT780370). "?" = position not covered. Nuclear partner residue: closest nuclear-subunit residue (three-letter + structure-numbering position) to the mt change, with the minimum mt-nuclear heavy-atom distance (interface ≤4 Å; shell ≤8 Å). "Derived allele at partner?": whether Denisova 3 / Denisova 25 / Neandertal carries a non-modern (derived) residue at that nuclear position. "anc" = ancestral / modern-matching, "n.c." = no confident call, "n/a" = no partner residue. All three archaics are ancestral at every partner residue tested: Denisova 3 and the Altai Neandertal by direct genotyping of the autosomal partners, and Denisova 25 (~200 ka, ~24× nuclear coverage) by direct genotyping of the per-chromosome VCFs, including *COX7B* Trp53 on chromosome X (Denisova 25 0/0, depth 9). The one gap is *ATP5MF* His84 in Denisova 25 (n.c.; those codon positions were filtered out). APOGEE2: intrinsic pathogenicity call for the mt variant; "not assessed" = outside the original eight-site intrinsic panel. ‡ *ATP6* Asn185→Ser: polarization ambiguous (apes carry both Asn and Ser); the site-calling code alternatively flags ND5 Ser515→Pro as the eighth derived site. ‡‡ ATP6 Ala7→Thr is derived in Denisova 2/3/8 but ancestral in the older Denisova 25, i.e. it arose after the oldest Denisovan (not present at the time of introgression).

### Three of the eight variant residues map to conserved OXPHOS nuclear interfaces

All thirteen mtDNA-encoded proteins are OXPHOS subunits, so the candidate DMI partners are the nuclear-encoded subunits contacting each variant residue. Three fixed Denisovan changes sit at true (≤4 Å) nuclear interfaces (Table S2.1): *CYTB* T70-*CYC1* (cytochrome $c_1$), *COX1* V456-*COX7B/COX4I1*, and *ATP6* A7-*ATP5MF*. A matched Neandertal-lineage control set (with interface changes *ND1* T168-*NDUFA13*, *CYTB* F245-*CYC1*, *ATP6* S176-*ATP5PB*) shows a comparable interface fraction (3/12 versus 3/8). Of these three interface changes, *CYTB* T70 and *COX1* V456 are present in the ~200 ka Denisova 25 mtDNA[20], which is not part of the fixation panel and so provides an independent temporal check. *ATP6* A7T is ancestral there, showing that it arose after the divergence of the lineage leading to the oldest Denisovan. This raises the possibility that it was not carried by the Denisovan-like mtDNA at the time of introgression.

### No evidence for Denisova-specific nuclear substitutions at positions that contact Denisovan-specific mtDNA substitutions at OXPHOS structural interfaces

The three archaic-versus-modern coding catalogs, covering between 789 and 2,002 genes, identify no fixed or high-frequency nonsynonymous change in any of the twelve nuclear partner genes (the six contacting the Denisovan-specific changes, and the six further partners of the Neandertal-lineage control set: *NDUFA13, ATP5PB, NDUFA1, NDUFC2, NDUFA11* and *ATP5MC1*; *CYC1* contacts interface changes in both sets), in any direction: modern-derived, archaic-derived, or Denisova-derived. gnomAD shows only ultra-rare singletons at contact residues. The absence of archaic-derived substitutions in the contacting nuclear residues provides evidence against a coding incompatibility of the kind predicted.

### Summary

Taken together, these results provide no evidence of the archaic-specific nuclear changes at mtDNA-contacting residues that a DMI would predict. In particular, the findings provide no evidence that the Denisovan mtDNA's failure to persist after introgression is explainable by a protein-coding mitochondrial-nuclear DMI at OXPHOS structural interfaces. This negative result is parallel to that of a previous study that searched for a mitonuclear incompatibility signature in the archaic ancestry retained in modern human genomes[7], which would reflect incompatibilities between archaic nuclear variants and modern human mtDNA arising during a later admixture event: the expansion of modern humans out of Africa and the Near East and admixture with Neandertals after around 50 ka.

The tests presented here are not definitive: the method I use is not perfectly sensitive to real DMIs, and research in swordtail fish hybrids identified a mitochondrial-nuclear DMI at the ATP synthase complex that was not mediated by contacting substitutions[21].

Overall, these results increase the weight of evidence for one of two conclusions: either no mtDNA-nuclear incompatibility drove the fixation of early modern human mtDNA in Neandertal ancestors, or such an incompatibility existed but acted through mechanisms not visible to this analysis, such as regulatory or expression-level mitochondrial-nuclear mismatch, mtDNA replication, or copy-number effects.

**Software and Reproducibility**

A zipped folder associated with this supplementary note (SI2codebase.zip) includes all the code necessary to replicate the results, using Python 3.11 (see also the README). The pipeline has two parts: a laptop analysis that produces the structural, catalog, and intrinsic-impact results, and two O2 (SLURM) pipelines that close the remaining gaps.

**Sequence data and fixation filter.** ncbi.py / panel.py / fetch_panel.py / analyze.py / analyze_per_individual.py: fetch the mtDNA sequences and OXPHOS panel, call and ape-polarize the coding changes, and build the multi-individual fixation filter that yields the eight Denisovan-specific nonsynonymous sites (analyze.py); analyze_per_individual.py extends the same alignment and polarization to every change derived in any single Denisovan, producing the per-individual genotypes and the fixed/non-fixed classification tabulated in Table S2.1.

**Structural interfaces.** contacts.py / lineage_table.py: compute inter-chain heavy-atom contacts (≤4 Å interface, ≤8 Å shell) from the cryo-EM structures, reconcile numbering to UniProt via SIFTS, and assemble the mt-residue→nuclear-partner contact table.

**Allele-level tests.** task3_workflow.js / task4_coevolution.py: the allele-level tests (catalog closure, gnomAD, intrinsic APOGEE2 impact, mammalian conservation) and the assembly of the matched Neandertal-lineage control set of mtDNA changes and their nuclear partners. The first of these is an agent-workflow definition rather than an analysis script. It specifies and dispatches the five convergent allele-level queries against the published catalogs, and its results are checked individually and, for the archaic genotypes, confirmed directly by the O2 nuclear scan below. extract_pdfs.py caches the text of the published catalog PDFs (data/papertext/) that the workflow queries.

**Cluster pipelines.** prep_o2_inputs.py and the o2/ folder: 6the SLURM pipelines: nuclear_scan/ (01_partner_regions.py, 02_slice_merge_annotate.sh, 03_classify.py: direct Denisova-vs-Neandertal-vs-modern genotyping of the partner genes), with environment.yml and a step-by-step README.

**Archaic genotyping.** o2/nuclear_scan/partner_archaic_alleles.py: genotypes the nuclear OXPHOS-partner contact residues in the high-coverage archaic genomes (Table S2.1, "Derived allele at partner?" column). Denisova 3 and the Altai Neandertal (local O2 VCFs) and Denisova 25 (the ~200 ka molar; EVA per-chromosome VCFs, read remotely, including COX7B on chromosome X) are all homozygous reference / ancestral at every partner residue (no archaic carries a derived partner allele). Run on the O2 cluster.

**Denisova 25 mtDNA.** I reconstructed the Denisova 25 (~200 ka) mtDNA genome to 382× depth from the EVA raw per-library BAM (16,569/16,569 positions covered; o2/den25/extract_den25_mt.sh, consensus_from_bam.py, genotype_den25_mt.py) and genotyped the fifteen Denisovan-derived sites. Seven of the eight fixed changes (all but *ATP6* A7T), together with the alternative site *ND5* S515P, are derived in Denisova 25, confirming that it lies on the Denisovan mt lineage. *ATP6* A7T is ancestral, so that interface change arose after the divergence of the Denisova 25 lineage and was probably absent at the introgression. All four Denisova 3+4 non-fixed changes are ancestral in Denisova 25 (a recent, post-introgression sub-lineage origin); *CYTB* L149M is shared by Denisova 8 and Denisova 25. Run on the O2 cluster.

**Dependencies and pipeline guide.** requirements.txt and environment.yml list the dependencies (Biopython, NumPy, pysam, plus bcftools/VEP/samtools for the O2 arm), and README.md gives the full script-by-script pipeline with the exact commands used for each reported number.
