## Supplementary Information section 2 code base for "Hypothesis: A modern human range expansion ∼300,000 years ago had a critical role in Neandertal origins": analyze.py

```
#!/usr/bin/env python3
"""Identify Denisovan-specific nonsynonymous mtDNA sites.

For each protein-coding gene, align every genome's translation to the rCRS
(modern human) translation and read positions in rCRS residue coordinates.
A 'Denisovan-specific' site = residue fixed across Denisovan individuals that
is absent from both modern human (rCRS) and ALL sampled Neandertals.
Polarized against great-ape outgroups (chimp/bonobo/gorilla).
"""
import os, sys, json
sys.path.insert(0, os.path.dirname(os.path.abspath(__file__)))
import panel

ROOT = os.path.dirname(os.path.dirname(os.path.abspath(__file__)))
G = json.load(open(os.path.join(ROOT, "data", "proteins.json")))

COMPLEX = {
    "ND1":"CI","ND2":"CI","ND3":"CI","ND4":"CI","ND4L":"CI","ND5":"CI","ND6":"CI",
    "CYTB":"CIII","COX1":"CIV","COX2":"CIV","COX3":"CIV","ATP6":"CV","ATP8":"CV",
}

def nw_align(ref, seq, match=2, mismatch=-1, gap=-2):
    """Global Needleman-Wunsch. Returns aligned (ref, seq) with '-' gaps."""
    n, m = len(ref), len(seq)
    # score & traceback
    S = [[0]*(m+1) for _ in range(n+1)]
    for i in range(1, n+1): S[i][0] = gap*i
    for j in range(1, m+1): S[0][j] = gap*j
    for i in range(1, n+1):
        ri = ref[i-1]
        Si, Sim1 = S[i], S[i-1]
        for j in range(1, m+1):
            d = Sim1[j-1] + (match if ri == seq[j-1] else mismatch)
            Si[j] = d if d >= Si[j-1]+gap and d >= Sim1[j]+gap else max(Si[j-1]+gap, Sim1[j]+gap)
    # traceback
    ar, asq = [], []
    i, j = n, m
    while i > 0 and j > 0:
        s = S[i][j]
        if s == S[i-1][j-1] + (match if ref[i-1]==seq[j-1] else mismatch):
            ar.append(ref[i-1]); asq.append(seq[j-1]); i-=1; j-=1
        elif s == S[i-1][j] + gap:
            ar.append(ref[i-1]); asq.append('-'); i-=1
        else:
            ar.append('-'); asq.append(seq[j-1]); j-=1
    while i > 0: ar.append(ref[i-1]); asq.append('-'); i-=1
    while j > 0: ar.append('-'); asq.append(seq[j-1]); j-=1
    return "".join(reversed(ar)), "".join(reversed(asq))

def ref_coord_residues(ref, seq):
    """Map seq onto ref columns -> list length len(ref); each = residue or None(gap)."""
    ar, asq = nw_align(ref, seq)
    out, ri = [], 0
    res = [None]*len(ref)
    for a, b in zip(ar, asq):
        if a != '-':
            res[ri] = (b if b != '-' else None)
            ri += 1
    return res

def main():
    REF = "rCRS_hg_H2a2a1"
    deni = list(panel.DENISOVAN)
    nean = list(panel.NEANDERTAL)
    outg = list(panel.OUTGROUP)
    candidates = []
    for g in panel.GENES:
        ref = G[REF]['genes'].get(g)
        if not ref: continue
        # map each genome onto ref coordinates
        mapped = {}
        for label in G:
            s = G[label]['genes'].get(g)
            # guard against the Denisova4 ATP8 misannotation (length far off)
            if not s or (g=="ATP8" and len(s) > 120):
                mapped[label] = [None]*len(ref); continue
            mapped[label] = ref_coord_residues(ref, s)
        for c in range(len(ref)):
            H = mapped[REF][c]
            if H is None: continue
            dres = [mapped[k][c] for k in deni if mapped[k][c] is not None]
            nres = [mapped[k][c] for k in nean if mapped[k][c] is not None]
            if len(dres) < 2 or len(nres) < 3: continue          # need coverage
            dset, nset = set(dres), set(nres)
            if len(dset) != 1: continue                          # fixed in Denisovans
            D = next(iter(dset))
            if D == H: continue                                  # differs from modern human
            if D in nset: continue                               # absent from Neandertals
            ores = [mapped[k][c] for k in outg if mapped[k][c] is not None]
            oset = set(ores)
            # polarization
            if D in oset:
                pol = "Denisova-ANCESTRAL (H/Nean derived)"
            elif (H in oset) or (nset & oset):
                pol = "Denisova-DERIVED"
            else:
                pol = "ambiguous"
            candidates.append({
                "gene": g, "complex": COMPLEX[g], "res": c+1,
                "human": H, "denisova": D,
                "neander": "".join(sorted(nset)),
                "outgroup": "".join(sorted(oset)),
                "polarization": pol,
                "deni_n": len(dres), "nean_n": len(nres),
            })
    # report
    der = [x for x in candidates if x["polarization"]=="Denisova-DERIVED"]
    print(f"Total Denisova-distinct nonsynonymous sites (fixed in Deni, absent in modern human + all Neandertals): {len(candidates)}")
    print(f"  of which Denisova-DERIVED (DMI-relevant): {len(der)}")
    print(f"  Denisova-ancestral / ambiguous: {len(candidates)-len(der)}\n")
    hdr = f"{'gene':5} {'cplx':4} {'res':>4}  {'chg':7} {'Nean':5} {'Out':5} polarization        cov(D/N)"
    print(hdr); print("-"*len(hdr))
    order = {"CI":0,"CIII":1,"CIV":2,"CV":3}
    for x in sorted(candidates, key=lambda r:(order[r['complex']], r['gene'], r['res'])):
        chg = f"{x['human']}{x['res']}{x['denisova']}"
        print(f"{x['gene']:5} {x['complex']:4} {x['res']:>4}  {chg:7} {x['neander']:5} {x['outgroup']:5} {x['polarization']:19} {x['deni_n']}/{x['nean_n']}")
    json.dump(candidates, open(os.path.join(ROOT,"results","denisova_specific_sites.json"),"w"), indent=2)
    print("\nwrote results/denisova_specific_sites.json")

if __name__ == "__main__":
    main()
```
