## Supplementary Information section 2 code base for "Hypothesis: A modern human range expansion ∼300,000 years ago had a critical role in Neandertal origins": analyze_per_individual.py

```
#!/usr/bin/env python3
"""Per-individual / any-Denisovan nonsynonymous mtDNA scan that reproduces Table S2.1.
Unlike analyze.py (which keeps only sites FIXED across Denisovans, yielding the eight
high-confidence sites), this reports every nonsynonymous change DERIVED in ANY of the
four sequenced Denisovans (Den2/Den3/Den4/Den8) and absent from all sampled Neandertals
and rCRS, with per-individual genotypes and fixed-vs-non-fixed status. Private premature
stop calls are flagged (probable aDNA/alignment artifacts) and excluded from the count.
Polarized against the great-ape outgroups (vertebrate-mitochondrial code, via proteins.json)."""
import os, sys, json
sys.path.insert(0, os.path.join(os.path.dirname(os.path.abspath(__file__))))
import panel
from analyze import ref_coord_residues
ROOT = os.path.dirname(os.path.dirname(os.path.abspath(__file__)))
G = json.load(open(os.path.join(ROOT, "data", "proteins.json")))
REF = "rCRS_hg_H2a2a1"
DEN = [("Den2","Denisova2"),("Den3","Denisova3_phalanx"),("Den4","Denisova4_molar"),("Den8","Denisova8")]
nean = list(panel.NEANDERTAL); outg = list(panel.OUTGROUP)
CPLX = {"ND1":"CI","ND2":"CI","ND3":"CI","ND4":"CI","ND4L":"CI","ND5":"CI","ND6":"CI",
        "CYTB":"CIII","COX1":"CIV","COX2":"CIV","COX3":"CIV","ATP6":"CV","ATP8":"CV"}
rows=[]
for g in panel.GENES:
    ref=G[REF]['genes'].get(g)
    if not ref: continue
    mapped={}
    for lab in G:
        s=G[lab]['genes'].get(g)
        mapped[lab]=([None]*len(ref)) if (not s or (g=="ATP8" and len(s)>120)) else ref_coord_residues(ref,s)
    for c in range(len(ref)):
        H=mapped[REF][c]
        if H is None: continue
        gt={sh:mapped[lab][c] for sh,lab in DEN}
        cov=[sh for sh in gt if gt[sh] is not None]
        if not cov: continue
        nres=[mapped[k][c] for k in nean if mapped[k][c] is not None]
        if len(nres)<3: continue
        nset=set(nres)
        for D in {gt[sh] for sh in cov if gt[sh]!=H and gt[sh] is not None}:
            if D in nset: continue
            oset=set(m for m in (mapped[k][c] for k in outg) if m)
            if D in oset: pol="ancestral"
            elif (H in oset) or (nset&oset): pol="DERIVED"
            else: pol="ambiguous"
            carriers=[sh for sh in cov if gt[sh]==D]
            rows.append(dict(gene=g,cplx=CPLX[g],res=c+1,H=H,D=D,pol=pol,carriers=carriers,
                fixed=(all(gt[sh]==D for sh in cov) and len(cov)>=2),
                gt="/".join(gt[sh] or "?" for sh,_ in DEN), stop=(D=="X" or D=="*")))
order={"CI":0,"CIII":1,"CIV":2,"CV":3}
der=[r for r in rows if r['pol']=="DERIVED"]
real=[r for r in der if not r['stop']]; stops=[r for r in der if r['stop']]
fixed=[r for r in real if r['fixed']]; nonf=[r for r in real if not r['fixed']]
print(f"DERIVED missense in any Denisovan: {len(real)}  ({len(fixed)} fixed + {len(nonf)} non-fixed)")
print("excluded private premature-stop calls:", len(stops), "->", [r['gene']+" "+str(r['res']) for r in stops])
print(f"\n{'site':10}{'H>D':6}{'fixed':6}{'Den2/Den3/Den4/Den8':22}carriers")
for r in sorted(real,key=lambda r:(0 if r['fixed'] else 1,order[r['cplx']],r['gene'],r['res'])):
    print(f"{r['gene']+' '+str(r['res']):10}{r['H']+'>'+r['D']:6}{('YES' if r['fixed'] else 'no'):6}{r['gt']:22}{','.join(r['carriers'])}")
```
