## Supplementary Information section 2 code base for "Hypothesis: A modern human range expansion ∼300,000 years ago had a critical role in Neandertal origins": contacts.py

```
#!/usr/bin/env python3
"""Map mt-encoded variant residues onto human OXPHOS structures and find
contacting nuclear-encoded subunits (+ cofactors).

For each target (gene, rCRS-residue) we:
  - locate the chain whose sequence matches that mt-encoded protein,
  - map the rCRS residue number to the PDB author numbering by alignment,
  - find every atom of OTHER chains/ligands within 8 A (heavy-atom),
  - classify each partner chain as nuclear-encoded / mt-encoded / cofactor,
  - report min distance and contacting partner residues at <=4 A and <=8 A.
"""
import os, sys, json, warnings
sys.path.insert(0, os.path.dirname(os.path.abspath(__file__)))
import numpy as np
from Bio.PDB import MMCIFParser, NeighborSearch
from Bio.PDB.MMCIF2Dict import MMCIF2Dict
from Bio.Align import PairwiseAligner, substitution_matrices
warnings.filterwarnings("ignore")

ROOT = os.path.dirname(os.path.dirname(os.path.abspath(__file__)))
RCRS = json.load(open(os.path.join(ROOT,"data","proteins.json")))["rCRS_hg_H2a2a1"]["genes"]

AA3to1 = {'ALA':'A','ARG':'R','ASN':'N','ASP':'D','CYS':'C','GLN':'Q','GLU':'E',
 'GLY':'G','HIS':'H','ILE':'I','LEU':'L','LYS':'K','MET':'M','PHE':'F','PRO':'P',
 'SER':'S','THR':'T','TRP':'W','TYR':'Y','VAL':'V','MSE':'M','SEC':'U'}
COFACTORS = {'HEM','HEA','HEB','HEC','HAS','FES','SF4','FMN','NDP','NAD','FAD',
 'ZN','CU','CU1','CUA','CUB','MG','CA','NA','K','CDL','PEE','PEF','PC1','PGV',
 'PLC','LMT','MC3','3PE','UQ','UQ1','DUM','SQD','LHG','CL','ACT','MN','HOH'}

aligner = PairwiseAligner()
aligner.substitution_matrix = substitution_matrices.load("BLOSUM62")
aligner.open_gap_score = -11; aligner.extend_gap_score = -1
aligner.mode = "global"

def chain_seq(chain):
    """list of (author_resseq:int, icode, one_letter, residue_obj) for std aa."""
    out=[]
    for res in chain:
        het, resseq, icode = res.id
        nm = res.resname.strip()
        if nm in AA3to1 and nm not in ('HOH',):
            out.append((resseq, icode, AA3to1[nm], res))
    return out

def best_gene(seq1):
    """return (gene, identity) of best matching mt protein, else (None,0)."""
    best=(None,0.0)
    for g,ref in RCRS.items():
        if abs(len(ref)-len(seq1))>0.4*len(ref) and len(seq1)>0:
            pass
        try:
            aln=aligner.align(ref, seq1)[0]
        except Exception:
            continue
        a,b = aln[0], aln[1]
        m=sum(1 for x,y in zip(a,b) if x==y and x!='-')
        ident=m/max(1,min(len(ref),len(seq1)))
        if ident>best[1]: best=(g,ident)
    return best

def map_rcrs_to_auth(gene, seq_tuples):
    """rCRS residue number(1-based) -> author resseq, via alignment."""
    ref=RCRS[gene]; chseq="".join(t[2] for t in seq_tuples)
    aln=aligner.align(ref, chseq)[0]
    a,b=aln[0],aln[1]
    ri=0; ci=0; mp={}
    for x,y in zip(a,b):
        if x!='-' and y!='-':
            mp[ri+1]=seq_tuples[ci][0]        # author resseq
        if x!='-': ri+=1
        if y!='-': ci+=1
    return mp

def entity_desc(cif):
    d=MMCIF2Dict(cif)
    def aslist(k):
        v=d.get(k);
        return v if isinstance(v,list) else ([v] if v is not None else [])
    ch=aslist('_atom_site.auth_asym_id'); ent=aslist('_atom_site.label_entity_id')
    c2e={}
    for c,e in zip(ch,ent): c2e.setdefault(c,e)
    eid=aslist('_entity.id'); des=aslist('_entity.pdbx_description')
    e2d=dict(zip(eid,des))
    return {c:e2d.get(e,"?") for c,e in c2e.items()}

def analyze(struct_id, targets):
    cif=os.path.join(ROOT,"data","pdb",f"{struct_id}.cif")
    model=MMCIFParser(QUIET=True).get_structure(struct_id, cif)[0]
    desc=entity_desc(cif)
    # classify chains
    chain_info={}   # cid -> ('mt:GENE'|'nuc', seq_tuples)
    target_chain={} # gene -> cid
    for ch in model:
        st=chain_seq(ch)
        if len(st)<20:
            chain_info[ch.id]=('other',st); continue
        g,ident=best_gene("".join(t[2] for t in st))
        if ident>0.9:
            chain_info[ch.id]=('mt:'+g,st); target_chain.setdefault(g,ch.id)
        else:
            chain_info[ch.id]=('nuc',st)
    atoms=[a for a in model.get_atoms()]
    ns=NeighborSearch(atoms)
    results=[]
    for gene,resnum,tag in targets:
        cid=target_chain.get(gene)
        if cid is None:
            results.append({"tag":tag,"gene":gene,"res":resnum,"struct":struct_id,"error":"chain not found"}); continue
        mp=map_rcrs_to_auth(gene, chain_info[cid][1])
        auth=mp.get(resnum)
        if auth is None:
            results.append({"tag":tag,"gene":gene,"res":resnum,"struct":struct_id,"error":"residue unmodeled"}); continue
        # target residue atoms
        tres=None
        for r in model[cid]:
            if r.id[1]==auth: tres=r; break
        tatoms=[a for a in tres] if tres else []
        partners={}   # (pcid, presseq, presname) -> min dist
        cofac={}
        for a in tatoms:
            for nb in ns.search(a.coord, 8.0, level='A'):
                pr=nb.get_parent(); pcid=pr.get_parent().id
                if pcid==cid: continue
                d=float(np.linalg.norm(a.coord-nb.coord))
                nm=pr.resname.strip()
                if nm in COFACTORS or pr.id[0].startswith('H_') and nm not in AA3to1:
                    key=(pcid,nm,pr.id[1]); cofac[key]=min(cofac.get(key,9),d)
                else:
                    key=(pcid,pr.id[1],nm); partners[key]=min(partners.get(key,9),d)
        # group by partner chain
        bychain={}
        for (pcid,pres,pnm),d in partners.items():
            kind,_=chain_info.get(pcid,('?',None))
            bychain.setdefault((pcid,kind),[]).append((pres,pnm,d))
        results.append({"tag":tag,"gene":gene,"res":resnum,"struct":struct_id,"auth":auth,
            "partners":{f"{pcid}|{kind}|{desc.get(pcid,'?')}":sorted(v,key=lambda x:x[2])
                        for (pcid,kind),v in bychain.items()},
            "cofactors":sorted([(nm,pcid,d) for (pcid,nm,_),d in cofac.items()],key=lambda x:x[2])})
    return results

# ---- targets ----
DENI=[("ND2",64,"DENI"),("ND2",164,"DENI"),("ND5",398,"DENI"),("CYTB",70,"DENI"),
      ("COX1",456,"DENI"),("COX2",27,"DENI"),("ATP6",7,"DENI"),("ATP6",185,"DENI")]
NEAN=[("ND1",10,"NEAN"),("ND1",168,"NEAN"),("ND2",346,"NEAN"),("ATP6",80,"NEAN"),
      ("ATP6",154,"NEAN"),("ATP6",176,"NEAN"),("COX3",84,"NEAN"),("ND5",489,"NEAN"),
      ("ND5",603,"NEAN"),("ND6",166,"NEAN"),("CYTB",245,"NEAN"),("CYTB",309,"NEAN")]
STRUCT={"ND1":"5XTD","ND2":"5XTD","ND5":"5XTD","ND6":"5XTD",
        "CYTB":"9HZL","COX1":"5Z62","COX2":"5Z62","COX3":"5Z62","ATP6":"8H9S","ATP8":"8H9S"}

def main():
    allt=DENI+NEAN
    bystruct={}
    for g,r,tag in allt: bystruct.setdefault(STRUCT[g],[]).append((g,r,tag))
    out=[]
    for sid,tg in bystruct.items():
        print(f"--- {sid}: {len(tg)} sites ---", flush=True)
        out+=analyze(sid,tg)
    json.dump(out, open(os.path.join(ROOT,"results","contacts.json"),"w"), indent=2)
    # print summary
    print("\n================ CONTACT SUMMARY ================")
    for r in out:
        head=f"[{r['tag']}] {r['gene']} {RCRS[r['gene']][r['res']-1]}{r['res']}  ({r['struct']})"
        if r.get("error"): print(head, "->", r["error"]); continue
        nucs=[]
        for k,v in r["partners"].items():
            pcid,kind,dsc=k.split("|",2)
            mind=min(x[2] for x in v)
            nucs.append((mind,kind,dsc,pcid,v))
        nucs.sort()
        print(f"\n{head}")
        for mind,kind,dsc,pcid,v in nucs:
            within4=[f"{nm}{pr}" for pr,nm,d in v if d<=4.0]
            label = "NUCLEAR" if kind=="nuc" else ("mt:"+kind.split(':')[1] if kind.startswith('mt') else kind)
            mark = "  <<< INTERFACE" if (kind=="nuc" and within4) else ""
            print(f"   {mind:4.1f}A {label:9} {dsc[:48]:48} contacts<=4A:{within4}{mark}")
        if r["cofactors"]:
            cof=[f"{nm}({d:.1f})" for nm,pcid,d in r["cofactors"][:4]]
            print(f"        cofactors<=8A: {cof}")
    json.dump(out, open(os.path.join(ROOT,"results","contacts.json"),"w"), indent=2)

if __name__=="__main__":
    main()
```
