## Supplementary Information section 2 code base for "Hypothesis: A modern human range expansion ∼300,000 years ago had a critical role in Neandertal origins": extract_pdfs.py

```
#!/usr/bin/env python3
"""Extract text from all paper PDFs to data/papertext/<name>.txt (cached)."""
import os, sys, glob
from pypdf import PdfReader
ROOT = os.path.dirname(os.path.dirname(os.path.abspath(__file__)))
OUT = os.path.join(ROOT, "data", "papertext"); os.makedirs(OUT, exist_ok=True)
for pdf in sorted(glob.glob(os.path.join(ROOT, "*.pdf"))):
    name = os.path.splitext(os.path.basename(pdf))[0]
    txt = os.path.join(OUT, name + ".txt")
    if os.path.exists(txt) and os.path.getsize(txt) > 0:
        print("cached", name); continue
    try:
        r = PdfReader(pdf)
        pages = []
        for i, p in enumerate(r.pages):
            try: pages.append(f"\n===PAGE {i+1}===\n" + (p.extract_text() or ""))
            except Exception as e: pages.append(f"\n===PAGE {i+1}=== [extract error {e}]")
        open(txt, "w").write("".join(pages))
        print(f"{name}: {len(r.pages)} pages -> {os.path.getsize(txt)} bytes")
    except Exception as e:
        print(f"{name}: FAILED {e}")
```
