## Supplementary Information section 2 code base for "Hypothesis: A modern human range expansion ∼300,000 years ago had a critical role in Neandertal origins": fetch_panel.py

```
#!/usr/bin/env python3
"""Fetch fasta_cds_aa for every genome in the panel, parse per-gene proteins."""
import os, re, json, sys
sys.path.insert(0, os.path.dirname(os.path.abspath(__file__)))
import ncbi, panel

def parse_cds_aa(text):
    """Return {GENE: protein_seq} from a fasta_cds_aa record."""
    out = {}
    name = None
    buf = []
    for line in text.splitlines():
        if line.startswith(">"):
            if name:
                out[name] = "".join(buf)
            m = re.search(r"\[gene=([^\]]+)\]", line)
            name = m.group(1).upper() if m else None
            buf = []
        else:
            buf.append(line.strip())
    if name:
        out[name] = "".join(buf)
    return out

def main():
    genomes = {}
    for label, acc in panel.ALL.items():
        txt = ncbi.efetch(acc, "fasta_cds_aa")
        genes = parse_cds_aa(txt)
        # strip trailing stop '*' if present
        genes = {g: s.rstrip("*") for g, s in genes.items()}
        genomes[label] = {"acc": acc, "genes": genes}
        present = [g for g in panel.GENES if g in genes]
        missing = [g for g in panel.GENES if g not in genes]
        print(f"{label:26s} {acc:14s} {len(present)}/13 genes" + (f"  MISSING:{missing}" if missing else ""))
    out = os.path.join(os.path.dirname(os.path.dirname(os.path.abspath(__file__))), "data", "proteins.json")
    json.dump(genomes, open(out, "w"), indent=0)
    print("wrote", out)

if __name__ == "__main__":
    main()
```
