## Supplementary Information section 2 code base for "Hypothesis: A modern human range expansion ∼300,000 years ago had a critical role in Neandertal origins": lineage_table.py

```
#!/usr/bin/env python3
"""Codon-aware partition of mtDNA coding substitutions by lineage.

One representative genome per group (Denisova 3, Vindija 33.19, rCRS), polarized
by great-ape majority (chimp/bonobo/gorilla). For every variable coding
nucleotide site we assign a lineage pattern and classify it synonymous vs
nonsynonymous under the VERTEBRATE MITOCHONDRIAL code (NCBI transl_table 2).
"""
import os, sys, json
sys.path.insert(0, os.path.dirname(os.path.abspath(__file__)))
from analyze import nw_align  # reuse generic NW aligner

ROOT = os.path.dirname(os.path.dirname(os.path.abspath(__file__)))
DATA = json.load(open(os.path.join(ROOT, "data", "cds_na.json")))
GENES = ["ND1","ND2","COX1","COX2","ATP8","ATP6","COX3","ND3","ND4L","ND4","ND5","ND6","CYTB"]

# ---- vertebrate mitochondrial genetic code (transl_table 2) ----
_bases = "TCAG"
_std = ("FFLLSSSSYY**CC*WLLLLPPPPHHQQRRRRIIMMTTTTNNKKSSRRVVVVAAAADDEEGGGG")
CODON = {}
i = 0
for a in _bases:
    for b in _bases:
        for c in _bases:
            CODON[a+b+c] = _std[i]; i += 1
# vertebrate-mito overrides
CODON.update({"AGA":"*","AGG":"*","ATA":"M","TGA":"W"})
def tr(codon):
    return CODON.get(codon.upper(), "X")

def map_to_ref(ref, seq):
    """Return list of len(ref): seq nucleotide per ref column ('-'/None = gap)."""
    if len(seq) == len(ref):
        return list(seq)                      # no coding indels among Homo
    ar, asq = nw_align(ref, seq, match=2, mismatch=-1, gap=-3)
    out, ri = [None]*len(ref), 0
    for a, b in zip(ar, asq):
        if a != '-':
            out[ri] = (b if b != '-' else None); ri += 1
    return out

def main():
    REF = "rCRS"
    apes = ["chimp","bonobo","gorilla"]
    cats = ["Denisova-only","Neandertal-only","Modern-human-only",
            "Shared Modern+Neandertal","Other/homoplasy/unpolarized"]
    counts = {c: {"NS":0, "S":0} for c in cats}
    deni_ns, nean_ns = [], []     # detailed nonsynonymous autapomorphy lists
    modern_ns, sharedmn_ns = [], []  # modern-only and shared Modern+Neandertal nonsyn
    multi = 0
    for g in GENES:
        ref = DATA[REF][g]
        ncod = len(ref)//3
        M = {t: map_to_ref(ref, DATA[t][g]) for t in ["Denisova3","Vindija33.19"]+apes}
        Mref = list(ref)
        for ci in range(ncod):
            cols = [3*ci, 3*ci+1, 3*ci+2]
            ref_codon = "".join(Mref[c] for c in cols)
            if "N" in ref_codon.upper() or "-" in ref_codon: continue
            nvar = 0
            for fp, p in enumerate(cols):
                d = M["Denisova3"][p]; n = M["Vindija33.19"][p]; m = Mref[p]
                if None in (d, n, m): continue
                d,n,m = d.upper(), n.upper(), m.upper()
                if d == n == m: continue              # invariant among ingroup
                nvar += 1
                ape_alleles = [M[a][p] for a in apes if M[a][p] not in (None,"-")]
                ape_alleles = [x.upper() for x in ape_alleles]
                anc = None
                if ape_alleles:
                    from collections import Counter
                    top, c = Counter(ape_alleles).most_common(1)[0]
                    if c >= 2 or len(ape_alleles)==1: anc = top
                # classify pattern + polarization
                cat, derived, ancestral = "Other/homoplasy/unpolarized", None, None
                if d != n and d != m and n == m:           # D differs ; N==M
                    if anc == n: cat, derived, ancestral = "Denisova-only", d, n
                    elif anc == d: cat, derived, ancestral = "Shared Modern+Neandertal", m, d
                elif n != d and n != m and d == m:         # N differs ; D==M
                    if anc == d: cat, derived, ancestral = "Neandertal-only", n, d
                elif m != d and m != n and d == n:         # M differs ; D==N
                    if anc == d: cat, derived, ancestral = "Modern-human-only", m, d
                # syn / nonsyn using rCRS codon context
                if derived is None:                        # unpolarized/all-diff
                    # still record syn/nonsyn vs rCRS state for the "other" bin
                    alt = d if (d!=m) else n
                    ancestral, derived = m, alt
                cod_a = list(ref_codon); cod_a[fp] = ancestral
                cod_d = list(ref_codon); cod_d[fp] = derived
                ns = tr("".join(cod_a)) != tr("".join(cod_d))
                counts[cat]["NS" if ns else "S"] += 1
                if ns and cat == "Denisova-only":
                    deni_ns.append((g, ci+1, tr("".join(cod_a))+str(ci+1)+tr("".join(cod_d)), anc))
                if ns and cat == "Neandertal-only":
                    nean_ns.append((g, ci+1, tr("".join(cod_a))+str(ci+1)+tr("".join(cod_d)), anc))
                if ns and cat == "Modern-human-only":
                    modern_ns.append((g, ci+1, tr("".join(cod_a))+str(ci+1)+tr("".join(cod_d)), anc))
                if ns and cat == "Shared Modern+Neandertal":
                    sharedmn_ns.append((g, ci+1, tr("".join(cod_a))+str(ci+1)+tr("".join(cod_d)), anc))
            if nvar > 1: multi += 1

    # ---- report ----
    print("Codon-aware lineage partition of mtDNA coding substitutions")
    print("(Denisova 3 vs Vindija 33.19 vs rCRS; polarized by great-ape majority; transl_table 2)\n")
    w = 30
    print(f"{'Lineage pattern':{w}} {'Nonsyn':>7} {'Syn':>5} {'Total':>6} {'NS/S':>6}")
    print("-"*(w+28))
    tot_ns=tot_s=0
    for c in cats:
        ns, s = counts[c]["NS"], counts[c]["S"]; tot_ns+=ns; tot_s+=s
        ratio = f"{ns/s:.2f}" if s else ("inf" if ns else "-")
        print(f"{c:{w}} {ns:>7} {s:>5} {ns+s:>6} {ratio:>6}")
    print("-"*(w+28))
    print(f"{'TOTAL':{w}} {tot_ns:>7} {tot_s:>5} {tot_ns+tot_s:>6} {tot_ns/tot_s:>6.2f}")
    print(f"\n(codons containing >1 variable site, syn/nonsyn approximated w/ rCRS context: {multi})")

    print("\nDenisova-only NONSYNONYMOUS sites (gene, residue, change, ape-ancestral nt):")
    for x in deni_ns: print("  ", x)
    print("\nNeandertal-only NONSYNONYMOUS sites (control):")
    for x in nean_ns: print("  ", x)
    print("\nModern-human-only NONSYNONYMOUS sites:")
    for x in modern_ns: print("  ", x)
    print("\nShared Modern+Neandertal NONSYNONYMOUS sites (present in modern AND Neandertal, absent Denisova):")
    for x in sharedmn_ns: print("  ", x)
    json.dump({"counts":counts,"deni_ns":deni_ns,"nean_ns":nean_ns,
               "modern_ns":modern_ns,"sharedmn_ns":sharedmn_ns},
              open(os.path.join(ROOT,"results","lineage_partition.json"),"w"), indent=2)

if __name__ == "__main__":
    main()
```
