## Supplementary Information section 2 code base for "Hypothesis: A modern human range expansion ∼300,000 years ago had a critical role in Neandertal origins": ncbi.py

```
#!/usr/bin/env python3
"""Minimal NCBI E-utilities helper (stdlib only). Caches fetches under data/."""
import sys, os, time, json, urllib.parse, urllib.request

BASE = "https://eutils.ncbi.nlm.nih.gov/entrez/eutils"
ROOT = os.path.dirname(os.path.dirname(os.path.abspath(__file__)))
CACHE = os.path.join(ROOT, "data")

def _get(url, tries=4):
    for i in range(tries):
        try:
            with urllib.request.urlopen(url, timeout=60) as r:
                return r.read().decode("utf-8", "replace")
        except Exception as e:
            if i == tries - 1:
                raise
            time.sleep(1.5 * (i + 1))

def esearch(term, retmax=100, db="nuccore"):
    q = urllib.parse.urlencode({"db": db, "term": term, "retmax": retmax, "retmode": "json"})
    d = json.loads(_get(f"{BASE}/esearch.fcgi?{q}"))
    return d["esearchresult"]["idlist"], d["esearchresult"]["count"]

def esummary(ids, db="nuccore"):
    q = urllib.parse.urlencode({"db": db, "id": ",".join(ids), "retmode": "json"})
    return json.loads(_get(f"{BASE}/esummary.fcgi?{q}"))

def efetch(acc, rettype, db="nuccore", cache=True):
    fn = os.path.join(CACHE, "genbank", f"{acc}.{rettype}")
    if cache and os.path.exists(fn) and os.path.getsize(fn) > 0:
        return open(fn).read()
    q = urllib.parse.urlencode({"db": db, "id": acc, "rettype": rettype, "retmode": "text"})
    txt = _get(f"{BASE}/efetch.fcgi?{q}")
    os.makedirs(os.path.dirname(fn), exist_ok=True)
    if cache:
        open(fn, "w").write(txt)
    time.sleep(0.4)
    return txt

if __name__ == "__main__":
    cmd = sys.argv[1]
    if cmd == "search":
        ids, cnt = esearch(sys.argv[2], retmax=int(sys.argv[3]) if len(sys.argv) > 3 else 100)
        print(f"count={cnt}")
        s = esummary(ids) if ids else {"result": {}}
        for i in ids:
            r = s["result"].get(i, {})
            print(f"{r.get('accessionversion','?'):16s} {r.get('slen','?'):>6} bp  {r.get('title','')}")
    elif cmd == "fetch":
        sys.stdout.write(efetch(sys.argv[2], sys.argv[3]))
```
