## Supplementary Information section 2 code base for "Hypothesis: A modern human range expansion ∼300,000 years ago had a critical role in Neandertal origins": consensus_from_bam.py

```
#!/usr/bin/env python3
"""Reference-free MT consensus from a sorted+indexed BAM (pysam pileup), for O2
samtools builds that lack `samtools consensus`. Majority base per rCRS position
1..16569 (the b37 'MT' contig = rCRS + 1000 bp wrap padding); positions with
depth<MINDP -> N.  Usage: consensus_from_bam.py in.sorted.bam out.fa [MINDP]"""
import sys
import pysam
from collections import Counter
bam=sys.argv[1]; out=sys.argv[2]; MINDP=int(sys.argv[3]) if len(sys.argv)>3 else 3
af=pysam.AlignmentFile(bam,"rb"); cons={}
for col in af.pileup("MT",0,16569,truncate=True,min_base_quality=20,max_depth=200000):
    c=Counter()
    for pr in col.pileups:
        if pr.is_del or pr.is_refskip or pr.query_position is None: continue
        c[pr.alignment.query_sequence[pr.query_position]]+=1
    if c:
        b,_=c.most_common(1)[0]
        cons[col.reference_pos+1]= b if sum(c.values())>=MINDP else "N"
af.close()
seq="".join(cons.get(i,"N") for i in range(1,16570))
open(out,"w").write(">Denisova25_MT\n"+seq+"\n")
print("wrote %s — %d/16569 covered"%(out,sum(1 for i in range(1,16570) if cons.get(i,'N')!='N')))
```
