## Supplementary Information section 2 code base for "Hypothesis: A modern human range expansion ∼300,000 years ago had a critical role in Neandertal origins": extract_den25_mt.sh

```
#!/bin/bash
# Extract the Denisova 25 mitochondrion from the EVA raw per-library BAMs and build
# a consensus. The rawBAMs have NO server-side .bai, so we stream each one over HTTP
# and keep only MT reads (no whole-file download, no sorting/indexing of the huge BAM).
#
# Reference = hs37d5/b37: the mt contig is "MT", length 17569 = rCRS 16569 + 1000 bp
# wrap-around padding, so positions 1..16569 are exactly rCRS coordinates.
#
# Run on O2:  export REPO=$HOME/dmi; conda activate dmi-o2; cd $REPO
#             bash o2/den25/extract_den25_mt.sh
set -euo pipefail
BASE="https://cdna.eva.mpg.de/denisova/Den25/rawBAM"
# Libraries, smallest-first (65,112,119,125 GB). mtDNA is high-copy, so ONE library is
# usually plenty of depth. Override with e.g. DEN25_LIBS="Denisova25.A45394.hg19.bam".
if [ -n "${DEN25_LIBS:-}" ]; then read -ra LIBS <<< "$DEN25_LIBS"
else LIBS=(Denisova25.A45394.hg19.bam Denisova25.A43615.hg19.bam Denisova25.A43617.hg19.bam Denisova25.A45393.hg19.bam); fi
OUT=o2/results/den25; mkdir -p "$OUT"

for L in "${LIBS[@]}"; do
  echo ">> streaming MT reads from $L (full-file HTTP stream; keeps only MT)"
  # -h keep header; awk keeps @headers + reads whose RNAME (col 3) is MT
  samtools view -h "$BASE/$L" \
    | awk 'BEGIN{OFS="\t"} /^@/ {print; next} $3=="MT"{print}' \
    | samtools view -b -o "$OUT/${L%.bam}.MT.bam" -
  echo "   $(samtools view -c "$OUT/${L%.bam}.MT.bam") MT reads"
done

echo ">> merge + sort + index"
samtools merge -f "$OUT/den25.MT.merged.bam" "$OUT"/*.MT.bam
samtools sort -o "$OUT/den25.MT.sorted.bam" "$OUT/den25.MT.merged.bam"
samtools index "$OUT/den25.MT.sorted.bam"
echo "   depth summary:"; samtools depth -a -r MT "$OUT/den25.MT.sorted.bam" | awk '{s+=$3;n++} END{printf "   mean MT depth = %.0f over %d positions\n", s/n, n}'

echo ">> consensus FASTA (rCRS 1..16569) via reference-free pysam pileup"
python3 o2/den25/consensus_from_bam.py "$OUT/den25.MT.sorted.bam" "$OUT/den25_MT.fa" 3
echo ">> done. consensus at $OUT/den25_MT.fa"
echo ">> next: python3 o2/den25/genotype_den25_mt.py $OUT/den25_MT.fa"
```
