## Supplementary Information section 2 code base for "Hypothesis: A modern human range expansion ∼300,000 years ago had a critical role in Neandertal origins": genotype_den25_mt.py

```
#!/usr/bin/env python3
"""Genotype the 15 Denisovan-derived mtDNA sites (8 fixed + 4 non-fixed, + the
ambiguous ATP6-185) in the Denisova 25 consensus produced by extract_den25_mt.sh.

Reads a single-record FASTA whose coordinates are rCRS (NC_012920) positions
1..16569, extracts the relevant codon per site (vertebrate-mitochondrial code,
transl_table 2; ND6 is on the light strand -> reverse-complement), and prints
Denisova 25's amino acid at each site next to the ancestral and Denisovan-derived
states. This is exactly the panel's site list, so the calls slot straight into
Table 1's Denisovan-genotype column.

Usage:  python3 o2/den25/genotype_den25_mt.py o2/results/den25/den25_MT.fa
"""
import sys

# standard rCRS (NC_012920) CDS coordinates, 1-based inclusive; strand
CDS = {
 "ND1":(3307,4262,'+'), "ND2":(4470,5511,'+'), "COX1":(5904,7445,'+'),
 "COX2":(7586,8269,'+'), "ATP8":(8366,8572,'+'), "ATP6":(8527,9207,'+'),
 "COX3":(9207,9990,'+'), "ND3":(10059,10404,'+'), "ND4L":(10470,10766,'+'),
 "ND4":(10760,12137,'+'), "ND5":(12337,14148,'+'), "ND6":(14149,14673,'-'),
 "CYTB":(14747,15887,'+'),
}
# vertebrate mitochondrial genetic code (transl_table 2)
BASES="TCAG"
STD=("FFLLSSSSYY**CC*WLLLLPPPPHHQQRRRRIIMMTTTTNNKKSSRRVVVVAAAADDEEGGGG")
CODONS=[a+b+c for a in BASES for b in BASES for c in BASES]
CODE=dict(zip(CODONS,STD))
CODE.update({"ATA":"M","TGA":"W","AGA":"*","AGG":"*"})  # mito differences
def comp(s): return s.translate(str.maketrans("ACGTNacgtn","TGCANtgcan"))
def rc(s): return comp(s)[::-1]

# (gene, residue, ancestral AA, Denisovan-derived AA, class)
SITES=[
 ("ND2",64,"A","T","fixed"),("ND2",164,"A","T","fixed"),("ND5",398,"T","A","fixed"),
 ("CYTB",70,"T","A","fixed"),("COX1",456,"V","A","fixed"),("COX2",27,"A","T","fixed"),
 ("ATP6",7,"A","T","fixed"),("ND5",515,"S","P","fixed (code); ATP6-185 in prose"),
 ("ATP6",185,"N","S","fixed (prose); ancestral by code — ambiguous"),
 ("ND1",153,"T","M","non-fixed (Den3,Den4)"),("ND2",163,"M","V","non-fixed (Den3,Den4)"),
 ("COX1",83,"V","I","non-fixed (Den3,Den4)"),("COX3",32,"A","T","non-fixed (Den3,Den4)"),
]

def main():
    fa=sys.argv[1] if len(sys.argv)>1 else "o2/results/den25/den25_MT.fa"
    seq="".join(l.strip() for l in open(fa) if not l.startswith(">")).upper()
    if len(seq)<16000: sys.exit(f"consensus too short ({len(seq)} bp) — check {fa}")
    def codon(gene,res):
        s,e,strand=CDS[gene]
        cds=seq[s-1:e]
        if strand=='-': cds=rc(cds)
        i=(res-1)*3
        return cds[i:i+3]
    print(f"{'site':10}{'anc>der':9}{'class':30}{'Den25 codon':12}{'Den25 AA':9}{'call'}")
    print("-"*90)
    for g,res,anc,der,cls in SITES:
        cod=codon(g,res); aa=CODE.get(cod,"?") if "N" not in cod and len(cod)==3 else "?"
        if aa=="?": call="no-call / low-cov (N in codon)"
        elif aa==der: call="DERIVED (Denisovan)"
        elif aa==anc: call="ancestral"
        else: call=f"other ({aa})"
        print(f"{g+' '+str(res):10}{anc+'>'+der:9}{cls:30}{cod:12}{aa:9}{call}")
    print("\nInterpretation: 'DERIVED' at a non-fixed site (ND1 153, ND2 163, COX1 83, COX3 32)")
    print("would mean the variant was already present in the 200-170 ka Denisovan (=> old);")
    print("'ancestral' confirms it arose later on the Denisova 3+4 sub-lineage, after the introgression.")

if __name__=="__main__":
    main()
```
