## Supplementary Information section 2 code base for "Hypothesis: A modern human range expansion ∼300,000 years ago had a critical role in Neandertal origins": environment.yml

```
name: dmi-o2
channels: [bioconda, conda-forge]
dependencies:
  - python=3.11
  - bcftools>=1.18
  - samtools>=1.18
  - htslib>=1.18          # tabix/bgzip with libcurl for remote VCF reads
  - ensembl-vep>=110      # GRCh37 cache: vep_install -a cf -s homo_sapiens -y GRCh37
  - pysam>=0.22
  - biopython>=1.83
  - numpy
  - pip
# After creating: conda activate dmi-o2
# VEP cache (once):  vep_install -a cf -s homo_sapiens -y GRCh37 -c $HOME/.vep
# GRCh37 fasta:      human_g1k_v37.fasta (set FASTA_GRCH37); index with `samtools faidx`
```
