## Supplementary Information section 2 code base for "Hypothesis: A modern human range expansion ∼300,000 years ago had a critical role in Neandertal origins": 01_partner_regions.py

```
#!/usr/bin/env python3
"""01 — Build GRCh37 CDS regions (BED) for the nuclear partner genes.

Output: o2/inputs/partner_cds_grch37.bed  (chrom  start0  end  gene  .  strand)
        o2/inputs/partner_gene_span_grch37.tsv (gene chrom start end strand ENSG ENSP)
Used by 02 (VCF slicing) and 03 (contact-residue flagging). Runs anywhere with
internet (Ensembl GRCh37 REST). Coordinates are 1-based inclusive from Ensembl;
BED is written 0-based half-open.
"""
import os, sys, json, time, urllib.request
ROOT = os.path.dirname(os.path.dirname(os.path.dirname(os.path.abspath(__file__))))
B = "https://grch37.rest.ensembl.org"

def get(url, tries=4):
    for i in range(tries):
        try:
            req = urllib.request.Request(url, headers={"Content-Type": "application/json"})
            return json.loads(urllib.request.urlopen(req, timeout=40).read())
        except Exception as e:
            if i == tries - 1: raise
            time.sleep(1.5 * (i + 1))

def load_partners():
    genes = []
    with open(os.path.join(ROOT, "o2", "inputs", "partner_genes_all.tsv")) as fh:
        next(fh)
        for ln in fh:
            g, uni, ens = (ln.rstrip("\n").split("\t") + ["", ""])[:3]
            genes.append((g, ens))
    return genes

def main():
    out_bed = open(os.path.join(ROOT, "o2", "inputs", "partner_cds_grch37.bed"), "w")
    out_span = open(os.path.join(ROOT, "o2", "inputs", "partner_gene_span_grch37.tsv"), "w")
    out_span.write("gene\tchrom\tstart\tend\tstrand\tENSG\tcanonical_ENST\tENSP\n")
    for gene, ens in load_partners():
        # Prefer stable Ensembl gene ID (GRCh37 server uses Ensembl-67-era symbols, so
        # current HGNC symbols like ATP5MF/ATP5PB fail symbol lookup).
        d = None
        if ens.startswith("ENSG"):
            try: d = get(f"{B}/lookup/id/{ens}?expand=1;content-type=application/json")
            except Exception: d = None
        if d is None:
            try: d = get(f"{B}/lookup/symbol/homo_sapiens/{gene}?expand=1;content-type=application/json")
            except Exception as e:
                print(f"{gene}: LOOKUP FAILED (id={ens}) {e}", file=sys.stderr); continue
        chrom = str(d["seq_region_name"]); strand = "+" if d["strand"] == 1 else "-"
        ensg = d["id"]
        tx = [t for t in d.get("Transcript", []) if t.get("is_canonical")] or d.get("Transcript", [])
        if not tx:
            print(f"{gene}: no transcript", file=sys.stderr); continue
        t = tx[0]; enst = t["id"]; ensp = t.get("Translation", {}).get("id", "")
        out_span.write(f"{gene}\t{chrom}\t{d['start']}\t{d['end']}\t{strand}\t{ensg}\t{enst}\t{ensp}\n")
        # CDS exon blocks (Translation gives genomic start/end of coding region per exon via /map)
        # simplest robust: use Exon list intersected with coding region [Translation.start, Translation.end]
        tr = t.get("Translation", {})
        cstart, cend = tr.get("start"), tr.get("end")
        for ex in t.get("Exon", []):
            s, e = ex["start"], ex["end"]
            if cstart and cend:               # clip to coding region
                s, e = max(s, cstart), min(e, cend)
                if s > e: continue
            out_bed.write(f"{chrom}\t{s-1}\t{e}\t{gene}\t.\t{strand}\n")
        print(f"{gene:8} chr{chrom}:{d['start']}-{d['end']} {strand} {ensg} {enst}")
        time.sleep(0.25)
    out_bed.close(); out_span.close()

if __name__ == "__main__":
    main()
```
