## Supplementary Information section 2 code base for "Hypothesis: A modern human range expansion ∼300,000 years ago had a critical role in Neandertal origins": 02_slice_merge_annotate.sh

```
#!/bin/bash
#SBATCH -J nuc_slice
#SBATCH -t 03:00:00
#SBATCH -c 4
#SBATCH --mem 12G
#SBATCH -p short
#SBATCH -o logs/02_slice_%j.out
# ---------------------------------------------------------------------------
# 02 — Slice the high-coverage archaic VCFs at the nuclear-partner CDS regions,
#      merge Denisova + Altai + Vindija into one multi-sample VCF, and annotate
#      coding consequences with Ensembl VEP (GRCh37). Output feeds 03_classify.py.
#
# Data: Prufer 2017 "Vindija" release, per-chromosome ALL-SITES VCFs
#   http://ftp.eva.mpg.de/neandertal/Vindija/VCF/{Altai,Denisova,Vindija33.19}/chrN_mq25_mapab100.vcf.gz
#   (internal contig names are '1'..'22','X'; INFO carries CAnc = chimp ancestral allele.)
# Requires: bcftools, htslib (tabix/bgzip built with libcurl for remote read), ensembl-vep
#           + a GRCh37 VEP cache (vep_install -a cf -s homo_sapiens -y GRCh37).
# Adjust module loads / paths for your O2 environment.
# ---------------------------------------------------------------------------
set -euo pipefail
cd "$(dirname "$0")/../.."                      # repo root
mkdir -p logs o2/results/nuclear_scan
OUT=o2/results/nuclear_scan
BED=o2/inputs/partner_cds_grch37.bed
BASE=http://ftp.eva.mpg.de/neandertal/Vindija/VCF
INDS=(Altai Denisova Vindija33.19)
VEP_CACHE=${VEP_CACHE:-$HOME/.vep}              # set to your GRCh37 cache dir
FASTA_GRCH37=${FASTA_GRCH37:-$HOME/ref/human_g1k_v37.fasta}  # GRCh37 reference

# module load gcc bcftools htslib ensembl-vep   # <-- uncomment / edit for O2

# tabix-style region list (1-based); BED is 0-based half-open so start+1
awk 'BEGIN{OFS=""}{print $1,":",($2+1),"-",$3}' "$BED" | sort -V -u > "$OUT/regions.txt"
CHRS=$(awk -F: '{print $1}' "$OUT/regions.txt" | sort -V -u)

for chr in $CHRS; do
  grep -E "^${chr}:" "$OUT/regions.txt" > "$OUT/reg_${chr}.txt"
  for ind in "${INDS[@]}"; do
    url="$BASE/$ind/chr${chr}_mq25_mapab100.vcf.gz"
    echo "[slice] $ind chr$chr"
    # -R reads only the indexed regions over HTTP; keep INFO (CAnc) and GT
    bcftools view -R "$OUT/reg_${chr}.txt" -Oz -o "$OUT/${ind}_chr${chr}.vcf.gz" "$url"
    # force a unique sample name so the merge keeps all three
    printf '%s\n' "$ind" > "$OUT/sn_${ind}.txt"
    bcftools reheader -s "$OUT/sn_${ind}.txt" "$OUT/${ind}_chr${chr}.vcf.gz" \
        -o "$OUT/${ind}_chr${chr}.rn.vcf.gz"
    bcftools index -f -t "$OUT/${ind}_chr${chr}.rn.vcf.gz"
  done
  # merge the three individuals for this chromosome
  bcftools merge -m all -Oz \
      -o "$OUT/merged_chr${chr}.vcf.gz" \
      "$OUT/Altai_chr${chr}.rn.vcf.gz" "$OUT/Denisova_chr${chr}.rn.vcf.gz" "$OUT/Vindija33.19_chr${chr}.rn.vcf.gz"
  bcftools index -f -t "$OUT/merged_chr${chr}.vcf.gz"
done

# concatenate all chromosomes (natural-sort so contigs are in karyotypic, not lexical, order)
bcftools concat -Oz -o "$OUT/partner_archaics.vcf.gz" $(ls "$OUT"/merged_chr*.vcf.gz | sort -V)
bcftools index -f -t "$OUT/partner_archaics.vcf.gz"

# annotate coding consequences (canonical transcripts; keep protein position + AA).
# --no_stats avoids the summary html; --fields ORDER must stay in sync with 03's CSQ parser.
vep --offline --cache --dir_cache "$VEP_CACHE" --assembly GRCh37 --fasta "$FASTA_GRCH37" \
    --vcf --force_overwrite --canonical --protein --symbol --no_stats \
    --fields "Consequence,SYMBOL,Gene,Feature,Protein_position,Amino_acids,CANONICAL" \
    -i "$OUT/partner_archaics.vcf.gz" -o "$OUT/partner_archaics.vep.vcf"
bgzip -f "$OUT/partner_archaics.vep.vcf"; tabix -f -p vcf "$OUT/partner_archaics.vep.vcf.gz"
echo "DONE -> $OUT/partner_archaics.vep.vcf.gz"
```
