## Supplementary Information section 2 code base for "Hypothesis: A modern human range expansion ∼300,000 years ago had a critical role in Neandertal origins": 03_classify.py

```
#!/usr/bin/env python3
"""03 — Classify the lineage pattern of every coding variant in the nuclear
partner genes, polarized by the chimp-ancestral allele (CAnc), and flag those
that fall on a residue contacting one of the lineage-specific mt sites.

The point of this scan (vs the published catalogs): it directly resolves the
Denisova-vs-Neandertal split, so it recovers **Neandertal-only** nuclear coding
changes (Neandertal-derived, modern + Denisova ancestral) — a phylogenetically
discordant pattern that the Prufer/Meyer/Kuhlwilm catalogs exclude by design.

Input : o2/results/nuclear_scan/partner_archaics.vep.vcf.gz  (from 02)
        o2/inputs/contact_pairs_all.tsv                      (nuclear contact residues)
Output: o2/results/nuclear_scan/partner_variant_classification.tsv
        + a console summary of any coding change present in modern/Neandertal &
          absent in Denisova that sits on a contact residue (the co-evolution hit).
Requires: pysam.
"""
import os, sys, csv, re
import pysam

ROOT = os.path.dirname(os.path.dirname(os.path.dirname(os.path.abspath(__file__))))
VCF = os.path.join(ROOT, "o2", "results", "nuclear_scan", "partner_archaics.vep.vcf.gz")
PAIRS = os.path.join(ROOT, "o2", "inputs", "contact_pairs_all.tsv")
OUT = os.path.join(ROOT, "o2", "results", "nuclear_scan", "partner_variant_classification.tsv")

A3to1 = {'ALA':'A','ARG':'R','ASN':'N','ASP':'D','CYS':'C','GLN':'Q','GLU':'E','GLY':'G','HIS':'H',
 'ILE':'I','LEU':'L','LYS':'K','MET':'M','PHE':'F','PRO':'P','SER':'S','THR':'T','TRP':'W','TYR':'Y','VAL':'V'}
# contact residues per nuclear gene -> set of protein positions (Ensembl precursor numbering),
# with the mt site(s) they touch and the expected residue one-letter (for the assertion).
contact = {}        # gene -> {protein_pos: [(mt_gene,mt_res,mt_lineage,dist,interface)]}
contact_aa = {}     # (gene, protein_pos) -> expected one-letter residue (from nuc_resname)
with open(PAIRS) as fh:
    for r in csv.DictReader(fh, delimiter="\t"):
        pos = int(r["nuc_res"])
        contact.setdefault(r["nuc_gene"], {}).setdefault(pos, []).append(
            (r["mt_gene"], r["mt_res"], r["lineage"], r["min_dist_A"], r["interface_4A"]))
        contact_aa[(r["nuc_gene"], pos)] = A3to1.get(r["nuc_resname"], "?")

def allele_of(gt_tuple, ref, alts):
    """Return the single ALT-or-REF base for a homozygous call, '.' if missing, 'HET' if het."""
    a = [x for x in gt_tuple if x is not None]
    if not a: return None
    if len(set(a)) > 1: return "HET"
    idx = a[0]
    return ref if idx == 0 else alts[idx - 1]

def classify(modern, nean, deni, anc):
    """Lineage pattern from per-lineage alleles vs ancestral (anc). All alleles and anc are
    upper-cased single bases. nean is the Altai/Vindija consensus; DISC handled by caller."""
    if None in (modern, deni, anc) or nean is None: return "missing"
    if "HET" in (modern, nean, deni): return "het(review)"
    der = lambda x: x.upper() != anc
    M, N, D = der(modern), der(nean), der(deni)
    if not (M or N or D): return "all-ancestral"
    if M and not N and not D: return "Modern-only"
    if N and not M and not D: return "Neandertal-only"
    if D and not M and not N: return "Denisova-only"
    if N and D and not M:     return "shared-archaic(Nean+Den)"
    if M and N and not D:     return "shared-Modern+Neandertal"   # Denisova-ancestral
    if M and D and not N:     return "Modern+Denisova"
    return "complex"

def main():
    if not os.path.exists(VCF):
        sys.exit(f"missing {VCF} — run 02 first")
    vf = pysam.VariantFile(VCF)
    samples = list(vf.header.samples)
    def s(rec, name):  # genotype tuple for a sample if present
        return rec.samples[name]["GT"] if name in rec.samples else (None,)
    rows = []
    for rec in vf:
        ref = rec.ref; alts = rec.alts or ()
        anc = rec.info.get("CAnc")
        anc = anc[0] if isinstance(anc, tuple) else anc
        anc = (anc or "").upper()                     # CAnc case-normalized (review fix M1)
        anc = anc if anc in ("A", "C", "G", "T") else None   # '.', 'N', multi-char -> unpolarizable
        modern = ref                                  # hg19 reference = modern baseline
        deni = allele_of(s(rec, "Denisova"), ref, alts)
        altai = allele_of(s(rec, "Altai"), ref, alts)
        vind = allele_of(s(rec, "Vindija33.19"), ref, alts)
        nean = altai if altai == vind else ("DISC" if (altai and vind) else (altai or vind))
        # parse VEP CSQ for coding consequence on canonical transcript
        csq = rec.info.get("CSQ")
        aa = ppos = sym = cons = ""
        if csq:
            for ent in csq:
                f = ent.split("|")
                # fields order set in 02: Consequence,SYMBOL,Gene,Feature,Protein_position,Amino_acids,CANONICAL
                if len(f) >= 7 and f[6] == "YES" and "missense" in f[0]:
                    cons, sym, ppos, aa = f[0], f[1], f[4], f[5]; break
        if "missense" not in cons:    # only coding-changing sites
            continue
        # keep Altai!=Vindija sites as a distinct category instead of silently dropping (review fix m4)
        pat = "Neandertal-DISC" if nean == "DISC" else classify(modern, nean, deni, anc)
        # contact flag (+ residue-name assertion: VEP wild-type AA must match the contact table)
        hit = ""
        try: pp = int(ppos.split("-")[0])
        except: pp = None
        if sym in contact and pp in contact.get(sym, {}):
            mt = contact[sym][pp]
            vep_wt = (aa.split("/")[0] if aa else "?")          # e.g. 'H/Q' -> 'H'
            expect = contact_aa.get((sym, pp), "?")
            mark = "" if vep_wt == expect else f"!NUMBERING-MISMATCH(VEP={vep_wt},table={expect})!"
            hit = mark + ";".join(f"{m[0]}{m[1]}({m[2]},{m[3]}A{'*' if m[4]=='True' else ''})" for m in mt)
        rows.append(dict(chrom=rec.chrom, pos=rec.pos, ref=ref, alt=",".join(alts), anc=anc,
                         gene=sym, prot_pos=ppos, aa=aa, modern=modern, altai=altai, vind=vind,
                         deni=deni, lineage_pattern=pat, contact_mt_site=hit))
    with open(OUT, "w") as fh:
        w = csv.DictWriter(fh, fieldnames=list(rows[0].keys()) if rows else
            ["chrom","pos","ref","alt","anc","gene","prot_pos","aa","modern","altai","vind","deni","lineage_pattern","contact_mt_site"],
            delimiter="\t")
        w.writeheader(); [w.writerow(r) for r in rows]
    print(f"wrote {OUT}: {len(rows)} missense variants in partner genes")
    # headline: present in modern/Neandertal, absent in Denisova, ON a contact residue
    tgt = {"Modern-only", "Neandertal-only", "shared-Modern+Neandertal"}
    hits = [r for r in rows if r["lineage_pattern"] in tgt and r["contact_mt_site"]]
    print(f"\nCO-EVOLUTION CANDIDATES (modern/Nean-derived, Denisova-ancestral, on a contact residue): {len(hits)}")
    for r in hits:
        print(f"  {r['gene']} {r['aa']}@{r['prot_pos']} [{r['lineage_pattern']}] contacts {r['contact_mt_site']}")
    # SYMMETRIC co-derived DM candidates (your hypothesis): a nuclear change that is shared-derived in
    # modern+Neandertal and ancestral in Denisova, sitting on a residue that contacts a Shared-M+N mt
    # change (so mt AND nuclear both arose on the modern+Neandertal stem; Denisovan-mt + M+N-nuclear is
    # the untested, potentially incompatible combination).
    sym = [r for r in rows if r["lineage_pattern"] == "shared-Modern+Neandertal"
           and "SHARED_MN" in (r["contact_mt_site"] or "")]
    print(f"\nSYMMETRIC SHARED-M+N CO-DERIVED PAIRS (nuclear M+N-derived/Den-ancestral at a Shared-M+N mt contact): {len(sym)}")
    for r in sym:
        print(f"  {r['gene']} {r['aa']}@{r['prot_pos']} [{r['lineage_pattern']}] contacts {r['contact_mt_site']}")
    # also report every nuclear shared-M+N coding change in a partner gene (contact or not), for context
    smn = [r for r in rows if r["lineage_pattern"] == "shared-Modern+Neandertal"]
    print(f"\nAll nuclear shared-Modern+Neandertal missense in partner genes: {len(smn)}")
    for r in smn[:40]:
        print(f"  {r['gene']} {r['aa']}@{r['prot_pos']}{' [CONTACT '+r['contact_mt_site']+']' if r['contact_mt_site'] else ''}")
    from collections import Counter
    print("\nlineage-pattern counts (all partner missense):", dict(Counter(r["lineage_pattern"] for r in rows)))

if __name__ == "__main__":
    main()
```
