## Supplementary Information section 2 code base for "Hypothesis: A modern human range expansion ∼300,000 years ago had a critical role in Neandertal origins": partner_archaic_alleles.py

```
#!/usr/bin/env python3
"""Genotype the NUCLEAR OXPHOS-partner contact residues across the high-coverage
archaic genomes, to fill column (b) of the extended Table 1: is any archaic
(Denisova 3, Denisova 25, Altai, Vindija, Chagyrskaya) DERIVED (i.e. non-reference,
= different from the modern-human hg19 residue) at a partner residue that contacts
a Denisovan mtDNA change?

Method: for each (partner protein, residue) map the residue to hg19 coordinates via
the Ensembl GRCh37 REST server, then tabix each archaic's local VCF and read the
genotype. Reference (hg19) = modern-human allele. GT 0/0 => ancestral/human-matching
(no derived allele); 1/1 or 0/1 => carries an alternate (report REF>ALT).

Run on O2 (login node has internet for the Ensembl mapping):
    export REPO=$HOME/dmi; conda activate dmi-o2; cd $REPO
    python3 o2/nuclear_scan/partner_archaic_alleles.py > o2/results/nuclear_scan/partner_archaic_alleles.tsv
Then rsync the TSV back to the Mac (run the rsync ON THE MAC):
    rsync -av:dmi/o2/results/nuclear_scan/partner_archaic_alleles.tsv ./
"""
import os, sys, json, time, re, urllib.request, warnings
warnings.filterwarnings("ignore")
import pysam

B = "https://grch37.rest.ensembl.org"

# ---- partner residues that contact the Denisovan-derived mt changes (ENSP, residue) ----
# (interface <=4 A: CYC1/COX7B/COX4I1/ATP5MF; shell <=8 A: NDUFB8/COX6C)
PARTNERS = [
    ("CYC1",   "ENSP00000317159", 199, "CYTB T70A (interface 3.0 A)"),
    ("COX7B",  "ENSP00000417656",  53, "COX1 V456A (interface 3.7 A)"),
    ("COX4I1", "ENSP00000457513", 118, "COX1 V456A (interface 3.8 A)"),
    ("ATP5MF", "ENSP00000292475",  84, "ATP6 A7T (interface 3.8 A)"),
    ("NDUFB8", "ENSP00000299166", 154, "ND5 T398A (shell 7.8 A)"),
    ("COX6C",  "ENSP00000428895",  37, "COX2 A27T (shell 6.4 A)"),
]

# ---- archaic genomes: name -> directory holding the VCF(s) ----
# CONFIRM these paths on your O2 filesystem. Altai + Denisova(Meyer=Den3) are the
# known-good local files used before; fill in Vindija, Chagyrskaya, Denisova 25.
DATA = "/n/groups/reich/DAVID/CURSOR_ENRICH/data"
SAMPLES = {
    "Denisova3_Meyer": os.path.join(DATA, "archaic_den"),   # den_meyer_autosomes.vcf.gz (= Denisova 3, high-cov)
    "AltaiNeandertal": os.path.join(DATA, "archaic_nea"),    # nea_altai_autosomes / chrN_mq25_mapab100.vcf.gz
    # ---- TODO: confirm/point these at the right local VCF directories on O2 ----
    "Vindija33.19":    os.environ.get("VINDIJA_DIR", ""),    # e.g. .../archaic_vindija
    "Chagyrskaya8":    os.environ.get("CHAGYRSKAYA_DIR", ""),
    "Denisova25":      os.environ.get("DEN25_DIR", ""),      # the Den25 nuclear VCF dir (2025 release)
}

def find_vcf(d, chrom):
    if not d or not os.path.isdir(d): return None
    files = [f for f in sorted(os.listdir(d)) if f.endswith(".vcf.gz")]
    if str(chrom).isdigit():
        for f in files:
            if "meyer" in f.lower() or "autosome" in f.lower(): return os.path.join(d, f)
    for f in files:
        if re.search(rf'(^|[._])(chr)?{chrom}[._]', f, re.I): return os.path.join(d, f)
    return None

def fetch(tb, chrom, start, end):
    for c in (str(chrom), "chr" + str(chrom)):
        try:
            return list(tb.fetch(c, start, end))
        except ValueError:
            continue
    return []

def get(u, tries=3):
    for i in range(tries):
        try:
            return json.loads(urllib.request.urlopen(urllib.request.Request(u, headers={"Content-Type":"application/json"}), timeout=40).read())
        except Exception:
            if i == tries - 1: raise
            time.sleep(1)

def call(d, chrom, s, e):
    """Return a compact per-sample string over the codon span [s,e] (hg19)."""
    vf = find_vcf(d, chrom)
    if not vf: return "NOFILE"
    try:
        tb = pysam.TabixFile(vf)
        outs = []
        for pos in range(s, e + 1):
            for row in fetch(tb, chrom, pos - 1, pos):
                f = row.split("\t")
                if len(f) < 10: continue
                gt = f[9].split(":")[0]
                if gt in ("0/0", "0|0", "."): continue
                outs.append(f"{f[1]}:{f[3]}>{f[4]}({gt})")
        tb.close()
        return "; ".join(outs) if outs else "0/0 (ref=modern)"
    except Exception as ex:
        return f"ERR:{str(ex)[:30]}"

def main():
    names = list(SAMPLES)
    print("partner_gene\tresidue\tcontacts_mt\thg19\t" + "\t".join(names))
    for gene, ensp, res, mtc in PARTNERS:
        try:
            m = get(f"{B}/map/translation/{ensp}/{res}-{res}?content-type=application/json")["mappings"][0]
            chrom, s, e = str(m["seq_region_name"]), m["start"], m["end"]
        except Exception as ex:
            print(f"{gene}\t{res}\t{mtc}\tMAP-ERR:{str(ex)[:30]}", file=sys.stderr); continue
        cells = [call(SAMPLES[n], chrom, s, e) for n in names]
        print(f"{gene}\t{res}\t{mtc}\t{chrom}:{s}-{e}\t" + "\t".join(cells))

if __name__ == "__main__":
    main()
```
