## Supplementary Information section 2 code base for "Hypothesis: A modern human range expansion ∼300,000 years ago had a critical role in Neandertal origins": README.md

# O2 pipelines — the two cluster arms of the Denisovan mito-nuclear DMI test

Two independent pipelines, each runnable on O2 (SLURM). They close the two gaps left by the
laptop analysis (see ../SUMMARY.md and ../results/):

1. **`nuclear_scan/`** — direct Denisova-vs-Neandertal-vs-modern genotyping of the nuclear partner
   genes, to recover **Neandertal-only** nuclear coding changes (the phylogenetically discordant
   pattern the Prüfer/Meyer/Kuhlwilm catalogs exclude by construction). Remote tabix of the Altai/
   Vindija VCFs failed on the laptop (htslib HTTP large-file BGZF bug); O2 reads them locally/over a
   working htslib.
2. **`den25/`** — reconstruct the Denisova 25 (~200 ka) mitochondrial genome from the EVA raw
   per-library BAMs (no server-side index) and genotype the Denisovan-derived mt sites, giving an
   independent temporal check outside the four-genome fixation panel.

## Setup
```
conda env create -f environment.yml && conda activate dmi-o2
vep_install -a cf -s homo_sapiens -y GRCh37 -c $HOME/.vep        # VEP cache (once)
export VEP_CACHE=$HOME/.vep  FASTA_GRCH37=$HOME/ref/human_g1k_v37.fasta
mkdir -p logs                          # REQUIRED: SLURM opens -o logs/... at launch
# run ALL `sbatch` calls from the repo root (the scripts cd there; -o is relative to submit dir)
# edit the `module load` lines in the .sh scripts to match O2, or rely on the conda env
```
**Before launching pipeline 1, verify two things on a real VCF (one-time, decisive):**
```
# (a) CAnc allele case — 03 polarizes on it; lowercase would garble classification (now upper-cased, verify anyway)
bcftools view -H http://ftp.eva.mpg.de/neandertal/Vindija/VCF/Altai/chr21_mq25_mapab100.vcf.gz 21:1-200000 | head
# (b) confirm `bcftools view -R <regions> <http-url>` actually streams on O2 before the full array (see M3 below)
```
Inputs are pre-generated in `inputs/` (no internet needed for them):
`partner_genes_all.tsv`, `partner_cds_grch37.bed`, `partner_gene_span_grch37.tsv`,
`contact_pairs_all.tsv` (193 mt↔nuclear residue pairs, 26 at ≤4 Å, 31 partner genes).

## Pipeline 1 — nuclear scan
```
python3 nuclear_scan/01_partner_regions.py        # (already run; regenerates the BED, needs internet)
sbatch  nuclear_scan/02_slice_merge_annotate.sh   # slice Altai/Denisova/Vindija + bcftools merge + VEP
python3 nuclear_scan/03_classify.py               # lineage classification + contact-residue flags
```
Output: `results/nuclear_scan/partner_variant_classification.tsv` and a console list of
**co-evolution candidates** = coding changes present in modern/Neandertal & absent in Denisova that
sit on a residue contacting a lineage-specific mt change. (Expectation, given the laptop result:
few/none — OXPHOS partners are deeply conserved — but this is the one pattern not yet excluded.)

## Pipeline 2 — Denisova 25 mtDNA
```
bash    den25/extract_den25_mt.sh                 # stream EVA raw BAMs, keep MT reads, merge/sort
python3 den25/consensus_from_bam.py <bam> <out.fa> # reference-free MT consensus (pysam pileup)
python3 den25/genotype_den25_mt.py <out.fa>       # amino acid at each Denisovan-derived mt site
```
Output: `results/den25/den25_mt_sites.tsv` — Denisova 25's state at the Denisovan-derived sites
(382× consensus, rCRS coords 1–16,569).

## Method notes / caveats (read before interpreting)
- mt genes use the **vertebrate-mitochondrial code** (transl_table 2); nuclear genes use the
  standard code.
- **Numbering**: contact residues are in human UniProt/structure numbering; `03` matches on VEP
  Protein_position (canonical transcript). Spot-check a couple of partners (e.g. CYC1, NDUFS7) that
  the residue identities line up before trusting hits.
- The Prüfer high-coverage VCFs are **all-sites** (explicit 0/0), so most CDS positions carry calls for
  all three archaics. `03` treats a genuinely absent / `./.` genotype after merge as **missing** (the
  site is dropped, not assumed reference) and reports Altai≠Vindija sites in a distinct `Neandertal-DISC`
  category. `02` keeps INFO **CAnc** (chimp ancestral) for polarization (`03` upper-cases it). "Modern"
  = hg19 reference; add a 1000G/gnomAD frequency join if you need modern-human allele frequencies.
- **Numbering is normalized at the source**: `contact_pairs_all.tsv` `nuc_res` is Ensembl-precursor
  numbering (auto-offset-corrected against UniProt; `nuc_res_struct` keeps the original structure number),
  and `03` asserts the residue at the mapped position equals `nuc_resname`, flagging
  `!NUMBERING-MISMATCH!` rather than silently mis-scoring.
