## Supplementary Information section 2 code base for "Hypothesis: A modern human range expansion ∼300,000 years ago had a critical role in Neandertal origins": panel.py

```
"""Curated mtDNA panel for the Denisovan mito-nuclear DMI analysis."""

# Denisovan individuals (the well-characterized ones). Variant must be present
# in ALL of these to be called "fixed in Denisovan mtDNA".
DENISOVAN = {
    "Denisova3_phalanx": "FN673705.1",   # Krause 2010 (== NC_013993)
    "Denisova4_molar":   "FR695060.1",   # Reich 2010 / Sawyer 2015
    "Denisova2":         "KX663333.1",   # Slon 2017
    "Denisova8":         "KT780370.1",   # Sawyer 2015
}

# Neandertal panel spanning the known mtDNA diversity (early -> late, E + W).
NEANDERTAL = {
    "Neander_RefSeq_Vi33.16": "NC_011137.1",  # Green 2008 (first complete)
    "Altai_KC879692":         "KC879692.1",
    "Chagyrskaya8":           "MK388903.1",
    "Mezmaiskaya1":           "FM865411.1",
    "Mezmaiskaya2":           "MG025537.1",
    "ElSidron1253":           "FM865409.1",
    "Feldhofer1":             "FM865407.1",
    "Vindija33.19":           "KJ533545.1",
    "LesCottes":              "MG025536.1",
    "Scladina_I4A":           "MK123269.1",  # ~127 ka, basal
    "Stajnia_S5000":          "MT795654.1",
}

# Modern human reference. (Polymorphism check vs MITOMAP/HmtVar done separately.)
MODERN_HUMAN = {
    "rCRS_hg_H2a2a1": "NC_012920.1",
}

# Outgroups for polarizing derived vs ancestral state.
OUTGROUP = {
    "Pan_troglodytes": "NC_001643.1",   # chimpanzee
    "Pan_paniscus":    "NC_001644.1",   # bonobo
    "Gorilla":         "NC_011120.1",   # gorilla
}

ALL = {}
for grp in (DENISOVAN, NEANDERTAL, MODERN_HUMAN, OUTGROUP):
    ALL.update(grp)

GENES = ["ND1","ND2","COX1","COX2","ATP8","ATP6","COX3","ND3","ND4L","ND4","ND5","ND6","CYTB"]
```
