## Supplementary Information section 2 code base for "Hypothesis: A modern human range expansion ∼300,000 years ago had a critical role in Neandertal origins": prep_o2_inputs.py

```
#!/usr/bin/env python3
"""Generate the master mt-residue <-> nuclear-residue contact-pair table across
ALL lineage-specific mt sites (Denisovan/Neandertal/Modern/Shared). This anchors
the O2 nuclear scan, which flags partner residues at these contacts."""
import os, sys, json, warnings
sys.path.insert(0, os.path.dirname(os.path.abspath(__file__)))
warnings.filterwarnings("ignore")
import contacts as C
from Bio.PDB.MMCIF2Dict import MMCIF2Dict

ROOT = os.path.dirname(os.path.dirname(os.path.abspath(__file__)))

# ---- the 43 lineage-specific mt nonsynonymous sites (validated sets) ----
SITES = {
 "DENISOVAN": [("ND2",64),("ND2",164),("ND5",398),("CYTB",70),("COX1",456),("COX2",27),("ATP6",7),("ATP6",185)],
 "NEANDERTAL":[("ND1",10),("ND1",168),("ND2",346),("ND5",489),("ND5",603),("ND6",166),("CYTB",245),("CYTB",309),("COX3",84),("ATP6",80),("ATP6",154),("ATP6",176)],
 "MODERN":    [("ND1",248),("COX2",22),("COX2",95),("COX2",146),("ATP6",112),("COX3",40),("ND3",114),("ND5",24),("ND5",257),("ND5",518),("ND5",573),("CYTB",7),("CYTB",233)],
 "SHARED_MN": [("ND1",39),("ND1",64),("COX1",155),("COX1",415),("ATP6",51),("ND4",86),("ND5",4),("ND5",531),("CYTB",4),("CYTB",122)],
}
STRUCT = {"ND1":"5XTD","ND2":"5XTD","ND3":"5XTD","ND4":"5XTD","ND4L":"5XTD","ND5":"5XTD","ND6":"5XTD",
          "CYTB":"9HZL","COX1":"5Z62","COX2":"5Z62","COX3":"5Z62","ATP6":"8H9S","ATP8":"8H9S"}
COMPLEX = {"ND1":"CI","ND2":"CI","ND3":"CI","ND4":"CI","ND4L":"CI","ND5":"CI","ND6":"CI",
           "CYTB":"CIII","COX1":"CIV","COX2":"CIV","COX3":"CIV","ATP6":"CV","ATP8":"CV"}
# UniProt -> (symbol, Ensembl)  (merged from Task2 + Task4)
UNI = json.load(open(os.path.join(ROOT,"data","task4_partner_uniprot_map.json")))
UNI.update({  # ensure originals present
 "O95169":("NDUFB8","ENSG00000166136"),"P08574":("CYC1","ENSG00000179091"),
 "P24311":("COX7B","ENSG00000131174"),"P13073":("COX4I1","ENSG00000131143"),
 "P09669":("COX6C","ENSG00000164919"),"P56134":("ATP5MF","ENSG00000241468"),
 "P24539":("ATP5PB","ENSG00000116459"),"Q9P0J0":("NDUFA13","ENSG00000186010"),
 "O15239":("NDUFA1","ENSG00000125356"),"O95298":("NDUFC2","ENSG00000087258"),
 "Q86Y39":("NDUFA11","ENSG00000174886"),"P05496":("ATP5MC1","ENSG00000159199"),
 "Q9UDW1":("UQCRH","ENSG00000173660")})

import urllib.request
A3to1={'ALA':'A','ARG':'R','ASN':'N','ASP':'D','CYS':'C','GLN':'Q','GLU':'E','GLY':'G','HIS':'H',
 'ILE':'I','LEU':'L','LYS':'K','MET':'M','PHE':'F','PRO':'P','SER':'S','THR':'T','TRP':'W','TYR':'Y','VAL':'V','MSE':'M'}
_useq={}
def uniprot_seq(acc):
    if acc not in _useq:
        try:
            t=urllib.request.urlopen(f"https://rest.uniprot.org/uniprotkb/{acc}.fasta",timeout=30).read().decode()
            _useq[acc]="".join(l for l in t.splitlines() if not l.startswith(">"))
        except Exception: _useq[acc]=None
    return _useq[acc]

def detect_offset(acc, pairs):
    """Find integer offset so that structure nuc_res -> UniProt precursor numbering
    (nuc_res+offset) maximizes residue-name agreement. pairs = [(res, aa1)]."""
    s=uniprot_seq(acc)
    if not s or not pairs: return 0, None
    best=(0,-1)
    for off in range(0,101):
        ok=sum(1 for res,aa in pairs if 1<=res+off<=len(s) and s[res+off-1]==aa)
        if ok>best[1]: best=(off,ok)
    return best[0], (best[1], len(pairs))

def chain_acc(sid):
    d=MMCIF2Dict(os.path.join(ROOT,"data","pdb",f"{sid}.cif"))
    al=lambda k:(d.get(k) if isinstance(d.get(k),list) else ([d.get(k)] if d.get(k) is not None else []))
    ch=al('_atom_site.auth_asym_id');ent=al('_atom_site.label_entity_id');c2e={}
    for c,e in zip(ch,ent):c2e.setdefault(c,e)
    e2u={}
    for e,a,db in zip(al('_struct_ref.entity_id'),al('_struct_ref.pdbx_db_accession'),al('_struct_ref.db_name')):
        if db=='UNP':e2u[e]=a
    return {c:e2u.get(e,'?') for c,e in c2e.items()}

rows=[]; partner_uni=set()
acc_cache={}
for lineage, sites in SITES.items():
    bystruct={}
    for g,r in sites: bystruct.setdefault(STRUCT[g],[]).append((g,r,lineage))
    for sid,tg in bystruct.items():
        if sid not in acc_cache: acc_cache[sid]=chain_acc(sid)
        for res in C.analyze(sid,tg):
            if res.get("error"):
                rows.append({"lineage":lineage,"mt_gene":res["gene"],"mt_res":res["res"],"struct":sid,"note":res["error"]}); continue
            for k,v in res["partners"].items():
                pcid,kind,dsc=k.split("|",2)
                if kind!="nuc": continue
                a=acc_cache[sid].get(pcid,"?"); sym,ens=UNI.get(a,(a,"?"))
                partner_uni.add(a)
                for pres,pnm,d in sorted(v,key=lambda x:x[2]):
                    rows.append({"lineage":lineage,"mt_gene":res["gene"],"mt_res":res["res"],"complex":COMPLEX[res["gene"]],
                                 "struct":sid,"nuc_gene":sym,"nuc_uniprot":a,"nuc_ens":ens,
                                 "nuc_res":pres,"nuc_resname":pnm,"min_dist_A":round(d,2),"interface_<=4A":d<=4.0})

# ---- normalize nuclear residue numbering to Ensembl/UniProt PRECURSOR (review fix B4) ----
from collections import defaultdict
bygene=defaultdict(list)
for r in rows:
    if r.get("note"): continue
    bygene[r["nuc_uniprot"]].append((r["nuc_res"], A3to1.get(r["nuc_resname"],"?")))
offsets={}
print("nuclear residue offset correction (structure -> UniProt precursor):")
for acc,prs in bygene.items():
    off,stat=detect_offset(acc, sorted(set(prs)))
    offsets[acc]=off
    gsym=UNI.get(acc,(acc,'?'))[0]
    if stat: print(f"  {gsym} ({acc}): offset +{off}  resname match {stat[0]}/{stat[1]}")
for r in rows:
    if r.get("note"): continue
    r["nuc_res_struct"]=r["nuc_res"]
    r["nuc_res"]=r["nuc_res"]+offsets.get(r["nuc_uniprot"],0)   # now precursor numbering

json.dump(rows, open(os.path.join(ROOT,"results","contact_pairs_all.json"),"w"), indent=1)
# TSV
import csv
with open(os.path.join(ROOT,"results","contact_pairs_all.tsv"),"w") as fh:
    w=csv.writer(fh,delimiter="\t",lineterminator="\n")
    w.writerow(["lineage","mt_gene","mt_res","complex","struct","nuc_gene","nuc_uniprot","nuc_ens","nuc_res","nuc_res_struct","nuc_resname","min_dist_A","interface_4A"])
    for r in rows:
        if r.get("note"): continue
        w.writerow([r["lineage"],r["mt_gene"],r["mt_res"],r.get("complex",""),r["struct"],r["nuc_gene"],r["nuc_uniprot"],r["nuc_ens"],r["nuc_res"],r.get("nuc_res_struct",r["nuc_res"]),r["nuc_resname"],r["min_dist_A"],r["interface_<=4A"]])
# partner gene list
genes={}
for r in rows:
    if r.get("note"): continue
    genes.setdefault(r["nuc_gene"],(r["nuc_uniprot"],r["nuc_ens"]))
with open(os.path.join(ROOT,"results","partner_genes_all.tsv"),"w") as fh:
    fh.write("gene\tuniprot\tensembl\n")
    for g,(u,e) in sorted(genes.items()): fh.write(f"{g}\t{u}\t{e}\n")
# summary
npair=sum(1 for r in rows if not r.get("note"))
niface=sum(1 for r in rows if not r.get("note") and r["interface_<=4A"])
print(f"contact residue-pairs: {npair} ({niface} at <=4A interface) across {len(genes)} nuclear partner genes")
print("partner genes:", ", ".join(sorted(genes)))
```
