## Supplementary Information section 2 code base for "Hypothesis: A modern human range expansion ∼300,000 years ago had a critical role in Neandertal origins": README.md

# Denisovan mtDNA–nuclear incompatibility test — code base

All code needed to replicate the results in the supplementary note *"No evidence for a
DMI between Denisovan mtDNA and modern human nuclear DNA."* Python 3.11. The analysis
has two parts: a **laptop pipeline** (this folder) that produces the structural,
catalog and intrinsic-impact results, and two **O2 (SLURM) pipelines**
(`o2/nuclear_scan/`, `o2/den25/`) that close the remaining gaps.

## Setup
```
pip install -r requirements.txt          # biopython, numpy, pysam, pypdf
# structures (mmCIF/PDB), mtDNA GenBank records, and the Prüfer/Meyer/Kuhlwilm catalogs
# are fetched by the scripts below or read from the paper supplements.
```

## Laptop pipeline — script by script
Run from this folder; each writes to a `results/` tree it creates.

- **`ncbi.py` / `panel.py` / `fetch_panel.py`** — fetch the mtDNA sequences (rCRS
  NC_012920, Denisova 3 FN673705, Vindija 33.19 KJ533545, and the great-ape outgroups) and
  assemble the 13-gene OXPHOS panel with the nuclear-encoded subunit list.
- **`analyze.py`** — call coding changes per gene from the NCBI CDS translations, align to
  rCRS, and polarize by great-ape majority under the vertebrate-mitochondrial code
  (`transl_table 2`). Applies the multi-individual fixation filter (4 Denisovan + 11
  Neandertal mtDNAs) that yields the **eight** Denisovan-specific nonsynonymous sites and
  the codon-aware dN/dS lineage partition.
- **`analyze_per_individual.py`** — extend the same alignment/polarization to every change derived in *any* single Denisovan (not only the fixed set); emits the per-individual Den2/Den3/Den4/Den8 genotypes and the fixed-vs-non-fixed classification tabulated in **Table S2.1** (eight fixed + seven non-fixed; two private premature-stop calls in Denisova 2 are flagged and excluded).
- **`contacts.py`** — compute inter-chain heavy-atom contacts from the human cryo-EM
  structures (5XTD, 9HZL, 5Z62, 8H9S, 5XTH); ≤4 Å = interface, ≤8 Å = shell. Reconciles
  residue numbering to UniProt via SIFTS and verifies by residue identity.
- **`lineage_table.py`** — join the mt changes to their nuclear structural partners and
  build the contact table (`contact_pairs_all.tsv`; 193 mt↔nuclear pairs, 26 at ≤4 Å).
- **`task3_workflow.js`** — the five convergent allele-level tests: catalog closure
  (Prüfer 2014 / Meyer 2012 / Kuhlwilm & Boeckx 2019), gnomAD v4 standing variation at the
  contact residues, the Denisova-specific contrast, intrinsic impact (MitImpact/APOGEE2),
  and mammalian conservation of the mtDNA variant residues.
- **`task4_coevolution.py`** — assemble the matched **Neandertal-lineage control set**:
  for the mt changes derived on the modern and shared modern+Neandertal lineages, find the
  nuclear subunits contacting them and emit the union of control-set partner genes
  (`results/task4_partners.json`, `results/task4_partner_genes.json`) for the catalog check.
- **`prep_o2_inputs.py`** — write the pre-generated inputs consumed by the O2 pipelines
  (`o2/inputs/`).
- **`extract_pdfs.py`** — pull the tables used from the paper supplements (e.g. the Krause
  2010 substitution counts).

## O2 pipelines — see `o2/README.md`
1. **`o2/nuclear_scan/`** — direct Denisova-vs-Neandertal-vs-modern genotyping of the
   nuclear partner genes, to recover any Neandertal-only nuclear coding change (the
   phylogenetically discordant pattern the fixed-difference catalogs exclude by
   construction). Uses local htslib to read the Altai/Vindija VCFs (remote tabix of the
   large BGZF files was unreliable on the laptop).
2. **`o2/den25/`** — reconstruct the Denisova 25 (~200 ka) mitochondrial genome from the
   EVA raw per-library BAMs and genotype the Denisovan-derived mt sites (see the Addendum).
   `o2/environment.yml` builds the conda environment (bcftools, VEP, samtools, pysam).

## Notes
- mt genes use the vertebrate-mitochondrial code (`transl_table 2`); nuclear genes use the
  standard code.
- Contact residues are in human UniProt/structure numbering; the allele scans assert the
  residue at each mapped position equals the expected identity and flag (rather than
  silently mis-score) any numbering mismatch.
- "Not detected" is bounded by the resources used — catalog sampling, gnomAD coverage,
  structure resolution, and predictor conservatism (see the note's Summary).

---
## Addendum (2026-07-31): archaic partner genotyping + Denisova 25

- `o2/nuclear_scan/partner_archaic_alleles.py` — genotypes the nuclear OXPHOS-partner
  contact residues (CYC1 199, COX7B 53, COX4I1 118, ATP5MF 84, NDUFB8 154, COX6C 37)
  in the high-coverage archaic genomes. Denisova 3 + Altai (local O2 VCFs) and
  Denisova 25 (EVA per-chromosome VCFs, read remotely — incl. COX7B on chrX) are all
  homozygous-reference / ancestral. Output: `o2/results/nuclear_scan/partner_archaic_alleles.tsv`
  (Den3/Altai) and `den25_partner_alleles.tsv` (Den25, from the EVA FTP).
- `o2/den25/extract_den25_mt.sh` — streams the EVA Denisova 25 raw per-library BAMs
  (no server-side index), keeps only MT reads, merges/sorts, and builds a consensus
  (rCRS coords 1–16569; the b37 "MT" contig is rCRS + 1000 bp wrap padding).
- `o2/den25/genotype_den25_mt.py` — reads that consensus and reports Denisova 25's
  amino acid at each of the 15 Denisovan-derived mt sites (vertebrate-mito code).
  Data source: https://cdna.eva.mpg.de/denisova/Den25/  (Den25 = 200–170 ka molar, ~24×).

- o2/den25/consensus_from_bam.py — reference-free MT consensus (pysam pileup) for O2 samtools builds lacking `samtools consensus`.
- o2/results/den25/den25_mt_sites.tsv — Den25 mtDNA genotypes at the 15 derived sites (382x).
