## Supplementary Information section 2 code base for "Hypothesis: A modern human range expansion ∼300,000 years ago had a critical role in Neandertal origins": task3_workflow.js

```
export const meta = {
  name: 'denisovan-dmi-task3',
  description: 'Task 3: validate Denisovan mt sites vs Krause2010, extract Osada2012 coevolution method/results, and determine whether nuclear OXPHOS partner proteins carry fixed modern-human-derived (or archaic) coding alleles across the Meyer2012 / Prufer2014 / Kuhlwilm2019 catalogs.',
  phases: [{ title: 'Extract' }, { title: 'Synthesize' }],
}

const ROOT = '/Users/reich/Documents/Claude_DM_incompatibilities'

const CONTEXT = `
BACKGROUND (results already computed in this project; do NOT recompute):
Hypothesis: a mito-nuclear Dobzhansky-Muller incompatibility blocked the deeply
divergent Denisovan-like mtDNA from persisting when modern-human-like populations
introgressed into Neandertal ancestors (~300-400 ka), so it was replaced by the
modern-human-like mtDNA (Posth 2017).

TASK 1 RESULT - 8 high-confidence Denisovan-specific, fixed, derived NONSYNONYMOUS
mtDNA changes (rCRS/UniProt residue numbering):
  ND2 A64T, ND2 A164T, ND5 T398A   (Complex I)
  CYTB T70A                         (Complex III)
  COX1 V456A, COX2 A27T             (Complex IV)
  ATP6 A7T, ATP6 N185S              (Complex V)
Matched NEANDERTAL-lineage control nonsynonymous changes:
  ND1 I10V, ND1 T168A, ND2 I346V, ND5 T489A, ND5 T603M, ND6 I166V (CI);
  CYTB F245L, CYTB M309V (CIII); COX3 I84V (CIV); ATP6 A80T, ATP6 M154V, ATP6 S176N (CV)

TASK 2 RESULT - nuclear-encoded subunits contacting these mt residues (human cryo-EM
structures, heavy-atom <=4 A = interface):
  DENISOVAN sites at nuclear interfaces:
    CYTB T70  -> CYC1 (cytochrome c1, UniProt P08574) residue Tyr199, 3.0 A
    COX1 V456 -> COX7B (P24311) Trp53  AND COX4I1 (P13073) Ile118, 3.7/3.8 A
    ATP6 A7   -> ATP5MF / "subunit f" (old symbol ATP5J2, P56134) His83, 3.8 A
    ND5 T398  -> NDUFB8 (O95169), proximal 7.8 A
    COX2 A27  -> COX6C (P09669), proximal 6.4 A
  NEANDERTAL-control sites at nuclear interfaces:
    ND1 T168  -> NDUFA13 / GRIM-19 (Q9P0J0) Arg57, 3.2 A
    CYTB F245 -> CYC1 (P08574), 2.2 A
    ATP6 S176 -> ATP5PB / "subunit b" (old symbol ATP5F1, P24539) Ala55, 3.3 A
    ND1 I10 -> NDUFA1/MWFE (O15239) proximal; ND2 I346 -> NDUFC2 (O95298) proximal;
    ND5 T603 -> NDUFA11 (Q86Y39) proximal; ATP6 M154 -> ATP5MC1/ATP5G1 (P05496) proximal

NUCLEAR PARTNER GENES TO CHECK IN CATALOGS (use BOTH new and old symbols):
  CYC1 ; COX7B ; COX4I1 (a.k.a. COX4) ; ATP5MF (a.k.a. ATP5J2) ;
  ATP5PB (a.k.a. ATP5F1) ; NDUFA13 (a.k.a. GRIM-19, GRIM19, CGI-39) ;
  NDUFB8 ; COX6C ; NDUFA1 (a.k.a. MWFE) ; NDUFC2 ; NDUFA11 ; ATP5MC1 (a.k.a. ATP5G1)
The KEY question of Task 3: do any of these nuclear partners carry a fixed
modern-human-derived (or archaic-specific) AMINO-ACID change? Such a change would be
the nuclear side of a candidate mito-nuclear incompatibility.
`

const CATALOG_SCHEMA = {
  type: 'object', additionalProperties: false,
  properties: {
    source: { type: 'string' },
    catalog_located: { type: 'boolean', description: 'Did you actually locate the list/table of fixed modern-human-derived (and/or archaic-derived) coding changes in this source (vs only a description of it)?' },
    where_found: { type: 'string', description: 'page numbers / table name / external URL / "list is external, not in PDF"' },
    how_genes_identified: { type: 'string', description: 'gene symbol / Ensembl ID / genomic coordinate / scanned image' },
    total_fixed_missense_reported: { type: 'string', description: 'the genome-wide count of fixed modern-human-derived missense/amino-acid changes reported, if stated' },
    partner_genes_hits: {
      type: 'array',
      items: { type: 'object', additionalProperties: false,
        properties: { gene: { type: 'string' }, change: { type: 'string' }, lineage: { type: 'string' }, context: { type: 'string' } },
        required: ['gene', 'change', 'lineage'] },
      description: 'Any of the listed NUCLEAR PARTNER genes (or their synonyms) found in the catalog of coding changes. Empty array if none.' },
    other_oxphos_or_mt_genes_hits: { type: 'array', items: { type: 'string' }, description: 'Any other OXPHOS / mitochondrially-targeted genes with fixed coding changes you noticed.' },
    coverage_caveat: { type: 'string', description: 'Could a partner gene have been present but missed (e.g. list is external/coordinate-based/image)? Be explicit.' },
    verdict: { type: 'string', description: 'Do any nuclear partner genes carry a fixed modern-human-derived (or archaic) coding change per THIS source? yes / no / cannot-determine, with one-line reason.' },
  },
  required: ['source', 'catalog_located', 'partner_genes_hits', 'verdict', 'coverage_caveat'],
}

phase('Extract')
const extracted = await parallel([
  // 1. Krause 2010 -> validate Task 1 mt sites
  () => agent(`${CONTEXT}
You are validating the 8 Denisovan-specific mtDNA nonsynonymous sites against the ORIGINAL Denisova mtDNA paper.
Files (read them):
  - ${ROOT}/Krause2010Main.pdf  (4 pp, text)
  - ${ROOT}/Krause2010Supplement.pdf  (16 pp, SCANNED IMAGES -> use Read with the pages parameter to view the tables as images; the table of amino-acid/coding differences between Denisova, human (CRS) and Neandertal mtDNA is in here)
  - extracted text: ${ROOT}/data/papertext/Krause2010Main.txt
Find Krause et al.'s reported list of amino-acid (nonsynonymous) differences on the Denisovan mtDNA relative to the human reference and Neandertal. Compare to our 8 sites. Report which of our 8 appear in their table, any we have that they don't (and vice versa), and the total number of amino-acid differences they report on the Denisovan lineage. Be precise about residue/gene.`,
    { label: 'krause-validate', phase: 'Extract', schema: {
      type: 'object', additionalProperties: false,
      properties: {
        krause_reported_aa_diffs: { type: 'array', items: { type: 'object', additionalProperties: false, properties: { gene: { type: 'string' }, change: { type: 'string' }, note: { type: 'string' } }, required: ['gene', 'change'] } },
        krause_total_aa_diff_count: { type: 'string' },
        our8_confirmed: { type: 'array', items: { type: 'string' } },
        our8_not_in_krause: { type: 'array', items: { type: 'string' } },
        krause_extra_not_in_ours: { type: 'array', items: { type: 'string' } },
        notes: { type: 'string' } },
      required: ['our8_confirmed', 'our8_not_in_krause', 'notes'] } }),

  // 2. Osada 2012 -> method + coevolution results
  () => agent(`${CONTEXT}
You are extracting the METHOD and RESULTS template from Osada & Akashi 2012 MBE "Mitochondrial-nuclear interactions and accelerated compensatory evolution: evidence from the primate cytochrome c oxidase complex". This is the canonical method for exactly our Task 2/3.
Files (read them):
  - ${ROOT}/Osada2012.pdf  (full text)
  - ${ROOT}/data/osada_si/Supplementary_Table_1.pdf
  - ${ROOT}/data/osada_si/Supplementary_Table_2.pdf
  - extracted text: ${ROOT}/data/papertext/Osada2012.txt
Extract precisely: (a) which crystal structure(s) of cytochrome c oxidase they used and the inter-atomic distance threshold defining mt<->nuclear contacts; (b) how they tested for mito-nuclear coevolution / accelerated compensatory evolution (the statistic, the phylogenetic method, the branches); (c) their KEY findings - did they find accelerated nonsynonymous/compensatory evolution in nuclear COX subunits at mt-contacting sites? which nuclear subunits (COX4I1/COX5A/COX5B/COX6A/6B/6C/COX7A/7B/7C)?; (d) any specific mt-residue <-> nuclear-residue coevolving PAIRS they list (esp. involving COX4I1, COX6C, COX7B which are our partners); (e) how directly their approach can be reused for our Denisovan COX1/COX2 sites. Quote table contents where relevant.`,
    { label: 'osada-method', phase: 'Extract', schema: {
      type: 'object', additionalProperties: false,
      properties: {
        structures_used: { type: 'string' },
        contact_distance_threshold: { type: 'string' },
        coevolution_method: { type: 'string' },
        key_findings: { type: 'string' },
        nuclear_cox_subunits_analyzed: { type: 'array', items: { type: 'string' } },
        coevolving_pairs: { type: 'array', items: { type: 'object', additionalProperties: false, properties: { mt: { type: 'string' }, nuclear: { type: 'string' }, note: { type: 'string' } }, required: ['mt', 'nuclear'] } },
        reusability_for_our_sites: { type: 'string' } },
      required: ['structures_used', 'contact_distance_threshold', 'coevolution_method', 'key_findings', 'reusability_for_our_sites'] } }),

  // 3. Meyer 2012 catalog
  () => agent(`${CONTEXT}
Search the high-coverage Denisovan genome paper (Meyer et al. 2012) for its catalog of fixed coding changes.
Files (read them):
  - ${ROOT}/Meyer2012Main.pdf
  - ${ROOT}/Meyer2012Supplement.pdf  -> "Note 19: A complete catalog of features unique to the human genome" begins around p.64 and "Note 20: Catalog of features unique to the Denisovan genome" around p.73. Use Read with the pages parameter to view those pages (they contain the list of amino-acid-changing / nonsynonymous SNCs and the affected genes). Also check the main paper's Table listing amino-acid substitutions.
  - extracted text: ${ROOT}/data/papertext/Meyer2012Supplement.txt and Meyer2012Main.txt (grep these first with Bash for the partner gene symbols and synonyms).
Determine: does Meyer 2012 list any of our NUCLEAR PARTNER genes (or synonyms) among fixed amino-acid (nonsynonymous) changes where present-day humans are derived vs Denisova? Report how genes are identified, the total count of such changes, and an explicit coverage caveat (e.g. if the genome-wide list is only available as an external file or image and the partner genes therefore can't be confirmed/excluded from the PDF alone).`,
    { label: 'meyer-catalog', phase: 'Extract', schema: CATALOG_SCHEMA }),

  // 4. Prufer 2014/2017 catalog
  () => agent(`${CONTEXT}
Search the Altai Neandertal genome paper (Prufer et al. 2014) and the Vindija paper (Prufer et al. 2017) for the list of genes carrying fixed modern-human-derived missense (amino-acid) changes vs ALL archaics.
Files (read them):
  - ${ROOT}/Prufer2014Main.pdf  (the main text / Extended Data lists the high-confidence genes with fixed missense SNCs; the famous ~96 changes / ~87 genes)
  - ${ROOT}/Prufer2014Supplement.pdf  (the "Modern-human-specific catalog" section ~p? - NOTE the full list is stated to be external at http://bioinf.eva.mpg.de/altai_catalog/ ; report this)
  - ${ROOT}/Prufer2017Main.pdf and ${ROOT}/Prufer2017Tables.xlsx (Table S52 is CNVs; check main text for the updated catalog of fixed AA changes)
  - extracted text under ${ROOT}/data/papertext/ (grep for the partner gene symbols + synonyms with Bash first).
Determine: do any of our NUCLEAR PARTNER genes appear among the fixed modern-human-derived missense changes? List the full set of genes with fixed missense changes if it's given in-PDF. Give an explicit coverage caveat (e.g. list is external/coordinate-based).`,
    { label: 'prufer-catalog', phase: 'Extract', schema: CATALOG_SCHEMA }),

  // 5. Kuhlwilm & Boeckx 2019 (fetch)
  () => agent(`${CONTEXT}
Retrieve the consolidated catalog of single-nucleotide changes distinguishing modern humans from archaic hominins: Kuhlwilm & Boeckx 2019, Scientific Reports 9:8463 (open access, DOI 10.1038/s41598-019-44877-x). Use WebSearch / WebFetch and/or the PubMed-Central full-text tools (load them via ToolSearch). Their supplementary dataset is a table of all SNCs with gene symbol + consequence (missense/synonymous) + which lineage is derived.
Goal: determine whether any of our NUCLEAR PARTNER genes (CYC1, COX7B, COX4I1, ATP5MF/ATP5J2, ATP5PB/ATP5F1, NDUFA13/GRIM-19, NDUFB8, COX6C, NDUFA1, NDUFC2, NDUFA11, ATP5MC1) carry a fixed modern-human-derived or archaic-specific MISSENSE change in their catalog. If you can download the supplementary table, save it to ${ROOT}/data/kuhlwilm2019/ and grep it. If you cannot retrieve it, say so clearly (catalog_located=false) and report what you could confirm from the abstract/text. Do NOT fabricate gene hits.`,
    { label: 'kuhlwilm-fetch', phase: 'Extract', schema: CATALOG_SCHEMA }),
])

const [krause, osada, meyer, prufer, kuhlwilm] = extracted

phase('Synthesize')
const synthesis = await agent(`${CONTEXT}

You are synthesizing Task 3. Here are the structured findings from five source-extraction agents (JSON):

KRAUSE (Task 1 validation): ${JSON.stringify(krause)}
OSADA (method + coevolution): ${JSON.stringify(osada)}
MEYER 2012 catalog: ${JSON.stringify(meyer)}
PRUFER 2014/2017 catalog: ${JSON.stringify(prufer)}
KUHLWILM 2019 catalog: ${JSON.stringify(kuhlwilm)}

Write a synthesis covering:
1. Task-1 validation verdict (do our 8 Denisovan mt sites agree with Krause 2010?).
2. The Task-3 core question: for each nuclear partner gene at a Denisovan-mt interface (CYC1, COX7B, COX4I1, ATP5MF) and each control-side partner (NDUFA13, ATP5PB, etc.), state whether any archaic-vs-modern FIXED coding allele exists per the catalogs, with the strongest available evidence and the coverage caveats. Be explicit when the answer is "no fixed coding difference found" vs "cannot be excluded from available sources."
3. What the Osada 2012 framework tells us we should do next to detect the nuclear side of the incompatibility even in the (likely) absence of a fixed modern-human amino-acid change - i.e. the primate coevolution analysis and the gnomAD standing-variation check at the contact residues.
4. A clear, honest bottom line on whether the literature/catalog evidence currently supports a protein-coding nuclear-side coevolved substitution for the Denisovan mt changes, and what compute (O2) would settle the open parts.
Keep it rigorous and concise. Flag any place a source's coverage caveat means the conclusion is provisional.`,
  { label: 'synthesize', phase: 'Synthesize' })

const verify = await agent(`Adversarially verify this Task-3 synthesis against the raw source findings. Your job is to catch (a) any claimed partner-gene coding hit that is NOT actually supported by a source, and (b) any "absence/no difference" claim that is stated more strongly than the source coverage caveats allow (false confidence). Also check the Krause validation claims.

SYNTHESIS:
${synthesis}

RAW SOURCE FINDINGS:
KRAUSE: ${JSON.stringify(krause)}
OSADA: ${JSON.stringify(osada)}
MEYER: ${JSON.stringify(meyer)}
PRUFER: ${JSON.stringify(prufer)}
KUHLWILM: ${JSON.stringify(kuhlwilm)}

Return your verdict.`,
  { label: 'verify', phase: 'Synthesize', schema: {
    type: 'object', additionalProperties: false,
    properties: {
      unsupported_claims: { type: 'array', items: { type: 'string' } },
      overconfident_absence_claims: { type: 'array', items: { type: 'string' } },
      krause_validation_ok: { type: 'string' },
      corrections: { type: 'string' },
      overall_reliability: { type: 'string' } },
    required: ['unsupported_claims', 'overconfident_absence_claims', 'overall_reliability'] } })

return { krause, osada, meyer, prufer, kuhlwilm, synthesis, verify }
```
