## Supplementary Information section 2 code base for "Hypothesis: A modern human range expansion ∼300,000 years ago had a critical role in Neandertal origins": task4_coevolution.py

```
#!/usr/bin/env python3
"""Co-evolved-pair test on the non-Denisovan lineages.
For mt nonsyn changes derived in modern/Neandertal/shared (Denisova ancestral),
find nuclear partner subunits (structural contacts) and report them for the
subsequent catalog lineage check."""
import os, sys, json, warnings
sys.path.insert(0, os.path.dirname(os.path.abspath(__file__)))
warnings.filterwarnings("ignore")
import contacts as C
from Bio.PDB.MMCIF2Dict import MMCIF2Dict

ROOT = os.path.dirname(os.path.dirname(os.path.abspath(__file__)))
lp = json.load(open(os.path.join(ROOT, "results", "lineage_partition.json")))

# route every mt gene to its structure
STRUCT = {"ND1":"5XTD","ND2":"5XTD","ND3":"5XTD","ND4":"5XTD","ND4L":"5XTD","ND5":"5XTD","ND6":"5XTD",
          "CYTB":"9HZL","COX1":"5Z62","COX2":"5Z62","COX3":"5Z62","ATP6":"8H9S","ATP8":"8H9S"}
SYM={'O95169':'NDUFB8','O15239':'NDUFA1','Q9P0J0':'NDUFA13','O95298':'NDUFC2','Q86Y39':'NDUFA11',
     'P08574':'CYC1','P24311':'COX7B','P13073':'COX4I1','P09669':'COX6C','P56134':'ATP5MF',
     'P05496':'ATP5MC1','P24539':'ATP5PB'}

def chain_acc(sid):
    d=MMCIF2Dict(os.path.join(ROOT,"data","pdb",f"{sid}.cif"))
    al=lambda k:(d.get(k) if isinstance(d.get(k),list) else ([d.get(k)] if d.get(k) is not None else []))
    ch=al('_atom_site.auth_asym_id'); ent=al('_atom_site.label_entity_id'); c2e={}
    for c,e in zip(ch,ent): c2e.setdefault(c,e)
    e2u={}
    for e,acc,db in zip(al('_struct_ref.entity_id'),al('_struct_ref.pdbx_db_accession'),al('_struct_ref.db_name')):
        if db=='UNP': e2u[e]=acc
    return {c:e2u.get(e,'?') for c,e in c2e.items()}

def run(setname, sites):
    bystruct={}
    for g,res,chg,anc in sites:
        bystruct.setdefault(STRUCT[g],[]).append((g,res,setname))
    accs={}
    results=[]
    for sid,tg in bystruct.items():
        if sid not in accs: accs[sid]=chain_acc(sid)
        for r in C.analyze(sid,tg):
            if r.get('error'):
                results.append({"site":f"{r['gene']}{r['res']}","err":r['error']}); continue
            partners=[]
            for k,v in r['partners'].items():
                pcid,kind,dsc=k.split('|',2)
                if kind!='nuc': continue
                mind=min(x[2] for x in v); w4=[f"{nm}{pr}" for pr,nm,d in v if d<=4]
                a=accs[sid].get(pcid,'?'); partners.append((mind,SYM.get(a,a),a,bool(w4)))
            partners.sort()
            results.append({"set":setname,"gene":r['gene'],"res":r['res'],
                "nuclear_partners":[{"gene":p[1],"uniprot":p[2],"min_dist":round(p[0],1),"interface_<=4A":p[3]} for p in partners[:5]]})
    return results

# build site lists (gene,res,chg,anc)
modern=[(x[0],x[1],x[2],x[3]) for x in lp["modern_ns"]]
shared=[(x[0],x[1],x[2],x[3]) for x in lp["sharedmn_ns"]]
out={"modern_lineage":run("MODERN",modern),"sharedMN_lineage":run("SHARED_MN",shared)}
json.dump(out, open(os.path.join(ROOT,"results","task4_partners.json"),"w"), indent=2)

# print + collect union of partner genes
allgenes=set()
for setname,res in out.items():
    print(f"\n===== {setname} =====")
    for r in res:
        if r.get('err'): print(f"  {r['site']}: {r['err']}"); continue
        ps=r['nuclear_partners']
        tag=f"{r['gene']} {r['res']}"
        if not ps: print(f"  {tag:12} -> (no nuclear partner <=8A)"); continue
        s=", ".join(f"{p['gene']}{'*' if p['interface_<=4A'] else ''}({p['min_dist']}A)" for p in ps)
        print(f"  {tag:12} -> {s}")
        for p in ps: allgenes.add(p['gene'])
print("\nUNION of nuclear partner genes (for catalog check):")
print(" ", sorted(allgenes))
json.dump(sorted(allgenes), open(os.path.join(ROOT,"results","task4_partner_genes.json"),"w"))
```
