## Supplementary Information section 3Supplementary Information section 1 for "Hypothesis: A modern human range expansion ∼300,000 years ago had a critical role in Neandertal origins"

### Supplementary Information section 3

Modern human range expansions predict fixation of modern mtDNA and Y in Neandertals

#### Motivation

This note identifies a solution to a previously unsolved puzzle: that sampled Neandertals carry mitochondrial DNA (mtDNA) and Y chromosomes related to living humans within a few hundred thousand years (ka)[1,2], even though the main split of modern humans from Neandertals and Denisovans occurred around 825-694 ka (range of dates from ref. 3).

The proximate cause is well understood: gene flow from early modern humans into Neandertals in the last few hundred thousand years, as supported not just by uniparental[4] but also by genetic data from chromosomes 1-22 and X[5,6,7]. In this study I simulate this event assuming ~300-200 ka for the date of this admixture[7], a range consistent with multiple estimates[4, 7,8].

But the ultimate cause of discordance between the nuclear genome and the uniparentally transmitted markers remains perplexing. In Neandertals, mtDNA and Y chromosomes closely related to those of modern humans occur despite the fact that on chromosomes 1-22, there is only a small proportion of such ancestry (~3-7%)[7,8,9.] In the absence of selection, and assuming x% pulse gene flow from early modern humans into the archaic ancestors of Neandertals, population genetic theory shows that the probability that lineages derived from modern humans rose to one hundred percent at any locus in the genome is x%[10]. Thus, the probability that ~3-7% modern human admixture caused both of these alleles to drift to fixation is tiny: at the midpoint of that range, it is the product of the two marginal probabilities: 0.25% (5% × 5%).

The puzzle posed by the discrepancy between mtDNA and genome-wide phylogenies was first appreciated in 2010[1,11]. That puzzle became even more challenging after it was reported in 2020 that the Y chromosome follows the same pattern[2]. The paper reporting the Y chromosome result recognized that these findings were too implausible to be due to chance, and argued that modern human lineages on the male side (tracked by the Y chromosome), the female side (tracked by mtDNA), or both, must have been selected relative to the archaic alleles[2].

However, the mechanism that the authors proposed cannot explain the data. Following previous modeling of this admixture[2,5,7,9], they simulated ~5% gene flow from a homogeneous modern human ancestral population into homogeneous archaic Europeans. I call this the "migration pulse model" to distinguish it from a geographically explicit "range expansion model". They coupled this demography with selection for the early modern human mtDNA or Y chromosomes due to an accumulation of more deleterious mutations on the archaic lineage since they separated. However, as I show in Supplementary Information section 1, biological constraints mean that the fitness gap between the early modern human and archaic lineages on mtDNA and the Y would not be large enough to drive this process. My simulations show that the probability of fixation of modern human lineages at both mtDNA and the Y is <2-3% given real biological constraints.

But a migration pulse model is an oversimplification, as it assumes that the geographic configuration in which the admixture occurs does not affect the process. In fact, in a range expansion, population genetic phenomena occur that are driven by the geography and are qualitatively different from those that are expected in other contexts.

One emergent phenomenon from range expansions that can help explain the observations is “massive introgression of local genes”[12]. Even a low rate of interbreeding between modern and archaic humans when they encounter each other is expected to be amplified at the wavefront of a range expansion, so the population that results ends up with little modern ancestry genome-wide, providing a natural explanation for the ~3-7% modern human ancestry on the autosomes.

A second way that range expansions can explain the data is at the uniparentally transmitted markers (mtDNA or the Y). The predominant pattern for human foragers and great apes is that dispersal across groups is mediated by one sex[13,14]. Modern ancestry in a range expansion is retained automatically at the marker tracking the non-dispersing sex (Figure S3.1), matching the data. This does not explain patterns at the other uniparentally inherited marker, but simplifies the challenge.

Third, range expansions can help to explain the data even for the other uniparentally transmitted marker. In a range expansion, genetic variants from the modern human population start out in the majority, whereas in a migration pulse they are in the minority. This means that for a range expansion, the challenge in fixing a modern lineage is different. For a range expansion, the challenge is to block archaic lineages from coming in despite a flood of migration, but in a migration pulse the challenge is to push them up from a low frequency, which is a qualitatively different problem. The rest of this note identifies plausible means to block archaic DNA at the other uniparentally inherited marker.

**Figure S3.1: Sex-biased range expansion naturally fixes a uniparentally transmitted marker.** (a) Migration pulse. 5% gene flow from early modern humans into archaic humans is followed by random genetic drift. Even under complete sex bias, fixation is unlikely (~10%; simulation framework in Appendix C of Supplementary Information section 1, 1000 simulations per datapoint). (b) Range expansion. Fixation for one marker is certain if there is full sex bias. This does not explain the pattern at the other uniparentally transmitted marker, where selection is still needed to block archaic lineages from coming into the population, but a range expansion helps explain this too (simulation framework in Appendices A and B, 1000 simulations per datapoint).

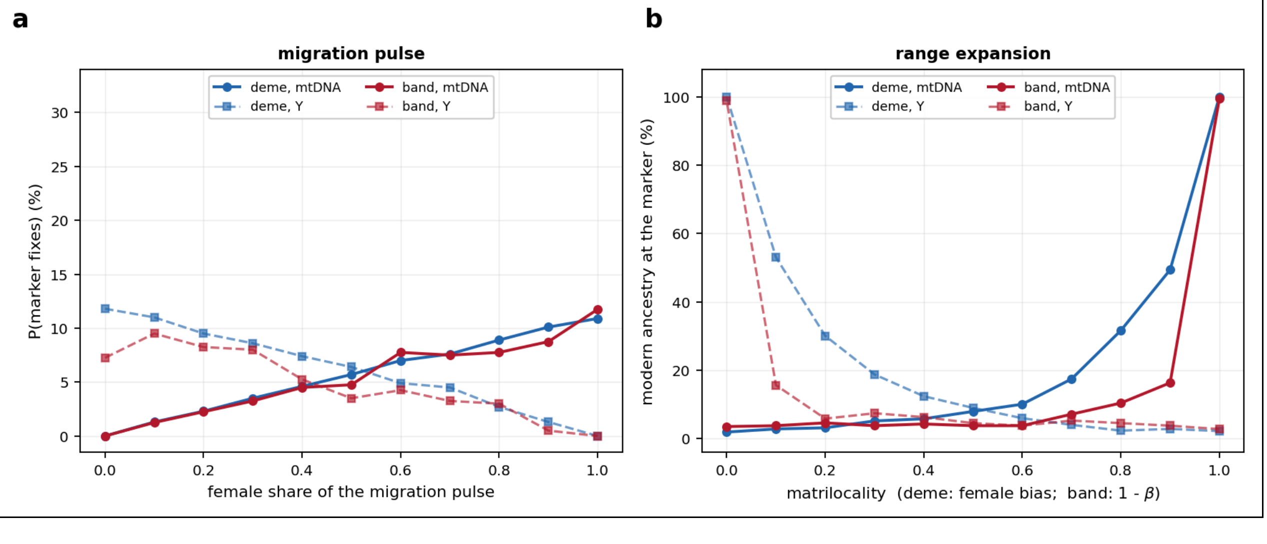

**Simulations of early modern human range expansion**

The simulations that follow are a product of my dialogue with AI coding agents of the Anthropic Claude family (Opus and Fable). I used the agents to write the simulations and ABC code in Python, to run diagnostics, and to help compose the text (any suggestions from the AI agents were edited by me). All quantitative results are produced by my execution of the cited scripts on the O2 computing cluster at Harvard Medical School, or my personal computer. I checked the robustness of the output, and reviewed the scientific interpretations, modeling choices, and final claims.

Throughout this note, I present results from two qualitatively different simulation frameworks, a "deme model" and a "band model", detailed in Appendices A and B.

In the deme model (Appendix A), inspired by the work of Currat and Excoffier studying range expansions in diverse ecological contexts[12,15], I model the European landscape as a square grid of demes each with a fixed carrying capacity, and in which individuals belong to a population that is either modern or archaic. I vary the rate of migration between demes, the propensity of individuals of different populations to interbreed, the relative rate of reproduction of individuals from modern and archaic populations when they compete, and the extent to which the population to which a person belongs (their "identity" which can be either modern or archaic in the model) is transmitted by mothers. In some cases, I make plots using a "baseline" set of simulation parameters (12×12 demes, deme carrying capacity K = 11, migration rate m = 0.10, rate of interbreeding with the other group $\gamma = 0.12$, and growth rate of moderns relative to archaics in the same deme r = 2). However, every scientific claim is based on running simulations over a wide grid of parameter combinations and verifying its validity across all of them.

In the band model (Appendix B), which aims to be more ecologically and anthropologically natural, I use a geographically explicit map of the part of Europe where Neandertals lived[16], which I seed at random latitudes and longitudes with bands of archaic humans, along with one modern human band in the southeast. In the simulations, bands always belong to one population (they cannot harbor a mixture of modern and archaic people as in the deme model), and I specify band size, band density, and the whole-band diffusion rate per generation. I also specify the dispersal rate of exogamous individuals who join different bands (if one band is modern and the other is archaic this will involve switching population), along with the degree of sex bias (e.g. only males switch bands). I specify competition for resources (when bands deviate from their optimal size or get too close to each other, their reproduction rate decreases), and the relative success of modern and archaic humans in competition. I make some plots using a set of baseline simulation parameters (70 bands, optimal band size K = 25, whole-band movement rate 0.45 per generation with a step scale of 60 km, exogamous dispersal scale 80 km, sex bias of exogamy $\beta = 0$ so only males disperse, growth rate of modern relative to archaic humans r = 2, rate of exogamy e = 0.19). Every claim is based on running simulations over a large grid of parameter combinations, and verifying its validity across all of them.

**Four scenarios whereby the other uniparentally transmitted marker would likely be retained**

Sex-biased range expansion automatically retains one uniparentally transmitted marker. But what of the marker that did not move with the expansion? Below I identify four scenarios that work.

*(Scenario 1) A range expansion where group label is transmitted by mothers, combined with an epistatic interaction between the archaic Y and modern partner alleles, can block the archaic Y chromosome from flooding into the expanding modern human population*

"Haldane's Rule" specifies that when two populations have been isolated for long enough that hybrids have compromised reproductive phenotypes, it is almost always the heterogametic sex (males for mammals) that is affected[17]. For a range expansion in which transmission of population label from generation to generation was through the mother, these modern human first generation (F1) hybrids would be males carrying archaic Y. If it was challenging for them to transmit their Y to subsequent generations due to reduced fertility or sex distortion leading them to produce primarily daughters, this has the potential to block archaic Y from the population.

Haldane's Rule often arises due to Dobzhansky-Muller Incompatibilities (DMIs): co-evolved variants that only work when they are on the same genetic background causing compromised reproduction in hybrids that do not carry both[18]. An important role for DMIs in the ~300-200 ka admixture event is not just theoretical: there is strong evidence for widespread DMIs as attested by reduced human modern ancestry at functionally important regions in Neandertals[5,7]. Systematic reduction of ancestry from one admixing population in theory can be explained by two processes: DMIs[18], or greater accumulation of deleterious mutations in populations of smaller size[19]. Here it is clearly DMIs, since the inferred smaller size of the archaic than of the modern population since they diverged[3] would be expected to increase modern ancestry, not decrease it as observed.

I first carried out simulations to explore "classic" DMI architectures that cause sterility in first-generation (F1) males, and found, to my surprise, that almost none fit. For each architecture, I tested 744 simulation parameter combinations for the deme model and 567 for the band model. The 744 are the survivors, out of a full Cartesian product of 11,424, of the requirement that the simulated autosomal effective population size fall within 10% of the inferred average from ~250-50 ka based on a demographic reconstruction in ref. 3 (Appendix A). (The band framework needs no such filter, because its census is fixed; its 567 combinations are the whole product; Appendix B.) For each tested parameter combination, I assessed if the results matched the data from the perspective of: (a) leading to a modern human autosomal proportion of 3-7%, (b) the range expansion completing in a time span of 100-1,000 generations, and (c) the expansion establishing rather than stalling in more than 50% of simulation instances. I repeated the simulations 10 times in the initial screen, and 100 or 200 times for every combination that passed the screen, the larger count for the families screened earliest. If many parameter combinations matched these criteria, I measured the fraction that retained modern human lineages in a way similar to empirical data by sampling 10 Neandertal males at the end of the simulation, and scored a match if all of them carried modern human Y and mtDNA (Figure S3.2). I obtained the following results:

- **DMIs between modern mtDNA and archaic Y provide the only DMI architecture that if it occurred would be highly likely to fit the data, but it is biologically unlikely.** Sterility due to such an interaction should immediately block archaic Y chromosomes from entering the population. When I searched for a possible mediator for such incompatibilities by testing whether any protein encoded by the male-specific region of the Y is included in MitoCarta3.0, an inventory of 1,136 human genes with strong support for mitochondrial localization[20], the only two annotated genes on the Y chromosome, *SLC25A6* and *GTPBP6*, lie in pseudoautosomal region 1, so neither is inherited in a male-specific way.

- **DMIs between modern mtDNA and a locus outside Y are unlikely to explain the data.** mtDNA is known to often participate in DMIs[21], and there are orders of magnitude more genes outside the Y chromosome than on it, so I carried out simulations of a range of such architectures (Figure S3.2). However, even in the most favorable of these architectures, the modern human partner allele tends to be rapidly replaced in the first generations of the range expansion by the archaic one due to selection against the DMI. I also searched for evidence of DMIs of this type through structural analysis of predicted physical interactions between mtDNA-encoded OXPHOS subunits and their nuclear-encoded partners (Supplementary Information section 2). I failed to find any such interaction, although a caveat is that mitonuclear incompatibilities need not involve physically interacting residues (for example, in swordtail fish hybrids, a mitonuclear incompatibility involving the ATP synthase complex does not involve substitutions at contacting positions[22]). Taken together, these analyses make it unlikely that a DMI involving modern mtDNA explains the data.

- **Compounding DMIs between modern mtDNA and archaic loci are unlikely to explain the data.** I carried out simulations in which the mtDNA partner was not a single locus, but multiple loci at which an archaic allele is incompatible with modern mtDNA in males[23,24,25]. Because an archaic father must now transmit modern alleles at all loci at once, the escape route that defeats the single-locus version is suppressed. However, this gives too-high autosomal ancestry proportions throughout the grid of simulation parameter settings I tried, because very large archaic gene flow rates are needed to overcome the reduced fertility.

- **DMIs not involving mtDNA are unlikely to explain the observed patterns.** I simulated incompatibilities that do not involve mtDNA, including X-Y incompatibility. This does not work: modern human X chromosome alleles flow into the archaic population and are highly selected for there as they do not participate in the DMI, ending the DMI in a few generations.

- **DMIs involving *PRDM9* are unlikely to explain the observed patterns.** I explored the possibility of a DMI at the *PRDM9* gene on chromosome 5, motivated by two lines of evidence suggesting it as a candidate for hybrid infertility. First, it is involved in infertility of house mice (*Mus musculus*) subspecies that interbreed in central Europe, due to an interaction of an allele at *PRDM9* with a partner locus on the X chromosome at *Hstx2*[26,27]. Second, there is evidence for a sweep in Neandertals for the modern human-like allele (*PRDM9*-A), making it natural to consider the possibility that this contributed to hybrid sterility[8]. The strength of selection for *PRDM9*-A relative to the archaic allele in Neandertals was estimated by ref. 8 to be $s = 0.5\%$[8], and I found that I could not obtain a fit with simulations with this selection coefficient (Figure S3.2). I also carried out the reverse simulations, with a maternally transmitted modern *PRDM9*-A allele and a paternally transmitted archaic partner, and this also does not result in retention of the modern Y with either 0.5% or 5% selection for modern *PRDM9*-A.

**Figure S3.2. Systematic survey of 43 candidate mechanisms for matching the data.** Orange band, purple deme, points full grid of parameter combinations, 10-200 replicates/point. Black rectangle is acceptance region (modern autosomal ancestry 3-7%, expansion 100-1,000 generations). Mean probability that all ten sampled males are modern at both mtDNA and the Y is shown for band top left, deme bottom left, averaged over admissible combinations (not FAIL). Dark points are where a sample of ten males is modern at both markers with probability ≥50%. Every point inside the box is plotted in the Figure 3b panel associated with the same scenario.

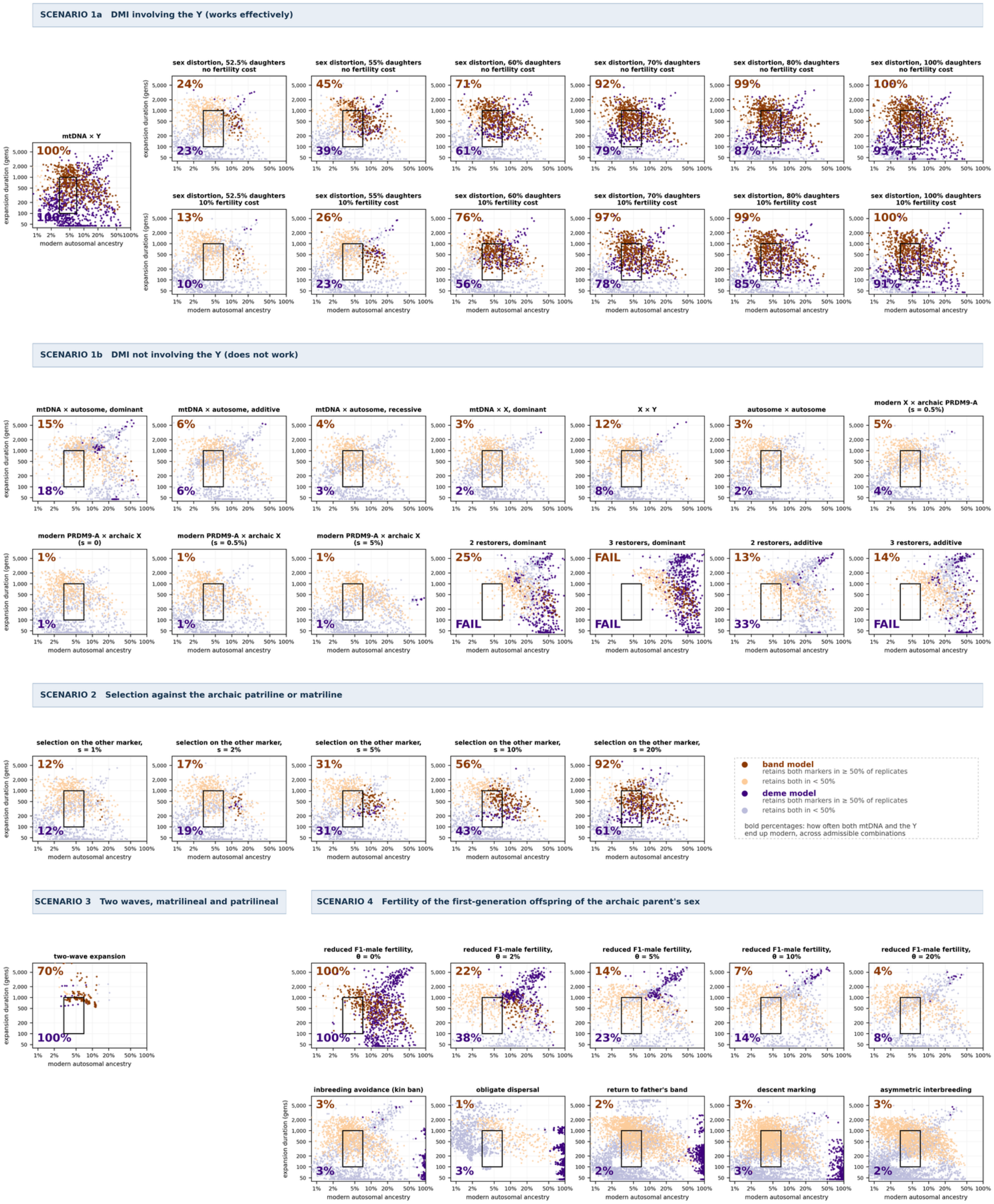

While classic DMIs are not a promising route to explaining the observed patterns, there is also an alternative type of DMI that is common in nature that affects the sex ratio of first generation male hybrids[28,29]. This arises because alleles on chromosome X and Y have different interests when it comes to sperm production: it is in the X chromosome's interest to produce a higher proportion of X-carrying sperm leading to more daughters, and in the Y chromosome's interest to produce more sons. Within a population, sex-distorting alleles on one of the sex chromosomes tend to be compensated by alleles on the other sex chromosome to restore sex balance to 50:50. The better-documented polarity is an X chromosome distorter compensated by a Y chromosome suppressor (the case considered here)[30,31]. When two divergent populations mix, and one harbors sex-distorting alleles, the suppressor is not present in first generation hybrids resulting in sex skew in the offspring. Real examples in nature include the Slx/Sly ampliconic arms race in hybrid mice, which distorts offspring sex ratios when copy numbers are mismatched[32], and *Overdrive* in *Drosophila*, which is both a hybrid-sterility gene and a segregation distorter[33].

A sex-distorting allele causing primarily daughters in first-generation male hybrids is an effective mechanism for blocking archaic Y from establishing itself within the population. A major challenge for "classic" DMIs explaining the observed patterns is that the F1 male sterility they produce causes strong selection against the archaic allele at the partner locus, which after the first few generations is replaced by the modern allele so the barrier to archaic Y disappears. In contrast, for sex distortion, only the Y is removed from the transmission stream—not the male who carries it—so this process surgically removes archaic ancestry at the Y, leaving the rest of the flow of archaic genes unopposed. The daughters of the F1 males receive a double dose of modern X, propagating the sex distortion mechanism so it is positively not negatively selected (in contrast to a classic DMI, where the partner allele on the modern human side is selected against).

I carried out simulations ranging from 52.5% to 100% daughters. Even at a 60% skew, the modern Y is retained in 84% of band and 50% of deme replicates (Figure 3b, Supplementary Table 1). At every drive strength from 70% daughters upward, the modern Y is retained in the great majority of instances: 97-100% in the band framework and 74-90% in the deme framework.

To probe the robustness of this mechanism, I also tested imposing a fitness cost of 0.1 to killing up to half of one's sperm (the Y-carrying ones), and found that it did not substantially compromise the mechanism (Figure S3.3). Another way to suppress this process would be for a new Y chromosome mutation suppressing the sex distortion to arise *de novo* during the range expansion. But this is biologically unlikely. The Y chromosome per base pair mutation rate is around $3.0\times10^{-8}$/bp/gen[34,35], the exonic span of coding genes in the X-degenerate sequence is around 100,000 bp[36], and assuming that the proportion of mutations changing the protein is 0.72, the estimated *de novo* deleterious input into genes per generation is $3.0\times10^{-8} \times 100{,}000 \times 0.72 \approx 0.22\%$. This is lower than the ~0.5% at which the mechanism is expected to stop working for the deme model and ~3% for the band model, so even in the implausible scenario that all *de novo* mutations in genes on the Y suppressed the X-linked sex-distorter, it would not be enough to stop the process of eliminating the archaic Y by sex distortion.

Speaking more broadly, the DMI architectures that match the data divide according to what locus the incompatibility acts on. A DMI architecture that acts on the Y chromosome directly retains the modern Y chromosome effectively: an mtDNA - Y incompatibility is near-certain to work across every admissible parameter combination, while X - Y sex distortion causes males to be ineffective at transmitting their Y chromosomes. In contrast, all the DMI architectures that do not involve the Y chromosome work poorly: there is strong selection against the archaic allele participating in the DMI, which is then rapidly replaced by modern human alleles, stopping the DMI.

**Figure S3.3. X-linked drive retains the modern Y.** *Left:* How the probability that an F1 male's offspring are daughters affects the probability that the modern human Y chromosome is fixed in an expansion where females transmit the population label to the next generation, with dashed lines showing a 10% fertility cost (400 replicates per point). *Right*: Stopping the process would require an implausibly high mutation rate: the rate of occurrence of new X-drive suppressing mutations must be >>0.5% per Y chromosome per generation, greater than the 0.22% of new mutations in genes per generation across the entire Y (vertical line).

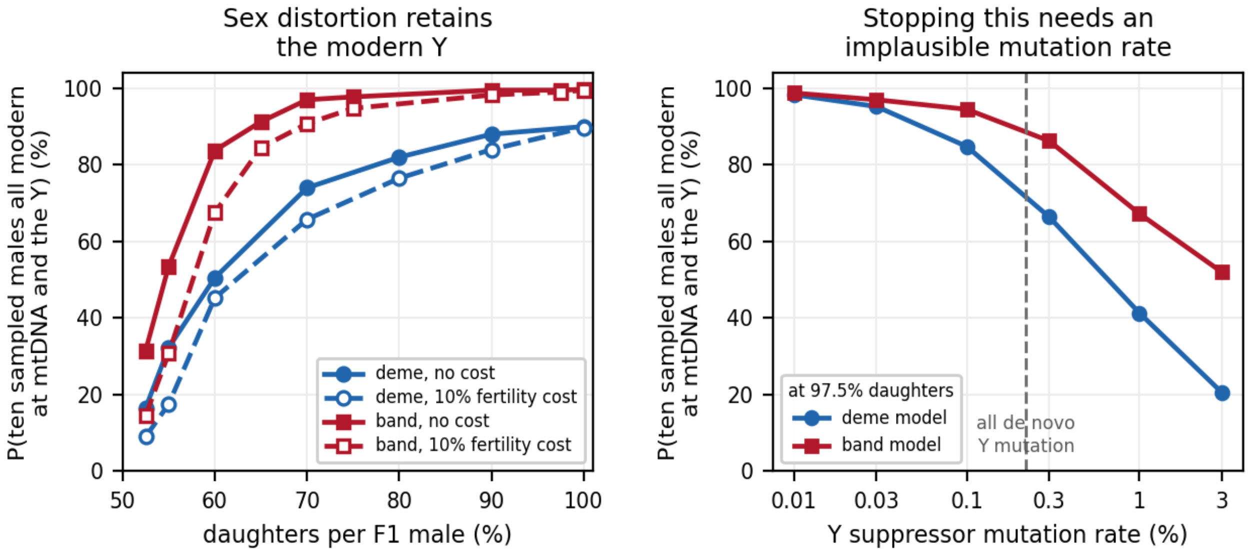


*(Scenario 2) Sex-biased modern human expansion, combined with selection for prestigious modern matrilines or patrilines, can lead to retention of the other uniparentally transmitted marker*

Biological selection is not a promising way to retain either the modern human mtDNA or Y in a sex-biased range expansion. In Figure S3.4, I show that for a migration pulse, a fitness gap of s ~ 0.9% on either mtDNA or the Y chromosome is enough to give the modern allele roughly even odds of being retained. Since the estimated gap for mtDNA is s = 0.32% and for Y it is s = 0.09-0.17%, retention is unlikely (Supplementary Information section 1). In the context of a range expansion, the population dynamics are even worse. Because modern uniparentally transmitted markers do not start at a substantial copy number as in a migration pulse, and instead enter as a handful of founding copies where drift is strong and selection works poorly, they tend to be lost before selection can begin to be effective. My simulations show that even odds of retaining modern copies of uniparentally transmitted markers requires a gap of about 6.6% in the band framework and 15.6% in the deme framework. This is so far outside the range that is consistent with the data, that this mechanism never works to retain modern mtDNA or Y (Figure S3.4).

**Figure S3.4: A simple mtDNA or Y chromosome fitness gap between the archaic and modern ancestors of Neandertals is unlikely to have been enough to drive these alleles to fixation during a range expansion.** Comparison of fixation of a uniparentally transmitted marker for a 5% migration pulse (black, 4,000 replicates per point), versus range expansions in a band (red, 1,000 replicates per point) or deme framework (blue, 346 replicates per point). The fitness gaps estimated for mtDNA (median 0.32%) and the Y (0.09-0.17%) are estimated in Supplementary Information section 1 and were already shown there to be not enough to drive the modern alleles to fixation (black curve). My simulations show that the challenge is far worse for a range expansion; intuitively, this is because the modern lineages enter the population in only a handful of copies, so the stochastic effects that the selection needs to overcome are stronger. Every curve here is a simulation under an unconditional biological fitness gap, the kind that arises from differential mutational load, and all are carried out with the mtDNA simulation machinery; the Y gap is therefore read against the mtDNA curves, which is informative only to the extent that the two markers share an effective population size and a transmission rule. The cultural advantage of Scenario 2's prestigious patrilines or matrilines is a different quantity and is not shown here.

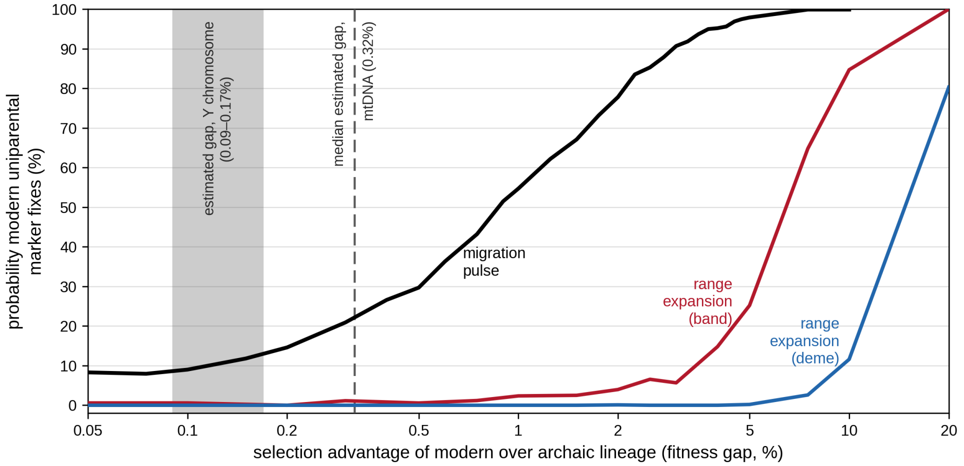


What process could lead to a reproductive advantage of this order for modern human patrilines or matrilines over archaic ones? Human cultures often track clan membership generations deep into the past either along the entire male or the entirely female line, and these would track either Y chromosome or mtDNA[37]. Consider a scenario where men from more modern patrilines tend to be better at competing for mates than archaic ones due to greater prestige in a community being attached to such a lineage. In the context of human societies where males typically have high variance in reproductive success whereas almost all females who can have a child do so, this can result in strong selection for some male patrilines (and thus Y chromosomes) over others. (A similar argument works for prestigious matrilines, although less well because of the lower variance in reproductive success for females.) Prestige of this kind is conferred by the community a man belongs to, so in the simulations the advantage is conditioned on group label: in each band (band model) or lineage (deme model), the patriline that matches the group's label is favored and the one that mismatches it is selected against, by the same coefficient s. Across the admissible parameter combinations, the median probability that ten sampled males carry modern Y and mtDNA crosses one half at s ≈ 9% in the band framework and s ≈ 14% in the deme framework (Figure 3b, Figure S3.2, Supplementary Table 1). These thresholds are not comparable to the 6.6% and 15.6% quoted above: those are for an unconditional biological gap acting everywhere; here the advantage acts only where a lineage matches its group label, and so has fewer opportunities to be expressed.

*(Scenario 3) Sex-biased range expansion can lead to retention of modern human mtDNA, preceded or succeeded by an expansion with the other bias leading to retention of the modern human Y*

Another scenario that could be consistent with the data is an expansion of modern humans where females are the sex that do not disperse leading to retention of modern mtDNA, followed or preceded by an expansion where males are the sex that do not disperse leading to retention of modern Y. My simulations confirm that this explains the data (Figure S3.2). In the band framework, the expansion has to be timed by when the uniparental markers reach 100% of one allele type rather than by when the last archaic-labelled band disappears: 49 band parameter combinations are then admissible, with a median probability of 73% of both markers' fixation.

*(Scenario 4) Sex-biased expansion, combined with a cultural mechanism, can lead to first-generation offspring of the same sex as their archaic parent not being raised*

Consider the possibility that in a sex-biased expansion, the same-sex children of incoming archaic males were unsuccessful in competing for modern mates. My simulations in Figure 3b of the main text and Figure S3.2 confirm that this works to retain modern human DNA at the uniparental marker not driving the expansion. But the simulations also show that the relative fertility of F1 males, $\theta$, needs to be extremely low to do this. Retention of modern human DNA becomes unlikely once $\theta$ rises above roughly 0.25% in the deme framework. In the band framework it is worse: the median never reaches a half at any non-zero $\theta$ tested, standing at 0.37 even at $\theta = 0.1\%$. In both frameworks, the mechanism therefore requires F1 fertility to be extremely low, not merely low.

The one cultural scenario I could identify that in theory could produce the effect of extreme infertility in first generation hybrids of the same sex as the archaic parent was one in which F1 females and males were treated extremely differently after birth. This could correspond to a band where female infants were more valued because descent and group membership passed through them and females are more limiting in reproduction than males[38]. Alternatively, male F1 hybrids could have been selectively incorporated into the next generation but females not. This has precedent in societies today, for example in central African rainforest hunter-gatherers and Bantu agriculturalist neighbors where male and female offspring of mixtures are incorporated into different groups depending on their sex[39,40]. Nevertheless, a 100-to-1 discrimination against hybrid offspring of one sex versus the other seems unlikely.

I carried out simulations of other cultural mechanisms I hypothesized might lead to very low $\theta$, but could find none. Below I list five failure modes, all simulated in Figure S3.2 in both frameworks, which shows that despite exploring an exhaustive array of parameter combinations, they could not match the data. Across all, the highest probability of retaining the uniparental marker not driving the expansion while holding autosomal ancestry at around 5% was 12% in the deme framework and 9% in the band framework:

1. I explored the hypothesis that F1 hybrids may have avoided mating with modern humans due to a taboo against mating with close relatives: inbreeding avoidance[41,42].
2. I explored the hypothesis that the sex that moves across groups practiced obligate dispersal from their natal band, also common in animals and in humans[43,44].
3. I explored a modified version of the second scenario, where in groups where females stayed within groups F1 sons who were offspring of modern mothers and archaic fathers had a strong preference for returning to their father's band, and in groups where males stayed within groups

F1 daughters who were offspring of modern fathers and archaic mothers preferred to return to their mother's band.

4. I explored the hypothesis that men of archaic paternal descent were marked as such under paternal descent reckoning and sought mates among archaic rather than modern bands, or that women of archaic maternal descent sought mates among archaic not modern bands.
5. I explored asymmetric interbreeding, in which archaic women took modern mates at a higher rate than modern women took archaic mates. This could conceivably cause the Y chromosomes of the archaic population to modernize more rapidly than the Y chromosomes of the modern population would become archaic, and thus might seem like a promising explanation for the observations. However, my simulations showed that this scenario failed, as the same matings that modernize the archaic Y pool also rapidly modernize the autosomes of the archaic population at the wavefront, so the ultimate modern proportion stays too high to fit the data. Flipping the sexes in the simulations also fails to account for the observed patterns.

**Summary**

In this note and in Supplementary Information section 1, I showed that the previously suggested scenario of a migration pulse with biological selection is unlikely to be able to explain the retention of both the Y chromosome and mtDNA in Neandertals[2]. However, in a range expansion, I have shown that there are multiple plausible scenarios where the observed patterns are highly likely.

At a high level, the advantage of a sex-biased range expansion lies in the fact that it makes early modern humans central to the process of formation of the resulting admixed population (Neandertal ancestors), despite the ultimately small proportion of nuclear ancestry they contribute. The retained modern human ancestry at one of the uniparentally inherited markers is then due to this central contribution of early modern humans.

#### Appendix A: Deme simulations on a square grid

Loosely following the framework of Currat and Excoffier[12,Error! Bookmark not defined.], I approximate Europe as a grid of demes, each with a carrying capacity of K people, a number that I hold to be the same for modern and archaic lineages. At generation t = 0, every deme except the corner deme in the southeast is seeded with Poisson draws of expected value K/2 archaic-lineage males and K/2 archaic-lineage females. The corner deme is seeded the same way with modern-lineage individuals. The baseline simulation model is a 12 × 12 grid with K = 11, and a census of 1,584. I tuned the effective population size across the landscape to match the inferred population size of the Neandertal lineage following mixture based on PSMC[3].

This model is an extreme oversimplification. Among other limitations, Europe is not a square grid of uniformly habitable cells. The goal of the simulations is an existence proof: that large-scale incorporation of local archaic genes at the autosomes can coexist with qualitatively different behavior on the Y chromosome and at mtDNA. The band model of Appendix B is more realistic in being geographically explicit and having a more anthropologically natural mechanism, and leads to qualitatively similar conclusions.

Each generation iterates four steps. What follows is a description of an expansion where women pass the group label to the next generation, but mathematical symmetry means the same results are obtained with the labels on mtDNA and Y exchanged, so I do not describe it separately.

(i) **Migration.** Each individual moves one deme in each adjacent direction with probability *m*, unless that would leave the grid.

(ii) **Counts of children.** A deme that contains no male, or no female, produces no children. Otherwise, the numbers of modern and archaic males and females in the next generation are Poisson draws with the expected values given in Table S3.1. The parameter *r* is the fitness of modern-lineage individuals relative to archaic-lineage individuals competing for the same deme. The assortative-mating parameter γ sets how readily cross-lineage pairings form.

**Table S3.1: Poisson expectations for eight offspring types in the next generation**

| **Father in gen. *t*** | **Mother in gen. *t* (passed to child)** | **Expected number in category in gen. *t*** | **Sex in gen. *t+1*** | **Contributes to which count in generation *t+1*** |
|---|---|---|---|---|
| Modern | Modern | $KrMM_{i,j,t}FM_{i,j,t}/Q$ | Male | $MM_{i,j,t+1}$ |
| Modern | Modern | $KrMM_{i,j,t}FM_{i,j,t}/Q$ | Female | $FM_{i,j,t+1}$ |
| Modern | Archaic | $K\gamma MM_{i,j,t}FA_{i,j,t}/Q$ | Male | $MA_{i,j,t+1}$ |
| Modern | Archaic | $K\gamma MM_{i,j,t}FA_{i,j,t}/Q$ | Female | $FA_{i,j,t+1}$ |
| Archaic | Modern | $Kr\gamma MA_{i,j,t}FM_{i,j,t}/Q$ | Male | $MM_{i,j,t+1}$ |
| Archaic | Modern | $Kr\gamma MA_{i,j,t}FM_{i,j,t}/Q$ | Female | $FM_{i,j,t+1}$ |
| Archaic | Archaic | $KMA_{i,j,t}FA_{i,j,t}/Q$ | Male | $MA_{i,j,t+1}$ |
| Archaic | Archaic | $KMA_{i,j,t}FA_{i,j,t}/Q$ | Female | $FA_{i,j,t+1}$ |

(iii) **Transmission**. For each child, I draw parents uniformly from the candidates of the required type in the same deme in the previous generation. I track 41 unlinked loci: mtDNA from the mother, Y from father to son, 36 autosomal loci with one allele from each parent (32 used to track ancestry and 4 reserved for selection, so no locus under selection is ever used for ancestry computations), and three X loci with females diploid and males haploid (one neutral locus used to track ancestry, one reserved as a selection or incompatibility partner, and one reserved for the sex-ratio distorter).

(iv) **Selection**. Additive selection removes an offspring with probability *s* per archaic allele at the selected locus, so an archaic homozygote at a diploid locus is removed with probability 2*s*. Selection may act on the Y, on one of the four reserved autosomal mechanism loci, or on the X selection-candidate locus; on the Y it is conditioned on group label, as described under the mechanism parameters below. Selected offspring are dropped rather than resampled, so a deme may fall below carrying capacity. Epistatic (DMI) selection, X-linked drive, polygenic restorers and the F1-male fertility parameter θ are layered on this step.

The parameter θ deserves special discussion because it tracks culture. It is keyed on the natal identities of the two parents: whether a male's mother and father come from different groups. It is not a locus, not heritable, and it carries no information about any biological incompatibility.

**Reseeding the origin when the expansion stalls.** The origin deme is repopulated with a modern individual if it is left without one, but only while no modern-lineage individual survives anywhere on the grid and archaic-lineage individuals remain.

**Restricting to parameters leading to 3-7% modern autosomal ancestry.** An empirical observation that I use to constrain the simulations is that sampled Neandertals are 3-7% modern in their nuclear genome. Autosomal ancestry is not automatically 3-7%; for example, reducing F1-male fertility removes the archaic autosomes those fathers would have carried, and strong selection on the Y chromosome removes archaic-Y-carrying-males along with their autosomes. I thus retune other parameters for each condition to get the autosomal ancestry proportions in the right range.

**Restricting to parameters leading to an effective population size matching estimates.** The post-admixture size is taken from PSMC trajectories of the three Neandertal-lineage genomes in ref. 3, digitized and averaged over the window between admixture and the age of the sequenced individuals (250-50 ka). I take a time-weighted harmonic mean, which is the appropriate average for drift, giving 1,447 diploids. I then measure effective sizes to calibrate the simulations: each configuration is run to a post-admixture steady state, I reset every marker to frequency one half, which for the three kinds of locus means three different draws only because they differ in ploidy: each individual's mtDNA is drawn Bernoulli(½) because mtDNA is a single haploid copy, each male's Y likewise, and each autosomal locus is drawn binomial(2, ½). I track the decay of expected heterozygosity, regressing $\ln(E[p(1 - p)])$ on time. I restrict analyses to parameter combinations that match the effective population size estimated by PSMC to within 10%, that is 1,302-1,592 diploids. The target is the time-weighted harmonic mean of the Neandertal-lineage effective size over the window between admixture and the age of the sequenced individuals, 250-50 ka, so the requirement is that the expansion's own effective size matches the average post-mixture size over the couple of hundred thousand years that follow it. It is of course possible that the effective size of humans during the expansion period was different from its long-term harmonic average afterward, but I do not explore scenarios like this.

**Restricting to parameters leading to a range expansion of reasonable duration.** I only use parameter combinations that at least 50% of the time lead to a range expansion duration (generations until the extinction of all people from the archaic group) of 100-1,000 generations. If the expansion fails and modern humans go extinct, I do not tabulate the simulation results.

**Interpolating the usable parameter region.** Screening every parameter combination at full length would be computationally wasteful, because most fail one of the constraints above and the constraints vary smoothly. I therefore run a coarse grid, stopping each replicate as soon as the archaic lineage goes extinct, and fit a smooth surface to the result. The surface is a local linear regression of log autosomal ancestry on the parameters, weighted by a Gaussian kernel in parameter space, and I invert it to read off the value of the calibrated knob ($\gamma$) at which autosomal ancestry hits its target for any combination of the others. Where it falls below 0.8, the grid is reported as too coarse and I carry out follow-up simulation iterations at higher density. Only combinations that survive this screen are run at full length and full replication.

**Simulations of a migration pulse.** The migration-pulse comparator used throughout this note is described in Supplementary Information section 1, Appendix C.

**Sampling time.** I sample the simulations up to 6,993 generations after admixture, which at 28.6 years per generation is the 200 thousand years between admixture at 250 ka and sequenced Neandertals at 50 ka. To speed simulation, I stop runs as soon as both uniparental markers are absorbed: either fixed or lost. After absorption, neither marker can change and autosomal ancestry is a martingale, so the expectations are unaffected.

**Variation of demographic parameters**

To test how the conclusions depend on the choice of parameters, I vary the following parameters, one at a time around the baseline (Figure S3.5). I do not vary grid, K and m independently. Instead, I only use combinations whose associated effective size falls within 10% of the PSMC target of 1,447 diploids. (1,584 is the deme model's census at the baseline geometry, a different quantity.)

- grid is the number of demes along each side of the square lattice (baseline 12, so 144 demes, varied over 6, 8, 10, 12, 14, 16 and 20).
- K is the carrying capacity of each deme, which is the same for both lineages (baseline 11). I vary it over 26 values from 6 to 46, of which 23 deliver an effective size within 10% of the PSMC value inferred empirically at one or more grid sizes. Across all three coupled axes, 56 of the 508 measured (grid, K, m) combinations (508 rather than the full 7 × 26 × 5 = 910 because the 26 K values and 5 m values are the union of the effective-size screen's successive ladders, not a single Cartesian pass; Supplementary Table 1, "Effective-size screen" tab) are admissible and are therefore searched.
- m is the probability that an individual moves one deme in each adjacent direction per generation (baseline 0.10, measured effective size 1,434 diploids; varied over 0.05, 0.08, 0.12 and 0.15). Migration is rejected and redrawn if it moves a person off the grid.
- $\gamma$ is the assortative-mating discount applied to cross-lineage pairings, which is the knob calibrated so that autosomal ancestry matches the data (baseline 0.12; varied over 8 values from 0.08 to 1.00, where 1.00 is random mating).
- r is the fitness of a modern-lineage individual relative to an archaic-lineage one competing for the same deme (baseline 2; varied over 1.5, 2 and 3).

**Variation of residence and wave structure parameters**

- Lineage determines whether a cross-lineage child joins its mother's group, its father's, or either at random.
- female_bias is the continuous form of the probability that such a child takes its mother's group, with 1.0 always determined by the mother, 0.5 unbiased, and 0.0 always determined by the father.
- wave2_gen is the generation at which a second, fresh modern group is founded under the opposite residence rule for the two-wave model. It is set to 500 for the deme results in Figure S3.2 and to 200, 250 or 300 for the band results, which expand more slowly, and to 0 for the single-wave scenarios.

**Variation of mechanism parameters**

Each of the mechanism parameters is set to zero or off unless the mechanism being tested requires it, and each defines a mechanism rather than perturbing one:

- s with sel_locus is the additive selection coefficient against the disfavored allele and the locus it acts on. In Scenario 1 it acts on one of the reserved autosomal loci, as independent positive selection on the modern partner of a DMI pair. In Scenario 2 it acts on the Y chromosome, and is conditioned on the child's group label. Thus, a son whose Y chromosome matches his group label is unaffected (this corresponds to a modern Y chromosome in a modern-lineage son, an archaic Y chromosome in an archaic-lineage son). A son whose Y mismatches his group label is removed with probability s.
- xdrive with xdrive_cost is the fertility cost to expressing males and xdrv_res specifies heritable resistance (Scenario 1).

- dm with dm_focal, dm_partner, dm_dom and dm_male_limited specify a DMI pair, which loci it joins, dominance, whether it acts only in males, and with restorers for the polygenic mitonuclear version and esc_eps for the escape stress test (Scenario 1).
- $\theta$ is the relative fertility of F1 males, which is cultural rather than genetic (Scenario 4).
- s_mt is an independent selection coefficient on mtDNA, kept separate from s so that a run can carry selection on both markers at once. Unlike s on the Y, s_mt is unconditional: it represents the biological fitness gap of Supplementary Information section 1, which does not depend on a group label.

**Figure S3.5. Sensitivity of the deme framework to each demographic parameter.** Each column varies one parameter across its tested range while holding all others at the baseline marked in a gray vertical line (grid 12, $K = 11$, $m = 0.10$, $\gamma = 0.12$, $r = 2$, group label always determined by the mother), with no mechanism acting ($s = 0$, $\theta = 1$). Rows give the modern proportion at the autosomes, at mtDNA and at the Y, and the number of generations to extinction of individuals with an archaic group label (2,000 replicates per point). mtDNA is at 100% throughout because the expanding matriline carries its own mtDNA.

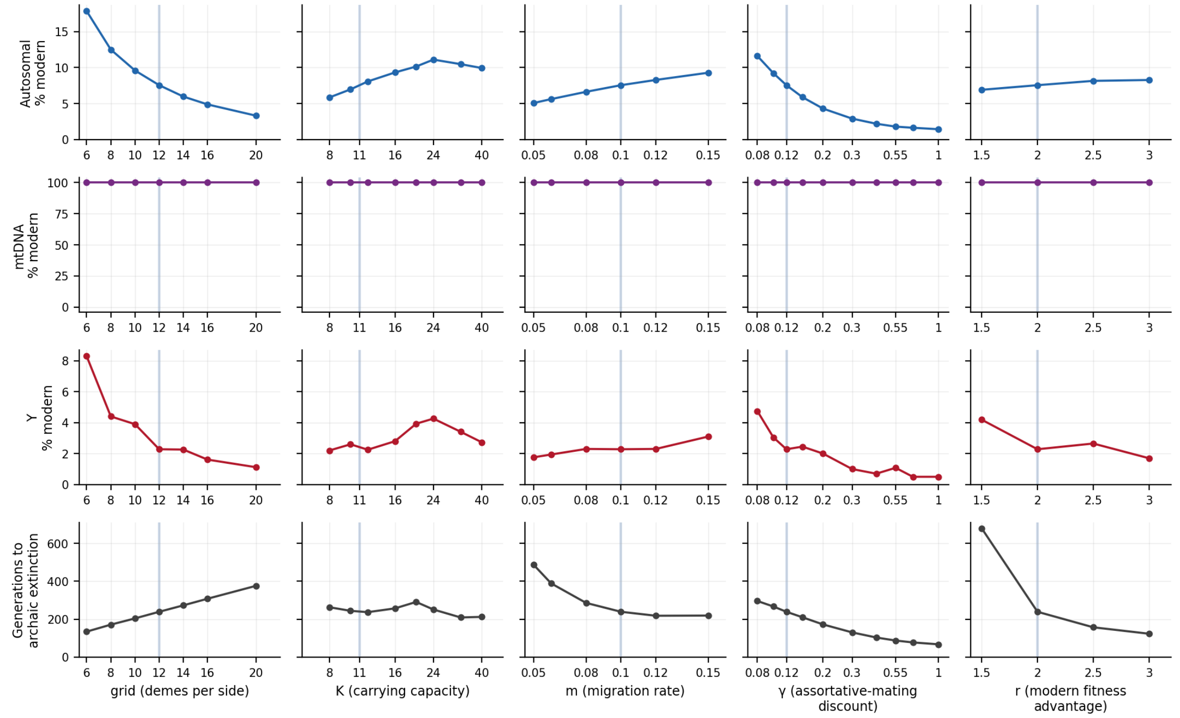


**Founding conditions and sampling, all fixed**

The modern lineage is founded in a single corner deme (seed_block = 1). The origin is re-founded if it loses its modern occupants, but only while archaic individuals remain anywhere on the grid (reseed_origin_after_extinction = false). Runs are sampled at generation 6,993, or as soon as both uniparental markers are absorbed if that comes first.

**Additional parameters built into simulations, some used to explore cultural scenarios**

Several of these were explored as candidate cultural mechanisms and are reported in the text as scenarios that failed.

- K_early and K_early_gens set a carrying capacity that contracts during the run. Neither is used in any result reported in this note: every cell of every figure and table runs at a constant carrying capacity, and the parameters are documented only because the engine accepts them.
- m_male and m_female set sex-specific migration rates.
- rho is a rule returning males to their father's natal deme.
- inbreed_ban captures inbreeding avoidance on the female relative side.
- cycle_T, cycle_R and cycle_phase correspond to periodic fluctuation in size.
- gamma_export and gamma_import are separate discounts for the two cross-mating directions.
- skew_frac is male reproductive skew.

**Appendix B: Roving band simulations**

The deme model is unrealistic in multiple ways, and I sought to assess the robustness of its inferences by using a simulation setup that was qualitatively different. Limitations of the deme framework included: (a) it made no attempt to model the geography of Europe, (b) it constrained everyone to a lattice, and (c) it did not capture the reality that hunter-gatherers are likely to have traveled in bands all with the same culture (modern or archaic group label).

To address these limitations, I designed a "roving band" simulation, in which every individual belongs to a band whose members live and move together and have a culture (modern or archaic) shared by every member. Bands have an optimal size K, and a position in continuous space at every generation. Bands fission on reaching twice their optimal size, splitting into two bands of about K, and disappear when they fall to nothing. A global fertility rescaling holds the total population constant. The baseline simulation setup is 70 bands of K = 25, leading to a census of 1,750 which, as in the deme model, I tuned to match the long-term effective population size. This is the size at which the measured autosomal drift matches the same PSMC anchor of 1,447 diploids (this is larger than the deme framework's 1,584 because the two frameworks are matched on effective size, not census). The optimal band size K = 25 is inspired by the literature: 25 has been argued to be reasonable for a co-resident band[45], and surveys have found a median of 27 across 263 forager societies[46] and a mean experienced band size of 28.2 adults across 32 societies[44].

**Two movement processes, tracked separately.** A whole band relocates with probability $p_{move}$ = 0.45 per generation, taking a Gaussian step of scale $\sigma_b$, with a baseline of 60 km. Independently, individuals leave their band and join another (exogamy) at per-capita rate e, diffusing a Gaussian distance of scale $\sigma_e$, with a baseline of 80 km, and joining whichever band is nearest where they land. Both scales are varied in the sensitivity analysis: $\sigma_b$ over 20, 30, 40, 60, 90, 120 and 180 km, and $\sigma_e$ over 30, 40, 50, 80, 120, 160, 180 and 240 km (Figure S3.6).

**Sex bias.** Sex-biased dispersal is parameterized as sex-specific per-capita rates of movement. Writing β for the female share of exogamy propensity, the rates are $2e\beta$ for women and $2e(1 - \beta)$ for men, so the mean per-capita rate is e whatever the value of β.

**Group label is carried by the band (exogamy leads to switching bands).** The group label, modern or archaic, belongs to the band, not the individual, so a migrant of the other group assimilates, contributing its genes to the local group. Exogamy therefore plays the role that γ and the female-bias parameter play in the deme model, and e is the calibrated knob.

**Density regulation.** A band's expected fertility is multiplied by $\exp(\alpha(1 - L/\bar{L}))$, where $\alpha$ is the strength of density regulation, L is the kernel-weighted local population around the band, and $\bar{L}$ its current mean across occupied space. Without this term, bands random-walk into clumps and the rest of the landscape empties. The global rescaling fixes how many people there are, and this local term fixes where they are. Bands with the modern group label have relative fitness r, which enters the same multiplier as a factor on the archaic side: an archaic band's fertility is multiplied by $\exp(\alpha(1 - L/\bar{L}))/r$, while a modern band's is multiplied by $\exp(\alpha(1 - L/\bar{L}))$, so the advantage is expressed against neighbors rather than globally. $\alpha$ has a baseline of 0.8, so a band sitting at twice the mean local density has its expected fertility multiplied by about 0.45 at the baseline. L is computed by weighting every band by a Gaussian kernel of scale $\sigma_e$ centered on it (the same 80 km scale that governs exogamous dispersal), so near neighbors count fully and distant ones hardly at all. $\bar{L}$ is the mean of L across occupied bands in the current generation, which makes the term relative rather than absolute. Because the global rescaling holds the total population constant, modern bands expand by taking a larger share of a fixed total.

**Transmission.** This is identical to that in the deme model of Appendix A.

**Range expansion.** Archaic bands are placed at random inside the habitable area with sizes drawn as Poisson(K), and a single modern band is seeded at the present-day location of Istanbul, on the European shore of the Bosphorus (41.05° N, 28.95° E). The lineage advantage r drives the expansion, and as in the deme model, the seed is re-founded if bands with a modern group label are lost entirely while archaic bands remain.

**Simulations of a second wave.** For the two-wave scenario, the landscape is relabeled at a chosen generation so every surviving individual counts as resident, whatever it was before, and a fresh modern group is founded at the origin. Only group label is reset; the residents keep the alleles the first wave left them. After a first wave in which females always stay with their group, the modern mtDNA is retained while their Y is archaic. In the second wave, males always stay within their group, and this spreads modern Y chromosomes.

**Simulations of a migration pulse.** The pulse in the band framework is described in Supplementary Information section 1, Appendix C.

**Restricting to parameter combinations that match the data.** As in the deme model, for many simulations, I tune the parameters to match the effective population size from PSMC, the duration of the range expansion (100-1,000 generations), and the probability that the range expansion proceeds rather than stalls (at least 50%).

**How the European landscape was generated.** The band model needs a bounded arena, and the maps in Figure 3a are drawn on it. Its outline follows a published reconstruction of the Neandertal niche[16]. I made three judgements to simplify the range definition for simulation purposes. First, where the niche model includes areas with little or no Neandertal fossil record (the east European plain and the Pontic steppe), I reduced the size of the range by a smooth curve along the northern limit together with a straight chord from the base of the Carpathians. Second, I excluded Crimea although Neandertals are believed to have lived there. Third, I hand-parameterized the interior boundary to the published figure. The map is plotted in Figure 3a, and the exact coordinates of the landscape I use are specified in Supplementary Table 1, in the tabs "Landscape definition", which lists the rules and their coordinates in the order they are applied, and "Landscape outline", which gives the resulting boundary as longitude and latitude.

**Projection and symbols used in the maps of Figure 3a.** The maps are drawn in an Albers equal-area conic projection with standard parallels at 38° and 52° N and origin at 17° E, 43° N, so no region is visually inflated relative to another. The mask itself is stored on the latitude/longitude lattice and is projection-free.

**Variation of demographic parameters**
As in the deme model, I evaluate how far the conclusions depend on choice of parameters:

- $N$ is the total census, held at 1,750 diploid individuals because that is the size at which the measured autosomal genetic drift matches the effective population size from PSMC.
- $K$ is the optimal band size, with a band fissioning when it reaches 2K and disappearing if it falls to nothing (baseline 25 gives about 70 bands).
- $e$ is the mean per-capita exogamy rate. This is the calibrated knob of this framework, playing the role $\gamma$ plays in the deme model (0.19 for an expansion where females always stay within the group; varied over 7 values from 0.08 to 0.56).
- $r$ is the fitness of a band with a modern group label relative to an archaic one (baseline 2; varied over 1.5, 2 and 3).
- $\alpha$ is the strength of local density regulation (baseline 0.8; sweep over 0.4, 0.8 and 1.6). A band at twice the mean local density has its expected fertility multiplied by $e^{-\alpha}$, about 0.45 at the baseline.
- $\beta$ is the female share of exogamy propensity and parameterizes sex bias. $\beta = 0$ when females always stay within the group and males disperse, and $\beta = 1$ when males stay within the group and females disperse.

**Variation of movement parameters**

- $\sigma_b$ is the scale of a band's Gaussian relocation step (baseline 60 km; varied over 20, 30, 40, 60, 90, 120 and 180 km).
- $p$ is the probability that a band relocates at all in a given generation (0.45, not varied).
- $\sigma_e$ is the scale over which exogamous individuals disperse to join the nearest band (baseline 80 km; varied over 30, 40, 50, 80, 120, 160, 180 and 240 km). It is also the kernel width used to compute local density, thus governing both gene flow between bands and competition.

I do not vary p independently of $\sigma_b$. Over many generations, only the diffusion coefficient $p\sigma_b^2$ matters, and simulations holding that product fixed at 1,620 while varying p seven-fold over 0.13, 0.45 and 0.90 give autosomal ancestry and median sweep completion that hardly change. I therefore hold *p* at 0.45 and treat $\sigma_b$ as the single movement parameter.

Movement is rejected and randomly redrawn if it would place a band outside the habitable mask (whether into the sea or beyond the modeled range, so the arena reflects rather than absorbs at its edge). The same applies to exogamous dispersal.

**Variation of mechanism parameters**
Nearly all of the mechanisms of the deme model are implemented in this framework as well, so each route to matching the observed data can be tested under both idealizations of space. Each is zero or off unless the mechanism being tested requires it.

- s is the additive selection coefficient on the Y, conditioned on the band's group label; a son with a Y chromosome mismatching the band he is born into (an archaic Y in a band with a modern group label, or a modern Y in a band with an archaic group label) is removed with probability s, while a son whose Y matches his band's group label is unaffected (Scenario 2).
- xdrive is X-linked drive, with xdrive_cost for the fertility cost to expressing males and xdrv_res for heritable resistance (Scenario 1).
- dm is the Dobzhansky-Muller pair, with dm_focal, dm_partner, dm_dom and dm_male_limited specifying which loci it joins, dominance, and whether it acts only in males. esc_eps is the escape stress test (Scenario 1).
- restorers specifies the number of polygenic mitonuclear restorer loci (Scenario 1).
- θ is the relative fertility of F1 males, which is cultural rather than genetic (Scenario 4).
- s_mt is an independent selection coefficient on mtDNA, kept separate from s so that a run can specify selection on both markers at once. Unlike s on the Y, s_mt is unconditional: it represents the biological fitness gap of Supplementary Information section 1, which does not depend on a group label.

**Figure S3.6. Sensitivity of the band framework to each demographic parameter**. The baseline is 70 bands of K = 25 at a census of 1,750, exogamy rate e = 0.19, β = 0 (females stay within the group and males disperse), r = 2, α = 0.8, band step $\sigma_b$ = 60 km and exogamous dispersal $\sigma_e$ = 80 km, with no mechanism acting (2,000 replicates per point, matching the deme framework). The duration row is the median generation at which the uniparental markers are absorbed, the band framework's analogue of the deme framework's time to archaic extinction.

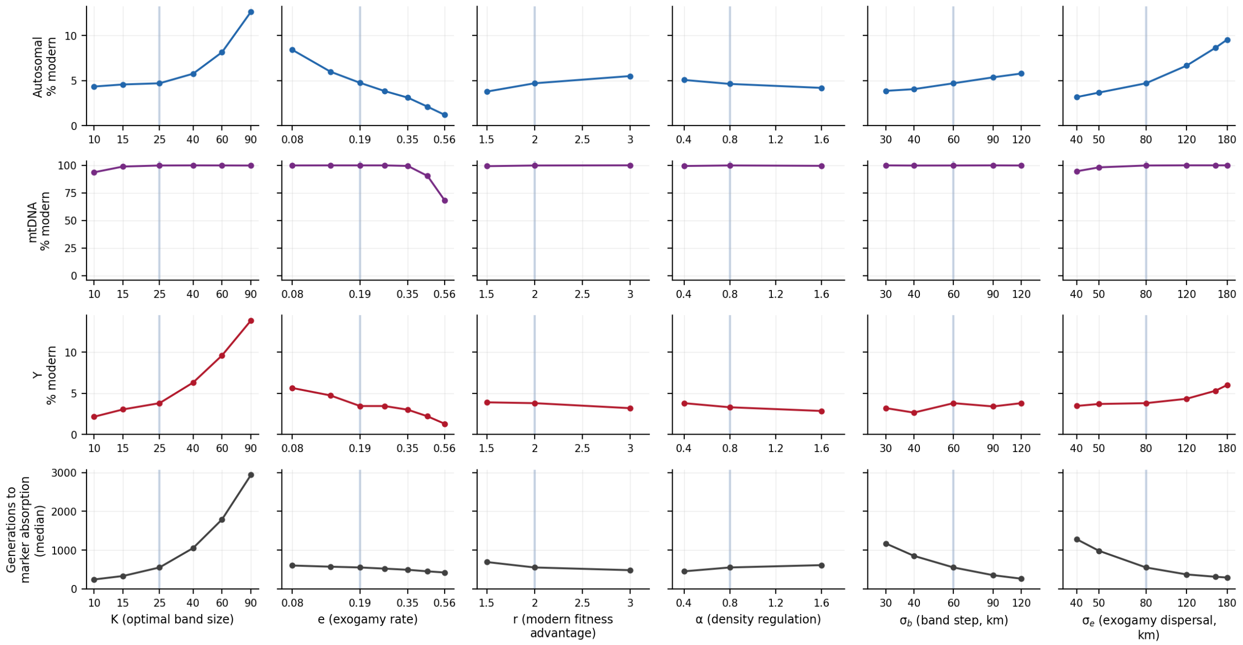


**Founding conditions and sampling, all fixed**

Archaic bands are placed at random inside the habitable area with sizes drawn Poisson(K), and the single modern band is founded at the European shore of the Bosphorus at the site of present-day Istanbul (41.05° N, 28.95° E). As in the deme model, the seed is re-founded if the bands with a modern group label are lost entirely while archaic bands remain, and never afterwards. For the migration-pulse comparator, pulse_frac (0.05) sets the fraction of individuals whose genome is replaced and pulse_beta its sex composition.

**Additional parameters built into simulations**
As in the deme model, I carried out a variety of simulations to explore candidate cultural mechanisms, and all of these are reported in the text as ones that failed.

- rho_return is a rule returning dispersers to their parent's natal band.
- inbreed_ban is inbreeding avoidance within a band.
- stable_boundary holds the two group labels at a frontier instead of letting one expand.
- hard_selection is selection that reduces the total population rather than rescaling away.
- skew_frac is male reproductive skew.
- e_export and e_import separate exogamy rates for the two interbreeding directions (modern to archaic, or archaic to modern).

**Software and Reproducibility**

A zipped folder associated with this note (SI3codebase.zip) contains all of the code needed to reproduce every simulation result reported here, using Python 3.9 or later. requirements.txt lists the dependencies and README.md gives the script-by-script pipeline together with the exact commands used to generate each reported number. The folder contains only code that produces a number, figure or table in this note. Every file in the archive is named below, so a reader can go from any figure to the code that drew it.

**Engines.** spatial_sim2.py is the deme range-expansion engine of Appendix A, in which gene flow occurs through explicit matings rather than through group-label-switching of adults, a child's lineage follows its mother, its father, or a random parent, and the simulation tracks mtDNA, the Y, 36 unlinked autosomes and three X loci under both additive selection and the general DMI rule, in which an offspring carrying a modern allele at a focal locus together with an archaic allele at a partner locus is removed, optionally only in males. spatial_sim3.py extends it with the mechanism loci, every new parameter defaulting off so that the engine is otherwise identical. bandsim.py is the roving-band engine of Appendix B, and make_range_mask.py builds its habitable arena from the published niche reconstruction. panmictic_sim.py is the migration-pulse comparator: no geography, a single instantaneous 5% pulse into a constant-size population reproducing under a Wright-Fisher model with separate sexes; the model it implements is set out in Supplementary Information section 1, Appendix C. ysupp_sim.py is the closed-population model of an X-linked distorter, a Y-linked suppressor and a *de novo* mutation rate on the Y, which is the model behind the Scenario 1 statement about suppressor mutations; it has no geography, and neither spatial engine can express it, because in both a suppressing Y is a lineage label fixed at initialization rather than something that can arise.

**Anchoring and calibration.** digitize_psmc.py digitizes the PSMC trajectories and takes the time-weighted harmonic mean that gives the 1,447-diploid anchor for the effective population size estimate, and anchors.py holds that figure and the censuses derived from it as the single source of truth the runners import. ne_screen.py measures the realized effective size of every (grid, K, m) triple in the deme framework and writes ne_screen.csv, the table that decides which geometries are admissible. On the band side, band_ne.py measures the realized uniparental effective size by the same neutral-tracer method, band_cal.py fits the exogamy rate that puts autosomal ancestry on target, bandcal_submit.py runs that calibration on the cluster.

**The Cartesian search.** This is the pipeline behind main-text Figure 3b, Figure S3.2 and Extended Data Figure 4. routes_build.py defines the mechanism sets and the demographic grid; routes_submit.py writes the main job array, rescreen_submit.py the 100-replicate re-screen of cells whose 10-replicate estimate is too close to the establishment threshold to resolve, cultural_submit.py the five cultural mechanisms, xdrvdeme_submit.py the anchored X-drive ladder with the assortative-mating discount refitted at every rung, and nexp_band_submit.py with demog_fill_submit.py the unbiased-expansion series. All of them emit tasks for robust_run.py, the worker that runs a single cell in either framework at either stage, with robust_submit.py as the generic array writer and robust_collect.py pooling the per-cell output. screen.py holds the single definition of admissibility—autosomal ancestry 3-7%, expansion completing in 100-1,000 generations, establishment above one half, decided once from the best estimate available for each cell—so that Figure 3b, Figure S3.2 and Extended Data Figure 4 cannot disagree about which cells they are drawing. Scenario 3, the two-wave scenario, has its own runners within the same screen: twowave_run.py and twowave_band.py run it in the two frameworks, tw5_run.py with tw5_submit.py score the grid, and tw6_run.py with tw6_submit.py rerun the band arm timed on the generation at which the uniparental markers are absorbed rather than on the disappearance of the last archaic-labelled band, with clock_check.py the comparison that establishes why those two clocks differ for a two-wave scenario and agree for a single wave.

**The migration pulse.** fixation_curves_mtdna.csv is the table of fixation probability against selective advantage and mtDNA copy number that Figure S3.4 reads; it is produced by the code base of Supplementary Information section 1, where the migration-pulse model is described (Appendix C there), and is shipped here as an input. panm4_run.py, panm4_submit.py and panm4_collect.py run the sex-biased pulse and the stable-boundary comparator behind Figure S3.1's left panel. The joint two-marker pulse is joint_run.py with joint_submit.py and joint_collect.py, extended by joint2_submit.py and joint2_collect.py and calibrated by jcal_submit.py and jcal_collect.py; mtypulse_submit.py runs the mtDNA-by-Y pulse, twpulse_submit.py the two-wave pulse and jdrive_submit.py the pulse carrying X-linked drive, all three of which Figure 3b draws as demographic comparison lines. mtexp_submit.py runs modern mtDNA under scenario where males always stay within their group and females disperse, which collect_mt_expansion.py tabulates as the range-expansion arms of Figure S3.4.

**Sweeps read directly by a figure.** grid5_run.py with grid5_submit.py and grid5_collect.py sweep the residence rule for Figure S3.1's right panel. scan4_run.py with scan4_submit.py and scan4_collect.py vary each demographic parameter in turn about the deme baseline and write figure3_full.csv, the source of Figure S3.5. xdrv2_run.py with xdrv2_submit.py and xdrv2_collect.py run the heritable-resistance blocks from whose raw per-seed output Figure S3.3's right panel is recomputed. Two further analyses are reported in the text without a figure: sf1_run.py with sf1_submit.py and sf1_collect.py test the counterpart of the Scenario 1 incompatibilities where males stay within the group and females disperse, in which a modern autosomal partner locus is incompatible with archaic mtDNA and is also under independent positive selection, and esc_run.py with esc_submit.py and esc_collect.py run the heritable-escape stress test of the mtDNA-by-Y incompatibility.

**The roving band pipeline.** band_reps.py is the band worker, running one condition for many replicates, and band_suite.py holds the constants the band scripts and figures share. bandsi_submit.py with bandsi_collect.py run and pool the sensitivity, residence-bias, pulse-bias and mtDNA-curve families into bandsi_summary.csv; bandmech_submit.py with bandmech_collect.py do the same for the mechanism sweeps into bandmech_summary.csv. bandprod_submit.py with bandprod_collect.py fit the exogamy rate per scenario and write bandprod_calibration.csv, the authority for e* that bandsi_submit.py and the Figure 3a script both read, and bandratchet_submit.py with bandratchet_collect.py run the production ensemble under the Muller's-ratchet posterior from which the replicate shown in Figure 3a was chosen. Every mechanism of the deme model is implemented in the band engine as well, except those requiring locus-specific positive selection or sex-specific migration, which it does not express.

**Figures.** figure2_full.py assembles main-text Figure 3 from its two panels. Panel (a) is drawn by figure2_bands.py, with figure2_seedscan.py choosing the replicate shown and site_stripes_figure.py the equal-area projection; panel (b) is drawn by figure2_merged.py, with figure2c_fountain.py supplying the admissible Cartesian combinations both panels use. make_si_figures.py draws both panels of Figure S3.1. Figure S3.4 is drawn by fig_fixation_gaps.py, from fixation_curves_mtdna.csv, the band pulse arm of bandsi_summary.csv and the range-expansion arms that collect_mt_expansion.py tabulates. fig_thumbnails_all.py, with figED_thumbnails.py setting the block order and the scenario numbering, draws Figure S3.2, fig_s34_drive_and_escape.py draws Figure S3.3, and fig_sensitivity.py draws Figures S3.5 and S3.6. The Extended Data figures are figED_sensitivity.py, which stacks S3.5 and S3.6 as Extended Data Figure 3, and figED_expansions.py, which draws the sample expansions of Extended Data Figure 2; Extended Data Figure 1 is Figure S3.1 and Extended Data Figure 4 is Figure S3.2, in each case the same figure rather than a redrawing. figure2a_movie.py renders the supplementary video animating Figure 3a: the same replicate as four published panels, at one frame every four generations, so the stills are frames of the film rather than a separate run.

**Analysis, checking and reproducibility.** collect_mt_expansion.py writes mt_expansion_curve.csv, and build_supp_workbook.py assembles Supplementary Table 1 from the simulation output, so that every tab of the workbook is regenerated rather than transcribed; it reads the per-cell simulation summaries through summary_cache.py, which gathers them once rather than file by file. audit_v66.py recomputes every simulation number this note states from the files on disk and prints what agrees and what does not, and reps_audit.py recounts the replicates behind every figure in which the two frameworks are drawn together. The *_submit.py scripts write the Harvard Medical School O2 SLURM job arrays used to produce the results reported here; the underlying workers take the same arguments and can equally be run on a single machine. Every summary table the figure scripts read is regenerated by the corresponding collect script, and the inputs a single run does not itself regenerate are shipped with the code, namely ne_screen.csv, fixation_curves_mtdna.csv, mt_gap_posterior_pooled_rna.txt, range_mask.npz and bandprod_calibration.csv. Two of those come from the code base of Supplementary Information section 1 rather than from this one: fixation_curves_mtdna.csv and mt_gap_posterior_pooled_rna.txt, the RNA-inclusive posterior draws of the mtDNA fitness gap that joint_run.py samples. One script is an exception and needs a file from outside this archive: digitize_psmc.py reads the published figure from which the PSMC trajectories are digitized. It is not needed to redraw any figure here, because the quantity it produces — the 1,447-diploid effective population size anchor — is already embedded in the scripts and data that ship. Replicates are seeded deterministically, so any reported value can be reproduced exactly.
