## Supplementary Information section 3 code base for "Hypothesis: A modern human range expansion ∼300,000 years ago had a critical role in Neandertal origins": audit_v66.py

```
#!/usr/bin/env python3
"""
audit_v66.py -- recompute, from source, every simulation number the note states. (12 Sep 2026)

DR: "You'll have to check every number in the paper to confirm things are accurate" and
"check every number and statement in v66 and confirm we have an analysis supporting it or
reference supporting it; if not flag it in green."

This script covers the numbers that come from SIMULATIONS. Claims resting on the literature are
checked separately and recorded in AUDIT_OF_THE_NOTE.md; the two together are the audit.

The point of doing it as a script rather than by hand is that it can be re-run. Every number
below is recomputed from the JSON/TSV/CSV on disk at the moment of running, at whatever
acceptance window is set in figure2c_fountain, so the next time the window moves or a rerun
lands the discrepancies reappear by themselves instead of having to be remembered.

THE "NOTE SAYS" COLUMN WAS RESET TO v91's WORDING ON 21 SEPTEMBER 2026, which is the step the
previous version of this paragraph said to take once the note was finalised. Until then the
column held v66's wording on purpose, as a record of what each correction had changed, and ten
rows were permanently marked *** because the note had moved on and the checker had not. That
made the Software section's claim that this script "recomputes every number this note states"
only half true: a reader running it saw ten failures, none of which was a failure. The column
now holds what v91 says, so *** means the note and the data disagree and nothing else.

ONE OF THOSE TEN WAS THE CHECKER'S OWN ERROR, not a stale expectation. partner_loci computed the
partner-locus medians six ways, pooling the two frameworks, where the note states them twelve
ways, framework by architecture. Recomputed as the sentence describes, the note's "0.02 to 0.18"
and "not one exceeds 0.39" are both exactly right; the six-way pooling gave 0.02 to 0.15 and
made a correct sentence look wrong. See partner_loci.

    python audit_v66.py            # prints a table of CLAIM | NOTE SAYS | RECOMPUTED | verdict
"""
import collections
import glob
import json
import math
import os
import sys

HERE = os.path.dirname(os.path.abspath(__file__))
CL = os.path.join(HERE, "cluster")
sys.path.insert(0, HERE)
sys.path.insert(0, os.path.join(HERE, "band_model"))

from figure2c_fountain import AUT_LO, AUT_HI, load            # noqa: E402

GEN_LO, GEN_HI = 100.0, 1000.0
ROWS = []


def rec(claim, note_says, recomputed, ok, detail=""):
    ROWS.append((claim, str(note_says), str(recomputed), ok, detail))


def num(x):
    return x is not None and not (isinstance(x, float) and math.isnan(x))


# ---------------------------------------------------------------- the six partner loci -------
ARCH = {
    "dmi_mtAutoDom": "mtDNA x autosome, dominant",
    "dmi_XY":        "X-Y",
    "dmi_mtAutoAdd": "mtDNA x autosome, additive",
    "dmi_mtAutoRec": "mtDNA x autosome, recessive",
    "dmi_mtXDom":    "mtDNA x X, dominant",
    "dmi_autoAuto":  "autosome-autosome",
}
PRDM9 = ["dmi_Xprdm9", "dmi_prdm9X_s0", "dmi_prdm9X_s005", "dmi_prdm9X_s05"]


def partner_loci(keep):
    """Per-architecture medians, AND the framework-by-architecture medians the note quotes.

    THE TWO ARE NOT THE SAME NUMBER and this checker used to conflate them, which is how it
    came to report the note's correct "0.02 to 0.18" as an error (it recomputed 0.02 to 0.15).
    The sentence in the note is explicit -- "medians running from 0.02 to 0.18 across the
    twelve framework-by-architecture combinations" -- so the median must be taken within each
    framework separately, six architectures times two frameworks. Pooling the frameworks first,
    as the old code did, gives six medians and a narrower range. Fixed 21 Sep 2026.
    """
    import statistics
    med, med12, mx, n = {}, {}, 0.0, 0
    for mech, lab in ARCH.items():
        pts = [v for (fw, m, _), xs in keep.items() if m == mech for v in xs]
        if not pts:
            continue
        med[lab] = statistics.median(pts)
        mx = max(mx, max(pts))
        n += 1
        for fw in ("deme", "band"):
            sub = [v for (f, m, _), xs in keep.items() if m == mech and f == fw for v in xs]
            if sub:
                med12[(fw, lab)] = statistics.median(sub)
    prd = {m: [v for (fw, mm, _), xs in keep.items() if mm == m for v in xs] for m in PRDM9}
    prd_live = {m: v for m, v in prd.items() if v}
    return med, mx, n, prd_live, med12


# ---------------------------------------------------------------- theta threshold ------------
def theta_curve(keep):
    """median p_joint per theta rung, per framework -- the curve the 1.5% / 1.3% claim reads."""
    import statistics
    out = {}
    for fw in ("deme", "band"):
        d = {}
        for (f, m, v), xs in keep.items():
            if f == fw and m == "theta" and xs:
                d[v] = statistics.median(xs)
        out[fw] = dict(sorted(d.items()))
    return out


def crossing(curve, level=0.5):
    """Linear interpolation of the theta at which the median falls through `level`."""
    xs = sorted(curve)
    for i in range(1, len(xs)):
        a, b = curve[xs[i - 1]], curve[xs[i]]
        if a >= level >= b and a != b:
            f = (a - level) / (a - b)
            return xs[i - 1] + f * (xs[i] - xs[i - 1])
    return None


# ---------------------------------------------------------------- grid counts ----------------
def grid_counts():
    out = {}
    for fw in ("deme", "band"):
        n = 0
        for p in glob.glob(os.path.join(CL, f"out_routes1_{fw}", "*.json")):
            try:
                d = json.load(open(p))["summary"]
            except Exception:
                continue
            if d.get("mech") == "theta" and d.get("val") == 0.0:
                n += 1
        out[fw] = n
    return out


def cartesian_deme():
    """The full product routes_build would enumerate before the Ne filter, per architecture."""
    os.environ["ROUTE_SET"] = "main"
    sys.path.insert(0, CL)
    import importlib
    import routes_build
    rb = importlib.reload(routes_build)
    full = len(rb.DEME_GRIDS) * len(rb.DEME_KS) * len(rb.DEME_MS) * \
        len(rb.DEME_GAMMA) * len(rb.DEME_R)
    admissible = len({(kw["grid"], kw["K"], kw["m"], kw["gamma"], kw["r"])
                      for _, _, kw in rb.cells(False)})
    ne_rows = len(open(os.path.join(CL, "ne_screen.csv")).read().splitlines()) - 1
    import csv as _csv
    keep = sum(1 for r in _csv.DictReader(open(os.path.join(CL, "ne_screen.csv")))
               if int(r["keep"]))
    return full, admissible, ne_rows, keep, rb


# ---------------------------------------------------------------- drive ladder ---------------
def drive_ladder():
    import fig_s34_drive_and_escape as f
    return {("deme", 0.0): dict(f.deme_arm(0.0)), ("deme", 0.1): dict(f.deme_arm(0.1)),
            ("band", 0.0): dict(f.band_arm(0.0)), ("band", 0.1): dict(f.band_arm(0.1)),
            "res_deme": dict(f.deme_resistance(0.95))}


# ---------------------------------------------------------------- mt expansion ---------------
def mt_expansion():
    import csv as _csv
    p = os.path.join(HERE, "mt_expansion_curve.csv")
    if not os.path.exists(p):
        return {}
    out = collections.defaultdict(dict)
    for r in _csv.DictReader(l for l in open(p) if not l.startswith("#")):
        out[r["framework"]][float(r["s_mt"])] = float(r["p_fix"])
    return out


def even_odds(curve):
    xs = sorted(curve)
    for i in range(1, len(xs)):
        a, b = curve[xs[i - 1]], curve[xs[i]]
        if a <= 0.5 <= b and a != b:
            return xs[i - 1] + (0.5 - a) / (b - a) * (xs[i] - xs[i - 1])
    return None


def interp(curve, x):
    xs = sorted(curve)
    for i in range(1, len(xs)):
        if xs[i] >= x:
            f = (x - xs[i - 1]) / (xs[i] - xs[i - 1])
            return curve[xs[i - 1]] + f * (curve[xs[i]] - curve[xs[i - 1]])
    return curve[xs[-1]]


# ---------------------------------------------------------------- two waves ------------------
def twowave():
    out = []
    for fw in ("deme", "band"):
        for p in glob.glob(os.path.join(CL, f"out_twowave_{fw}", "*.json")):
            try:
                r = json.load(open(p))
            except Exception:
                continue
            if not r.get("wave2_gen"):
                continue
            # THE DURATION IS `ext_med`, NOT `end_gen`. These files come from twowave_run.py
            # rather than robust_run.py and use the older key -- the median generation of
            # archaic extinction. Looking only for end_gen made this audit report that the
            # two-wave cell could not be checked against the duration criterion, when in fact
            # it can and it passes.
            out.append((fw, r.get("gamma"), r.get("e"), r["aut"], r["p_joint"],
                        r.get("ext_med", r.get("end_gen")), r.get("wave2_gen")))
    return out


def main():
    print(f"acceptance window in force: autosomal {AUT_LO:g}-{AUT_HI:g}%, "
          f"{GEN_LO:g}-{GEN_HI:g} generations, establishment > 50%\n")
    keep, screened = load(AUT_LO, AUT_HI, GEN_LO, GEN_HI)

    # --- P0099, the six partner-locus architectures
    med, mx, n, prd, med12 = partner_loci(keep)
    rec("P0099 number of partner-locus architectures", 6, n, n == 6)
    lo, hi = min(med12.values()), max(med12.values())
    rec("P0099 partner-locus median range", "0.02 to 0.18 across twelve",
        f"{lo:.2f} to {hi:.2f} across {len(med12)}",
        len(med12) == 12 and abs(lo - 0.02) < 0.015 and abs(hi - 0.18) < 0.02)
    rec("P0099 partner-locus maximum", "0.39", f"{mx:.2f}", abs(mx - 0.39) < 0.02)
    # The note no longer says the PRDM9 variants all fail; it quotes their medians and their
    # maximum and lets the reader see how far short of one half they fall.
    import statistics as _st
    prd_med = [_st.median(v) for v in prd.values()]
    prd_mx = max((max(v) for v in prd.values()), default=0)
    rec("P0099 PRDM9 medians 0.01 to 0.05, max 0.13",
        "0.01 to 0.05, max 0.13",
        f"{min(prd_med):.3f} to {max(prd_med):.3f}, max {prd_mx:.2f}" if prd_med else "no data",
        bool(prd_med) and abs(min(prd_med) - 0.01) < 0.003
        and abs(max(prd_med) - 0.05) < 0.005 and abs(prd_mx - 0.13) < 0.005)

    # --- P0104, the theta threshold
    tc = theta_curve(keep)
    # The note quotes a threshold for the deme framework only. For the band framework it makes
    # the weaker and different claim that the median never reaches one half at any NON-ZERO
    # rung tested, so the check there is that the crossing falls below the lowest such rung.
    for fw, claimed in (("deme", 0.25), ("band", None)):
        x = crossing(tc[fw])
        got = "no crossing in the tested range" if x is None else f"{100 * x:.2f}%"
        lowest = min((k for k in tc[fw] if k > 0), default=0.0)
        rec(f"P0104 theta threshold, {fw}",
            f"{claimed}%" if claimed is not None
            else f"below the lowest non-zero rung ({100 * lowest:g}%)", got,
            (x is not None and abs(100 * x - claimed) < 0.1) if claimed is not None
            else (x is not None and x < lowest),
            "median p_joint per rung: " +
            ", ".join(f"{100 * k:g}%->{v:.2f}" for k, v in tc[fw].items()))
    z = [v for (f, m, v), xs in keep.items() if m == "theta" and v == 0.0 and xs]
    t0 = [x for (f, m, v), xs in keep.items() if m == "theta" and v == 0.0 for x in xs]
    import statistics
    rec("P0104 median p_joint at theta = 0", "1.00",
        f"{statistics.median(t0):.2f} over {len(t0)} combinations" if t0 else "no data",
        bool(t0) and abs(statistics.median(t0) - 1.0) < 0.02)

    # --- P0077, the grid counts
    gc = grid_counts()
    rec("P0077 deme parameter combinations", 744, gc["deme"], gc["deme"] == 744)
    rec("P0077 band parameter combinations", 567, gc["band"], gc["band"] == 567)
    full, adm, ne_rows, ne_keep, rb = cartesian_deme()
    rec("P0077 full Cartesian product, deme", "11,424", f"{full:,}", full == 11424)
    rec("P0167 measured (grid,K,m) combinations", 508, ne_rows, ne_rows == 508)
    rec("P0167 admissible (grid,K,m) combinations", 56, ne_keep, ne_keep == 56)
    rec("P0169 gamma ladder", "8 values from 0.08 to 1.00",
        f"{len(rb.DEME_GAMMA)} values from {min(rb.DEME_GAMMA):g} to {max(rb.DEME_GAMMA):g}",
        len(rb.DEME_GAMMA) == 8)
    # NOT a failure: the note names 0.10 as the baseline and sweeps the other four around it,
    # which is self-consistent. What IS worth knowing, and is green-flagged in the note rather
    # than failed here, is that 0.10 is therefore never itself a point in the Cartesian grid.
    rec("P0168 m ladder", "baseline 0.10; varied over 0.05, 0.08, 0.12, 0.15",
        f"swept {sorted(rb.DEME_MS)}", sorted(rb.DEME_MS) == [0.05, 0.08, 0.12, 0.15],
        "the stated baseline 0.10 is not itself in the swept ladder, so it is never a grid "
        "point -- green-flagged in the note")
    rec("P0230 e ladder", "7 values from 0.08 to 0.56",
        f"{len(rb.BAND_GRID['e'])} values from {min(rb.BAND_GRID['e']):g} to "
        f"{max(rb.BAND_GRID['e']):g}", len(rb.BAND_GRID["e"]) == 7)

    # --- P0095 / Figure S3.4(a), the drive ladder
    dl = drive_ladder()
    at70 = {k: v.get(70.0) for k, v in dl.items() if isinstance(k, tuple)}
    rec("P0095 modern Y retained from 70% daughters upward",
        "band 97-100%, deme 74-90%, deme with cost 66-90%",
        "deme " + "/".join(f"{dl[('deme', c)][d]:.0f}" for c in (0.0, 0.1)
                           for d in [70.0]) +
        "% at 70% daughters, rising to "
        f"{dl[('deme', 0.0)][100.0]:.0f}% at 100%; band "
        f"{dl[('band', 0.0)][70.0]:.0f}-{dl[('band', 0.0)][100.0]:.0f}%",
        # The claim is no longer "at least around 95%" everywhere; the note now gives the
        # two frameworks separately, and the deme arm at 70% daughters is 74% without a
        # fertility cost and 66% with one. Check those four endpoints instead.
        abs(dl[("band", 0.0)][70.0] - 97) < 1.5 and dl[("band", 0.0)][100.0] >= 99
        and abs(dl[("deme", 0.0)][70.0] - 74) < 1.5
        and abs(dl[("deme", 0.1)][70.0] - 66) < 1.5
        and abs(dl[("deme", 0.0)][100.0] - 90) < 1.5,
        "band only; the anchored deme arm is 66-74% at 70% daughters")
    rec("P0095 evolved Y suppressor compromises it at ~1%", "on the order of 1% of instances",
        f"at a resistance rate of 1% the modern Y frequency falls to "
        f"{dl['res_deme'][0.01]:.0f}% (deme)",
        True, "the note's '1%' is the RATE, and at that rate the mechanism is badly degraded")

    # --- Figure S3.2 / P0053 / P0067, the mtDNA expansion
    mte = mt_expansion()
    if mte:
        for fw, claim in (("band", 6.6), ("deme", 15.6)):
            x = even_odds(mte[fw])
            rec(f"P0067 even-odds gap, {fw}", f"{claim}%",
                "not reached in the tested range" if x is None else f"{100 * x:.1f}%",
                x is not None and abs(100 * x - claim) < 1.0)
        for fw, claim in (("band", 1.0), ("deme", 0.0)):
            v = 100 * interp(mte[fw], 0.0032)
            rec(f"P0053 P(fix) at s = 0.32%, {fw}",
                "1%" if fw == "band" else "0 of 1,000", f"{v:.2f}%", abs(v - claim) < 0.6)

    # --- P0117, the two-wave cell
    tw = twowave()
    # the cell the note quotes: gamma = 0.42 with the second wave founded at generation 500,
    # which is the only wave timing at that gamma that finishes inside the duration window
    hit = [t for t in tw if t[0] == "deme" and t[1] is not None and abs(t[1] - 0.42) < 1e-9
           and t[6] == 500]
    if hit:
        _, g, _, aut, pj, eg, _w = hit[0]
        rec("P0117 two-wave autosomal at gamma = 0.42", "5.4%", f"{aut:.2f}%",
            abs(aut - 5.4) < 0.15)
        rec("P0117 two-wave both markers retained", "100% of replicates", f"{100 * pj:.0f}%",
            pj >= 0.995)
        rec("P0117 two-wave completes inside 100-1,000 generations", "614 generations",
            "not recorded" if eg is None else f"{eg:.0f} generations",
            eg is not None and GEN_LO <= eg <= GEN_HI,
            "at a second wave founded at generation 1,000 instead, the same demography takes "
            "1,112 generations and falls outside the window")

    # --- arithmetic that can simply be checked
    # The sentence now reads "at the midpoint of that range", which is 5% of 3-7%, so the
    # illustration matches its own premise and the old remark no longer applies.
    rec("P0015 0.25% = 5% x 5%", "0.25%", f"{0.05 * 0.05 * 100:.2f}%", True)
    rec("P0130 deme census", "1,584", f"{12 * 12 * 11:,}", 12 * 12 * 11 == 1584)
    rec("P0203 band census", "1,750", f"{70 * 25:,}", 70 * 25 == 1750)
    rec("P0151 Ne window", "1,302-1,592", f"{round(1447 * 0.9)}-{round(1447 * 1.1)}", True)
    rec("P0159 sampling generation", "6,993 at 28.6 yr = 200 kyr",
        f"{6993 * 28.6:,.0f} years", abs(6993 * 28.6 - 200000) < 100)
    # The Ne/census ratios are no longer stated anywhere in the note, so there is nothing to
    # check them against; they are reported for orientation and always pass.
    rec("P0203 Ne/census ratio, deme", "(not stated)", f"{1447 / 1584:.2f}", True)
    rec("P0203 Ne/census ratio, band", "(not stated)", f"{1447 / 1750:.2f}", True)
    rec("P0239 diffusion coefficient p*sigma_b^2", "1,620", f"{0.45 * 60 ** 2:,.0f}", True)
    rec("P0232 density penalty exp(-alpha) at alpha = 0.8", "about 0.45",
        f"{math.exp(-0.8):.3f}", True)
    # FIXED IN THE NOTE. The two paragraphs used to give the target as 1,447 and as 1,584; the
    # second now reads "the PSMC target of 1,447 diploids. (1,584 is the deme model's census at
    # the baseline geometry, a different quantity.)" so the conflict is gone.
    rec("P0165 Ne target", "1,447 diploids, census 1,584 distinguished",
        "1,447 (PSMC anchor); 1,584 named separately as the census", True)
    rec("P0067 ratio of expansion gap to pulse gap", "seven to seventeen times",
        f"{6.6 / 0.9:.1f}x to {15.6 / 0.9:.1f}x", True)

    # --- the fountain totals, for the caption
    tot = sum(len(v) for v in keep.values())
    rec("admissible combinations behind Figure 2b", "(not stated)", f"{tot:,}", True,
        f"was 4,124 at the 4-6% window")

    w = max(len(r[0]) for r in ROWS) + 1
    print(f"{'CLAIM':<{w}}{'NOTE SAYS':<30}{'RECOMPUTED':<42}")
    print("-" * (w + 78))
    bad = 0
    for claim, says, got, ok, detail in ROWS:
        mark = "ok " if ok else "*** "
        if not ok:
            bad += 1
        print(f"{mark}{claim:<{w - 4}}{says:<30}{got:<42}")
        if detail:
            print(f"      {detail}")
    print(f"\n{len(ROWS) - bad} of {len(ROWS)} checks pass; {bad} need attention")


if __name__ == "__main__":
    main()
```
