## Supplementary Information section 3 code base for "Hypothesis: A modern human range expansion ∼300,000 years ago had a critical role in Neandertal origins": band_cal.py

```
#!/usr/bin/env python3
"""
band_cal.py -- automated e-calibration for the band model: find the exogamy rate e at which
final modern autosomal ancestry = 5%, for a given condition. Same discipline as the deme
model's gamma calibration. Secant search on log(e) between two anchors, then one confirm run;
results cached in band_cal.json keyed by the condition signature.

Used as a library by band_suite.py; standalone:
    python band_cal.py --theta 0.05 --s 0.01 [--reps 24] [--target 0.05]
"""
import argparse
import json
import os
import subprocess
import sys

import numpy as np

HERE = os.path.dirname(os.path.abspath(__file__))
CACHE = os.path.join(HERE, "band_cal.json")


def _run_aut(flags, reps, procs, gens=800):
    """one calibration probe: returns mean final modern autosome fraction"""
    import csv as _csv
    tag = "cal_" + "_".join(f"{k.strip('-')}{v}" for k, v in sorted(flags.items()))
    path = os.path.join(HERE, f"band_reps_{tag}.csv")
    if not os.path.exists(path):
        cmd = [sys.executable, os.path.join(HERE, "band_reps.py"), "--reps", str(reps),
               "--gens", str(gens), "--procs", str(procs), "--tag", tag]
        for k, v in flags.items():
            cmd += [k, str(v)]
        subprocess.run(cmd, check=True, capture_output=True, cwd=HERE)
    with open(path) as f:
        rows = [r for r in _csv.DictReader(f) if r["died"] == "0"]
    est = [r for r in rows if float(r["sweep_gen"]) >= 0]   # condition on establishment
    rows = est if len(est) >= max(6, len(rows) // 3) else rows
    return float(np.mean([float(r["auto"]) for r in rows]))


def calibrate_e(cond_flags, reps=100, procs=4, target=0.05, verbose=True, e_guess=None,
                probes=(0.30, 0.40, 0.52)):
    """cond_flags: band_reps flags EXCLUDING --e. Three FIXED probes at `probes`, then
    log-linear least-squares fit of autosomal ancestry on log(e), inverted at `target`.
    Deterministic cost, no tolerance-chasing, and every probe informs the answer -- this
    replaces the secant search, whose stopping rule made e* jump around under probe noise.
    Matches the deme series' discipline (recalibrate per condition), just with a stabler
    estimator. Cached in band_cal.json."""
    key = json.dumps(sorted(cond_flags.items())) + f"|{target}|v2"
    cache = json.load(open(CACHE)) if os.path.exists(CACHE) else {}
    if key in cache:
        return cache[key]
    # beta = 0.5 dilutes ~3x faster per exogamy event, so the 5% point sits much lower;
    # start from a bracket appropriate to the scenario rather than extrapolating.
    if float(cond_flags.get("--beta", 0.0)) >= 0.25:
        probes = (0.08, 0.13, 0.22)
    xs, ys = [], []

    def probe(e):
        a = _run_aut({**cond_flags, "--e": round(e, 3)}, reps, procs)
        xs.append(np.log(round(e, 3))); ys.append(a)
        if verbose:
            print(f"    cal probe e={e:.3f} -> aut={100*a:.2f}%", flush=True)

    for e in probes:
        probe(e)
    for _round in range(2):
        b, c = np.polyfit(xs, ys, 1)
        if b >= -1e-6:
            e_star = float(np.exp(xs[int(np.argmin(np.abs(np.array(ys) - target)))]))
            break
        e_star = float(np.exp((target - c) / b))
        lo, hi = np.exp(min(xs)), np.exp(max(xs))
        if lo / 1.15 <= e_star <= hi * 1.15:     # inside (or just outside) the bracket: done
            break
        # extrapolated: re-bracket around the estimate and refit on all probes
        if verbose:
            print(f"    cal re-bracket: e*={e_star:.3f} outside [{lo:.3f},{hi:.3f}]", flush=True)
        for e in (max(0.05, e_star * 0.78), min(0.75, e_star * 1.28)):
            probe(e)
    e_star = round(float(np.clip(e_star, 0.05, 0.75)), 3)
    if verbose:
        print(f"    cal fit: slope {b:.4f}/log-e -> e* = {e_star}", flush=True)
    cache[key] = e_star
    json.dump(cache, open(CACHE, "w"), indent=1)
    return e_star


if __name__ == "__main__":
    ap = argparse.ArgumentParser()
    ap.add_argument("--s", type=float, default=0.0)
    ap.add_argument("--theta", type=float, default=1.0)
    ap.add_argument("--beta", type=float, default=0.0)
    ap.add_argument("--reps", type=int, default=100)
    ap.add_argument("--procs", type=int, default=4)
    ap.add_argument("--target", type=float, default=0.05)
    a = ap.parse_args()
    e = calibrate_e({"--s": a.s, "--theta": a.theta, "--beta": a.beta},
                    a.reps, a.procs, a.target)
    print(f"calibrated e = {e}")
```
