## Supplementary Information section 3 code base for "Hypothesis: A modern human range expansion ∼300,000 years ago had a critical role in Neandertal origins": band_ne.py

```
#!/usr/bin/env python3
"""
band_ne.py -- realised uniparental Ne for the band model, by the neutral-tracer method used
for the deme grid: run the model to a post-sweep steady state, RESET the uniparental markers
to frequency 1/2 (mtDNA across everyone; Y across males), then run on and fit

    ln E[p(1-p)]  ~  t * ln(1 - 1/Ne)          (E over replicates)

separately for mtDNA (female Ne) and Y (male Ne). Matching target from the Muller's-Ratchet
note: N_uniparental ~ 725 (bracket 500-1000), i.e. diploid ~1500.

    python band_ne.py [--reps 24] [--e 0.2] [--N 2000] [--burn 600] [--track 1500]
"""
import argparse
import os
from multiprocessing import Pool

import numpy as np

from bandsim import Config, Landscape, init_state, step

HERE = os.path.dirname(os.path.abspath(__file__))


def one(args):
    seed, kw, burn, track = args
    cfg = Config(gens=burn + track, seed=seed, **kw)
    rng = np.random.default_rng(cfg.seed)
    land = Landscape()
    P, pos, b_ident = init_state(cfg, land, rng)
    for g in range(burn):
        out = step(P, pos, b_ident, cfg, land, rng)
        if out[0] is None:
            return None
        P, pos, b_ident = out
    # reset tracers: mtDNA everywhere, Y among males, autosomal locus 0 -- all to p = 1/2
    P["mt"] = rng.integers(0, 2, len(P["mt"])).astype(np.int8)
    male = P["sex"] == 1
    yy = np.where(rng.random(len(P["Y"])) < 0.5, 1, -1).astype(np.int8)
    P["Y"] = np.where(male, yy, -1).astype(np.int8)
    P["auto"][:, 0] = rng.binomial(2, 0.5, len(P["mt"])).astype(np.int8)
    pm, py, pa = [], [], []
    for g in range(track):
        out = step(P, pos, b_ident, cfg, land, rng)
        if out[0] is None:
            return None
        P, pos, b_ident = out
        male = P["sex"] == 1
        pm.append(float((P["mt"] == 1).mean()))
        py.append(float((P["Y"][male] == 1).mean()) if male.any() else np.nan)
        pa.append(float(P["auto"][:, 0].mean() / 2.0))
    return pm, py, pa


def fit_ne(traj, lo_frac=0.05):
    """traj: (reps, T) of p. Fit slope of ln E[p(1-p)] over the window where E > lo_frac*E0."""
    het = traj * (1.0 - traj)
    E = np.nanmean(het, axis=0)
    E0 = E[:10].mean()
    good = np.where(E > lo_frac * E0)[0]
    t1 = good[-1] if len(good) else len(E) - 1
    t = np.arange(t1)
    y = np.log(np.maximum(E[:t1], 1e-12))
    b = np.polyfit(t, y, 1)[0]
    ne = -1.0 / b if b < 0 else np.inf
    return ne, t1


def main():
    ap = argparse.ArgumentParser()
    ap.add_argument("--reps", type=int, default=24)
    ap.add_argument("--e", type=float, default=0.2)
    ap.add_argument("--beta", type=float, default=0.0)
    ap.add_argument("--s", type=float, default=0.0)
    ap.add_argument("--N", type=int, default=1050)
    ap.add_argument("--K", type=int, default=25)
    ap.add_argument("--r", type=float, default=2.0)
    ap.add_argument("--burn", type=int, default=600)
    ap.add_argument("--track", type=int, default=1500)
    ap.add_argument("--procs", type=int, default=os.cpu_count())
    ap.add_argument("--stable-boundary", action="store_true")
    ap.add_argument("--sigma-e", type=float, default=80.0)
    ap.add_argument("--seed0", type=int, default=5000)
    a = ap.parse_args()

    kw = dict(N_target=a.N, K=a.K, beta=a.beta, s=a.s, e=a.e, r=a.r,
              sigma_e=a.sigma_e, stable_boundary=a.stable_boundary,
              record_every=10 ** 9)
    with Pool(a.procs) as pool:
        res = pool.map(one, [(a.seed0 + i, kw, a.burn, a.track) for i in range(a.reps)])
    res = [r for r in res if r is not None]
    pm = np.array([r[0] for r in res])
    py = np.array([r[1] for r in res])
    pa = np.array([r[2] for r in res])
    ne_f, t1f = fit_ne(pm)
    ne_m, t1m = fit_ne(py)
    ne_a, t1a = fit_ne(pa)
    print(f"\n{len(res)} tracer replicates (e={a.e}, beta={a.beta}, N={a.N}, K={a.K}, "
          f"burn {a.burn}, track {a.track})")
    print(f"  mtDNA (female) Ne : {ne_f:7.0f}   (fit window {t1f} gens)")
    print(f"  Y     (male)   Ne : {ne_m:7.0f}   (fit window {t1m} gens)")
    print(f"  autosomal      Ne : {ne_a:7.0f}   (diploid; fit window {t1a} gens)")
    print(f"  targets: diploid ~1500; uniparental bracket 500-1000")
    print(f"  census scaling hint (diploid anchor): N_target x {1500/max(ne_a,1):.2f}")


if __name__ == "__main__":
    main()
```
