## Supplementary Information section 3 code base for "Hypothesis: A modern human range expansion ∼300,000 years ago had a critical role in Neandertal origins": band_reps.py

```
#!/usr/bin/env python3
"""
band_reps.py -- replicate harness for the band model: run many seeds of one condition in
parallel, record per-replicate outcomes, and print the summary that matters:

  P(sweep)        fraction of replicates where the identity sweep completed by --gens
  sweep gen       median [IQR] completion generation among sweeps
  P(Y > 50%)      Y-establishment probability (global modern Y above half at the end)
  P(Y fixed)      Y at 100%
  mt / Y / aut    mean +- sd of final global modern fractions (post-sweep these equal the
                  within-modern-band fractions; both are recorded in the CSV)

Writes band_reps_<tag>.csv with one row per replicate.

    python band_reps.py --reps 100 --beta 0.0 --s 0.01 [--gens 800] [--e 0.08] [--procs 4]
"""
import argparse
import csv
import os
from multiprocessing import Pool

import numpy as np

from bandsim import AUTOS, Config, Landscape, init_state, step, stats

HERE = os.path.dirname(os.path.abspath(__file__))

# Loci reserved for mechanisms, never measured. Mirrors robust_run.MECH_LOCI and the
# spatial_sim2 layout: 0-31 neutral and measured, 32-35 available as selection targets.
MECH_LOCI = tuple(range(AUTOS - 4, AUTOS))


def excluded_loci(cfg):
    """Autosomal loci that must be dropped from the ancestry average for this condition.

    ADDED 11 Sep 2026 alongside bandsim.stats's `exclude` argument. band_reps called
    stats(P, b_ident) with no exclusions, so any condition that put an autosome under
    selection -- a DMI with an autosomal partner, or restorers -- averaged that locus,
    driven to ~100% modern, into the background estimate. At eight loci that added a fixed
    12.5 points per selected locus and the 2-15% screen was applied to the contaminated
    number, which is why the band arm of those families could not be repaired after the
    fact and had to be rerun. With the measurement panel now fixed at loci 0-31 the
    mechanism loci are excluded unconditionally, so this is belt and braces -- but a
    condition that points a mechanism at a measured locus is still a bug, and the assertion
    below refuses to run rather than producing a quietly wrong number.
    """
    excl = set(MECH_LOCI)
    if getattr(cfg, "dm", 0) > 0:
        if cfg.dm_focal == "auto":
            excl.add(cfg.dm_focal_locus)
        if cfg.dm_partner == "auto":
            excl.add(cfg.dm_partner_locus)
    if getattr(cfg, "restorers", 0) > 0:
        excl.update(range(AUTOS - cfg.restorers, AUTOS))
    stray = sorted(c for c in excl if c not in MECH_LOCI)
    if stray:
        raise SystemExit(
            f"band_reps: loci {stray} are under selection but sit inside the measured "
            f"panel 0-{AUTOS-5}. Point the mechanism at {MECH_LOCI} instead, or the "
            f"ancestry estimate will include the selected locus.")
    return tuple(sorted(excl))


def load_posterior(path, rna=1.0):
    """Accepted ABC draws of the mtDNA fitness gap (Muller's Ratchet note). One independent
    draw per replicate, exactly as joint_run.py does, so the reported probability integrates
    the gap posterior rather than conditioning on its median."""
    g = np.array([float(l) for l in open(path) if not l.startswith("#")])
    if g.size == 0:
        raise SystemExit(f"{path} contains no draws")
    return g * rna


def one_rep(args):
    seed, kw, gens = args
    cfg = Config(gens=gens, seed=seed, **kw)
    rng = np.random.default_rng(cfg.seed)
    land = Landscape()
    P, pos, b_ident = init_state(cfg, land, rng)
    sweep_gen = -1
    end_gen = gens
    for g in range(gens + 1):
        live = np.unique(P["band"])
        if sweep_gen < 0 and np.all(b_ident[live] == 1):
            sweep_gen = g
        # early stop (DR): sample at the deme-model horizon (~50 ka = gens) OR as soon as
        # both uniparental markers are absorbed (fixed or lost), whichever comes first.
        # Autosomal ancestry is neutral once band identities have settled, so its mean is
        # unaffected by stopping early; for expansions we additionally require the identity
        # sweep to be over (r > 1 still reshapes composition before that).
        if g % 10 == 0 and g > 0:
            male = P["sex"] == 1
            pm = float((P["mt"] == 1).mean())
            py = float((P["Y"][male] == 1).mean()) if male.any() else 0.0
            absorbed = pm in (0.0, 1.0) and py in (0.0, 1.0)
            settled = cfg.stable_boundary or sweep_gen >= 0
            if absorbed and settled:
                end_gen = g
                break
        out = step(P, pos, b_ident, cfg, land, rng)
        if out[0] is None:
            return dict(seed=seed, died=1, sweep_gen=-1, end_gen=g, mt=np.nan, Y=np.nan,
                        auto=np.nan, mt_mod=np.nan, Y_mod=np.nan, auto_mod=np.nan,
                        mtY_male=np.nan, p10=np.nan)
        P, pos, b_ident = out
    st = stats(P, b_ident, excluded_loci(cfg))
    return dict(seed=seed, died=0, sweep_gen=sweep_gen, end_gen=end_gen,
                mt=st["mt"], Y=st["Y"], auto=st["auto"],
                mt_mod=st["mt_mod"], Y_mod=st["Y_mod"], auto_mod=st["auto_mod"],
                mtY_male=st["mtY_male"], p10=float(st["mtY_male"] ** 10))


def main():
    ap = argparse.ArgumentParser()
    ap.add_argument("--reps", type=int, default=100)
    ap.add_argument("--gens", type=int, default=800)
    ap.add_argument("--beta", type=float, default=0.0)
    ap.add_argument("--s", type=float, default=0.01)
    ap.add_argument("--s-mt", type=float, default=0.0)
    ap.add_argument("--e", type=float, default=0.08)
    ap.add_argument("--r", type=float, default=2.0)
    ap.add_argument("--theta", type=float, default=1.0)
    ap.add_argument("--N", type=int, default=1050)
    ap.add_argument("--K", type=int, default=25)
    ap.add_argument("--sigma-e", type=float, default=80.0)
    ap.add_argument("--sigma-b", type=float, default=60.0)
    ap.add_argument("--p-move", type=float, default=0.45)
    ap.add_argument("--alpha-d", type=float, default=0.8)
    ap.add_argument("--s-mt-posterior", default=None,
                    help="file of accepted ABC draws; one s_mt drawn per replicate")
    ap.add_argument("--rna", type=float, default=1.0)
    ap.add_argument("--pulse-beta", type=float, default=0.5,
                    help="female share of the migration pulse (1 = all-female, 0 = all-male)")
    ap.add_argument("--pulse-mode", default="individual", choices=("individual", "band"))
    ap.add_argument("--no-reseed", action="store_true")
    ap.add_argument("--hard-selection", action="store_true",
                    help="refill slots vacated by selection (the deme engines' convention)")
    ap.add_argument("--stable-boundary", action="store_true")
    ap.add_argument("--pulse-frac", type=float, default=0.05)
    # ---- SI mechanism loci (see bandsim.Config for semantics; the sign and timing
    # conventions are copied from spatial_sim2/3 so band and deme cells are comparable)
    ap.add_argument("--dm", type=float, default=0.0)
    ap.add_argument("--dm-focal", default="mt", choices=("mt", "Y", "auto", "X"))
    ap.add_argument("--dm-partner", default="Y", choices=("mt", "Y", "auto", "X"))
    ap.add_argument("--dm-dom", default="dominant",
                    choices=("dominant", "additive", "recessive"))
    # Default moved 7 -> 35 on 11 Sep 2026 with the 32+4 locus layout. Locus 7 is now inside
    # the measured neutral panel (0-31); leaving the default there would have put a DMI
    # partner straight into the ancestry statistic. 35 is the last mechanism locus.
    ap.add_argument("--dm-focal-locus", type=int, default=AUTOS - 1)
    ap.add_argument("--dm-partner-locus", type=int, default=AUTOS - 1)
    ap.add_argument("--no-dm-male-limited", action="store_true")
    ap.add_argument("--restorers", type=int, default=0)
    ap.add_argument("--rest-pen", type=float, default=1.0)
    ap.add_argument("--xdrive", type=float, default=0.0)
    ap.add_argument("--xdrive-cost", type=float, default=0.0)
    ap.add_argument("--xdrv-res", type=float, default=0.0)
    ap.add_argument("--xdrv-all", action="store_true")
    ap.add_argument("--esc-eps", type=float, default=0.0)
    ap.add_argument("--inbreed-ban", type=int, default=0, choices=(0, 1, 2))
    ap.add_argument("--inbreed-tries", type=int, default=12)
    ap.add_argument("--rho-return", type=float, default=0.0)
    ap.add_argument("--skew-frac", type=float, default=1.0)
    ap.add_argument("--e-export", type=float, default=None)
    ap.add_argument("--e-import", type=float, default=None)
    ap.add_argument("--procs", type=int, default=os.cpu_count())
    ap.add_argument("--seed0", type=int, default=1000)
    ap.add_argument("--tag", default=None)
    ap.add_argument("--outfile", default=None)
    a = ap.parse_args()

    kw = dict(N_target=a.N, K=a.K, beta=a.beta, s=a.s, s_mt=a.s_mt, e=a.e, r=a.r,
              theta=a.theta, sigma_e=a.sigma_e, sigma_b=a.sigma_b, p_move=a.p_move, alpha_d=a.alpha_d,
              stable_boundary=a.stable_boundary, pulse_frac=a.pulse_frac,
              pulse_beta=a.pulse_beta, pulse_mode=a.pulse_mode,
              reseed=not a.no_reseed, hard_selection=a.hard_selection,
              dm=a.dm, dm_focal=a.dm_focal, dm_partner=a.dm_partner, dm_dom=a.dm_dom,
              dm_focal_locus=a.dm_focal_locus, dm_partner_locus=a.dm_partner_locus,
              dm_male_limited=not a.no_dm_male_limited,
              restorers=a.restorers, rest_pen=a.rest_pen,
              xdrive=a.xdrive, xdrive_cost=a.xdrive_cost, xdrv_res=a.xdrv_res,
              xdrv_all=a.xdrv_all, esc_eps=a.esc_eps,
              inbreed_ban=a.inbreed_ban, inbreed_tries=a.inbreed_tries,
              rho_return=a.rho_return, skew_frac=a.skew_frac,
              e_export=a.e_export, e_import=a.e_import,
              record_every=10 ** 9)
    # per-replicate s_mt draws (seeded on seed0 so batches of the same condition draw
    # DIFFERENT values instead of all repeating the first batch -- the joint_run.py rule)
    if a.s_mt_posterior:
        post = load_posterior(a.s_mt_posterior, a.rna)
        draw_rng = np.random.default_rng(abs(hash((a.tag or "", a.seed0))) % (2**32))
        s_mt_all = draw_rng.choice(post, size=a.reps, replace=True)
        tasks = [(a.seed0 + i, {**kw, "s_mt": float(s_mt_all[i])}, a.gens)
                 for i in range(a.reps)]
    else:
        s_mt_all = np.full(a.reps, a.s_mt)
        tasks = [(a.seed0 + i, kw, a.gens) for i in range(a.reps)]
    if a.procs <= 1:
        rows = [one_rep(t) for t in tasks]
    else:
        with Pool(a.procs) as pool:
            rows = pool.map(one_rep, tasks)

    for row, sv in zip(rows, s_mt_all):
        row["s_mt_drawn"] = float(sv)
    tag = a.tag or f"b{a.beta:g}_s{a.s:g}_e{a.e:g}_r{a.r:g}"
    out = a.outfile or os.path.join(HERE, f"band_reps_{tag}.csv")
    os.makedirs(os.path.dirname(out) or ".", exist_ok=True)
    with open(out, "w", newline="") as f:
        wr = csv.DictWriter(f, fieldnames=list(rows[0]))
        wr.writeheader()
        wr.writerows(rows)

    ok = [r for r in rows if not r["died"]]
    died = len(rows) - len(ok)
    sw = np.array([r["sweep_gen"] for r in ok])
    swept = sw >= 0
    Yv = np.array([r["Y"] for r in ok])
    mtv = np.array([r["mt"] for r in ok])
    auv = np.array([r["auto"] for r in ok])
    print(f"\n{a.reps} replicates  (beta={a.beta}, s={a.s}, s_mt={a.s_mt}, e={a.e}, "
          f"r={a.r}, theta={a.theta}, N={a.N}, K={a.K}, {a.gens} gens)")
    if died:
        print(f"  died out entirely : {died}")
    print(f"  P(sweep by gen {a.gens})   : {100 * swept.mean():.0f}%")
    if swept.any():
        q1, q2, q3 = np.percentile(sw[swept], [25, 50, 75])
        print(f"  sweep generation          : median {q2:.0f}  [IQR {q1:.0f}-{q3:.0f}]")
    print(f"  P(Y > 50% at end)         : {100 * (Yv > 0.5).mean():.0f}%")
    print(f"  P(Y fixed at end)         : {100 * (Yv > 0.999).mean():.0f}%")
    print(f"  final global modern fractions (mean +- sd over replicates):")
    print(f"      mt   {100 * mtv.mean():5.1f} +- {100 * mtv.std():4.1f}%")
    print(f"      Y    {100 * Yv.mean():5.1f} +- {100 * Yv.std():4.1f}%")
    print(f"      aut  {100 * auv.mean():5.1f} +- {100 * auv.std():4.1f}%")
    print(f"  wrote {out}")


if __name__ == "__main__":
    main()
```
