## Supplementary Information section 3 code base for "Hypothesis: A modern human range expansion ∼300,000 years ago had a critical role in Neandertal origins": band_suite.py

```
#!/usr/bin/env python3
"""
band_suite.py -- the full clan/band-model analysis suite: band analogues of every deme-model
scenario x mechanism cell, with per-condition e-calibration to 5% autosomes, run to the
2000-generation absorption horizon, collected into one summary table.

Scenarios (band analogues of the deme model's four):
  fexp  matrilocal expansion        beta=0,   Istanbul seed, r=2
  nexp  non-sex-biased expansion    beta=0.5, Istanbul seed, r=2
  fsb   matrilocal stable boundary  beta=0,   5% pulse, r=1
  nsb   non-sex-biased boundary     beta=0.5, 5% pulse, r=1

Mechanism sweeps per scenario:
  s      additive selection on the modern Y: 0 .002 .005 .01 .02 .04 .08 .16   (theta=1)
  theta  F1-male fertility: 1 .2 .05 0                                          (s=0)
Horizon: 6993 generations (the deme series' ~50 ka sampling point), with early stop the
moment both uniparental markers are absorbed (and, for expansions, the sweep is over).

Calibration: expansions get e calibrated PER CONDITION by band_cal.calibrate_e -- three fixed
probes (e = 0.30/0.40/0.52, 100 reps each) with a log-linear fit inverted at 5% autosomal
ancestry, conditioned on establishment. Stable-boundary scenarios keep e = 0.2; their
autosomal level is pinned by the 5% admixture pulse itself.

Outputs band_suite_summary.csv: scenario, sweep, value, e_used, n, P(mt fixed), P(Y fixed),
P(joint = both fixed), mean mt/Y/aut. Resumable: every condition cached as a CSV.

    python band_suite.py [--reps 100] [--procs 4] [--cal-reps 24] [--dry-run]
"""
import argparse
import csv
import os
import subprocess
import sys

import numpy as np

from band_cal import calibrate_e

HERE = os.path.dirname(os.path.abspath(__file__))

# SETTLED CENSUS (4 Sep 2026). N_target = 1750 is the value at which the measured autosomal
# effective size lands on the Peyregne anchor of 1,447 diploids (2,894 gene copies) in all four
# band configurations: fexp 2844, nexp 2777, fpulse 2807, npulse 3025, every one within 4%.
N_TARGET = 1750

SCEN = {
    "fexp": {"--beta": 0.0, "--r": 2.0, "--N": N_TARGET},   # K=25, p_move=0.45, sigma_b=60, sigma_e=80
    "nexp": {"--beta": 0.5, "--r": 2.0, "--N": N_TARGET},
    # migration pulse: 5% of individuals replaced at random, sex bias set by --pulse-beta
    # (female share; 1.0 = all-female, matching the deme comparator's male_frac = 0)
    "fsb":  {"--beta": 0.0, "--r": 1.0, "--stable-boundary": None, "--pulse-beta": 1.0,
             "--N": N_TARGET},
    "nsb":  {"--beta": 0.5, "--r": 1.0, "--stable-boundary": None, "--pulse-beta": 0.5,
             "--N": N_TARGET},
}
# extended to 0.32/0.64 to match the deme series, where the matrilocal expansion only reaches
# P(joint) = 1 at s >= 0.16; the band model needs a longer arm to show the same plateau
S_GRID = [0.0, 0.002, 0.005, 0.01, 0.02, 0.04, 0.08, 0.16, 0.32, 0.64]
# Extended toward zero on DR's instruction (5 Sep 2026): between theta = 0.05 and theta = 0
# the matrilocal expansion goes from 12% to 81% joint fixation, so the entire transition was
# being carried by a single segment. These rungs resolve it. Note the deme model puts its own
# threshold at theta < 4% (Figure S3.3), which falls inside the new spacing.
TH_GRID = [1.0, 0.2, 0.1, 0.05, 0.03, 0.02, 0.01, 0.005, 0.002, 0.0]
GENS = 6993   # deme-model horizon (~50 ka); early fix-or-lose stop in band_reps


def flags_to_cmd(flags):
    out = []
    for k, v in flags.items():
        out.append(k)
        if v is not None:
            out.append(str(v))
    return out


BATCH = 40   # reps per sub-batch: each fits in one supervision window, resumable


def run_cond(tag, flags, reps, procs):
    rows = []
    for b in range(0, reps, BATCH):
        n = min(BATCH, reps - b)
        path = os.path.join(HERE, f"band_reps_{tag}_b{b}.csv")
        if not os.path.exists(path):
            cmd = [sys.executable, os.path.join(HERE, "band_reps.py"), "--reps", str(n),
                   "--gens", str(GENS), "--procs", str(procs), "--tag", f"{tag}_b{b}",
                   "--seed0", str(1000 + b)] + flags_to_cmd(flags)
            subprocess.run(cmd, check=True, capture_output=True, cwd=HERE)
        with open(path) as f:
            rows += [r for r in csv.DictReader(f) if r["died"] == "0"]
    mt = np.array([float(r["mt"]) for r in rows])
    Y = np.array([float(r["Y"]) for r in rows])
    au = np.array([float(r["auto"]) for r in rows])
    sw = np.array([float(r["sweep_gen"]) for r in rows])
    sb = any(float(r["sweep_gen"]) < 0 and float(r["mt"]) > 0 for r in rows)  # crude sb flag
    est = (sw >= 0) if not sb else np.ones(len(rows), bool)
    fix = lambda x: x > 0.999
    joint = fix(mt) & fix(Y)
    return dict(n=len(rows), p_sweep=est.mean(),
                p_mt=fix(mt).mean(), p_Y=fix(Y).mean(), p_joint=joint.mean(),
                p_joint_est=joint[est].mean() if est.any() else np.nan,
                mt=mt.mean(), Y=Y.mean(), aut=au.mean(),
                aut_est=au[est].mean() if est.any() else np.nan)


def main():
    ap = argparse.ArgumentParser()
    ap.add_argument("--reps", type=int, default=100)
    ap.add_argument("--cal-reps", type=int, default=100)
    ap.add_argument("--procs", type=int, default=4)
    ap.add_argument("--scenarios", default="fexp,nexp,fsb,nsb")
    ap.add_argument("--dry-run", action="store_true")
    a = ap.parse_args()

    conds = []
    for sc in a.scenarios.split(","):
        base = SCEN[sc]
        for s in S_GRID:
            conds.append((sc, "s", s, {**base, "--s": s, "--theta": 1.0}))
        for th in TH_GRID:
            if th == 1.0:
                continue          # theta=1, s=0 duplicates the s-grid zero point
            conds.append((sc, "theta", th, {**base, "--s": 0.0, "--theta": th}))
    if a.dry_run:
        est = len(conds) * a.reps * GENS * 2.9e-3 / a.procs / 60
        print(f"{len(conds)} conditions x {a.reps} reps x {GENS} gens "
              f"~ {est:.0f} min on {a.procs} procs (+ calibration)")
        for c in conds:
            print("  ", c[0], c[1], c[2])
        return

    out_rows = []
    prev_e = {}
    for sc, sweep, val, flags in conds:
        expansion = "--stable-boundary" not in flags
        if expansion:
            cal_flags = {k: v for k, v in flags.items()}
            e = calibrate_e(cal_flags, reps=a.cal_reps, procs=a.procs)
        else:
            e = 0.2
        tag = f"suite_{sc}_{sweep}{val:g}"
        res = run_cond(tag, {**flags, "--e": e}, a.reps, a.procs)
        row = dict(scenario=sc, sweep=sweep, value=val, e_used=e, **res)
        out_rows.append(row)
        print(f"[{sc} {sweep}={val:g}] e={e:.3f} n={res['n']} P(sweep)={res['p_sweep']:.2f} "
              f"P(mt)={res['p_mt']:.2f} P(Y)={res['p_Y']:.2f} P(joint)={res['p_joint']:.2f} "
              f"(est: {res['p_joint_est']:.2f})  aut={100*res['aut']:.1f}% "
              f"(est: {100*res['aut_est']:.1f}%)", flush=True)
        with open(os.path.join(HERE, "band_suite_summary.csv"), "w", newline="") as f:
            w = csv.DictWriter(f, fieldnames=list(out_rows[0]))
            w.writeheader(); w.writerows(out_rows)
    print("wrote band_suite_summary.csv")


if __name__ == "__main__":
    main()
```
