## Supplementary Information section 3 code base for "Hypothesis: A modern human range expansion ∼300,000 years ago had a critical role in Neandertal origins": bandcal_submit.py

```
#!/usr/bin/env python3
"""
bandcal_submit.py -- STAGE 1 of the band-model suite on O2: the calibration probe array.

For every (scenario x mechanism) condition that needs calibration (the two expansion
scenarios), run the exogamy-rate probes. Stage 2 (bandprod_submit.py) fits e* per condition
from these probes and launches production. Exactly the jcal -> joint2 pattern.

    python bandcal_submit.py [reps] [batch] [concurrency] [partition] [--resume]
    sbatch bandcal_array.sh

Defaults: 200 probe reps per (condition, e), batch 50 -> 4 tasks each.
"""
import os
import sys

HERE = os.path.dirname(os.path.abspath(__file__))
sys.path.insert(0, HERE)
from band_suite import SCEN, S_GRID, TH_GRID, GENS, flags_to_cmd   # noqa: E402

# Probe ladders. These must BRACKET the 5% target or e* is an extrapolation.
#
# Moved down for beta = 0 on 4 Sep 2026 when the census went 1050 -> 1750. At the larger
# census there are ~70 bands rather than ~42, the wave makes more contacts on its way across,
# and modern autosomal ancestry at any given e is lower: the old ladder (0.26-0.56) produced
# 3.9% at its LOWEST rung, so the whole bracket sat above the target and every fexp e* was
# extrapolated. The old rungs are kept in out_bandcal and still feed the fit, so re-running
# with --resume simply widens the ladder to seven points rather than replacing it.
PROBES = {0.0: (0.08, 0.13, 0.19, 0.26),
          0.5: (0.10, 0.16, 0.26, 0.40)}


def conditions(scenarios=("fexp", "nexp")):
    """(scenario, sweep, value, flags) for every condition needing calibration"""
    out = []
    for sc in scenarios:
        base = SCEN[sc]
        for s in S_GRID:
            out.append((sc, "s", s, {**base, "--s": s, "--theta": 1.0}))
        for th in TH_GRID:
            if th == 1.0:
                continue
            out.append((sc, "theta", th, {**base, "--s": 0.0, "--theta": th}))
    return out


def main():
    args = [a for a in sys.argv[1:] if not a.startswith("--")]
    reps = int(args[0]) if args else 200
    batch = int(args[1]) if len(args) > 1 else 50
    cap = int(args[2]) if len(args) > 2 else 0
    part = args[3] if len(args) > 3 else "short"
    resume = "--resume" in sys.argv
    os.makedirs("out_bandcal", exist_ok=True)
    os.makedirs("logs", exist_ok=True)

    tasks, skipped = [], 0
    for sc, sweep, val, flags in conditions():
        beta = float(flags["--beta"])
        for e in PROBES[beta]:
            for s0 in range(0, reps, batch):
                n = min(batch, reps - s0)
                tag = f"{sc}_{sweep}{val:g}_e{e:g}_s{s0}"
                out = f"out_bandcal/{tag}.csv"
                if resume and os.path.exists(out) and sum(1 for _ in open(out)) >= n + 1:
                    skipped += 1
                    continue
                fl = " ".join(flags_to_cmd({**flags, "--e": e}))
                tasks.append(f"--reps {n} --seed0 {5000 + s0} --gens {GENS} "
                             f"--procs 1 --outfile {out} {fl}")
    if not tasks:
        print("nothing to do -- every probe output is already complete")
        return
    open("bandcal_params.txt", "w").write("\n".join(tasks) + "\n")
    n = len(tasks)
    cap = min(n, cap) if cap else 0
    open("bandcal_array.sh", "w").write(f"""#!/bin/bash
#SBATCH -J bandcal
#SBATCH -p {part}
#SBATCH -t 0-05:59
#SBATCH -c 1
#SBATCH --mem=2G
#SBATCH -o logs/bandcal_%A_%a.log
#SBATCH --array=0-{n - 1}{("%%%d" % cap) if cap else ""}

LINE=$(sed -n "$((SLURM_ARRAY_TASK_ID + 1))p" bandcal_params.txt)
echo "task $SLURM_ARRAY_TASK_ID : $LINE"
python band_reps.py $LINE
""")
    conds = len(conditions())
    print(f"wrote bandcal_params.txt ({n} tasks)")
    print(f"  {conds} conditions x 4 probes x {reps} reps (batch {batch})")
    if resume:
        print(f"  RESUME: skipped {skipped} complete tasks")
    print(f"  ~{batch * 1.6 / 60:.0f}-{batch * 6.0 / 60:.0f} min per task")
    print("submit:  sbatch bandcal_array.sh")
    print("then:    python bandprod_submit.py   (fits e*, writes the production array)")


if __name__ == "__main__":
    main()
```
