## Supplementary Information section 3 code base for "Hypothesis: A modern human range expansion ∼300,000 years ago had a critical role in Neandertal origins": bandmech_collect.py

```
#!/usr/bin/env python3
"""
bandmech_collect.py -- pool out_bandmech/ into one row per condition. (DR, 6 Sep 2026)

Writes bandmech_summary.csv and prints each family as a table, with the corresponding deme
number alongside wherever v27 quotes one, so the two frameworks can be compared row by row
rather than by eye across two documents.

Read the autosomal column first. The note's whole argument is that a mechanism which retains
the modern Y but drags autosomal ancestry far from the observed ~5% has not explained the data,
so a cell with a spectacular Y and 40% autosomes is a failure, not a success. Cells outside
4-6% are flagged, and the restorer family is expected to be among them -- that departure is the
result the deme runs already report (20-59%).

    python bandmech_collect.py [--family dmi]
"""
import csv
import glob
import os
import re
import sys

import numpy as np

HERE = os.path.dirname(os.path.abspath(__file__))

# Deme anchors quoted in v27 / the collected deme summaries, for the cells that correspond
# one-to-one. None where the deme grid has no directly matching cell.
DEME_ANCHOR = {
    "dmi_mtxY":        ("Table S3.1 mtDNA x Y", "Y 100%, auto 10%"),
    "dmi_mtxAutoDom":  ("Table S3.1 mtDNA x autoDom", "Y 53%, auto 23%"),
    "dmi_mtxAutoAdd":  ("Table S3.1 mtDNA x autoAdd", "Y 14%, auto 15%"),
    "dmi_mtxAutoRec":  ("Table S3.1 mtDNA x autoRec", "Y 5%, auto 11%"),
    "dmi_mtxXdom":     ("Table S3.1 mtDNA x Xdom", "Y 5%, auto 11%"),
    "dmi_XxY":         ("Table S3.1 X x Y", "Y 24%, auto 10%"),
    "esc_0":           ("Table S3.2 eps=0", "Y 100%"),
    "esc_0.1":         ("Table S3.2 eps=0.1", "Y 52.5%, escaped 47.5%"),
    "xdrv_d0.95_c0":   ("Table S3.3 d=0.95", "Y 100%, auto 10.5%"),
    "xdrv_d0.5_c0":    ("Table S3.3 d=0.5", "Y 99.8%, auto 9.1%"),
    "xdrv_res0.003":   ("Fig S3.5 right eps=0.003", "Y 60.4%, resistant 39.6%"),
    "xdrv_res0.03":    ("Fig S3.5 right eps=0.03", "Y 13.3%, resistant 86.7%"),
}

PATTERNS = [
    ("dmi",  r"^dmi_(?P<row>\w+)$"),
    ("rest", r"^rest_n(?P<nloci>\d+)_(?P<mode>\w+)$"),
    ("esc",  r"^esc_(?P<eps>[\d.e-]+)$"),
    ("xdrv", r"^xdrv_(?:d(?P<d>[\d.]+)_c(?P<cost>[\d.]+)|res(?P<res>[\d.e-]+))$"),
    ("sf1",  r"^sf1_s(?P<s>[\d.]+)$"),
    ("kin",  r"^kin_b(?P<ban>\d)_t(?P<theta>[\d.]+)$"),
    ("cult", r"^cult_r(?P<rho>[\d.]+)_b(?P<beta>[\d.]+)_k(?P<skew>[\d.]+)$"),
]


def parse(tag):
    for fam, pat in PATTERNS:
        m = re.match(pat, tag)
        if m:
            return fam, {k: v for k, v in m.groupdict().items() if v is not None}
    return None, {}


def main():
    only = sys.argv[sys.argv.index("--family") + 1] if "--family" in sys.argv else None
    by_tag = {}
    for path in sorted(glob.glob(os.path.join(HERE, "out_bandmech", "*.csv"))):
        tag = re.sub(r"_s\d+$", "", os.path.basename(path)[:-4])
        by_tag.setdefault(tag, []).extend(list(csv.DictReader(open(path))))
    if not by_tag:
        raise SystemExit("no outputs in out_bandmech/")

    rows, off = [], []
    for tag, raw in sorted(by_tag.items()):
        fam, params = parse(tag)
        if fam is None:
            continue
        live = [r for r in raw if r["died"] == "0"]
        if not live:
            continue
        g = lambda k: np.array([float(r[k]) for r in live])
        mt, Y, au = g("mt"), g("Y"), g("auto")
        fix = lambda x: x > 0.999
        p = float((fix(mt) & fix(Y)).mean())
        RESERVED = {"family", "tag", "n", "n_died", "mt", "Y", "aut", "p_mt", "p_Y",
                    "p_joint", "se_joint", "end_gen_med"}
        params = {(f"p_{k}" if k in RESERVED else k): v for k, v in params.items()}
        row = dict(family=fam, tag=tag, **params, n=len(live), n_died=len(raw) - len(live),
                   p_mt=round(float(fix(mt).mean()), 4), p_Y=round(float(fix(Y).mean()), 4),
                   p_joint=round(p, 4),
                   se_joint=round(float(np.sqrt(max(p * (1 - p), 1e-9) / len(live))), 4),
                   mt=round(float(mt.mean()), 4), Y=round(float(Y.mean()), 4),
                   aut=round(float(au.mean()), 4),
                   end_gen_med=int(np.median(g("end_gen"))))
        rows.append(row)
        if not 0.04 <= row["aut"] <= 0.06:
            off.append((fam, tag, row["aut"]))

    # NB: `rows` is always ALL families, so the summary written below is complete no matter
    # what --family was given; `only` is applied to the printed tables further down.
    keys = sorted({k for r in rows for k in r})
    order = ["family", "tag"] + [k for k in keys if k not in ("family", "tag")]
    with open(os.path.join(HERE, "bandmech_summary.csv"), "w", newline="") as f:
        w = csv.DictWriter(f, fieldnames=order)
        w.writeheader(); w.writerows(rows)
    print(f"wrote bandmech_summary.csv ({len(rows)} conditions)")

    for fam in sorted({r["family"] for r in rows}):
        if only and fam != only:
            continue
        sub = sorted([r for r in rows if r["family"] == fam], key=lambda r: r["tag"])
        print(f"\n=== {fam}  ({len(sub)} conditions) ===")
        print(f"  {'tag':24s} {'n':>5} {'mt%':>7} {'Y%':>7} {'joint%':>8} {'aut%':>7}   deme")
        for r in sub:
            anch = DEME_ANCHOR.get(r["tag"])
            note = f"   {anch[1]}" if anch else ""
            print(f"  {r['tag']:24s} {r['n']:5d} {100*r['mt']:7.1f} {100*r['Y']:7.1f} "
                  f"{100*r['p_joint']:8.1f} {100*r['aut']:7.1f}{note}")

    if off:
        print(f"\nautosomal ancestry outside 4-6% in {len(off)} cells "
              f"(expected for the restorers, which the deme runs put at 20-59%):")
        for fam, tag, a in off[:25]:
            print(f"  {fam:6s} {tag:24s} {100*a:6.1f}%")
        if len(off) > 25:
            print(f"  ... and {len(off) - 25} more")


if __name__ == "__main__":
    main()
```
