## Supplementary Information section 3 code base for "Hypothesis: A modern human range expansion ∼300,000 years ago had a critical role in Neandertal origins": bandmech_submit.py

```
#!/usr/bin/env python3
"""
bandmech_submit.py -- band-model analogues of the seven deme SI families that need the
mechanism loci in bandsim.py. (DR, 6 Sep 2026)

Companion to bandsi_submit.py, which covers the five families that needed no engine work.
Between them the twelve families give the band model a parallel analysis for every figure and
table in the introgression note.

    family    deme counterpart   v27 target          mechanism
    ------------------------------------------------------------------------------------
    dmi       dm2                Table S3.1          single-pair DMI, the scenario table
    rest      dm3                Table S3.1 (poly)   polygenic mito-nuclear restorers
    esc       esc                Table S3.2          heritable escape of the archaic Y
    xdrv      xdrv + xdrv2       Table S3.3, S3.5    X-drive, cost, heritable resistance
    sf1       sf1                Figure S3.4         partner locus under patrilocal expansion
    kin       kin2 + kin3        kin avoidance       co-resident kin ban, degree 1 and 2
    cult      mech3/4/5          cultural rules      sex-biased exogamy, return rule, skew

NO RATCHET POSTERIOR. Every deme SI family runs at s_mt = 0, so these do too. Getting this
wrong once already produced a band pulse cell reading 27.5% against a deme 10.9% that looked
like a framework disagreement and was only selection present on one side. The ratchet arm is a
separate production series (bandratchet), which is what Figure 2b uses.

CALIBRATION. Expansion families are calibrated to 5% autosomal ancestry by the two-stage probe
ladder, exactly as bandsi does. Mechanisms that themselves move autosomal ancestry a long way
(the restorers especially, which reach 20-59% in the deme runs) are NOT forced back to 5% --
that departure IS the result, and the collector reports realised autosomes so it stays visible.

    python bandmech_submit.py cal  [reps] [batch] [cap] [partition] [--only dmi,kin]
    sbatch bandmech_cal_array.sh
    python bandmech_submit.py prod [reps] [batch] [cap] [partition] [--resume]
    sbatch bandmech_prod_array.sh
    python bandmech_collect.py
"""
import csv
import glob
import os
import re
import sys

import numpy as np

HERE = os.path.dirname(os.path.abspath(__file__))
sys.path.insert(0, HERE)
from band_suite import GENS, N_TARGET, flags_to_cmd            # noqa: E402
from bandsi_submit import (BASE_EXP, BASE_PULSE, PROBES, TARGET,   # noqa: E402
                           fit_e_star, write_array, scan_base_e)

# ---------------------------------------------------------------- grids
# dm2's scenario table, restricted to the rows the band model can express. The deme rows that
# turn on PRDM9 or a second autosome are kept: bandsim carries the same locus layout as
# spatial_sim2 -- since 11 Sep 2026 that is 36 autosomes (0-31 neutral and measured, 32-35
# reserved for mechanisms) and 3 X loci (0 drive, 1 DMI candidate, 2 neutral), not the 8 and 2
# this comment used to claim. Any mechanism locus named below must sit in 32-35, or band_reps
# will refuse the run rather than average a selected locus into the ancestry estimate.
DMI_ROWS = [
    ("mtxY",      dict(dm_focal="mt",   dm_partner="Y")),
    ("mtxAutoDom", dict(dm_focal="mt",  dm_partner="auto", dm_dom="dominant")),
    ("mtxAutoAdd", dict(dm_focal="mt",  dm_partner="auto", dm_dom="additive")),
    ("mtxAutoRec", dict(dm_focal="mt",  dm_partner="auto", dm_dom="recessive")),
    # dm_partner_locus MUST be given for an X partner: the default is an autosomal index
    # (35) and X has only 3 loci -- leaving it unset raised IndexError in every task.
    # Locus 1 is the X's designated DMI/selection candidate; 2 is the neutral background.
    ("mtxXdom",   dict(dm_focal="mt",   dm_partner="X",    dm_dom="dominant",
                       dm_partner_locus=1)),
    ("XxY",       dict(dm_focal="X",    dm_partner="Y",    dm_focal_locus=1)),
    ("autoxmt",   dict(dm_focal="auto", dm_partner="mt")),
]
REST_N = [1, 2, 3, 4, 6]
REST_MODE = ["dominant", "additive"]
ESC_EPS = [0.0, 1e-4, 3e-4, 1e-3, 3e-3, 1e-2, 3e-2, 1e-1]
XDRV_D = [0.05, 0.1, 0.2, 0.3, 0.4, 0.5, 0.8, 0.95, 1.0]
XDRV_COST = [0.0, 0.1]
XDRV_RES = [1e-4, 3e-4, 1e-3, 3e-3, 1e-2, 3e-2]
SF1_S = [0.0, 0.005, 0.01, 0.02, 0.04, 0.08, 0.16, 0.32]
KIN_BANS = [0, 1, 2]
KIN_THETA = [0.0, 0.05, 0.1, 0.2, 0.4, 0.7, 1.0]
CULT_RHO = [0.0, 0.2, 0.6, 0.9, 1.0]
CULT_BETA = [0.0, 0.25, 0.5]
CULT_SKEW = [1.0, 0.5, 0.25]


def conditions(family):
    """(family, tag, needs_calibration, flags). Expansion families calibrate; the two that
    inherit the deme baseline geometry reuse the fexp e*."""
    out = []
    E = scan_base_e()
    if family == "dmi":
        for name, kw in DMI_ROWS:
            fl = {**BASE_EXP, "--beta": 0.0, "--s": 0.0, "--theta": 1.0, "--e": E, "--dm": 1.0}
            fl.update({f"--{k.replace('_', '-')}": v for k, v in kw.items()})
            out.append((family, f"dmi_{name}", False, fl))
    elif family == "rest":
        for n in REST_N:
            for mode in REST_MODE:
                out.append((family, f"rest_n{n}_{mode}", False,
                            {**BASE_EXP, "--beta": 0.0, "--s": 0.0, "--theta": 1.0, "--e": E,
                             "--restorers": n, "--rest-pen": 1.0, "--dm-dom": mode}))
    elif family == "esc":
        for eps in ESC_EPS:
            out.append((family, f"esc_{eps:g}", False,
                        {**BASE_EXP, "--beta": 0.0, "--s": 0.0, "--theta": 1.0, "--e": E,
                         "--dm": 1.0, "--dm-focal": "mt", "--dm-partner": "Y",
                         "--esc-eps": eps}))
    elif family == "xdrv":
        for d in XDRV_D:
            for c in XDRV_COST:
                out.append((family, f"xdrv_d{d:g}_c{c:g}", False,
                            {**BASE_EXP, "--beta": 0.0, "--s": 0.0, "--theta": 1.0, "--e": E,
                             "--xdrive": d, "--xdrive-cost": c}))
        for eps in XDRV_RES:                       # resistance block, at strong drive
            out.append((family, f"xdrv_res{eps:g}", False,
                        {**BASE_EXP, "--beta": 0.0, "--s": 0.0, "--theta": 1.0, "--e": E,
                         "--xdrive": 0.95, "--xdrv-res": eps}))
    elif family == "sf1":
        # patrilocal: beta = 1 means all-female exogamy, so men stay -- the band analogue of
        # the deme's lineage = "patrilineal". A mechanism locus under a DMI with archaic
        # mtDNA, with additive selection on the same locus, exactly as sf1_run.
        # Locus moved 7 -> 35 on 11 Sep 2026: under the 32+4 layout locus 7 is inside the
        # measured neutral panel, so the old value put the DMI focal locus -- driven to ~100%
        # modern -- directly into the autosomal ancestry statistic this family screens on.
        for s in SF1_S:
            out.append((family, f"sf1_s{s:g}", True,
                        {**BASE_EXP, "--beta": 1.0, "--theta": 1.0, "--s": s,
                         "--dm": 1.0, "--dm-focal": "auto", "--dm-focal-locus": 35,
                         "--dm-partner": "mt", "--dm-dom": "dominant"}))
    elif family == "kin":
        for ban in KIN_BANS:
            for th in KIN_THETA:
                out.append((family, f"kin_b{ban}_t{th:g}", False,
                            {**BASE_EXP, "--beta": 0.0, "--s": 0.0, "--theta": th, "--e": E,
                             "--inbreed-ban": ban}))
    elif family == "cult":
        for rho in CULT_RHO:
            for b in CULT_BETA:
                for sk in CULT_SKEW:
                    if rho == 0.0 and sk == 1.0 and b == 0.5:
                        pass                      # keep the plain baseline once
                    out.append((family, f"cult_r{rho:g}_b{b:g}_k{sk:g}", False,
                                {**BASE_EXP, "--beta": b, "--s": 0.0, "--theta": 1.0, "--e": E,
                                 "--rho-return": rho, "--skew-frac": sk}))
    else:
        raise SystemExit(f"unknown family {family}")
    return out


FAMILIES = ["dmi", "rest", "esc", "xdrv", "sf1", "kin", "cult"]


def probe_autosomes(tag):
    by_e = {}
    for path in glob.glob(os.path.join(HERE, "out_bandmech_cal", f"{tag}_e*_s*.csv")):
        m = re.search(r"_e([0-9.]+)_s\d+\.csv$", path)
        if not m:
            continue
        rows = [r for r in csv.DictReader(open(path)) if r["died"] == "0"]
        est = [r for r in rows if float(r["sweep_gen"]) >= 0]
        use = est if len(est) >= max(10, len(rows) // 3) else rows
        if use:
            by_e[float(m.group(1))] = float(np.mean([float(r["auto"]) for r in use]))
    return by_e


def main():
    if len(sys.argv) < 2 or sys.argv[1] not in ("cal", "prod"):
        raise SystemExit(__doc__)
    stage = sys.argv[1]
    VALUED = {"--only"}
    args, skip = [], False
    for a in sys.argv[2:]:
        if skip:
            skip = False
            continue
        if a.startswith("--"):
            skip = a in VALUED
            continue
        args.append(a)
    reps = int(args[0]) if args else (100 if stage == "cal" else 400)
    batch = int(args[1]) if len(args) > 1 else 50
    cap = int(args[2]) if len(args) > 2 else 0
    part = args[3] if len(args) > 3 else "short"
    resume = "--resume" in sys.argv
    only = set(sys.argv[sys.argv.index("--only") + 1].split(",")) if "--only" in sys.argv else None
    fams = [f for f in FAMILIES if only is None or f in only]

    for d in ("out_bandmech_cal", "out_bandmech", "logs"):
        os.makedirs(d, exist_ok=True)
    conds = [c for f in fams for c in conditions(f)]
    ncal = sum(1 for c in conds if c[2])
    print(f"families: {', '.join(fams)}")
    print(f"  {len(conds)} conditions, {ncal} needing e-calibration")

    if stage == "cal":
        tasks = []
        for fam, tag, needs, flags in conds:
            if not needs:
                continue
            for e in PROBES:
                for s0 in range(0, reps, batch):
                    n = min(batch, reps - s0)
                    out = f"out_bandmech_cal/{tag}_e{e:g}_s{s0}.csv"
                    if resume and os.path.exists(out) and sum(1 for _ in open(out)) >= n + 1:
                        continue
                    fl = " ".join(flags_to_cmd({**flags, "--e": e}))
                    tasks.append(f"--reps {n} --seed0 {51000 + s0} --gens {GENS} --procs 1 "
                                 f"--tag {tag}_e{e:g}_s{s0} --outfile {out} {fl}")
        if not tasks:
            print("nothing to calibrate")
            return
        n = write_array("bandmech_cal", tasks, cap, part)
        print(f"\nwrote bandmech_cal_params.txt ({n} tasks)")
        print("submit:  sbatch bandmech_cal_array.sh")
        return

    e_star, cal_rows = {}, []
    for fam, tag, needs, flags in conds:
        if not needs:
            continue
        est, slope = fit_e_star(probe_autosomes(tag))
        if est is not None:
            e_star[tag] = est
            cal_rows.append(dict(family=fam, tag=tag, e_star=est))
    if cal_rows:
        with open("bandmech_calibration.csv", "w", newline="") as f:
            w = csv.DictWriter(f, fieldnames=list(cal_rows[0]))
            w.writeheader(); w.writerows(cal_rows)
        print(f"\nwrote bandmech_calibration.csv ({len(cal_rows)} conditions)")
        for r in cal_rows:
            print(f"  {r['tag']:22s} e*={r['e_star']:.3f}")

    tasks, skipped = [], 0
    for fam, tag, needs, flags in conds:
        if needs:
            if tag not in e_star:
                continue
            flags = {**flags, "--e": e_star[tag]}
        for s0 in range(0, reps, batch):
            n = min(batch, reps - s0)
            out = f"out_bandmech/{tag}_s{s0}.csv"
            if resume and os.path.exists(out) and sum(1 for _ in open(out)) >= n + 1:
                skipped += 1
                continue
            fl = " ".join(flags_to_cmd(flags))
            tasks.append(f"--reps {n} --seed0 {61000 + s0} --gens {GENS} --procs 1 "
                         f"--tag {tag}_s{s0} --outfile {out} {fl}")
    if not tasks:
        print("nothing to submit")
        return
    n = write_array("bandmech_prod", tasks, cap, part)
    print(f"\nwrote bandmech_prod_params.txt ({n} tasks, {reps} reps per condition)")
    if resume:
        print(f"  RESUME: skipped {skipped}")
    for f in fams:
        print(f"  {f:6s} {len([c for c in conds if c[0] == f]):4d} conditions")
    print("submit:  sbatch bandmech_prod_array.sh")


if __name__ == "__main__":
    main()
```
