## Supplementary Information section 3 code base for "Hypothesis: A modern human range expansion ∼300,000 years ago had a critical role in Neandertal origins": bandprod_collect.py

```
#!/usr/bin/env python3
"""
bandprod_collect.py -- pool the O2 production outputs into one table.

Reads out_bandprod/<scenario>_<sweep><value>_s<seed0>.csv, pools batches per condition, and
writes band_suite_summary_o2.csv with, per condition:

    n, e_used, P(sweep), P(mt fixed), P(Y fixed), P(joint), P(joint | established),
    mean mt / Y / autosomes (unconditional and conditional on establishment),
    median end generation (how much the fix-or-lose early stop saved)

Prints the four scenarios as readable blocks.

    python bandprod_collect.py
"""
import csv
import glob
import os
import re
from collections import defaultdict

import numpy as np

HERE = os.path.dirname(os.path.abspath(__file__))
SCEN_LABEL = {
    "fexp": "(a) matrilocal range expansion        beta=0,   r=2",
    "nexp": "(b) non-sex-biased range expansion    beta=0.5, r=2",
    "fsb":  "(c) matrilocal stable boundary        beta=0,   r=1, 5% pulse",
    "nsb":  "(d) non-sex-biased stable boundary    beta=0.5, r=1, 5% pulse",
}


def main():
    e_used = {}
    cal = os.path.join(HERE, "bandprod_calibration.csv")
    if os.path.exists(cal):
        for r in csv.DictReader(open(cal)):
            e_used[(r["scenario"], r["sweep"], float(r["value"]))] = float(r["e_star"])

    pooled = defaultdict(list)
    for path in glob.glob(os.path.join(HERE, "out_bandprod", "*_s*.csv")):
        m = re.match(r"(fexp|nexp|fsb|nsb)_(s|theta)([0-9.]+)_s\d+\.csv$",
                     os.path.basename(path))
        if not m:
            continue
        key = (m.group(1), m.group(2), float(m.group(3)))
        with open(path) as f:
            pooled[key] += [r for r in csv.DictReader(f) if r["died"] == "0"]

    if not pooled:
        raise SystemExit("no outputs in out_bandprod/ -- has bandprod_array.sh finished?")

    rows = []
    for (sc, sweep, val), recs in sorted(pooled.items()):
        mt = np.array([float(r["mt"]) for r in recs])
        Y = np.array([float(r["Y"]) for r in recs])
        au = np.array([float(r["auto"]) for r in recs])
        sw = np.array([float(r["sweep_gen"]) for r in recs])
        eg = np.array([float(r["end_gen"]) for r in recs])
        est = sw >= 0 if sc in ("fexp", "nexp") else np.ones(len(recs), bool)
        fix = lambda x: x > 0.999                                   # noqa: E731
        joint = fix(mt) & fix(Y)
        n = len(recs)
        rows.append(dict(
            scenario=sc, sweep=sweep, value=val, n=n,
            e_used=e_used.get((sc, sweep, val), 0.2),
            p_sweep=round(float(est.mean()), 4),
            p_mt=round(float(fix(mt).mean()), 4),
            p_Y=round(float(fix(Y).mean()), 4),
            p_joint=round(float(joint.mean()), 4),
            se_joint=round(float(np.sqrt(max(joint.mean() * (1 - joint.mean()), 1e-9) / n)), 4),
            p_joint_est=round(float(joint[est].mean()), 4) if est.any() else float("nan"),
            mt=round(float(mt.mean()), 4), Y=round(float(Y.mean()), 4),
            aut=round(float(au.mean()), 4),
            aut_est=round(float(au[est].mean()), 4) if est.any() else float("nan"),
            end_gen_med=int(np.median(eg))))

    out = os.path.join(HERE, "band_suite_summary_o2.csv")
    with open(out, "w", newline="") as f:
        w = csv.DictWriter(f, fieldnames=list(rows[0]))
        w.writeheader(); w.writerows(rows)

    for sc in ("fexp", "nexp", "fsb", "nsb"):
        sub = [r for r in rows if r["scenario"] == sc]
        if not sub:
            continue
        print(f"\n  {SCEN_LABEL[sc]}")
        for sweep in ("s", "theta"):
            ss = sorted([r for r in sub if r["sweep"] == sweep], key=lambda r: -r["value"])
            if not ss:
                continue
            print(f"    sweep = {sweep}")
            print(f"      {'value':>7} {'e*':>6} {'n':>5} {'P(sweep)':>9} {'P(mt)':>7} "
                  f"{'P(Y)':>7} {'JOINT%':>8} {'+-':>5} {'aut%':>6} {'endgen':>7}")
            for r in ss:
                print(f"      {r['value']:>7g} {r['e_used']:>6.3f} {r['n']:>5d} "
                      f"{r['p_sweep']:>9.2f} {r['p_mt']:>7.2f} {r['p_Y']:>7.2f} "
                      f"{100*r['p_joint']:>8.1f} {100*r['se_joint']:>5.1f} "
                      f"{100*r['aut']:>6.1f} {r['end_gen_med']:>7d}")
    bad = [r for r in rows if r["scenario"] in ("fexp", "nexp")
           and not 0.04 <= r["aut"] <= 0.06]
    if bad:
        print(f"\n  WARNING: {len(bad)} calibrated conditions outside 4-6% autosomes:")
        for r in bad:
            print(f"    {r['scenario']} {r['sweep']}={r['value']:g} -> {100*r['aut']:.1f}%")
    print(f"\n  saved {os.path.basename(out)}  ({len(rows)} conditions)")


if __name__ == "__main__":
    main()
```
