## Supplementary Information section 3 code base for "Hypothesis: A modern human range expansion ∼300,000 years ago had a critical role in Neandertal origins": bandprod_submit.py

```
#!/usr/bin/env python3
"""
bandprod_submit.py -- STAGE 2 of the band-model suite on O2.

Reads the stage-1 probe outputs in out_bandcal/, fits e* per condition (log-linear fit of
autosomal modern ancestry on log(e), inverted at the 5% target, conditioned on establishment),
writes bandprod_calibration.csv for the record, and emits the production array: every
condition -- including the two stable-boundary scenarios, which need no calibration -- at
--reps replicates.

    python bandprod_submit.py [reps] [batch] [concurrency] [partition] [--resume]
    sbatch bandprod_array.sh
"""
import csv
import glob
import os
import re
import sys

import numpy as np

HERE = os.path.dirname(os.path.abspath(__file__))
sys.path.insert(0, HERE)
from band_suite import SCEN, S_GRID, TH_GRID, GENS, flags_to_cmd   # noqa: E402
from bandcal_submit import conditions                              # noqa: E402

# Settable from the command line as --target, because the autosomal proportion the calibration
# aims at is a modelling choice, not a constant: DR's relaxed analysis (9 Sep 2026) asks for
# 0.10 as well as 0.05. Refitting at a new target needs NO new probe runs -- fit_e_star reads
# the stage-1 probes already in out_bandcal/ -- so this is a seconds-long rerun of the fit.
TARGET = 0.05
if "--target" in sys.argv:
    TARGET = float(sys.argv[sys.argv.index("--target") + 1])
CAL_OUT = f"bandprod_calibration{'' if abs(TARGET - 0.05) < 1e-9 else '_%gpct' % (100 * TARGET)}.csv"


def probe_autosomes(sc, sweep, val):
    """{e: mean autosomal ancestry} from every stage-1 batch for this condition"""
    pat = os.path.join(HERE, "out_bandcal", f"{sc}_{sweep}{val:g}_e*_s*.csv")
    by_e = {}
    for path in glob.glob(pat):
        m = re.search(r"_e([0-9.]+)_s\d+\.csv$", path)
        if not m:
            continue
        with open(path) as f:
            rows = [r for r in csv.DictReader(f) if r["died"] == "0"]
        by_e.setdefault(float(m.group(1)), []).extend(rows)
    out = {}
    for e, rows in by_e.items():
        est = [r for r in rows if float(r["sweep_gen"]) >= 0]
        use = est if len(est) >= max(10, len(rows) // 3) else rows
        if use:
            out[e] = (float(np.mean([float(r["auto"]) for r in use])), len(use), len(rows))
    return out


def fit_e_star(by_e):
    """Log-linear fit of autosomal ancestry on log(e), inverted at TARGET.

    REFUSES TO EXTRAPOLATE (added 9 Sep 2026). The old version clipped the answer into
    [0.05, 0.9]. That is silent and dangerous: retargeting from 5% to 10% put the target ABOVE
    every probe for the whole s sweep -- those probes were chosen to bracket 5% and top out at
    6.5-8.4% -- so the inversion returned 0.02-0.048 and the clip turned all nineteen of them
    into an identical, entirely fictitious e* = 0.050. Nineteen rows agreeing exactly is the
    only reason it was caught.

    Now a target outside the probed range returns None, the condition is reported as
    uncalibrated, and bandratchet_submit skips it with a warning rather than running a sweep at
    a made-up exogamy rate."""
    if len(by_e) < 2:
        return None, None
    es = sorted(by_e)
    x = np.log(es)
    y = np.array([by_e[e][0] for e in es])
    b, c = np.polyfit(x, y, 1)
    if b >= -1e-6:
        return float(es[int(np.argmin(np.abs(y - TARGET)))]), b
    # the target must lie inside the range the probes actually reached
    if not (y.min() <= TARGET <= y.max()):
        return None, b
    est = float(np.exp((TARGET - c) / b))
    # ...and the answer must lie inside the range of e actually probed
    if not (min(es) <= est <= max(es)):
        return None, b
    return round(est, 3), b


def main():
    args, _skip = [], False
    for a in sys.argv[1:]:                     # --target takes a value; do not read it as reps
        if _skip:
            _skip = False; continue
        if a == "--target":
            _skip = True; continue
        if not a.startswith("--"):
            args.append(a)
    reps = int(args[0]) if args else 400
    batch = int(args[1]) if len(args) > 1 else 50
    cap = int(args[2]) if len(args) > 2 else 0
    part = args[3] if len(args) > 3 else "short"
    resume = "--resume" in sys.argv
    os.makedirs("out_bandprod", exist_ok=True)
    os.makedirs("logs", exist_ok=True)

    # ---- stage-1 fits
    cal_rows, e_star = [], {}
    missing = []
    for sc, sweep, val, flags in conditions():
        by_e = probe_autosomes(sc, sweep, val)
        est, slope = fit_e_star(by_e)
        if est is None:
            lo = min(v[0] for v in by_e.values()) if by_e else float("nan")
            hi = max(v[0] for v in by_e.values()) if by_e else float("nan")
            missing.append(f"{sc} {sweep}={val:g} (probes span {100*lo:.1f}-{100*hi:.1f}%, "
                           f"target {100*TARGET:g}% is outside)")
            continue
        e_star[(sc, sweep, val)] = est
        cal_rows.append(dict(scenario=sc, sweep=sweep, value=val, e_star=est,
                             slope=round(float(slope), 5), n_probes=len(by_e),
                             probes=";".join(f"{e:g}:{by_e[e][0]*100:.2f}%"
                                             for e in sorted(by_e))))
    if missing:
        print(f"WARNING: no usable probes for {len(missing)} conditions: "
              f"{', '.join(missing[:6])}{' ...' if len(missing) > 6 else ''}")
        print("  (stage 1 may still be running -- rerun this script when bandcal finishes)")
    with open(CAL_OUT, "w", newline="") as f:
        if cal_rows:
            w = csv.DictWriter(f, fieldnames=list(cal_rows[0]))
            w.writeheader(); w.writerows(cal_rows)
    print(f"wrote {CAL_OUT} ({len(cal_rows)} calibrated conditions, target {100*TARGET:g}% autosomal)")
    for r in cal_rows:
        print(f"  {r['scenario']:5s} {r['sweep']:5s}={r['value']:<6g} e*={r['e_star']:.3f}"
              f"   probes {r['probes']}")

    # ---- production tasks: calibrated expansions + uncalibrated stable boundaries
    all_conds = list(conditions())
    for sc in ("fsb", "nsb"):
        base = SCEN[sc]
        for s in S_GRID:
            all_conds.append((sc, "s", s, {**base, "--s": s, "--theta": 1.0}))
        for th in TH_GRID:
            if th == 1.0:
                continue
            all_conds.append((sc, "theta", th, {**base, "--s": 0.0, "--theta": th}))

    tasks, skipped = [], 0
    for sc, sweep, val, flags in all_conds:
        if sc in ("fsb", "nsb"):
            e = 0.2
        elif (sc, sweep, val) in e_star:
            e = e_star[(sc, sweep, val)]
        else:
            continue                      # uncalibrated: skip until stage 1 finishes
        for s0 in range(0, reps, batch):
            n = min(batch, reps - s0)
            out = f"out_bandprod/{sc}_{sweep}{val:g}_s{s0}.csv"
            if resume and os.path.exists(out) and sum(1 for _ in open(out)) >= n + 1:
                skipped += 1
                continue
            fl = " ".join(flags_to_cmd({**flags, "--e": e}))
            tasks.append(f"--reps {n} --seed0 {9000 + s0} --gens {GENS} "
                         f"--procs 1 --outfile {out} {fl}")
    if not tasks:
        print("nothing to submit")
        return
    open("bandprod_params.txt", "w").write("\n".join(tasks) + "\n")
    n = len(tasks)
    cap = min(n, cap) if cap else 0
    open("bandprod_array.sh", "w").write(f"""#!/bin/bash
#SBATCH -J bandprod
#SBATCH -p {part}
#SBATCH -t 0-05:59
#SBATCH -c 1
#SBATCH --mem=2G
#SBATCH -o logs/bandprod_%A_%a.log
#SBATCH --array=0-{n - 1}{("%%%d" % cap) if cap else ""}

LINE=$(sed -n "$((SLURM_ARRAY_TASK_ID + 1))p" bandprod_params.txt)
echo "task $SLURM_ARRAY_TASK_ID : $LINE"
python band_reps.py $LINE
""")
    print(f"\nwrote bandprod_params.txt ({n} tasks, {reps} reps per condition)")
    if resume:
        print(f"  RESUME: skipped {skipped} complete tasks")
    print("submit:  sbatch bandprod_array.sh")
    print("then:    python bandprod_collect.py")


if __name__ == "__main__":
    main()
```
