## Supplementary Information section 3 code base for "Hypothesis: A modern human range expansion ∼300,000 years ago had a critical role in Neandertal origins": bandratchet_collect.py

```
#!/usr/bin/env python3
"""
bandprod_collect.py -- pool the O2 production outputs into one table.

Reads out_bandratchet/<scenario>_<sweep><value>_s<seed0>.csv, pools batches per condition, and
writes band_ratchet_summary.csv with, per condition:

    n, e_used, P(sweep), P(mt fixed), P(Y fixed), P(joint), P(joint | established),
    mean mt / Y / autosomes (unconditional and conditional on establishment),
    median end generation (how much the fix-or-lose early stop saved)

Prints the four scenarios as readable blocks.

    python bandprod_collect.py
"""
import csv
import glob
import sys
import os
import re
from collections import defaultdict

import numpy as np

HERE = os.path.dirname(os.path.abspath(__file__))
SCEN_LABEL = {
    "fexp":   "(a) matrilocal range expansion       beta=0,   r=2",
    "nexp":   "(b) non-sex-biased range expansion   beta=0.5, r=2",
    "fpulse": "(c) MIGRATION PULSE, all-female      beta=0,   r=1, 5% pulse (no modern Y in)",
    "npulse": "(d) MIGRATION PULSE, unbiased        beta=0.5, r=1, 5% pulse",
}


def main():
    e_used = {}
    cal = os.path.join(HERE, "bandprod_calibration.csv")
    if os.path.exists(cal):
        for r in csv.DictReader(open(cal)):
            e_used[(r["scenario"], r["sweep"], float(r["value"]))] = float(r["e_star"])

    pooled = defaultdict(list)
    # --dir selects the calibration variant; the summary name follows it, so a 10%-calibrated
    # collect cannot overwrite band_ratchet_summary.csv
    _d = sys.argv[sys.argv.index("--dir") + 1] if "--dir" in sys.argv else "out_bandratchet"
    for path in glob.glob(os.path.join(HERE, _d, "*_s*.csv")):
        m = re.match(r"(fexp|nexp|fpulse|npulse)_(s|theta)([0-9.]+)_s\d+\.csv$",
                     os.path.basename(path))
        if not m:
            continue
        key = (m.group(1), m.group(2), float(m.group(3)))
        with open(path) as f:
            pooled[key] += [r for r in csv.DictReader(f) if r["died"] == "0"]

    if not pooled:
        raise SystemExit(f"no outputs in {_d}/ -- has the array finished?")

    rows = []
    for (sc, sweep, val), recs in sorted(pooled.items()):
        mt = np.array([float(r["mt"]) for r in recs])
        Y = np.array([float(r["Y"]) for r in recs])
        au = np.array([float(r["auto"]) for r in recs])
        sw = np.array([float(r["sweep_gen"]) for r in recs])
        eg = np.array([float(r["end_gen"]) for r in recs])
        smt = np.array([float(r.get("s_mt_drawn", 0) or 0) for r in recs])
        # DR's sampling metric: chance that 10 random males are all modern at mtDNA AND Y
        p10 = np.array([float(r["p10"]) for r in recs if r.get("p10") not in (None, "", "nan")])
        est = sw >= 0 if sc in ("fexp", "nexp") else np.ones(len(recs), bool)
        fix = lambda x: x > 0.999                                   # noqa: E731
        joint = fix(mt) & fix(Y)
        n = len(recs)
        rows.append(dict(
            scenario=sc, sweep=sweep, value=val, n=n,
            e_used=e_used.get((sc, sweep, val), 0.2),
            p_sweep=round(float(est.mean()), 4),
            p_mt=round(float(fix(mt).mean()), 4),
            p_Y=round(float(fix(Y).mean()), 4),
            p_joint=round(float(joint.mean()), 4),
            se_joint=round(float(np.sqrt(max(joint.mean() * (1 - joint.mean()), 1e-9) / n)), 4),
            p_joint_est=round(float(joint[est].mean()), 4) if est.any() else float("nan"),
            mt=round(float(mt.mean()), 4), Y=round(float(Y.mean()), 4),
            aut=round(float(au.mean()), 4),
            aut_est=round(float(au[est].mean()), 4) if est.any() else float("nan"),
            end_gen_med=int(np.median(eg)),
            s_mt_med=round(float(np.median(smt)), 5),
            s_mt_neg=round(float((smt <= 0).mean()), 3),
            p10=round(float(p10.mean()), 4) if p10.size else float("nan"),
            se_p10=round(float(p10.std() / max(np.sqrt(p10.size), 1)), 4)
            if p10.size else float("nan")))

    out = os.path.join(HERE, "band_ratchet_summary"
                   + _d.replace("out_bandratchet", "") + ".csv")
    with open(out, "w", newline="") as f:
        w = csv.DictWriter(f, fieldnames=list(rows[0]))
        w.writeheader(); w.writerows(rows)

    for sc in ("fexp", "nexp", "fpulse", "npulse"):
        sub = [r for r in rows if r["scenario"] == sc]
        if not sub:
            continue
        print(f"\n  {SCEN_LABEL[sc]}")
        for sweep in ("s", "theta"):
            ss = sorted([r for r in sub if r["sweep"] == sweep], key=lambda r: -r["value"])
            if not ss:
                continue
            print(f"    sweep = {sweep}")
            print(f"      {'value':>7} {'e*':>6} {'n':>5} {'P(sweep)':>9} {'P(mt)':>7} "
                  f"{'P(Y)':>7} {'P10%':>7} {'+-':>5} {'mt%':>6} {'Y%':>6} {'aut%':>6} "
                  f"{'endgen':>7} {'s_mt':>7}")
            for r in ss:
                print(f"      {r['value']:>7g} {r['e_used']:>6.3f} {r['n']:>5d} "
                      f"{r['p_sweep']:>9.2f} {r['p_mt']:>7.2f} {r['p_Y']:>7.2f} "
                      f"{100*r['p10']:>7.1f} {100*r['se_p10']:>5.1f} "
                      f"{100*r['mt']:>6.1f} {100*r['Y']:>6.1f} {100*r['aut']:>6.1f} "
                      f"{r['end_gen_med']:>7d} {r['s_mt_med']:>7.4f}")
    bad = [r for r in rows if r["scenario"] in ("fexp", "nexp")
           and not 0.04 <= r["aut"] <= 0.06]
    if bad:
        print(f"\n  WARNING: {len(bad)} calibrated conditions outside 4-6% autosomes:")
        for r in bad:
            print(f"    {r['scenario']} {r['sweep']}={r['value']:g} -> {100*r['aut']:.1f}%")
    print(f"\n  saved {os.path.basename(out)}  ({len(rows)} conditions)")


if __name__ == "__main__":
    main()
```
