## Supplementary Information section 3 code base for "Hypothesis: A modern human range expansion ∼300,000 years ago had a critical role in Neandertal origins": bandratchet_submit.py

```
#!/usr/bin/env python3
"""
bandratchet_submit.py -- the band suite rerun WITH the Muller's-ratchet mtDNA fitness gap.

Every replicate draws s_mt from the accepted ABC posterior (mt_gap_posterior_NA1500.txt),
exactly as cluster/joint_run.py does, so the reported probabilities integrate the gap
posterior rather than conditioning on its median. This is the arm that is comparable to the
deme series; the existing s_mt = 0 outputs in out_bandprod/ are the no-ratchet contrast.

Four scenarios, matching joint_run.py's definitions:
  fexp   matrilocal range expansion       beta=0,   Istanbul seed, r=2
  nexp   non-sex-biased range expansion   beta=0.5, Istanbul seed, r=2
  fpulse female-biased MIGRATION PULSE    beta=0,   r=1, 5% ALL-FEMALE pulse (pulse_beta=1)
  npulse unbiased MIGRATION PULSE         beta=0.5, r=1, 5% unbiased pulse (pulse_beta=0.5)

MIGRATION PULSE, as specified by DR (4 Sep 2026): the landscape starts entirely archaic; 5% of
INDIVIDUALS, drawn at random with no regard to the geographic position of their band, have
their genome replaced by a modern one. A single sex-bias parameter (pulse_beta, the same
female-share beta convention used for exogamy) sets the relative probability that a replaced
individual is female vs male, and applies to the initial condition only. The simulation then
runs, and proportions are counted globally once both mtDNA and Y are absorbed or at 6993
generations, whichever comes first -- identity is assumed lost by then, so there is no
subsetting by band. pulse_beta = 1 reproduces the deme comparator's fsb (male_frac = 0), in
which no modern Y ever enters.

Expansion e* is reused from bandprod_calibration.csv (s_mt ~ 0.2% shifts autosomes very
little); the collector reports realized autosomes so any cell that drifts off 5% is visible.
Pulse scenarios need no calibration -- the 5% pulse pins autosomal ancestry by construction.

    python bandratchet_submit.py [reps] [batch] [concurrency] [partition] [--resume]
    sbatch bandratchet_array.sh
    python bandratchet_collect.py
"""
import csv
import os
import sys

HERE = os.path.dirname(os.path.abspath(__file__))
sys.path.insert(0, HERE)
from band_suite import S_GRID, TH_GRID, GENS, N_TARGET, flags_to_cmd   # noqa: E402

POSTERIOR = "mt_gap_posterior_NA1500.txt"

SCEN = {
    "fexp":   {"--beta": 0.0, "--r": 2.0, "--N": N_TARGET},
    "nexp":   {"--beta": 0.5, "--r": 2.0, "--N": N_TARGET},
    "fpulse": {"--beta": 0.0, "--r": 1.0, "--stable-boundary": None,
               "--pulse-beta": 1.0, "--pulse-mode": "individual", "--N": N_TARGET},
    "npulse": {"--beta": 0.5, "--r": 1.0, "--stable-boundary": None,
               "--pulse-beta": 0.5, "--pulse-mode": "individual", "--N": N_TARGET},
}
# expansion cells reuse the stage-2 calibration; pulses are pinned by the 5% pulse itself
CAL_FROM = {"fexp": "fexp", "nexp": "nexp"}


# --cal lets the sweep run against a calibration fitted to a different autosomal target. The
# output directory is keyed off the same name, so a 10%-calibrated run cannot overwrite the
# 5%-calibrated one -- these differ ONLY in e, so identical filenames would silently mix two
# calibrations into one summary, which is the exact failure mode that produced the stale
# scanb numbers on 4 Sep.
CAL_FILE = "bandprod_calibration.csv"
if "--cal" in sys.argv:
    CAL_FILE = sys.argv[sys.argv.index("--cal") + 1]
SUFFIX = "" if CAL_FILE == "bandprod_calibration.csv" else \
    "_" + os.path.basename(CAL_FILE).replace("bandprod_calibration_", "").replace(".csv", "")
OUTDIR = "out_bandratchet" + SUFFIX


def load_cal():
    path = os.path.join(HERE, CAL_FILE)
    if not os.path.exists(path):
        raise SystemExit(f"{CAL_FILE} not found -- run bandprod_submit.py first")
    return {(r["scenario"], r["sweep"], float(r["value"])): float(r["e_star"])
            for r in csv.DictReader(open(path))}


def conditions():
    out = []
    for sc in SCEN:
        base = SCEN[sc]
        for s in S_GRID:
            out.append((sc, "s", s, {**base, "--s": s, "--theta": 1.0}))
        for th in TH_GRID:
            if th == 1.0:
                continue
            out.append((sc, "theta", th, {**base, "--s": 0.0, "--theta": th}))
    return out


def main():
    args, _skip = [], False
    for a in sys.argv[1:]:
        if _skip:
            _skip = False; continue
        if a == "--cal":
            _skip = True; continue
        if not a.startswith("--"):
            args.append(a)
    reps = int(args[0]) if args else 400
    batch = int(args[1]) if len(args) > 1 else 50
    cap = int(args[2]) if len(args) > 2 else 0
    part = args[3] if len(args) > 3 else "short"
    resume = "--resume" in sys.argv
    os.makedirs(OUTDIR, exist_ok=True)
    os.makedirs("logs", exist_ok=True)
    cal = load_cal()

    tasks, skipped, missing = [], 0, []
    for sc, sweep, val, flags in conditions():
        if sc in CAL_FROM:
            key = (CAL_FROM[sc], sweep, val)
            if key not in cal:
                missing.append(f"{sc} {sweep}={val:g}")
                continue
            e = cal[key]
        else:
            e = 0.2
        for s0 in range(0, reps, batch):
            n = min(batch, reps - s0)
            out = f"{OUTDIR}/{sc}_{sweep}{val:g}_s{s0}.csv"
            if resume and os.path.exists(out) and sum(1 for _ in open(out)) >= n + 1:
                skipped += 1
                continue
            fl = " ".join(flags_to_cmd({**flags, "--e": e}))
            tasks.append(f"--reps {n} --seed0 {20000 + s0} --gens {GENS} --procs 1 "
                         f"--s-mt-posterior {POSTERIOR} --tag {sc}_{sweep}{val:g}_s{s0} "
                         f"--outfile {out} {fl}")
    if missing:
        print(f"WARNING: no calibration for {len(missing)}: {', '.join(missing[:5])}")
    if not tasks:
        print("nothing to do")
        return
    open("bandratchet_params.txt", "w").write("\n".join(tasks) + "\n")
    n = len(tasks)
    cap = min(n, cap) if cap else 0
    open("bandratchet_array.sh", "w").write(f"""#!/bin/bash
#SBATCH -J bandratchet
#SBATCH -p {part}
#SBATCH -t 0-05:59
#SBATCH -c 1
#SBATCH --mem=2G
#SBATCH -o logs/bandratchet_%A_%a.log
#SBATCH --array=0-{n - 1}{("%%%d" % cap) if cap else ""}

LINE=$(sed -n "$((SLURM_ARRAY_TASK_ID + 1))p" bandratchet_params.txt)
echo "task $SLURM_ARRAY_TASK_ID : $LINE"
python band_reps.py $LINE
""")
    print(f"wrote bandratchet_params.txt ({n} tasks, {len(conditions())} conditions "
          f"x {reps} reps)")
    if resume:
        print(f"  RESUME: skipped {skipped} complete tasks")
    print(f"  every replicate draws s_mt from {POSTERIOR} (543 accepted ABC draws)")
    print("submit:  sbatch bandratchet_array.sh")
    print("then:    python bandratchet_collect.py")


if __name__ == "__main__":
    main()
```
