## Supplementary Information section 3 code base for "Hypothesis: A modern human range expansion ∼300,000 years ago had a critical role in Neandertal origins": bandsi_collect.py

```
#!/usr/bin/env python3
"""
bandsi_collect.py -- pool out_bandsi/ into one row per condition, with the deme counterpart
printed alongside wherever there is one. (DR, 5 Sep 2026)

Writes bandsi_summary.csv: family, tag, the swept parameter values parsed back out of the tag,
n, P(sweep completed), P(mt fixed), P(Y fixed), P(joint), its standard error, mean mt / Y /
autosomal ancestry, and the median stopping generation.

The autosomal column is the one to read first. Expansion cells are calibrated to 5% and any that
drift outside 4-6% are flagged; pulse cells are pinned at 5% by construction, so a pulse cell
outside that band means something is wrong with the run, not with the calibration.

    python bandsi_collect.py [--family beta]
"""
import csv
import glob
import os
import re
import sys

import numpy as np

HERE = os.path.dirname(os.path.abspath(__file__))

# tag -> (family, {parameter: value}). Kept in one place so the collector and any figure script
# agree on how a filename maps back to a cell.
PATTERNS = [
    ("beta",      r"^b(?P<beta>[-\d.]+)$"),
    ("pulsebeta", r"^pb(?P<pulse_beta>[-\d.]+)_N(?P<N>\d+)$"),
    ("mtcurve",   r"^smt(?P<s_mt>[-\d.e]+)_N(?P<N>\d+)$"),
    ("grid3b",    r"^t(?P<theta>[-\d.]+)_e(?P<e>[-\d.]+)_pm(?P<p_move>[-\d.]+)$"),
    # ALPHA ADDED 14 Sep, and its absence is why Figure S3.6's density-regulation column read
    # "runs pending" when the three cells had in fact been sitting finished in out_bandsi.
    # bandsi_submit gained the alpha panel on 13 Sep; this pattern did not, so every
    # alpha_*.csv fell through parse() and was dropped without a word. The panel list here and
    # SCAN_PANELS in bandsi_submit have to be kept in step -- they are the same set written
    # twice, which is the underlying fault.
    ("scanb",     r"^(?P<panel>e|p_move|sigma_b|sigma_e|alpha|K|r|s)_(?P<value>[-\d.]+)$"),
]
PULSE_FAMILIES = {"pulsebeta", "mtcurve"}


def parse(tag):
    for fam, pat in PATTERNS:
        m = re.match(pat, tag)
        if m:
            return fam, m.groupdict()
    return None, {}


def main():
    only = None
    if "--family" in sys.argv:
        only = sys.argv[sys.argv.index("--family") + 1]

    by_tag = {}
    for path in sorted(glob.glob(os.path.join(HERE, "out_bandsi", "*.csv"))):
        base = os.path.basename(path)[:-4]
        tag = re.sub(r"_s\d+$", "", base)
        with open(path) as f:
            by_tag.setdefault(tag, []).extend(list(csv.DictReader(f)))
    if not by_tag:
        raise SystemExit("no outputs in out_bandsi/")

    rows, off_target, unparsed = [], [], []
    for tag, raw in sorted(by_tag.items()):
        fam, params = parse(tag)
        if fam is None:
            # LOUD, NOT SILENT -- ADDED 14 Sep. A tag that matches no pattern used to be
            # skipped without a word, which is how three finished alpha cells stayed out of
            # the summary for a day while the figure said "runs pending".
            unparsed.append((tag, len(raw)))
            continue
        live = [r for r in raw if r["died"] == "0"]
        if not live:
            continue
        mt = np.array([float(r["mt"]) for r in live])
        Y = np.array([float(r["Y"]) for r in live])
        au = np.array([float(r["auto"]) for r in live])
        sw = np.array([float(r["sweep_gen"]) for r in live])
        # a pulse has no expanding wave, so "establishment" does not apply to it
        est = np.ones(len(live), bool) if fam in PULSE_FAMILIES else (sw >= 0)
        fix = lambda x: x > 0.999
        joint = fix(mt) & fix(Y)
        p = float(joint.mean())
        row = dict(family=fam, tag=tag, **params, n=len(live), n_died=len(raw) - len(live),
                   p_sweep=round(float(est.mean()), 4),
                   p_mt=round(float(fix(mt).mean()), 4), p_Y=round(float(fix(Y).mean()), 4),
                   p_joint=round(p, 4),
                   se_joint=round(float(np.sqrt(max(p * (1 - p), 1e-9) / len(live))), 4),
                   mt=round(float(mt.mean()), 4), Y=round(float(Y.mean()), 4),
                   aut=round(float(au.mean()), 4),
                   aut_est=round(float(au[est].mean()), 4) if est.any() else None,
                   end_gen_med=int(np.median([float(r.get("end_gen", 0)) for r in live])))
        rows.append(row)
        if not 0.04 <= row["aut"] <= 0.06:
            off_target.append((fam, tag, row["aut"]))

    if not rows:
        raise SystemExit("no rows matched (check --family)")
    # NB: `rows` is always ALL families, so the summary written below is complete no matter
    # what --family was given; `only` is applied to the printed tables further down.
    keys = sorted({k for r in rows for k in r})
    order = ["family", "tag"] + [k for k in keys if k not in ("family", "tag")]
    with open(os.path.join(HERE, "bandsi_summary.csv"), "w", newline="") as f:
        w = csv.DictWriter(f, fieldnames=order)
        w.writeheader()
        w.writerows(rows)
    print(f"wrote bandsi_summary.csv ({len(rows)} conditions)")

    for fam in sorted({r["family"] for r in rows}):
        if only and fam != only:
            continue
        sub = [r for r in rows if r["family"] == fam]
        print(f"\n=== {fam}  ({len(sub)} conditions) ===")
        print(f"  {'tag':24s} {'n':>5} {'P(mt)':>7} {'P(Y)':>7} {'P(joint)':>9} "
              f"{'+-':>5} {'aut%':>6}")
        for r in sorted(sub, key=lambda r: r["tag"]):
            print(f"  {r['tag']:24s} {r['n']:5d} {100*r['p_mt']:7.1f} {100*r['p_Y']:7.1f} "
                  f"{100*r['p_joint']:9.1f} {100*r['se_joint']:5.1f} {100*r['aut']:6.1f}")

    if unparsed:
        print(f"\n  !! {len(unparsed)} tag(s) matched no pattern and were DROPPED -- add them "
              f"to PATTERNS or they will never reach the summary:")
        for t, n in unparsed[:12]:
            print(f"       {t:<28} {n:,} replicate row(s)")

    if off_target:
        print(f"\nOFF-TARGET autosomal ancestry (outside 4-6%) in {len(off_target)} cells:")
        for fam, tag, a in off_target[:20]:
            why = ("pulse cell -- should be pinned at 5% by construction, so this indicates a "
                   "problem with the run" if fam in PULSE_FAMILIES else
                   "swept axis, not calibrated -- expected" if fam in ("grid3b", "scanb") else
                   "calibrated cell -- e* fit may have missed")
            print(f"  {fam:10s} {tag:24s} {100*a:5.1f}%   ({why})")
        if len(off_target) > 20:
            print(f"  ... and {len(off_target) - 20} more")


if __name__ == "__main__":
    main()
```
