## Supplementary Information section 3 code base for "Hypothesis: A modern human range expansion ∼300,000 years ago had a critical role in Neandertal origins": bandsi_submit.py

```
#!/usr/bin/env python3
"""
bandsi_submit.py -- band-model analogues of the deme-model SI families that need only
parameter sweeps of knobs bandsim.py already has. (DR, 5 Sep 2026)

Every figure and table in the introgression note is currently supported by the deme grid alone.
DR's instruction is that the band model carry a parallel analysis for each one, since the band
framework is likely to become primary. This script covers the five families that need NO new
engine code; the other five (partner locus, DMI + restorers, heritable escape, X-drive with
resistance, asymmetric DMI) plus the kin-avoidance and return-rule cultural families wait on the
mechanism loci going into bandsim.py, and are submitted by bandmech_submit.py.

    family      deme counterpart      v27 target        what is swept
    ---------------------------------------------------------------------------------
    beta        grid5                 Fig S3.1 left     exogamy sex bias, 0 -> 1
    pulsebeta   panm4                 Fig S3.1 right    migration-pulse sex bias, 0 -> 1
    mtcurve     panm3                 Fig S3.2          selection on mtDNA at the pulse
    grid3b      grid3                 Fig S3.3          F1 fertility x exogamy x mobility
    scanb       scan4                 Figure 3          sensitivity around the baseline

CALIBRATION. Expansion families need the exogamy rate e tuned per condition so that autosomal
ancestry lands on 5%; pulse families do not, because the 5% pulse pins it by construction. The
two-stage pattern is the proven one from bandcal_submit -> bandprod_submit: stage "cal" runs a
fixed probe ladder, stage "prod" fits e* by a log-linear regression of autosomal ancestry on
log(e) inverted at 5%, and emits production tasks.

Two families deliberately do NOT calibrate. grid3b sweeps e explicitly, exactly as the deme
grid3 sweeps gamma explicitly and then interpolates the value giving 5% afterwards -- calibrating
would collapse the axis the figure is about. scanb sweeps sensitivity AROUND a baseline, as
scan4 does, and reports realised autosomes so any cell that drifts off 5% is visible.

    python bandsi_submit.py cal  [reps] [batch] [cap] [partition] [--only beta,scanb]
    sbatch bandsi_cal_array.sh
    python bandsi_submit.py prod [reps] [batch] [cap] [partition] [--resume]
    sbatch bandsi_prod_array.sh
    python bandsi_collect.py
"""
import csv
import glob
import os
import re
import sys

import numpy as np

HERE = os.path.dirname(os.path.abspath(__file__))
sys.path.insert(0, HERE)
from band_suite import GENS, N_TARGET, flags_to_cmd            # noqa: E402

TARGET = 0.05
POSTERIOR = "mt_gap_posterior_NA1500.txt"

# Baseline, frozen 3-4 Sep 2026. Matches band_suite.SCEN and the census that puts measured
# autosomal Ne on the Peyregne anchor of 2,894 gene copies.
BASE_EXP = {"--r": 2.0, "--N": N_TARGET}
BASE_PULSE = {"--r": 1.0, "--stable-boundary": None, "--pulse-mode": "individual",
              "--N": N_TARGET}

# Census envelope for the pulse families, mirroring the deme side's [1000, 1500, 2000] ladder:
# the anchored value and a factor of two either side. The band anchor is 1750, not 1312, because
# bandsim and panmictic_sim reach the same measured Ne at different censuses.
N_ENVELOPE = [875, N_TARGET, 3500]

BETAS = [0.0, 0.1, 0.2, 0.3, 0.4, 0.5, 0.6, 0.7, 0.8, 0.9, 1.0]
# s_mt grid for the mtDNA fixation curves. Matches panm3_run.SVALS so the two frameworks can be
# drawn on one axis; negative values mean the ARCHAIC mtDNA is the fitter type.
S_MT = [-0.005, -0.002, -0.001, 0.0, 0.0005, 0.001, 0.002, 0.003,
        0.005, 0.0075, 0.01, 0.015, 0.02, 0.025, 0.03,
        # EXTENDED 21 Sep 2026 to match the panmictic ladder in fixation_curves_mtdna.csv.
        # The comment below says this grid "matches panm3_run.SVALS so the two frameworks can
        # be drawn on one axis"; that stopped being true when the panmictic side gained its
        # four top rungs, and Figure S3.2's band trace consequently stopped at 3% while the
        # deme trace ran to 10%. DR spotted the stub.
        0.04, 0.05, 0.075, 0.10]
# Extended toward zero on 6 Sep 2026. The whole transition sat in the 0.05 -> 0 segment:
# P(Y fixed) is 14.2% at theta = 0.05 and 89.5% at theta = 0, so the threshold the note needs
# to quote is somewhere inside a single unresolved interval. These rungs locate it. (The deme
# model has no such threshold to find -- no theta reaches 5% autosomes there at all.)
G3_THETAS = [0.0, 0.002, 0.005, 0.01, 0.02, 0.03, 0.04, 0.05, 0.07, 0.1, 0.2, 0.4, 0.7, 1.0]
# 0.10/0.16/0.22 added 9 Sep 2026. Reading the relaxed (10% autosomal) calibration off this
# grid currently lands on e = 0.13 at almost every theta, because the neighbouring probes at
# 0.08 and 0.19 straddle 10% too widely to resolve it. These three close the gap.
G3_E = [0.08, 0.10, 0.13, 0.16, 0.19, 0.22, 0.26, 0.35, 0.45, 0.56]
G3_PMOVE = [0.25, 0.45, 0.70]
# scanb panels: one knob at a time around the frozen baseline, the band counterpart of scan4's
# m / gamma / r / K / grid / sY panels.
SCAN_PANELS = {
    "e":       [0.08, 0.13, 0.19, 0.26, 0.35, 0.45, 0.56],
    "p_move":  [0.10, 0.20, 0.30, 0.45, 0.60, 0.80],
    # 30 AND 40/160 ADDED 13 Sep, AND alpha ADDED AT ALL. Appendix B states that sigma_b was
    # varied over 30, 60 and 120 km, sigma_e over 40, 80 and 160 km, and alpha over 0.4, 0.8
    # and 1.6 -- and NONE of those three sentences was true. The sweep ran sigma_b at
    # 20/40/60/90/120/180 and sigma_e at 30/50/80/120/180/240, which overlap the stated ladders
    # only at 60/120 and 80; and alpha had no panel here at all, so its whole column in
    # Figure S3.6 drew as "runs pending". The runner has always accepted --alpha-d; nobody
    # asked it to. Adding the three stated values makes the appendix's claims true rather than
    # rewriting the appendix to match what happened to run.
    "sigma_b": [20, 30, 40, 60, 90, 120, 180],
    "sigma_e": [30, 40, 50, 80, 120, 160, 180, 240],
    "alpha":   [0.4, 0.8, 1.6],
    "K":       [10, 15, 25, 40, 60, 90],
    "r":       [1.1, 1.25, 1.5, 2.0, 3.0, 4.0],
    "s":       [0.0, 0.002, 0.005, 0.01, 0.02, 0.04, 0.08, 0.16, 0.32, 0.64],
}
SCAN_FLAG = {"e": "--e", "p_move": "--p-move", "sigma_b": "--sigma-b",
             "sigma_e": "--sigma-e", "K": "--K", "r": "--r", "s": "--s",
             # the panel is named `alpha` because that is what Appendix B and
             # fig_sensitivity call it; the runner's flag is --alpha-d
             "alpha": "--alpha-d"}
def scan_base_e():
    """The calibrated fexp baseline, READ from bandprod_calibration.csv rather than written in.

    This was a literal (0.292) copied from a stale local copy of that file, and every scanb cell
    consequently ran at the wrong exogamy rate and landed at 3.2-3.8% autosomes instead of 5%.
    The file on the cluster is the only authority for e*, so take it from there and fail loudly
    if it is missing rather than falling back to a number that looks plausible."""
    path = os.path.join(HERE, "bandprod_calibration.csv")
    if not os.path.exists(path):
        raise SystemExit("bandprod_calibration.csv not found -- run bandprod_submit.py first")
    for r in csv.DictReader(open(path)):
        if r["scenario"] == "fexp" and r["sweep"] == "s" and float(r["value"]) == 0.0:
            return float(r["e_star"])
    raise SystemExit("no fexp s=0 row in bandprod_calibration.csv")

# Probe ladder for the calibrated family. Same rungs as bandcal_submit at beta = 0 / 0.5, which
# bracketed every e* inside 0.08-0.56 once the census moved to 1750.
PROBES = (0.08, 0.13, 0.19, 0.26, 0.40)


def conditions(family):
    """(family, tag, needs_calibration, flags) for every cell of a family."""
    out = []
    if family == "beta":
        for b in BETAS:
            out.append((family, f"b{b:g}", True, {**BASE_EXP, "--beta": b, "--s": 0.0,
                                                  "--theta": 1.0}))
    elif family == "pulsebeta":
        for pb in BETAS:
            for N in N_ENVELOPE:
                out.append((family, f"pb{pb:g}_N{N}", False,
                            {**BASE_PULSE, "--beta": 0.5, "--pulse-beta": pb,
                             "--s": 0.0, "--theta": 1.0, "--e": 0.2, "--N": N}))
    elif family == "mtcurve":
        for sm in S_MT:
            for N in N_ENVELOPE:
                out.append((family, f"smt{sm:g}_N{N}", False,
                            {**BASE_PULSE, "--beta": 0.5, "--pulse-beta": 0.5,
                             "--s": 0.0, "--s-mt": sm, "--theta": 1.0, "--e": 0.2, "--N": N}))
    elif family == "grid3b":
        for th in G3_THETAS:
            for e in G3_E:
                for pm in G3_PMOVE:
                    out.append((family, f"t{th:g}_e{e:g}_pm{pm:g}", False,
                                {**BASE_EXP, "--beta": 0.0, "--s": 0.0, "--theta": th,
                                 "--e": e, "--p-move": pm}))
    elif family == "scanb":
        base_e = scan_base_e()
        for panel, vals in SCAN_PANELS.items():
            for v in vals:
                fl = {**BASE_EXP, "--beta": 0.0, "--s": 0.0, "--theta": 1.0,
                      "--e": base_e}
                fl[SCAN_FLAG[panel]] = v
                out.append((family, f"{panel}_{v:g}", False, fl))
    else:
        raise SystemExit(f"unknown family {family}")
    return out


FAMILIES = ["beta", "pulsebeta", "mtcurve", "grid3b", "scanb"]


def probe_autosomes(tag):
    by_e = {}
    for path in glob.glob(os.path.join(HERE, "out_bandsi_cal", f"{tag}_e*_s*.csv")):
        m = re.search(r"_e([0-9.]+)_s\d+\.csv$", path)
        if not m:
            continue
        with open(path) as f:
            rows = [r for r in csv.DictReader(f) if r["died"] == "0"]
        by_e.setdefault(float(m.group(1)), []).extend(rows)
    out = {}
    for e, rows in by_e.items():
        est = [r for r in rows if float(r["sweep_gen"]) >= 0]
        use = est if len(est) >= max(10, len(rows) // 3) else rows
        if use:
            out[e] = float(np.mean([float(r["auto"]) for r in use]))
    return out


def fit_e_star(by_e):
    """Log-linear fit of autosomal ancestry on log(e), inverted at 5%. Returns (e*, slope)."""
    if len(by_e) < 2:
        return None, None
    es = sorted(by_e)
    b, c = np.polyfit(np.log(es), np.array([by_e[e] for e in es]), 1)
    if b >= -1e-6:      # non-monotone or flat: fall back to the nearest probe
        return float(es[int(np.argmin(np.abs([by_e[e] - TARGET for e in es])))]), b
    return round(float(np.clip(np.exp((TARGET - c) / b), 0.05, 0.9)), 3), b


def write_array(name, tasks, cap, part, mem="2G", hours="0-05:59", group=1):
    """GROUP consecutive tasks into one array element when the queue limit bites.

    O2 caps a user at 10,000 submitted array tasks across all jobs. Raising 400 replicates to
    1,000 across 556 conditions is 6,672 tasks on its own, and with the re-screen, the cultural
    sweep and the re-score already queued there is no room.

    The tempting fix -- a bigger --batch -- is WRONG here, and quietly so. Batch size sets the
    seed offset: band_reps is called with --seed0 41000+s0, so batches of 50 use seeds
    41000-41049, 41050-41099 and so on. Switching to batches of 200 would start tasks at
    41000, 41200, ... and the first of them would re-draw seeds 41000-41199, half of which the
    existing s0=0..150 files already contain. bandsi_collect pools every CSV in out_bandsi
    without checking seeds, so the duplicates would silently be counted twice and the "1,000
    replicates" would be perhaps 800 distinct ones.

    Grouping leaves the batch size, the seeds and the filenames exactly as they are and only
    changes how many of them share a SLURM task. Each still writes its own CSV and is still
    skipped by --resume, so a task that dies loses at most the batch it was running.
    """
    # `name` is the full stem, e.g. "cal" -> bandsi_cal, or "bandmech_prod" -> bandmech_prod.
    # bandmech_submit reuses this, so a hardcoded bandsi_ prefix would have written its arrays
    # over bandsi's.
    stem = name if name.startswith("bandmech") else f"bandsi_{name}"
    if group > 1:
        tasks = ["\u0000".join(tasks[i:i + group]) for i in range(0, len(tasks), group)]
    open(f"{stem}_params.txt", "w").write("\n".join(tasks) + "\n")
    n = len(tasks)
    cap = min(n, cap) if cap else 0
    open(f"{stem}_array.sh", "w").write(f"""#!/bin/bash
#SBATCH -J {stem.replace("_", "")}
#SBATCH -p {part}
#SBATCH -t {hours}
#SBATCH -c 1
#SBATCH --mem={mem}
#SBATCH -o logs/{stem}_%A_%a.log
#SBATCH --array=0-{n - 1}{("%%%d" % cap) if cap else ""}

LINE=$(sed -n "$((SLURM_ARRAY_TASK_ID + 1))p" {stem}_params.txt)
echo "task $SLURM_ARRAY_TASK_ID"
# one grouped task holds several band_reps invocations separated by NUL, split here so a
# failure in one batch does not abort the others.
# '%s\\n' AND NOT '%s' -- FIXED 13 Sep. `read` returns non-zero when it hits EOF without a
# trailing newline, and the shell tests that return value BEFORE running the loop body, so
# the final entry of every task was silently dropped. At --group 1 that is the ONLY entry:
# the task echoed its number, ran nothing, and exited COMPLETED in one second, which looks
# exactly like success in sacct. 150 tasks of the mtcurve top-up finished that way before
# anyone noticed no CSVs were appearing.
printf '%s\\n' "$LINE" | tr '\\0' '\\n' | while IFS= read -r ONE; do
  [ -z "$ONE" ] && continue
  echo "  $ONE"
  python band_reps.py $ONE
done
""")
    return n


def main():
    if len(sys.argv) < 2 or sys.argv[1] not in ("cal", "prod"):
        raise SystemExit(__doc__)
    stage = sys.argv[1]
    # Positional parsing must not be shifted by a flag's VALUE. "--only scanb" previously left
    # "scanb" in the positional list, where it was read as the concurrency cap and crashed on
    # int(). Drop each flag AND the token after it for flags that take a value.
    VALUED = {"--only", "--group"}
    args, skip = [], False
    for a in sys.argv[2:]:
        if skip:
            skip = False
            continue
        if a.startswith("--"):
            skip = a in VALUED
            continue
        args.append(a)
    reps = int(args[0]) if args else (100 if stage == "cal" else 400)
    batch = int(args[1]) if len(args) > 1 else 50
    cap = int(args[2]) if len(args) > 2 else 0
    part = args[3] if len(args) > 3 else "short"
    resume = "--resume" in sys.argv
    group = int(sys.argv[sys.argv.index("--group") + 1]) if "--group" in sys.argv else 1
    only = None
    if "--only" in sys.argv:
        only = set(sys.argv[sys.argv.index("--only") + 1].split(","))
    fams = [f for f in FAMILIES if only is None or f in only]

    os.makedirs("out_bandsi_cal", exist_ok=True)
    os.makedirs("out_bandsi", exist_ok=True)
    os.makedirs("logs", exist_ok=True)

    conds = [c for f in fams for c in conditions(f)]
    ncal = sum(1 for c in conds if c[2])
    print(f"families: {', '.join(fams)}")
    print(f"  {len(conds)} conditions, of which {ncal} need e-calibration")

    # ---------------------------------------------------------------- stage 1
    if stage == "cal":
        tasks = []
        for fam, tag, needs_cal, flags in conds:
            if not needs_cal:
                continue
            for e in PROBES:
                for s0 in range(0, reps, batch):
                    n = min(batch, reps - s0)
                    out = f"out_bandsi_cal/{tag}_e{e:g}_s{s0}.csv"
                    if resume and os.path.exists(out) and sum(1 for _ in open(out)) >= n + 1:
                        continue
                    fl = " ".join(flags_to_cmd({**flags, "--e": e}))
                    tasks.append(f"--reps {n} --seed0 {31000 + s0} --gens {GENS} --procs 1 "
                                 f"--tag {tag}_e{e:g}_s{s0} --outfile {out} {fl}")
        if not tasks:
            print("nothing to calibrate (no expansion families selected)")
            return
        n = write_array("cal", tasks, cap, part)
        print(f"\nwrote bandsi_cal_params.txt ({n} tasks: {ncal} conditions x "
              f"{len(PROBES)} probes x {reps} reps)")
        print("submit:  sbatch bandsi_cal_array.sh")
        print("then:    python bandsi_submit.py prod")
        return

    # ---------------------------------------------------------------- stage 2
    cal_rows, e_star, missing = [], {}, []
    for fam, tag, needs_cal, flags in conds:
        if not needs_cal:
            continue
        by_e = probe_autosomes(tag)
        est, slope = fit_e_star(by_e)
        if est is None:
            missing.append(tag)
            continue
        e_star[tag] = est
        cal_rows.append(dict(family=fam, tag=tag, e_star=est,
                             slope=round(float(slope), 5), n_probes=len(by_e),
                             probes=";".join(f"{e:g}:{by_e[e]*100:.2f}%" for e in sorted(by_e))))
    if cal_rows:
        with open("bandsi_calibration.csv", "w", newline="") as f:
            w = csv.DictWriter(f, fieldnames=list(cal_rows[0]))
            w.writeheader(); w.writerows(cal_rows)
        print(f"\nwrote bandsi_calibration.csv ({len(cal_rows)} conditions)")
        for r in cal_rows:
            print(f"  {r['tag']:20s} e*={r['e_star']:.3f}   probes {r['probes']}")
    if missing:
        print(f"WARNING: no usable probes for {len(missing)}: {', '.join(missing[:6])}")
        print("  (stage 1 may still be running -- rerun this script when bandsi_cal finishes)")

    tasks, skipped = [], 0
    for fam, tag, needs_cal, flags in conds:
        if needs_cal:
            if tag not in e_star:
                continue
            flags = {**flags, "--e": e_star[tag]}
        # NO ratchet posterior here. This was wrong in the first version: I drew s_mt from the
        # ABC posterior "as the deme series does", which is true of joint_run.py (the production
        # series) but NOT of the deme SI families these five are meant to parallel. scan4_run,
        # grid5_run, kin*_run, dm*_run, esc_run and xdrv*_run all run at s_mt = 0. With the
        # posterior on, the band pulsebeta cell reported P(modern mtDNA fixes) = 27.5% under an
        # all-female pulse against the deme's 10.9% -- a difference that looked like a framework
        # disagreement but was just selection present on one side and absent on the other. The
        # ratchet arm lives separately, in bandratchet, which is what Figure 2b uses.
        post = ""
        for s0 in range(0, reps, batch):
            n = min(batch, reps - s0)
            out = f"out_bandsi/{tag}_s{s0}.csv"
            if resume and os.path.exists(out) and sum(1 for _ in open(out)) >= n + 1:
                skipped += 1
                continue
            fl = " ".join(flags_to_cmd(flags))
            tasks.append(f"--reps {n} --seed0 {41000 + s0} --gens {GENS} --procs 1"
                         f"{post} --tag {tag}_s{s0} --outfile {out} {fl}")
    if not tasks:
        print("nothing to submit")
        return
    n = write_array("prod", tasks, cap, part, group=group)
    print(f"\nwrote bandsi_prod_params.txt ({n} tasks, {reps} reps per condition)")
    if resume:
        print(f"  RESUME: skipped {skipped} complete tasks")
    for f in fams:
        k = len([c for c in conds if c[0] == f])
        print(f"  {f:10s} {k:4d} conditions")
    print("submit:  sbatch bandsi_prod_array.sh")
    print("then:    python bandsi_collect.py")


if __name__ == "__main__":
    main()
```
