## Supplementary Information section 3 code base for "Hypothesis: A modern human range expansion ∼300,000 years ago had a critical role in Neandertal origins": bandsim.py

```
#!/usr/bin/env python3
"""
bandsim.py -- roving-band simulation of the modern expansion into Neandertal Europe, v0.

Implements BAND_MODEL_SPEC.md on the landscape of RANGE_MASK_METHODS.md (range_mask.npz).
Successor to cluster/clansim.py, from which it inherits two hard-won design lessons that are
deliberately preserved:
  * no size-peaked fertility (it silently blocks replacement: a shrinking band drifts back to
    the fertility peak and stabilises instead of dying) -- regulation here is a GLOBAL rescale
    plus local soft selection only;
  * daughter bands are placed at the characteristic band spacing, not at the dispersal scale
    (placing them at the dispersal scale collapsed the landscape into clumps).

MODEL (one generation)
  1. BAND MOVEMENT   each band, with prob p_move, steps N(0, sigma_b) in km; steps that leave
                     the habitable mask are rejected (the band stays put that generation).
  2. EXOGAMY         each individual independently leaves its band with the rate for its sex,
                       e_f = 2 e beta,   e_m = 2 e (1 - beta)
                     (beta = female share of exogamy propensity; no draw ever presupposes both
                     sexes are present). A leaver diffuses N(0, sigma_e) from its band's
                     position and joins the band NEAREST its landing point (its own band
                     excluded). Migrants assimilate: band identity never changes; the migrant
                     keeps its NATAL identity tag and its genes.
  3. REPRODUCTION    within-band, non-overlapping generations. Each woman's expected offspring
                     is 2 * lambda * b, where b is clansim's soft selection on LOCAL identity
                     composition (b = r/wbar for women in modern bands, 1/wbar in archaic
                     bands, wbar = r p + (1-p), p = kernel-weighted modern share within
                     sigma_e) and lambda is set each generation so the expected total equals
                     N_target -- the spec's "mechanism to keep total population size constant".
                     Fathers are drawn from the band's men, weighted by theta for F1 males
                     (cultural Haldane: an F1 is a child of parents with DIFFERENT natal
                     identities). A band lacking either sex produces nothing that generation
                     (it can be rescued by exogamy later). Loci are transmitted exactly as in
                     spatial_sim2: mtDNA maternal, Y paternal, 8 unlinked diploid autosomes,
                     2 X loci; optional additive selection s against the archaic Y and s_mt
                     against archaic mtDNA remove offspring (drop, not resample).
  4. FISSION/DEATH   a band over 2K splits into two bands of half its size, the daughter
                     displaced by the characteristic spacing; a band with no members dies.

Identity is band-level: children born into a band are members regardless of parentage, and the
child's NATAL tag is the band's identity (that is what "growing up in the band" means -- the
tag drives the cultural theta, not the genes).

Usage (smoke test):
    python bandsim.py --test
"""
import argparse
import os

import numpy as np

HERE = os.path.dirname(os.path.abspath(__file__))
AUTOS, XLOCI = 36, 3   # 32 neutral autosomes + 4 mechanism; X: drive, candidate,
                       # neutral. See spatial_sim2 for the variance argument.
KM_PER_DEG_LAT = 111.0
COS_REF = np.cos(np.deg2rad(47.0))          # local flat-earth projection reference


# ---------------------------------------------------------------- landscape
class Landscape:
    def __init__(self, path=os.path.join(HERE, "range_mask.npz")):
        z = np.load(path)
        self.lon, self.lat, self.mask = z["lon"], z["lat"], z["mask"]
        self.lon0, self.lat0 = self.lon[0], self.lat[0]
        self.step = float(self.lon[1] - self.lon[0])

    def ok(self, xy):
        """xy in km (n,2) -> boolean habitability. Positions off-grid are not habitable."""
        lon = xy[:, 0] / (KM_PER_DEG_LAT * COS_REF) + self.lon0
        lat = xy[:, 1] / KM_PER_DEG_LAT + self.lat0
        j = np.round((lon - self.lon0) / self.step).astype(int)
        i = np.round((lat - self.lat0) / self.step).astype(int)
        good = (i >= 0) & (i < self.mask.shape[0]) & (j >= 0) & (j < self.mask.shape[1])
        out = np.zeros(len(xy), bool)
        out[good] = self.mask[i[good], j[good]]
        return out

    def to_km(self, lonlat):
        return np.column_stack(((lonlat[:, 0] - self.lon0) * KM_PER_DEG_LAT * COS_REF,
                                (lonlat[:, 1] - self.lat0) * KM_PER_DEG_LAT))

    def habitable_lonlat(self):
        ii, jj = np.where(self.mask)
        return np.column_stack((self.lon[jj], self.lat[ii]))


# ---------------------------------------------------------------- config
class Config:
    def __init__(self, **kw):
        # SETTLED 4 Sep 2026. 1750 is the census at which the MEASURED autosomal effective size
        # lands on the Peyregne anchor of 1,447 diploids (2,894 gene copies) in all four band
        # configurations: fexp 2844, nexp 2777, fpulse 2807, npulse 3025, every one within 4%.
        # Was 1050, which predates the anchor and targeted 1500 by a different route. Production
        # scripts pass --N explicitly, but the default is what ad-hoc analysis picks up, so it
        # must not be the stale value (figure2_bands.py silently drew panel a at 1050 because of
        # exactly this).
        self.N_target = 1750
        self.K = 25                 # optimal band size; fission above 2K
        self.p_move = 0.45  # re-tuned at N=1050 (sweep median 495, IQR 460-580); raised from 0.1 (DR: wave must complete <=1000 gens, ideally ~500)           # band movement rate per generation
        self.sigma_b = 60   # raised from 40, same reason; completion now ~260-400 gens.0         # band step scale, km
        self.e = 0.08               # mean per-capita exogamy rate
        self.beta = 0.5             # female share of exogamy propensity
        self.sigma_e = 80   # frozen baseline (DR 3 Sep 2026): hunter-gatherer marital-distance scale; cite.0        # exogamy dispersal scale, km (also the composition kernel)
        self.r = 2.0                # modern:archaic soft-selection ratio (1.0 = neutral mixing)
        self.theta = 1.0            # relative fertility of F1 males (cultural, natal tags)
        # sel_locus ADDED 14 Sep 2026. `s` used to be selection against the archaic Y and
        # nothing else, hardwired into the survival test below. That made four architectures
        # -- the PRDM9 ones, which carry independent positive selection on PRDM9-A -- deme-only,
        # so their four thumbnails in Figure S3.3 had blue points and no red. DR: "In Figure
        # S3.3 the following thumbnails are missing red points: the four PRDM9", and then yes
        # to implementing it.
        #
        # SEMANTICS COPIED FROM spatial_sim2._archaic_count, so a band cell means what the
        # corresponding deme cell means: s is ADDITIVE in the number of archaic alleles at the
        # selected locus. A male carries one Y and one X; a female carries two X and no Y;
        # everyone carries two autosomes. "none" switches selection off entirely.
        #
        # THE DEFAULT IS "Y", WHICH REPRODUCES THE OLD BEHAVIOUR EXACTLY. At sel_locus = "Y"
        # the archaic-allele count is 1 for an archaic-Y male and 0 for everyone else, so the
        # survival test below reduces to the single Bernoulli draw it used to be -- not
        # approximately, identically. Every existing band result is therefore unaffected.
        self.s = 0.0                # additive selection against the archaic allele
        self.sel_locus = "Y"        # "Y" | "auto" | "X" | "none"
        self.auto_locus = 35        # which autosome, when sel_locus == "auto"
        self.x_locus = 1            # which X locus, when sel_locus == "X"
        self.s_mt = 0.0             # selection against archaic mtDNA

        # ---------------- SI mechanism loci (band analogues of spatial_sim2/3) -------------
        # Semantics are copied from the deme engine deliberately, so that a cell of a band
        # table means the same thing as the corresponding cell of a deme table. Two of these
        # are easy to get backwards and the difference is the whole argument of the note:
        #
        #   * the mt x Y DMI is tested on the CHILD's OWN mtDNA and Y, every generation. A
        #     grandson still carrying the archaic Y inside a modern-mtDNA matriline is struck
        #     again. This is the "sealed" architecture.
        #   * theta (above) is a PARENTAL-PAIRING property that is not inherited, so it taxes
        #     the archaic Y exactly once, as it enters. This is the "leaky" one.
        #
        # Implementing the DMI as a parental rule, or theta as a heritable flag, silently
        # swaps a sealed mechanism for a leaky one and produces entirely plausible wrong
        # numbers. See the deme spec in spatial_sim2._dm_penetrance.
        self.dm = 0.0               # penetrance of the single-pair DMI (0 = off, 1 = lethal)
        self.dm_focal = "mt"        # modern side: "mt" | "Y" | "auto" | "X"
        self.dm_partner = "Y"       # archaic side: "mt" | "Y" | "auto" | "X"
        self.dm_dom = "dominant"    # for auto/X partners: dominant | additive | recessive
        self.dm_male_limited = True
        self.dm_focal_locus = 35
        self.dm_partner_locus = 35

        # polygenic mito-nuclear restorers: the LAST n autosomal loci. Trigger is the child's
        # own modern mtDNA; dominant = OR across loci, additive = product over loci of
        # (1 - rest_pen * archaic_dose / 2). n = 1 dominant reproduces the single-pair
        # mtDNA x autosome-dominant row exactly.
        self.restorers = 0
        self.rest_pen = 1.0

        # X-linked drive at X locus 0. An expressing father (modern allele at X0 AND an
        # archaic Y) converts a fraction `xdrive` of his SONS into daughters -- transmission
        # distortion, not mortality, so the offspring count is unchanged. xdrive_cost is a flat
        # survival penalty on ALL his offspring. Resistance arises only among conceptions that
        # were about to be flipped, becomes Y == 2, and is inherited down the patriline.
        self.xdrive = 0.0
        self.xdrive_cost = 0.0
        self.xdrv_res = 0.0
        self.xdrv_all = False       # control: drive against every Y, not just the archaic one

        # heritable escape of the suppressed archaic Y. Drawn per conception but ONLY among
        # offspring the DMI was about to kill; once acquired the Y is allele 2, which is immune
        # to the DMI and to drive forever, in that individual and every patrilineal descendant.
        # Allele 2 counts as ARCHAIC in every readout.
        self.esc_eps = 0.0

        # ---------------- cultural mechanisms (band analogues of mech3/4/5 and kin2/3) -------
        # Kin avoidance. Relatedness is tracked exactly as spatial_sim2 does it and no further:
        # two integers per person, the row index of the mother (mat1) and of the maternal
        # grandmother (mat2), comparable only within a generation because generations do not
        # overlap. This is a strictly MATRILINEAL measure -- half-sibs through the father are
        # invisible to it. inbreed_ban = 1 forbids a pair sharing a mother, 2 a pair sharing a
        # maternal grandmother (which subsumes 1). After inbreed_tries failed redraws the
        # conception is ABANDONED, not substituted, so the band simply ends up smaller.
        self.inbreed_ban = 0
        self.inbreed_tries = 12

        # Return rule. rho is the probability that a MALE, having married out, instead joins the
        # band his FATHER was born in. Deme analogue: rho teleports a male to fnat, his father's
        # natal deme. Founders have no recorded paternal band and are never returned. This sorts
        # men by paternal origin, which is exactly what the Y tracks.
        self.rho_return = 0.0

        # Male reproductive skew: the fraction of men eligible to father children in a given
        # generation, redrawn each generation. 1.0 = off.
        self.skew_frac = 1.0

        # Asymmetric exogamy. The band counterpart of gamma_export / gamma_import: the relative
        # rate at which people move OUT of modern-identity bands into archaic ones (export) and
        # the reverse (import). Named from the modern side, as in spatial_sim3. None = symmetric.
        self.e_export = None
        self.e_import = None
        self.hard_selection = False  # fill slots vacated by selection with fresh draws from the
                                    # same band, as panmictic_sim/spatial_sim2 do, so that s
                                    # means the same thing in the band and deme frameworks.
                                    # False = the old drop-don't-resample behaviour, whose
                                    # realised coefficient is about half the nominal one.
        self.refill_rounds = 8
        self.alpha_d = 0.8          # local density regulation strength (0 = off). Without it
                                    # bands random-walk into stable clumps and the landscape
                                    # empties elsewhere -- the clansim clumping failure.
        self.reseed = True          # while archaic bands persist, re-found the Istanbul band
                                    # whenever no modern-identity band exists (the band-model
                                    # analogue of the deme model's guarded origin reseed: the
                                    # strait is a conduit from an off-map modern source)
        self.gens = 300
        self.seed = 1
        self.seed_lonlat = (28.95, 41.05)   # Istanbul, European shore
        self.pulse_beta = 0.5        # sex bias of the migration pulse, SAME beta convention as
                                     # exogamy: beta is the FEMALE share, so relative
                                     # probability female:male = beta : (1 - beta).
                                     # 1.0 = all-female pulse (the deme model's fsb, male_frac
                                     # = 0), 0.5 = unbiased, 0.0 = all-male. Used only to build
                                     # the initial condition; it does not affect the dynamics.
        self.pulse_mode = "individual"  # "individual": modern-origin PEOPLE replace a random
                                     # 5% of residents, ignoring geography (DR, 4 Sep 2026 --
                                     # the migration pulse is deliberately NOT spatially
                                     # explicit); "band": whole bands arrive modern (variant)
        self.stable_boundary = False # True: MIGRATION PULSE. No Istanbul seed -- the landscape
                                     # starts entirely archaic, then a fraction pulse_frac of
                                     # individuals, drawn at random regardless of where their
                                     # band sits, are replaced by modern people. r should be
                                     # set to 1 (no lineage advantage) per the spec.
        self.pulse_frac = 0.05
        self.record_every = 10
        for k, v in kw.items():
            if not hasattr(self, k):
                raise TypeError(f"unknown parameter {k!r}")
            setattr(self, k, v)


# ---------------------------------------------------------------- population state
# Individuals in flat arrays; band membership by index into the band tables.
def init_state(cfg, land, rng):
    n_bands = max(2, cfg.N_target // cfg.K)
    cells = land.habitable_lonlat()
    pos = land.to_km(cells[rng.choice(len(cells), n_bands, replace=False)])
    b_ident = np.zeros(n_bands, np.int8)        # 0 = archaic, 1 = modern
    if cfg.stable_boundary and cfg.pulse_mode == "band":
        # spatially clumped pulse: whole bands arrive modern, scattered over the range
        k = max(1, round(cfg.pulse_frac * n_bands))
        b_ident[rng.choice(n_bands, k, replace=False)] = 1
    elif cfg.stable_boundary:
        pass          # individual-mode pulse is applied below, after people exist
    else:
        # band 0 is the modern seed at Istanbul
        pos[0] = land.to_km(np.array([cfg.seed_lonlat]))[0]
        b_ident[0] = 1

    members = rng.poisson(cfg.K, n_bands).clip(min=2)
    n = members.sum()
    band = np.repeat(np.arange(n_bands), members)
    sex = rng.integers(0, 2, n).astype(np.int8)             # 1 = male
    ident = b_ident[band]                                    # natal identity = band identity
    P = dict(
        band=band, sex=sex, natal=ident.copy(), f1=np.zeros(n, bool),
        # distinct NEGATIVE sentinels so no two founders ever count as kin, and -1 for
        # "no recorded paternal band" so founders are never subject to the return rule
        mat1=-np.arange(1, n + 1, dtype=np.int64),
        mat2=-np.arange(1, n + 1, dtype=np.int64),
        fnat=np.full(n, -1, np.int64),
        mt=ident.copy(),
        Y=np.where(sex == 1, ident, -1).astype(np.int8),
        auto=np.repeat((2 * ident).astype(np.int8)[:, None], AUTOS, 1),
        X=(np.where(sex == 1, 1, 2)[:, None] * ident[:, None]).astype(np.int8)
          * np.ones((1, XLOCI), np.int8),
    )
    if cfg.stable_boundary and cfg.pulse_mode == "individual":
        # MIGRATION PULSE (DR, 4 Sep 2026). The landscape starts entirely archaic; a fraction
        # pulse_frac of INDIVIDUALS, drawn at random with no regard to where their band sits,
        # have their genome replaced by a modern one. Deliberately not spatially explicit: the
        # pulse is a one-off admixture event, and the spatial model then just carries it.
        # Sex composition follows the beta convention (beta = female share): each replaced
        # individual is female with probability pulse_beta, male with probability 1 - beta,
        # so beta = 1 reproduces the deme model's all-female fsb (male_frac = 0), in which no
        # modern Y ever enters. Band identity labels stay archaic throughout -- what is
        # measured afterwards is genome ancestry across everyone, not group membership.
        want = int(round(cfg.pulse_frac * n))
        male = sex == 1
        mi, fi = np.where(male)[0], np.where(~male)[0]
        nf = int(rng.binomial(want, float(np.clip(cfg.pulse_beta, 0.0, 1.0))))
        nf, nm = min(nf, len(fi)), min(want - nf, len(mi))
        pick = np.concatenate([rng.choice(fi, nf, replace=False),
                               rng.choice(mi, nm, replace=False)])
        P["mt"][pick] = 1
        P["natal"][pick] = 1
        ym = pick[P["sex"][pick] == 1]
        P["Y"][ym] = 1
        P["auto"][pick, :] = 2
        P["X"][pick, :] = np.where(P["sex"][pick] == 1, 1, 2)[:, None]
    return P, pos, b_ident


DRIVER_XLOCUS = 0        # X locus carrying the driver; locus 1 stays as neutral background


def _dm_penetrance(cfg, sx, mt, Yc, auto, X):
    """Probability the DMI removes this offspring, from its OWN genotype.

    Transcribed from spatial_sim2._dm_penetrance so that a band cell means the same as the
    corresponding deme cell. Two conventions that invert the result if mishandled:
      * auto and X store counts of MODERN alleles, so the archaic dose is (max - stored).
      * Y == 2 (escaped / drive-resistant) is NOT archaic here -- the test is Y == 0 -- even
        though it is reported as archaic. That asymmetry is the point of the escape mechanism.
    """
    n = len(sx)
    if auto is None or X is None:
        return np.zeros(n)

    def focal_modern():
        f = cfg.dm_focal
        if f == "mt":
            return mt == 1
        if f == "Y":
            return (sx == 1) & (Yc == 1)
        if f == "auto":
            return auto[:, cfg.dm_focal_locus] >= 1
        if f == "X":
            if not 0 <= cfg.dm_focal_locus < XLOCI:
                raise SystemExit(f"dm_focal='X' needs dm_focal_locus in 0..{XLOCI-1}, "
                                 f"got {cfg.dm_focal_locus}")
            return X[:, cfg.dm_focal_locus] >= 1
        raise ValueError(f"dm_focal={f}")

    def partner_archaic():
        p = cfg.dm_partner
        if p == "mt":
            return (mt == 0).astype(float)
        if p == "Y":
            return ((sx == 1) & (Yc == 0)).astype(float)
        if p == "auto":
            k = 2 - auto[:, cfg.dm_partner_locus].astype(float)
            maxd = np.full(n, 2.0)
        elif p == "X":
            if not 0 <= cfg.dm_partner_locus < XLOCI:
                raise SystemExit(f"dm_partner='X' needs dm_partner_locus in 0..{XLOCI-1}, "
                                 f"got {cfg.dm_partner_locus} (the default {AUTOS-1} is an "
                                 f"AUTOSOMAL index -- set it explicitly for an X partner)")
            maxd = np.where(sx == 0, 2.0, 1.0)
            k = maxd - X[:, cfg.dm_partner_locus].astype(float)
        else:
            raise ValueError(f"dm_partner={p}")
        if cfg.dm_dom == "dominant":
            return (k >= 1).astype(float)
        if cfg.dm_dom == "recessive":
            return (k == maxd).astype(float)
        return k / maxd                      # additive

    pen = cfg.dm * focal_modern().astype(float) * partner_archaic()
    if cfg.dm_male_limited:
        pen = pen * (sx == 1)
    return np.clip(pen, 0.0, 1.0)


def _restorer_survival(cfg, sx, mt, auto):
    """Survival multiplier from polygenic mito-nuclear restorers (spatial_sim3 semantics).

    The restorers are the LAST cfg.restorers autosomal loci. The trigger is the child's own
    MODERN mtDNA -- an archaic-mtDNA child is unaffected however many archaic restorer alleles
    it carries. dominant = OR across loci; additive = product ACROSS loci of the within-locus
    additive term, so with rest_pen = 1 a fully heterozygous child survives 0.5**n."""
    lo = AUTOS - cfg.restorers
    dose = (2 - auto[:, lo:]).astype(float)          # archaic allele count per restorer, 0..2
    if cfg.dm_dom == "dominant":
        surv = np.where((dose >= 1).any(1), 1.0 - cfg.rest_pen, 1.0)
    elif cfg.dm_dom == "additive":
        surv = np.prod(1.0 - cfg.rest_pen * dose / 2.0, axis=1)
    else:
        raise ValueError("restorers support dm_dom in {dominant, additive}")
    trig = (mt == 1)
    if cfg.dm_male_limited:
        trig = trig & (sx == 1)
    return np.where(trig, np.clip(surv, 0.0, 1.0), 1.0)


def _pair_transmit(P, mothers, fathers, rng, cfg=None):
    """Mendelian transmission for chosen parent index arrays; returns child locus arrays.

    If cfg carries an X-drive, a fraction of an expressing father's SONS are converted into
    DAUGHTERS here. That is transmission distortion, not mortality: the number of offspring is
    unchanged and only the sex ratio moves. A converted daughter necessarily inherits her
    father's driver X. Resistance arises only among conceptions that were about to be flipped,
    becomes Y == 2, and is then inherited down the patriline like any other Y."""
    nm = len(mothers)
    sex = rng.integers(0, 2, nm).astype(np.int8)
    mt = P["mt"][mothers]
    Y = np.where(sex == 1, P["Y"][fathers], -1).astype(np.int8)
    am, af = P["auto"][mothers], P["auto"][fathers]
    auto = (rng.random((nm, AUTOS)) < am / 2.0).astype(np.int8) + \
           (rng.random((nm, AUTOS)) < af / 2.0).astype(np.int8)
    xm = (rng.random((nm, XLOCI)) < P["X"][mothers] / 2.0).astype(np.int8)
    xf = P["X"][fathers]                     # father's single X, goes to daughters
    X = np.where(sex[:, None] == 1, xm, xm + xf)
    auto, X = auto.astype(np.int8), X.astype(np.int8)

    if cfg is not None and cfg.xdrive > 0:
        fa_x0 = P["X"][fathers][:, DRIVER_XLOCUS]
        fa_Y = P["Y"][fathers]
        # normal: the modern Y carries the coevolved suppressor and is immune, and so is the
        # resistant allele 2. xdrv_all is the control that drives against every Y.
        expressing = (fa_x0 == 1) & (fa_Y >= 0 if cfg.xdrv_all else fa_Y == 0)
        flip = (sex == 1) & expressing & (rng.random(nm) < cfg.xdrive)
        if cfg.xdrv_res > 0:
            res = flip & (rng.random(nm) < cfg.xdrv_res)
            if res.any():
                Y = np.where(res, 2, Y).astype(np.int8)
            flip = flip & ~res
        if flip.any():
            sex = np.where(flip, 0, sex).astype(np.int8)
            Y = np.where(flip, -1, Y).astype(np.int8)
            X = np.where(flip[:, None], (xm + xf).astype(np.int8), X).astype(np.int8)
    return sex, mt, Y, auto, X


def _archaic_count(cfg, sex, Y, auto, X, natal=None):
    """Archaic alleles at the selected locus, per individual, for additive selection.

    A transcription of spatial_sim2._archaic_count into the band engine's arrays, so that the
    same `s` means the same thing in both frameworks. The dosage rules are the ones biology
    imposes rather than a modelling choice: only males carry a Y, so the Y count is 0 or 1 and
    only for them; autosomes are diploid so the count runs 0 to 2; the X is diploid in females
    and haploid in males, so its maximum dose depends on sex.
    """
    loc = getattr(cfg, "sel_locus", "Y")
    if loc == "Y":
        # SELECTION ON THE Y IS CONDITIONED ON THE BAND'S IDENTITY (DR, 9 Oct 2026). The
        # mechanism is prestige, and prestige is conferred by the resident community, so it
        # attaches to the patriline that MATCHES the identity of the band a man lives in --
        # modern Y inside a modern-identity band, archaic Y inside an archaic-identity band.
        # It used to be unconditional: every archaic-Y male was selected against wherever he
        # was, so an incoming modern patriline was rewarded inside archaic bands it had no
        # standing in, which is what DR spotted in the film. Under matrilocality band identity
        # is carried by the women, so this is what "selection in the direction of the identity
        # of the sex that transmits identity" means operationally.
        #
        # Y == 2 is the heritable-escape allele: archaic for every purpose here, so the test
        # is on (Y == 1) rather than on (Y != 0).
        y_mod = (Y == 1)
        if natal is None:                      # no band context: fall back to unconditional
            return np.where((sex == 1) & ~y_mod, 1, 0)
        return np.where((sex == 1) & (y_mod != (natal == 1)), 1, 0)
    if loc == "auto":
        if auto is None:
            return np.zeros(sex.shape, np.int64)
        return 2 - auto[:, cfg.auto_locus]
    if loc == "X":
        if X is None:
            return np.zeros(sex.shape, np.int64)
        maxdose = np.where(sex == 0, 2, 1)
        return maxdose - X[:, cfg.x_locus]
    return np.zeros(sex.shape, np.int64)


def step(P, pos, b_ident, cfg, land, rng):
    n_bands = len(b_ident)
    alive_b = np.array([np.any(P["band"] == b) for b in range(n_bands)])

    # ---- 1. band movement
    movers = (rng.random(n_bands) < cfg.p_move) & alive_b
    if movers.any():
        prop = pos.copy()
        prop[movers] += rng.normal(0, cfg.sigma_b, (movers.sum(), 2))
        ok = land.ok(prop[movers])
        idx = np.where(movers)[0][ok]
        pos[idx] = prop[np.where(movers)[0][ok]]

    # ---- 2. exogamy  (mechanism helpers are defined at module scope, below step())
    e_f, e_m = 2 * cfg.e * cfg.beta, 2 * cfg.e * (1 - cfg.beta)
    rate = np.where(P["sex"] == 1, e_m, e_f)
    if cfg.e_export is not None or cfg.e_import is not None:
        # Asymmetric exogamy, the band counterpart of gamma_export / gamma_import. Named from
        # the modern side: EXPORT is a modern-band member marrying out into the archaic world,
        # IMPORT the reverse. The deme model applies the asymmetry to the mating rule; the band
        # model has no mating rule to modify, so it applies to the rate at which people leave,
        # scaled relative to the symmetric e so the two knobs remain interpretable as rates.
        in_modern = b_ident[P["band"]] == 1
        ex = cfg.e if cfg.e_export is None else cfg.e_export
        im = cfg.e if cfg.e_import is None else cfg.e_import
        rate = rate * np.where(in_modern, ex / max(cfg.e, 1e-12), im / max(cfg.e, 1e-12))
    leave = rng.random(len(rate)) < rate
    if leave.any() and alive_b.sum() > 1:
        li = np.where(leave)[0]
        land_pt = pos[P["band"][li]] + rng.normal(0, cfg.sigma_e, (len(li), 2))
        d2 = ((land_pt[:, None, :] - pos[None, :, :]) ** 2).sum(-1)
        d2[:, ~alive_b] = np.inf
        d2[np.arange(len(li)), P["band"][li]] = np.inf      # cannot rejoin own band
        P["band"][li] = np.argmin(d2, 1).astype(P["band"].dtype)
    if cfg.rho_return > 0:
        # band analogue of spatial_sim2's rho: a male returns to the band his FATHER was born
        # in. Overrides the exogamy destination, applies to all men with a recorded paternal
        # band (founders have -1), never to women. Bands that have since died are skipped.
        back = ((P["sex"] == 1) & (P["fnat"] >= 0)
                & (rng.random(len(P["band"])) < cfg.rho_return))
        if back.any():
            tgt = P["fnat"][back]
            ok = alive_b[np.clip(tgt, 0, n_bands - 1)] & (tgt < n_bands)
            idx = np.where(back)[0][ok]
            P["band"][idx] = tgt[ok].astype(P["band"].dtype)

    # ---- 3. reproduction (within band; global lambda keeps expected total at N_target)
    # local modern share p within the sigma_e kernel, from band positions and sizes
    sizes = np.bincount(P["band"], minlength=n_bands).astype(float)
    w = np.exp(-((pos[:, None, :] - pos[None, :, :]) ** 2).sum(-1) / (2 * cfg.sigma_e ** 2))
    w *= sizes[None, :]
    pmod = (w * b_ident[None, :]).sum(1) / np.maximum(w.sum(1), 1e-9)
    wbar = cfg.r * pmod + (1 - pmod)
    b_fac = np.where(b_ident == 1, cfg.r, 1.0) / wbar       # soft selection (clansim scheme)
    # local density regulation (clansim lesson): crowded neighbourhoods breed below
    # replacement, empty ones above, so bands spread instead of clumping. L is the
    # kernel-weighted local population; normalising by its current mean keeps the term
    # centred so the global lambda still sets the total.
    if cfg.alpha_d > 0:
        L = w.sum(1)
        b_fac = b_fac * np.exp(cfg.alpha_d * (1.0 - L / max(L[alive_b].mean(), 1e-9)))

    women = np.where(P["sex"] == 0)[0]
    men_mask = P["sex"] == 1
    if cfg.skew_frac < 1.0:
        # spatial_sim3 semantics: redrawn every generation, females untouched, and the SAME
        # mask feeds both the band man-weights and the father pools
        men_mask = men_mask & (rng.random(len(men_mask)) < cfg.skew_frac)
    # expected children per woman before global rescale
    exp_w = 2.0 * b_fac[P["band"][women]]
    # bands without any fertile man produce nothing
    man_wt_all = np.where(P["f1"] & men_mask, cfg.theta, 1.0) * men_mask
    band_manwt = np.zeros(n_bands)
    np.add.at(band_manwt, P["band"], man_wt_all)
    exp_w *= (band_manwt[P["band"][women]] > 0)
    lam = cfg.N_target / max(exp_w.sum(), 1e-9)
    kids_per_woman = rng.poisson(np.clip(exp_w * lam, 0, 8))
    mothers = np.repeat(women, kids_per_woman)
    if len(mothers) == 0:
        return None, pos, b_ident
    kband = P["band"][mothers]

    # fathers: sample per child from the band's men, weight theta for F1 males
    order = np.argsort(kband, kind="stable")
    mothers, kband = mothers[order], kband[order]
    fathers = np.empty(len(mothers), int)
    for b in np.unique(kband):
        men_b = np.where(men_mask & (P["band"] == b))[0]
        wts = man_wt_all[men_b]
        sl = kband == b
        if len(men_b) == 0 or wts.sum() <= 0:
            fathers[sl] = -1
            continue
        fathers[sl] = rng.choice(men_b, sl.sum(), p=wts / wts.sum())
    keep = fathers >= 0
    if cfg.inbreed_ban:
        # spatial_sim2 semantics: matrilineal relatedness only, both parents redrawn each try,
        # and after inbreed_tries failures the conception is DROPPED rather than substituted.
        anc = P["mat1"] if cfg.inbreed_ban == 1 else P["mat2"]
        bad = keep & (anc[fathers] == anc[mothers])
        for _ in range(cfg.inbreed_tries):
            if not bad.any():
                break
            for b in np.unique(kband[bad]):
                sl = bad & (kband == b)
                men_b = np.where(men_mask & (P["band"] == b))[0]
                wom_b = women[P["band"][women] == b]
                wts = man_wt_all[men_b]
                if not len(men_b) or not len(wom_b) or wts.sum() <= 0:
                    continue
                n_bad = int(sl.sum())
                fathers[sl] = rng.choice(men_b, n_bad, p=wts / wts.sum())
                mothers[sl] = rng.choice(wom_b, n_bad, replace=True)
            bad = bad & (anc[fathers] == anc[mothers])
        keep = keep & ~bad
    mothers, fathers, kband = mothers[keep], fathers[keep], kband[keep]

    sex, mt, Y, auto, X = _pair_transmit(P, mothers, fathers, rng, cfg)
    natal = b_ident[kband].astype(np.int8)                   # natal tag = band identity
    f1 = P["natal"][mothers] != P["natal"][fathers]          # cultural F1: mixed natal parents

    def _select(sex, mt, Y, auto=None, X=None):
        """Offspring viability. Every mechanism here acts on the CHILD's OWN genotype and is a
        DROP with no backfill (unless cfg.hard_selection), matching spatial_sim2. Returns
        (survivor mask, Y) -- Y is returned because heritable escape rewrites it to allele 2.

        Note Y == 2 is archaic for reporting but immune to the DMI: the tests below are
        `Y == 0`, which allele 2 fails."""
        n = len(sex)
        surv = np.ones(n, bool)
        if cfg.s > 0 and getattr(cfg, "sel_locus", "Y") != "none":
            k = _archaic_count(cfg, sex, Y, auto, X, natal=natal)
            surv &= ~(rng.random(n) < np.clip(cfg.s * k, 0.0, 1.0))
        if cfg.s_mt != 0:
            tgt = (mt == 0) if cfg.s_mt > 0 else (mt == 1)
            surv &= ~(tgt & (rng.random(n) < abs(cfg.s_mt)))
        if cfg.restorers > 0 and auto is not None:
            surv &= rng.random(n) < _restorer_survival(cfg, sex, mt, auto)
        if cfg.dm > 0:
            # separate Bernoulli rather than folded into a survival weight, because escape
            # needs to know which individuals the DMI actually struck (spatial_sim3 does the
            # same, and is statistically identical when esc_eps = 0)
            hit = rng.random(n) < _dm_penetrance(cfg, sex, mt, Y, auto, X)
            if cfg.esc_eps > 0 and hit.any():
                escaped = hit & (rng.random(n) < cfg.esc_eps)
                if escaped.any():
                    Y = np.where(escaped, 2, Y).astype(np.int8)
                hit = hit & ~escaped
            surv &= ~hit
        return surv, Y

    surv, Y = _select(sex, mt, Y, auto, X)
    if cfg.xdrive > 0 and cfg.xdrive_cost > 0:
        # applies to BOTH sexes and whether or not that conception was flipped. spatial_sim3
        # keeps the `== 0` test here even under xdrv_all; the asymmetry is reproduced so the
        # Block C control matches.
        expressing_fa = ((P["X"][fathers][:, DRIVER_XLOCUS] == 1) & (P["Y"][fathers] == 0))
        surv &= ~(expressing_fa & (rng.random(len(sex)) < cfg.xdrive_cost))
    kid = dict(band=kband, sex=sex, natal=natal, f1=f1, mt=mt, Y=Y, auto=auto, X=X,
               mat1=mothers.astype(np.int64),           # mother's row index this generation
               mat2=P["mat1"][mothers].astype(np.int64),  # maternal grandmother's
               fnat=P["band"][fathers].astype(np.int64))  # father's band = his natal band
    if cfg.hard_selection and (cfg.s > 0 or cfg.s_mt != 0) and not surv.all():
        # HARD SELECTION (matches the deme comparator). panmictic_sim.py and spatial_sim2.py
        # do not simply delete the disfavoured offspring: they keep drawing parent pairs until
        # the slot is filled, so a vacancy created by selection is taken by a fresh draw from
        # the same band -- a pool enriched for the favoured type. Deleting instead (soft
        # selection) shrinks the cohort, and the global lambda then raises EVERYONE's fertility
        # next generation, handing much of the advantage straight back. Measured effect: the
        # realised selection coefficient was about half the nominal one, which is why the band
        # model needed roughly twice the s of the deme model for the same outcome.
        target = np.bincount(kband, minlength=n_bands)
        parts = [{k: v[surv] for k, v in kid.items()}]
        for _ in range(cfg.refill_rounds):
            have = np.bincount(np.concatenate([p["band"] for p in parts]).astype(int),
                               minlength=n_bands)
            short = np.maximum(target - have, 0)
            if not short.any():
                break
            mb = []
            for b in np.where(short > 0)[0]:
                wb = women[P["band"][women] == b]
                if len(wb):
                    mb.append(rng.choice(wb, short[b], replace=True))
            if not mb:
                break
            m2 = np.concatenate(mb)
            k2 = P["band"][m2]
            o2 = np.argsort(k2, kind="stable")
            m2, k2 = m2[o2], k2[o2]
            f2 = np.empty(len(m2), int)
            for b in np.unique(k2):
                men_b = np.where(men_mask & (P["band"] == b))[0]
                wts = man_wt_all[men_b]
                sl = k2 == b
                f2[sl] = (rng.choice(men_b, sl.sum(), p=wts / wts.sum())
                          if len(men_b) and wts.sum() > 0 else -1)
            ok = f2 >= 0
            m2, f2, k2 = m2[ok], f2[ok], k2[ok]
            if not len(m2):
                break
            s2, mt2, Y2, au2, X2 = _pair_transmit(P, m2, f2, rng, cfg)
            sv2, Y2 = _select(s2, mt2, Y2, au2, X2)
            parts.append(dict(band=k2[sv2], sex=s2[sv2],
                              natal=b_ident[k2[sv2]].astype(np.int8),
                              f1=(P["natal"][m2] != P["natal"][f2])[sv2],
                              mt=mt2[sv2], Y=Y2[sv2], auto=au2[sv2], X=X2[sv2],
                              mat1=m2[sv2].astype(np.int64),
                              mat2=P["mat1"][m2][sv2].astype(np.int64),
                              fnat=P["band"][f2][sv2].astype(np.int64)))
        P2 = {k: np.concatenate([p[k] for p in parts]) for k in kid}
        P2["band"] = P2["band"].astype(P["band"].dtype)
    else:
        P2 = {k: v[surv] for k, v in kid.items()}
        P2["band"] = P2["band"].astype(P["band"].dtype)

    # ---- 4. fission and death
    sizes = np.bincount(P2["band"], minlength=n_bands)
    spacing = 0.5 / np.sqrt(max(alive_b.sum(), 1) /
                            max(np.ptp(pos[alive_b], 0).prod(), 1.0)) if alive_b.sum() > 2 else 60.0
    for b in np.where(sizes > 2 * cfg.K)[0]:
        mem = np.where(P2["band"] == b)[0]
        half = rng.permutation(mem)[: len(mem) // 2]
        dead = np.where(np.bincount(P2["band"], minlength=n_bands) == 0)[0]
        # reuse a dead slot if one exists, else append a new band
        if len(dead):
            nb = dead[0]
        else:
            nb = n_bands
            pos = np.vstack([pos, pos[b]])
            b_ident = np.append(b_ident, b_ident[b])
            n_bands += 1
        for _ in range(12):        # place the daughter at the characteristic spacing, on-mask
            cand = pos[b] + rng.normal(0, max(spacing, cfg.sigma_b), 2)
            if land.ok(cand[None, :])[0]:
                pos[nb] = cand
                break
        else:
            pos[nb] = pos[b]
        b_ident[nb] = b_ident[b]
        P2["band"][half] = nb
    return P2, pos, b_ident


def stats(P, b_ident, exclude=()):
    """exclude: autosomal loci to drop from the background ancestry average.

    ADDED 11 Sep 2026, and it was a real bug that it was missing. DR: "Are we measuring
    autosomal ancestry using the allele that's contributing to the DMI? I'm worried we are and
    we shouldn't be." We were. A DMI drives its own autosomal partner locus to ~100% modern, so
    including it in an eight-locus average adds a fixed 100/8 = 12.5 percentage points that has
    nothing to do with background ancestry. spatial_sim2.run() has always excluded these loci;
    bandsim.stats never did, and neither did robust_run's stage 2. Default is empty so existing
    callers are unchanged, but every caller running a DMI on an autosome must pass it.
    """
    male = P["sex"] == 1
    mod = b_ident[P["band"]] == 1          # members of modern-identity bands
    mm = mod & male
    return dict(
        n=len(P["sex"]),
        bands_mod=int(b_ident[np.unique(P["band"])].sum()),
        bands=len(np.unique(P["band"])),
        mt=float((P["mt"] == 1).mean()),
        # DR's sampling metric: the chance that 10 males drawn at random from the whole
        # landscape are ALL modern at both uniparental markers. q is the frequency of men
        # carrying modern mtDNA and a modern Y; the probability is q**10 per replicate,
        # averaged over replicates. Equals P(both fixed) whenever both markers are absorbed,
        # and is the honest number when they are not.
        mtY_male=float(((P["mt"][male] == 1) & (P["Y"][male] == 1)).mean())
        if male.any() else np.nan,
        Y=float((P["Y"][male] == 1).mean()) if male.any() else np.nan,
        auto=float(P["auto"][:, [c for c in range(AUTOS) if c not in exclude]].mean() / 2.0),
        # within modern-identity bands only (the tracked quantity for the expanding group)
        mt_mod=float((P["mt"][mod] == 1).mean()) if mod.any() else np.nan,
        Y_mod=float((P["Y"][mm] == 1).mean()) if mm.any() else np.nan,
        auto_mod=(float(P["auto"][mod][:, [c for c in range(AUTOS)
                                           if c not in exclude]].mean() / 2.0)
                  if mod.any() else np.nan),
        X=float(np.mean(np.concatenate([
            P["X"][~male].ravel() / 2.0, P["X"][male][:, :].ravel()]))) if len(P["sex"]) else np.nan,
    )


def run(cfg):
    rng = np.random.default_rng(cfg.seed)
    land = Landscape()
    P, pos, b_ident = init_state(cfg, land, rng)
    series, snaps = [], {}
    for g in range(cfg.gens + 1):
        if g % cfg.record_every == 0 or g == cfg.gens:
            s = stats(P, b_ident); s["gen"] = g
            series.append(s)
        if g in (0, cfg.gens // 3, cfg.gens):
            alive = np.unique(P["band"])
            snaps[g] = (pos[alive].copy(), b_ident[alive].copy(),
                        np.bincount(P["band"], minlength=len(b_ident))[alive])
        out = step(P, pos, b_ident, cfg, land, rng)
        if out[0] is None:
            break
        P, pos, b_ident = out
        # guarded reseed: only while archaic bands persist, and only when the modern
        # identity has no band at all (establishment failure, not ongoing propping)
        if cfg.reseed and not cfg.stable_boundary:
            live = np.unique(P["band"])
            has_mod = np.any(b_ident[live] == 1)
            has_arc = np.any(b_ident[live] == 0)
            if (not has_mod) and has_arc:
                k = max(2, int(rng.poisson(cfg.K)))
                dead = np.setdiff1d(np.arange(len(b_ident)), live)
                nb = dead[0] if len(dead) else len(b_ident)
                if nb == len(b_ident):
                    pos = np.vstack([pos, land.to_km(np.array([cfg.seed_lonlat]))[0][None]])
                    b_ident = np.append(b_ident, 1)
                else:
                    pos[nb] = land.to_km(np.array([cfg.seed_lonlat]))[0]
                    b_ident[nb] = 1
                sexn = rng.integers(0, 2, k).astype(np.int8)
                P = {key: np.concatenate([P[key], val]) for key, val in dict(
                    band=np.full(k, nb, P["band"].dtype), sex=sexn,
                    natal=np.ones(k, np.int8), f1=np.zeros(k, bool),
                    mt=np.ones(k, np.int8),
                    Y=np.where(sexn == 1, 1, -1).astype(np.int8),
                    auto=np.full((k, AUTOS), 2, np.int8),
                    X=(np.where(sexn == 1, 1, 2)[:, None]
                       * np.ones((1, XLOCI), np.int8)).astype(np.int8)).items()}
    return series, snaps, land, (P, pos, b_ident)


if __name__ == "__main__":
    ap = argparse.ArgumentParser()
    ap.add_argument("--test", action="store_true")
    a = ap.parse_args()
    if a.test:
        cfg = Config(N_target=2000, K=25, gens=240, record_every=20, seed=1)
        series, snaps, land, final = run(cfg)
        for s in series:
            print(f"  gen {s['gen']:>4}  N={s['n']:>5}  bands={s['bands']:>3} "
                  f"(modern {s['bands_mod']:>3})  mt={100*s['mt']:5.1f}%  Y={100*s['Y']:5.1f}%  "
                  f"auto={100*s['auto']:5.1f}%")
```
