## Supplementary Information section 3 code base for "Hypothesis: A modern human range expansion ∼300,000 years ago had a critical role in Neandertal origins": figure2_bands.py

```
#!/usr/bin/env python3
"""
figure2_bands.py -- Figure 2, band-model version.

  panel a  four maps of Europe from ONE matrilocal range expansion, at generations 1, 100,
           250 and 500. Each band is a small square carrying three bars -- the modern
           fraction at mtDNA (M), the Y, and the autosomes (A) -- with area proportional to
           the number of people in the band. There is deliberately NO separate identity mark:
           under matrilocality (beta = 0) women never leave their natal band, so the mtDNA bar
           being red or blue IS the band's ancestry status (DR, 4 Sep 2026).

  panel b  what it takes to fix the modern Y: P(modern mtDNA AND Y both fixed) against
           additive selection s on the modern Y (left) and against F1-male fertility theta
           (right), one curve per scenario. Read from a suite summary CSV, so the figure never
           contains a number that is not in a collected table.

    python figure2_bands.py [--data band_ratchet_summary.csv] [--seed 11] [--s 0.02]
                            [--panel a|b|both]
"""
import argparse
import sys
import csv
import os

import numpy as np
import matplotlib
matplotlib.use("Agg")
import matplotlib.pyplot as plt
from matplotlib.gridspec import GridSpec
# TWO TEXT CLASSES ACROSS BOTH PANELS OF FIGURE 2. DR: "make number labels and axis labels in
# bold same font and font size as each other, and not in bold same font and font size as each
# other." So every bold string in the figure is BOLD_PT and every plain one is PLAIN_PT, and
# figure2_merged.py imports the same two constants rather than keeping its own copies. The panel
# letters are deliberately outside both classes -- they are identifiers, not labels, and DR asked
# for them "significantly larger".
# Bumped one point on DR's request (was 13 / 11). LETTER_PT is left alone: it is already
# deliberately outsized and is not part of either class.
BOLD_PT, PLAIN_PT, LETTER_PT = 14, 12, 30

from matplotlib.patches import Rectangle

from bandsim import Config, Landscape, init_state, step, stats, KM_PER_DEG_LAT, COS_REF
from band_suite import N_TARGET
from site_stripes_figure import albers

HERE = os.path.dirname(os.path.abspath(__file__))
GENS = [0, 100, 300, 1000]   # the identity sweep completes at median gen 759 [IQR 642-886]
RED, BLUE = "#b2182b", "#2166ac"

# THE SQUARE PALETTE IS INDIRECT, so that --mono is a palette swap and nothing else.
# DR, 2 Oct 2026: "solid black is modern, gray is archaic, and the Europe map is either some
# other color (yellow?), or just an outline". The reason is panel b of the alternative figure:
# there red and blue have been reassigned to mean band and deme, so panel a must stop using
# the same two colours for modern and archaic or the reader carries the wrong key across the
# page. band_square() and the landmass fill read these three names at call time rather than
# RED/BLUE directly, so use_mono() flips the whole panel -- and the video, which imports
# band_square from here -- without touching any drawing code.
# PANEL A'S CONFIGURATION, IN ONE PLACE. figure2a_movie.py imports this dict rather than
# keeping its own copy: it used to carry hardcoded duplicates under a comment claiming they
# were read from here, and when panel a moved from s = 0.02 to s = 0.08 the film silently
# stayed at the old replicate. Importing the dict makes the drift impossible.
# MOVED TO s = 0.16 (10 Oct 2026), after selection on the Y was made conditional on band
# identity. Under the matching rule the band framework retains the modern Y in only 47% of
# production replicates at s = 0.08 (band_ratchet_summary.csv), so that rung is just under
# even odds; at s = 0.16 it is 73%. e* refit at 0.179 (bandprod_calibration.csv). Seed 20098
# is figure2_seedscan.py's pick at this cell: 5.06% autosomal against the ensemble's 5.05%,
# both markers fixed; 118 of 160 scanned seeds fix both, matching the ensemble rate.
PANEL_A = dict(seed=20098, s=0.16, s_mt=0.000651268, e=0.179, beta=0.0)

SQ_MOD, SQ_ARCH = RED, BLUE            # modern fill (from the bottom), archaic ground
LAND_C = "#e6e1d8"                     # the landmass wash; None draws a coastline instead
BAR_GAP = 0.0                          # fraction of each sub-bar's width left as a gap
MONO_MOD, MONO_ARCH, MONO_LAND = "#000000", "#bdbdbd", None


def use_mono():
    """Black squares on grey, and the landmass as an outline rather than a wash."""
    global SQ_MOD, SQ_ARCH, LAND_C, BAR_GAP
    SQ_MOD, SQ_ARCH, LAND_C = MONO_MOD, MONO_ARCH, MONO_LAND
    # THE BLACK LINE ALONG THE BOTTOM OF EVERY SQUARE. DR, 2 Oct 2026. It is not decoration
    # and not a border: it is the autosomal bar, which by generation 1000 is ~5% modern, so
    # its fill is one or two pixels high. In red/blue that sliver read as data; in black it
    # reads as a rule drawn under the square, and because the three sub-bars are contiguous
    # the slivers join into a single line across the whole width.
    #
    # The fix is a gap between the sub-bars, NOT a minimum bar height or a threshold below
    # which the fill is dropped -- either of those would hide the 5% autosomal signal, which
    # is the one number in panel a the martingale argument turns on. With a gap the sliver
    # is three short dashes under three separate bars, which reads as three small values.
    BAR_GAP = 0.18


def draw_land(ax, GX, GY, mask, zorder=0):
    """The landmass: a filled wash, or a coastline when LAND_C is None.

    Shared by the figure and the film so the two cannot drift apart -- the duplicated
    literal "#e6e1d8" in both files is what made this a two-place edit before.
    """
    if LAND_C is None:
        ax.contour(GX, GY, mask.astype(float), levels=[0.5],
                   colors=["0.55"], linewidths=0.7, zorder=zorder)
    else:
        ax.contourf(GX, GY, mask.astype(float), levels=[0.5, 1.5],
                    colors=[LAND_C], zorder=zorder)

# scenario -> (label, colour, marker)
CURVES = {
    "fexp":   ("range expansion, matrilineal",   "#b2182b", "o", "-"),
    "nexp":   ("range expansion, unbiased",     "#e08214", "^", "-"),
    "fpulse": ("migration pulse, all-female",   "#2166ac", "s", "--"),
    "npulse": ("migration pulse, unbiased",     "#7f7f7f", "D", "--"),
    "fsb":    ("migration pulse, all-female",   "#2166ac", "s", "--"),   # legacy names
    "nsb":    ("migration pulse, unbiased",     "#7f7f7f", "D", "--"),
}


# ------------------------------------------------------------------ panel a
def band_square(ax, x, y, fracs, w):
    """Three contiguous vertical bars: red from the bottom to the modern proportion, blue
    above. Two colours only, no internal lines, no identity bar."""
    bw = (w / 3) * (1.0 - BAR_GAP)          # drawn width of one sub-bar
    for k, f in enumerate(fracs):
        x0 = x - w / 2 + k * w / 3 + (w / 3 - bw) / 2
        ax.add_patch(Rectangle((x0, y - w / 2), bw, w, facecolor=SQ_ARCH,
                               edgecolor="none", zorder=6))
        ax.add_patch(Rectangle((x0, y - w / 2), bw, w * float(np.clip(f, 0, 1)),
                               facecolor=SQ_MOD, edgecolor="none", zorder=7))


def per_band(P, b_ident, n_slots):
    """alive band indices, sizes, and per-band modern fractions at mt / Y / autosomes"""
    bo = P["band"]
    male = P["sex"] == 1
    size = np.bincount(bo, minlength=n_slots).astype(float)
    mt = np.bincount(bo, weights=(P["mt"] == 1), minlength=n_slots)
    au = np.bincount(bo, weights=P["auto"].mean(1) / 2.0, minlength=n_slots)
    ym = np.bincount(bo[male], minlength=n_slots).astype(float)
    yv = np.bincount(bo[male], weights=(P["Y"][male] == 1), minlength=n_slots)
    alive = np.where(size > 0)[0]
    with np.errstate(invalid="ignore", divide="ignore"):
        return (alive, size[alive], mt[alive] / size[alive],
                np.where(ym[alive] > 0, yv[alive] / np.maximum(ym[alive], 1), 0.0),
                au[alive] / size[alive])


def run_panel_a(seed, s, e, beta, N, s_mt):
    # N is passed explicitly and defaults to the settled census, so panel a can never be drawn
    # at a different population size from panel b
    cfg = Config(gens=max(GENS), record_every=10 ** 9, beta=beta, s=s, e=e, r=2.0, seed=seed,
                 N_target=N, s_mt=s_mt)
    rng = np.random.default_rng(cfg.seed)
    land = Landscape()
    P, pos, b_ident = init_state(cfg, land, rng)
    snaps = {}
    for g in range(max(GENS) + 1):
        if g in GENS:
            alive, size, mtf, yf, auf = per_band(P, b_ident, len(b_ident))
            snaps[g] = (pos[alive].copy(), b_ident[alive].copy(), size,
                        # EXCLUDE THE FOUR RESERVED SELECTION LOCI, as figure2a_movie.py and
                        # spatial_sim2.run() do. Without it the readout averages loci
                        # that are not background ancestry, and the stills disagreed
                        # with the film they are supposed to be frames of (15% against
                        # 16% autosomes at generation 100).
                        stats(P, b_ident, exclude=tuple(range(32, 36))),
                        np.column_stack([mtf, yf, auf]))
        out = step(P, pos, b_ident, cfg, land, rng)
        if out[0] is None:
            print(f"  population lost at gen {g}")
            break
        P, pos, b_ident = out
    return snaps, land, cfg


def draw_panel_a(axes, snaps, land, cfg):
    # solid landmass: fill the habitable mask as one region rather than stippling its cells
    LON, LAT = np.meshgrid(land.lon, land.lat)
    GX, GY = albers(LON, LAT)
    ii, jj = np.where(land.mask)
    bx, by = albers(land.lon[jj][::12], land.lat[ii][::12])
    for ax, g in zip(axes, GENS):
        draw_land(ax, GX, GY, land.mask)
        if g not in snaps:
            ax.axis("off")
            continue
        p, idn, size, st, fr3 = snaps[g]
        lon = p[:, 0] / (KM_PER_DEG_LAT * COS_REF) + land.lon0
        lat = p[:, 1] / KM_PER_DEG_LAT + land.lat0
        X, Y = albers(lon, lat)
        WB = 0.0125
        # side = WB*sqrt(size/K), so AREA is proportional to band size, which is what the key
        # claims. The floor used to be 0.55*WB = 8 people, which clipped 14-17% of bands
        # (fission keeps making small ones) and drew them bigger than their share. 0.28*WB is
        # two people, so only singletons are enlarged, and the claim is true for ~99% of
        # squares. The ceiling is kept as a guard but never binds here: the largest band in
        # this replicate is 48 against a bound of 72.
        w = np.clip(WB * np.sqrt(size / float(cfg.K)), 0.28 * WB, 1.7 * WB)
        for i in range(len(X)):
            band_square(ax, X[i], Y[i], fr3[i], w[i])
        ax.set_title(f"generation {g}", fontsize=BOLD_PT, weight="bold", pad=6)
        # ANCESTRY BLOCK MOVED INSIDE THE MAP. DR: "change the ancestry labels to be in the white
        # space in eastern Europe, in four lines: header is %modern, then rows are like
        # mtDNA 100 / Y 3 / autos 21, with the justifying so the numbers always start in the same
        # place." It used to be one long line above the map, which forced pad=17 on the title to
        # clear it and wasted a strip of the figure on text that belongs over the sea.
        #
        # Two columns, both anchored in axes coordinates: labels right-aligned at LAB_X so they
        # end together, numbers LEFT-aligned at NUM_X so they all begin at the same place, which
        # is what DR asked for. The block sits over the eastern Mediterranean and Black Sea,
        # which is open water in every generation, so it never covers a band.
        # WHERE THE WHITE SPACE ACTUALLY IS, measured rather than eyeballed. Rasterising the
        # habitable cells onto a 20 x 20 grid of axes fractions shows one large contiguous empty
        # wedge, in the TOP RIGHT beyond the north-eastern edge of the range: every cell with
        # x > 0.80 and y > 0.62 is open. My first attempt put the block at (0.735, 0.375), which
        # the same map shows is solid land -- it landed across the Balkans and Ukraine and covered
        # several bands. A translucent backing panel is drawn underneath as well, so the block
        # stays readable even in a generation where a band diffuses into the corner.
        # NUMBERS LEFT, WORDS RIGHT. DR: "try putting the numbers on the left, words on the
        # right, again matching the justification for both." Numbers are RIGHT-aligned at NUM_X
        # so their digits line up regardless of width, and the words are LEFT-aligned at LAB_X so
        # they all begin together -- one edge each, both straight.
        # Y0 RAISED FROM 0.780 TO 0.855 -- the fix for the notch, see the patch below.
        NUM_X, LAB_X, Y0, DY = 0.835, 0.850, 0.855, 0.062
        # MOVED UP, NOT BOXED IN. DR: "the weird divet at the bottom left of each ancestry
        # proportion legend", then "I don't like the outlines... make it similar to how they
        # were. Maybe just move the box a little higher."
        #
        # The notch was the coastline showing through. At alpha 0.72 the map reads faintly
        # under the panel, which is fine over open sea but not over the north-eastern tip of
        # the landmass, which used to end INSIDE the panel's bottom-left corner: what came
        # through was one diagonal step of beige with nothing around it, and an isolated step
        # reads as damage to the panel rather than as land continuing underneath.
        #
        # Making the panel opaque and outlining it removed the notch but turned a light wash
        # into a hard-edged box sitting on the map, which is worse. Raising Y0 by 0.075 of the
        # axes lifts the whole block clear of the coastline instead, so the translucency and
        # the absent border both survive and there is no land under the panel to show through.
        # The block still sits in the empty north-eastern wedge -- every cell with x > 0.80 and
        # y > 0.62 is open -- and its top edge is at 0.95, clear of the panel title.
        ax.add_patch(Rectangle((0.775, Y0 - 2.5 * DY), 0.225, 4.0 * DY,
                               transform=ax.transAxes, facecolor="white", alpha=0.72,
                               edgecolor="none", zorder=4))
        # UNDERLINED RATHER THAN BOLD. DR: "'% modern' labels should not be in bold but
        # instead non-bold with an underline under them." Bold was doing two jobs at once --
        # marking the block heading and competing with the percentages below it, which are
        # the numbers the reader is actually here for. A rule under the words separates the
        # heading from its rows without adding weight.
        #
        # The rule is drawn rather than set as a font property because matplotlib has no
        # underline: it is a short horizontal line in axes coordinates, spanning the text,
        # placed just below the baseline. Its length is measured from the rendered text
        # rather than guessed, so it tracks the font size if that ever changes.
        hx, hy = NUM_X - 0.055, Y0 + DY
        t = ax.text(hx, hy, "% modern", transform=ax.transAxes,
                    ha="left", va="center", fontsize=BOLD_PT, zorder=5)
        fig = ax.get_figure()
        fig.canvas.draw()
        bb = t.get_window_extent(fig.canvas.get_renderer())
        w_ax = ax.transAxes.inverted().transform(
            [(bb.x0, bb.y0), (bb.x1, bb.y1)])
        # OVERSHOOT BY 3%. DR: "the underlining doesn't fully go under the 'n' at the end of
        # '% modern'." The text bbox is the LAYOUT box, which for most fonts ends a hair
        # inside the final glyph's right side bearing, so a rule drawn to exactly the bbox
        # width stops short of the last letter. 3% was not quite enough -- DR: "it should go
        # a little beyond the 'n'" -- so 7%, which puts the rule's end just past the glyph
        # rather than flush with it.
        ax.plot([hx, hx + 1.07 * (w_ax[1][0] - w_ax[0][0])], [hy - 0.030] * 2,
                transform=ax.transAxes, color="black", lw=0.9, zorder=5,
                solid_capstyle="butt")
        for i, (name, val) in enumerate((("mtDNA", st["mt_mod"]),
                                         ("Y", st["Y_mod"]),
                                         ("autos", st["auto_mod"]))):
            # Right-aligned WITH the % attached, so the percent signs form the straight edge
            # and the digits sit just left of it. DR asked for the sign on every number.
            ax.text(NUM_X, Y0 - i * DY, f"{100 * val:.0f}%", transform=ax.transAxes,
                    ha="right", va="center", fontsize=PLAIN_PT, zorder=5)
            ax.text(LAB_X, Y0 - i * DY, name, transform=ax.transAxes,
                    ha="left", va="center", fontsize=PLAIN_PT, zorder=5)
        ax.set_aspect("equal")
        # crop to the landscape so the map fills its cell instead of floating in whitespace
        mx, my = 0.02 * (bx.max() - bx.min()), 0.03 * (by.max() - by.min())
        ax.set_xlim(bx.min() - mx, bx.max() + mx)
        ax.set_ylim(by.min() - my, by.max() + my)
        ax.set_xticks([]); ax.set_yticks([])
        for sp in ax.spines.values():
            sp.set_visible(False)


def legend_square(ax, x, y, w=0.055):
    band_square(ax, x, y, (0.85, 0.5, 0.12), w)
    for k, lab in enumerate(("M", "Y", "A")):
        ax.text(x - w / 2 + (k + 0.5) * w / 3, y + w * 0.62, lab, ha="center", fontsize=7.5)


# ------------------------------------------------------------------ panel b
def load_summary(path):
    """Normalise the two column conventions (bandprod_collect vs the local suite writer)."""
    rows = []
    for r in csv.DictReader(open(path)):
        g = lambda *k: next((r[x] for x in k if x in r and r[x] not in ("", None)), None)
        rows.append(dict(
            scenario=r["scenario"], sweep=r["sweep"], value=float(r["value"]),
            n=int(float(g("n") or 0)),
            joint=float(g("p_joint", "joint")),
            se=float(g("se_joint", "se") or 0),
            aut=float(g("aut")),
            p_Y=float(g("p_Y") or "nan"), p_mt=float(g("p_mt") or "nan")))
    # 'joint' and 'aut' are fractions in the collector output, percents in the local writer
    if max(r["joint"] for r in rows) > 1.5:
        for r in rows:
            r["joint"] /= 100.0; r["se"] /= 100.0
    if max(r["aut"] for r in rows) > 1.5:
        for r in rows:
            r["aut"] /= 100.0
    return rows


def draw_panel_b(axes, rows, scenarios, errorbars=False):
    for ax, sweep, xlab, title in (
            (axes[0], "s", "additive selection $s$ on the modern Y ($\\theta$ = 1)", ""),
            (axes[1], "theta", "F1-male fertility $\\theta$ ($s$ = 0)", "")):
        for sc in scenarios:
            ss = sorted([r for r in rows if r["scenario"] == sc and r["sweep"] == sweep],
                        key=lambda r: r["value"])
            if sweep == "theta":
                # theta = 1 with s = 0 is the same cell as the s-sweep's zero point
                zero = [dict(r, value=1.0) for r in rows if r["scenario"] == sc
                        and r["sweep"] == "s" and r["value"] == 0]
                ss = sorted(zero + ss, key=lambda r: -r["value"])
            if not ss:
                continue
            lab, col, mk, ls = CURVES.get(sc, (sc, "black", "o", "-"))
            x = [r["value"] for r in ss]
            y = [r["joint"] for r in ss]
            ax.errorbar(x, y, yerr=[r["se"] for r in ss] if errorbars else None,
                        marker=mk, ms=4.5, color=col, lw=1.6, ls=ls,
                        capsize=2.5 if errorbars else 0, label=lab)
            # Hollow any point whose realised autosomal ancestry left the 4-6% window the
            # series is calibrated to. Two things push it out at the ends of the axes: strong
            # selection on the Y drags modern autosomal ancestry UP by hitchhiking (npulse
            # reaches 9.5% at s = 0.64), and F1 sterility pushes it DOWN (the theta arms of both
            # pulse scenarios sit at 2.4-3.1%). Those cells are not comparable to the calibrated
            # middle of the range, and saying so on the figure is better than a caption caveat.
            off = [(xi, yi) for xi, yi, r in zip(x, y, ss)
                   if not 0.04 <= r["aut"] <= 0.06]
            if off:
                ax.plot([p[0] for p in off], [p[1] for p in off], marker=mk, ms=4.5,
                        ls="none", mfc="white", mec=col, mew=1.3, zorder=8)
        ax.set_ylim(-0.04, 1.04)
        if sweep == "s":
            # symlog: s spans 0.002-0.64, two and a half decades, and the whole biological
            # question lives below 0.05. On a linear axis that region is four pixels wide and
            # the rest of the panel is a flat plateau. linthresh = 0.002 keeps the s = 0 point
            # on the same axis instead of dropping it or faking it as a small positive number.
            ax.set_xscale("symlog", linthresh=0.002, linscale=0.5)
            # thinned: labelling every grid value collides below 0.05
            ax.set_xticks([0, 0.002, 0.01, 0.05, 0.2])
            ax.set_xticklabels(["0", "0.002", "0.01", "0.05", "0.2"], fontsize=8.2)
            ax.set_xticks([0.005, 0.02, 0.04, 0.08, 0.16, 0.32, 0.64], minor=True)
            ax.set_xlim(-0.0006, 0.9)
        if sweep == "theta":
            # theta runs 1 -> 0.2 -> 0.05 -> 0, so nearly all of the response happens in the
            # last factor of four and a linear axis buries it against the right edge, exactly
            # as the linear s axis did. symlog rather than log because theta = 0 (complete F1
            # sterility) is a real and important point, not an asymptote to be dropped.
            ax.set_xscale("symlog", linthresh=0.05, linscale=0.6)
            ax.set_xticks([1.0, 0.5, 0.2, 0.1, 0.05, 0.0])
            ax.set_xticklabels(["1", "0.5", "0.2", "0.1", "0.05", "0"], fontsize=8.2)
            ax.set_xlim(1.45, -0.012)
        ax.set_xlabel(xlab, fontsize=9.5)
        ax.tick_params(labelsize=8.5)
        ax.grid(alpha=0.25, lw=0.6)
    axes[0].set_ylabel("P(10 random males all modern\nat both mtDNA and the Y)", fontsize=9.5)
    # upper left is the empty corner: every curve starts low and only the matrilocal one
    # climbs, and it does so on the right of the panel
    axes[0].legend(fontsize=8.2, loc="upper left", framealpha=0.95)
    axes[1].tick_params(labelleft=False)


# ------------------------------------------------------------------ main
def main():
    ap = argparse.ArgumentParser()
    ap.add_argument("--data", default="band_ratchet_summary.csv")
    # Seed 20063, chosen by figure2_seedscan.py. Among 240 replicates of the production
    # configuration, 107 (45%) fix both uniparental markers by generation 1000 -- which
    # reproduces the production ensemble's p_joint = 0.456 for this cell -- and 20063 is the
    # MEDIAN of those on generation-1000 autosomal ancestry, at 4.57% against the ensemble's
    # 4.65%. It is therefore representative twice over rather than cherry-picked.
    #
    # This replaces seed 20024, which was picked on autosomal ancestry at the END of the
    # production run and then displayed at generation 1000; those are different numbers, and
    # the panel showed 9% where the table says 4.65%.
    #
    # The autosomal distribution is WIDE and bimodal -- among fixers the quartiles are
    # 0.6 / 4.6 / 8.3% and 21 of 107 sit at exactly zero, because once the identity sweep
    # completes the origin stops being reseeded and the founding modern contribution keeps
    # diluting. No single replicate is representative of that spread; panel b carries the
    # ensemble, and panel a is an illustration of one history.
    # SEED CHOSEN ON WHAT THE PANEL ACTUALLY PRINTS. DR: "pick a replicate for the simulations
    # that ends up at 5% for the autosomes (and ideally also 25 for the Istanbul seed), just so
    # readers won't do double-takes." Scanning 346 seeds through run_panel_a itself -- not
    # through figure2_seedscan.csv, which disagrees with the figure (it lists 20063 at 4.57%
    # where the panel prints 2.10%, because it was collected under a different configuration) --
    # 151 fix both markers, and seed 20768 is the one that also lands on 4.91% autosomal with a
    # founding band of exactly 25, which is K. Previous default 20063: 2.10% and 28.
    #
    # MOVED FROM s = 0.02 TO s = 0.08 (DR, 9 Oct 2026). The panel was being drawn at a Y
    # selection coefficient of 2%, which is BELOW the band framework's even-odds threshold of
    # about 6.6% -- the production ensemble retains the modern Y in only 45.6% of replicates
    # at that cell (band_ratchet_summary.csv). So the illustrative replicate was showing an
    # outcome the text says is unlikely at that strength, which is the opposite of the point.
    # s = 0.08 is the next calibrated rung above the threshold (e* = 0.224, P(Y) = 85.0%), so
    # no new calibration or cluster run is needed. Seed 20030 is figure2_seedscan.py's pick at
    # that cell: 4.60% autosomal against the ensemble mean of 4.72%, both markers fixed.
    ap.add_argument("--seed", type=int, default=PANEL_A["seed"])
    ap.add_argument("--s", type=float, default=PANEL_A["s"])
    # The s_mt this replicate draws from the accepted ABC posterior. Panel a must run the same
    # configuration as panel b, and panel b integrates the posterior rather than conditioning
    # on its median -- fixing s_mt at the median gives P(joint) = 32% against 45.6%
    # integrated, because the right tail matters.
    ap.add_argument("--s-mt", type=float, default=PANEL_A["s_mt"])
    # e* for fexp at s = 0.08, from bandprod_calibration.csv. This is the value the production
    # ensemble ran at (band_ratchet_summary.csv records e_used = 0.224 for this cell). The
    # previous default of 0.307 was not the calibrated rate for ANY fexp cell -- it came from
    # a superseded local calibration -- so panel a was being drawn at a different exogamy rate
    # from the panel b curve directly beneath it.
    ap.add_argument("--e", type=float, default=PANEL_A["e"])
    ap.add_argument("--beta", type=float, default=PANEL_A["beta"])
    ap.add_argument("--N", type=int, default=N_TARGET)
    ap.add_argument("--panel", default="both", choices=("a", "b", "both"))
    # Declared rather than sniffed out of sys.argv: argparse rejects unknown flags, so the
    # sys.argv test alone made --only-a a usage error.
    ap.add_argument("--only-a", action="store_true",
                    help="with --panel a, drop the strip reserved for this script's own "
                         "panel b (used when composing main-text Figure 2)")
    # DR (5 Sep 2026): no error bars and no n in the legend -- the replicate count goes in
    # the caption instead. P(joint) does carry binomial sampling error (~1.8 points at
    # p = 0.15, n = 400; ~1.1 at n = 1000), so --errorbars is kept for checking whether a
    # wiggle is real before deciding it is a feature.
    ap.add_argument("--errorbars", action="store_true")
    # --mono BELONGS TO THE ALTERNATIVE FIGURE 2, not to the published one. Default off, so
    # every existing invocation draws exactly the red/blue panel it drew before.
    ap.add_argument("--mono", action="store_true",
                    help="black = modern, grey = archaic, pale yellow landmass")
    ap.add_argument("--out", default="figure2_bands.png")
    a = ap.parse_args()
    if a.mono:
        use_mono()

    data = a.data if os.path.exists(a.data) else os.path.join(HERE, a.data)
    if a.panel in ("b", "both") and not os.path.exists(data):
        alt = os.path.join(HERE, "band_o2_results.csv")
        print(f"  {a.data} not found -- falling back to {os.path.basename(alt)} "
              f"(NO-RATCHET run; swap in the ratchet summary when it lands)")
        data = alt
    rows = load_summary(data) if a.panel in ("b", "both") else []
    scen = [s for s in ("fexp", "nexp", "fpulse", "npulse", "fsb", "nsb")
            if any(r["scenario"] == s for r in rows)]
    if "fpulse" in scen:
        scen = [s for s in scen if s not in ("fsb", "nsb")]

    # --only-a DROPS THE RESERVED PANEL-B STRIP. DR: "what we've been calling Figure 2 is just
    # panel b, in figure include panel a (the four maps)". The composed Figure 2 takes its panel
    # b from figure2_merged.py -- the four-route panel -- not from this script's own panel b, so
    # when only the maps are wanted the 0.90-height row and the 0.16 key row must go, or the
    # stacked figure carries four inches of white space between the maps and the routes.
    only_a = a.panel == "a" and a.only_a
    if only_a:
        # ONE ROW OF FOUR, ACROSS THE FULL PAGE WIDTH. DR: "panel (a) should go across the whole
        # page". As a 2 x 2 block it was 9.26 in wide against panel b's 16.40, so the composite
        # had a narrow square of maps floating over a wide strip of curves -- the bad aspect
        # ratio he objected to. Laid out 1 x 4 at panel b's own width the two panels share an
        # edge, the composite comes out near 16.4 x 10.5 in, and each map is WIDER than it was
        # in the 2 x 2 version (3.9 in against 3.5), so nothing is lost by unstacking them.
        # FULL-WIDTH MARGINS, SET EXPLICITLY. DR: "Use the full left-right real-estate for panel
        # (a)". The GridSpec above inherited matplotlib's default left=0.125 / right=0.9, so a
        # fifth of the page was blank gutter before a single map was drawn -- which is why the
        # composite trimmed to 14.76 in against panel b's 16.40 and the outer maps stopped short
        # of the page edges. The height is cut to 3.9 in at the same time: the maps hold a fixed
        # geographic aspect, so with a taller figure they were height-centred in their axes and
        # the extra inch went into white space above and below rather than into the maps.
        # BACK TO 2 x 2, AND BIGGER. DR: "return Panel a to two rows, the maps are so beautiful
        # we should give the visual tracking of the simulation more space." The 1 x 4 row did use
        # the full width, but at four maps across each one was only about 3.9 in wide; two rows of
        # two at the same page width give roughly 8.0 in per map, so each is about four times the
        # area. The explicit full-width margins stay -- that was the fix for the blank gutter and
        # it applies to either arrangement.
        # HEIGHT, NOT WIDTH, WAS THE BINDING CONSTRAINT -- I had this backwards and said so.
        # The habitable range's projected bounding box has aspect w/h = 1.560, while the axes
        # cell at 9.6 in of figure height was 7.94 x 4.27 in, i.e. aspect 1.859. Matplotlib
        # holds the map's aspect with set_aspect("equal"), so the map fitted the HEIGHT and left
        # a band of sea down each side -- which is the indent DR spotted on Spain, and why
        # squeezing the left margin to 0.030 did nothing to move it. 7.94 / 1.560 = 5.09 in of
        # cell height is what fills the width, which needs 11.4 in of figure. The maps come out
        # 19% larger on a side, about 42% more area, and Iberia now reaches the left margin.
        fig = plt.figure(figsize=(16.4, 11.4))
        # key row 0.10 -> 0.15: the key text went from 8.8 to 13 pt and two lines of it no
        # longer fitted the strip, so it was overlapping the bottom row of maps.
        # left = 0.030 is where panel b's rotated y-axis label begins; right and wspace are
        # squeezed to give the maps the rest ("make the maps a bit bigger using more
        # horizontal space"). At this width each map is about 7.9 in and the maps are
        # WIDTH-limited, so this is the last of the horizontal slack there is to give.
        gs = GridSpec(3, 2, height_ratios=[1.0, 1.0, 0.12], hspace=0.055, wspace=0.0,
                      left=0.030, right=0.998, top=0.955, bottom=0.012, figure=fig)
        map_axes = [fig.add_subplot(gs[0, 0]), fig.add_subplot(gs[0, 1]),
                    fig.add_subplot(gs[1, 0]), fig.add_subplot(gs[1, 1])]
        KEY_ROW = 2
    else:
        fig = plt.figure(figsize=(10.2, 11.0))
        # row 2 is a dedicated strip for the key: placing it in figure coordinates next to
        # panel b made it collide with the axes and run off the right edge
        gs = GridSpec(4, 2, height_ratios=[1.0, 1.0, 0.16, 0.90], hspace=0.06, wspace=0.06,
                      figure=fig)
        map_axes = [fig.add_subplot(gs[0, 0]), fig.add_subplot(gs[0, 1]),
                    fig.add_subplot(gs[1, 0]), fig.add_subplot(gs[1, 1])]
        KEY_ROW = 2
    if a.panel in ("a", "both"):
        print(f"  panel a: one replicate, seed {a.seed}, beta={a.beta}, s={a.s}, "
          f"e={a.e}, N={a.N}, s_mt={a.s_mt:g}")
        snaps, land, cfg = run_panel_a(a.seed, a.s, a.e, a.beta, a.N, a.s_mt)
        draw_panel_a(map_axes, snaps, land, cfg)
        for g in GENS:
            if g in snaps:
                st = snaps[g][3]
                print(f"    gen {g:4d}: {st['bands_mod']:>3}/{st['bands']} modern bands, "
                      f"N={st['n']:>5}  mt {100*st['mt_mod']:5.1f}%  Y {100*st['Y_mod']:5.1f}%  "
                      f"aut {100*st['auto_mod']:5.2f}%  (within modern bands)")
    else:
        for ax in map_axes:
            ax.axis("off")

    seed_n = cfg.K if a.panel in ("a", "both") else 0
    if a.panel in ("a", "both") and 0 in snaps:
        _p, _idn, _size, _st, _fr = snaps[0]
        seed_n = int(_size[_idn == 1].sum())
        print(f"    modern seed band: {seed_n} individuals (K = {cfg.K})")

    bx = [] if only_a else [fig.add_subplot(gs[3, 0]), fig.add_subplot(gs[3, 1])]
    if a.panel in ("b", "both"):
        draw_panel_b(bx, rows, scen, errorbars=a.errorbars)
        print(f"  panel b: {os.path.basename(data)}, scenarios {', '.join(scen)}")
    else:
        for ax in bx:
            ax.axis("off")

    # panel letters and the panel-a key, placed in figure coordinates
    # PANEL LETTERS AT 30 pt. DR: "Make the a and b labels significantly larger, and in
    # general the legend fonts larger." This figure is 16.4 in wide and prints at about
    # 7, so everything on it is reduced by a factor of 2.3: the old 16 pt letter arrived
    # on the page at 7 pt and the 8.5 pt legends at under 4. At 30 and 13 they land at
    # about 13 and 5.7 pt, which is what the rest of the manuscript uses.
    fig.text(0.006, 0.992, "a", fontsize=LETTER_PT, weight="bold", va="top")
    # The "b" letter belongs to THIS script's panel b, which --only-a does not draw. Left
    # unconditional it printed a stray bold "b" floating beside the generation-300 map of the
    # composed Figure 2, where panel b is the four-route figure from figure2_merged.py and
    # carries its own letter.
    if not only_a:
        fig.text(0.012, 0.285, "b", fontsize=16, weight="bold")
    # Key, on its own full-width strip. The icon is a SQUARE: it gets an axes sized square
    # in DISPLAY inches (w_frac * fig_width == h_frac * fig_height), because drawing it inside
    # a wide flat axes stretched it into a rectangle.
    kax = fig.add_subplot(gs[KEY_ROW, :]); kax.axis("off")
    kpos = kax.get_position()
    FW, FH = fig.get_size_inches()
    # +30% IN AREA. DR: "increase the size of the caption legend for the square by 30% in
    # area, and also increase its associated legend font". Area scales as the square of the
    # side, so a 30% area increase is a factor of sqrt(1.3) = 1.14 on ih, not 1.3 -- setting
    # the side to 1.3x would have made it 69% bigger than asked.
    ih = 0.030 * 1.3 ** 0.5
    x0 = 0.145
    icon = fig.add_axes([x0, kpos.y0 + 0.004, ih * FH / FW, ih])
    icon.set_xlim(0, 1); icon.set_ylim(0, 1); icon.axis("off")
    band_square(icon, 0.5, 0.5, (0.85, 0.5, 0.12), 1.0)
    fig.text(x0 + ih * FH / FW + 0.014, kpos.y0 + 0.004 + ih / 2,
             "In each square, modern fraction given left-to-right for mtDNA-Y-autosomes. "
             "Area is band size",
             va="center", ha="left", fontsize=PLAIN_PT + 2)
    # NO TIGHT CROP IN --only-a. DR: "Spain should start at the same left position as the text
    # of the panel b figure." bbox_inches="tight" crops each panel to its own ink, so the two
    # panels' margins were destroyed at save time and no amount of composing could line them up.
    # Saved at the full 16.4 in width, panel a shares panel b's coordinate system and the
    # GridSpec left margin below is directly comparable to panel b's subplots_adjust(left=...).
    _bb = None if only_a else "tight"
    fig.savefig(os.path.join(HERE, a.out), dpi=160, bbox_inches=_bb)
    fig.savefig(os.path.join(HERE, os.path.splitext(a.out)[0] + ".pdf"), bbox_inches=_bb)
    print(f"  wrote {a.out}")


if __name__ == "__main__":
    main()
```
