## Supplementary Information section 3 code base for "Hypothesis: A modern human range expansion ∼300,000 years ago had a critical role in Neandertal origins": figure2_seedscan.py

```
#!/usr/bin/env python3
"""
figure2_seedscan.py -- choose the replicate that panel a of Figure 2 draws. (DR, 7 Sep 2026)

WHY THIS EXISTS
---------------
Panel a is one replicate, and which one it is was previously settled by a comment rather than by
a script. Two things went wrong as a result:

  * it ran at e = 0.307. That is not the calibrated exogamy rate for any fexp cell. For s = 0.02
    bandprod_calibration.csv gives e* = 0.192, and band_ratchet_summary.csv confirms the
    production ensemble ran at e_used = 0.192. Panel a was therefore not drawn from the same
    configuration as panel b, which is the one thing the figure's own docstring promises.

  * the seed was picked on autosomal ancestry AT THE END OF THE PRODUCTION RUN, but panel a
    stops at generation 1000 and prints the value there. Those are different numbers: at
    generation 1000 the wave has only just finished (end_gen_med = 590) and the founding modern
    contribution is still diluting. The published panel said 9%; the table says 4.65%.

So the selection criterion has to be evaluated at the generation the figure actually displays,
under the configuration the ensemble actually used. That is what this does.

CRITERION
---------
Among replicates that fix BOTH uniparental markers by generation 1000 (mtDNA and Y at 100%
within modern bands, which by then is the whole landscape), take the one whose generation-1000
autosomal ancestry is closest to the production ensemble mean of 4.65%. Ties are broken toward
the median band count so the map does not look unusually sparse or crowded.

s_mt is drawn per replicate from the same accepted ABC posterior the production run used. It
cannot be drawn the way band_reps.py draws it -- that uses hash((tag, seed0)), and Python
randomises string hashing per process, so those draws are not reproducible outside the job that
made them. Here the draw is a plain PCG64 stream seeded on the replicate seed, so the chosen
cell is exactly reproducible from the two numbers printed at the end.

    python figure2_seedscan.py [--n 600] [--procs 4]
"""
import argparse
import csv
import multiprocessing as mp
import os
import sys

import numpy as np

HERE = os.path.dirname(os.path.abspath(__file__))
sys.path.insert(0, HERE)

from bandsim import Config, Landscape, init_state, step, stats   # noqa: E402
from band_suite import N_TARGET                                   # noqa: E402

GEN = 1000                    # the generation panel a's last map shows
POSTERIOR = os.path.join(HERE, "mt_gap_posterior_NA1500.txt")


def calibrated_e(scenario="fexp", sweep="s", value=0.02):
    path = os.path.join(HERE, "bandprod_calibration.csv")
    for r in csv.DictReader(open(path)):
        if (r["scenario"], r["sweep"], float(r["value"])) == (scenario, sweep, value):
            return float(r["e_star"])
    raise SystemExit(f"no calibration for {scenario} {sweep}={value}")


def target_aut(scenario="fexp", sweep="s", value=0.02):
    """the production ensemble's autosomal ancestry for this cell, as a percent"""
    path = os.path.join(HERE, "band_ratchet_summary.csv")
    for r in csv.DictReader(open(path)):
        if (r["scenario"], r["sweep"], float(r["value"])) == (scenario, sweep, value):
            a = float(r["aut"])
            return 100 * a if a <= 1.5 else a
    raise SystemExit("no ensemble row")


def load_posterior():
    return np.array([float(l) for l in open(POSTERIOR) if not l.startswith("#") and l.strip()])


def one(args):
    seed, s, e, beta, N, s_mt = args
    cfg = Config(gens=GEN, record_every=10 ** 9, beta=beta, s=s, e=e, r=2.0,
                 seed=seed, N_target=N, s_mt=s_mt)
    rng = np.random.default_rng(cfg.seed)
    land = Landscape()
    P, pos, b_ident = init_state(cfg, land, rng)
    for g in range(GEN):
        out = step(P, pos, b_ident, cfg, land, rng)
        if out[0] is None:
            return dict(seed=seed, s_mt=s_mt, died=1)
        P, pos, b_ident = out
    st = stats(P, b_ident)
    return dict(seed=seed, s_mt=s_mt, died=0,
                mt=100 * st["mt_mod"], Y=100 * st["Y_mod"], aut=100 * st["auto_mod"],
                mt_all=100 * st["mt"], Y_all=100 * st["Y"], aut_all=100 * st["auto"],
                bands=st["bands"], bands_mod=st["bands_mod"], n=st["n"])


def main():
    ap = argparse.ArgumentParser()
    ap.add_argument("--n", type=int, default=600)
    ap.add_argument("--seed0", type=int, default=20000)
    ap.add_argument("--s", type=float, default=0.02)
    ap.add_argument("--beta", type=float, default=0.0)
    ap.add_argument("--N", type=int, default=N_TARGET)
    ap.add_argument("--procs", type=int, default=4)
    ap.add_argument("--out", default="figure2_seedscan.csv")
    a = ap.parse_args()

    e = calibrated_e(value=a.s)
    tgt = target_aut(value=a.s)
    post = load_posterior()
    print(f"  e* = {e} (bandprod_calibration.csv), ensemble autosomes = {tgt:.2f}% "
          f"(band_ratchet_summary.csv), N = {a.N}, s = {a.s}, beta = {a.beta}")

    tasks = []
    for i in range(a.n):
        seed = a.seed0 + i
        # one reproducible draw per seed, independent of process or tag
        s_mt = float(np.random.default_rng(seed).choice(post))
        tasks.append((seed, a.s, e, a.beta, a.N, s_mt))

    # Append, so the scan can be run in chunks. Seeds already present are skipped, which makes
    # the whole thing resumable -- necessary because each shell call here gets its own PID
    # namespace and a background job does not outlive the call that started it.
    path = os.path.join(HERE, a.out)
    have = set()
    if os.path.exists(path):
        have = {int(r["seed"]) for r in csv.DictReader(open(path))}
        tasks = [t for t in tasks if t[0] not in have]
        print(f"  {len(have)} seeds already in {a.out}; {len(tasks)} to run")

    if tasks:
        with mp.Pool(a.procs) as pool:
            rows = pool.map(one, tasks, chunksize=2)
        new = [r for r in rows if not r["died"]]
        cols = ["seed", "s_mt", "died", "mt", "Y", "aut", "mt_all", "Y_all", "aut_all",
                "bands", "bands_mod", "n"]
        fresh = not os.path.exists(path)
        with open(path, "a", newline="") as f:
            w = csv.DictWriter(f, fieldnames=cols)
            if fresh:
                w.writeheader()
            w.writerows(new)

    live = list(csv.DictReader(open(path)))
    for r in live:
        for k in ("s_mt", "mt", "Y", "aut", "mt_all", "Y_all", "aut_all"):
            r[k] = float(r[k])
        for k in ("seed", "bands", "bands_mod", "n"):
            r[k] = int(r[k])

    fixed = [r for r in live if r["mt"] >= 99.999 and r["Y"] >= 99.999]
    auts = np.array([r["aut"] for r in live])
    print(f"\n  {len(live)}/{a.n} survived; {len(fixed)} fix BOTH markers by generation {GEN}")
    print(f"  autosomes at generation {GEN}: mean {auts.mean():.2f}%  median "
          f"{np.median(auts):.2f}%  quartiles {np.percentile(auts,25):.2f} / "
          f"{np.percentile(auts,75):.2f}  range {auts.min():.2f}-{auts.max():.2f}")
    if not fixed:
        raise SystemExit("  no replicate fixes both markers -- widen the scan")

    fa = np.array([r["aut"] for r in fixed])
    print(f"  among the {len(fixed)} that fix both: mean {fa.mean():.2f}%  median "
          f"{np.median(fa):.2f}%  range {fa.min():.2f}-{fa.max():.2f}")

    med_bands = np.median([r["bands"] for r in fixed])
    fixed.sort(key=lambda r: (round(abs(r["aut"] - tgt), 3), abs(r["bands"] - med_bands)))
    print(f"\n  closest to the ensemble {tgt:.2f}%:")
    print(f"    {'seed':>6} {'aut%':>6} {'mt%':>6} {'Y%':>6} {'bands':>6} {'N':>6} {'s_mt':>12}")
    for r in fixed[:12]:
        print(f"    {r['seed']:>6} {r['aut']:>6.2f} {r['mt']:>6.1f} {r['Y']:>6.1f} "
              f"{r['bands']:>6} {r['n']:>6} {r['s_mt']:>12.9f}")
    b = fixed[0]
    print(f"\n  CHOSEN: --seed {b['seed']} --s-mt {b['s_mt']:.9f} --e {e} --s {a.s} --N {a.N}")
    print(f"          generation {GEN}: mtDNA {b['mt']:.0f}%  Y {b['Y']:.0f}%  "
          f"autosomes {b['aut']:.2f}%  ({b['bands']} bands, {b['n']} people)")


if __name__ == "__main__":
    main()
```
