## Supplementary Information section 3 code base for "Hypothesis: A modern human range expansion ∼300,000 years ago had a critical role in Neandertal origins": make_range_mask.py

```
#!/usr/bin/env python3
"""
make_range_mask.py -- APPROXIMATE habitable-range mask for the band model, drawn to the spirit
of Vidal-Cordasco et al. 2024 (Sci Rep) Figure 4a at a permissive suitability threshold (~0.2,
"green plus pink"), inside DR's chosen frame.

THIS IS A PLACEHOLDER GEOMETRY. The figure's underlying suitability rasters are deposited at
Zenodo (doi 10.5281/zenodo.10809139, Yaworsky, "The Neanderthal niche space of western Eurasia:
supplemental material"); when the band model is actually built, download that and threshold the
real raster instead of this hand-parameterised approximation. The coastline here is real
(GLOBE-derived land mask via global-land-mask); only the RANGE LIMITS within the land are
approximated.

Frame (DR): -11..32.5 E, 34..57 N (east margin trimmed so the space beyond the range's east
edge roughly matches the west margin) -- the map SHOWS Anatolia/Levant/Caucasus as context, but
the habitable range is EUROPE ONLY: everything east/south of the straits is excluded, and the
expansion is seeded at Istanbul on the European shore, so the entry across the Bosphorus is
the model's first event and no Levant/Anatolia preamble is simulated. Projection: Albers equal-area conic
(standard parallels 38/52 N) -- areas are exact, shapes far better at these latitudes than a
cylindrical equal-area.

Rules encoded, per DR's instructions:
  * permissive threshold: green AND pink of the suitability map both count as habitable
  * Great Britain reachable (kept, up to ~54 N); NO other island (Ireland, Balearics, Corsica,
    Sardinia, Sicily, Crete, Cyprus... all removed by connected-component filtering)
  * Africa excluded; ASIA excluded from habitability (Anatolia, Levant, Caucasus shown as
    pale context only), Thrace retained; the modern seed sits in Istanbul, European side
  * the northern limit is ONE smooth arc: Channel coast, northern Germany, then from the base
    of Denmark a single smooth curve southeast into the Black Sea WEST of Crimea; Belarus, most
    of Ukraine, the Pontic steppe AND Crimea fall northeast of it

Writes:  range_mask.npz  (lon, lat, mask -- 0.05 degree grid, for the simulator)
         range_mask_map.png  (the visual check)
    python make_range_mask.py
"""
import numpy as np
import matplotlib
matplotlib.use("Agg")
import matplotlib.pyplot as plt
from global_land_mask import globe
from scipy import ndimage
from scipy.interpolate import PchipInterpolator

LON0, LON1, LAT0, LAT1 = -11.0, 32.5, 34.0, 57.0
STEP = 0.05

lon = np.arange(LON0, LON1 + STEP, STEP)
lat = np.arange(LAT0, LAT1 + STEP, STEP)
LON, LAT = np.meshgrid(lon, lat)

land = globe.is_land(LAT, LON)

# ---- northern limit of the range: ONE smooth curve (PCHIP, C1-continuous, no kinks).
# Channel coast and northern Germany in the west; from the base of Denmark (~9.5 E, 54.6 N) it
# bends as a single smooth arc southeast and dives into the Black Sea WEST of Crimea: Crimea is excluded (DR -- though note
# Kiik-Koba and the Zaskalnaya cluster are conventionally attributed as Neandertal; revisit if
# desired). Limit stays below 44.3 across Crimean longitudes so no part of the peninsula,
# Sevastopol to Kerch, slips under.
_np_lon = [-11, -5,  0,   5,   9.5, 16.0, 22.0, 27.0, 30.5, 33.0, 34.0]
_np_lat = [ 47, 49, 51.5, 53.5, 54.6, 53.4, 51.5, 49.0, 46.6, 44.3, 44.0]
_arc = PchipInterpolator(_np_lon, _np_lat)
nlim = _arc(np.clip(LON, -11, 34))

habit = land & (LAT <= nlim)

# ---- straight-chord northeast cut (DR, 3 Sep 2026): "too much energy is spent on northeastern
# Europe". One straight line from the base of Denmark (9.5 E, 54.6 N) to the Bulgarian Black
# Sea coast just north of the seed (28.5 E, 42.0 N); everything northeast of it is out. This is
# strictly stronger than the arc east of Denmark: it removes most of Poland, eastern Slovakia,
# eastern Romania etc., focusing the arena on the fossil-bearing west and south.
# convex version (DR): the cut bows OUTWARD (northeast) rather than following the straight
# chord, via a parabolic bulge (+2.2 deg at midspan, zero at both endpoints).
_t = np.clip((LON - 9.5) / (28.5 - 9.5), 0, 1)
_chord = 54.6 + (42.0 - 54.6) * _t + 2.2 * 4 * _t * (1 - _t)
habit &= ~((LON >= 9.5) & (LAT > _chord))

# ---- Great Britain: reachable, up to ~54 N (no northern Britain per the frame)
gb = land & (LON >= -6.5) & (LON <= 2.0) & (LAT >= 49.8) & (LAT <= 54.0)
habit |= gb

# ---- Africa out. The Mediterranean's south shore: exclude land below a line that runs under
# Gibraltar across to Sinai, but keep the Levant strip.
africa = land & (LAT < 35.6) & (LON < 11.0)                       # Maghreb
africa |= land & (LAT < 34.0) & (LON >= 11.0) & (LON < 33.0)      # Libya/Egypt coast
habit &= ~africa

# ---- ASIA excluded from the habitable range (DR): the map still SHOWS Anatolia, the Levant
# and the Caucasus as pale land for context, but nothing east/south of the straits is
# habitable. The dividing rule: south of 42 N and east of 26 E is Asian, EXCEPT Thrace -- the
# European shore of the Marmara, north of a line from (26 E, 40.3 N) to the Bosphorus
# (29.05 E, ~41.0 N). The seed sits in Istanbul on the European side, so the expansion begins
# at the straits with all of Asia off-limits behind it.
thrace_line = 40.3 + 0.2295 * (LON - 26.0)
asia = land & (LON >= 26.0) & (LAT <= 42.0) & ((LAT < thrace_line) | (LON > 29.05))
habit &= ~asia

# ---- islands: drop every connected component except the continental mass and Great Britain.
# The Bosporus/Dardanelles pinch can disconnect Anatolia at this resolution, so bridge the
# straits before labelling (a 2-cell dilation across water at the two crossings).
bridge = ((LON >= 25.9) & (LON <= 26.6) & (LAT >= 40.0) & (LAT <= 40.4)) | \
         ((LON >= 28.9) & (LON <= 29.3) & (LAT >= 40.9) & (LAT <= 41.3)) | \
         ((LON >= 0.8) & (LON <= 1.8) & (LAT >= 50.7) & (LAT <= 51.3))    # Dover: GB counted
lab, n = ndimage.label(habit | bridge)


def comp_at(latv, lonv):
    j = int(round((lonv - LON0) / STEP)); i = int(round((latv - LAT0) / STEP))
    return lab[i, j]


keep = {comp_at(48, 5), comp_at(52.5, -1.5)}   # France (continent incl. Thrace), GB
keep.discard(0)
habit = np.isin(lab, list(keep)) & habit

# ---- plot: Albers equal-area conic, std parallels 38/52 N, centre 17 E. Land pale, range
# dark, dotted range boundary, no borders; removed islands still drawn as pale land.
P1, P2, LON_C, LAT_C = 38.0, 52.0, 17.0, 43.0
_n = (np.sin(np.deg2rad(P1)) + np.sin(np.deg2rad(P2))) / 2
_C = np.cos(np.deg2rad(P1)) ** 2 + 2 * _n * np.sin(np.deg2rad(P1))
_r0 = np.sqrt(_C - 2 * _n * np.sin(np.deg2rad(LAT_C))) / _n


def albers(lonv, latv):
    rho = np.sqrt(_C - 2 * _n * np.sin(np.deg2rad(latv))) / _n
    th = _n * np.deg2rad(lonv - LON_C)
    return rho * np.sin(th), _r0 - rho * np.cos(th)


X, Y = albers(LON, LAT)

fig, ax = plt.subplots(figsize=(12.5, 7.6))
ax.contourf(X, Y, land.astype(float), levels=[0.5, 1.5], colors=["#e8e4dd"])
ax.contourf(X, Y, habit.astype(float), levels=[0.5, 1.5], colors=["#8a8a8a"])
ax.contour(X, Y, habit.astype(float), levels=[0.5], colors="black",
           linewidths=1.3, linestyles="dotted")
sx, sy = albers(28.95, 41.05)     # Istanbul, European shore of the Bosphorus
ax.plot([sx], [sy], marker="s", ms=10, color="#b2182b", mec="black", zorder=5)
ax.annotate("modern seed", xy=(sx, sy), xytext=(12, 8),
            textcoords="offset points", fontsize=11)
ax.set_aspect("equal")
ax.set_xticks([]); ax.set_yticks([])
ax.set_title("Band-model landscape (APPROXIMATION of Vidal-Cordasco et al. 2024 Fig. 4a,\n"
             "permissive threshold; replace with the Zenodo raster 10.5281/zenodo.10809139)",
             fontsize=11.5)
for sp in ax.spines.values():
    sp.set_visible(False)
fig.tight_layout()
fig.savefig("band_model/range_mask_map.png", dpi=170, bbox_inches="tight")
np.savez_compressed("band_model/range_mask.npz", lon=lon, lat=lat, mask=habit)
frac = habit.sum() / land.sum()
print(f"wrote band_model/range_mask_map.png and range_mask.npz "
      f"({habit.sum():,} cells habitable = {100*frac:.0f}% of land in frame)")
```
