## Supplementary Information section 3 code base for "Hypothesis: A modern human range expansion ∼300,000 years ago had a critical role in Neandertal origins": site_stripes_figure.py

```
#!/usr/bin/env python3
"""
site_stripes_figure.py -- the band model's headline visualisation: square symbols at Neandertal
site locations, each carrying THREE VERTICAL STRIPES showing the local proportion of MODERN
ancestry at (left to right) mtDNA, Y, and autosomes, at the sampling generation.

Local ancestry at a site is the Gaussian-kernel-weighted average over bands (kernel scale =
the exogamy scale sigma_e, i.e. "the neighbourhood a person there would actually mate within").
Each stripe fills bottom-up in RED to the modern proportion (remainder BLUE = archaic), and the
square's heavy border encodes the local band identity in the same colors -- red = modern bands,
blue = archaic (kernel-weighted majority). A red middle stripe on a mostly-blue square is the
paper's signature pattern: modern mtDNA on an archaic genomic background.

Site coordinates are APPROXIMATE (plotting-grade, not gazetteer-grade -- verify before
publication). Demo defaults: male-biased exogamy (beta = 0.1, matrilocal bands) with modest
selection on the Y (s = 0.02).

    python site_stripes_figure.py [--gens 6993] [--beta 0.1] [--s 0.02]
"""
import argparse
import os

import numpy as np
import matplotlib
matplotlib.use("Agg")
import matplotlib.pyplot as plt
from matplotlib.patches import Rectangle

from bandsim import Config, run, KM_PER_DEG_LAT, COS_REF

HERE = os.path.dirname(os.path.abspath(__file__))

# classic western-European Neandertal (mtDNA-bearing) sites; coordinates approximate.
# Squares sit EXACTLY at the site location (no offsets, no leader lines); only the text label
# side varies for crowded clusters: b=below, t=above, l=left, r=right.
SITES = [
    ("El Sidron",        -5.33, 43.38, "b"),
    ("La Chapelle",       1.72, 44.98, "b"),
    ("Rochers-de-Villeneuve", 0.83, 46.40, "b"),
    ("Scladina",          5.02, 50.48, "l"),
    ("Engis",             5.40, 50.58, "t"),
    ("Feldhofer",         6.95, 51.23, "r"),
    ("Monte Lessini",    11.00, 45.60, "b"),
    ("Vindija",          16.07, 46.30, "b"),
]

# Albers (same parameters as the landscape map)
P1v, P2v, LON_C, LAT_C = 38.0, 52.0, 17.0, 43.0
_n = (np.sin(np.deg2rad(P1v)) + np.sin(np.deg2rad(P2v))) / 2
_C = np.cos(np.deg2rad(P1v)) ** 2 + 2 * _n * np.sin(np.deg2rad(P1v))
_r0 = np.sqrt(_C - 2 * _n * np.sin(np.deg2rad(LAT_C))) / _n


def albers(lon, lat):
    rho = np.sqrt(_C - 2 * _n * np.sin(np.deg2rad(lat))) / _n
    th = _n * np.deg2rad(lon - LON_C)
    return rho * np.sin(th), _r0 - rho * np.cos(th)


def local_ancestry(P, pos, b_ident, sites_km, sigma):
    """Kernel-weighted modern proportions (auto, mt, Y), head-count, and modern-identity
    weight share at each site."""
    band_of = P["band"]
    n_b = pos.shape[0]
    male = P["sex"] == 1
    size = np.bincount(band_of, minlength=n_b).astype(float)
    a_sum = np.zeros(n_b); np.add.at(a_sum, band_of, P["auto"].mean(1) / 2.0)
    m_sum = np.zeros(n_b); np.add.at(m_sum, band_of, (P["mt"] == 1).astype(float))
    y_cnt = np.zeros(n_b); np.add.at(y_cnt, band_of[male], 1.0)
    y_sum = np.zeros(n_b); np.add.at(y_sum, band_of[male], (P["Y"][male] == 1).astype(float))
    out = []
    for sx, sy in sites_km:
        w = np.exp(-((pos[:, 0] - sx) ** 2 + (pos[:, 1] - sy) ** 2) / (2 * sigma ** 2))
        w = w * (size > 0)
        wa = (w * a_sum).sum() / max((w * size).sum(), 1e-9)
        wm = (w * m_sum).sum() / max((w * size).sum(), 1e-9)
        wy = (w * y_sum).sum() / max((w * y_cnt).sum(), 1e-9)
        wid = (w * size * (b_ident == 1)).sum() / max((w * size).sum(), 1e-9)
        out.append((wa, wm, wy, float((w * size).sum()), wid))
    return out


def draw_stripes(ax, x, y, fracs, w, h, border):
    """Square of width w at (x, y): three vertical stripes, red fill bottom-up = modern
    proportion, blue = archaic; heavy border in the same colors gives local band identity."""
    for k, f in enumerate(fracs):
        x0 = x - w / 2 + k * w / 3
        ax.add_patch(Rectangle((x0, y - h / 2), w / 3, h, facecolor="#6a9fd8",
                               edgecolor="none", zorder=6))
        ax.add_patch(Rectangle((x0, y - h / 2), w / 3, h * float(np.clip(f, 0, 1)),
                               facecolor="#d6604d", edgecolor="none", zorder=7))
    for k in (1, 2):
        ax.plot([x - w / 2 + k * w / 3] * 2, [y - h / 2, y + h / 2],
                color="white", lw=0.5, zorder=8)
    ax.add_patch(Rectangle((x - w / 2, y - h / 2), w, h, facecolor="none",
                           edgecolor=border, linewidth=1.3, zorder=9))


def main():
    ap = argparse.ArgumentParser()
    ap.add_argument("--gens", type=int, default=6993)
    ap.add_argument("--beta", type=float, default=0.1)
    ap.add_argument("--s", type=float, default=0.02)
    ap.add_argument("--seed", type=int, default=11)
    a = ap.parse_args()

    cfg = Config(N_target=2000, K=25, gens=a.gens, record_every=max(a.gens // 20, 1),
                 beta=a.beta, s=a.s, r=2.0, seed=a.seed)
    series, snaps, land, (P, pos, b_ident) = run(cfg)
    last = series[-1]
    print(f"  final: N={last['n']}  modern bands {last['bands_mod']}/{last['bands']}  "
          f"mt={100*last['mt']:.0f}%  Y={100*last['Y']:.0f}%  auto={100*last['auto']:.1f}%")

    sites_lonlat = np.array([(lo, la) for _, lo, la, *_ in SITES])
    sites_km = land.to_km(sites_lonlat)
    fr = local_ancestry(P, pos, b_ident, sites_km, cfg.sigma_e)

    fig, ax = plt.subplots(figsize=(12.5, 8.2))
    ii, jj = np.where(land.mask)
    bx, by = albers(land.lon[jj][::6], land.lat[ii][::6])
    ax.scatter(bx, by, s=0.4, color="#dcd7cf", rasterized=True)
    # square AREA proportional to the local head-count at the site (kernel-weighted number of
    # individuals living around it in the sampling generation), DR request. The median site
    # gets the reference width; area scales linearly with count, so width goes as sqrt.
    W = 0.011
    counts = np.array([f[3] for f in fr])
    widths = np.clip(W * np.sqrt(counts / max(np.median(counts), 1e-9)), 0.55 * W, 1.7 * W)
    for (name, lo, la, side), f, wsz in zip(SITES, fr, widths):
        x, y = albers(lo, la)
        border = "#b2182b" if f[4] >= 0.5 else "#2166ac"
        draw_stripes(ax, x, y, (f[1], f[2], f[0]), wsz, wsz, border)
    # legend square
    lx, ly = albers(-8.5, 55.0)
    WL = 2.6 * W          # legend square drawn larger than map scale, for readability
    draw_stripes(ax, lx, ly, (0.7, 0.5, 0.3), WL, WL, "#b2182b")
    for k, lab in enumerate(("M", "Y", "A")):
        ax.annotate(lab, xy=(lx - WL / 2 + (k + 0.5) * WL / 3, ly + WL / 2),
                    xytext=(0, 3), textcoords="offset points",
                    ha="center", va="bottom", fontsize=9, zorder=11)
    ax.annotate("red = modern, blue = archaic\n"
                "fill = ancestry, border = band identity\n"
                "area = local population", xy=(lx, ly - WL / 2),
                xytext=(0, -6), textcoords="offset points", ha="center", va="top", fontsize=9)
    ax.set_aspect("equal"); ax.set_xticks([]); ax.set_yticks([])
    for sp in ax.spines.values():
        sp.set_visible(False)
    ax.set_title(f"Band model, generation {a.gens}:  local modern ancestry at classic sites\n"
                 f"(beta={a.beta}, s={a.s} on the Y, r={cfg.r}, e={cfg.e} — demo run, "
                 f"single replicate)", fontsize=12)
    fig.tight_layout()
    out = os.path.join(HERE, "site_stripes_demo.png")
    fig.savefig(out, dpi=150, bbox_inches="tight")
    print("wrote", out)
    for (name, *_), f in zip(SITES, fr):
        print(f"    {name:<24} auto={100*f[0]:5.1f}%  mt={100*f[1]:5.1f}%  Y={100*f[2]:5.1f}%")


if __name__ == "__main__":
    main()
```
