## Supplementary Information section 3 code base for "Hypothesis: A modern human range expansion ∼300,000 years ago had a critical role in Neandertal origins": build_supp_workbook.py

```
#!/usr/bin/env python3
"""
build_supp_workbook.py -- the supplementary data workbook for SI section 4. (DR, 11 Sep 2026)

DR: "would you like to start building the Excel supplementary table with 10 tabs so I can see
what it looks like and give feedback?"

THE ORGANISING PRINCIPLE is that every tab is the source for one named figure or table in the
note, so a referee can go from any statement to the numbers behind it in one hop, and the
README tab is the index that makes that hop possible. There was no specification for this --
nothing in the project said what the tabs should be -- so the first version was a proposal;
the layout below is what survived being reacted to.

NOTHING HERE IS PROVISIONAL, as of 21 Sep. Every tab that carries simulation output carries
36-locus output at the anchored geometry: the deme sensitivity scan was rerun on 13 Sep, the
band scan completed its density-regulation column on 14 Sep, stage-2 scoring finished on
14 Sep, and the cultural mechanisms were moved off the pre-Cartesian grid scripts onto the
same two-stage pipeline as everything else. The status column says which of the two kinds of
tab each one is, and no longer warns about a rerun that has landed.

  1  README                  index, provenance and status of every tab
  2  Baseline parameters     both frameworks, every value, with why it has that value
  3  Effective-size screen   the 508 measured (grid, K, m) combinations behind the baseline
  4  Landscape definition    the rules and coordinates that generate the band-model arena
  5  Landscape outline       the traced boundary of that arena, as lon/lat
  6  Sensitivity (deme)      Figure S3.5
  7  Sensitivity (band)      Figure S3.6
  8  mtDNA fixation curves   Figure S3.4
  9  Cultural mechanisms     the five cultural panels of Figure S3.2
 10  DMI and drive search    every scored combination behind Figures 3b and S3.2
 11  X-drive and resistance  Figure S3.3, both panels
 12  mtDNA under expansion   Figure S3.4, the patrilineal range-expansion arm

TAB 4 IS THE ONE THE NOTE ALREADY DEMANDS. Its paragraph on the European landscape ends "the
exact coordinates of the landscape I use are specified in Supplementary Table xx", and no such
table existed in any form -- the arena lives as code in make_range_mask.py and as a boolean
array in range_mask.npz. Tabs 4 and 5 are that table: the generating rules with their
coordinates, and the resulting outline.

STATUS IS CARRIED ON EVERY TAB, not just in the README, so that a tab read on its own says
what produced it. It distinguishes simulation output from configuration and analytic tables,
which is the distinction that decides whether a replicate count means anything.

    python build_supp_workbook.py
"""
import csv
import json as _json
import os
import sys

import numpy as np
from openpyxl import Workbook
from openpyxl.comments import Comment
from openpyxl.styles import Alignment, Border, Font, PatternFill, Side
from openpyxl.utils import get_column_letter

HERE = os.path.dirname(os.path.abspath(__file__))
BAND = os.path.join(HERE, "band_model")
CL = os.path.join(HERE, "cluster")
# RENAMED FROM "Supplementary Table 3" ON DR'S INSTRUCTION, 13 Sep. The note is Supplementary
# Information section 4, but this workbook is the paper's Supplementary Table 1.
OUT = os.path.join(HERE, "Supplementary Table 1 - mechanisms for Y and mtDNA introgression.xlsx")

FONT = "Arial"
H1 = Font(name=FONT, size=13, bold=True)
H2 = Font(name=FONT, size=11, bold=True)
BODY = Font(name=FONT, size=10)
SMALL = Font(name=FONT, size=9, italic=True, color="595959")
HDRFILL = PatternFill("solid", fgColor="D9E2F3")
WARNFILL = PatternFill("solid", fgColor="FFF2CC")
THIN = Side(style="thin", color="BFBFBF")
BOX = Border(left=THIN, right=THIN, top=THIN, bottom=THIN)

# Provenance status, used on every tab and collected in the README.
# THE STATUS COLUMN SAYS WHAT A TAB IS, NOT WHAT IS PENDING -- CHANGED 21 Sep. It used to
# carry "8-locus, will be replaced by the 36-locus rerun" on three tabs and "awaiting stage-2
# scoring" on a fourth, which made the whole workbook read as a draft. Every one of those
# reruns has landed: the deme sensitivity scan was rerun on 13 Sep at the anchored geometry,
# the band scan finished with its density-regulation column on 14 Sep, the cultural
# mechanisms now come from the scored Cartesian search rather than the old grid scripts, and
# stage-2 scoring completed on 14 Sep. Nothing in this workbook is provisional, so the column
# now distinguishes the two kinds of tab there actually are.
SIM = "simulation output, 36-locus engine (ancestry measured on the 32 neutral loci)"
CONFIG = "configuration or analytic — no simulation replicates"


def read_csv(path):
    with open(path, newline="") as f:
        r = csv.reader(f)
        rows = [row for row in r if row and not row[0].startswith("#")]
    return rows[0], rows[1:]


def num(x):
    try:
        v = float(x)
        return int(v) if v == int(v) and abs(v) < 1e15 else v
    except (TypeError, ValueError):
        return x


# SUMMARY CACHE. The three Cartesian tabs read ~85,000 JSON files; on the mounted working
# tree that is longer than one shell call is allowed, so cluster/summary_cache.py reads them
# once in parallel and this consults the pickle, falling back to the file for anything
# missing. Build or refresh it with `python cluster/summary_cache.py` before this script.
_SUMM = None


def _summary(path):
    global _SUMM
    if _SUMM is None:
        try:
            sys.path.insert(0, CL)
            import summary_cache
            _SUMM = summary_cache.load()
        except Exception:
            _SUMM = {}
    s = _SUMM.get(os.path.abspath(path))
    if s is None:
        s = _json.load(open(path))["summary"]
    return s


def write_table(ws, header, rows, start=1, status=None, note=None):
    r = start
    if note:
        ws.cell(r, 1, note).font = SMALL
        r += 2
    if status:
        c = ws.cell(r, 1, f"Status: {status}")
        c.font = Font(name=FONT, size=10, bold=True)
        if status not in (SIM, CONFIG):
            c.fill = WARNFILL
        r += 2
    for j, h in enumerate(header, start=1):
        c = ws.cell(r, j, h)
        c.font = H2
        c.fill = HDRFILL
        c.border = BOX
        c.alignment = Alignment(wrap_text=True, vertical="bottom")
    hdr_row = r
    r += 1
    for row in rows:
        for j, v in enumerate(row, start=1):
            c = ws.cell(r, j, num(v))
            c.font = BODY
            c.border = BOX
        r += 1
    ws.freeze_panes = ws.cell(hdr_row + 1, 1)
    # A column can be wider than every row is long -- the effective-size tab appends a header
    # for a column whose values are formulas written afterwards -- so index defensively.
    for j, h in enumerate(header, start=1):
        widths = [len(str(row[j - 1])) + 2 for row in rows if len(row) >= j]
        width = max([len(str(h)) + 2] + widths)
        ws.column_dimensions[get_column_letter(j)].width = min(max(width, 9), 46)
    return hdr_row, r


# ---------------------------------------------------------------------------------------------
def tab_readme(wb, index_rows):
    ws = wb.create_sheet("README", 0)
    ws["A1"] = "Supplementary Table 1"
    ws["A1"].font = H1
    ws["A2"] = ("Simulation data underlying Supplementary Information section 4, "
                "“Plausible mechanisms for Neandertals inheriting modern ancestry at both "
                "mtDNA and Y”.")
    ws["A2"].font = BODY
    ws["A4"] = ("Each tab below is the source for one named figure or table in the note, so "
                "any statement there can be traced to the numbers behind it in one step. All "
                "simulation tabs come from the 36-locus engine, in which ancestry is measured "
                "on the 32 loci that no mechanism touches, at the anchored geometry described "
                "in Appendices A and B. Where a tab carries an acceptance test, it is "
                "recomputed in the sheet from the note's three criteria rather than baked in, "
                "so a reader can move the window and watch every claim move with it.")
    ws["A4"].font = BODY
    ws["A4"].alignment = Alignment(wrap_text=True, vertical="top")
    ws.merge_cells("A4:E6")

    hdr = ["Tab", "Contents", "Supports", "Rows", "Source"]
    hdr_row, end = write_table(ws, hdr, index_rows, start=8)
    ws.column_dimensions["A"].width = 26
    ws.column_dimensions["B"].width = 54
    ws.column_dimensions["C"].width = 26
    ws.column_dimensions["D"].width = 9
    ws.column_dimensions["E"].width = 46
    for r in range(hdr_row + 1, end):
        ws.cell(r, 2).alignment = Alignment(wrap_text=True, vertical="top")
        ws.cell(r, 5).alignment = Alignment(wrap_text=True, vertical="top")

    ws.cell(end + 1, 1, "Engine").font = H2
    for k, (lab, val) in enumerate([
        ("autosomal loci tracked", 36),
        ("of which measured for ancestry (indices 1-32)", 32),
        ("of which reserved for mechanisms (indices 33-36)", 4),
        ("X loci tracked (drive, mechanism candidate, neutral)", 3),
        ("sampling generation", 6993),
        ("years per generation", 28.6),
        ("autosomal effective size anchor, diploid (PSMC, ref 3)", 1447),
    ]):
        ws.cell(end + 2 + k, 1, lab).font = BODY
        ws.cell(end + 2 + k, 2, val).font = BODY
    ws.cell(end + 2 + 8, 1, "Generated by build_supp_workbook.py.").font = SMALL
    return ws


def tab_baseline(wb):
    ws = wb.create_sheet("Baseline parameters")
    hdr = ["Framework", "Parameter", "Symbol", "Baseline", "Swept over",
           "How the baseline was arrived at"]
    rows = [
        ["Deme (Appendix A)", "demes per side", "grid", 12, "6, 8, 10, 12, 14, 16, 20",
         "Smallest lattice whose measured effective size falls inside the anchor window at more "
         "than one migration rate. See the Effective-size screen tab."],
        ["Deme (Appendix A)", "carrying capacity per deme", "K", 11,
         "26 values from 6 to 46; 23 admissible",
         "Set jointly with grid and m, not independently: only the 55 of 507 measured "
         "combinations within 10% of the anchor are searched."],
        ["Deme (Appendix A)", "migration rate", "m", 0.10, "0.05, 0.08, 0.12, 0.15",
         "Midpoint of the admissible range. NOTE: 0.10 is not itself on the measured ladder, so "
         "it has no directly measured effective size of its own."],
        ["Deme (Appendix A)", "assortative-mating discount", "gamma", 0.12,
         "8 values from 0.08 to 1.00",
         "Not chosen: calibrated per condition to bring modern autosomal ancestry onto target. "
         "0.12 is its value in the null model only."],
        ["Deme (Appendix A)", "modern fitness advantage", "r", 2, "1.5, 2, 3",
         "On the plateau of the response: autosomal ancestry is 7.6% at r = 2, 2.5 and 3 alike, "
         "and rises to 20% only at r = 1.5. See Sensitivity (deme)."],
        ["Deme (Appendix A)", "census", "", None, "",
         "Derived: grid squared times K."],
        ["Band (Appendix B)", "census", "N", 1750, "",
         "The size at which measured autosomal drift matches the same anchor of 1,447 diploids. "
         "Larger than the deme census because bands reproduce and die as units, which raises "
         "the variance in reproductive success and lowers the ratio of effective to census size."],
        ["Band (Appendix B)", "optimal band size", "K", 25, "10, 15, 25, 40, 60, 90",
         "Ethnographic rather than fitted: the minimal co-resident foraging band is "
         "conventionally about 25 people and observed forager bands cluster at 25-50."],
        ["Band (Appendix B)", "number of bands", "", None, "",
         "Derived: census divided by optimal band size."],
        ["Band (Appendix B)", "exogamy rate", "e", 0.19, "7 values from 0.08 to 0.56",
         "The calibrated knob of this framework, playing the role gamma plays in the deme "
         "model; reset per condition, so 0.19 is the matrilineal null value only."],
        ["Band (Appendix B)", "modern fitness advantage", "r", 2, "1.5, 2, 3",
         "Matched to the deme framework so the two are comparable."],
        ["Band (Appendix B)", "density regulation", "alpha", 0.8, "0.4, 0.8, 1.6",
         "A band at twice the mean local density has expected fertility multiplied by about "
         "0.45. Swept at all three values in the one-at-a-time scan: autosomal ancestry moves "
         "from 5.1% to 4.2% and Y retention from 3.8% to 2.9% across the range."],
        ["Band (Appendix B)", "band relocation step, km", "sigma_b", 60, "30, 60, 120",
         "Held with p at 0.45; over many generations only the diffusion coefficient p times "
         "sigma_b squared matters."],
        ["Band (Appendix B)", "exogamous dispersal, km", "sigma_e", 80, "40, 80, 160",
         "Also the kernel width for local density, so it sets both gene flow between bands and "
         "the spatial scale of competition."],
        ["Band (Appendix B)", "female share of exogamy", "beta", 0, "0 to 1",
         "0 is matrilineality, with only men dispersing; 1 is patrilineality."],
        ["Both", "effective-size anchor, diploid", "Ne", 1447, "window 1,302-1,592",
         "Time-weighted harmonic mean of PSMC trajectories of three Neandertal-lineage genomes "
         "over the window between admixture and sampling (250-50 ka), ref 3."],
    ]
    hdr_row, end = write_table(
        ws, hdr, rows, status=CONFIG,
        note="Every parameter of both frameworks, its baseline value, the range it is swept "
             "over, and the reason it has that value.")
    # the two derived cells are formulas, so they follow if a baseline is edited
    grid_r = hdr_row + 1
    k_r = hdr_row + 2
    census_r = hdr_row + 6
    ws.cell(census_r, 4).value = f"=D{grid_r}^2*D{k_r}"
    n_r = hdr_row + 7
    bandk_r = hdr_row + 8
    ws.cell(hdr_row + 9, 4).value = f"=ROUND(D{n_r}/D{bandk_r},0)"
    for r in (census_r, hdr_row + 9):
        ws.cell(r, 4).font = BODY
        ws.cell(r, 4).border = BOX
    ws.column_dimensions["F"].width = 70
    for r in range(hdr_row + 1, end):
        ws.cell(r, 6).alignment = Alignment(wrap_text=True, vertical="top")
        ws.cell(r, 5).alignment = Alignment(wrap_text=True, vertical="top")
    return ws, len(rows)


def tab_ne_screen(wb):
    hdr, rows = read_csv(os.path.join(CL, "ne_screen.csv"))
    ws = wb.create_sheet("Effective-size screen")
    ws["A1"] = "Anchor, diploid"
    ws["A1"].font = H2
    ws["B1"] = 1447
    ws["A2"] = "Tolerance"
    ws["A2"].font = H2
    ws["B2"] = 0.10
    ws["B2"].number_format = "0%"
    ws["A3"] = "Lower bound"
    ws["A3"].font = H2
    ws["B3"] = "=B1*(1-B2)"
    ws["A4"] = "Upper bound"
    ws["A4"].font = H2
    ws["B4"] = "=B1*(1+B2)"
    for a in ("A1", "A2", "A3", "A4", "B1", "B2", "B3", "B4"):
        if not ws[a].font.bold:
            ws[a].font = BODY
    ws["D1"] = "Combinations measured"
    ws["D1"].font = H2
    ws["D2"] = "Admissible on the measured size"
    ws["D2"].font = H2
    ws["D3"] = "Distinct K values measured"
    ws["D3"].font = H2

    out_hdr = hdr + ["within window (recomputed)"]
    hdr_row, end = write_table(
        ws, out_hdr, rows, start=6, status=CONFIG,
        note="Realized effective size of every (grid, K, m) combination, measured by resetting "
             "marker frequencies to one half and regressing the decay of expected "
             "heterozygosity. The last column is recomputed here from the bounds above, so "
             "editing the tolerance re-screens the table.")
    ne_col = get_column_letter(hdr.index("ne_auto_diploid") + 1)
    last = get_column_letter(len(out_hdr))
    for r in range(hdr_row + 1, end):
        ws.cell(r, len(out_hdr)).value = (
            f'=IF(AND({ne_col}{r}>=$B$3,{ne_col}{r}<=$B$4),"yes","no")')
        ws.cell(r, len(out_hdr)).font = BODY
        ws.cell(r, len(out_hdr)).border = BOX
    ws["E1"] = f"=COUNTA(A{hdr_row+1}:A{end-1})"
    ws["E2"] = f'=COUNTIF({last}{hdr_row+1}:{last}{end-1},"yes")'
    kcol = get_column_letter(hdr.index("K") + 1)
    ws["E3"] = f"=SUMPRODUCT((COUNTIF(OFFSET($A$1,0,{hdr.index('K')},1,1),1)>0))"
    ws["E3"] = 26          # a distinct-count needs an array function LibreOffice cannot spill
    ws["F3"] = "counted in Python; 23 of them are admissible at one or more grid sizes"
    ws["F3"].font = SMALL
    for a in ("E1", "E2", "E3"):
        ws[a].font = BODY
    ws.column_dimensions["D"].width = 32
    ws.column_dimensions["F"].width = 58
    return ws, len(rows)


def tab_landscape_rules(wb):
    ws = wb.create_sheet("Landscape definition")
    hdr = ["Step", "Rule", "Coordinates", "Reason"]
    rows = [
        [1, "Frame", "longitude -11.0 to 32.5 E, latitude 34.0 to 57.0 N, step 0.05 degrees",
         "The extent of the published Neandertal niche reconstruction (ref 53), on a regular "
         "lon/lat lattice."],
        [2, "Land", "land/sea from a global land mask at the same resolution",
         "Only land is habitable; the sea both bounds the arena and reflects movement."],
        [3, "Northern limit",
         "PCHIP spline through (-11, 47), (-5, 49), (0, 51.5), (5, 53.5), (9.5, 54.6), "
         "(16.0, 53.4), (22.0, 51.5), (27.0, 49.0), (30.5, 46.6), (33.0, 44.3), (34.0, 44.0)",
         "One smooth curve, so the boundary has no kinks. Follows the Channel coast and "
         "northern Germany in the west, then arcs southeast to enter the Black Sea west of "
         "Crimea."],
        [4, "Northeast cut",
         "convex chord from (9.5 E, 54.6 N) to (28.5 E, 42.0 N), bowed 2.2 degrees northeast "
         "at midspan",
         "The niche model includes the east European plain and Pontic steppe, which have no "
         "Neandertal fossil record. Removing them focuses the arena on the fossil-bearing west "
         "and south."],
        [5, "Great Britain included",
         "longitude -6.5 to 2.0 E, latitude 49.8 to 54.0 N",
         "Reachable and attested, up to about 54 N; northern Britain is outside the frame."],
        [6, "Africa excluded",
         "land below 35.6 N west of 11 E; land below 34.0 N between 11 and 33 E",
         "The Maghreb and the Libyan and Egyptian coasts are outside the range, while the "
         "Levant strip is kept."],
        [7, "Asia excluded",
         "land east of 26 E and below 42 N, except Thrace north of the line from (26 E, 40.3 N) "
         "to (29.05 E, 41.0 N)",
         "Anatolia, the Levant and the Caucasus are drawn as context but are not habitable. "
         "The expansion therefore begins at the straits with Asia closed behind it."],
        [8, "Crimea excluded", "northern limit held below 44.3 N across Crimean longitudes",
         "A simplification, noted as such: Kiik-Koba and the Zaskalnaya cluster are "
         "conventionally attributed to Neandertals."],
        [9, "Modern seed", "28.95 E, 41.05 N",
         "The European shore of the Bosphorus at the site of present-day Istanbul."],
        [10, "Projection for the maps", "Albers equal-area conic, standard parallels 38 and "
             "52 N, origin 17 E / 43 N",
         "So that no region is visually inflated relative to another. The mask itself is stored "
         "on the lon/lat lattice and is projection-free."],
    ]
    hdr_row, end = write_table(
        ws, hdr, rows, status=CONFIG,
        note="The rules that generate the band-model arena, in the order they are applied. This "
             "is the table the note refers to as holding “the exact coordinates of the "
             "landscape I use”. Generated by band_model/make_range_mask.py; the resulting "
             "boundary is on the next tab.")
    ws.column_dimensions["A"].width = 7
    ws.column_dimensions["B"].width = 24
    ws.column_dimensions["C"].width = 64
    ws.column_dimensions["D"].width = 64
    for r in range(hdr_row + 1, end):
        for c in (2, 3, 4):
            ws.cell(r, c).alignment = Alignment(wrap_text=True, vertical="top")
    return ws, len(rows)


def tab_landscape_outline(wb):
    """Trace the boundary of the stored mask so the arena is available as coordinates."""
    import matplotlib
    matplotlib.use("Agg")
    import matplotlib.pyplot as plt

    z = np.load(os.path.join(BAND, "range_mask.npz"))
    lon, lat, mask = z["lon"], z["lat"], z["mask"]
    fig, ax = plt.subplots()
    cs = ax.contour(lon, lat, mask.astype(float), levels=[0.5])
    segs = []
    for coll in cs.allsegs:
        for seg in coll:
            if len(seg) >= 20:
                segs.append(seg)
    plt.close(fig)
    segs.sort(key=len, reverse=True)

    rows = []
    for i, seg in enumerate(segs[:12], start=1):
        step = max(1, len(seg) // 400)          # thin to at most ~400 vertices per part
        for x, y in seg[::step]:
            rows.append([i, round(float(x), 3), round(float(y), 3)])
    ws = wb.create_sheet("Landscape outline")
    hdr = ["Part", "Longitude (E)", "Latitude (N)"]
    write_table(
        ws, hdr, rows, status=CONFIG,
        note="The boundary of the habitable arena, traced from band_model/range_mask.npz at the "
             "0.5 contour and thinned to at most 400 vertices per part. Part 1 is the mainland; "
             "later parts are islands, largest first. The stored mask is a 461 by 871 boolean "
             "lattice at 0.05 degrees, which is the object the simulation actually uses; this "
             "outline is for plotting and for checking the arena against a map.")
    return ws, len(rows)


def tab_sens_deme(wb):
    hdr, rows = read_csv(os.path.join(HERE, "figure3_full.csv"))
    ws = wb.create_sheet("Sensitivity (deme)")
    # THE BASELINE DISAGREEMENT IS GONE, AND IT WAS NOT A GEOMETRY DIFFERENCE. This tab used
    # to warn that "the grid and K panels disagree with the m, gamma and r panels at their
    # shared baseline by far more than replicate noise". The cause was found on 14 Sep: the
    # collector's glob was pooling the superseded 20x20 / K = 20 run from August with the
    # anchored September rerun, because the old files were never moved aside on this machine.
    # With the cutoff in place all five panels report exactly 7.55% autosomal, 2.28% Y and 239
    # generations at the shared baseline -- identical, as they must be, since they are the same
    # configuration sampled from the same 2,000 replicates.
    write_table(
        ws, hdr, rows, status=SIM,
        note="One parameter varied at a time about the deme baseline, all others held fixed, "
             "2,000 replicates per point. This is the source for Figure S3.5. The five panels "
             "that contain the baseline (grid 12, K, m = 0.10, gamma = 0.12, r = 2) agree "
             "there exactly, because they are the same configuration drawn from the same "
             "replicates. Columns: panel is the parameter varied, value its setting, "
             "auto/Y/mtDNA/X the modern proportions, P_* the fixation and ten-male sampling "
             "probabilities, ext_gen the generation of archaic extinction. A replicate in "
             "which no males survive to the sampling generation leaves the uniparental "
             "frequencies undefined and is excluded from those averages only; this affects "
             "one replicate of 2,000 at K = 20 and at K = 40 and three at K = 24.")
    return ws, len(rows)


def tab_sens_band(wb):
    hdr, rows = read_csv(os.path.join(BAND, "bandsi_summary.csv"))
    fam = hdr.index("family")
    rows = [r for r in rows if r[fam] == "scanb"]
    ws = wb.create_sheet("Sensitivity (band)")
    write_table(
        ws, hdr, rows, status=SIM,
        note="One parameter varied at a time about the band baseline, all others held fixed, "
             "2,000 replicates per point, matching the deme framework. This is the source for "
             "Figure S3.6, and every parameter of that figure is present, including density "
             "regulation alpha. end_gen_med is the median generation at which the uniparental "
             "markers are absorbed, which is this framework's analogue of the deme framework's "
             "time to archaic extinction rather than the identical quantity.")
    return ws, len(rows)


def tab_mtdna(wb):
    """Both solid arms of Figure S3.4, which used to be one.

    THE BAND ARM WAS MISSING FROM THIS TAB ENTIRELY. The figure draws two migration-pulse
    curves, a blue panmictic one and a red band one, and only the panmictic one was shipped;
    a reader could not check the red trace against anything. It is added here as extra rows
    with a `framework` column, which also makes the two ladders comparable at a glance --
    they now run over the same rungs, out to s = 10%, after the band sweep was extended on
    21 Sep 2026 (it had stopped at 3%, mid-rise, which is how the omission came to light).
    """
    hdr, rows = read_csv(os.path.join(HERE, "fixation_curves_mtdna.csv"))
    hdr = ["framework"] + hdr
    rows = [["panmictic (deme baseline)"] + r for r in rows]

    bh, br = read_csv(os.path.join(BAND, "bandsi_summary.csv"))
    ix = {k: bh.index(k) for k in ("family", "N", "s_mt", "p_mt", "n")}
    band = [r for r in br if r[ix["family"]] == "mtcurve"]
    band.sort(key=lambda r: (int(r[ix["N"]]), float(r[ix["s_mt"]])))
    for r in band:
        n = int(r[ix["n"]])
        pf = 100.0 * float(r[ix["p_mt"]])
        se = 100.0 * (float(r[ix["p_mt"]]) * (1 - float(r[ix["p_mt"]])) / n) ** 0.5
        rows.append(["band", int(r[ix["N"]]), None, float(r[ix["s_mt"]]),
                     round(pf, 1), n, round(se, 3)])

    ws = wb.create_sheet("mtDNA fixation curves")
    write_table(
        ws, hdr, rows, status=SIM,
        note="Probability that modern mtDNA replaces archaic mtDNA after a 5% migration "
             "pulse, against the fitness gap. Both solid curves of Figure S3.4 are here: the "
             "panmictic rows carry the mtDNA effective copy number and are the source of the "
             "conversion from fitness gap to probability of replacement used in Supplementary "
             "Information section 1; the band rows are the same experiment inside the roving-"
             "band landscape, indexed by census rather than by copy number, at 1,000 "
             "replicates a rung. The figure draws locus_Ne = 750 for the panmictic arm and "
             "census 1,750 for the band arm.")
    return ws, len(rows)


def tab_cultural(wb):
    """The five cultural mechanisms, from the scored Cartesian search.

    REBUILT 21 Sep, AND THE OLD VERSION WAS THE WORST TAB IN THE WORKBOOK. It pooled seven
    CSVs written by the pre-Cartesian grid scripts (kin2, kin3, mech3, mech4, mech5, asym,
    clan), which recorded ancestry and a success fraction but NO expansion duration, so the
    tab could not be tested for admissibility, could not be placed on Figure S3.2's axes, and
    carried a status line saying so. All five mechanisms have since been run through the same
    two-stage Cartesian pipeline as every other mechanism in the note -- out_cult1_* screened,
    out_cult2_* scored -- so this tab is now the same kind of object as the DMI and drive tab
    beside it: one row per scored parameter combination, with the acceptance test recomputed
    live.

    The clan-landscape sweep has no counterpart in the Cartesian search and is dropped rather
    than carried at the old precision; it is one of the scenarios the note reports as
    "simulations not shown", and shipping it in a form that cannot be checked against the
    others invites exactly the comparison it does not support.
    """
    import glob as _glob
    import json as _json
    import math as _math
    from figure2c_fountain import AUT_LO, AUT_HI

    LABEL = {"inbreed": "inbreeding avoidance (matrilineal kin ban)",
             "obligate": "obligate dispersal from the natal group",
             "rhoreturn": "return to the father's natal group",
             "skew": "descent marking (male reproductive skew)",
             "asym": "asymmetric interbreeding"}
    ws = wb.create_sheet("Cultural mechanisms")
    hdr = ["mechanism", "mechanism key", "mechanism value", "framework", "grid", "K", "m",
           "gamma", "r", "e", "sigma_b", "sigma_e", "alpha", "female bias", "beta",
           "replicates", "established", "modern autosomal %", "expansion generations",
           "p(10 males all modern at both)", "admissible"]
    out = []
    for fw in ("deme", "band"):
        for p in sorted(_glob.glob(os.path.join(CL, f"out_cult2_{fw}", "*.json"))):
            try:
                s2 = _summary(p)
            except Exception:
                continue
            if s2.get("joint") is None or s2.get("mech") not in LABEL:
                continue
            s1 = os.path.join(CL, f"out_cult1_{fw}", os.path.basename(p))
            g = None
            if os.path.exists(s1):
                try:
                    g = _summary(s1).get("end_gen")
                except Exception:
                    g = None
            aut = s2.get("aut")
            est = (s2.get("n_est") or 0) / max(s2.get("n") or 1, 1)
            okg = g is not None and not (isinstance(g, float) and _math.isnan(g)) \
                and 100 <= g <= 1000
            oka = aut is not None and AUT_LO <= aut <= AUT_HI
            out.append([
                LABEL[s2["mech"]], s2.get("mech"), s2.get("val"), fw, s2.get("grid"),
                s2.get("K"), s2.get("m"), s2.get("gamma"), s2.get("r"), s2.get("e"),
                s2.get("sigma_b"), s2.get("sigma_e"), s2.get("alpha_d"),
                s2.get("female_bias"), s2.get("beta"), s2.get("n"), round(est, 3),
                None if aut is None else round(aut, 3), g, round(s2["joint"], 4),
                "yes" if (oka and okg and est > 0.5) else "no"])
    n_ok = sum(1 for r in out if r[-1] == "yes")
    best = max((r[-2] for r in out if r[-1] == "yes"), default=0.0)
    write_table(
        ws, hdr, out, status=SIM,
        note=f"One row per scored parameter combination for each of the five cultural "
             f"mechanisms, in both frameworks, from the same two-stage Cartesian search that "
             f"produced the DMI and drive tab. {len(out):,} rows, of which {n_ok:,} are "
             f"admissible at the note's criteria (modern autosomal {AUT_LO:g}-{AUT_HI:g}%, "
             f"expansion completing in 100-1,000 generations, establishment above 50%). The "
             f"highest retention probability anywhere in the admissible set is {best:.3f}, "
             f"which is the quantitative form of the note's statement that none of these "
             f"mechanisms explains the observation. Reduced F1-male fertility, the one "
             f"cultural mechanism that does work, is a route in its own right and its cells "
             f"are in the DMI and drive tab under the mechanism key theta.")
    return ws, len(out)


# mechanism key -> (label, family, route). THE ROUTE COLUMN IS BACK, as of 21 Sep 2026.
# It was dropped on 14 Sep because Figure 2b had been reordered (incompatibilities became
# Route 1 and direct selection Route 2) while the note's prose still used the old numbering,
# so "Route 2" would have meant one thing against the figures and another against the text.
# v90 renumbered the prose and v91 renumbered the two appendix parameter lists that v90 had
# missed, so the two now agree and the number is unambiguous again. The family column stays:
# it is the same fact stated in a form that survives any future renumbering, and it is what
# the reader needs in order to see why the Y-acting architectures behave as a group.
ARCH_LABEL = {
    "sY": "selection on the Y conditioned on group identity (matching patriline favoured)",
    "xdrive": "X-linked sex distortion",
    "xdrive_c10": "X-linked sex distortion, 10% fertility cost",
    "dmi_mtY": "mtDNA × Y incompatibility",
    "dmi_mtAutoDom": "mtDNA × autosome, dominant",
    "dmi_mtAutoAdd": "mtDNA × autosome, additive",
    "dmi_mtAutoRec": "mtDNA × autosome, recessive",
    "dmi_mtXDom": "mtDNA × X, dominant",
    "dmi_XY": "X × Y incompatibility",
    "dmi_autoAuto": "autosome × autosome incompatibility",
    "dmi_Xprdm9": "modern X × archaic PRDM9-A (s = 0.5%)",
    "dmi_prdm9X_s0": "modern PRDM9-A × archaic X (s = 0)",
    "dmi_prdm9X_s005": "modern PRDM9-A × archaic X (s = 0.5%)",
    "dmi_prdm9X_s05": "modern PRDM9-A × archaic X (s = 5%)",
    "restorers": "polygenic mitonuclear restorers, dominant",
    "restorersAdd": "polygenic mitonuclear restorers, additive",
    "theta": "reduced F1-male fertility",
    "twowave": "two waves, matrilineal and patrilineal",
}
# Route 1 the incompatibilities (including X-linked sex distortion, which is an
# incompatibility acting on the Y), Route 2 selection on the marker itself, Route 3 two waves,
# Route 4 the cultural mechanism. This is Figure 2b's order and, since v90, the note's.
ARCH_ROUTE = {
    "sY": 2,
    "xdrive": 1, "xdrive_c10": 1,
    "dmi_mtY": 1, "dmi_mtAutoDom": 1, "dmi_mtAutoAdd": 1, "dmi_mtAutoRec": 1,
    "dmi_mtXDom": 1, "dmi_XY": 1, "dmi_autoAuto": 1,
    "dmi_Xprdm9": 1, "dmi_prdm9X_s0": 1, "dmi_prdm9X_s005": 1, "dmi_prdm9X_s05": 1,
    "restorers": 1, "restorersAdd": 1,
    "twowave": 3,
    "theta": 4,
}
ARCH_FAMILY = {
    "sY": "selection on the marker itself",
    "xdrive": "incompatibility acting on the Y (sex distortion)",
    "xdrive_c10": "incompatibility acting on the Y (sex distortion)",
    "dmi_mtY": "incompatibility acting on the Y",
    "dmi_mtAutoDom": "incompatibility through a partner locus",
    "dmi_mtAutoAdd": "incompatibility through a partner locus",
    "dmi_mtAutoRec": "incompatibility through a partner locus",
    "dmi_mtXDom": "incompatibility through a partner locus",
    "dmi_XY": "incompatibility through a partner locus",
    "dmi_autoAuto": "incompatibility through a partner locus",
    "dmi_Xprdm9": "incompatibility through a partner locus (PRDM9)",
    "dmi_prdm9X_s0": "incompatibility through a partner locus (PRDM9)",
    "dmi_prdm9X_s005": "incompatibility through a partner locus (PRDM9)",
    "dmi_prdm9X_s05": "incompatibility through a partner locus (PRDM9)",
    "restorers": "incompatibility through a partner locus (restorers)",
    "restorersAdd": "incompatibility through a partner locus (restorers)",
    "theta": "cultural",
    "twowave": "two waves",
}


def tab_architectures(wb):
    """The per-combination table that the note's architecture claims summarise.

    NO LONGER STRUCTURE ONLY. This tab held a single italicised example row and the word
    "pending" because stage-2 scoring had not finished. It has, so the tab is now the real
    thing: every scored parameter combination across all six sweeps and both frameworks, with
    the acceptance test recomputed live rather than baked in.

    The `admissible` column is computed here from the same three criteria the note states --
    autosomal 3-7%, expansion completing in 100-1,000 generations, establishment above half --
    so a reader can change the window in the spreadsheet and watch every claim move. That is
    the point of shipping the per-combination table rather than only the medians.
    """
    import glob as _glob
    import json as _json
    import math as _math
    from figure2c_fountain import AUT_LO, AUT_HI

    ws = wb.create_sheet("DMI and drive search")
    hdr = ["architecture", "route", "family", "framework", "mechanism", "mechanism value", "grid", "K",
           "m", "gamma", "r", "e", "sigma_b", "sigma_e", "alpha", "female bias", "beta", "s_mt",
           "replicates", "established", "modern autosomal %", "expansion generations",
           "p(10 males all modern at both)", "admissible"]
    out = []
    for pre in ("routes", "dmi", "dmi2", "xdc", "xd2", "tw"):
        for fw in ("deme", "band"):
            for p in sorted(_glob.glob(os.path.join(CL, f"out_{pre}2_{fw}", "*.json"))):
                try:
                    s = _summary(p)
                except Exception:
                    continue
                if s.get("joint") is None:
                    continue
                s1 = os.path.join(CL, f"out_{pre}1_{fw}", os.path.basename(p))
                g = None
                if os.path.exists(s1):
                    try:
                        g = _summary(s1).get("end_gen")
                    except Exception:
                        g = None
                aut = s.get("aut")
                est = (s.get("n_est") or 0) / max(s.get("n") or 1, 1)
                okg = g is not None and not (isinstance(g, float) and _math.isnan(g)) \
                    and 100 <= g <= 1000
                oka = aut is not None and AUT_LO <= aut <= AUT_HI
                mech = s.get("mech")
                out.append([
                    ARCH_LABEL.get(mech, mech), ARCH_ROUTE.get(mech),
                    ARCH_FAMILY.get(mech, ""), fw, mech,
                    s.get("val"), s.get("grid"), s.get("K"), s.get("m"), s.get("gamma"),
                    s.get("r"), s.get("e"), s.get("sigma_b"), s.get("sigma_e"),
                    s.get("alpha_d"), s.get("female_bias"), s.get("beta"), s.get("s_mt"),
                    s.get("n"), round(est, 3), None if aut is None else round(aut, 3),
                    g, round(s["joint"], 4),
                    "yes" if (oka and okg and est > 0.5) else "no"])
    n_ok = sum(1 for r in out if r[-1] == "yes")
    write_table(
        ws, hdr, out, status=SIM,
        note=f"One row per scored parameter combination in the Cartesian search, for every "
             f"incompatibility and drive architecture, in both frameworks. {len(out):,} rows, "
             f"of which {n_ok:,} are admissible at the acceptance window in force when this "
             f"workbook was built (modern autosomal {AUT_LO:g}-{AUT_HI:g}%, expansion "
             f"completing in 100-1,000 generations, establishment above 50%). Every "
             f"architecture claim in the note is a median or a maximum of the "
             f"p(10 males all modern) column over the admissible rows for that architecture, "
             f"so this tab is what makes those claims checkable. A blank cell means the "
             f"parameter does not apply to that framework: grid, K, m and gamma are the deme "
             f"model's, e, sigma_b, sigma_e and alpha the band model's.")
    return ws, len(out)


def tab_screen_only(wb):
    """Every cell that was screened at 10 replicates and never promoted.

    DR: "I never see the 10 rep examples in the Supplementary Notes (the ones we didn't raise
    higher because they failed to match)." He is right, and it was a real hole. The search is
    two-stage -- 10 replicates to screen, then 100-200 to score anything that lands in the
    acceptance box -- and until now the workbook shipped only the scored end of it. Figure
    S3.2's caption says the scatter is of mixed precision by construction, and a reader had no
    way to see the imprecise half or to check how far outside the box it sits.

    These are the cells the search LOOKED AT AND PUT DOWN: 66,929 of them, against 30,919
    promoted. Each carries its 10-replicate estimate of autosomal ancestry, expansion duration
    and establishment, and the reason it was not promoted, which is always that at least one
    of the three acceptance criteria failed at the screen. A handful sit just outside the box
    on one criterion; those are the cells the 100-replicate re-screen was built to catch, and
    where it caught them the cell appears in the scored tabs instead of here.

    TEN REPLICATES IS A COARSE NUMBER AND THE TAB SAYS SO. An autosomal mean from 10 runs has
    a standard error around a percentage point at these densities, which is why a cell near a
    boundary is re-screened rather than judged on it. Nothing in the note rests on a number
    from this tab; it is here so that the two-stage design is visible rather than asserted.
    """
    import glob as _glob
    import json as _json
    import math as _math
    from figure2c_fountain import AUT_LO, AUT_HI

    ws = wb.create_sheet("Screened, not scored")
    hdr = ["architecture", "route", "family", "framework", "mechanism", "mechanism value", "grid", "K",
           "m", "gamma", "r", "e", "sigma_b", "sigma_e", "alpha", "female bias", "beta",
           "replicates", "established", "modern autosomal %", "expansion generations",
           "why not promoted"]
    out, n_scored = [], 0
    for pre in ("routes", "dmi", "dmi2", "xdc", "xd2", "tw", "cult"):
        for fw in ("deme", "band"):
            scored = {os.path.basename(q) for q in
                      _glob.glob(os.path.join(CL, f"out_{pre}2_{fw}", "*.json"))}
            n_scored += len(scored)
            # ONE ROW PER CELL, AT ITS BEST ESTIMATE -- the same precedence screen.py applies.
            # A cell that was screened at 10 and re-screened at 100 without being promoted has
            # a file in both directories; listing both would double-count it and show the
            # coarser number beside the finer one as though they were different cells.
            best = {}
            for d in (f"out_{pre}1_{fw}", f"out_{pre}1hi_{fw}"):
                for q in _glob.glob(os.path.join(CL, d, "*.json")):
                    if os.path.basename(q) not in scored:
                        best[os.path.basename(q)] = q
            for p in sorted(best.values()):
                if True:
                    try:
                        s2 = _summary(p)
                    except Exception:
                        continue
                    aut, g = s2.get("aut"), s2.get("end_gen")
                    n = s2.get("n") or 0
                    est = (s2.get("n_est") or 0) / max(n, 1)
                    bad = []
                    if aut is None or aut != aut:
                        bad.append("no autosomal estimate")
                    elif not (AUT_LO <= aut <= AUT_HI):
                        bad.append(f"autosomal {aut:.1f}% outside {AUT_LO:g}-{AUT_HI:g}%")
                    if g is None or (isinstance(g, float) and _math.isnan(g)):
                        bad.append("expansion did not complete")
                    elif not (100 <= g <= 1000):
                        bad.append(f"{g:.0f} generations outside 100-1,000")
                    if est <= 0.5:
                        bad.append(f"established in {est:.0%} of replicates")
                    out.append([
                        ARCH_LABEL.get(s2.get("mech"), s2.get("mech")),
                        ARCH_ROUTE.get(s2.get("mech"), 4),
                        ARCH_FAMILY.get(s2.get("mech"), "cultural"), fw, s2.get("mech"),
                        s2.get("val"), s2.get("grid"), s2.get("K"), s2.get("m"),
                        s2.get("gamma"), s2.get("r"), s2.get("e"), s2.get("sigma_b"),
                        s2.get("sigma_e"), s2.get("alpha_d"), s2.get("female_bias"),
                        s2.get("beta"), n, round(est, 2),
                        None if aut is None else round(aut, 2),
                        None if g is None else g, "; ".join(bad) or "promoted elsewhere"])
    write_table(
        ws, hdr, out, status=SIM,
        note=f"The first stage of the two-stage search: {len(out):,} parameter combinations "
             f"that were screened and not carried forward, against the {n_scored:,} that "
             f"were. Almost all of these carry 10 replicates; the few at 100 are cells that "
             f"sat near a boundary and were re-screened before being put down, and the "
             f"replicates column says which is which. A cell is promoted to 100-200 "
             f"replicates only if its screen estimate "
             f"lands inside the acceptance box (modern autosomal {AUT_LO:g}-{AUT_HI:g}%, "
             f"expansion completing in 100-1,000 generations, establishment above 50%); the "
             f"last column gives the criterion or criteria that failed. Where a re-screen "
             f"promoted a cell it appears in the scored tabs rather than here. Ten "
             f"replicates is a "
             f"coarse estimate -- the autosomal standard error is of order a percentage point "
             f"-- so these rows show the shape of the search rather than supporting any "
             f"number in the note.")
    return ws, len(out)


def tab_drive(wb):
    """Figure S3.3: the sex-distortion ladder and the heritable-resistance ladder."""
    import fig_s34_drive_and_escape as f
    ws = wb.create_sheet("X-drive and resistance")
    hdr = ["panel", "framework", "fertility cost", "daughters among an F1 male's offspring (%)",
           "heritable resistance rate (%)", "value (%)", "quantity"]
    out = []
    for (fw, cost) in (("deme", 0.0), ("deme", 0.1), ("band", 0.0), ("band", 0.1)):
        arm = f.deme_arm(cost) if fw == "deme" else f.band_arm(cost)
        for d, v in arm:
            out.append(["(a) drive", fw, cost, d, "", round(v, 2),
                        "p(10 males all modern at both markers)"])
    for eps, v in f.deme_resistance(0.95):
        # THE QUANTITY LABEL WAS WRONG, and it described a measure this column stopped
        # carrying. deme_resistance returned the mean modern Y frequency among males when
        # this tab was written; it was changed to the ten-male joint statistic so that panel
        # (b) reports the same thing as panels (a) and (c), and the label did not follow.
        out.append(["(b) resistance", "deme", 0.0, 97.5, 100 * eps, round(v, 2),
                    "p(10 males all modern at both markers)"])
    write_table(
        ws, hdr, out, status=SIM,
        note="Figure S3.3. Panel (a) is the sex-distortion ladder: how strongly an X-linked "
             "distorter must skew the sex ratio of an F1 male's offspring for the modern Y to "
             "survive, with and without a 10% fertility cost to expressing males. It is "
             "computed from cluster/out_xdrvdeme for the deme framework and the band suite "
             "for the band framework, with gamma refitted at every rung so that modern "
             "autosomal ancestry lands on 5% and the rungs are therefore comparable with each "
             "other and with the rest of the note. Panel (b) is the heritable-resistance "
             "ladder, computed from the per-replicate output in cluster/out_xdrv2 at the "
             "anchored geometry. Both panels report the same quantity as every other "
             "retention number in the note: the probability that ten males sampled at random "
             "from the whole landscape are modern at both mtDNA and the Y. 400 replicates per "
             "point in panel (a), 900 in panel (b).")
    return ws, len(out)


def tab_band_production(wb):
    """The band framework's production sweeps: every (scenario, mechanism, value) cell of
    bandratchet_submit.py, 400 replicates each, s_mt drawn per replicate from the accepted
    ABC posterior. This is the table behind the Figure 3a replicate's cell (fexp, s = 0.16)
    and behind the Scenario 2 statements in Supplementary Information section 3 about how
    often the band framework retains the modern Y at a given prestige advantage. Selection on
    the Y is conditioned on band identity (10 Oct 2026); the s rows are from the rerun under
    that rule, the theta rows are unchanged by it."""
    hdr, rows = read_csv(os.path.join(BAND, "band_ratchet_summary.csv"))
    ws = wb.create_sheet("Band production sweep")
    write_table(
        ws, hdr, rows, status=SIM,
        note="Production ensembles of the band framework, 400 replicates per cell, exogamy "
             "rate e_used calibrated per cell so that modern autosomal ancestry lands at 5% "
             "(bandprod_calibration.csv). Scenarios: fexp = matrilocal range expansion, nexp = "
             "non-sex-biased expansion, and the two stable-boundary (migration-pulse) "
             "comparators. p_Y is the probability that the modern Y is fixed; p_joint that both "
             "uniparental markers are; p10 that ten sampled males are modern at both. Selection "
             "on the Y (sweep = s) is conditioned on band identity: the patriline matching the "
             "band's identity is favoured, the mismatch penalised. Figure 3a's replicate is the "
             "fexp, s = 0.16 cell (p_Y = 0.73).")
    return ws, len(rows)


def tab_mt_expansion(wb):
    """Figure S3.4's range-expansion arm: modern mtDNA fixing under a patrilineal range expansion."""
    p = os.path.join(HERE, "mt_expansion_curve.csv")
    if not os.path.exists(p):
        return None, 0
    hdr, rows = read_csv(p)
    # The label column still says "patrilocal", which is the wording the collector wrote and
    # the note no longer uses: residence language was replaced by lineal language throughout
    # the figures and the text. Rewritten on display so the workbook and the note agree; the
    # CSV is left alone because it is the raw collector output.
    rows = [[c.replace("patrilocal", "patrilineal").replace("matrilocal", "matrilineal")
             if isinstance(c, str) else c for c in r] for r in rows]
    ws = wb.create_sheet("mtDNA under expansion")
    write_table(
        ws, hdr, rows, status=SIM,
        note="The range-expansion arm of Figure S3.4: the probability that an introgressed modern mtDNA "
             "reaches fixation under a PATRILINEAL range expansion, against the selection "
             "coefficient favouring it. Patrilineality is the only rule for which the question "
             "is live, because under matrilineality the expanding matriline carries modern "
             "mtDNA by construction. Compare the 'mtDNA fixation curves' tab, which is the "
             "same quantity under a migration pulse: at the inferred gap of s = 0.32% the "
             "pulse fixes modern mtDNA 21.8% of the time in the deme framework and 18.4% in "
             "the band, against about 1% for the expansion in the band framework and not once "
             "in a thousand replicates in the deme. 1,000 replicates per point in the deme "
             "framework and 400 in the band.")
    return ws, len(rows)


def tab_ysupp(wb):
    """Comment 30: the de novo Y-suppressor mutation rate against a standing modern Y."""
    import json as _json
    p = os.path.join(CL, "ysupp_sweep.json")
    if not os.path.exists(p):
        return None, 0
    d = _json.load(open(p))
    ws = wb.create_sheet("Y suppressor waiting time")
    hdr = ["scenario", "de novo mutation rate per Y", "standing frequency of suppressing Y",
           "P(suppressor establishes)", "P(population extinct)", "P(distorter fixes)",
           "median generation suppressor appears", "replicates"]
    rows = []
    for r in d["A"]:
        rows.append(["A: no hybridisation", r["mu"], "", round(r["p_est"], 4),
                     round(r["p_ext"], 4), round(r["p_dfix"], 4), r.get("med_gen"), d["reps"]])
    for r in d["B"]:
        rows.append(["B: hybridisation", 0, r["f_S"], round(r["p_est"], 4),
                     round(r["p_ext"], 4), round(r["p_dfix"], 4), "", d["reps"]])
    write_table(
        ws, hdr, rows, status=SIM,
        note="Figure S3.3, right panel. A within-population Wright-Fisher "
             "model (cluster/ysupp_sim.py) of N = 1,500 diploids in which an X-linked "
             "distorter arises as a single copy and skews the offspring of expressing males "
             "to 95% daughters, against a Y-linked suppressor that restores an even sex "
             "ratio. Male fertility is capped at ten offspring per male per generation, so a "
             "population whose male share falls below 10% shrinks and can go extinct; without "
             "that cap a sex-ratio distorter cannot threaten a population at all and the "
             "question has nothing at stake. Scenario A reaches the suppressor only by "
             "mutation; scenario B starts with it present at frequency f, which is the modern "
             "Y arriving at the admixture proportion. Neither spatial engine can express this: "
             "in both, the suppressor is a lineage label fixed at initialisation, with no "
             "mutation process on the Y.")
    return ws, len(rows)


def main():
    wb = Workbook()
    wb.remove(wb.active)

    index = []
    ws, n = tab_baseline(wb)
    index.append(["Baseline parameters", "Every parameter of both frameworks, its baseline "
                  "value, its swept range, and why it has that value",
                  "Appendices A and B", n, CONFIG])
    ws, n = tab_ne_screen(wb)
    index.append(["Effective-size screen", "Measured effective size of every (grid, K, m) "
                  "combination, with the admissibility test recomputed live",
                  "Appendix A", n, CONFIG])
    ws, n = tab_landscape_rules(wb)
    index.append(["Landscape definition", "The rules and coordinates that generate the "
                  "band-model arena, in application order",
                  "Appendix B; this is the table the note's landscape paragraph points to",
                  n, CONFIG])
    ws, n = tab_landscape_outline(wb)
    index.append(["Landscape outline", "The traced boundary of that arena as longitude and "
                  "latitude, mainland and islands", "Figure 3a", n, CONFIG])
    ws, n = tab_sens_deme(wb)
    index.append(["Sensitivity (deme)", "One parameter varied at a time about the deme "
                  "baseline", "Figure S3.5", n, SIM])
    ws, n = tab_sens_band(wb)
    index.append(["Sensitivity (band)", "One parameter varied at a time about the band "
                  "baseline", "Figure S3.6", n, SIM])
    ws, n = tab_band_production(wb)
    index.append(["Band production sweep", "The band framework's calibrated production "
                  "ensembles for every scenario and mechanism value; the source of the Figure 3a "
                  "replicate's cell and of the Scenario 2 retention rates quoted in the note",
                  "Figure 3a; Scenario 2 text", n, SIM])
    ws, n = tab_mtdna(wb)
    index.append(["mtDNA fixation curves", "Probability that modern mtDNA replaces archaic "
                  "mtDNA after a 5% pulse, against fitness gap and copy number",
                  "Figure S3.4", n, CONFIG])
    ws, n = tab_cultural(wb)
    index.append(["Cultural mechanisms", "Every scored parameter combination for the five "
                  "cultural mechanisms, with admissibility recomputed live",
                  "Figure S3.2, cultural panels", n, SIM])
    ws, n = tab_architectures(wb)
    index.append(["DMI and drive search", "One row per scored parameter combination for every "
                  "incompatibility and drive architecture, with admissibility recomputed live",
                  "Figures 3b and S3.2", n, SIM])
    ws, n = tab_screen_only(wb)
    index.append(["Screened, not scored", "Every parameter combination the screen rejected, "
                  "at its screen estimate, with the criterion it failed",
                  "Figures 3b and S3.2, the mixed-precision statement", n, SIM])
    ws, n = tab_drive(wb)
    index.append(["X-drive and resistance", "The sex-distortion ladder and the "
                  "heritable-resistance ladder, both frameworks", "Figure S3.3", n, SIM])
    ws, n = tab_ysupp(wb)
    if ws is not None:
        index.append(["Y suppressor waiting time", "De novo suppressor mutation rate against "
                      "a standing modern Y, with extinction risk", "Figure S3.3, right panel",
                      n, SIM])
    ws, n = tab_mt_expansion(wb)
    if ws is not None:
        index.append(["mtDNA under expansion", "Probability that modern mtDNA fixes under a "
                      "patrilineal range expansion, against the fitness gap",
                      "Figure S3.4, range-expansion arm", n, SIM])

    tab_readme(wb, index)
    wb.save(OUT)
    print(f"wrote {os.path.basename(OUT)}")
    for row in index:
        print(f"  {row[0]:<24} {str(row[3]):>7} rows   {row[2]}")


if __name__ == "__main__":
    main()
```
