## Supplementary Information section 3 code base for "Hypothesis: A modern human range expansion ∼300,000 years ago had a critical role in Neandertal origins": anchors.py

```
#!/usr/bin/env python3
"""
anchors.py -- the single source of truth for population sizes and geometry. (DR, 5 Sep 2026)

WHY THIS FILE EXISTS
The post-admixture size appeared as a literal in at least nine places (panm2/3/4/5_run.py,
joint_submit.py, jcal_submit.py, and the commentary in consweep/gridsweep/mgsweep), written
variously as 1500, 725, 750, 1447 and 1312. Those are not all the same quantity and they drifted
apart: 1447/723 is an exactly 2:1 pair, while "1500/725" was the same pair rounded in opposite
directions. Every script now imports from here instead of restating a number.

THE ANCHOR
Peyregne et al. 2025, Fig. 4A, Neandertal-lineage PSMC curves, digitised by digitize_psmc.py and
reduced by a time-weighted HARMONIC mean over 250-50 ka -- i.e. over the post-admixture interval
that our simulations actually cover:

    autosomal   1,447 diploid   ( = 2,894 gene copies, which is what a tracer fit returns)
    uniparental   723 haploid   ( = exactly half; presumes no sex bias)

MATCHING RULE, settled by DR on 4 Sep 2026: tune every configuration so its measured AUTOSOMAL
effective size is 1,447 diploid, and REPORT the realised uniparental sizes rather than forcing
them. The uniparental anchor is the autosomal one halved, so it assumes no sex bias -- which is
precisely the assumption the sex-biased scenarios exist to violate. Forcing it would erase the
effect we are measuring.

CENSUS VALUES THAT DELIVER THE ANCHOR
These are measured, not assumed: ne_audit.py resets neutral tracers to p = 1/2 and fits the decay
of ln E[p(1-p)] in the production engines themselves.

    PANM_CENSUS = 1312   panmictic_sim: interpolated between N = 1265 (2,715 copies) and
                         N = 1390 (3,188 copies). Realised uniparental ~700 (mt) / ~590 (Y),
                         both below 723 -- report these, do not tune to them.
    BAND_N      = 1750   bandsim: measured 2,844 / 2,777 / 2,807 / 3,025 gene copies across the
                         four band configurations, every one within 4% of 2,894.
    DEME_GRID/K   12/11  spatial_sim2: post-wave census that lands on the same anchor. The
                         published 20x20 / K=20 baseline gives ~8,000, five times too large,
                         which inflated every spatial result that used it.

SENSITIVITY ENVELOPE
A factor of two either side of the anchor. This replaces the old [1000, 1500, 2000, 3000, 6000]
ladder, which mixed post-admixture sizes with ACCUMULATION-phase sizes from 750-250 ka -- a
different epoch, and not a quantity any post-admixture simulation should be run at.
"""

# ---------------------------------------------------------------- the anchor
NE_AUTO_DIPLOID = 1447          # Peyregne, time-weighted harmonic mean, 250-50 ka
NE_AUTO_COPIES = 2 * NE_AUTO_DIPLOID     # 2,894 -- the number a tracer fit returns
NE_UNIPARENTAL = 723            # = NE_AUTO_DIPLOID / 2, no-sex-bias reference only

# ------------------------------------------------- censuses that deliver it
PANM_CENSUS = 1312              # panmictic_sim.py  (migration-pulse comparator)
BAND_N = 1750                   # bandsim.py        (roving-band framework)
DEME_GRID = 12                  # spatial_sim2.py   (deme-grid framework)
DEME_K = 11

# --------------------------------------------------------------- envelope
ENVELOPE = [PANM_CENSUS // 2, PANM_CENSUS, PANM_CENSUS * 2]     # [656, 1312, 2624]

# ------------------------------------------------- the section-1 pairing
# TWO DIFFERENT CENSUSES ARE CORRECT HERE, and conflating them is the error this block exists
# to prevent (I made it on 5 Sep 2026).
#
#   PANM_CENSUS = 1312 is for scenarios calibrated on measured AUTOSOMAL Ne (jcal, joint2's
#   fsb/nsb). There the target is 1,447 diploid and the census that delivers it is 1312,
#   because panmictic_sim is not an ideal Wright-Fisher population.
#
#   SEC1_LADDER is for the mtDNA FIXATION CURVES (panm3) and the sex-bias pulse panel (panm4).
#   There the quantity that matters is the uniparental locus size, and the curves are consumed
#   by Supplementary Information section 1, which integrates each ABC posterior draw against
#   the curve at its PAIRED post-mixture size: locus_Ne = 500 / 750 / 1000, paired with the
#   isolation-phase archaic sizes N_(A-uniparental) = 1500 / 2250 / 3000. panm3's census maps
#   to locus_Ne as N/2, so the ladder must be [1000, 1500, 2000]. Substituting the autosomal
#   envelope here gives locus_Ne 328/656/1312 and section 1 can no longer find its curves.
#
# The central rung, locus_Ne = 750, is section 1's 725 rounded, which is in turn the Peyregne
# anchor of 723. The three numbers are the same quantity; 3000 and 6000 are NOT -- those were
# accumulation-phase sizes from 750-250 ka and have been dropped.
SEC1_LADDER = [1000, 1500, 2000]        # census -> locus_Ne 500 / 750 / 1000
SEC1_PRIMARY = 1500                     # census -> locus_Ne 750 = section 1's 725 = anchor 723

# ------------------------------------------------------------------ timing
GEN_YEARS = 28.6                # years per generation
SAMPLE_GEN = 6993               # 200 kyr: admixture at 250 ka -> sampling at 50 ka


def describe():
    """One line for a figure caption or a collector header."""
    return (f"autosomal Ne {NE_AUTO_DIPLOID:,} diploid (Peyregne et al. 2025, Fig. 4A, "
            f"time-weighted harmonic mean 250-50 ka); panmictic census {PANM_CENSUS:,}, "
            f"band N {BAND_N:,}, deme grid {DEME_GRID}x{DEME_GRID} K={DEME_K}; "
            f"sampled at generation {SAMPLE_GEN:,}")


if __name__ == "__main__":
    print(describe())
```
