## Supplementary Information section 3 code base for "Hypothesis: A modern human range expansion ∼300,000 years ago had a critical role in Neandertal origins": clock_check.py

```
#!/usr/bin/env python3
"""
clock_check.py -- are the two "expansion complete" clocks the same thing? (13 Sep 2026)

THE QUESTION. The note's admissibility criterion is that the expansion completes in 100-1,000
generations, tested against `end_gen`, which in both frameworks is when the last ARCHAIC-LABELLED
unit disappears. That is a label, not an allele frequency. For a single wave the distinction is
academic, because the archaic label dies out when the wave finishes. For a TWO-WAVE scenario it
is not, because at wave2_gen the whole landscape is relabelled archaic and only the new founder
group is modern -- so the label clock restarts on a population that is already genetically
modern, and it goes on ticking long after the genetics are settled.

So every two-wave cell needs both clocks measured before the criterion can be trusted:

    label   clock: the generation the last archaic-labelled deme/band disappears  (= end_gen)
    genetic clock: the generation whole-landscape mtDNA AND Y both reach 99% modern

A TRAP THAT HAS TO BE AVOIDED IN THE DEME ENGINE. spatial_sim2's trajectory fields mt_mod,
Y_mod and auto_mod are computed over lin == 1 only. After the wave-2 relabelling lin == 1 is
the founder deme alone, so Y_mod reads 1.000 from the very generation the second wave appears
while the landscape is at 0.001. Timing anything off mt_mod/Y_mod gives the wave-2 generation
back, plus the trajectory recording interval, and looks like a startlingly fast expansion. The
mt_all / Y_all / auto_all fields were added for this and are what this script reads. The
end-of-run SUMMARY was never affected: it is taken at sample_gen, thousands of generations
after the label absorbs, when lin == 1 is everyone again.

THE ANSWER, and the reason the two frameworks need different treatment:

    deme, gamma 0.42, wave2 500, 40 replicates
        label   615 (range 125-648)
        genetic 620 (range 590-650)        median paired difference +2 generations
        both clocks put 40 of 40 inside the window

    band, p_move 0.75, sigma_b 120, wave2 300, 3 replicates
        label   2,069, 5,100, and one that never completed in 6,000 generations
        genetic 1,038, 1,114, 1,148 -- including the replicate whose label never swept,
                which nevertheless ended at 100% modern mtDNA and 100% modern Y

In the deme model migration moves individuals between demes every generation, so labels and
alleles spread together and the two clocks coincide. In the band model identity is a band-level
attribute that changes only when a band is replaced, while alleles flow through exogamous
dispersal, so the two decouple completely. The criterion was therefore measuring something
reasonable in one framework and something meaningless in the other.

CONSEQUENCE FOR THE NOTE. Adopting the genetic clock changes no deme result -- which is the
point of running the deme arm here, since a metric change that moved the passing numbers would
need far more justification. On the band side it is the difference between a route that fails
and a route that works.

    python clock_check.py [--reps 40]
"""
import os
import sys

import numpy as np

HERE = os.path.dirname(os.path.abspath(__file__))
sys.path.insert(0, HERE)
sys.path.insert(0, os.path.join(os.path.dirname(HERE), "band_model"))

DEME_BASE = dict(grid=12, K=11, m=0.08, r=2.0, gens=2500, sample_gen=2490, record_every=10)


def flag(name, cast, default):
    return cast(sys.argv[sys.argv.index(name) + 1]) if name in sys.argv else default


def deme(gamma=0.42, wave2=500, reps=40):
    import spatial_sim2 as S2
    from spatial_sim2 import Config
    lab, gen = [], []
    for seed in range(reps):
        ts, ext = S2.run(Config(**dict(DEME_BASE, gamma=gamma, wave2_gen=wave2, seed=seed)))
        g, mt, y = ts["gen"], ts["mt_all"], ts["Y_all"]
        ok = np.where((mt >= 0.99) & (y >= 0.99) & (g > wave2))[0]
        gen.append(int(g[ok[0]]) if len(ok) else -1)
        lab.append(-1 if ext is None else int(ext))
    return np.array(lab), np.array(gen)


def report(name, lab, gen, lo=100, hi=1000):
    def s(a):
        v = a[a >= 0]
        if not len(v):
            return "never completed in any replicate"
        return (f"n={len(v)}/{len(a)}  median={np.median(v):.0f}  "
                f"range {v.min():.0f}-{v.max():.0f}  "
                f"inside {lo}-{hi}: {int(((v >= lo) & (v <= hi)).sum())}")
    print(f"{name}")
    print(f"  label   clock (end_gen)      {s(lab)}")
    print(f"  genetic clock (mt & Y >=99%) {s(gen)}")
    both = (lab >= 0) & (gen >= 0)
    if both.any():
        print(f"  paired: genetic later in {int((gen[both] > lab[both]).sum())}/{int(both.sum())}, "
              f"median difference {np.median(gen[both] - lab[both]):+.0f} generations")


def main():
    reps = flag("--reps", int, 40)
    lab, gen = deme(reps=reps)
    report(f"deme, gamma 0.42, wave2 500, {reps} replicates", lab, gen)
    print("\nband: see out_tw6_band, which records both clocks per cell "
          "(dur_med = label, arr_med = genetic)")


if __name__ == "__main__":
    main()
```
