## Supplementary Information section 3 code base for "Hypothesis: A modern human range expansion ∼300,000 years ago had a critical role in Neandertal origins": cultural_submit.py

```
#!/usr/bin/env python3
"""
cultural_submit.py -- the five cultural mechanisms on the Cartesian grid, in BOTH frameworks,
recording expansion duration. (12 Sep 2026)

WHY THIS RUN EXISTS. The audit of v66 turned up one claim with no data behind it at all:

    "Across all, the highest probability of retaining the unilineal marker not driving the
     expansion while holding autosomal ancestry at around 5% was 11% in the deme framework
     and 18% in the band framework"

There is no band framework number. kin2, kin3, mech3, mech4, mech5, clan and asym are every
one of them deme-only scripts. The 18% has no source on disk, and the deme figure recomputes
to 13.2% rather than 11%.

Those scripts also never recorded the duration of the expansion, which is why the five cultural
panels of Figure S3.3 are empty -- a point needs both coordinates to be placed on an (autosomal
ancestry, generations) plane -- and why those mechanisms have only ever been screened on one of
the note's three acceptance criteria instead of all three.

Routing them through robust_run fixes all of it at once: stage 1 records end_gen and
establishment, stage 2 records p_joint, and both engines already carry the parameters. They
then flow into the fountains, the thumbnails and the workbook with no special-casing anywhere,
because they are now just mechanisms like sY and theta.

ALL FIVE NOW RUN IN BOTH FRAMEWORKS -- CHANGED 13 Sep. Obligate dispersal used to be emitted
for the deme framework only, on the argument that it is sex-specific MIGRATION and bandsim's
only sex-specific rate is beta, which IS the residence rule and is the thing being held fixed.
That argument is right about beta on its own and wrong about the pair (e, beta). bandsim sets
e_f = 2 e beta and e_m = 2 e (1 - beta), so pinning the male rate to the ladder value and the
female rate to the same 0.005 the deme arm uses determines both outright:

    e = (e_m + e_f) / 2        beta = e_f / (e_m + e_f)

That moves the TOTAL exogamy rate as well as its sex bias, which varying beta at fixed e
cannot do, so it is the deme mechanism rather than the residence axis under another name.
Because e and beta are the mechanism here they are no longer swept for this one mechanism; see
demographies(). The difference that remains is one of baseline, not of capability, and the
note should say it: the band model is already male-exogamous at beta = 0, so its arm of this
panel asks at what RATE obligate dispersal helps, where the deme arm asks whether making
dispersal obligate helps at all.

    python cultural_submit.py 1     # screen,  10 reps      ~95 CPU-hours
    python cultural_submit.py 2     # score,  200 reps      depends on how many pass
    python cultural_submit.py status
"""
import glob
import itertools
import json
import os
import subprocess
import sys

HERE = os.path.dirname(os.path.abspath(__file__))
sys.path.insert(0, HERE)

from ne_screen import GRIDS as DEME_GRIDS, KS as DEME_KS, MS as DEME_MS   # noqa: E402

# The value ladders are the ones the original scripts swept, so a number here can be checked
# against the number the note already quotes.
MECHS = {
    "inbreed":   ([1, 2],                       True),   # ban sharing a mother / a grandmother
    # 13 Sep: NOW IN BOTH FRAMEWORKS. See the note in robust_run.CULTURAL -- bandsim has no
    # m_male/m_female, but (e, beta) fix the two per-sex exogamy rates exactly, so the deme
    # ladder is reproducible by inverting e_f = 2 e beta, e_m = 2 e (1 - beta).
    "obligate":  ([0.05, 0.10, 0.20, 0.40],     True),   # dispersing sex's migration rate
    "rhoreturn": ([0.25, 0.50, 0.75, 1.00],     True),   # P(son returns to father's band)
    "skew":      ([0.20, 0.40, 0.60, 0.80],     True),   # fraction of men who father children
    "asym":      ([0.04, 0.08, 0.16, 0.32],     True),   # export discount / export exogamy rate
}

DEME_GAMMA = [0.08, 0.12, 0.20, 0.30, 0.42, 0.55, 0.70, 1.00]
DEME_R = [1.5, 2.0, 3.0]
BAND_GRID = dict(e=[0.08, 0.13, 0.19, 0.26, 0.35, 0.45, 0.56],
                 sigma_b=[30.0, 60.0, 120.0], sigma_e=[40.0, 80.0, 160.0],
                 alpha_d=[0.4, 0.8, 1.6], r=[1.5, 2.0, 3.0],
                 p_move=[0.45], K=[25], N=[1750])

AUT_LO, AUT_HI = 2.0, 15.0          # the STAGE-1 screen, deliberately wide; 3-7% is applied later
GEN_LO, GEN_HI = 100.0, 1000.0
EST_MIN = 0.50
PART, CAP = "short", 900
OUT = "cult"

FLAG = {"grid": "--grid", "K": "--K", "m": "--m", "gamma": "--gamma", "r": "--r",
        "N": "--N", "e": "--e", "p_move": "--p-move", "sigma_b": "--sigma-b",
        "sigma_e": "--sigma-e", "alpha_d": "--alpha", "female_bias": "--female-bias",
        "beta": "--beta"}

_NE = {}


def ne_ok(grid, K, m):
    """Same admissibility manifold as routes_build: measured Ne within 10% of 1,447."""
    if not _NE:
        import csv
        with open(os.path.join(HERE, "ne_screen.csv")) as f:
            for row in csv.DictReader(f):
                _NE[(int(row["grid"]), int(row["K"]), float(row["m"]))] = int(row["keep"])
    return _NE.get((grid, K, m))


def demographies(band, mech=None):
    if band:
        grid = dict(BAND_GRID)
        # e AND beta ARE THE MECHANISM FOR obligate, so neither is swept and neither is passed
        # on the command line. The ladder pins the male exogamy rate to its value and the
        # female rate to 0.005, which determines the pair outright. Leaving e on its
        # seven-value ladder would emit seven cells per demography that robust_run then
        # overwrites with identical parameters: seven copies of one run, at seven times the
        # cost, under filenames that claim to differ and do not. Dropping beta too keeps the
        # filename honest -- a cell tagged beta0 that ran at beta = 0.024 is the kind of
        # mislabelling that takes a day to find later.
        fix = mech == "obligate"
        if fix:
            grid.pop("e")
        keys = list(grid)
        for combo in itertools.product(*(grid[k] for k in keys)):
            kw = dict(zip(keys, combo))
            if not fix:
                kw["beta"] = 0.0
            yield kw
    else:
        for g, k, m, ga, r in itertools.product(DEME_GRIDS, DEME_KS, DEME_MS,
                                                DEME_GAMMA, DEME_R):
            if ne_ok(g, k, m):
                yield dict(grid=g, K=k, m=m, gamma=ga, r=r, female_bias=1.0)


def tag(fw, mech, v, kw):
    s = f"{fw}_{mech}_{v:g}_" + "_".join(f"{k}{kw[k]:g}" for k in sorted(kw))
    return s.replace(".", "p").replace("-", "m")


def stage1_pass(d):
    n = d.get("n") or 1
    return (d.get("aut") is not None and AUT_LO <= d["aut"] <= AUT_HI
            and d.get("end_gen") is not None and GEN_LO <= d["end_gen"] <= GEN_HI
            and (d.get("n_est") or 0) / n > EST_MIN)


def build(stage, reps):
    tasks, per = [], {}
    for mech, (vals, in_band) in MECHS.items():
        for fw, band in (("deme", False), ("band", True)):
            if band and not in_band:
                continue
            outdir = f"out_{OUT}{stage}_{fw}"
            os.makedirs(os.path.join(HERE, outdir), exist_ok=True)
            n = 0
            for v in vals:
                for kw in demographies(band, mech):
                    name = tag(fw, mech, v, kw) + ".json"
                    if os.path.exists(os.path.join(HERE, outdir, name)):
                        continue
                    if stage == 2:
                        s1 = os.path.join(HERE, f"out_{OUT}1_{fw}", name)
                        if not os.path.exists(s1):
                            continue
                        try:
                            if not stage1_pass(json.load(open(s1))["summary"]):
                                continue
                        except Exception:
                            continue
                    fl = []
                    for k in sorted(kw):
                        fl += [FLAG[k], f"{kw[k]:g}"]
                    tasks.append(f"python robust_run.py {fw} {mech} {v:g} {stage} {reps} 0 "
                                 f"{os.path.join(outdir, name)} " + " ".join(fl))
                    n += 1
            per[(mech, fw)] = n
    return tasks, per


def emit(name, tasks, hours, batch=1):
    """BATCHED, because the grid is 21,330 cells and the submit limit is 10,000 arrays.

    The obvious response to hitting AssocMaxSubmitJobLimit is to narrow a ladder, and it is the
    wrong one here: the whole run is only 83 CPU-hours, so the limit being hit is a statement
    about task COUNT, not about cost. Cutting the grid would throw away coverage to satisfy a
    bookkeeping constraint. Instead each array task runs `batch` cells in sequence, which costs
    nothing and brings 21,330 cells down to about 2,700 tasks. Each cell still writes its own
    JSON and is still skipped by filename, so a task that times out loses at most the cell it
    was in the middle of.
    """
    if not tasks:
        print("  nothing to submit")
        return
    groups = [tasks[i:i + batch] for i in range(0, len(tasks), batch)]
    pf, sf = f"{name}_params.txt", f"{name}_array.sh"
    # one GROUP per line, cells joined by ';' so a failure does not abort the rest
    open(os.path.join(HERE, pf), "w").write(
        "\n".join(" ; ".join(g) for g in groups) + "\n")
    tasks = groups
    open(os.path.join(HERE, sf), "w").write(f"""#!/bin/bash
#SBATCH -J {name}
#SBATCH -p {PART}
#SBATCH -c 1
#SBATCH --mem=2G
#SBATCH -t {hours}
#SBATCH -a 0-{len(groups) - 1}%{CAP}
#SBATCH -o logs/{name}_%A_%a.log
cd {HERE}
CMD=$(sed -n "$((SLURM_ARRAY_TASK_ID + 1))p" {pf})
echo "$CMD"
eval "$CMD"
""")
    print(f"  {len(groups):,} array tasks of up to {batch} cells each -> {sf}")
    if "--no-go" in sys.argv:
        print(f"  NOT submitted (--no-go).  sbatch {sf}")
        return
    r = subprocess.run(["sbatch", sf], cwd=HERE, capture_output=True, text=True)
    print(f"  SUBMITTED: {r.stdout.strip()}" if r.returncode == 0
          else f"  !! sbatch FAILED: {(r.stderr or '').strip()}")


# measured 12 Sep: seconds per replicate, deme and band, stage 1 and stage 2
COST = {("deme", 1): 0.57, ("deme", 2): 0.60, ("band", 1): 2.80, ("band", 2): 2.37}


def main():
    what = sys.argv[1] if len(sys.argv) > 1 else "status"
    os.makedirs(os.path.join(HERE, "logs"), exist_ok=True)

    if what == "status":
        for stage in (1, 2):
            for fw in ("deme", "band"):
                d = os.path.join(HERE, f"out_{OUT}{stage}_{fw}")
                print(f"  out_{OUT}{stage}_{fw:<5} "
                      f"{len(glob.glob(os.path.join(d, '*.json'))):,}")
        return

    stage = int(what)
    reps = 10 if stage == 1 else 200
    tasks, per = build(stage, reps)
    print(f"{'mechanism':<12}{'framework':<11}{'cells':>8}{'CPU-h':>8}")
    tot = 0.0
    for (mech, fw), n in sorted(per.items()):
        h = n * COST[(fw, stage)] * reps / 3600
        tot += h
        print(f"{mech:<12}{fw:<11}{n:>8,}{h:>8.1f}")
    print(f"{'TOTAL':<23}{len(tasks):>8,}{tot:>8.1f}")
    # size the batch so the array stays under the submit limit with room to spare
    batch = 1
    while len(tasks) / batch > 3000:
        batch += 1
    print(f"  batching {batch} cells per array task")
    emit(f"{OUT}{stage}", tasks, "0-02:59" if stage == 1 else "0-11:59", batch=batch)


if __name__ == "__main__":
    main()
```
