## Supplementary Information section 3 code base for "Hypothesis: A modern human range expansion ∼300,000 years ago had a critical role in Neandertal origins": demog_fill_submit.py

```
#!/usr/bin/env python3
"""
demog_fill_submit.py -- fill the unbiased-expansion sweep at the MAIN sY ladder. (14 Sep 2026)

WHY. routes_build's `demog` set doubled above s = 0.02 -- 0.04, 0.08, 0.16, 0.32, 0.64 --
while the `main` set the red fountains come from uses 0.05, 0.10, 0.20, 0.35, 0.50. The two
sweeps therefore have no shared rung above 0.02, and any sentence comparing the matrilocal and
unbiased expansions at a given selection strength has to hedge. DR: "I don't like
interpolation and having to say something sloppy in the final text."

IT IS NOT, STRICTLY, INTERPOLATION -- and the note should not call it that. Nothing estimates
a value between two rungs. The problem is narrower and harder to write around: the two clouds
are measured at different s, so the only honest comparison is "at roughly comparable selection
strengths". Aligning the ladders removes the hedge rather than improving an estimate.

WHAT THIS SUBMITS, AND WHY NOT routes_submit DIRECTLY. With the ladder changed,
`ROUTE_SET=demog python routes_submit.py 1` enumerates the five new rungs across the WHOLE
demog grid -- both residence rules crossed with the five-rung mtDNA-selection ladder, roughly
ten times what is needed. The unbiased series is one slice of that:

    residence   beta = 0.5 (band) / female_bias = 0.5 (deme)
    mtDNA gap   s_mt = 0.00235, the posterior median the note quotes

which is the same slice nexp_band_submit scored for the existing rungs, so the new cells are
directly poolable with the 3,920 already on disk. Everything outside the slice is left alone.

    python demog_fill_submit.py 1            # screen,  deme
    python demog_fill_submit.py 1 --band     # screen,  band
    python demog_fill_submit.py 2            # score,   deme
    python demog_fill_submit.py 2 --band     # score,   band
    ... --no-go to enumerate and price without submitting
"""
import math
import os
import re
import subprocess
import sys

HERE = os.path.dirname(os.path.abspath(__file__))

S_MT = "0.00235"
# The residence flag differs by engine: beta is the band model's FEMALE share of exogamy, so
# 0.5 is unbiased; female_bias is the deme model's, and 0.5 is unbiased there too. Opposite
# ends of differently-oriented scales at their extremes, but they agree at the midpoint.
UNBIASED = {"band": "--beta 0.5", "deme": "--female-bias 0.5"}
# measured seconds per cell, from the arrays already run at these settings
SEC = {("deme", 1): 150.0, ("band", 1): 110.0, ("deme", 2): 200.0, ("band", 2): 3400.0}
CHUNK = {1: 12, 2: 3}
PART, MEM, TIME = "short", "3G", "0-11:59"


def newest(pat):
    fs = [f for f in os.listdir(HERE) if re.fullmatch(pat, f)]
    return max(fs, key=lambda f: os.path.getmtime(os.path.join(HERE, f))) if fs else None


def main():
    stage = int(sys.argv[1]) if len(sys.argv) > 1 and sys.argv[1] in ("1", "2") else 1
    band = "--band" in sys.argv
    fw = "band" if band else "deme"
    stem = f"dfill{stage}{fw[0]}"

    env = dict(os.environ, ROUTE_SET="demog")
    cmd = [sys.executable, "routes_submit.py", str(stage)] + (["--band"] if band else []) \
        + ["--no-go"]
    r = subprocess.run(cmd, cwd=HERE, env=env, capture_output=True, text=True)
    if r.returncode != 0:
        raise SystemExit(f"routes_submit failed:\n{r.stderr or r.stdout}")
    src = newest(rf"demog{stage}{fw[0]}_\d+_params\.txt")
    if src is None:
        print(f"  nothing pending for stage {stage}, {fw}")
        return

    cells = [ln.strip() for ln in open(os.path.join(HERE, src)) if ln.strip()]
    want = [c for c in cells if f"--s-mt {S_MT}" in c and UNBIASED[fw] in c]
    if not want:
        print(f"  {len(cells):,} cells pending, none in the unbiased / s_mt = {S_MT} slice "
              f"-- nothing to do for the orange sweep")
        return
    ch = CHUNK[stage]
    ntask = math.ceil(len(want) / ch)
    print(f"  stage {stage}, {fw}: {len(cells):,} demog cells pending in total")
    print(f"  {len(want):,} in the unbiased slice ({UNBIASED[fw]}, s_mt {S_MT})")
    print(f"  {ntask:,} array tasks of {ch}, "
          f"~{len(want) * SEC[(fw, stage)] / 3600:,.0f} CPU-hours")

    pf, sf = f"{stem}_params.txt", f"{stem}_array.sh"
    open(os.path.join(HERE, pf), "w").write("\n".join(want) + "\n")
    open(os.path.join(HERE, sf), "w").write(f"""#!/bin/bash
#SBATCH -J {stem}
#SBATCH -p {PART}
#SBATCH -t {TIME}
#SBATCH -c 1
#SBATCH --mem={MEM}
#SBATCH -a 0-{ntask - 1}
#SBATCH -o logs/{stem}_%A_%a.log
CHUNK={ch}
START=$((SLURM_ARRAY_TASK_ID * CHUNK + 1))
END=$((START + CHUNK - 1))
sed -n "${{START}},${{END}}p" {pf} | while read -r LINE; do
  [ -z "$LINE" ] && continue
  python robust_run.py $LINE
done
""")
    os.makedirs(os.path.join(HERE, "logs"), exist_ok=True)
    if "--no-go" in sys.argv:
        print(f"  NOT submitted (--no-go).  sbatch {sf}")
        return
    s = subprocess.run(["sbatch", sf], cwd=HERE, capture_output=True, text=True)
    print(f"  SUBMITTED: {s.stdout.strip()}" if s.returncode == 0
          else f"  !! sbatch FAILED: {(s.stderr or '').strip()}")


if __name__ == "__main__":
    main()
```
