## Supplementary Information section 3 code base for "Hypothesis: A modern human range expansion ∼300,000 years ago had a critical role in Neandertal origins": esc_collect.py

```
#!/usr/bin/env python3
"""
esc_collect.py -- pool out_esc/ and report the heritable-escape sweep for mtDNA x Y.

eps = 0 must reproduce the Table S2.1 mtDNA x Y row (modern Y 100%). The report is the
critical eps at which the sealed architecture starts to fail, with the escaped compatible
archaic Y tracked separately from the never-blocked one.

  python esc_collect.py        # -> esc_summary.csv, esc_panel.png
"""
import glob
import numpy as np
import matplotlib
matplotlib.use("Agg")
import matplotlib.pyplot as plt
from esc_run import EPSVALS

COL = dict(seed=0, mt=1, Y=2, Yesc=3, auto=4, X=5, ext=6)


def pool(eps):
    tag = f"e{eps:g}".replace(".", "p").replace("-", "m")
    rows = []
    for f in glob.glob(f"out_esc/out_{tag}_s*.txt"):
        for line in open(f).read().strip().split("\n")[1:]:
            p = line.split("\t")
            if len(p) >= 7 and "nan" not in line:
                rows.append([float(x) for x in p])
    return np.array(rows) if rows else None


def main():
    recs = []
    with open("esc_summary.csv", "w") as out:
        out.write("eps,nreps,mt,Y_modern,Y_escaped,auto,X,P_Y_fixed,P_all10,P_ext\n")
        print(f"{'eps':>8s} {'reps':>5s} | {'Y%':>6s} {'Yesc%':>6s} {'auto%':>6s} "
              f"{'X%':>6s} | {'P(Yfix)':>8s} {'P(all10)':>8s}")
        for eps in EPSVALS:
            a = pool(eps)
            if a is None:
                continue
            comp = 100 * (a[:, COL["ext"]] >= 0).mean()
            a = a[a[:, COL["ext"]] >= 0]               # conditional on completion (Table S2.1)
            if not len(a):
                continue
            Y = a[:, COL["Y"]]
            v = dict(eps=eps, reps=len(a), mt=100 * a[:, COL["mt"]].mean(),
                     Y=100 * Y.mean(), yesc=100 * a[:, COL["Yesc"]].mean(),
                     auto=100 * a[:, COL["auto"]].mean(), X=100 * a[:, COL["X"]].mean(),
                     fix=100 * (Y >= 0.999).mean(), p10=100 * (Y ** 10).mean(),
                     ext=comp)
            recs.append(v)
            out.write(f"{eps:g},{len(a)},{v['mt']:.2f},{v['Y']:.2f},{v['yesc']:.2f},"
                      f"{v['auto']:.2f},{v['X']:.2f},{v['fix']:.1f},{v['p10']:.1f},"
                      f"{v['ext']:.1f}\n")
            print(f"{eps:8g} {len(a):5d} | {v['Y']:6.1f} {v['yesc']:6.1f} {v['auto']:6.2f} "
                  f"{v['X']:6.2f} | {v['fix']:8.1f} {v['p10']:8.1f}")
    if not recs:
        print("no data yet"); return
    print("\nanchor: eps=0 is the Table S2.1 mtDNA x Y row (Y=100%)")

    fig, ax = plt.subplots(figsize=(6.4, 4.2))
    e = [max(r["eps"], 3e-5) for r in recs]     # plot eps=0 at the left edge of the log axis
    ax.plot(e, [r["p10"] for r in recs], "-o", color="#C44E52",
            label="P(all 10 sampled Y's modern)")
    ax.plot(e, [r["Y"] for r in recs], "--s", color="#4C72B0", label="mean modern Y")
    ax.plot(e, [r["yesc"] for r in recs], ":^", color="0.4", label="escaped compatible archaic Y")
    ax.set_xscale("log")
    ax.set_xlabel("heritable escape probability per blocked offspring, eps")
    ax.set_ylabel("percent"); ax.set_ylim(-3, 103)
    ax.grid(alpha=.3); ax.legend(fontsize=9, frameon=False)
    ax.set_title("How robust is the sealed mtDNA x Y DMI to heritable escape?", fontsize=11)
    fig.tight_layout()
    fig.savefig("esc_panel.png", dpi=160)
    print("saved esc_summary.csv and esc_panel.png")


if __name__ == "__main__":
    main()
```
