## Supplementary Information section 3 code base for "Hypothesis: A modern human range expansion ∼300,000 years ago had a critical role in Neandertal origins": esc_run.py

```
#!/usr/bin/env python3
"""
esc_run.py -- heritable-escape stress test of the sealed mtDNA x Y DMI, on spatial_sim3.py.

THE IDEA. The mtDNA x Y row of Table S2.1 is the unique sealed architecture, but its seal
assumes the archaic Y cannot BECOME compatible. Real escapes are heritable: a compensatory
mutation, or an archaic Y that acquires the modern pseudoautosomal region through one rare
successful pairing, founds a lineage that is immune forever after -- far more dangerous than
the i.i.d. escapes of the theta grids (where theta = 0.01 still retained the Y in 84% of
replicates). Here, each time the DMI would remove an offspring, with probability eps the
offspring instead survives with its Y converted to a heritably compatible allele, counted as
ARCHAIC in the output. The question is the critical eps below which the mechanism still holds
at the sampling generation -- a robustness number every sealed-DMI claim in the note should be
read against.

Baseline: grid=20, K=20, r=2, m=0.10, gamma=0.12, matrilineal, dm=1 mtDNA x Y male-limited,
sampled at generation 6,993, no early stopping. eps = 0 reproduces the Table S2.1 mtDNA x Y row.

Row layout:  seed  mt  Y  Yesc  auto  X  extgen   (Y = modern fraction; Yesc = escaped-archaic)
Usage:       python esc_run.py <eps> <reps> <seed0> <outfile>
"""
import os
import sys
import numpy as np
from dataclasses import replace
import spatial_sim2 as S2
from spatial_sim3 import Config, _repro

SAMPLE_GEN = 6993
EPSVALS = (0.0, 1e-4, 3e-4, 1e-3, 3e-3, 1e-2, 3e-2, 1e-1)

# --- expansion geometry, settable from the command line ---------------------------------
# The grid-20 / K-20 baseline has a post-wave census of ~8000, five times the post-mixture size
# the rest of the analysis assumes (Muller's Ratchet v32: NN-uniparental = 725-1000, diploid
# 1500-2000). Since the modern mtDNA's advantage is a ~0.3% coefficient whose potency scales
# with 2*Ne*s, that mismatch inflated every spatial result. r and m are NOT externally
# constrained (DR, 3 Sep 2026: "pulled from my head"), so they are exposed here too and their
# sensitivity is the honest thing to report rather than a fixed baseline.
#
# A flag only OVERRIDES a parameter this script holds fixed; where the script sweeps that
# parameter, the swept value still wins, so sensitivity panels keep their full range.
# K_early/K_early_gens are always ADDED, since no BASE dict carries them.
def _geom(kw):
    import sys as _s
    for flag, key, cast in (("--grid", "grid", int), ("--K", "K", int),
                            ("--m", "m", float), ("--r", "r", float)):
        if flag in _s.argv and key in kw:
            kw[key] = cast(_s.argv[_s.argv.index(flag) + 1])
    for flag, key in (("--K-early", "K_early"), ("--K-early-gens", "K_early_gens")):
        if flag in _s.argv:
            kw[key] = int(_s.argv[_s.argv.index(flag) + 1])
    if kw.get("K", 20) < 7:
        raise SystemExit("K < 7: deme reproductive failure collapses the archaic population")
    return kw


BASE = dict(grid=20, K=20, r=2.0, m=0.10, gamma=0.12, lineage="matrilineal",
            s=0.0, sel_locus="none", dm=1.0, dm_focal="mt", dm_partner="Y",
            dm_male_limited=True,
            gens=SAMPLE_GEN + 10, sample_gen=SAMPLE_GEN, stop_after_ext=0,
            record_every=100000)


def conditions():
    return list(EPSVALS)


def run_one(cfg):
    rng = np.random.default_rng(cfg.seed)
    _cap_early = S2.cfg_early(cfg)   # K_early phase, if any
    P = S2._init(_cap_early, rng)
    ext = None
    for g in range(cfg.gens):
        _c = S2.cap_at(cfg, _cap_early, g)
        S2._migrate(P, _c, rng)
        nxt = _repro(P, _c, rng)
        if nxt is None:
            break
        P = S2._reseed_origin(nxt, _c)
        if ext is None and (P["lin"] == 0).sum() == 0:
            ext = g
        if g == cfg.sample_gen:
            break
    st = S2._stats(P, exclude=(), exclude_x=())
    mmale = (P["lin"] == 1) & (P["sex"] == 1)
    yesc = float((P["Y"][mmale] == 2).mean()) if mmale.any() else np.nan
    return st, yesc, ext


def main():
    eps = float(sys.argv[1])
    reps, seed0, outfile = int(sys.argv[2]), int(sys.argv[3]), sys.argv[4]
    cfg0 = Config(**_geom(dict(BASE)), esc_eps=eps)
    os.makedirs(os.path.dirname(outfile) or ".", exist_ok=True)
    with open(outfile, "w") as f:
        f.write("seed\tmt\tY\tYesc\tauto\tX\textgen\n")
        for s in range(seed0, seed0 + reps):
            st, yesc, ext = run_one(replace(cfg0, seed=s))
            eg = int(ext) if ext is not None else -1
            f.write(f"{s}\t{st['mt_mod']:.6g}\t{st['Y_mod']:.6g}\t{yesc:.6g}\t"
                    f"{st['auto_mod']:.6g}\t{st['X_mod']:.6g}\t{eg}\n")
    print("wrote", outfile)


if __name__ == "__main__":
    main()
```
