## Supplementary Information section 3 code base for "Hypothesis: A modern human range expansion ∼300,000 years ago had a critical role in Neandertal origins": esc_submit.py

```
#!/usr/bin/env python3
"""esc_submit.py -- SLURM array for the heritable-escape sweep (esc_run.py).
Usage:  python esc_submit.py [reps] [batch]     # defaults 1000, 25
then:   sbatch esc_array.sh
"""
import os
import sys
from esc_run import conditions


GEOM_FLAGS = ("--grid", "--K", "--m", "--r", "--K-early", "--K-early-gens")


def _strip_flags(av):
    """Positional parsing must not be shifted by the geometry flags, so drop each flag AND its
    value before the positional args are read."""
    out, skip = [], False
    for a in av:
        if skip:
            skip = False; continue
        if a.startswith("--"):
            skip = a in GEOM_FLAGS or a == "--rna"; continue
        out.append(a)
    return out


def _geom_str():
    """Echo the geometry flags onto every task line. Without this the workers would silently run
    the grid-20 / K-20 default while the submit script had been told otherwise -- which produces
    plausible-looking numbers that are simply wrong."""
    import sys as _s
    return "".join(f" {f} {_s.argv[_s.argv.index(f) + 1]}"
                   for f in GEOM_FLAGS if f in _s.argv)


def main():
    GEOM = _geom_str()
    nums = [int(a) for a in sys.argv[1:] if a.isdigit()]
    total = nums[0] if len(nums) >= 1 else 1000
    batch = nums[1] if len(nums) >= 2 else 25
    os.makedirs("out_esc", exist_ok=True); os.makedirs("logs", exist_ok=True)
    tasks = []
    for eps in conditions():
        tag = f"e{eps:g}".replace(".", "p").replace("-", "m")
        for s0 in range(0, total, batch):
            reps = min(batch, total - s0)
            tasks.append(f"{eps} {reps} {s0} out_esc/out_{tag}_s{s0}.txt{GEOM}")
    open("esc_params.txt", "w").write("\n".join(tasks) + "\n")
    n = len(tasks)     # no throttle (standing instruction)
    open("esc_array.sh", "w").write(f"""#!/bin/bash
#SBATCH -J esc
#SBATCH -p short
#SBATCH -t 0-11:59
#SBATCH -c 1
#SBATCH --mem=4G
#SBATCH -o logs/esc_%A_%a.log
#SBATCH --array=0-{n - 1}

LINE=$(sed -n "$((SLURM_ARRAY_TASK_ID + 1))p" esc_params.txt)
echo "task $SLURM_ARRAY_TASK_ID : $LINE"
python esc_run.py $LINE
""")
    print(f"wrote esc_params.txt ({n} tasks over {len(conditions())} conditions, "
          f"{total} reps each) and esc_array.sh")
    print("submit:  sbatch esc_array.sh")


if __name__ == "__main__":
    main()
```
