## Supplementary Information section 3 code base for "Hypothesis: A modern human range expansion ∼300,000 years ago had a critical role in Neandertal origins": grid5_collect.py

```
#!/usr/bin/env python3
"""
grid5_collect.py -- pool out_grid5/ and map the parameter space that reproduces the empirical
observation.

    SUCCESS = 2% <= modern autosomal ancestry <= 15%  AND  modern Y fixed (100%)

Produces a panel of heatmaps, one per migration rate m, with gamma on the x-axis and s on
the y-axis, coloured by the fraction of replicates meeting the joint criterion. Also writes a
long-format CSV with the component criteria, so the two constraints can be inspected separately.

    python grid5_collect.py     # -> grid5_map.png, grid5_summary.csv
"""
import os
import glob
import sys
import numpy as np
import matplotlib
matplotlib.use("Agg")
import matplotlib.pyplot as plt
from matplotlib.colors import LinearSegmentedColormap
from grid5_run import BIASES, GAMMAS, MS

COL = dict(seed=0, mt=1, Y=2, auto=3, X=4, ext=5)

# ---- sample-based criterion -------------------------------------------------
# Population-wide fixation is far stricter than the data require. What is actually observed is
# a handful of Neandertals all carrying the modern-derived type (Petr et al. report three Y
# chromosomes; the mtDNA sample is a few dozen). If the modern type sits at frequency p in the
# final population, the chance that NSAMPLE individuals drawn from it ALL carry it is p**NSAMPLE
# (sampling with replacement; with ~8,000 individuals the finite-population correction is
# negligible). Averaging that over replicates gives the probability of reproducing the
# OBSERVATION rather than of achieving literal fixation. Override with:
#     NSAMPLE=3 python <collector>.py
NSAMPLE = int(os.environ.get("NSAMPLE", "10"))

# the autosomal acceptance window, as fractions. Override on the command line:
#     python grid5_collect.py 3 10      -> 3-10%
AUTO_LO, AUTO_HI = 0.02, 0.15
if len(sys.argv) >= 3:
    AUTO_LO, AUTO_HI = float(sys.argv[1]) / 100.0, float(sys.argv[2]) / 100.0
TAG = f"_{100*AUTO_LO:g}-{100*AUTO_HI:g}pct"   # keeps each window's outputs separate


def tag(th, ga, m):
    return f"t{th:g}_g{ga:g}_m{m:g}".replace(".", "p")


def pool(th, ga, m):
    rows = []
    for f in glob.glob(f"out_grid5/out_{tag(th, ga, m)}_s*.txt"):
        for line in open(f).read().strip().split("\n")[1:]:
            p = line.split("\t")
            if len(p) >= 6:
                rows.append([float(x) for x in p])
    return np.array(rows) if rows else None


def main():
    shape = (len(MS), len(BIASES), len(GAMMAS))
    succ = np.full(shape, np.nan)
    nrep = np.zeros(shape, int)
    auto_ok = np.full(shape, np.nan)
    yfix = np.full(shape, np.nan)
    with open(f"grid5_summary{TAG}.csv", "w") as out:
        out.write(f"female_bias,gamma,m,n,auto_mean,mt_mean,Y_mean,P_auto_in_range,P_mt_fixed,P_mt_sample{NSAMPLE},P_Y_fixed,P_success,P_success_sample{NSAMPLE}\n")
        for k, m in enumerate(MS):
            for i, th in enumerate(BIASES):
                for j, ga in enumerate(GAMMAS):
                    a = pool(th, ga, m)
                    if a is None:
                        continue
                    au = a[:, COL["auto"]]; Y = a[:, COL["Y"]]; MT = a[:, COL["mt"]]
                    ok_a = (au >= AUTO_LO) & (au <= AUTO_HI)
                    ok_y = Y  >= 0.999
                    ok_m = MT >= 0.999          # this grid's headline outcome
                    p_samp = MT ** NSAMPLE      # P(all NSAMPLE sampled carry modern mtDNA)
                    s = float((ok_a & ok_m).mean())   # autosomes right AND modern mtDNA fixed
                    s_samp = float((ok_a * p_samp).mean())
                    succ[k, i, j] = s
                    auto_ok[k, i, j] = float(ok_a.mean())
                    yfix[k, i, j] = float(ok_m.mean())
                    nrep[k, i, j] = len(a)
                    out.write(f"{th:g},{ga:g},{m:g},{len(a)},{100*np.nanmean(au):.2f},"
                              f"{100*np.nanmean(MT):.2f},{100*np.nanmean(Y):.2f},"
                              f"{100*ok_a.mean():.1f},{100*ok_m.mean():.1f},"
                              f"{100*p_samp.mean():.1f},{100*ok_y.mean():.1f},"
                              f"{100*s:.1f},{100*s_samp:.1f}\n")

    # ---- wide matrices for Excel: rows = (s, gamma), one column per m ----
    def write_matrix(fname, arr, label):
        with open(fname, "w") as fh:
            fh.write("s,gamma," + ",".join(f"m={m:g}" for m in MS) + "\n")
            for i, th in enumerate(BIASES):
                for j, ga in enumerate(GAMMAS):
                    vals = []
                    for k in range(len(MS)):
                        v = arr[k, i, j]
                        vals.append("" if np.isnan(v) else f"{100*v:.1f}")
                    fh.write(f"{th:g},{ga:g}," + ",".join(vals) + "\n")
        print(f"saved {fname}  ({label}; {len(BIASES)*len(GAMMAS)} rows x {len(MS)} m-columns)")

    write_matrix(f"grid5_matrix_success{TAG}.csv", succ, "% matching BOTH criteria (autosomes in window AND modern mtDNA fixed)")
    write_matrix(f"grid5_matrix_autoOK{TAG}.csv", auto_ok, f"% with autosomes in {100*AUTO_LO:g}-{100*AUTO_HI:g}%")
    write_matrix(f"grid5_matrix_mtfixed{TAG}.csv", yfix, "% with the modern mtDNA fixed")

    cmap = LinearSegmentedColormap.from_list("succ", ["#f7f7f7", "#fee391", "#fe9929", "#cc4c02"])
    ncol = 3; nrow = int(np.ceil(len(MS) / ncol))
    fig, axes = plt.subplots(nrow, ncol, figsize=(4.1 * ncol, 3.5 * nrow))
    for k, m in enumerate(MS):
        ax = axes.ravel()[k]
        im = ax.imshow(succ[k], origin="lower", aspect="auto", cmap=cmap, vmin=0, vmax=1)
        ax.set_xticks(range(len(GAMMAS))); ax.set_xticklabels([f"{g:g}" for g in GAMMAS], fontsize=7)
        ax.set_yticks(range(len(BIASES))); ax.set_yticklabels([f"{t:g}" for t in BIASES], fontsize=7)
        ax.set_title(f"m = {m:g}", fontsize=10)
        if k % ncol == 0: ax.set_ylabel("s (female bias in lineage inheritance)", fontsize=9)
        if k // ncol == nrow - 1: ax.set_xlabel(r"$\gamma$ (interbreeding rate)", fontsize=9)
        for i in range(len(BIASES)):
            for j in range(len(GAMMAS)):
                v = succ[k, i, j]
                if not np.isnan(v) and v >= 0.05:
                    ax.text(j, i, f"{100*v:.0f}", ha="center", va="center", fontsize=5.5,
                            color="white" if v > 0.55 else "black")
    for k in range(len(MS), nrow * ncol):
        axes.ravel()[k].axis("off")
    cb = fig.colorbar(im, ax=axes.ravel().tolist(), shrink=0.6, pad=0.02)
    cb.set_label("fraction of replicates matching the data\n"
                 f"({100*AUTO_LO:g}-{100*AUTO_HI:g}% autosomal AND modern mtDNA fixed)", fontsize=9)
    fig.suptitle("Parameter space reproducing the Neandertal observation\n"
                 f"matrilineal range expansion, no Haldane effect, r = 2, 20x20 demes, K = 20, {nrep.max()} replicates each",
                 fontsize=11)
    fig.savefig(f"grid5_map{TAG}.png", dpi=160, bbox_inches="tight")
    print(f"saved grid5_map{TAG}.png and grid5_summary{TAG}.csv")

    # ---- coverage: partial grids are fine, but say how complete they are ----
    total_cells = succ.size
    empty = int((nrep == 0).sum())
    partial = int(((nrep > 0) & (nrep < nrep.max())).sum())
    print(f"\ncoverage: {total_cells - empty}/{total_cells} cells have data "
          f"({empty} empty, {partial} with fewer replicates than the fullest cell); "
          f"replicates per cell range {nrep[nrep > 0].min() if (nrep > 0).any() else 0}"
          f"-{nrep.max()}")
    if empty:
        print("  empty cells are blank in the figure and left empty in the matrices")

    if not np.isfinite(succ).any():
        print("no data yet"); return
    best = np.unravel_index(np.nanargmax(succ), succ.shape)
    print(f"best cell: m={MS[best[0]]:g}, s={BIASES[best[1]]:g}, gamma={GAMMAS[best[2]]:g} "
          f"-> {100*succ[best]:.0f}% of replicates match")
    n_ok = int(np.nansum(succ >= 0.5))
    print(f"combinations with >=50% of replicates matching: {n_ok} of {succ.size}")


if __name__ == "__main__":
    main()
```
