## Supplementary Information section 3 code base for "Hypothesis: A modern human range expansion ∼300,000 years ago had a critical role in Neandertal origins": grid5_run.py

```
#!/usr/bin/env python3
"""
grid5_run.py -- the FEMALE-BIAS counterpart to grid3_run.py and grid4_run.py.

grid3 swept theta (F1 hybrid male fertility) and grid4 swept s (selection for the modern Y).
Here both are switched off (theta = 1, s = 0) and the swept parameter is instead

    female_bias = the probability that a CROSS-lineage child takes its MOTHER's lineage
                  (grows up in and belongs to her group) rather than its father's

An independent draw is made for every such child, so 1.0 is a fully matrilineal expansion,
0.5 is no bias at all, and 0.0 is fully patrilineal. Because mtDNA passes from mother to child,
a matrilineal group is a set of unbroken maternal lines and archaic mtDNA can never enter it;
as the bias falls, children of archaic mothers join the modern group and bring archaic mtDNA
with them. The question this grid answers is how strong the bias must be before modern mtDNA
reliably fixes, with autosomal ancestry held at the empirical value by recalibrating gamma.

Everything else is the vanilla baseline: grid 20x20, K = 20, r = 2, sampled at the sampling generation,
full-length runs (no early stopping) because the criterion is fixation.

Row layout:  seed  mt  Y  auto  X  extgen
Usage:       python grid5_run.py <female_bias> <gamma> <m> <reps> <seed0> <outfile>
"""
import os
import sys
import numpy as np
from dataclasses import replace
from spatial_sim2 import Config, run

SAMPLE_GEN = 6993
# --- expansion geometry, settable from the command line ---------------------------------
# The grid-20 / K-20 baseline has a post-wave census of ~8000, five times the post-mixture size
# the rest of the analysis assumes (Muller's Ratchet v32: NN-uniparental = 725-1000, diploid
# 1500-2000). Since the modern mtDNA's advantage is a ~0.3% coefficient whose potency scales
# with 2*Ne*s, that mismatch inflated every spatial result. r and m are NOT externally
# constrained (DR, 3 Sep 2026: "pulled from my head"), so they are exposed here too and their
# sensitivity is the honest thing to report rather than a fixed baseline.
#
# A flag only OVERRIDES a parameter this script holds fixed; where the script sweeps that
# parameter, the swept value still wins, so sensitivity panels keep their full range.
# K_early/K_early_gens are always ADDED, since no BASE dict carries them.
def _geom(kw):
    import sys as _s
    for flag, key, cast in (("--grid", "grid", int), ("--K", "K", int),
                            ("--m", "m", float), ("--r", "r", float)):
        if flag in _s.argv and key in kw:
            kw[key] = cast(_s.argv[_s.argv.index(flag) + 1])
    for flag, key in (("--K-early", "K_early"), ("--K-early-gens", "K_early_gens")):
        if flag in _s.argv:
            kw[key] = int(_s.argv[_s.argv.index(flag) + 1])
    if kw.get("K", 20) < 7:
        raise SystemExit("K < 7: deme reproductive failure collapses the archaic population")
    return kw


BASE = dict(grid=20, K=20, r=2.0, theta_f1m=1.0, s=0.0, sel_locus="none",
            gens=SAMPLE_GEN + 10, sample_gen=SAMPLE_GEN, stop_after_ext=0, record_every=1000)

BIASES = [0.0, 0.1, 0.2, 0.3, 0.4, 0.5, 0.6, 0.7, 0.8, 0.9, 1.0]
GAMMAS = [0.02, 0.04, 0.06, 0.08, 0.10, 0.12, 0.15, 0.20, 0.30]
MS     = [0.02, 0.03, 0.04, 0.05, 0.06, 0.08, 0.10, 0.12, 0.15]


def main():
    fb    = float(sys.argv[1]); gamma = float(sys.argv[2]); m = float(sys.argv[3])
    reps  = int(sys.argv[4]);   seed0 = int(sys.argv[5]);   outfile = sys.argv[6]
    os.makedirs(os.path.dirname(outfile) or ".", exist_ok=True)
    cfg0 = Config(**_geom(dict(BASE)), female_bias=fb, gamma=gamma, m=m)
    with open(outfile, "w") as f:
        f.write("seed\tmt\tY\tauto\tX\textgen\n")
        for seed in range(seed0, seed0 + reps):
            out, ext = run(replace(cfg0, seed=seed))
            i = int(np.argmin(np.abs(out["gen"] - SAMPLE_GEN)))
            eg = int(ext) if ext is not None else -1
            f.write(f"{seed}\t{out['mt_mod'][i]:.6g}\t{out['Y_mod'][i]:.6g}\t"
                    f"{out['auto_mod'][i]:.6g}\t{out['X_mod'][i]:.6g}\t{eg}\n")
    print("wrote", outfile)


if __name__ == "__main__":
    main()
```
