## Supplementary Information section 3 code base for "Hypothesis: A modern human range expansion ∼300,000 years ago had a critical role in Neandertal origins": grid5_submit.py

```
#!/usr/bin/env python3
"""grid5_submit.py -- SLURM array for the female_bias x gamma x m grid.

Usage:  python grid5_submit.py <reps> <batch> [concurrency] [partition] [--resume]
        python grid5_submit.py 20 5 300      # exploratory: 20 reps, small tasks, wide array
        python grid5_submit.py 100 50        # production
        python grid5_submit.py 20 5 500 --resume   # queue only the missing tasks
then:   sbatch grid5_array.sh

Wall time is set by concurrency, not by how the work is split, so raise the third argument as
high as the queue will allow. Output goes to out_grid5/.
"""
import sys, os
from grid5_run import BIASES, GAMMAS, MS


def tag(th, ga, m):
    return f"t{th:g}_g{ga:g}_m{m:g}".replace(".", "p")


GEOM_FLAGS = ("--grid", "--K", "--m", "--r", "--K-early", "--K-early-gens")


def _strip_flags(av):
    """Positional parsing must not be shifted by the geometry flags, so drop each flag AND its
    value before the positional args are read."""
    out, skip = [], False
    for a in av:
        if skip:
            skip = False; continue
        if a.startswith("--"):
            skip = a in GEOM_FLAGS or a == "--rna"; continue
        out.append(a)
    return out


def _geom_str():
    """Echo the geometry flags onto every task line. Without this the workers would silently run
    the grid-20 / K-20 default while the submit script had been told otherwise -- which produces
    plausible-looking numbers that are simply wrong."""
    import sys as _s
    return "".join(f" {f} {_s.argv[_s.argv.index(f) + 1]}"
                   for f in GEOM_FLAGS if f in _s.argv)


def main():
    GEOM = _geom_str()
    args = _strip_flags(sys.argv[1:])
    total = int(args[0]) if len(args) > 0 else 20
    batch = int(args[1]) if len(args) > 1 else 5
    want_cap = int(args[2]) if len(args) > 2 else 0   # 0 = no throttle
    part = args[3] if len(args) > 3 else "short"
    os.makedirs("out_grid5", exist_ok=True); os.makedirs("logs", exist_ok=True)
    resume = "--resume" in sys.argv          # skip tasks whose output already exists
    tasks, skipped = [], 0
    for th in BIASES:
        for ga in GAMMAS:
            for m in MS:
                for s0 in range(0, total, batch):
                    reps = min(batch, total - s0)
                    out = f"out_grid5/out_{tag(th, ga, m)}_s{s0}.txt"
                    if resume and os.path.exists(out) and sum(
                            1 for _ in open(out)) >= reps + 1:
                        skipped += 1
                        continue
                    tasks.append(f"{th} {ga} {m} {reps} {s0} {out}{GEOM}")
    open("grid5_params.txt", "w").write("\n".join(tasks) + "\n")
    if not tasks:
        print("nothing to do -- every output file is already complete"); return
    n = len(tasks); cap = min(n, want_cap) if want_cap else 0
    open("grid5_array.sh", "w").write(f"""#!/bin/bash
#SBATCH -J grid5
#SBATCH -p {part}
#SBATCH -t 0-11:59
#SBATCH -c 1
#SBATCH --mem=1G
#SBATCH -o logs/grid5_%A_%a.log
#SBATCH --array=0-{n - 1}{("%%%d" % cap) if cap else ""}

LINE=$(sed -n "$((SLURM_ARRAY_TASK_ID + 1))p" grid5_params.txt)
echo "task $SLURM_ARRAY_TASK_ID : $LINE"
python grid5_run.py $LINE
""")
    combos = len(BIASES) * len(GAMMAS) * len(MS)
    print(f"wrote grid5_params.txt ({n} tasks over {combos} parameter combinations)")
    if resume:
        print(f"  RESUME: skipped {skipped} tasks whose output is already complete")
    print(f"  female_bias {len(BIASES)} x gamma {len(GAMMAS)} x m {len(MS)}, {total} reps each")
    cpu_h = combos * total * 15.7 / 3600
    print(f"  ~{total * 15.7 / 60 * batch / total:.1f} min per task, {n} tasks")
    print(f"  estimated cost ~{cpu_h:.0f} CPU-hours "
          f"(~{60 * cpu_h / cap:.0f} min wall at {cap} concurrent)"
          if cap else "(no throttle)")
    print("submit:  sbatch grid5_array.sh")


if __name__ == "__main__":
    main()
```
