## Supplementary Information section 3 code base for "Hypothesis: A modern human range expansion ∼300,000 years ago had a critical role in Neandertal origins": jcal_collect.py

```
#!/usr/bin/env python3
"""
jcal_collect.py -- invert the calibration grid for the tuning value that puts autosomal
ancestry at 5%, and emit the production task list.

For each (scenario, sweep, value) the autosomal proportion is a monotone function of the
tuning knob -- gamma for the expansion (more interbreeding means MORE archaic introgression
and so LESS modern ancestry, i.e. decreasing), the pulse fraction f for the stable boundary
(increasing). The 5% crossing is found by linear interpolation in log(knob), the same
transform calibrate5.py uses, and is reported as knob_star.

Writes:
    joint_calibration.csv   one row per condition: knob_star, the bracketing grid points,
                            and a flag when 5% is not bracketed
    jcal_production.txt     ready-to-run joint_run.py argument lines at knob_star

    python jcal_collect.py [--reps 1000] [--batch 25] [--rna 1.25] [--grid G] [--K K] [--m M] [--r R]
"""
import glob
import os
import sys

import numpy as np

from joint_run import SVALS, THETAS, SCENARIOS, EXPANSIONS
from jcal_submit import GAMMAS, FVALS, SB_N

COL = dict(mt=2, Y=3, auto=4)


def pooled(sc, sweep, v, k):
    rows = []
    for f in glob.glob(f"out_jcal/out_{sc}_{sweep}_{v:g}_k{k:g}_s*.txt"):
        for line in open(f).read().strip().split("\n")[1:]:
            p = line.split("\t")
            if len(p) >= 7:
                rows.append([float(x) for x in p])
    if not rows:
        return None
    a = np.array(rows)
    return dict(n=len(a), auto=100 * a[:, COL["auto"]].mean(),
                joint=100 * (a[:, COL["mt"]] * a[:, COL["Y"]]).mean())


def solve(knobs, autos, target=5.0):
    """log-linear interpolation for the knob giving `target` % autosomal ancestry."""
    pts = [(k, a) for k, a in zip(knobs, autos) if a is not None]
    if len(pts) < 2:
        return None, None
    pts.sort(key=lambda t: t[0])
    ks = np.array([p[0] for p in pts]); au = np.array([p[1] for p in pts])
    for i in range(len(ks) - 1):
        lo, hi = au[i], au[i + 1]
        if (lo - target) * (hi - target) <= 0 and lo != hi:
            w = (target - lo) / (hi - lo)
            return float(np.exp(np.log(ks[i]) + w * (np.log(ks[i + 1]) - np.log(ks[i])))), \
                   (float(ks[i]), float(ks[i + 1]))
    return None, (float(ks[0]), float(ks[-1]))


def main():
    reps = int(sys.argv[sys.argv.index("--reps") + 1]) if "--reps" in sys.argv else 1000
    batch = int(sys.argv[sys.argv.index("--batch") + 1]) if "--batch" in sys.argv else 25
    rna = (sys.argv[sys.argv.index("--rna") + 1] if "--rna" in sys.argv else None)
    # The geometry must be echoed into the production lines, otherwise the production pass
    # would silently run the DEFAULT grid 20 / K 20 at gammas calibrated for a shrunken grid.
    grid = (sys.argv[sys.argv.index("--grid") + 1] if "--grid" in sys.argv else None)
    Kdeme = (sys.argv[sys.argv.index("--K") + 1] if "--K" in sys.argv else None)
    mmig = (sys.argv[sys.argv.index("--m") + 1] if "--m" in sys.argv else None)
    _ke  = (sys.argv[sys.argv.index("--K-early") + 1] if "--K-early" in sys.argv else None)
    _keg = (sys.argv[sys.argv.index("--K-early-gens") + 1]
            if "--K-early-gens" in sys.argv else None)
    rfit = (sys.argv[sys.argv.index("--r") + 1] if "--r" in sys.argv else None)
    geom = ((f" --grid {grid}" if grid else "") + (f" --K {Kdeme}" if Kdeme else "")
            + (f" --m {mmig}" if mmig else "") + (f" --r {rfit}" if rfit else "")
              + (f" --K-early {_ke}" if _ke else "")
              + (f" --K-early-gens {_keg}" if _keg else ""))

    prod, nfail = [], 0
    with open("joint_calibration.csv", "w") as fh:
        fh.write("scenario,sweep,value,knob,knob_star,bracket_lo,bracket_hi,"
                 "auto_at_lo,auto_at_hi,status\n")
        for sc in SCENARIOS:
            expansion = sc in EXPANSIONS
            knobs = GAMMAS if expansion else FVALS
            flag = "--gamma" if expansion else "--f"
            Narg = "-" if expansion else str(SB_N)
            b = batch if expansion else batch * 4
            print(f"\n  {sc}   knob = {'gamma' if expansion else 'f'}")
            for sweep, vals in (("s", SVALS), ("theta", THETAS)):
                for v in vals:
                    ds = [pooled(sc, sweep, v, k) for k in knobs]
                    autos = [d["auto"] if d else None for d in ds]
                    if all(a is None for a in autos):
                        continue
                    star, br = solve(knobs, autos)
                    lo, hi = (br if br else (float("nan"),) * 2)
                    alo = next((a for a in autos if a is not None), float("nan"))
                    ahi = next((a for a in reversed(autos) if a is not None), float("nan"))
                    st = "ok" if star else "NOT BRACKETED"
                    if not star:
                        nfail += 1
                    fh.write(f"{sc},{sweep},{v:g},{'gamma' if expansion else 'f'},"
                             f"{star if star else ''},{lo:g},{hi:g},{alo:.2f},{ahi:.2f},{st}\n")
                    print(f"    {sweep:>5}={v:<6g} auto {alo:5.1f}%..{ahi:5.1f}%  -> "
                          + (f"{'gamma' if expansion else 'f'}* = {star:.4f}" if star
                             else "NOT BRACKETED -- widen the grid"))
                    if star:
                        for s0 in range(0, reps, b):
                            r = min(b, reps - s0)
                            out = f"out_joint2/out_{sc}_{sweep}_{v:g}_s{s0}.txt"
                            extra = (f" {flag} {star:.6g}" + (f" --rna {rna}" if rna else "")
                                     + (geom if expansion else ""))
                            prod.append(f"{sc} {sweep} {v} {Narg} {r} {s0} {out}{extra}")
    open("jcal_production.txt", "w").write("\n".join(prod) + "\n")
    print(f"\nsaved joint_calibration.csv and jcal_production.txt ({len(prod)} tasks)")
    if nfail:
        print(f"  WARNING: {nfail} conditions did not bracket 5% -- widen GAMMAS/FVALS "
              f"in jcal_submit.py and rerun those")
    print("\nto run the production pass:")
    print("  cp jcal_production.txt joint2_params.txt")
    print("  # then submit an array over joint2_params.txt calling joint_run.py")


if __name__ == "__main__":
    main()
```
