## Supplementary Information section 3 code base for "Hypothesis: A modern human range expansion ∼300,000 years ago had a critical role in Neandertal origins": jcal_submit.py

```
#!/usr/bin/env python3
"""
jcal_submit.py -- calibration grid for series (b), (c) and (d) of main-text Figure 1B.

Autosomal ancestry is not automatically 5% once theta or s is switched on, so the four series
are not comparable at vanilla parameters. Observed drift in the uncalibrated runs:

    (b) nexp, theta sweep :  4.5% -> 8.7%   (removing F1 males removes their fathers'
                                             archaic autosomes, so modern ancestry rises)
    (c) fsb,  theta sweep :  5.0% -> 2.5%   (the mirror image under an all-female pulse:
                                             the F1 sons removed carry MODERN maternal DNA)
    (d) nsb,  s sweep     :  4.8% -> 11.3%  (strong selection on the Y removes archaic-Y
                                             males and the archaic autosomes they carry)

This sweeps the tuning parameter -- gamma for the expansion, the pulse fraction f for the
stable boundary -- across every condition at modest replication. jcal_collect.py then inverts
each condition for the value that puts autosomal ancestry at 5%, and the production pass reruns
at 1,000 replicates using those values.

Usage:  python jcal_submit.py <reps> <batch> [concurrency] [partition] [--resume]
                              [--only fexp|nexp|fsb|nsb] [--rna F] [--grid G] [--K K] [--m M] [--r R] [--m M] [--r R]
        python jcal_submit.py 200 25 500 short --rna 1.25 --grid 14 --K 8
then:   sbatch jcal_array.sh          (output -> out_jcal/)
"""
import os
import sys

from joint_run import SVALS, THETAS, SCENARIOS, EXPANSIONS

# gamma for the expansion: the vanilla 0.12 sits near the low end, and the calibrated gamma*
# of the matrilineal grid reaches 0.359, so the range has to run well above 0.12.
# Widened for the size-matched geometry. On the old grid 20 / K 20 the calibrated gamma* sat
# near 0.12-0.36; on a grid shrunk to a post-wave census of ~1500 the wave crosses fewer demes
# and so absorbs less archaic ancestry per unit gamma, pushing gamma* up to roughly 0.3-0.5.
# gamma is an assortative-mating discount on cross matings, so 1.0 (random mating) is its
# hard ceiling -- if 5% is not bracketed below 1.0 the geometry cannot reach the target at all.
# Reverted to the original ladder on 5 Sep 2026 after testing both directions.
# LOWER gamma means LESS cross-mating and therefore HIGHER modern ancestry, so extending
# downward was the wrong direction. Extending UPWARD past random mating (1.5, 2.0, 3.0) does
# not help either: at fexp theta = 0 autosomal ancestry RISES from 14.0% at gamma = 1 to 30.3%
# at gamma = 3, because in the Q-line formulas g multiplies the archaic-father x modern-mother
# term as well, and that pairing yields a modern-lineage child at rate r*g. More cross-mating
# feeds the expanding lineage as fast as it dilutes it.
#
# CONCLUSION: the nine cells that fail to bracket 5% (fexp theta <= 0.05, and s = 0.64 in both
# expansions) are not a grid problem. They are the contact-budget ceiling: on this landscape
# the wave cannot absorb enough archaic ancestry to end at 5% once hybrid sons are removed or
# archaic-Y males are strongly selected against. They are reported as unreachable in the deme
# framework. The band model calibrates the same cells without difficulty (theta = 0 at
# e* = 0.292, autosomes 4.4%, P(joint) = 77%) because exogamy contact does not require the two
# groups to coexist at a moving front.
GAMMAS = [0.06, 0.10, 0.15, 0.22, 0.30, 0.40, 0.55, 0.75, 1.0]
# pulse fraction for the stable boundary, bracketing the nominal 0.05 generously in both
# directions because theta pushes autosomal ancestry down and s pushes it up.
# 0.008-0.015 were added after the first pass: at nsb with s = 0.64, selection on the Y
# is so strong that it drags autosomal ancestry to 5.6% even at f = 0.02, so 5% was not
# bracketed from below. Everything else calibrated inside the original range.
FVALS = [0.008, 0.012, 0.016, 0.02, 0.03, 0.04, 0.05, 0.065, 0.08, 0.10, 0.13]
# Stable-boundary / migration-pulse census. 1312 is the interpolated value at which the
# measured autosomal effective size lands on the Peyregne anchor of 1,447 diploids
# (2,894 gene copies): N = 1265 gave 2715 and N = 1390 gave 3188. Was 1500.
from anchors import PANM_CENSUS as SB_N   # 1312; see anchors.py


def main():
    args   = [a for a in sys.argv[1:] if not a.startswith("--")]
    total  = int(args[0]) if len(args) > 0 else 200
    batch  = int(args[1]) if len(args) > 1 else 25
    cap    = int(args[2]) if len(args) > 2 else 500
    part   = args[3] if len(args) > 3 else "short"
    resume = "--resume" in sys.argv
    rna    = (sys.argv[sys.argv.index("--rna") + 1] if "--rna" in sys.argv else None)
    # Expansion geometry, passed straight through to joint_run.py. Omit to keep the published
    # grid 20 / K 20; set it to shrink the post-wave census to the post-mixture size the rest
    # of the analysis assumes (Muller's Ratchet v32: diploid 1500 / uniparental 725).
    grid   = (sys.argv[sys.argv.index("--grid") + 1] if "--grid" in sys.argv else None)
    Kdeme  = (sys.argv[sys.argv.index("--K") + 1] if "--K" in sys.argv else None)
    mmig   = (sys.argv[sys.argv.index("--m") + 1] if "--m" in sys.argv else None)
    _ke    = (sys.argv[sys.argv.index("--K-early") + 1] if "--K-early" in sys.argv else None)
    _keg   = (sys.argv[sys.argv.index("--K-early-gens") + 1]
              if "--K-early-gens" in sys.argv else None)
    rfit   = (sys.argv[sys.argv.index("--r") + 1] if "--r" in sys.argv else None)
    geom   = ((f" --grid {grid}" if grid else "") + (f" --K {Kdeme}" if Kdeme else "")
              + (f" --m {mmig}" if mmig else "") + (f" --r {rfit}" if rfit else "")
              + (f" --K-early {_ke}" if _ke else "")
              + (f" --K-early-gens {_keg}" if _keg else ""))
    only   = (sys.argv[sys.argv.index("--only") + 1] if "--only" in sys.argv else None)
    scen   = [s for s in SCENARIOS if only is None or s == only]
    os.makedirs("out_jcal", exist_ok=True); os.makedirs("logs", exist_ok=True)

    tasks, skipped = [], 0
    for sc in scen:
        expansion = sc in EXPANSIONS
        b = batch if expansion else batch * 4
        Narg = "-" if expansion else str(SB_N)
        knobs = GAMMAS if expansion else FVALS
        flag = "--gamma" if expansion else "--f"
        for sweep, vals in (("s", SVALS), ("theta", THETAS)):
            for v in vals:
                for k in knobs:
                    for s0 in range(0, total, b):
                        reps = min(b, total - s0)
                        out = f"out_jcal/out_{sc}_{sweep}_{v:g}_k{k:g}_s{s0}.txt"
                        if resume and os.path.exists(out) and \
                                sum(1 for _ in open(out)) >= reps + 1:
                            skipped += 1; continue
                        extra = (f" {flag} {k}" + (f" --rna {rna}" if rna else "")
                                 + (geom if expansion else ""))
                        tasks.append(f"{sc} {sweep} {v} {Narg} {reps} {s0} {out}{extra}")
    if not tasks:
        print("nothing to do -- every output file is already complete"); return
    open("jcal_params.txt", "w").write("\n".join(tasks) + "\n")
    n = len(tasks); cap = min(n, cap)
    open("jcal_array.sh", "w").write(f"""#!/bin/bash
#SBATCH -J jcal
#SBATCH -p {part}
#SBATCH -t 0-11:59
#SBATCH -c 1
#SBATCH --mem=4G
#SBATCH -o logs/jcal_%A_%a.log
#SBATCH --array=0-{n - 1}{("%%%d" % cap) if cap else ""}

LINE=$(sed -n "$((SLURM_ARRAY_TASK_ID + 1))p" jcal_params.txt)
echo "task $SLURM_ARRAY_TASK_ID : $LINE"
python joint_run.py $LINE
""")
    print(f"wrote jcal_params.txt ({n} tasks), {total} reps each")
    if resume: print(f"  RESUME: skipped {skipped} complete tasks")
    print(f"  scenarios: {', '.join(scen)}" + (f"   RNA x{rna}" if rna else "")
          + (f"   geometry:{geom}" if geom else "   geometry: default grid 20 / K 20"))
    print("submit:  sbatch jcal_array.sh")


if __name__ == "__main__":
    main()
```
