## Supplementary Information section 3 code base for "Hypothesis: A modern human range expansion ∼300,000 years ago had a critical role in Neandertal origins": jdrive_submit.py

```
#!/usr/bin/env python3
"""
jdrive_submit.py -- the missing drive sweep for Route 2 of Figure 2. (12 Sep 2026)

DR: "Could you add the range expansion unbiased / migration pulse all female / migration pulse
all biased curves to Route 2 in Figure 2, and also add a trace for the range expansion,
matrilocal?"

WHY ROUTE 2 HAS NO LINES TODAY. Figure 2's four demographic scenario curves come from
joint2_summary.csv, which carries exactly two sweeps -- `s` (Route 1) and `theta` (Route 3).
There has never been a drive sweep for any scenario, so that panel is points-only. joint_run.py
now has a third sweep, `drive`, and this script runs it.

WHAT THIS COVERS, AND WHAT IT CANNOT

    range expansion, matrilocal   fexp   -- run here, both cost arms
    range expansion, unbiased     nexp   -- run here, cost-free
    migration pulse, all-female   fsb    -- NOT RUN, AND NOT NEEDED. No modern male enters an
                                           all-female pulse, so the modern Y starts at zero and
                                           stays there whatever the drive does to archaic-Y
                                           transmission. The curve is identically zero and can
                                           be drawn as such, exactly as it already is in Routes
                                           1 and 3.
    migration pulse, unbiased     nsb    -- BLOCKED. panmictic_sim.py has no X-linked drive at
                                           all; joint_run raises rather than silently returning
                                           a no-mechanism control. Needs xdrive implemented in
                                           that engine first.

THE fexp COST ARM DOES DOUBLE DUTY. It is also the deme curve for panel (a) of the new Figure
S3.4. The band framework already has that ladder at nine rungs from bandmech_summary.csv, but
the deme side only had the old dedicated run at d = 0.5/0.8/0.95/1 -- nothing below 75%
daughters, which is the interesting region where band climbs from 0.15 to 0.98. Running the
full seven-rung ladder here fills it.

TWO STAGES, BECAUSE gamma HAS TO BE RETUNED PER RUNG. Removing archaic-Y males removes their
autosomes too, so modern autosomal ancestry rises with drive strength and the baseline gamma of
0.12 no longer lands on 5%. Uncalibrated, a smoke test at d = 0.4 came back at 18-20% autosomal.
Stage `cal` runs a gamma ladder at 30 replicates per point; `prod` fits gamma* per condition by
regressing autosomal ancestry on log(gamma) and inverting at 5%, then runs 400 replicates there.
That is the same pattern jcal -> joint2 used for the s and theta sweeps.

    python jdrive_submit.py cal        # ~13 CPU-hours
    python jdrive_submit.py prod       # ~35 CPU-hours, after cal has drained
    python jdrive_submit.py prod --resume
"""
import glob
import math
import os
import subprocess
import sys

HERE = os.path.dirname(os.path.abspath(__file__))

# cfg.xdrive = d is the probability a Y-bearing conception of an expressing male is converted to
# a daughter, so daughters = 50*(1+d). This is the ladder Figure 2's Route 2 axis uses.
# d = 0 ADDED 13 Sep. DR: "connect red line to 50% in Route 2". The panel's x rungs start at
# d = 0, i.e. 50% daughters and no distortion at all, and the Cartesian cloud is drawn there --
# but the sweep began at d = 0.05, so the expansion LINES started one rung in and left the 50%
# cloud unconnected. d = 0 is the null: the line should arrive at essentially zero, which is
# what makes the rise from it meaningful.
DVALS = [0.0, 0.05, 0.1, 0.2, 0.4, 0.6, 0.8, 1.0]
# (scenario, xdrive_cost, label) -- the three series that can be run at all.
SERIES = [("fexp", 0.0, "matrilocal expansion, no cost"),
          ("fexp", 0.10, "matrilocal expansion, 10% fertility cost"),
          ("nexp", 0.0, "unbiased expansion, no cost")]
GAMMAS = [0.12, 0.20, 0.30, 0.45, 0.65]     # probe ladder; gamma rises with drive strength
CAL_REPS, PROD_REPS = 30, 400
GEOM = ["--grid", "12", "--K", "11"]        # the anchored geometry, not EXP_BASE's 20/20
TARGET = 5.0                                # percent modern autosomal ancestry
OUT_CAL, OUT_PROD = "out_jdrive_cal", "out_jdrive"
PART, CAP = "short", 400


def tag(sc, cost, d, g=None):
    t = f"{sc}_c{cost:g}_d{d:g}".replace(".", "p")
    return t + (f"_g{g:g}".replace(".", "p") if g is not None else "")


def autosomal(path):
    """Mean autosomal percentage over established replicates of one joint_run tsv."""
    vals = []
    for line in open(path).read().splitlines()[1:]:
        f = line.split("\t")
        if len(f) < 5:
            continue
        try:
            vals.append(100.0 * float(f[4]))
        except ValueError:
            continue
    return sum(vals) / len(vals) if vals else None


def fit_gamma(sc, cost):
    """gamma* solving autosomal(gamma) = 5%, by least squares on log gamma."""
    pts = []
    for g in GAMMAS:
        for d in DVALS:
            p = os.path.join(HERE, OUT_CAL, tag(sc, cost, d, g) + ".tsv")
            if os.path.exists(p):
                a = autosomal(p)
                if a:
                    pts.append((d, g, a))
    out = {}
    for d in DVALS:
        sub = [(math.log(g), a) for dd, g, a in pts if dd == d]
        if len(sub) < 2:
            continue
        n = len(sub)
        mx = sum(x for x, _ in sub) / n
        my = sum(y for _, y in sub) / n
        den = sum((x - mx) ** 2 for x, _ in sub)
        if den == 0:
            continue
        b = sum((x - mx) * (y - my) for x, y in sub) / den
        a0 = my - b * mx
        if b == 0:
            continue
        # clamp into the probed range rather than extrapolating off the end of the ladder
        g = math.exp((TARGET - a0) / b)
        out[d] = min(max(g, GAMMAS[0]), GAMMAS[-1])
    return out


def write_and_submit(name, tasks, hours="0-05:59"):
    if not tasks:
        print("  nothing to submit")
        return
    pf, sf = f"{name}_params.txt", f"{name}_array.sh"
    open(os.path.join(HERE, pf), "w").write("\n".join(tasks) + "\n")
    open(os.path.join(HERE, sf), "w").write(f"""#!/bin/bash
#SBATCH -J {name}
#SBATCH -p {PART}
#SBATCH -c 1
#SBATCH --mem=2G
#SBATCH -t {hours}
#SBATCH -a 0-{len(tasks) - 1}%{CAP}
#SBATCH -o logs/{name}_%A_%a.log
cd {HERE}
CMD=$(sed -n "$((SLURM_ARRAY_TASK_ID + 1))p" {pf})
echo "$CMD"
eval "$CMD"
""")
    print(f"  {len(tasks)} tasks -> {sf}")
    if "--no-go" in sys.argv:
        print(f"  NOT submitted (--no-go).  sbatch {sf}")
        return
    r = subprocess.run(["sbatch", sf], cwd=HERE, capture_output=True, text=True)
    print(f"  SUBMITTED: {r.stdout.strip()}" if r.returncode == 0
          else f"  !! sbatch FAILED: {(r.stderr or '').strip()}")


def main():
    stage = sys.argv[1] if len(sys.argv) > 1 else "cal"
    reps = int(sys.argv[sys.argv.index("--reps") + 1]) if "--reps" in sys.argv else PROD_REPS
    os.makedirs(os.path.join(HERE, "logs"), exist_ok=True)

    if stage == "cal":
        os.makedirs(os.path.join(HERE, OUT_CAL), exist_ok=True)
        tasks = []
        for sc, cost, _ in SERIES:
            for d in DVALS:
                for g in GAMMAS:
                    t = tag(sc, cost, d, g)
                    out = os.path.join(OUT_CAL, t + ".tsv")
                    if os.path.exists(os.path.join(HERE, out)):
                        continue
                    tasks.append(f"python joint_run.py {sc} drive {d:g} - {CAL_REPS} 0 {out} "
                                 + " ".join(GEOM) + f" --gamma {g:g} --xcost {cost:g}")
        print(f"CALIBRATION  {len(SERIES)} series x {len(DVALS)} rungs x {len(GAMMAS)} gammas "
              f"x {CAL_REPS} reps")
        write_and_submit("jdcal", tasks)
        return

    if stage == "prod":
        os.makedirs(os.path.join(HERE, OUT_PROD), exist_ok=True)
        n_cal = len(glob.glob(os.path.join(HERE, OUT_CAL, "*.tsv")))
        want = len(SERIES) * len(DVALS) * len(GAMMAS)
        print(f"CALIBRATION on disk: {n_cal}/{want}")
        if n_cal < want and "--resume" not in sys.argv:
            print("  incomplete. Wait for jdcal to drain, or pass --resume to fit on what is "
                  "there.")
            return
        tasks = []
        for sc, cost, lab in SERIES:
            gs = fit_gamma(sc, cost)
            print(f"  {lab}")
            for d in DVALS:
                if d not in gs:
                    print(f"    d={d:<5g} ({50 * (1 + d):5.1f}% daughters)  no gamma fit")
                    continue
                print(f"    d={d:<5g} ({50 * (1 + d):5.1f}% daughters)  gamma* = {gs[d]:.3f}")
                out = os.path.join(OUT_PROD, tag(sc, cost, d) + ".tsv")
                # --reps RAISES THE COUNT, AND SO HAS TO BEAT THE FILENAME SKIP. DR: the
                # unbiased (nexp) curve is visibly jagged. That is sampling noise and nothing
                # else: p_joint for nexp runs between 0.008 and 0.04, so at 400 replicates the
                # standard error is a third to a half of the value itself and neighbouring
                # rungs bounce around each other.
                #
                # joint_run writes one TSV per point holding every replicate at seeds
                # 0..reps-1, with no seed offset, so a higher count REPLACES the file rather
                # than appending to it -- the earlier 400 draws are discarded, not pooled. A
                # point is skipped only when its file already holds at least the requested
                # count, which makes the flag idempotent: running it twice at the same value
                # submits nothing the second time.
                have = 0
                full = os.path.join(HERE, out)
                if os.path.exists(full):
                    have = max(sum(1 for _ in open(full)) - 1, 0)   # minus the header
                if have >= reps:
                    continue
                tasks.append(f"python joint_run.py {sc} drive {d:g} - {reps} 1000 {out} "
                             + " ".join(GEOM) + f" --gamma {gs[d]:.4f} --xcost {cost:g}")
        print(f"\nPRODUCTION  {reps} replicates per point")
        write_and_submit("jdprod", tasks, hours="0-11:59")
        return

    raise SystemExit("stage must be 'cal' or 'prod'")


if __name__ == "__main__":
    main()
```
