## Supplementary Information section 3 code base for "Hypothesis: A modern human range expansion ∼300,000 years ago had a critical role in Neandertal origins": joint2_collect.py

```
#!/usr/bin/env python3
"""
joint2_collect.py -- pool out_joint2/ (the autosome-calibrated production pass) into
joint2_summary.csv.

Same quantity as joint_collect.py -- JOINT is the per-replicate product of modern Y and modern
mtDNA frequency at the sampling generation, averaged over replicates, not the product of the
two marginal means -- but read from the calibrated runs and annotated with the knob value that
was used, so the calibration can be audited from the summary alone.

The autosomal column is the check that the calibration worked: every row should now sit near
5%. Rows outside 4-6% are flagged, since the whole point of the pass was to make the four
series comparable at a common autosomal ancestry.

    python joint2_collect.py
"""
import csv
import glob
import os

import numpy as np

from joint_run import SVALS, THETAS, SCENARIOS, EXPANSIONS

COL = dict(s_mt=1, mt=2, Y=3, auto=4, X=5, ext=6)
LABEL = {"fexp": "(a) female-biased range expansion",
         "nexp": "(b) non-sex-biased range expansion",
         "fsb":  "(c) female-biased gene flow, stable boundary",
         "nsb":  "(d) non-sex-biased gene flow, stable boundary"}


def knobs():
    """scenario/sweep/value -> (knob name, calibrated value), from jcal_collect.py."""
    out = {}
    if not os.path.exists("joint_calibration.csv"):
        return out
    for r in csv.DictReader(open("joint_calibration.csv")):
        if r.get("knob_star"):
            out[(r["scenario"], r["sweep"], float(r["value"]))] = \
                (r["knob"], float(r["knob_star"]))
    return out


def pooled(sc, sweep, v, calibrated=None):
    # A condition whose calibration did not bracket 5% gets NO production line, so any output
    # files sitting in out_joint2/ for it are LEFTOVERS FROM A PREVIOUS GEOMETRY. They pool
    # silently and look like real data -- nexp s=0.64 came through as the old grid-20 value of
    # 37.99% this way. Refuse anything without a calibrated knob.
    if calibrated is not None and (sc, sweep, float(v)) not in calibrated:
        return None
    rows = []
    for f in glob.glob(f"out_joint2/out_{sc}_{sweep}_{v:g}_s*.txt"):
        for line in open(f).read().strip().split("\n")[1:]:
            p = line.split("\t")
            if len(p) >= 7:
                rows.append([float(x) for x in p])
    if not rows:
        return None
    a = np.array(rows)
    mt, Y = a[:, COL["mt"]], a[:, COL["Y"]]
    return dict(n=len(a),
                joint=100 * (mt * Y).mean(),
                mt=100 * mt.mean(), Y=100 * Y.mean(),
                auto=100 * a[:, COL["auto"]].mean(), X=100 * a[:, COL["X"]].mean(),
                bothfix=100 * ((mt >= 0.999) & (Y >= 0.999)).mean(),
                s_mt=a[:, COL["s_mt"]].mean())


def main():
    kn = knobs()
    # the set of conditions that actually have a calibrated knob
    cal = {k for k, v in kn.items() if v[1] == v[1]}
    stale = 0
    bad = []
    with open("joint2_summary.csv", "w") as fh:
        fh.write("scenario,sweep,value,knob,knob_star,n,joint_pct,mt_mean,Y_mean,auto_mean,"
                 "X_mean,P_both_fixed,s_mt_mean\n")
        for sc in SCENARIOS:
            for sweep, vals in (("s", SVALS), ("theta", THETAS)):
                for v in vals:
                    d = pooled(sc, sweep, v, cal)
                    if d is None:
                        continue
                    nm, ks = kn.get((sc, sweep, float(v)), ("", float("nan")))
                    fh.write(f"{sc},{sweep},{v:g},{nm},{ks:.6g},{d['n']},{d['joint']:.2f},"
                             f"{d['mt']:.2f},{d['Y']:.2f},{d['auto']:.2f},{d['X']:.2f},"
                             f"{d['bothfix']:.2f},{d['s_mt']:.6g}\n")
                    if not 4.0 <= d["auto"] <= 6.0:
                        bad.append((sc, sweep, v, d["auto"]))
    print("saved joint2_summary.csv\n")
    for sc in SCENARIOS:
        shown = False
        for sweep, vals in (("s", SVALS), ("theta", THETAS)):
            rows = [(v, pooled(sc, sweep, v, cal)) for v in vals]
            rows = [(v, d) for v, d in rows if d is not None]
            if not rows:
                continue
            if not shown:
                print(f"  {LABEL[sc]}"); shown = True
            print(f"    sweep = {sweep}")
            print(f"      {'value':>7} {'knob*':>8} {'n':>6} {'JOINT%':>8} {'mt%':>7} "
                  f"{'Y%':>7} {'auto%':>7}")
            for v, d in rows:
                _, ks = kn.get((sc, sweep, float(v)), ("", float("nan")))
                print(f"      {v:>7g} {ks:>8.4f} {d['n']:>6} {d['joint']:>8.2f} "
                      f"{d['mt']:>7.2f} {d['Y']:>7.2f} {d['auto']:>7.2f}")
        if shown:
            print()
    if bad:
        print(f"WARNING: {len(bad)} rows have autosomal ancestry outside 4-6%:")
        for sc, sweep, v, au in bad[:12]:
            print(f"    {sc} {sweep}={v:g} -> {au:.2f}%")
    else:
        print("autosomal ancestry is within 4-6% at every point: the four series are "
              "comparable")
    # the cell that reconciles against the note's stable-boundary mtDNA number
    d = pooled("nsb", "theta", 1, cal)
    if d:
        print(f"\nreconciliation cell -- (d) at theta = 1, no selection on the Y:")
        print(f"    mtDNA = {d['mt']:.1f}%   (Supplementary Information section 1 gives ~22% "
              f"with the RNA correction; 21.1% by curve integration)")


if __name__ == "__main__":
    main()
```
