## Supplementary Information section 3 code base for "Hypothesis: A modern human range expansion ∼300,000 years ago had a critical role in Neandertal origins": joint2_submit.py

```
#!/usr/bin/env python3
"""
joint2_submit.py -- production pass for series (b), (c) and (d) at the calibrated knobs.

jcal_collect.py writes jcal_production.txt: one joint_run.py argument line per task, already
carrying the calibrated --gamma or --f and the --rna factor. This wraps that list in a SLURM
array. Output goes to out_joint2/, kept separate from the uncalibrated out_joint/ so the two
can be compared rather than one silently overwriting the other.

Usage:  python joint2_submit.py [concurrency] [partition] [--resume]
then:   sbatch joint2_array.sh
"""
import os
import sys


def main():
    args = [a for a in sys.argv[1:] if not a.startswith("--")]
    cap = int(args[0]) if len(args) > 0 else 500
    part = args[1] if len(args) > 1 else "short"
    resume = "--resume" in sys.argv
    if not os.path.exists("jcal_production.txt"):
        raise SystemExit("jcal_production.txt not found -- run jcal_collect.py first")
    os.makedirs("out_joint2", exist_ok=True); os.makedirs("logs", exist_ok=True)

    tasks, skipped = [], 0
    for ln in open("jcal_production.txt"):
        ln = ln.strip()
        if not ln:
            continue
        t = ln.split()
        idx = max(i for i, x in enumerate(t) if x.endswith(".txt"))
        out, reps = t[idx], int(t[idx - 2])
        if resume and os.path.exists(out) and sum(1 for _ in open(out)) >= reps + 1:
            skipped += 1; continue
        tasks.append(ln)
    if not tasks:
        print("nothing to do -- every output file is already complete"); return
    open("joint2_params.txt", "w").write("\n".join(tasks) + "\n")
    n = len(tasks); cap = min(n, cap)
    open("joint2_array.sh", "w").write(f"""#!/bin/bash
#SBATCH -J joint2
#SBATCH -p {part}
#SBATCH -t 0-11:59
#SBATCH -c 1
#SBATCH --mem=4G
#SBATCH -o logs/joint2_%A_%a.log
#SBATCH --array=0-{n - 1}{("%%%d" % cap) if cap else ""}

LINE=$(sed -n "$((SLURM_ARRAY_TASK_ID + 1))p" joint2_params.txt)
echo "task $SLURM_ARRAY_TASK_ID : $LINE"
python joint_run.py $LINE
""")
    print(f"wrote joint2_params.txt ({n} tasks)")
    if resume: print(f"  RESUME: skipped {skipped} complete tasks")
    print("submit:  sbatch joint2_array.sh")


if __name__ == "__main__":
    main()
```
