## Supplementary Information section 3 code base for "Hypothesis: A modern human range expansion ∼300,000 years ago had a critical role in Neandertal origins": joint_collect.py

```
#!/usr/bin/env python3
"""
joint_collect.py -- pool out_joint/ into joint_summary.csv.

The plotted quantity is JOINT: the per-replicate product of modern Y and modern mtDNA
frequency at the sampling generation, averaged over replicates. It is computed per replicate
and then averaged, NOT as the product of the two marginal means -- those differ whenever the
two markers are correlated across replicates, which they are as soon as anything acts on both.

Also written, for checking rather than plotting:
  * mt_mean, Y_mean, auto_mean, X_mean -- the marginals. auto_mean should sit near 5%; if it
    does not, the scenario is not comparable to the others and the joint value is misleading.
  * P_both_fixed -- both markers fixed modern in the same replicate.
  * s_mt_mean    -- the mean posterior draw actually used, as a check that the integration
                    sampled the intended distribution.
  * P_ext        -- fraction of expansion replicates that ran to archaic extinction.

    python joint_collect.py
"""
import glob
import os
import re

import numpy as np

from joint_run import SVALS, THETAS, SCENARIOS

COL = dict(s_mt=1, mt=2, Y=3, auto=4, X=5, ext=6)
LABEL = {"fexp": "(a) female-biased (matrilocal) range expansion",
         "nexp": "(b) non-sex-biased range expansion",
         "fsb":  "(c) female-biased gene flow, stable boundary",
         "nsb":  "(d) non-sex-biased gene flow, stable boundary"}


def pooled(sc, sweep, v):
    rows = []
    for f in glob.glob(f"out_joint/out_{sc}_{sweep}_{v:g}_s*.txt"):
        for line in open(f).read().strip().split("\n")[1:]:
            p = line.split("\t")
            if len(p) >= 7:
                rows.append([float(x) for x in p])
    if not rows:
        return None
    a = np.array(rows)
    mt, Y = a[:, COL["mt"]], a[:, COL["Y"]]
    ext = a[:, COL["ext"]]
    return dict(n=len(a),
                joint=100 * (mt * Y).mean(),
                mt=100 * mt.mean(), Y=100 * Y.mean(),
                auto=100 * a[:, COL["auto"]].mean(), X=100 * a[:, COL["X"]].mean(),
                bothfix=100 * ((mt >= 0.999) & (Y >= 0.999)).mean(),
                s_mt=a[:, COL["s_mt"]].mean(),
                pext=100 * (ext >= 0).mean())


def main():
    with open("joint_summary.csv", "w") as fh:
        fh.write("scenario,sweep,value,n,joint_pct,mt_mean,Y_mean,auto_mean,X_mean,"
                 "P_both_fixed,s_mt_mean,P_ran_to_extinction\n")
        for sc in SCENARIOS:
            for sweep, vals in (("s", SVALS), ("theta", THETAS)):
                for v in vals:
                    d = pooled(sc, sweep, v)
                    if d is None:
                        continue
                    fh.write(f"{sc},{sweep},{v:g},{d['n']},{d['joint']:.2f},{d['mt']:.2f},"
                             f"{d['Y']:.2f},{d['auto']:.2f},{d['X']:.2f},{d['bothfix']:.2f},"
                             f"{d['s_mt']:.6g},{d['pext']:.1f}\n")
    print("saved joint_summary.csv\n")
    for sc in SCENARIOS:
        any_rows = False
        for sweep, vals in (("s", SVALS), ("theta", THETAS)):
            rows = [(v, pooled(sc, sweep, v)) for v in vals]
            rows = [(v, d) for v, d in rows if d is not None]
            if not rows:
                continue
            if not any_rows:
                print(f"  {LABEL[sc]}")
                any_rows = True
            print(f"    sweep = {sweep}")
            print(f"      {'value':>7} {'n':>6} {'JOINT%':>8} {'mt%':>7} {'Y%':>7} "
                  f"{'auto%':>7} {'s_mt':>9}")
            for v, d in rows:
                print(f"      {v:>7g} {d['n']:>6} {d['joint']:>8.2f} {d['mt']:>7.2f} "
                      f"{d['Y']:>7.2f} {d['auto']:>7.2f} {d['s_mt']:>9.5f}")
        if any_rows:
            bad = [(sweep, v, d['auto']) for sweep, vals in (("s", SVALS), ("theta", THETAS))
                   for v in vals if (d := pooled(sc, sweep, v)) and not 2.0 <= d['auto'] <= 15.0]
            if bad:
                print(f"      NOTE: autosomal ancestry outside 2-15% at {len(bad)} points, "
                      f"e.g. {bad[0][0]}={bad[0][1]:g} -> {bad[0][2]:.1f}%")
            print()


if __name__ == "__main__":
    main()
```
