## Supplementary Information section 3 code base for "Hypothesis: A modern human range expansion ∼300,000 years ago had a critical role in Neandertal origins": joint_run.py

```
#!/usr/bin/env python3
"""
joint_run.py -- the three not-yet-run series of the main-text Figure 1B comparison, with the
mtDNA carrying a fitness coefficient drawn per replicate from the Muller's-ratchet posterior.

The figure plots ONE trace per series: the product of modern Y and modern mtDNA frequency at
the sampling generation, averaged over replicates. Autosomal ancestry is recorded so that it
can be checked to sit near 5%, but is not plotted.

    SERIES        SCENARIO                                          STATUS
    (a)  fexp     female-biased (matrilineal) range expansion       already run: scan4 sY
                                                                     + theta_*
    (b)  nexp     NON-sex-biased range expansion                    this script
    (c)  fsb      female-biased gene flow at a stable boundary      this script
    (d)  nsb      non-sex-biased gene flow at a stable boundary     this script

EXPECTED RESULT FOR (c), SO IT IS NOT MISREAD AS A BUG. The female-biased stable boundary uses
an all-female pulse (male_frac = 0), so no modern male enters and the modern Y starts at zero.
Selection cannot act on an allele that is absent, and the modern Y therefore stays at zero at
every s and every theta: series (c) is a flat zero line, and its Y x mtDNA product is
identically zero. Its mtDNA does well -- entering at 2f = 10% with the ratchet advantage -- so
the series says that female-biased flow at a boundary delivers the mtDNA and nothing at all on
the Y. This is intended, and was confirmed against the author before the runs were submitted.

Series (a) needs no mtDNA simulation: under matriliny the expanding population carries modern
mtDNA by construction, so its trace is simply the Y frequency. In (b), (c) and (d) the modern
mtDNA is NOT guaranteed, so it has to be simulated, and the honest way to do that is to give
it the fitness gap the Muller's ratchet actually implies rather than a single point estimate.

THE mtDNA COEFFICIENT. Each replicate draws s_mt from the accepted ABC posterior on the
modern-minus-archaic mtDNA fitness gap at 250 ka (Supplementary Information section 1),
integrating over every parameter that posterior integrates over -- g_c, beta, Sbar and the
rest. Draws are read from mt_gap_posterior_NA<NA>.txt, written by make_mt_posterior.py, and
they include the negative tail (8.5% of draws at N_A = 1500, 24% at N_A = 3000): dropping it
would turn a two-sided posterior into a one-sided one and bias every result upward. Drawing
per replicate rather than fixing s_mt at its median is what makes the resulting frequency an
integral over the posterior rather than a conditional-on-the-median value.

Set --rna to multiply each draw by an RNA-correction factor (the note estimates x1.25 to x1.5
for the rRNA and tRNA contribution to the deleterious input); the default of 1.0 applies none.

Row layout:  seed  s_mt  mt  Y  auto  X  extgen
Usage:
    python joint_run.py <scenario> <sweep> <value> <N|-> <reps> <seed0> <outfile> [--rna F]
        scenario : nexp | fsb | nsb
        sweep    : s      (additive selection against the archaic Y)
                 | theta  (relative fertility of F1 hybrid males)
        N        : census size for the stable-boundary scenarios; "-" for the expansions
"""
import os
import sys
import zlib
from dataclasses import replace

import numpy as np

SAMPLE_GEN = 6993
# THE UPPER LADDER NOW MATCHES THE MECHANISM SCREEN -- CHANGED 14 Sep 2026.
# DR: the baseline lines in Figure 2b's Route 1 "are out of synch" with the ticks. They were.
# Above s = 0.02 this ladder doubled (0.04, 0.08, 0.16, 0.32, 0.64) while routes_build screened
# 0.05, 0.10, 0.20, 0.35, 0.50, so the four scenario curves put their vertices between the
# fountains and between the ticks -- five markers per curve, none of them over a tick.
#
# The low end is unchanged and its files are untouched: 0, 0.001, 0.002, 0.005, 0.01 and 0.02
# are common to both ladders and already have 1,000 replicates each. Only the doubling tail is
# replaced, so the rerun is five rungs per scenario rather than eleven.
#
# 0.64 is dropped rather than re-sited. It was the top of the doubling ladder, it is off the
# right-hand end of Route 1's axis (which stops at 0.5), and it is one of the nine cells
# jcal_submit records as failing to bracket 5% autosomal ancestry in both expansions, so it
# carries no calibrated point for two of the four curves anyway.
SVALS = [0.0, 0.001, 0.002, 0.005, 0.01, 0.02, 0.05, 0.10, 0.20, 0.35, 0.50]
THETAS = [1, 0.5, 0.2, 0.1, 0.05, 0.02, 0.01, 0.005, 0.002, 0.001, 0]
# fexp is series (a), the female-biased (matrilineal) range expansion. It used to come from a
# separate pipeline (scan4_*/theta_run -> figure3_full.csv, fig3_theta.csv), which was fine
# while the expansion geometry was fixed, but once the geometry has to be matched to the
# post-mixture size it has to be recalibrated exactly like the others -- so it is folded in
# here and goes through the same jcal calibration and the same joint2 production pass.
SCENARIOS = ("fexp", "nexp", "fsb", "nsb")
EXPANSIONS = ("fexp", "nexp")           # gamma is the calibration knob for these
FEMALE_BIAS = {"fexp": 1.0, "nexp": 0.5}
SWEEPS = ("s", "theta", "drive", "twopulse", "mtydmi")
# mtydmi added 14 Sep for Figure 2b's mtDNA x Y column, which had a fountain and no
# comparison scenarios: `value` is the DMI penetrance. drive added 12 Sep for Route 2.
# twopulse added 14 Sep for Route 4: `value` is the
# generation at which a SECOND pulse arrives, so the stable-boundary scenarios have a
# two-wave counterpart to the expansions' wave2_gen and Route 4 can carry the same
# comparison curves as the other three panels.
POST_NA = 1500          # which posterior file to draw s_mt from

# vanilla range-expansion baseline, matching theta_run.py / scan4_run.py exactly
EXP_BASE = dict(grid=20, K=20, r=2.0, m=0.10, gamma=0.12,
                gens=SAMPLE_GEN + 10, sample_gen=SAMPLE_GEN,
                stop_after_ext=0, record_every=100)


def load_posterior(na=POST_NA, rna=1.0):
    """Draws of the mtDNA fitness gap, preferring the RNA-INCLUSIVE pooled posterior.

    CHANGED 2 Oct 2026, and this was a real error in everything built from these runs.
    Until now the only file this looked for was mt_gap_posterior_NA<NA>.txt, which is the
    CODING-ONLY ABC posterior: 543 accepted draws, median 0.235%, mean 0.252%. Section 1
    headlines the RNA-inclusive posterior -- 11,698 accepted draws, median 0.320%, mean
    0.386% -- and that is the one every other figure now uses. The production joint2 sweep
    ran with --rna 1.0, so its recorded s_mt_mean is 0.2528%: the coding-only mean, with no
    correction. Figure 2b's curves were therefore drawn with an mtDNA advantage about a
    third too small, and since p_joint can never exceed the mtDNA's own fixation
    probability, its unbiased arms were capped at roughly 18% where they should approach
    24%. That is why the unbiased migration pulse plateaued near 15% instead of ~24%.

    The pooled file has two columns -- gap and the N_A the draw came from. The N_A column is
    ignored here: these runs are at a single census, so there is no size to pair against,
    and pooling is what the main-text and Extended Data figures do.

    The old file is still accepted as a fallback so that an archive extracted without the
    pooled draws still runs, but it warns loudly, because silently reverting to it is
    exactly the failure this docstring exists to record.
    """
    here = os.path.dirname(os.path.abspath(__file__))
    names = ["mt_gap_posterior_pooled_rna.txt",
             f"mt_gap_posterior_NA{na}_rna.txt",
             f"mt_gap_posterior_NA{na}.txt"]
    for nm in names:
        for p in (nm, os.path.join(here, nm), os.path.join(here, "..", nm)):
            if os.path.exists(p):
                a = np.loadtxt(p, comments="#")
                g = a[:, 0] if a.ndim == 2 else a
                if g.size == 0:
                    raise SystemExit(f"{p} contains no draws")
                if nm.endswith(f"NA{na}.txt"):
                    sys.stderr.write(
                        f"WARNING: falling back to the SUPERSEDED coding-only posterior "
                        f"{nm} (median 0.235%). Section 1 headlines the RNA-inclusive "
                        f"posterior (median 0.320%); results will understate mtDNA "
                        f"retention. Copy mt_gap_posterior_pooled_rna.txt alongside this "
                        f"script to use it.\n")
                return g * rna
    raise SystemExit("no mt_gap_posterior file found -- expected one of: "
                     + ", ".join(names))


def main():
    args = [a for a in sys.argv[1:] if not a.startswith("--")]
    scenario, sweep, value, Narg = args[0], args[1], float(args[2]), args[3]
    reps, seed0, outfile = int(args[4]), int(args[5]), args[6]
    rna = 1.0
    if "--rna" in sys.argv:
        rna = float(sys.argv[sys.argv.index("--rna") + 1])
    # Calibration overrides. Autosomal ancestry is NOT automatically 5% once theta or s is
    # switched on: reducing F1 male fertility removes the archaic autosomes those fathers
    # carried (so modern ancestry rises in an expansion, and falls under an all-female pulse),
    # and strong selection on the Y removes archaic-Y males along with their autosomes. To
    # keep the four series comparable, gamma (expansion) or f (stable boundary) is retuned per
    # condition so that autosomal ancestry lands at 5%. These flags set the value to use.
    gamma_ov = float(sys.argv[sys.argv.index("--gamma") + 1]) if "--gamma" in sys.argv else None
    # --xcost c: the fertility cost paid by a drive-EXPRESSING male (modern X driver plus
    # archaic Y), offspring removed with probability c. Only meaningful with sweep=drive.
    # Figure S3.4(a) needs both arms of the ladder, and the note's 10% is the arbitrary
    # allowance for destroying half of one's sperm.
    xcost = float(sys.argv[sys.argv.index("--xcost") + 1]) if "--xcost" in sys.argv else 0.0
    f_ov = float(sys.argv[sys.argv.index("--f") + 1]) if "--f" in sys.argv else None
    # Expansion geometry. The published runs used grid = 20, K = 20, whose post-wave census is
    # ~8000 (uniparental Ne ~2000-4000) -- five times the post-mixture size the rest of the
    # analysis assumes (Muller's Ratchet v32: NN-uniparental = 725, diploid 1500), and that
    # size difference, not the geography, is what let the expansion beat the stable boundary.
    # These flags shrink the grid to match. K must stay >= 7: a deme makes no offspring if it
    # lacks either sex, a Poisson(K/2) = 0 event, and below K = 7 that failure rate (>10%)
    # collapses the archaic population before it can be absorbed, leaving autosomal ancestry
    # stuck above 80% modern at ANY gamma.
    grid_ov = int(sys.argv[sys.argv.index("--grid") + 1]) if "--grid" in sys.argv else None
    K_ov = int(sys.argv[sys.argv.index("--K") + 1]) if "--K" in sys.argv else None
    # Per-generation migration probability per cardinal direction. Shrinking the grid at fixed
    # physical landscape size makes each deme wider, so preserving the real dispersal distance
    # (and hence the physical wave speed) means preserving D ~ m * w^2 with w ~ 1/grid, i.e.
    # m scaling as grid^2: m = 0.10 * (grid/20)^2. Lower m also raises Ne by strengthening
    # spatial structure, which works against the whole point of matching the post-mixture size,
    # so it is swept rather than assumed.
    m_ov = float(sys.argv[sys.argv.index("--m") + 1]) if "--m" in sys.argv else None
    # r, the modern:archaic lineage fitness ratio. Raising r LOWERS final modern autosomal
    # ancestry (a faster lineage sweep pulls more archaic genomes into the modern lineage per
    # cross mating), so it trades against gamma: higher r lets the same 5% target be hit at a
    # lower, more biologically natural assortative-mating discount.
    r_ov = float(sys.argv[sys.argv.index("--r") + 1]) if "--r" in sys.argv else None
    # Carrying capacity during the expansion, before it contracts to K. A single constant K
    # forces the expansion-phase size and the post-expansion Ne to be the same number, but they
    # do different jobs: the first sets how many modern x archaic contact events the wave gets,
    # the second sets how effective the mtDNA fitness gap is. Matching the post-mixture Ne
    # therefore starves the expansion, which is what makes the theta mechanism unreachable.
    ke_ov = int(sys.argv[sys.argv.index("--K-early") + 1]) if "--K-early" in sys.argv else None
    keg_ov = (int(sys.argv[sys.argv.index("--K-early-gens") + 1])
              if "--K-early-gens" in sys.argv else None)
    if K_ov is not None and K_ov < 7:
        raise SystemExit(f"--K {K_ov} is below the K >= 7 floor: at K < 7 the deme "
                         f"reproductive-failure rate collapses the archaic population")
    if scenario not in SCENARIOS:
        raise SystemExit(f"scenario must be one of {SCENARIOS}")
    if sweep not in SWEEPS:
        raise SystemExit(f"sweep must be one of {SWEEPS}")
    os.makedirs(os.path.dirname(outfile) or ".", exist_ok=True)

    post = load_posterior(rna=rna)
    # One independent posterior draw per replicate. The stream is seeded from the condition
    # and from seed0, via crc32 rather than hash(): Python randomises string hashing per
    # process, so hash() would give different draws on every run and the results would not be
    # reproducible. Keying on seed0 as well means separate batches of the same condition draw
    # different s_mt values instead of all repeating the first batch.
    tag = f"{scenario}|{sweep}|{value:.10g}|{seed0}".encode()
    draw_rng = np.random.default_rng(zlib.crc32(tag))
    s_mt_all = draw_rng.choice(post, size=reps, replace=True)

    with open(outfile, "w") as fh:
        fh.write("seed\ts_mt\tmt\tY\tauto\tX\textgen\n")
        for k, seed in enumerate(range(seed0, seed0 + reps)):
            s_mt = float(s_mt_all[k])
            if scenario in EXPANSIONS:
                # X-LINKED DRIVE LIVES IN spatial_sim3, NOT spatial_sim2. The s and theta
                # sweeps have always used sim2, which has no xdrive field at all -- passing
                # one would be silently dropped by Config and the run would come back as a
                # no-mechanism control that looks like "drive does nothing". sim3 is sim2 plus
                # the restorer / X-drive / escape machinery, so the two agree exactly where
                # the mechanism is off.
                if sweep == "drive":
                    import spatial_sim3 as S
                else:
                    import spatial_sim2 as S
                kw = dict(EXP_BASE, female_bias=FEMALE_BIAS[scenario], s_mt=s_mt)
                if gamma_ov is not None:
                    kw["gamma"] = gamma_ov
                if grid_ov is not None:
                    kw["grid"] = grid_ov
                if K_ov is not None:
                    kw["K"] = K_ov
                if m_ov is not None:
                    kw["m"] = m_ov
                if r_ov is not None:
                    kw["r"] = r_ov
                if ke_ov is not None:
                    kw["K_early"] = ke_ov
                if keg_ov is not None:
                    kw["K_early_gens"] = keg_ov
                if sweep == "s":
                    kw.update(s=value, sel_locus="Y")
                elif sweep == "drive":
                    # `value` is cfg.xdrive = d, the probability that a Y-bearing conception
                    # of an expressing male is CONVERTED to a daughter -- not the daughter
                    # proportion itself. Proportion daughters = 0.5 + 0.5*d, so the ladder
                    # d = 0.05/0.1/0.2/0.4/0.6/0.8/1.0 is the 52.5/55/60/70/80/90/100% that
                    # Figure 2's Route 2 axis and Table S3.2 report, and d = 0 is no
                    # distortion at all. The submitter converts; do not double-convert here.
                    kw.update(s=0.0, sel_locus="none", xdrive=value, xdrive_cost=xcost)
                else:
                    kw.update(s=0.0, sel_locus="none", theta_f1m=value)
                out, ext = S.run(replace(S.Config(**kw), seed=seed))
                i = int(np.argmin(np.abs(out["gen"] - SAMPLE_GEN)))
                mt, Y = out["mt_mod"][i], out["Y_mod"][i]
                au, xx = out["auto_mod"][i], out["X_mod"][i]
                eg = int(ext) if ext is not None else -1
            else:
                import panmictic_sim as PS
                N = int(Narg)
                kw = dict(N=N, f=(0.05 if f_ov is None else f_ov), s_mt=s_mt,
                          male_frac=(0.0 if scenario == "fsb" else 0.5))
                if sweep == "drive":
                    # IMPLEMENTED 14 Sep 2026. This branch used to raise: panmictic_sim had no
                    # X-linked drive, so Route 2 of Figure 2b could say nothing about either
                    # pulse scenario and its black curve was simply absent. DR: "where is the
                    # black curve at route 2?" panmictic_sim now carries xdrive / xdrive_cost
                    # with the same semantics as spatial_sim3 (see the Config comment there
                    # for why it is a father weight here and a sex flip in the spatial engine).
                    #
                    # fsb IS STILL NOT SIMULATED, AND STILL SHOULD NOT BE. Under an all-female
                    # pulse no modern male ever enters, so the modern Y starts at zero and
                    # stays there; p_joint is identically zero at every d, analytically. A run
                    # would spend CPU confirming an identity and would report a noisy zero.
                    if scenario == "fsb":
                        raise SystemExit(
                            "joint_run: sweep 'drive' is analytically zero for fsb -- no "
                            "modern male enters an all-female pulse, so there is no modern Y "
                            "for a driver to favour. Figure 2b draws that line without a run.")
                    kw.update(s=0.0, sel_locus="none", xdrive=value, xdrive_cost=xcost)
                elif sweep == "twopulse":
                    # ROUTE 4 AT A STABLE BOUNDARY. `value` is the generation of the second
                    # pulse. The sex bias of each pulse is the scenario's own: fsb is
                    # all-female twice, nsb is all-female then ALL-MALE -- the stable-boundary
                    # reading of "matrilocal then patrilocal", which is the comparison Route 4
                    # actually needs. Two unbiased pulses deliver more of the same and cannot
                    # produce the asymmetry the route turns on.
                    #
                    # f IS HALVED, ANALYTICALLY RATHER THAN BY CALIBRATION. Two pulses of
                    # fraction f leave 2f - f^2 modern ancestry, so hitting the 5% anchor
                    # needs f = 1 - sqrt(0.95) = 0.02532 in each. Measured across the ladder
                    # that lands at 5.1-5.9%, inside the 3-7% window at every rung, so no
                    # probe grid is required -- unlike the expansions, where gamma has to be
                    # retuned per rung because ancestry is built up at a moving front.
                    f2 = (1.0 - (1.0 - 0.05) ** 0.5) if f_ov is None else f_ov
                    kw.update(s=0.0, sel_locus="none", f=f2, male_frac=0.0,
                              pulse2_gen=int(value), pulse2_f=f2,
                              pulse2_male_frac=(0.0 if scenario == "fsb" else 1.0))
                elif sweep == "mtydmi":
                    # THE mtDNA x Y INCOMPATIBILITY AT A STABLE BOUNDARY. Figure 2b's mtDNA x
                    # Y column carried a fountain and no comparison scenarios; this is the
                    # pulse arm of them. `value` is the penetrance, so 1.0 is the lethal
                    # single-pair DMI the column is drawn for.
                    #
                    # fsb IS ANALYTIC AND NOT RUN, for the third time in this file and for the
                    # same reason: an all-female pulse admits no modern male, so there is no
                    # modern Y for the incompatibility to act on and p_joint is identically
                    # zero however penetrant the DMI is.
                    if scenario == "fsb":
                        raise SystemExit(
                            "joint_run: sweep 'mtydmi' is analytically zero for fsb -- an "
                            "all-female pulse never introduces a modern Y. Figure 2b draws "
                            "that marker without a run.")
                    kw.update(s=0.0, sel_locus="none", dm=value,
                              dm_focal="mt", dm_partner="Y")
                elif sweep == "s":
                    kw.update(s=value, sel_locus=("none" if value == 0 else "Y"))
                else:
                    kw.update(s=0.0, sel_locus="none", f1_male_theta=value)
                r = PS.run(replace(PS.Config(**kw), seed=seed))
                mt, Y, au, xx, eg = r["mt_mod"], r["Y_mod"], r["auto_mod"], r["X_mod"], -1
            fh.write(f"{seed}\t{s_mt:.8g}\t{mt:.6g}\t{Y:.6g}\t{au:.6g}\t{xx:.6g}\t{eg}\n")
    print("wrote", outfile)


if __name__ == "__main__":
    main()
```
