## Supplementary Information section 3 code base for "Hypothesis: A modern human range expansion ∼300,000 years ago had a critical role in Neandertal origins": joint_submit.py

```
#!/usr/bin/env python3
"""
joint_submit.py -- SLURM array for joint_run.py (series b, c and d of main-text Figure 1B).

Usage:  python joint_submit.py <reps> <batch> [concurrency] [partition] [--resume]
                               [--only nexp|fsb|nsb]
        python joint_submit.py 1000 25 500 short
then:   sbatch joint_array.sh          (output -> out_joint/)

The two stable-boundary scenarios are cheap (a few seconds per replicate) and the expansion is
not (about a minute), so batch sizes are set per scenario rather than globally: the expansion
uses the batch given, the stable boundary uses four times it.
"""
import os
import sys

from joint_run import SVALS, THETAS, SCENARIOS

# SUPERSEDED 5 Sep 2026: this was 1500, which predates the Peyregne anchor. joint2/jcal are
# the live chain; this script is kept only for provenance. Do not resubmit from it.
from anchors import PANM_CENSUS as SB_N   # 1312; see anchors.py


def main():
    args   = [a for a in sys.argv[1:] if not a.startswith("--")]
    total  = int(args[0]) if len(args) > 0 else 1000
    batch  = int(args[1]) if len(args) > 1 else 25
    cap    = int(args[2]) if len(args) > 2 else 500
    part   = args[3] if len(args) > 3 else "short"
    resume = "--resume" in sys.argv
    only   = (sys.argv[sys.argv.index("--only") + 1] if "--only" in sys.argv else None)
    scen   = [s for s in SCENARIOS if only is None or s == only]
    if not scen:
        raise SystemExit(f"--only must name one of {SCENARIOS}")
    os.makedirs("out_joint", exist_ok=True); os.makedirs("logs", exist_ok=True)

    tasks, skipped = [], 0
    for sc in scen:
        b = batch if sc == "nexp" else batch * 4
        Narg = "-" if sc == "nexp" else str(SB_N)
        for sweep, vals in (("s", SVALS), ("theta", THETAS)):
            for v in vals:
                for s0 in range(0, total, b):
                    reps = min(b, total - s0)
                    out = f"out_joint/out_{sc}_{sweep}_{v:g}_s{s0}.txt"
                    if resume and os.path.exists(out) and sum(1 for _ in open(out)) >= reps + 1:
                        skipped += 1; continue
                    tasks.append(f"{sc} {sweep} {v} {Narg} {reps} {s0} {out}")
    if not tasks:
        print("nothing to do -- every output file is already complete"); return
    open("joint_params.txt", "w").write("\n".join(tasks) + "\n")
    n = len(tasks); cap = min(n, cap)
    open("joint_array.sh", "w").write(f"""#!/bin/bash
#SBATCH -J joint
#SBATCH -p {part}
#SBATCH -t 0-11:59
#SBATCH -c 1
#SBATCH --mem=4G
#SBATCH -o logs/joint_%A_%a.log
#SBATCH --array=0-{n - 1}{("%%%d" % cap) if cap else ""}

LINE=$(sed -n "$((SLURM_ARRAY_TASK_ID + 1))p" joint_params.txt)
echo "task $SLURM_ARRAY_TASK_ID : $LINE"
python joint_run.py $LINE
""")
    ncond = len(scen) * (len(SVALS) + len(THETAS))
    print(f"wrote joint_params.txt ({n} tasks over {ncond} conditions), {total} reps each")
    if resume: print(f"  RESUME: skipped {skipped} complete tasks")
    print(f"  scenarios: {', '.join(scen)}")
    print("submit:  sbatch joint_array.sh")


if __name__ == "__main__":
    main()
```
