## Supplementary Information section 3 code base for "Hypothesis: A modern human range expansion ∼300,000 years ago had a critical role in Neandertal origins": mtexp_submit.py

```
#!/usr/bin/env python3
"""
mtexp_submit.py -- extend Figure S3.2's range-expansion traces to the pulse's full s ladder.
(12 Sep 2026)

DR: "Could you extend the range expansion series for this figure to the full set of selection
coefficients run for the migration pulse?"

WHY THIS IS A NEW RUN AND NOT A QUERY. The existing dashed traces come from the residence axis
of the `demog` screen, whose SMT_LADDER stops at s = 0.64% because it was built from the
quartiles of the ABC posterior, not to span a figure. There is nothing on disk above 0.64%.

AND WHY IT IS CHEAP ANYWAY -- ~80 CPU-HOURS, NOT ~750. The temptation is to extend `demog`
itself, pooling thousands of admissible demographic combinations per rung as the current traces
do. That would cost about 750 CPU-hours for ten more rungs, and it would also be the wrong
comparison: the SOLID pulse curves are not pooled over a grid, they are one baseline demography
at 1,000 (deme) and 400 (band) replicates per point. Matching that -- one baseline, many
replicates, the same fifteen s values -- is both apples-to-apples and two orders of magnitude
cheaper. The pooled 5-rung version stays in the CSV as a cross-check; where the two overlap they
should agree within sampling error, and if they do not, the pooled trace was being driven by
which combinations happen to be admissible rather than by s.

BASELINE, AND PATRILOCAL. robust_run's defaults are already the note's baseline (grid 12,
K 11, m 0.10, r 2 for the deme; N 1,750, K 25, p_move 0.45 for the band), so the only flags
passed are the residence rule, s_mt, and the recalibrated interbreeding rate explained at PLAN
below. Under full MATRILOCALITY the mtDNA is already 100% modern by construction and the
question is vacuous; under PATRILOCALITY the residence rule carries the Y instead, and the
mtDNA has to be rescued by selection and luck alone. That is the scenario this panel is about.

NEGATIVE s IS SIMULATED, NOT CLIPPED. Both engines branch on the sign of s_mt (spatial_sim2
line 511, bandsim line 535) and select against the modern type when it is negative, so the
three negative rungs of the pulse ladder are honest points rather than forced zeros.

    python mtexp_submit.py            # writes the array and submits it
    python mtexp_submit.py --no-go    # write only
"""
import os
import subprocess
import sys

HERE = os.path.dirname(os.path.abspath(__file__))

# Exactly the pulse ladder inside Figure S3.2's x range, so the dashed traces land on the same
# abscissae as the solid ones and the eye can read them off against each other.
# EXTENDED RIGHT TO 20%. DR: "We should extend the selection coefficient series to the right,
# I want to see the probability of modern mtDNA fixing start to climb." Under a PATRILOCAL
# expansion the mtDNA is the mirror of Route 1's Y, and Route 1 needed s ~ 4% (band) to reach
# even odds, so nothing was ever going to move below 3%. The first fifteen rungs are the pulse
# ladder, for overlay; the last five carry the curve up to where it has to climb.
S_LADDER = [-0.005, -0.002, -0.001, 0.0, 0.0005, 0.001, 0.002, 0.003,
            0.005, 0.0075, 0.01, 0.015, 0.02, 0.025, 0.03,
            0.04, 0.05, 0.075, 0.10, 0.20]

# Replicates per point, matching the pulse curves, split into batches so no task runs long.
#
# PATRILOCAL, NOT UNBIASED -- CORRECTED. The first run used female_bias 0.5 / beta 0.5, which is
# the NON-sex-biased expansion. DR: "Is this a patrilocal or matrilocal range expansion we're
# simulating here? For this version it should be a patrilocal expansion." He is right, and it is
# the whole point of the panel: under patrilocality the residence rule carries the Y for free and
# the mtDNA is the marker that has to be rescued, which is the question Figure S3.2 asks. Under
# no sex bias neither marker is carried, so the answer was about a different scenario.
#
# female_bias 0.0 is patrilineal in spatial_sim2 (1.0 matrilineal, 0.5 unbiased); beta 1.0 is
# patrilocal in bandsim (0 matrilocal, so only males disperse).
#
# RECALIBRATED FOR IT, because the interbreeding rate that lands on 5% autosomal moves again.
# Measured at 20 replicates, s_mt = 0:
#     deme   gamma 0.12 -> 7.57%   0.16 -> 6.43%   0.22 -> 4.45%    => gamma* ~ 0.20
#     band   e     0.13 -> 6.22%   0.19 -> 3.17%   0.26 -> 1.54%    => e*     ~ 0.155
RESIDENCE = {"deme": ("--female-bias", "0.0"), "band": ("--beta", "1.0")}
# BAND RAISED 400 -> 1,000 TO MATCH THE DEME ARM. DR: "everywhere in the note where we have
# different number of reps for bands and demes, we should raise the lower one to the higher
# one." Figure S3.2 drew its dashed deme curve from 1,000 replicates per point and its dashed
# band curve from 400, so the two traces carried different error and the caption had to say so.
# They are now both 1,000. Batches are summed per (framework, s) by collect_mt_expansion, and
# existing batch files are skipped by name, so this re-runs only the missing 600.
PLAN = {"deme": dict(reps=1000, batch=200, extra=["--gamma", "0.20"]),
        "band": dict(reps=1000, batch=200, extra=["--e", "0.155"])}
OUT = "out_mtexp_pat"   # patrilocal; the unbiased run stays in out_mtexp
PART = "short"
CAP = 400


def tag(fw, s, b):
    return f"{fw}_smt{s:g}_b{b}".replace("-", "m").replace(".", "p")


def main():
    os.makedirs(os.path.join(HERE, OUT), exist_ok=True)
    os.makedirs(os.path.join(HERE, "logs"), exist_ok=True)
    tasks, skipped = [], 0
    for fw, p in PLAN.items():
        nb = p["reps"] // p["batch"]
        for s in S_LADDER:
            for b in range(nb):
                t = tag(fw, s, b)
                out = os.path.join(OUT, t + ".json")
                if os.path.exists(os.path.join(HERE, out)):
                    skipped += 1
                    continue
                # mech `none` with value 0: no DMI, no drive, no Y selection -- the only thing
                # acting is s_mt, which is the whole point.
                rflag, rval = RESIDENCE[fw]
                tasks.append(f"python robust_run.py {fw} none 0 1 {p['batch']} "
                             f"{b * p['batch']} {out} {rflag} {rval} --s-mt {s:g} "
                             + " ".join(p["extra"]))
    if not tasks:
        print(f"  nothing to submit ({skipped} cells already on disk)")
        return

    pf, sf = "mtexp_params.txt", "mtexp_array.sh"
    with open(os.path.join(HERE, pf), "w") as f:
        f.write("\n".join(tasks) + "\n")
    with open(os.path.join(HERE, sf), "w") as f:
        f.write(f"""#!/bin/bash
#SBATCH -J mtexp
#SBATCH -p {PART}
#SBATCH -c 1
#SBATCH --mem=2G
#SBATCH -t 0-05:59
#SBATCH -a 0-{len(tasks) - 1}%{CAP}
#SBATCH -o logs/mtexp_%A_%a.log
cd {HERE}
CMD=$(sed -n "$((SLURM_ARRAY_TASK_ID + 1))p" {pf})
echo "$CMD"
eval "$CMD"
""")
    print(f"  {len(tasks)} tasks ({skipped} already on disk)")
    est = sum(200 * (15.0 if t.split()[2] == 'deme' else 11.0) for t in tasks) / 3600
    print(f"  ~{est:.0f} CPU-hours; {len(S_LADDER)} s values x "
          f"{'/'.join(str(v['reps']) for v in PLAN.values())} replicates")
    print(f"  wrote {pf} and {sf}")
    if "--no-go" in sys.argv:
        print(f"  NOT submitted (--no-go).  sbatch {sf}")
        return
    r = subprocess.run(["sbatch", sf], cwd=HERE, capture_output=True, text=True)
    print(f"  SUBMITTED: {r.stdout.strip()}" if r.returncode == 0
          else f"  !! sbatch FAILED: {(r.stderr or '').strip()}")


if __name__ == "__main__":
    main()
```
