## Supplementary Information section 3 code base for "Hypothesis: A modern human range expansion ∼300,000 years ago had a critical role in Neandertal origins": mtypulse_submit.py

```
#!/usr/bin/env python3
"""
mtypulse_submit.py -- the mtDNA x Y incompatibility at a stable boundary. (14 Sep 2026)

DR: "There are no blue-orange-black lines for mtDNA-Y incompatibility." Figure 2b's mtDNA x Y
column carried a fountain and nothing to compare it against. Of the three comparison
scenarios:

  ORANGE  the unbiased expansion, from the `demog` sweep, which already carries dmi_mtY at
          beta = 0.5. No run needed.
  BLUE    the all-female pulse. Analytic and deliberately not run: no modern male enters, so
          there is never a modern Y for the incompatibility to act on and p_joint is
          identically zero however penetrant the DMI is. joint_run raises rather than
          spending CPU to confirm an identity.
  BLACK   the unbiased pulse. This script.

f = 0.091, MEASURED RATHER THAN ASSUMED, and this is the one place in the pulse family where
that was necessary. For a plain pulse the autosomal proportion IS f, so the 5% anchor needs no
calibration; for the two-pulse rows it is 2f - f^2 and the anchor is analytic. Here the DMI
kills modern-mtDNA males and takes their modern autosomes with them, so ancestry ends well
BELOW f and the relation has no closed form. Measured at N = 1312, 40 replicates per rung:

    f = 0.05  ->  2.96%        f = 0.12  ->  6.99%
    f = 0.08  ->  4.23%        f = 0.18  -> 12.66%
                               f = 0.25  -> 20.02%

Interpolating between the two rungs that bracket the target gives f = 0.091, which is what is
used. The value is recorded here rather than in a calibration file because it is one number
for one figure column, and a jcal-style probe grid would be four hundred lines of machinery
to find it.

    python mtypulse_submit.py              # 1 cell x 2,000 replicates, minutes
    python mtypulse_submit.py --no-go
"""
import os
import subprocess
import sys

HERE = os.path.dirname(os.path.abspath(__file__))
sys.path.insert(0, HERE)

from anchors import PANM_CENSUS as SB_N      # 1312

F_STAR = 0.091      # see the docstring; lands autosomal ancestry on ~5%
DM = 1.0            # lethal single-pair DMI, matching the column Figure 2b draws
REPS = 2000
OUT = "out_mtypulse"
PART = "short"


def main():
    reps = int(sys.argv[sys.argv.index("--reps") + 1]) if "--reps" in sys.argv else REPS
    os.makedirs(os.path.join(HERE, OUT), exist_ok=True)
    os.makedirs(os.path.join(HERE, "logs"), exist_ok=True)

    out = os.path.join(OUT, f"nsb_dm{DM:g}".replace(".", "p") + ".tsv")
    full = os.path.join(HERE, out)
    if os.path.exists(full) and max(sum(1 for _ in open(full)) - 1, 0) >= reps:
        print(f"  nothing to do -- already {reps:,} replicates")
        return
    task = (f"python joint_run.py nsb mtydmi {DM:g} {SB_N} {reps} 1000 {out} "
            f"--f {F_STAR}")
    print(f"  1 cell, {reps:,} replicates, N = {SB_N}, f = {F_STAR}")

    pf, sf = "mtypulse_params.txt", "mtypulse_array.sh"
    open(os.path.join(HERE, pf), "w").write(task + "\n")
    open(os.path.join(HERE, sf), "w").write(f"""#!/bin/bash
#SBATCH -J mtypulse
#SBATCH -p {PART}
#SBATCH -t 0-11:59
#SBATCH -c 1
#SBATCH --mem=4G
#SBATCH -a 0-0
#SBATCH -o logs/mtypulse_%A_%a.log
CMD=$(sed -n "$((SLURM_ARRAY_TASK_ID + 1))p" {pf})
echo "$CMD"
eval "$CMD"
""")
    if "--no-go" in sys.argv:
        print(f"  NOT submitted (--no-go).  sbatch {sf}")
        return
    r = subprocess.run(["sbatch", sf], cwd=HERE, capture_output=True, text=True)
    print(f"  SUBMITTED: {r.stdout.strip()}" if r.returncode == 0
          else f"  !! sbatch FAILED: {(r.stderr or '').strip()}")


if __name__ == "__main__":
    main()
```
