## Supplementary Information section 3 code base for "Hypothesis: A modern human range expansion ∼300,000 years ago had a critical role in Neandertal origins": ne_screen.py

```
#!/usr/bin/env python3
"""
ne_screen.py -- which (grid, K, m) geometries actually deliver Ne ~ 1447? (DR, 10 Sep 2026)

DR: "only run parameters that constrain Ne to be close to 1500, and explore varying deme
numbers too. I really want to evaluate all parameters."

Geometry is not a free parameter: 12x12 with K=11 was chosen because its MEASURED autosomal
effective size matches the Peyregne anchor. Other geometries can be admitted, but only after
measuring, because census and Ne are not the same thing -- spatial structure inflates Ne, and
it does so more when there are many small demes and little migration.

WHY THIS IS NOT ne_audit.py
---------------------------
Two reasons, both discovered by reading it rather than trusting the docstring:

  1. `ne_audit.py deme nsb` does NOT measure the deme grid. "nsb" is absent from DEME_FB, so it
     falls through to the PANMICTIC engine. Only "fexp" and "nexp" reach spatial_sim2.
  2. Its documented `--grid` and `--K` flags are never parsed. Geometry is read from the NEGRID
     and NEK environment variables, so the flags are silently ignored and you measure 12x12/K=10
     whatever you type. (Same failure mode as the xdrive kwarg: an unrecognised option that
     changes nothing and reports a plausible number.)

and one reason of substance: ne_audit hardcodes m = 0.10, but migration is precisely what sets
the spatial structure that makes Ne differ from census. It has to be swept.

The ESTIMATOR here is identical to ne_audit's, deliberately: burn in, reset mtDNA over everyone,
the Y over males and all eight autosomal loci to binomial(2, 1/2), then track and fit

    ln E[p(1-p)]  ~  -t / Ne

averaging the fit over the eight unlinked autosomal loci.

WHAT DOES NOT NEED SWEEPING. r and gamma are irrelevant here and are left at their defaults.
Both act only while the two lineages coexist -- r is the lineage advantage and gamma the
assortative-mating discount -- and this measurement is made after burn-in, when the population
is of one lineage. So Ne is a function of (grid, K, m) alone, which is what makes the screen
cheap enough to be worth doing properly.

    python ne_screen.py [--reps 96] [--out ne_screen.csv] [--budget 0]
"""
import csv
import os
import sys
import time
from concurrent.futures import ProcessPoolExecutor

import numpy as np

HERE = os.path.dirname(os.path.abspath(__file__))
sys.path.insert(0, HERE)

BURN, TRACK = 400, 1500
ANCHOR = 1447.0                     # diploid, Peyregne time-weighted harmonic mean 250-50 ka
# Tolerance is set by what the estimator can actually resolve. Measured repeatability at
# grid 12 / K 11 / m 0.08, three independent blocks of 96 replicates: 1477, 1477, 1510 --
# SD 19, i.e. 1.3%. At 24 replicates the same configuration scattered by ~20%, which is why the
# first pilot marked everything KEEP: the tolerance was wider than the noise, so the filter was
# measuring nothing. At 96 replicates a 10% window is ~7 SD wide and genuinely discriminates.
TOL = 0.10                          # keep |Ne/ANCHOR - 1| <= TOL  (1302 - 1592 diploid)

# Geometries spanning few-large-demes to many-small-demes at a census near 1500. For each grid,
# K is the census-matching value and its neighbours, so the screen can find the K that lands on
# the anchor rather than assuming census == Ne.
# DR, 10 Sep: "I want the full cartesian product of independent lists."
#
# So grid and K are now independent ranges rather than K being chosen per grid to hit a target
# census. That deliberately includes combinations that cannot possibly match the anchor -- 6x6
# with K=4 is 144 people, 20x20 with K=42 is 16,800 -- because the point is that the Ne
# constraint is a RESULT of the screen, not an assumption built into the grid. The reported
# "parameter combinations searched" is then the honest full product, and the Ne filter is one of
# the data-matching criteria applied afterwards, alongside autosomal ancestry and timing.
#
# Cost of the waste is acceptable: Ne does not depend on gamma or r (both act only while the two
# lineages coexist, and this is measured after burn-in), so the whole product needs only
# len(GRIDS) x len(KS) x len(MS) measurements, about 10 CPU-hours including the large censuses.
GRIDS = [6, 8, 10, 12, 14, 16, 20]
# K must be a LADDER, not a handful of round numbers. The carrying capacity that reaches the
# anchor differs about tenfold across the grid range -- roughly K = 7 at 16x16 but K = 42 at
# 6x6 -- so a coarse list lands on the admissible band for one or two grids and misses it for
# the rest. The first attempt, [4, 6, 8, 11, 15, 23, 42], retained only 18 of the 47 admissible
# (grid, K, m) triples already measured: the other 29 sat at K values the list simply skipped.
# This ladder steps by about 15%, which is finer than the width of the admissible band at every
# grid size, so the product covers it rather than sampling it.
#
# 11 ADDED 11 Sep 2026, and it is not a rounding choice. K = 11 at 12x12 is the BASELINE deme
# geometry -- the one every production figure and every number in Figure 2b was computed at --
# and ne_screen.csv already measures it as admissible (Ne = 1488 against the 1447 anchor). But
# 11 fell between 10 and 12 on this ladder, so cells() never generated it, and the baseline
# geometry was absent from the Cartesian search. That is fatal to DR's proposed merge: he
# expects the Figure 2b baseline to be "a subset of Figure 2c", and on the deme side it was not
# a member of the set at all. No new measurement is needed; the row is already in the CSV.
KS = [6, 7, 8, 9, 10, 11, 12, 13, 15, 17, 20, 23, 26, 30, 35, 40, 46]
GEOMS = [(g, k) for g in GRIDS for k in KS]
MS = [0.05, 0.08, 0.12, 0.15]


def fit_ne(traj, lo=0.05):
    het = np.asarray(traj) * (1.0 - np.asarray(traj))
    E = np.nanmean(het, axis=0)
    E0 = E[:10].mean()
    good = np.where(E > lo * E0)[0]
    t1 = good[-1] if len(good) else len(E) - 1
    if t1 < 20:
        return float("nan")
    b = np.polyfit(np.arange(t1), np.log(np.maximum(E[:t1], 1e-12)), 1)[0]
    return -1.0 / b if b < 0 else float("inf")


def one(args):
    grid, K, m, seed = args
    import spatial_sim2 as S
    from spatial_sim2 import Config
    cfg = Config(grid=grid, K=K, m=m, gamma=0.12, r=2.0, female_bias=1.0,
                 gens=BURN + TRACK, record_every=10 ** 9, seed=seed)
    rng = np.random.default_rng(cfg.seed)
    P = S._init(cfg, rng)

    def advance(P):
        S._migrate(P, cfg, rng)
        nxt = S._repro(P, cfg, rng)
        return None if nxt is None else S._reseed_origin(nxt, cfg)

    for _ in range(BURN):
        P = advance(P)
        if P is None:
            return None
    P["mt"] = rng.integers(0, 2, len(P["mt"])).astype(np.int8)
    male = P["sex"] == 1
    P["Y"] = np.where(male, np.where(rng.random(len(P["Y"])) < 0.5, 1, -1), -1).astype(np.int8)
    P["auto"][:, :] = rng.binomial(2, 0.5, P["auto"].shape).astype(np.int8)

    pm, py, pa = [], [], []
    for _ in range(TRACK):
        P = advance(P)
        if P is None:
            return None
        male = P["sex"] == 1
        pm.append(float((P["mt"] == 1).mean()))
        py.append(float((P["Y"][male] == 1).mean()) if male.any() else np.nan)
        pa.append(list(P["auto"].mean(axis=0) / 2.0))
    return pm, py, pa


def flag(name, cast, default):
    return cast(sys.argv[sys.argv.index(name) + 1]) if name in sys.argv else default


OUTDIR = os.path.join(HERE, "out_ne_screen")


def measure(g, k, m, reps, workers=4):
    """One configuration -> a result row, or None if every replicate died."""
    with ProcessPoolExecutor(workers) as ex:
        res = [r for r in ex.map(one, [(g, k, m, 7000 + i) for i in range(reps)]) if r]
    if not res:
        return None
    pm = np.array([r[0] for r in res]); py = np.array([r[1] for r in res])
    pa = np.array([r[2] for r in res])
    ne_a = float(np.mean([fit_ne(pa[:, :, j]) for j in range(pa.shape[2])]))
    row = dict(grid=g, K=k, m=m, census=g * g * k, reps=len(res),
               ne_auto_diploid=round(ne_a / 2, 1),
               ne_mt=round(fit_ne(pm), 1), ne_Y=round(fit_ne(py), 1))
    row["ratio"] = round(row["ne_auto_diploid"] / ANCHOR, 3)
    row["keep"] = int(abs(row["ratio"] - 1) <= TOL)
    return row


def cfg_name(g, k, m):
    return f"ne_g{g}_K{k}_m{m:g}".replace(".", "p") + ".json"


def worker():
    """--only g,k,m : measure one configuration and write its own file. Array-task safe."""
    import json
    g, k, m = sys.argv[sys.argv.index("--only") + 1].split(",")
    g, k, m = int(g), int(k), float(m)
    reps = flag("--reps", int, 96)
    os.makedirs(OUTDIR, exist_ok=True)
    out = os.path.join(OUTDIR, cfg_name(g, k, m))
    if os.path.exists(out):
        print(f"  {os.path.basename(out)} exists, skipping")
        return
    row = measure(g, k, m, reps, workers=int(os.environ.get("NEPROCS", 1)))
    if row is None:
        print(f"  grid {g} K {k} m {m}: every replicate died")
        return
    with open(out, "w") as f:
        json.dump(row, f)
    print(f"  grid {g:2d} K {k:3d} m {m:.2f}  census {row['census']:5d}  "
          f"Ne_auto {row['ne_auto_diploid']:7.0f}  ratio {row['ratio']:.2f}  "
          f"{'KEEP' if row['keep'] else 'drop'}")


def collect():
    import glob
    import json
    rows = [json.load(open(p)) for p in sorted(glob.glob(os.path.join(OUTDIR, "*.json")))]
    if not rows:
        raise SystemExit("no per-config output in out_ne_screen/")
    rows.sort(key=lambda r: (r["grid"], r["K"], r["m"]))
    fn = os.path.join(HERE, "ne_screen.csv")
    with open(fn, "w", newline="") as f:
        w = csv.DictWriter(f, fieldnames=list(rows[0]))
        w.writeheader(); w.writerows(rows)
    nk = sum(r["keep"] for r in rows)
    print(f"  wrote ne_screen.csv: {len(rows)} configurations, {nk} within "
          f"{TOL:.0%} of the {ANCHOR:.0f} anchor")
    return rows


def main():
    if "--list" in sys.argv:
        for (g, k) in GEOMS:
            for m in MS:
                print(f"{g},{k},{m:g}")
        return
    if "--only" in sys.argv:
        return worker()
    if "collect" in sys.argv:
        return collect()
    reps = flag("--reps", int, 96)
    outfn = os.path.join(HERE, flag("--out", str, "ne_screen.csv"))
    budget = flag("--budget", int, 0)

    done = {}
    if os.path.exists(outfn):
        with open(outfn) as f:
            for r in csv.DictReader(f):
                done[(int(r["grid"]), int(r["K"]), float(r["m"]))] = r

    todo = [(g, k, m) for (g, k) in GEOMS for m in MS if (g, k, m) not in done]
    print(f"  {len(GEOMS)} geometries x {len(MS)} migration rates = {len(GEOMS)*len(MS)} "
          f"configurations; {len(done)} already measured, {len(todo)} to go")
    if not todo:
        print("  nothing left")
    rows = list(done.values())
    t0 = time.time()
    with ProcessPoolExecutor(4) as ex:
        for (g, k, m) in todo:
            res = [r for r in ex.map(one, [(g, k, m, 7000 + i) for i in range(reps)]) if r]
            if not res:
                print(f"  grid {g:2d} K {k:3d} m {m:.2f}: every replicate died")
                continue
            pm = np.array([r[0] for r in res]); py = np.array([r[1] for r in res])
            pa = np.array([r[2] for r in res])
            ne_a = float(np.mean([fit_ne(pa[:, :, j]) for j in range(pa.shape[2])]))
            row = dict(grid=g, K=k, m=m, census=g * g * k, reps=len(res),
                       ne_auto_diploid=round(ne_a / 2, 1),
                       ne_mt=round(fit_ne(pm), 1), ne_Y=round(fit_ne(py), 1))
            row["ratio"] = round(row["ne_auto_diploid"] / ANCHOR, 3)
            row["keep"] = int(abs(row["ratio"] - 1) <= TOL)
            rows.append(row)
            print(f"  grid {g:2d} K {k:3d} m {m:.2f}  census {row['census']:5d}  "
                  f"Ne_auto {row['ne_auto_diploid']:7.0f}  ratio {row['ratio']:.2f}  "
                  f"{'KEEP' if row['keep'] else 'drop'}")
            with open(outfn, "w", newline="") as f:
                w = csv.DictWriter(f, fieldnames=list(rows[0]))
                w.writeheader(); w.writerows(rows)
            if budget and time.time() - t0 > budget:
                print(f"  budget reached; {len(todo)-len(rows)+len(done)} left, run again")
                break
    nk = sum(int(r["keep"]) for r in rows)
    print(f"\n  {len(rows)} measured, {nk} within {TOL:.0%} of the {ANCHOR:.0f} anchor")
    print(f"  wrote {os.path.basename(outfn)}")


if __name__ == "__main__":
    main()
```
