## Supplementary Information section 3 code base for "Hypothesis: A modern human range expansion ∼300,000 years ago had a critical role in Neandertal origins": nexp_band_submit.py

```
#!/usr/bin/env python3
"""
nexp_band_submit.py -- the band arm of the UNBIASED expansion, so the orange curve in
Figure 2b can be the mean of its own fountains. (14 Sep 2026)

WHY THIS EXISTS. Figure 2b's red curve is now the mean of the admissible combinations drawn
beneath it rather than a single calibrated parameter set (see figure2_merged.spine). DR:
"Fountains are points that fall in our admissible box, and the red curves are the averages of
our fountains" -- and then, correctly: "this is being inconsistent because the yellow curve
would then need to be average of its fountains and we haven't made yellow curve fountains".

He is right, with one qualification. A Cartesian sweep for the unbiased expansion DOES exist:
the `demog` set, which crosses the mechanisms with the residence rule and the mtDNA-selection
ladder precisely so that Figure 2b's demographic scenarios sit inside the same search as the
fountains. Its DEME arm is scored -- 23,092 cells in out_demog2_deme. Its BAND arm was
screened (91,854 cells in out_demog1_band) and never scored, so there is nothing to average.
This script scores it.

WHY NOT SIMPLY `ROUTE_SET=demog python routes_submit.py 2 --band`. That emits 26,476 cells at
about 25,000 CPU-hours, and most of them are not needed here:

    by residence   beta 0    (matrilocal)   7,073     already covered by the red fountains
                   beta 0.5  (unbiased)    19,403     what the orange curve needs
    by mtDNA gap   s_mt 0                 11,023     an unbiased expansion with no mtDNA
                                                     selection is a martingale; the curve is
                                                     flat at the autosomal proportion by
                                                     construction and needs no search
                   s_mt 0.0011 .. 0.0064  15,453     the posterior ladder, four extra rungs

Fixing s_mt at 0.00235 -- the median of the Muller's-ratchet posterior, the value the note
quotes -- leaves 3,920 cells and about 3,700 CPU-hours. Every one of them is beta 0.5, because
the matrilocal arm was only ever run at s_mt = 0.

THE ASYMMETRY THIS INTRODUCES IS REAL AND MUST BE IN THE CAPTION. The red cloud is at
s_mt = 0 and the orange cloud will be at s_mt = 0.00235. That is not a bookkeeping slip, it is
the note's own argument -- an unbiased expansion cannot retain modern mtDNA without selection
on it, while a matrilocal one carries its own mtDNA by construction -- but a reader comparing
two clouds is entitled to be told that one of them has a mechanism the other does not.

    python nexp_band_submit.py            # enumerate, price, submit
    python nexp_band_submit.py --no-go    # enumerate and price only
"""
import math
import os
import re
import subprocess
import sys

HERE = os.path.dirname(os.path.abspath(__file__))

S_MT = "0.00235"          # posterior median; see the docstring
CHUNK = 3                 # cells per array task, as routes_submit uses for this set
SEC_PER_CELL = 3600 * 0.944     # measured: 25,005 CPU-h over 26,476 cells
PART, MEM, TIME = "short", "3G", "0-11:59"
STEM = "nexpband"


def newest(pattern):
    fs = [f for f in os.listdir(HERE) if re.fullmatch(pattern, f)]
    if not fs:
        return None
    return max(fs, key=lambda f: os.path.getmtime(os.path.join(HERE, f)))


def main():
    # ENUMERATE FRESH RATHER THAN REUSING A params FILE LYING AROUND. A stale one would be
    # from a different screen state and would re-run cells already scored, or miss new ones.
    env = dict(os.environ, ROUTE_SET="demog")
    r = subprocess.run([sys.executable, "routes_submit.py", "2", "--band", "--no-go"],
                       cwd=HERE, env=env, capture_output=True, text=True)
    if r.returncode != 0:
        raise SystemExit(f"routes_submit failed:\n{r.stderr or r.stdout}")
    src = newest(r"demog2b_\d+_params\.txt")
    if src is None:
        print("  nothing to do -- routes_submit emitted no cells, so every demog band cell "
              "that passed the screen is already scored")
        return

    cells = [ln.strip() for ln in open(os.path.join(HERE, src)) if ln.strip()]
    want = [c for c in cells if f"--s-mt {S_MT}" in c]
    # A guard, not decoration: if the flag were ever renamed or the ladder re-spaced this
    # filter would silently match nothing and submit an empty array that looks like success.
    if not want:
        raise SystemExit(f"no cells matched '--s-mt {S_MT}' among {len(cells):,} -- has the "
                         f"s_mt ladder or the flag name changed?")
    other = len(cells) - len(want)
    ntask = math.ceil(len(want) / CHUNK)
    print(f"  {len(cells):,} demog band cells pending in total")
    print(f"  {len(want):,} at s_mt = {S_MT} (the orange fountains); {other:,} left for later")
    print(f"  {ntask:,} array tasks of {CHUNK}, ~{len(want) * SEC_PER_CELL / 3600:,.0f} CPU-hours")

    pf, sf = f"{STEM}_params.txt", f"{STEM}_array.sh"
    open(os.path.join(HERE, pf), "w").write("\n".join(want) + "\n")
    open(os.path.join(HERE, sf), "w").write(f"""#!/bin/bash
#SBATCH -J {STEM}
#SBATCH -p {PART}
#SBATCH -t {TIME}
#SBATCH -c 1
#SBATCH --mem={MEM}
#SBATCH -a 0-{ntask - 1}
#SBATCH -o logs/{STEM}_%A_%a.log
CHUNK={CHUNK}
START=$((SLURM_ARRAY_TASK_ID * CHUNK + 1))
END=$((START + CHUNK - 1))
sed -n "${{START}},${{END}}p" {pf} | while read -r LINE; do
  [ -z "$LINE" ] && continue
  python robust_run.py $LINE
done
""")
    os.makedirs(os.path.join(HERE, "logs"), exist_ok=True)
    if "--no-go" in sys.argv:
        print(f"  NOT submitted (--no-go).  sbatch {sf}")
        return
    s = subprocess.run(["sbatch", sf], cwd=HERE, capture_output=True, text=True)
    print(f"  SUBMITTED: {s.stdout.strip()}" if s.returncode == 0
          else f"  !! sbatch FAILED: {(s.stderr or '').strip()}")


if __name__ == "__main__":
    main()
```
