## Supplementary Information section 3 code base for "Hypothesis: A modern human range expansion ∼300,000 years ago had a critical role in Neandertal origins": panm4_collect.py

```
#!/usr/bin/env python3
"""
panm4_collect.py -- mean modern ancestry at each locus against the sex bias of the gene flow,
for the stable-boundary model. Writes sexbias_stable_boundary.csv.
"""
import os, glob
import numpy as np
from panm4_run import MFRACS, NVALS, PULSE

COL = dict(seed=0, mt=1, Y=2, auto=3, X=4)


def pool(mf, N):
    rows = []
    for f in glob.glob(f"out_panm4/out_{mf:g}_{N}_s*.txt"):
        for line in open(f).read().strip().split("\n")[1:]:
            p = line.split("\t")
            if len(p) >= 5:
                rows.append([float(x) for x in p])
    return np.array(rows) if rows else None


def main():
    with open("sexbias_stable_boundary.csv", "w") as fh:
        fh.write("# Stable-boundary model: single instantaneous pulse, modern fraction "
                 f"{PULSE:g}, scored at the sampling generation.\n"
                 "# male_frac = fraction of the MODERN CONTRIBUTION that is male "
                 "(0 = all gene flow through women).\n"
                 "# female_frac = 1 - male_frac. census_N is diploid individuals; the "
                 "uniparental effective size is census_N/2.\n"
                 "census_N,uniparental_Ne,male_frac,female_frac,n,auto_pct,mt_pct,Y_pct,"
                 "auto_se,mt_se,Y_se,P_mt_fixed,P_Y_fixed,P_both_fixed\n")
        print(f"{'N':>6} {'male_frac':>9} {'n':>5} {'auto%':>7} {'mtDNA%':>8} {'Y%':>7}")
        for N in NVALS:
            for mf in MFRACS:
                a = pool(mf, N)
                if a is None:
                    continue
                au, mt, Y = a[:, COL["auto"]], a[:, COL["mt"]], a[:, COL["Y"]]
                se = lambda v: 100 * v.std(ddof=1) / np.sqrt(len(v))
                # fixation is all-or-nothing by the sampling generation, so these are simply
                # the fraction of replicates at 1.0; the joint is the fraction with BOTH
                fm, fy = mt >= 0.999, Y >= 0.999
                fh.write(f"{N},{N//2},{mf:g},{1-mf:g},{len(a)},{100*au.mean():.3f},"
                         f"{100*mt.mean():.3f},{100*Y.mean():.3f},"
                         f"{se(au):.3f},{se(mt):.3f},{se(Y):.3f},"
                         f"{100*fm.mean():.2f},{100*fy.mean():.2f},"
                         f"{100*(fm & fy).mean():.2f}\n")
                print(f"{N:>6} {mf:>9g} {len(a):>5} {100*au.mean():>7.2f} "
                      f"{100*mt.mean():>8.1f} {100*Y.mean():>7.1f}")
    print("\nwrote sexbias_stable_boundary.csv")


if __name__ == "__main__":
    main()
```
