## Supplementary Information section 3 code base for "Hypothesis: A modern human range expansion ∼300,000 years ago had a critical role in Neandertal origins": panm4_run.py

```
#!/usr/bin/env python3
"""
panm4_run.py -- sex-biased gene flow at a stable boundary.

The stable-boundary counterpart to the female_bias sweep of the range expansion (grid5). Note
the two parameters are NOT the same quantity and should not share an axis label:

  * here, `male_frac` is the fraction of the MODERN CONTRIBUTION that is male. It is a property
    of who migrates. At male_frac = 0 every modern immigrant is a woman, so modern mtDNA enters
    at 2f and modern Y at 0.
  * in the range expansion, `female_bias` is the probability that a cross-lineage CHILD joins
    its mother's group rather than its father's. It is a residence rule; the gene flow itself is
    symmetric.

Autosomal ancestry is f by construction at every male_frac, so the 5% target needs no tuning.

Row layout:  seed  mt  Y  auto  X
Usage:       python panm4_run.py <male_frac> <N> <reps> <seed0> <outfile>
"""
import os
import sys
from dataclasses import replace
import panmictic_sim as PS
from anchors import SEC1_LADDER, SEC1_PRIMARY  # single source of truth: see anchors.py

MFRACS = [0.0, 0.1, 0.2, 0.3, 0.4, 0.5, 0.6, 0.7, 0.8, 0.9, 1.0]
# Section-1 pairing ladder (anchors.py): census -> locus_Ne 500/750/1000, which is what
# SI section 1 integrates its ABC posterior draws against. 3000 and 6000 are dropped:
# those were ACCUMULATION-phase sizes from 750-250 ka, a different epoch.
NVALS  = list(SEC1_LADDER)
PULSE  = 0.05


def main():
    mf = float(sys.argv[1]); N = int(sys.argv[2])
    reps, seed0, outfile = int(sys.argv[3]), int(sys.argv[4]), sys.argv[5]
    os.makedirs(os.path.dirname(outfile) or ".", exist_ok=True)
    cfg0 = PS.Config(N=N, f=PULSE, male_frac=mf, s=0.0, sel_locus="none")
    with open(outfile, "w") as f:
        f.write("seed\tmt\tY\tauto\tX\n")
        for seed in range(seed0, seed0 + reps):
            r = PS.run(replace(cfg0, seed=seed))
            f.write(f"{seed}\t{r['mt_mod']:.6g}\t{r['Y_mod']:.6g}\t"
                    f"{r['auto_mod']:.6g}\t{r['X_mod']:.6g}\n")
    print("wrote", outfile)


if __name__ == "__main__":
    main()
```
