## Supplementary Information section 3 code base for "Hypothesis: A modern human range expansion ∼300,000 years ago had a critical role in Neandertal origins": panm4_submit.py

```
#!/usr/bin/env python3
"""
panm4_submit.py -- SLURM array for panm4_run.py.
Usage:  python panm4_submit.py <reps> <batch> [concurrency] [partition] [--resume]
        python panm4_submit.py 1000 100 500 short
then:   sbatch panm4_array.sh          (output -> out_panm4/)
"""
import os
import sys
from panm4_run import MFRACS, NVALS


def main():
    args   = [a for a in sys.argv[1:] if not a.startswith("--")]
    total  = int(args[0]) if len(args) > 0 else 1000
    batch  = int(args[1]) if len(args) > 1 else 100
    cap    = int(args[2]) if len(args) > 2 else 500
    part   = args[3] if len(args) > 3 else "short"
    resume = "--resume" in sys.argv
    os.makedirs("out_panm4", exist_ok=True); os.makedirs("logs", exist_ok=True)
    tasks, skipped = [], 0
    for s_ in MFRACS:
        for N in NVALS:
            for s0 in range(0, total, batch):
                reps = min(batch, total - s0)
                out = f"out_panm4/out_{s_:g}_{N}_s{s0}.txt"
                if resume and os.path.exists(out) and sum(1 for _ in open(out)) >= reps + 1:
                    skipped += 1; continue
                tasks.append(f"{s_} {N} {reps} {s0} {out}")
    if not tasks:
        print("nothing to do -- every output file is already complete"); return
    open("panm4_params.txt", "w").write("\n".join(tasks) + "\n")
    n = len(tasks); cap = min(n, cap)
    open("panm4_array.sh", "w").write(f"""#!/bin/bash
#SBATCH -J panm4
#SBATCH -p {part}
#SBATCH -t 0-11:59
#SBATCH -c 1
#SBATCH --mem=1G
#SBATCH -o logs/panm4_%A_%a.log
#SBATCH --array=0-{n - 1}{("%%%d" % cap) if cap else ""}

LINE=$(sed -n "$((SLURM_ARRAY_TASK_ID + 1))p" panm4_params.txt)
echo "task $SLURM_ARRAY_TASK_ID : $LINE"
python panm4_run.py $LINE
""")
    ncond = len(MFRACS) * len(NVALS)
    print(f"wrote panm4_params.txt ({n} tasks over {ncond} conditions), {total} reps each")
    if resume: print(f"  RESUME: skipped {skipped} complete tasks")
    print(f"  {ncond} (male_frac, N) combinations")
    print("submit:  sbatch panm4_array.sh")


if __name__ == "__main__":
    main()
```
