## Supplementary Information section 3 code base for "Hypothesis: A modern human range expansion ∼300,000 years ago had a critical role in Neandertal origins": panmictic_sim.py

```
"""
panmictic_sim.py -- the NON-SPATIAL alternative to spatial_sim2.py.

Here the gene exchange between early modern humans and Neandertal ancestors happens at a
stable geographic boundary rather than through a range expansion, so there is no need to model
geography. The exchange is a single INSTANTANEOUS PULSE into one randomly mating (panmictic)
Neandertal population, which then evolves for `gens` generations.

Model
  * One population of N diploid individuals, N/2 males and N/2 females, constant through time.
  * At generation 0 a fraction `f` of the individuals are early modern humans and the rest are
    archaic. `male_frac` sets what fraction of the modern contribution is male, so the pulse
    can be unbiased (0.5), entirely male (1.0), or entirely female (0.0). Note the resulting
    starting frequencies:
        autosomes        f            (regardless of the sex bias)
        Y (among males)  2*f*male_frac
        mtDNA            2*f*(1-male_frac)
    so an all-male pulse starts the modern Y at 2f and the modern mtDNA at 0, and vice versa.
  * Each generation is Wright-Fisher with separate sexes: every offspring draws a father
    uniformly from the males and a mother uniformly from the females.
  * Loci: mtDNA (maternal), Y (paternal, males only), 8 unlinked autosomes (diploid), 2 X loci
    (mother transmits one of her two; father transmits his X to daughters only) -- the same
    complement as the spatial engine, so results are directly comparable.
  * Selection is viability selection with population regulation back to N (candidates are
    oversampled and non-survivors discarded). This differs from the spatial engine, where a
    removed offspring simply lowers local density; in a single panmictic population of fixed
    size the standard treatment is to regulate back to N.

Contrast with the range-expansion model
  Under a pulse the autosomal modern proportion starts at f and stays there in expectation --
  it is set by the pulse, not built up by repeated local introgression. The fixation
  probability of a neutral uniparental marker equals its starting frequency, so with an
  unbiased 5% pulse each of the Y and mtDNA fixes about 5% of the time and both fix about
  0.25% of the time. That is the baseline this model exists to quantify.
"""

import numpy as np
from dataclasses import dataclass, replace

AUTOS = 8
XLOCI = 2
DRIVER_XLOCUS = 0     # X locus carrying the driver; locus 1 stays as background
                      # (same index as spatial_sim3, so the two engines agree)


@dataclass
class Config:
    N: int = 1000              # diploid individuals (Petr et al. use Ne ~1000 for Neandertals)
    f: float = 0.05            # modern human fraction in the instantaneous pulse
    male_frac: float = 0.5     # fraction of the modern contribution that is male
    # --- additive selection against the archaic allele ---
    s: float = 0.0         # >0 favours the modern allele, <0 favours the archaic one
    sel_locus: str = "none"    # "none", "Y", "mt", "auto", "X"
    auto_locus: int = 7
    x_locus: int = 1
    # --- CRYPTIC X-LINKED DRIVE AGAINST THE ARCHAIC Y, same semantics as spatial_sim3 -------
    # ADDED 14 Sep 2026. DR: "where is the black curve at route 2?" It was absent because this
    # engine had no X-drive, so the unbiased migration pulse -- the one scenario in Figure 2b
    # that is a pulse rather than an expansion -- could not be evaluated against the mechanism
    # Route 2 varies. The panel therefore showed three of its four scenarios with no statement
    # about the fourth, which a reader meeting the colour scheme in Route 1 reads as data loss.
    #
    # THE MECHANISM, copied from spatial_sim3 so the two engines answer the same question:
    # X locus 0 is a driver. In a male carrying the MODERN allele there together with an
    # ARCHAIC Y, each Y-bearing conception is converted to an X-bearing one with probability
    # xdrive. The modern Y carries the coevolved suppressor and is immune. xdrive_cost is the
    # standard fertility cost of sperm-killing drive, removing an expressing male's offspring
    # with that probability regardless of sex.
    #
    # WHY IT IS A WEIGHT HERE AND A SEX FLIP THERE. spatial_sim3 draws conceptions and flips
    # some from son to daughter. This engine fills a FIXED quota of each sex by rejection
    # sampling, so flipping is not available -- a flipped son would have to be handed to the
    # daughter pass, which does not exist. The equivalent is to weight fathers: an expressing
    # male fathers sons at (1 - d) times the rate of a non-expressing one, and daughters at
    # (1 + d) times. Weights are acceptance probabilities and must not exceed 1, so the
    # daughter pass is written as 1 for expressing and 1/(1 + d) for everyone else, which is
    # the same ratio. The realised sex ratio stays at the configured N/2, which is what this
    # engine assumes throughout; in spatial_sim3 the drive perturbs it slightly. That is the
    # one behavioural difference between the two implementations and it is second order --
    # the quantity Route 2 plots is the joint retention probability, not the sex ratio.
    xdrive: float = 0.0
    xdrive_cost: float = 0.0
    # --- epistatic (Dobzhansky-Muller) selection, same semantics as spatial_sim2 ---
    dm: float = 0.0
    dm_focal: str = "mt"
    dm_focal_locus: int = 7
    dm_partner: str = "Y"
    dm_partner_locus: int = 7
    dm_dom: str = "dominant"
    dm_male_limited: bool = True
    # F1 male sterility: sons produced in the FIRST generation after the pulse by a modern
    # mother and an archaic father leave no offspring. With an all-female pulse every son of
    # a modern mother is such an F1, and is identified by his modern mtDNA.
    # s_mt: selection on the mtDNA, INDEPENDENT of s/sel_locus, so a run can carry selection
    # on the Y and on the mtDNA simultaneously. Positive favours the modern type; negative
    # favours the archaic one, which is needed because the Muller's-ratchet posterior on the
    # mtDNA fitness gap has 8.5% (N_A = 1500) to 24% (N_A = 3000) of its mass below zero.
    s_mt: float = 0.0
    f1_male_sterile: bool = False
    # f1_male_theta: the CONTINUOUS version of the flag above, and the stable-boundary
    # counterpart of spatial_sim2's theta_f1m. An F1 hybrid male -- a son of an ARCHAIC
    # father and a MODERN mother, matching spatial_sim2's f1m flag exactly -- reproduces at
    # relative rate theta. Unlike f1_male_sterile it identifies the father as well as the
    # mother (modern mtDNA AND an archaic Y), which f1_male_sterile does not: that flag tests
    # mt alone and so is correct only for an all-female pulse, where every son of a modern
    # mother necessarily has an archaic father. Like that flag it fires only in the generation
    # bred from the pulse, because a single pulse produces genuine F1s exactly once.
    f1_male_theta: float = 1.0
# 6993 generations = 200 kyr at 28.6 yr/generation: mixture at 250 ka to the age of the
# sequenced Neandertals at 50 ka. The previous value of 9091 encoded 260 kyr, i.e.
# sampling at the present day, which predates the timeline being pinned down. The two
# give identical results (trajectories are fixed or lost long before either), so this is
# a consistency change rather than a substantive one.
    gens: int = 6993
    # fix-or-lose early stop (DR, 4 Sep 2026): stop as soon as BOTH uniparental markers are
    # absorbed -- fixed or lost. Nothing changes at either marker afterwards and autosomal
    # ancestry is neutral, so the reported means are identical; only the runtime differs.
    # There is no lineage sweep and no origin reseed here, so no further guard is needed.
    stop_when_absorbed: bool = True
    # --- PERIODIC SIZE FLUCTUATION (DR, 4 Sep 2026) -----------------------------------------
    # Climatic worsenings on a Heinrich-event cadence drive the population through crashes and
    # recoveries. Drift is set by the HARMONIC mean of size over time, but a beneficial allele
    # escapes loss whenever size is momentarily large, so a cycling population can hold the
    # coalescent Ne that the PSMC curves require while giving selection far more purchase than
    # a constant population of that size. PSMC has no resolution at millennial scales at a
    # 200 kyr time depth, so this is unconstrained by the data rather than contradicted by it.
    #
    # size(t) = base * (m + a*sin(2*pi*t/T + phase)) / sqrt(R),  m = (R+1)/2, a = (R-1)/2
    # The sqrt(R) divisor is what pins the HARMONIC mean of the cycle to `base`, so turning the
    # fluctuation on does not change the drift anchor -- only how the size is distributed in
    # time. phase = pi/2 starts the run at a peak.
    #
    # Period in generations at 28.6 yr/gen: Dansgaard-Oeschger ~1,470 yr = 51; Heinrich
    # ~6-7 kyr = 210-245. cycle_T = 0 disables the fluctuation entirely.
    # --- A SECOND PULSE (DR, 14 Sep 2026) ---------------------------------------------------
    # Route 4 of Figure 2b is the two-wave scenario: a matrilocal range expansion preceded or
    # followed by a patrilocal one, so that one wave delivers the modern mtDNA and the other
    # the modern Y. The panel had no comparison curves at all, because the stable-boundary
    # scenarios had no second event to give them.
    #
    # These three parameters add one. At generation pulse2_gen a further fraction pulse2_f of
    # the population is replaced by modern individuals, of whom pulse2_male_frac are male.
    # pulse2_gen = 0 disables it, so every existing run is unaffected.
    #
    # WHY THE SEX BIAS IS A FREE PARAMETER RATHER THAN FIXED. Two UNBIASED pulses deliver more
    # of the same and cannot produce the asymmetry Route 4 turns on. The informative control is
    # an all-female pulse followed by an all-male one -- the stable-boundary reading of
    # "matrilocal then patrilocal" -- which tests whether the two-wave logic needs a range
    # expansion at all, or whether a pair of sex-biased pulses at a stationary boundary does
    # the same work. Leaving male_frac free covers both readings, so the sweep chooses and the
    # engine does not commit the figure to either.
    #
    # The replacement is by individual, matching the first pulse: the chosen individuals become
    # wholly modern at every locus. Drawing them uniformly WITHIN each sex keeps the population
    # sex ratio at N/2 and avoids preferentially overwriting whatever the first wave left,
    # which would understate how much of wave one survives wave two.
    pulse2_gen: int = 0
    pulse2_f: float = 0.05
    pulse2_male_frac: float = 1.0
    cycle_T: int = 0
    cycle_R: float = 1.0           # peak/trough ratio
    cycle_phase: float = 1.5707963267948966   # pi/2: start at the peak

    oversample: float = 3.0    # candidate offspring per slot when selection is acting
    seed: int = 0


# ---------------------------------------------------------------- initialization
def _init(cfg, rng):
    N = cfg.N
    nm = N // 2
    nf = N - nm
    n_mod_m = int(round(cfg.f * N * cfg.male_frac))
    n_mod_f = int(round(cfg.f * N * (1.0 - cfg.male_frac)))
    if n_mod_m > nm or n_mod_f > nf:
        raise ValueError("modern pulse exceeds the number of individuals of that sex")
    sex = np.concatenate([np.ones(nm, np.int8), np.zeros(nf, np.int8)])
    mod = np.zeros(N, bool)
    mod[:n_mod_m] = True                       # modern males occupy the first slots
    mod[nm:nm + n_mod_f] = True                # modern females the first female slots
    v = mod.astype(np.int8)
    P = dict(
        sex=sex,
        mt=v.copy(),
        Y=np.where(sex == 1, v, -1).astype(np.int8),
        auto=np.repeat((2 * v)[:, None], AUTOS, axis=1).astype(np.int8),
        X=np.repeat((np.where(sex == 0, 2, 1) * v)[:, None], XLOCI, axis=1).astype(np.int8),
    )
    return P


def _second_pulse(P, cfg, rng):
    """Replace a fraction of the population with modern individuals, in place.

    Sampled without replacement within each sex, so the sex ratio is untouched and the number
    of modern arrivals of each sex is exactly what pulse2_f and pulse2_male_frac specify --
    not a binomial draw around it, which at f = 0.05 and N = 1312 would add noise of the same
    order as the signal the two-wave comparison is trying to measure.
    """
    N = P["sex"].size
    males = np.nonzero(P["sex"] == 1)[0]
    females = np.nonzero(P["sex"] == 0)[0]
    n_m = min(int(round(cfg.pulse2_f * N * cfg.pulse2_male_frac)), males.size)
    n_f = min(int(round(cfg.pulse2_f * N * (1.0 - cfg.pulse2_male_frac))), females.size)
    idx = np.concatenate([rng.choice(males, n_m, replace=False),
                          rng.choice(females, n_f, replace=False)]).astype(int)
    if idx.size == 0:
        return P
    P["mt"][idx] = 1
    male_idx = idx[P["sex"][idx] == 1]
    P["Y"][male_idx] = 1
    P["auto"][idx, :] = 2
    P["X"][idx, :] = np.where(P["sex"][idx, None] == 0, 2, 1)
    return P


# ---------------------------------------------------------------- selection
def _dm_penetrance(cfg, sx, mt, Yc, auto, X):
    n = sx.shape[0]

    def focal_modern():
        f = cfg.dm_focal
        if f == "mt":   return (mt == 1)
        if f == "Y":    return (sx == 1) & (Yc == 1)
        if f == "auto": return auto[:, cfg.dm_focal_locus] >= 1
        if f == "X":    return X[:, cfg.dm_focal_locus] >= 1
        raise ValueError(cfg.dm_focal)

    def partner_archaic():
        p, dom = cfg.dm_partner, cfg.dm_dom
        if p == "mt":   return (mt == 0).astype(float)
        if p == "Y":    return ((sx == 1) & (Yc == 0)).astype(float)
        if p == "auto":
            k = 2 - auto[:, cfg.dm_partner_locus]; maxd = np.full(n, 2)
        elif p == "X":
            maxd = np.where(sx == 0, 2, 1); k = maxd - X[:, cfg.dm_partner_locus]
        else:
            raise ValueError(p)
        if dom == "dominant":  return (k >= 1).astype(float)
        if dom == "recessive": return (k == maxd).astype(float)
        return k / maxd

    p = cfg.dm * focal_modern().astype(float) * partner_archaic()
    if cfg.dm_male_limited:
        p = p * (sx == 1)
    return np.clip(p, 0.0, 1.0)


def _disfavoured_count(cfg, sx, mt, Yc, auto, X):
    """Copies of the DISFAVOURED allele carried by each candidate.

    cfg.s > 0 selects against the ARCHAIC allele (the modern one is favoured); cfg.s < 0
    reverses this, selecting against the MODERN allele with strength |s|. Allowing negative s
    matters when a posterior on the fitness gap puts mass on the archaic type being the fitter
    of the two, which must not be silently treated as neutral.
    """
    n = _archaic_count(cfg, sx, mt, Yc, auto, X)
    if cfg.s >= 0:
        return n
    if cfg.sel_locus == "Y":    return np.where(sx == 1, 1 - n, 0)   # males only
    if cfg.sel_locus == "mt":   return 1 - n
    if cfg.sel_locus == "auto": return 2 - n
    if cfg.sel_locus == "X":    return np.where(sx == 0, 2, 1) - n
    return n


def _archaic_count(cfg, sx, mt, Yc, auto, X):
    if cfg.sel_locus == "Y":
        return np.where((sx == 1) & (Yc == 0), 1, 0)
    if cfg.sel_locus == "mt":
        return (mt == 0).astype(np.int64)
    if cfg.sel_locus == "auto":
        return 2 - auto[:, cfg.auto_locus]
    if cfg.sel_locus == "X":
        return np.where(sx == 0, 2, 1) - X[:, cfg.x_locus]
    return np.zeros(sx.shape, np.int64)


# ---------------------------------------------------------------- one generation
def _breed(P, cfg, rng, n_needed, sex_val, gen=1):
    """Draw `n_needed` surviving offspring of sex `sex_val`."""
    males = np.nonzero(P["sex"] == 1)[0]
    females = np.nonzero(P["sex"] == 0)[0]
    if males.size == 0 or females.size == 0:
        return None
    selecting = ((cfg.s != 0 and cfg.sel_locus != "none") or cfg.dm > 0 or cfg.s_mt != 0
                 or cfg.xdrive > 0 or cfg.xdrive_cost > 0
                 or (cfg.f1_male_sterile and gen == 0 and sex_val == 1)
                 or (cfg.f1_male_theta < 1.0 and gen == 0 and sex_val == 1))
    out = {k: [] for k in ("mt", "Y", "auto", "X")}
    got = 0
    tries = 0
    drawn = 0          # adaptive oversampling: track the realised acceptance rate so that
    kept_tot = 0       # strong selection (e.g. s=1, where nearly every male is culled) does
    while got < n_needed and tries < 200:   # not require hundreds of undersized batches
        tries += 1
        if selecting:
            if drawn > 0 and kept_tot > 0:      # use the realised acceptance rate once known
                factor = min(max(1.1 / max(kept_tot / drawn, 1e-3), 1.05), 500.0)
            else:                                # first batch: fall back to the configured guess
                factor = cfg.oversample
        else:
            factor = 1.0
        B = int(np.ceil((n_needed - got) * factor))
        fa = males[rng.integers(0, males.size, B)]
        mo = females[rng.integers(0, females.size, B)]
        mt = P["mt"][mo].astype(np.int8)
        Yc = (P["Y"][fa] if sex_val == 1 else np.full(B, -1)).astype(np.int8)
        auto = ((rng.random((B, AUTOS)) < P["auto"][fa] / 2.0).astype(np.int8)
                + (rng.random((B, AUTOS)) < P["auto"][mo] / 2.0).astype(np.int8))
        mo_tx = (rng.random((B, XLOCI)) < P["X"][mo] / 2.0).astype(np.int8)
        X = (mo_tx + P["X"][fa]).astype(np.int8) if sex_val == 0 else mo_tx.astype(np.int8)
        sx = np.full(B, sex_val, np.int8)
        if selecting:
            w = np.ones(B)
            if cfg.s != 0 and cfg.sel_locus != "none":
                w *= np.clip(1.0 - abs(cfg.s)
                             * _disfavoured_count(cfg, sx, mt, Yc, auto, X), 0.0, 1.0)
            if cfg.s_mt != 0:
                dis = (mt == 0) if cfg.s_mt > 0 else (mt == 1)
                w *= np.clip(1.0 - abs(cfg.s_mt) * dis, 0.0, 1.0)
            if cfg.dm > 0:
                w *= (1.0 - _dm_penetrance(cfg, sx, mt, Yc, auto, X))
            if cfg.xdrive > 0 or cfg.xdrive_cost > 0:
                # expressing = modern driver allele at X locus 0 AND an archaic Y. Males carry
                # one X, coded 1 for modern and 0 for archaic, so == 1 is the modern allele.
                expressing = (P["X"][fa][:, DRIVER_XLOCUS] == 1) & (P["Y"][fa] == 0)
                if cfg.xdrive > 0:
                    d = cfg.xdrive
                    w *= (np.where(expressing, 1.0 - d, 1.0) if sex_val == 1
                          else np.where(expressing, 1.0, 1.0 / (1.0 + d)))
                if cfg.xdrive_cost > 0:
                    w *= np.where(expressing, 1.0 - cfg.xdrive_cost, 1.0)
            if cfg.f1_male_sterile and gen == 0 and sex_val == 1:
                w = w * (mt != 1)      # F1 sons of modern mothers do not reproduce
            if cfg.f1_male_theta < 1.0 and gen == 0 and sex_val == 1:
                # modern mother (modern mtDNA) AND archaic father (archaic Y)
                f1 = (mt == 1) & (Yc == 0)
                w = w * np.where(f1, cfg.f1_male_theta, 1.0)
            keep = np.nonzero(rng.random(B) < w)[0]
        else:
            keep = np.arange(B)
        drawn += B; kept_tot += keep.size
        if keep.size:
            take = keep[:n_needed - got]
            out["mt"].append(mt[take]); out["Y"].append(Yc[take])
            out["auto"].append(auto[take]); out["X"].append(X[take])
            got += take.size
    if got < n_needed:
        return None
    return {k: (np.concatenate(v, 0) if v[0].ndim > 1 else np.concatenate(v))
            for k, v in out.items()}


def _step(P, cfg, rng, gen=1):
    nm = cfg.N // 2
    nf = cfg.N - nm
    sons = _breed(P, cfg, rng, nm, 1, gen)
    if sons is None: return None
    daughters = _breed(P, cfg, rng, nf, 0, gen)
    if daughters is None: return None
    return dict(
        sex=np.concatenate([np.ones(nm, np.int8), np.zeros(nf, np.int8)]),
        mt=np.concatenate([sons["mt"], daughters["mt"]]),
        Y=np.concatenate([sons["Y"], daughters["Y"]]),
        auto=np.concatenate([sons["auto"], daughters["auto"]], 0),
        X=np.concatenate([sons["X"], daughters["X"]], 0),
    )


# ---------------------------------------------------------------- stats & driver
def _stats(P, exclude=(), exclude_x=()):
    male = P["sex"] == 1
    xcols = [c for c in range(XLOCI) if c not in exclude_x]
    acols = [c for c in range(AUTOS) if c not in exclude]

    def xfrac(mask, diploid):
        if not mask.any() or not xcols:
            return np.nan
        cap = 2.0 if diploid else 1.0
        return float((P["X"][mask][:, xcols].sum(1) / (len(xcols) * cap)).mean())

    return dict(
        mt_mod=float((P["mt"] == 1).mean()),
        Y_mod=float((P["Y"][male] == 1).mean()) if male.any() else np.nan,
        auto_mod=float(P["auto"][:, acols].mean() / 2.0),
        X_mod=float(np.nanmean([v for v in (xfrac(~male, True), xfrac(male, False)) if v == v])),
        pop=int(P["sex"].size))


def run(cfg):
    rng = np.random.default_rng(cfg.seed)
    excl = set(); excl_x = set()
    if cfg.sel_locus == "auto": excl.add(cfg.auto_locus)
    if cfg.sel_locus == "X": excl_x.add(cfg.x_locus)
    if cfg.dm > 0 and cfg.dm_focal == "auto": excl.add(cfg.dm_focal_locus)
    if cfg.dm > 0 and cfg.dm_partner == "auto": excl.add(cfg.dm_partner_locus)
    if cfg.dm > 0 and cfg.dm_focal == "X": excl_x.add(cfg.dm_focal_locus)
    if cfg.dm > 0 and cfg.dm_partner == "X": excl_x.add(cfg.dm_partner_locus)
    # The driver locus is under selection, so it is excluded from the ancestry measurement --
    # DR's rule that "all measurement should be done on markers not affected by selection".
    if cfg.xdrive > 0: excl_x.add(DRIVER_XLOCUS)
    P = _init(cfg, rng)
    for g in range(cfg.gens):
        cur = cfg
        if cfg.cycle_T > 0 and cfg.cycle_R > 1.0:
            m, a = (cfg.cycle_R + 1) / 2.0, (cfg.cycle_R - 1) / 2.0
            f = (m + a * np.sin(2 * np.pi * g / cfg.cycle_T + cfg.cycle_phase)) \
                / np.sqrt(cfg.cycle_R)
            cur = replace(cfg, N=max(20, int(round(cfg.N * f))))
        nxt = _step(P, cur, rng, g)
        if nxt is None:
            break
        P = nxt
        if cfg.pulse2_gen and g == cfg.pulse2_gen:
            P = _second_pulse(P, cfg, rng)
        # THE ABSORPTION STOP IS GUARDED AGAINST THE SECOND PULSE. Without `g > pulse2_gen`
        # the run would halt as soon as wave one's markers absorbed -- which is usually long
        # before pulse2_gen -- and return the SINGLE-wave answer under a two-wave label. This
        # is the same trap that voided every two-wave deme cell in the spatial engine until
        # 13 Sep; it is cheaper to guard it here than to find it in a figure later.
        if (cfg.stop_when_absorbed and g % 10 == 0 and g > 0
                and (not cfg.pulse2_gen or g > cfg.pulse2_gen)):
            male = P["sex"] == 1
            pm = float((P["mt"] == 1).mean())
            py = float((P["Y"][male] == 1).mean()) if male.any() else 0.0
            if pm in (0.0, 1.0) and py in (0.0, 1.0):
                break
    return _stats(P, tuple(excl), tuple(excl_x))


if __name__ == "__main__":
    import sys
    n = int(sys.argv[1]) if len(sys.argv) > 1 else 20
    for tag, kw in [("unbiased 5% pulse", {}),
                    ("all-male 5% pulse", dict(male_frac=1.0)),
                    ("all-female 5% pulse", dict(male_frac=0.0))]:
        res = [run(replace(Config(**kw), seed=s)) for s in range(n)]
        mt = np.array([r["mt_mod"] for r in res]); Y = np.array([r["Y_mod"] for r in res])
        au = np.array([r["auto_mod"] for r in res])
        print(f"{tag:22s} n={n}  auto={100*au.mean():5.2f}%  "
              f"P(Y fixed)={100*(Y >= 1).mean():5.1f}%  P(mt fixed)={100*(mt >= 1).mean():5.1f}%")
```
