## Supplementary Information section 3 code base for "Hypothesis: A modern human range expansion ∼300,000 years ago had a critical role in Neandertal origins": rescreen_submit.py

```
#!/usr/bin/env python3
"""
rescreen_submit.py -- re-screen the borderline cells at 100 replicates instead of 10, then
score whichever newly pass. (12 Sep 2026)

WHY. Figure 2b's Route 4 deme column holds ONE point. That looked like a coverage gap in the
gamma ladder and it is not. Of the 19 deme cells at theta = 0 whose autosomal proportion lands
in 3-7% with the expansion completing in 100-1,000 generations, sixteen are rejected by the
establishment criterion:

    aut    end_gen   established     scored?
    3.01      79        2 of 10        no
    5.40     405        4 of 10        no
    6.01     160        5 of 10        no       <- rejected
    6.06      50        5 of 10        no       <- rejected
    6.65     617       10 of 10        YES

`stage1_pass` requires n_est / n > 0.50 and stage 1 runs TEN replicates, so the gate is
"6 or more heads in 10 tosses". A cell whose true establishment probability is 0.55 fails this
screen about 40% of the time, and four of the sixteen rejected cells sit exactly at 5 of 10.
Route 4's near-empty deme column is therefore substantially an artefact of the screen's
replicate count, not a statement about the mechanism -- which is the opposite of what the
figure's "an empty column is a result" note currently claims for it.

At 100 replicates the standard error on establishment falls from 0.158 to 0.050, and the gate
stops being a coin flip.

SCOPE: BORDERLINE CELLS ONLY. A cell is re-screened if it already passes the autosomal and
duration criteria (3-7%, 100-1,000 generations) and its establishment estimate is between 0.2
and 0.8 -- i.e. only where the 10-replicate estimate is imprecise enough to plausibly be on the
wrong side of 0.5. Cells at 0 of 10 or 10 of 10 are left alone: more replicates cannot move
them across the threshold, and re-running everything would cost 20x for no change in the
figure. 2,120 cells qualify across the six sets and both frameworks.

    measured per-cell cost      stage 1 @ 100      stage 2 @ 200
        deme                         57 s              120 s
        band                        280 s              475 s

    re     222 deme + 1,898 band   ~152 CPU-hours
    score  whichever newly pass    ~130 CPU-hours if half of them do

RESULTS GO TO A PARALLEL DIRECTORY, out_{prefix}1hi_{fw}, NOT over the 10-replicate files.
Overwriting would destroy the provenance of every number already in the note and make the
"combinations searched" totals unreconstructable. Readers of the screen (figure2c_fountain.py,
routes_build.build stage 2) prefer the hi-replicate file when one exists and fall back to the
original otherwise, so the two can coexist while the figures are rebuilt.

    python rescreen_submit.py re       # stage 1 at 100 reps, borderline cells only
    python rescreen_submit.py stale    # stage 1 at 100 reps, EVERY cell of a stale directory
    python rescreen_submit.py score    # stage 2 at 200 reps, for cells that now pass
    python rescreen_submit.py status   # what is on disk, no submission
"""
import glob
import importlib
import json
import math
import os
import subprocess
import sys

HERE = os.path.dirname(os.path.abspath(__file__))
sys.path.insert(0, HERE)

SETS = ["main", "dmi", "dmi2", "xdc", "xd2", "tw"]
# routes_build's set NAMES differ from its output PREFIXes for one set: `main` writes `routes`.
# STALE is keyed on set name, the directories on prefix, so the two must be translated or the
# lookup silently never matches -- the same mismatch that kept the `main` set from running at
# all for two days.
SET_PREFIX = {"main": "routes"}
AUT_LO, AUT_HI = 3.0, 7.0          # the widened acceptance window
GEN_LO, GEN_HI = 100.0, 1000.0
EST_BAND = (0.20, 0.80)            # only cells the 10-rep screen cannot resolve
RE_REPS, SCORE_REPS = 100, 200
PART, CAP = "short", 900


def load_set(name):
    """Import routes_build under one ROUTE_SET. The module reads the variable at import, so
    it has to be reloaded per set rather than imported once."""
    os.environ["ROUTE_SET"] = name
    import routes_build
    return importlib.reload(routes_build)


def num(x):
    return x is not None and not (isinstance(x, float) and math.isnan(x))


def summary_of(path):
    try:
        with open(path) as f:
            return json.load(f)["summary"]
    except Exception:
        return None


def est_of(d):
    n, ne = d.get("n"), d.get("n_est")
    if not n:
        return None
    return ne / n if ne is not None else None


def borderline(rb, fw, band):
    """Cells whose 10-replicate establishment estimate cannot resolve the > 50% gate."""
    out = []
    s1dir = os.path.join(HERE, f"out_{rb.PREFIX}1_{fw}")
    hidir = os.path.join(HERE, f"out_{rb.PREFIX}1hi_{fw}")
    for mech, v, kw in rb.cells(band):
        name = rb.tag(fw, mech, v, kw) + ".json"
        if os.path.exists(os.path.join(hidir, name)):
            continue
        d = summary_of(os.path.join(s1dir, name))
        if d is None:
            continue
        a, g, e = d.get("aut"), d.get("end_gen"), est_of(d)
        if not (num(a) and num(g)) or e is None:
            continue
        if not (AUT_LO <= a <= AUT_HI and GEN_LO <= g <= GEN_HI):
            continue
        if not (EST_BAND[0] <= e <= EST_BAND[1]):
            continue
        out.append((mech, v, kw, name))
    return out


def all_cells(rb, fw, band):
    """Every cell of a set, borderline or not -- for re-running a directory that is STALE.

    WHY THIS MODE EXISTS. out_tw1_band was written on 12 Sep between 00:26 and 02:55, before
    the wave-2 guard landed in robust_run. Without that guard, stage 1 breaks at the FIRST
    archaic extinction, which under a two-wave scenario is the end of wave ONE -- so every one
    of those 1,701 files records the duration of the first wave and labels it the duration of
    the two-wave process. The re-screen caught it by accident, on the 238 cells it happened to
    cover:

        wave2_gen      old (stale)   correctly run
            500            404           2,859
           1000            658           3,585
           2000            749           4,182

    A factor of four to seven. And the consequence is not cosmetic: at the correct durations,
    ZERO of 224 band two-wave cells complete inside the note's 100-1,000 generation window,
    where the stale files put most of them comfortably inside it. Route 4's band column in
    Figure 2b and its panel in Figure S3.3 are both drawn from that screen.

    out_tw1_deme is NOT affected -- it was written at 08:09-08:59, after the fix -- and it
    behaves quite differently: at a second wave founded at generation 500, 657 of its 744 cells
    do finish inside the window. So this is a genuine difference between the frameworks once
    both are measured correctly, not a bug on both sides.

    Results go to out_{prefix}1hi_{fw} like the rest of the re-screen, so the readers that
    already prefer that directory pick them up with no further change, and the stale files stay
    on disk for provenance rather than being deleted.
    """
    # ONLY CELLS THAT WERE ACTUALLY SCREENED, not the whole Cartesian product. Enumerating
    # rb.cells() gives every admissible (grid, K, m, gamma, r) crossed with every wave-2 rung
    # -- 34,272 for the deme tw set -- where the screen that went stale covered 2,232 of them.
    # Re-running the other 32,040 would not be a re-run, it would be a new and much larger
    # sweep costing 540 CPU-hours, and it would silently change the "combinations searched"
    # totals the note quotes. `stale` means redo what is there.
    out = []
    hidir = os.path.join(HERE, f"out_{rb.PREFIX}1hi_{fw}")
    s1dir = os.path.join(HERE, f"out_{rb.PREFIX}1_{fw}")
    for mech, v, kw in rb.cells(band):
        name = rb.tag(fw, mech, v, kw) + ".json"
        if os.path.exists(os.path.join(hidir, name)):
            continue
        if not os.path.exists(os.path.join(s1dir, name)):
            continue
        out.append((mech, v, kw, name))
    return out


# Stage-2 files older than this were scored against the pre-fix screen and, for two-wave
# cells, under the 2,500-generation budget that cannot fit two sweeps. They must be REPLACED,
# not skipped -- which is what `score` would otherwise do, since it treats the existence of a
# stage-2 file as proof the cell is done. 463 of the 740 cells in out_tw2_band are in this
# state.
import datetime as _dt
FIXTIME = _dt.datetime(2026, 9, 12, 7, 35, 8).timestamp()


def newly_passing(rb, fw, band):
    """Cells whose 100-replicate screen passes and which need a stage-2 file."""
    out = []
    hidir = os.path.join(HERE, f"out_{rb.PREFIX}1hi_{fw}")
    s2dir = os.path.join(HERE, f"out_{rb.PREFIX}2_{fw}")
    for mech, v, kw in rb.cells(band):
        name = rb.tag(fw, mech, v, kw) + ".json"
        s2 = os.path.join(s2dir, name)
        # A SET LISTED IN `STALE` IS ALWAYS RE-SCORED, whatever its timestamp. The mtime test
        # catches files written before 12 Sep 07:35, but the two-wave DEME stage-2 files were
        # written at 08:38-09:53 -- after that cutoff and so "clean" by date -- while being
        # void for a quite different reason: the second wave never switched residence rule.
        # Trusting the timestamp here would have left 447 wrong cells in place and the chained
        # scoring pass would have reported nothing to do.
        stale_set = (rb.PREFIX, fw) in STALE_PREFIXES
        if os.path.exists(s2) and not stale_set and os.path.getmtime(s2) >= FIXTIME:
            continue
        d = summary_of(os.path.join(hidir, name))
        if d is None:
            continue
        a, g, e = d.get("aut"), d.get("end_gen"), est_of(d)
        if not (num(a) and num(g)) or e is None or e <= 0.50:
            continue
        if not (AUT_LO <= a <= AUT_HI and GEN_LO <= g <= GEN_HI):
            continue
        out.append((mech, v, kw, name))
    return out


def cmd_for(rb, fw, stage, reps, mech, v, kw, outrel):
    fl = []
    for k in sorted(kw):
        fl += [rb.FLAG[k], f"{kw[k]:g}"]
    return (f"python robust_run.py {fw} {mech} {v:g} {stage} {reps} 0 {outrel} "
            + " ".join(fl))


def emit(name, tasks, hours):
    """Returns the SLURM job id, or None if nothing was submitted."""
    if not tasks:
        print("  nothing to submit")
        return None
    pf, sf = f"{name}_params.txt", f"{name}_array.sh"
    with open(os.path.join(HERE, pf), "w") as f:
        f.write("\n".join(tasks) + "\n")
    with open(os.path.join(HERE, sf), "w") as f:
        f.write(f"""#!/bin/bash
#SBATCH -J {name}
#SBATCH -p {PART}
#SBATCH -c 1
#SBATCH --mem=2G
#SBATCH -t {hours}
#SBATCH -a 0-{len(tasks) - 1}%{CAP}
#SBATCH -o logs/{name}_%A_%a.log
cd {HERE}
CMD=$(sed -n "$((SLURM_ARRAY_TASK_ID + 1))p" {pf})
echo "$CMD"
eval "$CMD"
""")
    print(f"  {len(tasks):,} tasks -> {sf}")
    if "--no-go" in sys.argv:
        print(f"  NOT submitted (--no-go).  sbatch {sf}")
        return
    r = subprocess.run(["sbatch", sf], cwd=HERE, capture_output=True, text=True)
    if r.returncode != 0:
        print(f"  !! sbatch FAILED: {(r.stderr or '').strip()}")
        return None
    print(f"  SUBMITTED: {r.stdout.strip()}")
    return r.stdout.strip().split()[-1]


def chain(dep):
    """Queue the scoring pass behind the re-screen, so one command covers both.

    afterany, not afterok: an array whose last task times out still leaves every other cell's
    JSON on disk, and `score` selects on what is there. Making the follow-up conditional on a
    clean exit would throw away 2,000 good cells because one of them ran long.
    """
    sf = "rescore_chain.sh"
    with open(os.path.join(HERE, sf), "w") as f:
        f.write(f"""#!/bin/bash
#SBATCH -J rechain
#SBATCH -p {PART}
#SBATCH -c 1
#SBATCH --mem=2G
#SBATCH -t 0-00:20
#SBATCH -o logs/rechain_%j.log
cd {HERE}
python rescreen_submit.py score
""")
    r = subprocess.run(["sbatch", f"--dependency=afterany:{dep}", sf],
                       cwd=HERE, capture_output=True, text=True)
    print(f"  CHAINED scoring pass: {r.stdout.strip()}" if r.returncode == 0
          else f"  !! chain FAILED: {(r.stderr or '').strip()}")


# per-cell seconds, measured 12 Sep on one deme and one band cell, linear in replicates
COST = {("deme", 1): 0.57, ("deme", 2): 0.60, ("band", 1): 2.80, ("band", 2): 2.37}


# Directories known to predate a correctness fix, and therefore re-run in full rather than
# only at the borderline. (set, framework) -> why.
STALE = {
    ("tw", "band"): "written before the wave-2 guard; end_gen is wave ONE's duration",
    # 13 Sep. The DEME two-wave cells are void for a different and worse reason: spatial_sim2
    # switched cfg.lineage at wave 2 but not cfg.female_bias, and _repro ignores cfg.lineage
    # whenever female_bias is set -- which robust_run does unconditionally, at 1.0. So the
    # second wave ran under the SAME matrilocal rule as the first, the modern Y never had a
    # route in, and every cell returned Y at 0-2%. Fixed in spatial_sim2 by switching both;
    # at grid 12 / K 11 / m 0.08 / gamma 0.42 the corrected cell gives Y 100%, mtDNA 100% and
    # 4.35% autosomal, against Y 0.0% before. The band path was never affected: robust_run's
    # band_cell sets cfg.beta = 1.0 at wave 2 explicitly.
    ("tw", "deme"): "second wave never switched residence rule; see spatial_sim2 wave-2 block",
}

# STALE keyed on output PREFIX rather than set name, computed once so the test below is a
# plain membership check.
STALE_PREFIXES = {(SET_PREFIX.get(_st, _st), _fw) for (_st, _fw) in STALE}


def run(stage):
    which = {"re": borderline, "stale": all_cells}.get(stage, newly_passing)
    st, reps = (1, RE_REPS) if stage in ("re", "stale") else (2, SCORE_REPS)
    tasks, secs = [], 0.0
    print(f"{'set':<8}{'fw':<6}{'cells':>8}{'CPU-h':>8}")
    for name in SETS:
        rb = load_set(name)
        for fw, band in (("deme", False), ("band", True)):
            if band and not (rb.BAND_SUPPORTS & set(rb.ROUTES)):
                continue
            if stage == "stale" and (name, fw) not in STALE:
                continue
            sel = which(rb, fw, band)
            if not sel:
                continue
            outdir = f"out_{rb.PREFIX}{'1hi' if st == 1 else '2'}_{fw}"
            os.makedirs(os.path.join(HERE, outdir), exist_ok=True)
            for mech, v, kw, fn in sel:
                tasks.append(cmd_for(rb, fw, st, reps, mech, v, kw,
                                     os.path.join(outdir, fn)))
            s = len(sel) * COST[(fw, st)] * reps
            secs += s
            print(f"{name:<8}{fw:<6}{len(sel):>8,}{s / 3600:>8.1f}")
    print(f"{'TOTAL':<14}{len(tasks):>8,}{secs / 3600:>8.1f}")
    if stage in ("re", "stale"):
        jid = emit("rescr" if stage == "re" else "restale", tasks, "0-02:59")
        if jid and "--chain" in sys.argv:
            chain(jid)
    else:
        emit("rescore", tasks, "0-11:59")


def status():
    print(f"{'set':<8}{'fw':<6}{'10-rep':>9}{'100-rep':>9}{'stage2':>9}")
    for name in SETS:
        rb = load_set(name)
        for fw in ("deme", "band"):
            g = lambda d: len(glob.glob(os.path.join(HERE, d, "*.json")))  # noqa: E731
            a = g(f"out_{rb.PREFIX}1_{fw}")
            b = g(f"out_{rb.PREFIX}1hi_{fw}")
            c = g(f"out_{rb.PREFIX}2_{fw}")
            if a or b or c:
                print(f"{name:<8}{fw:<6}{a:>9,}{b:>9,}{c:>9,}")


def main():
    what = sys.argv[1] if len(sys.argv) > 1 else "status"
    os.makedirs(os.path.join(HERE, "logs"), exist_ok=True)
    if what == "status":
        status()
    elif what in ("re", "stale", "score"):
        run(what)
    else:
        raise SystemExit("usage: rescreen_submit.py re|stale|score|status")


if __name__ == "__main__":
    main()
```
