## Supplementary Information section 3 code base for "Hypothesis: A modern human range expansion ∼300,000 years ago had a critical role in Neandertal origins": robust_collect.py

```
#!/usr/bin/env python3
"""
robust_collect.py -- pool the robustness sweep and fit the ridge. (DR, 10 Sep 2026)

Stage 1: writes robust1_<fw>.csv (one row per cell) and, for the deme grid, fits the
calibrate5-style surface -- local linear regression of log(autosomal ancestry) on
(theta, log gamma, log m) with a Gaussian kernel -- inverts it for the gamma that hits the
autosomal target, and reports the leave-one-out R-squared so you can see whether the coarse
grid was dense enough to interpolate on. The inverted gammas are the ridge, and they are what
stage 2 should be run at.

Stage 2: writes robust2_<fw>.csv with the joint statistic per cell, in the shape
figure2_mockup.py reads (route, sublabel, framework, p_joint).

    python robust_collect.py 1 [--band] [--target 5] [--aut-lo 2 --aut-hi 15]
                               [--gen-lo 100 --gen-hi 1000]
    python robust_collect.py 2 [--band]
"""
import csv
import glob
import json
import os
import sys

import numpy as np

HERE = os.path.dirname(os.path.abspath(__file__))


def flag(name, cast, default):
    return cast(sys.argv[sys.argv.index(name) + 1]) if name in sys.argv else default


def main():
    stage = int(sys.argv[1]) if len(sys.argv) > 1 else 1
    band = "--band" in sys.argv
    fw = "band" if band else "deme"
    target = flag("--target", float, 5.0)
    lo, hi = flag("--aut-lo", float, 2.0), flag("--aut-hi", float, 15.0)
    glo, ghi = flag("--gen-lo", float, 100.0), flag("--gen-hi", float, 1000.0)

    rows = []
    for p in glob.glob(os.path.join(HERE, f"out_robust{stage}_{fw}", "*.json")):
        with open(p) as f:
            rows.append(json.load(f)["summary"])
    if not rows:
        raise SystemExit(f"no output in out_robust{stage}_{fw}/")

    # pool the seed batches of each cell
    key = lambda r: (r["mech"], r["val"], r.get("gamma"), r.get("m"), r.get("e"),
                     r.get("p_move"), r["r"], r["K"], r.get("grid"), r.get("N"))
    pooled = {}
    for r in rows:
        k = key(r)
        d = pooled.setdefault(k, dict(r, n=0, n_est=0, _aut=[], _gen=[], _j=[]))
        d["n"] += r["n"]; d["n_est"] += r["n_est"]
        if r["n_est"]:
            d["_aut"].append(r["aut"]); d["_gen"].append(r["end_gen"])
            if r.get("joint") is not None:
                d["_j"].append(r["joint"])
    out = []
    for k, d in pooled.items():
        if not d["_aut"]:
            continue
        d["aut"] = float(np.mean(d["_aut"])); d["end_gen"] = float(np.mean(d["_gen"]))
        d["joint"] = float(np.mean(d["_j"])) if d["_j"] else None
        d["estab"] = d["n_est"] / max(d["n"], 1)
        d["passes"] = int(lo <= d["aut"] <= hi and glo <= d["end_gen"] <= ghi
                          and d["estab"] > 0.999)
        for x in ("_aut", "_gen", "_j"):
            d.pop(x)
        out.append(d)
    out.sort(key=lambda d: (d["mech"], d["val"], d.get("gamma") or d.get("e")))

    cols = [c for c in ("framework", "mech", "val", "gamma", "m", "e", "p_move", "r", "K",
                        "grid", "N", "beta", "lineage", "n", "estab", "aut", "end_gen",
                        "joint", "passes") if c in out[0]]
    fn = os.path.join(HERE, f"robust{stage}_{fw}.csv")
    with open(fn, "w", newline="") as f:
        w = csv.DictWriter(f, fieldnames=cols, extrasaction="ignore")
        w.writeheader(); w.writerows(out)
    npass = sum(d["passes"] for d in out)
    print(f"  wrote {os.path.basename(fn)}: {len(out)} cells, {npass} pass "
          f"(autosomes {lo:g}-{hi:g}%, expansion {glo:g}-{ghi:g} generations, all established)")

    if stage != 1 or band:
        return

    # ---------------- fit the ridge (calibrate5's smoother, on whatever grid we have)
    # the surface is fitted on the mechanism-free and theta cells: those are the ones whose
    # autosomal proportion is a pure function of (theta, gamma, m), which is what the ridge is
    R = [d for d in out if d["mech"] in ("none", "theta")]
    if len(R) < 20:
        print("  too few cells to fit the surface; run a denser grid")
        return
    th = np.array([d["val"] if d["mech"] == "theta" else 1.0 for d in R])
    lga = np.log([d["gamma"] for d in R])
    lm = np.log([d["m"] for d in R])
    lau = np.log([max(d["aut"], 1e-3) for d in R])
    H_TH, H_GA, H_M = 1.6, 0.42, 0.42

    def fit(qt, qg, qm, y, excl=None):
        d2 = ((th - qt) / H_TH) ** 2 + ((lga - qg) / H_GA) ** 2 + ((lm - qm) / H_M) ** 2
        w = np.exp(-0.5 * d2)
        if excl is not None:
            w[excl] = 0.0
        k = w > 1e-4
        if k.sum() < 8:
            return np.nan
        Wt = w[k]
        X = np.column_stack([np.ones(k.sum()), th[k] - qt, lga[k] - qg, lm[k] - qm])
        try:
            return float(np.linalg.solve(X.T @ (Wt[:, None] * X), X.T @ (Wt * y[k]))[0])
        except np.linalg.LinAlgError:
            return np.nan

    res = [lau[i] - fit(th[i], lga[i], lm[i], lau, excl=i) for i in range(len(R))]
    res = np.array([x for x in res if x == x])
    print(f"  surface fit: leave-one-out R^2 = {1 - res.var() / lau.var():.3f} "
          f"on {len(res)} cells (residual SD {res.std():.3f} log units)")
    if 1 - res.var() / lau.var() < 0.8:
        print("  ** R^2 below 0.8: the grid is too coarse to interpolate on; densify before "
              "trusting the ridge **")

    # invert for gamma at the target, per (theta, m)
    lgrid = np.linspace(lga.min(), lga.max(), 241)
    ridge = []
    for t in sorted(set(th)):
        for mm in sorted(set(np.exp(lm))):
            pred = np.array([fit(t, g, np.log(mm), lau) for g in lgrid])
            ok = np.isfinite(pred)
            if ok.sum() < 5:
                continue
            j = np.argmin(np.abs(np.exp(pred[ok]) - target))
            gstar = float(np.exp(lgrid[ok][j]))
            hit = float(np.exp(pred[ok][j]))
            if abs(hit - target) < 0.35 * target:
                ridge.append(dict(theta=t, m=mm, gamma_star=round(gstar, 4),
                                  aut_at_star=round(hit, 2)))
    fn2 = os.path.join(HERE, "robust1_ridge.csv")
    if ridge:
        with open(fn2, "w", newline="") as f:
            w = csv.DictWriter(f, fieldnames=list(ridge[0])); w.writeheader(); w.writerows(ridge)
        print(f"  wrote robust1_ridge.csv: {len(ridge)} ridge points at {target:g}% autosomes")
        print("  these are the (theta, m, gamma*) triples to run at in stage 2")
    else:
        print(f"  no ridge found at {target:g}% -- the target may be outside the grid's reach")


if __name__ == "__main__":
    main()
```
