## Supplementary Information section 3 code base for "Hypothesis: A modern human range expansion ∼300,000 years ago had a critical role in Neandertal origins": robust_run.py

```
#!/usr/bin/env python3
"""
robust_run.py -- one cell of the robustness sweep. (DR, 10 Sep 2026)

Stage 1 (screen)  stop at archaic extinction, few reps, record the autosomal proportion and the
                  completion time. 24x faster than a full run in the deme model, because the
                  deme engine otherwise grinds through ~6,800 generations of an all-modern
                  population. (In the band model the saving is only ~10%: it already stops at
                  marker absorption around generation 900.)

Stage 2 (score)   run until BOTH uniparental markers are absorbed, or generation 6993, and
                  record how often ten sampled males are all modern at both.

A cell passes the screen when the mean autosomal proportion is inside --aut-lo/--aut-hi AND the
mean completion time is inside --gen-lo/--gen-hi AND every replicate established.

WHY BOTH FILTERS. DR, 10 Sep: matching the autosomal proportion alone admits combinations whose
expansion takes 3,000 generations, which is not the archaeological scenario. 100-1000 generations
is the window.

RESEEDING. Uses the settled `guarded` policy: the origin is re-founded only if the modern lineage
goes extinct entirely while archaic demes remain. The old unconditional rule pumped modern
genomes in for ~6,800 generations after the expansion finished and inflated every deme autosomal
number; see the note in spatial_sim2.Config.

    python robust_run.py <framework> <mech> <mechval> <stage> <reps> <seed0> <out> [--flags]
"""
import json
import os
import sys

import numpy as np
from dataclasses import replace

HERE = os.path.dirname(os.path.abspath(__file__))
# ORDER MATTERS, AND IT WAS WRONG. These were three insert(0, ...) calls in the order HERE,
# parent, band_model -- so each one pushed the previous down and the PARENT directory ended up
# ahead of `cluster/` on sys.path. There is a second copy of spatial_sim2.py at the project
# root, so every local run silently imported that one instead of the engine sitting next to
# this script. The two were byte-identical until today, which is exactly why it went unnoticed
# for so long; the moment cluster/spatial_sim2.py was edited to carry 32 loci, the runs kept
# using the 8-locus root copy and failed with "index 31 is out of bounds for axis 1 with
# size 8". O2 has a flat layout with only one copy, so no cluster result was ever affected.
# Appended in priority order now: this directory first, always.
for _p in (HERE, os.path.join(os.path.dirname(HERE), "band_model"), os.path.dirname(HERE)):
    if _p not in sys.path:
        sys.path.append(_p)
sys.path.insert(0, HERE)

NSAMPLE = 10
CAP = 6993


def flag(name, cast, default):
    return cast(sys.argv[sys.argv.index(name) + 1]) if name in sys.argv else default


# RETIRED, 13 Sep 2026, and kept only as a record of what it used to hold.
#
# SIM3_ONLY named the mechanisms that only spatial_sim3 implements, and deme_cell picked its
# engine from it. That made correctness depend on a human remembering to extend a list, and
# when the five cultural mechanisms were added nobody did: `skew` needs skew_frac and `asym`
# needs gamma_export, both spatial_sim3-only, so both were sent to spatial_sim2, the
# missing-field guard fired, and 5,952 deme cells exited without writing a single file. Nothing
# alerted us, because a mechanism producing no output is indistinguishable from one not yet
# submitted; it surfaced only when the per-mechanism file counts were tallied by hand.
#
# deme_cell now asks the engines which of them can express the mechanism -- see the selection
# block there -- so the list cannot be wrong because there is no list.
_RETIRED_SIM3_ONLY = {"xdrive", "xdrive_c10", "restorers", "restorersAdd"}


# EVERY architecture of Table S3.2, in one registry shared by both frameworks. (11 Sep 2026.)
#
# DR: "Ideally we don't just do baseline, but full Cartesian grid search to see if any parameter
# combinations work." Until now only three of the fifteen Table S3.2 rows had been put through
# the Cartesian pipeline -- mt x Y, mt x autosome dominant, and the restorers -- and the other
# twelve were single-geometry numbers inherited from dm2_run.py. A row that fails at one
# geometry is not a row that fails, so the table could not honestly say FAIL against any of
# them. This registry puts all of them on the same grid, the same filters and the same replicate
# count as Routes 1, 2, 4 and 5.
#
# The kwargs are copied verbatim from dm2_run.SCEN so that a row means the same thing it meant
# in the published table, but the LOCUS INDICES were renumbered on 11 Sep 2026 for the 32+4
# layout: autosomal index 35 = PRDM9-A and the single-pair DMI partner, autosomal index 34 =
# the second autosomal partner, X index 1 = "X locus 2" (X index 0 is the drive locus, 2 the
# neutral background). The old numbering put these at autosomal 31 and 30, which are now
# inside the measured neutral panel. Note that the default dm_partner_locus is an AUTOSOMAL
# index (35): leaving it in place while setting dm_partner="X" indexes past the end of a
# 3-column X array, so every X-partner architecture must name its locus explicitly. bandsim
# raises on this; spatial_sim2 would raise IndexError.
DMI_ARCH = {
    "dmi_mtY":        dict(dm=1.0, dm_focal="mt", dm_partner="Y"),
    "dmi_mtAutoDom":  dict(dm=1.0, dm_focal="mt", dm_partner="auto",
                           dm_partner_locus=35, dm_dom="dominant"),
    "dmi_mtAutoAdd":  dict(dm=1.0, dm_focal="mt", dm_partner="auto",
                           dm_partner_locus=35, dm_dom="additive"),
    "dmi_mtAutoRec":  dict(dm=1.0, dm_focal="mt", dm_partner="auto",
                           dm_partner_locus=35, dm_dom="recessive"),
    "dmi_mtXDom":     dict(dm=1.0, dm_focal="mt", dm_partner="X",
                           dm_partner_locus=1, dm_dom="dominant"),
    # modern X x archaic Y -- the Campbell & Nachman configuration. It sterilises F1 males but
    # is invadable through their SISTERS, who carry the paternal archaic X into the modern
    # matriline; from the next generation an archaic-fathered son can be archaic-X + archaic-Y
    # and fully fertile. This is the architecture the note's own Table S3.2 records as "does not
    # retain Y", which is why the merged Route 2 cannot be labelled an X-Y DMI route.
    "dmi_XY":         dict(dm=1.0, dm_focal="X", dm_focal_locus=1, dm_partner="Y"),
    "dmi_autoAuto":   dict(dm=1.0, dm_focal="auto", dm_focal_locus=35, dm_partner="auto",
                           dm_partner_locus=34, dm_dom="dominant"),
    # modern X locus 2 x archaic PRDM9-A, the orientation that sterilises F1 males.
    "dmi_Xprdm9":     dict(dm=1.0, dm_focal="X", dm_focal_locus=1, dm_partner="auto",
                           dm_partner_locus=35, dm_dom="dominant", s=0.005, sel_locus="auto"),
    # modern PRDM9-A x archaic X locus 2, three strengths of independent positive selection.
    "dmi_prdm9X_s0":  dict(dm=1.0, dm_focal="auto", dm_focal_locus=35, dm_partner="X",
                           dm_partner_locus=1, dm_dom="dominant", s=0.0, sel_locus="none"),
    "dmi_prdm9X_s005": dict(dm=1.0, dm_focal="auto", dm_focal_locus=35, dm_partner="X",
                            dm_partner_locus=1, dm_dom="dominant", s=0.005, sel_locus="auto"),
    "dmi_prdm9X_s05": dict(dm=1.0, dm_focal="auto", dm_focal_locus=35, dm_partner="X",
                           dm_partner_locus=1, dm_dom="dominant", s=0.05, sel_locus="auto"),
}
# The polygenic architectures take the restorer count as their mechanism value, so they are
# handled separately rather than in the registry above.
RESTORER_ARCH = {"restorers": "dominant", "restorersAdd": "additive"}

# ---------------------------------------------------------------------------------------------
# THE FIVE CULTURAL MECHANISMS, ROUTED THROUGH THIS RUNNER. 12 Sep 2026.
#
# These have always existed as their own scripts -- kin2_run, mech3_run, mech4_run, mech5_run,
# clan_run -- each with its own output format, and NONE of them records the duration of the
# range expansion. That single omission has three consequences the note is currently living
# with:
#
#   1. the five cultural panels of Figure S3.3 are empty, because a point cannot be placed on
#      an (autosomal ancestry, generations) plane without the second coordinate;
#   2. the note's claim that the best of them reaches "11% in the deme framework and 18% in the
#      band framework" has no band number behind it at all -- every one of those scripts is
#      deme-only;
#   3. their acceptance can only be tested on autosomal ancestry, so they are screened on one
#      of the note's three criteria rather than all three.
#
# Routing them through robust_run fixes all three at once, because stage 1 records end_gen and
# establishment and stage 2 records p_joint, in whichever framework is asked for. Both engines
# already have the parameters; what was missing was only the dispatch.
#
# THE PARAMETER VALUES ARE THE ONES THE ORIGINAL SCRIPTS SWEPT, not new inventions. `val` is
# the one quantity each mechanism varies, so that a mechanism occupies one column of the
# thumbnail grid exactly as sY and theta do.
CULTURAL = {
    # val = 1 forbids a pair sharing a mother, 2 a pair sharing a grandmother (kin2/kin3)
    "inbreed": dict(deme=lambda v: dict(inbreed_ban=int(v)),
                    band=lambda v: dict(inbreed_ban=int(v))),
    # val = the dispersing sex's migration rate, against 0.005 for the philopatric sex (mech3)
    #
    # THE BAND ARM, WHICH USED TO BE None. bandsim has no m_male/m_female, and the earlier
    # reading was that this makes the mechanism deme-only: its one sex-specific rate is beta,
    # which is the residence rule and is the thing being held fixed. That is right about beta
    # ALONE and wrong about the pair (e, beta), which between them fix the two per-sex rates
    # exactly. bandsim's step() sets
    #
    #     e_f = 2 e beta          e_m = 2 e (1 - beta)
    #
    # so any (e_f, e_m) is reachable by inverting it: e = (e_f + e_m)/2 and
    # beta = e_f / (e_f + e_m). Pinning e_m to the ladder value and e_f to the same 0.005 the
    # deme arm uses therefore reproduces the deme mechanism rather than renaming the residence
    # axis -- the sweep moves the total exogamy rate as well as its sex bias, which varying
    # beta at fixed e cannot do.
    #
    # WHAT THE BAND ARM IS ACTUALLY TESTING, WHICH IS NOT QUITE THE DEME ARM. The band baseline
    # is already e = 0.08, beta = 0, i.e. obligate male exogamy at a male rate of 0.16 and a
    # female rate of exactly zero. So where the deme arm asks "does it help to make male
    # dispersal obligate?", the band arm asks "at what RATE?", against a residence rule that is
    # male-exogamous to begin with. Both are the same one-parameter family and the ladder is
    # the deme arm's, so the panels are comparable; the note should say which question each
    # framework is answering rather than implying they are identical.
    "obligate": dict(deme=lambda v: dict(m_male=float(v), m_female=0.005),
                     band=lambda v: dict(e=(float(v) + 0.005) / 2.0,
                                         beta=0.005 / (float(v) + 0.005))),
    # val = the probability a son returns to his father's natal group (mech4)
    "rhoreturn": dict(deme=lambda v: dict(rho=float(v)),
                      band=lambda v: dict(rho_return=float(v))),
    # val = the fraction of men who father any children at all (mech5's skew arm)
    "skew": dict(deme=lambda v: dict(skew_frac=float(v)),
                 band=lambda v: dict(skew_frac=float(v))),
    # val = the export discount, with import held at the baseline (asym)
    "asym": dict(deme=lambda v: dict(gamma_export=float(v)),
                 band=lambda v: dict(e_export=float(v))),
}


# WHERE MECHANISMS LIVE, AND WHERE ANCESTRY IS MEASURED. DR, 11 Sep: "all measurement should
# be done on markers not affected by selection, and this should apply to all runs", and then
# "we should always make the measurement on all the loci that were not used for selection".
#
# There are 36 autosomal loci. The TOP FOUR -- 32, 33, 34, 35 -- are reserved for mechanisms
# and are never measured: a single-pair DMI uses locus 35, autosome x autosome uses 34 and 35,
# PRDM9 uses 35, and the restorers take the last n, which at the maximum n = 4 is exactly
# 32-35. Ancestry is measured on the fixed panel of loci 0-31, which no architecture in the
# registry touches. FIXED_MEASUREMENT_SET below makes that panel constant across
# architectures, so a 4-restorer cell and an mt x Y cell are measured on the same 32 loci and
# their autosomal percentages are directly comparable -- the earlier design measured on
# "everything not under selection", which was also unbiased but gave each architecture a
# different-sized panel and therefore a different sampling variance.
#
# FOUR IS THE HARD CEILING ON RESTORERS under this layout. routes_build caps the ladder at 4
# for that reason. A 5- or 6-restorer cell would spill into loci 30-31 and quietly re-create
# the measurement bug; band_reps.excluded_loci raises rather than allowing it, and the legacy
# bandmech_submit REST_N list still contains a 6 that will now refuse to run. Raising the
# ceiling means raising AUTOS, not reusing a measured locus.
#
# All 36 loci segregate independently (spatial_sim3 line 222 draws each separately from each
# parent), so the measured loci are unlinked from the mechanism throughout.
MECH_LOCI = (32, 33, 34, 35)      # reserved; loci 0-31 are the neutral panel
FIXED_MEASUREMENT_SET = True      # DR's decision, 11 Sep: measure on EVERY locus not used for
                                  # selection. Unbiased either way, and using all available
                                  # unaffected loci is strictly the less noisy choice; the
                                  # varying number of measurement loci is described in the
                                  # methods rather than engineered away. MECH_LOCI is retained
                                  # only to document that mechanisms live on chromosomes 5-8.


def _excluded_loci(cfg, mod):
    """Autosomal and X loci that must NOT count towards background ancestry.

    DR, 11 Sep 2026: "DMI should be mediated by chromosome 8 and autosomal ancestry measured
    on chromosomes 1-7." That is exactly what spatial_sim2.run() does -- it builds this set
    and passes it to _stats. But stage 2 here does NOT call run(); it drives the generation
    loop itself so that it can stop at marker absorption, and it was calling _stats(P) with no
    exclusions at all. A DMI drives its own autosomal partner to ~100% modern, so including
    that locus in an eight-locus mean adds a fixed 12.5 percentage points of pure artefact.
    That is the whole of the "2.2x drift after the sweep" I reported: back-correcting it moves
    the stage-1 to stage-2 ratio from 2.12 to 0.97, i.e. the martingale holds exactly, as it
    always should have.
    """
    excl, excl_x = set(), set()
    if FIXED_MEASUREMENT_SET:
        excl.update(MECH_LOCI)
    if getattr(cfg, "sel_locus", "none") == "auto":
        excl.add(cfg.auto_locus)
    if getattr(cfg, "dm", 0) > 0:
        if cfg.dm_focal == "auto":
            excl.add(cfg.dm_focal_locus)
        if cfg.dm_partner == "auto":
            excl.add(cfg.dm_partner_locus)
        if cfg.dm_focal == "X":
            excl_x.add(cfg.dm_focal_locus)
        if cfg.dm_partner == "X":
            excl_x.add(cfg.dm_partner_locus)
    if getattr(cfg, "sel_locus", "none") == "X":
        excl_x.add(cfg.x_locus)
    if getattr(cfg, "restorers", 0) > 0:
        excl.update(range(mod.AUTOS - cfg.restorers, mod.AUTOS))
    if getattr(cfg, "xdrive", 0) > 0:
        excl_x.add(getattr(mod, "DRIVER_XLOCUS", 0))
    return tuple(sorted(excl)), tuple(sorted(excl_x))


# ------------------------------------------------------------------ deme
def _mech_kwargs(mech, val):
    """The Config keyword arguments a deme mechanism sets. ONE definition, two callers.

    deme_cell needs this twice: once before choosing an engine, to ask which engine can express
    the mechanism, and once after, to apply it. Writing the mapping out twice is how the
    engine-selection list went stale and lost 5,952 cells, so it is written once here.
    """
    if mech == "sY":
        return dict(s=val, sel_locus="Y")
    if mech == "theta":
        return dict(theta_f1m=val)
    if mech == "xdrive":
        return dict(xdrive=val)
    if mech == "xdrive_c10":
        # Table S3.1 reports every drive strength both with and without a 10% fertility cost to
        # expressing males, but only the cost-free arm was ever put on the Cartesian grid.
        return dict(xdrive=val, xdrive_cost=0.10)
    if mech in DMI_ARCH:
        return dict(DMI_ARCH[mech])
    if mech in RESTORER_ARCH:
        return dict(dm=1.0, dm_focal="mt", dm_partner="auto", restorers=int(val),
                    dm_dom=RESTORER_ARCH[mech])
    if mech == "twowave":
        # Route 5: relabel the landscape at generation `val` and found a fresh modern group
        # under the opposite residence rule. See the wave-2 block in spatial_sim2.
        return dict(wave2_gen=int(val), wave2_lineage="patrilineal")
    if mech in CULTURAL:
        return dict(CULTURAL[mech]["deme"](val))
    if mech == "none":
        return {}
    raise SystemExit(f"unknown mechanism {mech}")


def deme_cell(mech, val, stage, reps, seed0, kw):
    # THE ENGINE IS CHOSEN BY WHAT THE MECHANISM NEEDS, NOT BY A HAND-KEPT LIST.
    #
    # SIM3_ONLY was that hand-kept list, and it went out of date the moment the five cultural
    # mechanisms were added: `skew` needs skew_frac and `asym` needs gamma_export, both of
    # which exist only in spatial_sim3, and neither was in the list. So both were routed to
    # spatial_sim2, the missing-field guard below fired, and all 5,952 deme cells exited
    # without writing a file. The guard did exactly its job -- it failed loudly instead of
    # silently running the null, which is the failure it was written for after the X-drive
    # episode -- but nothing had to notice, because a mechanism that produces no output looks
    # identical to one that has not been submitted yet. It took counting files per mechanism
    # to see it.
    #
    # Asking the engines which one can express the mechanism removes the list, and with it the
    # possibility of the list being wrong. spatial_sim3 is a strict superset of spatial_sim2,
    # so preferring sim2 when it suffices keeps every existing result bit-identical.
    _needs = set(_mech_kwargs(mech, val))
    import spatial_sim2 as _s2
    import spatial_sim3 as _s3
    if _needs <= set(_s2.Config.__dataclass_fields__):
        S2, Config = _s2, _s2.Config
    elif _needs <= set(_s3.Config.__dataclass_fields__):
        S2, Config = _s3, _s3.Config
    else:
        raise SystemExit(
            f"mechanism {mech!r} needs Config field(s) "
            f"{sorted(_needs - set(_s3.Config.__dataclass_fields__))}, which neither engine "
            f"has")
    # NB no `reseed` key: spatial_sim2/3 spell it `reseed_origin_after_extinction` and both
    # default to False, which is the settled no-reseeding policy. An earlier draft passed
    # reseed="guarded", which is not a field and was silently dropped -- harmless, because the
    # default was already what we wanted, but it made the docstring a lie. Left explicit now.
    base = dict(grid=kw["grid"], K=kw["K"], m=kw["m"], gamma=kw["gamma"], r=kw["r"],
                lineage=kw.get("lineage", "matrilineal"), gens=CAP + 10, sample_gen=CAP,
                record_every=10 ** 9, reseed_origin_after_extinction=False,
                # THE RESIDENCE AND mtDNA-SELECTION AXES, added 11 Sep 2026. female_bias is the
                # probability that a cross-lineage child takes its mother's lineage: 1.0 is
                # full matrilocality (Figure 2b's "fexp"), 0.5 is the unbiased expansion
                # ("nexp"). s_mt is selection against archaic mtDNA, drawn from the Muller's-
                # ratchet posterior. DR: "for the non-sex-biased scenarios mtDNA selection a la
                # Muller's Ratchet is required" -- correct, and it is why the unbiased arm needs
                # this axis to appear in the Cartesian search at all: at s_mt = 0 the mtDNA of
                # an unbiased expansion is a pure martingale and p_joint cannot exceed ~5%
                # however strong the mechanism acting on the Y.
                female_bias=kw.get("female_bias", 1.0), s_mt=kw.get("s_mt", 0.0))
    mech_kw = _mech_kwargs(mech, val)
    base.update(mech_kw)
    mech_keys = set(mech_kw)

    # THE GUARD. Fail loudly rather than silently running the null.
    missing = sorted(k for k in mech_keys if k not in Config.__dataclass_fields__)
    if missing:
        raise SystemExit(f"mechanism {mech!r} needs Config field(s) {missing}, which "
                         f"{Config.__module__} does not have -- this cell would silently run "
                         f"the null. Add the field or route {mech!r} to an engine that has it.")

    rows = []
    for s in range(seed0, seed0 + reps):
        cfg = Config(**{k: v for k, v in dict(base, seed=s).items()
                        if k in Config.__dataclass_fields__})
        if stage == 1:
            cfg = Config(**{**{k: v for k, v in dict(base, seed=s, stop_after_ext=1).items()
                               if k in Config.__dataclass_fields__}})
            ts, ext = S2.run(cfg)
            if ext is None:
                rows.append(dict(seed=s, established=0)); continue
            # Y and mtDNA means are recorded even in the screen: DR's Supplementary Table 3
            # wants mean Y proportion and mean mtDNA proportion as columns alongside the
            # autosomal one, for every row.
            rows.append(dict(seed=s, established=1, end_gen=ext,
                             aut=100 * float(ts["auto_mod"][-1]),
                             Y=100 * float(ts["Y_mod"][-1]),
                             mt=100 * float(ts["mt_mod"][-1])))
        else:
            rng = np.random.default_rng(cfg.seed)
            ex, ex_x = _excluded_loci(cfg, S2)
            P = S2._init(cfg, rng); ext = None; g = 0
            cur = cfg
            w2 = int(getattr(cfg, "wave2_gen", 0) or 0)
            for g in range(CAP):
                # THE WAVE-2 BLOCK, WHICH THIS LOOP NEVER HAD (13 Sep 2026). Stage 2 does not
                # call S2.run(); it reimplements the generation loop out of _migrate/_repro/
                # _reseed_origin so it can stop on marker absorption rather than on a fixed
                # sampling generation. That reimplementation omitted the second wave entirely,
                # so EVERY deme two-wave cell was scored as a plain single matrilocal wave --
                # mtDNA 100%, Y 0%, joint 0 -- in all 200 replicates of all 448 cells. The
                # band path has carried this block since 12 Sep (see band_cell); the deme path
                # never did, which is why stage 1 gives sensible two-wave durations (~587
                # generations, via S2.run) while stage 2 disagreed with it completely.
                #
                # Mirrors spatial_sim2.run's own block: relabel everyone resident, keeping
                # their alleles, and found a fresh modern group at the origin under the
                # opposite residence rule. female_bias must be switched as well as lineage --
                # _repro ignores cfg.lineage whenever female_bias is set, and robust_run
                # always sets it, which was the 12 Sep bug in the engine's own copy.
                if w2 and g == w2:
                    P["lin"][:] = 0
                    # P is a dict of arrays in spatial_sim2, not a structured array -- the
                    # band engine's P is the structured one, and copying its idiom across
                    # raised AttributeError on the first replicate.
                    if "f1m" in P:
                        P["f1m"][:] = False
                    here = P["pos"] == 0
                    if here.any():
                        P["lin"][here] = 1
                        P["mt"][here] = 1
                        male = here & (P["sex"] == 1)
                        P["Y"][male] = 1
                        P["auto"][here, :] = 2
                        P["X"][here, :] = np.where(P["sex"][here, None] == 0, 2, 1)
                    fb2 = S2._FB_OF_MODE.get(getattr(cfg, "wave2_lineage", "patrilineal"))
                    cur = replace(cur, lineage=getattr(cfg, "wave2_lineage", "patrilineal"))
                    if getattr(cfg, "female_bias", None) is not None and fb2 is not None:
                        cur = replace(cur, female_bias=fb2)
                    ext = None          # the first wave's extinction is not this run's answer
                S2._migrate(P, cur, rng)
                nxt = S2._repro(P, cur, rng)
                if nxt is None:
                    break
                P = S2._reseed_origin(nxt, cur)
                if ext is None and (P["lin"] == 0).sum() == 0:
                    ext = g
                # THE SAME WAVE-2 GUARD THE OTHER THREE EARLY STOPS CARRY. Without it the
                # absorption test fires on wave one's absorbed markers and the run ends before
                # generation w2, which is the single-wave answer by another route.
                if ext is not None and g % 25 == 0 and (not w2 or g > w2):
                    st = S2._stats(P, ex, ex_x)
                    if st["mt_mod"] in (0.0, 1.0) and st["Y_mod"] in (0.0, 1.0):
                        break
            st = S2._stats(P, ex, ex_x)
            q = (st["mt_mod"] if st["mt_mod"] == st["mt_mod"] else 0.0)
            qy = (st["Y_mod"] if st["Y_mod"] == st["Y_mod"] else 0.0)
            rows.append(dict(seed=s, established=int(ext is not None), end_gen=g,
                             aut=100 * float(st["auto_mod"]),
                             Y=100 * float(qy), mt=100 * float(q),
                             joint=float((q * qy) ** NSAMPLE)))
    return rows


# ------------------------------------------------------------------ band
def band_cell(mech, val, stage, reps, seed0, kw):
    import bandsim as _bandmod
    from bandsim import Config, Landscape, init_state, step, stats
    # sigma_b, sigma_e and alpha_d were fixed at their defaults until 10 Sep. DR: "we'll want
    # to start testing all the parameters we weren't varying before". Note that sigma_e does
    # double duty -- it is both the exogamous dispersal scale and the kernel width for density
    # regulation -- so varying it moves two things at once, which matters for interpretation.
    base = dict(N_target=kw["N"], K=kw["K"], e=kw["e"], p_move=kw["p_move"], r=kw["r"],
                beta=kw.get("beta", 0.0), sigma_b=kw.get("sigma_b", 60),
                sigma_e=kw.get("sigma_e", 80), alpha_d=kw.get("alpha_d", 0.8),
                # beta is the FEMALE share of exogamy propensity, so 0.0 is full matrilocality
                # (Figure 2b's "fexp") and 0.5 is the unbiased expansion ("nexp"). The deme
                # equivalent is female_bias = 1.0 and 0.5 respectively -- opposite ends of
                # differently-oriented scales, which is easy to get backwards.
                s_mt=kw.get("s_mt", 0.0),
                # gens: 2,500 is ample for ONE band-model sweep (median completion ~590) but
                # not for two. The two-wave route founds its second expansion at generation 500,
                # 1,000 or 2,000 and then needs another full sweep, so at 2,500 the later rungs
                # had no room and every replicate came back unestablished once the wave-2 guard
                # stopped them exiting early. 6,000 leaves a full sweep after the latest rung.
                # The deme engine has always run to 7,003 for the same reason.
                gens=(6000 if mech == "twowave" else 2500), record_every=10 ** 9)
    if mech == "sY":
        base.update(s=val)
    elif mech == "theta":
        base.update(theta=val)
    elif mech == "xdrive":
        base.update(xdrive=val)
    elif mech == "xdrive_c10":
        base.update(xdrive=val, xdrive_cost=0.10)
    elif mech in DMI_ARCH:
        # LOCUS-SPECIFIC SELECTION IS NOW AVAILABLE IN THE BAND ENGINE -- CHANGED 14 Sep 2026.
        # bandsim's `s` used to mean selection against the archaic Y and nothing else, so the
        # four PRDM9 architectures -- the ones carrying independent positive selection on
        # PRDM9-A -- could not be expressed here and this branch raised rather than passing
        # kwargs that would have been silently reinterpreted as selection on the Y, which is
        # Route 1 and not a DMI at all. bandsim now carries sel_locus / auto_locus / x_locus
        # with spatial_sim2's semantics (see its _archaic_count), so the architecture
        # dictionary passes through unchanged and a band cell means what the deme cell means.
        arch = DMI_ARCH[mech]
        base.update(arch)
    elif mech in RESTORER_ARCH:
        base.update(dm=1.0, dm_focal="mt", dm_partner="auto", restorers=int(val),
                    dm_dom=RESTORER_ARCH[mech])
    elif mech in CULTURAL:
        maker = CULTURAL[mech]["band"]
        if maker is None:
            # Obligate dispersal is sex-specific MIGRATION, and the band model has no such
            # parameter: movement is by whole bands, and the only sex-specific rate it has is
            # beta, the sex bias of exogamy -- which IS the residence rule and is therefore
            # already the thing being held fixed. Expressing "obligate dispersal" as beta
            # would silently re-run the residence axis under a different name.
            raise SystemExit(f"{mech!r} is sex-specific migration, which bandsim does not have "
                             f"(its only sex-specific rate, beta, is the residence rule "
                             f"itself). Deme-only mechanism.")
        got = maker(val)
        base.update(got)
        # THE SUMMARY MUST REPORT WHAT RAN, NOT WHAT WAS ASKED FOR. main() writes every swept
        # axis into the summary straight out of `kw`, which is the command line. A mechanism
        # that OVERRIDES a swept axis -- obligate sets both e and beta -- would therefore be
        # recorded under the command-line defaults, and every figure that colours or filters
        # band cells by exogamy rate would be reading a number the simulation never used.
        kw.update({k: v for k, v in got.items() if k in kw})
    elif mech not in ("none", "twowave"):
        # Without this, an unrecognised mechanism silently runs the null and returns a
        # plausible number -- the failure that voided every X-drive result on 10 Sep.
        raise SystemExit(f"band_cell does not implement mechanism {mech!r}")

    wave2_gen = int(val) if mech == "twowave" else 0
    rows = []
    land = Landscape()
    for s in range(seed0, seed0 + reps):
        cfg = Config(**dict(base, seed=s))
        rng = np.random.default_rng(cfg.seed)
        P, pos, b = init_state(cfg, land, rng)
        sweep = None
        ex_b, _ = _excluded_loci(cfg, _bandmod)
        # `failed` must be a per-REPLICATE flag. It used to be inferred from
        # `rows[-1].get("established") == 0`, which is the previous replicate's row at the point
        # the test runs, because this replicate's row has not been appended yet. So after the
        # first establishment failure the test stayed true for every later seed and the loop
        # skipped the rest of the replicates forever. The signature is n_est == n - 1, which
        # held in 48/48 band stage-2 cells and 35/70 stage-1 cells before this fix.
        failed = False
        seed_xy = np.array(cfg.seed_lonlat, float)
        for g in range(cfg.gens):
            if wave2_gen and g == wave2_gen:
                # Route 5: relabel every band resident, keeping alleles, and found a fresh
                # modern band at the Bosphorus under the opposite residence rule. Only
                # identity is reset -- the residents carry forward what the first wave left.
                b[:] = False
                j = int(np.argmin(((pos - seed_xy) ** 2).sum(1)))
                b[j] = True
                mem = P["band"] == j
                if mem.any():
                    P["mt"][mem] = 1
                    P["Y"][mem & (P["sex"] == 1)] = 1
                    P["auto"][mem, :] = 2
                    P["X"][mem, :] = np.where(P["sex"][mem] == 1, 1, 2)[:, None]
                import copy as _c
                cfg = _c.copy(cfg); cfg.beta = 1.0
                sweep = None
            out = step(P, pos, b, cfg, land, rng)
            if out[0] is None:
                rows.append(dict(seed=s, established=0)); failed = True; break
            P, pos, b = out
            if sweep is None and b.all():
                sweep = g
                # THE WAVE-2 GUARD, which the band path never had. spatial_sim2 carries it on
                # both of its early stops (lines 697 and 720) precisely because without it the
                # two-wave route cannot score above zero however the parameters are set: wave 1
                # completes, the loop exits, and wave 2 never fires. The band path breaks here
                # at stage 1 and at the absorption test below, and did so unguarded -- so every
                # `tw` band cell was screened, and then scored, as a SINGLE matrilocal wave.
                # That is why Route 4's band triangles all sat at zero: under matrilocality
                # alone mtDNA fixes and the Y is lost, so mtY_male is 0 and joint is 0**10.
                if stage == 1 and (not wave2_gen or g > wave2_gen):
                    break
            if (stage == 2 and sweep is not None and g % 25 == 0
                    and (not wave2_gen or g > wave2_gen)):
                st = stats(P, b, ex_b)
                if st["mt"] in (0.0, 1.0) and st["Y"] in (0.0, 1.0):
                    break
        if failed:
            continue
        st = stats(P, b, ex_b)
        rec = dict(seed=s, established=int(sweep is not None), end_gen=(sweep or g),
                   aut=100 * float(st["auto_mod"]),
                   Y=100 * float(st["Y_mod"]), mt=100 * float(st["mt_mod"]))
        if stage == 2:
            rec["joint"] = float(st["mtY_male"] ** NSAMPLE)
        rows.append(rec)
    return rows


def main():
    fw, mech, val, stage, reps, seed0, out = (
        sys.argv[1], sys.argv[2], float(sys.argv[3]), int(sys.argv[4]),
        int(sys.argv[5]), int(sys.argv[6]), sys.argv[7])
    kw = dict(grid=flag("--grid", int, 12), K=flag("--K", int, 11),
              m=flag("--m", float, 0.10), gamma=flag("--gamma", float, 0.12),
              r=flag("--r", float, 2.0), N=flag("--N", int, 1750),
              e=flag("--e", float, 0.19), p_move=flag("--p-move", float, 0.45),
              beta=flag("--beta", float, 0.0),
              sigma_b=flag("--sigma-b", float, 60.0),
              sigma_e=flag("--sigma-e", float, 80.0),
              alpha_d=flag("--alpha", float, 0.8),
              female_bias=flag("--female-bias", float, 1.0),
              s_mt=flag("--s-mt", float, 0.0),
              lineage=("patrilineal" if "--patri" in sys.argv else "matrilineal"))
    rows = (deme_cell if fw == "deme" else band_cell)(mech, val, stage, reps, seed0, kw)
    os.makedirs(os.path.dirname(out) or ".", exist_ok=True)
    est = [r for r in rows if r.get("established")]
    summary = dict(framework=fw, mech=mech, val=val, stage=stage, reps=reps,
                   n_est=len(est), n=len(rows),
                   aut=(float(np.mean([r["aut"] for r in est])) if est else float("nan")),
                   end_gen=(float(np.mean([r["end_gen"] for r in est])) if est else float("nan")),
                   joint=(float(np.mean([r.get("joint", 0.0) for r in est])) if est and stage == 2
                          else None),
                   # every swept axis is recorded, so a figure can split on any of them without
                   # having to re-derive it from the filename. sigma_b/sigma_e/alpha_d and the
                   # two new demographic axes were missing, which meant the band scatter could
                   # not be coloured by dispersal scale even though it had been swept.
                   **{k: kw[k] for k in ("grid", "K", "m", "gamma", "r", "N", "e", "p_move",
                                         "beta", "lineage", "sigma_b", "sigma_e", "alpha_d",
                                         "female_bias", "s_mt")})
    with open(out, "w") as f:
        json.dump(dict(summary=summary, rows=rows), f)
    print(f"wrote {out}: est {len(est)}/{len(rows)}  aut {summary['aut']:.2f}%  "
          f"gen {summary['end_gen']:.0f}" +
          (f"  joint {summary['joint']:.3f}" if summary["joint"] is not None else ""))


if __name__ == "__main__":
    main()
```
