## Supplementary Information section 3 code base for "Hypothesis: A modern human range expansion ∼300,000 years ago had a critical role in Neandertal origins": robust_submit.py

```
#!/usr/bin/env python3
"""
robust_submit.py -- SLURM array for the robustness sweep. (DR, 10 Sep 2026)

DR, 10 Sep: "start with very few reps and possibly cruder search grids so we can do fast
exploration". So the defaults here are deliberately thin: 4 replicates, a coarse grid, stage 1
only. Turn both up once the shape of the answer is visible.

THE SEARCH IS OVER A RIDGE, NOT A VOLUME
----------------------------------------
DR's Figure S2.3 made the point that matters for cost: once gamma is calibrated so that the
autosomal proportion hits its target, the dependence of Y fixation on theta is essentially
INDEPENDENT of m -- all the m curves lie on top of one another. The allowable parameter space is
a two-dimensional ridge inside the three-dimensional (theta, gamma, m) box, and calibrate5.py
already knows how to find it: local linear regression of log(autosomal ancestry) on
(theta, log gamma, log m) with a Gaussian kernel, inverted for gamma at the target. Leave-one-out
R-squared was 0.97.

So the workflow is: crude grid -> fit the surface -> invert to get the ridge -> only then spend
replicates, and spend them along the ridge. That is what makes the whole thing affordable.

    CAVEAT. The published ridge was fitted at the superseded 20x20 / K=20 geometry AND before
    the origin-reseeding fix, so it has to be refitted. The METHOD carries over; the numbers do
    not.

STAGES
------
  1  screen   stop at archaic extinction; record autosomal % and completion generation.
              24x faster than a full run in the deme model. This is the grid that feeds the
              surface fit.
  2  score    run to absorption of both markers; record P(10 sampled males all modern at both).
              Run only at ridge points that passed the screen.

GRANULARITY. The pilot is ~90 seconds of compute for the whole deme grid, so one array task per
cell is nonsense: SLURM's per-task overhead (~20 s of queueing and start-up) is larger than the
work. --chunk packs many cells into one task. The default aims at roughly five minutes of work
per task, which is the point where overhead stops mattering and jobs still schedule quickly.

    python robust_submit.py 1 [reps] [batch] [cap] [partition] [--band] [--resume]
                              [--chunk N]      # cells per array task; 0 = auto
                              [--serial]       # one job for everything, no array
    sbatch robust_array.sh
    python robust_collect.py 1
"""
import itertools
import os
import sys

HERE = os.path.dirname(os.path.abspath(__file__))

# ---- grids. Widened after the 10 Sep pilot (156 cells, 4 reps), which showed two things:
#
#   * gamma had to go higher. At the reduced-fertility settings the autosomal proportion at the
#     old top of the grid (gamma = 0.55) was still 5.9-21.9%, so the 5% ridge lay OUTSIDE the
#     box and the inversion found ridge points only for theta = 1. That is the contact-budget
#     ceiling showing up as a grid-boundary artefact. gamma now runs to 1.0 (random mating).
#   * m = 0.03 is not worth carrying. With F1 males removed the wave crawls -- 4341 generations
#     at theta = 0.02, 4981 at theta = 0 -- so those cells fail the 100-1000 generation window
#     no matter what gamma does.
#
# The surface fit was already good enough on the crude grid (leave-one-out R^2 = 0.933), so the
# extra density here is for the inversion to have somewhere to land, not for the smoother.
DEME = dict(
    gamma=[0.04, 0.08, 0.12, 0.20, 0.30, 0.42, 0.55, 0.70, 0.85, 1.00],
    m=[0.05, 0.08, 0.12, 0.15],
    r=[2.0],
    K=[11],
    grid=[12],
)
BAND = dict(
    e=[0.08, 0.13, 0.19, 0.26, 0.35, 0.45, 0.56],
    p_move=[0.45],
    r=[2.0],
    K=[25],
    N=[1750],
)

# mechanism, values. "none" is the null that calibrates the autosomal surface itself.
MECHS = [
    ("none", [0.0]),
    ("sY", [0.01, 0.05]),
    ("theta", [0.0, 0.02, 0.1, 1.0]),
    ("xdrive", [0.2, 0.8]),
    ("dmi_mtY", [1.0]),
    ("dmi_mtAutoDom", [1.0]),
    ("restorers", [2, 3]),
]

GEOM = ("--grid", "--K", "--m", "--gamma", "--r", "--N", "--e", "--p-move", "--beta")


def cells(band):
    grid = BAND if band else DEME
    keys = list(grid)
    for mech, vals in MECHS:
        for v in vals:
            for combo in itertools.product(*(grid[k] for k in keys)):
                yield mech, v, dict(zip(keys, combo))


def tag(fw, mech, v, kw):
    s = f"{fw}_{mech}_{v:g}_" + "_".join(f"{k}{kw[k]:g}" for k in sorted(kw))
    return s.replace(".", "p").replace("-", "m")


def main():
    stage = int(sys.argv[1]) if len(sys.argv) > 1 else 1
    args = [a for a in sys.argv[2:] if not a.startswith("--")]
    reps = int(args[0]) if args else 4
    batch = int(args[1]) if len(args) > 1 else 1
    cap = int(args[2]) if len(args) > 2 else 0
    part = args[3] if len(args) > 3 else "short"
    band = "--band" in sys.argv
    resume = "--resume" in sys.argv
    serial = "--serial" in sys.argv
    chunk = int(sys.argv[sys.argv.index("--chunk") + 1]) if "--chunk" in sys.argv else 0
    fw = "band" if band else "deme"
    outdir = f"out_robust{stage}_{fw}"
    os.makedirs(outdir, exist_ok=True)
    os.makedirs("logs", exist_ok=True)

    flagmap = {"grid": "--grid", "K": "--K", "m": "--m", "gamma": "--gamma", "r": "--r",
               "N": "--N", "e": "--e", "p_move": "--p-move", "beta": "--beta"}
    tasks, skipped = [], 0
    for mech, v, kw in cells(band):
        t = tag(fw, mech, v, kw)
        for s0 in range(0, reps, batch):
            n = min(batch, reps - s0)
            out = f"{outdir}/{t}_s{s0}.json"
            if resume and os.path.exists(out):
                skipped += 1
                continue
            fl = " ".join(f"{flagmap[k]} {kw[k]:g}" for k in sorted(kw))
            tasks.append(f"{fw} {mech} {v:g} {stage} {n} {s0} {out} {fl}")
    if not tasks:
        print("nothing to do"); return

    # pack cells into chunks so each array task is a sensible size
    per_rep = 0.14 if (stage == 1 and not band) else (1.7 if band else 0.55)
    if serial:
        chunk = len(tasks)
    elif chunk == 0:
        secs_per_task = per_rep * batch
        chunk = max(1, min(len(tasks), int(300 / max(secs_per_task, 0.01))))
    groups = [tasks[i:i + chunk] for i in range(0, len(tasks), chunk)]
    # One CELL per line, no delimiters -- an earlier draft packed several cells per line with a
    # ';;' separator, which needs IFS parsing in bash, and IFS is a set of single characters so
    # ';;' silently means ';'. Each array task instead runs a contiguous RANGE of lines.
    with open("robust_params.txt", "w") as f:
        f.write("\n".join(tasks) + "\n")

    ncell = len(list(cells(band)))
    per = per_rep
    cpu = ncell * reps * per / 3600.0
    n = len(groups)
    capn = min(n, cap) if cap else 0
    hdr = f"""#!/bin/bash
#SBATCH -J robust{stage}{fw[0]}
#SBATCH -p {part}
#SBATCH -t 0-02:59
#SBATCH -c 1
#SBATCH --mem=3G
#SBATCH -a 0-{n-1}{('%' + str(capn)) if capn else ''}
#SBATCH -o logs/robust_%A_%a.log
"""
    body = ("CHUNK=" + str(chunk) + "\n"
            "START=$((SLURM_ARRAY_TASK_ID * CHUNK + 1))\n"
            "END=$((START + CHUNK - 1))\n"
            "sed -n \"${START},${END}p\" robust_params.txt | while read -r LINE; do\n"
            "  [ -z \"$LINE\" ] && continue\n"
            "  python robust_run.py $LINE\n"
            "done\n")
    open("robust_array.sh", "w").write(hdr + body)
    print(f"  framework {fw}, stage {stage}, {reps} reps")
    print(f"  {ncell} cells -> {len(tasks)} cell-jobs packed into {n} array task(s), "
          f"{chunk} per task" + (f" ({skipped} skipped by --resume)" if skipped else ""))
    print(f"  ~{per:.2f} s per replicate -> {cpu*3600:.0f} CPU-seconds total "
          f"(~{cpu*3600/max(n,1):.0f} s per task)")
    print(f"  wrote robust_params.txt and robust_array.sh")
    print(f"  submit:  sbatch robust_array.sh")
    print(f"  then:    python robust_collect.py {stage} {'--band' if band else ''}")


if __name__ == "__main__":
    main()
```
