## Supplementary Information section 3 code base for "Hypothesis: A modern human range expansion ∼300,000 years ago had a critical role in Neandertal origins": routes_build.py

```
#!/usr/bin/env python3
"""
routes_build.py -- simulations behind the Routes 1/2/4 scatter in Figure 2b. (DR, 10 Sep 2026)

DR wants the three cleanest routes first, in the panel order already settled for Figure 2b:

    Route 1  additive selection on the modern Y          (left)     mech "sY"
    Route 2  X-linked sex distorters in F1 males         (middle)   mech "xdrive"
    Route 4  cultural mechanism, F1 males few offspring  (right)    mech "theta"

Route 2 is the mechanism that `robust_run.py` was silently running as the null until today: it
lives in spatial_sim3, and the Config kwarg filter dropped it. Every X-drive number produced
before this run is void. That is why the rerun and the figure are the same job.

TWO STAGES, RESUMABLE
---------------------
Each cell writes one JSON and is skipped if that JSON exists, because a bash call here is capped
at ~2 minutes and background jobs do not survive the call boundary. Run the script repeatedly
until it reports nothing left.

  stage 1  screen   stop at archaic extinction; keep cells with autosomal ancestry in the
                    window, expansion completing in 100-1000 generations, and establishment
                    above 50% (DR: "we should require that they go forward >50% of the time").
  stage 2  score    run to absorption of BOTH uniparental markers -- monomorphic in either
                    direction, which is where the run stops being informative because the
                    autosomes are a martingale from then on. Records p_joint, the fraction of
                    replicates in which ten sampled males are all modern at both markers.

BAND COVERAGE. bandsim implements s and theta but NOT xdrive, so Route 2 has no band points
until that is added. The script says so rather than quietly emitting a half-empty column.

    python routes_build.py 1 [--band] [--reps 10]
    python routes_build.py 2 [--band] [--reps 40]
    python routes_build.py collect
"""
import csv
import glob
import itertools
import json
import os
import subprocess
import sys
import time
from concurrent.futures import ProcessPoolExecutor

HERE = os.path.dirname(os.path.abspath(__file__))

# Mechanism sets. ROUTE_SET picks one; it is an environment variable rather than an argument so
# that routes_submit.py, which imports this module, cannot disagree with routes_build.py about
# which set is in play. Each set has its OWN output directories, so a job for one set can never
# overwrite the params file or the results of a job for another that is still running.
SETS = {
    "main": ({
        # FOUR RUNGS ADDED 13 Sep, TWO AT EACH END. DR, on Figure 2b: "could you give me sims
        # to give red point clouds below 0.01 and above 0.2". The four demographic LINES in
        # that panel already run from s = 0.001 to 0.5; only the Cartesian clouds stopped at
        # 0.01 and 0.20, so the panel showed scatter over the middle of a curve whose ends
        # were drawn but unsupported. 0.002 and 0.005 sit where the matrilocal line is still
        # near zero and 0.35 and 0.50 where it has saturated, so the new clouds bracket the
        # rise rather than extending it.
        # TWO MORE AT THE BOTTOM, 14 Sep. DR: "for Route 1 the red fountains don't exist at
        # 0.001 and 0.0005 but the other curves (yellow/black/blue) exist there, so we either
        # remove the other curves, or add the red fountains." Adding them is the right way
        # round: s = 0 is the null -- no selection on the Y at all -- and dropping the three
        # comparison curves' left ends to hide the gap would remove the one place on the panel
        # where all four scenarios can be compared with the mechanism switched OFF.
        #
        # s = 0 also makes Route 1 symmetric with Route 3, whose ladder has carried its null
        # (theta = 1) and its zero from the start. Both panels then run from "no mechanism" to
        # "mechanism as strong as it can be", and the reader can see where each curve starts.
        "sY":     ("Route 1", "additive selection on Y",
                   [0.0, 0.001, 0.002, 0.005, 0.01, 0.02, 0.05, 0.10, 0.20, 0.35, 0.50]),
        "xdrive": ("Route 2", "X-linked sex distortion", [0.1, 0.2, 0.4, 0.6, 0.8]),
        # FOUR LOWER RUNGS ADDED 12 Sep. DR: "Route 3 dots and traces need to be extended to
        # lower thetas." The baseline curves in joint2_summary already ran down to 0.001; only
        # the Cartesian points stopped at 0.02, so the panel showed a cloud that ended exactly
        # where the mechanism starts to work. Scored medians over admissible combinations are
        # 1.00 at theta = 0 and already 0.34 (deme) / 0.22 (band) by 0.02, so the transition is
        # entirely inside the gap these rungs fill.
        # 0.5 AND 1.0 ADDED 13 Sep. DR: "Give me sims for 0.5 and 1 point clouds". theta = 1 is
        # the null -- F1 males at full fertility, no mechanism at all -- so the cloud there is
        # the control the rest of the panel is read against, and it was the one rung with a
        # line but no points.
        "theta":  ("Route 4", "reduced F1-male fertility",
                   [0.0, 0.001, 0.002, 0.005, 0.01, 0.02, 0.05, 0.10, 0.20, 0.50, 1.00]),
    # ALL of these run in bandsim. An earlier version of this set listed only {"sY","theta"},
    # on my mistaken belief that the band engine lacked the others -- but xdrive, the DM rule,
    # the restorers and the escape test are all implemented there (bandsim.py lines 335, 344,
    # 382, 543). The restriction was mine, not the engine's, and it silently kept Routes 2 and 3
    # out of the band framework entirely.
    }, {"sY", "theta", "xdrive"}, "routes"),
    # Route 3. Both DMI variants and the polygenic restorers live in spatial_sim3; `restorers`
    # was being dropped by the Config kwarg filter in exactly the way `xdrive` was, so every
    # restorer number before 10 Sep was the null model. None of this is implemented in bandsim,
    # so Route 3 is deme-only for now.
    "dmi": ({
        "dmi_mtY":       ("Route 3a", "mt-Y incompatibility", [1.0]),
        "dmi_mtAutoDom": ("Route 3b", "mt-autosome, dominant", [1.0]),
        "restorers":     ("Route 3c", "polygenic restorers", [1, 2, 3, 4]),
    }, {"dmi_mtY", "dmi_mtAutoDom", "restorers"}, "dmi"),
    # EVERY REMAINING TABLE S3.2 ARCHITECTURE, 11 Sep 2026. The `dmi` set above covers three of
    # the fifteen rows; the other twelve were single-geometry numbers carried over from
    # dm2_run.py, so the table could not honestly write FAIL against any of them -- failing at
    # one geometry is not failing. This set puts them on the same Cartesian grid, filters and
    # replicate count as everything else. Separate PREFIX, so it cannot collide with a `dmi`
    # job still on the queue.
    #
    # The three PRDM9 architectures and the modern-X x archaic-PRDM9 orientation carry
    # independent positive selection on a named autosomal locus. That USED to be deme-only,
    # because bandsim's `s` acted on the archaic Y and nothing else; since 14 Sep it has
    # sel_locus and they run in both frameworks. See the note on BAND_SUPPORTS below.
    "dmi2": ({
        "dmi_mtAutoAdd":   ("Route 3", "mt-autosome, additive", [1.0]),
        "dmi_mtAutoRec":   ("Route 3", "mt-autosome, recessive", [1.0]),
        "dmi_mtXDom":      ("Route 3", "mt-X, dominant", [1.0]),
        "dmi_XY":          ("Route 3", "X-Y incompatibility", [1.0]),
        "dmi_autoAuto":    ("Route 3", "autosome-autosome", [1.0]),
        "dmi_Xprdm9":      ("Route 3", "modern X x archaic PRDM9-A", [1.0]),
        "dmi_prdm9X_s0":   ("Route 3", "modern PRDM9-A x archaic X, s=0", [1.0]),
        "dmi_prdm9X_s005": ("Route 3", "modern PRDM9-A x archaic X, s=0.5%", [1.0]),
        "dmi_prdm9X_s05":  ("Route 3", "modern PRDM9-A x archaic X, s=5%", [1.0]),
        "restorersAdd":    ("Route 3", "polygenic restorers, additive", [1, 2, 3, 4]),
    # THE FOUR PRDM9 ARCHITECTURES JOIN THE BAND WHITELIST, 14 Sep 2026. They were excluded
    # because bandsim's `s` was hardwired to the archaic Y, so an architecture carrying
    # independent positive selection on PRDM9-A could not be expressed and robust_run raised
    # on it. The consequence was four thumbnails in Figure S3.3 with blue points and no red --
    # DR: "In Figure S3.3 the following thumbnails are missing red points: the four PRDM9."
    # bandsim now has sel_locus / auto_locus / x_locus with spatial_sim2's semantics, verified
    # to reproduce the old Y-only behaviour bit for bit at sel_locus = "Y", so the four can be
    # screened in the band framework like everything else.
    }, {"dmi_mtAutoAdd", "dmi_mtAutoRec", "dmi_mtXDom", "dmi_XY", "dmi_autoAuto",
        "dmi_Xprdm9", "dmi_prdm9X_s0", "dmi_prdm9X_s005", "dmi_prdm9X_s05",
        "restorersAdd"}, "dmi2"),
    # THE MERGED-FIGURE SET. One mechanism from each surviving route, crossed with the residence
    # rule and the mtDNA-selection ladder, so that all four of Figure 2b's demographic scenarios
    # (or at least both range expansions -- the migration pulses are a different engine) live
    # inside the same Cartesian search as Figure 2c's fountains. That is what lets the two
    # figures be overlaid instead of juxtaposed.
    #
    # The matrilocal arm duplicates cells already computed under the `main` / `dmi` / `tw`
    # prefixes, because the filename tag now carries female_bias and s_mt and so no longer
    # matches. That is deliberate: one sweep, one tagging convention, one figure, rather than a
    # figure stitched from three sweeps whose cells cannot be checked against each other.
    # THE MECHANISM LADDERS ARE THE FIGURE 2b LADDERS. DR: "to make the figures merge properly,
    # we might have to do the full Cartesian grid for a few more values, e.g. down to s = 0,
    # 0.005, 0.002, 0.001 for selection, and maybe another point or two for cultural F1
    # fertility." The reason this matters is not coverage but registration: if the fountains sit
    # at s = 0.05, 0.10, 0.20 and the baseline curves are drawn through s = 0.04, 0.08, 0.16,
    # the reader cannot read a fountain against the curve behind it, and the overlay is
    # decorative. So the s ladder here is exactly the ten values of the band production sweep
    # plus DR's 0.001, and the theta ladder extends past 0.20 to find where the route dies.
    # s = 0 is the null: a fountain at s = 0 IS the no-mechanism scenario, which is what makes
    # the baseline a point in the search rather than an annotation on it.
    "demog": ({
        # BROUGHT ONTO THE MAIN LADDER, 14 Sep. This set used to double above s = 0.02 --
        # 0.04, 0.08, 0.16, 0.32, 0.64 -- while the mechanism screen the red fountains come
        # from uses 0.05, 0.10, 0.20, 0.35, 0.50. Every quantity the two sets are compared on
        # therefore had to be interpolated across the mismatch above s = 0.02. DR: "I don't
        # like interpolation and having to say something sloppy in the final text." 0.64 is
        # dropped with the rest of the doubling tail; it is off the right-hand end of Route
        # 1's axis and is one of the cells jcal_submit records as failing to bracket 5%.
        "sY":      ("Route 1", "additive selection on Y",
                    [0.0, 0.001, 0.002, 0.005, 0.01, 0.02, 0.05, 0.10, 0.20, 0.35, 0.50]),
        # X-DRIVE AND THETA BROUGHT ONTO THE MAIN LADDER TOO, 14 Sep, for the same reason
        # the sY ladder above was. Route 2's unbiased curve ran from 0.1 to 0.8 while its
        # fountains run from 0 to 1.0, so the curve stopped short of the panel at both ends
        # -- and the two ends are where the route is decided: 0 is the null and 1.0 is
        # every child a daughter. Route 3's unbiased curve stopped at theta = 0.02 while the
        # fountains go down to 0.001, which is the whole interesting part of that route.
        "xdrive":  ("Route 2", "X-linked sex distortion",
                    [0.0, 0.05, 0.1, 0.2, 0.4, 0.6, 0.8, 1.0]),
        "dmi_mtY": ("Route 2", "mt-Y incompatibility", [1.0]),
        "theta":   ("Route 3", "reduced F1-male fertility",
                    [0.0, 0.001, 0.002, 0.005, 0.01, 0.02, 0.05, 0.10, 0.20, 0.35, 0.50]),
        "twowave": ("Route 4", "two waves", [500, 1000, 2000]),
    }, {"sY", "xdrive", "dmi_mtY", "theta", "twowave"}, "demog"),
    # The three daughter-proportion rungs DR asked for after seeing the collapsed Route 2
    # panel: d = 0 is 50% daughters (no distortion at all -- the null, and the control that
    # says whether 55% really is doing the work), d = 0.05 is 52.5%, d = 1 is 100% daughters.
    "xd2": ({
        "xdrive": ("Route 2", "X-linked sex distortion", [0.0, 0.05, 1.0]),
    }, {"xdrive"}, "xd2"),
    # The cost-bearing arm of Table S3.1's X-drive rows, which the `main` set omits.
    # LADDER EXTENDED 12 Sep to 0.05 and 1.0 (52.5% and 100% daughters) so that the cost arm
    # covers the same rungs as the cost-free one. Figure S3.3 stacks the two arms in matched
    # columns and DR asked for those two columns filled; without them the comparison has holes
    # at both ends of the range, which are exactly the ends that matter -- the threshold and
    # the ceiling. The five existing values are unchanged, so their cells are already on disk
    # and routes_submit will emit only the two new ones.
    "xdc": ({
        "xdrive_c10": ("Route 2", "X-linked sex distortion, 10% cost",
                       [0.05, 0.1, 0.2, 0.4, 0.6, 0.8, 1.0]),
    }, {"xdrive_c10"}, "xdc"),
    # Route 5. Until now this lived in its own gamma-sweep scripts at a single geometry, which
    # is why its column in Figure 2c had 8 points against ~140 for Routes 1-3. Putting it
    # through the same pipeline puts it on the same Cartesian grid as everything else. The
    # "value" is the generation at which the second wave is founded.
    "tw": ({
        "twowave": ("Route 5", "two waves", [500, 1000, 2000]),
    }, {"twowave"}, "tw"),
}
_set = os.environ.get("ROUTE_SET", "main")
if _set not in SETS:
    raise SystemExit(f"ROUTE_SET={_set!r} unknown; choose from {sorted(SETS)}")
ROUTES, BAND_SUPPORTS, PREFIX = SETS[_set]

# ROUTE_ONLY restricts a set to named mechanisms, so one mechanism can be rerun without
# re-queuing its siblings. Added 9 Oct 2026 for the Scenario 2 rerun: selection on the Y
# became conditional on lineage identity (spatial_sim2._archaic_count), which changes every
# sY cell and no xdrive or theta cell. The old sY outputs must be DELETED first -- cells are
# skipped by filename, so a rerun over existing files does nothing:
#     rm out_routes1_{deme,band}/*_sY_* out_routes1hi_{deme,band}/*_sY_* out_routes2_{deme,band}/*_sY_*
#     ROUTE_SET=main ROUTE_ONLY=sY python routes_submit.py 1 ...
_only = os.environ.get("ROUTE_ONLY")
if _only:
    _keep = set(_only.split(","))
    # A set with none of the named mechanisms is filtered to NOTHING rather than rejected:
    # rescreen_submit.py reloads this module once per set, so a hard error here would abort
    # it on every set that does not contain sY. An empty ROUTES emits zero cells, which is
    # the intent. Typos are still caught, because at least one set must match.
    ROUTES = {k: v for k, v in ROUTES.items() if k in _keep}
    BAND_SUPPORTS = BAND_SUPPORTS & _keep
    if not ROUTES:
        _all = {m for _r, _b, _p in SETS.values() for m in _r}
        if not (_keep & _all):
            raise SystemExit(f"ROUTE_ONLY names {sorted(_keep)} match no mechanism in any "
                             f"set; choose from {sorted(_all)}")

# r added 10 Sep. It is the one genuinely free parameter that was never varied, and it is safe
# to vary WITHOUT waiting on the Ne screen: r is the lineage advantage, so it acts only while
# modern and archaic lineages coexist, whereas Ne is measured after burn-in when the population
# is of one lineage. Geometry (grid, K) stays at the validated 12x12 / K=11, which measures
# Ne = 1488 diploid against the 1447 anchor; widening THAT axis does need the screen.
# The r=2.0 cells already exist and are skipped by name, so this only queues the new r values.
DEME_GRID = dict(gamma=[0.08, 0.12, 0.20, 0.30, 0.42, 0.55, 0.70, 1.00],
                 m=[0.05, 0.08, 0.12, 0.15], r=[1.5, 2.0, 3.0], K=[11], grid=[12])
# p_move is NOT varied: measured at matched p*sigma_b^2 it is redundant with sigma_b, so only
# sigma_b is swept. K and N stay at the anchored values, which are the band-model equivalent of
# the deme geometry constraint and would need their own Ne screen to widen.
BAND_GRID = dict(e=[0.08, 0.13, 0.19, 0.26, 0.35, 0.45, 0.56],
                 sigma_b=[30.0, 60.0, 120.0], sigma_e=[40.0, 80.0, 160.0],
                 alpha_d=[0.4, 0.8, 1.6], r=[1.5, 2.0, 3.0],
                 p_move=[0.45], K=[25], N=[1750])

AUT_LO, AUT_HI = 2.0, 15.0
GEN_LO, GEN_HI = 100.0, 1000.0
EST_MIN = 0.50

FLAG = {"grid": "--grid", "K": "--K", "m": "--m", "gamma": "--gamma", "r": "--r",
        "N": "--N", "e": "--e", "p_move": "--p-move",
        "sigma_b": "--sigma-b", "sigma_e": "--sigma-e", "alpha_d": "--alpha",
        "female_bias": "--female-bias", "beta": "--beta", "s_mt": "--s-mt"}

# THE RESIDENCE / mtDNA-SELECTION AXES. 11 Sep 2026.
#
# DR: "you can merge Figure 2b and 2c by simply using the results of the simulation with the
# baseline parameters for current Figure 2b and overlaying them on current Figure 2c. The
# baseline parameters are a subset of Figure 2c so it should work, right"
#
# Right for ONE of Figure 2b's four scenarios. The Cartesian grid fixes full matrilocality, so
# only the sex-biased range expansion lives inside it; the unbiased expansion and the two
# migration pulses are a different demography and have no representation in it at all. Adding
# the residence axis fixes that for the expansions.
#
# And DR again: "for the non-sex-biased scenarios mtDNA selection a la Muller's Ratchet is
# required". Exactly so, and it is not optional -- with an unbiased residence rule the mtDNA is
# an unselected martingale, so p_joint is capped near the autosomal proportion whatever is done
# to the Y, and an unbiased arm run at s_mt = 0 would search a region that cannot work by
# construction. The s_mt ladder is 0 plus the quartiles and the 95th percentile of the accepted
# ABC posterior in mt_gap_posterior_NA1500.txt (median 0.0024, IQR 0.0011-0.0036).
#
# s_mt is NOT crossed with full matrilocality: there the mtDNA is already 100% modern by
# construction, so the axis would multiply the cost by five and change nothing. The residence
# axis therefore contributes 1 + 5 = 6 settings, not 2 x 5 = 10.
# WHICH RUNG IS THE CENTRAL ESTIMATE CHANGED ON 11 Sep, THE RUNGS DID NOT.
#
# The ladder was built from the quartiles of mt_gap_posterior_NA1500.txt, which is the
# CODING-ONLY ABC posterior (median 0.235%). The Muller's Ratchet note headlines the
# RNA-INCLUSIVE ABC posterior (median 0.32%, giving its 23% fixation probability), and section 3
# should be quoting that one -- the two notes were describing the same event with fitness gaps
# 40% apart. mt_gap_posterior_NA1500_rna.txt now holds it.
#
# Fortunately this needs no rerun. The RNA-inclusive median is 0.33% and its upper quartile
# 0.50%, so the existing rungs 0.0036 and 0.0064 already straddle the central estimate and the
# ladder spans the posterior either way. What changes is only which rung is reported as
# central: 0.0036, not 0.00235. Left as-is so that cells already on disk and the arrays in
# flight stay valid; rebuild from the RNA quartiles only if the whole sweep is ever rerun.
SMT_LADDER = [0.0, 0.0011, 0.00235, 0.0036, 0.0064]
SMT_CENTRAL = 0.0036      # RNA-inclusive ABC median 0.33%, the Muller's Ratchet headline
DEME_RESIDENCE = ([dict(female_bias=1.0, s_mt=0.0)]
                  + [dict(female_bias=0.5, s_mt=v) for v in SMT_LADDER])
BAND_RESIDENCE = ([dict(beta=0.0, s_mt=0.0)]
                  + [dict(beta=0.5, s_mt=v) for v in SMT_LADDER])


def admissible_triples():
    """(grid, K, m) combinations whose MEASURED autosomal Ne is within 10% of the 1447 anchor.

    Geometry is not a free axis: census and Ne are different quantities, and at small K the
    frequent local extinction and recolonisation of demes drives Ne well below census (20x20
    cannot reach the anchor at all). So the demographic grid is a full factorial over
    (grid, K, m) x gamma x r RESTRICTED to the measured-admissible manifold, rather than a
    product of independent ranges. ne_screen.csv is the measurement; see ne_screen.py.
    """
    fn = os.path.join(HERE, "ne_screen.csv")
    if not os.path.exists(fn):
        raise SystemExit("ne_screen.csv missing -- run ne_screen.py first; the deme grid is "
                         "defined by which geometries actually deliver Ne ~ 1447")
    out = []
    with open(fn) as f:
        for row in csv.DictReader(f):
            if int(row["keep"]):
                out.append((int(row["grid"]), int(row["K"]), float(row["m"])))
    return sorted(set(out))


# DR, 10 Sep: "I want the full cartesian product of independent lists." So the five demographic
# parameters are five independent ranges, and the Ne constraint is applied afterwards as a
# data-matching filter rather than being baked into the grid. That makes the reported
# "combinations searched" the honest full product.
#
# Fixed by DR's decision, not swept: female_bias = 1.0 (full matrilocality), s_mt = 0,
# K_early off, rho / inbreeding ban / cyclical K out of scope.
# IMPORTED, not restated. These three lists define the product that ne_screen.py measures Ne
# over, and the product this module then filters. Writing them out twice let them drift: the
# K ladder was refined in ne_screen.py to 16 values and this module kept the old 7, so the
# product silently shrank to 196 combinations and only 18 survived the Ne filter, when the
# real product is 448. One definition, imported.
from ne_screen import GRIDS as DEME_GRIDS, KS as DEME_KS, MS as DEME_MS  # noqa: E402
DEME_GAMMA = [0.08, 0.12, 0.20, 0.30, 0.42, 0.55, 0.70, 1.00]
DEME_R = [1.5, 2.0, 3.0]


# Sets that additionally sweep the residence rule and mtDNA selection. Kept as an opt-in list
# rather than switched on everywhere, because turning the axis on for a set whose cells are
# already on disk would not re-run them -- they are skipped by filename -- but would silently
# renumber the "combinations searched" total that the note quotes.
RESIDENCE_SETS = {"demog"}


def demographies(band):
    """The full product. Use ne_ok() to test whether a combination is worth spending on."""
    res = ((BAND_RESIDENCE if band else DEME_RESIDENCE) if _set in RESIDENCE_SETS
           else [{}])
    if band:
        keys = list(BAND_GRID)
        for combo in itertools.product(*(BAND_GRID[k] for k in keys)):
            for extra in res:
                yield dict(zip(keys, combo), **extra)
    else:
        for g, k, m, ga, r in itertools.product(DEME_GRIDS, DEME_KS, DEME_MS,
                                                DEME_GAMMA, DEME_R):
            for extra in res:
                yield dict(grid=g, K=k, m=m, gamma=ga, r=r, **extra)


_NE_CACHE = {}


def ne_ok(kw):
    """Did this (grid, K, m) deliver a measured autosomal Ne within 10% of 1447?

    Ne does not depend on gamma or r -- both act only while the two lineages coexist, and the
    measurement is made after burn-in when the population is of one lineage -- so one lookup
    per (grid, K, m) covers the whole gamma x r plane above it. Returns None if that triple has
    not been measured yet, which the submit script reports rather than silently skipping.
    """
    if not _NE_CACHE:
        fn = os.path.join(HERE, "ne_screen.csv")
        if not os.path.exists(fn):
            raise SystemExit("ne_screen.csv missing -- run ne_screen.py first")
        with open(fn) as f:
            for row in csv.DictReader(f):
                _NE_CACHE[(int(row["grid"]), int(row["K"]), float(row["m"]))] = int(row["keep"])
    return _NE_CACHE.get((kw["grid"], kw["K"], kw["m"]))


def cells(band):
    for mech, (_, _, vals) in ROUTES.items():
        if band and mech not in BAND_SUPPORTS:
            continue
        for v in vals:
            for kw in demographies(band):
                yield mech, v, kw


def tag(fw, mech, v, kw):
    s = f"{fw}_{mech}_{v:g}_" + "_".join(f"{k}{kw[k]:g}" for k in sorted(kw))
    return s.replace(".", "p").replace("-", "m")


def work(job):
    cmd, out = job
    if os.path.exists(out):
        return 0
    r = subprocess.run(cmd, capture_output=True, text=True, cwd=HERE)
    if r.returncode != 0:
        sys.stderr.write(f"  FAILED {os.path.basename(out)}: {r.stderr.strip()[:200]}\n")
        return -1
    return 1


def stage1_pass(d):
    return (AUT_LO <= d["aut"] <= AUT_HI and GEN_LO <= d["end_gen"] <= GEN_HI
            and d["n_est"] / max(d["n"], 1) > EST_MIN)


def build(stage, band, reps, budget):
    fw = "band" if band else "deme"
    outdir = os.path.join(HERE, f"out_{PREFIX}{stage}_{fw}")
    os.makedirs(outdir, exist_ok=True)

    jobs = []
    for mech, v, kw in cells(band):
        if stage == 2:
            s1 = os.path.join(HERE, f"out_{PREFIX}1_{fw}", tag(fw, mech, v, kw) + ".json")
            if not os.path.exists(s1):
                continue
            with open(s1) as f:
                if not stage1_pass(json.load(f)["summary"]):
                    continue
        out = os.path.join(outdir, tag(fw, mech, v, kw) + ".json")
        if os.path.exists(out):
            continue
        fl = []
        for k in sorted(kw):
            fl += [FLAG[k], f"{kw[k]:g}"]
        jobs.append(([sys.executable, "robust_run.py", fw, mech, f"{v:g}", str(stage),
                      str(reps), "0", out] + fl, out))

    total = len(list(cells(band)))
    done = len(glob.glob(os.path.join(outdir, "*.json")))
    if not jobs:
        print(f"  stage {stage} {fw}: nothing left ({done} cells on disk)")
        return 0
    print(f"  stage {stage} {fw}: {len(jobs)} cells to run, {done} already done "
          f"(of {total} in the grid)")
    t0 = time.time()
    n = 0
    with ProcessPoolExecutor(4) as ex:
        for r in ex.map(work, jobs):
            n += 1
            if time.time() - t0 > budget:
                print(f"  budget reached after {n} cells")
                break
    print(f"  ran {n} cells in {time.time()-t0:.0f} s; "
          f"{len(glob.glob(os.path.join(outdir,'*.json')))} on disk")
    return len(jobs) - n


def collect():
    rows = []
    for fw in ("deme", "band"):
        s1dir = os.path.join(HERE, f"out_{PREFIX}1_{fw}")
        s2dir = os.path.join(HERE, f"out_{PREFIX}2_{fw}")
        for p in sorted(glob.glob(os.path.join(s2dir, "*.json"))):
            with open(p) as f:
                d = json.load(f)["summary"]
            s1p = os.path.join(s1dir, os.path.basename(p))
            aut1 = None
            if os.path.exists(s1p):
                with open(s1p) as f:
                    aut1 = json.load(f)["summary"]["aut"]
            route, label, _ = ROUTES[d["mech"]]
            rows.append(dict(route=route, mech=d["mech"], label=label, val=d["val"],
                             framework=fw, gamma=d.get("gamma"), m=d.get("m"),
                             e=d.get("e"), K=d.get("K"), N=d.get("N"),
                             aut_screen=aut1, aut=d["aut"], end_gen=d["end_gen"],
                             n=d["n"], estab=d["n_est"] / max(d["n"], 1),
                             p_joint=d.get("joint")))
    if not rows:
        raise SystemExit("no stage-2 output yet")
    rows.sort(key=lambda r: (r["route"], r["val"], r["framework"]))
    fn = os.path.join(HERE, f"{PREFIX}124.csv")
    cols = ["route", "mech", "label", "val", "framework", "gamma", "m", "e", "K", "N",
            "aut_screen", "aut", "end_gen", "n", "estab", "p_joint"]
    with open(fn, "w", newline="") as f:
        w = csv.DictWriter(f, fieldnames=cols, extrasaction="ignore")
        w.writeheader(); w.writerows(rows)
    print(f"  wrote {PREFIX}124.csv: {len(rows)} scored parameter combinations")
    for route in sorted({r["route"] for r in rows}):
        sub = [r for r in rows if r["route"] == route]
        for fw in ("deme", "band"):
            s = [r for r in sub if r["framework"] == fw]
            if s:
                pj = [r["p_joint"] for r in s if r["p_joint"] is not None]
                print(f"    {route:8s} {fw:5s}: {len(s):3d} combinations, "
                      f"p_joint {min(pj):.3f}-{max(pj):.3f}" if pj else
                      f"    {route:8s} {fw:5s}: {len(s):3d} combinations")
        if not any(r["framework"] == "band" for r in sub):
            print(f"    {route:8s} band : none -- mechanism not implemented in bandsim")


def main():
    if sys.argv[1] == "collect":
        return collect()
    stage = int(sys.argv[1])
    band = "--band" in sys.argv
    reps = int(sys.argv[sys.argv.index("--reps") + 1]) if "--reps" in sys.argv else (
        10 if stage == 1 else 40)
    budget = int(sys.argv[sys.argv.index("--budget") + 1]) if "--budget" in sys.argv else 95
    left = build(stage, band, reps, budget)
    if left:
        print(f"  {left} cells still to do -- run again")


if __name__ == "__main__":
    main()
```
