## Supplementary Information section 3 code base for "Hypothesis: A modern human range expansion ∼300,000 years ago had a critical role in Neandertal origins": routes_submit.py

```
#!/usr/bin/env python3
"""
routes_submit.py -- SLURM arrays for the Routes 1/2/4 figure. (DR, 10 Sep 2026)

Emits a params file (one cell per line) and an array script that runs a contiguous RANGE of
lines per task. One cell per array task would be absurd here -- SLURM's per-task overhead is
~20 s and a deme cell is under a minute -- so cells are packed to roughly five minutes each.

    python routes_submit.py <stage> [--band] [--reps N] [--chunk N] [--cap N] [--part short]
    sbatch routes_<stage><fw>_array.sh

Stage 2 only emits cells that PASSED the stage-1 screen, so stage 1 for that framework must be
collected first. The script refuses rather than silently emitting an empty array.
"""
import glob
import json
import os
import subprocess
import sys
import time

HERE = os.path.dirname(os.path.abspath(__file__))
sys.path.insert(0, HERE)
from routes_build import cells, tag, stage1_pass, FLAG, PREFIX, ne_ok  # noqa: E402

# measured on this machine, per replicate: deme screen ~0.10 s, deme score ~0.55 s,
# band screen ~65 s (the band engine is far heavier per replicate), band score ~70 s.
# RECALIBRATED 11 Sep from sacct on the demog run, and the old deme numbers were an order of
# magnitude too low. They were measured on the matrilocal null; the demog set adds twowave and
# the unbiased s_mt arm, which run for thousands more generations because nothing goes extinct
# early. The consequence was real: stage 1 was chunked at 300 cells on a 5-minute estimate,
# tasks took 52 minutes on average and the slow tail ran past two hours against a 2:59 limit.
# 1.0 s/replicate is the measured mean (52 min / 300 cells / 10 reps).
# RECALIBRATED AGAIN, 11 Sep, from sacct on the 36-locus rerun. The deme stage-1 figure has
# now been wrong twice: 0.10 s/replicate (measured on the matrilocal null, which goes extinct
# early), then 1.0 s after the demog set added twowave and the unbiased s_mt arm. The true
# cost at 36 loci is 12-15 s -- de1d ran 58 min per 30-cell task and dm1d 76 min. Using 15.
# RECALIBRATED AGAIN, 12 Sep, from sacct on the stage-2 round. Stage 2 runs to marker
# absorption and most cells absorb early, so it is far cheaper per replicate than stage 1's
# fixed-length screen: measured 5.3 min/cell for ro2d and 9.7 for dm2d at 200 replicates,
# i.e. ~2 s/replicate, against the 20 s I had assumed. The band side is the opposite way --
# 109-131 min/cell, ~34 s/replicate.
PER_REP = {(1, "deme"): 15.0, (2, "deme"): 2.0, (1, "band"): 11.0, (2, "band"): 34.0}

# HOW LONG ONE ARRAY TASK SHOULD TAKE. Five minutes is right for stage 1, where a cell is
# under a minute and SLURM's ~20 s per-task overhead would otherwise dominate. It is wrong for
# stage 2, where a single cell already exceeds it: chunk then collapses to 1 cell per task and
# a set like demog, with ~10,000 cells passing the screen, becomes 10,000 queued jobs. That is
# what produced AssocMaxSubmitJobLimit -- the cap is on jobs QUEUED ACROSS ALL ARRAYS, not on
# array size, so MAX_TASKS below does not protect against it. An hour per task keeps the job
# count in the hundreds.
# STAGE 1 RAISED FROM 300 s TO 1800 s, 12 Sep. A deme stage-1 cell is ~22 s at 36 loci, so a
# five-minute target packs only 2 cells per task and dmi2 alone became 4,836 queued jobs --
# which then starved xdc and demog of submit slots entirely. The cap is on jobs across all
# arrays, so array size is the thing to control. Half an hour per task takes demog deme from
# 9,272 tasks to about 1,600 and leaves room for everything else.
PACK_SECONDS = {1: 1800.0, 2: 3600.0, 3: 3600.0}

# O2 runs MaxArraySize = 10001, so the largest legal array index is 10000. Leave a little
# headroom rather than sitting exactly on the boundary.
MAX_TASKS = 9500


def flag(name, cast, default):
    return cast(sys.argv[sys.argv.index(name) + 1]) if name in sys.argv else default


def active_submission(prefix, stage, fw):
    """The job id of a live array already covering this (set, stage, framework), or None.

    WHY THIS EXISTS. routes_submit emits every cell whose output json is absent -- which is
    correct for a cold start and WRONG for a retry, because a cell being computed right now by
    a running task also has no json yet. Re-running the submitter while an array is in flight
    therefore re-emits the entire in-flight set. On 12 Sep a fifteen-minute retry loop produced
    four concurrent tw2b arrays and three xd2b arrays, all racing to write the same files.
    Wasted cores are the mild consequence; two tasks writing one json is the real one.

    A stamp file per (set, stage, framework) records the job id at submission. If that job is
    still known to SLURM, this submission is redundant and is refused.
    """
    stamp = os.path.join(HERE, f".submitted_{prefix}{stage}{fw}")
    if not os.path.exists(stamp):
        return None
    jid = open(stamp).read().strip()
    if not jid:
        return None
    r = subprocess.run(["squeue", "-h", "-j", jid, "-o", "%A"],
                       capture_output=True, text=True)
    return jid if r.returncode == 0 and r.stdout.strip() else None


def main():
    stage = int(sys.argv[1])
    band = "--band" in sys.argv
    fw = "band" if band else "deme"
    reps = flag("--reps", int, 10 if stage == 1 else 100)
    cap = flag("--cap", int, 0)
    part = flag("--part", str, "short")

    live = active_submission(PREFIX, stage, fw)
    if live and "--force" not in sys.argv:
        raise SystemExit(f"  already submitted as job {live}, still in the queue -- nothing to "
                         f"do. Cells in flight have no json yet, so resubmitting now would "
                         f"duplicate them. Use --force only if you know that job is dead.")

    outdir = f"out_{PREFIX}{stage}_{fw}"
    os.makedirs(os.path.join(HERE, outdir), exist_ok=True)

    tasks = []
    skipped_missing = 0
    searched = ne_fail = ne_unmeasured = 0
    for mech, v, kw in cells(band):
        searched += 1
        # The Ne filter. Applied here rather than in the grid definition so that `searched` is
        # the honest size of the full Cartesian product -- the number DR wants to quote as
        # "parameter combinations" -- while replicates are only spent where the configuration
        # could match the anchor at all.
        if not band:
            ok = ne_ok(kw)
            if ok is None:
                ne_unmeasured += 1
                continue
            if not ok:
                ne_fail += 1
                continue
        t = tag(fw, mech, v, kw)
        if stage == 2:
            s1 = os.path.join(HERE, f"out_{PREFIX}1_{fw}", t + ".json")
            if not os.path.exists(s1):
                skipped_missing += 1
                continue
            with open(s1) as f:
                if not stage1_pass(json.load(f)["summary"]):
                    continue
        out = f"{outdir}/{t}.json"
        if os.path.exists(os.path.join(HERE, out)):
            continue
        fl = " ".join(f"{FLAG[k]} {kw[k]:g}" for k in sorted(kw))
        tasks.append(f"{fw} {mech} {v:g} {stage} {reps} 0 {out} {fl}")

    if stage == 2 and skipped_missing:
        print(f"  !! {skipped_missing} cells have no stage-1 result for {fw}. Run stage 1 "
              f"first, or those combinations will be missing from the figure.")
    if not tasks:
        raise SystemExit("  nothing to submit (all cells already on disk, or none passed)")

    per_cell = PER_REP[(stage, fw)] * reps
    # --pack SECONDS overrides the packing target. Added 12 Sep for the fairshare hole: with
    # FairShare floored at 0.000000 the main scheduler never selects these jobs, so they run
    # only by backfill -- and backfill looks at a bounded number of jobs PER USER per cycle.
    # 6,222 pending tasks from one user therefore means most are never evaluated at all, while
    # 68 nodes sit idle. Far fewer, far longer tasks get proportionally more work out of each
    # backfill selection. Pair with --part medium, whose 5-day limit allows it.
    target = flag("--pack", float, PACK_SECONDS.get(stage, 300.0))
    chunk = flag("--chunk", int, max(1, min(len(tasks), int(target / max(per_cell, 0.01)))))

    # SLURM'S MaxArraySize IS A HARD WALL, and hitting it is not a graceful failure: sbatch
    # rejects the whole script with "Invalid job array specification" and nothing runs. O2 is
    # configured at MaxArraySize = 10001, so the highest legal index is 10000.
    #
    # The five-minute packing target above sets the chunk from the per-cell cost alone, which
    # is right when the cell count is moderate and wrong when it is not. The band demog screen
    # is 91,854 cells at ~65 s each, giving chunk = 4 and 22,964 array tasks -- more than twice
    # the wall. Chunk size is the free variable, so raise it until the task count fits and let
    # the tasks run longer, rather than letting sbatch refuse the job.
    want = -(-len(tasks) // chunk)
    if want > MAX_TASKS:
        chunk = -(-len(tasks) // MAX_TASKS)
        print(f"  chunk raised to {chunk} so the array fits under MaxArraySize "
              f"({want} tasks would have been rejected outright)")

    groups = [tasks[i:i + chunk] for i in range(0, len(tasks), chunk)]
    n = len(groups)
    capn = min(n, cap) if cap else 0

    # UNIQUE PARAMS FILE PER SUBMISSION. The array reads its slice of the params file lazily,
    # with `sed`, at the moment each task STARTS. So regenerating a params file of the same name
    # while an earlier array is still running silently renumbers the lines under it: some cells
    # then run twice and others are never run at all, and two tasks can write the same output
    # file at once. That happened on 10 Sep when ro1b/ro1d were each submitted twice.
    # A timestamp in the name makes collision impossible; --suffix still works for hand-naming.
    sfx = flag("--suffix", str, "") or time.strftime("_%H%M%S")
    pf = f"{PREFIX}{stage}{fw[0]}{sfx}_params.txt"
    sf = f"{PREFIX}{stage}{fw[0]}{sfx}_array.sh"
    with open(os.path.join(HERE, pf), "w") as f:
        f.write("\n".join(tasks) + "\n")

    # ASK FOR THE LONG WINDOW MUCH EARLIER. The old threshold gave a 2:59 limit to any task
    # estimated under 100 minutes, which sounds safe until the distribution has a tail: tasks
    # averaging 58 and 76 minutes produced 12 timeouts, and a timeout silently abandons the
    # cells after the kill point. `short` allows 12 hours at no scheduling cost worth caring
    # about, so anything over half an hour now takes the long window.
    hours = 2 if per_cell * chunk < 1800 else 11
    if per_cell * chunk > 0.9 * hours * 3600:
        print(f"  !! each task is estimated at {per_cell*chunk/3600:.1f} h against a {hours} h "
              f"limit. Raise --cap and submit in waves, or pass a smaller --chunk, or these "
              f"tasks will be killed mid-chunk and the cells in them silently left undone.")
    hdr = f"""#!/bin/bash
#SBATCH -J {PREFIX[:2]}{stage}{fw[0]}
#SBATCH -p {part}
#SBATCH -t 0-{hours:02d}:59
#SBATCH -c 1
#SBATCH --mem=3G
#SBATCH -a 0-{n-1}{('%' + str(capn)) if capn else ''}
#SBATCH -o logs/{PREFIX[:2]}{stage}{fw[0]}{sfx}_%A_%a.log
"""
    body = (f"CHUNK={chunk}\n"
            f"START=$((SLURM_ARRAY_TASK_ID * CHUNK + 1))\n"
            f"END=$((START + CHUNK - 1))\n"
            f'sed -n "${{START}},${{END}}p" {pf} | while read -r LINE; do\n'
            f'  [ -z "$LINE" ] && continue\n'
            f"  python robust_run.py $LINE\n"
            f"done\n")
    with open(os.path.join(HERE, sf), "w") as f:
        f.write(hdr + body)

    cpu = len(tasks) * per_cell / 3600.0
    print(f"  stage {stage}, {fw}, {reps} replicates per cell")
    if not band:
        print(f"  full Cartesian product searched : {searched:6d} parameter combinations")
        print(f"    excluded, Ne outside 10% of 1447: {ne_fail:6d}")
        if ne_unmeasured:
            print(f"    ** not yet measured by ne_screen : {ne_unmeasured:6d} "
                  f"-- rerun ne_screen.py or these are silently missing **")
        print(f"    remaining after the Ne filter   : {searched-ne_fail-ne_unmeasured:6d}")
    print(f"  {len(tasks)} cells -> {n} array task(s) of {chunk} cells "
          f"(~{per_cell*chunk/60:.0f} min each)")
    print(f"  ~{cpu:.1f} CPU-hours total")
    print(f"  wrote {pf} and {sf}")

    # SUBMIT IT. DR: "The system of writing a time stamped sbatch is a little hard for me,
    # since I have to wait for a run to go and then meticulously copy and paste the sbatch
    # command. Is it okay for me going forward to just type a single thing?" Yes -- the
    # timestamp exists to stop a DIFFERENT bug (an array reads its slice of the params file
    # lazily with sed at task start, so regenerating a file of the same name under a running
    # job renumbers its work), and a unique name is all that needs. Nothing about it required
    # a human in the loop, and putting one there is how dmi2d got written and never run.
    #
    # --no-go writes the script without submitting, for when you want to look first.
    if "--no-go" in sys.argv:
        print(f"  NOT submitted (--no-go).  sbatch {sf}")
        return
    r = subprocess.run(["sbatch", sf], cwd=HERE, capture_output=True, text=True)
    if r.returncode == 0:
        print(f"  SUBMITTED: {r.stdout.strip()}")
        jid = r.stdout.strip().split()[-1]
        with open(os.path.join(HERE, f".submitted_{PREFIX}{stage}{fw}"), "w") as f:
            f.write(jid)
    else:
        err = (r.stderr or "").strip()
        print(f"  !! sbatch FAILED: {err}")
        if "AssocMaxSubmitJobLimit" in err:
            print("     You are at the 10,000-job cap across all arrays. Let the queue drain "
                  "and rerun this same command; already-finished cells are skipped, so "
                  "nothing is lost or repeated.")
        print(f"     retry by hand with:  sbatch {sf}")


if __name__ == "__main__":
    main()
```
