## Supplementary Information section 3 code base for "Hypothesis: A modern human range expansion ∼300,000 years ago had a critical role in Neandertal origins": scan4_collect.py

```
#!/usr/bin/env python3
"""
scan4_collect.py -- build Figure 3 from the FULL-LENGTH runs in out4/. Six panels (m, gamma, r, K,
grid, sY), each showing mean % modern-lineage ancestry at Autosomes / Y / mtDNA / X plus mean
generations to archaic extinction (right axis), sampled at the sampling generation. Legend on the right; no
title; no per-panel n annotation (matching the finalized Figure-3 style).

Identical in style to collect_scan3.py, with two additions: the runs behind it are full
length (see scan4_run.py for why that matters in the sY panel), and a dashed grey line gives
the FIXATION probability of the modern Y -- the quantity the note actually argues about, and
one that early stopping made impossible to estimate.

  python scan4_collect.py     # -> figure3_full.png, figure3_full.csv

Seventh panel: theta (the fertility of F1 hybrid males) at the standard m = 0.10, with gamma
recalibrated at every theta to hold autosomes at 5% -- the m = 0.10 slice of the calibrated
grid3 analysis. Its data come from fig3_theta.csv (written by fig3_theta.py after the grid3
rerun is collected); if that file is absent the panel is skipped with a note, so this
collector can be run before grid3 finishes.
"""
import os
import datetime as _dt
import glob
import numpy as np
import matplotlib
matplotlib.use("Agg")
import matplotlib.pyplot as plt
from scan4_run import PANELS

ORDER = ["m", "gamma", "r", "K", "grid", "sY"]
# 4x2 layout. The bottom row holds the two analyses that act directly on the Y -- the
# calibrated-theta panel and the additive-selection panel -- and the legend takes the
# free top-right cell.
#     0 m        1 LEGEND
#     2 gamma    3 r
#     4 K        5 grid
#     6 theta    7 sY
SLOT = {"m": 0, "gamma": 2, "r": 3, "K": 4, "grid": 5, "sY": 7}
LEGEND_SLOT = 1
THETA_SLOT = 6
LABELS = {"m": "m (migration rate)", "gamma": "γ (interbreeding rate)",
          "r": "r (modern:archaic fitness)", "K": "K (carrying capacity)",
          "grid": "grid dimension", "sY": "s (additive selection against archaic Y)"}
LOGX = {"gamma", "sY"}
# columns in out3: seed, mt, Y, auto, X, extgen
NSAMPLE = int(os.environ.get('NSAMPLE', '10'))
COL = {"mt": 1, "Y": 2, "auto": 3, "X": 4, "ext": 5}
SER = [("auto", "Autosomes", "#1f77b4"), ("Y", "Y", "#ff7f0e"),
       ("mt", "mtDNA", "#2ca02c"), ("X", "X chromosome", "#9467bd")]


# THE SUPERSEDED GEOMETRY IS STILL SITTING IN out4 -- FOUND 14 Sep, and it is the scan4
# geometry incident again. The 13 Sep rerun at the anchored grid 12 / K 11 wrote seed blocks
# at stride 100 (s0, s100, ... s1900, 100 replicates each); the original August run at grid
# 20 / K 20 wrote blocks at the odd fifties (s50, s150, ...) and was never moved aside on
# this machine. Both match out_{panel}_{v}_s*.txt, so a plain glob pools 2,000 anchored
# replicates with 500 superseded ones and every number moves: K = 8 autosomal ancestry reads
# 11.7% pooled against 5.9% anchored, because the 20x20 lattice has five times the census.
# Files written before the cutoff are skipped, and the count of what was skipped is printed
# so this cannot pass unnoticed again.
SCAN4_FIX = _dt.datetime(2026, 9, 1, 0, 0, 0).timestamp()


def pooled(panel, v):
    rows, skipped = [], 0
    for f in glob.glob(f"out4/out_{panel}_{v}_s*.txt"):
        if os.path.getmtime(f) < SCAN4_FIX:
            skipped += 1
            continue
        for line in open(f).read().strip().split("\n")[1:]:
            p = line.split("\t")
            if len(p) >= 6:
                rows.append([float(x) for x in p])
    if skipped:
        print(f"    {panel} = {v}: skipped {skipped} pre-September file(s) at the superseded "
              f"grid 20 / K 20 geometry")
    if not rows:
        return None
    a = np.array(rows)
    ext = a[:, COL["ext"]]; ext = ext[ext >= 0]
    # nanmean, NOT mean -- FIXED 14 Sep. DR: "why are there missing points for Y % modern (K)
    # in demes?" Because a replicate in which no males are alive at the sampling generation
    # records the Y frequency among males as NaN, which is the honest value for that
    # replicate, and a plain .mean() then propagates it to the whole cell. One replicate in
    # 2,500 did that at K = 20 and at K = 40, and three at K = 24, and those three points
    # vanished from the published figure while their neighbours, drawn from the same 2,500,
    # stayed. The fraction dropped is recorded so the caption can state it rather than the
    # figure hiding it.
    Yc, mtc = a[:, COL["Y"]], a[:, COL["mt"]]
    n_bad = int(np.isnan(Yc).sum() + np.isnan(mtc).sum())
    if n_bad:
        print(f"    {panel} = {v}: {n_bad} of {len(a)} replicate(s) with an undefined "
              f"uniparental frequency (no males alive at sampling); averaged over the rest")
    return dict(n=len(a), n_undefined=n_bad,
                mt=100 * np.nanmean(mtc), Y=100 * np.nanmean(Yc),
                auto=100 * np.nanmean(a[:, COL["auto"]]),
                X=100 * np.nanmean(a[:, COL["X"]]),
                ext=(ext.mean() if ext.size else np.nan),
                Yfix=100 * np.nanmean(Yc >= 0.999),
                mtfix=100 * np.nanmean(mtc >= 0.999),
                Ysamp=100 * np.nanmean(Yc ** NSAMPLE),
                mtsamp=100 * np.nanmean(mtc ** NSAMPLE))


def read_theta_panel():
    """fig3_theta.csv rows -> the same dict shape pooled() returns, plus gamma_star."""
    if not os.path.exists("fig3_theta.csv"):
        return None
    import csv
    out = []
    for r in csv.DictReader(open("fig3_theta.csv")):
        if not r.get("Y"):
            continue
        out.append((float(r["theta"]),
                    dict(auto=float(r["auto"]), Y=float(r["Y"]), mt=float(r["mt"]),
                         X=float(r["X"]), ext=float(r["ext"]) if r["ext"] else np.nan,
                         Yfix=float(r["Yfix"]), Ysamp=float(r["Yall"]),
                         gstar=float(r["gamma_star"]))))
    return out or None


def main():
    theta = read_theta_panel()
    fig, axes = plt.subplots(4, 2, figsize=(11, 17.3))
    flat = axes.ravel()
    if theta is None:
        print("note: fig3_theta.csv not found -- writing the six-panel figure; run "
              "fig3_theta.py after the grid3 rerun, then re-run this collector")
    handles = None
    for panel in ORDER:
        ax = flat[SLOT[panel]]
        xs = [float(v) for v in PANELS[panel]]
        data = [pooled(panel, v) for v in PANELS[panel]]
        keep = [(x, d) for x, d in zip(xs, data) if d is not None]
        ax.set_xlabel(LABELS[panel]); ax.set_ylabel("Percent modern"); ax.set_ylim(-3, 103)
        if not keep:
            ax.set_title("(no data yet)"); continue
        X = [k[0] for k in keep]; D = [k[1] for k in keep]
        lh = []
        for key, lab, col in SER:
            l, = ax.plot(X, [d[key] for d in D], "-o", ms=3, color=col, label=lab)
            lh.append(l)
        lf, = ax.plot(X, [d["Yfix"] for d in D], "--", lw=1.2, color="#ff7f0e", alpha=.75,
                      label="P(modern Y fixed)")
        ls_, = ax.plot(X, [d["Ysamp"] for d in D], ":", lw=1.4, color="#8c3b00", alpha=.85,
                       label=f"P(all {NSAMPLE} sampled Y modern)")
        lh += [lf, ls_]
        if panel in LOGX:
            ax.set_xscale("log")
        ax2 = ax.twinx()
        le, = ax2.plot(X, [d["ext"] for d in D], "-o", ms=3, color="0.6",
                       label="generations to extinction")
        ax2.set_ylabel("Generations to extinction"); ax2.set_ylim(0, 1000)
        ax.grid(alpha=0.3)
        if handles is None:
            handles = lh + [le]

    # ---- seventh panel: calibrated theta at m = 0.10 ----
    ax = flat[THETA_SLOT]
    if theta is not None:
        X = [t for t, _ in theta]; D = [d for _, d in theta]
        xr = np.arange(len(X))            # rank spacing, as in the calibrated-theta figure
        for key, lab, col in SER:
            ax.plot(xr, [d[key] for d in D], "-o", ms=3, color=col)
        ax.plot(xr, [d["Yfix"] for d in D], "--", lw=1.2, color="#ff7f0e", alpha=.75)
        ax.plot(xr, [d["Ysamp"] for d in D], ":", lw=1.4, color="#8c3b00", alpha=.85)
        ax.set_xticks(xr); ax.set_xticklabels([f"{t:g}" for t in X], fontsize=8)
        # two lines: the single-line version overflows the axes and gets clipped
        ax.set_xlabel("θ (fertility of F1 hybrid males)\n"
                      "γ recalibrated to 5% autosomes, m = 0.10")
        ax.set_ylabel("Percent modern"); ax.set_ylim(-3, 103); ax.grid(alpha=0.3)
        ax2 = ax.twinx()
        ax2.plot(xr, [d["ext"] for d in D], "-o", ms=3, color="0.6")
        ax2.set_ylabel("Generations to extinction"); ax2.set_ylim(0, 1000)
    else:
        ax.axis("off")

    # The legend goes in the free top-right cell rather than in a reserved right-hand strip,
    # so the seven panels get the full width of the page.
    legend_ax = flat[LEGEND_SLOT]
    legend_ax.axis("off")
    if handles:
        legend_ax.legend(handles=handles, labels=[h.get_label() for h in handles],
                         loc="center", ncol=1, fontsize=11, frameon=False)
    fig.tight_layout()
    fig.savefig("figure3_full.png", dpi=140, bbox_inches="tight")
    print("saved figure3_full.png")
    with open("figure3_full.csv", "w") as fh:
        fh.write(f"panel,value,n,auto,Y,mtDNA,X,P_Y_fixed,P_mt_fixed,P_Y_sample{NSAMPLE},P_mt_sample{NSAMPLE},ext_gen\n")
        for panel in ORDER:
            for v in PANELS[panel]:
                d = pooled(panel, v)
                if d is None:
                    continue
                fh.write(f"{panel},{v},{d['n']},{d['auto']:.2f},{d['Y']:.2f},{d['mt']:.2f},"
                         f"{d['X']:.2f},{d['Yfix']:.1f},{d['mtfix']:.1f},"
                         f"{d['Ysamp']:.1f},{d['mtsamp']:.1f},{d['ext']:.0f}\n")
    if theta is not None:
        with open("figure3_full.csv", "a") as fh:
            for t, d in theta:
                fh.write(f"theta_cal,{t:g},,{d['auto']:.2f},{d['Y']:.2f},{d['mt']:.2f},"
                         f"{d['X']:.2f},{d['Yfix']:.1f},,{d['Ysamp']:.1f},,{d['ext']:.0f}\n")
    print("saved figure3_full.csv")


if __name__ == "__main__":
    main()
```
