## Supplementary Information section 3 code base for "Hypothesis: A modern human range expansion ∼300,000 years ago had a critical role in Neandertal origins": scan4_run.py

```
#!/usr/bin/env python3
"""
scan4_run.py -- Figure 3 parameter scan, FULL LENGTH (no early stopping).
ONE cluster task = one (panel, value) x `reps` replicates, sampled at the sampling generation.

Vanilla baseline: K=20, r=2, m=0.10, gamma=0.12, grid=20, s=0, matrilineal, NO DM
(the epistatic/DM machinery is not implemented, so Figure 3 has no DM panel).

Panels scanned (one parameter varied from baseline at a time):
    m, gamma, r, K, grid                     (demographic; s=0)
    sY  -> ADDITIVE selection against the archaic Y (sel_locus="Y", s=value)

WHY THIS REPLACES scan_run3.py
  scan_run3 stopped each replicate 50 generations after archaic extinction, on the argument
  that post-extinction ancestry is neutral and therefore stationary. That argument holds only
  when s = 0. In the sY panel selection on the Y keeps acting for the whole remaining run, and
  since extinction typically occurs by generation ~300-700, early stopping discarded roughly
  8,400 generations of selection. Checked directly at s = 0.005, gamma = 0.12, m = 0.10:
  the modern Y averages 3.1% at extinction+50 but 33.1% at the sampling generation, and individual
  replicates go from ~0% to 98.7%. The demographic panels are unaffected (their loci really
  are neutral post-extinction) but are rerun here too so that every panel of the figure is
  produced the same way.

  Fixation probabilities are also reported now, which early stopping made meaningless:
  drift continues for thousands of generations after the archaics are gone.

Writes one row per replicate:  seed  mt  Y  auto  X  extgen

Usage:  python scan4_run.py <panel> <value> <reps> <seed0> <outfile>
"""
import os
import sys
import numpy as np
from dataclasses import replace
from spatial_sim2 import Config, run

SAMPLE_GEN = 6993
BASE = dict(grid=20, K=20, r=2.0, m=0.10, gamma=0.12, s=0.0, sel_locus="none",
            gens=SAMPLE_GEN + 10, sample_gen=SAMPLE_GEN, stop_after_ext=0, record_every=1000)

PANELS = {
    "m":     [0.02, 0.03, 0.04, 0.05, 0.06, 0.08, 0.10, 0.12, 0.15],
    # EXTENDED TO 1.00 ON 13 Sep. Appendix A's gamma ladder is "8 values from 0.08 to 1.00",
    # and the Cartesian search does run all eight -- but this sensitivity sweep stopped at
    # 0.30, so Figure S3.5 drew open grey circles at 0.42, 0.55, 0.70 and 1.00 meaning "on the
    # ladder, never run". 0.42 is the value the two-wave result in Route 4 is quoted at, which
    # is the one place where having no sensitivity point is actually awkward.
    "gamma": [0.02, 0.04, 0.06, 0.08, 0.10, 0.12, 0.15, 0.20, 0.30, 0.42, 0.55, 0.70, 1.00],
    "r":     [1.1, 1.25, 1.5, 2.0, 2.5, 3.0, 4.0],
    "K":     [4, 6, 8, 10, 12, 16, 20, 24, 32, 40],
    # 14 ADDED: it is on the note's geometry ladder and is the grid kearly's cells use, so
    # leaving it out of the sensitivity sweep meant the one geometry with new results had no
    # baseline point to be read against.
    "grid":  [4, 6, 8, 10, 12, 14, 16, 20, 24, 28],
    # finer at the low end, where the old panel was most badly wrong
    "sY":    [0.0, 0.001, 0.002, 0.005, 0.01, 0.02, 0.04, 0.08, 0.16, 0.32, 0.64],
}


# Expansion geometry overrides. The published baseline (grid 20, K 20) has a post-wave census
# of ~8000, five times the post-mixture size the rest of the analysis assumes (Muller's Ratchet
# v32: NN-uniparental = 725, diploid 1500). Since the modern mtDNA's advantage is a ~0.3%
# selection coefficient whose potency scales with 2*Ne*s, that size mismatch inflated every
# spatial result. These flags shift the BASELINE; where a panel sweeps the same parameter, the
# swept value still wins, so the panel shows sensitivity AROUND the new baseline.
def _geom(kw):
    import sys as _s
    for flag, key, cast in (("--grid", "grid", int), ("--K", "K", int),
                            ("--m", "m", float), ("--r", "r", float)):
        if flag in _s.argv:
            if key not in kw:
                continue          # this script sweeps that parameter; do not pin it
            kw[key] = cast(_s.argv[_s.argv.index(flag) + 1])
    for flag, key in (("--K-early", "K_early"), ("--K-early-gens", "K_early_gens")):
        if flag in _s.argv:
            kw[key] = int(_s.argv[_s.argv.index(flag) + 1])
    if kw.get("K", 20) < 7:
        raise SystemExit("K < 7: deme reproductive failure collapses the archaic population")
    return kw


def make_cfg(panel, value):
    kw = _geom(dict(BASE))
    if panel == "m":       kw["m"] = float(value)
    elif panel == "gamma": kw["gamma"] = float(value)
    elif panel == "r":     kw["r"] = float(value)
    elif panel == "K":     kw["K"] = int(value)
    elif panel == "grid":  kw["grid"] = int(value)
    elif panel == "sY":    kw["s"] = float(value); kw["sel_locus"] = "Y"
    else: raise ValueError(f"unknown panel {panel}")
    return Config(**kw)


def main():
    panel, value = sys.argv[1], sys.argv[2]
    reps, seed0, outfile = int(sys.argv[3]), int(sys.argv[4]), sys.argv[5]
    cfg0 = make_cfg(panel, value)
    os.makedirs(os.path.dirname(outfile) or ".", exist_ok=True)  # so a moved/renamed
    # output directory cannot silently fail every task in the array
    with open(outfile, "w") as f:
        f.write("seed\tmt\tY\tauto\tX\textgen\n")
        for s in range(seed0, seed0 + reps):
            out, ext = run(replace(cfg0, seed=s))
            i = int(np.argmin(np.abs(out["gen"] - SAMPLE_GEN)))
            eg = int(ext) if ext is not None else -1
            f.write(f"{s}\t{out['mt_mod'][i]:.6g}\t{out['Y_mod'][i]:.6g}\t"
                    f"{out['auto_mod'][i]:.6g}\t{out['X_mod'][i]:.6g}\t{eg}\n")
    print("wrote", outfile)


if __name__ == "__main__":
    main()
```
