## Supplementary Information section 3 code base for "Hypothesis: A modern human range expansion ∼300,000 years ago had a critical role in Neandertal origins": scan4_submit.py

```
#!/usr/bin/env python3
"""
scan4_submit.py -- SLURM array for the full-length Figure 3 scan (scan4_run.py).

Usage:  python scan4_submit.py <reps> <batch> [concurrency] [partition] [--resume]
        python scan4_submit.py 1000 50 500 short
then:   sbatch scan4_array.sh          (output goes to out4/)
"""
import os
import sys
from scan4_run import PANELS


def _strip_flags(av):
    """Drop --flag and its VALUE, so positional parsing is not shifted by the geometry flags."""
    out, skip = [], False
    for a in av:
        if skip:
            skip = False; continue
        if a.startswith("--"):
            skip = a in GEOM_FLAGS or a == "--rna"; continue
        out.append(a)
    return out


GEOM_FLAGS = ("--grid", "--K", "--m", "--r")


def _geom_str():
    """Echo the geometry flags into every task line, so the workers cannot silently run at the
    published grid 20 / K 20 while the submit script was told otherwise."""
    import sys as _s
    return "".join(f" {f} {_s.argv[_s.argv.index(f) + 1]}" for f in GEOM_FLAGS if f in _s.argv)


def main():
    GEOM = _geom_str()
    args   = _strip_flags(sys.argv[1:])
    total  = int(args[0]) if len(args) > 0 else 1000
    batch  = int(args[1]) if len(args) > 1 else 50
    cap    = int(args[2]) if len(args) > 2 else 500
    part   = args[3] if len(args) > 3 else "short"
    resume = "--resume" in sys.argv

    os.makedirs("out4", exist_ok=True); os.makedirs("logs", exist_ok=True)
    tasks, skipped = [], 0
    for panel, values in PANELS.items():
        for v in values:
            for s0 in range(0, total, batch):
                reps = min(batch, total - s0)
                out = f"out4/out_{panel}_{v}_s{s0}.txt"
                if resume and os.path.exists(out) and sum(1 for _ in open(out)) >= reps + 1:
                    skipped += 1
                    continue
                tasks.append(f"{panel} {v} {reps} {s0} {out}{GEOM}")
    if not tasks:
        print("nothing to do -- every output file is already complete"); return

    open("scan4_params.txt", "w").write("\n".join(tasks) + "\n")
    n = len(tasks); cap = min(n, cap)
    open("scan4_array.sh", "w").write(f"""#!/bin/bash
#SBATCH -J scan4
#SBATCH -p {part}
#SBATCH -t 0-11:59
#SBATCH -c 1
#SBATCH --mem=1G
#SBATCH -o logs/scan4_%A_%a.log
#SBATCH --array=0-{n - 1}{("%%%d" % cap) if cap else ""}

LINE=$(sed -n "$((SLURM_ARRAY_TASK_ID + 1))p" scan4_params.txt)
echo "task $SLURM_ARRAY_TASK_ID : $LINE"
python scan4_run.py $LINE
""")
    ncond = sum(len(v) for v in PANELS.values())
    print(f"  geometry:{GEOM or ' default grid 20 / K 20'}")
    print(f"wrote scan4_params.txt ({n} tasks over {ncond} panel-values), {total} reps each")
    if resume:
        print(f"  RESUME: skipped {skipped} already-complete tasks")
    print(f"  ~{batch * 15.5 / 60:.1f} min per task, {ncond * total * 15.5 / 3600:,.0f} CPU-hours total")
    print("submit:  sbatch scan4_array.sh")


if __name__ == "__main__":
    main()
```
