## Supplementary Information section 3 code base for "Hypothesis: A modern human range expansion ∼300,000 years ago had a critical role in Neandertal origins": sf1_collect.py

```
#!/usr/bin/env python3
"""
sf1_collect.py -- pool out_sf1/ into Supplementary Figure 1. Patrilineal expansion with a
modern-partner-autosome x archaic-mtDNA DM plus independent additive positive selection s on
the partner locus.

Ancestry is reported CONDITIONAL ON A SUCCESSFUL EXPANSION -- i.e. averaged only over the
replicates in which the archaic population actually went extinct (extgen >= 0). This removes
the stalled replicates, in which the modern lineage never took over and its small near-origin
sample is trivially ~100% modern, and which otherwise inflate the low-s means. The fraction of
replicates that expanded successfully is drawn as a separate line so the reader can see how
often the expansion stalls.

Outputs: suppfig1.png (+ suppfig1_data.csv with both conditional means and expansion fraction).

    python sf1_collect.py
"""
import glob
import numpy as np
import matplotlib
matplotlib.use("Agg")
import matplotlib.pyplot as plt

COL = dict(seed=0, mt=1, partner=2, autobg=3, Y=4, X=5, ext=6)


def pool(tag):
    rows = []
    for f in glob.glob(f"out_sf1/out_{tag}_s*.txt"):
        for line in open(f).read().strip().split("\n")[1:]:
            p = line.split("\t")
            if len(p) >= 7:
                rows.append([float(x) for x in p])
    return np.array(rows) if rows else None


def main():
    svals = []
    for f in glob.glob("out_sf1/out_*_s0.txt"):
        tag = f.split("out_")[-1].rsplit("_s", 1)[0]
        svals.append((float(tag.replace("p", ".")), tag))
    svals = sorted(set(svals))

    xs, mt, part, bg, frac, nsucc = [], [], [], [], [], []
    with open("suppfig1_data.csv", "w") as out:
        out.write("s,mt_success,partner_success,autobg_success,expansion_fraction,n_success,n_total\n")
        for s, tag in svals:
            a = pool(tag)
            if a is None:
                continue
            succ = a[a[:, COL["ext"]] >= 0]          # archaic went extinct
            n_tot = len(a); n_s = len(succ)
            xs.append(s); frac.append(100 * n_s / n_tot); nsucc.append(n_s)
            if n_s:
                mt.append(100 * succ[:, COL["mt"]].mean())
                part.append(100 * succ[:, COL["partner"]].mean())
                bg.append(100 * succ[:, COL["autobg"]].mean())
            else:
                mt.append(np.nan); part.append(np.nan); bg.append(np.nan)
            out.write(f"{s},{mt[-1]:.2f},{part[-1]:.2f},{bg[-1]:.2f},{frac[-1]:.1f},{n_s},{n_tot}\n")

    fig, ax = plt.subplots(figsize=(8, 5.5))
    ax.plot(xs, mt, "-o", color="#d62728", label="mtDNA")
    ax.plot(xs, part, "-s", color="#1f77b4", label="partner locus")
    ax.plot(xs, bg, "-^", color="0.5", label="background autosomes")
    ax.plot(xs, frac, "--D", color="#2ca02c", ms=4, label="replicates expanding successfully")
    ax.set_xscale("symlog", linthresh=0.005)
    ax.set_xlabel("positive selection coefficient s at the partner locus")
    ax.set_ylabel("Percent modern")
    ax.set_ylim(-3, 103); ax.grid(alpha=0.3); ax.legend(frameon=False, loc="center left", fontsize=9)
    ax.set_xticks(xs); ax.set_xticklabels([("0" if v == 0 else f"{v:g}") for v in xs])
    fig.tight_layout()
    fig.savefig("suppfig1.png", dpi=160, bbox_inches="tight")
    print("saved suppfig1.png and suppfig1_data.csv")
    print("s / mt% / partner% / autobg% (successful only) / %expanded / n_success")
    for i, s in enumerate(xs):
        print(f"  {s:<6g} {mt[i]:5.1f} {part[i]:6.1f} {bg[i]:6.1f}   {frac[i]:5.1f}   n={nsucc[i]}")


if __name__ == "__main__":
    main()
```
