## Supplementary Information section 3 code base for "Hypothesis: A modern human range expansion ∼300,000 years ago had a critical role in Neandertal origins": sf1_run.py

```
#!/usr/bin/env python3
"""
sf1_run.py -- Supplementary Figure 1 worker (new engine). Patrilineal range expansion with a
male-limited dominant Dobzhansky-Muller incompatibility between a MODERN autosomal partner locus
(locus 8 = index 7) and ARCHAIC mtDNA, PLUS independent additive positive selection of
coefficient s favouring the modern allele at that same partner locus.

ONE cluster task = one s-value x `reps` replicates, sampled at the sampling generation (6,993; no early stop). Row: seed  mt  partner  autobg  Y  X  extgen
(partner = modern fraction at locus 8; autobg = loci 1-7).

Usage:  python sf1_run.py <s> <reps> <seed0> <outfile>
"""
import os
import sys
import numpy as np
from dataclasses import replace
import spatial_sim2 as S

SAMPLE_GEN = 6993


# --- expansion geometry, settable from the command line (see grid5_run.py for the rationale) ---
def _geom(kw):
    import sys as _s
    for flag, key, cast in (("--grid", "grid", int), ("--K", "K", int),
                            ("--m", "m", float), ("--r", "r", float)):
        if flag in _s.argv and key in kw:
            kw[key] = cast(_s.argv[_s.argv.index(flag) + 1])
    for flag, key in (("--K-early", "K_early"), ("--K-early-gens", "K_early_gens")):
        if flag in _s.argv:
            kw[key] = int(_s.argv[_s.argv.index(flag) + 1])
    if kw.get("K", 20) < 7:
        raise SystemExit("K < 7: deme reproductive failure collapses the archaic population")
    return kw


def cfg_for(s):
    return S.Config(**_geom(dict(grid=20, K=20, m=0.10, gamma=0.12, r=2.0, lineage="patrilineal",
                    s=float(s), sel_locus="auto", auto_locus=7,
                    dm=1.0, dm_focal="auto", dm_focal_locus=7, dm_partner="mt",
                    dm_male_limited=True, dm_dom="dominant",
                    gens=SAMPLE_GEN + 10, sample_gen=SAMPLE_GEN,
            # stop_after_ext was 50. Rerunning Table 1 at the sampling generation showed
            # that rule can catch a still-moving trajectory: modern PRDM9-A went from 28%
            # to 96% once given the full run, because a sweep at s = 0.005 needs several
            # thousand generations. Sampling every analysis at 6,993 removes that risk and
            # makes all results in the note mutually comparable.
                    stop_after_ext=0, record_every=200)))


def run_one(cfg):
    rng = np.random.default_rng(cfg.seed)
    excl = (7,)
    _cap_early = S.cfg_early(cfg)   # K_early phase, if any
    P = S._init(_cap_early, rng); ext = None; res = None
    for g in range(cfg.gens):
        _c = S.cap_at(cfg, _cap_early, g)
        S._migrate(P, _c, rng)
        nxt = S._repro(P, _c, rng)
        if nxt is None:
            break
        P = S._reseed_origin(nxt, _c)
        if ext is None and (P["lin"] == 0).sum() == 0:
            ext = g
        if g == cfg.sample_gen:
            res = _snap(P, excl); break
        if ext is not None and cfg.stop_after_ext > 0 and g >= ext + cfg.stop_after_ext:
            res = _snap(P, excl); break
    if res is None:
        res = _snap(P, excl)
    return res, ext


def _snap(P, excl):
    st = S._stats(P, excl)
    mod = P["lin"] == 1
    partner = float(P["auto"][mod][:, 7].mean() / 2.0) if mod.any() else np.nan
    return st, partner


def main():
    s = sys.argv[1]; reps = int(sys.argv[2]); seed0 = int(sys.argv[3]); outfile = sys.argv[4]
    cfg0 = cfg_for(s)
    os.makedirs(os.path.dirname(outfile) or ".", exist_ok=True)  # so a moved/renamed
    # output directory cannot silently fail every task in the array
    with open(outfile, "w") as f:
        f.write("seed\tmt\tpartner\tautobg\tY\tX\textgen\n")
        for seed in range(seed0, seed0 + reps):
            (st, pm), ext = run_one(replace(cfg0, seed=seed))
            eg = int(ext) if ext is not None else -1
            f.write(f"{seed}\t{st['mt_mod']:.6g}\t{pm:.6g}\t{st['auto_mod']:.6g}\t"
                    f"{st['Y_mod']:.6g}\t{st['X_mod']:.6g}\t{eg}\n")
    print("wrote", outfile)


if __name__ == "__main__":
    main()
```
