## Supplementary Information section 3 code base for "Hypothesis: A modern human range expansion ∼300,000 years ago had a critical role in Neandertal origins": sf1_submit.py

```
#!/usr/bin/env python3
"""
sf1_submit.py -- SLURM job array for Supplementary Figure 1 (sf1_run.py). One task per
(s-value x batch). Pooled by sf1_collect.py.

Usage:  python sf1_submit.py            # 100 reps/s, batch 25
        python sf1_submit.py 100 25
then:   sbatch sf1_array.sh
"""
import sys, os

S_VALUES = [0.0, 0.005, 0.01, 0.02, 0.04, 0.08, 0.16, 0.32]


GEOM_FLAGS = ("--grid", "--K", "--m", "--r", "--K-early", "--K-early-gens")


def _strip_flags(av):
    """Positional parsing must not be shifted by the geometry flags, so drop each flag AND its
    value before the positional args are read."""
    out, skip = [], False
    for a in av:
        if skip:
            skip = False; continue
        if a.startswith("--"):
            skip = a in GEOM_FLAGS or a == "--rna"; continue
        out.append(a)
    return out


def _geom_str():
    """Echo the geometry flags onto every task line. Without this the workers would silently run
    the grid-20 / K-20 default while the submit script had been told otherwise -- which produces
    plausible-looking numbers that are simply wrong."""
    import sys as _s
    return "".join(f" {f} {_s.argv[_s.argv.index(f) + 1]}"
                   for f in GEOM_FLAGS if f in _s.argv)


def main():
    GEOM = _geom_str()
    nums = [a for a in sys.argv[1:]]
    total = int(nums[0]) if len(nums) >= 1 else 100
    batch = int(nums[1]) if len(nums) >= 2 else 25
    os.makedirs("out_sf1", exist_ok=True); os.makedirs("logs", exist_ok=True)
    tasks = []
    for s in S_VALUES:
        for s0 in range(0, total, batch):
            reps = min(batch, total - s0)
            tag = str(s).replace(".", "p")
            tasks.append(f"{s} {reps} {s0} out_sf1/out_{tag}_s{s0}.txt{GEOM}")
    open("sf1_params.txt", "w").write("\n".join(tasks) + "\n")
    n = len(tasks); cap = 0        # no throttle
    open("sf1_array.sh", "w").write(f"""#!/bin/bash
#SBATCH -J sf1new
#SBATCH -p short
#SBATCH -t 0-11:59
#SBATCH -c 1
#SBATCH --mem=4G
#SBATCH -o logs/sf1_%A_%a.log
#SBATCH --array=0-{n - 1}{("%%%d" % cap) if cap else ""}

LINE=$(sed -n "$((SLURM_ARRAY_TASK_ID + 1))p" sf1_params.txt)
echo "task $SLURM_ARRAY_TASK_ID : $LINE"
python sf1_run.py $LINE
""")
    print(f"wrote sf1_params.txt ({n} tasks) and sf1_array.sh -- submit: sbatch sf1_array.sh")


if __name__ == "__main__":
    main()
```
