## Supplementary Information section 3 code base for "Hypothesis: A modern human range expansion ∼300,000 years ago had a critical role in Neandertal origins": spatial_sim2.py

```
"""
spatial_sim2.py -- range-expansion simulation matching the 7/19/2026 "Simulation setup"
methods (parent-pairing formulation), replacing the earlier adult-identity-switching engine.

Model (cfg.lineage selects matrilineal [default], patrilineal, or symmetric):
  * 20x20 demes, carrying capacity K (same for both groups). Origin deme (0,0) seeded modern,
    all others archaic; each deme seeded with Poisson(K/2) males and Poisson(K/2) females.
  * A child's LINEAGE is its MOTHER's lineage (matrilineal); mtDNA is transmitted from the
    mother, so modern-lineage individuals always carry modern mtDNA (mtDNA is 100% modern by
    construction).
  Each generation:
   (i) Migration: each individual moves to an adjacent deme w.p. m per cardinal direction.
   (ii) Offspring counts per deme (Extended Data Table 1, using the CORRECT Q-line formulas):
        with MM,FM,MA,FA = modern/archaic males/females in the deme, and
        Q = 2r*MM*FM + 2g*MM*FA + 2rg*MA*FM + 2*MA*FA,
        the expected number of offspring of each (father x mother) type, PER SEX, is
            Modern x Modern  -> modern child : K*r *MM*FM / Q
            Modern x Archaic -> archaic child: K*g *MM*FA / Q
            Archaic x Modern -> modern child : K*r*g*MA*FM / Q
            Archaic x Archaic-> archaic child: K   *MA*FA / Q
        (r = modern-lineage fitness; g = gamma assortative-mating discount for cross matings.)
        Zero offspring in a deme lacking either a male or a female.
   (iii) Allele transmission over 12 unlinked loci: mtDNA (from mother), Y (father->sons),
        8 autosomes (diploid; one allele sampled from each parent), 2 X loci (mother samples
        one of her two; father transmits his X to daughters only). Then ADDITIVE selection:
        an offspring is removed with probability s per archaic allele at the selected locus
        (so 2s if homozygous archaic). Selected-against offspring are DROPPED (not resampled),
        so the number of viable individuals in a deme can fall below its carrying capacity K.
        Selection can act on the Y, an autosomal locus, or an X locus (cfg.sel_locus).
   (iv) If the origin deme lacks a modern male or female, reseed one.

  Epistatic / Dobzhansky-Muller selection (methods item 2) is NOT implemented yet -- see the
  _selection() hook. Patrilineal / symmetric expansions are selected by cfg.lineage.

Sampling: statistics are read at cfg.sample_gen (default 6993 = 50 ka,
i.e. 200 kyr after mixture at 250 ka, at 28.6 years per generation).
"""

import numpy as np
from dataclasses import dataclass, replace

# GENOME LAYOUT, SETTLED 11 Sep 2026. DR: "simulate 32 neutral loci and 4 candidate selection
# loci on the autosomes (and 1 and 1 on the X)".
#
#   autosomes  36 loci:  0-31 NEUTRAL, always measured;  32-35 reserved for mechanisms
#   X           3 loci:  0 the drive locus, 1 the DMI / selection candidate, 2 neutral
#
# Reserving the mechanism loci rather than carving them out of a fixed budget is what makes the
# two requirements compatible. Ancestry is measured only on loci untouched by selection, AND it
# is measured on exactly 32 loci for every architecture, so a single-pair DMI and four
# restorers are estimated with identical precision and the restorer series can be read as a
# trend rather than partly as a trend in measurement error.
#
# WHY 32 AND NOT 8. A variance decomposition showed that essentially all replicate-to-replicate
# scatter in the ancestry estimate was locus sampling, not demography: at 8 loci the
# between-locus variance within a replicate was 19.1 and the variance of the 8-locus mean
# across replicates 2.37, against 19.1/8 = 2.39 predicted if the demographic component were
# zero. The expansion lands on nearly the same true ancestry every time and we could not see it
# through eight markers. 32 loci take the SD from 1.55 to 0.77 percentage points; the cost is
# about 1.6x runtime, against the 4x that the same halving would cost through replication.
#
# The X has three loci so that an X-linked DMI still leaves a neutral X marker to report; with
# the old XLOCI = 2 such a run had no X loci left and returned nan.
#
# Nothing demographic changes, so ne_screen.csv remains valid: effective size is a property of
# the grid, K and m, not of how many markers are carried.
AUTOS = 36      # 32 neutral (0-31) + 4 mechanism (32-35)
XLOCI = 3       # 0 drive, 1 DMI/selection candidate, 2 neutral


@dataclass
class Config:
    grid: int = 20
    K: int = 20
    m: float = 0.10           # vanilla baseline (tuned so autosomes ~5% modern at sampling)
    gamma: float = 0.12       # vanilla baseline
    r: float = 2.0
    lineage: str = "matrilineal"  # "matrilineal" (child=mother, Y introgresses),
                                  # "patrilineal" (child=father, mtDNA introgresses), or "symmetric"
    # --- continuous version of the above ---
    # female_bias is the probability that a CROSS-lineage child takes its MOTHER's lineage
    # (i.e. grows up in and belongs to her group) rather than its father's. A separate draw is
    # made for each such child. 1.0 is matrilineal, 0.5 is no bias, 0.0 is patrilineal, and it
    # reproduces those three named modes exactly. Leave as None to use cfg.lineage instead.
    female_bias: float = None
    s: float = 0.0             # additive selection coefficient against the archaic allele
    sel_locus: str = "none"    # "none", "Y", "auto", or "X" (which locus s acts on)
    # s_mt: selection on the mtDNA, INDEPENDENT of s/sel_locus, so that a run can carry
    # selection on the Y and on the mtDNA at the same time. Positive favours the modern type
    # (archaic carriers survive at 1 - s_mt); negative favours the archaic type, which must be
    # supported because the Muller's-ratchet posterior on the mtDNA fitness gap puts 8.5% of
    # its mass (N_A = 1500) below zero, and silently flooring that at neutrality would bias
    # the integrated answer upward.
    s_mt: float = 0.0
    # Whether the origin deme may still be reseeded with a modern individual AFTER the archaic
    # lineage has gone extinct. False is correct (see _reseed_origin for why, and for how badly
    # it bites at small K); True reproduces the pre-guard behaviour exactly, for reproducing
    # earlier results.
    reseed_origin_after_extinction: bool = False
    # Reseed the origin ONLY when the modern lineage is absent from the whole grid, rather than
    # whenever it is absent from deme 0. See _reseed_origin for the measurement: the old rule
    # inflates modern autosomal ancestry by 1.44 points at the baseline, a quarter of the 4-6%
    # acceptance window, and always in the direction that flatters a mechanism.
    #
    # DEFAULT FLIPPED TO TRUE, 12 Sep 2026. It was False for one day only, so that the screens
    # then in flight stayed comparable with each other; that run is finished and its deme half
    # is being discarded rather than corrected. Set False to reproduce a pre-12-Sep deme
    # result, and expect it to read about 1.4 points high.
    reseed_only_if_globally_extinct: bool = True
    # --- size of the modern founding population -------------------------------------------
    # seed_block = n founds the modern lineage in the n x n block of demes at the origin corner
    # rather than in the single corner deme. DR, 10 Sep: "we could potentially go back to having
    # a larger modern human seed if that helps".
    #
    # WHAT IT CHANGES, AND WHAT IT DOES NOT. A single-deme seed carries ~K/2 = 5 modern males,
    # so the modern Y starts as one of ~5 haplotypes in the founding wave and most of the
    # stochasticity in which Y ends up fixing is decided in the first few dozen generations.
    # Widening the seed raises the number of founding modern lineages ~n^2-fold, which reduces
    # the chance of losing the modern marker to early drift. It does NOT change the equilibrium
    # autosomal proportion much, because that is set by the contact budget along the wave front,
    # not by the founding size -- so this lever should buy determinism without moving the 5%
    # calibration. That prediction is worth checking rather than assuming.
    seed_block: int = 1
    # --- SECOND WAVE (Route 5) ---------------------------------------------------------
    # DR, 10 Sep: the two-wave scenario is "a fresh migration pulse", not the same population
    # changing its residence rule. So at generation wave2_gen the landscape is RELABELLED as
    # resident -- every surviving individual becomes lineage 0, whatever it was -- and a new
    # modern group is founded at the origin under wave2_lineage.
    #
    # The point of relabelling rather than switching a flag is that it makes the second wave a
    # genuine expansion into an occupied landscape, exactly like the first. What differs is what
    # the residents are carrying: after a matrilocal first wave their mtDNA is already modern,
    # so the second wave has nothing to change there, while their Y is archaic and the incoming
    # patrilocal wave retains its own. That is the whole mechanism, and it needs the ALLELIC
    # state of the residents to be preserved across the relabelling -- only `lin` is reset.
    #
    # wave2_gen = 0 disables it. It must be later than the first wave completes (extinction of
    # the original archaic lineage, typically generation 100-500) or the two waves interfere.
    wave2_gen: int = 0
    wave2_lineage: str = "patrilineal"
    # --- carrying capacity that CONTRACTS partway through the run -------------------------
    # K_early (if set) is the per-deme carrying capacity for the first K_early_gens
    # generations; afterwards it drops to cfg.K. This decouples two things that a single
    # constant K forces to be equal:
    #   * the size DURING the expansion, which sets how many modern x archaic contact events
    #     the wave gets, and therefore how much archaic ancestry it can absorb;
    #   * the size AFTER the expansion, which sets the long-run Ne and hence how effective the
    #     mtDNA fitness gap is (the quantity matched to NN-uniparental = 725-1000).
    # With one constant K, shrinking the landscape to match the post-mixture Ne also starves
    # the expansion of contact events -- which is what makes the reduced-F1-fertility (theta)
    # mechanism unreachable, since removing F1 sons already closes off the paternal route for
    # archaic autosomes. A larger K during the expansion restores those events while leaving the
    # long-run Ne almost untouched: at grid 14, K 10 with K_early = 50 for 500 of 6993
    # generations, the harmonic-mean census is ~2080 against ~1960 for constant K, a 6% change.
    K_early: int = None
    K_early_gens: int = 500
    auto_locus: int = 35       # 0-based index of the autosomal locus under selection
                               # ("chromosome 32"; mechanisms live on the top four)
    x_locus: int = 1           # 0-based index of the X locus under selection ("locus 2")
    # --- general epistatic (Dobzhansky-Muller) selection ---
    # An offspring carrying a MODERN allele at the focal locus AND an ARCHAIC allele at the
    # partner locus is removed with probability dm * (partner-archaic factor), optionally only
    # in males (Haldane's rule). Removal is a DROP (the individual is deleted, NOT resampled) --
    # so a 100%-incompatible parent pairing simply yields fewer offspring, never an infinite loop.
    dm: float = 0.0            # DM penetrance (0 = off)
    dm_focal: str = "mt"       # focal (modern-derived) locus: "mt", "Y", "auto", or "X"
    dm_focal_locus: int = 35   # 0-based index if dm_focal is "auto"/"X" (PRDM9-A = chr 32)
    dm_partner: str = "Y"      # partner (archaic-derived) locus: "mt", "Y", "auto", or "X"
    dm_partner_locus: int = 35 # 0-based index if dm_partner is "auto"/"X"
    dm_dom: str = "dominant"   # dominance at a diploid partner: "dominant", "additive", "recessive"
    dm_male_limited: bool = True  # if True, only males are affected (Haldane's rule)
    # --- reduced fertility of F1 hybrid males (Haldane's Rule as a continuous parameter) ---
    # theta is the fertility of an F1 hybrid male relative to a normal male. An "F1 hybrid
    # male" is a son of an ARCHAIC-lineage father and a MODERN-lineage mother -- exactly the
    # individuals who carry an archaic Y into the matrilineally expanding modern population.
    # theta = 1 is no effect; theta = 0 means such males never reproduce, so archaic Y
    # chromosomes reach the modern population but are never transmitted onward.
    theta_f1m: float = 1.0
    # --- obligate male dispersal with return to the father's natal deme -------------------
    # Sex-specific migration. Leave both at None to use cfg.m for everyone, which reproduces
    # the legacy behaviour bit-for-bit (the RNG draw sequence is untouched in that case).
    # m_male = 1.0 makes male dispersal obligate: every male leaves his deme each generation.
    m_male: float = None
    m_female: float = None
    # rho is the probability that a male, instead of dispersing at random, moves to the deme
    # where his FATHER was born -- the "homeland" he is familiar with through his father. This
    # is the mechanism that lets an archaic Y exit a modern matriline: an archaic male disperses
    # into a modern deme and sires F1 sons there, but those sons return to their father's natal
    # (archaic) deme to breed, carrying the archaic Y back out of the modern population.
    # rho = 0 disables it. Founders have no recorded father and are never returned.
    rho: float = 0.0
    # --- matrilineal inbreeding avoidance ---
    # 0 = off; 1 = a pair may not share a mother; 2 = may not share a maternal grandmother
    # (which subsumes sharing a mother). Generations do not overlap, so ancestor-descendant
    # pairings cannot arise and only shared-ancestor tests are needed. The ban applies to
    # everyone, archaic and modern alike: it is inbreeding avoidance, not a preference.
    # The point of the rule is that it falls asymmetrically in a matrilineal expansion. An F1
    # male sits inside his mother's band, where at the wavefront most modern women are his
    # maternal kin; a fully archaic male is related to none of them. So the individuals barred
    # from reproducing are exactly those carrying an archaic Y into the modern matriline, and a
    # low effective theta becomes a consequence of band structure rather than an assumption.
    inbreed_ban: int = 0
    inbreed_tries: int = 12    # redraw attempts before an offspring is simply not produced
# 6993 generations = 200 kyr at 28.6 yr/generation: mixture at 250 ka to the age of the
# sequenced Neandertals at 50 ka. The previous value of 9091 encoded 260 kyr, i.e.
# sampling at the present day, which predates the timeline being pinned down. The two
# give identical results (trajectories are fixed or lost long before either), so this is
# a consistency change rather than a substantive one.
    gens: int = 7003
    sample_gen: int = 6993
    # fix-or-lose early stop: sample as soon as both uniparental markers are absorbed and the
    # archaic lineage is extinct, instead of always running to sample_gen. Set False to
    # reproduce the pre-4-Sep-2026 behaviour exactly.
    stop_when_absorbed: bool = True
    # --- PERIODIC SIZE FLUCTUATION (DR, 4 Sep 2026) -----------------------------------------
    # Climatic worsenings on a Heinrich-event cadence drive the population through crashes and
    # recoveries. Drift is set by the HARMONIC mean of size over time, but a beneficial allele
    # escapes loss whenever size is momentarily large, so a cycling population can hold the
    # coalescent Ne that the PSMC curves require while giving selection far more purchase than
    # a constant population of that size. PSMC has no resolution at millennial scales at a
    # 200 kyr time depth, so this is unconstrained by the data rather than contradicted by it.
    #
    # size(t) = base * (m + a*sin(2*pi*t/T + phase)) / sqrt(R),  m = (R+1)/2, a = (R-1)/2
    # The sqrt(R) divisor is what pins the HARMONIC mean of the cycle to `base`, so turning the
    # fluctuation on does not change the drift anchor -- only how the size is distributed in
    # time. phase = pi/2 starts the run at a peak.
    #
    # Period in generations at 28.6 yr/gen: Dansgaard-Oeschger ~1,470 yr = 51; Heinrich
    # ~6-7 kyr = 210-245. cycle_T = 0 disables the fluctuation entirely.
    cycle_T: int = 0
    cycle_R: float = 1.0           # peak/trough ratio
    cycle_phase: float = 1.5707963267948966   # pi/2: start at the peak

    cap: float = 4.0           # per-deme offspring cap = cap*K (guards against blowups)
    record_every: int = 50
    stop_after_ext: int = 0    # if >0, stop this many gens AFTER archaic extinction. Valid for
                               # reading ancestry MEANS: post-extinction (s=0) ancestry is neutral,
                               # so E[auto%] is stationary and equals the sampling-generation expectation.
    seed: int = 0


# ---------------------------------------------------------------- initialization
def cycle_factor(cfg, t):
    """Multiplier on the carrying capacity at generation t; 1.0 when the cycle is off. The
    harmonic mean of this factor over a full cycle is 1 by construction."""
    if cfg.cycle_T <= 0 or cfg.cycle_R <= 1.0:
        return 1.0
    m, a = (cfg.cycle_R + 1) / 2.0, (cfg.cycle_R - 1) / 2.0
    return float((m + a * np.sin(2 * np.pi * t / cfg.cycle_T + cfg.cycle_phase))
                 / np.sqrt(cfg.cycle_R))


def cfg_early(cfg):
    """The config that applies during the pre-contraction phase (K_early), or cfg itself."""
    return replace(cfg, K=cfg.K_early) if cfg.K_early is not None else cfg


def cap_at(cfg, early, ggen):
    """Which config a given generation runs under, given the precomputed `early`."""
    return early if (cfg.K_early is not None and ggen < cfg.K_early_gens) else cfg


def _init(cfg, rng):
    G, K = cfg.grid, cfg.K
    cols = dict(pos=[], sex=[], lin=[], mt=[], Y=[], auto=[], X=[])
    # f1m marks F1 hybrid males; nobody qualifies in the founding generation
    for i in range(G):
        for j in range(G):
            d = i * G + j
            modern = (i < cfg.seed_block and j < cfg.seed_block)
            v = 1 if modern else 0
            for sx in (1, 0):                       # males then females
                n = int(rng.poisson(K / 2.0))
                if n == 0:
                    continue
                cols["pos"].append(np.full(n, d, np.int32))
                cols["sex"].append(np.full(n, sx, np.int8))
                cols["lin"].append(np.full(n, v, np.int8))
                cols["mt"].append(np.full(n, v, np.int8))
                cols["Y"].append(np.full(n, v if sx == 1 else -1, np.int8))
                cols["auto"].append(np.full((n, AUTOS), 2 * v, np.int8))
                xval = (2 if sx == 0 else 1) * v     # females diploid, males haploid
                cols["X"].append(np.full((n, XLOCI), xval, np.int8))
    P = {}
    for k, v in cols.items():
        P[k] = np.concatenate(v, 0) if v[0].ndim > 1 else np.concatenate(v)
    n0 = P["pos"].size
    P["f1m"] = np.zeros(n0, bool)
    # matrilineal bookkeeping: mat1 = mother's index in the previous generation, mat2 = mother's
    # own mat1 (i.e. the maternal grandmother). Founders get distinct negative sentinels so that
    # no two of them count as kin. Indices are comparable only within a generation, which is all
    # the ban needs since generations do not overlap.
    P["mat1"] = -np.arange(1, n0 + 1, dtype=np.int64)
    P["mat2"] = -np.arange(1, n0 + 1, dtype=np.int64)
    # natal deme, and the father's natal deme ("homeland"). Founders have no father, so
    # fnat = -1 and they are never subject to the rho return rule.
    P["natal"] = P["pos"].astype(np.int32).copy()
    P["fnat"] = np.full(n0, -1, np.int32)
    return P


# ---------------------------------------------------------------- migration
def _migrate(P, cfg, rng):
    G = cfg.grid
    pos = P["pos"]; N = pos.size
    i = pos // G; j = pos % G
    delta = np.array([-G, G, -1, 1])
    valid = np.stack([i > 0, i < G - 1, j > 0, j < G - 1], axis=1)

    if cfg.m_male is None and cfg.m_female is None and cfg.rho <= 0:
        # legacy path, left untouched so the RNG stream is bit-identical to earlier runs
        mv = rng.random(N) < (1.0 - (1.0 - cfg.m) ** valid.sum(1))
        r4 = rng.random((N, 4)); r4[~valid] = -1.0
        P["pos"] = np.where(mv, pos + delta[r4.argmax(1)], pos).astype(np.int32)
        return

    male = P["sex"] == 1
    mm = cfg.m if cfg.m_male is None else cfg.m_male
    mf = cfg.m if cfg.m_female is None else cfg.m_female
    rate = np.where(male, mm, mf)
    # (1 - rate)**n_valid is the chance of staying when each of the n_valid neighbours is
    # offered independently; with rate = 1 and at least one neighbour this is always a move.
    mv = rng.random(N) < (1.0 - (1.0 - rate) ** valid.sum(1))
    r4 = rng.random((N, 4)); r4[~valid] = -1.0
    newpos = np.where(mv, pos + delta[r4.argmax(1)], pos)

    if cfg.rho > 0:
        # A male returns to his father's natal deme instead of dispersing at random. The
        # father's natal deme is one step from the son's birth deme (the father made a single
        # hop before breeding), so this stays within the stepping-stone neighbourhood.
        # Founders carry fnat = -1 and are excluded.
        back = male & (P["fnat"] >= 0) & (rng.random(N) < cfg.rho)
        newpos = np.where(back, P["fnat"], newpos)

    P["pos"] = newpos.astype(np.int32)


# ---------------------------------------------------------------- selection hook
def _dm_penetrance(cfg, sx, mt, Yc, auto, X):
    """Per-offspring probability of removal under the epistatic (Dobzhansky-Muller) rule:
    dm * [modern allele present at the focal locus] * [archaic dose factor at the partner locus],
    optionally male-limited. Returned as a probability in [0, 1]. Removal is a DROP (see run)."""
    n = sx.shape[0]

    def focal_modern():
        f = cfg.dm_focal
        if f == "mt":   return (mt == 1)
        if f == "Y":    return (sx == 1) & (Yc == 1)
        if f == "auto": return auto[:, cfg.dm_focal_locus] >= 1        # carrier of modern allele
        if f == "X":    return X[:, cfg.dm_focal_locus] >= 1
        raise ValueError(cfg.dm_focal)

    def partner_archaic():
        p, dom = cfg.dm_partner, cfg.dm_dom
        if p == "mt":   return (mt == 0).astype(float)
        if p == "Y":    return ((sx == 1) & (Yc == 0)).astype(float)   # hemizygous archaic Y
        if p == "auto":
            k = 2 - auto[:, cfg.dm_partner_locus]; maxd = np.full(n, 2)
        elif p == "X":
            maxd = np.where(sx == 0, 2, 1); k = maxd - X[:, cfg.dm_partner_locus]
        else:
            raise ValueError(p)
        if dom == "dominant":  return (k >= 1).astype(float)
        if dom == "recessive": return (k == maxd).astype(float)
        return k / maxd                                                # additive (per archaic allele)

    p = cfg.dm * focal_modern().astype(float) * partner_archaic()
    if cfg.dm_male_limited:
        p = p * (sx == 1)
    return np.clip(p, 0.0, 1.0)


def _archaic_count(cfg, sx, Yc, auto, X, ident=None):
    """Number of selected-against alleles at the selected locus, per candidate.

    ident: the CHILD's lineage label (1 modern, 0 archaic). Used only for sel_locus == "Y".

    SELECTION ON THE Y IS CONDITIONED ON LINEAGE IDENTITY (DR, 9 Oct 2026). The Scenario 2
    mechanism is prestige, which is conferred by the community a man belongs to, so it
    attaches to the patriline that MATCHES his lineage label: modern Y in a modern-lineage
    individual, archaic Y in an archaic-lineage one. It used to be unconditional -- every
    archaic-Y male was selected against regardless of lineage -- so an incoming modern
    patriline was rewarded inside the archaic population it had no standing in. Under
    matrilineality the label is carried by mothers, so this is "selection in the direction
    of the identity of the sex that transmits identity", stated operationally. The band
    engine (bandsim._archaic_count) applies the identical rule with the band's identity in
    place of the individual's label.
    """
    if cfg.sel_locus == "Y":
        y_mod = (Yc == 1)
        if ident is None:                                        # no label: unconditional
            return np.where((sx == 1) & ~y_mod, 1, 0)            # only males carry a Y
        return np.where((sx == 1) & (y_mod != (np.asarray(ident) == 1)), 1, 0)
    if cfg.sel_locus == "auto":
        return 2 - auto[:, cfg.auto_locus]                     # diploid: 0,1,2 archaic
    if cfg.sel_locus == "X":
        # daughters diploid (0..2 archaic), sons haploid (0..1 archaic)
        maxdose = np.where(sx == 0, 2, 1)
        return maxdose - X[:, cfg.x_locus]
    return np.zeros(sx.shape, np.int64)


# ---------------------------------------------------------------- reproduction
# The female_bias value equivalent to each named residence mode. _repro reproduces the named
# modes bit for bit at these values (see the rule table), so this is an identity rather than
# an approximation, and it is what lets the second wave switch residence rule whichever
# parameterisation the caller chose.
_FB_OF_MODE = {"matrilineal": 1.0, "symmetric": 0.5, "patrilineal": 0.0}


def _repro(P, cfg, rng):
    G2 = cfg.grid ** 2
    pos = P["pos"]; sex = P["sex"]; lin = P["lin"]
    male = sex == 1; female = ~male
    MM = np.bincount(pos[male & (lin == 1)], minlength=G2).astype(float)
    MA = np.bincount(pos[male & (lin == 0)], minlength=G2).astype(float)
    FM = np.bincount(pos[female & (lin == 1)], minlength=G2).astype(float)
    FA = np.bincount(pos[female & (lin == 0)], minlength=G2).astype(float)
    r, g, K = cfg.r, cfg.gamma, cfg.K

    # sorted-order index into male / female pools by (deme, lineage) group
    def pool(mask):
        idx = np.nonzero(mask)[0]
        key = pos[idx] * 2 + lin[idx]
        order = np.argsort(key, kind="stable")
        cnt = np.bincount(key, minlength=2 * G2)
        start = np.zeros(2 * G2, np.int64); start[1:] = np.cumsum(cnt)[:-1]
        return idx[order], cnt, start
    m_ord, m_cnt, m_start = pool(male)
    f_ord, f_cnt, f_start = pool(female)

    # Production rules (father_lin, mother_lin, child_lin, weight). A rule's weight carries r if
    # the CHILD is modern-lineage and gamma for a cross-lineage mating. Child lineage follows the
    # mother (matrilineal), the father (patrilineal), or a random parent 50/50 (symmetric).
    def W(fl, ml, cl, base):
        return base * (r if cl == 1 else 1.0) * (g if fl != ml else 1.0)
    fb = cfg.female_bias
    if fb is not None:
        if not 0.0 <= fb <= 1.0:
            raise ValueError(f"female_bias must lie in [0, 1], got {fb}")
        # A cross-lineage child follows its mother with probability fb, its father otherwise.
        # Rules whose weight is identically zero are skipped below, so fb = 1 / 0.5 / 0 consume
        # the RNG in exactly the same order as the named modes and reproduce them bit for bit.
        rules = [(1, 1, 1, W(1, 1, 1, MM * FM)),
                 (1, 0, 1, W(1, 0, 1, (1.0 - fb) * MM * FA)),
                 (1, 0, 0, W(1, 0, 0, fb         * MM * FA)),
                 (0, 1, 1, W(0, 1, 1, fb         * MA * FM)),
                 (0, 1, 0, W(0, 1, 0, (1.0 - fb) * MA * FM)),
                 (0, 0, 0, W(0, 0, 0, MA * FA))]
        mode = None
    else:
        mode = cfg.lineage
    if mode == "matrilineal":
        rules = [(1, 1, 1, W(1, 1, 1, MM * FM)), (1, 0, 0, W(1, 0, 0, MM * FA)),
                 (0, 1, 1, W(0, 1, 1, MA * FM)), (0, 0, 0, W(0, 0, 0, MA * FA))]
    elif mode == "patrilineal":
        rules = [(1, 1, 1, W(1, 1, 1, MM * FM)), (1, 0, 1, W(1, 0, 1, MM * FA)),
                 (0, 1, 0, W(0, 1, 0, MA * FM)), (0, 0, 0, W(0, 0, 0, MA * FA))]
    elif mode == "symmetric":
        rules = [(1, 1, 1, W(1, 1, 1, MM * FM)),
                 (1, 0, 1, W(1, 0, 1, 0.5 * MM * FA)), (1, 0, 0, W(1, 0, 0, 0.5 * MM * FA)),
                 (0, 1, 1, W(0, 1, 1, 0.5 * MA * FM)), (0, 1, 0, W(0, 1, 0, 0.5 * MA * FM)),
                 (0, 0, 0, W(0, 0, 0, MA * FA))]
    elif mode is not None:
        raise ValueError(f"unknown lineage mode {mode}")
    if fb is not None:
        # Only for the continuous parameterisation: drop rules that carry no weight at all, so
        # that fb = 1 / 0.5 / 0 consume the RNG exactly as the named modes do. This filter is
        # deliberately NOT applied to the named modes, whose draw sequence must not change.
        rules = [t for t in rules if np.any(t[3] > 0)]
    Q = 2.0 * sum(rule[3] for rule in rules)
    has_pair = ((MM + MA) > 0) & ((FM + FA) > 0) & (Q > 0)

    capn = int(cfg.cap * K)
    out = {k: [] for k in ("pos", "sex", "lin", "mt", "Y", "auto", "X", "f1m", "mat1", "mat2",
                           "natal", "fnat")}

    for fl, ml, cl, w in rules:
        lam = np.where(has_pair, K * w / np.where(Q > 0, Q, 1), 0.0)   # per-sex expected count
        for sx in (1, 0):
            tgt = np.minimum(rng.poisson(np.clip(lam, 0, None)), capn).astype(np.int64)
            demes = np.nonzero(tgt > 0)[0]
            if demes.size == 0:
                continue
            cand_deme = np.repeat(demes, tgt[demes])       # exactly `target` per deme (no oversample)
            B = cand_deme.size
            mk = cand_deme * 2 + fl                          # father from (deme, fl) male pool
            fk = cand_deme * 2 + ml                          # mother from (deme, ml) female pool
            fa = m_ord[m_start[mk] + (rng.random(B) * m_cnt[mk]).astype(np.int64)]
            mo = f_ord[f_start[fk] + (rng.random(B) * f_cnt[fk]).astype(np.int64)]

            keep_pair = np.ones(B, bool)
            if cfg.inbreed_ban:
                # Redraw pairs that share a mother (ban 1) or a maternal grandmother (ban 2).
                # A pair still related after inbreed_tries attempts simply produces no offspring,
                # matching the DROP semantics used for selection below: local density falls
                # rather than the pair being replaced by an unrelated one.
                key = "mat1" if cfg.inbreed_ban == 1 else "mat2"
                anc = P[key]
                bad = anc[fa] == anc[mo]
                for _ in range(cfg.inbreed_tries):
                    nb = int(bad.sum())
                    if nb == 0:
                        break
                    idx = np.nonzero(bad)[0]
                    fa[idx] = m_ord[m_start[mk[idx]]
                                    + (rng.random(nb) * m_cnt[mk[idx]]).astype(np.int64)]
                    mo[idx] = f_ord[f_start[fk[idx]]
                                    + (rng.random(nb) * f_cnt[fk[idx]]).astype(np.int64)]
                    bad[idx] = anc[fa[idx]] == anc[mo[idx]]
                keep_pair = ~bad

            # transmit the genetic loci (tracked independently of the lineage label cl)
            childmt = P["mt"][mo].astype(np.int8)                          # mtDNA from the mother
            childY = np.where(sx == 1, P["Y"][fa], -1).astype(np.int8)      # Y from father (sons only)
            fa_auto = P["auto"][fa]; mo_auto = P["auto"][mo]
            childauto = ((rng.random((B, AUTOS)) < fa_auto / 2.0).astype(np.int8)
                         + (rng.random((B, AUTOS)) < mo_auto / 2.0).astype(np.int8))
            mo_tx = (rng.random((B, XLOCI)) < P["X"][mo] / 2.0).astype(np.int8)
            childX = (mo_tx + P["X"][fa]).astype(np.int8) if sx == 0 else mo_tx.astype(np.int8)

            # viability selection as a DROP (no resampling / backfill): additive against archaic
            # alleles and the epistatic (DM) rule. Removed offspring are simply gone, so a
            # 100%-incompatible parent pairing yields fewer offspring and never loops.
            sxarr = np.full(B, sx, np.int8)
            w_surv = np.ones(B)
            if cfg.theta_f1m < 1.0:
                # an F1 hybrid male sires offspring at relative rate theta
                w_surv = w_surv * np.where(P["f1m"][fa], cfg.theta_f1m, 1.0)
            if cfg.s > 0 and cfg.sel_locus != "none":
                k_arch = _archaic_count(cfg, sxarr, childY, childauto, childX, ident=cl)
                w_surv *= np.clip(1.0 - cfg.s * k_arch, 0.0, 1.0)
            if cfg.s_mt != 0.0:
                # haploid, maternally inherited: childmt is 1 modern, 0 archaic
                dis = (childmt == 0) if cfg.s_mt > 0 else (childmt == 1)
                w_surv *= np.clip(1.0 - abs(cfg.s_mt) * dis, 0.0, 1.0)
            if cfg.dm > 0:
                w_surv = w_surv * (1.0 - _dm_penetrance(cfg, sxarr, childmt, childY, childauto, childX))
            sel = np.nonzero((rng.random(B) < w_surv) & keep_pair)[0]
            if sel.size == 0:
                continue
            out["pos"].append(cand_deme[sel].astype(np.int32))
            out["sex"].append(np.full(sel.size, sx, np.int8))
            out["lin"].append(np.full(sel.size, cl, np.int8))
            out["mt"].append(childmt[sel])
            out["Y"].append(childY[sel])
            out["auto"].append(childauto[sel])
            out["X"].append(childX[sel])
            # this cohort are F1 hybrid males iff sons of an archaic father and a
            # modern mother (rule fl=0, ml=1)
            out["f1m"].append(np.full(sel.size, (sx == 1 and fl == 0 and ml == 1), bool))
            out["mat1"].append(mo[sel].astype(np.int64))          # mother's index this generation
            out["mat2"].append(P["mat1"][mo[sel]].astype(np.int64))   # maternal grandmother
            out["natal"].append(cand_deme[sel].astype(np.int32))      # deme of birth
            out["fnat"].append(P["natal"][fa[sel]].astype(np.int32))  # father's natal deme

    if not out["pos"]:
        return None
    P2 = {k: np.concatenate(out[k])
          for k in ("pos", "sex", "lin", "mt", "Y", "f1m", "mat1", "mat2",
                    "natal", "fnat")}
    P2["auto"] = np.concatenate(out["auto"], 0)
    P2["X"] = np.concatenate(out["X"], 0)
    return P2


def _reseed_origin(P, cfg):
    # Reseeding exists for ONE purpose: to stop the founding modern lineage, which starts as a
    # handful of individuals in deme 0, from being lost to chance before the wave is
    # established. Once the archaic lineage is extinct the wave is over and reseeding has no
    # job left -- but the trigger below is "deme 0 lacks a modern-lineage individual of this
    # sex", which after extinction is simply a Poisson(K/2) = 0 event. It therefore keeps
    # firing for the rest of the run, each time injecting a 100% modern individual (mt = 1,
    # Y = 1, autosomes all modern) into a population whose whole point is that it has been
    # archaicised. That is a steady, artefactual mutation pressure towards modern alleles, and
    # it scales steeply as K falls:
    #
    #     K            20      12       8       7       4
    #     injections   0.6    34.7   256.2   422.3  1892.8    (per 6993-generation run)
    #
    # Negligible at the K = 20 the published runs used, which is why it went unnoticed, but it
    # dominates any attempt to shrink K to lower the effective population size: it is what made
    # NEUTRAL modern mtDNA drift UPWARDS after the wave at K = 7 and K = 10. Guard it.
    if cfg.reseed_origin_after_extinction is False and (P["lin"] == 0).sum() == 0:
        return P
    # SECOND, INDEPENDENT LEAK, found by DR on 11 Sep 2026: "Why do we do this when there are
    # still modern individuals elsewhere in the landscape? The point of reseeding was to prevent
    # extinction, but if moderns are persisting in other grid points I don't think we need to
    # reseed." He is right, and it is not a small effect. The guard above only asks whether the
    # ARCHAIC lineage still exists; it never asks whether the MODERN lineage needs rescuing. So
    # the reseed also fires whenever deme 0 happens to lose its modern occupants during a
    # perfectly healthy expansion, injecting a fully modern individual into a landscape that is
    # supposed to be archaicising.
    #
    # Measured at the baseline (grid 12, K 11, m 0.10, gamma 0.12, r 2), 300 replicates each:
    #
    #     guard                              autosomal ancestry      Y        generations
    #     current (archaic-extinction only)    9.07% +- 0.21       4.33%          223
    #     also require global modern loss      7.63% +- 0.23       3.33%          238
    #
    # 1.44 points, z = 4.7. That is a quarter of the acceptance window, so it decides which
    # cells pass a 4-6% screen. Per run the reseed fires 0-51 times, of which 6-24 occur while
    # the modern lineage is alive elsewhere on the grid.
    #
    # DEFAULT LEFT AT THE OLD BEHAVIOUR ON PURPOSE. Flipping it silently would invalidate the
    # 36-locus screens currently on the queue and make them incomparable with each other. Set
    # reseed_only_if_globally_extinct=True to get the correct behaviour, and rerun.
    if getattr(cfg, "reseed_only_if_globally_extinct", False) and (P["lin"] == 1).any():
        return P
    for sx in (1, 0):
        if not ((P["pos"] == 0) & (P["lin"] == 1) & (P["sex"] == sx)).any():
            P["pos"] = np.append(P["pos"], 0)
            P["sex"] = np.append(P["sex"], sx)
            P["lin"] = np.append(P["lin"], 1)
            P["mt"] = np.append(P["mt"], 1)
            P["Y"] = np.append(P["Y"], 1 if sx == 1 else -1)
            P["auto"] = np.vstack([P["auto"], np.full((1, AUTOS), 2, np.int8)])
            P["X"] = np.vstack([P["X"], np.full((1, XLOCI), 2 if sx == 0 else 1, np.int8)])
            P["f1m"] = np.append(P["f1m"], False)
            tag = -(int(P["mat1"].min()) * 0 + abs(int(P["mat1"].min())) + 1)
            P["mat1"] = np.append(P["mat1"], tag)
            P["mat2"] = np.append(P["mat2"], tag)
            P["natal"] = np.append(P["natal"], 0).astype(np.int32)
            P["fnat"] = np.append(P["fnat"], -1).astype(np.int32)
    return P


# ---------------------------------------------------------------- stats & driver
def _stats(P, exclude=(), exclude_x=()):
    # ancestry is reported AMONG MODERN-LINEAGE individuals (the expanding population),
    # matching the manuscript's framing and the old engine. After archaic extinction the
    # whole population is modern-lineage, so this is just the surviving population.
    # `exclude` drops autosomal loci and `exclude_x` drops X loci from the reported averages,
    # so that a locus under selection or participating in a DM is not counted in the
    # "background" ancestry estimate (the remaining loci are unlinked to it).
    mod = P["lin"] == 1
    male = P["sex"] == 1
    mmale = mod & male
    modf = mod & (P["sex"] == 0)
    xcols = [c for c in range(XLOCI) if c not in exclude_x]
    def xfrac(mask, diploid):
        if not mask.any() or not xcols:
            return np.nan
        cap = 2.0 if diploid else 1.0
        return float((P["X"][mask][:, xcols].sum(1) / (len(xcols) * cap)).mean())
    return dict(
        mt_mod=float((P["mt"][mod] == 1).mean()) if mod.any() else np.nan,
        Y_mod=float((P["Y"][mmale] == 1).mean()) if mmale.any() else np.nan,
        auto_mod=(float(P["auto"][mod][:, [c for c in range(AUTOS) if c not in exclude]].mean() / 2.0)
                  if mod.any() else np.nan),
        X_mod=(np.nanmean([v for v in (xfrac(modf, True), xfrac(mmale, False)) if v == v])
               if mod.any() else np.nan),
        # WHOLE-LANDSCAPE frequencies, added 13 Sep 2026 alongside the _mod ones rather than
        # replacing them. mt_mod/Y_mod/auto_mod are computed over lin == 1 only. For a single
        # wave that is harmless: the archaic lineage goes extinct, so lin == 1 is everyone by
        # the sampling generation. After a TWO-WAVE relabelling it is not -- lin is reset to 0
        # for the whole landscape and only the new founder group carries lin == 1 -- so the
        # trajectory between wave 2 and label absorption describes the incoming group, and
        # reads 100% at both markers from the generation the second wave is founded. Timing
        # the expansion off that is meaningless, which is exactly the mistake these fields
        # exist to prevent. The end-of-run summary is unaffected, because it is taken at
        # sample_gen, long after the label has absorbed and lin == 1 is everyone again.
        mt_all=float((P["mt"] == 1).mean()) if P["pos"].size else np.nan,
        Y_all=(float((P["Y"][male] == 1).mean()) if male.any() else np.nan),
        auto_all=(float(P["auto"][:, [c for c in range(AUTOS) if c not in exclude]].mean() / 2.0)
                  if P["pos"].size else np.nan),
        arc=int((P["lin"] == 0).sum()), pop=int(P["pos"].size))


def run(cfg):
    rng = np.random.default_rng(cfg.seed)
    excl = set(); excl_x = set()
    if cfg.sel_locus == "auto": excl.add(cfg.auto_locus)
    if cfg.dm > 0 and cfg.dm_focal == "auto": excl.add(cfg.dm_focal_locus)
    if cfg.dm > 0 and cfg.dm_partner == "auto": excl.add(cfg.dm_partner_locus)
    if cfg.sel_locus == "X": excl_x.add(cfg.x_locus)
    if cfg.dm > 0 and cfg.dm_focal == "X": excl_x.add(cfg.dm_focal_locus)
    if cfg.dm > 0 and cfg.dm_partner == "X": excl_x.add(cfg.dm_partner_locus)
    excl = tuple(excl); excl_x = tuple(excl_x)
    # The expansion is founded at whatever capacity applies in generation 0.
    cfg_early = replace(cfg, K=cfg.K_early) if cfg.K_early is not None else cfg
    P = _init(cfg_early, rng)
    ts = {k: [] for k in ("gen", "mt_mod", "Y_mod", "auto_mod", "X_mod",
                          "mt_all", "Y_all", "auto_all", "arc", "pop")}
    ext = None
    for ggen in range(cfg.gens):
        # capacity for THIS generation: the contraction is a step change at K_early_gens
        cur = cfg_early if (cfg.K_early is not None and ggen < cfg.K_early_gens) else cfg
        if cfg.cycle_T > 0 and cfg.cycle_R > 1.0:
            cur = replace(cur, K=max(2, int(round(cur.K * cycle_factor(cfg, ggen)))))
        # SECOND WAVE. Relabel everyone as resident, keeping their alleles, and found a fresh
        # modern group at the origin under the second residence rule. Done before migration so
        # the new founders disperse in the same generation they appear, as the first wave's do.
        if cfg.wave2_gen and ggen == cfg.wave2_gen:
            P["lin"][:] = 0
            P["f1m"][:] = False
            d0 = 0                                   # origin deme, as in _init
            here = P["pos"] == d0
            if here.any():
                P["lin"][here] = 1
                P["mt"][here] = 1
                male = here & (P["sex"] == 1)
                P["Y"][male] = 1
                P["auto"][here, :] = 2
                P["X"][here, :] = np.where(P["sex"][here, None] == 0, 2, 1)
                # THE SWITCH MUST APPLY TO WHICHEVER PARAMETERISATION IS IN USE.
                #
                # _repro reads cfg.lineage ONLY when cfg.female_bias is None; when female_bias
                # is set it takes the continuous path and never consults cfg.lineage at all.
                # So flipping lineage alone silently did nothing for any caller that passes a
                # female_bias -- which robust_run does unconditionally, at 1.0. Every two-wave
                # cell in the Cartesian sweep therefore ran its SECOND wave under the SAME
                # matrilocal rule as the first, the modern Y never had a route in, and the
                # cells came back with Y at 0-2% and p_joint ~0.02. The standalone
                # twowave_run.py leaves female_bias at None, takes the named-mode path, and
                # gets Y at 100% from the identical geometry -- which is how the two disagreed
                # by a factor of fifty on the same cell.
                #
                # Switching both keeps the two parameterisations in step.
                cur = replace(cur, lineage=cfg.wave2_lineage)
                cfg = replace(cfg, lineage=cfg.wave2_lineage)
                cfg_early = replace(cfg_early, lineage=cfg.wave2_lineage)
                if cfg.female_bias is not None:
                    fb2 = _FB_OF_MODE.get(cfg.wave2_lineage)
                    if fb2 is None:
                        raise ValueError(
                            f"wave2_lineage={cfg.wave2_lineage!r} has no female_bias "
                            f"equivalent, and female_bias is set, so the second wave's "
                            f"residence rule cannot be expressed")
                    cur = replace(cur, female_bias=fb2)
                    cfg = replace(cfg, female_bias=fb2)
                    cfg_early = replace(cfg_early, female_bias=fb2)
                ext = None                           # the second wave has its own extinction
        elif cfg.wave2_gen and ggen > cfg.wave2_gen:
            cur = replace(cur, lineage=cfg.wave2_lineage)
            if cfg.female_bias is not None:
                cur = replace(cur, female_bias=_FB_OF_MODE[cfg.wave2_lineage])

        _migrate(P, cur, rng)
        nxt = _repro(P, cur, rng)
        if nxt is None:
            break
        P = _reseed_origin(nxt, cur)
        if ggen % cfg.record_every == 0 or ggen == cfg.gens - 1:
            s = _stats(P, excl, excl_x); ts["gen"].append(ggen)
            for k in s:
                ts[k].append(s[k])
        if ext is None and (P["lin"] == 0).sum() == 0:
            ext = ggen
        # FIX-OR-LOSE EARLY STOP (DR, 4 Sep 2026). Sample at cfg.sample_gen OR as soon as both
        # uniparental markers are absorbed -- fixed or lost -- whichever comes first. Once both
        # are absorbed nothing further changes at either marker, and autosomal ancestry is
        # neutral, so the reported means are unaffected. Two guards: the archaic lineage must
        # already be extinct (ext is not None), because until then the lineage sweep is still
        # reshaping the modern-lineage composition that _stats averages over, AND because the
        # origin reseed can reintroduce a lost modern marker while archaic demes persist.
        # The early stop must not fire BEFORE the second wave arrives, or the run ends while
        # the landscape is still single-wave and wave2_gen never happens -- which is exactly
        # what made the first two-wave test return the single-wave answer.
        # AND THE SAME GUARD FOR K_early, MISSING FOR THE SAME REASON (13 Sep 2026). This is
        # the third scheduled event an unguarded absorption stop has jumped in front of, after
        # wave2_gen here and the wave-2 guard in the band screen. K_early raises the per-deme
        # capacity for the first K_early_gens generations and then drops it to cfg.K; the whole
        # point is that the long-run census, and so Ne, is the anchored one. Absorption
        # typically completes around generation 100-200, well before a K_early_gens of 300-500,
        # so without this clause the run ends DURING the high-capacity phase and reports its
        # statistics at a census several times the anchor. Measured at grid 12 / K 11 /
        # K_early 50 / 500 generations, theta = 0, gamma = 0.42:
        #
        #     stop unguarded   autosomal 7.72%   census at sampling 7,291
        #     stop suppressed  autosomal 5.88%   census at sampling 1,550
        #
        # The second is the cell the note would want to report -- inside the 3-7% window at the
        # anchored census -- and the first is the same cell measured in the wrong world.
        if (cfg.stop_when_absorbed and ext is not None and ggen % 10 == 0
                and ggen > 0 and ggen < cfg.gens - 1
                and (cfg.K_early is None or ggen > cfg.K_early_gens)
                and (not cfg.wave2_gen or ggen > cfg.wave2_gen)):
            male = P["sex"] == 1
            pm = float((P["mt"] == 1).mean())
            py = float((P["Y"][male] == 1).mean()) if male.any() else 0.0
            if pm in (0.0, 1.0) and py in (0.0, 1.0):
                s = _stats(P, excl, excl_x)
                ts["gen"].append(cfg.sample_gen)     # label it as the sampling point
                for k in s:
                    ts[k].append(s[k])
                break
        # THE SAME WAVE-2 GUARD AS THE ABSORPTION STOP ABOVE, AND IT WAS MISSING HERE.
        # (Found 12 Sep 2026 while explaining why Route 4 scored near zero.) The screen runs
        # with stop_after_ext set, so a replicate ends a few generations after the FIRST wave
        # finishes -- typically generation 80-150. wave2_gen is 500, 1,000 or 2,000. Every
        # two-wave cell in the stage-1 screen therefore terminated before the second wave was
        # ever founded, and reported the single-wave answer: mtDNA modern, Y archaic, autosomal
        # ancestry at the one-wave value. The comment on the absorption stop above warns about
        # exactly this failure and the guard was added there but not here.
        #
        # Without the guard the two-wave route cannot score above zero no matter what the
        # parameters are, because the mechanism never runs.
        if (ext is not None and cfg.stop_after_ext > 0
                and ggen >= ext + cfg.stop_after_ext
                and (not cfg.wave2_gen or ggen > cfg.wave2_gen)):
            s = _stats(P, excl, excl_x); ts["gen"].append(ggen)
            for k in s:
                ts[k].append(s[k])
            break
    return {k: np.array(v) for k, v in ts.items()}, ext


if __name__ == "__main__":
    import sys
    quick = "--quick" in sys.argv
    cfg = Config(grid=(8 if quick else 20), gens=(1200 if quick else 7003),
                 sample_gen=(1190 if quick else 6993))
    out, ext = run(cfg)
    print(f"grid={cfg.grid} gens={cfg.gens} extinction={ext}")
    for k in ("mt_mod", "Y_mod", "auto_mod", "X_mod", "pop"):
        print(f"  {k:9s} {out[k][-1]:.3f}")
```
