## Supplementary Information section 3 code base for "Hypothesis: A modern human range expansion ∼300,000 years ago had a critical role in Neandertal origins": spatial_sim3.py

```
"""
spatial_sim3.py -- spatial_sim2 extended with three mechanisms that evade the leak problem.

Motivation. For a DMI to keep sterilizing archaic-Y carriers for ~7,000 generations, the modern
trigger must sit on something maternally transmitted that cannot introgress (under matriliny,
only mtDNA), and the archaic partner must be co-inherited with the archaic Y in every prospective
father forever (only the Y itself). mtDNA x Y is therefore the unique SEALED two-locus
architecture, and Table S2.1 already contains it. This engine adds the mechanisms that TOLERATE
leakage instead of being sealed against it:

  (1) POLYGENIC MITO-NUCLEAR RESTORERS (cfg.restorers = n > 0). Modern mtDNA is incompatible
      with archaic alleles at the last n autosomal loci ("restorers"), male-limited. The
      biological reading is the cytoplasmic-male-sterility / mother's-curse architecture:
      male-harming variants accumulate on mtDNA unopposed (selection never sees mtDNA in males),
      each lineage evolves its own nuclear compensators, and a hybrid male inherits the modern
      mtDNA without its restorers. The father is the leak, and now he must transmit modern
      alleles at ALL n loci at once, so the escape rate falls geometrically with n.
        dm_dom = "dominant": one archaic allele at any restorer sterilizes (OR-logic).
        dm_dom = "additive": survival = prod_l (1 - dm * dose_l / 2); a pure-archaic father's
                 F1 son is heterozygous at all n loci, so F1 fertility = (1-dm/2)^n -- with
                 dm = 1, theta_F1 = 0.5^n, crossing the 4% line at n = 5.
      restorers = 1 with "dominant" is EXACTLY the mtDNA x autosome (dominant) row of
      Table S2.1 (locus 7), which anchors the family to published numbers.

  (2) CRYPTIC X-LINKED DRIVE AGAINST THE ARCHAIC Y (cfg.xdrive = d > 0). X locus 0 is a driver:
      in a male carrying the MODERN allele there together with an ARCHAIC Y, each Y-bearing
      conception is converted to an X-bearing one (a daughter) with probability d. The modern Y
      carries the coevolved suppressor and is immune. This is the Hurst-Pomiankowski route to
      Haldane's Rule (hybrid dysfunction as unmasked drive; cf. mouse Slx/Sly, Phadnis & Orr's
      Overdrive). Unlike a sterility DMI, the F1 male still reproduces -- his "failure" is
      converted into wavefront daughters -- and the driver X gains a transmission advantage
      exactly when paired with an archaic Y, so it is pulled toward the front rather than
      diluted: a self-repairing barrier rather than a sealed one.
      cfg.xdrive_cost = c removes offspring of drive-EXPRESSING males (modern X0 + archaic Y)
      with probability c, the standard fertility cost of sperm-killing drive.

  (3) HERITABLE ESCAPE FROM THE SEALED DMI (cfg.esc_eps > 0, with the usual dm=1 mtDNA x Y
      incompatibility). Each time the DMI would remove an offspring, with probability eps the
      offspring instead survives and its Y converts to allele 2, a heritably COMPATIBLE archaic
      Y (a compensatory mutation, or a Y that acquired the modern pseudoautosomal region through
      a rare successful pairing). Allele 2 is immune to the DMI forever after and is counted as
      archaic in the output. This stress-tests every mechanism above: an escape that becomes
      heritable is far more dangerous than the i.i.d. escapes simulated elsewhere.

Everything else -- demography, migration, the parent-pairing reproduction rules, additive
selection, the single-pair DM machinery, theta_f1m, sex-specific migration, rho, the inbreeding
ban -- is inherited unchanged from spatial_sim2 (same defaults, so with the three new knobs off
this engine is statistically identical to spatial_sim2; the RNG draw sequence differs only when
a new mechanism is switched on).

Sampling conventions follow the house standard: sample_gen 6993, stop_after_ext 0.
"""

import numpy as np
from dataclasses import replace
from dataclasses import dataclass, replace

import spatial_sim2 as S2
from spatial_sim2 import AUTOS, XLOCI, _init, _migrate, _reseed_origin, _stats, _dm_penetrance, \
    _archaic_count


@dataclass
class Config(S2.Config):
    # (1) polygenic mito-nuclear restorers: 0 = off; n > 0 uses the LAST n autosomal loci.
    # rest_pen is the restorer rule's own penetrance knob -- deliberately separate from cfg.dm,
    # which would otherwise also switch on the single-pair DM machinery.
    restorers: int = 0
    rest_pen: float = 1.0
    # (2) X-linked drive: probability that a Y-bearing conception of an expressing male
    # (modern allele at X locus 0, archaic Y) becomes a daughter instead. 0 = off.
    xdrive: float = 0.0
    xdrive_cost: float = 0.0   # fertility cost paid by expressing males (offspring removed w.p. c)
    # xdrv_res: heritable RESISTANCE stress test. Each time a Y-bearing conception would be
    # flipped, with this probability the archaic Y instead goes through and converts to allele
    # 2, a driver-resistant archaic Y that is immune to the drive forever after (counted as
    # archaic in the output). The drive analogue of esc_eps.
    xdrv_res: float = 0.0
    # xdrv_all: no-suppressor CONTROL. The driver kills Y-bearing sperm regardless of which Y
    # the male carries (the modern Y loses its immunity), so the coevolved-suppressor asymmetry
    # is removed and only raw drive remains.
    xdrv_all: bool = False
    # (3) heritable escape from the DM removal: probability that a removed offspring instead
    # survives with its Y converted to the heritably compatible allele 2. 0 = off.
    esc_eps: float = 0.0
    # (4) asymmetric interbreeding. gamma_export scales MODERN-father x ARCHAIC-mother matings
    # (under matriliny the children join the archaic side, carrying modern Y's out with them);
    # gamma_import scales ARCHAIC-father x MODERN-mother matings (the channel that brings
    # archaic autosomes and archaic Y's into the matriline). None -> cfg.gamma for that
    # direction, reproducing the symmetric model. The point: one export event converts an
    # archaic patriline's Y wholesale and forever, while its autosomal contribution halves
    # every archaic-side generation -- so export modernizes the archaic pool's Y's
    # geometrically faster than its autosomes, decoupling what the import channel later
    # delivers.
    gamma_export: float = None
    gamma_import: float = None
    # (5) male reproductive skew: each generation only a random fraction skew_frac of males is
    # eligible to father children (paternity concentration; females untouched; eligibility is
    # redrawn every generation). This collapses the Y's effective population size relative to
    # the autosomes', so drift finishes whatever majority the other mechanisms achieve.
    # 1.0 = off.
    skew_frac: float = 1.0


DRIVER_XLOCUS = 0              # X locus carrying the driver (locus 1 stays as background)


def _restorer_survival(cfg, sx, mt, auto):
    """Survival multiplier for the polygenic mito-nuclear DMI. Trigger: modern mtDNA
    (male-limited if dm_male_limited). Partners: archaic alleles at the last cfg.restorers
    autosomal loci. 'dominant' = any archaic allele at any restorer is lethal to the male
    offspring (with probability dm); 'additive' = per-locus survival (1 - dm*dose/2),
    multiplied across loci as independent causes of sterility."""
    lo = AUTOS - cfg.restorers
    dose = (2 - auto[:, lo:]).astype(float)          # archaic allele count per restorer, 0..2
    if cfg.dm_dom == "dominant":
        surv = np.where((dose >= 1).any(1), 1.0 - cfg.rest_pen, 1.0)
    elif cfg.dm_dom == "additive":
        surv = np.prod(1.0 - cfg.rest_pen * dose / 2.0, axis=1)
    else:
        raise ValueError(f"restorers supports dominant/additive, got {cfg.dm_dom}")
    trig = (mt == 1)
    if cfg.dm_male_limited:
        trig = trig & (sx == 1)
    return np.where(trig, surv, 1.0)


# ---------------------------------------------------------------- reproduction (forked)
def _repro(P, cfg, rng):
    G2 = cfg.grid ** 2
    pos = P["pos"]; sex = P["sex"]; lin = P["lin"]
    male = sex == 1; female = ~male
    # (5) reproductive skew: ineligible males are excluded from BOTH the mating-weight counts
    # and the father pools, so the two stay consistent and no empty pool is ever sampled.
    if cfg.skew_frac < 1.0:
        male = male & (rng.random(pos.size) < cfg.skew_frac)
    MM = np.bincount(pos[male & (lin == 1)], minlength=G2).astype(float)
    MA = np.bincount(pos[male & (lin == 0)], minlength=G2).astype(float)
    FM = np.bincount(pos[female & (lin == 1)], minlength=G2).astype(float)
    FA = np.bincount(pos[female & (lin == 0)], minlength=G2).astype(float)
    r, g, K = cfg.r, cfg.gamma, cfg.K
    g_exp = g if cfg.gamma_export is None else cfg.gamma_export   # modern father x archaic mother
    g_imp = g if cfg.gamma_import is None else cfg.gamma_import   # archaic father x modern mother

    def pool(mask):
        idx = np.nonzero(mask)[0]
        key = pos[idx] * 2 + lin[idx]
        order = np.argsort(key, kind="stable")
        cnt = np.bincount(key, minlength=2 * G2)
        start = np.zeros(2 * G2, np.int64); start[1:] = np.cumsum(cnt)[:-1]
        return idx[order], cnt, start
    m_ord, m_cnt, m_start = pool(male)
    f_ord, f_cnt, f_start = pool(female)

    def W(fl, ml, cl, base):
        gdir = 1.0 if fl == ml else (g_exp if fl == 1 else g_imp)
        return base * (r if cl == 1 else 1.0) * gdir
    fb = cfg.female_bias
    if fb is not None:
        if not 0.0 <= fb <= 1.0:
            raise ValueError(f"female_bias must lie in [0, 1], got {fb}")
        rules = [(1, 1, 1, W(1, 1, 1, MM * FM)),
                 (1, 0, 1, W(1, 0, 1, (1.0 - fb) * MM * FA)),
                 (1, 0, 0, W(1, 0, 0, fb * MM * FA)),
                 (0, 1, 1, W(0, 1, 1, fb * MA * FM)),
                 (0, 1, 0, W(0, 1, 0, (1.0 - fb) * MA * FM)),
                 (0, 0, 0, W(0, 0, 0, MA * FA))]
        rules = [t for t in rules if np.any(t[3] > 0)]
    elif cfg.lineage == "matrilineal":
        rules = [(1, 1, 1, W(1, 1, 1, MM * FM)), (1, 0, 0, W(1, 0, 0, MM * FA)),
                 (0, 1, 1, W(0, 1, 1, MA * FM)), (0, 0, 0, W(0, 0, 0, MA * FA))]
    elif cfg.lineage == "patrilineal":
        rules = [(1, 1, 1, W(1, 1, 1, MM * FM)), (1, 0, 1, W(1, 0, 1, MM * FA)),
                 (0, 1, 0, W(0, 1, 0, MA * FM)), (0, 0, 0, W(0, 0, 0, MA * FA))]
    elif cfg.lineage == "symmetric":
        rules = [(1, 1, 1, W(1, 1, 1, MM * FM)),
                 (1, 0, 1, W(1, 0, 1, 0.5 * MM * FA)), (1, 0, 0, W(1, 0, 0, 0.5 * MM * FA)),
                 (0, 1, 1, W(0, 1, 1, 0.5 * MA * FM)), (0, 1, 0, W(0, 1, 0, 0.5 * MA * FM)),
                 (0, 0, 0, W(0, 0, 0, MA * FA))]
    else:
        raise ValueError(f"unknown lineage mode {cfg.lineage}")
    Q = 2.0 * sum(rule[3] for rule in rules)
    has_pair = ((MM + MA) > 0) & ((FM + FA) > 0) & (Q > 0)

    capn = int(cfg.cap * K)
    out = {k: [] for k in ("pos", "sex", "lin", "mt", "Y", "auto", "X", "f1m", "mat1", "mat2",
                           "natal", "fnat")}

    for fl, ml, cl, w in rules:
        lam = np.where(has_pair, K * w / np.where(Q > 0, Q, 1), 0.0)
        for sx in (1, 0):
            tgt = np.minimum(rng.poisson(np.clip(lam, 0, None)), capn).astype(np.int64)
            demes = np.nonzero(tgt > 0)[0]
            if demes.size == 0:
                continue
            cand_deme = np.repeat(demes, tgt[demes])
            B = cand_deme.size
            mk = cand_deme * 2 + fl
            fk = cand_deme * 2 + ml
            fa = m_ord[m_start[mk] + (rng.random(B) * m_cnt[mk]).astype(np.int64)]
            mo = f_ord[f_start[fk] + (rng.random(B) * f_cnt[fk]).astype(np.int64)]

            keep_pair = np.ones(B, bool)
            if cfg.inbreed_ban:
                key = "mat1" if cfg.inbreed_ban == 1 else "mat2"
                anc = P[key]
                bad = anc[fa] == anc[mo]
                for _ in range(cfg.inbreed_tries):
                    nb = int(bad.sum())
                    if nb == 0:
                        break
                    idx = np.nonzero(bad)[0]
                    fa[idx] = m_ord[m_start[mk[idx]]
                                    + (rng.random(nb) * m_cnt[mk[idx]]).astype(np.int64)]
                    mo[idx] = f_ord[f_start[fk[idx]]
                                    + (rng.random(nb) * f_cnt[fk[idx]]).astype(np.int64)]
                    bad[idx] = anc[fa[idx]] == anc[mo[idx]]
                keep_pair = ~bad

            childmt = P["mt"][mo].astype(np.int8)
            childY = np.where(sx == 1, P["Y"][fa], -1).astype(np.int8)
            fa_auto = P["auto"][fa]; mo_auto = P["auto"][mo]
            childauto = ((rng.random((B, AUTOS)) < fa_auto / 2.0).astype(np.int8)
                         + (rng.random((B, AUTOS)) < mo_auto / 2.0).astype(np.int8))
            mo_tx = (rng.random((B, XLOCI)) < P["X"][mo] / 2.0).astype(np.int8)
            childX = (mo_tx + P["X"][fa]).astype(np.int8) if sx == 0 else mo_tx.astype(np.int8)

            sxarr = np.full(B, sx, np.int8)

            # ---- (2) X-linked drive: an expressing father's Y-bearing conception becomes a
            # daughter with probability xdrive. The child then needs the father's X as well.
            if cfg.xdrive > 0 and sx == 1:
                if cfg.xdrv_all:
                    expressing = (P["X"][fa][:, DRIVER_XLOCUS] == 1) & (P["Y"][fa] >= 0)
                else:
                    expressing = (P["X"][fa][:, DRIVER_XLOCUS] == 1) & (P["Y"][fa] == 0)
                flip = expressing & (rng.random(B) < cfg.xdrive)
                if cfg.xdrv_res > 0:
                    res = flip & (rng.random(B) < cfg.xdrv_res)
                    if res.any():
                        childY = np.where(res, 2, childY).astype(np.int8)
                        flip = flip & ~res
                if flip.any():
                    sxarr = np.where(flip, 0, sxarr).astype(np.int8)
                    childY = np.where(flip, -1, childY).astype(np.int8)
                    childX = np.where(flip[:, None],
                                      (mo_tx + P["X"][fa]).astype(np.int8), childX)

            w_surv = np.ones(B)
            if cfg.theta_f1m < 1.0:
                w_surv = w_surv * np.where(P["f1m"][fa], cfg.theta_f1m, 1.0)
            if cfg.xdrive > 0 and cfg.xdrive_cost > 0:
                expressing_fa = (P["X"][fa][:, DRIVER_XLOCUS] == 1) & (P["Y"][fa] == 0)
                w_surv = w_surv * np.where(expressing_fa, 1.0 - cfg.xdrive_cost, 1.0)
            if cfg.s > 0 and cfg.sel_locus != "none":
                k_arch = _archaic_count(cfg, sxarr, childY, childauto, childX, ident=cl)
                w_surv *= np.clip(1.0 - cfg.s * k_arch, 0.0, 1.0)
            # ---- (1) polygenic mito-nuclear restorers
            if cfg.restorers > 0:
                w_surv = w_surv * _restorer_survival(cfg, sxarr, childmt, childauto)

            alive = (rng.random(B) < w_surv) & keep_pair
            # ---- single-pair DM, with (3) the heritable escape channel
            if cfg.dm > 0:
                dmp = _dm_penetrance(cfg, sxarr, childmt, childY, childauto, childX)
                hit = rng.random(B) < dmp
                if cfg.esc_eps > 0:
                    escp = hit & (rng.random(B) < cfg.esc_eps)
                    if escp.any():
                        childY = np.where(escp, 2, childY).astype(np.int8)
                    hit = hit & ~escp
                alive = alive & ~hit

            sel = np.nonzero(alive)[0]
            if sel.size == 0:
                continue
            out["pos"].append(cand_deme[sel].astype(np.int32))
            out["sex"].append(sxarr[sel])
            out["lin"].append(np.full(sel.size, cl, np.int8))
            out["mt"].append(childmt[sel])
            out["Y"].append(childY[sel])
            out["auto"].append(childauto[sel])
            out["X"].append(childX[sel])
            # F1 hybrid males: sons of an archaic father and a modern mother. Under drive a
            # flipped child is a daughter and does not qualify.
            out["f1m"].append((sxarr[sel] == 1) & (fl == 0) & (ml == 1))
            out["mat1"].append(mo[sel].astype(np.int64))
            out["mat2"].append(P["mat1"][mo[sel]].astype(np.int64))
            out["natal"].append(cand_deme[sel].astype(np.int32))
            out["fnat"].append(P["natal"][fa[sel]].astype(np.int32))

    if not out["pos"]:
        return None
    P2 = {k: np.concatenate(out[k])
          for k in ("pos", "sex", "lin", "mt", "Y", "f1m", "mat1", "mat2", "natal", "fnat")}
    P2["auto"] = np.concatenate(out["auto"], 0)
    P2["X"] = np.concatenate(out["X"], 0)
    return P2


# ---------------------------------------------------------------- driver
def run(cfg):
    """Same contract as spatial_sim2.run, using the forked _repro."""
    rng = np.random.default_rng(cfg.seed)
    excl = set(); excl_x = set()
    if cfg.sel_locus == "auto": excl.add(cfg.auto_locus)
    if cfg.dm > 0 and cfg.dm_focal == "auto": excl.add(cfg.dm_focal_locus)
    if cfg.dm > 0 and cfg.dm_partner == "auto": excl.add(cfg.dm_partner_locus)
    if cfg.restorers > 0: excl.update(range(AUTOS - cfg.restorers, AUTOS))
    if cfg.sel_locus == "X": excl_x.add(cfg.x_locus)
    if cfg.dm > 0 and cfg.dm_focal == "X": excl_x.add(cfg.dm_focal_locus)
    if cfg.dm > 0 and cfg.dm_partner == "X": excl_x.add(cfg.dm_partner_locus)
    if cfg.xdrive > 0: excl_x.add(DRIVER_XLOCUS)
    excl = tuple(excl); excl_x = tuple(excl_x)
    # Config is inherited from spatial_sim2, so K_early/K_early_gens are ACCEPTED here whether
    # or not this run loop honours them. Honour them, identically to spatial_sim2.run: a larger
    # carrying capacity for the first K_early_gens generations, then a step contraction to K.
    cfg_early = replace(cfg, K=cfg.K_early) if cfg.K_early is not None else cfg
    P = _init(cfg_early, rng)
    # KEYS COME FROM _stats, NOT FROM A LIST HERE (13 Sep 2026). This module imports _stats
    # from spatial_sim2 but used to hardcode the key list, so the moment mt_all / Y_all /
    # auto_all were added to _stats for the two-clock diagnostic, every spatial_sim3 run died
    # with KeyError: 'mt_all' on its first recorded generation -- which is what killed all 15
    # jdcal tasks. Deriving the keys from the function that produces them means the two engines
    # cannot drift apart again.
    _probe = _stats(_init(replace(cfg, gens=1), np.random.default_rng(0)), (), ())
    ts = {k: [] for k in ("gen", *_probe)}
    ext = None
    for ggen in range(cfg.gens):
        cur = cfg_early if (cfg.K_early is not None and ggen < cfg.K_early_gens) else cfg
        _migrate(P, cur, rng)
        nxt = _repro(P, cur, rng)
        if nxt is None:
            break
        P = _reseed_origin(nxt, cur)
        if ggen % cfg.record_every == 0 or ggen == cfg.gens - 1:
            s = _stats(P, excl, excl_x); ts["gen"].append(ggen)
            for k in s:
                ts[k].append(s[k])
        if ext is None and (P["lin"] == 0).sum() == 0:
            ext = ggen
        if ext is not None and cfg.stop_after_ext > 0 and ggen >= ext + cfg.stop_after_ext:
            s = _stats(P, excl, excl_x); ts["gen"].append(ggen)
            for k in s:
                ts[k].append(s[k])
            break
    return {k: np.array(v) for k, v in ts.items()}, ext


if __name__ == "__main__":
    import sys
    quick = "--quick" in sys.argv
    base = dict(grid=(8 if quick else 20), gens=(1200 if quick else 7003),
                sample_gen=(1190 if quick else 6993))
    for name, kw in (("restorers n=3 dom", dict(restorers=3, dm_dom="dominant")),
                     ("xdrive d=1.0", dict(xdrive=1.0)),
                     ("esc eps=0.01", dict(dm=1.0, dm_focal="mt", dm_partner="Y",
                                           esc_eps=0.01))):
        cfg = Config(**base, **kw)
        out, ext = run(cfg)
        print(f"{name:20s} ext={ext}  " +
              "  ".join(f"{k}={out[k][-1]:.3f}" for k in ("mt_mod", "Y_mod", "auto_mod",
                                                          "X_mod")))
```
