## Supplementary Information section 3 code base for "Hypothesis: A modern human range expansion ∼300,000 years ago had a critical role in Neandertal origins": summary_cache.py

```
#!/usr/bin/env python3
"""summary_cache.py -- one pickle of every out_*/*.json "summary" block, built in parallel.

build_supp_workbook.py reads ~24,000 stage-2 and ~60,000 stage-1 JSON files one at a time,
which at a few milliseconds each on the mounted working tree is longer than a single shell
call here is allowed to run. This builds the same information once, eight files at a time,
and is RESUMABLE per directory: each out_* directory is cached to its own pickle, so an
interrupted run loses only the directory it was on. build_supp_workbook consults the cache
through _summary() and falls back to reading the file if a path is missing from it.

    python summary_cache.py            # (re)build whatever is missing or stale
    python summary_cache.py --force    # rebuild everything
"""
import glob
import json
import multiprocessing as mp
import os
import pickle
import sys

HERE = os.path.dirname(os.path.abspath(__file__))
CACHE = os.path.join(HERE, "_summary_cache")


def _one(p):
    try:
        return p, json.load(open(p))["summary"]
    except Exception:
        return p, None


WANT = [f"out_{pre}{st}_{fw}" for pre in ("routes", "dmi", "dmi2", "xdc", "xd2", "tw", "cult")
        for st in ("1", "1hi", "2") for fw in ("deme", "band")]


def build(force=False, budget_s=150):
    """Directory by directory, in a time budget, so repeated calls finish the job. Staleness
    is judged on the file COUNT only: a stat per file on the mounted tree costs more than the
    read it was meant to save."""
    import time
    t0 = time.time()
    os.makedirs(CACHE, exist_ok=True)
    for name in WANT:
        d = os.path.join(HERE, name)
        if not os.path.isdir(d):
            continue
        pk = os.path.join(CACHE, name + ".pkl")
        files = glob.glob(os.path.join(d, "*.json"))
        if not files:
            continue
        if not force and os.path.exists(pk):
            try:
                if len(pickle.load(open(pk, "rb"))) == len(files):
                    continue
            except Exception:
                pass
        if time.time() - t0 > budget_s:
            print(f"  budget reached before {name}; run again to continue")
            return False
        with mp.Pool(8) as pool:
            out = dict(pool.map(_one, files, chunksize=64))
        pickle.dump(out, open(pk, "wb"))
        print(f"  {name:22s} built ({len(files):,})  {time.time() - t0:5.0f}s")
    print("  cache complete")
    return True


def load():
    """{absolute json path: summary dict} for every cached directory."""
    out = {}
    for pk in glob.glob(os.path.join(CACHE, "*.pkl")):
        out.update(pickle.load(open(pk, "rb")))
    return out


if __name__ == "__main__":
    build("--force" in sys.argv)
```
