## Supplementary Information section 3 code base for "Hypothesis: A modern human range expansion ∼300,000 years ago had a critical role in Neandertal origins": tw5_run.py

```
#!/usr/bin/env python3
"""
tw5_run.py -- Route 5 (two waves), re-run so that both frameworks report the same things.
(13 Sep 2026)

WHY THIS EXISTS. Route 5 was the one route whose two frameworks were not measured alike, and
the mismatch was not visible in the figure because each side printed a number.

  1. THE STATISTIC WAS FIXATION, NOT THE SAMPLE OF TEN. twowave_run.py and twowave_band.py both
     report p_joint = mean(mt >= 0.999 AND Y >= 0.999) -- the fraction of replicates in which
     both markers went to fixation. Everywhere else the note now reports P(ten sampled males
     all modern at both markers), per DR: "throughout the note, we should be using not Y
     chromosome, but the 10 males retaining both Y and mtDNA (except in Figure 2 panel a)".
     For the deme cells that happen to fix in every replicate the two agree to within 2%, but
     they are different quantities and only one of them is comparable with the rest of Figure 2.
     Per-replicate mt and Y were not retained, so the ten-male statistic could not be
     recovered from the JSON and the cells had to be re-run.

  2. THE BAND SIDE HAD NO DURATION AT ALL. twowave_run.py records ext_med, the median
     generation of archaic extinction, which is what the note's 100-1,000 generation criterion
     is tested against. twowave_band.py records `swept` per replicate and then throws it away,
     keeping only est = fraction that swept. So the band Route 5 point in Figure 2b was never
     filtered on duration, and figure2c_fountain.load does not filter it either -- Route 5 is
     the one route whose loader branch has no generation test. dur_med below closes that.

  3. THE BAND SIDE WAS AT TEN REPLICATES AND ONE SECOND-WAVE RUNG. ES x WAVE2 = 7 x [0, 1200]
     at 20 reps requested and 10 delivered, against 8 x [0, 500, 1000, 2000] at 100 on the deme
     side. DR: "everywhere in the note where we have different number of reps for bands and
     demes, we should raise the lower one to the higher one." And 1200 is a perverse rung to
     have chosen as the only one: a second wave founded at generation 1,200 cannot finish
     inside a 1,000-generation window no matter how fast it expands, so the single band rung
     was disqualified by its own parameter before any simulation ran.

RESULTS GO TO out_tw5_{fw}, NOT over out_twowave_{fw}. The old files stay for provenance, and
the readers prefer out_tw5_* where it exists, the same convention out_*1hi_* already uses.

    python tw5_run.py deme 0.42 500 100 out_tw5_deme/tw_g0p42_w500.json
    python tw5_run.py band 0.45 500 100 out_tw5_band/tw_e0p45_w500.json
"""
import copy
import json
import os
import sys

import numpy as np

HERE = os.path.dirname(os.path.abspath(__file__))
sys.path.insert(0, HERE)
sys.path.insert(0, os.path.join(os.path.dirname(HERE), "band_model"))

NSAMPLE = 10

# The deme baseline, unchanged from twowave_run.py so the two are comparable cell for cell.
DEME_BASE = dict(grid=12, K=11, m=0.08, r=2.0, gens=7003, sample_gen=6993,
                 record_every=10 ** 9)
GAMMAS = [0.08, 0.12, 0.20, 0.30, 0.42, 0.55, 0.70, 1.00]
ES = [0.08, 0.13, 0.19, 0.26, 0.35, 0.45, 0.56]
WAVE2 = [0, 500, 1000, 2000]          # identical rungs in both frameworks now


def one_deme(g, w, seed):
    import spatial_sim2 as S2
    from spatial_sim2 import Config
    ts, ext = S2.run(Config(**dict(DEME_BASE, gamma=g, wave2_gen=w, seed=seed)))
    return (float(ts["auto_mod"][-1]), float(ts["mt_mod"][-1]), float(ts["Y_mod"][-1]),
            -1 if ext is None else int(ext))


def one_band(e, w, seed):
    from bandsim import Config, Landscape, init_state, step, stats
    cfg = Config(N_target=1750, K=25, e=e, p_move=0.45, r=2.0, beta=0.0,
                 gens=6000, seed=seed, record_every=10 ** 9)
    land = Landscape()
    rng = np.random.default_rng(seed)
    P, pos, b = init_state(cfg, land, rng)
    seed_xy = np.array(cfg.seed_lonlat, float)
    swept = None
    for gen in range(cfg.gens):
        if w and gen == w:
            b[:] = False
            j = int(np.argmin(((pos - seed_xy) ** 2).sum(1)))
            b[j] = True
            mem = P["band"] == j
            if mem.any():
                P["mt"][mem] = 1
                P["Y"][mem & (P["sex"] == 1)] = 1
                P["auto"][mem, :] = 2
                P["X"][mem, :] = np.where(P["sex"][mem] == 1, 1, 2)[:, None]
            cfg = copy.copy(cfg)
            cfg.beta = 1.0                  # second wave is patrilocal
            swept = None
        out = step(P, pos, b, cfg, land, rng)
        if out[0] is None:
            return None
        P, pos, b = out
        if swept is None and b.all():
            swept = gen
            if not w or gen > w:
                break
    st = stats(P, b)
    # Whole-landscape statistics, for the reason twowave_band.py documents: after relabelling,
    # the _mod columns describe only the second wave's own descendants.
    return (float(st["auto"]), float(st["mt"]), float(st["Y"]),
            -1 if swept is None else int(swept))


def main():
    fw, val, w, reps, outrel = (sys.argv[1], float(sys.argv[2]), int(sys.argv[3]),
                                int(sys.argv[4]), sys.argv[5])
    f = one_deme if fw == "deme" else one_band
    res = [r for r in (f(val, w, s) for s in range(reps)) if r]
    path = os.path.join(HERE, outrel)
    os.makedirs(os.path.dirname(path), exist_ok=True)
    if not res:
        json.dump(dict(framework=fw, val=val, wave2_gen=w, reps=0, died=True),
                  open(path, "w"))
        print("  every replicate died")
        return
    a = np.array([r[0] for r in res])
    mt = np.array([r[1] for r in res])
    y = np.array([r[2] for r in res])
    d = np.array([r[3] for r in res])
    fm, fy = mt >= 0.999, y >= 0.999
    # THE NOTE'S STATISTIC. P(all ten sampled males modern at BOTH markers) in one replicate is
    # (mt * Y) ** 10 under independent sampling; the cell's value is the mean over replicates.
    p10 = (mt * y) ** NSAMPLE
    row = dict(framework=fw, val=val, wave2_gen=w, reps=len(res),
               aut=round(100 * float(a.mean()), 3),
               p10=round(float(p10.mean()), 4),
               p10_med=round(float(np.median(p10)), 4),
               mt_mean=round(float(mt.mean()), 4), Y_mean=round(float(y.mean()), 4),
               # kept so the new files can be checked against the old ones cell for cell
               p_mt=round(float(fm.mean()), 3), p_Y=round(float(fy.mean()), 3),
               p_joint=round(float((fm & fy).mean()), 3),
               est=round(float((d >= 0).mean()), 3),
               dur_med=float(np.median(d[d >= 0])) if (d >= 0).any() else None)
    json.dump(row, open(path, "w"))
    print(f"  {fw} val {val:<5g} wave2 {w:<5d} aut {row['aut']:6.2f}%  p10 {row['p10']:.3f}  "
          f"p_joint {row['p_joint']:.2f}  dur_med {row['dur_med']}  est {row['est']:.2f}")


if __name__ == "__main__":
    main()
```
