## Supplementary Information section 3 code base for "Hypothesis: A modern human range expansion ∼300,000 years ago had a critical role in Neandertal origins": tw5_submit.py

```
#!/usr/bin/env python3
"""
tw5_submit.py -- submit the Route 5 re-run. (13 Sep 2026)

60 cells: deme 8 gammas x 4 second-wave rungs, band 7 escape rates x the same 4 rungs, all at
100 replicates. One array task per cell, because the band cells are the expensive ones at about
280 s and the deme cells at about 60 s, so grouping would only make the tail longer.

    python tw5_submit.py          # submits
    python tw5_submit.py dry      # prints the cell list and the cost, submits nothing
"""
import os
import subprocess
import sys

HERE = os.path.dirname(os.path.abspath(__file__))
sys.path.insert(0, HERE)

from tw5_run import ES, GAMMAS, WAVE2       # noqa: E402

REPS = 100
PART, CAP = "short", 400
COST = {"deme": 0.57, "band": 2.80}         # seconds per replicate, measured 12 Sep


def tag(fw, v, w):
    k = "g" if fw == "deme" else "e"
    return f"tw_{k}{v:g}_w{w}".replace(".", "p") + ".json"


def cells():
    out = []
    for w in WAVE2:
        for g in GAMMAS:
            out.append(("deme", g, w))
        for e in ES:
            out.append(("band", e, w))
    return out


def main():
    dry = len(sys.argv) > 1 and sys.argv[1] == "dry"
    tasks, secs = [], 0.0
    for fw, v, w in cells():
        outrel = os.path.join(f"out_tw5_{fw}", tag(fw, v, w))
        if os.path.exists(os.path.join(HERE, outrel)):
            continue
        tasks.append(f"python tw5_run.py {fw} {v:g} {w} {REPS} {outrel}")
        secs += COST[fw] * REPS
    for fw in ("deme", "band"):
        os.makedirs(os.path.join(HERE, f"out_tw5_{fw}"), exist_ok=True)
    print(f"  {len(tasks)} cells, {secs / 3600:.1f} CPU-h, {REPS} replicates each")
    if not tasks:
        print("  nothing to submit")
        return
    if dry:
        for t in tasks[:6]:
            print("   ", t)
        print("    ...")
        return
    with open(os.path.join(HERE, "tw5_params.txt"), "w") as f:
        f.write("\n".join(tasks) + "\n")
    with open(os.path.join(HERE, "tw5_array.sh"), "w") as f:
        f.write(f"""#!/bin/bash
#SBATCH -J tw5
#SBATCH -p {PART}
#SBATCH -c 1
#SBATCH --mem=2G
#SBATCH -t 0-02:59
#SBATCH -a 0-{len(tasks) - 1}%{CAP}
#SBATCH -o logs/tw5_%A_%a.log
cd {HERE}
sed -n "$((SLURM_ARRAY_TASK_ID + 1))p" tw5_params.txt | bash
""")
    os.makedirs(os.path.join(HERE, "logs"), exist_ok=True)
    r = subprocess.run(["sbatch", "tw5_array.sh"], cwd=HERE, capture_output=True, text=True)
    print(f"  {r.stdout.strip()}" if r.returncode == 0 else f"  !! {(r.stderr or '').strip()}")


if __name__ == "__main__":
    main()
```
