## Supplementary Information section 3 code base for "Hypothesis: A modern human range expansion ∼300,000 years ago had a critical role in Neandertal origins": tw6_run.py

```
#!/usr/bin/env python3
"""
tw6_run.py -- Route 4/5 in the band framework at expansion speeds the grid never tested.
(13 Sep 2026)

DR: "The issue you found is that the 1000 generation gap is not enough to complete the two
waves in the band model. Could we do something like, pick a parameter combination that makes
the band expansion go faster? ... it's obviously right. So why not change the parameters here
to make it go faster?"

He is right, and the screened grid could not have answered this, for two reasons I should have
checked before writing the failure into v71.

  1. p_move WAS NEVER SWEPT. Every one of the 1,701 band two-wave combinations ran at
     p_move = 0.45. The whole-band movement rate is the primary determinant of how fast a wave
     of advance crosses the map and it was held fixed while sigma_b, sigma_e, e, r and alpha_d
     were all varied. So "no band combination completes in time" was a statement about one
     movement rate, not about the band model.

  2. THE SECOND-WAVE RUNGS START TOO LATE. The tested rungs are 500, 1,000 and 2,000
     generations. The note's own archaeological anchor is tighter than all of them: IUP moderns
     mixing with Neandertals from ~45 ka and the founder population of later Europeans present
     by ~37 ka is a gap of ~8 kyr, which at 28.6 years per generation is ~280 generations. The
     empirically motivated rung was below the bottom of the tested range.

AND THE SPEED RESPONSE IS STEEP, which is what makes this worth running rather than arguing.
For a SINGLE band wave, across the screened grid:

        sigma_b      n      min    median
          30       3,437    193     1,080
          60       3,678    149       665
         120       3,843     86       357

A single wave at sigma_b = 120 completes in a median of 357 generations. Two of those with a
280-generation gap is ~640, inside the window, before p_move is touched at all.

WHAT MUST NOT BE FUDGED. p_move and sigma_b are not ethnographic quantities -- bandsim's own
comments record that both were tuned upward to make the wave complete inside the window in the
first place ("DR: wave must complete <=1000 gens, ideally ~500"). sigma_e = 80 km IS anchored,
to hunter-gatherer marital distance, and is left alone here. So raising p_move and sigma_b for
the two-wave case is the same operation that set them, applied to a scenario that needs two
crossings instead of one -- but the note has to say that plainly, and it has to report the
SINGLE-wave duration at the same parameters, which must also stay inside the window. A
parameter set that makes two waves fit by making one wave absurdly fast has proved nothing.
That is why wave2_gen = 0 is in the grid below: it is the control.

    python tw6_run.py 0.75 120 300 0.26 100 out_tw6_band/pm0p75_sb120_w300_e0p26.json
      argv: p_move sigma_b wave2_gen e reps outrel
"""
import copy
import json
import os
import sys

import numpy as np

HERE = os.path.dirname(os.path.abspath(__file__))
sys.path.insert(0, HERE)
sys.path.insert(0, os.path.join(os.path.dirname(HERE), "band_model"))

NSAMPLE = 10

P_MOVE = [0.45, 0.60, 0.75, 0.90]
SIGMA_B = [60.0, 120.0, 180.0]
# 0 is the SINGLE-WAVE CONTROL and is not optional: see the docstring.
WAVE2 = [0, 250, 300, 400, 500]
ES = [0.13, 0.19, 0.26, 0.35, 0.45]


def one(p_move, sigma_b, w, e, seed, beta2=1.0):
    from bandsim import Config, Landscape, init_state, step, stats
    cfg = Config(N_target=1750, K=25, e=e, p_move=p_move, sigma_b=sigma_b, sigma_e=80.0,
                 r=2.0, beta=0.0, gens=6000, seed=seed, record_every=10 ** 9)
    land = Landscape()
    rng = np.random.default_rng(seed)
    P, pos, b = init_state(cfg, land, rng)
    seed_xy = np.array(cfg.seed_lonlat, float)
    swept = None
    # TWO CLOCKS, because they are not the same event and the note's criterion needs the
    # second one. `swept` is when the last archaic-LABELLED band disappears, which is what
    # end_gen has always recorded. `arrive` is when the landscape's mtDNA and Y are both at
    # least 99% modern -- the expansion completing in the sense the note cares about. For a
    # single wave the two nearly coincide. After a wave-2 relabelling they do not: every
    # surviving "archaic" band is genetically modern already, so `swept` goes on ticking for
    # thousands of generations after the genetic outcome is settled, and in some replicates
    # never fires at all while mtDNA and Y sit at 100%.
    arrive = None
    for gen in range(cfg.gens):
        if w and gen == w:
            b[:] = False
            j = int(np.argmin(((pos - seed_xy) ** 2).sum(1)))
            b[j] = True
            mem = P["band"] == j
            if mem.any():
                P["mt"][mem] = 1
                P["Y"][mem & (P["sex"] == 1)] = 1
                P["auto"][mem, :] = 2
                P["X"][mem, :] = np.where(P["sex"][mem] == 1, 1, 2)[:, None]
            cfg = copy.copy(cfg)
            # beta2 = 1.0 is the scenario (second wave patrilocal). beta2 = 0.0 is a DIAGNOSTIC
            # that keeps the first wave's rule, to separate "the second wave is slow because it
            # is patrilocal" from "the second wave is slow because the landscape is already
            # full". Those call for different sentences in the note.
            cfg.beta = beta2
            swept = None
        out = step(P, pos, b, cfg, land, rng)
        if out[0] is None:
            return None
        P, pos, b = out
        if gen > w and arrive is None:
            _s = stats(P, b)
            if _s["mt"] >= 0.99 and _s["Y"] >= 0.99:
                arrive = gen
        if swept is None and b.all():
            swept = gen
        # STOP SHORTLY AFTER THE GENETIC CLOCK FIRES. Waiting for the label to sweep costs
        # 2,000-5,000 generations per replicate and, on the evidence, sometimes never happens;
        # 200 generations of margin is enough to confirm the state is stable and cuts the run
        # to about a third of its length.
        if arrive is not None and gen > arrive + 200:
            break
        if swept is not None and gen > w:
            break
    st = stats(P, b)
    return (float(st["auto"]), float(st["mt"]), float(st["Y"]),
            -1 if swept is None else int(swept),
            -1 if arrive is None else int(arrive))


def main():
    p_move, sigma_b, w, e = (float(sys.argv[1]), float(sys.argv[2]),
                             int(sys.argv[3]), float(sys.argv[4]))
    reps, outrel = int(sys.argv[5]), sys.argv[6]
    beta2 = float(sys.argv[7]) if len(sys.argv) > 7 else 1.0
    res = [r for r in (one(p_move, sigma_b, w, e, s, beta2) for s in range(reps)) if r]
    path = os.path.join(HERE, outrel)
    os.makedirs(os.path.dirname(path), exist_ok=True)
    base = dict(framework="band", p_move=p_move, sigma_b=sigma_b, sigma_e=80.0,
                wave2_gen=w, e=e, K=25, N=1750, r=2.0, beta2=beta2)
    if not res:
        json.dump(dict(base, reps=0, died=True), open(path, "w"))
        print("  every replicate died")
        return
    a = np.array([r[0] for r in res])
    mt = np.array([r[1] for r in res])
    y = np.array([r[2] for r in res])
    d = np.array([r[3] for r in res])
    ar = np.array([r[4] for r in res])
    p10 = (mt * y) ** NSAMPLE
    row = dict(base, reps=len(res),
               aut=round(100 * float(a.mean()), 3),
               p10=round(float(p10.mean()), 4),
               mt_mean=round(float(mt.mean()), 4), Y_mean=round(float(y.mean()), 4),
               est=round(float((d >= 0).mean()), 3),
               # label-absorption clock, what end_gen has always been
               dur_med=float(np.median(d[d >= 0])) if (d >= 0).any() else None,
               dur_min=float(d[d >= 0].min()) if (d >= 0).any() else None,
               swept_frac=round(float((d >= 0).mean()), 3),
               # genetic clock, the one the duration criterion should be applied to
               arr_med=float(np.median(ar[ar >= 0])) if (ar >= 0).any() else None,
               arr_min=float(ar[ar >= 0].min()) if (ar >= 0).any() else None,
               arr_frac=round(float((ar >= 0).mean()), 3))
    json.dump(row, open(path, "w"))
    print(f"  p_move {p_move:<5g} sigma_b {sigma_b:<6g} wave2 {w:<5d} e {e:<5g}  "
          f"aut {row['aut']:6.2f}%  p10 {row['p10']:.3f}  "
          f"arrive_med {row['arr_med']}  label_med {row['dur_med']}  "
          f"arr_frac {row['arr_frac']:.2f}")


if __name__ == "__main__":
    main()
```
