## Supplementary Information section 3 code base for "Hypothesis: A modern human range expansion ∼300,000 years ago had a critical role in Neandertal origins": tw6_submit.py

```
#!/usr/bin/env python3
"""
tw6_submit.py -- the band two-wave speed search. (13 Sep 2026)

DR: "pick a parameter combination that makes the band expansion go faster ... it's obviously
right." He is right about the biology and the runs below test it properly. See tw6_run.py for
why the screened grid could not answer this: p_move was never swept, the earliest second-wave
rung was later than the archaeology implies, and the duration being tested was the wrong clock.

120 cells x 50 replicates. p_move 0.45 is dropped -- it is the value the 1,701 screened
combinations already ran at -- and sigma_b 60 is kept as the slow anchor. wave2_gen 0 is the
single-wave control at each speed and is what stops a parameter set from "succeeding" by
making one wave implausibly fast.

    python tw6_submit.py [dry]
"""
import json
import os
import subprocess
import sys

HERE = os.path.dirname(os.path.abspath(__file__))
REPS = 50
PART, CAP = "short", 400
P_MOVE = [0.60, 0.75, 0.90]
SIGMA_B = [60.0, 120.0, 180.0]
WAVE2 = [0, 200, 250, 300]
ES = [0.13, 0.19, 0.26, 0.35, 0.45]
SEC_PER_REP = 45.0          # measured locally after the early-stop change


def tag(pm, sb, w, e):
    return f"pm{pm:g}_sb{sb:g}_w{w}_e{e:g}".replace(".", "p") + ".json"


def main():
    dry = len(sys.argv) > 1 and sys.argv[1] == "dry"
    # --reps RAISES THE COUNT AND THEREFORE HAS TO DEFEAT THE FILENAME SKIP. DR: "put in runs
    # that replace the lower count with a higher one so we can say the same number of reps for
    # each." These band cells are the Route 4 arm of Figure 2b and Figure S3.3, drawn beside a
    # deme arm scored at 200 replicates, and they were run at 50.
    #
    # tw6_run writes ONE JSON per cell containing every replicate, with seeds 0..reps-1, so
    # there is no seed offset to append at the way bandsi_submit has: a 200-replicate cell
    # simply redraws the 50 and adds 150. The existing file therefore has to be rerun rather
    # than skipped, and a cell is skipped only when the file on disk ALREADY holds at least
    # the requested count. That makes the flag idempotent -- running it twice at 200 submits
    # nothing the second time -- while a plain filename test would either refuse to top up at
    # all or redo the whole sweep on every invocation.
    reps = int(sys.argv[sys.argv.index("--reps") + 1]) if "--reps" in sys.argv else REPS
    os.makedirs(os.path.join(HERE, "out_tw6_band"), exist_ok=True)
    tasks, redo = [], 0
    for pm in P_MOVE:
        for sb in SIGMA_B:
            for w in WAVE2:
                for e in ES:
                    rel = os.path.join("out_tw6_band", tag(pm, sb, w, e))
                    full = os.path.join(HERE, rel)
                    if os.path.exists(full):
                        try:
                            have = int(json.load(open(full)).get("reps") or 0)
                        except Exception:
                            have = 0
                        if have >= reps:
                            continue
                        redo += 1
                    tasks.append(f"python tw6_run.py {pm:g} {sb:g} {w} {e:g} {reps} {rel}")
    print(f"  {len(tasks)} cells, {len(tasks)*reps*SEC_PER_REP/3600:.0f} CPU-h, "
          f"{reps} replicates each")
    if redo:
        print(f"  {redo} of them REPLACE a file with fewer than {reps} replicates "
              f"(the earlier draws are discarded, not pooled)")
    if not tasks or dry:
        for t in tasks[:5]:
            print("   ", t)
        return
    open(os.path.join(HERE, "tw6_params.txt"), "w").write("\n".join(tasks) + "\n")
    open(os.path.join(HERE, "tw6_array.sh"), "w").write(f"""#!/bin/bash
#SBATCH -J tw6
#SBATCH -p {PART}
#SBATCH -c 1
#SBATCH --mem=2G
#SBATCH -t 0-02:59
#SBATCH -a 0-{len(tasks) - 1}%{CAP}
#SBATCH -o logs/tw6_%A_%a.log
cd {HERE}
sed -n "$((SLURM_ARRAY_TASK_ID + 1))p" tw6_params.txt | bash
""")
    os.makedirs(os.path.join(HERE, "logs"), exist_ok=True)
    r = subprocess.run(["sbatch", "tw6_array.sh"], cwd=HERE, capture_output=True, text=True)
    print(f"  {r.stdout.strip()}" if r.returncode == 0 else f"  !! {(r.stderr or '').strip()}")


if __name__ == "__main__":
    main()
```
