## Supplementary Information section 3 code base for "Hypothesis: A modern human range expansion ∼300,000 years ago had a critical role in Neandertal origins": twowave_band.py

```
#!/usr/bin/env python3
"""
twowave_band.py -- Route 5 in the roving-band framework. (DR, 10 Sep 2026)

The relabelling is done here rather than inside bandsim, because it needs nothing the engine
does not already expose: `step` takes the identity array and the config afresh every
generation, so a driver can flip both between calls. Keeping it out of the engine also means
the band result cannot be contaminated by a change to the shared code path.

At wave2_gen:
  * every band is relabelled archaic (b_ident[:] = False), keeping every allele, so the second
    wave expands into an occupied landscape exactly as the first did;
  * the band nearest the Bosphorus seed point is relabelled modern and its members given modern
    alleles at every locus;
  * beta flips to 1.0, so the second wave is patrilocal -- women disperse, men stay, and the
    incoming modern Y is retained.

    python twowave_band.py [--reps 20] [--budget 100]
"""
import copy
import glob
import json
import os
import sys
import time
from concurrent.futures import ProcessPoolExecutor

import numpy as np

HERE = os.path.dirname(os.path.abspath(__file__))
sys.path.insert(0, HERE)
sys.path.insert(0, os.path.join(os.path.dirname(HERE), "band_model"))

OUT = os.path.join(HERE, "out_twowave_band")
ES = [0.08, 0.13, 0.19, 0.26, 0.35, 0.45, 0.56]
WAVE2 = [0, 1200]
NSAMPLE = 10


def flag(name, cast, default):
    return cast(sys.argv[sys.argv.index(name) + 1]) if name in sys.argv else default


def one(args):
    e, w, seed = args
    from bandsim import Config, Landscape, init_state, step, stats
    cfg = Config(N_target=1750, K=25, e=e, p_move=0.45, r=2.0, beta=0.0,
                 gens=6000, seed=seed, record_every=10 ** 9)
    land = Landscape()
    rng = np.random.default_rng(seed)
    P, pos, b = init_state(cfg, land, rng)
    seed_xy = np.array(cfg.seed_lonlat, float)
    swept = None
    for g in range(cfg.gens):
        if w and g == w:
            b[:] = False
            # nearest band to the Bosphorus becomes the second wave's founder
            j = int(np.argmin(((pos - seed_xy) ** 2).sum(1)))
            b[j] = True
            mem = P["band"] == j
            if mem.any():
                P["mt"][mem] = 1
                P["Y"][mem & (P["sex"] == 1)] = 1
                P["auto"][mem, :] = 2
                P["X"][mem, :] = np.where(P["sex"][mem] == 1, 1, 2)[:, None]
            cfg = copy.copy(cfg)
            cfg.beta = 1.0                      # second wave is patrilocal
            swept = None
        out = step(P, pos, b, cfg, land, rng)
        if out[0] is None:
            return None
        P, pos, b = out
        if swept is None and b.all():
            swept = g
            if not w or g > w:
                break
    st = stats(P, b)
    # WHOLE-LANDSCAPE statistics, not the _mod ones. mt_mod/Y_mod/auto_mod are computed within
    # modern-IDENTITY bands only. For a single wave that is harmless, because the archaic
    # identity goes extinct and the two coincide. For the two-wave scenario it is wrong: after
    # the landscape is relabelled, "modern identity" means only the second wave's descendants,
    # so the _mod statistics describe the incoming group rather than the population that would
    # actually be sampled -- and they return nan outright if that band dies. The observable
    # here is ancestry across everyone alive at the end.
    return (float(st["auto"]), float(st["mt"]), float(st["Y"]),
            -1 if swept is None else int(swept))


def main():
    reps = flag("--reps", int, 20)
    budget = flag("--budget", int, 100)
    os.makedirs(OUT, exist_ok=True)
    todo = [(e, w) for w in WAVE2 for e in ES
            if not os.path.exists(os.path.join(OUT, f"tw_e{e:g}_w{w}".replace(".", "p")
                                               + ".json"))]
    print(f"  {len(ES)*len(WAVE2)} cells, {len(glob.glob(os.path.join(OUT,'*.json')))} done, "
          f"{len(todo)} to go, {reps} reps each")
    t0 = time.time()
    with ProcessPoolExecutor(4) as ex:
        for (e, w) in todo:
            res = [r for r in ex.map(one, [(e, w, s) for s in range(reps)]) if r]
            if not res:
                print(f"    e {e:<5g} wave2 {w:<5d}  every replicate died")
                continue
            a = np.array([r[0] for r in res]); mt = np.array([r[1] for r in res])
            y = np.array([r[2] for r in res]); sw = np.array([r[3] for r in res])
            fm, fy = mt >= 0.999, y >= 0.999
            row = dict(e=e, wave2_gen=w, reps=len(res),
                       aut=round(100 * float(a.mean()), 3),
                       p_mt=round(float(fm.mean()), 3), p_Y=round(float(fy.mean()), 3),
                       p_joint=round(float((fm & fy).mean()), 3),
                       est=round(float((sw >= 0).mean()), 3))
            with open(os.path.join(OUT, f"tw_e{e:g}_w{w}".replace(".", "p") + ".json"), "w") as f:
                json.dump(row, f)
            print(f"    e {e:<5g} wave2 {w:<5d} aut {row['aut']:6.2f}%  p_mt {row['p_mt']:.2f}  "
                  f"p_Y {row['p_Y']:.2f}  p_joint {row['p_joint']:.2f}  est {row['est']:.2f}")
            if time.time() - t0 > budget:
                print("  budget reached; run again")
                break


if __name__ == "__main__":
    main()
```
