## Supplementary Information section 3 code base for "Hypothesis: A modern human range expansion ∼300,000 years ago had a critical role in Neandertal origins": twowave_run.py

```
#!/usr/bin/env python3
"""
twowave_run.py -- Route 5 production runs, for Figure 2c. (DR, 10 Sep 2026)

DR settled the semantics: the second wave is "a fresh migration pulse", not the same population
changing residence rule. The landscape is relabelled at wave2_gen so every survivor counts as
resident, and a new modern group is founded at the origin under the opposite rule; only group
identity is reset, so the residents keep the alleles the first wave left them.

Resumable: one JSON per cell, skipped if present, because a shell call here is capped at about
two minutes and background jobs do not survive between calls.

    python twowave_run.py [--reps 100] [--budget 100]
"""
import glob
import json
import os
import sys
import time
from concurrent.futures import ProcessPoolExecutor

import numpy as np

HERE = os.path.dirname(os.path.abspath(__file__))
sys.path.insert(0, HERE)

OUT = os.path.join(HERE, "out_twowave_deme")
BASE = dict(grid=12, K=11, m=0.08, r=2.0, gens=7003, sample_gen=6993, record_every=10 ** 9)
GAMMAS = [0.08, 0.12, 0.20, 0.30, 0.42, 0.55, 0.70, 1.00]
# 0 is the single-wave control, which must reproduce p_Y at the martingale (~5%) or the
# comparison in the figure is meaningless.
WAVE2 = [0, 500, 1000, 2000]


def flag(name, cast, default):
    return cast(sys.argv[sys.argv.index(name) + 1]) if name in sys.argv else default


def one(args):
    g, w, seed = args
    import spatial_sim2 as S2
    from spatial_sim2 import Config
    ts, ext = S2.run(Config(**dict(BASE, gamma=g, wave2_gen=w, seed=seed)))
    return (float(ts["auto_mod"][-1]), float(ts["mt_mod"][-1]), float(ts["Y_mod"][-1]),
            -1 if ext is None else int(ext))


def main():
    reps = flag("--reps", int, 100)
    budget = flag("--budget", int, 100)
    os.makedirs(OUT, exist_ok=True)
    todo = [(g, w) for w in WAVE2 for g in GAMMAS
            if not os.path.exists(os.path.join(OUT, f"tw_g{g:g}_w{w}".replace(".", "p") + ".json"))]
    have = len(glob.glob(os.path.join(OUT, "*.json")))
    print(f"  {len(GAMMAS)*len(WAVE2)} cells, {have} done, {len(todo)} to go, {reps} reps each")
    t0 = time.time()
    with ProcessPoolExecutor(4) as ex:
        for (g, w) in todo:
            res = list(ex.map(one, [(g, w, s) for s in range(reps)]))
            a = np.array([r[0] for r in res]); mt = np.array([r[1] for r in res])
            y = np.array([r[2] for r in res]); e = np.array([r[3] for r in res])
            fx_mt, fx_y = mt >= 0.999, y >= 0.999
            row = dict(gamma=g, wave2_gen=w, reps=reps,
                       aut=round(100 * float(a.mean()), 3),
                       p_mt=round(float(fx_mt.mean()), 3),
                       p_Y=round(float(fx_y.mean()), 3),
                       p_joint=round(float((fx_mt & fx_y).mean()), 3),
                       est=round(float((e >= 0).mean()), 3),
                       ext_med=float(np.median(e[e >= 0])) if (e >= 0).any() else None)
            fn = os.path.join(OUT, f"tw_g{g:g}_w{w}".replace(".", "p") + ".json")
            with open(fn, "w") as f:
                json.dump(row, f)
            print(f"    gamma {g:<5g} wave2 {w:<5d} aut {row['aut']:6.2f}%  "
                  f"p_mt {row['p_mt']:.2f}  p_Y {row['p_Y']:.2f}  p_joint {row['p_joint']:.2f}")
            if time.time() - t0 > budget:
                print("  budget reached; run again")
                break


if __name__ == "__main__":
    main()
```
