## Supplementary Information section 3 code base for "Hypothesis: A modern human range expansion ∼300,000 years ago had a critical role in Neandertal origins": twpulse_submit.py

```
#!/usr/bin/env python3
"""
twpulse_submit.py -- Route 4's comparison curves: TWO PULSES at a stable boundary. (14 Sep 2026)

DR: "write me sweeps to generate the blue, black and yellow curves for Route 4."

Route 4 -- the two-wave scenario -- was the one panel of Figure 2b with no comparison curves at
all, because the two stable-boundary scenarios had no second event to give them. The three are
in three different states, and only two of them need running:

  YELLOW (nexp, unbiased range expansion)   NO RUN NEEDED. The `demog` set already carries
        twowave at beta = 0.5: 2,096 scored deme cells at wave2 = 500 and 326 band cells
        across 500/1000/2000. The curve is computed from those.

  BLUE (fsb, all-female flow)               RUN, though the answer is close to foregone. Two
        all-female pulses deliver no modern male either time, so the modern Y never enters and
        p_joint should be identically zero. It is cheap enough to measure rather than assert,
        and a measured zero beside a measured non-zero is worth more in a figure than an
        asterisk -- but if it comes back non-zero, something is wrong with the engine and I
        would want to know that before the note goes out.

  BLACK (nsb, non-sex-biased flow)          RUN, AND THIS IS THE INFORMATIVE ONE. Its two
        pulses are all-female then ALL-MALE: the stable-boundary reading of "matrilocal then
        patrilocal". Two UNBIASED pulses would deliver more of the same and could not produce
        the asymmetry Route 4 turns on, so they would answer a question nobody asked. Posed
        this way the point is substantive: if a pair of sex-biased pulses at a stationary
        boundary retains both markers, then Route 4 does not depend on the range expansion,
        and the note should say so.

f IS HALVED, ANALYTICALLY. Two pulses of fraction f leave 2f - f^2 modern ancestry, so the 5%
anchor needs f = 1 - sqrt(0.95) = 0.02532 in each. Measured across the ladder that lands at
5.1-5.9% autosomal -- inside the 3-7% window at every rung -- so unlike the expansions, which
need gamma retuned per rung because ancestry accumulates at a moving front, no probe grid is
required here. joint_run applies it; this script only chooses the rungs.

    python twpulse_submit.py              # 6 cells x 2,000 replicates, well under 1 CPU-hour
    python twpulse_submit.py --reps 400
    python twpulse_submit.py --no-go
"""
import os
import subprocess
import sys

HERE = os.path.dirname(os.path.abspath(__file__))
sys.path.insert(0, HERE)

from anchors import PANM_CENSUS as SB_N      # 1312, the census matching the Peyregne anchor

# The wave2 ladder the expansions use in routes_build's demog set, so the comparison points
# sit at the same second-wave timings as the cloud they are being compared with.
WAVE2 = [500, 1000, 2000]
SCEN = ["fsb", "nsb"]
REPS = 2000
OUT = "out_twpulse"
PART, CAP = "short", 200


def main():
    reps = int(sys.argv[sys.argv.index("--reps") + 1]) if "--reps" in sys.argv else REPS
    os.makedirs(os.path.join(HERE, OUT), exist_ok=True)
    os.makedirs(os.path.join(HERE, "logs"), exist_ok=True)

    tasks = []
    for sc in SCEN:
        for g in WAVE2:
            out = os.path.join(OUT, f"{sc}_tp{g}.tsv")
            full = os.path.join(HERE, out)
            # Idempotent on the replicate count, like tw6_submit and nsbdrive_submit.
            if os.path.exists(full) and max(sum(1 for _ in open(full)) - 1, 0) >= reps:
                continue
            tasks.append(f"python joint_run.py {sc} twopulse {g} {SB_N} {reps} 1000 {out}")
    if not tasks:
        print(f"  nothing to do -- every cell already has {reps:,} replicates")
        return
    print(f"  {len(tasks)} cells ({len(SCEN)} scenarios x {len(WAVE2)} second-wave timings) "
          f"at {reps:,} replicates, N = {SB_N}")

    pf, sf = "twpulse_params.txt", "twpulse_array.sh"
    open(os.path.join(HERE, pf), "w").write("\n".join(tasks) + "\n")
    open(os.path.join(HERE, sf), "w").write(f"""#!/bin/bash
#SBATCH -J twpulse
#SBATCH -p {PART}
#SBATCH -t 0-11:59
#SBATCH -c 1
#SBATCH --mem=4G
#SBATCH -a 0-{len(tasks) - 1}%{CAP}
#SBATCH -o logs/twpulse_%A_%a.log
CMD=$(sed -n "$((SLURM_ARRAY_TASK_ID + 1))p" {pf})
echo "$CMD"
eval "$CMD"
""")
    if "--no-go" in sys.argv:
        print(f"  NOT submitted (--no-go).  sbatch {sf}")
        return
    r = subprocess.run(["sbatch", sf], cwd=HERE, capture_output=True, text=True)
    print(f"  SUBMITTED: {r.stdout.strip()}" if r.returncode == 0
          else f"  !! sbatch FAILED: {(r.stderr or '').strip()}")


if __name__ == "__main__":
    main()
```
