## Supplementary Information section 3 code base for "Hypothesis: A modern human range expansion ∼300,000 years ago had a critical role in Neandertal origins": xdrv2_collect.py

```
#!/usr/bin/env python3
"""
xdrv2_collect.py -- pool out_xdrv2/ and report the three X-drive follow-ups.

  T: the drive-strength floor (main grid worked at every d >= 0.5).
  R: heritable driver-resistance (allele 2, reported as Yres) -- the stress test.
  C: no-suppressor control -- should FAIL to fix the modern Y if the mechanism's success
     comes from the coevolved asymmetry rather than from drive per se.

  python xdrv2_collect.py        # -> xdrv2_summary.csv, xdrv2_panel.png
"""
import glob
import numpy as np
import matplotlib
matplotlib.use("Agg")
import matplotlib.pyplot as plt
from xdrv2_run import conditions, BLOCK_T, BLOCK_R, BLOCK_C
from xdrv2_submit import tag

COL = dict(seed=0, mt=1, Y=2, Yres=3, auto=4, X=5, X0=6, ext=7)


def pool(block, d, par):
    rows = []
    for f in glob.glob(f"out_xdrv2/out_{tag(block, d, par)}_s*.txt"):
        for line in open(f).read().strip().split("\n")[1:]:
            p = line.split("\t")
            if len(p) >= 8 and "nan" not in line:
                rows.append([float(x) for x in p])
    return np.array(rows) if rows else None


def main():
    recs = []
    with open("xdrv2_summary.csv", "w") as out:
        out.write("block,d,par,nreps,mt,Y,Yres,auto,X_bg,X0,P_Y_fixed,P_all10,P_ext\n")
        print(f"{'blk':>3s} {'d':>5s} {'par':>7s} {'reps':>5s} | {'Y%':>6s} {'Yres%':>6s} "
              f"{'auto%':>6s} {'Xbg%':>6s} {'X0%':>6s} | {'P(Yfix)':>8s} {'P(all10)':>8s} "
              f"{'ext%':>6s}")
        for block, d, par in conditions():
            a = pool(block, d, par)
            if a is None:
                continue
            comp = 100 * (a[:, COL["ext"]] >= 0).mean()
            a = a[a[:, COL["ext"]] >= 0]       # conditional on completion (Table S2.1)
            if not len(a):
                print(f"{block:>3s} {d:5g} {par:7g}     0 |  (no completed replicates; "
                      f"completion {comp:.1f}%)")
                recs.append(dict(block=block, d=d, par=par, reps=0, Y=np.nan, yres=np.nan,
                                 auto=np.nan, X=np.nan, X0=np.nan, fix=np.nan, p10=np.nan,
                                 ext=comp))
                continue
            Y = a[:, COL["Y"]]
            v = dict(block=block, d=d, par=par, reps=len(a),
                     mt=100 * a[:, COL["mt"]].mean(), Y=100 * Y.mean(),
                     yres=100 * a[:, COL["Yres"]].mean(),
                     auto=100 * a[:, COL["auto"]].mean(), X=100 * a[:, COL["X"]].mean(),
                     X0=100 * a[:, COL["X0"]].mean(),
                     fix=100 * (Y >= 0.999).mean(), p10=100 * (Y ** 10).mean(), ext=comp)
            recs.append(v)
            out.write(f"{block},{d:g},{par:g},{len(a)},{v['mt']:.2f},{v['Y']:.2f},"
                      f"{v['yres']:.2f},{v['auto']:.2f},{v['X']:.2f},{v['X0']:.2f},"
                      f"{v['fix']:.1f},{v['p10']:.1f},{v['ext']:.1f}\n")
            print(f"{block:>3s} {d:5g} {par:7g} {len(a):5d} | {v['Y']:6.1f} {v['yres']:6.1f} "
                  f"{v['auto']:6.2f} {v['X']:6.2f} {v['X0']:6.1f} | {v['fix']:8.1f} "
                  f"{v['p10']:8.1f} {v['ext']:6.1f}")
    if not recs:
        print("no data yet"); return
    print("\nreference: the main grid gives P(all10) 99-100% at every d in 0.5-1.0, "
          "auto ~4.6-5.1%, X ~9%")

    fig, axes = plt.subplots(1, 2, figsize=(11, 4.4))
    tsel = [r for r in recs if r["block"] == "T" and r["reps"]]
    axes[0].plot([r["d"] for r in tsel], [r["p10"] for r in tsel], "-o", color="#C44E52",
                 label="threshold block")
    axes[0].axhline(99.3, ls=":", c="0.5", lw=1)
    axes[0].text(0.06, 95, "main grid at d ≥ 0.5", fontsize=8, color="0.35")
    axes[0].set_xlabel("drive strength d"); axes[0].set_ylabel("P(all 10 Y modern) (%)")
    axes[0].set_ylim(-3, 103)
    rsel = [r for r in recs if r["block"] == "R" and r["d"] == 0.95 and r["reps"]]
    axes[1].plot([r["par"] for r in rsel], [r["p10"] for r in rsel], "-o", color="#4C72B0",
                 label="P(all 10 Y modern)")
    axes[1].plot([r["par"] for r in rsel], [r["yres"] for r in rsel], ":^", color="0.4",
                 label="resistant archaic Y (%)")
    axes[1].set_xscale("log")
    axes[1].set_xlabel("resistance probability per suppressed transmission, eps_r  (d = 0.95)")
    axes[1].set_ylim(-3, 103)
    for ax in axes:
        ax.grid(alpha=.3); ax.legend(fontsize=8, frameon=False)
    fig.suptitle("X-drive follow-ups: threshold and heritable resistance", fontsize=11)
    fig.tight_layout()
    fig.savefig("xdrv2_panel.png", dpi=160)
    print("saved xdrv2_summary.csv and xdrv2_panel.png")


if __name__ == "__main__":
    main()
```
