## Supplementary Information section 3 code base for "Hypothesis: A modern human range expansion ∼300,000 years ago had a critical role in Neandertal origins": xdrv2_run.py

```
#!/usr/bin/env python3
"""
xdrv2_run.py -- three follow-ups to the X-drive grid (xdrv_run.py), which reproduced the full
empirical pattern in every cell (Y ~100%, autosomes ~5%, X ~9% > autosomes, mtDNA 100%,
driver polymorphic) at every drive strength tested down to d = 0.5.

Block T, THRESHOLD: d in {0.05, 0.1, 0.2, 0.3, 0.4}, cost 0. The mechanism worked at every d
    tested so far, so the informative number is the floor. A floor near d ~ 0.1-0.2 would put
    the requirement inside the transmission-distortion range documented for real, currently
    segregating systems (mouse Slx/Sly copy-number mismatches shift litter sex ratios by tens
    of percent), making the anthropological ask minimal.

Block R, HERITABLE RESISTANCE: the one result that could still undermine the mechanism. Each
    suppressed Y-transmission escapes with probability eps_r, converting to a driver-resistant
    archaic Y that is immune forever after -- a resistance mutation, the standard fate of drive
    systems, and the drive analogue of the esc_eps stress test on the sealed mtDNA x Y DMI.
    Run at d = 0.95 across six orders of eps_r, plus two cells at d = 0.5.

Block C, NO-SUPPRESSOR CONTROL: the driver kills Y-bearing sperm regardless of which Y the
    male carries, removing the coevolved immunity of the modern Y. If the mechanism's success
    came from drive per se this should still work; if -- as the design intends -- it comes from
    the ASYMMETRY (only the archaic Y is unprotected), this control should fail to fix the
    modern Y and should distort the sex ratio wherever the driver is common (Hamilton 1967).

Baseline as xdrv_run: grid=20, K=20, r=2, m=0.10, gamma=0.12, matrilineal, no sterility DMI,
sampled at 6,993, no early stopping. X locus 0 = driver (excluded from the X aggregate).

Row layout:  seed  mt  Y  Yres  auto  X  X0  extgen   (Yres = resistant archaic Y, allele 2)
Usage:       python xdrv2_run.py <block> <d> <par> <reps> <seed0> <outfile>
             block T: par ignored; block R: par = eps_r; block C: par ignored
"""
import os
import sys
import numpy as np
from dataclasses import replace
import spatial_sim2 as S2
from spatial_sim3 import Config, _repro, DRIVER_XLOCUS

SAMPLE_GEN = 6993

BLOCK_T = [("T", d, 0.0) for d in (0.05, 0.1, 0.2, 0.3, 0.4)]
BLOCK_R = ([("R", 0.95, e) for e in (1e-4, 3e-4, 1e-3, 3e-3, 1e-2, 3e-2)]
           + [("R", 0.5, 1e-3), ("R", 0.5, 1e-2)])
BLOCK_C = [("C", 0.5, 0.0), ("C", 1.0, 0.0)]

# --- expansion geometry, settable from the command line ---------------------------------
# The grid-20 / K-20 baseline has a post-wave census of ~8000, five times the post-mixture size
# the rest of the analysis assumes (Muller's Ratchet v32: NN-uniparental = 725-1000, diploid
# 1500-2000). Since the modern mtDNA's advantage is a ~0.3% coefficient whose potency scales
# with 2*Ne*s, that mismatch inflated every spatial result. r and m are NOT externally
# constrained (DR, 3 Sep 2026: "pulled from my head"), so they are exposed here too and their
# sensitivity is the honest thing to report rather than a fixed baseline.
#
# A flag only OVERRIDES a parameter this script holds fixed; where the script sweeps that
# parameter, the swept value still wins, so sensitivity panels keep their full range.
# K_early/K_early_gens are always ADDED, since no BASE dict carries them.
def _geom(kw):
    import sys as _s
    for flag, key, cast in (("--grid", "grid", int), ("--K", "K", int),
                            ("--m", "m", float), ("--r", "r", float)):
        if flag in _s.argv and key in kw:
            kw[key] = cast(_s.argv[_s.argv.index(flag) + 1])
    for flag, key in (("--K-early", "K_early"), ("--K-early-gens", "K_early_gens")):
        if flag in _s.argv:
            kw[key] = int(_s.argv[_s.argv.index(flag) + 1])
    if kw.get("K", 20) < 7:
        raise SystemExit("K < 7: deme reproductive failure collapses the archaic population")
    return kw


BASE = dict(grid=20, K=20, r=2.0, m=0.10, gamma=0.12, lineage="matrilineal",
            s=0.0, sel_locus="none", dm=0.0, xdrive_cost=0.0,
            gens=SAMPLE_GEN + 10, sample_gen=SAMPLE_GEN, stop_after_ext=0,
            record_every=100000)


def conditions():
    return BLOCK_T + BLOCK_R + BLOCK_C


def run_one(cfg):
    rng = np.random.default_rng(cfg.seed)
    _cap_early = S2.cfg_early(cfg)   # K_early phase, if any
    P = S2._init(_cap_early, rng)
    ext = None
    for g in range(cfg.gens):
        _c = S2.cap_at(cfg, _cap_early, g)
        S2._migrate(P, _c, rng)
        nxt = _repro(P, _c, rng)
        if nxt is None:
            break
        P = S2._reseed_origin(nxt, _c)
        if ext is None and (P["lin"] == 0).sum() == 0:
            ext = g
        if g == cfg.sample_gen:
            break
    st = S2._stats(P, exclude=(), exclude_x=(DRIVER_XLOCUS,))
    mod = P["lin"] == 1
    mmale = mod & (P["sex"] == 1)
    yres = float((P["Y"][mmale] == 2).mean()) if mmale.any() else np.nan
    if mod.any():
        Xm = P["X"][mod]; sxm = P["sex"][mod]
        maxd = np.where(sxm == 0, 2.0, 1.0)
        x0 = float((Xm[:, DRIVER_XLOCUS] / maxd).mean())
    else:
        x0 = np.nan
    return st, yres, x0, ext


def main():
    block = sys.argv[1]; d = float(sys.argv[2]); par = float(sys.argv[3])
    reps, seed0, outfile = int(sys.argv[4]), int(sys.argv[5]), sys.argv[6]
    kw = dict(BASE, xdrive=d)
    if block == "R":
        kw["xdrv_res"] = par
    elif block == "C":
        kw["xdrv_all"] = True
    elif block != "T":
        raise SystemExit(f"unknown block {block}")
    cfg0 = Config(**_geom(kw))
    os.makedirs(os.path.dirname(outfile) or ".", exist_ok=True)
    with open(outfile, "w") as f:
        f.write("seed\tmt\tY\tYres\tauto\tX\tX0\textgen\n")
        for s in range(seed0, seed0 + reps):
            st, yres, x0, ext = run_one(replace(cfg0, seed=s))
            eg = int(ext) if ext is not None else -1
            f.write(f"{s}\t{st['mt_mod']:.6g}\t{st['Y_mod']:.6g}\t{yres:.6g}\t"
                    f"{st['auto_mod']:.6g}\t{st['X_mod']:.6g}\t{x0:.6g}\t{eg}\n")
    print("wrote", outfile)


if __name__ == "__main__":
    main()
```
