## Supplementary Information section 3 code base for "Hypothesis: A modern human range expansion ∼300,000 years ago had a critical role in Neandertal origins": xdrv2_submit.py

```
#!/usr/bin/env python3
"""xdrv2_submit.py -- SLURM array for the X-drive follow-ups (xdrv2_run.py).
Usage:  python xdrv2_submit.py [reps] [batch] [--resume]     # defaults 1000, 25
then:   sbatch xdrv2_array.sh
"""
import os
import sys
from xdrv2_run import conditions


def tag(block, d, par):
    t = f"{block}_d{d:g}"
    if par > 0:
        t += f"_e{par:g}"
    return t.replace(".", "p").replace("-", "m")


GEOM_FLAGS = ("--grid", "--K", "--m", "--r", "--K-early", "--K-early-gens")


def _strip_flags(av):
    """Positional parsing must not be shifted by the geometry flags, so drop each flag AND its
    value before the positional args are read."""
    out, skip = [], False
    for a in av:
        if skip:
            skip = False; continue
        if a.startswith("--"):
            skip = a in GEOM_FLAGS or a == "--rna"; continue
        out.append(a)
    return out


def _geom_str():
    """Echo the geometry flags onto every task line. Without this the workers would silently run
    the grid-20 / K-20 default while the submit script had been told otherwise -- which produces
    plausible-looking numbers that are simply wrong."""
    import sys as _s
    return "".join(f" {f} {_s.argv[_s.argv.index(f) + 1]}"
                   for f in GEOM_FLAGS if f in _s.argv)


def main():
    GEOM = _geom_str()
    nums = [int(a) for a in sys.argv[1:] if a.isdigit()]
    total = nums[0] if len(nums) >= 1 else 1000
    batch = nums[1] if len(nums) >= 2 else 25
    resume = "--resume" in sys.argv
    os.makedirs("out_xdrv2", exist_ok=True); os.makedirs("logs", exist_ok=True)
    tasks, skipped = [], 0
    for block, d, par in conditions():
        for s0 in range(0, total, batch):
            reps = min(batch, total - s0)
            out = f"out_xdrv2/out_{tag(block, d, par)}_s{s0}.txt"
            if resume and os.path.exists(out) and sum(1 for _ in open(out)) >= reps + 1:
                skipped += 1
                continue
            tasks.append(f"{block} {d} {par} {reps} {s0} {out}{GEOM}")
    if resume:
        print(f"  RESUME: skipped {skipped} complete tasks")
    if not tasks:
        print("nothing to do"); return
    open("xdrv2_params.txt", "w").write("\n".join(tasks) + "\n")
    n = len(tasks)     # no throttle (standing instruction)
    open("xdrv2_array.sh", "w").write(f"""#!/bin/bash
#SBATCH -J xdrv2
#SBATCH -p short
#SBATCH -t 0-11:59
#SBATCH -c 1
#SBATCH --mem=4G
#SBATCH -o logs/xdrv2_%A_%a.log
#SBATCH --array=0-{n - 1}

LINE=$(sed -n "$((SLURM_ARRAY_TASK_ID + 1))p" xdrv2_params.txt)
echo "task $SLURM_ARRAY_TASK_ID : $LINE"
python xdrv2_run.py $LINE
""")
    print(f"wrote xdrv2_params.txt ({n} tasks over {len(conditions())} conditions, "
          f"{total} reps each) and xdrv2_array.sh")
    print("submit:  sbatch xdrv2_array.sh")


if __name__ == "__main__":
    main()
```
