## Supplementary Information section 3 code base for "Hypothesis: A modern human range expansion ∼300,000 years ago had a critical role in Neandertal origins": xdrvdeme_submit.py

```
#!/usr/bin/env python3
"""
xdrvdeme_submit.py -- the deme drive ladder down to 52.5% daughters, for Figure S3.4(a).
(12 Sep 2026)

DR: "For left panel of this, run the deme framework down to 52.5% to complete the series."

WHY A RUN, WHEN THE NUMBERS APPEAR TO EXIST ALREADY. xdrv2_summary.csv block T does carry the
deme ladder at d = 0.05 to 0.4 -- 52.5% to 70% daughters -- at 1,000 replicates, and it would
have been tempting to read it. It cannot be used, for two independent reasons:

  1. IT IS AT THE SUPERSEDED GEOMETRY. Both xdrv_run.py and xdrv2_run.py set
     BASE = dict(grid=20, K=20, ...), a post-wave census near 8,000 against the ~1,584 the
     Peyregne anchor implies. The potency of any coefficient scales with 2*Ne*s, so the drive
     does far more work there than at the anchored size. Block T reports P(all ten modern) =
     82% at 52.5% daughters; the anchored Cartesian data in out_xdc2_deme gives a median of
     10.5% at the same rung. An eight-fold gap is what that census difference buys.
  2. IT PREDATES THE RESEED FIX. All 255 files in out_xdrv2 were written before 07:35 on
     12 September, so every one carries the origin-reseeding inflation of about 1.44 points.

  The internal tell that the file is the OLD run rather than the anchored rerun: its block R
  reproduces the note's 29.3% / 12.3% resistance figures, where the anchored rerun gave
  41% / 20%.

WHY robust_run AND NOT joint_run. Panel (a)'s y axis is p_joint -- the probability that ten
sampled males are all modern at both markers. robust_run computes exactly that at stage 2 and
reports it as `joint`; joint_run reports population frequencies, from which p_joint could only
be approximated. Using the same runner as the Cartesian sweep also means the new points are
directly comparable to the band curve rather than nearly so.

TWO STAGES, BECAUSE gamma MOVES WITH DRIVE STRENGTH. Removing archaic-Y males removes their
autosomes with them, so modern autosomal ancestry climbs as the drive strengthens: at the
baseline gamma = 0.12 a d = 0.05 cell comes back at 8.1% autosomal, not 5%. Stage `cal` probes a
gamma ladder at 20 replicates; `prod` inverts a log-linear fit at 5% and runs 400 replicates,
matching the band arm's replicate count.

    python xdrvdeme_submit.py cal      # ~6 CPU-hours
    python xdrvdeme_submit.py prod     # ~23 CPU-hours, once cal has drained
"""
import glob
import math
import os
import subprocess
import sys

HERE = os.path.dirname(os.path.abspath(__file__))

# d is the probability a Y-bearing conception of an expressing male becomes a daughter, so
# daughters = 50*(1+d). This ladder is 52.5% through 100%, matching the band curve's rungs.
DVALS = [0.05, 0.1, 0.2, 0.4, 0.6, 0.8, 1.0]
ARMS = [("xdrive", 0.0, "no fertility cost"), ("xdrive_c10", 0.10, "10% fertility cost")]
GAMMAS = [0.12, 0.18, 0.26, 0.36, 0.50]
CAL_REPS, PROD_REPS = 20, 400
TARGET = 5.0
OUT_CAL, OUT_PROD = "out_xdrvdeme_cal", "out_xdrvdeme"
PART, CAP = "short", 400


def tag(mech, d, g=None):
    t = f"{mech}_d{d:g}".replace(".", "p")
    return t + (f"_g{g:g}".replace(".", "p") if g is not None else "")


def aut_of(path):
    import json
    try:
        return json.load(open(path))["summary"].get("aut")
    except Exception:
        return None


def fit_gamma(mech):
    out = {}
    for d in DVALS:
        pts = []
        for g in GAMMAS:
            a = aut_of(os.path.join(HERE, OUT_CAL, tag(mech, d, g) + ".json"))
            if a:
                pts.append((math.log(g), a))
        if len(pts) < 2:
            continue
        n = len(pts)
        mx = sum(x for x, _ in pts) / n
        my = sum(y for _, y in pts) / n
        den = sum((x - mx) ** 2 for x, _ in pts)
        if den == 0:
            continue
        b = sum((x - mx) * (y - my) for x, y in pts) / den
        if b == 0:
            continue
        a0 = my - b * mx
        # clamped to the probed range: extrapolating off the end of the ladder is how you get a
        # gamma that was never tested and an autosomal proportion nobody checked
        out[d] = min(max(math.exp((TARGET - a0) / b), GAMMAS[0]), GAMMAS[-1])
    return out


def emit(name, tasks, hours="0-05:59"):
    if not tasks:
        print("  nothing to submit")
        return
    pf, sf = f"{name}_params.txt", f"{name}_array.sh"
    open(os.path.join(HERE, pf), "w").write("\n".join(tasks) + "\n")
    open(os.path.join(HERE, sf), "w").write(f"""#!/bin/bash
#SBATCH -J {name}
#SBATCH -p {PART}
#SBATCH -c 1
#SBATCH --mem=2G
#SBATCH -t {hours}
#SBATCH -a 0-{len(tasks) - 1}%{CAP}
#SBATCH -o logs/{name}_%A_%a.log
cd {HERE}
CMD=$(sed -n "$((SLURM_ARRAY_TASK_ID + 1))p" {pf})
echo "$CMD"
eval "$CMD"
""")
    print(f"  {len(tasks)} tasks -> {sf}")
    if "--no-go" in sys.argv:
        print(f"  NOT submitted (--no-go).  sbatch {sf}")
        return
    r = subprocess.run(["sbatch", sf], cwd=HERE, capture_output=True, text=True)
    print(f"  SUBMITTED: {r.stdout.strip()}" if r.returncode == 0
          else f"  !! sbatch FAILED: {(r.stderr or '').strip()}")


def main():
    stage = sys.argv[1] if len(sys.argv) > 1 else "cal"
    os.makedirs(os.path.join(HERE, "logs"), exist_ok=True)

    if stage == "cal":
        os.makedirs(os.path.join(HERE, OUT_CAL), exist_ok=True)
        tasks = []
        for mech, _, _ in ARMS:
            for d in DVALS:
                for g in GAMMAS:
                    out = os.path.join(OUT_CAL, tag(mech, d, g) + ".json")
                    if os.path.exists(os.path.join(HERE, out)):
                        continue
                    tasks.append(f"python robust_run.py deme {mech} {d:g} 1 {CAL_REPS} 0 {out} "
                                 f"--female-bias 1.0 --gamma {g:g}")
        print(f"CALIBRATION  {len(ARMS)} arms x {len(DVALS)} rungs x {len(GAMMAS)} gammas "
              f"x {CAL_REPS} reps")
        emit("xdcal", tasks)
        return

    if stage == "prod":
        os.makedirs(os.path.join(HERE, OUT_PROD), exist_ok=True)
        have = len(glob.glob(os.path.join(HERE, OUT_CAL, "*.json")))
        want = len(ARMS) * len(DVALS) * len(GAMMAS)
        print(f"CALIBRATION on disk: {have}/{want}")
        if have < want and "--resume" not in sys.argv:
            print("  incomplete. Wait for xdcal, or pass --resume to fit on what is there.")
            return
        tasks = []
        for mech, cost, lab in ARMS:
            gs = fit_gamma(mech)
            print(f"  {lab}")
            for d in DVALS:
                if d not in gs:
                    print(f"    {50 * (1 + d):5.1f}% daughters  no gamma fit")
                    continue
                print(f"    {50 * (1 + d):5.1f}% daughters  gamma* = {gs[d]:.3f}")
                out = os.path.join(OUT_PROD, tag(mech, d) + ".json")
                if os.path.exists(os.path.join(HERE, out)):
                    continue
                tasks.append(f"python robust_run.py deme {mech} {d:g} 2 {PROD_REPS} 1000 {out} "
                             f"--female-bias 1.0 --gamma {gs[d]:.4f}")
        print(f"\nPRODUCTION  stage 2, {PROD_REPS} replicates, p_joint reported as `joint`")
        emit("xdprod", tasks, hours="0-11:59")
        return

    raise SystemExit("stage must be 'cal' or 'prod'")


if __name__ == "__main__":
    main()
```
