## Supplementary Information section 3 code base for "Hypothesis: A modern human range expansion ∼300,000 years ago had a critical role in Neandertal origins": ysupp_sim.py

```
#!/usr/bin/env python3
"""
ysupp_sim.py -- how fast must a de novo Y suppressor arise to stop an X-linked distorter?
(13 Sep 2026)

DR, comment 30: "Can we simulate the Y-chromosome mutation rate needed to generate a suppressor
under the scenarios that we are just within an archaic population without hybridization and an
X-distorter arises, or alternatively we are in a hybridization context where a suppressor is
already available: the modern Y?"

THE CLAIM THIS TESTS. Route 2 rests on a sentence in the note:

    "When an X-linked distorter arises, there is high positive selection for de novo mutations
     on chromosome Y to suppress it. But the situation here is different: a suppressor is
     present from the modern Y, so evolutionary constraints are satisfied by spreading the
     modern Y rather than rescuing the archaic one."

That is an argument about WAITING TIMES, and until now it had no numbers behind it. A
population facing a new distorter has two ways out: wait for a suppressing mutation on the Y,
or already have one. This model measures both and asks how large the de novo mutation rate
would have to be for the first route to compete with the second.

NEITHER EXISTING ENGINE CAN DO THIS. spatial_sim3 and bandsim both carry an X-drive locus and a
Y that either suppresses it or does not, but in both the suppressor is a LINEAGE LABEL -- the
modern Y suppresses, the archaic Y does not -- fixed at initialisation. Neither has a mutation
process on the Y, and neither is a within-population model: they are spatial expansions in
which the distorter's fate is entangled with the wavefront. The question here is the ordinary
population-genetic one, so the model is deliberately the simplest thing that can answer it: one
panmictic Wright-Fisher population, no geography, no lineages.

THE MODEL. N diploid individuals, discrete generations, separate sexes.

  X locus     `+` or `D`. Females carry two, males one. A male carrying D distorts his own
              sperm so that a fraction `d` of what would have been sons become daughters --
              the same parameterisation as the spatial engines, offspring being (1 + d)/2
              daughters -- unless his Y suppresses.
  Y locus     `+` or `S`. S fully restores the even sex ratio in a D-carrying male. Each Y
              transmitted mutates + -> S with probability mu.
  costs       D carries a fitness cost c_D where expressed, S a cost c_S. Both default to 0
              and are swept, because a cost-free suppressor is the most favourable case for
              the de novo route and the note should not rest on the most favourable case.

  extinction  a generation with no males of any kind produces no children and the population
              is recorded as extinct. This is the outcome the suppressor is rescuing FROM, and
              it is why the waiting time matters rather than the equilibrium.

SCENARIO A (no hybridisation): one new D on one X, no S anywhere, S reachable only by mutation.
SCENARIO B (hybridisation): the same new D, but a fraction f of Y chromosomes already carry S
            -- the modern Y, entering at the admixture proportion. No waiting is required.

WHAT IS MEASURED, per replicate. The headline statistic is p_S10, the probability that ten
sampled males ALL carry the suppressing Y -- which is the same quantity Figure S3.4a plots,
and deliberately so. DR: "maybe instead you should plot the fraction of 10 males carrying both
mtDNA and Y."

There is no mtDNA in this model, and there could not be: it has no modern and archaic
lineages, so a mitochondrion would have nothing to be modern WITH RESPECT TO. That costs
nothing, because in the spatial models p_joint already EQUALS the Y term -- mtDNA fixes in
100% of established replicates at every drive strength, since under full matrilocality the
expanding matriline carries its own mtDNA by construction. So "ten males all modern at both
markers" and "ten males all carrying the suppressing Y" are the same number there, and the
second is what this model can measure.

Extinct replicates score p_S10 = 0: with no males left, ten modern males cannot be sampled.
That folds the two failure modes -- no suppressor, and no population -- into one axis, which
is what makes the panel commensurable with (a) and (b).

Also recorded: whether a suppressor ever establishes (reaches 5% of Ys), the generation it
does, whether the distorter fixes, whether the population goes extinct, and the minimum male
fraction reached.

    python ysupp_sim.py --scenario A --mu 1e-6 --reps 1000
    python ysupp_sim.py --scenario B --f 0.05 --reps 1000
"""
import argparse
import json
import math
import os
import sys

import numpy as np


def _p_s10(males):
    """P(ten sampled males all carry the suppressing Y). Zero if there are no males."""
    n = int(males.sum())
    if n == 0:
        return 0.0
    return float((males[:, 1].sum() / n) ** 10)


def simulate(N=1500, d=0.9, mu=1e-6, f_S=0.0, c_D=0.0, c_S=0.0,
             gens=5000, seed=0, establish=0.05, male_cap=10.0):
    """One replicate. Returns a dict of outcomes.

    STATE IS ALLELE COUNTS, NOT INDIVIDUALS. The population is panmictic and the loci are two,
    so nothing is gained by tracking individuals and a great deal of speed is lost. Females are
    represented by their two X alleles, males by (X allele, Y allele).
    """
    rng = np.random.default_rng(seed)
    nf = nm = N // 2

    # females: count carrying 0, 1, 2 copies of D. males: (X is D?) x (Y is S?)
    fD = np.array([nf, 0, 0], dtype=np.int64)          # D copies 0, 1, 2
    mXD = np.array([nm, 0], dtype=np.int64)            # X is +, X is D   (Y ignored for now)
    # split males by Y type: rows = X(+ / D), cols = Y(+ / S)
    males = np.zeros((2, 2), dtype=np.int64)
    n_S0 = rng.binomial(nm, f_S)
    males[0, 0] = nm - n_S0
    males[0, 1] = n_S0

    # seed ONE distorter on ONE X, in a female, which is where a new X mutation would arise
    # two thirds of the time and where it is not immediately exposed to selection
    if fD[0] > 0:
        fD[0] -= 1
        fD[1] += 1

    first_S = None
    ever_est = False
    min_male_frac = 1.0
    min_pop = N
    dfix = False

    for g in range(1, gens + 1):
        nf = int(fD.sum())
        nm = int(males.sum())
        tot = nf + nm
        if tot == 0 or nm == 0 or nf == 0:
            return dict(extinct=True, gen_extinct=g, established=ever_est,
                        gen_establish=first_S, distorter_fixed=dfix,
                        min_male_frac=min_male_frac, final_male_frac=0.0, min_pop=min_pop,
                        p_S10=_p_s10(males))
        min_male_frac = min(min_male_frac, nm / tot)
        min_pop = min(min_pop, tot)

        # ---- maternal gametes: D frequency among X's contributed by mothers ----------------
        # relative fertility of a female is 1 - c_D per D copy expressed (additive)
        wf = np.array([1.0, 1.0 - c_D, 1.0 - 2 * c_D]).clip(0.0)
        pf = fD * wf
        if pf.sum() <= 0:
            return dict(extinct=True, gen_extinct=g, established=ever_est,
                        gen_establish=first_S, distorter_fixed=dfix,
                        min_male_frac=min_male_frac, final_male_frac=nm / tot, min_pop=min_pop,
                        p_S10=_p_s10(males))
        pf = pf / pf.sum()
        # a mother of class k transmits D with probability k/2
        q_mat = float(pf[1] * 0.5 + pf[2] * 1.0)

        # ---- paternal gametes, and the sex ratio they imply ---------------------------------
        # a male distorts only if his X is D AND his Y is not S
        wm = np.array([[1.0, 1.0 - c_S],
                       [1.0 - c_D, (1.0 - c_D) * (1.0 - c_S)]]).clip(0.0)
        pm = males * wm
        if pm.sum() <= 0:
            return dict(extinct=True, gen_extinct=g, established=ever_est,
                        gen_establish=first_S, distorter_fixed=dfix,
                        min_male_frac=min_male_frac, final_male_frac=nm / tot, min_pop=min_pop,
                        p_S10=_p_s10(males))
        pm = pm / pm.sum()
        # per father class, the fraction of his offspring that are daughters
        daughters = np.array([[0.5, 0.5],
                              [(1.0 + d) / 2.0, 0.5]])
        frac_daughters = float((pm * daughters).sum())

        # ---- form the next generation -------------------------------------------------------
        # MALE FERTILITY IS CAPPED, AND WITHOUT THIS THE MODEL CANNOT GO EXTINCT.
        # Holding the population at N regardless of the sex ratio means that as males become
        # rare each surviving male simply fathers unboundedly more children, so a distorter can
        # drive the male share to 5% and the population never notices. With no extinction
        # pressure the race the note describes -- damage from the distorter against the arrival
        # of a suppressor -- has nothing at stake, and every replicate ends in a comfortable
        # female-biased equilibrium. That is an artefact of the bookkeeping, not biology.
        #
        # male_cap is the most offspring one male can father in a generation. Production is
        # min(N, male_cap * males), so the population is female-limited at balanced sex ratios
        # and male-limited once the male share falls below 1/male_cap -- 10% at the default
        # cap of 10. Below that the population shrinks geometrically and goes extinct, which
        # is the outcome a suppressor has to arrive in time to prevent.
        n_next = int(min(N, male_cap * nm))
        if n_next < 2:
            return dict(extinct=True, gen_extinct=g, established=ever_est,
                        gen_establish=first_S, distorter_fixed=dfix,
                        min_male_frac=min_male_frac, final_male_frac=nm / tot,
                        min_pop=n_next, p_S10=_p_s10(males))
        n_daughters = rng.binomial(n_next, frac_daughters)
        n_sons = n_next - n_daughters

        # daughters: one X from mother (D with prob q_mat), one from father (D iff his X is D)
        #
        # THE PATERNAL DRAW MUST BE WEIGHTED BY DAUGHTER OUTPUT, AND THIS IS THE DRIVE ITSELF.
        # Using the fitness-weighted father frequency pm[1] -- which is what this line did at
        # first -- says a distorting father sires the same share of daughters as any other,
        # and the distorter then does not spread at all: in a trace at a 20% starting
        # frequency it sat flat at 0.20-0.22 for fifteen generations while the sex ratio
        # skewed to 40% male, which is the signature of a drive that skews sex but transmits
        # no faster. A distorting father's offspring are (1 + d)/2 daughters instead of 1/2,
        # every one of them receives his D-bearing X, and that excess IS how an X-linked
        # distorter increases in frequency. Weighting by the daughter column restores it.
        dau_w = pm * daughters
        dau_w = dau_w / dau_w.sum()
        p_pat_D = float(dau_w[1].sum())
        # A DAUGHTER'S TWO X's ARE INDEPENDENT DRAWS, SO THE THREE GENOTYPE CLASSES ARE ONE
        # MULTINOMIAL. The first version of this drew the maternal and paternal D counts
        # separately and then tried to reconcile them into 0/1/2-copy classes with
        #     both = Binomial(min(kD_mat, kD_pat), 0.5)
        # and a clip, `one = min(one, n_daughters - two)`. That clip is not a safety net, it
        # is a leak: once q_mat and p_pat_D are both high the implied one-copy count exceeds
        # the daughters available, and the excess D copies are silently discarded. It caps the
        # distorter at an artificial ceiling -- in a trace the frequency plateaued at 0.67 and
        # sat there for forty generations while the deterministic recursion for the same
        # parameters goes to fixation and drives the male share to 5%. The population then
        # never becomes male-limited and never goes extinct, which is exactly the behaviour
        # this model exists to measure.
        pz = (1.0 - q_mat) * (1.0 - p_pat_D)
        po = q_mat * (1.0 - p_pat_D) + (1.0 - q_mat) * p_pat_D
        pt = q_mat * p_pat_D
        fD = rng.multinomial(n_daughters, [pz, po, pt]).astype(np.int64)

        # sons: X from mother, Y from father. A distorting father's surviving sons are the
        # ones whose Y-bearing sperm escaped, which does not change WHICH Y they carry.
        kX = rng.binomial(n_sons, q_mat)
        # Y type: drawn from fathers weighted by how many SONS each class contributes
        son_w = pm * (1.0 - daughters)
        if son_w.sum() <= 0:
            return dict(extinct=True, gen_extinct=g, established=ever_est,
                        gen_establish=first_S, distorter_fixed=dfix,
                        min_male_frac=min_male_frac, final_male_frac=0.0, min_pop=min_pop,
                        p_S10=_p_s10(males))
        son_w = son_w / son_w.sum()
        p_Y_S = float(son_w[:, 1].sum())
        kY = rng.binomial(n_sons, p_Y_S)
        # de novo suppressor mutations on the + Ys
        kY += rng.binomial(n_sons - kY, mu)

        males = np.zeros((2, 2), dtype=np.int64)
        # X and Y assort independently
        xD = rng.binomial(kY, kX / n_sons) if n_sons else 0
        males[1, 1] = xD
        males[0, 1] = kY - xD
        rem = n_sons - kY
        xD2 = rng.binomial(rem, kX / n_sons) if n_sons else 0
        males[1, 0] = xD2
        males[0, 0] = rem - xD2

        nS = int(males[:, 1].sum())
        nmales = int(males.sum())
        if nS and first_S is None:
            first_S = g
        if nmales and nS / nmales >= establish:
            ever_est = True
        nD_total = int(fD[1] + 2 * fD[2] + males[1].sum())
        nX_total = int(2 * fD.sum() + males.sum())
        if nX_total and nD_total == nX_total:
            dfix = True
        if nX_total and nD_total == 0:
            # the distorter is gone: nothing left to suppress, stop early
            return dict(extinct=False, gen_extinct=None, established=ever_est,
                        gen_establish=first_S, distorter_fixed=False,
                        min_male_frac=min_male_frac,
                        final_male_frac=nmales / (nmales + int(fD.sum())),
                        distorter_lost=True, gen_end=g, min_pop=min_pop,
                        p_S10=_p_s10(males))

    nmales = int(males.sum())
    nfem = int(fD.sum())
    return dict(extinct=False, gen_extinct=None, established=ever_est,
                gen_establish=first_S, distorter_fixed=dfix,
                min_male_frac=min_male_frac,
                final_male_frac=nmales / max(nmales + nfem, 1),
                distorter_lost=False, gen_end=gens, min_pop=min_pop,
                p_S10=_p_s10(males))


def main():
    ap = argparse.ArgumentParser()
    ap.add_argument("--scenario", default="A", choices=["A", "B"])
    ap.add_argument("--N", type=int, default=1500)
    ap.add_argument("--d", type=float, default=0.9)
    ap.add_argument("--mu", type=float, default=1e-6)
    ap.add_argument("--f", type=float, default=0.0, help="standing S frequency (scenario B)")
    ap.add_argument("--cD", type=float, default=0.0)
    ap.add_argument("--cS", type=float, default=0.0)
    ap.add_argument("--gens", type=int, default=5000)
    ap.add_argument("--male-cap", type=float, default=10.0,
                    help="most offspring one male can father per generation")
    ap.add_argument("--reps", type=int, default=200)
    ap.add_argument("--seed0", type=int, default=0)
    ap.add_argument("--out", default="")
    a = ap.parse_args()

    f_S = a.f if a.scenario == "B" else 0.0
    rows = []
    for s in range(a.seed0, a.seed0 + a.reps):
        rows.append(simulate(N=a.N, d=a.d, mu=a.mu, f_S=f_S, c_D=a.cD, c_S=a.cS,
                             gens=a.gens, seed=s, male_cap=a.male_cap))
    n = len(rows)
    est = sum(1 for r in rows if r["established"])
    ext = sum(1 for r in rows if r["extinct"])
    fix = sum(1 for r in rows if r["distorter_fixed"])
    gs = [r["gen_establish"] for r in rows if r["gen_establish"]]
    summary = dict(scenario=a.scenario, N=a.N, d=a.d, mu=a.mu, f_S=f_S, cD=a.cD, cS=a.cS,
                   reps=n, p_establish=est / n, p_extinct=ext / n, p_dfix=fix / n,
                   median_gen_establish=(float(np.median(gs)) if gs else None),
                   male_cap=a.male_cap,
                   mean_min_male_frac=float(np.mean([r["min_male_frac"] for r in rows])),
                   mean_min_pop=float(np.mean([r.get("min_pop", a.N) for r in rows])),
                   p_S10=float(np.mean([r.get("p_S10", 0.0) for r in rows])))
    print(f"scenario {a.scenario}  N={a.N} d={a.d} mu={a.mu:g} f_S={f_S:g} "
          f"cD={a.cD} cS={a.cS}  reps={n}")
    print(f"  suppressor established   {summary['p_establish']:.3f}")
    print(f"  population extinct       {summary['p_extinct']:.3f}")
    print(f"  distorter fixed          {summary['p_dfix']:.3f}")
    print(f"  median gen to suppressor {summary['median_gen_establish']}")
    print(f"  mean lowest male share   {summary['mean_min_male_frac']:.3f}")
    print(f"  mean lowest population   {summary['mean_min_pop']:.0f} of {a.N}")
    print(f"  P(10 males all suppress) {summary['p_S10']:.3f}")
    if a.out:
        os.makedirs(os.path.dirname(a.out) or ".", exist_ok=True)
        json.dump(dict(summary=summary, rows=rows), open(a.out, "w"))
        print(f"  wrote {a.out}")


if __name__ == "__main__":
    main()
```
