## Supplementary Information section 3 code base for "Hypothesis: A modern human range expansion ∼300,000 years ago had a critical role in Neandertal origins": collect_mt_expansion.py

```
#!/usr/bin/env python3
"""
collect_mt_expansion.py -- P(modern mtDNA fixes) against s under a RANGE EXPANSION, in both
frameworks, for the right-hand panel of Figure S3.2. (12 Sep 2026)

DR: "we also want to show the results for the range expansion, where I expect the mtDNA fitness
gap won't be enough to drive the process."

NO NEW SIMULATIONS WERE NEEDED. The cells already exist inside the `demog` set, which is the
only set routes_build crosses with its residence axis: DEME_RESIDENCE and BAND_RESIDENCE pair
full sex bias at s_mt = 0 with a NON-SEX-BIASED expansion (female_bias = 0.5, beta = 0.5) at
each rung of SMT_LADDER. This script aggregates those cells into one curve per framework.

WHY THE NON-SEX-BIASED ARM IS THE ONLY ONE THAT CAN ANSWER THE QUESTION. Under a matrilocal
expansion the mtDNA is ~100% modern by construction -- the residence rule carries it -- so
asking whether a fitness gap could drive it to fixation is asking about a race already won, and
routes_build deliberately does not cross s_mt with full matrilocality for exactly that reason.
The meaningful comparison is an expansion with no sex bias, where the mtDNA has to get there on
selection and luck alone, which is also the patrilocal case with the markers exchanged.

AGGREGATION. One point per s_mt rung, pooling every admissible demographic combination:
autosomal ancestry 4-6%, expansion completing in 100-1,000 generations, establishing in more
than half of replicates -- the same acceptance region as Table S3.1 and Figure S3.3. Within a
combination only established replicates count, and fixation means modern mtDNA at 100%. Pooling
across combinations is what makes 10 replicates per cell enough: each rung carries thousands.

THE DEME ARM IS PROVISIONAL AND THE SCRIPT SAYS SO. The local out_demog1_deme was written on
11 Sep, before the reseed guard landed at 07:35 on 12 Sep, so its autosomal values are inflated
by about 1.44 points and the 4-6% filter therefore selects slightly the wrong combinations. The
conclusion is two orders of magnitude away from anything that matters, so it will not change,
but the numbers will move a little. Refresh with:

    scp -r o2:/n/groups/reich/DAVID/CLAUDE_ARCHAICSIMS/out_demog1_deme cluster/

    python collect_mt_expansion.py        # -> mt_expansion_curve.csv
"""
import collections
import datetime
import csv
import glob
import json
import os

HERE = os.path.dirname(os.path.abspath(__file__))
CL = os.path.join(HERE, "cluster")
OUT = os.path.join(HERE, "mt_expansion_curve.csv")

AUT_LO, AUT_HI = 3.0, 7.0
GEN_LO, GEN_HI = 100, 1000
ESTABLISH = 0.5
FIXED = 99.9                      # modern mtDNA percentage counted as fixation

# (framework, directory, glob, the key that marks the non-sex-biased arm, its value)
ARMS = [
    ("deme", "out_demog1_deme", "*female_bias0p5*.json", "female_bias", 0.5),
    ("band", "out_demog1_band", "*_beta0p5_*.json",      "beta",        0.5),
]

# The engine fix: cells older than this are pre-reseed-guard. Built from the date rather than
# hardcoded, because the hardcoded constant I first used was exactly 24 h early and so declared
# an entirely pre-fix deme directory "clean" -- the one thing this check exists to catch.
ENGINE_FIX = datetime.datetime(2026, 9, 12, 7, 35, 8).timestamp()


def collect(fw, d, pat, key, val):
    agg = collections.defaultdict(lambda: [0, 0, 0])      # s_mt -> [fixed, reps, cells]
    files = glob.glob(os.path.join(CL, d, pat))
    if not files:
        print(f"  {fw}: no files under cluster/{d} -- skipped")
        return {}, 0, 0
    stale = sum(1 for f in files if os.path.getmtime(f) < ENGINE_FIX)
    for f in files:
        try:
            js = json.load(open(f))
        except Exception:
            continue
        s = js["summary"]
        if abs(s.get(key, 1.0) - val) > 1e-9:
            continue
        rows = js.get("rows") or []
        est = [r for r in rows if r.get("established")]
        if not est or len(est) / len(rows) <= ESTABLISH:
            continue
        aut, eg = s.get("aut"), s.get("end_gen")
        if aut is None or not (AUT_LO <= aut <= AUT_HI):
            continue
        if eg is None or not (GEN_LO <= eg <= GEN_HI):
            continue
        a = agg[s["s_mt"]]
        a[0] += sum(1 for r in est if r["mt"] >= FIXED)
        a[1] += len(est)
        a[2] += 1
    return agg, len(files), stale


DIRNAME = "cluster/out_mtexp_pat"


def collect_mtexp(dirname=DIRNAME):
    """The dedicated full-ladder run from cluster/mtexp_submit.py, if it has landed.

    READS out_mtexp_pat, THE PATRILOCAL RUN. The earlier out_mtexp holds the NON-sex-biased
    expansion, which answers a different question: with no sex bias neither marker is carried by
    the residence rule. Under PATRILOCALITY the Y is carried for free and the mtDNA is the marker
    that has to be rescued, which is what this panel asks about. out_mtexp stays on disk as the
    unbiased comparison but is no longer what the figure draws.

    PREFERRED OVER THE POOLED demog ARM when present, for two reasons. It spans the pulse's whole
    s ladder rather than stopping at 0.64%, and it is one baseline demography at 1,000 / 400
    replicates per point, which is what the solid pulse curves are -- so the dashed and solid
    traces become the same kind of measurement instead of a pooled screen set against a single
    baseline. Batches are summed per (framework, s).
    """
    agg = collections.defaultdict(lambda: [0, 0, 0])
    for f in glob.glob(os.path.join(HERE, dirname, "*.json")):
        try:
            js = json.load(open(f))
        except Exception:
            continue
        s = js["summary"]
        fw = s.get("framework") or ("band" if "band" in os.path.basename(f) else "deme")
        est = [r for r in (js.get("rows") or []) if r.get("established")]
        if not est:
            continue
        a = agg[(fw, s["s_mt"])]
        a[0] += sum(1 for r in est if r["mt"] >= FIXED)
        a[1] += len(est)
        a[2] += 1
    return agg


def main():
    rows = []
    mte = collect_mtexp()
    if mte:
        print(f"using the dedicated full-ladder run in {DIRNAME}")
        print(f"    {'fw':<5} {'s_mt':>8} {'batches':>7} {'reps':>7} {'fixed':>6}   P(mt fixes)")
        for (fw, s) in sorted(mte, key=lambda k: (k[0], k[1])):
            fx, reps, nb = mte[(fw, s)]
            p = fx / reps if reps else float("nan")
            print(f"    {fw:<5} {s:8.5f} {nb:7d} {reps:7d} {fx:6d}      {p:.4f}")
            rows.append(dict(framework=fw, s_mt=s, combos=nb, reps=reps, fixed=fx,
                             p_fix=round(p, 5), provenance="patrilocal baseline full ladder"))
        if not rows:
            raise SystemExit(f"no rows to write -- refusing to truncate {OUT}")
        with open(OUT, "w", newline="") as fh:
            fh.write("# P(modern mtDNA fixes) under a PATRILOCAL range expansion at the\n")
            fh.write(f"# baseline demography, from {DIRNAME} (mtexp_submit.py). Under\n")
            fh.write("# patrilocality the residence rule carries the Y and the mtDNA must be\n")
            fh.write("# rescued by selection, which is the question Figure S3.2 asks.\n")
            w = csv.DictWriter(fh, fieldnames=list(rows[0]))
            w.writeheader(); w.writerows(rows)
        print(f"\nwrote {os.path.basename(OUT)}: {len(rows)} points")
        return

    print(f"{DIRNAME} is empty -- falling back to the pooled 5-rung demog arm")
    print("aggregating the non-sex-biased expansion arm of the demog screen")
    for fw, d, pat, key, val in ARMS:
        agg, n, stale = collect(fw, d, pat, key, val)
        if not agg:
            continue
        flag = "PRE-FIX" if stale > n / 2 else "clean"
        print(f"  {fw}: {n:,} files, {stale:,} written before the reseed guard  [{flag}]")
        print(f"    {'s_mt':>8} {'combos':>7} {'reps':>8} {'fixed':>6}   P(mt fixes)")
        for s in sorted(agg):
            fx, reps, cells = agg[s]
            p = fx / reps if reps else float("nan")
            print(f"    {s:8.5f} {cells:7d} {reps:8d} {fx:6d}      {p:.4f}")
            rows.append(dict(framework=fw, s_mt=s, combos=cells, reps=reps, fixed=fx,
                             p_fix=round(p, 5), provenance=flag))

    # SAME GUARD AS panm3_collect, and for the same reason: OUT is shipped with the code base
    # and opening it for writing truncates it, so a run with no simulation output on disk used
    # to leave the reader with a three-line stub where the table had been.
    if not rows:
        raise SystemExit(
            f"neither {DIRNAME} nor the demog fallback holds any output -- run mtexp_submit.py "
            f"first.\nRefusing to overwrite {os.path.basename(OUT)}, which ships with the "
            "code base.")
    with open(OUT, "w", newline="") as fh:
        fh.write("# P(modern mtDNA fixes) under a non-sex-biased RANGE EXPANSION, pooled over\n")
        fh.write(f"# admissible combinations: autosomal {AUT_LO}-{AUT_HI}%, "
                 f"{GEN_LO}-{GEN_HI} generations, establishment > {ESTABLISH:.0%}.\n")
        fh.write("# Written by collect_mt_expansion.py from cluster/out_demog1_{deme,band}.\n")
        w = csv.DictWriter(fh, fieldnames=list(rows[0]))
        w.writeheader()
        w.writerows(rows)
    print(f"\nwrote {os.path.basename(OUT)}: {len(rows)} points")


if __name__ == "__main__":
    main()
```
