## Supplementary Information section 3 code base for "Hypothesis: A modern human range expansion ∼300,000 years ago had a critical role in Neandertal origins": digitize_psmc.py

```
#!/usr/bin/env python3
"""
digitize_psmc.py -- extract the PSMC trajectories from Figure 4A of Peyregne et al. 2025
(Perygene.pdf, page 8) and summarise them over the periods relevant to the ratchet analysis.

Method: the page is rendered at 450 dpi; the y axis is calibrated from the centres of the
"30,000 / 20,000 / 10,000 / 0" tick labels and the x axis from the "0 / 500,000 / 1,000,000 /
1,500,000" labels. The five archaic genomes are separated by nearest-colour assignment to their
legend swatches; the five modern genomes are black/grey and are taken as an envelope, after
discarding dark pixels lying within 3 rows of a coloured curve (anti-aliasing).

Reported per period: the envelope across genomes, and the TIME-WEIGHTED HARMONIC MEAN, which is
the appropriate average when Ne varies over a period. Haploid (mtDNA / Y) size is half the
diploid value, since only one sex transmits.

    python digitize_psmc.py
"""
import numpy as np
from PIL import Image
import fitz

PDF = "Perygene.pdf"
PAGE, DPI = 7, 450
Y0, PX_PER_10K = 1763.0, 237.67
X0, PX_PER_500K = 676.2, 255.8
LEGEND_ROWS, LEGEND_FROM_X = (1010, 1400), 900
ARCHAIC = {"Denisova 5":     ((16, 78, 139),   "Neandertal"),
           "Chagyrskaya 8":  ((30, 165, 211),  "Neandertal"),
           "Vindija 33.19":  ((0, 206, 209),   "Neandertal"),
           "Denisova 25":    ((255, 134, 134), "Denisovan"),
           "Denisova 3":     ((255, 99, 71),   "Denisovan")}

ne   = lambda y: (Y0 - y) / PX_PER_10K * 10000.0
xof  = lambda ka: X0 + ka * 1000.0 / 500000.0 * PX_PER_500K
kaof = lambda x: (x - X0) / PX_PER_500K * 500.0


def load():
    fitz.open(PDF)[PAGE].get_pixmap(dpi=DPI).save("/tmp/_psmc.png")
    a = np.asarray(Image.open("/tmp/_psmc.png").convert("RGB")).astype(float)
    mx, mn = a.max(2), a.min(2); sat = mx - mn
    coloured = (sat > 45) & (mx > 90)
    near = np.zeros_like(coloured)
    for dy in range(-3, 4):
        near |= np.roll(coloured, dy, 0)
    dark = (mx < 150) & (sat < 40) & (~near)
    return a, coloured, dark


def hmean(pts):
    pts = sorted(pts)
    t = np.array([p[0] for p in pts]); v = np.clip([p[1] for p in pts], 1, None)
    w = np.abs(np.gradient(t))
    return w.sum() / np.sum(w / v)


def amean_pts(pts):
    pts = sorted(pts)
    t = np.array([p[0] for p in pts]); v = np.array([p[1] for p in pts], float)
    w = np.abs(np.gradient(t))
    return float((v * w).sum() / w.sum())


def _unused_amean(pts):
    """Time-weighted ARITHMETIC mean. The harmonic mean is the right average for DRIFT (it is
    dominated by the smallest sizes, which is where coalescence happens). It is NOT the right
    average for the EFFICACY OF SELECTION: a beneficial allele makes progress whenever N is
    large, and the arithmetic mean weights those intervals properly. Reporting both makes the
    choice explicit instead of implicit."""
    import numpy as _np
    v = _np.array([p[1] for p in pts], float)
    w = _np.array([p[2] for p in pts], float) if len(pts[0]) > 2 else _np.ones(len(v))
    return float((v * w).sum() / w.sum())


def main():
    a, coloured, dark = load()

    def archaic(name, lo, hi):
        tgt = np.array(ARCHAIC[name][0], float); out = []
        others = [np.array(v[0], float) for k, v in ARCHAIC.items() if k != name]
        for x in range(int(xof(lo)), int(xof(hi)) + 1):
            ys = np.nonzero(coloured[1000:1795, x])[0] + 1000
            if x > LEGEND_FROM_X:
                ys = ys[(ys < LEGEND_ROWS[0]) | (ys > LEGEND_ROWS[1])]
            if not len(ys): continue
            px = a[ys, x, :]
            d0 = np.linalg.norm(px - tgt, axis=1)
            dm = np.min([np.linalg.norm(px - o, axis=1) for o in others], axis=0)
            k = (d0 < 60) & (d0 <= dm)                 # nearest-colour assignment
            if k.any():
                out.append((kaof(x), ne(float(np.median(ys[k])))))
        return out

    def modern(lo, hi):
        mid, vals = [], []
        for x in range(int(xof(lo)), int(xof(hi)) + 1):
            ys = np.nonzero(dark[1000:1790, x])[0] + 1000
            if x > LEGEND_FROM_X:
                ys = ys[(ys < LEGEND_ROWS[0]) | (ys > LEGEND_ROWS[1])]
            g = []
            for y in ys:
                if g and y - g[-1][-1] <= 4: g[-1].append(y)
                else: g.append([y])
            v = [ne(float(np.mean(r))) for r in g if len(r) >= 2]
            v = [q for q in v if 8000 < q < 30000]
            if v:
                mid.append((kaof(x), (min(v) + max(v)) / 2)); vals += v
        return mid, vals

    def show(tag, pts_by_curve, pooled):
        env = [v for s in pts_by_curve.values() for _, v in s]
        h = hmean(pooled)
        print(f"  envelope across genomes : {min(env):,.0f} - {max(env):,.0f} diploid")
        print(f"  time-weighted harmonic mean: {h:,.0f} diploid  ->  {h/2:,.0f} haploid (mtDNA/Y)")
        return h

    print("=" * 74)
    print("(a) EARLY MODERN HUMANS, 750-250 ka")
    mid, vals = modern(250, 750)
    hm = hmean(mid)
    print(f"  envelope across the 5 modern genomes: {min(vals):,.0f} - {max(vals):,.0f} diploid")
    print(f"  time-weighted harmonic mean: {hm:,.0f} diploid  ->  {hm/2:,.0f} haploid (mtDNA/Y)")

    print("\n(b) ARCHAIC, 750-250 ka")
    byc = {n: archaic(n, 250, 750) for n in ARCHAIC}
    for n, s in byc.items():
        if s:
            v = [p[1] for p in s]
            print(f"    {n:15s} ({ARCHAIC[n][1]:10s}) {min(v):7,.0f} - {max(v):7,.0f}"
                  f"   hmean {hmean(s):7,.0f}")
    nea = [p for n in byc if ARCHAIC[n][1] == "Neandertal" for p in byc[n]]
    allp = [p for s in byc.values() for p in s]
    show("archaic", byc, allp)
    hn = hmean(nea)
    print(f"  Neandertal-lineage genomes only: {hn:,.0f} diploid  ->  {hn/2:,.0f} haploid")

    print("\n(c) NEANDERTAL ANCESTORS POST-ADMIXTURE, 250-50 ka")
    byc2 = {n: archaic(n, 50, 250) for n in ARCHAIC if ARCHAIC[n][1] == "Neandertal"}
    for n, s in byc2.items():
        if s:
            v = [p[1] for p in s]
            print(f"    {n:15s} {min(v):7,.0f} - {max(v):7,.0f}   hmean {hmean(s):7,.0f}")
    pooled2 = [p for s in byc2.values() for p in s]
    h2 = hmean(pooled2)
    env2 = [v for _, v in pooled2]
    print(f"  envelope: {min(env2):,.0f} - {max(env2):,.0f} diploid")
    print(f"  time-weighted harmonic mean: {h2:,.0f} diploid  ->  {h2/2:,.0f} haploid (mtDNA/Y)")
    a2 = amean_pts(pooled2)
    print(f"  time-weighted ARITHMETIC mean: {a2:,.0f} diploid  ->  {a2/2:,.0f} haploid")
    print(f"    (harmonic is the drift average; arithmetic is the selection-efficacy average)")


if __name__ == "__main__":
    main()
```
