## Supplementary Information section 3 code base for "Hypothesis: A modern human range expansion ∼300,000 years ago had a critical role in Neandertal origins": fig_fixation_gaps.py

```
#!/usr/bin/env python3
"""fig_fixation_gaps.py -- the fixation curves alone, with both estimated gaps marked.

DR, 8 Oct 2026: the curves panel on its own, relabelled for uniparental markers generally,
with the median estimated fitness gaps shown for mtDNA (0.32%) and for the Y (0.09-0.17%).

THE CURVES AND THEIR DATA ARE UNCHANGED. Everything here is loaded by the same readers, from
the same two files, as panel b of fig_s42_double.py -- fixation_curves_mtdna.csv for the
migration pulse and mt_expansion_curve.csv for the two range expansions -- and drawn with the
same colours, widths and in-plot labels. Only the frame, the axis titles and the two gap
markers are new, so this figure cannot disagree with Figure S4.2 about what the curves do.

A LINEAR X AXIS, as in the --linear variant: the gaps being marked are all under half a
percent, and on the symlog axis that region is stretched so wide that the two markers sit a
third of the way across the panel and stop reading as "both are tiny". The cost is that the
pulse curve is compressed into the left, which is the point being made.

WHAT THE TWO MARKERS MEAN, AND THE ONE CAVEAT WORTH A CAPTION LINE. The curves are
simulations of MODERN mtDNA: the pulse enters at 5% of transmitting females, the expansions
seed a handful of founding copies. Dropping the Y's estimated gap onto them reads the Y's
selection coefficient against the mtDNA fixation curve, which is only fair to the extent the
two markers share an effective population size and a transmission rule. They do not exactly
-- which is the asymmetry the paper is about -- so the Y marker is drawn as the interval it
is, and the caption should say the curves are the mtDNA simulations.

    python fig_fixation_gaps.py            -> Figure_fixation_gaps.png / .pdf
"""
import csv
import os

import numpy as np
import matplotlib
matplotlib.use("Agg")
matplotlib.rcParams["font.family"] = ["Liberation Sans", "Arial", "DejaVu Sans"]
import matplotlib.pyplot as plt

HERE = os.path.dirname(os.path.abspath(__file__))

DEME_C = "#2166ac"
BAND_C = "#b2182b"
PULSE_C = "black"
GRID_C = "0.88"
DEME_LOCUS_NE = 750
EXP = "mt_expansion_curve.csv"
# A PLAIN LOG X AXIS, cut off at 0.01 (DR, 8 Oct). The point of this version is to show
# where the two estimated gaps fall, and they are 0.09-0.32% on an axis running to 20: on a
# linear axis they sit on top of one another against the left spine with no room to label
# either, which is what defeated the first two attempts. Log spreads them across a third of
# the panel. Cutting at 0.05 drops the s = 0 and negative rungs, so plain log replaces the
# earlier symlog -- there is no longer anything at or below zero to keep on the figure.
XLO, XHI = 0.05, 20.0
XTICKS = [0.05, 0.1, 0.2, 0.5, 1, 2, 5, 10, 20]
GRID_Y = list(range(10, 101, 10))

LABELS = [(PULSE_C, "migration\npulse", 36.0),
          (BAND_C, "range\nexpansion\n(band)", 26.0),
          (DEME_C, "range\nexpansion\n(deme)", 16.0)]

# The estimated gaps, in percent. mtDNA is section 1's RNA-inclusive posterior median; the Y
# is an interval rather than a point, so it is drawn as a band.
MT_MEDIAN = 0.32
Y_LO, Y_HI = 0.09, 0.17
MARK_C = "0.35"


def rd(p):
    return list(csv.DictReader(l for l in open(p) if not l.startswith("#")))


def interp(xs, ys, x):
    return float(np.interp(x, xs, ys))


def crossing(xs, ys, level):
    for i in range(1, len(xs)):
        if ys[i - 1] < level <= ys[i]:
            f = (level - ys[i - 1]) / (ys[i] - ys[i - 1])
            return xs[i - 1] + f * (xs[i] - xs[i - 1])
    return None


def load_pulse():
    rows = [r for r in rd(os.path.join(HERE, "fixation_curves_mtdna.csv"))
            if int(r["locus_Ne"]) == DEME_LOCUS_NE]
    rows.sort(key=lambda r: float(r["s"]))
    return ([100 * float(r["s"]) for r in rows], [float(r["pfix_pct"]) for r in rows])


def load_expansion(fw):
    rows = sorted([r for r in rd(os.path.join(HERE, EXP)) if r["framework"] == fw],
                  key=lambda r: float(r["s_mt"]))
    return ([100 * float(r["s_mt"]) for r in rows], [100 * float(r["p_fix"]) for r in rows])


def main():
    # 8.6 x 4.4 in is 1.96:1, the aspect of the figure DR sent (940 x 480).
    fig, ax = plt.subplots(figsize=(8.6, 4.4))
    fig.subplots_adjust(left=0.105, right=0.975, top=0.965, bottom=0.165)

    curves = {}
    px, py = load_pulse()
    ax.plot(px, py, "-", color=PULSE_C, lw=2.4, zorder=5)
    curves[PULSE_C] = (px, py)
    for fw, col in (("band", BAND_C), ("deme", DEME_C)):
        xs, ys = load_expansion(fw)
        ax.plot(xs, ys, "-", color=col, lw=2.2, zorder=4)
        curves[col] = (xs, ys)

    ax.set_xscale("log")
    ax.set_xlim(XLO, XHI)
    ax.set_ylim(-2, 100)
    ax.set_xticks(XTICKS)
    ax.set_xticklabels([f"{t:g}" for t in XTICKS])
    ax.set_xticks([], minor=True)
    ax.set_yticks(GRID_Y + [0])
    for yv in GRID_Y:
        ax.axhline(yv, color=GRID_C, lw=0.7, zorder=0)
    ax.set_axisbelow(True)
    for sp in ("top", "right"):
        ax.spines[sp].set_visible(True)
    ax.tick_params(labelsize=9.5)
    ax.set_xlabel("selection advantage of modern over archaic lineage "
                  "(fitness gap, %)", fontsize=11)
    ax.set_ylabel("probability modern uniparental\nmarker fixes (%)",
                  fontsize=11, linespacing=1.4)

    # ---- the two estimated gaps --------------------------------------------------------
    # Both are under half a percent, so on this axis they land almost on top of one another
    # at the far left. They are therefore marked at the axis and labelled out in the open
    # gap between the pulse curve and the two expansion curves, with leaders back to the
    # marks, rather than written where they would overlap each other and the black curve.
    ax.axvspan(Y_LO, Y_HI, color=MARK_C, alpha=0.30, lw=0, zorder=1)
    ax.axvline(MT_MEDIAN, color=MARK_C, lw=1.3, ls=(0, (6, 3)), zorder=3)

    # Each label sits ON its own mark (DR, 8 Oct): the Y label centred inside the band, and
    # the mtDNA label split so that one line runs down each side of the dashed rule. No
    # leader lines -- every path from here to open space crossed the pulse or band curve,
    # which is what made the linear version unworkable.
    #
    # Centred GEOMETRICALLY, since the axis is log: the arithmetic midpoint of 0.09 and
    # 0.17 is 0.13, which sits visibly right of centre in the drawn band.
    y_mid = (Y_LO * Y_HI) ** 0.5
    lab = dict(rotation=90, ha="center", va="top", fontsize=9.0, color="0.15",
               linespacing=1.25, zorder=7)
    ax.text(y_mid, 97, "estimated gap, Y chromosome\n(0.09–0.17%)", **lab)
    ax.text(MT_MEDIAN * 0.90, 97, "median estimated gap,", **lab)
    ax.text(MT_MEDIAN * 1.11, 97, "mtDNA (0.32%)", **lab)

    # ---- the curve labels, as in Figure S4.2 -------------------------------------------
    for col, lab, y_lab in LABELS:
        cx_, cy_ = curves[col]
        x_at = crossing(cx_, cy_, y_lab) or cx_[len(cx_) // 2]
        # Back off MULTIPLICATIVELY: on a log axis a fixed step of 0.35 is a huge move at
        # x = 0.5 and an invisible one at x = 15, so the linear version's stepping either
        # overshot past the spine or never cleared the curve.
        x_lab, side = x_at * 0.88, "right"
        for _ in range(40):
            if interp(cx_, cy_, x_lab) < y_lab - 6.0:
                break
            x_lab *= 0.90
        if x_lab < 0.6:                       # no room left of the pulse curve
            x_lab, side = x_at * 1.14, "left"
        ax.text(x_lab, y_lab, lab, color=col, fontsize=9.6, linespacing=1.25,
                ha="right" if side == "right" else "left", va="center", zorder=6)

    base = os.path.join(HERE, "Figure_fixation_gaps")
    for ext in ("png", "pdf"):
        fig.savefig(f"{base}.{ext}", dpi=300)
    print(f"wrote Figure_fixation_gaps.png / .pdf  "
          f"({fig.get_figwidth():.2f} x {fig.get_figheight():.2f} in, "
          f"aspect {fig.get_figwidth() / fig.get_figheight():.2f})")
    for name, x in (("mtDNA 0.32%", MT_MEDIAN), ("Y 0.09%", Y_LO), ("Y 0.17%", Y_HI)):
        print(f"  at {name:12s}  pulse {interp(*curves[PULSE_C], x):5.1f}%   "
              f"band {interp(*curves[BAND_C], x):4.1f}%   "
              f"deme {interp(*curves[DEME_C], x):4.1f}%")


if __name__ == "__main__":
    main()
```
