## Supplementary Information section 3 code base for "Hypothesis: A modern human range expansion ∼300,000 years ago had a critical role in Neandertal origins": fig_s34_drive_and_escape.py

```
#!/usr/bin/env python3
"""
fig_s34_drive_and_escape.py -- the note's Figure S3.4 as TWO panels. (12 Sep 2026)

DR: "merges Table S3.2's content into a LEFT panel of Figure S3.4 which is about the same
drive process (the current one would then show escape)."

  (a)  the sex-distortion ladder that was Table S3.2: how strongly the X has to skew the sex
       ratio for the modern Y to survive, with and without a 10% fertility cost.
  (b)  heritable resistance on the archaic Y eroding the same drive (this was the note's
       Figure S3.4, produced by fig_resistance.py).
  (c)  whether such a suppressor could arise DE NOVO at all, from cluster/ysupp_sim.py.
       Added 13 Sep on DR's instruction: "the new analyses showing that a Y chromosome
       suppressor doesn't work should be added as a new panel in the Y chromosome suppressor /
       driver figure (three figures side by side)".

So the figure now reads: the drive works -> it can be escaped by a resistant Y -> but such a Y
cannot be invented at any plausible mutation rate, which is why the escape has to be inherited
and why an introgressing modern Y, which already suppresses, is the thing that satisfies the
evolutionary pressure. That is the argument the surrounding text makes, in order.

WHICH FIGURE IS WHICH, because the filenames in this folder have drifted and cost time:
  * word/media/image5.png  == Fig_resistance.png            == the note's Figure S3.4  <-- here
  * word/media/image4.png  == figS3_4_thumbnails_embed.png  == the note's Figure S3.3
  * figS3_4_both_frameworks.png (make_si_figures.fig_s34) is embedded nowhere in v64.
  * make_si_figures.fig_s35() draws a two-panel version of the drive/resistance material and is
    also not the embedded figure. This script does not touch any of those.

WHERE THE LEFT PANEL'S NUMBERS COME FROM. Every point is recomputed from the CSVs that Table
S3.2 was built from, not typed out of the docx:
  * deme  ->  xdrv_summary.csv, column P_all10, rows cost = 0 / 0.1, 400 replicates per row.
  * band  ->  band_model/bandmech_summary.csv, family "xdrv", column p_joint, rows cost = 0 /
              0.1, 400 replicates per row.
Both columns are already the quantity the y axis wants: the probability that ten randomly
sampled males are all modern at both mtDNA and the Y. Reproducing the docx table from these
files is exact to the rounding the table shows, so there is no table-versus-CSV conflict to
adjudicate in the panel itself. (The scored-Cartesian-grid p_joint series quoted in the Table
S3.2 note is a DIFFERENT data product -- 200 replicates per admissible parameter combination,
median across combinations -- and does disagree with a recomputation; that is recorded in the
caption file, not plotted, because overlaying a second data product would put seven curves in
a 3.25-inch panel.)

d IS NOT THE X AXIS. The simulations sweep drive strength d; the table and this panel use the
quantity a reader can interpret, the percentage of an F1 male's offspring that are daughters,
which is 50*(1 + d). The mapping is checked against the docx table at import time by
DAUGHTER_PCT below rather than asserted.

THE DEME ARM IS NOW ANCHORED, AND THE PANEL CHANGES SHAPE BECAUSE OF IT. Until 12 Sep the deme
arm came from xdrv_summary.csv at 8.9-10.5% modern autosomal ancestry -- about twice the
proportion the note conditions on -- and reported 99.8-100% at every rung it covered. It now
comes from cluster/out_xdrvdeme, where gamma is refitted per rung so autosomal ancestry lands
at 5%, and it covers the full 52.5-100% ladder rather than starting at 75%. The anchored curve
is far lower:

    daughters      70%    80%    90%    100%
    deme, anchored  74     82     88      90       (was 99.8-100 at the inflated proportion)
    band            97     98     99.5    99.8

So the mechanism is still strong, but "at least around 95%" is true of the band framework and
not of the deme one, and the note's text has to say which.

THE RIGHT PANEL IS A VERBATIM PORT of fig_resistance.py's drawing code -- same two series, same
log x axis, same axis labels, same colours (#1f4e79 / #c0504d, which are fig_resistance.py's
own, NOT the #2166ac / #b2182b used elsewhere in the note; see the note in the caption file).
Only the title wrapping and font sizes change, because the original was drawn at 5.4 inches
wide and this panel is 3.1.

    python fig_s34_drive_and_escape.py
"""
import csv
import os

import matplotlib
matplotlib.use("Agg")
import matplotlib.pyplot as plt

HERE = os.path.dirname(os.path.abspath(__file__))
BAND = os.path.join(HERE, "band_model")

# Left panel: the project's standard framework colours (make_si_figures.py).
DEME_C, BAND_C = "#2166ac", "#b2182b"
# Right panel: fig_resistance.py's own colours, kept so that panel is unchanged.
# ONE PALETTE ACROSS BOTH PANELS. fig_resistance.py used #1f4e79 / #c0504d, a different
# blue and red from the project standard, which was invisible while it was a figure on
# its own and obvious the moment it sat beside panel (a). Same framework must mean the
# same colour within a figure, so the right panel adopts the standard pair.
DEME_R, BAND_R = DEME_C, BAND_C

OUT = os.path.join(HERE, "FigS3_4_drive_and_escape")


def rd(p):
    return list(csv.DictReader(l for l in open(p) if not l.startswith("#")))


def daughter_pct(d):
    """Drive strength -> percentage of an F1 male's offspring that are daughters."""
    return 50.0 * (1.0 + d)


# The rungs the docx Table S3.2 prints, used only to confirm the d -> %daughters mapping.
DAUGHTER_PCT = {0.05: 52.5, 0.1: 55.0, 0.2: 60.0, 0.3: 65.0, 0.4: 70.0,
                0.5: 75.0, 0.8: 90.0, 0.95: 97.5, 1.0: 100.0}
for _d, _p in DAUGHTER_PCT.items():
    assert abs(daughter_pct(_d) - _p) < 1e-9, (_d, _p)


def deme_arm(cost):
    """(daughters %, P(all 10 modern at both markers) %) from cluster/out_xdrvdeme.

    THE OLD SOURCE, xdrv_summary.csv, IS AT THE WRONG AUTOSOMAL PROPORTION and is no longer
    read. Its four rows sit at 8.9-10.5% modern autosomal ancestry -- roughly twice the target
    the whole note is conditioned on -- because gamma was never retuned as the drive strengthened.
    Removing archaic-Y males removes their autosomes too, so modern autosomal ancestry climbs
    with d unless gamma is pulled down to compensate; at the baseline gamma = 0.12 a d = 0.05
    cell comes back at 8.1%. At that inflated proportion the mechanism looks nearly perfect:
    P_all10 = 99.8-100% at every rung it covered, which is where the note's "at least around
    95% of instances" came from.

    out_xdrvdeme (xdrvdeme_submit.py) is the replacement DR asked for -- "For left panel of
    this, run the deme framework down to 52.5% to complete the series" -- with gamma refitted
    per rung so that autosomal ancestry lands at 5%, at 400 replicates through robust_run
    stage 2, which reports p_joint directly. It lands at 3.5-5.4% autosomal and covers the same
    52.5-100% ladder as the band arm, so the two arms are finally comparable and the panel no
    longer stops at 75% on one side.

    The anchored numbers are much lower and the text must follow them: at 70% daughters the
    modern Y is retained in 74% of replicates (66% with the fertility cost), not 99.8%.

    mtDNA fixes in 100% of established replicates at every rung, as it must under full
    matrilocality, so p_joint here IS the probability the modern Y is retained.
    """
    import glob
    import json
    mech = "xdrive" if cost == 0.0 else "xdrive_c10"
    out = []
    for f in glob.glob(os.path.join(HERE, "cluster", "out_xdrvdeme", "*.json")):
        s = json.load(open(f))["summary"]
        if s.get("mech") != mech:
            continue
        out.append((daughter_pct(float(s["val"])), 100.0 * float(s["joint"])))
    return sorted(out)


def band_arm(cost):
    """(daughters %, p_joint %) from bandmech_summary.csv, family xdrv.

    The xdrv family in that file also holds the heritable-resistance rows, which carry a value
    in `res` and no `d`; filtering on a non-empty `d` keeps only the drive ladder.
    """
    rows = [r for r in rd(os.path.join(BAND, "bandmech_summary.csv"))
            if r["family"] == "xdrv" and r["d"] not in ("", None)
            and abs(float(r["cost"] or 0) - cost) < 1e-9
            and not r.get("res")]
    return sorted((daughter_pct(float(r["d"])), 100 * float(r["p_joint"])) for r in rows)


def panel_a(ax):
    series = [
        (deme_arm(0.0),  DEME_C, "o", "-",  "deme, no cost"),
        (deme_arm(0.1),  DEME_C, "o", "--", "deme, 10% fertility cost"),
        (band_arm(0.0),  BAND_C, "s", "-",  "band, no cost"),
        (band_arm(0.1),  BAND_C, "s", "--", "band, 10% fertility cost"),
    ]
    for pts, c, mk, ls, lab in series:
        # Dashed arms get hollow markers as well as a dashed line: the cost and cost-free arms
        # converge above 90% daughters and at print size line style alone is not separable
        # there.
        face = c if ls == "-" else "white"
        ax.plot([x for x, _ in pts], [y for _, y in pts], marker=mk, ls=ls, color=c,
                mfc=face, mew=1.2, lw=1.7, ms=4.6, label=lab, clip_on=False, zorder=3)

    ax.set_xlim(50, 101)
    ax.set_ylim(0, 104)
    ax.set_xticks([50, 60, 70, 80, 90, 100])
    ax.set_xlabel("daughters per F1 male (%)", fontsize=8.2)
    ax.set_ylabel("P(ten sampled males all modern\nat mtDNA and the Y) (%)", fontsize=8.0)
    ax.set_title("Sex distortion retains\n the modern Y", fontsize=9.0)
    ax.legend(fontsize=6.0, loc="lower right", framealpha=0.95, handlelength=2.3,
              borderpad=0.4, labelspacing=0.3)
    ax.tick_params(labelsize=7.4)
    ax.grid(color="0.93", lw=0.7)
    ax.set_axisbelow(True)


def deme_resistance(d=0.95):
    """(resistance rate, P(ten sampled males all modern at both markers) %) from out_xdrv2.

    THE SAME STATISTIC AS PANELS (a) AND (c). DR: "throughout the note, we should be using not
    Y chromosome, but the 10 males retaining both Y and mtDNA." This panel used to plot the
    mean modern Y frequency among males, which is a different quantity from its neighbours, so
    a reader comparing the three panels was comparing two things. The raw files carry mt and Y
    per seed, so the joint sample-of-ten statistic is computable directly: mean over seeds of
    (mt * Y)^10.

    It barely moves the curve -- 98.3/95.2/84.8/66.4/41.3/20.6 against 98.4/95.5/85.4/67.3/
    42.6/21.3 for the mean frequency -- because runs stop at marker absorption, so most
    replicates end with each marker at 0 or 1 and the two measures coincide. The point is the
    axis, not the numbers.

    xdrv2_summary.csv IS STALE AND MUST NOT BE READ FOR THIS PANEL. The CSV was written from
    the first, superseded xdrv2 run and was never regenerated after the anchored rerun landed;
    the raw per-seed files in cluster/out_xdrv2 ARE the rerun. They disagree substantially:

        resistance rate      0.01%   0.03%   0.1%   0.3%    1%     3%
        raw files (rerun)     98.4    95.5   85.4   67.3   42.6   21.3
        xdrv2_summary.csv     98.4    93.1   79.4   60.4   30.9   13.4

    The gap widens with the resistance rate and reaches 11 points at the last rung, so the CSV
    makes resistance look about half again as destructive as it is. The 41% / 20% pair at the
    top two rungs is the recorded signature of the anchored rerun; 29.3% / 12.3% is the
    signature of the superseded one, and that is the pair the CSV -- and therefore the figure
    and the note -- has been carrying.

    Recomputing from the raw files rather than regenerating the CSV keeps the provenance in one
    place and means this panel cannot silently go stale again the next time a rerun lands.

    900 replicates per point, not the 1,000 the CSV claims. Modern autosomal ancestry across
    these cells is 6.3-6.9%, which sits inside the 3-7% acceptance window but outside the old
    4-6% one -- so under the previous window this entire panel was drawn from cells the note's
    own acceptance criterion would have rejected.
    """
    import glob
    agg = {}
    for f in glob.glob(os.path.join(HERE, "cluster", "out_xdrv2", "out_R_*.txt")):
        b = os.path.basename(f)
        dd = float(b.split("_d")[1].split("_e")[0].replace("p", "."))
        if abs(dd - d) > 1e-9:
            continue
        eps = float(b.split("_e")[1].rsplit("_s", 1)[0].replace("p", "."))
        tot, n = agg.get(eps, (0.0, 0))
        for line in open(f).read().splitlines()[1:]:
            p = line.split("\t")
            if len(p) < 8:
                continue
            try:
                mt, yy = float(p[1]), float(p[2])
            except ValueError:
                continue
            tot += (mt * yy) ** 10
            n += 1
        agg[eps] = (tot, n)
    return sorted((e, 100.0 * t / n) for e, (t, n) in agg.items() if n)


def panel_c(ax):
    """Can a suppressing Y arise de novo? Two scenarios on one axis, honestly.

    ONE x AXIS, NOT TWO. The two scenarios are parameterised differently -- scenario A by a
    mutation rate, scenario B by a standing frequency -- so they cannot share an x axis without
    lying about one of them. What they DO share is the outcome: the probability the population
    is driven extinct. So A is drawn as a curve against mu and B enters as horizontal reference
    lines at the extinction risk it achieves, which makes the comparison a reading across
    rather than a forced common axis.

    The comparison is exact at one point and that is what the panel is for: scenario A reaches
    1% extinction at mu = 3e-4, and a modern Y standing at 5% reaches 1% extinction with no
    mutation at all. Four orders of magnitude above a per-site rate to match what introgression
    supplies for free.
    """
    import json
    d = json.load(open(os.path.join(HERE, "cluster", "ysupp_sweep.json")))
    A = [r for r in d["A"] if r["mu"] > 0]
    mus = [r["mu"] for r in A]
    pS = [100 * r["p_S10"] for r in A]
    ext = [100 * r["p_ext"] for r in A]
    B = {r["f_S"]: (100 * r["p_S10"], 100 * r["p_ext"]) for r in d["B"]}

    # PRIMARY AXIS: the sample-of-ten statistic, matching (a) and (b). There is no mtDNA in
    # this model -- it has no modern and archaic lineages, so a mitochondrion would have
    # nothing to be modern with respect to -- but that costs nothing, because in the spatial
    # runs mtDNA is fixed modern in 100% of established replicates under full matrilocality,
    # so "ten males modern at both" and "ten males carrying the suppressing Y" are the same
    # number. Extinct replicates score zero: with no males left, ten cannot be sampled.
    ax.plot(mus, pS, "o-", color=DEME_C, lw=1.7, ms=4.4, label="de novo only")
    # ONE reference line, not two. At 5% and 10% the standing-suppressor extinction risks are
    # 1.0% and 0.3%, which at this scale are the same line; drawing both added a legend entry
    # and no information. 5% is the proportion the rest of the note conditions on.
    if 0.05 in B:
        ax.axhline(B[0.05][0], color=BAND_C, lw=1.6, ls="--",
                   label="modern Y at 5%")
    ax.set_xscale("log")
    ax.set_xlabel("de novo rate per Y", fontsize=8.0)
    ax.set_ylabel("P(ten sampled males all modern\nat mtDNA and the Y) (%)", fontsize=8.0)
    ax.set_title("A suppressor cannot\narise fast enough", fontsize=9.0)
    ax.axvline(1e-8, color="0.6", lw=1.0, ls=":")
    ax.text(1.3e-8, 2.0, "per-site rate", fontsize=6.2, color="0.4",
            ha="left", va="bottom", rotation=90)
    # LOWER LEFT. The curve is a high flat plateau on the left falling to zero on the right,
    # so the empty quadrant is bottom-left -- and that plateau IS the result, three rungs at
    # 22% showing the de novo route doing nothing at any plausible rate. An upper-right legend
    # looked clear but its left edge covered exactly those three points.
    ax.set_xlim(5e-9, 2e-3)
    ax.set_ylim(-1.5, 32)
    # RIGHT AXIS: extinction, which DR asked to keep visible. It is the other half of the
    # story -- the de novo route fails both by not supplying a suppressor and by losing the
    # population -- and putting it opposite rather than dropping it means the panel carries
    # both without either axis having to serve two quantities.
    axr = ax.twinx()
    axr.plot(mus, ext, "s:", color="0.45", lw=1.4, ms=3.6, mfc="white",
             label="population extinct")
    axr.set_ylabel("% of populations extinct", fontsize=7.4, color="0.35")
    axr.set_ylim(-1.5, 32)
    axr.tick_params(labelsize=6.8, colors="0.35")
    axr.grid(False)
    h1, l1 = ax.get_legend_handles_labels()
    h2, l2 = axr.get_legend_handles_labels()
    # CENTRE LEFT. Upper left sat on the red reference line and the grey extinction plateau at
    # once; the blue curve is flat at zero across the left half, so the band between it and
    # those two is the only clear space.
    ax.legend(h1 + h2, l1 + l2, fontsize=5.6, loc="center left", framealpha=0.95,
              handlelength=1.8, borderpad=0.3, labelspacing=0.22)
    ax.grid(color="0.93", lw=0.7)
    ax.set_axisbelow(True)
    ax.tick_params(labelsize=7.4)


def panel_b(ax):
    deme = deme_resistance(0.95)
    band = sorted(((float(r["res"]), 100 * float(r["Y"]))
                   for r in rd(os.path.join(BAND, "bandmech_summary.csv"))
                   if r.get("family") == "xdrv"
                   and r.get("res") not in (None, "", "0", "0.0")))
    ax.plot([x for x, _ in deme], [y for _, y in deme], "o-", color=DEME_R, lw=1.7, ms=4.6,
            label="deme model")
    ax.plot([x for x, _ in band], [y for _, y in band], "s-", color=BAND_R, lw=1.7, ms=4.6,
            label="band model")
    ax.set_xscale("log")
    # NO SCIENTIFIC NOTATION, AND NO RAW epsilon. DR: "Express things as percentage of
    # daughters as in the other plots, and on the x-axis don't use scientific notation."
    # The resistance rate is a probability per transmission, so it reads as a percentage on
    # the same footing as every other rate in the figure, and the six rungs 1e-4 ... 3e-2
    # become 0.01% ... 3%. The axis stays logarithmic -- the rungs are half-decade spaced and
    # linear would pile five of the six against the left spine -- but the labels are now plain
    # decimals, which is what the scientific notation was obscuring.
    TICKS = [1e-4, 3e-4, 1e-3, 3e-3, 1e-2, 3e-2]
    ax.set_xticks(TICKS)
    ax.set_xticklabels(["0.01", "0.03", "0.1", "0.3", "1", "3"])
    ax.set_xticks([], minor=True)
    ax.xaxis.set_major_formatter(plt.FixedFormatter(
        ["0.01", "0.03", "0.1", "0.3", "1", "3"]))
    # WHAT THIS AXIS ACTUALLY IS. DR: "could you change panel (b)'s axis to de novo mutation
    # rate needed to do this? That's more informative than heritable suppressor rate." Agreed,
    # and it is what the engine implements -- but the denominator matters. spatial_sim3's
    # xdrv_res is applied EACH TIME a Y-bearing conception in a distorting male would be
    # flipped: with that probability the Y instead survives and converts to a permanently
    # drive-resistant archaic Y. So it is a de novo origination rate per drive-exposed
    # conception, not per generation and not per site, and labelling it "mutation rate" alone
    # would invite a reader to compare it directly with a per-site rate of ~1e-8, which is a
    # different quantity by however many drive events a lineage experiences.
    # SHORT ON THE AXIS, DEFINED IN THE CAPTION. DR: "come up with a shorter x-axis label for
    # panel b that is less jargonny and explain it in the legend." The denominator still has to
    # be stated somewhere -- xdrv_res is applied per conception exposed to the drive, not per
    # generation and not per site, so an unqualified rate invites comparison with a per-site
    # 1e-8 that is a different quantity -- but the caption is the place for it, not a
    # two-line axis label on a 3-inch panel.
    ax.set_xlabel("Y suppressor mutation rate (%)", fontsize=8.2)
    ax.set_ylabel("P(ten sampled males all modern\nat mtDNA and the Y) (%)", fontsize=8.0)
    ax.set_ylim(0, 104)
    ax.tick_params(labelsize=7.4)
    ax.grid(color="0.93", lw=0.7)
    ax.set_axisbelow(True)
    # BOTH curves are at 97.5% daughters (the engines were both run at xdrive = 0.95),
    # so the strength belongs in the legend title rather than on one entry or in floating
    # text. An earlier label named only the deme curve, leaving the band curve's drive
    # strength unstated, and two attempts to place it as free text collided with the band
    # curve along the top and the legend along the bottom left.
    ax.legend(fontsize=6.0, loc="lower left", framealpha=0.95,
              title="at 97.5% daughters", title_fontsize=6.0)
    # DR comment 29: "Add this 0.22% number (if you agree with it) as a vertical line in panel
    # b if you agree with the argument." I do agree with it, and the line is the whole point of
    # the panel: 0.22% is the note's estimate of the TOTAL de novo deleterious input into genes
    # anywhere on the Y per generation (3.0e-8 per bp x ~100,000 bp of X-degenerate exonic span
    # x 0.72 of changes being functional). Every one of those mutations would have to suppress
    # this particular distorter for the de novo route to reach the rate the curves require --
    # which is why the line sits to the LEFT of where either curve has fallen far.
    # THE AXIS IS IN RAW RATE, NOT PERCENT. The tick LABELS read 0.01 to 3 but the plotted x
    # values are the rates themselves, 1e-4 to 3e-2, relabelled. So 0.22% is x = 0.0022, and
    # drawing the line at 0.22 puts it at 220% -- off the right-hand edge, which is exactly
    # where the first attempt landed.
    ax.axvline(0.0022, color="0.45", lw=1.1, ls="--", zorder=2)
    # LABEL AT THE FOOT OF THE LINE. Placing it top-left of the line put it straight through
    # the band curve, which is still at 95% there; the top is occupied by the band curve out to
    # 0.3% and the lower left by the legend. The strip below y = 15 is empty everywhere right
    # of the legend, and centring on the line makes clear which line it names.
    ax.text(0.0022, 4, "all de novo\nY mutation", fontsize=6.4, color="0.35",
            ha="center", va="bottom", linespacing=1.25)
    ax.set_title("Stopping this needs an\nimplausible mutation rate", fontsize=9.0)
    # The deme curve is drawn at ONE drive strength and the reader must be told which. It was
    # labelled "d = 0.95", the engine's internal parameter, which appears nowhere else in the
    # note; 50*(1 + d) = 97.5% daughters is the same quantity on panel (a)'s x axis.
    # THE DRIVE STRENGTH GOES IN THE LEGEND, not in floating text. Each curve is drawn at a
    # single drive strength and the reader has to be told which, but this panel has no empty
    # corner: the band curve occupies the top, the deme curve the diagonal, and the legend the
    # bottom left. Two successive placements of a free-standing label collided with one of the
    # three. Naming the strength in the legend entry it belongs to costs no space at all, and
    # it also fixes the older problem that the label only ever named the deme curve, leaving
    # the band curve's drive strength unstated.


def main():
    # THREE PANELS ACROSS A 6.5 in TEXT BLOCK, so each is about 2.05 in wide against the 3.1 in
    # the two-panel version gave them. Height goes up a little to keep the axes from becoming
    # letterbox-shaped, and the per-panel font sizes come down; the alternative -- keeping the
    # old sizes -- overruns every axis label into its neighbour.
    # TWO PANELS, NOT THREE. The third panel was dropped on 13 Sep after DR asked for it:
    # what he wanted plotted -- P(ten males modern at both markers) against the rate at which
    # a suppressing Y arises, starting high and falling -- is precisely what panel (b) already
    # shows, and shows better, because (b) is a range expansion in both frameworks at the
    # note's baseline while the panmictic ysupp_sim model is neither. A second curve of the
    # same shape measuring nearly the same thing would have been worse than none. What the
    # third panel contributed instead is one sentence of body text: a suppressor arising de
    # novo in a CLOSED archaic population, with no hybridisation, rescues nothing at any
    # plausible rate. ysupp_sim.py and its sweep stay in the code base and the workbook.
    fig, (axa, axb) = plt.subplots(1, 2, figsize=(6.5, 3.05))
    panel_a(axa)
    panel_b(axb)
    fig.tight_layout(rect=(0.015, 0, 1, 0.95), w_pad=2.0)
    for i, L in enumerate("ab"):
        fig.text(0.004 + 0.518 * i, 0.978, L, fontsize=12, weight="bold", va="top")
    fig.savefig(OUT + ".png", dpi=200)
    fig.savefig(OUT + ".pdf")
    print("  wrote", os.path.basename(OUT) + ".png / .pdf")
    for cost in (0.0, 0.1):
        print(f"  deme cost={cost}: ", ", ".join(f"{x:g}%->{y:.1f}" for x, y in deme_arm(cost)))
        print(f"  band cost={cost}: ", ", ".join(f"{x:g}%->{y:.1f}" for x, y in band_arm(cost)))


if __name__ == "__main__":
    main()
```
