## Supplementary Information section 3 code base for "Hypothesis: A modern human range expansion ∼300,000 years ago had a critical role in Neandertal origins": fig_sensitivity.py

```
#!/usr/bin/env python3
"""
fig_sensitivity.py -- the old Figure 3, rebuilt as two one-parameter-at-a-time sensitivity
panels, one per framework. (11 Sep 2026)

DR: "Add a version of what we used to call Figure 3, showing, for both Appendix A and B, how
varying each parameter and holding the others constant affects: autosome proportion, mt
proportion, Y proportion, and sweep duration."

    figS3_5_sensitivity_deme.png/.pdf    Appendix A, the deme framework: grid, K, m, gamma, r
    figS3_6_sensitivity_band.png/.pdf    Appendix B, the band framework: K, e, r, alpha,
                                         sigma_b, sigma_e

Parameters are columns, the four response variables are rows, y-axes are shared across a row,
and the note's baseline for each parameter is drawn as a light vertical line so the reader can
see where the other four (or five) parameters are pinned.


WHERE THE NUMBERS COME FROM, AND WHY NOT FROM THE CARTESIAN SWEEP
-----------------------------------------------------------------
There are exactly two one-parameter-at-a-time scans on disk, and they are the two halves of
this figure:

    deme   figure3_full.csv          written by cluster/scan4_collect.py from scan4_run.py.
                                     1,500 replicates per point. Baseline s = 0, theta = 1,
                                     i.e. NO mechanism -- pure demography.
    band   band_model/bandsi_summary.csv, family "scanb", written by bandsi_collect.py.
                                     400 replicates per point. Baseline beta = 0 (matrilocal),
                                     e = 0.193 (the calibrated fexp value), s = 0, theta = 1.

The obvious alternative, slicing the big Cartesian sweep in cluster/out_routes{1,2}_{deme,band},
does NOT work, for three independent reasons, each of which was checked rather than assumed:

  1. There is no null cell in it. `theta` is the RELATIVE FERTILITY of F1 males in both engines
     (spatial_sim2.Config.theta_f1m = 1.0, bandsim.Config.theta = 1.0 are the defaults, meaning
     "no effect"), but routes_build's theta ladder is [0.0, 0.02, 0.05, 0.10, 0.20]. So the
     "theta = 0" cells are F1 males with ZERO fertility -- the strongest possible form of the
     cultural mechanism, not the null. The sY ladder starts at 0.01 and the xdrive ladder at
     0.1. Every cell of that sweep carries an active mechanism.
  2. The deme geometry axes are not independent there. The note says so itself: "I do not vary
     grid, K and m independently of one another. Instead, I only use combinations whose
     measured effective size falls within 10% of the anchor." Holding K = 11 and m at baseline
     while varying grid leaves exactly one admissible cell.
  3. cluster/out_demog1_deme and out_demog1_band -- the ROUTE_SET="demog" directories named in
     the brief -- do not exist. That set has not been started.

MISSING DATA IS SHOWN, NOT PAPERED OVER. Nothing is interpolated across a gap. A parameter value
with no run gets no marker and the line breaks. A value that is on the note's stated ladder but
was never run is marked with a small open grey circle on the axis floor. A parameter with no
data at all still gets its column of four panels, empty, labelled "runs pending".

    python fig_sensitivity.py
"""
import csv
import os

import numpy as np
import matplotlib
matplotlib.use("Agg")
import matplotlib.pyplot as plt

HERE = os.path.dirname(os.path.abspath(__file__))

FOOTNOTE = ("Light vertical line = baseline value at which the other parameters are held. "
            "Open grey circles on the axis floor = values on the note's ladder with no run "
            "yet; lines break at gaps and nothing is interpolated across them.")
BANDDIR = os.path.join(HERE, "band_model")

# ---------------------------------------------------------------- the four response variables
# (key, row label, colour). Order is the order DR asked for.
ROWS = [("auto", "Autosomal\n% modern",  "#2166ac"),
        ("mt",   "mtDNA\n% modern",      "#762a83"),
        ("Y",    "Y\n% modern",          "#b2182b"),
        ("gen",  "Sweep duration\n(generations)", "#404040")]

GREY = "0.62"


# ---------------------------------------------------------------- parameter definitions
# `ladder` is the range the NOTE says was tested (v54, "Variation of demographic parameters").
# `base` is the note's baseline. `log` selects a log x-axis. Values outside `span` are dropped
# so that each panel shows the range the note claims to have explored.
DEME_PARAMS = [
    dict(key="grid",  label="grid (demes per side)", base=12,
         ladder=[6, 8, 10, 12, 14, 16, 20], span=(5.4, 22), log=False,
         ticks=[6, 8, 10, 12, 14, 16, 20]),
    # K has NO one-at-a-time ladder in the note. The note's "17 values from 6 to 46" is
    # ne_screen.KS, and that ladder exists only inside the (grid, K, m) manifold -- "I do not
    # vary grid, K and m independently of one another". Treating those 17 rungs as gaps in a
    # one-at-a-time scan would shatter this curve into eight isolated dots and imply a sparsity
    # that is not there, so the ladder here is scan4's own. (ne_screen.csv in fact holds 26
    # distinct K, 23 of them admissible, not 17 and 14; see PROVENANCE_AUDIT.md §2.4.)
    dict(key="K",     label="K (carrying capacity)", base=11,
         ladder=[8, 10, 12, 16, 20, 24, 32, 40],
         span=(6, 50), log=True, ticks=[8, 11, 16, 24, 40]),
    dict(key="m",     label="m (migration rate)", base=0.10,
         ladder=[0.05, 0.08, 0.10, 0.12, 0.15], span=(0.045, 0.16), log=False,
         ticks=[0.05, 0.08, 0.10, 0.12, 0.15]),
    dict(key="gamma", label="γ (assortative-mating\ndiscount)", base=0.12,
         ladder=[0.08, 0.12, 0.20, 0.30, 0.42, 0.55, 0.70, 1.00], span=(0.07, 1.15), log=True,
         ticks=[0.08, 0.12, 0.20, 0.30, 0.55, 1.00]),
    dict(key="r",     label="r (modern fitness\nadvantage)", base=2.0,
         ladder=[1.5, 2.0, 3.0], span=(1.4, 3.2), log=False, ticks=[1.5, 2.0, 2.5, 3.0]),
]

BAND_PARAMS = [
    dict(key="K",       label="K (optimal band size)", base=25,
         ladder=[10, 15, 25, 40, 60, 90], span=(9, 100), log=True,
         ticks=[10, 15, 25, 40, 60, 90]),
    dict(key="e",       label="e (exogamy rate)", base=0.19,
         ladder=[0.08, 0.13, 0.19, 0.26, 0.35, 0.45, 0.56], span=(0.07, 0.63), log=True,
         ticks=[0.08, 0.19, 0.35, 0.56]),
    dict(key="r",       label="r (modern fitness\nadvantage)", base=2.0,
         ladder=[1.5, 2.0, 3.0], span=(1.4, 3.2), log=False, ticks=[1.5, 2.0, 2.5, 3.0]),
    dict(key="alpha",   label="α (density regulation)", base=0.8,
         ladder=[0.4, 0.8, 1.6], span=(0.3, 1.8), log=False, ticks=[0.4, 0.8, 1.2, 1.6]),
    dict(key="sigma_b", label="σ$_b$ (band step, km)", base=60,
         ladder=[30, 60, 120], span=(25, 145), log=True, ticks=[30, 40, 60, 90, 120]),
    dict(key="sigma_e", label="σ$_e$ (exogamy dispersal,\nkm)", base=80,
         ladder=[40, 80, 160], span=(35, 195), log=True, ticks=[40, 50, 80, 120, 180]),
]


def _f(x):
    try:
        v = float(x)
    except (TypeError, ValueError):
        return np.nan
    return v


# ---------------------------------------------------------------- deme data (scan4)
def load_deme():
    """figure3_full.csv -> {param: {value: {auto, mt, Y, gen, n}}}.

    K < 7 is dropped. scan4_run.py refuses to run it ("K < 7: deme reproductive failure
    collapses the archaic population") and ne_screen.csv marks no K = 6 geometry admissible;
    the two rows that survive in the CSV predate that guard, sit at 94-96% autosomal ancestry
    because the archaic population collapses rather than being replaced, and have an undefined
    Y. They are an artefact of a broken configuration, not a sensitivity result.
    """
    path = os.path.join(HERE, "figure3_full.csv")
    out = {p["key"]: {} for p in DEME_PARAMS}
    for r in csv.DictReader(open(path)):
        panel = r["panel"]
        if panel not in out:
            continue
        v = _f(r["value"])
        if panel == "K" and v < 7:
            continue
        out[panel][v] = dict(auto=_f(r["auto"]), mt=_f(r["mtDNA"]), Y=_f(r["Y"]),
                             gen=_f(r["ext_gen"]), n=_f(r["n"]))
    return out


# ---------------------------------------------------------------- band data (scanb)
def load_band():
    """bandsi_summary.csv family scanb -> the same shape. Proportions are stored 0-1."""
    path = os.path.join(BANDDIR, "bandsi_summary.csv")
    out = {p["key"]: {} for p in BAND_PARAMS}
    # scanb's panel names. ALPHA IS NOW MAPPED. It was left out with the comment "there is no
    # alpha panel", which was true of the DATA and became untrue of the submit script on
    # 13 Sep, when alpha was added to bandsi_submit's SCAN_PANELS -- so the reader would have
    # gone on dropping the column even once the cells existed. Mapped now; the column fills
    # the moment out_bandsi carries alpha_0.4 / alpha_0.8 / alpha_1.6.
    panel_map = {"K": "K", "e": "e", "r": "r", "alpha": "alpha",
                 "sigma_b": "sigma_b", "sigma_e": "sigma_e"}
    for r in csv.DictReader(open(path)):
        if r["family"] != "scanb":
            continue
        key = panel_map.get(r["panel"])
        if key is None:
            continue
        out[key][_f(r["value"])] = dict(auto=100 * _f(r["aut"]), mt=100 * _f(r["mt"]),
                                        Y=100 * _f(r["Y"]), gen=_f(r["end_gen_med"]),
                                        n=_f(r["n"]))
    return out


# ---------------------------------------------------------------- drawing
def pick_ticks(cands, base, maxn=5):
    """At most `maxn` labelled ticks per panel, always keeping the ends and the baseline.

    Labelling every rung crams 20 numbers into a 2-inch panel; tight_layout then steals the
    vertical space to fit them and the panels collapse to a few pixels tall.
    """
    cands = sorted(cands)
    if len(cands) <= maxn:
        return cands
    idx = sorted(set(np.linspace(0, len(cands) - 1, maxn).round().astype(int).tolist()))
    if base in cands:
        b = cands.index(base)
        if b not in idx:                      # swap in the baseline for its nearest neighbour
            idx[min(range(len(idx)), key=lambda k: abs(idx[k] - b))] = b
    return sorted({cands[i] for i in idx})


def build(params, data, title, subtitle, stem, dur_label):
    ncol, nrow = len(params), len(ROWS)
    fig, axes = plt.subplots(nrow, ncol, figsize=(1.95 * ncol + 1.15, 7.4),
                             sharey="row", squeeze=False)

    for j, p in enumerate(params):
        d = data[p["key"]]
        xs = sorted(d)
        # restrict to the range the note says it explored
        lo, hi = p["span"]
        xs = [x for x in xs if lo <= x <= hi]
        # ladder values inside the plotted span that were never run
        missing = [v for v in p["ladder"] if lo <= v <= hi and v not in d]

        for i, (key, _lab, col) in enumerate(ROWS):
            ax = axes[i][j]
            ax.axvline(p["base"], color="#c9d6e8", lw=1.6, zorder=0)
            if p["log"]:
                ax.set_xscale("log")
            ax.set_xlim(*p["span"])
            ax.tick_params(labelsize=7.5)
            ax.grid(alpha=0.25, lw=0.5)
            for s in ("top", "right"):
                ax.spines[s].set_visible(False)

            ticks = p.get("ticks") or pick_ticks(
                set(xs) | set(v for v in p["ladder"] if lo <= v <= hi), p["base"])
            ax.set_xticks(ticks)
            ax.set_xticklabels([f"{t:g}" for t in ticks])
            ax.minorticks_off()

            if not xs:
                ax.text(0.5, 0.5, "runs\npending", transform=ax.transAxes, ha="center",
                        va="center", fontsize=8, style="italic", color=GREY)
                continue

            # A ladder value with no run becomes an explicit NaN, so the line BREAKS there
            # instead of being drawn straight through it. Nothing is interpolated across a
            # gap. (The same happens where a response is undefined at a value that has runs,
            # e.g. a Y proportion in a configuration where the modern Y never established.)
            gx = sorted(set(xs) | {v for v in missing if xs[0] < v < xs[-1]})
            gy = [d[x][key] if x in d else np.nan for x in gx]
            ax.plot(gx, gy, "-o", ms=3.4, lw=1.4, color=col, zorder=3)

        axes[-1][j].set_xlabel(p["label"], fontsize=8.5)

    # Row y-limits are fixed only AFTER every column has been drawn: freezing them column by
    # column (which is what plotting the "not run" markers at get_ylim()[0] used to do) turns
    # autoscaling off for the shared row and clips every later column's data off the top.
    for i, (key, lab, _c) in enumerate(ROWS):
        axes[i][0].set_ylabel(dur_label if key == "gen" else lab, fontsize=8.5)
        ax0 = axes[i][0]
        if key == "mt":
            ax0.set_ylim(-4, 104)
        else:
            top = max((a.get_ylim()[1] for a in axes[i]), default=1.0)
            ax0.set_ylim(0, top)

    # "not run" ladder values, on the floor of every panel, drawn last so they sit on the spine
    for j, p in enumerate(params):
        d = data[p["key"]]
        lo, hi = p["span"]
        missing = [v for v in p["ladder"] if lo <= v <= hi and v not in d]
        if not missing:
            continue
        for i in range(nrow):
            ax = axes[i][j]
            yl = ax.get_ylim()
            ax.plot(missing, [yl[0]] * len(missing), "o", ms=4, mfc="none", mec=GREY,
                    mew=0.9, clip_on=False, zorder=4)
            ax.set_ylim(*yl)

    fig.suptitle(title, fontsize=11.5, y=0.995)
    fig.text(0.5, 0.963, subtitle, ha="center", fontsize=7.6, color="0.32")
    fig.tight_layout(rect=(0, 0.018, 1, 0.955))
    # FOOTNOTE is a module constant so a caller can drop it. The Extended Data version does:
    # DR asked for it out of that figure, where the same two sentences belong in the caption.
    # The supplementary figures keep it, because they are read away from their captions.
    if FOOTNOTE:
        fig.text(0.012, 0.004, FOOTNOTE, fontsize=6.8, color="0.42")

    for ext in ("png", "pdf"):
        fig.savefig(os.path.join(HERE, f"{stem}.{ext}"), dpi=200, bbox_inches="tight")
    plt.close(fig)
    print(f"  wrote {stem}.png / {stem}.pdf")


# ---------------------------------------------------------------- coverage report
def report(params, data, name):
    print(f"\n{name}")
    print(f"  {'parameter':10s} {'pts':>4}  {'values with data':52s} missing from the note's "
          f"ladder")
    for p in params:
        d = data[p["key"]]
        lo, hi = p["span"]
        xs = [x for x in sorted(d) if lo <= x <= hi]
        miss = [v for v in p["ladder"] if v not in d]
        print(f"  {p['key']:10s} {len(xs):4d}  "
              f"{', '.join(f'{x:g}' for x in xs)[:52]:52s} "
              f"{', '.join(f'{v:g}' for v in miss) if miss else '-'}")
    print(f"\n  {'param':8s} {'value':>7} {'n':>6} {'auto%':>7} {'mt%':>7} {'Y%':>7} {'gens':>7}")
    for p in params:
        d = data[p["key"]]
        lo, hi = p["span"]
        for x in sorted(d):
            if not lo <= x <= hi:
                continue
            row = d[x]
            def s(v, f="%7.2f"):
                return "     --" if v != v else f % v
            print(f"  {p['key']:8s} {x:7g} {s(row['n'], '%6.0f'):>6} {s(row['auto'])} "
                  f"{s(row['mt'])} {s(row['Y'])} {s(row['gen'], '%7.0f')}")


def main():
    deme, band = load_deme(), load_band()

    build(DEME_PARAMS, deme,
          "Figure S3.5  Sensitivity of the deme framework (Appendix A) to each "
          "demographic parameter",
          "One parameter varied at a time about the baseline; all others held fixed. "
          "scan4 (figure3_full.csv), 2,000 replicates per point, no mechanism (s = 0, θ = 1), "
          "full matrilineality.",
          "figS3_5_sensitivity_deme",
          "Generations to\narchaic extinction")

    build(BAND_PARAMS, band,
          "Figure S3.6  Sensitivity of the band framework (Appendix B) to each "
          "demographic parameter",
          "One parameter varied at a time about the baseline; all others held fixed. "
          "scanb (bandsi_summary.csv), 2,000 replicates per point, no mechanism (s = 0, θ = 1), "
          "β = 0 (matrilineal), e* = 0.193.",
          "figS3_6_sensitivity_band",
          "Generations to\nmarker absorption\n(median)")

    report(DEME_PARAMS, deme, "DEME (Appendix A) -- source: figure3_full.csv, family scan4")
    report(BAND_PARAMS, band, "BAND (Appendix B) -- source: band_model/bandsi_summary.csv, "
                              "family scanb")


if __name__ == "__main__":
    main()
```
