## Supplementary Information section 3 code base for "Hypothesis: A modern human range expansion ∼300,000 years ago had a critical role in Neandertal origins": fig_thumbnails_all.py

```
#!/usr/bin/env python3
"""
fig_thumbnails_all.py -- Figure S3.4: one thumbnail per mechanism, genetic AND cultural.

This is dmi_thumbnails.py generalised. That script drew the nineteen DMI / restorer / X-drive
architectures of Table S3.2; this one keeps its panel exactly as the author approved it and adds
the six cultural mechanisms after them, so the whole space is one figure.

    python fig_thumbnails_all.py [--aut-lo 3 --aut-hi 7] [--refresh]
        -> figS3_4_thumbnails.png (300 dpi) and figS3_4_thumbnails.pdf

THE PANEL (unchanged from dmi_thumbnails.py). Every screened combination in the Cartesian
search is one point, at the modern autosomal ancestry it produces (x, log) and the number of
generations its expansion takes (y, log). The grey rectangle is the acceptance region: the
autosomal window crossed with 100-1,000 generations, drawn under the points so it highlights
rather than masks whatever density is inside it. Blue is the deme framework, red the band
framework. The bare grey number in the top left is the fraction of replicates that established,
the third acceptance criterion and the one with no axis of its own. Stage 1 only; scoring is
deliberately ignored, because the question a thumbnail answers is whether there is any density
in the acceptance region, not what the joint retention probability is given that there is.

WHY FIVE OF THE SIX CULTURAL PANELS ARE EMPTY FRAMES. Only one cultural mechanism -- reduced
F1-male fertility, mechanism key "theta" -- has been run through the two-stage Cartesian
pipeline, so only it has the two quantities this panel needs. The other five were run under the
earlier grid scripts, whose summaries (kin2/kin3/mech3/mech4/mech5/clan/asym _summary csvs)
record auto_mean, Y_mean and a success percentage but NO expansion duration: there is no
end_gen or ext_gen column anywhere in them. Sweep duration is the y-axis, so those mechanisms
cannot be placed on these axes at all. Rather than substitute some other quantity onto y -- which
would silently make five panels mean something different from the other twenty -- the frames are
drawn empty, with the acceptance box in place and the words "runs pending", and they will fill
in when those mechanisms are rerun through the Cartesian pipeline.
"""
import datetime as _dt
import glob
import json
import os
import sys

import matplotlib
matplotlib.use("Agg")
import matplotlib.pyplot as plt
import numpy as np
from matplotlib.patches import Rectangle

HERE = os.path.dirname(os.path.abspath(__file__))
CL = os.path.join(HERE, "cluster")

import screen  # noqa: E402
# Deme two-wave cells scored before this moment ran their second wave under the first
# wave's residence rule and are void; figure2c_fountain carries the same constant.
TW_DEME_FIX = _dt.datetime(2026, 9, 13, 0, 0, 0).timestamp()
# `demog` IS DELIBERATELY EXCLUDED. DR: "why does reduced F1 male fertility seem to have so
# many [more] points than the others?" Because of this set. `demog` crosses three of the
# mechanisms -- sY, xdrive, theta -- and dmi_mtY with TWO EXTRA AXES that no other architecture
# is run over: the residence rule and the five-rung mtDNA-selection ladder. It exists so that
# the four demographic scenarios of Figure 2b sit inside the same Cartesian search as the
# fountains, which is a different question from the one these panels ask.
#
# Including it made mtDNA x Y (5,706 points), X-linked drive (5,721) and theta (36,618) look
# five to thirty times denser than architectures run only on the base grid (~1,250). A reader
# reads cloud density as weight of evidence, so that is not a cosmetic problem: it made three
# panels look better supported than their neighbours for a reason that has nothing to do with
# the mechanism. Excluded, every panel is the same grid and the counts are comparable.
# "cult" ADDED 13 Sep. The five cultural mechanisms were screened into out_cult1_{fw} and
# scored into out_cult2_{fw}, but this list -- which is what _scan_all walks -- never learned
# about them, so their five panels drew as empty "runs pending" frames while the data sat on
# disk. A mechanism with no cells looks exactly like a mechanism not yet run.
SETS = ["routes", "dmi", "dmi2", "xdc", "xd2", "tw", "cult"]
GEN_LO, GEN_HI = 100.0, 1000.0
MIRROR = "/tmp/s1"                      # optional fast local copy of the stage-1 dirs
# v4: prefers the stage-2 score over the 100-replicate re-screen over the 10-replicate
# screen, cell by cell -- the best estimate that exists for each cell. See _scan_all.
# THE VERSION NUMBER IS THE INVALIDATION MECHANISM. This cache keys on nothing but the file
# name, so a v2 cache written before the re-screen landed would be reused verbatim and the new
# data silently ignored -- the same class of failure as the figure reading two of six output
# directories. Bump the suffix whenever what gets scanned changes.
CACHE_FILE = "/tmp/thumb_cache_v4.json"   # shared with dmi_thumbnails.py; same scan, same format

# (label, mechanism key, val or None for "any val", source)
# source "sim"     -> stage-1 cells, plotted
# source "pending" -> empty frame, see the module docstring
# EVERY MECHANISM-VALUE COMBINATION GETS ITS OWN PANEL. DR: "For all the ones where there is
# a series of parameters, show the range (so for the points plotted in Figure 2b, all should be
# represented as thumbnails)."
#
# So the panel list is GENERATED from the ladders rather than hand-picked. A mechanism with one
# setting gets one panel; a mechanism swept over a series gets one panel per rung, in ladder
# order, so the reader can watch a route strengthen or fail as its parameter moves. All four
# routes of Figure 2b are here: additive selection on the Y (Scenario 1), sex distortion and the
# incompatibilities (Scenario 2), reduced F1-male fertility (Scenario 3) and the two-wave expansion
# (Scenario 4).
#
# d is the probability that an F1 male's Y-bearing sperm are inviable. Half of conceptions are
# Y-bearing to begin with, so his offspring are (1 + d)/2 daughters, and the panels are titled
# in daughters rather than in d -- DR: "don't use 'd', use percentage of daughters."
def _daughters(d):
    return f"{(1.0 + d) / 2 * 100:g}% daughters"


NCOL = 7            # see main(): chosen so the figure fits a Letter text block
BANNER_H = 0.30     # banner row height, as a fraction of a panel row
TARGET_ASPECT = 9.0 / 6.5     # height / width of a 6.5 x 9.0 in text block
# PANEL ASPECT, FROZEN AT THE APPROVED VALUE. This is exactly the height/width the old
# "solve PANEL_H so the figure lands on TARGET_ASPECT" line produced for the ten-and-a-half
# rows of the previous layout: 18.9 * (9/6.5) / 10.5 / 2.7 = 12/13. It is a constant now
# because TARGET_ASPECT is a CEILING rather than a target -- see main().
#
# NOTE that PANEL_ASPECT is the ROW PITCH, panel plus the gap below it, not the aspect of the
# drawn axes rectangle; hspace decides how the pitch is split between the two.
PANEL_ASPECT = 12.0 / 13.0
HSPACE_MIN = 0.55   # the inter-row gap of the approved layout; only ever increased, see main()
# THE APPROVED PAGE SHAPE, HELD FIXED WHILE ROWS COME AND GO -- NEW. 9.2 is sum(heights) for
# the layout DR signed off on: four banner rows at BANNER_H plus eight panel rows. Dropping
# four Scenario 2b thumbnails took that block from three rows to two, and with the page height
# left as PANEL_H * sum(heights) the figure would simply have come out shorter (aspect 1.08)
# with the panels in exactly the same tight grid. DR: "this will allow you to space out the
# figure vertically a bit more keeping the same aspect ratio" -- so the page keeps this shape
# and the row that was saved is spent on the gaps between rows instead. See main().
APPROVED_ASPECT = PANEL_ASPECT * 9.2 / NCOL      # = 1.2132 -> 7.89 in tall at 6.5 in wide
# Cells that span two rows of their block so they can be centred between them. The value is
# how many COLUMNS the cell spans: 1 for mtDNA x Y, which sits in column 0 of Scenario 2a, and 2
# for Scenario 4's single panel, which is centred in the two columns Scenario 3's five leave free.
SPAN = {"sim_span": 1, "rerun_span": 2}
# Ink clearance, in inches, left between a span panel's outermost tick label and the ink of
# whatever panel is to its right. See the deferred-geometry pass in main().
CLEAR_IN = 0.10


def _rows(items, n):
    """Chunk a flat list into rows of width n, padding the last row with blanks."""
    out = []
    for i in range(0, len(items), n):
        row = items[i:i + n]
        out.append(row + [None] * (n - len(row)))
    return out


def _build_groups():
    """The panels, grouped by route, each group an explicit GRID OF ROWS.

    DR: "Organize the thumbnails into categories, and indicating which are Scenario 1/2/3/4", and
    then "in Scenario 2a, I suggest removing the 50% and 90% daughter numbers, and stacking the no
    cost [and] 10% cost on top of each other in two rows".

    A group is a list of rows and a row is a list of panels, with None for a deliberate blank.
    That is what lets the sex-distortion block put the cost-free arm directly above the 10%-cost
    arm IN THE SAME COLUMN, so the cost is read by looking down rather than by hunting for a
    matching title. The two arms do not cover the same rungs -- the cost arm was only ever run
    at 55-90% daughters -- so the columns without a cost run are left blank rather than filled
    with a neighbouring value.

    Scenario numbers follow the note's headings: 1 = reduced fitness of the archaic Y, 2 =
    epistatic interaction with the archaic Y, 3 = a cultural mechanism, 4 = two waves. Within
    Scenario 2 the split between mechanisms acting on the Y itself and mechanisms acting through a
    partner locus is the figure's main point, so they are separate blocks.
    """
    def sx(d, cost=False):
        mech = "xdrive_c10" if cost else "xdrive"
        tail = "\n10% fertility cost" if cost else "\nno fertility cost"
        return (f"sex distortion, {_daughters(d)}{tail}", mech, d, "sim")

    G = []
    # SCENARIO 1 IS STATED WITHOUT NAMING THE Y. DR: "Scenario 1 thumbnails should not specify Y.
    # Instead they should be titled things like 'Sex bias and selection on the other
    # unilineally inherited marker'." The engine runs the matrilocal case, in which the marker
    # under selection is the Y -- but the model is symmetric under exchanging the sexes, so the
    # same runs describe a patrilocal expansion with selection on mtDNA. Naming the Y in the
    # title asserts a residence rule the result does not depend on.
    G.append(("SCENARIO 1   Sex bias and selection on the other unilineally inherited marker",
              # five panels in a seven-wide grid leaves two columns spare, which is where
              # the key goes -- DR: "the legend could go to the right of the top row".
              [[(f"selection on the other marker,\ns = {v * 100:g}%", "sY", v, "sim")
                for v in (0.01, 0.02, 0.05, 0.1, 0.2)]
               + [("__key__", None, None, "key_wide"), None]]))

    # SCENARIO 2a: mtDNA x Y and sex distortion back in ONE block under the pre-split title.
    # DR: "Scenario 2a and 2b should be merged back into one section with the same title we had
    # before, even if mtDNAxY is orphaned on one line." The orphan line then has five columns
    # of empty space, which is the natural home for the colour key -- so the reader meets the
    # red/blue convention on the first row that uses it rather than hunting in the caption.
    DAU = (0.05, 0.1, 0.2, 0.4, 0.6, 1.0)          # 52.5, 55, 60, 70, 80, 100% daughters
    G.append(("SCENARIO 2a   Matrilocality and incompatibility acting DIRECTLY ON THE Y — no escape "
              "route, because the Y itself cannot be replaced",
              # A ROW SAVED. DR: "You can save a row by moving the mtDNA x Y below to the
              # next row (next to sex distortion)... It could even be halfway between the next
              # two rows." The sex-distortion ladder is six wide in a seven-wide grid, so the
              # last column is free in both rows; mtDNA x Y takes it and is centred vertically
              # across the pair. It sits with the other mechanism that acts on the Y itself,
              # which is where it belongs, and the block drops from three rows to two.
              # mtDNA x Y sits in the FIRST column, one panel tall, centred between the two
              # sex-distortion rows. DR: "Should be on the left. Make it the same size as the
              # others and centered half-way between the rows." The two arms of the ladder
              # then occupy columns 2-7 in matched pairs, so the cost is still read by looking
              # straight down.
              [[("mtDNA \u00d7 Y", "dmi_mtY", 1.0, "sim_span")] + [sx(d) for d in DAU],
               [None] + [sx(d, cost=True) for d in DAU]]))

    partner = [(lab, mech, 1.0, "sim") for lab, mech in [
        ("mtDNA \u00d7 autosome, dominant", "dmi_mtAutoDom"),
        ("mtDNA \u00d7 autosome, additive", "dmi_mtAutoAdd"),
        ("mtDNA \u00d7 autosome, recessive", "dmi_mtAutoRec"),
        ("mtDNA \u00d7 X, dominant", "dmi_mtXDom"),
        ("X \u00d7 Y", "dmi_XY"),
        ("autosome \u00d7 autosome", "dmi_autoAuto"),
        ("modern X \u00d7 archaic PRDM9-A\n(s = 0.5%)", "dmi_Xprdm9"),
        ("modern PRDM9-A \u00d7 archaic X\n(s = 0)", "dmi_prdm9X_s0"),
        ("modern PRDM9-A \u00d7 archaic X\n(s = 0.5%)", "dmi_prdm9X_s005"),
        ("modern PRDM9-A \u00d7 archaic X\n(s = 5%)", "dmi_prdm9X_s05"),
    ]]
    # ONLY THE 2- AND 3-RESTORER RUNGS -- CHANGED. The ladder used to run n = 1, 2, 3, 4 for
    # both the dominant and the additive restorer, eight panels, which put Scenario 2b at
    # 10 + 8 = 18 thumbnails and so 7 + 7 + 4 across three rows with three quarters of the
    # last row empty. DR: drop "1 restorer, dominant", "4 restorers, dominant",
    # "1 restorer, additive" and "4 restorers, additive". The remaining four rungs bring the
    # block to 14 thumbnails, which packs into exactly 7 + 7 -- a whole row saved, and the
    # saving is handed to the vertical spacing (see APPROVED_ASPECT and main()).
    #
    # The positions are still packed by _rows() rather than listed by hand, so the next edit to
    # this list reflows on its own.
    rest = [(f"{n} restorer{'s' if n > 1 else ''}, {word}", mech, float(n), "sim")
            for mech, word in (("restorers", "dominant"), ("restorersAdd", "additive"))
            for n in (2, 3)]
    flat = partner + rest
    G.append(("SCENARIO 2b   Matrilocality and incompatibility acting THROUGH A PARTNER LOCUS — "
              "introgression replaces the archaic partner and the barrier lapses",
              _rows(flat, NCOL)))

    # THETA IS PRINTED AS A PER CENT -- CHANGED 14 Sep. DR spotted the unit clash: "in
    # Thumbnails Scenario 4, it looks like we get reasonable probability of Y retention for
    # theta=0.02, but the Figure 2b fountains say otherwise. Is the theta unit a percent or a
    # proportion?" It is a PROPORTION in the engine -- the relative fertility weight an F1
    # male carries in the father draw, 1 = full fertility, 0 = sterile -- and the two figures
    # agree on the numbers (at theta = 0.02, mean retention over admissible cells is 0.379
    # deme and 0.224 band; the panels print 38% and 22%). What clashed was the LABELLING:
    # Figure 2b's axis now reads "% offspring raised" with ticks multiplied by 100, so the
    # same cell appeared as "2" there and "0.02" here. The panels now print per cent too, and
    # keep the theta symbol so the tie to the glossary and to the note's "0.2%" survives.
    cult = [(f"reduced F1-male fertility,\n\u03b8 = {th * 100:g}%", "theta", th, "sim")
            for th in (0.0, 0.02, 0.05, 0.1, 0.2)]
    # val=None pools every rung of each mechanism, which points_for already supports; the
    # panel is one per mechanism, not one per rung (asym 0.04-0.32, inbreed 1-2, obligate
    # 0.05-0.4, rhoreturn 0.25-1, skew 0.2-0.8). These drew as "runs pending" only because
    # `cult` was missing from SETS above, not because the runs were missing.
    cult += [(lab, mech, None, "sim") for lab, mech in
             (("inbreeding avoidance (kin ban)", "inbreed"),
              ("obligate dispersal", "obligate"),
              ("return to father's band", "rhoreturn"),
              ("descent marking", "skew"),
              ("asymmetric interbreeding", "asym"))]
    # SCENARIO 3 IS FIVE COLUMNS WIDE, NOT SEVEN, AND SCENARIO 4 RIDES BESIDE IT. DR: "save about
    # 1.5 rows of vertical space", so that the key lands on the same page. Previously Scenario 3
    # was 7 + 3 across two rows and Scenario 4 had a third panel row and a second banner all to
    # itself: 3 panel rows + 2 banner rows. Two rows of FIVE leave columns 5-6 free in both
    # rows, which is exactly where Scenario 4's one remaining panel goes, so the two blocks share
    # ONE banner row: 2 panel rows + 1 banner row, i.e. 1.3 panel rows less.
    #
    # Five columns is also the honest split of Scenario 3's ten panels: the five theta rungs,
    # which are simulated, end up on the first row and the five cultural rules that have no
    # duration recorded yet end up together on the second, instead of the boundary falling
    # mid-row as it did at seven wide.
    r3 = _rows(cult, 5)

    # SCENARIO 4 IS BLANKED, NOT PLOTTED, AND THIS MARKER KEEPS GETTING LOST IN REFACTORS --
    # it has now been reinstated twice. Every two-wave cell on disk terminated before the
    # second wave was founded (spatial_sim2's stop_after_ext had no wave2 guard), so those
    # numbers are single-wave runs mislabelled. Plotting them asserts something false: the
    # route in fact retains both markers in 100% of replicates once the guard is in place.
    # Restore "sim" only after the tw set has been rerun with the guarded engine.
    #
    # ONE PANEL, NOT THREE, AND NO GENERATION IN THE TITLE. DR: keep the gen-1,000 run only.
    # The three second-wave timings were three copies of one empty frame -- all three are
    # "rerunning" for the reason above -- so the ladder showed the reader nothing that the
    # single frame does not, and gen 1,000 is the timing Figure 2b plots. The generation is
    # dropped from the title because with one panel there is no ladder to place it on.
    #
    # "rerun_span" rather than "rerun": a two-row, two-column cell, shrunk back to one panel's
    # size and centred in it by the deferred pass in main(), so the panel sits in the middle of
    # the space Scenario 3 leaves free rather than being stretched over it or shoved against
    # Scenario 3's last column.
    # 13 Sep: the rerun is in. Every two-wave cell used to be a mislabelled single wave --
    # robust_run's stage-2 loop reimplements the generation loop and omitted the wave-2 block
    # entirely -- so this panel was deliberately blanked, by mech=None AND by the "rerun_span"
    # source tag. out_tw2_deme now gives joint = 1.000 across all 173 rescored cells, so both
    # go: the mechanism is named so the lookup finds its cells, and the tag becomes "sim".
    # NOTE the tag is changed HERE ONLY. "rerun_span" also appears in the layout code, where
    # it drives the shrink-and-centre pass for a spanned cell; replacing it globally mangles
    # the block layout, which is exactly what happened on the first attempt.
    r3[0] = r3[0] + [("two-wave expansion", "twowave", None, "sim"), None]
    r3[1] = r3[1] + [None, None]

    # TWO BANNERS, ONE BANNER ROW. A block whose title is a LIST carries one
    # (title, first column, last column + 1) per banner, so each pale blue bar covers only the
    # columns of the block it names -- Scenario 3 over columns 0-4, Scenario 4 over columns 5-6 --
    # rather than one bar running the width of the figure over both.
    #
    # SCENARIO 4'S TITLE IS SHORT SO IT CAN BE SET AT THE SAME SIZE AS EVERY OTHER BANNER --
    # CHANGED. It used to read "A matrilocal expansion preceded or followed by a patrilocal
    # one", which needs 7.3 in at 12.5 pt and its bar is only two columns, 4.79 in, wide. The
    # per-banner auto-fit therefore stepped it down to about 9.5 pt while the full-width
    # banners stayed at 12.5 pt, and Scenario 4's section header looked like a different kind of
    # heading from all the others. Shortened to 4.60 in of text, it fits its own bar at the
    # common size; the ordering ("preceded or followed") is deliberately dropped rather than
    # abbreviated, because the route is symmetric in which wave comes first and the caption is
    # where that belongs.
    G.append(([("SCENARIO 3   A cultural mechanism removing the offspring of the archaic "
                "parent's sex", 0, 5),
               ("SCENARIO 4   Two waves, matrilineal and patrilineal", 5, NCOL)],
              r3))
    return G


GROUPS = _build_groups()
# The output stem, as a module constant so that a caller can render the same figure under a
# different name and a different block order without copying main(). figED_thumbnails.py is
# that caller: it reorders GROUPS to the Extended Data route numbering and sets OUT_BASE.
OUT_BASE = "figS3_4_thumbnails"
# DR, 8 Oct: drop the running head. It says what the colours and the bold numbers mean,
# which is caption material. (figED_thumbnails.py already set this to None for the Extended
# Data version; now both figures agree.)
HEADING = None
PANELS = [pn for _, rows in GROUPS for row in rows for pn in row if pn]
# TWO SHADES PER FRAMEWORK. DR: "color the points that have >=50% mtDNAxY fixation rate
# darker (dark red and dark blue) than those at <50% (pale red and pale blue)". This puts the
# result on the points themselves rather than only in the corner number, so a panel where the
# admissible region is full of pale points reads as a failure at a glance even though it has
# plenty of density in the box -- which is exactly the case for the partner-locus
# architectures, and the single most important thing this figure has to convey.
FWCOL = {"deme": "#1f4e79", "band": "#c0504d"}
FWDARK = {"deme": "#0d2d4d", "band": "#8c1d18"}
FWPALE = {"deme": "#a9c8e6", "band": "#f0b4ae"}

# --alt: PURPLE AND ORANGE INSTEAD OF BLUE AND RED. DR, 2 Oct 2026, for the alternative
# Figure 2, where panel a keeps its red/blue squares (red = modern, blue = archaic) and
# panel b now carries the framework in colour. Reusing red and blue there would hand the
# reader two incompatible keys on one page, so the frameworks move to a pair that appears
# nowhere else in the figure -- and this file gets the same pair, so the thumbnails and
# Figure 2b agree.
#
# Purple #6a51a3 and orange #d95f02 are chosen to survive red-green colour blindness: they
# differ in hue along the blue-yellow axis, which both deuteranopia and protanopia preserve,
# AND in lightness, so they stay distinct in greyscale and to a monochromat. A red/green or
# red/blue pair fails the first test; a teal/green pair fails the second.
ALT = "--alt" in sys.argv
if ALT:
    FWCOL = {"deme": "#6a51a3", "band": "#d95f02"}
    FWDARK = {"deme": "#3f007d", "band": "#8c3a00"}
    FWPALE = {"deme": "#bcbddc", "band": "#fdcc9a"}
    # AND THE HEADING, which names the colours in words. The first --alt render came out
    # purple and orange under a banner that still read "blue the deme model, red the band
    # model" -- the same failure as the label keys in figure_pulse.py: one fact (which
    # colour means which framework) written down in two places, so changing one leaves the
    # other quietly wrong. Reassigned here, next to the palette it describes.
    HEADING = None
FIXCUT = 0.50
FWMARK = {"deme": "o", "band": "o"}


def flag(name, cast, default):
    return cast(sys.argv[sys.argv.index(name) + 1]) if name in sys.argv else default


def load(mech, val):
    """{framework: [(autosomal %, duration, establishment, retention)]} for every cell.

    REWRITTEN 14 Sep 2026 TO READ screen.py, which is now the single definition of what a cell
    is and whether it is admissible. This function used to walk the output directories itself,
    with its own precedence rule between the ten-replicate screen and the hundred-replicate
    re-screen, its own cache, and its own copy of the two-wave genetic-clock exception --
    while figure2c_fountain did the same job differently. The two disagreed on 1,911 of the
    10,399 points Figure 2b plots, because this figure positioned a cell by its stage-1
    autosomal mean and that one judged it by its stage-2 one.

    Each cell now arrives at the best estimate that exists for it, so the coordinate a point is
    DRAWN at is the coordinate it was JUDGED on, and every fountain point in Figure 2b can be
    found inside the box drawn here.

    RETENTION prefers the scored p_joint over the fraction of screen replicates that fixed both
    markers. They measure the same thing -- the note's own definition says p_joint "equals
    P(both fixed) whenever both markers are absorbed", and runs stop at absorption -- but the
    scored one rests on 100-200 replicates against the screen's 10.
    """
    d = screen.by_key(mech, val)
    out = {}
    for fw in ("deme", "band"):
        out[fw] = [(c["aut"], c["gen"], (c.get("est") if c.get("est") is not None else 1.0),
                    (c["joint"] if c.get("joint") is not None else c.get("both")))
                   for c in d[fw]]
    return out


def frame(ax, label, aut_lo, aut_hi, show_ylabel=True):
    """The axes, the grey acceptance box and the title -- everything but the data."""
    # On linear axes the acceptance region is a sliver a few percent wide near the origin. That
    # IS the result -- the data is a very small target -- so the axes are logarithmic and the box
    # is drawn at true scale.
    #
    # UNFILLED, BLACK, AND ON TOP. DR: "the gray triangles in the thumbnails end up covered with
    # points. Maybe indicate the box just with a black rectangle." A filled patch underneath the
    # data loses the one contest the panel exists to settle -- whether there is density inside
    # the box -- because the points that land there are exactly what hides it. An open outline
    # drawn above everything marks the region without competing with it.
    ax.add_patch(Rectangle((aut_lo, GEN_LO), aut_hi - aut_lo, GEN_HI - GEN_LO,
                           facecolor="none", edgecolor="black", lw=1.3, zorder=6))
    ax.set_xscale("log"); ax.set_yscale("log")
    ax.set_xlim(0.8, 100); ax.set_ylim(40, 9000)
    ax.set_xticks([1, 2, 5, 10, 20, 50, 100])
    ax.set_xticklabels(["1%", "2%", "5%", "10%", "20%", "50%", "100%"], fontsize=7.5)
    ax.set_yticks([50, 100, 200, 500, 1000, 2000, 5000])
    ax.set_yticklabels(["50", "100", "200", "500", "1,000", "2,000", "5,000"], fontsize=7)
    ax.set_xticks([], minor=True); ax.set_yticks([], minor=True)
    ax.set_title(label, fontsize=8.4, fontweight="bold")
    ax.grid(color="0.95", lw=0.6); ax.set_axisbelow(True)
    if show_ylabel:
        ax.set_ylabel("expansion duration (gens)", fontsize=8)


def panel(ax, label, mech, val, aut_lo, aut_hi, show_ylabel=True):
    data = load(mech, val)
    tot = sum(len(v) for v in data.values())
    frame(ax, label, aut_lo, aut_hi, show_ylabel)
    if tot == 0:
        ax.text(0.5, 0.5, "not searched", transform=ax.transAxes, ha="center", va="center",
                fontsize=8, color="0.5", style="italic")
        return 0, None, 0, None
    n_in = 0
    inbox = {"deme": [], "band": []}
    for fw in ("deme", "band"):
        v = data[fw]
        if not v:
            continue
        a = np.array([x[0] for x in v])
        g = np.clip(np.array([x[1] for x in v]), 45, 8800)
        e = np.array([x[2] for x in v])
        b = np.array([x[3] if len(x) > 3 else np.nan for x in v])
        # ADMISSIBLE = inside the box AND establishing more than half the time: all three
        # acceptance criteria, applied to every screened combination.
        ok = ((a >= aut_lo) & (a <= aut_hi) & (g >= GEN_LO) & (g <= GEN_HI) & (e > 0.5))
        n_in += int(ok.sum())
        inbox[fw] = b[ok & ~np.isnan(b)].tolist()
        # Dark where this combination retains BOTH markers at least half the time, pale where
        # it does not. Cells with no per-replicate rows recorded count as pale rather than
        # being dropped, so the point count still matches the screen.
        hi = (b >= FIXCUT) & ~np.isnan(b)
        ax.scatter(a[~hi], g[~hi], s=5, marker=FWMARK[fw], c=[FWPALE[fw]], alpha=0.75,
                   edgecolors="none", zorder=3)
        ax.scatter(a[hi], g[hi], s=6, marker=FWMARK[fw], c=[FWDARK[fw]], alpha=0.85,
                   edgecolors="none", zorder=5)
    est = float(np.mean([x[2] for v in data.values() for x in v]))

    # ONE NUMBER PER FRAMEWORK, IN THAT FRAMEWORK'S COLOUR. DR: "instead of a single number in
    # the top left, you have two numbers in blue and red corresponding to the band and deme
    # results... maybe one in the top left (red) and one in the bottom left (blue)".
    #
    # Band red at the top, deme blue at the bottom, which is also where each framework's points
    # tend to sit -- the band cloud runs long and the deme cloud short, so the numbers land near
    # their own data. Pooling them hid a real disagreement: the two frameworks differ by more
    # than sampling noise on several architectures, and a reader is entitled to see that rather
    # than an average of the two.
    #
    # Averaged over the ADMISSIBLE combinations of that framework only, and printed as FAIL
    # where that framework has none, since then there is no population of runs to average over.
    def corner(fw, y, va):
        v = inbox[fw]
        txt = "FAIL" if not v else f"{float(np.mean(v)):.0%}"
        ax.text(0.04, y, txt, transform=ax.transAxes, ha="left", va=va,
                fontsize=15, fontweight="bold", color=FWDARK[fw])
        return None if not v else float(np.mean(v))

    joint_band = corner("band", 0.96, "top")
    joint_deme = corner("deme", 0.05, "bottom")
    joint = (None if not (inbox["deme"] + inbox["band"])
             else float(np.mean(inbox["deme"] + inbox["band"])))
    return tot, est, n_in, joint


def key_panel(ax):
    """The colour key, drawn into the empty space beside the orphaned mtDNA x Y panel.

    DR: "Somewhere, maybe in the orphan line containing mtDNA x Y, add a key indicating what
    red and blue mean." Putting it inside the grid rather than in the caption means the reader
    meets the convention at the moment they first need it.
    """
    ax.axis("off")
    ax.set_xlim(0, 1); ax.set_ylim(0, 1)
    rows = [
        (FWDARK["band"], "band model", "retains both markers in \u2265 50% of replicates"),
        (FWPALE["band"], None, "retains both in < 50%"),
        (FWDARK["deme"], "deme model", "retains both markers in \u2265 50% of replicates"),
        (FWPALE["deme"], None, "retains both in < 50%"),
    ]
    y = 0.90
    for colour, head, sub in rows:
        ax.plot([0.06], [y], marker="o", ms=7, color=colour, clip_on=False)
        if head:
            ax.text(0.13, y, head, va="center", ha="left", fontsize=10.5, fontweight="bold",
                    color=colour)
            ax.text(0.13, y - 0.085, sub, va="center", ha="left", fontsize=9, color="0.35")
            y -= 0.20
        else:
            ax.text(0.13, y, sub, va="center", ha="left", fontsize=9, color="0.35")
            y -= 0.14
    ax.text(0.06, 0.13, "bold percentages: how often both mtDNA and the Y\nend up modern, "
                        "across admissible combinations",
            va="center", ha="left", fontsize=9, color="0.25")
    ax.add_patch(Rectangle((0.02, 0.03), 0.96, 0.95, transform=ax.transAxes, facecolor="none",
                           edgecolor="0.75", lw=0.8, ls=(0, (3, 3))))


def rerun_panel(ax, label, aut_lo, aut_hi, show_ylabel=True):
    """Frame only: the cells on disk are invalid and are being recomputed."""
    frame(ax, label, aut_lo, aut_hi, show_ylabel)
    ax.text(0.5, 0.5, "rerunning", transform=ax.transAxes, ha="center", va="center",
            fontsize=9, color="#b03030", style="italic", fontweight="bold")


def pending_panel(ax, label, aut_lo, aut_hi, show_ylabel=True):
    """Same axes, same box, no points: the mechanism has no sweep-duration record yet."""
    frame(ax, label, aut_lo, aut_hi, show_ylabel)
    ax.text(0.5, 0.5, "runs pending", transform=ax.transAxes, ha="center", va="center",
            fontsize=8, color="0.5", style="italic")


def main():
    # 4-6%, matching Figure 2. These thumbnails are that figure's visual companion, so the box
    # they draw has to be the box it uses.
    aut_lo, aut_hi = flag("--aut-lo", float, 3.0), flag("--aut-hi", float, 7.0)

    ncol = NCOL
    # One block per category, each with its own banner row, and each block an explicit grid of
    # rows so that panels meant to be compared sit in the same column.
    blocks = [(title, rows) for title, rows in GROUPS]
    heights = []
    for _, rows in blocks:
        heights.append(BANNER_H)              # the banner
        heights.extend([1.0] * len(rows))

    # SIZED TO A LETTER PAGE. DR: "see if you can change the dimension of the thumbnail plot
    # maybe by adding another column to see if you can get the aspect ratio proper on a Letter
    # Sized page." A Letter text block at 1 in margins is 6.5 x 9.0, so the figure has to come
    # out at height/width = 1.385 or Word scales it down to fit the narrower dimension and
    # wastes the rest of the page. Seven columns rather than six is most of the way there --
    # it does not change the ROW count, because the row structure is set by the blocks, so it
    # is pure width.
    #
    # TARGET_ASPECT IS A CEILING, NOT A TARGET -- CHANGED. PANEL_H used to be solved as
    # fig_w * TARGET_ASPECT / sum(heights), which pinned the figure at exactly 9.0 in tall
    # whatever the block list contained. That made saved vertical space invisible: dropping a
    # row just made every panel taller and the figure came out the same height, which is the
    # opposite of what dropping a row is for. 1.385 matters only as a LIMIT (a Letter text
    # block is 6.5 x 9.0; a taller figure is scaled down to fit the width and the rest of the
    # page is wasted), so the panel keeps the aspect it had when the layout was approved and
    # the figure is only squeezed if the block list ever grows tall enough to break the limit.
    # TOP AND BOTTOM MARGINS IN INCHES, NOT IN FIGURE FRACTIONS -- CHANGED with the ceiling
    # above. Both reserve room for TEXT of a fixed point size (the running head at the top,
    # the bottom row's x-axis label below the last panels), so the reservation cannot scale
    # with the figure: the old fractions 0.985 / 0.012 / y=0.998 were tuned at the 26.17 in
    # height the solved layout always produced. These are those same three gaps in inches --
    # except the bottom one, which is 0.046 in bigger: at the old 0.012 the bottom row's
    # "modern autosomal ancestry" was cut off 0.04 in short of its baseline, in this layout
    # and in every previous one. 0.36 in is the measured depth of a bottom-row panel's tick
    # labels plus its x-axis label, so nothing hangs off the page.
    #
    # 0.3750, NOT 0.3600 -- CHANGED. At 0.36 the x-axis label fitted to within a pixel or two
    # and the DESCENDERS still did not: the tail of the "y" in "ancestry" reached the last
    # scanline of the canvas and its tip was sliced off, in this layout and in every earlier
    # one (0.36 was measured from a cap-height bbox, which does not include descenders).
    # 0.015 in is four pixels at 300 dpi, which clears the deepest tail with room to spare and
    # costs nothing but four pixels of the inter-row gap solved below.
    #
    # They are set HERE, above the figure, rather than beside subplots_adjust where they used
    # to be, because the hspace solved just below has to subtract them -- they are the part of
    # the page height that is not gridspec.
    TOP_PAD_IN, BOT_PAD_IN, HEAD_PAD_IN = 0.3925, 0.3750, 0.0523

    PANEL_W = 2.7
    fig_w = PANEL_W * ncol
    nrows, sum_h = len(heights), sum(heights)
    PANEL_H = min(PANEL_W * PANEL_ASPECT, fig_w * TARGET_ASPECT / sum_h)

    # THE SAVED ROW BECOMES WHITE SPACE, NOT A SHORTER PAGE -- NEW. fig_h used to be simply
    # PANEL_H * sum(heights), so losing a row lost 2.49 in of page and left the panels packed
    # exactly as tightly as before. The page now holds APPROVED_ASPECT (see the constant) --
    # floored, so a future block that pushes the natural height past it is not stretched, and
    # ceilinged at TARGET_ASPECT, so the figure can never outgrow a Letter text block.
    fig_h = min(max(PANEL_H * sum_h, fig_w * APPROVED_ASPECT), fig_w * TARGET_ASPECT)

    # ... AND hspace IS SOLVED SO THAT ALL OF THE EXTRA HEIGHT GOES INTO THE GAPS. matplotlib
    # gives a height-ratio-1.0 row the drawn height
    #     (fig_h - pads) * nrows / ((nrows + hspace * (nrows - 1)) * sum_h),
    # so `ink_h` below is the panel height the old formula would have produced at HSPACE_MIN,
    # i.e. the size the panels are today, and hspace is then inverted out of the same
    # expression at the taller fig_h. The panels therefore keep their approved size and their
    # approved aspect and only the space between rows grows. max() because hspace is only ever
    # relaxed: if a future layout is naturally taller than APPROVED_ASPECT there is nothing to
    # spend and the gap stays at the value the approved figure used.
    ink_h = ((PANEL_H * sum_h - TOP_PAD_IN - BOT_PAD_IN) * nrows
             / ((nrows + HSPACE_MIN * (nrows - 1)) * sum_h))
    hspace = max(HSPACE_MIN,
                 ((fig_h - TOP_PAD_IN - BOT_PAD_IN) * nrows / (ink_h * sum_h) - nrows)
                 / (nrows - 1))

    fig = plt.figure(figsize=(fig_w, fig_h))
    gs = fig.add_gridspec(len(heights), ncol, height_ratios=heights, hspace=hspace,
                          wspace=0.22)

    report, r, deferred, ref_h, banners = [], 0, [], {}, []
    for title, rows in blocks:
        # One banner per block normally, but Scenario 3 and Scenario 4 share a banner ROW, so a
        # block's title may be a list of (title, first column, last column + 1).
        for btitle, bc0, bc1 in (title if isinstance(title, list) else [(title, 0, ncol)]):
            ban = fig.add_subplot(gs[r, bc0:bc1])
            ban.axis("off")
            ban.add_patch(Rectangle((0, 0.04), 1, 0.92, transform=ban.transAxes,
                                    facecolor="#e8eef5", edgecolor="#9bb3cc", lw=0.8))
            banners.append((ban, ban.text(0.008, 0.5, btitle, transform=ban.transAxes,
                                          ha="left", va="center", fontsize=12.5,
                                          fontweight="bold", color="#16344f")))
        r += 1
        for ri, row in enumerate(rows):
            # With a spanning cell in column 0, the leftmost ORDINARY panel of a row is not
            # at ci = 0, and that is the one that needs the y-axis label.
            lead = next((k for k, c in enumerate(row)
                         if c is not None and c[3] not in set(SPAN) | {"key", "key_wide"}), 0)
            for ci, pn in enumerate(row):
                if pn is None:
                    continue
                lab, mech, val, src = pn
                # A cell may span the two rows of its block and sit centred between them, and
                # one or two columns with it. Both are laid out here and the spanning ones are
                # nudged into place after the figure geometry is final -- see `deferred` below.
                if src in SPAN:
                    ax = fig.add_subplot(gs[r + ri:r + ri + 2, ci:ci + SPAN[src]])
                    deferred.append((ax, r + ri))
                elif src == "key_wide":
                    ax = fig.add_subplot(gs[r + ri, ci:ci + 2])
                else:
                    ax = fig.add_subplot(gs[r + ri, ci])
                    ref_h.setdefault(r + ri, ax)
                if src in ("key", "key_wide"):
                    key_panel(ax)
                    continue
                if src == "sim_span":
                    tot, est, n_in, joint = panel(ax, lab, mech, val, aut_lo, aut_hi,
                                                  show_ylabel=True)
                    report.append((lab, tot, est, n_in, joint))
                    ax.set_xlabel("modern autosomal ancestry", fontsize=8)
                    continue
                if src == "rerun_span":
                    # Scenario 4's one panel. It is the only panel of its block, so it carries
                    # both axis labels itself rather than borrowing the leftmost panel's:
                    # `lead` is column 0, which belongs to Scenario 3.
                    rerun_panel(ax, lab, aut_lo, aut_hi, show_ylabel=True)
                    report.append((lab, None, None, None, None))
                    ax.set_xlabel("modern autosomal ancestry", fontsize=8)
                    continue
                if src == "rerun":
                    rerun_panel(ax, lab, aut_lo, aut_hi, show_ylabel=(ci == 0))
                    report.append((lab, None, None, None, None))
                    continue
                if src == "pending":
                    pending_panel(ax, lab, aut_lo, aut_hi, show_ylabel=(ci == lead))
                    report.append((lab, None, None, None, None))
                else:
                    tot, est, n_in, joint = panel(ax, lab, mech, val, aut_lo, aut_hi,
                                                  show_ylabel=(ci == lead))
                    report.append((lab, tot, est, n_in, joint))
                # x-label on the bottom-most occupied panel of this column WITHIN the block, so
                # each block reads on its own and no label lands in the middle of one
                if not any(len(rr) > ci and rr[ci] for rr in rows[ri + 1:]):
                    ax.set_xlabel("modern autosomal ancestry", fontsize=8)
        r += len(rows)

    # NO BAKED-IN CAPTION. An earlier version put the whole explanatory paragraph in a
    # suptitle, where it overflowed both margins at this figure width and could not be edited
    # without regenerating the figure. The caption belongs in the document, where it is part of
    # the tracked text; the figure carries only a one-line running head.
    #
    # TOP_PAD_IN / BOT_PAD_IN / HEAD_PAD_IN are set at the top of main(), because the page
    # height and the solved hspace both need them; see the comment there for why inches.
    # HEADING is a module constant so a caller can drop it: the Extended Data version does,
    # because in that figure the same sentence is in the caption underneath.
    if HEADING:
        fig.suptitle(HEADING, fontsize=12, y=1.0 - HEAD_PAD_IN / fig_h)
    # right 0.990, NOT 0.995 -- CHANGED. The last x-tick label of the rightmost column,
    # "100%", is centred on the right spine and so overhangs it by half a label; at 0.995 its
    # final "%" ran 0.05 in past the right edge of the canvas and was sliced off in every
    # version of this figure. 0.990 buys 0.09 in, which clears it. It costs 0.01 in of panel
    # width per column and nothing at all in height, so the page shape is untouched.
    fig.subplots_adjust(left=0.045, right=0.990,
                        top=1.0 - TOP_PAD_IN / fig_h, bottom=BOT_PAD_IN / fig_h)

    # AFTER subplots_adjust, NOT BEFORE -- it recomputes every subplot's position from the
    # gridspec and silently threw away an earlier version of this fix.
    fig.canvas.draw()
    rend = fig.canvas.get_renderer()
    inv = fig.transFigure.inverted()

    # ONE FONT SIZE FOR ALL BANNERS, FITTED TO THE TIGHTEST OF THEM -- CHANGED. The fit used
    # to be run per banner: each title was stepped down until it fitted its own bar. That is
    # correct as far as overflow goes but it made the SIZES disagree, and because Scenario 4's
    # bar is only two of the seven columns wide its heading came out at about 9.5 pt against
    # 12.5 pt for the full-width ones -- DR read that as a different kind of heading rather
    # than the same kind in a narrower box. So the step-down is now a single shared size:
    # every banner is measured against its own width, the smallest size that satisfies ALL of
    # them wins, and that one size is applied to all. Scenario 4's title has also been shortened
    # (see _build_groups) so that the shared size stays at the full 12.5 pt; this loop is what
    # guarantees the sizes stay equal, and no title overflows, if a title is ever edited long.
    size = banners[0][1].get_fontsize() if banners else 12.5
    for ban, txt in banners:
        avail = ban.get_position().width * fig_w * 0.985      # 0.008 of it is the left inset
        txt.set_fontsize(size)
        while size > 7.0 and txt.get_window_extent(rend).width / fig.dpi > avail:
            size -= 0.25
            txt.set_fontsize(size)
    for _, txt in banners:
        txt.set_fontsize(size)

    # SPANNING CELLS, SIZED FROM A MEASURED SIBLING.
    #
    # HEIGHT: a two-row span is twice a panel's height, which would distort it, and half of a
    # two-row span is not one row either -- the span swallows the hspace between the rows, so
    # halving leaves the panel 1.5x too tall. Measure a genuine single-row panel from the same
    # row and match it exactly, then centre: "halfway between the next two rows".
    #
    # WIDTH: taken from the same sibling and centred in the span, so Scenario 4's two-column cell
    # comes out one panel wide in the middle of the two columns Scenario 3 leaves free instead of
    # being stretched across them. For a one-column span (mtDNA x Y) this is a no-op.
    for ax, row_idx in deferred:
        ref = ref_h.get(row_idx)
        if ref is None:
            continue
        one = ref.get_position()
        pos = ax.get_position()
        x0 = pos.x0 + (pos.width - one.width) / 2.0
        w = one.width

        # RIGHT-HAND CLEARANCE -- FIXES THE mtDNA x Y PANEL RUNNING INTO ITS NEIGHBOURS.
        # Previously the span simply kept its gridspec width, which is one full column, and
        # the mtDNA x Y panel's right border overlapped the panels beside it. The cause is not
        # the spine but the ink either side of it: this panel's last x-tick label ("100%")
        # overhangs its own right spine by half a label width, and the panel to its right --
        # the leftmost sex-distortion panel, which is the first ORDINARY panel of the row and
        # therefore the one that prints the y-axis label -- puts that rotated label about
        # 0.028 of the figure width to the LEFT of its own spine, more than the 0.025
        # inter-column gap itself. So at full column width the two texts were drawn on top of
        # each other. Every other column-0 panel is safe because its right-hand neighbour has
        # no y-axis label.
        #
        # Shrink (never grow) until this panel's ink clears the neighbour's ink by CLEAR_IN,
        # measuring both bboxes rather than hardcoding a width, and keep the panel anchored at
        # its column's left edge so it stays in line with the column-0 panels of the other
        # blocks.
        over = inv.transform_bbox(ax.get_tightbbox(rend)).x1 - pos.x1   # tick-label overhang
        right = [inv.transform_bbox(a.get_tightbbox(rend)).x0
                 for a in (ref_h.get(row_idx), ref_h.get(row_idx + 1))
                 if a is not None and a.get_position().x0 > pos.x0]
        if right:
            w = min(w, min(right) - CLEAR_IN / fig_w - over - x0)

        ax.set_position([x0, pos.y0 + (pos.height - one.height) / 2.0, w, one.height])

    # The suffix is applied HERE rather than to OUT_BASE itself, because figED_thumbnails.py
    # imports this module and overwrites OUT_BASE for the Extended Data ordering; suffixing
    # at save time means --alt works through that caller too and cannot overwrite the
    # published thumbnails either way.
    stem = OUT_BASE + ("_alt" if ALT else "")
    base = os.path.join(HERE, stem)
    fig.savefig(base + ".png", dpi=300)
    fig.savefig(base + ".pdf")
    print(f"  wrote {stem}.png / .pdf")
    for lab, tot, est, n_in, joint in report:
        flat = lab.replace("\n", " ")
        if tot is None:
            print(f"    {flat:<46} runs pending (no duration recorded)")
        elif tot == 0:
            print(f"    {flat:<46} not searched")
        else:
            res = "FAIL" if joint is None else f"{joint:.0%}"
            print(f"    {flat:<46} n={tot:<6d} est={est:.0%}  in box={n_in:<5d} "
                  f"both markers fixed (admissible only) = {res}")


if __name__ == "__main__":
    main()
```
