## Supplementary Information section 3 code base for "Hypothesis: A modern human range expansion ∼300,000 years ago had a critical role in Neandertal origins": figED_expansions.py

```
#!/usr/bin/env python3
"""
figED_expansions.py -- Extended Data: sample expansions for every scenario in Figure 2b.
(14 Sep 2026)

DR: "show sample expansions for all the scenarios shown in the fountain plot (Figure 2). The
page gets divided into two, on the left the deme and on the right the band (with a vertical
line between them). In each row, we have three deme plots for sample expansions at the left
100, 300 and 1000 generations showing square grids with the three stripe squares using the
same convention as in Figure 2b; and three band plots on the right ... at 100, 300 and 1000
generations."

WHAT THIS FIGURE IS FOR. Figure 2b reduces each parameter combination to one number, and
Figure S3.3 reduces it to one point. Both answer "does the route work"; neither shows what
working looks like. A reader who sees p_joint = 0.93 at s = 0.5 has no picture of the
landscape that produces it, and no way to see WHY a route that works in the band model can
fail in the deme model at the same mechanism strength. These are the same simulations, drawn.

THE SQUARE CONVENTION IS FIGURE 2a's, REUSED RATHER THAN REIMPLEMENTED. Each unit -- a deme
on the left, a band on the right -- is three contiguous vertical bars, mtDNA / Y / autosomes,
filled red from the bottom in proportion to how modern that marker is and blue above.
band_square() is imported from figure2_bands, so the two figures cannot drift apart: if the
convention changes there it changes here.

WHAT DIFFERS BETWEEN THE TWO SIDES, AND WHY IT HAS TO. The deme model is a fixed square
lattice, so its panels are a grid with one square per deme at its lattice position. The band
model has bands that move, fission and die on a real map of Europe, so its panels are the
Albers projection with one square per surviving band at its position, sized by band size --
exactly Figure 2a. Forcing either into the other's layout would misrepresent the model. The
vertical rule between them says they are different spaces rather than two halves of one.

EACH ROW IS DRAWN AT ITS OWN FOUNTAIN POINT, NOT AT A COMMON BASELINE. This is the one
decision in the figure that is not cosmetic, and getting it wrong the first time produced a
figure that contradicted Figure 2b: at the band baseline the Route 3 rows ended with the
modern Y at 0% after a thousand generations while the fountain at the same theta reports
p_joint above 0.9. A fountain point is not "the baseline with a mechanism switched on" -- it
is a cell of the contact sweep that lands in the acceptance box, and the contact rate is part
of the scenario. Every row therefore looks up the admissible cell for its own mechanism and
value whose autosomal ancestry is closest to 5%, preferring the note's baseline geometry.
FigureED_expansions_cells.txt, written beside the figure, names the cell behind every panel.

ROWS, as specified, and two notes on them:

    1       baseline, no mechanism            s = 0, 50% daughters, theta = 1
    2-5     Route 1, selection on the Y       s = 0, 0.0005, 0.05, 0.5
    6-9     Route 2, sex distortion           55%, 70%, 90% daughters; mtDNA x Y
    10-13   Route 3, reduced F1 fertility     theta = 0.1, 0.01, 0.001, 0
    14      Route 4, two waves

  ROWS 1 AND 2 ARE THE SAME SIMULATION, so the figure draws 13 rows, not 14. "No mechanism"
  and "Route 1 at s = 0" are the same parameter set at the same seed, and the two rows came
  out pixel-identical, which on the page reads as a printing fault rather than as the point
  being made. Row 1 is labelled as both. `--all-rows` draws the specified 14 if the separate
  row is wanted after all.

  s = 0.0005 IS NOT ON THE SCREENED LADDER, which runs 0.001 as its smallest non-zero rung.
  The row is still run -- this figure simulates rather than reading the screen -- but it has
  no counterpart fountain in Figure 2b, and it borrows its cell from s = 0.001, the nearest
  rung. It is the only row whose panels cannot be matched to a point in the other figure.

  TWO ROWS ARE NOT AT THE BASELINE GEOMETRY, and both are findings rather than accidents.
  The deme model at theta = 0 has exactly one admissible cell in the entire screen and it is
  a 6x6 lattice of 40-person demes; obligate dispersal simply does not fit inside the
  baseline lattice's contact budget. Every admissible band two-wave cell has p_move >= 0.6
  and sigma_b >= 120 against the baseline's 0.45 and 60: the second wave has to overtake a
  landscape that is already full, and only a more mobile, wider-ranging population manages
  it inside a thousand generations. The cell file flags both.

ONE SEED FOR THE WHOLE FIGURE. Every panel uses the same seed, so differences down a column
are the mechanism and not the draw. That is the only way a reader can compare rows at all,
and it is also the figure's main limitation: one replicate is one replicate, and the caption
should say that the quantitative claims live in Figure 2b.

    python figED_expansions.py [--seed 0] [--dpi 200]
"""
import glob
import json
import os
import sys

import matplotlib
matplotlib.use("Agg")
import matplotlib.pyplot as plt
import numpy as np
from matplotlib.lines import Line2D
from matplotlib.patches import Rectangle

HERE = os.path.dirname(os.path.abspath(__file__))
sys.path.insert(0, os.path.join(HERE, "cluster"))
sys.path.insert(0, os.path.join(HERE, "band_model"))

import screen as SC                                                # noqa: E402
import spatial_sim3 as S3                                           # noqa: E402
from spatial_sim3 import Config as DConfig                          # noqa: E402
from dataclasses import replace as _replace                         # noqa: E402
from spatial_sim2 import _init, _migrate, _reseed_origin            # noqa: E402
import bandsim as BS                                                # noqa: E402
from bandsim import Config as BConfig, Landscape, init_state, step  # noqa: E402
from figure2_bands import band_square, per_band, albers, \
    KM_PER_DEG_LAT, COS_REF, BOLD_PT, PLAIN_PT                      # noqa: E402

# DR: "maybe change the time points to 30, 300, 3000". A decade of the expansion, its
# middle, and long after it has finished -- the last column now shows where the landscape
# SETTLES rather than catching several rows mid-sweep.
GENS = [30, 300, 3000]
# The seed search still judges a replicate at generation 1,000, because that is where the
# acceptance window and the fountain's retention probability are defined. Drawing at 3,000
# does not change what the cell claims.
MEASURE_GEN = 1000

# The note's baselines, so a sample expansion is a sample of the thing the rest of the note
# describes rather than of some other landscape. Deme: the Ne-anchored geometry. Band: the
# frozen baseline of Appendix B.
DEME_BASE = dict(grid=12, K=11, m=0.10, gamma=0.12, r=2.0,
                 gens=max(GENS) + 1, sample_gen=max(GENS), record_every=10 ** 9)
BAND_BASE = dict(N_target=1750, K=25, e=0.19, p_move=0.45, r=2.0, beta=0.0,
                 sigma_b=60.0, sigma_e=80.0, alpha_d=0.8, gens=max(GENS) + 1,
                 record_every=10 ** 9)

# (row label, deme mechanism kwargs, band mechanism kwargs, (mech, val) key into the
# screen). The mechanism kwargs are the row's identity; everything else -- grid, K,
# contact rate, dispersal kernel -- comes from the admissible cell that (mech, val)
# resolves to, via row_kwargs(). Empty mechanism dicts mean the cell already carries the
# setting (the two-wave rows) or there is nothing to set (the null row).
# d is the fraction of Y-bearing sperm made inviable, so the daughter proportion is (1+d)/2:
# 55% -> d = 0.1, 70% -> d = 0.4, 90% -> d = 0.8.
ROWS = [
    # NO NULL ROW. DR: "remove the no mechanism row", twice -- first for the row labelled as
    # such and then for the selection row at s = 0, which is the same simulation under
    # another name. The ladder now starts where the mechanism starts.
    #
    # SCENARIO NUMBERING AND ORDER FOLLOW THE NOTE (10 Oct 2026): 1 = Dobzhansky-Muller
    # incompatibilities including X-linked drive, 2 = selection for prestigious patrilines,
    # 3 = two waves, 4 = reduced fertility of F1 offspring of the archaic parent's sex. The
    # labels used to say "Route" in an older numbering (selection first).
    ("Scenario 1\n55% daughters", dict(xdrive=0.1), dict(xdrive=0.1), ("xdrive", 0.1)),
    ("Scenario 1\n90% daughters", dict(xdrive=0.8), dict(xdrive=0.8), ("xdrive", 0.8)),
    ("Scenario 1\nmtDNA × Y", dict(dm=1.0, dm_focal="mt", dm_partner="Y"),
     dict(dm=1.0, dm_focal="mt", dm_partner="Y"), ("dmi_mtY", 1.0)),
    # Selection on the Y is conditioned on group identity (bandsim / spatial_sim2, 9 Oct
    # 2026): the patriline matching the band's or lineage's identity is favoured, the
    # mismatch penalised. These two rows were re-simulated and re-seeded under that rule.
    ("Scenario 2\n$s$ = 0.005", dict(s=0.005, sel_locus="Y"), dict(s=0.005, sel_locus="Y"),
     ("sY", 0.005)),
    ("Scenario 2\n$s$ = 0.5", dict(s=0.5, sel_locus="Y"), dict(s=0.5, sel_locus="Y"),
     ("sY", 0.5)),
    ("Scenario 3\ntwo waves", {}, {}, ("twowave", None)),
    ("Scenario 4\nθ = 0.1", dict(theta_f1m=0.1), dict(theta=0.1), ("theta", 0.1)),
    ("Scenario 4\nθ = 0.01", dict(theta_f1m=0.01), dict(theta=0.01), ("theta", 0.01)),
]


# ---------------------------------------------------------------------------------------
# WHICH CELL EACH ROW IS DRAWN AT
# ---------------------------------------------------------------------------------------
# The first version ran every row at the note's baseline contact rate and let only the
# mechanism vary. That was wrong, and visibly so: at the band baseline the Route 3 rows came
# out with the modern Y at 0% at generation 1,000, while Figure 2b's fountain at the same
# theta reaches p_joint = 0.95. The figure would have contradicted the figure it illustrates.
#
# The reason is that a fountain point is not the baseline with a mechanism switched on. It is
# a cell of the gamma (deme) / e (band) sweep that lands in the acceptance box -- 3-7%
# autosomal, 100-1,000 generations, establishment above 50%. The contact rate is part of the
# scenario, not a fixed background, and at the baseline rate most of these mechanisms are
# being asked to work under a contact budget that no admissible cell uses.
#
# So each row now looks up its own cell: among the admissible cells for that mechanism and
# that value, the one whose autosomal ancestry is closest to 5%, which is the anchor the rest
# of the note calibrates to. Every panel is therefore a draw from a point that Figure 2b
# plots, and the two figures cannot disagree.
AUT_ANCHOR = 5.0
AUT_LO, AUT_HI = SC.AUT_LO, SC.AUT_HI

# THE CANONICAL GEOMETRY IS PREFERRED, AND ALMOST ALWAYS AVAILABLE. The screen sweeps the
# lattice shape and the dispersal kernel as nuisance axes, so "the admissible cell closest to
# 5%" on its own picked a 6x6 lattice of 40-person demes for one row and a 16x16 lattice of
# 7-person demes for the next -- both legitimate fountain points, but a page of panels that
# change their unit of measurement from row to row is unreadable. Candidates are therefore
# restricted first to the note's baseline geometry, which leaves only the contact rate and the
# growth rate free; every row but one has between three and sixteen admissible cells there.
# The exception is the deme model at theta = 0, which has exactly one admissible cell in the
# whole screen and it is a 6x6 lattice. That row falls back, and says so.
CANON_DEME = dict(grid=12, K=11)
CANON_BAND = dict(K=25, p_move=0.45, sigma_b=60.0, sigma_e=80.0, alpha_d=0.8)

_DEME_KEYS = ("grid", "K", "m", "gamma", "r", "female_bias", "s_mt", "lineage")
_BAND_KEYS = ("K", "e", "p_move", "beta", "r", "sigma_b", "sigma_e", "alpha_d", "s_mt")


_SUMS = {}
# Scanning the screen directories means opening fifty thousand small JSON files, which takes
# a minute and a half of the two minutes this figure otherwise needs. The fields this module
# reads are a dozen numbers per cell, so they are distilled once into a single file and read
# back from there; --rescan rebuilds it.
SUM_CACHE = os.path.join(HERE, "figED_summaries.json")
_KEEP = ("mech", "val", "aut", "end_gen", "n_est", "n", "joint", "N", "lineage",
         "grid", "K", "m", "gamma", "r", "female_bias", "s_mt",
         "e", "p_move", "beta", "sigma_b", "sigma_e", "alpha_d")


def _cell_summaries(fw):
    """[summary] for one framework, at the precedence screen.py applies."""
    if fw in _SUMS:
        return _SUMS[fw]
    if not _SUMS and os.path.exists(SUM_CACHE) and "--rescan" not in sys.argv:
        try:
            _SUMS.update(json.load(open(SUM_CACHE)))
        except Exception:
            _SUMS.clear()
        if fw in _SUMS:
            return _SUMS[fw]
    for f in ("deme", "band"):
        files = {}
        for pref in SC.SETS:
            for d in (f"out_{pref}1_{f}", f"out_{pref}1hi_{f}", f"out_{pref}2_{f}"):
                for q in glob.glob(os.path.join(SC.CL, d, "*.json")):
                    files[os.path.basename(q)] = q
        # READ THROUGH cluster/summary_cache.py WHEN IT EXISTS. Opening fifty thousand JSON
        # files one at a time on the mounted tree takes longer than one shell call here is
        # allowed; the cache holds the same "summary" blocks, gathered once in parallel.
        _cached = {}
        try:
            sys.path.insert(0, SC.CL)
            import summary_cache
            _cached = summary_cache.load()
        except Exception:
            _cached = {}
        out = []
        for q in files.values():
            try:
                sm = _cached.get(os.path.abspath(q)) or json.load(open(q)).get("summary", {})
            except Exception:
                continue
            if sm.get("mech") is None or sm.get("val") is None:
                continue
            out.append({k: sm[k] for k in _KEEP if sm.get(k) is not None})
        _SUMS[f] = out
    json.dump(_SUMS, open(SUM_CACHE, "w"))
    return _SUMS[fw]


def _cell_of(sm):
    """The screen's Cell for one summary, without re-reading the file it came from.

    screen._read recomputes `both` from the per-replicate rows, which this module never
    looks at; everything pick_params tests is in the summary itself.
    """
    a, g, n = sm.get("aut"), sm.get("end_gen"), sm.get("n") or 0
    if a is None or g is None or a != a or g != g:
        return None
    return SC.Cell(aut=float(a), gen=float(g),
                   est=((sm.get("n_est") or 0) / n) if n else None,
                   joint=sm.get("joint"), reps=sm.get("reps") or n or None,
                   mech=sm.get("mech"), val=sm.get("val"))


def _tw_files(fw):
    return ([os.path.join(SC.CL, f"out_tw5_{fw}", "*.json")]
            + ([os.path.join(SC.CL, "out_tw6_band", "*.json")] if fw == "band" else []))


def _canon_ok(sm, canon):
    return all(sm.get(k) is not None and abs(float(sm[k]) - v) < 1e-9 for k, v in canon.items())


def pick_params(fw, mech, val):  # noqa: D401
    """Engine kwargs for the admissible cell of (fw, mech, val) closest to 5% autosomal.

    Returns ({}, None) when the mechanism value was never screened -- s = 0.0005 is not on the
    ladder, and the row falls back to the baseline with a note on the figure.
    """
    # THE DEME LATTICE NEVER CHANGES. DR: "make all the grid sizes the same, why is Route 3
    # at 6x6?" Because the screen sweeps the lattice as a nuisance axis and theta = 0 has
    # exactly one admissible cell in the whole sweep, a 6x6 of 40-person demes. A page of
    # panels that changes its unit of measurement halfway down cannot be read, so the deme
    # side now stays at grid 12 / K 11 come what may: if no cell there is admissible, the
    # closest one to 5% autosomal at the right geometry is used and the cell file says so.
    # The band side has no such constraint -- its panels are a map, and the kernel does not
    # change the size of anything -- so there the box comes first and the baseline second.
    canon = CANON_DEME if fw == "deme" else CANON_BAND
    passes = ([(CANON_DEME, True), (CANON_DEME, False)] if fw == "deme"
              else [(canon, True), ({}, True), ({}, False)])
    for cn, want_box in passes:
        got = _pick(fw, mech, val, cn, want_box)
        if got[1] is not None:
            return got[0], got[1], bool(cn) and want_box
    return {}, None, False


def _pick(fw, mech, val, canon, want_box=True):
    best = None
    if mech == "twowave":
        for pat in _tw_files(fw):
            for q in glob.glob(pat):
                try:
                    r = json.load(open(q))
                except Exception:
                    continue
                if r.get("died") or not r.get("reps") or not r.get("wave2_gen"):
                    continue
                # tw5_run fixes the deme geometry at grid 12 / K 11 and does not record it,
                # so there is nothing to test and the absence of the fields is not evidence
                # of an off-baseline cell. On the band side the kernel IS swept and IS
                # recorded, so the test applies.
                if fw == "band" and not _canon_ok(
                        r, {k: v for k, v in canon.items()
                            if k in ("p_move", "sigma_b", "sigma_e", "K")}):
                    continue
                dur = r.get("arr_med", r.get("dur_med"))
                # p10 IS CARRIED, and it was not at first: without it the two-wave rows had
                # no retention probability, the seed search read that as "below a half" and
                # went looking for a replicate in which the Y does NOT fix -- the opposite of
                # what the route claims.
                c = SC.Cell(aut=float(r["aut"]), gen=(1e9 if dur is None else float(dur)),
                            est=r.get("arr_frac", r.get("est")),
                            joint=r.get("p10"), reps=r.get("reps"))
                if want_box and not SC.in_box(c):
                    continue
                d = abs(c["aut"] - AUT_ANCHOR)
                if best is None or d < best[0]:
                    kw = dict(wave2_gen=int(r["wave2_gen"]))
                    if fw == "deme":
                        # tw5 deme files record the swept gamma as "val" and run at m = 0.08
                        kw.update(gamma=float(r["val"]), m=0.08, K=11, grid=12)
                    else:
                        kw.update(e=float(r.get("e", r.get("val"))), K=int(r.get("K", 25)),
                                  N_target=int(r.get("N", 1750)),
                                  p_move=float(r.get("p_move", 0.45)),
                                  r=float(r.get("r", 2.0)),
                                  sigma_b=float(r.get("sigma_b", 60.0)),
                                  sigma_e=float(r.get("sigma_e", 80.0)))
                    best = (d, kw, c)
    else:
        for sm in _cell_summaries(fw):
            if sm.get("mech") != mech:
                continue
            if abs(float(sm["val"]) - float(val)) > 1e-9:
                continue
            if not _canon_ok(sm, canon):
                continue
            c = _cell_of(sm)
            # `joint` is required, not merely nice to have: a cell without one is a cell the
            # screen accepted but never scored, so it is not a point in Figure 2b and this
            # figure should not present it as one.
            if c is None or c.get("joint") is None:
                continue
            if want_box and not SC.in_box(c):
                continue
            d = abs(c["aut"] - AUT_ANCHOR)
            if best is None or d < best[0]:
                keys = _DEME_KEYS if fw == "deme" else _BAND_KEYS
                kw = {k: sm[k] for k in keys if sm.get(k) is not None}
                if fw == "band":
                    kw["N_target"] = int(sm.get("N", 1750))
                    kw.pop("N", None)
                best = (d, kw, c)
    return (({}, None) if best is None else (best[1], best[2]))


# ---------------------------------------------------------------------------------------
# WHICH REPLICATE EACH ROW SHOWS
# ---------------------------------------------------------------------------------------
# DR: "search for seeds where the autosome proportion ends between 3-7%" and "restrict to
# 1000 gen outcomes that match the fountain plot, so if it is <50% probability of fixing make
# the Y not fix, and if it is >[50%] make it fix."
#
# One replicate is a draw, and a draw can be unrepresentative in either direction: the first
# version's Route 3 theta = 0.1 deme panel showed the modern Y at 38% and falling, in a cell
# whose scored retention is 11% -- not wrong, but a reader would take the picture for the
# result. The seed is therefore chosen rather than fixed at zero, on two conditions that are
# both properties the cell already claims:
#
#   1. autosomal ancestry at the last sampled generation inside the 3-7% acceptance window,
#      so the panel is a replicate the screen would have accepted;
#   2. the Y fixed if the cell's p_joint is at least a half, not fixed if it is below,
#      so the panel agrees with the fountain point it is drawn from.
#
# Seeds are tried in order and the first that satisfies both is used, so the choice is
# reproducible and is the lowest qualifying seed rather than a hand-picked one. The answers
# are cached in figED_seeds.json; --reseed recomputes them.
SEED_TRIES = 40
SEED_CACHE = os.path.join(HERE, "figED_seeds.json")
FIX = 0.99          # "the marker is fixed": modern at 99% of the whole landscape or more


def _final(fw, kw, gens, seed):
    """(mt, Y, autosome) modern fractions at the last sampled generation, or None if dead."""
    snaps = (deme_snapshots(kw, seed, gens) if fw == "deme"
             else band_snapshots(kw, seed, gens)[0])   # band_snapshots also returns the map
    snap = snaps.get(max(gens))
    return None if snap is None else snap[-1]


def _trial(job):
    """2 = window and Y outcome both match, 1 = window only, 0 = neither."""
    fw, kw, gens, seed, want_fix = job
    try:
        w = _final(fw, kw, gens, seed)
    except Exception:
        return seed, 0
    if w is None:
        return seed, 0
    mt, y, au = w
    if au != au or not (AUT_LO <= 100 * au <= AUT_HI):
        return seed, 0
    return seed, (2 if bool(y == y and y >= FIX) == want_fix else 1)


def find_seed(fw, kw, gens, want_fix, tag, cache, tries=SEED_TRIES):
    """The lowest seed that lands in the window and matches the fountain's Y outcome.

    Falls back to the lowest seed that only lands in the window, and finally to seed 0. The
    cache stores [seed, grade] so a later run knows how good the answer was without redoing
    the search.
    """
    if tag in cache:
        return tuple(cache[tag])
    from concurrent.futures import ProcessPoolExecutor
    scores = {}
    with ProcessPoolExecutor(max_workers=4) as ex:
        for sd, g in ex.map(_trial, [(fw, kw, gens, sd, want_fix) for sd in range(tries)]):
            scores[sd] = g
    best = ([sd for sd in sorted(scores) if scores[sd] == 2]
            or [sd for sd in sorted(scores) if scores[sd] == 1])
    cache[tag] = [best[0], scores[best[0]]] if best else [0, 0]
    return tuple(cache[tag])


def row_kwargs(fw, mech, val, mech_kw):
    """The row's mechanism settings on top of its own admissible cell's geometry."""
    got, cell, canon = pick_params(fw, mech, val)
    base = dict(DEME_BASE if fw == "deme" else BAND_BASE)
    base.update(got)
    base.update(mech_kw)
    return base, cell, canon


def deme_snapshots(kw, seed, gens=None):
    """{generation: (grid, per-deme [mt, Y, auto] modern fractions)}.

    The generation loop is hand-rolled rather than calling S3.run() because run() returns
    summary trajectories and this figure needs the STATE at three moments. The steps are
    S3's own -- migrate, reproduce, reseed -- so nothing is reimplemented, only stopped at.

    spatial_sim3 is used for every row even where spatial_sim2 would do, because sim3 is a
    strict superset and picking per row would mean maintaining a list of which mechanism
    lives where. That list is exactly what went wrong in robust_run in September.
    """
    gens = list(GENS if gens is None else gens)
    cfg = DConfig(**dict(DEME_BASE, seed=seed, **kw))
    cfg = _replace(cfg, gens=max(gens) + 1, sample_gen=max(gens))
    rng = np.random.default_rng(seed)
    P = _init(cfg, rng)
    w2 = int(getattr(cfg, "wave2_gen", 0) or 0)
    out = {}
    n = cfg.grid ** 2
    for g in range(max(gens) + 1):
        if g in gens:
            out[g] = (cfg.grid, _per_deme(P, n), _whole_deme(P))
        if w2 and g == w2:
            # THE WAVE-2 RELABELLING, WHICH THE ENGINE DOES NOT DO. `wave2_gen` appears
            # nowhere in spatial_sim3: robust_run performs this externally, and the one time
            # a caller forgot to, every deme two-wave cell silently became a single-wave run
            # under a two-wave label. Copied from robust_run's stage-2 loop rather than
            # paraphrased.
            #
            # Only IDENTITY is reset. The residents keep the alleles the first wave left
            # them, which is the whole point of the route: after a matrilocal first wave they
            # already carry modern mtDNA, and the patrilocal second wave supplies the Y.
            P["lin"][:] = 0
            if "f1m" in P:
                P["f1m"][:] = False
            here = P["pos"] == 0
            if here.any():
                P["lin"][here] = 1
                P["mt"][here] = 1
                male = here & (P["sex"] == 1)
                P["Y"][male] = 1
                P["auto"][here, :] = 2
                P["X"][here, :] = np.where(P["sex"][here, None] == 0, 2, 1)
            cfg = _replace(cfg, lineage="patrilineal", female_bias=0.0)
        _migrate(P, cfg, rng)
        nxt = S3._repro(P, cfg, rng)
        if nxt is None:
            break
        P = _reseed_origin(nxt, cfg)
    return out


def _whole_deme(P):
    """Whole-landscape modern fractions, the numbers the panel label quotes.

    Computed over the WHOLE population rather than averaged over the per-deme fractions:
    the two differ whenever demes differ in size, and it is the population proportion a
    reader means by "the Y is 9% modern".
    """
    male = P["sex"] == 1
    mt = float((P["mt"] == 1).mean()) if P["mt"].size else np.nan
    y = float((P["Y"][male] == 1).mean()) if male.any() else np.nan
    au = float(P["auto"][:, :32].mean() / 2.0) if P["auto"].size else np.nan
    return mt, y, au


def _per_deme(P, n):
    """Per-deme modern fractions at mtDNA, Y and the measured autosomes; NaN where empty."""
    pos = P["pos"]
    male = P["sex"] == 1
    size = np.bincount(pos, minlength=n).astype(float)
    mt = np.bincount(pos, weights=(P["mt"] == 1), minlength=n)
    # the measured panel is loci 0-31; 32-35 are reserved for mechanisms and are under
    # selection in several rows, so including them would make the autosomal bar report the
    # mechanism rather than ancestry
    au = np.bincount(pos, weights=P["auto"][:, :32].mean(1) / 2.0, minlength=n)
    ym = np.bincount(pos[male], minlength=n).astype(float)
    yv = np.bincount(pos[male], weights=(P["Y"][male] == 1), minlength=n)
    with np.errstate(invalid="ignore", divide="ignore"):
        return np.column_stack([np.where(size > 0, mt / size, np.nan),
                                np.where(ym > 0, yv / ym, np.nan),
                                np.where(size > 0, au / size, np.nan)])


def band_snapshots(kw, seed, gens=None):
    """{generation: (positions, sizes, per-band [mt, Y, auto])}, as Figure 2a builds them."""
    gens = list(GENS if gens is None else gens)
    cfg = BConfig(**{k: v for k, v in dict(BAND_BASE, **kw).items()
                     if k != "wave2_gen"})
    cfg.seed = seed
    cfg.gens = max(gens) + 1
    rng = np.random.default_rng(seed)
    land = Landscape()
    P, pos, b_ident = init_state(cfg, land, rng)
    out = {}
    w2 = int(kw.get("wave2_gen", 0) or 0)
    seed_xy = np.array(cfg.seed_lonlat, float)
    for g in range(max(gens) + 1):
        if w2 and g == w2:
            # The band counterpart, also performed by robust_run rather than by step():
            # every resident band is relabelled archaic, a fresh modern band is founded at
            # the Bosphorus, and the residence rule flips to the opposite bias.
            b_ident[:] = False
            j = int(np.argmin(((pos - seed_xy) ** 2).sum(1)))
            b_ident[j] = True
            mem = P["band"] == j
            if mem.any():
                P["mt"][mem] = 1
                P["Y"][mem & (P["sex"] == 1)] = 1
                P["auto"][mem, :] = 2
                P["X"][mem, :] = np.where(P["sex"][mem] == 1, 1, 2)[:, None]
            import copy as _c
            cfg = _c.copy(cfg)
            cfg.beta = 1.0
        if g in gens:
            # per_band averages all 36 autosomal loci, including the four mechanism loci that
            # several rows put under selection; restrict to the neutral panel by handing it a
            # view of P whose "auto" field is loci 0-31, so the bars measure ancestry.
            Pn = dict(P, auto=P["auto"][:, :32])
            alive, size, mtf, yf, auf = per_band(Pn, b_ident, len(b_ident))
            # WHOLE LANDSCAPE, AND EXCLUDING THE MECHANISM LOCI -- the same two choices
            # _whole_deme makes, so the number under a band panel means what the number under
            # the deme panel beside it means. The first version read mt_mod / Y_mod /
            # auto_mod, which are measured WITHIN MODERN-IDENTITY BANDS only. That is the
            # right quantity for tracking an expanding group and the wrong one for a caption
            # that says "the Y is 9% modern", and it is also why the two-wave row's last band
            # panel printed "mt -- Y -- au --": after the wave-2 relabelling every band is
            # archaic-identity bar one, and when that one does not survive to generation
            # 1,000 the modern-only denominator is zero.
            st = BS.stats(P, b_ident, exclude=tuple(range(32, BS.AUTOS)))
            out[g] = (pos[alive].copy(), size, np.column_stack([mtf, yf, auf]),
                      (st["mt"], st["Y"], st["auto"]))
        res = step(P, pos, b_ident, cfg, land, rng)
        if res is None or res[0] is None:
            break
        P, pos, b_ident = res
    return out, land


def _label_text(whole, gen=None):
    """The three whole-population percentages, on one line BELOW the panel.

    DR: "Will have to label ancestry proportions for each replicate: '% modern: mtDNA=100%,
    Y=9%, autos=7%'." Per PANEL rather than per row, because all three move with generation
    and a row-level number would be true of at most one of the three columns.

    BELOW THE PANEL, NOT INSIDE IT. The first version put a boxed three-line label in the
    top-left corner, which in the deme panels sits on top of nine demes -- every cell of the
    lattice is data, so there is no corner to put it in -- and in the band panels covered the
    North Sea. Under the axes it occupies space that was empty anyway, and one line fits
    because "mt" and "au" are unambiguous once the caption has said what the three bars are.
    """
    if whole is None:
        return ""
    mt, y, au = whole
    def pct(v):
        return "--" if v != v else f"{100 * v:.0f}"
    # The two-wave row is sampled on its own clock, so its panels carry the generation --
    # on a line of its own, because the two strings together are wider than a deme panel.
    head = "" if gen is None else f"generation {gen:,}\n"
    return f"{head}M={pct(mt)}%, Y={pct(y)}%, A={pct(au)}%"


def draw_deme(ax, snap):
    """One square per deme at its lattice position; empty demes are left blank."""
    ax.set_xticks([]); ax.set_yticks([])
    for sp in ax.spines.values():
        sp.set_visible(False)
    if snap is None:
        ax.set_xlim(0, 1); ax.set_ylim(0, 1)
        return None
    grid, fr, whole = snap
    ax.set_xlim(-0.6, grid - 0.4); ax.set_ylim(-0.6, grid - 0.4)
    ax.set_aspect("equal")
    for i in range(grid * grid):
        f = fr[i]
        if not np.isfinite(f).any():
            continue
        # NaN at the Y means the deme has no males this generation, which is a real state in
        # a small deme and not missing data. It is drawn as an empty bar rather than skipped,
        # so the square still marks an occupied deme.
        # 0.80 RATHER THAN 0.86. At 0.86 on unit spacing the gap between squares is thinner
        # than one of the three bars, so a deme at mtDNA = 100% put its red left bar flush
        # against its neighbour's blue right bar and a fully modern lattice read as blue
        # squares ruled with red lines. The extra gap costs nothing and makes the unit of the
        # picture the square rather than the stripe.
        # MIRRORED IN x SO THE ORIGIN IS THE SOUTH-EAST CORNER. DR: "the deme grids look
        # like the migrations are coming from the southwest not the southeast". Deme 0 is
        # the founding deme in spatial_sim2 and sits at lattice (0, 0), which drawn plainly
        # is bottom-LEFT; the expansion this note is about comes out of the Near East. The
        # lattice has no intrinsic handedness -- migration is symmetric in x -- so mirroring
        # the drawing is a relabelling of the axis and not a change to the simulation.
        band_square(ax, (grid - 1) - (i % grid), i // grid,
                    np.nan_to_num(f, nan=0.0), 0.80)
    return whole


_LIM = {}


def band_limits(land):
    """(x0, x1, y0, y1) of the habitable landmass in projected coordinates, with margins.

    CROPPED TO THE LANDMASS. The projected grid runs well beyond the habitable mask, so an
    uncropped panel spends most of its area on empty sea and the bands come out too small to
    read the three bars. Taken from the mask rather than typed in, so it follows the
    landscape if that is ever changed -- and computed ONCE, because main() sizes the band
    columns from the same numbers that draw_band draws inside.
    """
    if not _LIM:
        ii, jj = np.where(land.mask)
        lx, ly = albers(land.lon[jj], land.lat[ii])
        mx, my = 0.02 * (lx.max() - lx.min()), 0.03 * (ly.max() - ly.min())
        _LIM.update(v=(lx.min() - mx, lx.max() + mx, ly.min() - my, ly.max() + my))
    return _LIM["v"]


def draw_band(ax, snap, land):
    """The Albers map with one square per surviving band, sized by band size -- Figure 2a."""
    ax.set_xticks([]); ax.set_yticks([])
    for sp in ax.spines.values():
        sp.set_visible(False)
    LON, LAT = np.meshgrid(land.lon, land.lat)
    GX, GY = albers(LON, LAT)
    ax.contourf(GX, GY, land.mask.astype(float), levels=[0.5, 1.5],
                colors=["#e6e1d8"], zorder=0)
    ax.set_aspect("equal")
    x0, x1, y0, y1 = band_limits(land)
    ax.set_xlim(x0, x1)
    ax.set_ylim(y0, y1)
    if snap is None:
        return None
    p, size, fr, whole = snap
    lon = p[:, 0] / (KM_PER_DEG_LAT * COS_REF) + land.lon0
    lat = p[:, 1] / KM_PER_DEG_LAT + land.lat0
    X, Y = albers(lon, lat)
    # DR: "for the band plots, make the squares smaller so the landscape is not as
    # cluttered". 0.030 -> 0.021 keeps the three bars legible at print size while letting
    # the coastline show through a crowded landscape.
    WB = 0.021
    for k in range(len(X)):
        w = WB * float(np.clip(np.sqrt(max(size[k], 1.0) / 25.0), 0.28, 2.0))
        band_square(ax, X[k], Y[k], np.nan_to_num(fr[k], nan=0.0), w)
    return whole


def main():
    dpi = int(sys.argv[sys.argv.index("--dpi") + 1]) if "--dpi" in sys.argv else 200
    cache = {}
    if os.path.exists(SEED_CACHE) and "--reseed" not in sys.argv:
        try:
            cache = json.load(open(SEED_CACHE))
        except Exception:
            cache = {}

    # THE SEED SEARCH IS RESUMABLE. It is a few minutes of simulation, so it runs in slices
    # -- `--seed-rows 0 4` fills the cache for the first four rows and exits -- and the cache
    # is written after every row. A full render with a warm cache is under a minute.
    seeds_only = "--seed-rows" in sys.argv
    only = None
    if seeds_only:
        k = sys.argv.index("--seed-rows")
        only = (int(sys.argv[k + 1]), int(sys.argv[k + 2]))

    rows = ROWS
    nrow, ncol = len(rows), 7

    # EVERY PANEL IS DRAWN AT THE SAME NORTH-SOUTH SCALE, AND EVERY PANEL FILLS ITS SLOT.
    # DR: "the roving band model plots should be rescaled so the southernmost-to-northernmost
    # -point-of-Europe dimension is the same as that of the squares in the deme model. Then
    # the text underneath can line up." Both kinds of panel hold their aspect ratio, so
    # matplotlib shrinks whichever axes box does not match its data and the two halves ended
    # up different heights with their captions at different places. The fix is to give each
    # column a slot of exactly the right shape and let nothing shrink:
    #
    #   * the deme lattice is square, so its slot is square;
    #   * the map's slot is its own width-to-height ratio, measured from the landmass.
    #
    # The figure width is then solved for, rather than guessed, so that one unit of slot
    # height is one unit of slot width in inches. That also answers "make deme plots bigger":
    # the lattice now fills the full row height, and the page widens to let it.
    x0, x1, y0, y1 = band_limits(Landscape())
    asp = (x1 - x0) / (y1 - y0)
    # A SPACER COLUMN DOWN THE MIDDLE. DR: "make more space on either side of the middle
    # vertical line to get the aspect ratio a bit better." wspace is uniform across a
    # gridspec, so the only way to widen one gutter is to give it a column of its own: an
    # empty axes, 0.42 of a deme panel wide, with the divider drawn down its centre. It also
    # widens the sheet, which is what the aspect ratio wanted.
    SPACER = 0.42
    wr = [1.0, 1.0, 1.0, SPACER, asp, asp, asp]
    row_h = 1.72
    fig, axes = plt.subplots(nrow, ncol, squeeze=False,
                             figsize=(row_h * sum(wr) + 2.0, row_h * nrow + 0.75),
                             gridspec_kw=dict(width_ratios=wr))
    # bottom leaves a strip for the key; hspace leaves room for the ancestry line, which is
    # two lines deep in the two-wave row.
    fig.subplots_adjust(left=0.112, right=0.996, top=0.962, bottom=0.050,
                        wspace=0.07, hspace=0.185)
    # One exact correction: the fractional geometry above is fixed, so the figure width that
    # makes the deme slot square in inches is figh * (slot height fraction / width fraction).
    pos = axes[0][0].get_position()
    fig.set_size_inches(fig.get_figheight() * pos.height / pos.width, fig.get_figheight())

    land, table = None, []
    for i, (label, dkw, bkw, key) in enumerate(rows):
        if only is not None and not (only[0] <= i < only[1]):
            continue
        dfull, dcell, dcan = row_kwargs("deme", key[0], key[1], dkw)
        bfull, bcell, bcan = row_kwargs("band", key[0], key[1], bkw)

        # (9) THE TWO-WAVE ROW IS SAMPLED ON ITS OWN CLOCK. DR: "uniquely for the two-wave
        # plots, we may need to modify the generation we're sampling at and label it as
        # such." The second wave is founded at generation 200 (band) or 500 (deme), so at
        # the common 100 / 300 / 1,000 the row would show two panels of wave one and one
        # panel of the aftermath. Sampling at the founding of the second wave and 300 and
        # 1,000 generations after it puts the three panels at the same points in the
        # scenario's own history as every other row, and each panel says what generation it
        # is. Note the first panel is the state the second wave arrives INTO.
        dg = ([dfull["wave2_gen"] + d for d in (0, 300, 3000)]
              if key[0] == "twowave" else list(GENS))
        bg = ([bfull["wave2_gen"] + d for d in (0, 300, 3000)]
              if key[0] == "twowave" else list(GENS))

        dfix = bool(dcell is not None and (dcell.get("joint") or 0) >= 0.5)
        bfix = bool(bcell is not None and (bcell.get("joint") or 0) >= 0.5)
        dmg = [dfull["wave2_gen"] + MEASURE_GEN] if key[0] == "twowave" else [MEASURE_GEN]
        bmg = [bfull["wave2_gen"] + MEASURE_GEN] if key[0] == "twowave" else [MEASURE_GEN]
        dseed, dok = find_seed("deme", dfull, dmg, dfix, f"deme|{key[0]}|{key[1]}", cache)
        bseed, bok = find_seed("band", bfull, bmg, bfix, f"band|{key[0]}|{key[1]}", cache)
        json.dump(cache, open(SEED_CACHE, "w"), indent=1, sort_keys=True)
        if seeds_only:
            print(f"  row {i + 1:>2}  {label.replace(chr(10), ' '):<24}"
                  f" deme seed {dseed} (grade {dok})   band seed {bseed} (grade {bok})")
            continue

        dsn = deme_snapshots(dfull, dseed, dg)
        bsn, land = band_snapshots(bfull, bseed, bg)
        wholes = []
        for j in range(3):
            wholes.append(draw_deme(axes[i][j], dsn.get(dg[j])))
        for j in range(3):
            wholes.append(draw_band(axes[i][j + 4], bsn.get(bg[j]), land))
        axes[i][3].axis("off")

        # (5) THE SIX ANCESTRY LINES OF A ROW SIT ON ONE LINE. DR: "in the band plots, align
        # the mt/Y/au labels to be on the same rows as the deme for the same scenario." They
        # used to be placed in each panel's own axes coordinates, and because every panel
        # sets an equal aspect, matplotlib shrinks each axes box to its data -- by a
        # different amount for a square lattice and for a map -- so the two halves' labels
        # drifted apart. Placed in FIGURE coordinates off the subplot slot, which does not
        # shrink, they cannot.
        pos = axes[i][0].get_position()
        y_lab = pos.y0 - 0.0035
        for k in (0, 1, 2, 4, 5, 6):
            bx = axes[i][k].get_position()
            g = (dg[k] if k < 3 else bg[k - 4])
            fig.text((bx.x0 + bx.x1) / 2, y_lab,
                     _label_text(wholes[k if k < 3 else k - 1],
                                 g if key[0] == "twowave" else None),
                     ha="center", va="top", fontsize=6.6, color="0.15")
        fig.text(0.016, (pos.y0 + pos.y1) / 2, label, fontsize=PLAIN_PT - 1.0,
                 va="center", ha="left", linespacing=1.35)

        table.append((label.replace("\n", " "), dfull, dcell, dcan, dseed, dok, dfix,
                      bfull, bcell, bcan, bseed, bok, bfix))
        print(f"  row {i + 1:>2}  {label.replace(chr(10), ' '):<24}"
              f" deme seed {dseed:>2}{'' if dok else '!'}"
              f" gamma={dfull.get('gamma')} grid={dfull.get('grid')}"
              f"{'' if dcan else ' [cell not in box]'}"
              f" | band seed {bseed:>2}{'' if bok else '!'} e={bfull.get('e')}"
              f"{'' if bcan else ' [off-baseline kernel]'}")

    if seeds_only:
        json.dump(cache, open(SEED_CACHE, "w"), indent=1, sort_keys=True)
        print("seed cache written")
        return

    # (2) COLUMN HEADERS, in figure coordinates rather than as titles: every panel sets an
    # equal aspect, so a title hangs off a box that has been shrunk by a different amount in
    # each column and the band headers sat visibly below the deme ones.
    y_hdr = axes[0][0].get_position().y1 + 0.004
    for j, g in enumerate(GENS):
        for k in (j, j + 4):
            bx = axes[0][k].get_position()
            fig.text((bx.x0 + bx.x1) / 2, y_hdr, f"{g:,} gens", ha="center", va="bottom",
                     fontsize=PLAIN_PT, fontweight="bold")

    # THE DIVIDER, in figure coordinates, midway between column 3 and column 4. Drawn as a
    # figure line rather than a spine so it spans every row without belonging to any panel.
    x_div = (axes[0][3].get_position().x0 + axes[0][3].get_position().x1) / 2
    fig.add_artist(Line2D([x_div, x_div], [0.046, 0.948], color="0.35", lw=1.1))

    xa = (axes[0][0].get_position().x0 + axes[0][2].get_position().x1) / 2
    xb = (axes[0][4].get_position().x0 + axes[0][6].get_position().x1) / 2
    fig.text(xa, 0.992, "Deme model", ha="center", va="top", fontsize=BOLD_PT,
             fontweight="bold")
    fig.text(xb, 0.992, "Roving band model", ha="center", va="top", fontsize=BOLD_PT,
             fontweight="bold")

    # THE KEY TO THE SQUARE, in Figure 2a's words rather than paraphrased, so a reader who
    # has met the convention in the main text meets the same sentence here. DR: "add
    # somewhere a key for how to interpret the square using the same language as Figure 2a."
    # Drawn in the empty margin under the row labels, which is the only white space on the
    # sheet big enough to hold it.
    FW, FH = fig.get_size_inches()
    ih = 0.0165
    kx, ky = 0.016, 0.008
    icon = fig.add_axes([kx, ky, ih * FH / FW, ih])
    icon.set_xlim(0, 1); icon.set_ylim(0, 1); icon.axis("off")
    band_square(icon, 0.5, 0.5, (0.85, 0.5, 0.12), 1.0)
    fig.text(kx + ih * FH / FW + 0.006, ky + ih / 2,
             "In each square, modern fraction given left-to-right for mtDNA-Y-autosomes; "
             "area is band size.   M / Y / A: whole-landscape percentages.",
             va="center", ha="left", fontsize=PLAIN_PT + 1.0)

    json.dump(cache, open(SEED_CACHE, "w"), indent=1, sort_keys=True)

    # THE CELL TABLE, written beside the figure. Every panel is a draw from a specific
    # screened cell at a specific seed, and the caption has to be able to name both.
    with open(os.path.join(HERE, "FigureED_expansions_cells.txt"), "w") as fh:
        fh.write("Row-by-row provenance for Extended Data: sample expansions.\n\n"
                 "Each row is drawn at the cell of the contact sweep whose autosomal\n"
                 "ancestry is closest to 5%, among those admissible (3-7% autosomal,\n"
                 "100-1,000 generations, establishment > 50%) at the note's baseline\n"
                 "geometry. The replicate is the lowest seed whose own autosomal ancestry\n"
                 "ends in 3-7% AND whose Y outcome matches the cell: fixed where the cell's\n"
                 "p_joint is at least 0.5, unfixed where it is below. Both are judged at\n"
                 "generation 1,000, where the acceptance window is defined; the panels are\n"
                 "drawn at 30, 300 and 3,000.\n\n")
        for (lab, dk, dc, dcan, ds, dok, dfx,
             bk, bc, bcan, bs, bok, bfx) in table:
            fh.write(f"{lab}\n")
            for tag, kw, c, can, sd, ok, fx in (("deme", dk, dc, dcan, ds, dok, dfx),
                                                ("band", bk, bc, bcan, bs, bok, bfx)):
                if c is None:
                    fh.write(f"    {tag}: no scored cell; drawn at the baseline, seed {sd}\n")
                    continue
                keys = (("grid", "K", "m", "gamma", "r") if tag == "deme"
                        else ("N_target", "K", "e", "p_move", "r", "sigma_b", "sigma_e",
                              "alpha_d"))
                ps = ", ".join(f"{k} = {kw[k]:g}" for k in keys if k in kw)
                off = "" if can else ("   [no admissible cell at this geometry; the closest "
                                      "one to 5% is used]")
                pj = "--" if c.get("joint") is None else format(c["joint"], ".3f")
                w2 = "" if "wave2_gen" not in kw else f", second wave at {kw['wave2_gen']:g}"
                fh.write(f"    {tag}: {ps}{w2}{off}\n"
                         f"          cell: autosomal {c['aut']:.2f}%, {c['gen']:.0f} "
                         f"generations, p_joint {pj}\n"
                         f"          shown: seed {sd}, judged at generation "
                         f"{MEASURE_GEN:,}, chosen for Y "
                         f"{'fixed' if fx else 'not fixed'}"
                         f"{'' if ok else '  [NO SEED IN THE FIRST ' + str(SEED_TRIES) + ' QUALIFIED; SEED 0 SHOWN]'}\n")
    for ext in ("pdf", "png"):
        fig.savefig(os.path.join(HERE, f"FigureED_expansions.{ext}"), dpi=dpi)
    print(f"\nwrote FigureED_expansions.pdf / .png  "
          f"{fig.get_figwidth():.1f} x {fig.get_figheight():.1f} in")


if __name__ == "__main__":
    main()
```
