## Supplementary Information section 3 code base for "Hypothesis: A modern human range expansion ∼300,000 years ago had a critical role in Neandertal origins": figED_sensitivity.py

```
#!/usr/bin/env python3
"""
figED_sensitivity.py -- Extended Data: the two sensitivity figures stacked. (14 Sep 2026)

DR: "a stacked version of the two figures in the supplement showing simulation robustness to
parameter variations, deme on top and band on the bottom. It should be the exact same content,
just stacked with a and b labels."

    a   Figure S3.5, the deme framework: grid, K, m, gamma, r
    b   Figure S3.6, the band framework: K, e, r, alpha, sigma_b, sigma_e

COMPOSED FROM THE RENDERED PDFs RATHER THAN REDRAWN, which is the whole point of doing it this
way. "Exact same content" is a guarantee if the panels are the very files the supplement uses,
and merely a hope if they are redrawn from a second code path that has to be kept in step.
This is the same approach figure2_full.py takes for Figure 2, and the same reason.

WIDTHS DIFFER AND THE PANELS ARE THEREFORE CENTRED, not left-aligned. The deme framework has
five swept parameters and the band framework six, so at 1.95 in per column panel b is about
two inches wider. figure2_full can left-align because both its panels are saved at the same
width; here that would hang panel a off to one side. Centring puts the two grids on a common
vertical axis, which is what the eye uses to compare a row across the two frameworks.

THE PANEL LETTERS ARE A SEPARATE VECTOR OVERLAY. fig_sensitivity draws no letters -- it has
never needed them -- and adding them there would put "a" and "b" into the supplement figures
too. The overlay is a transparent matplotlib page the size of the composed sheet carrying two
characters, merged on top, so both the supplement and this figure keep exactly the artwork
they should have and neither constrains the other.

    python figED_sensitivity.py                 # rebuild both panels, then compose
    python figED_sensitivity.py --compose       # compose from the PDFs already on disk
"""
import os
import subprocess
import sys

import matplotlib
matplotlib.use("Agg")
import matplotlib.pyplot as plt
from pypdf import PageObject, PdfReader, PdfWriter
from pypdf.generic import RectangleObject

HERE = os.path.dirname(os.path.abspath(__file__))
# ED-SPECIFIC PANELS, not the supplement's own files. DR: "remove the detailed titles, and
# just change to 'a  Deme model parameter sensitivity' and 'b  Roving band model parameter
# sensitivity'." The supplement captions carry their figure numbers and a long provenance
# subtitle, both wrong inside a paper. fig_sensitivity.build() takes the title and subtitle as
# arguments, so the panels are redrawn with those blank and the headings supplied by the
# overlay instead -- the artwork is still the one code path, which is what keeps the two
# versions from drifting.
A_PDF = os.path.join(HERE, "figED_a_deme.pdf")
B_PDF = os.path.join(HERE, "figED_b_band.pdf")
TITLE_A = "Deme model parameter sensitivity"
TITLE_B = "Roving band model parameter sensitivity"
TITLE_PT = 13.0
TITLE_GAP_PT = 26.0      # from the letter's left edge to the start of the heading
OUT_PDF = os.path.join(HERE, "FigureED_sensitivity.pdf")
OUT_PNG = os.path.join(HERE, "FigureED_sensitivity.png")

GAP_PT = 14.0        # vertical space between the two panels, in points
# DR: "put the title 'Deme model' and 'Roving band model' in bold and centered and spaced
# more from the actual plots." The headings used to be drawn INSIDE each panel's own top
# margin, beside its letter, which is why they sat almost on the first row of axes. Each
# panel now gets a band of its own above it, HEAD_PT tall, and the heading is centred in it.
HEAD_PT = 40.0
LETTER_PT = 17.0     # panel letter size; the supplement's own panel letters are 30 at
                     # Figure 2's 16.4 in width, which scales to about this at 12 in
PAD_PT = 4.0         # inset of the letter from the top-left corner of its panel


def rebuild():
    """Redraw both panels untitled, through fig_sensitivity's own build()."""
    sys.path.insert(0, HERE)
    import fig_sensitivity as FS
    # DR: "remove the 'Light vertical line...' captions in gray." Both sentences move to the
    # figure caption; the supplementary versions of these panels keep them.
    FS.FOOTNOTE = None
    print("rebuilding both sensitivity panels, untitled and unfootnoted")
    deme, band = FS.load_deme(), FS.load_band()
    FS.build(FS.DEME_PARAMS, deme, "", "", "figED_a_deme",
             "Generations to\narchaic extinction")
    FS.build(FS.BAND_PARAMS, band, "", "", "figED_b_band",
             "Generations to\nmarker absorption\n(median)")


def letters_overlay(w, h, positions):
    """A transparent page of the composed size carrying the panel letters and headings.

    BOTH LETTERS SIT AT THE SAME x. DR: "the (a) label in the top should be left justified at
    the same point as the (b) one at the bottom." They did not, because the two panels have
    different widths and each letter was insetting from its own panel's left edge -- so the
    narrower deme panel pushed its letter inward. The common x is the left edge of the wider
    panel, which is the left edge of the sheet.
    """
    fig = plt.figure(figsize=(w / 72.0, h / 72.0))
    fig.patch.set_alpha(0.0)
    for ch, (x, y, title) in positions.items():
        # figure coordinates, measured from the bottom left as PDF space is. The letter stays
        # at the common left edge; the heading is centred on the sheet and bold.
        fig.text(x / w, y / h, ch, fontsize=LETTER_PT, fontweight="bold",
                 va="top", ha="left")
        fig.text(0.5, y / h, title, fontsize=TITLE_PT, fontweight="bold",
                 va="top", ha="center")
    tmp = os.path.join(HERE, "_ed_letters.pdf")
    fig.savefig(tmp, transparent=True)
    plt.close(fig)
    return tmp


def compose():
    for p in (A_PDF, B_PDF):
        if not os.path.exists(p):
            raise SystemExit(f"missing {p} -- run without --compose first")
    a = PdfReader(A_PDF).pages[0]
    b = PdfReader(B_PDF).pages[0]
    aw, ah = float(a.mediabox.width), float(a.mediabox.height)
    bw, bh = float(b.mediabox.width), float(b.mediabox.height)

    W = max(aw, bw)
    # A HEADING BAND ABOVE EACH PANEL. Top to bottom the page is now
    #     [head a][panel a][gap][head b][panel b]
    # so the two headings have HEAD_PT of clear space to sit in and neither crowds the first
    # row of its own axes.
    H = HEAD_PT + ah + GAP_PT + HEAD_PT + bh
    ax0 = (W - aw) / 2.0        # centred; see the module docstring
    bx0 = (W - bw) / 2.0

    page = PageObject.create_blank_page(width=W, height=H)
    # pypdf composes bottom-up, so panel b (band) goes on first and panel a (deme) above it
    # -- which is DR's "deme on top and band on the bottom".
    page.merge_translated_page(b, bx0, 0)
    page.merge_translated_page(a, ax0, bh + HEAD_PT + GAP_PT)

    x_letter = min(ax0, bx0) + PAD_PT        # common left edge; see letters_overlay
    y_a = H - PAD_PT                          # top of the sheet
    y_b = bh + HEAD_PT - PAD_PT               # top of panel b's heading band
    tmp = letters_overlay(W, H, {"a": (x_letter, y_a, TITLE_A),
                                 "b": (x_letter, y_b, TITLE_B)})
    page.merge_page(PdfReader(tmp).pages[0])
    os.remove(tmp)

    page.mediabox = RectangleObject((0, 0, W, H))
    w = PdfWriter()
    w.add_page(page)
    with open(OUT_PDF, "wb") as fh:
        w.write(fh)
    print(f"\nwrote {os.path.basename(OUT_PDF)}   {W / 72:.2f} x {H / 72:.2f} in")
    print(f"   a  deme  {aw / 72:.2f} x {ah / 72:.2f} in")
    print(f"   b  band  {bw / 72:.2f} x {bh / 72:.2f} in")

    try:
        import pymupdf
    except ImportError:
        print("   (no pymupdf, so no PNG preview)")
        return
    d = pymupdf.open(OUT_PDF)
    d[0].get_pixmap(dpi=200).save(OUT_PNG)
    print(f"wrote {os.path.basename(OUT_PNG)} at 200 dpi")


def main():
    if "--compose" not in sys.argv:
        rebuild()
    compose()


if __name__ == "__main__":
    main()
```
