## Supplementary Information section 3 code base for "Hypothesis: A modern human range expansion ∼300,000 years ago had a critical role in Neandertal origins": figED_thumbnails.py

```
#!/usr/bin/env python3
"""
figED_thumbnails.py -- the thumbnail survey in the NEW route order. (14 Sep 2026)

DR: "make a version of the Extended Data Figure with the thumbnails, reordered with our new
order", the order being the one Figure 2b now uses:

    Scenario 1   Dobzhansky-Muller incompatibilities causing selection against the archaic Y
    Scenario 2   Selection against the archaic patriline or matriline
    Scenario 3   Two waves
    Scenario 4   Fertility of the first-generation offspring of the mixing archaic's sex

NOTHING IS RECOMPUTED. Every panel is the same panel fig_thumbnails_all.py draws, from the
same screen.py call on the same cells; this module reorders the blocks, renumbers the banners
and re-titles the two that changed. Keeping it as a wrapper rather than a copy means the two
figures cannot drift: a change to how a thumbnail is drawn, or to what counts as admissible,
lands in both.

THE FOUR BLOCKS MAP ONTO THE NEW NUMBERING TWO-TO-ONE IN ONE PLACE. Figure 2b's Scenario 1 is a
single panel whose x-axis is the daughter proportion, with the mtDNA x Y column beside it; the
thumbnail figure splits the same territory into incompatibilities that act on the Y itself and
incompatibilities that act through a partner locus, because that split is the figure's main
point. They become Scenario 1a and Scenario 1b rather than being merged, so the split survives and
the numbering still matches.

THE LAST BANNER ROW SWAPS SIDES. Scenarios 3 and 4 share one banner row, two waves in a narrow
block beside the cultural mechanisms in a wide one. Under the old numbering the cultural block
was Scenario 3 and sat on the left; under the new one two waves is Scenario 3, so it moves to the
left and the cultural block to the right, and the banners follow. Reading order and numbering
agree again.

THE COLOUR KEY TRAVELS WITH THE SELECTION BLOCK, which is now Scenario 2 rather than Scenario 1, so
the reader meets it in the middle of the page rather than at the top. That is the one thing
this reordering makes worse. The alternative is to move the key into the Scenario 1a block, which
has no free cell -- its two rows are full at seven columns -- so it would cost a row.

    python figED_thumbnails.py [--aut-lo 3 --aut-hi 7] [--refresh]
        -> FigureED_thumbnails.png (300 dpi) and FigureED_thumbnails.pdf
"""
import os
import sys

HERE = os.path.dirname(os.path.abspath(__file__))
sys.path.insert(0, HERE)

import fig_thumbnails_all as T                                     # noqa: E402

NCOL = T.NCOL


def reordered():
    """The blocks of fig_thumbnails_all, in the new route order and renumbered."""
    sel, direct, partner, (cult_banners, r3) = T._build_groups()

    # The shared last row, rebuilt with the two-wave panel on the left. r3's rows are five
    # cultural panels followed by the two-wave cell and a blank; they become the two-wave
    # cell and a blank followed by the five cultural panels.
    tw_cell = next(c for c in r3[0][5:] if c)
    rows34 = [[tw_cell, None] + r3[0][:5],
              [None, None] + r3[1][:5]]

    return [
        # SCENARIO 1a and 1b, retitled on DR's wording. The old banners explained the mechanism
        # -- "no escape route, because the Y itself cannot be replaced" against "introgression
        # replaces the archaic partner and the barrier lapses" -- and the new ones state the
        # result instead, which is what the reader is looking at: the first block's numbers
        # run to 100% and the second's top out at 18%.
        ("SCENARIO 1a   DMI involving the Y (works effectively)", direct[1]),
        ("SCENARIO 1b   DMI not involving the Y (does not work)", partner[1]),
        # SCENARIO 2: selection on the marker itself, stated without naming the Y for the reason
        # given in fig_thumbnails_all -- the model is symmetric under exchanging the sexes.
        ("SCENARIO 2   Selection against the archaic patriline or matriline", sel[1]),
        ([("SCENARIO 3   Two waves, matrilineal and patrilineal", 0, 2),
          ("SCENARIO 4   Fertility of the first-generation offspring of the archaic parent's sex",
           2, NCOL)],
         rows34),
    ]


def main():
    T.GROUPS = reordered()
    T.PANELS = [pn for _, rows in T.GROUPS for row in rows for pn in row if pn]
    T.OUT_BASE = "FigureED_thumbnails"
    # DR: "remove the header 'Each point is one searched...'". It says what the colours and
    # the bold numbers mean, which is caption material; in the supplementary figure it is a
    # running head because that figure is read away from its caption.
    T.HEADING = None
    n_old = len([pn for _, rows in T._build_groups() for row in rows for pn in row if pn])
    # A COUNT CHECK, because a reordering that silently drops a thumbnail looks fine.
    if len(T.PANELS) != n_old:
        raise SystemExit(f"panel count changed: {n_old} -> {len(T.PANELS)}")
    print(f"  {len(T.PANELS)} thumbnails, four blocks, new route order")
    T.main()


if __name__ == "__main__":
    main()
```
