## Supplementary Information section 3 code base for "Hypothesis: A modern human range expansion ∼300,000 years ago had a critical role in Neandertal origins": figure2_full.py

```
#!/usr/bin/env python3
"""
figure2_full.py -- the whole of main-text Figure 2: panel a above panel b. (12 Sep 2026)

DR: "what we've been calling Figure 2 is just panel b, in figure include panel a (the four
maps)."

    panel a   four maps of Europe from ONE matrilocal range expansion, at generations 0, 100,
              300 and 1,000, each band a square whose three bars give the modern fraction at
              mtDNA, the Y and the autosomes and whose area is the number of people in it.
              Drawn by band_model/figure2_bands.py --panel a --only-a.
    panel b   the four routes: p_joint against mechanism strength, four demographic scenarios
              as lines and every admissible Cartesian combination as points.
              Drawn by figure2_merged.py.

STACKED AS VECTORS, NOT AS IMAGES. Both sources already emit PDF, and pypdf can place one page
inside another with a transform, so the composite keeps live text and line art at any zoom. The
obvious alternative -- rendering both to PNG and concatenating -- would have upscaled panel a
from 1,632 px to panel b's 3,279 px, a 2x blur on the only panel in the figure that a reader
looks at closely.

PANEL LETTERS COME FROM THE SOURCES, not from here: figure2_bands draws its own "a", and
figure2_merged now draws a "b". That keeps them as vector text positioned in each panel's own
coordinates, so they stay put whatever width the composite is scaled to.

WIDTH IS SET BY PANEL B, which is the wider of the two, and panel a is centred under it. Panel a
is 10.2 in wide against panel b's 16.4, so scaling panel a UP to match would blow the maps up
past the point where the 70-odd band squares stay distinguishable. Centring it at its native size
keeps both panels legible and reads as a deliberate layout rather than a stretch.

    python figure2_full.py            # rebuilds both panels, then composes
    python figure2_full.py --compose  # compose from the PDFs already on disk
"""
import os
import subprocess
import sys

from pypdf import PageObject, PdfReader, PdfWriter
from pypdf.generic import RectangleObject

HERE = os.path.dirname(os.path.abspath(__file__))
BAND = os.path.join(HERE, "band_model")
# --alt BUILDS THE ALTERNATIVE VERSION (DR, 2 Oct 2026) and writes to its own four
# filenames, so the published Figure 2 is never overwritten and the two can be compared
# side by side. Panel a goes monochrome (black modern, grey archaic, pale yellow land) and
# panel b drops three of its four scenarios and recodes deme/band as blue/red dots.
ALT = "--alt" in sys.argv
_SFX = "_alt" if ALT else ""
A_PDF = os.path.join(BAND, f"figure2_panelA{_SFX}.pdf")
B_PDF = os.path.join(HERE, f"Figure2_merged{_SFX}.pdf")
OUT_PDF = os.path.join(HERE, f"Figure2_full{_SFX}.pdf")
OUT_PNG = os.path.join(HERE, f"Figure2_full{_SFX}.png")
GAP_PT = 14.0            # breathing room between the panels, in points


def build():
    # PANEL a IS NOT TOUCHED BY --alt. DR, 2 Oct 2026, having seen the monochrome version:
    # "Let's go back to the blue/red and yellow background in the top panel a, exactly as
    # before." So the variant is now a panel-b-only change, and --mono survives only as an
    # option nothing calls. The red/blue key in panel a is also why panel b must not use red
    # or blue for the two frameworks -- see FW_COL in figure2_merged.py.
    print("panel a  (band_model/figure2_bands.py --panel a --only-a)")
    r = subprocess.run([sys.executable, "figure2_bands.py", "--panel", "a", "--only-a",
                        "--out", f"figure2_panelA{_SFX}.png"],
                       cwd=BAND, capture_output=True, text=True)
    for line in (r.stdout or "").strip().splitlines():
        print("   ", line.strip())
    if r.returncode != 0:
        raise SystemExit((r.stderr or "").strip()[-1500:])

    print("\npanel b  (figure2_merged.py" + (" --alt)" if ALT else ")"))
    r = subprocess.run([sys.executable, "figure2_merged.py"] + (["--alt"] if ALT else []),
                       cwd=HERE, capture_output=True, text=True)
    if r.returncode != 0:
        raise SystemExit((r.stderr or "").strip()[-1500:])
    print(f"    rebuilt Figure2_merged{_SFX}.pdf")


def compose():
    for p in (A_PDF, B_PDF):
        if not os.path.exists(p):
            raise SystemExit(f"missing {p} -- run without --compose first")
    a = PdfReader(A_PDF).pages[0]
    b = PdfReader(B_PDF).pages[0]
    aw, ah = float(a.mediabox.width), float(a.mediabox.height)
    bw, bh = float(b.mediabox.width), float(b.mediabox.height)

    W = max(aw, bw)
    H = ah + GAP_PT + bh
    page = PageObject.create_blank_page(width=W, height=H)
    # pypdf composes bottom-up, so panel b goes on first at y = 0 and panel a above it.
    # LEFT-ALIGNED, NOT CENTRED. Both panels are now saved uncropped at the same 16.4 in width,
    # so stacking them at x = 0 makes their internal margins line up -- which is what puts Spain
    # directly above panel b's y-axis label. Centring was what broke the alignment before.
    page.merge_translated_page(b, 0, 0)
    page.merge_translated_page(a, 0, bh + GAP_PT)
    page.mediabox = RectangleObject((0, 0, W, H))

    w = PdfWriter()
    w.add_page(page)
    with open(OUT_PDF, "wb") as fh:
        w.write(fh)
    print(f"\nwrote {os.path.basename(OUT_PDF)}   {W / 72:.2f} x {H / 72:.2f} in")
    print(f"   panel a {aw / 72:.2f} x {ah / 72:.2f} in, centred")
    print(f"   panel b {bw / 72:.2f} x {bh / 72:.2f} in")

    try:
        import pymupdf
    except ImportError:
        print("   (no pymupdf, so no PNG preview)")
        return
    d = pymupdf.open(OUT_PDF)
    d[0].get_pixmap(dpi=160).save(OUT_PNG)
    print(f"wrote {os.path.basename(OUT_PNG)} at 160 dpi")


def main():
    if "--compose" not in sys.argv:
        build()
    compose()


if __name__ == "__main__":
    main()
```
