## Supplementary Information section 3 code base for "Hypothesis: A modern human range expansion ∼300,000 years ago had a critical role in Neandertal origins": figure2_merged.py

```
#!/usr/bin/env python3
"""
figure2_merged.py -- Figures 2b and 2c as ONE figure. (DR, 11 Sep 2026)

DR, twice: "I still don't believe Figure 2b and 2c merge. I think they can be made to merge and
it would be better."

He is right that they can, and right that the first attempt did not. That attempt
(figure2bc_combined.py) collapsed 2b's four scenarios to four horizontal reference lines and
threw away the mechanism-strength axis, which is the only thing 2b ever had to say. Three of the
four lines then coincided at zero and had to be pooled into a grey band, so the panel asserted
"the baselines are all at zero" without showing the reader anything they could check.

WHAT MAKES THE MERGE WORK is keeping the mechanism-strength axis and overlaying the two things
on it, rather than flattening one onto the other:

  2b was  p_joint vs mechanism strength, four demographic scenarios, BASELINE parameters.
  2c is   p_joint at the Cartesian-admissible parameter combinations, sex-biased range
          expansion only, one fountain per mechanism setting.

Same y, same x. So: the four scenarios stay as lines, and the fountain is drawn on top of the
line for the scenario it belongs to (sex-biased range expansion). The reader sees at a glance
that the fountain sits where the matrilocal curve is, that the other three scenarios never leave
the martingale band whatever the mechanism does, and how wide the parameter uncertainty is at
each mechanism strength. None of that survives in a bar of horizontal lines.

THE MARTINGALE BAND is drawn across every panel at the autosomal proportion. It is the ceiling
for any scenario in which the marker is not exempted by the residence rule, and it is why the
three non-matrilocal curves are flat: no strength of selection on the Y can rescue the mtDNA.

    python figure2_merged.py [--aut-lo 3 --aut-hi 7] [--gen-lo 100 --gen-hi 1000]
"""
import csv
import math
import os
import sys

import matplotlib
matplotlib.use("Agg")
import matplotlib.pyplot as plt

# FONTS MATCHED TO PANEL a. DR: "Make the font sizes in general bigger in panel b and matched if
# possible to the font sizes of panel a." Panel a runs at 14 pt for the generation titles, 12 for
# the ancestry block and 14 for its key, so panel b's titles go to 14, its axis labels to 12 and
# its ticks to 11. Set through rcParams rather than call by call, so an axis added later inherits
# them instead of quietly reverting to the 10 / 8 defaults; the handful of hand-set sizes that
# remain are the ones that have to be smaller to fit.

import numpy as np
from matplotlib.lines import Line2D

HERE = os.path.dirname(os.path.abspath(__file__))
sys.path.insert(0, HERE)
import screen  # noqa: E402
from figure2c_fountain import load, flag  # noqa: E402
# The two text classes come from figure2_bands so panels a and b cannot drift apart. Imported
# after HERE is defined, which is why this sits below the fountain import rather than with the
# other top-level imports.
sys.path.insert(0, os.path.join(HERE, "band_model"))
from figure2_bands import BOLD_PT, PLAIN_PT, LETTER_PT        # noqa: E402

plt.rcParams.update({"axes.labelsize": PLAIN_PT, "xtick.labelsize": PLAIN_PT,
                     "ytick.labelsize": PLAIN_PT, "legend.fontsize": PLAIN_PT})

# COLOUR IS THE SCENARIO, SHAPE IS THE FRAMEWORK. DR: "we are going to want to have the same
# color scheme as in our original Figure 2b, and use only shape to separate band and deme."
#
# The palette is lifted verbatim from band_model/figure2_bands.py, which draws the published
# Figure 2, so a curve here is the same colour as the same curve there. The earlier draft of
# this figure spent colour on deme-versus-band, which is the least interesting distinction on
# the page and left nothing to distinguish the four scenarios with.
#
# This also makes the figure ready for the `demog` sweep. Right now every plotted point is a
# matrilocal range expansion, so every point is fexp red; once that sweep lands, the unbiased
# combinations enter the same fountains in nexp orange, and the scheme needs no change.
SCEN = {
    # The red series is no longer one calibrated parameter set but the MEAN over the
    # admissible combinations drawn beneath it, so the label has to say which it is --
    # otherwise the reader takes it for a scenario like the other three.
    # DR, 27 Sep: "matrilocal" -> "sex-biased" in the legend, and "all-female" likewise.
    # The four entries now read as two pairs -- range expansion and migration pulse, each
    # sex-biased or unbiased -- which is the contrast the panel is actually drawn to make,
    # and it no longer commits the key to a residence rule the curves do not distinguish.
    "fexp": ("range expansion, sex-biased", "#b2182b", "-",  "o"),
    "nexp": ("range expansion, unbiased",   "#e08214", "-",  "^"),
    "fsb":  ("migration pulse, sex-biased", "#2166ac", "--", "s"),
    # Darkened from the published #7f7f7f on DR's instruction. In the original Figure 2 this
    # series was a control sitting at zero and mid-grey was right for it; with mtDNA selection
    # switched on it is the second-highest curve on the page and carries the 16% result, so a
    # colour that reads as "ignore me" now actively misleads.
    "nsb":  ("migration pulse, unbiased",   "#3f3f3f", "--", "D"),
}
SCEN_COL = {k: v[1] for k, v in SCEN.items()}

# ------------------------------------------------------------------ ALTERNATIVE VERSION
# --alt, DR 2 Oct 2026: "(1) In panel b remove the range expansion unbiased and migration
# pulse lines in the four routes (so remove three lines in each panel). (2) instead of using
# triangles and squares for the two model classes, use dots with two different colors, the
# same red/blue code as in the thumbnail plots."
#
# The two changes are connected, and that is the point of the variant. In the published
# figure colour means SCENARIO and shape means FRAMEWORK; dropping three of the four
# scenarios leaves colour with nothing to say, so colour is handed to the framework instead
# -- deme blue, band red, as in the thumbnails and in Extended Data Figure 1 -- and both get
# the same round marker.
#
# Which forces one decision DR did not specify: the surviving fexp curve was red, and red now
# means the band framework. A red line over red-and-blue points would read as a band-model
# curve, which it is not -- it is the mean over both. So in --alt the curve is black. Flagged
# because it is my choice, not an instruction.
ALT = "--alt" in sys.argv
DRAW = {"fexp"} if ALT else set(SCEN)        # which scenarios get drawn at all
# NOT RED AND BLUE. DR, 2 Oct 2026: panel a stays exactly as published -- red squares for
# modern, blue for archaic, yellow landmass -- so if panel b used red and blue for band and
# deme the page would carry two contradictory keys in the same two colours. Purple and
# orange appear nowhere else in the figure, and are a red-green-safe pair: they separate
# along the blue-yellow axis, which deuteranopia and protanopia both preserve, and they
# differ in lightness so they survive greyscale printing too. The same pair is used by
# fig_thumbnails_all.py --alt, so the two figures agree.
FW_COL = {"deme": "#6a51a3", "band": "#d95f02"}
CURVE_C = "#000000"                          # the fexp mean curve in --alt
# Framework is carried by shape alone: filled circle for the deme lattice, open triangle for
# the roving bands. Open-versus-filled does half the work, so the two are legible even where
# the fountains overlap and even in greyscale.
FW_MARK = {"deme": "o", "band": "o"} if ALT else {"deme": "o", "band": "^"}
FW_FILL = {"deme": True, "band": True} if ALT else {"deme": True, "band": False}
MARTINGALE = 0.05        # the autosomal proportion; see the note's martingale argument


def baseline_curves(src="joint2"):
    """p_joint vs mechanism strength for the four scenarios, at baseline parameters.

    THE SOURCE MATTERS, and getting it wrong was my error. Three datasets carry these four
    scenarios and they disagree by a factor of four, for two separate reasons:

      band_suite_summary_o2.csv   band engine, s_mt = 0.       nexp 4%, nsb 6.8%
      joint2_summary.csv          deme, size-matched geometry, s_mt drawn per replicate from
                                  the Muller's-ratchet posterior.   nexp 12%, nsb 15.8%
      joint_summary.csv           deme, SUPERSEDED grid 20 / K 20, same posterior.
                                                                    nexp 21-42%, nsb 19%

    The first jump is DR's point: "for the non-sex-biased scenarios mtDNA selection a la
    Muller's Ratchet is required". Without it the mtDNA of an unbiased expansion is an
    unselected martingale and p_joint cannot exceed the autosomal proportion whatever is done
    to the Y, so the band-suite curves are flat by construction. I drew the first version of
    this figure from that file, which is why its unbiased curves hugged the martingale and I
    could not reproduce the ~23% in the current Figure 2b.

    The second jump is the geometry. joint_summary.csv predates the Ne recalibration and sits
    at grid 20 / K 20, whose post-wave census is ~8,000 against the ~1,500 the Peyregne anchor
    implies. The potency of a mtDNA coefficient scales with 2*Ne*s_mt, so a 0.25% coefficient
    does two to four times as much work there as it does at the anchored size. That is where
    ~23% comes from, and it is not a number the note can still quote.

    Default is joint2: the deme framework, so it matches the fountains, at the size-matched
    geometry, with the mtDNA fitness gap switched on.
    """
    fn = {"joint2": "joint2_summary.csv", "joint": "joint_summary.csv"}[src]
    out = {}
    for r in csv.DictReader(open(os.path.join(HERE, fn))):
        out.setdefault((r["scenario"], r["sweep"]), []).append(
            (float(r["value"]), float(r["joint_pct"]) / 100.0))
    for k in out:
        out[k].sort()
    return out


def panm_pulse():
    """The panmictic migration-pulse sweep, which is NOT a plain CSV read.

    Its scenario names contain a comma -- "(a) + selection for modern Y, s=0.001" -- and the
    field is unquoted, so every selection row has ten fields against nine headers and
    csv.DictReader silently shifts the whole row one column left. Reading "JOINT%" off it
    returns P(Y fixed), which is 100% at strong selection; that is the source of the
    "unbiased pulse reaches 100%" figure I flagged as a discrepancy against DR's ~16%. It was
    never a result. Parsed positionally from the right, the true joint probability at s = 0.5
    is 3%, and P(mt fixed) never leaves 5% -- the martingale, exactly as the theorem requires.
    """
    fn = os.path.join(HERE, "panm_summary_N1500.csv")
    pts = []
    for line in open(fn).read().splitlines()[1:]:
        f = line.split(",")
        if "selection for modern Y" not in f[0]:
            continue
        s = float(f[1].split("=")[1])
        pts.append((s, float(f[-2]) / 100.0))
    return sorted(pts)


def drive_curves(dirname="cluster/out_jdrive"):
    """{scenario: [(d, p_joint)]} from the drive sweep, or {} until cluster/out_jdrive exists.

    Returns nothing rather than raising when the directory is absent, so this figure can be
    rebuilt before jdrive_submit.py has run and simply omits the two expansion lines. Drawing a
    line that is not backed by a file is the one thing this figure has always refused to do.
    """
    import glob as _glob
    out = {}
    for f in _glob.glob(os.path.join(HERE, dirname, "*.tsv")):
        base = os.path.basename(f)
        sc = base.split("_")[0]
        # nsb ADMITTED 14 Sep 2026. The unbiased migration pulse was absent from Scenario 2
        # because panmictic_sim had no X-linked drive; it has one now, so the black curve can
        # be read here like the other two. fsb stays out and stays analytic: no modern male
        # enters an all-female pulse, so p_joint is identically zero at every d and the flat
        # line is drawn without a run.
        if sc not in ("fexp", "nexp", "nsb"):
            continue
        # only the cost-free arm belongs on this panel; the 10% cost arm is Figure S3.4(a)
        if "_c0.1" in base or "_c0p1" in base:
            continue
        try:
            d = float(base.split("_d")[1].split(".tsv")[0].replace("p", "."))
        except (IndexError, ValueError):
            continue
        # A REPLICATE WITH NO SURVIVING MALES WRITES nan AT THE Y, AND MUST BE DROPPED RATHER
        # THAN AVERAGED. One row of the 8,400 in this sweep -- seed 1139 of fexp at d = 0.8,
        # i.e. 90% daughters -- ends with mt = 1, Y = nan, autosomes = 0. At that drive strength
        # the male pool can vanish before absorption, and the Y frequency is then undefined, not
        # zero. Summing it made p_joint nan for the whole rung and blanked the 90% point of the
        # fexp line, which is how this was noticed. Averaging it AS zero would be worse: it
        # would silently report a 1-in-400 demographic accident as a failure of the mechanism.
        # The rung is reported over the replicates in which the statistic exists, and the count
        # is returned so the caption can say so.
        mt, y, n, dropped = [], [], 0, 0
        for line in open(f).read().splitlines()[1:]:
            fl = line.split("\t")
            if len(fl) < 4:
                continue
            try:
                a, b = float(fl[2]), float(fl[3])
            except ValueError:
                dropped += 1
                continue
            if math.isnan(a) or math.isnan(b):
                dropped += 1
                continue
            mt.append(a); y.append(b); n += 1
        if dropped:
            print(f"    {base}: dropped {dropped} of {n + dropped} replicates with an "
                  f"undefined marker frequency (no surviving males)")
        if not n:
            continue
        # p_joint for ten sampled males, the same statistic the fountains carry
        p = sum((m * yy) ** 10 for m, yy in zip(mt, y)) / n
        out.setdefault(sc, []).append((d, p))
    for k in out:
        out[k].sort()
    return out


def pts(keep, mech, val):
    """Every admissible Cartesian combination for one mechanism setting, by framework."""
    out = {}
    for fw in ("deme", "band"):
        if val is None:
            v = [x for (f, m, _), xs in keep.items() if f == fw and m == mech for x in xs]
        else:
            v = list(keep.get((fw, mech, val), []))
        out[fw] = v
    return out


def spine(ax, keep, pts_xy, scen="fexp"):
    """The red curve: the MEAN of each fountain, joined rung to rung.

    DR: "the red curves should be the average of the fountains... Fountains are points that
    fall in our admissible box, and the red curves are the averages of our fountains."

    WHAT THIS REPLACES, AND WHY IT IS BETTER. The red curve used to be a different object from
    the cloud it ran through: one baseline parameter set, with gamma tuned so that autosomal
    ancestry landed on 5% exactly, read from joint2_summary.csv. That created three problems
    at once, all of which this removes.

      1. IT STOPPED. Below theta = 0.1 no gamma in [0, 1] holds the matrilocal expansion at 5%
         -- jcal_submit calls it the contact-budget ceiling -- so the calibration had no
         solution and the curve simply ended, leaving Scenario 3 with seven fountains and no line
         over them. A mean of the fountains exists wherever a fountain does.
      2. IT LANDED BETWEEN THE TICKS. The calibration sweep's ladder doubles (0.04, 0.08, 0.16,
         0.32) while the mechanism screen uses 0.05, 0.10, 0.20, 0.35, 0.50, so above s = 0.02
         every vertex fell between two fountains. A mean of the fountains is by construction at
         the fountain's own x.
      3. IT ANSWERED A DIFFERENT QUESTION. The cloud is "what does the admissible parameter
         space do"; the old line was "what does one particular parameter set do". Running the
         second through the middle of the first invited the reader to treat them as the same
         statement. They are not, and at theta = 0 they disagreed completely.

    MEAN, NOT MEDIAN, because DR asked for the average and because the quantity being averaged
    is already a probability per cell -- the mean over admissible cells is the expected
    retention of a combination drawn uniformly from the admissible set, which is a statement
    the caption can make. The median is printed alongside so the two can be compared; where
    they diverge the cloud is skewed and that is worth knowing.

    POOLED ACROSS FRAMEWORKS, matching the cloud, which is drawn in one colour with shape
    carrying the framework. Per-framework means go to stdout.
    """
    xs, ys = [], []
    for xpos, mech, val in pts_xy:
        p = pts(keep, mech, val)
        allp = p["deme"] + p["band"]
        if not allp:
            continue
        xs.append(xpos)
        ys.append(float(np.mean(allp)))
        d, b = p["deme"], p["band"]
        print(f"      {mech} {val if val is not None else '':<6} mean {np.mean(allp):.3f}  "
              f"median {np.median(allp):.3f}  "
              f"(deme n={len(d)} mean {np.mean(d):.3f}; band n={len(b)} mean {np.mean(b):.3f})"
              if d and b else
              f"      {mech} {val if val is not None else '':<6} mean {np.mean(allp):.3f}  "
              f"median {np.median(allp):.3f}  (n={len(allp)})")
    if xs:
        ax.plot(xs, ys, "-", color=(CURVE_C if ALT else SCEN_COL[scen]),
                lw=1.8, marker="o", ms=4.2, zorder=6)
    return list(zip(xs, ys))


def pulse_dmi_curve(sc="nsb"):
    """p_joint for a migration pulse carrying an mtDNA x Y incompatibility.

    Returns None until out_mtypulse exists, so the figure omits the marker rather than
    inventing one -- the same rule every other series in this figure follows.
    """
    import glob as _glob
    fs = sorted(_glob.glob(os.path.join(HERE, "cluster", "out_mtypulse", f"{sc}_*.tsv")))
    mt, y = [], []
    for f in fs:
        for ln in open(f).read().splitlines()[1:]:
            t = ln.split("\t")
            if len(t) < 5:
                continue
            try:
                a, b = float(t[2]), float(t[3])
            except ValueError:
                continue
            if a != a or b != b:
                continue
            mt.append(a); y.append(b)
    if not mt:
        return None
    return sum((m * yy) ** 10 for m, yy in zip(mt, y)) / len(mt)


def twopulse_curve(sc):
    """p_joint for a stable-boundary scenario under TWO pulses, pooled over wave-2 timings.

    Scenario 4's comparison points. The expansions get their two-wave counterpart from the
    `demog` sweep's twowave mechanism; the two pulse scenarios had none until panmictic_sim
    gained a second pulse, because a stable boundary has no second event to give them.

    fsb is all-female twice, nsb is all-female then ALL-MALE -- the stable-boundary reading
    of "matrilocal then patrilocal". Pooled over second-wave generations 500, 1000 and 2000
    because Scenario 4's cloud pools over its own wave2 ladder and the two must be comparable.
    """
    import glob as _glob
    vals = []
    for f in sorted(_glob.glob(os.path.join(HERE, "cluster", "out_twpulse", f"{sc}_tp*.tsv"))):
        mt, y = [], []
        for ln in open(f).read().splitlines()[1:]:
            t = ln.split("\t")
            if len(t) < 5:
                continue
            try:
                a, b = float(t[2]), float(t[3])
            except ValueError:
                continue
            if a != a or b != b:      # a replicate with no surviving males; see drive_curves
                continue
            mt.append(a); y.append(b)
        if mt:
            vals.append(sum((m * yy) ** 10 for m, yy in zip(mt, y)) / len(mt))
    return float(np.mean(vals)) if vals else None


def spine_demog(ax, mech, xmap=None, only=None):
    """The orange curve: the mean of the UNBIASED expansion's admissible cells.

    DR settled that the unbiased series is "a single comparison curve like blue and black,
    with red the only fountain series" -- so the cloud is NOT drawn. But the curve should
    still be the same KIND of object as the red one, or the panel carries two lines meaning
    different things: red the mean over an admissible set, orange one calibrated parameter
    point. This computes orange the same way and plots only the line.

    NO LONGER PROVISIONAL, AS OF 15 Sep. The `demog` sweep used to double above s = 0.02
    (0.04, 0.08, 0.16, 0.32) where the main screen uses 0.05, 0.10, 0.20, 0.35, 0.50, so the
    two series shared no rung above 0.02 and no sentence could compare them at a given
    selection strength without hedging. The realigned rungs are scored and on disk -- sY and
    theta at eleven shared values each, xdrive at eight -- so red and orange are now measured
    at the same x everywhere they are drawn. The superseded doubling cells are still on disk
    and are excluded by the `only` filter below, not deleted.
    """
    d = screen.demog_nexp(mech)
    xs, ys = [], []
    vals = sorted(set(d["deme"]) | set(d["band"]))
    if only is not None:
        # THE OLD DOUBLING RUNGS ARE STILL ON DISK AND MUST NOT BE DRAWN. demog's sY ladder
        # was realigned on 14 Sep from 0.04/0.08/0.16/0.32/0.64 to the mechanism screen's
        # 0.05/0.10/0.20/0.35/0.50, but the superseded cells were not deleted -- they are
        # valid runs at parameter values the figure no longer uses. Plotting both interleaves
        # two sweeps and the curve zig-zags between them, which reads as noise in the model
        # rather than as two ladders. Only the current ladder is drawn.
        keep = {round(float(x), 6) for x in only}
        vals = [v for v in vals if round(float(v), 6) in keep]
    for v in vals:
        p = [c["joint"] for fw in ("deme", "band") for c in d[fw].get(v, [])
             if c.in_box and c.get("joint") is not None]
        if not p:
            continue
        x = xmap(v) if xmap else v
        if x is None:
            continue
        xs.append(x); ys.append(float(np.mean(p)))
        print(f"      nexp {mech} {v:<7} n={len(p):<4} mean {np.mean(p):.4f}")
    if xs and "nexp" in DRAW:
        _, c, ls, mk = SCEN["nexp"]
        ax.plot(xs, ys, ls, color=c, lw=1.5, marker=mk, ms=3.4, zorder=4)


def fountain(ax, keep, xpos, mech, val, rng, width=0.13, log=False, scen="fexp",
             symlog_jit=False):
    """One fountain at x = xpos, coloured by scenario and shaped by framework.

    Every combination currently in the sweep is a matrilocal range expansion, so `scen` is
    fexp throughout and the points take fexp's red. The argument exists so that the unbiased
    combinations from the `demog` sweep can be dropped into the same call in nexp's orange
    without touching anything else.
    """
    n = 0
    col = SCEN_COL[scen]
    for fw in ("deme", "band"):
        p = pts(keep, mech, val)[fw]
        if not p:
            continue
        if log:
            jit = xpos * np.exp(rng.uniform(-width, width, len(p)))
        elif symlog_jit:
            # On a symlog axis the spacing between neighbouring theta values is wildly uneven,
            # so a fixed jitter that reads well at theta = 0.2 spills across three ticks at
            # theta = 0.001. Scale it with the value, and give theta = 0 a small linear spread
            # since multiplying zero achieves nothing.
            jit = (rng.uniform(-2.3e-4, 2.3e-4, len(p)) if xpos == 0
                   else xpos * np.exp(rng.uniform(-0.22, 0.22, len(p))))
        else:
            jit = xpos + rng.uniform(-width, width, len(p))
        # In --alt the point colour carries the FRAMEWORK, not the scenario: both frameworks
        # get a filled dot, blue for deme and red for band. `col` (the scenario colour) is
        # ignored here, which is safe because --alt draws only one scenario.
        pc = FW_COL[fw] if ALT else col
        if FW_FILL[fw]:
            ax.scatter(jit, p, s=13, marker=FW_MARK[fw], facecolors=pc, edgecolors="white",
                       linewidths=0.3, alpha=0.55, zorder=5)
        else:
            ax.scatter(jit, p, s=15, marker=FW_MARK[fw], facecolors="none", edgecolors=pc,
                       linewidths=0.7, alpha=0.65, zorder=5)
        n += len(p)
    return n


def martingale(ax):
    """Removed on DR's instruction, 11 Sep. The band was drawn at a fixed 5% to mark the
    ceiling for a marker the residence rule does not exempt. It is now misleading rather than
    helpful: with the mtDNA fitness gap switched on the unbiased curves legitimately sit above
    it, so a reader sees curves crossing what looks like a hard ceiling. The theorem it stood
    for belongs in the text, not as a band the data now exceed. Kept as a no-op so the call
    sites read the same and the decision is recorded rather than silently reverted."""
    return


def main():
    aut_lo, aut_hi = flag("--aut-lo", float, 3.0), flag("--aut-hi", float, 7.0)
    gen_lo, gen_hi = flag("--gen-lo", float, 100.0), flag("--gen-hi", float, 1000.0)
    keep, screened = load(aut_lo, aut_hi, gen_lo, gen_hi)
    base = baseline_curves()
    rng = np.random.default_rng(0)

    # PANEL ORDER, AND IT IS THE SCENARIO NUMBERING THAT MOVED, NOT THE CONTENT. DR: "switch the
    # order of the Scenarios ... Scenario 1 epistasis, Scenario 2 selection against patri/matrilines,
    # Scenario 3 two waves, and Scenario 4 first generation same-sex offspring not raised."
    #
    # Each block below still draws the mechanism it always drew, from the same data; what
    # changed is which column it lands in and what the title calls it. The width ratios move
    # with the panels -- epistasis is the widest because it carries the mtDNA x Y sub-column,
    # two waves the narrowest because it has a single x position -- so the new order is
    # 2.5 (epistasis), 2.2 (selection), 0.65 (two waves), 1.9 (fertility).
    fig, axes = plt.subplots(1, 4, figsize=(16.4, 5.9), sharey=True,
                             gridspec_kw=dict(width_ratios=[2.5, 2.2, 0.65, 1.9],
                                              wspace=0.07))
    counts = {}

    # ------------------------------------------- Scenario 2 (column 2): selection on the Y
    ax = axes[1]
    martingale(ax)
    # fexp IS EXCLUDED FROM THIS LOOP -- its curve is now the mean of the fountains drawn
    # below rather than a single calibrated parameter set. See spine().
    for sc, (lab, c, ls, mk) in SCEN.items():
        if sc in ("fexp", "nexp"):        # both are means of their admissible sets now
            continue
        if sc not in DRAW:
            continue
        xy = base.get((sc, "s"))
        if not xy:
            continue
        # NO CLAMP. The axis is symlog now, so s = 0 has a real position and does not have
        # to be faked at 8e-4 to keep it off a logarithmic axis's missing left edge.
        ax.plot([v for v, _ in xy], [p for _, p in xy], ls, color=c, lw=1.5,
                marker=mk, ms=3.4, zorder=3)
    # THE FULL LADDER, NOT THE ORIGINAL FIVE -- CHANGED 13 Sep. DR: "could you give me sims to
    # give red point clouds below 0.01 and above 0.2". Those runs landed
    # (out_routes1_deme / out_routes1_band both carry s = 0.002, 0.005, 0.35 and 0.50) but this
    # tuple was never extended to match routes_build's ladder, so four fountains' worth of
    # completed simulation was sitting on disk invisible. The list is now the ladder itself:
    # anything screened is drawn, and a fountain missing from the panel means missing data
    # rather than a missing line of code.
    # s = 0 AND s = 0.001 ADDED. DR: "In Scenario 1, points below 0.002 don't have red
    # fountains." They had none because the ladder started at 0.002 while the three
    # comparison curves ran down to zero, so the panel showed lines over a region the cloud
    # did not cover. routes_build now screens both rungs and they are scored: 217 and 208
    # admissible deme cells, with the band arm alongside. s = 0 is the null -- no selection
    # on the Y at all -- and is the one place all four scenarios can be compared with the
    # mechanism switched off.
    SY = (0.0, 0.001, 0.002, 0.005, 0.01, 0.02, 0.05, 0.10, 0.20, 0.35, 0.50)
    for v in SY:
        counts[("sY", v)] = fountain(ax, keep, v, "sY", v, rng, log=True)
    print("    Scenario 1, red curve = mean of each fountain:")
    spine(ax, keep, [(v, "sY", v) for v in SY])
    print("    Scenario 1, orange curve = mean of the unbiased sweep:")
    spine_demog(ax, "sY", only=SY)
    # SYMLOG, NOT LOG -- CHANGED with the null rungs. A logarithmic axis cannot place s = 0
    # at all, which is why the three comparison curves used to clamp their zero to 8e-4 and
    # why the fountains could not start there. symlog is linear inside +/- linthresh and
    # logarithmic outside, so zero sits at its true position and every other interval keeps
    # its ratio. Same settings as Scenario 3's theta axis, so the two panels read alike.
    ax.set_xscale("symlog", linthresh=0.001, linscale=0.45)
    ax.set_xlim(-0.00028, 0.62)
    # Plain decimals rather than 10^-3: these are selection coefficients a reader wants to
    # compare with quoted values like 0.9% or 0.32%, and scientific notation makes that
    # a mental conversion on every glance.
    # THE TICKS ARE THE FOUNTAIN RUNGS, EXACTLY. DR: "the labels in Scenario 1 don't line up with
    # the points in the graph." They did not, in both directions: there was a tick at s = 0.001
    # and at s = 1 where no simulation exists, and a fountain at s = 0.35 with no tick under it.
    # The ticks are now the nine values `routes_build` actually screened, so every tick has a
    # cloud above it and every cloud has a tick below it.
    #
    # THE LINES STILL DO NOT LAND ON THE TICKS, AND SHOULD NOT. The four baseline scenario
    # curves come from the joint2 sweep, whose ladder doubles -- 0.04, 0.08, 0.16, 0.32 --
    # while the mechanism screen uses 0.05, 0.10, 0.20, 0.35, 0.50. Above s = 0.02 the two
    # ladders interleave. The curves are continuous functions of s sampled at their own rungs,
    # so their vertices falling between ticks is correct; forcing them onto the tick positions
    # would mean plotting them at s values they were never run at.
    ax.set_xticks([0.0, 0.001, 0.002, 0.005, 0.01, 0.02, 0.05, 0.10, 0.20, 0.35, 0.50])
    # FEWER LABELS RATHER THAN A SMALLER FONT. Ten of them across this width collided at
    # PLAIN_PT ("0.001 0.002" merged), and shrinking just this row would have broken the rule
    # that every plain label in the figure is one size. Every other tick keeps its mark, so
    # nothing is hidden -- only alternate labels are dropped.
    # EVERY TICK LABELLED -- CHANGED. DR: "In figure 2b x-axes, label all ticks." The
    # alternate-label thinning was introduced when ten labels at PLAIN_PT collided
    # ("0.001 0.002" merged) and the rule that every plain label in the figure is one size
    # ruled out shrinking this row alone. Rotating them 45 degrees buys the width instead: the
    # labels no longer compete for horizontal space at all, they keep PLAIN_PT, and every tick
    # says what it is. Right-anchored so each label ends under its own tick rather than
    # drifting left of it.
    # HORIZONTAL AND CENTRED ON THE TICK, WITH THE LEADING ZERO DROPPED. DR: the labels
    # "should also be all labeled exactly below the ticks... If it helps to remove the 0
    # leading the decimal we can do that." Rotating them did free the width but it also slid
    # each label off its own tick, which is the complaint. Dropping the leading zero buys the
    # same width without moving anything: ".001" is four characters where "0.001" is five, and
    # the tightest tick gap on this axis is 0.301 decades = 0.48 in against a 0.36 in label at
    # PLAIN_PT, so all ten fit centred.
    ax.set_xticklabels(["0", ".001", ".002", ".005", ".01", ".02", ".05", ".1", ".2",
                        ".35", ".5"], fontsize=PLAIN_PT)
    ax.set_xticks([], minor=True)
    # DR asked for this as "Scenario 3", but Scenario 3 is the two-wave panel, which has no
    # x-axis at all -- one scenario at one x position. The selection coefficient is this
    # panel's axis, so the relabelling is applied here.
    ax.set_xlabel("Selection coefficient")
    # DR: "change title of Scenario 1 to 'Selection against archaic patri or matriline' since
    # this applies to both matrilocality and patrilocality" -- "patri" abbreviated on the
    # second pass so the line fits the panel at the same size as its three neighbours. The simulations behind the panel
    # put the selection coefficient on the Y, because the observed pattern is an archaic Y
    # that failed to introgress into a matrilocal expansion; the mechanism is indifferent to
    # which uniparental marker it acts on, and under patrilocality the same argument runs with
    # mtDNA in the Y's place. The title now says the general thing the panel is evidence for.
    # DR: "allow to go to two lines, 'Selection against archaic patriline or matriline'."
    # The abbreviation was only ever there to fit one line; with the subtitle wrapped the
    # full word fits and the shared title size is unaffected.
    ax.set_title("Scenario 2\nSelection against archaic\npatriline or matriline",
                 fontsize=BOLD_PT, fontweight="bold")

    # ------------------------------------------------- Scenario 2: everything acting on the Y
    # DR: the two Scenario 2 panels collapse into one. Left block is the X-Y conflict, with the
    # axis relabelled as the OBSERVABLE -- the proportion of an F1 male's children that are
    # daughters -- rather than the model parameter d. They are the same axis: a distorter that
    # makes a fraction d of Y-bearing sperm inviable leaves (1-d)/2 sons, so the daughter
    # proportion is (1+d)/2. Quoting d asks the reader to do that conversion; quoting the
    # daughter proportion is what the mechanism actually looks like in a pedigree.
    #
    # The failing architectures -- mt x autosome dominant, the polygenic restorers -- are no
    # longer drawn. Their result is that no admissible combination exists, which is a table
    # row, not a column of white space.
    ax = axes[0]
    martingale(ax)
    # DR: "Is 55% daughters really enough of a skew... We may want to include a 50% point,
    # and possibly one intermediate between 50% and 55%. Also, include a 100% daughters point."
    # d is the fraction of Y-bearing sperm made inviable, so the daughter proportion is
    # (1+d)/2: d = 0 is 50% (the null, no distortion at all), d = 0.05 is 52.5%, d = 1 is 100%
    # (no sons at all). The three new rungs are queued in the `xd2` sweep; until it lands they
    # simply plot empty, which is honest -- an absent column is visibly absent.
    XD = [0.0, 0.05, 0.1, 0.2, 0.4, 0.6, 0.8, 1.0]
    for i, v in enumerate(XD):
        counts[("xdrive", v)] = fountain(ax, keep, i, "xdrive", v, rng, width=0.2)

    # THE FOUR BASELINE SCENARIO LINES, as in Scenarios 1 and 3. DR: "add a trace for the baseline
    # parameters for the four types of processes, as in Scenario 1."
    #
    # These come from the `drive` sweep of joint_run.py, which did not exist until 12 Sep --
    # joint2_summary.csv carries only the `s` and `theta` sweeps, which is exactly why this panel
    # has been points-only. cluster/jdrive_submit.py runs it. Of the four scenarios:
    #
    #   fexp, nexp  read from out_jdrive once it lands; drawn only if present, so this figure
    #               never invents a line it does not have.
    #   fsb         ANALYTICALLY ZERO, so it is drawn without a run. No modern male enters an
    #               all-female pulse, the modern Y starts and stays absent, and no amount of
    #               drive against an archaic Y can conjure a modern one. Flat at zero, exactly
    #               as it already is in Scenarios 1 and 3.
    #   nsb         NOT DRAWN. panmictic_sim.py has no X-linked drive, so there is nothing to
    #               read and nothing that can honestly be interpolated.
    # x is the INDEX of the rung in XD, because this panel is categorical -- the drive rungs are
    # not evenly spaced in d and the fountains are drawn at integer positions.
    xpos = {v: i for i, v in enumerate(XD)}
    dv = drive_curves()
    # NEITHER EXPANSION COMES FROM drive_curves ANY MORE. DR: "There are two orange curves in
    # Figure 2b, Scenario 2." There were: I switched the unbiased series to the mean of its
    # admissible set (spine_demog, below) and left the old calibrated line from out_jdrive
    # drawn as well, so the panel carried the same scenario twice under one colour. Only the
    # mean is drawn, matching Scenarios 1 and 3. out_jdrive still supplies the PULSE curves.
    print("    Scenario 2, red curve = mean of each fountain:")
    spine(ax, keep, [(xpos[v], "xdrive", v) for v in XD])
    print("    Scenario 2, orange curve = mean of the unbiased sweep:")
    spine_demog(ax, "xdrive", xmap=lambda v: xpos.get(v))
    # THE UNBIASED PULSE, WHICH I DROPPED BY ACCIDENT AN HOUR AGO. Removing the two-orange-
    # curves bug meant deleting the loop that read drive_curves -- and that loop was also the
    # only thing drawing nsb, the curve panmictic_sim's new X-drive was implemented to
    # produce. The expansions come from their own fountains now; the PULSE still comes from
    # out_jdrive, and always will, because a pulse has no Cartesian sweep to average over.
    xy = [(d, p) for d, p in dv.get("nsb", []) if d in xpos]
    if xy and "nsb" in DRAW:
        _, c, ls, mk = SCEN["nsb"]
        ax.plot([xpos[d] for d, _ in xy], [p for _, p in xy], ls, color=c, lw=1.5,
                marker=mk, ms=3.4, zorder=3)
        print(f"    Scenario 2, unbiased pulse (nsb): {len(xy)} rungs from out_jdrive")
    if "fsb" in DRAW:
        _, c, ls, mk = SCEN["fsb"]
        ax.plot(list(range(len(XD))), [0.0] * len(XD), ls, color=c, lw=1.5,
                marker=mk, ms=3.4, zorder=3)
    MT_X = len(XD) + 1.0
    MT_W = 0.28          # the fountain's jitter half-width, reused by the markers below
    counts[("dmi_mtY", None)] = fountain(ax, keep, MT_X, "dmi_mtY", None, rng, width=MT_W)
    # THE COMPARISON SCENARIOS FOR THIS COLUMN. DR: "There are no blue-orange-black lines for
    # mtDNA-Y incompatibility." Like Scenario 4 this is one categorical column, so they are
    # markers rather than curves.
    #
    # ORANGE comes from the demog sweep, which carries dmi_mtY at beta = 0.5. BLUE is
    # analytic and needs no run, for the same reason it is analytic in the drive block: an
    # all-female pulse admits no modern male, so there is never a modern Y for the
    # incompatibility to act on and p_joint is identically zero. BLACK needs a pulse carrying
    # an mtDNA x Y DMI, which panmictic_sim can express -- it has the dm machinery -- but
    # which no sweep has yet run; see mtypulse_submit.py.
    print("    Scenario 2, mtDNA x Y comparison scenarios:")
    dmt = screen.demog_nexp("dmi_mtY")
    pj = [c["joint"] for fw in ("deme", "band") for v in dmt[fw] for c in dmt[fw][v]
          if c.in_box and c.get("joint") is not None]
    mty = {"nexp": (float(np.mean(pj)) if pj else None),
           "fsb": 0.0,
           "nsb": pulse_dmi_curve()}
    for sc, val in mty.items():
        if sc not in DRAW:
            continue
        if val is None:
            print(f"      {sc:<5} no data")
            continue
        # DR: "make the points the same size as the ones in the other panel, and extend
        # lines the same width as the red points on top." ms 3.4 is the size every scenario
        # marker carries on the curves in Scenarios 1-3, and MT_W is the fountain's own jitter
        # half-width, so the rule is exactly as wide as the cloud it sits under.
        _, c, ls, mk = SCEN[sc]
        ax.plot([MT_X - MT_W, MT_X + MT_W], [val, val], ls, color=c, lw=1.5, zorder=7)
        ax.plot([MT_X], [val], marker=mk, color=c, ms=3.4, ls="none", zorder=8,
                markeredgecolor=c)
        print(f"      {sc:<5} p_joint {val:.4f}")
    # The divider runs BELOW the axis as well as above it, so that the mt-Y column is visibly
    # not the last rung of the daughter-proportion series. DR: "extend a line further down to
    # its left making it clear that it's not part of the daughter distortion series."
    # DIVIDER CLIPPED TO THE AXES. DR: "stop extending the gray line below the plot". It was
    # drawn with clip_on=False and a ymax beyond 1, which let it run down past the x axis and
    # through the tick labels into the caption space. It only ever needed to separate the drive
    # block from the mtDNA x Y column INSIDE the plot, so it is now clipped and stops at the
    # axes frame.
    ax.axvline(len(XD) - 0.35, color="0.55", lw=1.0, zorder=0, clip_on=True)
    ax.set_xlim(-0.6, MT_X + 0.7)
    # REMEMBERED FOR THE --alt LEGEND, which has to stop at this line. The mtDNA x Y column
    # is NOT a separate axes -- it lives in this one, beyond the divider -- so matplotlib's
    # "lower right" puts a legend over the column, straddling the line. Stashing the data
    # coordinate here lets the legend be placed from the divider's real position instead of
    # a guessed offset, and it follows automatically if the ladder gains a rung.
    ax._mt_divider_x = len(XD) - 0.35
    ax.set_xticks(list(range(len(XD))))
    ax.set_xticklabels([f"{(1 + d) / 2:.1%}".replace(".0%", "%") for d in XD],
                       fontsize=PLAIN_PT)
    # Two sub-labels placed under their own blocks, in axis coordinates. A single multi-line
    # xlabel cannot do this: it is centred on the whole axis, so the right-hand caption drifts
    # away from the column it describes as soon as the panel is resized.
    ax.set_xlabel("")
    xl = (len(XD) - 1) / 2.0
    # SAME SIZE AS THE OTHER PANELS' X LABELS. Scenarios 1 and 3 use set_xlabel, which takes
    # rcParams["axes.labelsize"] = 10.0 pt; these two were hand-set at 8.5, so Scenario 2's caption
    # read as a footnote next to its neighbours. DR also asked to drop "due to unsuppressed X-Y
    # conflict", which leaves one line and needs no linespacing.
    ax.text(xl, -0.075, "% daughters in first generation males",
            transform=ax.get_xaxis_transform(), ha="center", va="top", fontsize=PLAIN_PT)
    # Shifted left on DR's instruction: centred on MT_X the second line overhung the
    # panel's right edge, and the column it names sits left of its own tick once the
    # comparison markers are added.
    ax.text(MT_X - 0.30, -0.028, "mtDNA-Y\nincompatibility",
            transform=ax.get_xaxis_transform(), ha="center", va="top",
            fontsize=PLAIN_PT, linespacing=1.35)
    # DR: drop "causing discrimination against archaic Y transmission". Two lines is also what
    # the other three panels use, so the four titles now sit on the same baseline.
    # Wrapped onto three lines: at 12 pt the single long line still overhung the panel and
    # collided with Scenario 1's title, which is the widest in the figure.
    ax.set_title("Scenario 1\nDobzhansky-Muller incompatibilities\n"
                 "causing selection against archaic Y",
                 fontsize=BOLD_PT, fontweight="bold", linespacing=1.35)

    # --------------------------------------- Scenario 4 (column 4): F1 same-sex offspring
    ax = axes[3]
    martingale(ax)
    # SYMLOG, not log. DR wants a log axis with theta = 0 on it, and 0 has no place on a log
    # axis. symlog is linear inside +/- linthresh and logarithmic outside, so theta = 0 sits at
    # its true position, the action between 0.001 and 0.05 is spread out instead of being
    # crushed against the left edge, and the null end at theta = 1 still fits.
    for sc, (lab, c, ls, mk) in SCEN.items():
        if sc in ("fexp", "nexp"):
            continue
        if sc not in DRAW:
            continue
        xy = base.get((sc, "theta"))
        if not xy:
            continue
        ax.plot([v for v, _ in xy], [p for _, p in xy], ls, color=c, lw=1.5, marker=mk,
                ms=3.4, zorder=3)
    # THE FULL LADDER -- CHANGED 13 Sep, as in Scenario 1. DR: "give me sims for 0.5 and 1 point
    # clouds, plus 0.02 and smaller to be comparable to the yellow curve -- I want to connect
    # the line up to 0". Both ends were run and neither end was plotted: theta = 0.5 and 1.0
    # are the weak-mechanism end the baseline curves run out to, and 0.001, 0.002 and 0.005
    # are where the band framework's retention actually collapses (median 1.00 at theta = 0
    # and 0.21 by theta = 0.002), which at five rungs fell entirely into the gap between 0 and
    # 0.02. This is also what the theta threshold quoted in the text is read off, so the
    # figure and the sentence are now looking at the same points.
    TH = (0.0, 0.001, 0.002, 0.005, 0.01, 0.02, 0.05, 0.10, 0.20, 0.50, 1.00)
    for v in TH:
        counts[("theta", v)] = fountain(ax, keep, v, "theta", v, rng, width=0.0,
                                        symlog_jit=True)
    # AND THE CURVE NOW REACHES theta = 0. The calibrated version could not: below theta = 0.1
    # no gamma in [0, 1] holds the matrilocal expansion at 5% autosomal ancestry, so the line
    # stopped and left seven fountains with nothing over them. A mean exists wherever a
    # fountain does.
    print("    Scenario 3, red curve = mean of each fountain:")
    spine(ax, keep, [(v, "theta", v) for v in TH])
    print("    Scenario 3, orange curve = mean of the unbiased sweep:")
    spine_demog(ax, "theta")
    ax.set_xscale("symlog", linthresh=0.001, linscale=0.45)
    # AXIS INVERTED, on DR's instruction, so every panel reads "stronger mechanism to the
    # right". theta is a relative FERTILITY, so it runs the opposite way to s and to the
    # daughter proportion: theta = 1 is no mechanism at all and theta = 0 is complete F1
    # sterility. Left as drawn, Scenario 3 was the one panel whose curves fell as the others rose.
    ax.set_xlim(1.35, -0.00028)
    # THE FULL LADDER, EVERY TICK LABELLED -- CHANGED with Scenario 1. The ticks are now the
    # eleven theta values that actually have fountains, so a reader can place any cloud
    # against its own tick instead of interpolating between every other one, and the labels
    # are rotated for the same reason as Scenario 1's.
    ax.set_xticks([0.0, 0.001, 0.002, 0.005, 0.01, 0.02, 0.05, 0.1, 0.2, 0.5, 1.0])
    # Same convention as Scenario 1: horizontal, centred on the tick, no leading zero. This panel
    # is narrower (1.9 of the 7.25 width units against Scenario 1's 2.2) and carries eleven
    # labels rather than ten, so it is the one place in the figure where PLAIN_PT does not
    # fit -- the tightest pair would overlap by about a character. The row is stepped down by
    # the smallest amount that clears it, measured after the draw rather than guessed; see the
    # fit pass at the end of main(). Every other plain label in the figure keeps PLAIN_PT.
    # TICKS IN PER CENT, because the axis label now is. The positions are unchanged -- they
    # are still theta on a symlog scale -- but a label reading "% offspring raised" over a
    # tick reading ".01" would be read as one hundredth of a per cent when the datum is one
    # per cent. Multiplying the printed labels by 100 leaves the data alone and makes the
    # axis and its label agree.
    ax.set_xticklabels(["0", ".1", ".2", ".5", "1", "2", "5",
                        "10", "20", "50", "100"], fontsize=PLAIN_PT)
    ax.set_xticks([], minor=True)
    ax.set_xlabel("% offspring raised")
    # WRAPPED, and the wrap is what keeps every title at one readable size. On one line this
    # subtitle is 46 characters over the second-narrowest panel, and the shared fit drove all
    # four titles down to 10.75 pt against panel a's 14. Broken in two it clears at full size.
    ax.set_title("Scenario 4\nFertility rate of first gen offspring\n"
                 "of same sex as mixing archaic",
                 fontsize=BOLD_PT, fontweight="bold")

    # ------------------------------------------------ Scenario 3 (column 3): two waves
    ax = axes[2]
    martingale(ax)
    TW_W = 0.25          # the fountain's jitter half-width, reused by the markers below
    counts[("twowave", None)] = fountain(ax, keep, 0, "twowave", None, rng, width=TW_W)

    # THE OTHER THREE SCENARIOS. DR: "Two-waves only have the red points, not the other three
    # lines." This panel is one categorical column, so they are markers rather than curves,
    # drawn at the same x as the cloud and in the same colours and shapes as their lines
    # elsewhere, so a reader who has learned the key in Scenarios 1 to 3 needs no second key.
    #
    # THE RESULT IS WORTH THE SPACE, because it is the one place the note can say whether
    # Scenario 4 needs a range expansion. It does: a pair of sex-biased pulses at a stationary
    # boundary -- all-female then all-male, the stable-boundary reading of "matrilocal then
    # patrilocal" -- reaches p_joint = 0.009, and the unbiased expansion 0.007, against a
    # matrilocal cloud averaging near 0.7. The all-female pulse is exactly zero, as it must
    # be: no modern male enters either wave.
    print("    Scenario 4, comparison scenarios:")
    r4 = {"nexp": None, "fsb": twopulse_curve("fsb"), "nsb": twopulse_curve("nsb")}
    dn = screen.demog_nexp("twowave")
    pj = [c["joint"] for fw in ("deme", "band") for v in dn[fw] for c in dn[fw][v]
          if c.in_box and c.get("joint") is not None]
    if pj:
        r4["nexp"] = float(np.mean(pj))
    for sc, val in r4.items():
        if sc not in DRAW:
            continue
        if val is None:
            print(f"      {sc:<5} no data")
            continue
        # DR asked first for lines across the whole panel, then: "shorten the lines at the
        # bottom to have the same width as the red points at the top." TW_W is the fountain's
        # own jitter half-width, so a comparison rule is exactly as wide as the cloud it is
        # being compared with, in this panel and in the mtDNA x Y column alike.
        _, c, ls, mk = SCEN[sc]
        ax.plot([-TW_W, TW_W], [val, val], ls, color=c, lw=1.5, zorder=7)
        ax.plot([0], [val], marker=mk, color=c, ms=3.4, ls="none", zorder=8,
                markeredgecolor=c)
        print(f"      {sc:<5} p_joint {val:.4f}")
    ax.set_xlim(-0.7, 0.7)
    ax.set_xticks([0])
    # DR: the "second wave" tick label is redundant -- the panel is titled "two waves",
    # and a single categorical column needs no name of its own.
    ax.set_xticks([])
    # DR, 27 Sep: an x-axis title for Scenario 3, "in the same font size as other Figure 2b
    # x-axis titles" -- so no fontsize argument, which leaves it on rcParams["axes.labelsize"]
    # = PLAIN_PT, the same 12 pt set_xlabel gives Scenarios 2 and 4. BROKEN OVER TWO LINES because
    # this panel is 0.65 width units against its neighbours' 1.9 to 2.5: on one line the phrase
    # is wider than the panel and runs under Scenario 4.
    # DR, 8 Oct: name the two waves by the bias they carry rather than by residence
    # rule, which is what the simulation actually varies.
    ax.set_xlabel("one male-biased,\none female-biased", linespacing=1.35)
    # DR: "for Scenario 3, put 'Two' and 'waves' on different lines." The panel is 0.65
    # width units against its neighbours' 1.9 to 2.5, and the two words on one line were
    # wider than the panel they title.
    ax.set_title("Scenario 3\nTwo\nwaves", fontsize=BOLD_PT, fontweight="bold")

    for ax in axes:
        ax.set_ylim(-0.03, 1.05)
        ax.grid(axis="y", color="0.94", lw=0.7)
        ax.set_axisbelow(True)
    axes[0].set_ylabel("Probability modern Y and mtDNA fix", fontsize=PLAIN_PT)

    # Two legends, because they say different things. The first is the colour key -- which
    # demographic scenario -- and reuses the published Figure 2 palette. The second is the
    # shape key, and is drawn in neutral grey precisely to make the point that shape carries
    # the framework independently of colour.
    # TWO SEPARATE LEGENDS IN DIFFERENT PLACES, on DR's instruction. They answer different
    # questions -- colour says which demographic scenario, shape says which spatial framework --
    # and a single combined block invites the reader to think the six entries are six series.
    if ALT:
        # NO SCENARIO LEGEND. DR, 2 Oct 2026: with three of the four series gone there is
        # only one curve on the page, so a one-entry colour key says nothing the caption
        # cannot -- and a lone key invites the reader to look for the other three.
        # The framework key stays, because it is now a COLOUR key and carries the variant.
        h_scen = []
        h_fw = [Line2D([], [], color=FW_COL["deme"], marker="o", ls="none", ms=5,
                       label="deme model"),
                Line2D([], [], color=FW_COL["band"], marker="o", ls="none", ms=5,
                       label="band model")]
    else:
        h_scen = [Line2D([], [], color=c, ls=ls, marker=mk, ms=4, lw=1.5, label=lab)
                  for lab, c, ls, mk in SCEN.values()]
        h_fw = [Line2D([], [], color="0.35", marker="o", ls="none", ms=5,
                       label="deme model (filled)"),
                Line2D([], [], markerfacecolor="none", markeredgecolor="0.35", marker="^",
                       ls="none", ms=6, label="band model (open)")]
    # subplots_adjust, NOT tight_layout: tight_layout does not know about a figure-level legend,
    # so it hands back the space the legend needs and the legend lands on top of the x labels.
    # BOTTOM MARGIN RECLAIMED IN --alt. DR, 2 Oct 2026: "Put the 'Deme model / band model'
    # red/blue key in the Scenario 1 box in the lower right, and clip off the bottom of the
    # figure with the saved space." The 0.235 margin existed to hold two figure-level
    # legends below the axes; with the scenario key gone and the framework key moved inside
    # Scenario 1, all that is left down there is the x tick labels and their axis title.
    fig.subplots_adjust(left=0.068, right=0.992, top=0.845,
                        bottom=0.125 if ALT else 0.235)
    if h_scen:
        fig.legend(handles=h_scen, fontsize=PLAIN_PT, loc="lower left", ncol=2, frameon=False,
                   bbox_to_anchor=(0.055, 0.005), columnspacing=2.4, handlelength=2.6,
                   alignment="left")
    # MOVED OUT OF THE SCENARIO 4 AXES. It used to be an axes legend at "lower right" of the
    # narrow Scenario 4 panel, where its box sat on top of the panel's own frame and its rows ran
    # over the two-wave points -- the one panel too narrow to host it. It is now a figure legend
    # in the bottom margin, opposite the scenario legend at lower left, so the two keys read as
    # a pair and neither touches an axes.
    # No title. DR: "in panel b remove 'framework (shape)'" -- the two entries say "deme model
    # (filled)" and "band model (open)", which is self-describing.
    # CENTRED UNDER THE TWO-WAVE PANEL, not flushed to the page edge. DR: "move the 'deme
    # model (filled)...' legend more left, under 'Two waves'." Taken from that panel's own
    # position rather than typed in, so it follows the column if the width ratios change.
    if ALT:
        # INSIDE SCENARIO 1, lower right. That corner is empty in this panel -- the curve
        # climbs from bottom-left to the top-right plateau -- so the key costs no data.
        # An axes legend rather than a figure legend, so it travels with the panel.
        # RIGHT-JUSTIFIED AGAINST THE DIVIDER. DR, 2 Oct 2026: "justified right against the
        # vertical line separating daughter skew from mtDNA-Y incompatibility". That line is
        # the right-hand edge of this axes -- the incompatibility column is its own narrow
        # axes beyond it -- so "lower right" with a small inset puts the legend's right edge
        # on the divider without any hand-typed coordinate.
        #
        # THE OPAQUE PATCH STAYS. The thing that was bisecting the legend earlier was one of
        # the horizontal grey rules, which a frameless legend lets straight through; that is
        # a property of the legend, not of where it sits, so it would come back here too.
        _a = axes[0]
        _lo, _hi = _a.get_xlim()
        _frac = (getattr(_a, "_mt_divider_x", _hi) - _lo) / (_hi - _lo)
        _a.legend(handles=h_fw, fontsize=PLAIN_PT, loc="lower right", frameon=True,
                  facecolor="white", edgecolor="none", framealpha=1.0,
                  handletextpad=0.4, labelspacing=0.35, borderpad=0.3,
                  bbox_to_anchor=(_frac - 0.015, 0.012), bbox_transform=_a.transAxes)
    else:
        _tw = axes[2].get_position()
        fig.legend(handles=h_fw, fontsize=PLAIN_PT, loc="lower center", frameon=False,
                   bbox_to_anchor=((_tw.x0 + _tw.x1) / 2, 0.005))
    # Panel letter, because this figure is now panel b of the composed Figure 2 (the four
    # band maps are panel a). Drawn here rather than by the composer so that the letter is
    # vector text in the PDF and lands in the same place at any page size.
    # 30 pt to match panel a; see the note there on the 2.3x reduction at print size.
    fig.text(0.004, 0.988, "b", fontsize=LETTER_PT, fontweight="bold", va="top", ha="left")
    # NO SUPTITLE. DR: "Remove the text for panel b about lines and points". It said what the
    # lines and the points are and what the acceptance window is -- all of which belongs in the
    # figure caption, where it is not competing with the panel titles for the same strip of page.
    # The acceptance numbers are still echoed to stdout below, so they stay checkable.
    # ---- x-tick fit pass, MEASURED RATHER THAN GUESSED --------------------------------------
    # DR asked for every tick labelled and for each label to sit exactly under its own tick,
    # which rules out both thinning the labels and rotating them. What is left is the font
    # size, and the size that works is a property of the panel width, the number of ticks and
    # the tick spacing -- three things that have all changed more than once in this figure.
    # So it is measured: draw, take each label's rendered bbox, and step the row down only
    # while two of them actually overlap. A panel whose labels already fit keeps PLAIN_PT
    # untouched, which is why Scenarios 1 and 2 are unaffected and only Scenario 3's eleven labels
    # in the narrowest panel step down at all.
    # ONE SIZE FOR ALL THREE PANELS, FITTED TO THE TIGHTEST. DR: "drop down the numbers in
    # Scenario 1 and Scenario 2 x-axis label panels too to match Scenario 3 font size for numbers."
    # Fitting each panel independently is correct about overflow and wrong about appearance:
    # Scenario 3 is the narrowest panel carrying the most labels, so it came out at 9.25 pt
    # against 12 for its neighbours, and three x-axes of a single figure read at three
    # different sizes. So the loop finds the largest size at which EVERY panel clears, and
    # applies that one size everywhere -- the same rule the thumbnail figure uses for its
    # banners, and for the same reason.
    fig.canvas.draw()
    rend = fig.canvas.get_renderer()
    rows = [[t for t in ax.get_xticklabels() if t.get_text()] for ax in axes]
    rows = [r for r in rows if len(r) > 1]

    def clears(labs):
        bb = sorted((t.get_window_extent(rend) for t in labs), key=lambda b: b.x0)
        # 1 px of clearance: touching labels read as one token at print size
        return all(b.x1 + 1.0 <= nb.x0 for b, nb in zip(bb, bb[1:]))

    size = PLAIN_PT
    while size > 6.0 and not all(clears(r) for r in rows):
        size -= 0.25
        for r in rows:
            for t in r:
                t.set_fontsize(size)
        fig.canvas.draw()
    if size < PLAIN_PT:
        print(f"    x-tick labels fitted at {size:g} pt across all panels "
              f"(PLAIN_PT is {PLAIN_PT}); the binding panel is the narrowest one")

    # TITLES SHRUNK TO THEIR OWN PANEL, measured rather than guessed. Scenario 1's title grew
    # from "Direct selection against the archaic Y" to "Selection against archaic patriline
    # or matriline", which at BOLD_PT is wider than its panel and ran into Scenario 2's title.
    # Each title is measured against the axes it sits over and shrunk only if it overflows,
    # so the three titles that already fit keep their size and nothing moves but the one that
    # did not.
    fig.canvas.draw()
    rend = fig.canvas.get_renderer()
    fitted = []
    for ax in fig.axes:
        t = ax.title
        if not t.get_text():
            continue
        avail = ax.get_window_extent().width * 1.06      # a title may overhang slightly
        size = t.get_fontsize()
        while size > 6.0 and t.get_window_extent(rend).width > avail:
            size -= 0.25
            t.set_fontsize(size)
            fig.canvas.draw()
        fitted.append((t.get_text().splitlines()[0], size,
                       t.get_window_extent(rend).width / ax.get_window_extent().width))
    # ONE SIZE FOR ALL FOUR, which is the whole point of the pass. DR: "make font match."
    # Fitting each title independently left Scenario 1 at 13.5 pt beside three at 14, and a
    # single title half a point off its neighbours reads as a mistake rather than as a fit.
    # The tick labels on this figure already work this way: measure the binding panel, then
    # apply its size everywhere.
    if fitted:
        one = min(sz for _, sz, _ in fitted)
        for ax in fig.axes:
            if ax.title.get_text():
                ax.title.set_fontsize(one)
        fig.canvas.draw()
        fitted = [(n, one, ax.title.get_window_extent(rend).width
                   / ax.get_window_extent().width)
                  for (n, _, _), ax in zip(fitted, [a for a in fig.axes
                                                    if a.title.get_text()])]
    print("    panel titles: " + ",  ".join(f"{n} {sz:g} pt ({fr:.0%} of panel)"
                                            for n, sz, fr in fitted))

    # SEPARATE FILENAMES, so the variant can never overwrite the published panel b and the
    # two can be put side by side.
    stem = "Figure2_merged_alt" if ALT else "Figure2_merged"
    for ext in ("png", "pdf"):
        fig.savefig(os.path.join(HERE, f"{stem}.{ext}"), dpi=200)
    print(f"  wrote {stem}.png / .pdf")
    for k, v in counts.items():
        print(f"    {str(k):<28s} {v:4d} admissible combinations plotted")
    print("  panmictic pulse, joint probability parsed positionally:")
    for s, p in panm_pulse():
        print(f"    s = {s:<6g} joint {p:.2f}")


if __name__ == "__main__":
    main()
```
