## Supplementary Information section 3 code base for "Hypothesis: A modern human range expansion ∼300,000 years ago had a critical role in Neandertal origins": figure2a_movie.py

```
#!/usr/bin/env python3
"""
figure2a_movie.py -- Figure 2a's range expansion as a film. (15 Sep 2026)

DR: "can you make a video of the figure 2a range expansion?"

THE SAME REPLICATE, NOT A NEW ONE. Seed 20098 with s = 0.16, e = 0.179, beta = 0, N = 1,750
and s_mt = 0.000651268: the exact configuration figure2_bands.py draws as panel a, taken
from that script's own defaults rather than retyped, so the four published stills are four
frames of this film. The square convention is figure2_bands' band_square, imported, so the
bars mean here what they mean there -- red from the bottom in proportion to how modern that
marker is, mtDNA / Y / autosomes left to right, area proportional to band size.

WHAT THE FILM SHOWS THAT THE STILLS CANNOT. The four panels are generations 0, 100, 300 and
1,000, and between the second and the third the interesting thing happens: the modern mtDNA
bar goes red across the whole landscape while the Y bar stays blue for another few hundred
generations, and the archaic-identity bands are squeezed into the north-west before they go.
At one frame per 4 generations that reads as a wave rather than as four unrelated maps.

THE COASTLINE IS DRAWN ONCE. contourf on a 1,000-frame film is most of the runtime, so the
landmass goes down as a single collection at setup and only the squares are cleared and
redrawn per frame. The generation counter and the three whole-landscape percentages are
updated in place for the same reason.

    python figure2a_movie.py                 # 1,000 generations, every 4th, -> .mp4
    python figure2a_movie.py --every 2       # smoother, twice the frames
    python figure2a_movie.py --gens 3000     # keep going past the sweep
    python figure2a_movie.py --gif           # also write an animated GIF

RENDERED IN SLICES WHEN THERE ARE A LOT OF FRAMES. One frame costs about 0.4 s, so the 5x
slower cut -- every generation, 1,001 frames -- is six minutes of drawing and has to be done
in pieces. --slice i j renders frames [i, j) into the frame directory and leaves it there;
--assemble turns whatever is in that directory into the film and cleans up. The simulation is
re-run each time and is deterministic from the seed, so a sliced render is frame-for-frame
identical to a single-pass one.

    python figure2a_movie.py --every 1 --fps 20 --out Figure2a_expansion_slow --slice 0 350
    python figure2a_movie.py --every 1 --fps 20 --out Figure2a_expansion_slow --slice 350 700
    python figure2a_movie.py --every 1 --fps 20 --out Figure2a_expansion_slow --slice 700 1001
    python figure2a_movie.py --every 1 --fps 20 --out Figure2a_expansion_slow --assemble
"""
import argparse
import glob
import os
import subprocess
import sys

import matplotlib
matplotlib.use("Agg")
import matplotlib.pyplot as plt
import numpy as np

HERE = os.path.dirname(os.path.abspath(__file__))
BAND = os.path.join(HERE, "band_model")
sys.path.insert(0, BAND)

from bandsim import Config, Landscape, init_state, step, stats      # noqa: E402
from figure2_bands import (band_square, per_band, albers,           # noqa: E402
                           KM_PER_DEG_LAT, COS_REF, BOLD_PT, PLAIN_PT)
import figure2_bands as F2B                                         # noqa: E402

# Panel a's configuration, IMPORTED from figure2_bands so the two cannot drift. These used to
# be hardcoded here under a comment that said they were read from there, and the film was left
# on the superseded s = 0.02 replicate when panel a moved to s = 0.08.
SEED = F2B.PANEL_A["seed"]
S = F2B.PANEL_A["s"]
E = F2B.PANEL_A["e"]
BETA = F2B.PANEL_A["beta"]
S_MT = F2B.PANEL_A["s_mt"]
N = 1750
WB = 0.0125                      # panel a's square side at band size K; area scales with size


def frames(gens, every):
    """[(generation, positions, sizes, per-band fractions, whole-landscape stats)]."""
    cfg = Config(gens=gens + 1, record_every=10 ** 9, beta=BETA, s=S, e=E, r=2.0,
                 seed=SEED, N_target=N, s_mt=S_MT)
    rng = np.random.default_rng(cfg.seed)
    land = Landscape()
    P, pos, b_ident = init_state(cfg, land, rng)
    out = []
    for g in range(gens + 1):
        if g % every == 0:
            alive, size, mtf, yf, auf = per_band(P, b_ident, len(b_ident))
            st = stats(P, b_ident, exclude=tuple(range(32, 36)))
            out.append((g, pos[alive].copy(), size,
                        np.column_stack([mtf, yf, auf]),
                        # WITHIN MODERN-IDENTITY BANDS, as figure2_bands' stills are (it
                        # reads mt_mod / Y_mod / auto_mod). This used the whole-landscape
                        # mt / Y / auto, so the film and the four stills it is supposed to
                        # contain disagreed: at generation 100 the film said mtDNA 5%, Y 85%,
                        # autosomes 5% where the still said 100 / 100 / 15. The mtDNA line is
                        # the giveaway -- under full matrilocality women never leave their
                        # natal band, so every modern-identity band carries modern mtDNA by
                        # construction and the number is 100% at every generation.
                        (st["mt_mod"], st["Y_mod"], st["auto_mod"]),
                        int(b_ident[np.unique(P["band"])].sum()),
                        len(np.unique(P["band"]))))
        res = step(P, pos, b_ident, cfg, land, rng)
        if res is None or res[0] is None:
            print(f"  population lost at generation {g}")
            break
        P, pos, b_ident = res
    return out, land, cfg


def render(fr, land, cfg, outdir, dpi, lo=0, hi=None):
    """One PNG per frame, for frames [lo, hi). The map is drawn once; only the squares move."""
    LON, LAT = np.meshgrid(land.lon, land.lat)
    GX, GY = albers(LON, LAT)
    ii, jj = np.where(land.mask)
    lx, ly = albers(land.lon[jj], land.lat[ii])
    mx, my = 0.02 * (lx.max() - lx.min()), 0.03 * (ly.max() - ly.min())

    fig, ax = plt.subplots(figsize=(7.2, 5.4))
    # ROOM AT THE BOTTOM FOR THE KEY. DR, 28 Sep: "for the Supplementary Video, could you put
    # the key showing how to interpret the three vertical stripes in the box at the bottom?"
    # The film had no key at all -- the stills in Figure 2a carry one and the video did not, so
    # a viewer meeting the film first had no way to read the squares.
    fig.subplots_adjust(left=0.01, right=0.99, top=0.93, bottom=0.135)
    F2B.draw_land(ax, GX, GY, land.mask)
    ax.set_xlim(lx.min() - mx, lx.max() + mx)
    ax.set_ylim(ly.min() - my, ly.max() + my)
    ax.set_aspect("equal")
    ax.set_xticks([]); ax.set_yticks([])
    for sp in ax.spines.values():
        sp.set_visible(False)
    title = ax.set_title("", fontsize=BOLD_PT, weight="bold", pad=6)
    # The ancestry block sits over the north-eastern sea, the one large area of open water in
    # every frame -- the same place panel a puts it, and for the same measured reason.
    txt = ax.text(0.985, 0.975, "", transform=ax.transAxes, ha="right", va="top",
                  fontsize=PLAIN_PT, linespacing=1.45, family="monospace")
    foot = ax.text(0.01, 0.02, "", transform=ax.transAxes, ha="left", va="bottom",
                   fontsize=PLAIN_PT - 2, color="0.35")

    # ---- the key, drawn once: an example square, then the sentence Figure 2a uses ----------
    # SAME WORDING AND SAME ICON as figure2_bands' key, and the icon comes from the same
    # band_square call, so the film cannot end up explaining the squares differently from the
    # figure. Broken over two lines because the sentence is about 8 inches long at 12 pt and
    # the frame is 7.2 inches wide; at 10 pt on two lines it clears with room to spare.
    FW, FH = fig.get_size_inches()
    KEY_PT = PLAIN_PT - 2
    ih = 0.058                                   # icon height, as a fraction of figure height
    iw = ih * FH / FW                            # equal in DISPLAY inches, so it stays square
    GAP = 0.012
    TXT = ("In each square, modern fraction given left-to-right\n"
           "for mtDNA-Y-autosomes. Area is band size")
    # Centre the icon-plus-text group under the map. The text block is measured rather than
    # guessed: a renderer pass gives its width in figure fractions, so the group stays centred
    # if the wording or the font size changes.
    fig.canvas.draw()
    probe = fig.text(0, -1, TXT, fontsize=KEY_PT, linespacing=1.35)
    tw = (probe.get_window_extent(fig.canvas.get_renderer()).width
          / fig.bbox.width)
    probe.remove()
    x0 = 0.5 - (iw + GAP + tw) / 2.0
    ycen = 0.068
    icon = fig.add_axes([x0, ycen - ih / 2, iw, ih])
    icon.set_xlim(0, 1); icon.set_ylim(0, 1); icon.axis("off")
    band_square(icon, 0.5, 0.5, (0.85, 0.5, 0.12), 1.0)
    fig.text(x0 + iw + GAP, ycen, TXT, va="center", ha="left",
             fontsize=KEY_PT, linespacing=1.35)

    os.makedirs(outdir, exist_ok=True)
    hi = len(fr) if hi is None else min(hi, len(fr))
    paths = []
    for k in range(lo, hi):
        g, p, size, fr3, whole, n_mod, n_all = fr[k]
        for coll in list(ax.patches):
            coll.remove()
        lon = p[:, 0] / (KM_PER_DEG_LAT * COS_REF) + land.lon0
        lat = p[:, 1] / KM_PER_DEG_LAT + land.lat0
        X, Y = albers(lon, lat)
        w = np.clip(WB * np.sqrt(size / float(cfg.K)), 0.28 * WB, 1.7 * WB)
        for i in range(len(X)):
            band_square(ax, X[i], Y[i], np.nan_to_num(fr3[i], nan=0.0), w[i])
        mt, y, au = whole
        def pct(v):
            return "  --" if v != v else f"{100 * v:3.0f}%"
        title.set_text(f"generation {g:,}")
        txt.set_text(f"% modern\nmtDNA {pct(mt)}\nY     {pct(y)}\nautos {pct(au)}")
        foot.set_text(f"{n_mod} of {n_all} bands carry the modern identity")
        path = os.path.join(outdir, f"f{k:05d}.png")
        fig.savefig(path, dpi=dpi)
        paths.append(path)
    plt.close(fig)
    return paths


def main():
    ap = argparse.ArgumentParser()
    ap.add_argument("--gens", type=int, default=1000)
    ap.add_argument("--every", type=int, default=4)
    ap.add_argument("--fps", type=int, default=25)
    ap.add_argument("--dpi", type=int, default=110)
    ap.add_argument("--gif", action="store_true")
    ap.add_argument("--out", default="Figure2a_expansion")
    # --mono: black = modern, grey = archaic, pale yellow landmass, for the alternative
    # Figure 2 in which red and blue have been reassigned to the deme and band frameworks.
    # The frame cache is keyed on --out, so give the mono render its own --out or it will
    # reuse the red/blue frames already on disk and silently assemble a mixed film.
    ap.add_argument("--mono", action="store_true")
    ap.add_argument("--slice", nargs=2, type=int, metavar=("LO", "HI"))
    ap.add_argument("--assemble", action="store_true")
    # RETIMING RATHER THAN RE-RENDERING. Once the film is one frame per generation there is
    # nothing finer to sample -- a slower cut can only hold each generation longer -- so
    # "slow it down another 2x" is a change of frame rate, not of frames. --retime takes an
    # existing mp4 and re-encodes it at 1/FACTOR of its rate, which is exact and takes
    # seconds, where re-rendering the 1,001 frames takes six minutes and would produce
    # pixel-identical images.
    ap.add_argument("--retime", nargs=2, metavar=("SRC_MP4", "FACTOR"))
    a = ap.parse_args()
    if a.mono:
        F2B.use_mono()

    if a.retime:
        src, factor = a.retime[0], float(a.retime[1])
        src = src if os.path.exists(src) else os.path.join(HERE, src)
        probe = subprocess.run(["ffprobe", "-v", "error", "-select_streams", "v:0",
                                "-show_entries", "stream=r_frame_rate,nb_frames",
                                "-of", "csv=p=0", src], capture_output=True, text=True)
        rate, nframes = probe.stdout.strip().split(",")
        num, den = (float(x) for x in rate.split("/"))
        fps_in = num / den
        fps_out = fps_in / factor
        dst = os.path.join(HERE, f"{a.out}.mp4")
        r = subprocess.run(["ffmpeg", "-y", "-loglevel", "error", "-i", src,
                            "-vf", f"setpts={factor}*PTS", "-r", f"{fps_out:g}",
                            "-pix_fmt", "yuv420p", "-c:v", "libx264", "-crf", "20", dst],
                           capture_output=True, text=True)
        if r.returncode:
            raise SystemExit("ffmpeg failed:\n" + (r.stderr or "")[:800])
        print(f"retimed {os.path.basename(src)} ({fps_in:g} fps, {nframes} frames) "
              f"to {fps_out:g} fps")
        print(f"wrote {a.out}.mp4   {float(nframes) / fps_out:.1f} s, "
              f"{os.path.getsize(dst) / 1e6:.1f} MB")
        return

    outdir = os.path.join(HERE, f"_f2a_frames_{a.out}")
    if a.assemble:
        paths = sorted(glob.glob(os.path.join(outdir, "f*.png")))
        if not paths:
            raise SystemExit(f"no frames in {outdir} -- render some slices first")
        print(f"assembling {len(paths)} frames from {os.path.basename(outdir)}")
    else:
        print(f"panel a's replicate: seed {SEED}, s={S}, e={E}, beta={BETA}, N={N}, "
              f"s_mt={S_MT:g}")
        fr, land, cfg = frames(a.gens, a.every)
        print(f"  {len(fr)} frames, generation 0 to {fr[-1][0]:,} every {a.every}")
        lo, hi = a.slice if a.slice else (0, None)
        paths = render(fr, land, cfg, outdir, a.dpi, lo, hi)
        print(f"  rendered {len(paths)} frames at {a.dpi} dpi "
              f"({len(os.listdir(outdir))} now in {os.path.basename(outdir)})")
        if a.slice:
            print("  slice only -- run again with --assemble when every slice is done")
            return

    mp4 = os.path.join(HERE, f"{a.out}.mp4")
    cmd = ["ffmpeg", "-y", "-loglevel", "error", "-framerate", str(a.fps),
           "-i", os.path.join(outdir, "f%05d.png"),
           # yuv420p and an even-dimension filter, or QuickTime and PowerPoint refuse it
           "-vf", "pad=ceil(iw/2)*2:ceil(ih/2)*2", "-pix_fmt", "yuv420p",
           "-c:v", "libx264", "-crf", "20", mp4]
    r = subprocess.run(cmd, capture_output=True, text=True)
    if r.returncode:
        print("  !! ffmpeg failed:\n" + (r.stderr or "")[:800])
    else:
        mb = os.path.getsize(mp4) / 1e6
        print(f"wrote {a.out}.mp4   {len(paths) / a.fps:.1f} s at {a.fps} fps, {mb:.1f} MB")

    if a.gif:
        from PIL import Image
        ims = [Image.open(p).convert("P", palette=Image.ADAPTIVE) for p in paths]
        gif = os.path.join(HERE, f"{a.out}.gif")
        ims[0].save(gif, save_all=True, append_images=ims[1:],
                    duration=int(1000 / a.fps), loop=0, optimize=True)
        print(f"wrote {a.out}.gif   {os.path.getsize(gif) / 1e6:.1f} MB")

    # CLEANUP IS BEST-EFFORT. On a read-only or sync-managed directory the frames cannot be
    # deleted, and an unhandled PermissionError here used to abort the run AFTER the film had
    # already been written -- which reads like a failure when the only thing left undone is
    # housekeeping. Report what could not be removed and exit cleanly.
    left = []
    for p in paths:
        try:
            os.remove(p)
        except OSError:
            left.append(p)
    if not left:
        try:
            os.rmdir(outdir)
        except OSError:
            left.append(outdir)
    if left:
        print(f"note: could not delete {len(left)} intermediate file(s) in "
              f"{os.path.basename(outdir)} -- the film is complete; remove the directory "
              f"by hand if you want the space back")


if __name__ == "__main__":
    main()
```
