## Supplementary Information section 3 code base for "Hypothesis: A modern human range expansion ∼300,000 years ago had a critical role in Neandertal origins": figure2c_fountain.py

```
#!/usr/bin/env python3
"""
figure2c_fountain.py -- Figure 2c, one fountain per mechanism setting. (DR, 11 Sep 2026)

DR's revisions to the first draft:
  Route 1  separate fountains for a few s values
  Route 2  separate fountains for a few sex-distortion values
  Route 3  show mt-Y, mt-autosomal-dominant and restorers separately
  Route 4  restrict to complete infertility (theta = 0)
  and show BOTH frameworks, not deme alone.

Each point is one parameter combination that survived all four constraints, plotted at the
probability that ten sampled males are all modern at both markers. Deme and band are
interspersed within each sub-column and separated by colour.

THE JOIN. Expansion duration comes from the stage-1 screen (the generation of archaic
extinction); the autosomal proportion and p_joint come from stage 2. A cell is admitted on its
stage-1 timing and stage-2 autosomal proportion, matched by filename, and plotted at its
stage-2 p_joint.

AN EMPTY COLUMN IS A RESULT. Route 4 at theta = 0 has no surviving deme combinations out of 627
screened, while the band model has some. Drawing the column empty, with the screened count
underneath, says that; dropping the column would hide it.

    python figure2c_fountain.py [--aut-lo 3 --aut-hi 7] [--gen-lo 100 --gen-hi 1000]
"""
import glob
import json
import os
import sys

import matplotlib
matplotlib.use("Agg")
import matplotlib.pyplot as plt
import numpy as np

HERE = os.path.dirname(os.path.abspath(__file__))
import screen  # noqa: E402

CL = os.path.join(HERE, "cluster")

# THE ACCEPTANCE WINDOW, WIDENED FROM 4-6% TO 3-7% ON 12 Sep 2026.
#
# DR: "do you want to change the allowance window for all our plots from 4-6% to 3-7%, matching
# the number from the papers?"
#
# The old window was a symmetric bracket around a "~5%" that appears in neither cited source.
# Hubisz et al. report 3% of the Neandertal genome as putatively introgressed and about 7%
# after correcting for a detection power of 30-55%; Harris et al. report about 6% for the Altai
# Neandertal. So 3-7% is the span the literature actually supports and 4-6% was narrower than
# the evidence warrants.
#
# ONE CAVEAT, STATED SO IT IS NOT FORGOTTEN: 3-7% is the range spanned by two papers' POINT
# estimates, not a confidence interval on a single estimate. It is a defensible acceptance
# region for a screen -- arguably more defensible than the old one, since it does not pretend
# to a precision no source claims -- but it should not be described anywhere as a confidence
# interval on the modern proportion.
#
# WHAT IT CHANGES: admissible combinations go from 4,124 to 9,039, and no column median moves
# by more than 0.03. Route 1 at s = 0.01 goes 0.244 -> 0.219 and Route 2 at d = 0.8 goes
# 1.000 -> 0.995; every other column moves less. So the conclusions are unchanged and the
# sample behind each of them slightly more than doubles.
AUT_LO, AUT_HI = 3.0, 7.0

# Stage-2 two-wave DEME files written before this are void; see the guard in load().
import datetime as _dt
TW_DEME_FIX = _dt.datetime(2026, 9, 13, 0, 0, 0).timestamp()

# THE ROUTE NUMBERS MUST BE THE NOTE'S, AND THEY WERE NOT (13 Sep 2026). DR: "Why do you have
# a 'Route 5' at all? Route 5 isn't referred to in our note anymore." He is right, and the error
# is older than the two-wave work: the note has FOUR routes, because sex distortion and
# epistatic incompatibility were merged into a single route back at v50, and this list still
# separated them. Every column from sex distortion onward therefore carried a label one higher
# than the section it illustrates -- theta appeared as "Route 4" where the note's Route 4 is the
# two-wave scenario, and two waves appeared as a "Route 5" that does not exist in the note.
#
#   Route 1  sex-biased expansion + reduced fitness of archaic lineages of the other sex
#   Route 2  matrilocal expansion + epistatic interaction archaic Y x modern partner alleles,
#            sex distortion included (merged in v50)
#   Route 3  sex-biased expansion + cultural mechanism, F1 of the archaic parent's sex not raised
#   Route 4  matrilocal expansion preceded or followed by a patrilocal one -- two waves
#
# (route label, sub-label, mech, value or None for any)
COLUMNS = [
    ("Route 1\nselection on Y", "0.01", "sY", 0.01),
    ("", "0.02", "sY", 0.02),
    ("", "0.05", "sY", 0.05),
    ("", "0.10", "sY", 0.10),
    ("", "0.20", "sY", 0.20),
    ("Route 2\nincompatibility and\nsex distortion", "0.1", "xdrive", 0.1),
    ("", "0.2", "xdrive", 0.2),
    ("", "0.4", "xdrive", 0.4),
    ("", "0.6", "xdrive", 0.6),
    ("", "0.8", "xdrive", 0.8),
    ("", "mt-Y", "dmi_mtY", None),
    ("", "mt-auto\ndominant", "dmi_mtAutoDom", None),
    ("", "restorers", "restorers", None),
    ("Route 3\nF1 not raised", "θ = 0", "theta", 0.0),
    ("Route 4\ntwo waves", "", "twowave", None),
]
COL = {"deme": "#1f4e79", "band": "#c0504d"}
MARK = {"deme": "o", "band": "^"}


def flag(name, cast, default):
    return cast(sys.argv[sys.argv.index(name) + 1]) if name in sys.argv else default


def load(aut_lo, aut_hi, gen_lo, gen_hi):
    """{(framework, mech, val): [p_joint]} for the ADMISSIBLE cells, plus the screened counts.

    REWRITTEN 14 Sep 2026 TO READ screen.py. DR: "one shared admissibility function: a cell is
    in the box or it isn't, decided once from the best estimate available for that cell, and
    both figures read the answer." This function used to do its own directory walk, its own
    precedence between the screen and the re-screen, its own two-wave clock handling and its
    own K_early injection; Figure S3.3 did the same job separately and the two disagreed on
    1,911 of 10,399 points. All of that now lives in screen.py and both figures call it, so a
    fountain point is by construction a point inside the box S3.3 draws.

    TWO THINGS WENT OUT WITH THE REWRITE, both deliberate:

      * THE K_early CELLS. DR: "K_early (changing density during the expansion) is out! We
        decided that and write that in the text." Route 3's theta = 0 deme column drops from
        11 admissible combinations to 1 and theta = 0.02 loses about 19 of 42. That is the
        honest number for a screen run with constant K, which is what routes_build declares.
      * THE STAGE-1 AUTOSOMAL FILTER. Admissibility is now decided on the best estimate of
        each quantity rather than on stage-1 duration crossed with stage-2 ancestry.
    """
    keep, screened = {}, {}
    for key, cells in screen.scan().items():
        screened[key] = len(cells)
        p = [c["joint"] for c in cells
             if screen.in_box(c, aut_lo, aut_hi, gen_lo, gen_hi) and c.get("joint") is not None]
        if p:
            keep[key] = p
    return keep, screened


def pick(store, fw, mech, val):
    """val None means AGGREGATE over every value of that mechanism.

    Route 3's columns are mechanisms rather than parameter settings -- mt-Y and
    mt-autosomal-dominant have a single penetrance, and `restorers` spans 1 to 4 loci -- so
    those columns pool. Keying them on a literal None silently matched nothing, which is why
    the first build of this figure showed Route 3 as 0 of 0 rather than as data.
    """
    if val is not None:
        return store.get((fw, mech, val), [] if isinstance(store.get((fw, mech, val), []), list)
                         else 0)
    out = []
    tot = 0
    for (f, m, v), x in store.items():
        if f == fw and m == mech:
            if isinstance(x, list):
                out.extend(x)
            else:
                tot += x
    return out if out or not tot else tot


def main():
    aut_lo, aut_hi = flag("--aut-lo", float, AUT_LO), flag("--aut-hi", float, AUT_HI)
    gen_lo, gen_hi = flag("--gen-lo", float, 100.0), flag("--gen-hi", float, 1000.0)
    keep, screened = load(aut_lo, aut_hi, gen_lo, gen_hi)

    fig, ax = plt.subplots(figsize=(12.4, 4.8))
    rng = np.random.default_rng(0)
    counts = []

    for i, (route, sub, mech, val) in enumerate(COLUMNS):
        tot_keep = tot_scr = 0
        for fw in ("deme", "band"):
            pts = pick(keep, fw, mech, val)
            pts = pts if isinstance(pts, list) else []
            scr = pick(screened, fw, mech, val)
            scr = scr if isinstance(scr, int) else (screened.get((fw, mech, val), 0)
                                                    if val is not None else
                                                    sum(v for (f, m, _), v in screened.items()
                                                        if f == fw and m == mech))
            tot_keep += len(pts)
            tot_scr += scr
            if not pts:
                continue
            x = i + rng.uniform(-0.20, 0.20, len(pts)) + (0.09 if fw == "band" else -0.09)
            ax.scatter(x, pts, s=22, marker=MARK[fw], c=COL[fw], alpha=0.72,
                       edgecolors="white", linewidths=0.4, zorder=3,
                       label=fw if i == 0 else None)
        allp = [v for fw in ("deme", "band")
                for v in (pick(keep, fw, mech, val) if isinstance(pick(keep, fw, mech, val), list) else [])]
        if allp:
            med = float(np.median(allp))
            ax.plot([i - 0.3, i + 0.3], [med, med], color="0.25", lw=1.8, zorder=4)
        else:
            ax.text(i, 0.5, "none\nsurvive", ha="center", va="center", fontsize=7,
                    color="0.45", style="italic")
        counts.append((tot_keep, tot_scr))
        if route:
            ax.axvline(i - 0.5, color="0.85", lw=0.9, zorder=0)

    ax.set_xticks(range(len(COLUMNS)))
    ax.set_xticklabels([c[1] for c in COLUMNS], fontsize=8)
    for i, (route, _, _, _) in enumerate(COLUMNS):
        if route:
            ax.text(i, -0.20, route, ha="left", va="top", fontsize=9, fontweight="bold",
                    transform=ax.get_xaxis_transform(), linespacing=1.4)
    ax.set_xlim(-0.6, len(COLUMNS) - 0.4)
    ax.set_ylim(-0.05, 1.08)
    ax.set_ylabel("P(10 sampled males all modern\nat both mtDNA and Y)")
    ax.axhline(0.5, color="0.88", lw=0.8, zorder=1)
    ax.grid(axis="y", color="0.94", lw=0.7)
    ax.set_axisbelow(True)
    ax.legend(fontsize=8, loc="lower left", framealpha=0.95, title="framework",
              title_fontsize=8)
    ax.set_title(f"Parameter combinations matching autosomal {aut_lo:g}-{aut_hi:g}%, expansion "
                 f"{gen_lo:g}-{gen_hi:g} generations, establishment > 50%", fontsize=10)
    fig.text(0.5, 0.008, "surviving / scored per column:  " +
             "  ".join(f"{k}/{n}" for k, n in counts), ha="center", fontsize=7,
             color="0.35")
    fig.tight_layout(rect=(0, 0.13, 1, 1))
    for ext in ("png", "pdf"):
        fig.savefig(os.path.join(HERE, f"Figure2c_fountain.{ext}"), dpi=200)
    print("  wrote Figure2c_fountain.png / .pdf")
    for (route, sub, mech, val), (k, n) in zip(COLUMNS, counts):
        lab = (route.split("\n")[0] + " " if route else "   ") + sub.replace("\n", " ")
        dd = pick(keep, "deme", mech, val); bb = pick(keep, "band", mech, val)
        d = len(dd) if isinstance(dd, list) else 0
        b = len(bb) if isinstance(bb, list) else 0
        print(f"    {lab:<28s} keep {k:4d} of {n:4d}   (deme {d:3d}, band {b:3d})")


if __name__ == "__main__":
    main()
```
