## Supplementary Information section 3 code base for "Hypothesis: A modern human range expansion ∼300,000 years ago had a critical role in Neandertal origins": make_si_figures.py

```
#!/usr/bin/env python3
"""
make_si_figures.py -- Figures S3.1, S3.2, S3.4 and S3.5 rebuilt with both frameworks.
(DR, 6 Sep 2026. Figure S3.3 has its own script, make_figS3_3.py, because the deme model
produces a null result there and the figure had to change shape.)

Each figure puts the deme curve and the band curve on the same axes. Where the two knobs are
not the same quantity the caption has to say so, and the two places that applies are noted in
the code below rather than left for a reader to infer:

  S3.1 left   deme sweeps female_bias, which is the probability that a cross-lineage child
              takes its MOTHER's lineage. The band sweeps beta, the female share of exogamy --
              who physically leaves. The extremes correspond (deme fb=1 is band beta=0, both
              matrilocal) but intermediate values are only loosely comparable, and the band
              transition is much sharper.
  S3.1 right  deme sweeps the MALE fraction of the pulse; band pulse_beta is the FEMALE share.
              The band axis is reversed relative to the deme one.

    python make_si_figures.py
"""
import csv
import os

import numpy as np
import matplotlib
matplotlib.use("Agg")
import matplotlib.pyplot as plt

HERE = os.path.dirname(os.path.abspath(__file__))
BAND = os.path.join(HERE, "band_model")
DEME_C, BAND_C = "#2166ac", "#b2182b"
WIN = (4.0, 6.0)


def rd(p):
    return list(csv.DictReader(l for l in open(p) if not l.startswith("#")))


def bandrows(f, fam):
    return [r for r in rd(os.path.join(BAND, f)) if r["family"] == fam]


def style(ax, xlab, ylab, title):
    ax.set_xlabel(xlab, fontsize=9.5)
    ax.set_ylabel(ylab, fontsize=9.5)
    ax.set_title(title, fontsize=9.8, weight="bold")
    ax.grid(alpha=0.25, lw=0.6)
    ax.tick_params(labelsize=8.5)


# ---------------------------------------------------------------- S3.1
def fig_s31():
    fig, ax = plt.subplots(1, 2, figsize=(10.4, 4.2))

    d = {float(r["female_bias"]): (float(r["mt_mean"]), float(r["Y_mean"]))
         for r in rd(os.path.join(HERE, "grid5_summary_2-15pct.csv"))
         if r["gamma"] == "0.12" and r["m"] == "0.1"}
    b = {float(r["beta"]): (100 * float(r["mt"]), 100 * float(r["Y"]))
         for r in bandrows("bandsi_summary.csv", "beta")}
    x = sorted(d)
    ax[0].plot(x, [d[v][0] for v in x], "o-", color=DEME_C, ms=4.5, lw=1.7, label="deme, mtDNA")
    ax[0].plot(x, [d[v][1] for v in x], "s--", color=DEME_C, ms=4.5, lw=1.4, alpha=0.65,
               label="deme, Y")
    # band beta is reversed relative to deme female_bias: beta = 0 is matrilocal = fb 1
    xb = sorted(b)
    ax[0].plot([1 - v for v in xb], [b[v][0] for v in xb], "o-", color=BAND_C, ms=4.5, lw=1.7,
               label="band, mtDNA")
    ax[0].plot([1 - v for v in xb], [b[v][1] for v in xb], "s--", color=BAND_C, ms=4.5, lw=1.4,
               alpha=0.65, label="band, Y")
    style(ax[0], "matrilocality  (deme: female bias;  band: 1 - $\\beta$)",
          "modern ancestry at the marker (%)", "range expansion")
    ax[0].legend(fontsize=7.6, loc="upper center", ncol=2, framealpha=0.95)
    ax[0].set_ylim(-4, 108)

    d2 = {float(r["male_frac"]): (float(r["P_mt_fixed"]), float(r["P_Y_fixed"]))
          for r in rd(os.path.join(HERE, "sexbias_stable_boundary.csv"))
          if r["census_N"] == "1500"}
    b2 = {float(r["pulse_beta"]): (100 * float(r["p_mt"]), 100 * float(r["p_Y"]))
          for r in bandrows("bandsi_summary.csv", "pulsebeta") if r["N"] == "1750"}
    xf = sorted(b2)                      # female share
    ax[1].plot([1 - v for v in sorted(d2)], [d2[v][0] for v in sorted(d2)], "o-",
               color=DEME_C, ms=4.5, lw=1.7, label="deme, mtDNA")
    ax[1].plot([1 - v for v in sorted(d2)], [d2[v][1] for v in sorted(d2)], "s--",
               color=DEME_C, ms=4.5, lw=1.4, alpha=0.65, label="deme, Y")
    ax[1].plot(xf, [b2[v][0] for v in xf], "o-", color=BAND_C, ms=4.5, lw=1.7,
               label="band, mtDNA")
    ax[1].plot(xf, [b2[v][1] for v in xf], "s--", color=BAND_C, ms=4.5, lw=1.4, alpha=0.65,
               label="band, Y")
    # THE DOTTED GREY LINE AND ITS LABEL ARE GONE. DR, comment 20: "remove the dotted gray line
    # and 'neutral expectation' caption on the right panel." The line marked 10%, the neutral
    # fixation probability for modern mtDNA when an all-female 5% pulse puts it in 10% of
    # transmitting females. That is worth stating, but it is a fact about one end of one curve
    # and the caption already says it, so drawing it across the whole panel implied a baseline
    # for every series -- including the Y, whose neutral expectation under an all-female pulse
    # is zero, not 10%.
    style(ax[1], "female share of the migration pulse",
          "P(marker fixes) (%)", "migration pulse")
    ax[1].legend(fontsize=7.6, loc="upper center", ncol=2, framealpha=0.95)
    ax[1].set_ylim(-1.5, 34)

    for i, L in enumerate("ab"):
        fig.text(0.012 + 0.5 * i, 0.965, L, fontsize=15, weight="bold")
    fig.tight_layout(rect=(0.01, 0, 1, 0.965))
    fig.savefig(os.path.join(HERE, "figS3_1_both_frameworks.png"), dpi=170, bbox_inches="tight")
    print("  wrote figS3_1_both_frameworks.png")


# ---------------------------------------------------------------- S3.2
def fig_s32():
    fig, ax = plt.subplots(figsize=(5.8, 4.3))
    rows = rd(os.path.join(HERE, "fixation_curves_mtdna.csv"))
    for ne, ls, lab in ((500, ":", "500"), (750, "-", "750 (anchor)"), (1000, "--", "1000")):
        s = sorted([r for r in rows if int(r["locus_Ne"]) == ne], key=lambda r: float(r["s"]))
        ax.plot([float(r["s"]) for r in s], [float(r["pfix_pct"]) for r in s], ls,
                color=DEME_C, lw=1.8, label=f"deme, locus $N_e$ = {lab}")
    for ne, ls in ((1500, (0, (5, 1, 1, 1))), (3000, (0, (1, 1)))):
        s = sorted([r for r in rows if int(r["locus_Ne"]) == ne], key=lambda r: float(r["s"]))
        if s:
            ax.plot([float(r["s"]) for r in s], [float(r["pfix_pct"]) for r in s], ls=ls,
                    color="#7f7f7f", lw=1.3,
                    label=f"deme, no contraction ($N_e$ = {ne})")
    b = sorted(bandrows("bandsi_summary.csv", "mtcurve"), key=lambda r: float(r["s_mt"]))
    b = [r for r in b if r["N"] == "1750"]
    ax.plot([float(r["s_mt"]) for r in b], [100 * float(r["p_mt"]) for r in b], "o-",
            color=BAND_C, ms=4, lw=1.8, label="band, N = 1750")
    ax.axhline(50, color="grey", lw=0.8, ls=":")
    style(ax, "selection coefficient on modern mtDNA, $s$",
          "P(modern mtDNA fixes) (%)", "migration pulse, 5% unbiased")
    ax.legend(fontsize=7.4, loc="upper left", framealpha=0.95)
    ax.set_xlim(-0.006, 0.032)
    fig.tight_layout()
    fig.savefig(os.path.join(HERE, "figS3_2_both_frameworks.png"), dpi=170, bbox_inches="tight")
    print("  wrote figS3_2_both_frameworks.png")


# ---------------------------------------------------------------- S3.4
def fig_s34():
    fig, ax = plt.subplots(1, 2, figsize=(10.2, 4.2))
    d = sorted(rd(os.path.join(HERE, "suppfig1_data.csv")), key=lambda r: float(r["s"]))
    b = sorted(bandrows("bandmech_summary.csv", "sf1"), key=lambda r: float(r["s"]))
    ax[0].plot([float(r["s"]) for r in d], [float(r["mt_success"]) for r in d], "o-",
               color=DEME_C, ms=4.5, lw=1.8, label="deme")
    ax[0].plot([float(r["s"]) for r in b], [100 * float(r["mt"]) for r in b], "s-",
               color=BAND_C, ms=4.5, lw=1.8, label="band")
    ax[0].axhline(50, color="grey", lw=0.8, ls=":")
    style(ax[0], "selection on the partner locus, $s$",
          "modern mtDNA retained (%)", "does the mechanism retain mtDNA?")
    ax[0].legend(fontsize=8, loc="upper left")

    ax[1].axhspan(*WIN, color="#cfe3f2", alpha=0.8, zorder=0, label="observed autosomal level")
    ax[1].plot([float(r["s"]) for r in d], [float(r["autobg_success"]) for r in d], "o-",
               color=DEME_C, ms=4.5, lw=1.8, label="deme")
    ax[1].plot([float(r["s"]) for r in b], [100 * float(r["aut"]) for r in b], "s-",
               color=BAND_C, ms=4.5, lw=1.8, label="band")
    style(ax[1], "selection on the partner locus, $s$",
          "background autosomal ancestry (%)", "what it costs on the autosomes")
    ax[1].legend(fontsize=8, loc="upper left")

    for i, L in enumerate("ab"):
        fig.text(0.012 + 0.5 * i, 0.965, L, fontsize=15, weight="bold")
    fig.tight_layout(rect=(0.01, 0, 1, 0.965))
    fig.savefig(os.path.join(HERE, "figS3_4_both_frameworks.png"), dpi=170, bbox_inches="tight")
    print("  wrote figS3_4_both_frameworks.png")


# ---------------------------------------------------------------- S3.5
def fig_s35():
    fig, ax = plt.subplots(1, 2, figsize=(10.2, 4.2))
    x2 = rd(os.path.join(HERE, "xdrv2_summary.csv"))
    T = sorted([r for r in x2 if r["block"] == "T"], key=lambda r: float(r["d"]))
    bt = sorted([r for r in bandrows("bandmech_summary.csv", "xdrv")
                 if r.get("cost") == "0"], key=lambda r: float(r["d"]))
    ax[0].plot([float(r["d"]) for r in T], [float(r["P_all10"]) for r in T], "o-",
               color=DEME_C, ms=4.5, lw=1.8, label="deme")
    ax[0].plot([float(r["d"]) for r in bt], [100 * float(r["p_Y"]) for r in bt], "s-",
               color=BAND_C, ms=4.5, lw=1.8, label="band")
    style(ax[0], "drive strength $d$", "P(all 10 sampled Y modern) (%)",
          "how weak can the drive be?")
    ax[0].legend(fontsize=8, loc="lower right")
    ax[0].set_ylim(-4, 104)

    R = sorted([r for r in x2 if r["block"] == "R" and float(r["d"]) == 0.95],
               key=lambda r: float(r["par"]))
    br = sorted([r for r in bandrows("bandmech_summary.csv", "xdrv") if r.get("res")],
                key=lambda r: float(r["res"]))
    ax[1].plot([float(r["par"]) for r in R], [float(r["Y"]) for r in R], "o-",
               color=DEME_C, ms=4.5, lw=1.8, label="deme, modern Y")
    # DR comment 31 (10 Sep 2026): "remove the dotted line on the right". The dotted series was
    # the resistant archaic Y, which is just 100 minus the modern Y on the same axes and so
    # carried no information the solid curve did not already give.
    ax[1].plot([float(r["res"]) for r in br], [100 * float(r["Y"]) for r in br], "s-",
               color=BAND_C, ms=4.5, lw=1.8, label="band, modern Y")
    ax[1].set_xscale("log")
    style(ax[1], "heritable resistance rate $\\epsilon$", "frequency among males (%)",
          "resistance erodes it")
    ax[1].legend(fontsize=7.6, loc="center left")
    ax[1].set_ylim(-4, 104)

    for i, L in enumerate("ab"):
        fig.text(0.012 + 0.5 * i, 0.965, L, fontsize=15, weight="bold")
    fig.tight_layout(rect=(0.01, 0, 1, 0.965))
    fig.savefig(os.path.join(HERE, "figS3_5_both_frameworks.png"), dpi=170, bbox_inches="tight")
    print("  wrote figS3_5_both_frameworks.png")


if __name__ == "__main__":
    for f in (fig_s31, fig_s32, fig_s34, fig_s35):
        try:
            f()
        except Exception as e:
            print(f"  {f.__name__} FAILED: {type(e).__name__}: {e}")
```
