## Supplementary Information section 3 code base for "Hypothesis: A modern human range expansion ∼300,000 years ago had a critical role in Neandertal origins": README.md

# Simulation code for Supplementary Information section 3

*Plausible mechanisms for Neandertals inheriting modern ancestry at both mtDNA and Y*

Generated by `code_release.py` on 2026-10-10. Python 3.9 or later; `pip install -r requirements.txt`.

This archive contains only code that produces a number, figure or table in the note. Superseded pipelines have been removed, and the Code and Software Availability statement in the note names every file here and nothing else -- `code_release.py --check` verifies that correspondence.

The layout mirrors the working tree the scripts were written in, because each one resolves its inputs relative to its own location: the figure and analysis scripts sit at the top level, the deme pipeline in `cluster/`, the band pipeline in `band_model/`. Extract and run in place. The role headings below are for navigation; they are not directories.

## Reproducing the figures

```
python cluster/ne_screen.py                 # which geometries are admissible
ROUTE_SET=main python cluster/routes_submit.py 1
ROUTE_SET=main python cluster/routes_submit.py 2
python cluster/rescreen_submit.py re --chain

python figure2_full.py                      # main-text Figure 2
python fig_fixation_gaps.py                 # Figure S3.4
python figED_thumbnails.py                  # Figure S3.2, the thumbnail survey
python fig_s34_drive_and_escape.py          # Figure S3.3
python fig_sensitivity.py                   # Figures S3.5 and S3.6
python figED_sensitivity.py                 # Extended Data, stacked sensitivity
python figED_expansions.py                  # Extended Data, sample expansions
python figure2a_movie.py                    # the supplementary video of Figure 2a

python audit_v66.py                         # recompute every number the note states
```

## engines

Selection on the Y (Scenario 2) is conditioned on group identity in both engines (10 Oct 2026): the patriline matching the identity of the band (bandsim) or lineage label (spatial_sim2/3) is favoured and the mismatch penalised. The Cartesian `sY` cells and the band `fexp, s` family were rerun under this rule; `ROUTE_ONLY=sY` in routes_build.py is the switch that restricted the rerun to them.

The five simulation engines. The deme and band engines are the two independent idealisations of space whose agreement is the note's robustness argument; the panmictic engine is the migration-pulse comparator; ysupp_sim is the closed-population model behind the Y-suppressor calculation.

- `cluster/spatial_sim2.py` — deme range-expansion engine (Appendix A)
- `cluster/spatial_sim3.py` — deme engine extended with the mechanism loci
- `band_model/bandsim.py` — roving-band engine (Appendix B)
- `band_model/make_range_mask.py` — builds the habitable arena from the niche reconstruction
- `cluster/panmictic_sim.py` — Wright-Fisher migration-pulse engine
- `cluster/ysupp_sim.py` — X-distorter / Y-suppressor model, Figure S3.3 right panel

## anchors

Everything that fixes a population size or a calibrated knob before any mechanism is switched on. The PSMC anchor of 1,447 diploids is measured here, and both frameworks are tuned to it.

- `digitize_psmc.py` — digitises the PSMC curves and takes the time-weighted harmonic mean (1,447)
- `cluster/anchors.py` — the single source of truth for census and effective sizes
- `cluster/ne_screen.py` — measures the effective size of every (grid, K, m) triple; writes ne_screen.csv
- `band_model/band_ne.py` — realised uniparental Ne for the band model by the neutral-tracer method
- `band_model/band_cal.py` — fits the exogamy rate e that puts autosomal ancestry at the target
- `band_model/bandcal_submit.py` — SLURM array for the band calibration

## search

The Cartesian search that Figure 2b and Figure S3.2 (which also serves as the thumbnail Extended Data figure) are drawn from. routes_build defines the mechanism sets and the demographic grid; the submitters emit tasks; robust_run runs one cell at stage 1 (screen to archaic extinction) or stage 2 (score to marker absorption); screen.py decides admissibility once, for everything.

- `cluster/routes_build.py` — mechanism sets and the demographic grid
- `cluster/routes_submit.py` — the main SLURM array
- `cluster/rescreen_submit.py` — the 100-replicate re-screen of borderline cells
- `cluster/cultural_submit.py` — the five cultural mechanisms, on the same grid
- `cluster/xdrvdeme_submit.py` — the anchored X-drive ladder, gamma refitted per rung
- `cluster/nexp_band_submit.py` — the unbiased-expansion series, band side
- `cluster/demog_fill_submit.py` — fills the rungs the unbiased series was missing
- `cluster/robust_run.py` — the worker: one cell, either framework, either stage
- `cluster/robust_submit.py` — generic array writer for the worker
- `cluster/robust_collect.py` — pools the per-cell JSON
- `screen.py` — the single definition of admissibility, read by every figure

## twowave

Route 3. The second wave is founded under the opposite residence rule, and the band arm has to be timed on the genetic clock rather than the label clock, which clock_check establishes.

- `cluster/twowave_run.py` — two-wave runner, deme
- `cluster/twowave_band.py` — two-wave runner, band
- `cluster/tw5_run.py` — the scored two-wave grid
- `cluster/tw5_submit.py` — its SLURM array
- `cluster/tw6_run.py` — the band two-wave grid on the genetic clock
- `cluster/tw6_submit.py` — its SLURM array
- `cluster/clock_check.py` — shows the label and genetic clocks differ only for two waves

## pulse

The migration-pulse comparator in all its forms: neutral, selected, sex-biased, and the two-marker joint pulse. These are what Figure S3.1's right panel and Figure S3.2's solid curves are made of, and what the note contrasts the range expansion against.

- `cluster/panm4_run.py` — sex-biased pulse and the stable-boundary comparator
- `cluster/panm4_submit.py` — its array
- `cluster/panm4_collect.py` — writes sexbias_stable_boundary.csv
- `cluster/joint_run.py` — the joint mtDNA-and-Y pulse
- `cluster/joint_submit.py` — its array
- `cluster/joint_collect.py` — writes joint_summary.csv
- `cluster/joint2_submit.py` — the second joint series
- `cluster/joint2_collect.py` — writes joint2_summary.csv
- `cluster/jcal_submit.py` — calibrates the joint series
- `cluster/jcal_collect.py` — writes joint_calibration.csv
- `cluster/mtypulse_submit.py` — the mtDNA x Y pulse
- `cluster/twpulse_submit.py` — the two-wave pulse comparator
- `cluster/jdrive_submit.py` — the pulse with X-linked drive
- `cluster/mtexp_submit.py` — modern mtDNA under a patrilineal range expansion

## sweeps

Four named sweeps that are read directly by a figure rather than through the Cartesian search, and two analyses the note reports without showing a figure.

- `cluster/grid5_run.py` — the residence-rule sweep, Figure S3.1 left panel
- `cluster/grid5_submit.py` — its array
- `cluster/grid5_collect.py` — writes grid5_summary_2-15pct.csv
- `cluster/scan4_run.py` — one-at-a-time sensitivity around the deme baseline
- `cluster/scan4_submit.py` — its array
- `cluster/scan4_collect.py` — writes figure3_full.csv, the source of Figure S3.5
- `cluster/xdrv2_run.py` — heritable resistance to the X-drive
- `cluster/xdrv2_submit.py` — its array
- `cluster/xdrv2_collect.py` — pools it; Figure S3.4's right panel reads the raw files
- `cluster/sf1_run.py` — the patrilineal mtDNA incompatibility (figure not shown)
- `cluster/sf1_submit.py` — its array
- `cluster/sf1_collect.py` — writes suppfig1_data.csv
- `cluster/esc_run.py` — the heritable-escape stress test of the mtDNA x Y incompatibility
- `cluster/esc_submit.py` — its array
- `cluster/esc_collect.py` — pools it

## band

The band framework's own production pipeline: one worker, and two submit/collect pairs for the sensitivity sweeps and the mechanism sweeps.

- `band_model/band_reps.py` — the band worker: one condition, many replicates
- `band_model/band_suite.py` — shared band constants, imported by the figures
- `band_model/bandsi_submit.py` — sensitivity, beta, pulse-beta and mtDNA-curve families
- `band_model/bandsi_collect.py` — writes bandsi_summary.csv
- `band_model/bandmech_submit.py` — the mechanism sweeps in the band framework
- `band_model/bandmech_collect.py` — writes bandmech_summary.csv
- `band_model/bandprod_submit.py` — calibrates e* per scenario; writes bandprod_calibration.csv
- `band_model/bandprod_collect.py` — pools that calibration
- `band_model/bandratchet_submit.py` — the production ensemble under the ratchet posterior
- `band_model/bandratchet_collect.py` — writes band_ratchet_summary.csv

## figures

One script per figure in the note and the main text, each regenerating from the simulation output rather than from a transcribed table.

- `figure2_full.py` — assembles main-text Figure 2 from panels (a) and (b)
- `band_model/figure2_bands.py` — Figure 2a, the four band maps
- `band_model/figure2_seedscan.py` — chooses the replicate Figure 2a shows
- `band_model/site_stripes_figure.py` — the Albers projection, imported by Figure 2a
- `figure2_merged.py` — Figure 2b, the four routes
- `figure2c_fountain.py` — supplies the admissible Cartesian combinations both panels draw
- `make_si_figures.py` — Figure S3.1, both panels
- `fig_fixation_gaps.py` — Figure S3.4, the fixation curves with the two estimated gaps
- `fig_thumbnails_all.py` — draws the thumbnail panels and their layout
- `figED_thumbnails.py` — sets the block order and scenario numbering; the two together draw Figure S3.2, which also serves as the thumbnail Extended Data figure
- `fig_s34_drive_and_escape.py` — Figure S3.3, the drive ladder and the resistance ladder
- `fig_sensitivity.py` — Figures S3.5 and S3.6
- `figED_sensitivity.py` — stacks S3.5 and S3.6 into one Extended Data figure
- `figED_expansions.py` — the sample expansions
- `figure2a_movie.py` — renders the Figure 2a replicate as the supplementary video

## analysis

What turns the simulation output into the numbers the note states, and what checks them.

- `collect_mt_expansion.py` — writes mt_expansion_curve.csv, Figure S3.4's range-expansion arms
- `build_supp_workbook.py` — assembles Supplementary Table 1 (15 tabs)
- `cluster/summary_cache.py` — gathers every out_*/*.json summary once, in parallel, for the workbook builder
- `audit_v66.py` — recomputes every simulation number the note states
- `reps_audit.py` — recounts the replicates behind every two-framework figure

## data

The only files the code reads that it does not itself produce in the course of a run.

- `cluster/ne_screen.csv` — the measured effective size of every geometry
- `fixation_curves_mtdna.csv` — the pulse fixation table, produced by section 1
- `mt_gap_posterior_pooled_rna.txt` — the RNA-inclusive mtDNA fitness-gap posterior, produced by section 1 and sampled by joint_run.py
- `band_model/range_mask.npz` — the habitable arena as a boolean lattice
- `band_model/bandprod_calibration.csv` — the calibrated exogamy rates, read as an input by three scripts
