## Supplementary Information section 3 code base for "Hypothesis: A modern human range expansion ∼300,000 years ago had a critical role in Neandertal origins": reps_audit.py

```
#!/usr/bin/env python3
"""
reps_audit.py -- where do the two frameworks report DIFFERENT replicate counts? (13 Sep 2026)

DR: "go through all the figures in the note where the deme and band have different numbers of
reps shown, and put in runs that replace the lower count with a higher one so we can say the
same number of reps for each."

WHY THIS IS WORTH A SCRIPT RATHER THAN A ONE-OFF GREP. The replicate count is not a cosmetic
detail in a figure where the two frameworks are drawn on the same axes: a band curve at 400
replicates and a deme curve at 1,000 have standard errors differing by a factor of 1.6, so the
band arm looks noisier, and a reader comparing the two is partly reading the sampling effort
rather than the models. The counts are also scattered across four different storage formats --
per-cell JSON with a `reps` field, two collected CSVs with an `n` column, a third with
`n_replicates`, and a fourth with `reps` meaning ESTABLISHED replicates rather than attempted
-- so the only way to keep the claim "the same number for each" true after the next rerun is
to recompute it rather than to remember it.

WHAT IT FOUND, and note that three of the four are band-low, which is the expected direction:
the band engine costs roughly 45 s per replicate against the deme engine's 2 s, so every
earlier decision about how many to run went the same way.

    figure   what                        deme            band           verdict
    S3.2     migration pulse             1,000           400            band low
    S3.2     patrilocal expansion        5 combos x 200  2 combos x 200  band low, in COMBOS
    S3.5/6   one-at-a-time sensitivity   2,000 *         400            band low
    S3.3/2b  two-wave expansion          200 (tw2)       50 (tw6)       band low

    * and RAGGED WITHIN THE DEME ARM: 55 of the 60 scan4 points are at 2,000 and five are at
      1,500 -- gamma 0.42, 0.55, 0.70, 1.00 and grid 14, which are exactly the rungs added on
      12 Sep to fill the open grey circles. The caption says 1,500 for all of them, so it is
      wrong about the 55 as well as silent about the split.

THE EXPANSION ROW IS A DIFFERENT KIND OF MISMATCH and cannot be fixed by adding replicates.
Both frameworks ran 200 replicates per parameter combination; the deme arm pools five
admissible combinations and the band arm two, so the deme curve rests on 1,000 draws and the
band curve on 400. Raising the band arm means finding three more admissible band demographies,
or running its two at 500 each and saying so. That is a judgement about what the point
represents, not arithmetic, which is why this script reports it and does not queue it.

    python reps_audit.py            # the table
    python reps_audit.py --cmds     # the table, then the commands that level it up
"""
import csv
import collections
import glob
import json
import os
import sys

HERE = os.path.dirname(os.path.abspath(__file__))
CL = os.path.join(HERE, "cluster")
BAND = os.path.join(HERE, "band_model")

# Measured per-replicate seconds, from sacct. The band engine is ~20x the deme engine, which is
# both why the counts drifted apart and why levelling them up is not free.
SEC = {"deme": 2.0, "band": 45.0}


def _rows(path):
    """A CSV that may carry leading '#' provenance lines."""
    if not os.path.exists(path):
        return []
    return list(csv.DictReader([l for l in open(path) if not l.startswith("#")]))


def _json_reps(pattern):
    c = collections.Counter()
    for p in glob.glob(os.path.join(CL, pattern)):
        try:
            s = json.load(open(p))
            s = s.get("summary", s)
        except Exception:
            continue
        r = s.get("reps") or s.get("n")
        if r:
            c[int(r)] += 1
    return c


def _csv_reps(path, family=None, col="n"):
    c = collections.Counter()
    for r in _rows(path):
        if family is not None and r.get("family") != family:
            continue
        v = r.get(col)
        if v:
            c[int(float(v))] += 1
    return c


def fmt(c):
    if not c:
        return "no data"
    return ", ".join(f"{k:,}x{v}" for k, v in sorted(c.items(), reverse=True))


def main():
    pulse_d = _csv_reps(os.path.join(HERE, "fixation_curves_mtdna.csv"), col="n_replicates")
    pulse_b = _csv_reps(os.path.join(BAND, "bandsi_summary.csv"), "mtcurve")
    scan_d = _csv_reps(os.path.join(HERE, "figure3_full.csv"))
    scan_b = _csv_reps(os.path.join(BAND, "bandsi_summary.csv"), "scanb")
    tw_d = _json_reps("out_tw2_deme/*.json")
    tw_b = _json_reps("out_tw6_band/*.json")

    exp = _rows(os.path.join(HERE, "mt_expansion_curve.csv"))
    exp_d = sum(int(r["combos"]) for r in exp if r["framework"] == "deme")
    exp_b = sum(int(r["combos"]) for r in exp if r["framework"] == "band")
    nd = len([r for r in exp if r["framework"] == "deme"]) or 1
    nb = len([r for r in exp if r["framework"] == "band"]) or 1

    print(f"\n{'figure':<10}{'quantity':<30}{'deme':<26}{'band':<26}verdict")
    print("-" * 108)
    rows = [
        ("S3.2", "migration pulse", fmt(pulse_d), fmt(pulse_b)),
        ("S3.2", "patrilocal expansion",
         f"{exp_d // nd} combos x 200", f"{exp_b // nb} combos x 200"),
        ("S3.5/6", "one-at-a-time sensitivity", fmt(scan_d), fmt(scan_b)),
        ("S3.3/2b", "two-wave expansion", fmt(tw_d), fmt(tw_b)),
    ]
    for fig, q, d, b in rows:
        def top(s):
            return int(s.split("x")[0].replace(",", "").split()[0]) if s != "no data" else 0
        v = "MATCHED" if top(d) == top(b) else ("band low" if top(b) < top(d) else "deme low")
        print(f"{fig:<10}{q:<30}{d:<26}{b:<26}{v}")

    # The within-deme raggedness is not a framework mismatch and would be invisible above,
    # since the table prints the modal count. It still breaks "the same number for each".
    if len(scan_d) > 1:
        lo = min(scan_d)
        pts = [f"{r['panel']} {r['value']}" for r in _rows(os.path.join(HERE, "figure3_full.csv"))
               if r.get("n") and int(r["n"]) == lo]
        print(f"\n  within the DEME sensitivity arm: {scan_d[lo]} of "
              f"{sum(scan_d.values())} points are at {lo:,} rather than {max(scan_d):,} -- "
              + ", ".join(pts))

    if "--cmds" not in sys.argv:
        print("\n  (--cmds for the levelling runs)")
        return

    tgt_scan, tgt_pulse, tgt_tw = max(scan_d or [2000]), max(pulse_d or [1000]), max(tw_d or [200])
    n_scan, n_pulse, n_tw = sum(scan_b.values()), sum(pulse_b.values()), sum(tw_b.values())
    cost = {
        "scanb": n_scan * (tgt_scan - (max(scan_b) if scan_b else 0)) * SEC["band"] / 3600,
        "mtcurve": n_pulse * (tgt_pulse - (max(pulse_b) if pulse_b else 0)) * SEC["band"] / 3600,
        # tw6 has no seed-offset top-up, so the whole cell is redrawn rather than extended
        "tw6": n_tw * tgt_tw * SEC["band"] / 3600,
    }
    print(f"""
LEVELLING RUNS -- on O2, from the directory named in each block.

  band_model/     the two bandsi families. --resume tops up by SEED OFFSET rather than
                  redoing the cell: batches of 50 already on disk are skipped and only the
                  new ones are emitted, so raising 400 to {tgt_scan:,} costs the difference
                  and not the whole thing. Do NOT raise --batch to make the arrays smaller;
                  the batch size IS the seed offset and changing it silently re-draws seeds
                  that existing files already contain (see write_array's docstring).

    python bandsi_submit.py prod {tgt_scan} 50 0 short --only scanb --resume    # ~{cost['scanb']:.0f} CPU-h
    python bandsi_submit.py prod {tgt_pulse} 50 0 short --only mtcurve --resume  # ~{cost['mtcurve']:.0f} CPU-h

  cluster/        the two-wave band cells. tw6_run writes one JSON per cell holding every
                  replicate, with seeds 0..reps-1, so there is no way to append: the cell is
                  rerun at the higher count and the 50-replicate file is replaced. That
                  wastes the {n_tw * 50 * SEC['band'] / 3600:.0f} CPU-h already spent, which is
                  the price of not building seed-offset machinery for one sweep.

    python tw6_submit.py --reps {tgt_tw}                                  # ~{cost['tw6']:.0f} CPU-h

  cluster/        the five deme sensitivity points still at 1,500. scan4 batches by seed
                  offset into out4/ exactly as bandsi does, so --resume tops these up too:
                  only s0 = 1500 .. 1950 are emitted for the five short points and the 55
                  already at {tgt_scan:,} are skipped. Nothing needs deleting first.

    python scan4_submit.py {tgt_scan} 50 500 short --resume       # ~{5 * 500 * SEC['deme'] / 3600:.0f} CPU-h

TOTAL ~{sum(cost.values()):.0f} CPU-hours, almost all of it band.

NOT QUEUED: the S3.2 expansion curve. See the module docstring -- the band arm is short on
COMBINATIONS, not on replicates, and levelling it is a decision rather than a rerun.""")


if __name__ == "__main__":
    main()
```
