## Supplementary Information section 3 code base for "Hypothesis: A modern human range expansion ∼300,000 years ago had a critical role in Neandertal origins": screen.py

```
#!/usr/bin/env python3
"""
screen.py -- ONE definition of "admissible", shared by Figure 2b and Figure S3.3. (14 Sep 2026)

DR: "one shared admissibility function: a cell is in the box or it isn't, decided once from the
best estimate available for that cell, and both figures read the answer. S3.3 then plots the
same coordinate it was judged on, and every fountain point is by construction a point you can
find in the S3.3 box."

WHY THIS MODULE EXISTS. The two figures answer the same question about the same cells and used
to compute it separately, which meant they could disagree and did:

  * Figure 2b filtered on the STAGE-2 autosomal mean while Figure S3.3 plotted the STAGE-1 one.
    Same criterion, same threshold, different estimate of the same quantity -- 1,911 of the
    10,399 points in Figure 2b sat outside the box as S3.3 drew it, almost all of them cells
    straddling 3% or 7% where the ten-replicate screen is noisy.
  * Figure 2b read out_kearly_deme into two of Route 3's columns and Figure S3.3 did not, so
    the same mechanism at the same theta had eleven admissible deme cells in one figure and one
    in the other.
  * The two-wave band arm is timed on the genetic clock, and each figure had grown its own
    copy of that exception.

Those are three instances of one mistake: admissibility was a property of the plotting script
rather than of the cell. Here it is a property of the cell.

THE RULE, in full:

  BEST ESTIMATE PER CELL. A cell may have up to three measurements -- the 10-replicate screen,
  the 100-replicate re-screen of borderline cells, and the stage-2 score. The last one that
  exists wins, for every quantity: autosomal ancestry, duration, establishment and retention.
  DR: "Isn't our process to always use the largest number?" It is.

  THE CONSEQUENCE IS A MIXED-PRECISION SCATTER, DELIBERATELY. Only cells that passed the screen
  were promoted to stage 2, so near the box boundary a cell that passed carries a sharper
  estimate than one that failed. That is what "use the best number you have" implies, and it
  beats discarding 28,856 better measurements to keep the scatter uniform -- but it is a fact
  about the figure that the caption should state.

  K_early IS EXCLUDED. DR: "K_early (changing density during the expansion) is out! We decided
  that and write that in the text." routes_build.py has always said so -- "K_early off, rho /
  inbreeding ban / cyclical K out of scope" -- and out_kearly_deme was a diagnostic that
  answered whether theta = 0 is unreachable in the deme model or was merely run with the knob
  switched off. It should never have reached the published figure. Nothing here reads it.

  THE TWO-WAVE EXCEPTION IS OWNED HERE, ONCE. For a two-wave scenario the ordinary duration --
  the generation the last archaic-LABELLED unit disappears -- restarts at the second wave on a
  population that is already genetically modern, so it keeps ticking long after the genetics
  have settled. Those cells are timed instead on the generation whole-landscape mtDNA and Y
  both reach 99% modern. In the deme framework the two clocks agree to within two generations
  (clock_check.py: 615 against 620 over 40 replicates); in the band framework they diverge
  completely, and it is the difference between the route failing and working.
"""
import datetime as _dt
import glob
import json
import os

HERE = os.path.dirname(os.path.abspath(__file__))
CL = os.path.join(HERE, "cluster")

# The mechanism families, in the order their directories are read. A set missing from this list
# is a set the figures cannot see, which is how four of them were invisible until 12 Sep.
SETS = ["routes", "dmi", "dmi2", "xdc", "xd2", "tw", "cult"]

# The acceptance region, in one place. Every figure that draws a box draws this box.
AUT_LO, AUT_HI = 3.0, 7.0
GEN_LO, GEN_HI = 100.0, 1000.0
EST_MIN = 0.5

# Deme two-wave cells scored before this moment switched cfg.lineage at the second wave but not
# cfg.female_bias, so they ran wave two under wave one's residence rule and returned Y at 0-2%.
# They are still on disk for provenance and are skipped everywhere.
TW_DEME_FIX = _dt.datetime(2026, 9, 13, 0, 0, 0).timestamp()

_CACHE = {}


class Cell(dict):
    """One parameter combination, at the best estimate available for it.

    A dict rather than a namedtuple so that a caller can ask for a field the scan did not
    happen to fill without the whole figure raising.
    """

    @property
    def in_box(self):
        return in_box(self)


def in_box(c, aut_lo=AUT_LO, aut_hi=AUT_HI, gen_lo=GEN_LO, gen_hi=GEN_HI, est_min=EST_MIN):
    """The acceptance test, and the only one. Both figures call this.

    The bounds are arguments so that a caller can widen them for a sensitivity check, but the
    defaults are the note's criteria and nothing in the figures passes anything else.
    """
    return (c.get("aut") is not None
            and aut_lo <= c["aut"] <= aut_hi
            and c.get("gen") is not None
            and gen_lo <= c["gen"] <= gen_hi
            and (c.get("est") is None or c["est"] > est_min))


def _both_fixed(rows):
    """Fraction of replicates ending modern at BOTH markers, or None if none were recorded."""
    fix = [r for r in rows if r.get("mt") is not None and r.get("Y") is not None]
    if not fix:
        return None
    return sum(1 for r in fix if r["mt"] >= 99.999 and r["Y"] >= 99.999) / len(fix)


def _read(path):
    try:
        d = json.load(open(path))
    except Exception:
        return None
    s = d.get("summary", d)
    a, g = s.get("aut"), s.get("end_gen")
    if a is None or g is None or a != a or g != g:
        return None
    n = s.get("n") or 0
    return Cell(aut=float(a), gen=float(g),
                est=((s.get("n_est") or 0) / n) if n else None,
                joint=s.get("joint"),
                both=_both_fixed(d.get("rows") or []),
                reps=s.get("reps") or n or None,
                mech=s.get("mech"), val=s.get("val"), fw=s.get("framework"))


def _twowave(fw):
    """The two-wave cells, timed on the genetic clock. See the module docstring."""
    out = []
    pats = [f"out_tw5_{fw}/*.json"] + ([f"out_tw6_band/*.json"] if fw == "band" else [])
    for pat in pats:
        for p in glob.glob(os.path.join(CL, pat)):
            try:
                r = json.load(open(p))
            except Exception:
                continue
            if r.get("died") or not r.get("reps") or not r.get("wave2_gen"):
                continue
            # arr_med is the genetic clock and dur_med the label clock; prefer the former and
            # fall back only where the file predates it. A cell that never converged keeps a
            # duration beyond the top of any axis rather than being dropped, so it is visibly
            # outside the box instead of silently absent.
            dur = r.get("arr_med")
            if dur is None:
                dur = r.get("dur_med")
            out.append(Cell(aut=float(r["aut"]), gen=(1e9 if dur is None else float(dur)),
                            # arr_frac, not est: tw6's `est` is a synonym for swept_frac, the
                            # LABEL clock again, and only 5 of 135 cells clear it.
                            est=r.get("arr_frac", r.get("est")),
                            joint=r.get("p10"), both=r.get("p10"),
                            reps=r.get("reps"), mech="twowave", val=None, fw=fw))
    return out


def scan(refresh=False):
    """{(framework, mech, val): [Cell, ...]} for every screened combination.

    Precedence per cell: stage-2 score > 100-replicate re-screen > 10-replicate screen.
    """
    if _CACHE and not refresh:
        return _CACHE
    _CACHE.clear()
    for fw in ("deme", "band"):
        for pref in SETS:
            files = {}
            for d in (f"out_{pref}1_{fw}", f"out_{pref}1hi_{fw}", f"out_{pref}2_{fw}"):
                for p in glob.glob(os.path.join(CL, d, "*.json")):
                    if (pref == "tw" and fw == "deme" and d.endswith("2_deme")
                            and os.path.getmtime(p) < TW_DEME_FIX):
                        continue
                    files[os.path.basename(p)] = p
            for p in files.values():
                c = _read(p)
                if c is None or c.get("mech") is None:
                    continue
                if c["mech"] == "twowave":     # handled below, on the genetic clock
                    continue
                _CACHE.setdefault((fw, c["mech"], round(float(c["val"]), 6)), []).append(c)
        for c in _twowave(fw):
            _CACHE.setdefault((fw, "twowave", None), []).append(c)
    return _CACHE


def demog_nexp(mech):
    """{framework: {value: [Cell]}} for the UNBIASED expansion, from the `demog` sweep.

    KEPT OUT OF scan() ON PURPOSE. `demog` crosses the mechanisms with two axes no other set
    has -- the residence rule and the five-rung mtDNA-selection ladder -- so pooling it into
    the ordinary scan would make three mechanisms look five to thirty times denser than their
    neighbours for a reason that has nothing to do with the mechanism. It is read here, by
    itself, for one purpose: the unbiased-expansion series in Figure 2b.

    THE SLICE. residence unbiased (beta = 0.5 in the band engine, female_bias = 0.5 in the
    deme one) and s_mt = 0.00235, the median of the Muller's-ratchet posterior. Fixing s_mt
    matters: an unbiased expansion has no residence rule carrying its mtDNA, so without
    selection on mtDNA the marker is an unselected martingale and the whole series is flat at
    the autosomal proportion by construction. The matrilocal cells it is compared against run
    at s_mt = 0, which is not an oversight but the same asymmetry the note argues for -- and
    which the caption has to state, because two clouds at different s_mt are not otherwise
    comparable.

    THE LADDERS ARE ALIGNED, as of 15 Sep 2026. This sweep used to double above s = 0.02
    (0.04, 0.08, 0.16, 0.32) where the main screen uses 0.05, 0.10, 0.20, 0.35, 0.50, so the
    two series shared no rung above 0.02 and any sentence comparing them at a given s had to
    hedge. The realigned cells are scored: sY and theta at eleven shared rungs each and
    xdrive at eight, including the ends (s = 0, 50% and 100% daughters, theta = 0) that the
    curves used to stop short of. The superseded doubling cells remain on disk; callers that
    draw the curve filter to the main ladder rather than deleting them.
    """
    out = {"deme": {}, "band": {}}
    for fw in ("deme", "band"):
        key = "female_bias" if fw == "deme" else "beta"
        for p in glob.glob(os.path.join(CL, f"out_demog2_{fw}", "*.json")):
            try:
                s = json.load(open(p))["summary"]
            except Exception:
                continue
            if s.get("mech") != mech:
                continue
            if abs(float(s.get(key) or 0) - 0.5) > 1e-9:
                continue
            if abs(float(s.get("s_mt") or 0) - 0.00235) > 1e-9:
                continue
            c = _read(p)
            if c is not None:
                out[fw].setdefault(round(float(s["val"]), 6), []).append(c)
    return out


def by_key(mech, val, refresh=False):
    """{framework: [Cell]} for one mechanism setting; val None pools every rung."""
    c = scan(refresh)
    out = {}
    for fw in ("deme", "band"):
        if val is None:
            out[fw] = [x for (f, m, _v), xs in c.items() if m == mech and f == fw for x in xs]
        else:
            out[fw] = list(c.get((fw, mech, round(float(val), 6)), []))
    return out


if __name__ == "__main__":
    c = scan()
    tot = sum(len(v) for v in c.values())
    box = sum(1 for v in c.values() for x in v if x.in_box)
    j = sum(1 for v in c.values() for x in v if x.in_box and x.get("joint") is not None)
    print(f"{tot:,} screened cells in {len(c)} (framework, mechanism, value) groups")
    print(f"{box:,} inside the acceptance box "
          f"({AUT_LO}-{AUT_HI}% autosomal, {GEN_LO:.0f}-{GEN_HI:.0f} generations, "
          f"establishment > {EST_MIN:.0%})")
    print(f"{j:,} of those also carry a scored retention probability, so they can be plotted "
          f"as a fountain point in Figure 2b")
```
