## Supplementary Information section 4 for "Hypothesis: A modern human range expansion ∼300,000 years ago had a critical role in Neandertal origins"

A catalog of mutations that may have been important in uniquely modern human biology

**Findings**

- I identify 141 loci spanning 1.82 Mb where 16 African haplotypes and 4 Neandertal haplotypes are likely to share a common ancestor more recently than their common ancestors with Denisovans. These are candidates for loci important in early modern human biology, which were then retained by positive selection in Neandertals.
- One of the candidate regions overlaps *FADS1/2*, critical for conversion of dietary fats and with repeated evidence of natural selection.
- I identify nine fixed derived single nucleotide substitutions at functional sites where nearly all modern humans and analyzed Neandertals carry the derived allele while analyzable Denisovans are ancestral. Particularly notable is Glu1489Asp in *ARHGAP32*, a gene controlling the structure and strength of neuronal connections.
- I tested for enrichment of the catalog in particular functional classes and identified an excess of genes involved in proteasome-mediated protein degradation. The excess is absent from a control catalog built by exchanging the Neandertal and Denisovan labels. It is also stable to the choice of catalog threshold.
- The loci I single out for discussion are chosen based on a mixture of statistical support and literature about the genes. Genes that were the subject of selective sweeps during early modern human evolution may lie among the loci I passed over, and all 141 are listed in Table S4.1.

**Motivation**

Assembling catalogs of genetic variants that are present in all living humans and absent in sequenced archaic humans has the potential to shed light on uniquely modern human evolution[1,2]. Such catalogs may include mutations that occurred in early modern human ancestors, conferred distinctive traits, and rose to high frequency due to natural selection.

The primary approach to building such catalogs has been to use all available Neandertal and Denisovan genomes to represent archaic humans. However, the present study raises the possibility that a catalog where the individuals defined as archaic are Denisovans, and the fixed derived alleles in modern humans are shared with Neandertals, could also be of interest. While Neandertals derive only around 3-7% of their autosomal DNA from modern humans[3,4,5], this is a significant heritage, as it is derived from a probable early modern human range expansion into Europe. If a population is the product of a range expansion, using ancestry proportion to estimate the significance of the originally expanding population to its formation can be deceptive. The centrality of early modern humans to Neandertal origins is independently supported by shared archaeological features between West Eurasians and Africans after ~400-250 ka that were hardly present or even absent in East Asians around the same time. This includes use of Levallois point stone tools[6,7], improved hafting technology, a dramatic expansion in the use of fire, use of ochre, and larger and more organized campsites[8]. Could a handful of biological adaptations derived from early modern human ancestors have been important for allowing Neandertals to retain these distinctive features?

To identify mutations shared recently by all modern humans and Neandertals to the exclusion of Denisovans, I leveraged an Ancestral Recombination Graph (ARG) reconstructed using ARGweaver-D[4]. I analyzed 8 Africans (16 haplotypes), 2 Neandertals (4 haplotypes), and 1 Denisovan (2 haplotypes). I restricted to locations where the ARG inferred the time to the most recent common ancestor (TMRCA) of all African haplotypes and all Neandertal haplotypes to be more recent than the main split with Denisovans. As a control catalog, I repeated this analysis flipping the identity of Neandertals and Denisovans. I optimized catalog size (larger is better) while retaining enrichment for signal (the control catalog is only a small fraction of the case one).

The analyses and writing that follow are the product of a dialogue between the author and Artificial Intelligence (AI) agents of the Anthropic Claude family (Sonnet, Opus, and Fable models), for the most part running inside the Cursor Integrated Development Environment. I used the agents to write the simulations and ABC code in Python, to run diagnostics, and to help compose the text (any suggestions from the AI agents were reviewed and edited by me). All quantitative results are produced by my execution of the cited scripts on the O2 computing cluster at Harvard Medical School, or on my personal computer. I checked the robustness of the output, and reviewed the scientific interpretations, modeling choices, and final claims.

**Building the catalog**

I used the publicly available round 2 ARGweaver-D[4] 4ARG, shared by Melissa Hubisz (all positions in GRCh37 / hg19)[9]. At every kilobase (kb), I interpolated a TMRCA: the time to the most recent common ancestor of a set of haplotypes, converted to thousands of years assuming 29 years per generation, the value used by ARGweaver-D and retained here so that dates stay on its published scale (the forward simulations of Supplementary Information sections 1 and 3 instead use 28.6 years per generation). The ARGweaver-D files provide 2.5% and 97.5% percentiles from Markov Chain Monte Carlo replicates, which allowed me to interpolate intermediate percentiles of $p_{cut}$. I did this by exploiting ARGweaver-D's discretization of time on a log-spaced grid. I mapped the mean and the two reported percentiles onto that grid, fit a normal distribution in grid-index coordinates, and read off the percentile. ARGweaver-D was run in overlapping windows, and 4% of grid points thus carry two different gene genealogy inferences for the same position. At these positions I keep the one whose 95% posterior interval is narrower in absolute time.

The modern panel comprises eight diploid sub-Saharan African genomes (16 haplotypes) from the Mandenka, Khomani San, Mbuti, Mende, Yoruba, Luhya, Bantu from Kenya and Gambian populations. Archaic genomes are the Altai (Denisova 5)[10] and Vindija 33.19 (four haplotypes) Neandertals[11], and the Denisovan pinky (Denisova 3; two haplotypes)[12]. The three TMRCAs used for the catalog are $T_{Afr}$ (Africans only), $T_{Afr+Nea}$ (Africans + Neandertals), and $T_{Afr+Den}$ (Africans + Denisovans). I record these quantities at every autosomal grid point; results are in Supplementary Table 2, which has 2,440,090 rows.

To build catalogs, I identified grid points where $T_{Afr+Nea}$ falls below a cutoff $x$ near the modern-Neandertal split while $T_{Afr+Den}$ lies at $y \geq x$. To obtain insight into what fraction are real, I also built a control catalog with the two archaic genomes interchanged, and treated the ratio of control to case size as an estimate of the false-discovery proportion. The two archaic labels are not truly exchangeable, among other reasons because Neandertals carry around 5% modern human ancestry and Denisovans carry none, and because there are more Neandertals than Denisovans used in the catalog-building and their demographic histories differ. Nevertheless, the ratio of counts becomes <<1 when $x \leq 675$ ka, suggesting real signal.

For percentile cut $p_{cut}$, let $u$ be the $p_{cut}$$^{th}$ percentile of $T_{Afr+Nea}$, and $v$ the (100-$p_{cut}$)$^{th}$ of $T_{Afr+Den}$. I define a grid point as a case if $u$ falls below a threshold $x$ and $v$ above a threshold $y$ (interchanging the two archaic labels gives the controls). After experimentation, my final catalog used $p_{cut} = 50$ (the median), with $x = 625$ ka and $y = 700$ ka (I set y at 700 ka, near the recent end of the 825-694 ka estimate of the most recent population separation of Denisovan from modern humans[13]). The mutation rate, generation interval, and data filters used in the ARG construction differ from those in the study that estimated the population split[13]. However, the catalog definition itself is not based on the absolute timings.

Figure S4.1A shows that case and control counts separate sharply below 675 ka, so that *P(real)* rises there, implying enrichment for true signal. The two directions are not equally likely even where no sweep signal is possible: above the published modern-archaic split, where a coalescence on either side carries no sweep evidence, the Neandertal direction is in excess by a factor $\omega_0$ = 1.0125 (95% CI 1.005-1.020, measured over 254,809 grid points between 825 and 1,100 ka). The excess is stable across neighboring time ranges and flat across B-statistic deciles in both B maps, so a single global value is used and *P(real)* is computed throughout as 1 - (control count)/(case count)/$\omega_0$. The adjustment is small: the catalog's *P(real)* is 0.774 with it and 0.777 without. To test if the choice of parameters I used for catalog building (x = 625 ka, y = 700 ka, $p_{cut} = 50$) could be optimized further, I varied each of the three parameters in turn. Figure S4.1B plots *P(real)* against catalog size for variations of each of the three parameters. Between 200 and 1,600 grid points the traces approximately coincide. Above 1,600, P(real) falls steeply whichever parameter is changed. The thresholds used for the catalog are thus close to providing the largest catalog available at a *P(real)* similar to the best one. I chose a parameter combination that included more grid points while retaining a low estimated false-positive rate. The false-positive rate remained low even in a cross-validation analysis based on training the model on half the chromosomes and measuring on the other half. A caveat is that because I chose the curve's percentile cut after the held-out result was seen, it is not a fully unbiased estimate of out-of-sample performance.

**Figure S4.1: Sensitivity of the catalog to each of the three parameters used to build it.** (A) Grid points in case and control catalogs against threshold x, with $y = 700$ ka and $p_{cut} = 50$. The right axis gives *P(real)*, one minus the control-to-case ratio divided by $\omega_0$. (B) *P(real)* against catalog size. Each series varies one parameter while holding the other two fixed (star). Higher is better. Green series is a fitted surface, trained on one half of the chromosomes, scored on the other. Shaded bands 95% confidence intervals on *P(real),* from binomial counts.

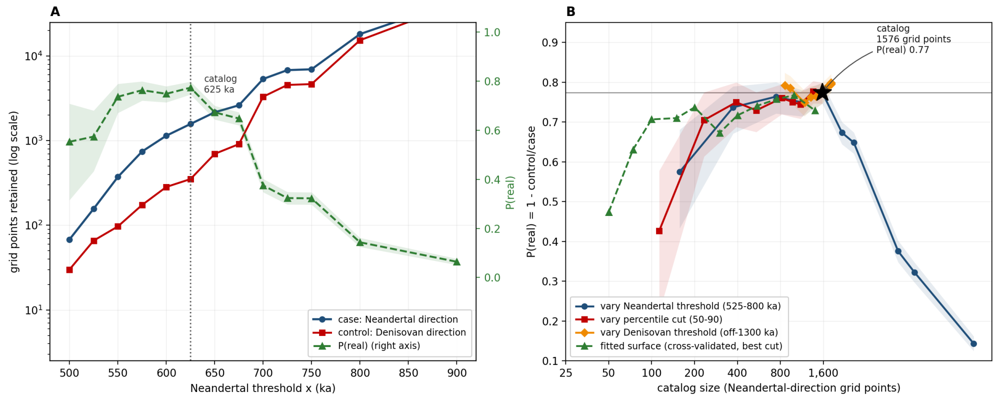

I attempted to extend the catalog to chromosome X. However, the control count there is at least 57% of that for the cases across the range of parameters I varied, so I was not able to distinguish it clearly from noise. Whether this reflects a genuinely different history on chromosome X is unclear. A more sensitive catalog-building algorithm might identify candidate modern human adaptations in this important part of the genome.

Coalescence times vary due to linked selection around deleterious mutations that has nothing to do with the sweeps of interest, a process known as background selection. I explored whether a "B statistic" measuring the strength of this process carries information that I could use to improve the catalog. I tested this in both the CADD- and phastCons-calibrated Murphy B-statistic maps[14], comparing the pre-specified single-threshold rule against one whose threshold shifts with B decile and against a free threshold per decile. At matched held-out catalog size, I could not improve the results with B-statistic conditioning.

I validated some of the catalog findings with an independent ARG-inference software, Relate, which reconstructs genealogies by a different algorithm that can evaluate many more samples, but is less accurate and does not produce estimates of uncertainty[15,16]. I used the Relate ARG to validate the modern-human TMRCA at the catalog loci, not to rebuild the catalog. Specifically, I took a Relate genealogy shared by Leo Speidel for 302 individuals of the 1000 Genomes phase 3 dataset (Yoruba, Han and British), inferred jointly with three Neandertals and a Denisovan. From it, I read the TMRCA of the 216 Yoruba haplotypes, and compared the mean estimate across each locus to the autosome-wide distribution of the same quantity averaged over blocks of the same span, expressed as a percentile.

### Functional annotation

To shed light on the biological impact of the catalog loci, I merged catalog grid points lying within 5 kb of one another, extended each by 5 kb, and intersected them with GENCODE v19 gene bodies, including all biotypes and introns[17]. For loci with no overlapping gene, I recorded the distance to the nearest gene. I used functional summaries from MyGene.info[18].

I also identified sites that alter a protein or fall in an element independently annotated as functional, that are fixed derived in Neandertals and modern humans (or nearly so), and absent in Denisovans. Because of the *a priori* importance of such positions, I built a new catalog to maximize the probability of including candidates, relaxing the Neandertal threshold from 625 to 675 ka, holding the percentile cut and the Denisovan threshold at their catalog values, and scanning for every fixed derived variant within ±5 kb. This led to 2,621 grid points in 287 loci, corresponding to 2.98 Mb rather than 1.82 Mb in the main catalog, both measured before the process of joining neighboring loci described below (padding). For each, I took every 1000 Genomes phase 3 biallelic single nucleotide variant[19], polarized it against the Ensembl six-primate ancestral sequence, and retained sites where the derived allele is at a frequency of >99% in 2,504 modern samples, fixed derived in all sampled Neandertals, and fixed ancestral in all sampled Denisovans.

After restricting to these sites, I separated out ones with strong predictions of functionality. From GENCODE v19, I identified predicted protein-altering changes—missense, stop gain, stop loss and start loss—along with the two intronic bases flanking each internal splice donor and acceptor, the untranslated regions of protein-coding transcripts, and the kilobase upstream of each protein-coding transcription start. I used all available protein-coding transcripts. I also used four further sources of annotation: the phastCons 46-way placental conserved elements[20], the per-base phyloP scores from the same alignment[21], the CADD v1.6 score[22], and the ENCODE DNase and transcription-factor binding clusters[23].

#### A catalog of 141 loci that are candidates for specifically modern human adaptation

The main catalog ($x$ = 625 ka, $y$ = 700 ka, $p_{cut}$ = 50) contains 1,576 of the 2,440,090 autosomal grid points. The estimated proportion of these grid points reflecting real signal is 77%, based on a ratio of 352 controls to 1,576 cases, adjusted by $\omega_0$ as above. Median TMRCAs at these points are 568 ka among Africans, 639 ka for Africans plus Neandertals, and 1,218 ka for Africans plus Denisovans. Merging grid points within 5 kb of one another yields 141 loci spanning 1.82 Mb, of which 24 are single grid points, and the largest spans 108 kb (Table S4.1). Relate places these loci among the most recently coalescing parts of the genome, with a median percentile of 10 using Yoruba haplotypes (however, 19 of 141 loci sit above the 50th percentile, and so do not replicate in the Relate ARG). A caution is that the catalog is incomplete in multiple ways due to missing parts of the genome that were not screened by ARGweaver-D, including parts of the genome likely to contain variants important in human evolution. For example, the catalog misses many regions of segmental duplication, which are sources of evolutionary innovation[24] and yet are challenging for ARG inference especially in a joint analysis with ancient genomes carrying short, difficult-to-map reads. The screen reaches 85% of autosomal bases, so 15% of the autosomes are absent altogether; even within autosomal genes, 16% of coding bases are missing.

Many of the 141 loci are clustered: merging adjacent ones within 100 kb causes them to collapse to 66 clusters (Table S4.1). For example, there is a chromosome 3 region spanning 11 loci and 398 kb and containing *SETD2*, *KLHL18*, *KIF9*, *PTPN23*, *SCAP* and *ELP6*, and a chromosome 11 region spanning 8 loci over 232 kb and containing *RHOD* and *KDM2A*.

I highlight ten clusters of loci that I judge to be notable candidates for distinctively modern human biology, weighing *P(real)*, the posterior probability that the modern-plus-Neandertal coalescence really is below the 700 ka threshold that defines the catalog, whether a fixed derived site there carries strong functional evidence, and whether the evidence in the literature points to the locus being of evolutionary interest (Figure S4.2). TMRCA values for individual loci below are the posterior medians reported in Table S4.1.

a. Eleven loci fall within 398 kb on chromosome 3p21, including *SETD2*, *KLHL18*, *KIF9*, *PTPN23*, *SCAP* and *ELP6*, with a maximum *P(real)* of 0.87. This is one of the most gene-rich clusters in the catalog, and it harbors two protein-altering changes in the gene *SETD2*, encoding the histone H3K36 trimethyltransferase required for transcriptional fidelity and DNA repair. *KLHL18*, also in this cluster, encodes a BTB-Kelch substrate adaptor for a Cullin-3 ubiquitin ligase and is one of the thirteen genes underlying the protein degradation enrichment identified by the gene ontology analysis (below). *KIF9* stands out due to its role in mitosis and due to the association of other mitotic spindle proteins (encoded by KIF18A and KNL1) with evidence of modern human distinctiveness in neocortex development[25].

b. *MED12L* on chromosome 3 is at *P(real)* = 0.83 and spans 25 kb, with $T_{Afr}$ = 508 ka, $T_{Afr+Nea}$ = 566 ka and $T_{Afr+Den}$ = 1607 ka. Heterozygous loss of function causes Nizon-Isidor syndrome, characterized by general developmental delay and impaired intellectual development including speech and language deficits[26].

c. *GLTSCR1L* on chromosome 6 spans 40 kb with *P(real)* = 0.83, $T_{Afr}$ = 525 ka, $T_{Afr+Nea}$ = 534 ka, and $T_{Afr+Den}$ = 1346 ka. Of the ten highlighted clusters, it is the one where $T_{Afr+Nea}$, the modern-plus-Neandertal coalescence, is most confidently <700 ka (probability near 1.00). It encodes a subunit of the non-canonical BAF chromatin remodeling complex, and includes a fixed derived site in a 5′ untranslated region inside a DNase cluster and a conserved element.

d. *SPATA31D1* on chromosome 9 spans 67 kb and is associated with a very confident signal: *P(real)* = 0.98. Its African-plus-Neandertal TMRCA is 384 ka against 1007 ka for Africans plus Denisovans, and it carries a protein-altering fixed derived change, albeit not one of high predicted functional significance based on conservation in multi-species alignments (phyloP -0.59). A caveat is that *SPATA31D1*, associated with spermatogenesis, lies within a segmental duplication, so mismapping and inaccurate estimation of ARG TMRCAs are concerns.

e. The FADS1/FADS2 locus on chromosome 11 spans 44 kb and has *P(real)* = 0.87, with $T_{Afr}$ = 436 ka, $T_{Afr+Nea}$ = 544 ka, and $T_{Afr+Den}$ = 1583 ka. It includes 15 fixed derived sites, and while none changes coding sequence, variants of known significance in this region are regulatory[27]. The genes encode the fatty acid desaturases that catalyze the rate-limiting steps in converting 18-carbon precursors into long-chain polyunsaturated fatty acids[28]. The derived haplogroup is associated with more efficient conversion and metabolism of plant-based fatty acids, and it is known to have swept to near-fixation in Africa[29]. It is not a new observation that Neandertals and most modern humans are more closely related here than either is to Denisovans[30]. However, the appearance of *FADS1/2* in this scan raises the possibility that adaptation to a more diverse and less exclusively carnivorous diet, facilitated by the more efficient haplotype better able to digest plant fats, may have been important in early modern human adaptation. The less efficient, more carnivorously adapted allele that later rose to high frequency in some non-Africans was not lost by this sweep, and may have persisted in the African archaic ancestors of modern humans. These results highlight *FADS1*/*FADS2* as a locus where the efficiency of fatty acid conversion has been a labile target of selection in humans[30], going back at least to the formation of the early modern human lineage.

f. *ARHGAP32* on chromosome 11 spans 20 kb and is associated with *P(real)* = 0.87. At this locus, $T_{Afr}$ = 490 ka, $T_{Afr+Nea}$ = 524 ka, and $T_{Afr+Den}$ = 1538 ka. The gene regulates dendritic spine morphology and glutamate receptor trafficking in neurons. The catalog peak contains within it a fixed derived site Glu1489Asp, the only site in the catalog that is both at a conserved base (phyloP 1.08) and inside a conserved element.

g. *MYCBP2* on chromosome 13 is within a cluster of five merged catalog loci totalling 35 kb. The modern-plus-Neandertal coalescence is near 600 ka across this region while the Denisovan value runs from 1425 to 1703 ka. *MYCBP2* encodes an atypical RING E3 ubiquitin ligase of the PHR family and is one of the thirteen genes behind the protein degradation enrichment identified by the gene ontology analysis (below). The PHR proteins direct axon guidance, axon termination and synapse formation. Mice lacking the orthologue show partial agenesis of the corpus callosum, and in humans loss of function causes developmental delay with corpus callosum defects, intellectual disability, epilepsy and autistic features.

h. A run of ten loci on chromosome 14 totals 96 kb and spans the genes *ACTR10*, *PSMA3*, *ARID4A*, *TOMM20L*, *TIMM9* and *KIAA0586*. Over the 80 grid points in this locus cluster that qualify for inclusion in the catalog, the joint probability that the Neandertal coalescence lies below 700 ka while the Denisovan one stays above reaches 0.99. The Table S4.2 site in this cluster is a protein-altering change in *TOMM20L* that also falls inside a DNase cluster, a TFBS cluster, and a conserved element. *PSMA3* encodes a core alpha subunit of the 20S proteasome and is one of the thirteen genes behind the protein degradation enrichment identified by the gene ontology analysis (below). *KIAA0586* encodes the centrosomal protein *TALPID3*, required for primary cilium formation and Hedgehog signaling, and its loss causes Joubert syndrome, with cerebellar and brainstem malformation and developmental delay.

i. *PTPN9* falls in a 68 kb locus on chromosome 15 and is at *P(real)* = 0.86. It encodes a phosphatase antagonizing hepatic insulin signaling, and is a candidate drug target for type 2 diabetes[31]. The same locus contains *SIN3A*, a chromatin co-repressor whose haploinsufficiency causes intellectual disability with microcephaly by disrupting cortical development[32].

j. A cluster of two loci on chromosome 17 totals 45 kb and reaches *P(real)* = 0.87. The stronger signal is at *SMURF2* with $T_{Afr+Nea}$ = 567 ka against $T_{Afr+Den}$ = 1353 ka (a weaker peak in this cluster contains *CEP95*). *SMURF2* encodes a HECT domain E3 ubiquitin ligase that terminates TGF-β and BMP signaling by ubiquitinating the SMADs and the type I receptor for degradation. Mice lacking it have reduced bone mass and raised bone resorption, and overexpression in cartilage accelerates chondrocyte maturation and endochondral ossification, so this is the one highlighted locus pointing at skeletal biology, where the contrast between modern humans and Neandertals is most visible in the skeletal record. It is the strongest representative of the protein degradation pathway that the gene ontology analysis highlights (below).

I discuss but do not visualize in Figure S4.2 four further signals that seem like particularly good candidates for follow-up analysis but not as compelling by themselves as the first ten. First, *HTR2B* at chr2:231.981-231.995 is associated with *P(real)* = 0.59, and encodes the serotonin 2B receptor, where a population-specific stop codon predisposes to severe impulsivity and knockout raises impulsive behavior in mice[33]; the same locus contains *PSMD1*, a 19S regulatory particle subunit and one of the thirteen genes behind the protein degradation enrichment identified by the gene ontology analysis (below). Second, a signal at chr17:74.063-74.070 (*P(real)* = 0.83) includes *GALR2*, which is a galanin receptor acting on feeding behavior and anxiety (the gene *ZACN* is also at the peak). Third, *WDFY3* on chromosome 4 regulates neural progenitor division and brain size. Fourth, a locus at chr3:83.244-83.351 spans 108 kb at *P(real)* = 0.86, with an African TMRCA of 285 ka against a Denisovan value of 1.2 million years, but the nearest annotated feature is the *CYP51P1* processed pseudogene 387 kb away.

**Table S4.1: Catalog of candidate loci for early modern human adaptation.** 141 runs of 1 kb autosomal grid points (points within 5 kb of one another merged into a single locus) where 16 sampled modern human and 4 Neandertal haplotypes are inferred to derive from a common ancestor ≤625 ka while 2 Denisovan haplotypes diverge ≥700 ka based on ARGweaver-D analysis. I pool loci within 100 kb to form 66 clusters (gray striping). Each TMRCA is the posterior median with its 90% central interval. "P(<700 ka)" is the posterior probability that the modern-plus-Neandertal coalescence is below the 700 ka threshold at the strongest gridpoint of the locus. "Relate percentile" is the inferred time to the most recent ancestor compared to the autosome-wide distribution in a larger panel of 216 West African Yoruba haplotypes. "P(real)" from the 1 kb grid point in each locus showing the strongest posterior signal of enrichment.

| chr | interval (Mb) in hg19/GRCh37 | span (kb) | *P* *(Real)* | $T_{Afr}$, ka (90% CI) (Argweaver) | $T_{Afr}$ percentile (Relate) | $T_{Afr+Nea}$, ka (90% CI) (Argweaver) | P($T_{Afr+Nea}$ <700 ka) (Argweaver) | $T_{Afr+Den}$, ka (90% CI) (Argweaver) | genes within 5 kb |
|---|---|---|---|---|---|---|---|---|---|
| 1 | 16.804-16.806 | 3 | 0.73 | 521 (271-925) | 9 | 596 (431-1125) | 0.80 | 1400 (813-2437) | *CROCCP3* (non-coding) |
| 1 | 16.815 | 1 | 0.73 | 550 (424-804) | 4 | 575 (428-987) | 0.80 | 1400 (813-2437) | *CROCCP3* (non-coding) |
| 1 | 35.762-35.775 | 14 | 0.77 | 565 (434-874) | 3 | 584 (464-909) | 0.88 | 1479 (983-2416) | *ZMYM4* (+2 non-coding) |
| 1 | 35.821-35.824 | 4 | 0.67 | 600 (558-684) | 3 | 600 (558-684) | 0.97 | 1500 (986-2674) | *ZMYM4* (+2 non-coding) |
| 1 | 35.831-35.835 | 5 | 0.67 | 603 (558-690) | 3 | 603 (558-690) | 0.97 | 1535 (960-3049) | *ZMYM4* (+1 non-coding) |
| 1 | 35.859 | 1 | 0.67 | 550 (424-804) | 4 | 600 (516-836) | 0.84 | 1400 (980-1879) | *ZMYM4* |
| 1 | 155.212-155.215 | 4 | 0.56 | 589 (430-1091) | 8 | 589 (430-1091) | 0.80 | 862 (629-1167) | *GBA*, *FAM189B* |
| 2 | 22.123-22.124 | 2 | 0.40 | 538 (276-1047) | 13 | 600 (432-1171) | 0.69 | 1981 (1541-3668) | *AC067959*.1 (non-coding) |
| 2 | 81.377-81.415 | 39 | 0.81 | 543 (388-841) | 90 | 555 (413-874) | 0.99 | 895 (644-1221) | none within 5 kb; *AC105342*.1 at 6 kb |
| 2 | 107.414 | 1 | 0.40 | 550 (424-804) | 10 | 600 (516-836) | 0.84 | 1500 (996-2518) | *ST6GAL2* |
| 2 | 162.102-162.110 | 9 | 0.74 | 526 (273-966) | 4 | 581 (429-1033) | 0.90 | 1172 (654-1837) | *AC009299*.2, *AC009299*.3 (non-coding) |
| 2 | 162.117-162.127 | 11 | 0.66 | 494 (262-741) | 29 | 598 (469-989) | 0.85 | 1236 (754-1826) | none within 5 kb; *AC009299*.2 at 6 kb |
| 2 | 172.735-172.738 | 4 | 0.43 | 527 (273-967) | 3 | 606 (475-1027) | 0.74 | 1488 (932-2717) | *SLC25A12* |
| 2 | 195.306 | 1 | 0.48 | 550 (280-1139) | 51 | 575 (372-1155) | 0.74 | 1600 (1180-2598) | none within 5 kb; *AC106883*.1 at 13 kb |
| 2 | 195.330 | 1 | 0.46 | 556 (282-1185) | 99 | 581 (374-1201) | 0.72 | 7563 (3850-12545) | none within 5 kb; *AC106883*.1 at 10 kb |
| 2 | 195.339-195.344 | 6 | 0.74 | 521 (271-925) | 97 | 560 (367-1040) | 0.90 | 1413 (884-2345) | none within 5 kb; *AC106883*.1 at 19 kb |
| 2 | 230.808 | 1 | 0.55 | 556 (425-850) | 11 | 613 (517-882) | 0.79 | 825 (720-930) | *FBXO36* (+1 non-coding) |
| 2 | 231.981-231.995 | 15 | 0.59 | 599 (474-996) | 9 | 599 (474-996) | 0.85 | 1429 (975-2106) | *PSMD1*, *HTR2B* (+1 non-coding) |
| 3 | 27.164-27.165 | 2 | 0.77 | 566 (427-918) | 68 | 566 (427-918) | 0.88 | 1700 (1196-3518) | *NEK10* |
| 3 | 27.180-27.200 | 21 | 0.79 | 571 (490-725) | 36 | 573 (492-729) | 0.97 | 2308 (1704-4662) | *NEK10* |
| 3 | 40.959-40.969 | 11 | 0.79 | 569 (427-947) | 72 | 580 (446-953) | 0.90 | 1631 (1114-3201) | none within 5 kb; *RP11-520A21*.1 at 43 kb |
| 3 | 47.157-47.174 | 18 | 0.79 | 557 (413-888) | 2 | 564 (428-899) | 0.91 | 1147 (664-1768) | *SETD2* |
| 3 | 47.180-47.212 | 33 | 0.83 | 550 (364-973) | 5 | 569 (370-1104) | 0.92 | 1233 (733-1906) | *SETD2* (+1 non-coding) |
| 3 | 47.218 | 1 | 0.61 | 581 (374-1201) | 1 | 613 (382-1385) | 0.63 | 1200 (656-1831) | *KIF9-AS1* (non-coding) |
| 3 | 47.224-47.229 | 6 | 0.66 | 600 (380-1339) | 1 | 600 (380-1339) | 0.65 | 1250 (660-2052) | *KIF9-AS1* (non-coding) |
| 3 | 47.236-47.266 | 31 | 0.81 | 531 (284-982) | 1 | 563 (368-1063) | 0.90 | 1212 (696-1884) | *KIF9* (+1 non-coding) |
| 3 | 47.272-47.295 | 24 | 0.81 | 520 (271-920) | 0 | 564 (369-1074) | 0.90 | 1236 (753-1781) | *KIF9* (+2 non-coding) |
| 3 | 47.305-47.315 | 11 | 0.71 | 557 (358-1008) | 4 | 592 (431-1110) | 0.80 | 1219 (702-1807) | *KIF9* |
| 3 | 47.326-47.353 | 28 | 0.87 | 508 (267-839) | 2 | 544 (362-935) | 0.97 | 1153 (691-1686) | *KIF9*, *KLHL18* |
| 3 | 47.441-47.478 | 38 | 0.79 | 465 (253-654) | 3 | 577 (428-998) | 0.90 | 1500 (1004-2584) | *PTPN23*, *SCAP* |
| 3 | 47.522-47.531 | 10 | 0.69 | 477 (257-700) | 0 | 602 (485-962) | 0.84 | 1496 (877-3074) | *SCAP* |
| 3 | 47.550-47.554 | 5 | 0.73 | 561 (426-886) | 0 | 589 (472-912) | 0.88 | 1770 (1185-3376) | *ELP6* |
| 3 | 83.244-83.351 | 108 | 0.86 | 285 (216-343) | 4 | 557 (437-794) | 0.97 | 1212 (810-1892) | none within 5 kb; *CYP51P1* at 387 kb |
| 3 | 150.865-150.889 | 25 | 0.83 | 508 (267-853) | 10 | 566 (427-932) | 0.97 | 1607 (1075-3084) | *MED12L* |
| 3 | 182.984-182.992 | 9 | 0.83 | 558 (412-902) | 96 | 558 (412-902) | 0.92 | 1828 (1213-3002) | *MCF2L2*, *B3GNT5* |
| 3 | 183.003-183.004 | 2 | 0.67 | 600 (516-836) | 99 | 600 (516-836) | 0.84 | 4000 (3161-11973) | *MCF2L2*, *B3GNT5* |
| 4 | 44.455-44.460 | 6 | 0.62 | 539 (361-887) | 1 | 539 (361-887) | 0.90 | 1608 (1126-2975) | *KCTD8* (+1 non-coding) |
| 4 | 54.153-54.157 | 5 | 0.64 | 580 (479-823) | 2 | 580 (479-823) | 0.88 | 785 (623-1024) | *SCFD2* |
| 4 | 85.821-85.854 | 34 | 0.69 | 498 (264-762) | 1 | 587 (483-859) | 0.88 | 1395 (910-2135) | *WDFY3* |
| 4 | 85.864 | 1 | 0.40 | 506 (266-817) | 2 | 600 (474-1004) | 0.74 | 1500 (996-2518) | *WDFY3* |
| 4 | 85.892-85.909 | 18 | 0.78 | 486 (277-730) | 4 | 554 (366-999) | 0.92 | 1137 (651-1716) | *WDFY3* (+1 non-coding) |

| | | | | | | | | | |
|---|---|---|---|---|---|---|---|---|---|
| 4 | 85.918-85.928 | 11 | 0.77 | 494 (262-750) | 6 | 541 (362-909) | 0.92 | 1509 (956-2921) | *WDFY3-AS2* (non-coding) |
| 4 | 139.977-139.998 | 22 | 0.67 | 370 (230-473) | 15 | 561 (435-849) | 0.96 | 1431 (894-2500) | *ELF2* (+1 non-coding) |
| 4 | 168.893 | 1 | 0.35 | 613 (517-882) | 21 | 613 (517-882) | 0.79 | 1500 (996-2518) | *RP11-310I9*.1 (non-coding) |
| 5 | 23.548-23.555 | 8 | 0.46 | 597 (474-984) | 51 | 597 (474-984) | 0.85 | 789 (623-1032) | none within 5 kb; *PRDM9* at 19 kb |
| 6 | 27.503-27.509 | 7 | 0.40 | 533 (295-966) | 41 | 602 (474-1010) | 0.74 | 1589 (1107-2859) | none within 5 kb; *HNRNPA1P1* at 11 kb |
| 6 | 27.530-27.544 | 15 | 0.64 | 569 (464-799) | 33 | 569 (464-799) | 0.96 | 1419 (952-2130) | XXbac-*BPG34I8*.3 (non-coding) |
| 6 | 27.560-27.604 | 45 | 0.76 | 542 (434-712) | 28 | 555 (455-738) | 0.97 | 1350 (831-2101) | *RNU6-471P*, *TRNAI6* (non-coding) |
| 6 | 27.639 | 1 | 0.40 | 600 (474-1004) | 42 | 600 (474-1004) | 0.74 | 1500 (955-2727) | none within 5 kb; *RP1-15D7*.1 at 18 kb |
| 6 | 27.754-27.755 | 2 | 0.69 | 597 (378-1293) | 16 | 597 (378-1293) | 0.72 | 1012 (641-1443) | none within 5 kb; *RSL24D1P1* at 5 kb |
| 6 | 42.771-42.810 | 40 | 0.83 | 525 (330-873) | 19 | 534 (359-879) | 1.00 | 1346 (774-2360) | *GLTSCR1L* |
| 6 | 83.303-83.332 | 30 | 0.64 | 531 (369-788) | 1 | 563 (454-805) | 0.97 | 1833 (1306-3334) | none within 5 kb; *TPBG* at 222 kb |
| 6 | 83.338-83.342 | 5 | 0.40 | 600 (474-1004) | 3 | 600 (474-1004) | 0.74 | 2300 (1636-5156) | none within 5 kb; *TPBG* at 257 kb |
| 6 | 83.348-83.355 | 8 | 0.48 | 596 (473-964) | 3 | 596 (473-964) | 0.88 | 1607 (1098-3083) | none within 5 kb; *UBE3D* at 247 kb |
| 7 | 97.737-97.740 | 4 | 0.82 | 553 (394-910) | 42 | 572 (398-1048) | 0.90 | 1450 (821-2897) | *LMTK2* |
| 7 | 109.185-109.197 | 13 | 0.79 | 559 (420-881) | 3 | 559 (420-881) | 0.90 | 1798 (1336-3413) | *AC073071*.1 (non-coding) |
| 7 | 142.876 | 1 | 0.54 | 613 (433-1217) | 51 | 613 (433-1217) | 0.66 | 1000 (640-1420) | *TAS2R39* |
| 7 | 142.892-142.895 | 4 | 0.64 | 559 (315-1145) | 47 | 575 (372-1155) | 0.74 | 947 (636-1322) | none within 5 kb; *TAS2R39* at 10 kb |
| 8 | 81.158 | 1 | 0.48 | 550 (280-1139) | 24 | 575 (372-1155) | 0.74 | 1575 (967-3417) | *MIR5708* (non-coding) |
| 8 | 85.006-85.007 | 2 | 0.65 | 550 (364-971) | 95 | 600 (474-1004) | 0.74 | 1062 (645-1535) | *RP11-120I21*.2 (non-coding) |
| 8 | 86.117 | 1 | 0.65 | 601 (464-1046) | 26 | 601 (464-1046) | 0.73 | 1100 (764-1436) | *E2F5* |
| 8 | 86.304-86.321 | 18 | 0.70 | 597 (515-808) | 27 | 597 (515-808) | 0.97 | 1050 (666-1588) | *CA3* |
| 9 | 84.505-84.524 | 20 | 0.76 | 580 (473-854) | 49 | 581 (476-854) | 0.97 | 916 (633-1268) | *RNA5SP287*, *SNORD95* & 1 other |
| 9 | 84.565-84.631 | 67 | 0.98 | 349 (250-495) | 10 | 384 (320-502) | 1.00 | 1007 (641-1486) | *SPATA31D1* (+3 non-coding) |
| 9 | 84.641-84.642 | 2 | 0.96 | 363 (310-405) | 2 | 363 (310-405) | 1.00 | 787 (623-1029) | none within 5 kb; *SPATA31B1* at 31 kb |
| 9 | 84.659-84.679 | 21 | 0.75 | 536 (360-887) | 83 | 536 (360-887) | 0.97 | 832 (627-1111) | *SPATA31B1* (non-coding) |
| 9 | 84.687-84.708 | 22 | 0.71 | 552 (369-980) | 11 | 552 (369-980) | 0.97 | 852 (628-1147) | *DDX10P2* (non-coding) |
| 9 | 84.726-84.727 | 2 | 0.71 | 503 (349-663) | 20 | 503 (349-663) | 0.97 | 725 (618-914) | none within 5 kb; *RP11-388B24*.4 at 19 kb |
| 9 | 84.737-84.747 | 11 | 0.62 | 577 (378-1150) | 11 | 577 (378-1150) | 0.84 | 897 (632-1229) | *RP11-388B24*.4 (non-coding) |
| 9 | 84.780-84.784 | 5 | 0.60 | 564 (386-1014) | 2 | 579 (424-1023) | 0.80 | 820 (626-1089) | none within 5 kb; *RP11-388B24*.4 at 33 kb |
| 9 | 84.792-84.799 | 8 | 0.68 | 529 (357-821) | 18 | 543 (396-828) | 0.97 | 770 (622-997) | none within 5 kb; *RP11-388B24*.4 at 45 kb |
| 9 | 84.813-84.814 | 2 | 0.57 | 538 (360-879) | 22 | 563 (426-895) | 0.90 | 762 (621-983) | none within 5 kb; *RP11-388B24*.4 at 66 kb |
| 9 | 84.820 | 1 | 0.53 | 575 (372-1155) | 15 | 600 (422-1213) | 0.68 | 900 (632-1236) | none within 5 kb; *RP11-15B24*.5 at 68 kb |
| 9 | 84.899-84.918 | 20 | 0.73 | 575 (470-820) | 60 | 586 (472-892) | 0.88 | 1697 (1196-3229) | *RP11-15B24*.5 (non-coding) |
| 9 | 135.801 | 1 | 0.40 | 601 (506-878) | 29 | 601 (506-878) | 0.81 | 725 (618-914) | *TSC1* |
| 10 | 4.737-4.741 | 5 | 0.59 | 555 (414-882) | 49 | 558 (415-901) | 0.91 | 1535 (1052-2588) | none within 5 kb; *LINC00704* at 17 kb |
| 10 | 109.119-109.132 | 14 | 0.79 | 537 (276-1048) | 97 | 564 (369-1072) | 0.92 | 1021 (662-1426) | none within 5 kb; *RNA5SP326* at 89 kb |
| 10 | 109.251-109.254 | 4 | 0.57 | 563 (436-812) | 25 | 600 (516-836) | 0.84 | 800 (716-884) | none within 5 kb; *RNA5SP326* at 30 kb |
| 11 | 1.020-1.023 | 4 | 0.77 | 569 (435-910) | 29 | 569 (435-910) | 0.90 | 1325 (842-1861) | *MUC6* |
| 11 | 14.555 | 1 | 0.58 | 556 (282-1185) | 53 | 600 (380-1339) | 0.65 | 1000 (640-1420) | *PSMA1* |
| 11 | 14.566-14.573 | 8 | 0.69 | 577 (359-1167) | 1 | 602 (425-1182) | 0.76 | 1378 (836-2328) | *PSMA1* |
| 11 | 14.579-14.587 | 9 | 0.73 | 585 (470-891) | 1 | 585 (470-891) | 0.88 | 1378 (898-2018) | *PSMA1* |
| 11 | 57.372-57.381 | 10 | 0.67 | 588 (440-995) | 37 | 602 (474-1011) | 0.74 | 1533 (982-3145) | *SERPING1* |
| 11 | 61.055-61.071 | 17 | 0.58 | 581 (468-869) | 75 | 581 (468-869) | 0.97 | 792 (624-1038) | *VWCE*, *DDB1* |
| 11 | 61.541-61.548 | 8 | 0.71 | 481 (258-718) | 11 | 588 (438-1048) | 0.90 | 1156 (653-1774) | *MYRF*, *TMEM258* |
| 11 | 61.554-61.597 | 44 | 0.87 | 436 (253-620) | 1 | 544 (399-835) | 0.97 | 1583 (1064-2948) | *MYRF*, *TMEM258*, *FEN1*, *FADS2*, *FADS1* |
| 11 | 66.822-66.857 | 36 | 0.68 | 554 (380-961) | 9 | 555 (380-967) | 0.92 | 819 (626-1087) | *SYT12*, *RHOD* |
| 11 | 66.870-66.907 | 38 | 0.73 | 567 (427-934) | 3 | 567 (427-934) | 0.97 | 891 (631-1220) | *KDM2A* |
| 11 | 66.915-66.920 | 6 | 0.54 | 595 (431-1125) | 5 | 595 (431-1125) | 0.80 | 875 (630-1190) | *KDM2A* |
| 11 | 66.928-66.968 | 41 | 0.81 | 544 (423-775) | 4 | 544 (423-775) | 0.97 | 1020 (642-1464) | *KDM2A*, *AP001885*.1 |
| 11 | 66.974-66.987 | 14 | 0.67 | 538 (422-726) | 4 | 538 (422-726) | 0.97 | 758 (621-975) | *KDM2A* (+1 non-coding) |
| 11 | 66.996-67.009 | 14 | 0.69 | 571 (449-877) | 1 | 571 (449-877) | 0.90 | 890 (631-1217) | *KDM2A* |
| 11 | 67.022 | 1 | 0.49 | 613 (517-882) | 35 | 613 (517-882) | 0.79 | 900 (632-1236) | *KDM2A* |

| | | | | | | | | | |
|---|---|---|---|---|---|---|---|---|---|
| 11 | 67.054 | 1 | 0.62 | 613 (517-882) | 26 | 613 (517-882) | 0.79 | 1300 (796-1855) | *ADRBK1*, *ANKRD13D* |
| 11 | 128.831-128.850 | 20 | 0.87 | 490 (261-743) | 6 | 524 (356-805) | 0.97 | 1538 (1023-2764) | *ARHGAP32* |
| 12 | 3.767 | 1 | 0.64 | 600 (432-1171) | 58 | 600 (432-1171) | 0.69 | 1175 (734-1616) | *EFCAB4B* |
| 12 | 53.852-53.857 | 6 | 0.55 | 565 (426-911) | 11 | 591 (469-949) | 0.88 | 1442 (847-2681) | *PCBP2* (+1 non-coding) |
| 12 | 74.343-74.370 | 28 | 0.59 | 499 (316-695) | 2 | 565 (476-734) | 0.97 | 1863 (1363-3337) | none within 5 kb; *RP11-711C17*.1 at 30 kb |
| 12 | 85.564-85.581 | 18 | 0.46 | 594 (480-927) | 11 | 597 (486-929) | 0.85 | 1826 (1351-3585) | *LRRIQ1* |
| 12 | 85.612-85.618 | 7 | 0.59 | 577 (470-833) | 45 | 601 (513-848) | 0.85 | 1525 (1024-2658) | *LRRIQ1* |
| 12 | 85.630-85.668 | 39 | 0.65 | 582 (490-805) | 7 | 584 (492-806) | 0.97 | 1574 (1039-2702) | *LRRIQ1* (+1 non-coding) |
| 13 | 77.631-77.640 | 10 | 0.67 | 580 (477-833) | 4 | 603 (516-847) | 0.84 | 1609 (1198-2614) | *MYCBP2* |
| 13 | 77.729-77.741 | 13 | 0.73 | 598 (537-744) | 7 | 598 (537-744) | 0.97 | 1703 (1240-3126) | *MYCBP2* |
| 13 | 77.748 | 1 | 0.67 | 569 (501-674) | 20 | 600 (558-684) | 0.97 | 1425 (817-2667) | *MYCBP2* |
| 13 | 77.761-77.765 | 5 | 0.72 | 484 (259-687) | 30 | 595 (515-801) | 0.95 | 1544 (1161-2249) | *MYCBP2* |
| 13 | 77.783-77.788 | 6 | 0.67 | 600 (516-836) | 43 | 604 (516-851) | 0.84 | 1696 (1357-2675) | *MYCBP2* |
| 14 | 21.360-21.362 | 3 | 0.65 | 600 (516-836) | 46 | 600 (516-836) | 0.84 | 1054 (645-1534) | *RNASE3* (+1 non-coding) |
| 14 | 21.379-21.386 | 8 | 0.75 | 566 (419-924) | 43 | 566 (419-924) | 0.97 | 1048 (644-1516) | *RP11-219E7*.4, *RP11-84C10*.3 & 1 other |
| 14 | 21.392-21.407 | 16 | 0.87 | 415 (246-534) | 5 | 498 (348-687) | 1.00 | 877 (630-1193) | *RP11-219E7*.4, *RP11-84C10*.3 & 1 other |
| 14 | 58.701-58.702 | 2 | 0.62 | 516 (269-886) | 9 | 606 (475-1027) | 0.74 | 1162 (653-1752) | C14orf37, *ACTR10* |
| 14 | 58.711-58.726 | 16 | 0.59 | 362 (234-500) | 1 | 591 (443-1043) | 0.88 | 1467 (865-2939) | C14orf37, *PSMA3* |
| 14 | 58.743-58.746 | 4 | 0.65 | 200 (200-200) | 1 | 578 (429-1010) | 0.80 | 1300 (796-1855) | C14orf37, *PSMA3* (+1 non-coding) |
| 14 | 58.757-58.758 | 2 | 0.59 | 231 (205-257) | 2 | 600 (432-1171) | 0.69 | 1300 (796-1855) | C14orf37, *AL132989*.1 (+3 non-coding) |
| 14 | 58.773-58.802 | 30 | 0.66 | 375 (235-488) | 9 | 560 (414-904) | 0.92 | 1596 (1064-3114) | *ARID4A* |
| 14 | 58.813 | 1 | 0.46 | 506 (266-817) | 0 | 581 (419-1075) | 0.76 | 1700 (1196-3518) | *ARID4A* |
| 14 | 58.826-58.834 | 9 | 0.64 | 541 (416-779) | 2 | 545 (423-781) | 0.97 | 2181 (1593-3288) | *ARID4A* (+1 non-coding) |
| 14 | 58.845-58.849 | 5 | 0.64 | 469 (254-627) | 1 | 553 (425-827) | 0.90 | 1550 (1005-2977) | *ARID4A* (+1 non-coding) |
| 14 | 58.857-58.862 | 6 | 0.83 | 451 (249-578) | 0 | 487 (344-613) | 1.00 | 1392 (803-2563) | *TOMM20L* (+2 non-coding) |
| 14 | 58.872-58.892 | 21 | 0.71 | 462 (252-614) | 10 | 513 (352-727) | 0.97 | 1666 (1160-3287) | *TOMM20L*, *TIMM9*, *KIAA0586* |
| 14 | 102.184-102.185 | 2 | 0.65 | 588 (420-1121) | 12 | 588 (420-1121) | 0.78 | 1300 (796-1855) | *RP11-1029J19*.5 (non-coding) |
| 15 | 72.713-72.720 | 8 | 0.78 | 536 (276-1036) | 30 | 581 (374-1201) | 0.84 | 1256 (756-1861) | *RP11-1007O24*.3 (non-coding) |
| 15 | 72.763-72.769 | 7 | 0.73 | 580 (471-852) | 9 | 580 (471-852) | 0.88 | 1236 (699-1952) | *ARIH1* (+2 non-coding) |
| 15 | 72.797-72.804 | 8 | 0.79 | 508 (351-694) | 6 | 586 (514-747) | 0.97 | 1388 (889-2135) | *ARIH1* |
| 15 | 72.859-72.867 | 9 | 0.72 | 588 (491-838) | 1 | 588 (491-838) | 0.88 | 1128 (769-1494) | *ARIH1* |
| 15 | 74.952-74.962 | 11 | 0.55 | 581 (429-1025) | 22 | 588 (441-1029) | 0.80 | 830 (626-1106) | *EDC3* |
| 15 | 74.972-74.975 | 4 | 0.54 | 586 (482-847) | 27 | 586 (482-847) | 0.88 | 806 (625-1063) | *EDC3* |
| 15 | 74.983 | 1 | 0.44 | 613 (475-1050) | 30 | 613 (475-1050) | 0.71 | 825 (626-1098) | *EDC3* |
| 15 | 75.738-75.805 | 68 | 0.86 | 515 (272-875) | 3 | 550 (368-963) | 0.97 | 1041 (643-1517) | *SIN3A*, *PTPN9* (+2 non-coding) |
| 15 | 78.310-78.311 | 2 | 0.67 | 600 (558-684) | 13 | 600 (558-684) | 0.97 | 1313 (882-1773) | *TBC1D2B* |
| 17 | 62.513-62.532 | 20 | 0.80 | 562 (368-1035) | 0 | 574 (405-1043) | 0.91 | 1516 (1011-2695) | *CEP95* (+1 non-coding) |
| 17 | 62.610-62.634 | 25 | 0.87 | 567 (386-1041) | 50 | 567 (386-1041) | 0.97 | 1353 (803-2192) | *SMURF2* |
| 17 | 67.847 | 1 | 0.67 | 575 (428-987) | 8 | 600 (474-1004) | 0.74 | 1400 (813-2437) | *AC003051*.1 (non-coding) |
| 17 | 74.063-74.070 | 8 | 0.83 | 545 (385-864) | 4 | 556 (390-933) | 0.92 | 1563 (1001-3119) | *SRP68*, *ZACN*, *GALR2* |
| 18 | 42.233-42.253 | 21 | 0.62 | 603 (558-690) | 96 | 603 (558-690) | 0.97 | 1697 (1189-2999) | *RP11-456K23*.1 (non-coding) |
| 18 | 76.991 | 1 | 0.40 | 575 (428-987) | 21 | 600 (474-1004) | 0.74 | 1700 (1196-3518) | *ATP9B* |
| 18 | 76.998-77.004 | 7 | 0.52 | 602 (474-1010) | 83 | 602 (474-1010) | 0.74 | 1514 (1023-2610) | *ATP9B* |
| 18 | 77.034-77.046 | 13 | 0.65 | 541 (383-827) | 15 | 555 (425-835) | 0.90 | 1583 (1068-3009) | *ATP9B* |
| 18 | 77.053-77.081 | 29 | 0.66 | 579 (468-855) | 31 | 579 (468-855) | 0.89 | 1538 (1055-2613) | *ATP9B* |
| 20 | 32.635-32.653 | 19 | 0.76 | 532 (358-845) | 1 | 545 (373-895) | 0.92 | 1317 (795-2147) | *RALY* (+1 non-coding) |
| 20 | 46.186-46.198 | 13 | 0.65 | 600 (474-1004) | 18 | 602 (474-1009) | 0.74 | 1031 (643-1488) | *NCOA3* (+1 non-coding) |
| 22 | 29.723-29.726 | 4 | 0.63 | 575 (384-1124) | 1 | 600 (432-1171) | 0.69 | 906 (633-1247) | *AP1B1* (+1 non-coding) |

**Figure S4.2: Ten notable genomic regions where modern humans and Neandertals share an extraordinarily recent ancestor making them candidates for harboring evolutionary adaptations important to early modern humans.** The estimated probability of modern humans and Neandertals sharing a common ancestor more recently than the population split from Denisovans is shown on the left axis as *P(real)*. The right axis shows the inferred mean time to the most recent common ancestor of both Africans and Neandertals ($T_{Afr+Nea}$), as well as Africans and Denisovans ($T_{Afr+Den}$); parentheses give percentile in the genome-wide distribution of $T_{Afr+Nea}$, dotted lines genome-wide averages. Below the graphs, I show the grid points that qualify for inclusion in the catalog, fixed derived sites (starred if predicted to be highly functional), and GENCODE v19 genes with coding sequence drawn thickly.

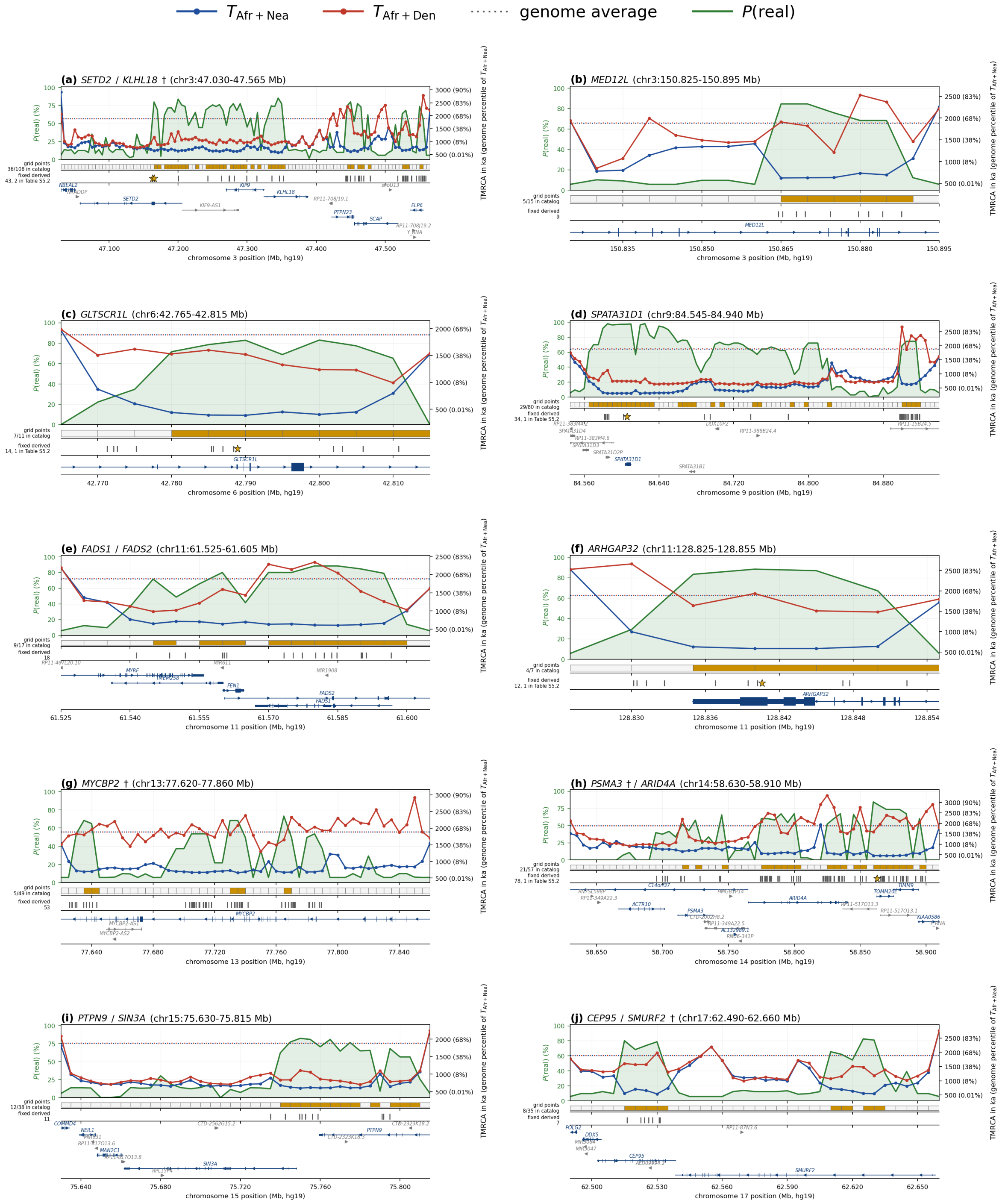

### Substitutions of high predicted functional significance

Using a looser catalog defined by the parameters ($x$ = 675 ka, $y$ = 700 ka, $p_{cut}$ = 50), I searched for single nucleotide substitutions that were candidates for having been driven to high frequency on the early modern human lineage. After defining the ancestral allele based on the Ensembl six-primate ancestral sequence, I restricted to sites in the catalog where the derived allele frequency in the 1000 Genomes Project dataset was >99%, all available Neandertal haplotypes from 5 high-coverage genomes[10,11,34,35,36] were derived (minimum coverage of 2 haplotypes and maximum coverage of 10), and where all available Denisovan haplotypes from 2 high-coverage genomes[12,13] were ancestral (minimum coverage of 2 haplotypes and maximum coverage of 4).

**Table S4.2: Candidate variants of high predicted functional significance.**

| chr | position hg19/ GRCh37 | gene | functional evidence (6 protein-coding changes highlighted) | CADD | *P (real)* | moderns | Nea | Den |
|---|---|---|---|---|---|---|---|---|
| 2 | 162,101,812 | *TANK* (9 kb) | conserved, DNase cluster, CADD top 1% | 20.2 | 0.71 | 5008/5008 | 10/10 | 0/4 |
| 3 | 47,162,659 | *SETD2* | His1156Arg | 19.2 | 0.76 | 5008/5008 | 10/10 | 0/4 |
| 3 | 47,165,351 | *SETD2* | Val259Ile | 16.7 | 0.79 | 5008/5008 | 10/10 | 0/4 |
| 3 | 182,994,744 | *MCF2L2* | Asn593Ser, splice region, conserved element | 16.2 | 0.00 | 4872/5008 | 10/10 | 0/4 |
| 6 | 42,788,926 | *GLTSCR1L* | 5′ untranslated, conserved, DNase cluster | 14.7 | 0.75 | 4970/5008 | 10/10 | 0/2 |
| 9 | 84,605,717 | *SPATA31D1* | Leu111Arg | 10.9 | 0.97 | 5000/5008 | 10/10 | 0/2 |
| 11 | 67,053,028 | *ADRBK1* | 3′ UTR, conserved, TFBS cluster | 15.8 | 0.52 | 5008/5008 | 10/10 | 0/4 |
| 11 | 128,840,599 | *ARHGAP32* | Glu1489Asp, 3′ UTR, conserved, DNase cluster | 18.3 | 0.87 | 5008/5008 | 10/10 | 0/4 |
| 14 | 58,862,763 | *TOMM20L* | Asp30Asn, conserved, DNase & TFBS clusters | 16.7 | 0.00 | 5008/5008 | 8/8 | 0/2 |

Notes: The amino-acid-changing substitutions in Table 1 of the main text are a subset of the 9 variants here. CADD measures predicted deleteriousness by combining functional annotations into one score on the PHRED scale of $-10 \times \log_{10}$(percentile), for this table running from 10.9 (top 10%) through 20.2 (top 1%). Haplotype columns give derived over total count for 5,008 modern human haplotypes in the 1000 Genomes Project, up to 10 haplotypes from high-coverage Neandertals, and up to 4 haplotypes from high-coverage Denisovans. Gene column gives the (nearest) gene body. P(real) is evaluated at the substitution itself rather than across the locus around it, and is 0.00 at MCF2L2 and TOMM20L because the modern-plus-Neandertal coalescence inferred at those exact positions is older than the 625 ka catalog ceiling even though the surrounding locus qualifies; both loci are at P(real) = 0.83 in Table S4.1.

This procedure identified 991 fixed derived sites at >99% derived frequency in the 1000 Genomes project dataset, together with 232 more recovered in a >95% sensitivity run (Supplementary Table 3). The nine predicted to be of strongest functional significance are listed in Table S4.2: the eight that meet the criteria below among the 991, together with one protein-altering site at *MCF2L2* that appears only in the sensitivity run. These change the coding sequence of a gene, occur at an essential splice position, [22]score in the top 1% of possible substitutions by CADD[22], or fall in a conserved element coinciding with an untranslated region or promoter. I excluded synonymous changes even when they scored highly by CADD. A synonymous change acting through splicing or codon usage should have left conservation at the base; where it has not, the score instead reflects the exon around the site, which the conserved element scan would already catch.

(a) *ARHGAP32* Glu1489Asp was listed in the catalog of candidate variants for modern human adaptation in the paper that first published a high-coverage archaic genome, a Denisovan[12]. It was screened out in later catalogs because Neandertals carry the derived allele, but the present study shows it to be of even greater interest than the original study suggested. In that study, it was one of 260 substitutions at which modern humans are fixed for the derived state while the Denisovan carries the ancestral one, and one of only 23 such positions with a primate conservation score of at least 0.95 on that study's own primate-alignment scale. The protein is a GTPase-activating protein involved in NMDA receptor signaling, controlling dendritic spine morphology and axon growth. It is constrained in humans today, with a gnomAD loss-of-function intolerance of 1[37].

(b) *SETD2* Val259Ile falls in a gene encoding a single-copy histone H3K36 trimethyltransferase, essential for transcriptional fidelity and DNA repair. Its haploinsufficiency causes an overgrowth and neurodevelopmental syndrome[38]. Residue 259 lies in the N-terminal low-complexity region, outside the catalytic SET domain and outside the AWS and post-SET domains[39]. The substitution is not predicted to be functionally particularly important, as it is between chemically similar residues, but it still merits follow-up analysis.

(c) *SPATA31D1* Leu111Arg falls in a gene family that is a well-documented target of positive selection in humans and African apes[40]. The family expanded from a single copy in mouse, to two in macaque, to many copies in hominoids. In the primate lineage, it acquired new protein domains implicated in ultraviolet damage response and DNA repair[41]. A caution is that copy number variation raises concerns about mismapping (a particular problem for ancient DNA reads) and errors in ARG reconstruction, raising the possibility that this signal is a false positive.

**The catalog is significantly enriched in genes involved in proteasome biology**

To test whether the catalog is enriched for particular categories of gene function, I used the regulatory domain model of GREAT[42]. This assigns each locus not only to the genes it overlaps, but to genes whose potential regulatory domain it falls in, with a domain defined as running from 5 kb upstream to 1 kb downstream of the transcription start site, then outward to the nearest neighboring domain, capped at 1 Mb. Over GENCODE v19 protein-coding genes, such domains cover 91% of the autosomes, so most non-coding intervals can be associated with gene function. The 141 loci fall into regulatory domains of 110 genes carrying a Gene Ontology (GO) biological process annotation[43], less than one percent of the 14,515 GO-annotated genes. I tested 536 GO terms that contain between 10 and 1,000 annotated genes and at least two catalog genes, using a hypergeometric test over genes and a binomial test over loci.

Table S4.3 shows all the categories that are significant after correcting for multiple hypothesis testing. Nine categories pass a Bonferroni correction for the 200 categories tested by permutation, and all largely reflect the same signal. Seven are nested descriptions of protein catabolism, with a particularly notable case being proteasome-mediated ubiquitin-dependent protein catabolism: 13 genes (*CTNNB1*, *FBXL3*, *KDM2A*, *KLHL18*, *MYCBP2*, *PCBP2*, *PSMA1*, *PSMA3*, *PSMD1*, *PSMD14*, *SMURF2*, *TRIP12* and *UBE3D*), compared with an expected 3.0 (nominal $p = 9.5 \times 10^{-6}$, false discovery rate of $q = 0.0016$). Four of these genes (*PSMA1*, *PSMA3*, *PSMD1* and *PSMD14*) also fall in the two type I interferon response categories, so those categories are correlated observations (those categories carry 7 and 6 catalog genes; Table S4.3).

**Table S4.3: Gene Ontology biological categories most strongly enriched in the catalog**

| GO biological process | genes | catalog | exp. | p-value (nominal) | q-value (FDR) | p-value ($10^5$ perm.) |
|---|---|---|---|---|---|---|
| macromolecule catabolic process | 960 | 22 | 7.3 | $2.5 \times 10^{-6}$ | 0.0010 | $< 1 \times 10^{-5}$ |
| response to type I interferon | 85 | 7 | 0.6 | $3.6 \times 10^{-6}$ | 0.0010 | $< 1 \times 10^{-5}$ |
| proteasome-mediated ubiquitin-dependent protein catabolic | 397 | 13 | 3.0 | $9.5 \times 10^{-6}$ | 0.0016 | $< 1 \times 10^{-5}$ |
| ubiquitin-dependent protein catabolic process | 541 | 15 | 4.1 | $1.4 \times 10^{-5}$ | 0.0016 | $< 1 \times 10^{-5}$ |
| modification-dependent protein catabolic process | 551 | 15 | 4.2 | $1.7 \times 10^{-5}$ | 0.0016 | $< 1 \times 10^{-5}$ |
| modification-dependent macromolecule catabolic process | 553 | 15 | 4.2 | $1.8 \times 10^{-5}$ | 0.0016 | $< 1 \times 10^{-5}$ |
| cellular response to type I interferon | 78 | 6 | 0.6 | $2.7 \times 10^{-5}$ | 0.0021 | $4 \times 10^{-5}$ |
| proteasomal protein catabolic process | 477 | 13 | 3.6 | $6.4 \times 10^{-5}$ | 0.0043 | $2 \times 10^{-5}$ |
| protein catabolic process | 722 | 16 | 5.5 | $1.0 \times 10^{-4}$ | 0.0062 | $5 \times 10^{-5}$ |

Notes: "genes" is the genes in the category of the total of 14,515; "catalog" those whose regulatory domain contains one of the 141 loci; "exp." is the hypergeometric expectation. "p-value (nominal)" reports a hypergeometric test over genes and "q-value (FDR)" the Benjamini-Hochberg false discovery rate across all 536 categories that contain between 10 and 1,000 annotated genes and at least two catalog genes. "p-value ($10^5$ permutations)" is from a test that relocates all 66 clusters to blocks of equal length and similar mean B, and reports the fraction of replicates that reached the observed enrichment. I ran the permutation only for the 200 most significant categories rather than all 536; the Bonferroni threshold over those 200 is $2.5 \times 10^{-4}$; all nine clear this threshold. The nine are not independent tests: seven are nested descriptions of protein catabolism and two of type I interferon response with overlapping genes, so this is a single finding.

Surprised by this result, I considered the possibility that it might be an artifact since a scan for depressed TMRCA finds regions of low diversity, and background selection also reduces diversity. I relocated each of the 66 clusters to a block of equal length and similar mean B statistic[44] from the Murphy et al. map[14]. Across 100,000 such block permutations, the null reproduces the number of clusters and annotated genes closely, and none of the 100,000 permutations reaches the observed enrichment for six of the nine categories, including proteasome-mediated ubiquitin-dependent protein catabolism (Table S4.3).

Because the prior against an enrichment for protein catabolism is high, I also asked whether the result depends on where the catalog threshold is set. Holding the Denisovan floor y = 700 ka and the percentile cut $p_{cut} = 50$ fixed, I rebuilt the catalog at six values of the modern-plus-Neandertal ceiling x and repeated the identical test on each rebuild (Table S4.4). The enrichment is strongest in the smaller and highest-confidence catalogs and weakens as the threshold admits more loci, but every one of the nine categories stays at or below a matched permutation $p = 2 \times 10^{-3}$ at every rung from x = 575 to 675 ka, a 3.5-fold range in catalog size, and below $4 \times 10^{-4}$ at every rung from 600 to 675 ka. The signal is lost only at x = 700 ka, where the catalog grows to 829 loci and P(real) falls to 0.38 so that most loci are expected to be false. The loci added beyond the published threshold carry catabolism genes at close to the background rate.

As a second robustness test, I repeated the analysis in a control catalog of similar size obtained by flipping the Neandertal and Denisovan labels in the ascertainment, and relaxing the time thresholds ($T_{Afr+Den} \leq 680$ ka, $T_{Afr+Nea} \geq 700$ ka, $p_{cut} = 50$). This produced a control catalog of 97 clusters and 136 annotated genes. The number of proteasome genes in it was 4 against 3.7 expected, a slight excess rather than a deficit. The 22 exact control regions, before any relaxation, contain no gene in the category against 0.8 expected.

Proteasome-associated genes are a surprising target of distinctively early modern human adaptation, as this is not a functional category that one might expect to be especially shaped by selection in this period, unlike genes involved in neural and behavioral function, skeletal biology, or diet and energetics. But the proteasome has already been singled out as evolving in an unusual way on the human lineage, as in humans its subunits carry high-frequency nonsynonymous polymorphism at sites conserved across mammals, a pattern absent in chimpanzees that was previously hypothesized to reflect a human-specific relaxation of constraint (rather than the positive selection implied here)[45]. A caution is that the previously reported signal lies in the catalytic beta subunits of the 20S core, whereas the subunits here are two alpha subunits (*PSMA1* and *PSMA3*) and two of the 19S regulatory particle (*PSMD1* and *PSMD14*). A signal on the proteasome is suggestive of viral interactions, as it both degrades viral proteins and is extensively subverted by viruses[46]. Virus-interacting proteins have been estimated to account for around 30% of adaptive amino acid substitutions in the mammalian-conserved part of the human proteome[47], and this analysis supports their having shaped unique modern-human biology in a notable way.

**Table S4.4: Stability of the gene ontology enrichment across catalog thresholds**

| **GO biological process** | **x = 575** | **x = 600** | **x = 625** | **x = 650** | **x = 675** | **x = 700** |
|---|---|---|---|---|---|---|
| loci in catalog | **81** | **116** | **141** | **242** | **287** | **829** |
| P(real) | **0.76** | **0.75** | **0.77** | **0.67** | **0.65** | **0.38** |
| macromolecule catabolic process | 3.2<br>($2\times10^{-5}$) | 3.5<br>($<10^{-5}$) | 3.0<br>($<10^{-5}$) | 2.3<br>($2\times10^{-5}$) | 2.0<br>($8\times10^{-5}$) | 1.2<br>(n.t.) |
| response to type I interferon | 12.9<br>($3.7\times10^{-4}$) | 13.7<br>($<10^{-5}$) | 10.9<br>($4\times10^{-5}$) | 6.9<br>($1.0\times10^{-4}$) | 5.7<br>($1.7\times10^{-4}$) | 1.9<br>(n.t.) |
| proteasome-mediated ubiquitin-dependent protein catabolic process | 3.9<br>($5.1\times10^{-4}$) | 5.0<br>($<10^{-5}$) | 4.3<br>($<10^{-5}$) | 3.0<br>($2\times10^{-5}$) | 2.6<br>($1.2\times10^{-4}$) | 1.3<br>(n.t.) |
| ubiquitin-dependent protein catabolic process | 3.7<br>($2.6\times10^{-4}$) | 4.3<br>($<10^{-5}$) | 3.7<br>($<10^{-5}$) | 2.6<br>($2\times10^{-5}$) | 2.3<br>($1.8\times10^{-4}$) | 1.2<br>(n.t.) |
| modification-dependent protein catabolic process | 3.6<br>($2.6\times10^{-4}$) | 4.2<br>($<10^{-5}$) | 3.6<br>($<10^{-5}$) | 2.5<br>($3\times10^{-5}$) | 2.2<br>($1.9\times10^{-4}$) | 1.2<br>(n.t.) |
| modification-dependent macromolecule catabolic process | 3.6<br>($2.6\times10^{-4}$) | 4.2<br>($<10^{-5}$) | 3.6<br>($<10^{-5}$) | 2.5<br>($3\times10^{-5}$) | 2.2<br>($1.9\times10^{-4}$) | 1.2<br>(n.t.) |
| cellular response to type I interferon | 11.3<br>($1.2\times10^{-3}$) | 12.8<br>($4\times10^{-5}$) | 10.2<br>($1.8\times10^{-4}$) | 6.6<br>($1.6\times10^{-4}$) | 5.5<br>($3.3\times10^{-4}$) | 1.8<br>(n.t.) |
| proteasomal protein catabolic process | 3.2<br>($2.0\times10^{-3}$) | 4.2<br>($2\times10^{-5}$) | 3.6<br>($2\times10^{-5}$) | 2.8<br>($5\times10^{-5}$) | 2.4<br>($6\times10^{-5}$) | 1.3<br>(n.t.) |
| protein catabolic process | 2.7<br>($1.8\times10^{-3}$) | 3.2<br>($<10^{-5}$) | 2.9<br>($2\times10^{-5}$) | 2.3<br>($<10^{-5}$) | 2.0<br>($1.9\times10^{-4}$) | 1.1<br>(n.t.) |

Notes: Columns are values of the modern-plus-Neandertal ceiling x in ka, with the Denisovan floor y = 700 ka and the percentile cut $p_{cut}$ = 50 held fixed; x = 625 ka is the published catalog and reproduces Table S4.3 exactly. Categories are the nine of Table S4.3, in the same order. Each cell gives the fold enrichment over the hypergeometric expectation with the B-matched block permutation p beneath it. The permutation is the same 100,000 replicates used for Table S4.3, so "$<10^{-5}$" means that no replicate reached the observed enrichment. "n.t.", not tested: at x = 700 ka none of these categories falls among the 200 most significant that the permutation is run over, and all have nominal $p > 0.05$.

### Software and Reproducibility

A zipped folder associated with this supplementary note (SI4codebase.zip) contains all of the code needed to reproduce the results reported here, using Python 3.9 or later. requirements.txt lists the dependencies and README.md gives the script-by-script pipeline together with the exact commands used for each reported number; every file in the archive is named below. All coordinates are GRCh37 / hg19 and the grid is every 1 kb on the autosomes throughout. Three external inputs are not redistributed: the ARGweaver-D round 2 ARG, the Relate genealogy inferred over 1000 Genomes together with three

Neandertals and a Denisovan, and the high-coverage archaic VCFs; paths to all three are arguments, collected for the archaic genomes in archaic_panel.json.
**The TMRCA scan.** hubisz_out_bed_tmrca_scan.py parses the ARGweaver-D local trees and writes the posterior mean and percentiles of the TMRCA of each haplotype set at every grid point, submitted on the cluster by submit_hubisz_tmrca_slurm.sh with its array task slurm_hubisz_tmrca_array_task.sh and collated by hubisz_tmrca_collate_parts.sh. The haplotype sets are defined in tmrca_queries.hubisz_8afr.json, the eight African diploids behind this note, and are passed to the submitter together with the 1 kb grid step. tmrca_queries.hubisz_8modern.json is the earlier four-modern-genome panel, which was run at 5 kb by the fixed-parameter pair submit_hubisz_tmrca_slurm_8modern.sh and slurm_hubisz_tmrca_array_task_8modern.sh and is retained only for the comparison below. resolve_by_interval.py keeps the read whose 95% posterior interval is narrower in absolute time where ARGweaver-D windows overlap, which is what leaves 2,440,090 autosomal grid points, and approx_percentiles.py reconstructs percentiles between the two reported ones by mapping the mean and bounds onto the model's own log-spaced time grid (times2.txt). compare_hubisz_tmrca_runs.py checks the eight-genome run against the earlier two-genome run, audit_argweaver_consistency.py is the positive control that the haplotype sets nest as the scan assumed, and screen_coverage.py measures the fraction of the autosomes carrying ARG trees.

**Building the catalog.** discriminator_grid.py and threshold_landscape.py produce the case and control counts over the grid of the two archaic thresholds and the posterior cut; den_threshold_test.py checks at matched catalog size whether decoupling the two archaic thresholds improves the contrast, and optimize_rule_by_size.py runs the cross-validated search over all three parameters by chromosome parity, so that no grid point is scored by a model fitted on its own chromosome. make_catalog_2080.py assembles the catalog at the reported rule and make_loci_table.py writes Table S4.1. fit_fdr_surface.py fits the joint binomial logistic regression by Fisher scoring with internal standardization of predictors and a deviance-based line search, build_preal_grid.py evaluates *P(real)* from that surface at every grid point and flags those evaluated outside the fitted region, and p_nea_lt700.py reports the posterior probability that $T_{Afr+Nea}$ is below 700 ka.

**The null, and background selection.** null_ratio_audit.py asks whether the control catalog is an honest null, bstat_to_grid.py places both B statistic maps on the 1 kb grid, null_ratio_shell_bstat.py measures the control-to-case ratio within B shells, and bstat_incremental_value.py tests rules whose threshold shifts with B decile against the flat rule. At matched held-out catalog size none of them improves on it, which is the result reported above.

**Gene and functional annotation.** annotate_catalog_regions.py merges catalog grid points within 5 kb, intersects them with GENCODE v19 gene bodies and adds functional summaries from MyGene.info, with annotate_gene_list_function.py fetching the per-gene prose used in the text; annotate_functional_classes.py classifies the screened sites, reading GENCODE v19 locally and querying phastCons, phyloP, CADD and the ENCODE clusters at the candidate coordinates.

**The fixed derived screen.** preal30_extension.py builds the deliberately looser region set that the screen runs on, sbatch_extract_chagyrskaya.sh re-blocks the Chagyrskaya 8 VCFs, which are plain gzip rather than BGZF; sbatch_index_archaics.sh builds the tabix indices the screen requires and o2_build_venv.sh the pysam environment its job scripts run in; and the screen itself is o2_screen_fixed_derived.py under the job script o2_sbatch_fixed_derived.sh, run once at >99% and once at >95% modern derived frequency. archaic_gt_check_derived_pattern.py and compare_screen_processings.py verify the genotype logic and that the two callers agree where they overlap. fetch_cadd_regions.py pulls CADD v1.6 by region rather than site by site, assemble_fd_annotation.py joins the annotation and resolves the overlap duplicates, annotate_fixed_derived_consequences.py applies the Neandertal condition and resolves codon and amino-acid consequences, canonicalize_protein_positions.py places the residue numbers on canonical transcripts, and make_functional_table.py writes Table S4.2 and holds the single definition of its rule, which build_supp_table2.py imports rather than restates. P(real) for these sites is read at the substitution itself rather than at the nearest lattice point: submit_hubisz_tmrca_sites.sh with slurm_hubisz_tmrca_sites_task.sh runs the same scan at a list of positions, resolve_by_interval.py again chooses between overlapping ARG windows, and site_exact_preal.py evaluates the fitted surface there.

**Relate replication.** sbatch_relate_archaic.sh, or cluster_relate_one_chr.sh for a single chromosome, infers the genealogy, and sbatch_relate_tmrca_chr_array.sh with relate_ts_tmrca_grid.py takes the TMRCA of the modern haplotypes only from the resulting tree sequence onto a fixed grid, with relate_marginal_tmrca_grid.py doing the same within the per-chromosome path, over the chromosomes and lengths in relate_chrom_list_autosomes_22.txt and relate_hg19_chrom_sizes.tsv. relate_locus_percentiles.py places each catalog locus in the genome-wide distribution of locus-sized block means of that quantity, relate_modern_vs_argweaver.py reports the concordance between the two methods, and optimize_relate_rule.py is the ceiling calculation establishing that a Relate-only case and control rule has no discriminating power in that ARG, which is what makes the replication rate interpretable.

**Figures.** fig_s51_model_curve.py supplies the cross-validated surface that fig_s51_composite.py draws as Figure S4.1, both counting cases, controls and confidence intervals with the helpers in fig_s51_sensitivity.py, and fig_locus_zoom.py draws the ten locus panels of Figure S4.2 over the per-grid-point P(real) map from build_preal_grid.py, running each window outward until the African-plus-Neandertal TMRCA recovers above 1,500 ka. recalibrate_peyregne.py puts the ARGweaver-D dates onto the published calibration of ref. 13 and fig_calibration.py draws the calibration panels; both are reported for orientation only, and no result in this note depends on the recalibration.

**Supplementary tables.** build_supp_table1.py writes Supplementary Table 2, a tab-delimited file with 2,440,090 rows, one per autosomal grid point, giving the posterior mean and the 5 and 95 percentile bounds for seven haplotype sets, *P(real)* and whether the point is in the catalog, and the Relate modern-only TMRCA with its genome-wide percentile for a 216-haplotype Yoruba panel and a 604-haplotype Yoruba, British and Han panel. The Relate columns are filled at all but 1,694 of the grid points (0.07%), which lie where the

Relate genealogy stops short of the ARGweaver-D grid, mostly within 200 kb of a chromosome end. build_supp_table2.py writes Supplementary Table 3, a tab-delimited file with 1,223 rows, giving derived-over-observed haplotype counts for modern humans, Neandertals and Denisovans, the per-genome calls behind them, 90% intervals on the three TMRCAs, and the functional annotation. The two script names predate the renumbering of the supplementary tables, so they are offset by one against the tables they write.

**Gene ontology.** great_domains.py builds the regulatory domains from GENCODE v19 and great_test.py runs the hypergeometric and binomial tests together with the B-matched block permutation, with great_matched_100k.py, driven by run_great_100k.sh, repeating the matched permutation at 100,000 replicates over parallel chunks; great_annotation_bias.py provides the annotation-depth controls and great_negative_control.py the Denisovan-side mirror catalog. great_threshold_ladder.sh rebuilds the catalog across the ladder of thresholds in Table S4.4 and repeats the same test on each rebuild. These read go-basic.obo and goa_human.gaf.gz from the Gene Ontology Consortium (release 2026-07-26), which are archived alongside the code.
