## Supplementary Information section 4 code base for "Hypothesis: A modern human range expansion ∼300,000 years ago had a critical role in Neandertal origins": annotate_catalog_regions.py

```
#!/usr/bin/env python3
"""
Annotate a TMRCA catalog (chrom, pos rows on a 5 kb grid) with genes.

Pure-Python replacement for the cluster bedtools pipeline
(overlap_hg19_positions_to_genes.sh + annotate_gene_list_function.py),
with knobs for playing with alternative annotation strategies:

  --cluster-gap BP   merge catalog sites within this many bp into one region
                     (default 5000, i.e. adjacent 5 kb grid hits)
  --flank BP         extend each region by +/- BP before intersecting genes
                     (default 5000, matching the original ±5 kb run)
  --biotypes LIST    comma-separated GENCODE gene_type filter
                     (e.g. protein_coding or protein_coding,lincRNA);
                     default: all biotypes
  --nearest          when a region overlaps no gene (after biotype filter),
                     also report the nearest gene and its distance
  --summaries        fetch functional summaries for overlapping gene symbols
                     from MyGene.info (needs network; summaries align with
                     NCBI Gene). Cached in --summary-cache TSV.

Outputs (TSV, to --out-prefix):
  <prefix>.regions.tsv  one row per region: chrom, start, end, n_sites,
                        mean TMRCAs, gene symbols with biotype and distance
  <prefix>.genes.tsv    one row per (gene, region): symbol, gene_id, biotype,
                        region, distance_bp, plus summary columns if requested

Example:
  python3 scripts/annotate_catalog_regions.py \
      hubisz_8afr_filter_modern_nea_le700_den_gt700.tsv \
      refs/gencode.v19.annotation.gtf.gz \
      --flank 5000 --nearest --summaries \
      --out-prefix catalog_annotation/catalog_929
"""

from __future__ import annotations

import argparse
import csv
import gzip
import json
import os
import sys
import time
import urllib.parse
import urllib.request
from collections import defaultdict

MYGENE_QUERY = "https://mygene.info/v3/query"
MYGENE_FIELDS = "symbol,name,summary,type,map_location,entrezgene,ensembl.gene"
MYGENE_CHUNK = 80


def norm_chrom(c: str) -> str:
    c = c.strip()
    if c.startswith("chr"):
        c = c[3:]
    if c == "23":
        c = "X"
    if c == "24":
        c = "Y"
    if c in ("25", "MT"):
        c = "M"
    return c


def read_catalog(path: str):
    """Return list of dict rows with chrom (normalized), pos (int), and means."""
    rows = []
    with open(path, encoding="utf-8") as f:
        reader = csv.DictReader(f, delimiter="\t")
        for r in reader:
            try:
                pos = int(float(r["pos"]))
            except (KeyError, ValueError):
                continue
            rows.append(
                {
                    "chrom": norm_chrom(r["chrom"]),
                    "pos": pos,
                    "t_afr": r.get("01_modern_only_mean_kya", ""),
                    "t_afr_nea": r.get("02_modern_plus_both_Nea_mean_kya", ""),
                    "t_afr_den": r.get("03_modern_plus_Den_mean_kya", ""),
                }
            )
    return rows


def cluster_sites(rows, gap: int):
    """Group rows into regions: same chrom, successive positions <= gap apart."""
    by_chr = defaultdict(list)
    for r in rows:
        by_chr[r["chrom"]].append(r)
    regions = []
    for chrom, rs in by_chr.items():
        rs.sort(key=lambda r: r["pos"])
        cur = [rs[0]]
        for r in rs[1:]:
            if r["pos"] - cur[-1]["pos"] <= gap:
                cur.append(r)
            else:
                regions.append(cur)
                cur = [r]
        regions.append(cur)

    def chrom_key(region):
        c = region[0]["chrom"]
        return (0, int(c)) if c.isdigit() else (1, c)

    regions.sort(key=lambda reg: (chrom_key(reg), reg[0]["pos"]))
    return regions


def read_genes(gtf_path: str, biotypes: set[str] | None):
    """Parse GENCODE GTF gene lines -> per-chrom sorted list of
    (start, end, symbol, gene_id, biotype). Coordinates 1-based inclusive."""
    genes = defaultdict(list)
    opener = gzip.open if gtf_path.endswith(".gz") else open
    with opener(gtf_path, "rt", encoding="utf-8") as f:
        for line in f:
            if line.startswith("#"):
                continue
            fields = line.rstrip("\n").split("\t")
            if len(fields) < 9 or fields[2] != "gene":
                continue
            attrs = {}
            for item in fields[8].split(";"):
                item = item.strip()
                if not item:
                    continue
                k, _, v = item.partition(" ")
                attrs[k] = v.strip('"')
            btype = attrs.get("gene_type", "")
            if biotypes and btype not in biotypes:
                continue
            genes[norm_chrom(fields[0])].append(
                (int(fields[3]), int(fields[4]), attrs.get("gene_name", ""),
                 attrs.get("gene_id", ""), btype)
            )
    for chrom in genes:
        genes[chrom].sort()
    return genes


def overlap_genes(genes_chr, start: int, end: int):
    """Genes overlapping [start, end] (1-based inclusive). Linear scan is fine
    at this scale (365 regions x <6k genes/chrom)."""
    return [g for g in genes_chr if g[0] <= end and g[1] >= start]


def nearest_gene(genes_chr, start: int, end: int):
    best, best_d = None, None
    for g in genes_chr:
        if g[0] <= end and g[1] >= start:
            return g, 0
        d = g[0] - end if g[0] > end else start - g[1]
        if best_d is None or d < best_d:
            best, best_d = g, d
    return best, best_d


def fetch_summaries(symbols: list[str], cache_path: str | None, sleep: float = 0.15):
    """symbol -> dict with official_name, map_location, entrez_id, summary."""
    cache: dict[str, dict] = {}
    if cache_path and os.path.exists(cache_path):
        with open(cache_path, encoding="utf-8") as f:
            for r in csv.DictReader(f, delimiter="\t"):
                cache[r["symbol"].upper()] = r
    todo = [s for s in dict.fromkeys(symbols) if s.upper() not in cache]
    for i in range(0, len(todo), MYGENE_CHUNK):
        chunk = todo[i : i + MYGENE_CHUNK]
        q = " OR ".join(f"symbol:{s}" for s in chunk)
        params = {"q": q, "species": "human", "fields": MYGENE_FIELDS,
                  "size": max(len(chunk) * 3, 50)}
        url = MYGENE_QUERY + "?" + urllib.parse.urlencode(params)
        req = urllib.request.Request(url, headers={"User-Agent": "annotate_catalog_regions.py"})
        with urllib.request.urlopen(req, timeout=60) as resp:
            hits = json.loads(resp.read().decode("utf-8")).get("hits") or []
        by_sym = defaultdict(list)
        for h in hits:
            s = (h.get("symbol") or "").upper()
            if s:
                by_sym[s].append(h)
        for s in chunk:
            cands = by_sym.get(s.upper(), [])
            if not cands:
                cache[s.upper()] = {"symbol": s, "official_name": "", "map_location": "",
                                    "entrez_id": "", "summary": "", "note": "no_match"}
                continue
            best = max(cands, key=lambda h: len(h.get("summary") or ""))
            cache[s.upper()] = {
                "symbol": s,
                "official_name": " ".join((best.get("name") or "").split()),
                "map_location": best.get("map_location") or "",
                "entrez_id": str(best.get("entrezgene") or ""),
                "summary": " ".join((best.get("summary") or "").split()),
                "note": "ok",
            }
        if i + MYGENE_CHUNK < len(todo):
            time.sleep(sleep)
    if cache_path:
        os.makedirs(os.path.dirname(cache_path) or ".", exist_ok=True)
        with open(cache_path, "w", encoding="utf-8", newline="") as f:
            w = csv.DictWriter(f, fieldnames=["symbol", "official_name", "map_location",
                                              "entrez_id", "summary", "note"],
                               delimiter="\t", extrasaction="ignore")
            w.writeheader()
            for k in sorted(cache):
                w.writerow(cache[k])
    return cache


def mean_of(rows, key):
    vals = [float(r[key]) for r in rows if r[key] not in ("", None)]
    return round(sum(vals) / len(vals)) if vals else ""


def main() -> int:
    ap = argparse.ArgumentParser(description=__doc__.splitlines()[1])
    ap.add_argument("catalog")
    ap.add_argument("gtf")
    ap.add_argument("--cluster-gap", type=int, default=5000)
    ap.add_argument("--flank", type=int, default=5000)
    ap.add_argument("--grid-step", type=int, default=5000,
                    help="bp a grid point stands for; 1000 for the 1 kb scan. "
                         "Only affects span_kb, which has to cover the last "
                         "grid point's territory. --flank stays 5000 because "
                         "'genes within 5 kb' is a biological choice, not a "
                         "property of the lattice.")
    ap.add_argument("--biotypes", default="",
                    help="comma-separated gene_type filter; empty = all")
    ap.add_argument("--nearest", action="store_true")
    ap.add_argument("--summaries", action="store_true")
    ap.add_argument("--summary-cache", default="catalog_annotation/mygene_cache.tsv")
    ap.add_argument("--out-prefix", required=True)
    args = ap.parse_args()

    biotypes = set(b.strip() for b in args.biotypes.split(",") if b.strip()) or None
    rows = read_catalog(args.catalog)
    regions = cluster_sites(rows, args.cluster_gap)
    genes = read_genes(args.gtf, biotypes)
    print(f"{len(rows)} sites -> {len(regions)} regions "
          f"(gap {args.cluster_gap}); GTF genes kept: "
          f"{sum(len(v) for v in genes.values())}", file=sys.stderr)

    region_rows, gene_rows = [], []
    for reg in regions:
        chrom = reg[0]["chrom"]
        start, end = reg[0]["pos"], reg[-1]["pos"]
        qs, qe = start - args.flank, end + args.flank
        hits = overlap_genes(genes.get(chrom, []), qs, qe)
        gene_field = []
        for (gs, ge, sym, gid, btype) in hits:
            d = 0 if (gs <= end and ge >= start) else (gs - end if gs > end else start - ge)
            gene_field.append(f"{sym}|{btype}|{d}")
            gene_rows.append({"chrom": chrom, "region_start": start, "region_end": end,
                              "n_sites": len(reg), "symbol": sym, "gene_id": gid,
                              "biotype": btype, "distance_bp": d, "relation": "overlap_flank"})
        near_field = ""
        if args.nearest and not hits:
            g, d = nearest_gene(genes.get(chrom, []), qs, qe)
            if g:
                near_field = f"{g[2]}|{g[4]}|{d + args.flank}"
                gene_rows.append({"chrom": chrom, "region_start": start, "region_end": end,
                                  "n_sites": len(reg), "symbol": g[2], "gene_id": g[3],
                                  "biotype": g[4], "distance_bp": d + args.flank,
                                  "relation": "nearest"})
        region_rows.append({
            "chrom": chrom, "start": start, "end": end, "n_sites": len(reg),
            "span_kb": (end - start + args.grid_step) // 1000,
            "mean_t_afr": mean_of(reg, "t_afr"),
            "mean_t_afr_nea": mean_of(reg, "t_afr_nea"),
            "mean_t_afr_den": mean_of(reg, "t_afr_den"),
            "genes": ";".join(gene_field),
            "nearest_if_none": near_field,
        })

    if args.summaries:
        syms = [g["symbol"] for g in gene_rows if g["symbol"]]
        cache = fetch_summaries(syms, args.summary_cache)
        for g in gene_rows:
            info = cache.get(g["symbol"].upper(), {})
            g["official_name"] = info.get("official_name", "")
            g["map_location"] = info.get("map_location", "")
            g["entrez_id"] = info.get("entrez_id", "")
            g["summary"] = info.get("summary", "")

    os.makedirs(os.path.dirname(args.out_prefix) or ".", exist_ok=True)
    with open(args.out_prefix + ".regions.tsv", "w", encoding="utf-8", newline="") as f:
        w = csv.DictWriter(f, fieldnames=list(region_rows[0].keys()), delimiter="\t")
        w.writeheader()
        w.writerows(region_rows)
    gene_fields = ["chrom", "region_start", "region_end", "n_sites", "symbol", "gene_id",
                   "biotype", "distance_bp", "relation"]
    if args.summaries:
        gene_fields += ["official_name", "map_location", "entrez_id", "summary"]
    with open(args.out_prefix + ".genes.tsv", "w", encoding="utf-8", newline="") as f:
        w = csv.DictWriter(f, fieldnames=gene_fields, delimiter="\t", extrasaction="ignore")
        w.writeheader()
        w.writerows(gene_rows)
    print(f"wrote {args.out_prefix}.regions.tsv ({len(region_rows)} regions), "
          f"{args.out_prefix}.genes.tsv ({len(gene_rows)} gene rows)", file=sys.stderr)
    return 0


if __name__ == "__main__":
    raise SystemExit(main())
```
