## Supplementary Information section 4 code base for "Hypothesis: A modern human range expansion ∼300,000 years ago had a critical role in Neandertal origins": annotate_fixed_derived_consequences.py

```
#!/usr/bin/env python3
"""
Post-process the fixed-derived screen: apply the Neandertal condition and
resolve protein consequences for coding sites.

Two things the screen itself does not do.

(1) The Neandertal requirement. screen_fixed_derived_in_clusters.py enforces
    only "derived and near-fixed in modern humans, ancestral in Denisova", and
    records the Altai and Vindija states as columns without filtering on them.
    The catalog premise is stronger: the derived allele should be shared by
    modern humans AND Neandertals to the exclusion of Denisova. This script
    applies that condition, with --neandertal controlling how strictly:

      both  (default) Altai and Vindija both homozygous derived
      any             at least one homozygous derived, the other not ancestral
      none            no requirement, i.e. the screen's own output

(2) Protein consequences. The screen labels genomic context (CDS / exon /
    intron / intergenic) but never asks whether a CDS hit changes an amino
    acid. There is no local hg19 reference to build codons from, so coding
    sites are sent to the Ensembl GRCh37 VEP REST endpoint, which returns the
    consequence, the amino acid pair and the protein position.

    Direction of the query. At these sites the DERIVED allele is the hg19
    reference base and the ANCESTRAL allele is the alternate. VEP therefore
    annotates the reverse of the mutation that actually occurred, and its
    "amino_acids" field reads reference/alternate = derived/ancestral. The
    codon change and its missense-versus-synonymous status are symmetric, so
    the consequence is valid as returned; only the labels are swapped, and the
    output columns below name them derived_aa and ancestral_aa accordingly.

Usage:
  python3 scripts/annotate_fixed_derived_consequences.py \
      --sites catalog_annotation/fixed_derived_preal50.sites.tsv \
      -o catalog_annotation/fixed_derived_preal50.consequences.tsv
"""

from __future__ import annotations

import argparse
import csv
import json
import os
import sys
import time
import urllib.error
import urllib.request

VEP_URL = "https://grch37.rest.ensembl.org/vep/human/region"
BATCH = 180

# The screen writes these labels for each Neandertal.
DERIVED = "derived_hom"
ANCESTRAL = "ancestral_hom"


def neandertal_ok(row, mode):
    a, v = row.get("altai", ""), row.get("vindija", "")
    if mode == "none":
        return True
    if mode == "both":
        return a == DERIVED and v == DERIVED
    if mode == "any":
        return (a == DERIVED or v == DERIVED) and ANCESTRAL not in (a, v)
    raise ValueError(mode)


def vep_post(variants, tries=4, delay=4):
    """POST a batch of 'chrom pos . ref alt . . .' strings to GRCh37 VEP."""
    body = json.dumps({"variants": variants}).encode()
    req = urllib.request.Request(
        VEP_URL, data=body,
        headers={"Content-Type": "application/json",
                 "Accept": "application/json"})
    for i in range(tries):
        try:
            with urllib.request.urlopen(req, timeout=120) as r:
                return json.load(r)
        except (urllib.error.URLError, urllib.error.HTTPError, TimeoutError) as e:
            if i == tries - 1:
                sys.stderr.write(f"WARN: VEP request failed: {e}\n")
                return []
            time.sleep(delay * (i + 1))
    return []


def pick_consequence(entry):
    """Most severe coding consequence across transcripts, preferring canonical."""
    tcs = entry.get("transcript_consequences") or []
    coding = [t for t in tcs if t.get("amino_acids") or
              any("synonymous" in c or "missense" in c or "stop" in c or "start" in c
                  for c in (t.get("consequence_terms") or []))]
    if not coding:
        return None
    rank = {"stop_gained": 0, "start_lost": 1, "stop_lost": 1, "missense_variant": 2,
            "synonymous_variant": 3, "stop_retained_variant": 3}

    def key(t):
        terms = t.get("consequence_terms") or []
        best = min((rank.get(c, 9) for c in terms), default=9)
        return (0 if t.get("canonical") else 1, best)

    coding.sort(key=key)
    return coding[0]


def main() -> int:
    ap = argparse.ArgumentParser()
    ap.add_argument("--sites", default="catalog_annotation/fixed_derived_preal50.sites.tsv")
    ap.add_argument("--neandertal", choices=("both", "any", "none"), default="both")
    ap.add_argument("--no-vep", action="store_true",
                    help="skip the Ensembl queries and only apply the filters")
    ap.add_argument("-o", "--out",
                    default="catalog_annotation/fixed_derived_preal50.consequences.tsv")
    args = ap.parse_args()

    if not os.path.exists(args.sites):
        sys.stderr.write(f"sites file not found: {args.sites}\n"
                         "(the screen may still be running)\n")
        return 1

    with open(args.sites, encoding="utf-8") as f:
        rows = list(csv.DictReader(f, delimiter="\t"))
    print(f"{len(rows):,} sites from the screen "
          f"(moderns derived and near-fixed, Denisova ancestral)")

    # ---- Neandertal state breakdown, before filtering ----
    combos = {}
    for r in rows:
        combos[(r.get("altai", ""), r.get("vindija", ""))] = \
            combos.get((r.get("altai", ""), r.get("vindija", "")), 0) + 1
    print("\nNeandertal states (Altai, Vindija):")
    for k, n in sorted(combos.items(), key=lambda kv: -kv[1]):
        print(f"  {k[0]:<14} {k[1]:<14} {n:>6,}")

    kept = [r for r in rows if neandertal_ok(r, args.neandertal)]
    print(f"\n--neandertal {args.neandertal}: {len(kept):,} of {len(rows):,} sites retained")

    ctx = {}
    for r in kept:
        ctx[r["context"]] = ctx.get(r["context"], 0) + 1
    print(f"  by context: {ctx}")

    coding = [r for r in kept if r["context"] == "CDS"]
    print(f"  coding (CDS): {len(coding):,}")

    # ---- protein consequences for coding sites ----
    ann = {}
    if coding and not args.no_vep:
        variants = []
        for r in coding:
            # derived == hg19 reference; ancestral == the alternate we query
            variants.append(f"{r['chrom']} {r['pos']} . "
                            f"{r['derived_ref']} {r['ancestral']} . . .")
        print(f"\nquerying Ensembl GRCh37 VEP for {len(variants)} coding sites ...")
        for i in range(0, len(variants), BATCH):
            chunk = variants[i:i + BATCH]
            for entry in vep_post(chunk):
                inp = entry.get("input", "")
                parts = inp.split()
                if len(parts) >= 2:
                    ann[(parts[0], parts[1])] = entry
            time.sleep(0.4)
        print(f"  annotated {len(ann)} of {len(variants)}")

    # ---- output ----
    cols = ["chrom", "pos", "ancestral", "derived_ref", "g1k_alt_af",
            "g1k_polymorphic", "denisova_gt", "altai", "vindija",
            "context", "gene", "cluster_chrom", "cluster_start", "cluster_end",
            "consequence", "derived_aa", "ancestral_aa", "protein_position",
            "transcript", "canonical", "sift", "polyphen"]
    os.makedirs(os.path.dirname(args.out) or ".", exist_ok=True)
    n_missense = n_syn = 0
    with open(args.out, "w", newline="", encoding="utf-8") as f:
        w = csv.writer(f, delimiter="\t")
        w.writerow(cols)
        for r in kept:
            e = ann.get((r["chrom"], str(r["pos"])))
            t = pick_consequence(e) if e else None
            cons = aa_d = aa_a = ppos = tx = canon = sift = pph = ""
            if t:
                terms = t.get("consequence_terms") or []
                cons = ",".join(terms)
                aas = (t.get("amino_acids") or "")
                if "/" in aas:
                    aa_d, aa_a = aas.split("/", 1)
                elif aas:
                    aa_d = aa_a = aas
                ppos = t.get("protein_start") or ""
                tx = t.get("transcript_id") or ""
                canon = "yes" if t.get("canonical") else ""
                sift = t.get("sift_prediction") or ""
                pph = t.get("polyphen_prediction") or ""
                if "missense_variant" in terms:
                    n_missense += 1
                if "synonymous_variant" in terms:
                    n_syn += 1
            w.writerow([r["chrom"], r["pos"], r["ancestral"], r["derived_ref"],
                        r.get("g1k_alt_af", ""), r.get("g1k_polymorphic", ""),
                        r.get("denisova_gt", ""), r.get("altai", ""),
                        r.get("vindija", ""), r["context"], r.get("gene", ""),
                        r.get("cluster_chrom", ""), r.get("cluster_start", ""),
                        r.get("cluster_end", ""),
                        cons, aa_d, aa_a, ppos, tx, canon, sift, pph])

    print(f"\nwrote {args.out}")
    if coding and not args.no_vep:
        print(f"  missense: {n_missense}, synonymous: {n_syn}")
        print("  amino acid columns read derived -> ancestral, i.e. the residue "
              "carried by modern humans and Neandertals versus the one Denisova "
              "retains.")
    return 0


if __name__ == "__main__":
    raise SystemExit(main())
```
