## Supplementary Information section 4 code base for "Hypothesis: A modern human range expansion ∼300,000 years ago had a critical role in Neandertal origins": annotate_functional_classes.py

```
#!/usr/bin/env python3
"""
Classify fixed-derived sites by function, and mark the ones worth following up.

WHY THIS IS A SEPARATE STEP
The screen answers a question about ancestry -- is this site fixed derived in
modern humans and Neandertals while Denisova is ancestral -- and the great
majority of the sites it returns are intronic or intergenic, where the ancestry
pattern is all there is to say. The interesting subset is the one that can
plausibly do something. Splitting the classification out means the ancestry
screen can keep running on O2 against the genotype data, which is where that
data lives, while this step runs anywhere with a network connection.

WHY NOTHING LARGE IS DOWNLOADED
CADD v1.6 for GRCh37 is about 80 GB and the phyloP alignment is several more.
Both are indexed, so both can be read at a coordinate over HTTPS: a CADD lookup
costs roughly a second, which for a few hundred candidate sites is a couple of
minutes against days of transfer and a permanent disk cost. The two small
interval sets, phastCons elements and the ENCODE clusters, are cached locally
on first use because they are tens of megabytes and are scanned repeatedly.

THE CLASSES
Gene structure, from GENCODE v19 against GRCh37:
  protein_altering   missense, stop gain, stop loss, start loss
  synonymous         coding but not protein-altering; recorded, and qualifying
                     only with --include-synonymous
  splice_essential   the two intronic bases at a donor or acceptor boundary
  splice_region      three exonic or eight intronic bases around a boundary
  utr5, utr3         untranslated regions of a protein-coding transcript
  promoter           within 1 kb upstream of a protein-coding TSS
Independent of gene structure:
  phastcons          inside a phastCons 46-way placental conserved element
  phylop             phyloP 46-way placental score above --phylop-min
  cadd               CADD v1.6 PHRED at or above --cadd-min for the observed
                     substitution, not the site's maximum over alternates
  encode_dnase       inside an ENCODE DNase hypersensitivity cluster (hg19)
  encode_tfbs        inside an ENCODE TFBS cluster (hg19)

Everything is hg19-native. No liftover is performed anywhere, which is the
reason the ENCODE clusters are used rather than the current cCRE registry: that
is built on hg38, and lifting regulatory intervals onto the coordinates of a
different assembly introduces exactly the kind of quiet error this catalog
cannot afford.

Classes are assigned over all protein-coding transcripts, not a single
canonical one, so a position can be both CDS in one transcript and 3'UTR in
another and will carry both labels. That is a property of the annotation rather
than an inconsistency, and keeping both is more informative than picking one.

A site qualifies if it belongs to any qualifying class. Every class it belongs
to is recorded, so the definition can be narrowed afterwards from the output
table without paying for the lookups again.

Example:
  python3 annotate_functional_classes.py \
      -i catalog_annotation/fixed_derived_preal50.sites.tsv \
      -o catalog_annotation/fixed_derived_functional.tsv
"""

from __future__ import annotations

import argparse
import bisect
import csv
import gzip
import os
import sys
import urllib.request
from collections import defaultdict

REF = os.path.join(os.path.dirname(os.path.dirname(os.path.abspath(__file__))),
                   "refdata")
GTF = os.path.join(REF, "gencode.v19.annotation.gtf.gz")
UCSC = "https://hgdownload.soe.ucsc.edu/goldenPath/hg19/database"
# Overridable so the screen can read a local tabix subset instead of seeking
# into the remote 80 GB file for every site. That matters more than it sounds:
# a remote per-site fetch costs ~11 s against ~0.0001 s locally, because the
# cost is the seek rather than the bytes, which is the difference between a
# day and a minute over the screened sites. scripts/fetch_cadd_regions.py
# builds the subset; set ANNOT_CADD_URL to its output. Only do that when every
# site being annotated lies inside the fetched regions, since a site outside
# them gets no CADD score rather than an error.
CADD = os.environ.get(
    "ANNOT_CADD_URL",
    "https://kircherlab.bihealth.org/download/CADD/v1.6/GRCh37/"
    "whole_genome_SNVs.tsv.gz")
# Overridable because the phyloP track is read by random access over HTTP, and
# some networks resolve hgdownload.soe.ucsc.edu but refuse it, or the reverse.
# Where DNS is the only thing blocked, pointing this at UCSC's address over
# plain HTTP works and is far cheaper than fetching the whole 8.6 GB file:
#   ANNOT_PHYLOP_URL=http://128.114.119.163/goldenPath/hg19/phyloP46way/\
#   placentalMammals.phyloP46way.bw
# A local copy of the .bw is equally valid and is what to prefer if there is
# one, since the address above is not guaranteed to stay put.
PHYLOP = os.environ.get(
    "ANNOT_PHYLOP_URL",
    "https://hgdownload.soe.ucsc.edu/goldenPath/hg19/phyloP46way/"
    "placentalMammals.phyloP46way.bw")

GENE_CLASSES = ["protein_altering", "synonymous", "splice_essential",
                "splice_region", "utr5", "utr3", "promoter"]
EXTRA_CLASSES = ["phastcons", "phylop", "cadd", "encode_dnase", "encode_tfbs"]
DEFAULT_QUALIFY = [c for c in GENE_CLASSES if c != "synonymous"] + EXTRA_CLASSES

CODING_ALTERING = {"missense", "nonsense", "stop_gain", "stop_loss",
                   "start_loss", "stoploss", "startloss"}


def cache(url, name):
    """Fetch a small interval file once and keep it."""
    path = os.path.join(REF, name)
    if not os.path.exists(path):
        os.makedirs(REF, exist_ok=True)
        print(f"  fetching {name} ...", file=sys.stderr)
        urllib.request.urlretrieve(url, path)
    return path


class Intervals:
    """Sorted per-chromosome intervals with a point-membership test."""

    def __init__(self):
        self.starts = defaultdict(list)
        self.ends = defaultdict(list)

    def add(self, c, s, e):
        self.starts[c].append(s)
        self.ends[c].append(e)

    def finish(self):
        for c in self.starts:
            order = sorted(range(len(self.starts[c])),
                           key=lambda i: self.starts[c][i])
            self.starts[c] = [self.starts[c][i] for i in order]
            ends = [self.ends[c][i] for i in order]
            # running maximum, so a binary search on starts is conclusive
            run, m = [], 0
            for e in ends:
                m = max(m, e)
                run.append(m)
            self.ends[c] = run
        return self

    def hit(self, c, p):
        ss = self.starts.get(c)
        if not ss:
            return False
        i = bisect.bisect_right(ss, p) - 1
        return i >= 0 and self.ends[c][i] >= p


def load_ucsc_bed(name, chrom_col=1, start_col=2, end_col=3):
    """UCSC database dumps are BED-like with a leading bin column."""
    path = cache(f"{UCSC}/{name}.txt.gz", f"{name}.txt.gz")
    iv = Intervals()
    with gzip.open(path, "rt", errors="replace") as f:
        for line in f:
            fl = line.rstrip("\n").split("\t")
            if len(fl) <= end_col:
                continue
            c = fl[chrom_col].replace("chr", "")
            try:
                iv.add(c, int(fl[start_col]) + 1, int(fl[end_col]))
            except ValueError:
                continue
    return iv.finish()


def load_gencode(path):
    """Splice, UTR and promoter intervals for protein-coding transcripts."""
    splice_ess, splice_reg = Intervals(), Intervals()
    utr5, utr3, promoter = Intervals(), Intervals(), Intervals()
    exons = defaultdict(list)          # tx -> [(start, end)]
    meta = {}                          # tx -> (chrom, strand, type)
    utr_raw = defaultdict(list)
    cds_span = {}
    op = gzip.open if path.endswith(".gz") else open
    with op(path, "rt", errors="replace") as f:
        for line in f:
            if line.startswith("#"):
                continue
            fl = line.rstrip("\n").split("\t")
            if len(fl) < 9:
                continue
            kind, c, s, e, strand = fl[2], fl[0].replace("chr", ""), \
                int(fl[3]), int(fl[4]), fl[6]
            if kind not in ("exon", "UTR", "CDS", "transcript"):
                continue
            attrs = fl[8]
            if 'transcript_type "protein_coding"' not in attrs:
                continue
            tx = ""
            for item in attrs.split(";"):
                item = item.strip()
                if item.startswith("transcript_id "):
                    tx = item[len("transcript_id "):].strip('"')
                    break
            if not tx:
                continue
            meta[tx] = (c, strand)
            if kind == "exon":
                exons[tx].append((s, e))
            elif kind == "UTR":
                utr_raw[tx].append((s, e))
            elif kind == "CDS":
                lo, hi = cds_span.get(tx, (s, e))
                cds_span[tx] = (min(lo, s), max(hi, e))
            elif kind == "transcript":
                tss = s if strand == "+" else e
                if strand == "+":
                    promoter.add(c, max(1, tss - 1000), tss)
                else:
                    promoter.add(c, tss, tss + 1000)

    for tx, ex in exons.items():
        c, strand = meta[tx]
        ex.sort()
        for n, (s, e) in enumerate(ex):
            # A boundary is a splice site only if it is internal to the
            # transcript: the outer edges of the first and last exon are the
            # transcript ends, not junctions.
            if n > 0:
                splice_ess.add(c, s - 2, s - 1)
                splice_reg.add(c, s - 8, s + 2)
            if n < len(ex) - 1:
                splice_ess.add(c, e + 1, e + 2)
                splice_reg.add(c, e - 2, e + 8)

    # A UTR record is 5' or 3' according to which side of the CDS it lies on.
    for tx, blocks in utr_raw.items():
        if tx not in cds_span:
            continue
        c, strand = meta[tx]
        lo, hi = cds_span[tx]
        for s, e in blocks:
            upstream = e < lo if strand == "+" else s > hi
            (utr5 if upstream else utr3).add(c, s, e)

    return {"splice_essential": splice_ess.finish(),
            "splice_region": splice_reg.finish(),
            "utr5": utr5.finish(), "utr3": utr3.finish(),
            "promoter": promoter.finish()}


def main():
    ap = argparse.ArgumentParser()
    ap.add_argument("-i", "--infile", required=True)
    ap.add_argument("-o", "--outfile", required=True)
    ap.add_argument("--gtf", default=GTF)
    ap.add_argument("--cadd-min", type=float, default=20.0)
    ap.add_argument("--phylop-min", type=float, default=2.0)
    ap.add_argument("--include-synonymous", action="store_true")
    ap.add_argument("--qualify", default=",".join(DEFAULT_QUALIFY),
                    help="comma-separated class names that make a site qualify")
    ap.add_argument("--no-remote", action="store_true",
                    help="gene-structure classes only; skip network lookups")
    a = ap.parse_args()

    qual = set(a.qualify.split(","))
    if a.include_synonymous:
        qual.add("synonymous")
    else:
        qual.discard("synonymous")

    with open(a.infile, newline="") as f:
        rows = list(csv.DictReader(f, delimiter="\t"))
    print(f"{len(rows):,} sites from {os.path.basename(a.infile)}")

    print("loading GENCODE v19 ...")
    iv = load_gencode(a.gtf)
    if not a.no_remote:
        print("loading interval sets ...")
        iv["phastcons"] = load_ucsc_bed("phastConsElements46way")
        iv["encode_dnase"] = load_ucsc_bed("wgEncodeRegDnaseClusteredV3")
        iv["encode_tfbs"] = load_ucsc_bed("wgEncodeRegTfbsClusteredV3")

    bw = tb = None
    if not a.no_remote:
        import pyBigWig
        import pysam
        print("opening remote phyloP and CADD ...")
        bw = pyBigWig.open(PHYLOP)
        tb = pysam.TabixFile(CADD)

    counts = defaultdict(int)
    for n, r in enumerate(rows, 1):
        c = r["chrom"].replace("chr", "")
        p = int(r["pos"])
        cls = []

        # Coding consequence, taken from whichever column the screen wrote.
        cons = (r.get("consequence") or "").strip().lower()
        ctx = (r.get("context") or "").strip()
        if cons in CODING_ALTERING:
            cls.append("protein_altering")
        elif cons == "synonymous":
            cls.append("synonymous")
        elif ctx == "CDS" and not cons:
            cls.append("coding_unresolved")

        for name in ("splice_essential", "splice_region", "utr5", "utr3",
                     "promoter", "phastcons", "encode_dnase", "encode_tfbs"):
            if name in iv and iv[name].hit(c, p):
                cls.append(name)

        phylop = cadd = cadd_max = ""
        if bw is not None:
            try:
                v = bw.values(f"chr{c}", p - 1, p)[0]
                if v is not None and v == v:
                    phylop = round(float(v), 3)
                    if phylop >= a.phylop_min:
                        cls.append("phylop")
            except (RuntimeError, IndexError):
                pass
        if tb is not None:
            # Every site here is fixed derived in all modern humans, so the
            # derived allele is the hg19 reference and CADD, which scores
            # ref->alt, has no row for ancestral->derived. The row that does
            # exist is the reverse substitution, derived->ancestral, and its
            # score is the closest thing available to a score for the change
            # that happened. The maximum over all three alternates is recorded
            # alongside it as a measure of how constrained the position is
            # regardless of direction, and that is what qualifies a site: the
            # question being asked is whether the position matters, not whether
            # one particular substitution at it is damaging.
            anc = (r.get("ancestral") or "").strip().upper()
            best = None
            try:
                for line in tb.fetch(c, p - 1, p):
                    fl = line.split("\t")
                    if len(fl) < 6:
                        continue
                    v = float(fl[5])
                    best = v if best is None else max(best, v)
                    if anc and fl[3].upper() == anc:
                        cadd = v
            except (ValueError, OSError):
                pass
            cadd_max = "" if best is None else round(best, 3)
            if best is not None and best >= a.cadd_min:
                cls.append("cadd")

        r["classes"] = ";".join(cls)
        r["phylop"] = phylop
        r["cadd_reversion"] = cadd
        r["cadd_max"] = cadd_max
        r["qualifying"] = "yes" if (set(cls) & qual) else "no"
        for k in cls:
            counts[k] += 1
        if n % 25 == 0:
            print(f"  {n}/{len(rows)}", file=sys.stderr)

    keep = sum(1 for r in rows if r["qualifying"] == "yes")
    fields = list(rows[0].keys())
    with open(a.outfile, "w", newline="") as f:
        w = csv.DictWriter(f, fieldnames=fields, delimiter="\t",
                           extrasaction="ignore")
        w.writeheader()
        w.writerows(rows)

    print(f"\nqualifying: {keep} of {len(rows)} sites")
    print("by class:")
    for k in GENE_CLASSES + EXTRA_CLASSES + ["coding_unresolved"]:
        if counts.get(k):
            mark = "*" if k in qual else " "
            print(f" {mark} {k:<18} {counts[k]:>5}")
    print(f"\n(* counts toward qualifying)\nwrote {a.outfile}")
    return 0


if __name__ == "__main__":
    raise SystemExit(main())
```
