## Supplementary Information section 4 code base for "Hypothesis: A modern human range expansion ∼300,000 years ago had a critical role in Neandertal origins": annotate_gene_list_function.py

```
#!/usr/bin/env python3
"""
Fetch short functional annotation for a list of human gene symbols.

Uses the public MyGene.info API (https://mygene.info/) — summary text is
typically aligned with NCBI Gene. No API key required; be gentle with large lists
(run once, cache output).

Input: one gene symbol per line (or first column of TSV). Lines starting with #
and empty lines are skipped.

Output: TSV to stdout with columns:
  symbol, official_name, gene_type, cytogenetic_location, entrez_id,
  ensembl_gene_id, summary_paragraph

Example:
  python3 annotate_gene_list_function.py genes.txt > genes_annotated.tsv
  python3 annotate_gene_list_function.py genes.txt --sleep 0.15 -o genes_annotated.tsv
"""

from __future__ import annotations

import argparse
import csv
import json
import sys
import time
import urllib.error
import urllib.parse
import urllib.request
from typing import Any

MYGENE_QUERY = "https://mygene.info/v3/query"
DEFAULT_FIELDS = (
    "symbol,name,summary,type,map_location,entrezgene,ensembl.gene"
)
CHUNK = 80  # symbols per request (keep URL size reasonable)
DEFAULT_SLEEP = 0.12  # seconds between chunk requests


def read_symbols(path: str) -> list[str]:
    symbols: list[str] = []
    seen: set[str] = set()
    opener = open(path, encoding="utf-8", newline="")
    with opener as f:
        for line in f:
            line = line.strip()
            if not line or line.startswith("#"):
                continue
            sym = line.split()[0].strip()
            if not sym:
                continue
            key = sym.upper()
            if key in seen:
                continue
            seen.add(key)
            symbols.append(sym)
    return symbols


def fetch_chunk(symbols: list[str], fields: str, timeout: float) -> list[dict[str, Any]]:
    """OR-query MyGene for a batch of symbols; return raw hits."""
    q = " OR ".join(f"symbol:{s}" for s in symbols)
    params = {
        "q": q,
        "species": "human",
        "fields": fields,
        "size": max(len(symbols) * 3, 50),
    }
    url = MYGENE_QUERY + "?" + urllib.parse.urlencode(params)
    req = urllib.request.Request(
        url,
        headers={"User-Agent": "annotate_gene_list_function.py (research; contact lab)"},
    )
    with urllib.request.urlopen(req, timeout=timeout) as resp:
        data = json.loads(resp.read().decode("utf-8"))
    return list(data.get("hits") or [])


def _one_line(s: str) -> str:
    """Single line safe for TSV (no tabs or newlines)."""
    if not s:
        return ""
    return " ".join(s.replace("\t", " ").split())


def normalize_hit(hit: dict[str, Any]) -> dict[str, Any]:
    sym = (hit.get("symbol") or "").strip()
    ensembl = hit.get("ensembl")
    ensembl_id = ""
    if isinstance(ensembl, dict):
        ensembl_id = str(ensembl.get("gene") or "")
    elif isinstance(ensembl, list) and ensembl:
        g = ensembl[0].get("gene") if isinstance(ensembl[0], dict) else None
        ensembl_id = str(g or "")
    summary = hit.get("summary") or ""
    summary = _one_line(summary)
    return {
        "symbol": sym,
        "official_name": _one_line((hit.get("name") or "").strip()),
        "gene_type": _one_line((hit.get("type") or "").strip()),
        "cytogenetic_location": _one_line((hit.get("map_location") or "").strip()),
        "entrez_id": str(hit.get("entrezgene") or ""),
        "ensembl_gene_id": _one_line(ensembl_id),
        "summary_paragraph": summary,
    }


def match_hits_to_symbols(
    symbols: list[str], hits: list[dict[str, Any]]
) -> dict[str, dict[str, Any]]:
    """Map requested symbol -> best matching hit (exact symbol, human)."""
    by_upper: dict[str, list[dict[str, Any]]] = {}
    for h in hits:
        row = normalize_hit(h)
        s = row["symbol"]
        if not s:
            continue
        by_upper.setdefault(s.upper(), []).append(row)

    out: dict[str, dict[str, Any]] = {}
    for sym in symbols:
        u = sym.upper()
        candidates = by_upper.get(u) or []
        if not candidates:
            out[sym] = {
                "symbol": sym,
                "official_name": "",
                "gene_type": "",
                "cytogenetic_location": "",
                "entrez_id": "",
                "ensembl_gene_id": "",
                "summary_paragraph": "",
                "note": "no_match",
            }
            continue
        # prefer longest summary if multiple
        best = max(candidates, key=lambda r: len(r["summary_paragraph"]))
        best = dict(best)
        best["note"] = "ok" if len(candidates) == 1 else "ambiguous_multiple_hits"
        out[sym] = best
    return out


def main() -> int:
    ap = argparse.ArgumentParser(
        description="Annotate human gene symbols with names and function summaries (MyGene.info)."
    )
    ap.add_argument(
        "gene_list",
        help="File with one symbol per line (or TSV; first column used)",
    )
    ap.add_argument(
        "-o",
        "--output",
        help="Write TSV here instead of stdout",
    )
    ap.add_argument(
        "--sleep",
        type=float,
        default=DEFAULT_SLEEP,
        help=f"Seconds to sleep between API chunk requests (default {DEFAULT_SLEEP})",
    )
    ap.add_argument(
        "--timeout",
        type=float,
        default=60.0,
        help="HTTP timeout seconds (default 60)",
    )
    args = ap.parse_args()

    symbols = read_symbols(args.gene_list)
    if not symbols:
        print("No gene symbols found in input.", file=sys.stderr)
        return 1

    all_hits_by_symbol: dict[str, dict[str, Any]] = {}
    for i in range(0, len(symbols), CHUNK):
        chunk = symbols[i : i + CHUNK]
        try:
            hits = fetch_chunk(chunk, DEFAULT_FIELDS, args.timeout)
        except urllib.error.HTTPError as e:
            print(f"MyGene HTTP error: {e}", file=sys.stderr)
            return 1
        except urllib.error.URLError as e:
            print(f"Network error: {e}", file=sys.stderr)
            return 1
        merged = match_hits_to_symbols(chunk, hits)
        all_hits_by_symbol.update(merged)
        if i + CHUNK < len(symbols) and args.sleep > 0:
            time.sleep(args.sleep)

    # preserve input order
    rows = [all_hits_by_symbol[s] for s in symbols]

    fieldnames = [
        "symbol",
        "official_name",
        "gene_type",
        "cytogenetic_location",
        "entrez_id",
        "ensembl_gene_id",
        "summary_paragraph",
        "note",
    ]
    out_fp = open(args.output, "w", encoding="utf-8", newline="") if args.output else sys.stdout
    try:
        w = csv.DictWriter(
            out_fp,
            fieldnames=fieldnames,
            delimiter="\t",
            extrasaction="ignore",
            quoting=csv.QUOTE_MINIMAL,
        )
        w.writeheader()
        w.writerows(rows)
    finally:
        if args.output:
            out_fp.close()

    missing = sum(1 for r in rows if r.get("note") == "no_match")
    if missing:
        print(f"Warning: {missing} symbol(s) had no MyGene hit.", file=sys.stderr)
    return 0


if __name__ == "__main__":
    raise SystemExit(main())
```
