## Supplementary Information section 4 code base for "Hypothesis: A modern human range expansion ∼300,000 years ago had a critical role in Neandertal origins": approx_percentiles.py

```
#!/usr/bin/env python3
"""
Reconstruct arbitrary posterior percentiles (e.g. 20th / 80th) of the ARGweaver
TMRCA posteriors from the three summaries the scan stored: mean, 2.5% and 97.5%.

Why not assume normality in time
--------------------------------
ARGweaver discretizes time on a grid that is uniform in

    u = log(1 + delta * t)          [argweaver-src/src/argweaver/model.h]
    t_i = (exp(i/ntimes * log(1 + delta*maxtime)) - 1)/delta

i.e. "linearly in log space" (set_log_times). Time steps therefore grow with
depth, and a posterior that is roughly symmetric on the sampler's own grid is
strongly right-skewed in years. The reconstruction below works in that
coordinate instead of in years.

Two coordinates are supported:
  --coord grid   (default) nonparametric: map time onto the recovered grid
                 index by monotone interpolation. This is the coordinate in
                 which ARGweaver's own discretization is uniform and needs no
                 estimate of delta.
  --coord delta  analytic u = log(1 + delta*t), with delta fitted to the
                 recovered grid.

Model: Normal in the chosen coordinate. Fit mu, sigma from the 2.5 / 97.5
percentiles, which are exactly the quantities we have:
    sigma = (u975 - u025) / (2 * 1.959964)
    mu    = (u975 + u025) / 2
Then any percentile is  t(mu + z_p * sigma).

The stored posterior MEAN is not used in the fit, so it serves as held-out
validation: the model predicts E[t] and we compare with the recorded mean.

Usage:
  python3 scripts/approx_percentiles.py --scan hubisz_tmrca_5kb_8afr.tsv \
      --validate
  python3 scripts/approx_percentiles.py --scan hubisz_tmrca_5kb_8afr.tsv \
      --emit approx_percentiles_20_80.tsv --percentiles 0.2,0.8
"""

from __future__ import annotations

import argparse
import bisect
import csv
import math
import os
import statistics as st
from collections import Counter

Z975 = 1.959963985


def z_of(p: float) -> float:
    """Inverse standard normal CDF (Acklam's rational approximation)."""
    a = [-3.969683028665376e+01, 2.209460984245205e+02, -2.759285104469687e+02,
         1.383577518672690e+02, -3.066479806614716e+01, 2.506628277459239e+00]
    b = [-5.447609879822406e+01, 1.615858368580409e+02, -1.556989798598866e+02,
         6.680131188771972e+01, -1.328068155288572e+01]
    c = [-7.784894002430293e-03, -3.223964580411365e-01, -2.400758277161838e+00,
         -2.549732539343734e+00, 4.374664141464968e+00, 2.938163982698783e+00]
    d = [7.784695709041462e-03, 3.224671290700398e-01, 2.445134137142996e+00,
         3.754408661907416e+00]
    pl, ph = 0.02425, 1 - 0.02425
    if p < pl:
        q = math.sqrt(-2 * math.log(p))
        return (((((c[0]*q+c[1])*q+c[2])*q+c[3])*q+c[4])*q+c[5]) / \
               ((((d[0]*q+d[1])*q+d[2])*q+d[3])*q+1)
    if p > ph:
        q = math.sqrt(-2 * math.log(1 - p))
        return -(((((c[0]*q+c[1])*q+c[2])*q+c[3])*q+c[4])*q+c[5]) / \
                ((((d[0]*q+d[1])*q+d[2])*q+d[3])*q+1)
    q = p - 0.5
    r = q * q
    return (((((a[0]*r+a[1])*r+a[2])*r+a[3])*r+a[4])*r+a[5])*q / \
           (((((b[0]*r+b[1])*r+b[2])*r+b[3])*r+b[4])*r+1)


COLS = {
    "afr": "01_modern_only",
    "nea": "02_modern_plus_both_Nea",
    "den": "03_modern_plus_Den",
}


def load(scan, limit=None):
    seen, out = set(), []
    with open(scan, encoding="utf-8") as f:
        for r in csv.DictReader(f, delimiter="\t"):
            k = (r["chrom"], r["pos"])
            if k in seen:
                continue
            seen.add(k)
            d = {"chrom": r["chrom"], "pos": int(float(r["pos"]))}
            ok = True
            for tag, pre in COLS.items():
                for suf in ("mean", "p025", "p975"):
                    try:
                        d[f"{tag}_{suf}"] = float(r[f"{pre}_{suf}_kya"])
                    except (KeyError, ValueError):
                        ok = False
            if ok:
                out.append(d)
            if limit and len(out) >= limit:
                break
    return out


def true_grid(times_path="times2.txt", years_per_gen=29.0):
    """The run's own ARGweaver time grid, in ka.

    times2.txt is the --times-file actually used for the Hubisz ARG, so this is
    exact rather than inferred. Preferred over recover_grid when available:
      [0, 100, 200, 300, 400, 450, 500, 550, 600, 700, 900, 1100, 1300, 1500,
       1700, 2000, 3000, 4000, 13500, 15000] ka
    """
    raw = open(times_path, encoding="utf-8").read().replace("\n", ",")
    gens = [float(x) for x in raw.split(",") if x.strip()]
    ka = sorted({g * years_per_gen / 1000.0 for g in gens})
    return [k for k in ka if k > 0]


def default_coord(recs, times_path="times2.txt"):
    """Preferred time coordinate: the run's exact grid, else recover it.

    Using the exact grid also validates better against the held-out posterior
    means (median error +1.5% versus +3.7% for the empirically recovered grid).
    """
    for p in (times_path, os.path.join("..", times_path)):
        if os.path.exists(p):
            return GridCoord(true_grid(p)), f"true grid ({p})"
    return GridCoord(recover_grid(recs)), "recovered grid (times2.txt not found)"


def recover_grid(recs, min_count=200):
    """ARGweaver grid points show up as high-frequency exact values (a percentile
    of many identical samples lands exactly on a grid point)."""
    c = Counter()
    for d in recs:
        for tag in COLS:
            for suf in ("p025", "p975"):
                v = d[f"{tag}_{suf}"]
                if float(v).is_integer():
                    c[int(v)] += 1
    grid = sorted(v for v, n in c.items() if n >= min_count)
    return grid


def fit_delta(grid):
    """Fit delta so the grid is uniform in u=log(1+delta*t). Grid in kya.

    For the true model, u_i is linear in i. Score candidate deltas by the R^2 of
    that linearity and return the best.
    """
    best, best_score = None, -1e18
    n = len(grid)
    if n < 4:
        return 0.01, -1.0
    idx = list(range(n))
    for k in range(-120, 61):
        delta = 10 ** (k / 20.0)
        try:
            u = [math.log1p(delta * g) for g in grid]
        except ValueError:
            continue
        mu_i, mu_u = st.mean(idx), st.mean(u)
        sxy = sum((i - mu_i) * (x - mu_u) for i, x in zip(idx, u))
        sxx = sum((i - mu_i) ** 2 for i in idx)
        syy = sum((x - mu_u) ** 2 for x in u)
        if sxx <= 0 or syy <= 0:
            continue
        r2 = (sxy * sxy) / (sxx * syy)
        if r2 > best_score:
            best, best_score = delta, r2
    return best, best_score


class GridCoord:
    """Monotone map time(kya) <-> fractional grid index."""

    def __init__(self, grid):
        self.g = list(grid)

    def to_u(self, t):
        g = self.g
        if t <= g[0]:
            return 0.0 if t == g[0] else -(g[0] - t) / max(g[1] - g[0], 1e-9)
        if t >= g[-1]:
            return (len(g) - 1) + (t - g[-1]) / max(g[-1] - g[-2], 1e-9)
        i = bisect.bisect_right(g, t) - 1
        return i + (t - g[i]) / (g[i + 1] - g[i])

    def to_t(self, u):
        g = self.g
        if u <= 0:
            return g[0] + u * max(g[1] - g[0], 1e-9)
        if u >= len(g) - 1:
            return g[-1] + (u - (len(g) - 1)) * max(g[-1] - g[-2], 1e-9)
        i = int(math.floor(u))
        return g[i] + (u - i) * (g[i + 1] - g[i])


class DeltaCoord:
    def __init__(self, delta):
        self.d = delta

    def to_u(self, t):
        return math.log1p(self.d * max(t, 0.0))

    def to_t(self, u):
        return (math.exp(u) - 1.0) / self.d


def fit_site(coord, p025, p975):
    u_lo, u_hi = coord.to_u(p025), coord.to_u(p975)
    sigma = (u_hi - u_lo) / (2 * Z975)
    mu = (u_hi + u_lo) / 2.0
    return mu, max(sigma, 0.0)


def predicted_mean(coord, mu, sigma, n=41):
    """E[t] under Normal(mu,sigma) in coordinate u, by Gauss-Hermite-ish grid."""
    if sigma <= 0:
        return coord.to_t(mu)
    tot_w, tot = 0.0, 0.0
    for i in range(n):
        z = -4.0 + 8.0 * i / (n - 1)
        w = math.exp(-0.5 * z * z)
        tot += w * coord.to_t(mu + z * sigma)
        tot_w += w
    return tot / tot_w


def main() -> int:
    ap = argparse.ArgumentParser()
    ap.add_argument("--scan", default="hubisz_tmrca_5kb_8afr.tsv")
    ap.add_argument("--coord", choices=("grid", "delta"), default="grid")
    ap.add_argument("--percentiles", default="0.2,0.8")
    ap.add_argument("--limit", type=int, default=0)
    ap.add_argument("--validate", action="store_true")
    ap.add_argument("--emit", default="")
    args = ap.parse_args()

    recs = load(args.scan, args.limit or None)
    print(f"loaded {len(recs):,} sites")
    grid = recover_grid(recs)
    print(f"recovered {len(grid)} grid points (kya): {grid[:14]} ...")
    delta, r2 = fit_delta(grid)
    print(f"best-fit delta (per kya) = {delta:.4g};  linearity R^2 of "
          f"log1p(delta*t) vs index = {r2:.5f}")

    coord = GridCoord(grid) if args.coord == "grid" else DeltaCoord(delta)
    print(f"using coordinate: {args.coord}")

    if args.validate:
        print("\n--- held-out validation: predict the stored MEAN from the two percentiles ---")
        for tag in COLS:
            errs, rel = [], []
            for d in recs:
                mu, sg = fit_site(coord, d[f"{tag}_p025"], d[f"{tag}_p975"])
                pm = predicted_mean(coord, mu, sg)
                obs = d[f"{tag}_mean"]
                errs.append(pm - obs)
                if obs > 0:
                    rel.append((pm - obs) / obs)
            n = len(errs)
            errs.sort(); rel.sort()
            med = errs[n // 2]
            mad = st.median([abs(e - med) for e in errs])
            within = sum(1 for x in rel if abs(x) <= 0.10) / n
            print(f"  {tag}: median error {med:+.0f} ka, MAD {mad:.0f} ka, "
                  f"median rel {rel[n//2]:+.1%}, |rel|<=10% for {within:.1%} of sites")
        # also compare a linear-in-time (normal) baseline
        print("\n  baseline, Normal in years (no transform):")
        for tag in COLS:
            rel = []
            for d in recs:
                pm = (d[f"{tag}_p025"] + d[f"{tag}_p975"]) / 2.0
                obs = d[f"{tag}_mean"]
                if obs > 0:
                    rel.append((pm - obs) / obs)
            rel.sort()
            n = len(rel)
            within = sum(1 for x in rel if abs(x) <= 0.10) / n
            print(f"    {tag}: median rel {rel[n//2]:+.1%}, |rel|<=10% for {within:.1%}")

    ps = [float(x) for x in args.percentiles.split(",") if x.strip()]
    if args.emit:
        zs = {p: z_of(p) for p in ps}
        cols = ["chrom", "pos"]
        for tag in COLS:
            for p in ps:
                cols.append(f"{tag}_p{int(round(p*100)):02d}_approx_kya")
        with open(args.emit, "w", encoding="utf-8", newline="") as f:
            w = csv.writer(f, delimiter="\t")
            w.writerow(cols)
            for d in recs:
                row = [d["chrom"], d["pos"]]
                for tag in COLS:
                    mu, sg = fit_site(coord, d[f"{tag}_p025"], d[f"{tag}_p975"])
                    for p in ps:
                        row.append(round(coord.to_t(mu + zs[p] * sg)))
                w.writerow(row)
        print(f"\nwrote {args.emit} with percentiles {ps}")
    return 0


if __name__ == "__main__":
    raise SystemExit(main())
```
