## Supplementary Information section 4 code base for "Hypothesis: A modern human range expansion ∼300,000 years ago had a critical role in Neandertal origins": archaic_gt_check_derived_pattern.py

```
#!/usr/bin/env python3
"""
For each candidate site, query archaic VCFs with bcftools and test:

  Neandertal: homozygous for the *derived* allele (“fixed” in one diploid genome)
  Denisovan: homozygous for the *ancestral* allele

Also reports looser flags (Neandertal carries derived at least once).

Input TSV must include header matching output of ensembl_vep_grch37_derived_missense.py:
  chrom, pos, ref, alt, ... , ancestral, derived_allele, ...

Requires: bcftools on PATH; archaic VCFs tabix-indexed. Chrom names must match
the VCF (often 1 vs chr1 — use --region-chr-prefix chr when needed).

Example:
  python3 archaic_gt_check_derived_pattern.py hits.tsv \\
    --nea-vcf nea.vcf.gz --nea-sample Vindija33.25 \\
    --den-vcf den.vcf.gz --den-sample Denisova3 \\
    -o hits_with_archaic.tsv
"""

from __future__ import annotations

import argparse
import csv
import re
import subprocess
import sys
from typing import Optional


def parse_gt_alleles(ref: str, alt_field: str, gt: str) -> Optional[list[str]]:
    if not gt or "." in gt:
        return None
    ref = ref.upper()
    alts = [a.strip().upper() for a in alt_field.split(",") if a.strip()]
    alleles = [ref] + alts
    g = re.sub(r"\|", "/", gt)
    parts = g.split("/")
    out: list[str] = []
    try:
        for p in parts:
            idx = int(p)
            if idx < 0 or idx >= len(alleles):
                return None
            out.append(alleles[idx])
    except ValueError:
        return None
    return out


def bcftools_query_gt(
    vcf: str,
    sample: str,
    chrom: str,
    pos: int,
    timeout: float,
) -> tuple[str, str, str, str]:
    region = f"{chrom}:{pos}-{pos}"
    cmd = [
        "bcftools",
        "query",
        "-f",
        "%CHROM\t%POS\t%REF\t%ALT[\t%GT]\n",
        "-s",
        sample,
        "-r",
        region,
        vcf,
    ]
    p = subprocess.run(
        cmd,
        check=False,
        capture_output=True,
        text=True,
        timeout=timeout,
    )
    if p.returncode != 0:
        return "", "", "", ""
    line = (p.stdout or "").strip().split("\n")
    if not line or not line[0].strip():
        return "", "", "", ""
    cols = line[0].rstrip("\n").split("\t")
    if len(cols) < 5:
        return "", "", "", ""
    c, po, r, a, gt = cols[0], cols[1], cols[2], cols[3], cols[4]
    return c, po, r, a, gt


def carries_allele(alleles: Optional[list[str]], want: str) -> bool:
    if not alleles:
        return False
    w = want.upper()
    return any(a.upper() == w for a in alleles)


def homozygous_for(alleles: Optional[list[str]], want: str) -> bool:
    if not alleles or len(alleles) < 2:
        return False
    w = want.upper()
    return all(a.upper() == w for a in alleles)


def main() -> int:
    ap = argparse.ArgumentParser(description="Archaic bcftools genotype check for candidate sites.")
    ap.add_argument("hits_tsv", help="VEP output TSV with header")
    ap.add_argument("--nea-vcf", required=True)
    ap.add_argument("--nea-sample", required=True)
    ap.add_argument("--den-vcf", required=True)
    ap.add_argument("--den-sample", required=True)
    ap.add_argument("-o", "--output", required=True)
    ap.add_argument(
        "--region-chr-prefix",
        default="",
        help='If archaic VCFs use chr1-style contigs, pass "chr" to query chr1:pos-pos.',
    )
    ap.add_argument("--timeout", type=float, default=30.0)
    args = ap.parse_args()

    with open(args.hits_tsv, encoding="utf-8", newline="") as f:
        r = csv.DictReader(f, delimiter="\t")
        rows = list(r)

    if not rows:
        print("No rows in input.", file=sys.stderr)
        return 1
    fieldnames = list(rows[0].keys())
    extra = [
        "nea_gt",
        "nea_ref_obs",
        "nea_alt_obs",
        "den_gt",
        "den_ref_obs",
        "den_alt_obs",
        "nea_carries_derived",
        "nea_homozygous_derived",
        "den_homozygous_ancestral",
        "pattern_nea_hom_der_den_hom_anc",
        "pattern_modern_like",
    ]
    out_fields = fieldnames + [k for k in extra if k not in fieldnames]

    out_rows: list[dict[str, str]] = []
    for row in rows:
        chrom = row.get("chrom", "").strip()
        pos_s = row.get("pos", "").strip()
        derived = row.get("derived_allele", "").strip()
        ancestral = row.get("ancestral", "").strip()
        if not chrom or not pos_s or not derived or not ancestral:
            continue
        pos = int(pos_s)
        rchrom = f"{args.region_chr_prefix}{chrom}" if args.region_chr_prefix else chrom

        nc, npo, nr, na, ngt = bcftools_query_gt(
            args.nea_vcf, args.nea_sample, rchrom, pos, args.timeout
        )
        dc, dpo, dr, da, dgt = bcftools_query_gt(
            args.den_vcf, args.den_sample, rchrom, pos, args.timeout
        )

        nea_als = parse_gt_alleles(nr, na, ngt) if nr and na else None
        den_als = parse_gt_alleles(dr, da, dgt) if dr and da else None

        nea_carries = carries_allele(nea_als, derived)
        nea_hom_der = homozygous_for(nea_als, derived)
        den_hom_anc = homozygous_for(den_als, ancestral)

        # Strict: Neandertal homozygous derived, Denisovan homozygous ancestral
        pattern_strict = nea_hom_der and den_hom_anc
        # Legacy loose name: at least one derived read in Nea + hom anc in Den
        pattern_loose = nea_carries and den_hom_anc

        row_out = dict(row)
        row_out["nea_gt"] = ngt
        row_out["nea_ref_obs"] = nr
        row_out["nea_alt_obs"] = na
        row_out["den_gt"] = dgt
        row_out["den_ref_obs"] = dr
        row_out["den_alt_obs"] = da
        row_out["nea_carries_derived"] = str(int(bool(nea_carries)))
        row_out["nea_homozygous_derived"] = str(int(bool(nea_hom_der)))
        row_out["den_homozygous_ancestral"] = str(int(bool(den_hom_anc)))
        row_out["pattern_nea_hom_der_den_hom_anc"] = str(int(bool(pattern_strict)))
        row_out["pattern_modern_like"] = str(int(bool(pattern_loose)))
        out_rows.append(row_out)

    with open(args.output, "w", encoding="utf-8", newline="") as f:
        w = csv.DictWriter(f, fieldnames=out_fields, delimiter="\t", extrasaction="ignore")
        w.writeheader()
        w.writerows(out_rows)

    n_strict = sum(1 for r in out_rows if r.get("pattern_nea_hom_der_den_hom_anc") == "1")
    n_loose = sum(1 for r in out_rows if r.get("pattern_modern_like") == "1")
    print(
        f"Wrote {args.output} ({len(out_rows)} rows; "
        f"nea_hom_der+den_hom_anc={n_strict}; loose_carry+den_hom_anc={n_loose})",
        file=sys.stderr,
    )
    return 0


if __name__ == "__main__":
    raise SystemExit(main())
```
