## Supplementary Information section 4 code base for "Hypothesis: A modern human range expansion ∼300,000 years ago had a critical role in Neandertal origins": assemble_fd_annotation.py

```
#!/usr/bin/env python3
"""Collect the fixed-derived sites of the 1 kb screen with their annotation.

The screen is run per padded region, so a site inside two overlapping regions
is emitted twice, identical except for the cluster_start/cluster_end that say
which locus it was found under. That did not happen on the 5 kb grid, where no
two padded regions overlapped, so the earlier pipeline had no reason to handle
it; on the 1 kb grid six sites at the >99% tier and seven overall are affected.
Leaving the duplicates in would double-count them in Supplementary Table 3 and
would make a site appear under two loci in Table S4.2.

A duplicate is resolved to the stronger region: higher P(real) first, then a
catalog region over a non-catalog one, then the tighter region. Every case
here is a weak non-catalog region overlapping a catalog one, or two catalog
regions, so the rule is picking the locus the site most plausibly belongs to
rather than making a fine judgement. The choice only sets which locus the site
is reported under; the P(real) printed for the site itself is read at the base
by site_exact_preal.py and does not depend on it.

Annotation is joined in from whichever files carry it, the 5 kb run's output
included, because the 1 kb screen is a strict superset of the 5 kb one and
re-annotating sites that already have an identical annotation would only cost
time. Sites must not go missing in the join, so the script fails loudly rather
than emitting a short table.
"""

import argparse
import csv
import os
import sys
from collections import defaultdict


def read(fn):
    with open(fn, encoding="utf-8") as f:
        return list(csv.DictReader(f, delimiter="\t"))


def passing(fn):
    """Sites matching the fixed-derived pattern, keyed by position."""
    by_pos = defaultdict(list)
    for r in read(fn):
        if r.get("archaic_pattern") == "yes":
            by_pos[(r["chrom"], int(r["pos"]))].append(r)
    return by_pos


def region_index(fn):
    """(chrom, padded start, padded end) -> region record."""
    idx = {}
    for r in read(fn):
        idx[(r["chrom"], int(r["start"]), int(r["end"]))] = r
    return idx


def pick(cands, regions):
    """Strongest region for a site emitted more than once."""
    def key(r):
        reg = regions.get((r["chrom"], int(r["cluster_start"]),
                           int(r["cluster_end"])))
        if reg is None:
            return (-1.0, 0, 0)
        return (float(reg["set_preal_at_min_x"]),
                1 if reg["in_catalog"] == "yes" else 0,
                -int(reg["span_kb"]))
    return max(cands, key=key)


def main():
    ap = argparse.ArgumentParser()
    ap.add_argument("--af99", default="k1/fd/fd_k1_af99.all.tsv")
    ap.add_argument("--af95", default="k1/fd/fd_k1_af95.all.tsv")
    ap.add_argument("--regions", default="k1/screen_k1.padded.regions.tsv")
    ap.add_argument("--annot", nargs="+", default=[
        "tight/fd_af99.functional.canonical.tsv",
        "catalog_annotation/fd_af95.added135.functional.tsv",
        "k1/fd/fd_k1_af99.newsites.functional.tsv",
        "k1/fd/fd_k1_af95.addedonly.functional.tsv"])
    ap.add_argument("--out99", default="k1/fd/fd_k1_af99.functional.tsv")
    ap.add_argument("--out95", default="k1/fd/fd_k1_af95.added.functional.tsv")
    args = ap.parse_args()

    regions = region_index(args.regions)
    p99, p95 = passing(args.af99), passing(args.af95)

    dup99 = sum(1 for v in p99.values() if len(v) > 1)
    dup95 = sum(1 for v in p95.values() if len(v) > 1)
    print(f"af99 pass: {sum(len(v) for v in p99.values())} rows -> "
          f"{len(p99)} sites ({dup99} resolved)")
    print(f"af95 pass: {sum(len(v) for v in p95.values())} rows -> "
          f"{len(p95)} sites ({dup95} resolved)")

    keep99 = {k: (v[0] if len(v) == 1 else pick(v, regions))
              for k, v in p99.items()}
    keep95 = {k: (v[0] if len(v) == 1 else pick(v, regions))
              for k, v in p95.items()}
    added = {k: v for k, v in keep95.items() if k not in keep99}
    print(f"added by the >95% tier: {len(added)}")

    annot = {}
    for fn in args.annot:
        if not os.path.exists(fn):
            print(f"  (missing, skipped) {fn}")
            continue
        n = 0
        for r in read(fn):
            annot[(r["chrom"], int(r["pos"]))] = r
            n += 1
        print(f"  annotation from {os.path.basename(fn):<44} {n:>6}")

    def emit(keep, out):
        miss = [k for k in keep if k not in annot]
        if miss:
            sys.exit(f"ERROR: {len(miss)} sites have no annotation, "
                     f"e.g. {miss[:3]}")
        rows = []
        for k in sorted(keep, key=lambda x: (int(x[0]) if x[0].isdigit()
                                             else 99, x[1])):
            rec = dict(annot[k])
            # The annotation was produced per input file and carries whichever
            # cluster that file happened to hold; the resolved one wins.
            rec["cluster_start"] = keep[k]["cluster_start"]
            rec["cluster_end"] = keep[k]["cluster_end"]
            rows.append(rec)
        with open(out, "w", newline="", encoding="utf-8") as f:
            w = csv.DictWriter(f, fieldnames=list(rows[0]), delimiter="\t")
            w.writeheader()
            w.writerows(rows)
        q = sum(1 for r in rows if r.get("qualifying") == "yes")
        print(f"wrote {out}  {len(rows)} sites, {q} qualifying")

    emit(keep99, args.out99)
    emit(added, args.out95)


if __name__ == "__main__":
    sys.exit(main())
```
