## Supplementary Information section 4 code base for "Hypothesis: A modern human range expansion ∼300,000 years ago had a critical role in Neandertal origins": audit_argweaver_consistency.py

```
#!/usr/bin/env python3
"""Positive control: does the screen reproduce what the ARGweaver-D scan assumed?

The scan conditioned on 16 African + 4 Neandertal haplotypes carrying the
derived allele and 2 Denisovan haplotypes carrying the ancestral one. Its four
Neandertal haplotypes are Altai and Vindija 33.19; its two Denisovan haplotypes
are Denisova 3. Those three genomes are therefore NOT independent evidence in
the confirmation screen -- the scan already required this of them, so if the
screen disagrees about them the cause is a pipeline fault, not biology.

The genuinely new evidence is Chagyrskaya 8, Goyet Q56-1 and Denisova 25, plus
the jump from 16 to 5,024 modern haplotypes.

This reads the --audit output, which lists every candidate position including
the ones the pass rule rejects. Reading the passing sites alone proves nothing:
the rule requires unanimity, so a scan genome contradicting the pattern is
dropped before it can be seen.

Usage:
  python3 audit_argweaver_consistency.py out_audit/fixed_derived.audit.chr*.tsv
"""

from __future__ import annotations

import csv
import sys
from collections import Counter, defaultdict

SCAN_NEA = ["Altai", "Vindija33.19"]      # 4 haplotypes the scan conditioned on
SCAN_DEN = ["Denisova3"]                  # 2 haplotypes the scan conditioned on
NEW_NEA = ["Chagyrskaya8", "GoyetQ56-1"]
NEW_DEN = ["Denisova25"]
NO_EVIDENCE = ("no_data", "no_call", "low_qual")


def main(paths):
    rows = []
    for p in paths:
        try:
            with open(p, encoding="utf-8") as f:
                rows.extend(csv.DictReader(f, delimiter="\t"))
        except (OSError, csv.Error) as e:
            print(f"  skipped {p}: {e}", file=sys.stderr)
    if not rows:
        sys.exit("no audit rows found -- run the screen with --audit first")

    print(f"candidate positions examined: {len(rows)}")
    print(f"  pass the catalog rule:      "
          f"{sum(1 for r in rows if r['passes'] == 'yes')}")
    present = [g for g in SCAN_NEA + SCAN_DEN + NEW_NEA + NEW_DEN
               if f"{g}_state" in rows[0]]
    print(f"  genomes in this run:        {', '.join(present)}")

    print("\n=== HARD CHECK: genomes the scan already conditioned on ===")
    print("A violation here cannot be biology. Expected states are")
    print("derived_hom for Altai/Vindija and ancestral_hom for Denisova 3.\n")
    bad_total = 0
    for g, want in ([(g, "derived_hom") for g in SCAN_NEA]
                    + [(g, "ancestral_hom") for g in SCAN_DEN]):
        key = f"{g}_state"
        if key not in rows[0]:
            print(f"  {g:<14} not in this run")
            continue
        c = Counter(r[key] for r in rows)
        informative = sum(v for k, v in c.items() if k not in NO_EVIDENCE)
        agree = c.get(want, 0)
        contra = informative - agree
        bad_total += contra
        pct = 100.0 * agree / informative if informative else float("nan")
        print(f"  {g:<14} {agree}/{informative} = {pct:.1f}% {want}"
              + (f"   <-- {contra} CONTRADICT" if contra else "   ok"))
        for k, v in sorted(c.items(), key=lambda x: -x[1]):
            flag = "  <-- unexpected" if (k not in NO_EVIDENCE and k != want) else ""
            print(f"      {k:<15} {v}{flag}")

    print("\n=== modern allele frequency ===")
    print("The scan required the derived allele fixed in its 16 African")
    print("haplotypes. In 5,008 modern haplotypes it should be common; the")
    print("ancestral allele frequency should sit near zero.\n")
    afs = []
    for r in rows:
        try:
            afs.append(float(r["g1k_anc_af"]))
        except (KeyError, ValueError):
            pass
    if afs:
        afs.sort()
        n = len(afs)
        print(f"  ancestral-allele frequency over {n} candidates:")
        for q in (0.5, 0.9, 0.99, 1.0):
            print(f"    {int(q*100):>3}th percentile: {afs[min(int(q*n), n-1)]:.4f}")
        hi = sum(1 for a in afs if a > 0.05)
        print(f"  above 0.05: {hi} ({100.0*hi/n:.2f}%)")
        poly = Counter(r.get("g1k_polymorphic", "?") for r in rows)
        print(f"  polymorphic in 1000G: {dict(poly)}")

    print("\n=== NEW evidence: genomes the scan never saw ===")
    print("These may legitimately disagree -- that is information, not a bug.\n")
    for g, want in ([(g, "derived_hom") for g in NEW_NEA]
                    + [(g, "ancestral_hom") for g in NEW_DEN]):
        key = f"{g}_state"
        if key not in rows[0]:
            print(f"  {g:<14} not in this run")
            continue
        c = Counter(r[key] for r in rows)
        informative = sum(v for k, v in c.items() if k not in NO_EVIDENCE)
        agree = c.get(want, 0)
        pct = 100.0 * agree / informative if informative else float("nan")
        print(f"  {g:<14} {agree}/{informative} = {pct:.1f}% {want}")
        for k, v in sorted(c.items(), key=lambda x: -x[1]):
            print(f"      {k:<15} {v}")

    print("\n=== why candidates were rejected ===")
    reasons = Counter()
    for r in rows:
        if r["passes"] == "yes":
            continue
        for part in r["reason"].split(";"):
            reasons[part.split(":")[0]] += 1
    for k, v in reasons.most_common(12):
        print(f"  {k:<20} {v}")

    print("\n=== VERDICT ===")
    if bad_total == 0:
        print("  PASS: every scan genome behaves as the scan assumed.")
        print("  The screen is consistent with its input; differences from the")
        print("  ARGweaver catalog come from the added genomes and haplotypes,")
        print("  not from a bioinformatics fault.")
    else:
        print(f"  FAIL: {bad_total} scan-genome states contradict the scan.")
        print("  Investigate before using these counts. Likely causes, in the")
        print("  order worth checking: ancestral-allele polarity (is the FASTA")
        print("  being read on the right strand and build?), a coordinate")
        print("  off-by-one between the region file and the VCFs, the wrong")
        print("  sample column in a multi-sample VCF, or a genome build mismatch.")
    return 0 if bad_total == 0 else 1


if __name__ == "__main__":
    raise SystemExit(main(sys.argv[1:]))
```
