## Supplementary Information section 4 code base for "Hypothesis: A modern human range expansion ∼300,000 years ago had a critical role in Neandertal origins": bstat_incremental_value.py

```
#!/usr/bin/env python3
"""Does the B statistic buy a larger catalog at the same purity?

This replaces the wrong question with the right one. The earlier B analysis
asked whether omega0 varies across deciles of B -- whether background selection
confounds the null ratio. It does not (decile odds 0.985-1.039 against a pooled
1.011, no Cochran-Armitage trend). But omega0 is ~1 and is no longer a free
parameter of interest, so that test could only ever return "no harm done".

The question that matters for the catalog is whether B has incremental
discriminative value: at a fixed control/case ratio, does a rule that knows B
admit more grid points than one that does not? These come apart because
background selection shortens genealogies on the Neandertal and Denisovan sides
alike. A symmetric enrichment of shallow TMRCAs leaves the ratio untouched --
which is exactly what the decile test saw -- while still filling the catalog
with grid points that are shallow for reasons unrelated to a sweep. Flat odds
across deciles does not imply B is uninformative for the rule.

Three rules are compared, all with the Denisovan floor held at the catalog's
700 ka and all evaluated on held-out chromosomes:

  blind   one global threshold x
  shift   x_d = x + beta * (decile - 4.5); two parameters, so it can express
          "be stricter where background selection is strong" and little else
  greedy  a free threshold per decile, grown one step at a time by whichever
          decile offers the purest next increment. This is not a rule anyone
          would publish; it is an upper bound on what any B-aware rule of this
          shape can achieve, so if it fails to beat blind out of sample, the
          answer is settled rather than merely unproven.

Selection is on odd chromosomes and scoring on even. With ten deciles and
twenty thresholds the greedy has enough freedom to fit noise comfortably, and
an in-sample gain would prove nothing.
"""

from __future__ import annotations

import argparse
import csv
import os
import sys

import numpy as np

sys.path.insert(0, os.path.dirname(os.path.abspath(__file__)))
from approx_percentiles import default_coord, fit_site, load  # noqa: E402
from optimize_rule_by_size import NULL_RATIO  # noqa: E402

Y_FLOOR = 700.0


def load_scan():
    recs = load("hubisz_tmrca_5kb_8afr.tsv")
    coord, _ = default_coord(recs)
    ch, pos, nea, den = [], [], [], []
    for d in recs:
        c = d["chrom"].replace("chr", "")
        if c in ("X", "Y", "23", "24"):
            continue
        mn, _ = fit_site(coord, d["nea_p025"], d["nea_p975"])
        md, _ = fit_site(coord, d["den_p025"], d["den_p975"])
        ch.append(int(c))
        pos.append(int(d["pos"]))
        nea.append(coord.to_t(mn))
        den.append(coord.to_t(md))
    return (np.array(ch), np.array(pos), np.array(nea), np.array(den))


def load_b(path):
    out = {}
    with open(path, encoding="utf-8") as f:
        for r in csv.DictReader(f, delimiter="\t"):
            v = r.get("bstat", "NA")
            if v in ("NA", ""):
                continue
            # Partly-covered grid points carry a B averaged over less than the
            # full 5 kb, so their decile assignment is the least reliable part
            # of the input; requiring most of the window to be covered costs
            # few points and removes that noise from the comparison.
            try:
                if float(r.get("bstat_covered_frac", 1.0)) < 0.5:
                    continue
                out[(int(r["chrom"]), int(r["pos"]))] = float(v)
            except ValueError:
                continue
    return out


def ratio_of(case, ctrl):
    return (ctrl / case) if case else float("nan")


def preal(case, ctrl):
    r = ratio_of(case, ctrl)
    return max(0.0, 1.0 - r / NULL_RATIO) if case else float("nan")


def blind_trace(nea, den, xs):
    out = []
    for x in xs:
        case = int(((nea <= x) & (den > Y_FLOOR)).sum())
        ctrl = int(((den <= x) & (nea > Y_FLOOR)).sum())
        out.append((x, case, ctrl))
    return out


def eval_vector(nea, den, dec, thr):
    """Counts for a per-decile threshold vector."""
    x_of = thr[dec]
    case = int(((nea <= x_of) & (den > Y_FLOOR)).sum())
    ctrl = int(((den <= x_of) & (nea > Y_FLOOR)).sum())
    return case, ctrl


def main():
    ap = argparse.ArgumentParser()
    ap.add_argument("--bstat", default="catalog_annotation/bstat.CADD_bestfit.5kb.tsv")
    ap.add_argument("--x-grid", default="500:1300:10")
    ap.add_argument("--deciles", type=int, default=10)
    ap.add_argument("--sizes", default="150,300,600,1500,4000,10000",
                    help="held-out catalog sizes at which to compare purity")
    args = ap.parse_args()

    ch, pos, nea, den = load_scan()
    bmap = load_b(args.bstat)
    bv = np.array([bmap.get((c, p), np.nan) for c, p in zip(ch, pos)])
    keep = ~np.isnan(bv)
    print(f"scan grid points (autosomal): {len(nea):,}")
    print(f"with a usable B value:        {keep.sum():,} "
          f"({100 * keep.mean():.1f}%)")
    ch, nea, den, bv = ch[keep], nea[keep], den[keep], bv[keep]

    edges = np.quantile(bv, np.linspace(0, 1, args.deciles + 1)[1:-1])
    dec = np.searchsorted(edges, bv)
    print(f"B deciles from {bv.min():.3f} to {bv.max():.3f}; "
          f"sizes {np.bincount(dec, minlength=args.deciles)}")

    lo, hi, step = (float(v) for v in args.x_grid.split(":"))
    xs = np.arange(lo, hi + step, step)

    odd = ch % 2 == 1
    even = ~odd
    print(f"odd (fit): {odd.sum():,}   even (score): {even.sum():,}\n")

    # ---- blind, scored on even -------------------------------------------
    blind = []
    for x, _c, _k in blind_trace(nea[odd], den[odd], xs):
        ce = int(((nea[even] <= x) & (den[even] > Y_FLOOR)).sum())
        ke = int(((den[even] <= x) & (nea[even] > Y_FLOOR)).sum())
        blind.append((x, ce, ke))

    # ---- shift, fitted on odd, scored on even ----------------------------
    betas = np.arange(-40, 41, 5.0)
    shift = []
    for x in xs:
        for b in betas:
            thr = x + b * (np.arange(args.deciles) - (args.deciles - 1) / 2.0)
            co, ko = eval_vector(nea[odd], den[odd], dec[odd], thr)
            ce, ke = eval_vector(nea[even], den[even], dec[even], thr)
            shift.append((x, b, co, ko, ce, ke))

    # ---- water-filling per-decile, fitted on odd, scored on even ---------
    # The optimal nested per-decile rule at any size equalises MARGINAL purity
    # across deciles: pick a cutoff lambda, then advance each decile's
    # threshold while its next increment is at least that pure. Sweeping lambda
    # traces the whole frontier, and unlike a one-decile-at-a-time greedy it
    # contains the near-uniform solutions, so it cannot score worse than blind
    # by construction and is a genuine upper bound on rules of this shape.
    d_case = np.zeros((args.deciles, len(xs)), dtype=int)
    d_ctrl = np.zeros((args.deciles, len(xs)), dtype=int)
    for d in range(args.deciles):
        m = odd & (dec == d)
        nm, dm = nea[m], den[m]
        for i, x in enumerate(xs):
            d_case[d, i] = int(((nm <= x) & (dm > Y_FLOOR)).sum())
            d_ctrl[d, i] = int(((dm <= x) & (nm > Y_FLOOR)).sum())
    water = []
    for lam in np.unique(np.concatenate([np.arange(0, 1.01, 0.01), [1e9]])):
        ptr = np.zeros(args.deciles, dtype=int)
        for d in range(args.deciles):
            while ptr[d] + 1 < len(xs):
                i = ptr[d]
                dc = d_case[d, i + 1] - d_case[d, i]
                dk = d_ctrl[d, i + 1] - d_ctrl[d, i]
                if dc <= 0 or dk / dc <= lam:
                    ptr[d] += 1
                else:
                    break
        thr = xs[ptr]
        co, ko = eval_vector(nea[odd], den[odd], dec[odd], thr)
        ce, ke = eval_vector(nea[even], den[even], dec[even], thr)
        water.append((thr.copy(), co, ko, ce, ke))

    # ---- comparison at matched held-out size -----------------------------
    # Matched size rather than matched ratio: sizes are integers and can be
    # matched closely, whereas a coarse threshold grid steps over any target
    # ratio and manufactures spurious differences. The Wilson interval is what
    # decides the question -- at the catalog's own size there are only ~165
    # grid points per half genome, so this test can only detect a large gain,
    # which is why larger sizes are included where it has real power.
    from optimize_rule_by_size import wilson_hi, wilson_lo

    def nearest(cands, n):
        ok = [c for c in cands if c[0] > 0]
        return min(ok, key=lambda c: abs(c[0] - n)) if ok else None

    blind_pts = [(ce, ke) for _x, ce, ke in blind]
    shift_pts = [(ce, ke) for _x, _b, _co, _ko, ce, ke in shift]
    water_pts = [(ce, ke) for _t, _co, _ko, ce, ke in water]

    print(f"{'target':>7} | {'blind':>28} | {'shift (2 par)':>28} "
          f"| {'water-fill (10 par)':>28}")
    print(f"{'size':>7} | {'size':>6} {'ratio':>6} {'95% CI':>15} | "
          f"{'size':>6} {'ratio':>6} {'95% CI':>15} | "
          f"{'size':>6} {'ratio':>6} {'95% CI':>15}")
    for n in (int(v) for v in args.sizes.split(",")):
        cells = []
        for pts in (blind_pts, shift_pts, water_pts):
            r = nearest(pts, n)
            if not r:
                cells.append(f"{'-':>6} {'-':>6} {'-':>15}")
                continue
            c, k = r
            cells.append(f"{c:>6,} {ratio_of(c, k):>6.3f} "
                         f"[{wilson_lo(k, c):.3f},{wilson_hi(k, c):.3f}]".rjust(15))
        print(f"{n:>7,} | " + " | ".join(cells))

    print("\nall sizes and ratios are held-out (even-chromosome). A B-aware "
          "rule earns its keep only if its ratio is lower than blind's at "
          "matched size by more than the intervals allow.")
    return 0


if __name__ == "__main__":
    sys.exit(main())
```
