## Supplementary Information section 4 code base for "Hypothesis: A modern human range expansion ∼300,000 years ago had a critical role in Neandertal origins": bstat_to_grid.py

```
#!/usr/bin/env python3
"""
Project sellalab/Murphy-style B-maps onto the 5-kb ARGweaver grid.

The bmaps are run-length encoded, two whitespace-separated columns per line:

    B  segment_length_bp

Coordinates are implicit: segment k covers [cum, cum + length), where cum is
the sum of all preceding lengths. So the file has to be walked in order, and
summing the length column recovers the chromosome length -- which is also the
genome-build check (hg19 chr1 = 249,250,621).

B is stored scaled by 1000 in the files I have seen (values like 635 meaning
0.635), but McVicker-era maps and some rebuilds differ, so the scale is
auto-detected from the observed maximum unless --scale is given explicitly.
Getting this wrong silently would put B on the wrong footing in the
regression, so the resolved scale is always reported.

Output is one row per grid window with the length-weighted mean B over
[pos, pos + grid_step), plus the covered fraction so that windows that are
only partly spanned by the map can be filtered downstream.

Example:
  python3 bstat_to_grid.py --bmap chr1.bmap.txt --chrom 1 \
      --chrom-len 249250621 -o chr1.bstat.5kb.tsv
"""

from __future__ import annotations

import argparse
import gzip
import sys
from typing import Optional, TextIO

# hg19/hg18/hg38 chromosome-1 lengths, for build identification.
BUILD_BY_CHR1_LEN = {
    249250621: "hg19/GRCh37",
    247249719: "hg18/NCBI36",
    248956422: "hg38/GRCh38",
}


def _open_text(path: str) -> TextIO:
    if path.endswith(".gz"):
        return gzip.open(path, "rt", encoding="utf-8", errors="replace")
    return open(path, encoding="utf-8", errors="replace")


def main() -> None:
    ap = argparse.ArgumentParser(
        description=__doc__,
        formatter_class=argparse.RawDescriptionHelpFormatter)
    ap.add_argument("--bmap", required=True, help="run-length encoded bmap file")
    ap.add_argument("--chrom", required=True)
    ap.add_argument("--chrom-len", type=int, required=True,
                    help="expected chromosome length, for the build check")
    ap.add_argument("--grid-step", type=int, default=5000)
    ap.add_argument("--scale", type=float, default=None,
                    help="divide raw B by this (default: auto-detect 1 or 1000)")
    ap.add_argument("--column-name", default="bstat")
    ap.add_argument("-o", "--out", required=True)
    args = ap.parse_args()

    segs: list = []
    total = 0
    raw_max = 0.0
    with _open_text(args.bmap) as fh:
        for lineno, line in enumerate(fh, 1):
            line = line.strip()
            if not line or line.startswith("#"):
                continue
            f = line.split()
            if len(f) < 2:
                sys.stderr.write(f"ERROR: {args.bmap}:{lineno}: expected 2 "
                                 f"columns, got {len(f)}: {line!r}\n")
                sys.exit(1)
            try:
                b = float(f[0])
                ln = int(f[1])
            except ValueError:
                sys.stderr.write(f"ERROR: {args.bmap}:{lineno}: unparseable: "
                                 f"{line!r}\n")
                sys.exit(1)
            if ln <= 0:
                continue
            segs.append((total, total + ln, b))
            total += ln
            raw_max = max(raw_max, b)

    if not segs:
        sys.stderr.write(f"ERROR: no usable segments in {args.bmap}\n")
        sys.exit(1)

    build = BUILD_BY_CHR1_LEN.get(total)
    sys.stderr.write(f"segments={len(segs)}  total_bp={total:,}  raw_max_B={raw_max:g}\n")
    if str(args.chrom) in ("1", "chr1"):
        sys.stderr.write(f"build by chr1 length: {build or 'UNRECOGNISED'}\n")
    if total != args.chrom_len:
        sys.stderr.write(
            f"WARNING: total_bp {total:,} != --chrom-len {args.chrom_len:,} "
            f"(difference {total - args.chrom_len:+,}). Wrong build or a "
            f"partial map; not joining blindly.\n")

    if args.scale is not None:
        scale = args.scale
        how = "given"
    elif raw_max > 1.5:
        scale = 1000.0
        how = "auto (raw max > 1.5)"
    else:
        scale = 1.0
        how = "auto (raw max <= 1.5)"
    sys.stderr.write(f"B scale = {scale:g} [{how}] -> B range up to "
                     f"{raw_max / scale:.4f}\n")
    if raw_max / scale > 1.0001:
        sys.stderr.write(
            f"WARNING: scaled B exceeds 1 ({raw_max / scale:.4f}); the scale "
            f"is probably wrong.\n")

    step = args.grid_step
    n_win = (args.chrom_len + step - 1) // step
    wsum = [0.0] * n_win   # length-weighted sum of B
    wlen = [0] * n_win     # bp of this window covered by the map

    for s, e, b in segs:
        if e <= 0:
            continue
        first = s // step
        last = (e - 1) // step
        for w in range(first, min(last, n_win - 1) + 1):
            ws, we = w * step, (w + 1) * step
            ov = min(e, we) - max(s, ws)
            if ov > 0:
                wsum[w] += b * ov
                wlen[w] += ov

    col = args.column_name
    n_out = 0
    with open(args.out, "w") as out:
        out.write(f"chrom\tpos\t{col}\t{col}_covered_frac\n")
        for w in range(n_win):
            pos = w * step
            if wlen[w] == 0:
                out.write(f"{args.chrom}\t{pos}\tNA\t0.0000\n")
                continue
            b = (wsum[w] / wlen[w]) / scale
            out.write(f"{args.chrom}\t{pos}\t{b:.4f}\t{wlen[w] / step:.4f}\n")
            n_out += 1

    sys.stderr.write(f"Wrote {args.out}: {n_out:,}/{n_win:,} windows with B\n")


if __name__ == "__main__":
    main()
```
