## Supplementary Information section 4 code base for "Hypothesis: A modern human range expansion ∼300,000 years ago had a critical role in Neandertal origins": build_preal_grid.py

```
#!/usr/bin/env python3
"""Cache P(real) at every autosomal grid point, plus the two TMRCA means.

The locus figures need P(real) at ordinary grid points, not just at catalog
members, because the plotted window is defined by walking outward until the
signal dies (P(real) < 0.2). Recomputing the surface per figure would refit it
per figure, so it is fitted once here and cached.

The surface is exactly the one behind Table S4.1's P(real) column: binomial
logistic fitted to case and control counts, cross-validated by chromosome
parity so no grid point is scored by a model trained on its own chromosome,
and evaluated at the boundary of its support for the minority of points whose
TMRCAs fall outside the fitted region. That last step matters more here than
in the table: walking away from a locus quickly reaches shallow-Neandertal or
deep-Denisovan values outside the support, and clamping keeps the trace
monotone there instead of letting it wander.
"""

import argparse
import os
import sys

import numpy as np

sys.path.insert(0, os.path.dirname(os.path.abspath(__file__)))
from approx_percentiles import default_coord, fit_site, load, z_of  # noqa: E402
from fig_s51_model_curve import U_EDGES, V_EDGES, fit_on  # noqa: E402
from fit_fdr_surface import PCUTS, design  # noqa: E402
from make_loci_table import EPS, RULE_CUT, preal_from_theta  # noqa: E402


def main():
    ap = argparse.ArgumentParser()
    ap.add_argument("--scan", default="hubisz_tmrca_5kb_8afr.tsv")
    ap.add_argument("-o", "--out",
                    default="catalog_annotation/preal_gridpoints.tsv")
    a = ap.parse_args()

    recs = load(a.scan)
    coord, desc = default_coord(recs)
    recs = [d for d in recs
            if d["chrom"].replace("chr", "") not in ("X", "Y", "23", "24")]
    for d in recs:
        for tag in ("nea", "den"):
            mu, sg = fit_site(coord, d[f"{tag}_p025"], d[f"{tag}_p975"])
            for p in PCUTS:
                d[f"{tag}_hi{p}"] = coord.to_t(mu + z_of(p) * sg)
                d[f"{tag}_lo{p}"] = coord.to_t(mu + z_of(1 - p) * sg)
    sys.stderr.write(f"time coordinate: {desc}; {len(recs):,} grid points\n")

    odd = [d for d in recs if int(d["chrom"].replace("chr", "")) % 2 == 1]
    even = [d for d in recs if int(d["chrom"].replace("chr", "")) % 2 == 0]
    beta_for_even, n1 = fit_on(odd)
    beta_for_odd, n2 = fit_on(even)
    sys.stderr.write(f"surface fitted on {n1:,} odd and {n2:,} even cells\n")

    u_lo, u_hi = U_EDGES[0] + EPS, U_EDGES[-1]
    v_lo, v_hi = V_EDGES[0] + EPS, V_EDGES[-1]
    n_cl = 0
    with open(a.out, "w", encoding="utf-8") as f:
        f.write("chrom\tpos\tnea_mean_ka\tden_mean_ka\tafr_mean_ka\t"
                "preal\tclamped\n")
        for d in recs:
            c = int(d["chrom"].replace("chr", ""))
            u, v = d[f"nea_hi{RULE_CUT}"], d[f"den_lo{RULE_CUT}"]
            cu, cv = min(max(u, u_lo), u_hi), min(max(v, v_lo), v_hi)
            tag = ("u" if cu != u else "") + ("v" if cv != v else "")
            n_cl += bool(tag)
            beta = beta_for_odd if c % 2 == 1 else beta_for_even
            eta = design(np.array([cu / 100.0]), np.array([cv / 100.0]),
                         np.array([RULE_CUT]))[0] @ beta
            pr = preal_from_theta(1.0 / (1.0 + np.exp(-eta)))
            f.write(f"{c}\t{d['pos']}\t{d['nea_mean']:.0f}\t"
                    f"{d['den_mean']:.0f}\t{d['afr_mean']:.0f}\t"
                    f"{pr:.4f}\t{tag or '-'}\n")

    nea = np.array([d["nea_mean"] for d in recs])
    den = np.array([d["den_mean"] for d in recs])
    afr = np.array([d["afr_mean"] for d in recs])
    sys.stderr.write(
        f"wrote {a.out}\n"
        f"  {n_cl:,} of {len(recs):,} scored at a support boundary\n"
        f"  genome mean TAfr+Nea {nea.mean():.0f} ka (median {np.median(nea):.0f})\n"
        f"  genome mean TAfr+Den {den.mean():.0f} ka (median {np.median(den):.0f})\n"
        f"  genome mean TAfr     {afr.mean():.0f} ka (median {np.median(afr):.0f})\n")
    return 0


if __name__ == "__main__":
    sys.exit(main())
```
