## Supplementary Information section 4 code base for "Hypothesis: A modern human range expansion ∼300,000 years ago had a critical role in Neandertal origins": build_supp_table1.py

```
#!/usr/bin/env python3
"""Supplementary Table 2: the whole-genome 1-kb scan, with Relate percentiles.

The file is Supplementary Table 2 of the note and the scan behind it is on a
1 kb lattice; the script keeps its original name because the note's Software
section names it, and both the 5 kb and the 1 kb table are built by it -- the
grid is whatever --argweaver is pointed at.

Emits every grid point in the ARGweaver-D scan, carrying its ARGweaver-D dates
through unchanged -- no Peyregne recalibration, as requested -- and adding, per
grid point, where Relate's modern-only TMRCA at that point falls in the
genome-wide Relate distribution.

The percentile is the useful form rather than the raw Relate date because the
two methods do not share a timescale: ARGweaver-D's modern-only median is
1588 ka against Relate's 1499 ka, and the two disagree by more than that at
individual points. A percentile is invariant to any monotone rescaling, so it
can be compared with the ARGweaver columns without either having to be
recalibrated onto the other.

A caveat travels with the broader of the two Relate columns, and it is the
reason that column is named for its populations rather than for 1000 Genomes.
The released Relate ARG containing archaic genomes has 604 modern haplotypes
drawn from three populations only -- Yoruba, British and Han, 302 individuals
-- not the full 5,008 of phase 3. The full-panel run exists as tree sequences
on O2 but its TMRCA grids were never computed, so the broadest panel available
genome-wide is 604. Calling it a "1000 Genomes" panel invites the reader to
assume the whole phase 3 sample, so the columns are relate_ygc604_*.

Panel breadth turns out not to matter: the per-locus median percentile moves
from the 10th to the 8th between 216 and 604 haplotypes, and on chromosome 22,
where a 1,322-haplotype African grid does exist, concordance with ARGweaver is
0.444 against 0.472 for 216 haplotypes.
"""

import argparse
import csv
import glob
import gzip
import os
import sys

import numpy as np

sys.path.insert(0, os.path.dirname(os.path.abspath(__file__)))
from approx_percentiles import default_coord, fit_site, z_of  # noqa: E402


REL = {"yri216": "yri_01_modern_tmrca_years",
       "ygc604": "ygc_01_modern_tmrca_years"}


def rebound(fields, coord):
    """Rewrite the 2.5/97.5 bounds as 5/95, returning a per-row converter.

    ARGweaver-D reports the 2.5th and 97.5th percentiles and nothing else, so
    the 5th and 95th have to be reconstructed the same way every other derived
    percentile in this project is: map the two reported bounds onto the run's
    log-spaced time grid, fit a normal in grid-index coordinates, and read the
    new pair off the fit. Nothing is lost by the substitution. Two percentiles
    of a normal determine its mean and standard deviation, so the 5/95 pair
    carries exactly the information the 2.5/97.5 pair did and either can be
    recovered from the other.

    The point of doing it is consistency: Table S4.1 quotes 90% central
    intervals, because the 95th percentile of the modern-plus-Neandertal
    coalescence and the 5th of the modern-plus-Denisovan one are the two
    numbers the catalog rule actually turns on, and a reader checking a locus
    against the genome-wide file should find the same quantity there.
    """
    pairs = [(f[:-len("_p025_kya")], f) for f in fields
             if f.endswith("_p025_kya")]
    cache = {}

    def convert(row):
        for pre, lo_f in pairs:
            hi_f = f"{pre}_p975_kya"
            lo, hi = row.pop(lo_f), row.pop(hi_f)
            key = (lo, hi)
            if key not in cache:
                try:
                    mu, sg = fit_site(coord, float(lo), float(hi))
                    cache[key] = (f"{coord.to_t(mu + z_of(0.05) * sg):.0f}",
                                  f"{coord.to_t(mu + z_of(0.95) * sg):.0f}")
                except (TypeError, ValueError):
                    cache[key] = ("NA", "NA")
            row[f"{pre}_p05_kya"], row[f"{pre}_p95_kya"] = cache[key]
        return row

    out = []
    for f in fields:
        if f.endswith("_p025_kya"):
            out.append(f"{f[:-len('_p025_kya')]}_p05_kya")
        elif f.endswith("_p975_kya"):
            out.append(f"{f[:-len('_p975_kya')]}_p95_kya")
        else:
            out.append(f)
    return out, convert, cache


def load_relate(paths):
    """(chrom,pos) -> {panel: ka}, plus the genome-wide sorted values."""
    per, pools = {}, {k: [] for k in REL}
    for p in sorted(paths):
        with open(p, encoding="utf-8") as f:
            rd = csv.DictReader(f, delimiter="\t")
            for k, c in REL.items():
                if c not in rd.fieldnames:
                    sys.exit(f"{p}: no column {c}")
            for r in rd:
                c = r["chrom"].replace("chr", "")
                if c in ("X", "Y", "23", "24"):
                    continue
                rec = {}
                for k, col in REL.items():
                    v = r[col]
                    if v in ("", "NA"):
                        continue
                    rec[k] = float(v) / 1000.0
                    pools[k].append(rec[k])
                if rec:
                    per[(int(c), int(r["pos"]))] = rec
    ref = {k: np.sort(np.array(v)) for k, v in pools.items()}
    sys.stderr.write(f"Relate: {len(per):,} grid points; "
                     + ", ".join(f"{k} n={len(v):,}" for k, v in ref.items())
                     + "\n")
    return per, ref


def main():
    ap = argparse.ArgumentParser()
    ap.add_argument("--argweaver", default="hubisz_tmrca_5kb_8afr.tsv")
    ap.add_argument("--relate-glob", default="relate_archaic_1kb/chr*.tsv")
    ap.add_argument("--preal",
                    default="catalog_annotation/preal_gridpoints.tsv")
    ap.add_argument("--catalog-sites",
                    default="catalog_annotation/"
                            "catalog_final_median_x625_y700_auto.sites.tsv")
    ap.add_argument("-o", "--out",
                    default="catalog_annotation/"
                            "supplementary_table1_scan_5kb.tsv.gz")
    ap.add_argument("--bounds", choices=("2.5-97.5", "5-95"),
                    default="5-95",
                    help="percentile bounds to print for each haplotype set. "
                         "2.5-97.5 passes through what ARGweaver-D reported; "
                         "5-95 reconstructs the 90%% central interval used by "
                         "Table S4.1. The two are interchangeable, since "
                         "either pair determines the fitted normal.")
    args = ap.parse_args()

    rel, ref = load_relate(sorted(glob.glob(args.relate_glob)))

    cat = set()
    with open(args.catalog_sites, encoding="utf-8") as f:
        for r in csv.DictReader(f, delimiter="\t"):
            cat.add((int(r["chrom"].replace("chr", "")), int(r["pos"])))
    sys.stderr.write(f"catalog: {len(cat):,} grid points\n")

    preal = {}
    with open(args.preal, encoding="utf-8") as f:
        for r in csv.DictReader(f, delimiter="\t"):
            preal[(int(r["chrom"].replace("chr", "")), int(r["pos"]))] = (
                r["preal"], r["clamped"])
    sys.stderr.write(f"P(real): {len(preal):,} grid points\n")

    n_out = n_rel = 0
    with open(args.argweaver, encoding="utf-8") as fi, \
            gzip.open(args.out, "wt", newline="", encoding="utf-8") as fo:
        rd = csv.DictReader(fi, delimiter="\t")
        extra = ["preal", "preal_extrapolated", "in_catalog",
                 "relate_yri216_tmrca_ka", "relate_yri216_percentile",
                 "relate_ygc604_tmrca_ka", "relate_ygc604_percentile"]
        if args.bounds == "5-95":
            coord, cdesc = default_coord([])
            sys.stderr.write(f"bounds: 5/95 reconstructed on {cdesc}\n")
            names, convert, cache = rebound(list(rd.fieldnames), coord)
        else:
            names, convert, cache = list(rd.fieldnames), lambda r: r, {}
        w = csv.DictWriter(fo, delimiter="\t",
                           fieldnames=names + extra)
        w.writeheader()
        seen = set()
        for r in rd:
            c = r["chrom"].replace("chr", "")
            if c in ("X", "Y", "23", "24"):
                continue          # autosomes only, as described in the note
            key = (int(c), int(r["pos"]))
            if key in seen:
                continue          # the collated scan overlaps at chunk edges;
            seen.add(key)         # keep the first, as the analysis did
            out = convert(dict(r))
            pr = preal.get(key)
            out["preal"] = pr[0] if pr else "NA"
            out["preal_extrapolated"] = ("yes" if pr and pr[1] == "c"
                                         else "no" if pr else "NA")
            out["in_catalog"] = "yes" if key in cat else "no"
            got = rel.get(key, {}) if key else {}
            for k in ("yri216", "ygc604"):
                if k in got:
                    v = got[k]
                    pct = np.searchsorted(ref[k], v, "right") / len(ref[k])
                    out[f"relate_{k}_tmrca_ka"] = f"{v:.0f}"
                    out[f"relate_{k}_percentile"] = f"{100 * pct:.1f}"
                else:
                    out[f"relate_{k}_tmrca_ka"] = "NA"
                    out[f"relate_{k}_percentile"] = "NA"
            if got:
                n_rel += 1
            w.writerow(out)
            n_out += 1

    sys.stderr.write(f"wrote {args.out}\n  {n_out:,} grid points, "
                     f"{n_rel:,} with Relate values "
                     f"({100 * n_rel / n_out:.1f}%)\n")
    if cache:
        # ARGweaver-D discretizes time, so the reported bounds take only a few
        # thousand distinct values and the fit is done once per distinct pair.
        sys.stderr.write(f"  bounds rewritten as 5/95 from "
                         f"{len(cache):,} distinct percentile pairs\n")
    return 0


if __name__ == "__main__":
    sys.exit(main())
```
