## Supplementary Information section 4 code base for "Hypothesis: A modern human range expansion ∼300,000 years ago had a critical role in Neandertal origins": build_supp_table2.py

```
#!/usr/bin/env python3
"""Supplementary Table 2: the fixed-derived sites, with haplotype counts.

Reports, per site, how many haplotypes carry the derived allele out of how many
were actually observed, separately for 1000 Genomes moderns, Neandertals and
Denisovans. Writing it as derived/total rather than as a frequency is the point:
the archaic denominators are small and vary site to site as coverage comes and
goes, and a bare frequency would hide whether "all Neandertals derived" rested
on five genomes or one.

The panel is five high-coverage Neandertals (Altai, Vindija 33.19, Chagyrskaya
8, Goyet Q56-1, Denisova 17) and two high-coverage Denisovans (Denisova 3,
Denisova 25), so the denominators cap at 10 and 4. Per-genome state columns are
carried through unchanged so a reader can see exactly which genome supplied
which call, and the two callers (GATK-era and snpAD) are reported separately
because they mask and filter differently.

The functional columns (classes, phyloP, CADD, and the "qualifying" flag that
selects the Table S4.2 rows) come from annotate_functional_classes.py.

TWO FREQUENCY TIERS
  The screen proper is the 801 sites at >99% derived in 1000 Genomes. The note
  also reports what happens when that threshold is relaxed to >95%, which adds
  135 sites, and it prints one of them -- MCF2L2 Asn593Ser -- in Table S4.2 as
  an exception. Those 135 were originally left out of this file, so a reader
  following the note to Table S4.2 could not find the MCF2L2 row anywhere in
  the release. They are now included and separated by modern_freq_tier, which
  also makes the "adds 135 sites" claim checkable.

TWO P(REAL) COLUMNS, WHICH DO NOT AGREE AND SHOULD NOT
  catalog_locus_preal is the value of the catalog locus containing the site,
  from the 5 kb scan. preal_site is read at the substitution itself from the
  local ARGweaver-D tree, which is what Table S4.2 prints. They diverge where a
  site sits near a locus edge: chr2:162,101,812 is 0.09 on the nearest grid
  point and 0.73 at the base, and chr14:58,862,763 runs the other way, 0.79 on
  the grid and 0.00 at the base. Carrying only the locus value, as this file
  used to, left the numbers in Table S4.2 unreproducible from the release.

TWO FLAGS, BECAUSE THE RULE AND THE PRINTED TABLE ARE NOT THE SAME SET
  meets_table_s5_2_rule applies qualifies() mechanically to every row in both
  tiers. in_table_s5_2 records what the note actually prints. They differ at
  one row: chr3:182,990,363, a 3' UTR base in a conserved element from the >95%
  tier, meets the rule but is not in the table. Keeping both columns makes that
  visible instead of quietly resolving it in either direction.
"""

import argparse
import csv
import os
import sys


G1K_HAPS = 5008  # 2,504 individuals, phase 3

# The Table S4.2 rule is imported, not restated. It had been mirrored here and
# the copy drifted twice over -- ENCODE tracks crept into PAIR_A and the
# synonymous partner was missing from PAIR_B -- so this file flagged 24 sites
# where the table in the note has 13.
sys.path.insert(0, os.path.dirname(os.path.abspath(__file__)))
from approx_percentiles import default_coord, fit_site, z_of  # noqa: E402
from make_functional_table import qualifies  # noqa: E402

NEANDERTALS = ["Altai", "Vindija33.19", "Chagyrskaya8", "GoyetQ56-1",
               "Denisova17"]
DENISOVANS = ["Denisova3", "Denisova25"]


def state_counts(state):
    """(derived, observed) haplotypes from the screen's per-genome label."""
    if state == "derived_hom":
        return 2, 2
    if state == "ancestral_hom":
        return 0, 2
    if state == "het":
        return 1, 2
    return 0, 0  # no_data, no_call, low_qual: absence of evidence


def load_preal(path):
    """(chrom, start, end) -> P(real), from the catalog locus table."""
    out = []
    with open(path, encoding="utf-8") as f:
        for r in csv.DictReader(f, delimiter="\t"):
            iv = r["interval (Mb)"]
            a = float(iv.split("-")[0])
            b = float(iv.split("-")[1]) if "-" in iv else a
            out.append((r["chr"], a * 1e6, b * 1e6, r["P (real)"],
                        r["genes within 5 kb"]))
    return out


def preal_at(loci, chrom, pos, pad=5000):
    for c, a, b, p, g in loci:
        if c == chrom and a - pad <= pos <= b + pad:
            return p, g
    return "NA", ""


def load_tmrca_reads(path):
    """(chrom, pos) -> the resolved TMRCA read, for the 95% interval bounds.

    site_exact_preal.py carries only posterior means, but a mean is not enough
    to judge a site: a substitution whose Neandertal coalescence is placed at
    674 ka with an interval of 600-1300 ka is telling the reader something very
    different from one placed at 582 ka with 300-700 ka, even though both sit
    near the 625 ka threshold. Emitting the bounds lets the wide ones be
    discounted rather than read at face value.
    """
    if not os.path.exists(path):
        sys.stderr.write(f"note: {path} absent, interval columns will be NA\n")
        return {}
    out = {}
    with open(path, encoding="utf-8") as f:
        for r in csv.DictReader(f, delimiter="\t"):
            key = (r["chrom"].replace("chr", ""), str(int(float(r["pos"]))))
            out[key] = r
    sys.stderr.write(f"TMRCA reads with intervals: {len(out):,} sites\n")
    return out


def ci90(read, tag, coord):
    """'600-1300', the 5th to 95th percentile of a resolved read, or 'NA'.

    ARGweaver-D reports only the 2.5th and 97.5th percentiles, so the pair
    printed here is reconstructed from them through the same log-grid normal
    fit the rest of the project uses. The 90% interval rather than the 95% is
    what Table S4.1 and Supplementary Table 2 quote, because the 95th
    percentile of the Neandertal-inclusive coalescence and the 5th of the
    Denisovan-inclusive one are the two bounds the catalog rule turns on. A
    reader comparing a site here against its locus in Table S4.1 should not
    have to convert between two different interval widths.
    """
    if not read:
        return "NA"
    pre = {"afr": "01_modern_only", "nea": "02_modern_plus_both_Nea",
           "den": "03_modern_plus_Den"}[tag]
    try:
        lo = float(read[f"{pre}_p025_kya"])
        hi = float(read[f"{pre}_p975_kya"])
    except (KeyError, ValueError):
        return "NA"
    mu, sg = fit_site(coord, lo, hi)
    return (f"{coord.to_t(mu + z_of(0.05) * sg):.0f}-"
            f"{coord.to_t(mu + z_of(0.95) * sg):.0f}")


def load_site_exact(path):
    """(chrom, pos) -> the site-exact read, or {} if the file is absent.

    site_exact_preal.py evaluates the fitted surface at the base itself using
    the local tree, rather than at the nearest point of the 5 kb lattice. These
    are the values Table S4.2 prints.
    """
    if not os.path.exists(path):
        sys.stderr.write(f"note: {path} absent, site-exact columns will be NA\n")
        return {}
    out = {}
    with open(path, encoding="utf-8") as f:
        for r in csv.DictReader(f, delimiter="\t"):
            out[(r["chrom"].replace("chr", ""), r["pos"])] = r
    sys.stderr.write(f"site-exact P(real): {len(out):,} sites\n")
    return out


# Rows the note prints in Table S4.2. The >99% tier is selected by qualifies(),
# but MCF2L2 is there by an explicit authorial exception and chr3:182,990,363
# meets the rule without being printed, so neither can be derived from the rule
# alone. Listing the exceptions by position keeps this file honest about which
# is which rather than tuning qualifies() until the counts happen to match.
TABLE_S52_EXTRA = {("3", "182994744")}      # printed, though not >99%
TABLE_S52_OMIT = {("3", "182990363")}       # meets the rule, not printed


def main():
    ap = argparse.ArgumentParser()
    ap.add_argument("--sites",
                    default="catalog_annotation/fd_af99.functional.canonical.tsv")
    ap.add_argument("--sites95",
                    default="catalog_annotation/"
                            "fd_af95.added135.functional.tsv",
                    help="the sites the >95%% relaxation adds, annotated by "
                         "the same pipeline; carried at modern_freq_tier=95")
    ap.add_argument("--site-exact",
                    default="catalog_annotation/site_exact_preal.tsv")
    ap.add_argument("--tmrca-reads",
                    default="catalog_annotation/"
                            "hubisz_tmrca_sites_8afr.tight.tsv",
                    help="resolved site reads, for the 95%% interval bounds")
    ap.add_argument("--loci",
                    default="catalog_annotation/table_s51_loci.tsv")
    ap.add_argument("-o", "--out",
                    default="catalog_annotation/"
                            "supplementary_table2_fixed_derived.tsv")
    args = ap.parse_args()

    loci = load_preal(args.loci)
    exact = load_site_exact(args.site_exact)
    reads = load_tmrca_reads(args.tmrca_reads)
    coord, cdesc = default_coord([])
    sys.stderr.write(f"site intervals: 5/95 reconstructed on {cdesc}\n")

    rows = [(r, "99") for r in csv.DictReader(
        open(args.sites, encoding="utf-8"), delimiter="\t")]
    if not rows:
        sys.exit("no sites")
    n99 = len(rows)
    if args.sites95 and os.path.exists(args.sites95):
        rows += [(r, "95") for r in csv.DictReader(
            open(args.sites95, encoding="utf-8"), delimiter="\t")]
    else:
        sys.stderr.write(f"note: {args.sites95} absent, "
                         "the >95% tier is omitted\n")
    sys.stderr.write(f"sites: {n99:,} at >99%, "
                     f"{len(rows) - n99:,} added at >95%\n")

    out = []
    for r, tier in rows:
        anc_af = float(r["g1k_anc_af"] or 0.0)
        mod_der = int(round((1.0 - anc_af) * G1K_HAPS))

        nd = nt = 0
        for g in NEANDERTALS:
            d, t = state_counts(r.get(f"{g}_state", ""))
            nd += d
            nt += t
        dd = dt = 0
        for g in DENISOVANS:
            d, t = state_counts(r.get(f"{g}_state", ""))
            dd += d
            dt += t

        # Table S4.2 takes the strongest subset; see qualifies() for the rule.
        strong = qualifies(r["classes"])
        key = (r["chrom"], r["pos"])
        # Meeting the rule only puts a site in the table if it is also in the
        # >99% tier. The tier test used to be implicit: on the 5 kb screen the
        # only >95% sites that qualified were the two named above, so a bare
        # `strong` happened to give the right answer. The 1 kb screen adds
        # chr1:37,249,786, a conserved non-coding base 11 kb from GRIK3 with
        # the highest CADD of any site here, which would then have walked into
        # Table S4.2 without anyone choosing to put it there. Anything from
        # the >95% tier now has to be named in TABLE_S52_EXTRA, as MCF2L2 is.
        printed = ((strong and tier == "99") or key in TABLE_S52_EXTRA) \
            and key not in TABLE_S52_OMIT

        pr, genes = preal_at(loci, r["chrom"], int(r["pos"]))
        rec = {
            "chrom": r["chrom"], "pos": r["pos"],
            "modern_freq_tier": tier,
            "ancestral": r["ancestral"], "derived": r["derived"],
            "modern_derived_over_total": f"{mod_der}/{G1K_HAPS}",
            "modern_derived_freq": f"{1.0 - anc_af:.5f}",
            "nea_derived_over_observed": f"{nd}/{nt}",
            "den_derived_over_observed": f"{dd}/{dt}",
            "nea_genomes_called": r["nea_called"],
            "den_genomes_called": r["den_called"],
            "callers_nea": r["callers_nea"], "callers_den": r["callers_den"],
        }
        for g in NEANDERTALS + DENISOVANS:
            rec[f"{g}_state"] = r.get(f"{g}_state", "")
        rec.update({
            "gene": r["gene"], "transcript": r["transcript"],
            "consequence": r["consequence"],
            "aa_ancestral": r["aa_ancestral"], "aa_derived": r["aa_derived"],
            "protein_position": r["protein_position"],
            "functional_classes": r["classes"],
            "phylop_46way": r["phylop"], "cadd_v16_phred": r["cadd_max"],
            "functionally_qualifying": r["qualifying"],
            "meets_table_s5_2_rule": "yes" if strong else "no",
            "in_table_s5_2": "yes" if printed else "no",
            "catalog_locus_genes": genes,
            "catalog_locus_preal": pr,
        })
        ex = exact.get((r["chrom"], r["pos"]))
        rd = reads.get((r["chrom"], r["pos"]))
        rec.update({
            "preal_site": ex["preal_site"] if ex else "NA",
            "preal_nearest_grid": ex["preal_nearest_grid"] if ex else "NA",
            "afr_mean_ka_site": ex["afr_mean_ka"] if ex else "NA",
            "afr_ci90_ka_site": ci90(rd, "afr", coord),
            "nea_mean_ka_site": ex["nea_mean_ka"] if ex else "NA",
            "nea_ci90_ka_site": ci90(rd, "nea", coord),
            "den_mean_ka_site": ex["den_mean_ka"] if ex else "NA",
            "den_ci90_ka_site": ci90(rd, "den", coord),
        })
        out.append(rec)

    out.sort(key=lambda r: (int(r["chrom"]), int(r["pos"])))
    with open(args.out, "w", newline="", encoding="utf-8") as f:
        w = csv.DictWriter(f, delimiter="\t", fieldnames=list(out[0]))
        w.writeheader()
        w.writerows(out)

    nea_full = sum(1 for r in out
                   if r["nea_derived_over_observed"].split("/")[1] == "10")
    den_full = sum(1 for r in out
                   if r["den_derived_over_observed"].split("/")[1] == "4")
    print(f"wrote {args.out}")
    print(f"  sites                              {len(out)}")
    for tier in ("99", "95"):
        n = sum(1 for r in out if r["modern_freq_tier"] == tier)
        if n:
            print(f"    at >{tier}% derived in 1000 Genomes  {n}")
    print(f"  functionally qualifying            "
          f"{sum(1 for r in out if r['functionally_qualifying'] == 'yes')}")
    print(f"  meets the Table S4.2 rule          "
          f"{sum(1 for r in out if r['meets_table_s5_2_rule'] == 'yes')}")
    print(f"  printed in Table S4.2              "
          f"{sum(1 for r in out if r['in_table_s5_2'] == 'yes')}")
    print(f"  with a site-exact P(real)          "
          f"{sum(1 for r in out if r['preal_site'] != 'NA')}")
    print(f"  all 10 Neandertal haplotypes seen  {nea_full}")
    print(f"  all 4 Denisovan haplotypes seen    {den_full}")
    print(f"  protein-altering                   "
          f"{sum(1 for r in out if 'missense' in r['consequence'] or 'stop' in r['consequence'] or 'start' in r['consequence'])}")
    return 0


if __name__ == "__main__":
    sys.exit(main())
```
