## Supplementary Information section 4 code base for "Hypothesis: A modern human range expansion ∼300,000 years ago had a critical role in Neandertal origins": canonicalize_protein_positions.py

```
#!/usr/bin/env python3
"""Rewrite the protein-level columns of the fixed-derived screen onto canonical
transcripts.

The O2 screen records whichever transcript its annotation source returned first,
which is often not the canonical one. That is harmless for deciding whether a
site is coding, but it makes the reported residue numbers unusable: ARHGAP32
Glu1489Asp was emitted as Glu1140Asp on ARHGAP32-206, SETD2 Val259Ile as
Val215Ile on SETD2-203, and TOMM20L Asp30Asn was called against a
nonsense-mediated-decay transcript. Residue numbers that do not match the
canonical protein cannot be checked against the domain literature, which is
what they are mainly used for.

This queries Ensembl VEP on GRCh37 -- the assembly the screen used -- and takes
the consequence on the canonical protein-coding transcript.

Orientation matters and is easy to get backwards. These sites are fixed derived
in modern humans, so the human reference carries the DERIVED allele. Querying
the derived allele as the alternate returns "synonymous_variant" for every site,
because the alternate equals the reference. The ancestral allele is therefore
submitted as the alternate and the resulting change is inverted for reporting.
"""

import argparse
import csv
import json
import sys
import urllib.request

SERVER = "https://grch37.rest.ensembl.org"
CODING = ("missense", "synonymous", "stop", "start", "coding")

# The note writes residues in three-letter form, so match it rather than
# emitting VEP's single-letter codes and leaving the table internally mixed.
AA3 = {"A": "Ala", "R": "Arg", "N": "Asn", "D": "Asp", "C": "Cys",
       "Q": "Gln", "E": "Glu", "G": "Gly", "H": "His", "I": "Ile",
       "L": "Leu", "K": "Lys", "M": "Met", "F": "Phe", "P": "Pro",
       "S": "Ser", "T": "Thr", "W": "Trp", "Y": "Tyr", "V": "Val",
       "*": "Ter", "X": "Ter"}


def three(a):
    return AA3.get(a, a)


def vep(variants):
    """POST a batch to VEP, return input -> canonical transcript consequence."""
    req = urllib.request.Request(
        f"{SERVER}/vep/human/region?canonical=1",
        data=json.dumps({"variants": variants}).encode(),
        headers={"Content-type": "application/json",
                 "Accept": "application/json"})
    with urllib.request.urlopen(req, timeout=120) as fh:
        out = json.load(fh)
    best = {}
    for v in out:
        for t in v.get("transcript_consequences", []):
            if (t.get("canonical") == 1
                    and t.get("biotype") == "protein_coding"
                    and t.get("amino_acids")):
                best[v["input"]] = t
                break
    return best


def main():
    ap = argparse.ArgumentParser()
    ap.add_argument("-i", "--infile",
                    default="catalog_annotation/fd_af99.functional.tsv")
    ap.add_argument("-o", "--out",
                    default="catalog_annotation/"
                            "fd_af99.functional.canonical.tsv")
    a = ap.parse_args()

    rows = list(csv.DictReader(open(a.infile, encoding="utf-8"),
                               delimiter="\t"))
    coding = [r for r in rows
              if any(k in (r.get("consequence") or "").lower() for k in CODING)]
    sys.stderr.write(f"{len(rows):,} sites, {len(coding)} with a coding "
                     f"consequence\n")
    if not coding:
        sys.exit("nothing to canonicalise")

    # reference carries the derived allele, so the ancestral allele is the alt
    key = {}
    variants = []
    for r in coding:
        s = f"{r['chrom'].replace('chr','')} {r['pos']} . {r['derived']} {r['ancestral']} . . ."
        key[s] = r
        variants.append(s)

    got = {}
    for i in range(0, len(variants), 150):
        got.update(vep(variants[i:i + 150]))
    sys.stderr.write(f"canonical consequence returned for {len(got)}/"
                     f"{len(variants)}\n")

    changed = []
    for s, t in got.items():
        r = key[s]
        aa = t["amino_acids"].split("/")
        # A synonymous change returns one residue rather than a pair; the
        # ancestral and derived residues are then the same. Skipping those
        # would leave synonymous sites on the old non-canonical numbering.
        if len(aa) == 2:
            # VEP reports reference/alternate = derived/ancestral; invert it
            anc, der = three(aa[1]), three(aa[0])
        elif len(aa) == 1:
            anc = der = three(aa[0])
        else:
            continue
        pos = str(t["protein_start"])
        old = (r.get("transcript"), r.get("protein_position"))
        if (r.get("protein_position") != pos
                or r.get("transcript") != t["transcript_id"]):
            changed.append((r["chrom"], r["pos"], r.get("gene"),
                            f"{r.get('aa_ancestral','')}{r.get('protein_position','')}"
                            f"{r.get('aa_derived','')}", old[0],
                            f"{anc}{pos}{der}", t["transcript_id"]))
        r["transcript"] = t["transcript_id"]
        r["aa_ancestral"], r["aa_derived"] = anc, der
        r["protein_position"] = pos
        r["consequence"] = ",".join(t["consequence_terms"])

    with open(a.out, "w", newline="", encoding="utf-8") as f:
        w = csv.DictWriter(f, delimiter="\t", fieldnames=list(rows[0]))
        w.writeheader()
        w.writerows(rows)

    sys.stderr.write(f"wrote {a.out}\n  {len(changed)} residue numbers or "
                     f"transcripts corrected\n")
    for c in changed:
        sys.stderr.write(f"    chr{c[0]}:{int(c[1]):,} {c[2] or '-':<11} "
                         f"{c[3]:<12} ({c[4]})  ->  {c[5]:<12} ({c[6]})\n")
    return 0


if __name__ == "__main__":
    sys.exit(main())
```
