## Supplementary Information section 4 code base for "Hypothesis: A modern human range expansion ∼300,000 years ago had a critical role in Neandertal origins": cluster_relate_one_chr.sh

```
#!/usr/bin/env bash
# Run on a cluster compute node: RelateExtract AncToNewick for one chromosome, then
# relate_marginal_tmrca_grid.py. Intended to be called from Slurm or manually.
#
# Required environment variables:
#   CHROM          Chromosome label for output TSV (e.g. 1, 22, X) — must match a line in relate_hg19_chrom_sizes.tsv
#   RELATE_BIN     Path to RelateExtract binary (on O2: .../relate-build/bin/RelateExtract)
#   RELATE_DATA    Directory containing per-chromosome .anc.gz / .mut.gz
#   ANC_BASENAME   Filename pattern: use {CHR} placeholder, e.g. "1000GP_phase3_chr{CHR}.anc.gz"
#   MUT_BASENAME   Same for .mut, e.g. "1000GP_phase3_chr{CHR}.mut.gz"
#   REPO_ROOT      Path to Cursor_superarchaic (contains scripts/)
#   OUTNW          Where to write chrN.newick and chrN.pos
#   OUTTSV         Where to write chrN_full_sample_tmrca_5kb.tsv
#   VENV           Optional Python venv (no longer required: the grid script
#                  has no third-party dependencies as of Sep 2026)
#
#   POPLABELS      Relate .poplabels for the dataset (needed for subset TMRCAs)
#
# Optional:
#   YEARS_PER_GEN  default 28
#   GRID_STEP      default 5000
#   SUBSETS        Space-separated PREFIX=POPS specs, default:
#                    "relate_afr1322=AFR relate_yri216=YRI"
#                  All subsets are computed in a single pass over the newick,
#                  because parsing 5,008-tip trees is what costs, not the
#                  MRCA lookups.

set -euo pipefail

: "${CHROM:?}"
: "${RELATE_BIN:?}"
: "${RELATE_DATA:?}"
: "${ANC_BASENAME:?}"
: "${MUT_BASENAME:?}"
: "${REPO_ROOT:?}"
: "${OUTNW:?}"
: "${OUTTSV:?}"

: "${POPLABELS:?}"

YEARS_PER_GEN="${YEARS_PER_GEN:-28}"
GRID_STEP="${GRID_STEP:-5000}"
SUBSETS="${SUBSETS:-relate_afr1322=AFR relate_yri216=YRI}"

if [[ ! -f "$POPLABELS" ]]; then
  echo "ERROR: missing POPLABELS=$POPLABELS"
  exit 1
fi

SIZES="$REPO_ROOT/scripts/relate_hg19_chrom_sizes.tsv"
CHROM_LEN="$(awk -v c="$CHROM" '!/^#/ && $1 == c { print $2; exit }' "$SIZES")"
if [[ -z "${CHROM_LEN}" ]]; then
  echo "ERROR: unknown CHROM=$CHROM (not in $SIZES)"
  exit 1
fi

# Expand {CHR} in basename patterns (e.g. chr{CHR} or {CHR})
expand_pat() {
  local p="$1"
  echo "${p//\{CHR\}/$CHROM}"
}

ANC_FILE="$RELATE_DATA/$(expand_pat "$ANC_BASENAME")"
MUT_FILE="$RELATE_DATA/$(expand_pat "$MUT_BASENAME")"

if [[ ! -f "$ANC_FILE" ]]; then
  echo "ERROR: missing $ANC_FILE"
  exit 1
fi
if [[ ! -f "$MUT_FILE" ]]; then
  echo "ERROR: missing $MUT_FILE"
  exit 1
fi

mkdir -p "$OUTNW" "$OUTTSV"

# Every intermediate is namespaced by VARIANT so the pre-Ne and post-Ne runs can
# proceed concurrently without overwriting each other. This is not hypothetical:
# an earlier hand-launched `sbatch --wrap` AncToNewick spent 10+ hours writing
# chr22.newick from another node while this script was also targeting that exact
# path, and the collision was invisible because both writers produce a file with
# the same name and a plausible size.
VARIANT="${VARIANT:-postne}"
PREFIX="$OUTNW/chr${CHROM}.${VARIANT}"

echo "=== CHROM=$CHROM len=$CHROM_LEN ==="
echo "ANC=$ANC_FILE"
echo "MUT=$MUT_FILE"

# ---------------------------------------------------- branch-length re-inference
# The O2 trees are pre-Ne: basename 1000GP_Phase3_mask_prene, and results/ holds
# no post-EstimatePopulationSize set. Their branch lengths were therefore fitted
# under a constant-Ne assumption, which distorts TMRCA in a depth-dependent way
# -- not by a constant factor, so it cannot be absorbed into a single calibration
# multiplier downstream. A .coal file IS distributed with the dataset, so the
# lengths can be re-inferred conditional on the estimated size history.
#
# Set COAL to enable this. Left unset, the script behaves as before and uses the
# pre-Ne lengths, which is only appropriate for a topology-only sanity check.
#
# The mode name differs across Relate releases, so it is discovered from --help
# rather than assumed, and the script stops with the help text if neither
# candidate is present.
COAL="${COAL:-}"
MU="${MU:-1.25e-8}"
if [[ -n "$COAL" ]]; then
  if [[ ! -f "$COAL" ]]; then
    echo "ERROR: COAL=$COAL not found"
    exit 1
  fi
  BINDIR="$(dirname "$RELATE_BIN")"
  RCR="$BINDIR/RelateCoalescentRate"
  REL="$BINDIR/Relate"

  MODE=""
  TOOL=""
  if [[ -x "$RCR" ]] && "$RCR" --help 2>&1 | grep -q "ReEstimateBranchLengths"; then
    TOOL="$RCR"; MODE="ReEstimateBranchLengths"
  elif [[ -x "$REL" ]] && "$REL" --help 2>&1 | grep -q "InferBranchLengths"; then
    TOOL="$REL"; MODE="InferBranchLengths"
  fi

  if [[ -z "$MODE" ]]; then
    echo "ERROR: no branch-length re-inference mode found in this Relate build."
    echo "--- RelateCoalescentRate --help ---"
    [[ -x "$RCR" ]] && "$RCR" --help 2>&1 | head -40
    echo "--- Relate --help ---"
    [[ -x "$REL" ]] && "$REL" --help 2>&1 | head -40
    exit 1
  fi

  echo "re-inferring branch lengths: $(basename "$TOOL") --mode $MODE"
  echo "  coal=$COAL  mu=$MU"

  # RelateCoalescentRate takes -i/-o as an EXTENSIONLESS PREFIX and expects
  # <prefix>.anc and <prefix>.mut side by side; it has no --anc/--mut options
  # (passing them aborts with cxxopts option_not_exists). The dataset keeps anc
  # and mut in separate directories, so there is no such prefix on disk and one
  # has to be manufactured. Symlinks rather than copies: these are ~500 MB each.
  STAGE="$OUTNW/stage_chr${CHROM}.${VARIANT}"
  mkdir -p "$STAGE"
  IN="$STAGE/chr${CHROM}"
  ln -sfn "$ANC_FILE" "${IN}.anc.gz"
  ln -sfn "$MUT_FILE" "${IN}.mut.gz"

  REPREFIX="$OUTNW/chr${CHROM}_reinferred"
  run_reinfer() {
    "$TOOL" --mode "$MODE" -i "$1" -o "$REPREFIX" \
            --coal "$COAL" -m "$MU" --years_per_gen "$YEARS_PER_GEN"
  }
  found_anc() {
    for cand in "$REPREFIX.anc.gz" "$REPREFIX.anc"; do
      [[ -f "$cand" ]] && { echo "$cand"; return 0; }
    done
    return 1
  }

  # Try the gzipped symlinks first. Relate handles .gz in recent builds, but not
  # all of them; if nothing is emitted, decompress once and retry rather than
  # failing the job over a compression detail.
  run_reinfer "$IN" || true
  if ! found_anc >/dev/null; then
    echo "  gz input produced nothing; decompressing and retrying"
    gunzip -c "$ANC_FILE" > "${IN}.anc"
    gunzip -c "$MUT_FILE" > "${IN}.mut"
    rm -f "${IN}.anc.gz" "${IN}.mut.gz"
    run_reinfer "$IN" || true
  fi

  NEW_ANC="$(found_anc || true)"
  if [[ -z "$NEW_ANC" ]]; then
    echo "ERROR: re-inference produced no anc file at $REPREFIX.anc[.gz]"
    echo "--- $OUTNW ---"; ls -l "$OUTNW" | head -20
    echo "--- $STAGE ---"; ls -lL "$STAGE" | head -10
    exit 1
  fi
  ANC_FILE="$NEW_ANC"
  for cand in "$REPREFIX.mut.gz" "$REPREFIX.mut"; do
    [[ -f "$cand" ]] && { MUT_FILE="$cand"; break; }
  done
  echo "  now using ANC=$ANC_FILE"
  echo "  now using MUT=$MUT_FILE"
else
  echo "WARNING: COAL unset -- using pre-Ne branch lengths. TMRCAs from these"
  echo "         are not coalescent-calibrated; set COAL for real output."
fi

# AncToNewick is the expensive step (10+ h for a 5,008-tip chromosome), so an
# already-exported newick can be reused -- but only when asked for explicitly by
# NEWICK_IN, and only after checking that its tree count matches its .pos index.
# A truncated newick from a killed job has a perfectly plausible size and would
# otherwise yield a TMRCA track that is simply short, with no error raised.
if [[ -n "${NEWICK_IN:-}" ]]; then
  NIN_NW="$NEWICK_IN"
  NIN_POS="${NEWICK_IN%.newick}.pos"
  if [[ ! -f "$NIN_NW" || ! -f "$NIN_POS" ]]; then
    echo "ERROR: NEWICK_IN set but $NIN_NW / $NIN_POS missing"
    exit 1
  fi
  n_nw=$(grep -c ';' "$NIN_NW" || true)
  n_pos=$(( $(wc -l < "$NIN_POS") ))
  echo "reusing newick: $NIN_NW"
  echo "  trees=$n_nw  pos_lines=$n_pos"
  if [[ "$n_nw" -eq 0 || "$n_pos" -eq 0 ]]; then
    echo "ERROR: empty newick or pos"
    exit 1
  fi
  # Allow one line of slack for a header or trailing newline.
  if (( n_nw < n_pos - 1 || n_nw > n_pos + 1 )); then
    echo "ERROR: tree count $n_nw does not match pos count $n_pos --"
    echo "       the newick looks truncated (killed job?). Refusing to use it."
    exit 1
  fi
  PREFIX="${NEWICK_IN%.newick}"
else
  "$RELATE_BIN" \
    --mode AncToNewick \
    --anc "$ANC_FILE" \
    --mut "$MUT_FILE" \
    --first_bp 1 \
    --last_bp "$CHROM_LEN" \
    -o "$PREFIX"
fi

if [[ -n "${VENV:-}" && -f "$VENV/bin/activate" ]]; then
  # shellcheck source=/dev/null
  source "$VENV/bin/activate"
fi

SUBSET_ARGS=()
for s in $SUBSETS; do
  SUBSET_ARGS+=(--subset "$s")
done

python3 "$REPO_ROOT/scripts/relate_marginal_tmrca_grid.py" \
  --pos "${PREFIX}.pos" \
  --newick "${PREFIX}.newick" \
  --chrom "$CHROM" \
  --chrom-len "$CHROM_LEN" \
  --grid-step "$GRID_STEP" \
  --years-per-gen "$YEARS_PER_GEN" \
  --poplabels "$POPLABELS" \
  "${SUBSET_ARGS[@]}" \
  -o "$OUTTSV/chr${CHROM}.${VARIANT}_subset_tmrca_5kb.tsv"

echo "Done chr${CHROM}"
```
