## Supplementary Information section 4 code base for "Hypothesis: A modern human range expansion ∼300,000 years ago had a critical role in Neandertal origins": compare_hubisz_tmrca_runs.py

```
#!/usr/bin/env python3
"""
Compare two hubisz_out_bed_tmrca_scan.py genome TSVs and plot overlapping 5 kb sites.

(a) Print header equality, column counts, row counts, and inner-join count on (chrom, pos).
(b) Archaic: Neandertal-only and Denisovan-only mean kya (columns 04_* and 05_*), both runs.
(c) Modern: 01_modern_only mean kya, both runs (semantics depend on which queries JSON each run used).

Requires: pip install pandas matplotlib

Example:
  python3 scripts/compare_hubisz_tmrca_runs.py \\
    --tsv-a /n/groups/reich/DAVID/CURSOR_ARGWEAVER/tmrca_scan_out/hubisz_tmrca_5kb.tsv \\
    --label-a A_round1_2026.05.08 \\
    --tsv-b /n/groups/reich/DAVID/CURSOR_ARGWEAVER/tmrca_scan_out_B_round2_2026.05.12/hubisz_tmrca_5kb.tsv \\
    --label-b B_round2_2026.05.12 \\
    --chrom 4 --start-bp 71000000 --end-bp 72000000 \\
    --out /tmp/chr4_71_72_tmrca_compare
"""

from __future__ import annotations

import argparse
import sys
from pathlib import Path

import pandas as pd


def read_tmrca_tsv(path: Path) -> pd.DataFrame:
    df = pd.read_csv(path, sep="\t", dtype={"chrom": int, "pos": int})
    return df


def na_rate(s: pd.Series) -> float:
    return float(s.isna().mean()) if len(s) else 0.0


def to_numeric_mean(df: pd.DataFrame, col: str) -> pd.Series:
    """Mean kya column may contain 'NA' strings."""
    if col not in df.columns:
        raise SystemExit(f"Missing column {col!r} in dataframe")
    return pd.to_numeric(df[col], errors="coerce")


def main() -> None:
    ap = argparse.ArgumentParser(description=__doc__)
    ap.add_argument("--tsv-a", type=Path, required=True, help="First collated TSV (e.g. older run)")
    ap.add_argument("--tsv-b", type=Path, required=True, help="Second collated TSV (e.g. B.round2)")
    ap.add_argument("--label-a", default="run_A")
    ap.add_argument("--label-b", default="run_B")
    ap.add_argument("--chrom", type=int, default=4)
    ap.add_argument("--start-bp", type=int, default=71_000_000)
    ap.add_argument("--end-bp", type=int, default=72_000_000)
    ap.add_argument(
        "--out",
        type=Path,
        required=True,
        help="Output image path without suffix (writes .png and .pdf)",
    )
    args = ap.parse_args()

    try:
        import matplotlib.pyplot as plt
    except ImportError:
        sys.stderr.write("Install matplotlib: pip install matplotlib\n")
        raise

    p_a, p_b = args.tsv_a, args.tsv_b
    if not p_a.is_file() or not p_b.is_file():
        raise SystemExit("One or both TSV paths are not readable files.")

    df_a = read_tmrca_tsv(p_a)
    df_b = read_tmrca_tsv(p_b)

    key = ["chrom", "pos"]
    dup_a = int(df_a.duplicated(subset=key).sum())
    dup_b = int(df_b.duplicated(subset=key).sum())
    if dup_a:
        print(f"WARNING: dropping {dup_a:,} duplicate (chrom,pos) rows in A (keeping first)")
        df_a = df_a.drop_duplicates(subset=key, keep="first")
    if dup_b:
        print(f"WARNING: dropping {dup_b:,} duplicate (chrom,pos) rows in B (keeping first)")
        df_b = df_b.drop_duplicates(subset=key, keep="first")

    cols_a = list(df_a.columns)
    cols_b = list(df_b.columns)

    print("=== (a) Format check ===")
    print(f"A rows: {len(df_a):,}  path: {p_a}")
    print(f"B rows: {len(df_b):,}  path: {p_b}")
    if cols_a == cols_b:
        print(f"Headers: IDENTICAL ({len(cols_a)} columns)")
    else:
        print("Headers: DIFFER")
        sa, sb = set(cols_a), set(cols_b)
        print("  Only in A:", sorted(sa - sb)[:30], ("..." if len(sa - sb) > 30 else ""))
        print("  Only in B:", sorted(sb - sa)[:30], ("..." if len(sb - sa) > 30 else ""))
        raise SystemExit("Stop: align query JSON / collate so headers match before plotting.")

    merged = df_a[key].merge(df_b[key], on=key, how="inner")
    print(f"Inner join on chrom,pos: {len(merged):,} sites (both runs, unique keys)")

    sub_a = df_a.set_index(key)
    sub_b = df_b.set_index(key)
    common = sub_a.index.intersection(sub_b.index)
    sub_a = sub_a.loc[common].sort_index()
    sub_b = sub_b.loc[common].sort_index()
    assert sub_a.index.equals(sub_b.index)

    reg = (sub_a.index.get_level_values("chrom") == args.chrom) & (
        sub_a.index.get_level_values("pos") >= args.start_bp
    ) & (sub_a.index.get_level_values("pos") < args.end_bp)
    xa = sub_a.index.get_level_values("pos")[reg].to_numpy()
    if len(xa) == 0:
        raise SystemExit(
            f"No overlapping rows in chr{args.chrom} [{args.start_bp}, {args.end_bp}). "
            "Check chrom numbering and coordinates."
        )

    col_mod = "01_modern_only_mean_kya"
    col_nea = "04_both_Nea_only_mean_kya"
    col_den = "05_Den_only_mean_kya"
    for c in (col_mod, col_nea, col_den):
        if c not in sub_a.columns or c not in sub_b.columns:
            raise SystemExit(f"Expected column {c!r} missing — was a different queries JSON used?")

    ya_mod = to_numeric_mean(sub_a.loc[reg], col_mod)
    yb_mod = to_numeric_mean(sub_b.loc[reg], col_mod)
    ya_nea = to_numeric_mean(sub_a.loc[reg], col_nea)
    yb_nea = to_numeric_mean(sub_b.loc[reg], col_nea)
    ya_den = to_numeric_mean(sub_a.loc[reg], col_den)
    yb_den = to_numeric_mean(sub_b.loc[reg], col_den)

    mask_plot = ya_mod.notna() & yb_mod.notna() & ya_nea.notna() & yb_nea.notna() & ya_den.notna() & yb_den.notna()
    print(
        f"Region chr{args.chrom} {args.start_bp}-{args.end_bp}: {mask_plot.sum()} / {len(mask_plot)} "
        f"overlap sites with non-NA {col_mod}, {col_nea}, {col_den} in BOTH runs"
    )
    print(f"  NA fraction A: mod {na_rate(ya_mod):.3f} nea {na_rate(ya_nea):.3f} den {na_rate(ya_den):.3f}")
    print(f"  NA fraction B: mod {na_rate(yb_mod):.3f} nea {na_rate(yb_nea):.3f} den {na_rate(yb_den):.3f}")

    x = xa[mask_plot.to_numpy()]
    fig, axes = plt.subplots(2, 1, figsize=(10, 7), sharex=True, constrained_layout=True)

    axes[0].plot(x / 1e6, ya_nea[mask_plot], label=f"{args.label_a} Neandertal-only", lw=1.2)
    axes[0].plot(x / 1e6, yb_nea[mask_plot], label=f"{args.label_b} Neandertal-only", lw=1.2)
    axes[0].plot(x / 1e6, ya_den[mask_plot], ls="--", label=f"{args.label_a} Denisovan-only", lw=1.2)
    axes[0].plot(x / 1e6, yb_den[mask_plot], ls="--", label=f"{args.label_b} Denisovan-only", lw=1.2)
    axes[0].set_ylabel("TMRCA mean (kya)")
    axes[0].set_title(
        "Archaic TMRCA (Neandertal-only vs Denisovan-only; same columns as hubisz_2afr queries)"
    )
    axes[0].legend(loc="best", fontsize=8)

    axes[1].plot(x / 1e6, ya_mod[mask_plot], label=args.label_a, lw=1.2)
    axes[1].plot(x / 1e6, yb_mod[mask_plot], label=args.label_b, lw=1.2)
    axes[1].set_ylabel("TMRCA mean (kya)")
    axes[1].set_xlabel(f"chr{args.chrom} position (Mb)")
    axes[1].set_title("Modern-only TMRCA (column 01_modern_only_mean_kya)")
    axes[1].legend(loc="best")

    out = args.out
    out.parent.mkdir(parents=True, exist_ok=True)
    png = out.with_suffix(".png")
    pdf = out.with_suffix(".pdf")
    fig.savefig(png, dpi=150)
    fig.savefig(pdf)
    print(f"Wrote {png} and {pdf}")
    plt.close(fig)


if __name__ == "__main__":
    main()
```
