## Supplementary Information section 4 code base for "Hypothesis: A modern human range expansion ∼300,000 years ago had a critical role in Neandertal origins": compare_screen_processings.py

```
#!/usr/bin/env python3
"""Do the newly located archaic VCFs agree with the earlier processing?

Three independent processings of the same catalog regions now exist:

  old    catalog_annotation/fixed_derived_preal50.sites.tsv -- the earlier
         streaming screen, which RECORDED the Neandertal state rather than
         requiring it (hence 3 rows where both Neandertals are ancestral_hom)
  panel  out/fixed_derived.chr*.tsv -- the 6-genome panel reading the
         per-chromosome VCFs located on the filesystem in September 2026
  legacy out_legacy_pair/*.tsv -- the staged merged GATK-era VCFs

Agreement between them at shared positions tests the bioinformatics rather than
the biology: allele polarity, coordinate alignment, sample-column selection and
1000 Genomes frequency lookup are all shared assumptions that would break
loudly here. Disagreement about which SITES appear is expected, because the
processings apply different rules; disagreement about the STATE of a genome at
a position both agree on is not.

Usage:
  python3 compare_screen_processings.py --old catalog_annotation/fixed_derived_preal50.sites.tsv \\
      --new 'out/fixed_derived.chr*.tsv'
"""

from __future__ import annotations

import argparse
import csv
import glob
import sys
from collections import Counter


def read_tsv(paths):
    rows = []
    for p in sorted(glob.glob(paths) if isinstance(paths, str) else paths):
        with open(p, encoding="utf-8") as f:
            rows.extend(csv.DictReader(f, delimiter="\t"))
    return rows


def gt_to_state(gt, anc, der):
    """Old file stores a genotype like 'G/G'; convert to the new state vocabulary."""
    if not gt or gt in ("", "./."):
        return "no_data"
    b = set(gt.replace("|", "/").split("/"))
    if b == {anc}:
        return "ancestral_hom"
    if b == {der}:
        return "derived_hom"
    return "het"


def main():
    ap = argparse.ArgumentParser()
    ap.add_argument("--old", required=True)
    ap.add_argument("--new", required=True)
    ap.add_argument("--af-tol", type=float, default=0.002,
                    help="tolerated absolute difference in 1000G frequency")
    args = ap.parse_args()

    old = {(r["chrom"], int(r["pos"])): r for r in read_tsv([args.old])}
    new = {(r["chrom"], int(r["pos"])): r for r in read_tsv(args.new)}
    shared = sorted(set(old) & set(new))

    print(f"old processing : {len(old)} sites")
    print(f"new processing : {len(new)} sites")
    print(f"shared positions: {len(shared)}")
    print(f"  only in old: {len(set(old) - set(new))}    "
          f"only in new: {len(set(new) - set(old))}")
    if not shared:
        sys.exit("no shared positions -- coordinate systems may differ")

    print("\n=== 1. allele polarity (the assumption everything rests on) ===")
    pol = Counter()
    examples = []
    for k in shared:
        o, n = old[k], new[k]
        same_anc = o["ancestral"].upper() == n["ancestral"].upper()
        same_der = o["derived_ref"].upper() == n["derived"].upper()
        pol["both agree" if (same_anc and same_der) else
            "ancestral differs" if not same_anc else "derived differs"] += 1
        if not (same_anc and same_der) and len(examples) < 5:
            examples.append(f"    {k[0]}:{k[1]} old {o['ancestral']}>{o['derived_ref']}"
                            f"  new {n['ancestral']}>{n['derived']}")
    for k, v in pol.most_common():
        print(f"  {k:<20} {v}")
    for e in examples:
        print(e)

    print("\n=== 2. archaic genotype agreement at shared positions ===")
    print("  Old columns 'altai'/'vindija' are states; 'denisova_gt' is a genotype.")
    pairs = [("altai", "Altai_state"), ("vindija", "Vindija33.19_state")]
    for ocol, ncol in pairs:
        if ocol not in next(iter(old.values())) or ncol not in next(iter(new.values())):
            print(f"  {ocol} -> {ncol}: column absent, skipped")
            continue
        agree = disagree = 0
        detail = Counter()
        for k in shared:
            a, b = old[k][ocol], new[k][ncol]
            if a == b:
                agree += 1
            else:
                disagree += 1
                detail[f"old={a} new={b}"] += 1
        tot = agree + disagree
        print(f"  {ocol:<8} -> {ncol:<20} {agree}/{tot} identical "
              f"({100.0*agree/tot:.1f}%)")
        for k, v in detail.most_common(6):
            print(f"      {k}  x{v}")

    ncol = "Denisova3_state"
    if ncol in next(iter(new.values())):
        agree = disagree = 0
        detail = Counter()
        for k in shared:
            o, n = old[k], new[k]
            want = gt_to_state(o.get("denisova_gt", ""), o["ancestral"].upper(),
                               o["derived_ref"].upper())
            if want == n[ncol]:
                agree += 1
            else:
                disagree += 1
                detail[f"old={want} new={n[ncol]}"] += 1
        tot = agree + disagree
        print(f"  denisova_gt -> {ncol:<20} {agree}/{tot} identical "
              f"({100.0*agree/tot:.1f}%)")
        for k, v in detail.most_common(6):
            print(f"      {k}  x{v}")

    print("\n=== 3. 1000 Genomes frequency agreement ===")
    print("  Old column is g1k_alt_af, new is g1k_anc_af. These should be the")
    print("  SAME number: the screen requires the reference to carry the derived")
    print("  allele, so ALT is the ancestral allele. A systematic difference")
    print("  would mean that identification is wrong somewhere.")
    dif = []
    for k in shared:
        try:
            a = float(old[k]["g1k_alt_af"]); b = float(new[k]["g1k_anc_af"])
        except (KeyError, ValueError):
            continue
        dif.append((abs(a - b), k, a, b))
    if dif:
        dif.sort(reverse=True)
        within = sum(1 for d, *_ in dif if d <= args.af_tol)
        print(f"  within {args.af_tol}: {within}/{len(dif)} "
              f"({100.0*within/len(dif):.1f}%)")
        print("  largest discrepancies:")
        for d, k, a, b in dif[:5]:
            print(f"    {k[0]}:{k[1]}  old={a:.6f}  new={b:.6f}  diff={d:.6f}")

    print("\n=== 4. sites the old screen kept that contradict the scan ===")
    print("  The earlier screen recorded rather than required the Neandertal")
    print("  state, so it retained rows a strict rule would drop. These are the")
    print("  informative cases: present in the catalog regions, yet not matching")
    print("  the pattern the ARGweaver run assumed.")
    bad = [k for k in old
           if old[k].get("altai") == "ancestral_hom"
           or old[k].get("vindija") == "ancestral_hom"]
    print(f"  found {len(bad)}:")
    for k in sorted(bad)[:12]:
        o = old[k]
        n = new.get(k)
        print(f"    {k[0]}:{k[1]} {o['ancestral']}>{o['derived_ref']} "
              f"altai={o.get('altai')} vindija={o.get('vindija')} "
              f"af={o.get('g1k_alt_af')} gene={o.get('gene','')}")
        print(f"        new screen: "
              + (f"Altai={n['Altai_state']} Vindija={n['Vindija33.19_state']}"
                 if n else "position absent (dropped by the unanimity rule)"))
    return 0


if __name__ == "__main__":
    raise SystemExit(main())
```
