## Supplementary Information section 4 code base for "Hypothesis: A modern human range expansion ∼300,000 years ago had a critical role in Neandertal origins": den_threshold_test.py

```
#!/usr/bin/env python3
"""Does a separate Denisovan threshold buy anything the Neandertal one cannot?

THE QUESTION
The catalog rule is

    case     nea_hi(p) <= x   and   den_lo(p) >  y
    control  den_hi(p) <= x   and   nea_lo(p) >  y      (archaics swapped)

and the operating point leaves y at 400 ka, where it almost never binds. That
looks odd: requiring the Denisovan coalescence to be deep ought to help.

WHY THE NAIVE COMPARISON IS WRONG
Raising y removes windows, so it lowers the ratio for the same reason that
raising x does -- the catalog gets smaller and what remains is more extreme.
Any ratio improvement from y therefore has to be judged against the ratio you
would have got by tightening x to the same catalog size. Only a curve that
lies BELOW the x-only curve in the (size, ratio) plane is evidence that y
carries independent information.

WHAT IS COMPUTED
Three sweeps from the operating point, one parameter at a time, plus a
matched-size comparison of the y sweep against a densely sampled x sweep.

Usage:
  python3 scripts/den_threshold_test.py
"""

from __future__ import annotations

import argparse
import os
import sys

sys.path.insert(0, os.path.dirname(os.path.abspath(__file__)))
from approx_percentiles import default_coord, load  # noqa: E402
from fig_s51_sensitivity import counts, site_percentiles, wilson  # noqa: E402

# The operating point actually used for the catalog.
BASE_X, BASE_Y, BASE_CUT = 625.0, 400.0, 50.0

# Dense x sweep, used both as its own curve and as the matched-size reference.
X_DENSE = list(range(500, 901, 5))
Y_SWEEP = [400, 450, 500, 550, 600, 625, 650, 700, 750, 800, 900, 1000, 1200]
CUT_SWEEP = [50, 55, 60, 65, 70, 75, 80, 85, 90, 95]


def main():
    ap = argparse.ArgumentParser()
    ap.add_argument("--scan", default="hubisz_tmrca_5kb_8afr.tsv")
    ap.add_argument("-o", "--out",
                    default="catalog_annotation/den_threshold_test.tsv")
    a = ap.parse_args()

    recs = load(a.scan)
    coord, desc = default_coord(recs)
    recs = [d for d in recs if d["chrom"].replace("chr", "") not in ("X", "23")]
    sites = site_percentiles(recs, coord)
    print(f"time coordinate: {desc}")
    print(f"{len(sites):,} autosomal windows usable\n")

    bc, bk = counts(sites, coord, BASE_X, BASE_Y, BASE_CUT)
    print(f"operating point  x={BASE_X:.0f} y={BASE_Y:.0f} cut={BASE_CUT:.0f}:"
          f"  n={bc}  ctrl={bk}  ratio={bk/bc:.4f}\n")

    # ---- the x-only reference curve, densely sampled ----
    xcurve = []
    for x in X_DENSE:
        c, k = counts(sites, coord, x, BASE_Y, BASE_CUT)
        if c >= 5:
            xcurve.append((c, k / c, x))
    xcurve.sort()
    print(f"x sweep (y off, cut {BASE_CUT:.0f}): {len(xcurve)} points, "
          f"sizes {xcurve[0][0]}-{xcurve[-1][0]}\n")

    def matched(n_target):
        """Ratio the x-only rule achieves at the closest catalog size."""
        best = min(xcurve, key=lambda t: abs(t[0] - n_target))
        return best  # (n, ratio, x)

    # ---- the y sweep, and the matched-size verdict ----
    rows = []
    hdr = (f"{'y (ka)':>7} {'n':>6} {'ctrl':>5} {'ratio':>7} {'95% CI':>15} "
           f"{'x-only n':>9} {'x-only r':>9} {'at x':>5} {'verdict':>9}")
    print("VARYING THE DENISOVAN THRESHOLD, x and cut held at the operating point")
    print(hdr)
    print("-" * len(hdr))
    for y in Y_SWEEP:
        c, k = counts(sites, coord, BASE_X, y, BASE_CUT)
        if c < 5:
            print(f"{y:>7} {c:>6} {k:>5}   catalog too small to compare")
            continue
        r = k / c
        lo, hi = wilson(k, c)
        mn, mr, mx = matched(c)
        # y helps only if it beats the x-only curve at the same size, by more
        # than the noise on this ratio.
        verdict = verdict_for(lo, hi, mr)
        print(f"{y:>7} {c:>6} {k:>5} {r:>7.4f} "
              f"[{lo:>6.3f},{hi:>6.3f}] {mn:>9} {mr:>9.4f} {mx:>5} "
              f"{verdict:>9}")
        rows.append({"sweep": "den_threshold", "value": y, "n": c, "ctrl": k,
                     "ratio": round(r, 5), "matched_n": mn,
                     "matched_ratio": round(mr, 5), "matched_x": mx,
                     "verdict": verdict})

    # ---- the cut sweep, same matched-size treatment ----
    print("\nVARYING THE PERCENTILE CUT, x and y held at the operating point")
    print(hdr)
    print("-" * len(hdr))
    for cut in CUT_SWEEP:
        c, k = counts(sites, coord, BASE_X, BASE_Y, cut)
        if c < 5:
            continue
        r = k / c
        lo, hi = wilson(k, c)
        mn, mr, mx = matched(c)
        verdict = verdict_for(lo, hi, mr)
        print(f"{cut:>7} {c:>6} {k:>5} {r:>7.4f} "
              f"[{lo:>6.3f},{hi:>6.3f}] {mn:>9} {mr:>9.4f} {mx:>5} "
              f"{verdict:>9}")
        rows.append({"sweep": "percentile_cut", "value": cut, "n": c, "ctrl": k,
                     "ratio": round(r, 5), "matched_n": mn,
                     "matched_ratio": round(mr, 5), "matched_x": mx,
                     "verdict": verdict})

    # ---- is there ANY (x, y) pair that beats x alone at matched size? ----
    print("\nJOINT SEARCH: best (x, y) at each size band against x alone")
    print(f"{'size band':>14} {'best x':>7} {'best y':>7} {'n':>6} "
          f"{'ratio':>7} {'x-only r':>9} {'gain':>7}")
    print("-" * 64)
    joint = []
    for x in range(500, 901, 25):
        for y in range(400, 1301, 50):
            c, k = counts(sites, coord, x, y, BASE_CUT)
            if c >= 20:
                joint.append((c, k / c, x, y))
    for lo_n, hi_n in ((20, 50), (50, 100), (100, 200), (200, 400),
                       (400, 800), (800, 2000)):
        band = [t for t in joint if lo_n <= t[0] < hi_n]
        if not band:
            continue
        c, r, x, y = min(band, key=lambda t: t[1])
        mn, mr, mx = matched(c)
        print(f"{lo_n:>6}-{hi_n:<7} {x:>7} {y:>7} {c:>6} {r:>7.4f} "
              f"{mr:>9.4f} {mr - r:>+7.4f}")
        rows.append({"sweep": "joint_best", "value": f"{lo_n}-{hi_n}", "n": c,
                     "ctrl": int(round(r * c)), "ratio": round(r, 5),
                     "matched_n": mn, "matched_ratio": round(mr, 5),
                     "matched_x": x, "verdict": f"y={y}"})

    os.makedirs(os.path.dirname(a.out) or ".", exist_ok=True)
    keys = ["sweep", "value", "n", "ctrl", "ratio", "matched_n",
            "matched_ratio", "matched_x", "verdict"]
    with open(a.out, "w", encoding="utf-8") as f:
        f.write("\t".join(keys) + "\n")
        for r in rows:
            f.write("\t".join(str(r[k]) for k in keys) + "\n")
    print(f"\nwrote {a.out}")
    return 0


def verdict_for(lo, hi, matched_ratio):
    """Whether a rule beats the x-only curve at the same catalog size.

    The comparison is between two noisy ratios, so a bare point comparison
    would call a coin flip a result. The matched x-only ratio is treated as the
    reference and asked whether it falls outside the confidence interval of the
    rule under test: entirely above it means the rule genuinely helps, entirely
    below means it hurts, and overlap means the data cannot separate them.
    """
    if hi < matched_ratio:
        return "helps"
    if lo > matched_ratio:
        return "worse"
    return "ties"


if __name__ == "__main__":
    raise SystemExit(main())
```
