## Supplementary Information section 4 code base for "Hypothesis: A modern human range expansion ∼300,000 years ago had a critical role in Neandertal origins": discriminator_grid.py

```
#!/usr/bin/env python3
"""
Grid of discriminating conditions x modern-sweep thresholds for the sweep catalog.

Rows (5 confidence conditions). For a condition written q/(1-q), the archaic that
is supposed to JOIN the moderns must have its q-th posterior percentile below the
threshold, and the archaic that is supposed to stay OUT must have its (1-q)-th
percentile above it:

    condition      join-side test        outgroup-side test
    mean           mean      <= x        mean      >  x
    80/20          p80       <= x        p20       >  x
    90/10          p90       <= x        p10       >  x
    95/5           p95       <= x        p05       >  x
    97.5/2.5       p975      <= x        p025      >  x     <- stored, exact

Columns (13 thresholds): 500, 525, ..., 800 ka.

Each cell reports the number of Neandertal points (case: Neandertals join,
Denisova stays out) and the Denisova/Neandertal ratio, which estimates the false
discovery rate because the two archaics are sister lineages and present-day
Africans are therefore equidistant from both.

Percentiles other than 2.5 / 97.5 are reconstructed by approx_percentiles.py,
assuming the posterior is Normal in ARGweaver's log-spaced time coordinate. The
97.5/2.5 row uses the stored values directly, so it is exact by construction.

The grid is computed separately for the autosomes and for the X chromosome
rather than pooling them, because the X has a smaller effective population
size and a different expected sweep density; pooling would mix two signals.
Each output row carries a chrom_set field of "autosomes", "X", or "all".

Usage:
  python3 scripts/discriminator_grid.py --scan hubisz_tmrca_5kb_8afr.tsv \
      -o catalog_annotation/discriminator_grid.tsv
"""

from __future__ import annotations

import argparse
import csv
import os

from approx_percentiles import default_coord, fit_site, load, z_of

# (label, join-side quantile, outgroup-side quantile); None => use the mean
CONDITIONS = [
    ("mean", None, None),
    ("80/20", 0.80, 0.20),
    ("90/10", 0.90, 0.10),
    ("95/5", 0.95, 0.05),
    ("97.5/2.5", 0.975, 0.025),
]

THRESHOLDS = list(range(500, 801, 25))


def main() -> int:
    ap = argparse.ArgumentParser()
    ap.add_argument("--scan", default="hubisz_tmrca_5kb_8afr.tsv")
    ap.add_argument("-o", "--out", default="catalog_annotation/discriminator_grid.tsv")
    args = ap.parse_args()

    recs = load(args.scan)
    for d in recs:
        c = d["chrom"].replace("chr", "")
        d["is_x"] = c in ("X", "23")

    coord, grid_desc = default_coord(recs)
    print(f"time coordinate: {grid_desc}")

    # Reconstruct every needed percentile once per site.
    quants = sorted({q for _, a, b in CONDITIONS for q in (a, b) if q is not None})
    zs = {q: z_of(q) for q in quants}
    for d in recs:
        for tag in ("nea", "den"):
            mu, sg = fit_site(coord, d[f"{tag}_p025"], d[f"{tag}_p975"])
            for q in quants:
                d[f"{tag}_q{q}"] = coord.to_t(mu + zs[q] * sg)

    def key(tag, q):
        return f"{tag}_mean" if q is None else f"{tag}_q{q}"

    SETS = [
        ("autosomes", [d for d in recs if not d["is_x"]]),
        ("X", [d for d in recs if d["is_x"]]),
        ("all", recs),
    ]
    for name, sub in SETS:
        print(f"{name:<10} grid points: {len(sub):,}")

    rows = []
    for set_name, sub in SETS:
        for label, qin, qout in CONDITIONS:
            for x in THRESHOLDS:
                nea = sum(1 for d in sub
                          if d[key("nea", qin)] <= x and d[key("den", qout)] > x)
                den = sum(1 for d in sub
                          if d[key("den", qin)] <= x and d[key("nea", qout)] > x)
                rows.append({
                    "chrom_set": set_name,
                    "condition": label,
                    "threshold_ka": x,
                    "nea_points": nea,
                    "den_points": den,
                    "den_over_nea": round(den / nea, 4) if nea else None,
                    "nea_Mb": round(nea * 5 / 1000, 3),
                })

    os.makedirs(os.path.dirname(args.out) or ".", exist_ok=True)
    with open(args.out, "w", newline="", encoding="utf-8") as f:
        w = csv.DictWriter(f, fieldnames=list(rows[0].keys()), delimiter="\t")
        w.writeheader()
        w.writerows(rows)

    def cell(set_name, label, x, field):
        return next(r[field] for r in rows
                    if r["chrom_set"] == set_name and r["condition"] == label
                    and r["threshold_ka"] == x)

    for set_name, _ in SETS:
        for field, title in (("nea_points", "Neandertal points (case count)"),
                             ("den_over_nea", "Denisova/Neandertal ratio (FDR estimate)")):
            print(f"\n===== [{set_name}] {title} =====")
            print(f"{'condition':<10}" + "".join(f"{x:>8}" for x in THRESHOLDS))
            for label, _, _ in CONDITIONS:
                cells = []
                for x in THRESHOLDS:
                    v = cell(set_name, label, x, field)
                    if v is None:
                        cells.append(f"{'-':>8}")
                    elif field == "nea_points":
                        cells.append(f"{v:>8}")
                    else:
                        cells.append(f"{v:>8.2f}")
                print(f"{label:<10}" + "".join(cells))

    print(f"\nwrote {args.out}")
    return 0


if __name__ == "__main__":
    raise SystemExit(main())
```
