## Supplementary Information section 4 code base for "Hypothesis: A modern human range expansion ∼300,000 years ago had a critical role in Neandertal origins": fetch_cadd_regions.py

```
#!/usr/bin/env python3
"""Pull CADD v1.6 for the screened regions into one local, tabix-indexed file.

annotate_functional_classes.py needs a CADD score at every candidate site, and
reading them one site at a time from the remote 80 GB whole_genome_SNVs.tsv.gz
is unusable: each fetch re-seeks into the file and costs about 11 seconds, so
the 7,597 sites of the 1 kb screen would take roughly a day. Fetching a whole
screen region at once costs about the same 1.5 seconds whether it returns one
row or thirty thousand, because the price is the seek and not the bytes. The
287 regions therefore come down in a few minutes, after which annotation reads
CADD off local disk.

The output is the same format as the source, so the annotator cannot tell the
difference; point it at the result with

  ANNOT_CADD_URL=k1/cadd_screen_regions.tsv.gz

The file only covers the screened regions, which is the point and also the one
trap: using it for any site outside those regions silently yields no CADD score
rather than an error. It is a cache for this screen, not a general CADD subset.
"""

import argparse
import csv
import os
import sys
import time


def main():
    ap = argparse.ArgumentParser()
    ap.add_argument("--regions", default="k1/screen_k1.padded.regions.tsv",
                    help="TSV with chrom/start/end; the screened territory")
    ap.add_argument("--cadd",
                    default="https://kircherlab.bihealth.org/download/CADD/"
                            "v1.6/GRCh37/whole_genome_SNVs.tsv.gz")
    ap.add_argument("-o", "--out", default="k1/cadd_screen_regions.tsv")
    args = ap.parse_args()

    import pysam

    with open(args.regions) as f:
        regs = [(r["chrom"], int(r["start"]), int(r["end"]))
                for r in csv.DictReader(f, delimiter="\t")]
    # Merge overlaps so no row is fetched or written twice; the screen regions
    # are padded and neighbouring loci can run into one another.
    by_chrom = {}
    for c, s, e in regs:
        by_chrom.setdefault(c, []).append([s, e])
    merged = []
    for c in sorted(by_chrom, key=lambda x: (len(x), x)):
        iv = sorted(by_chrom[c])
        out = [iv[0]]
        for s, e in iv[1:]:
            if s <= out[-1][1]:
                out[-1][1] = max(out[-1][1], e)
            else:
                out.append([s, e])
        merged += [(c, s, e) for s, e in out]
    span = sum(e - s for _, s, e in merged)
    print(f"{len(regs)} regions -> {len(merged)} merged, {span/1e6:.2f} Mb")

    tb = pysam.TabixFile(args.cadd)
    header = [l for l in tb.header]
    t0 = time.time()
    n = 0
    with open(args.out, "w") as out:
        for h in header:
            out.write(h + "\n")
        for i, (c, s, e) in enumerate(merged, 1):
            for line in tb.fetch(c, s, e):
                out.write(line + "\n")
                n += 1
            if i % 25 == 0 or i == len(merged):
                el = time.time() - t0
                print(f"  {i}/{len(merged)} regions  {n:,} rows  "
                      f"{el:.0f}s elapsed  eta {el/i*(len(merged)-i):.0f}s",
                      flush=True)

    print(f"compressing and indexing {args.out} ...")
    gz = args.out + ".gz"
    if os.path.exists(gz):
        os.remove(gz)
    pysam.tabix_compress(args.out, gz, force=True)
    # CADD is 1-based with chrom in col 1 and pos in col 2, and its header
    # lines start with '#'; preset="vcf" would mis-read it, so be explicit.
    pysam.tabix_index(gz, seq_col=0, start_col=1, end_col=1, meta_char="#",
                      force=True)
    os.remove(args.out)
    print(f"wrote {gz} ({os.path.getsize(gz)/1e6:.0f} MB) and {gz}.tbi")
    print(f"{n:,} CADD rows in {time.time()-t0:.0f}s")


if __name__ == "__main__":
    sys.exit(main())
```
