## Supplementary Information section 4 code base for "Hypothesis: A modern human range expansion ∼300,000 years ago had a critical role in Neandertal origins": fig_calibration.py

```
#!/usr/bin/env python3
"""
Figure for the calibration section of SI section 5.

Four panels, in the order the argument is made:

  (a) The mirrored control. Applying the identical test with the two archaics
      swapped counts how often the discriminator fires in a direction that
      cannot be a modern-human sweep. Case and control counts are shown
      against the crossing threshold.
  (b) The null is not symmetric. The background Neandertal-direction fraction
      theta0 sits above 0.5 at every percentile cut, because TMRCA(Afr+Nea)
      pools four archaic haplotypes from two Neandertals while TMRCA(Afr+Den)
      pools two from one Denisovan, and the archaics differ in effective size
      and in their migration bands. P(real) is normalized to theta0, not 0.5.
  (c) Pooling stabilizes the estimate. Raw per-cell skew against the value
      fitted by the joint binomial logistic surface; small cells are dragged
      to the trend, large cells are left alone.
  (d) The resulting per-site probabilities. Loci retained against the P(real)
      cutoff for autosomes and for the X, which never reaches 0.5.

Usage:
  PYTHONPATH=scripts:.pylibs MPLCONFIGDIR=/tmp/mpl \
      python3 scripts/fig_calibration.py -o figures/calibration.png
"""

from __future__ import annotations

import argparse
import csv
import math
import os
from collections import defaultdict

import matplotlib
matplotlib.use("Agg")
import matplotlib.pyplot as plt
import numpy as np

from approx_percentiles import default_coord, fit_site, load, z_of
from fit_fdr_surface import PCUTS, U_EDGES, V_EDGES, binof, design, irls

np.seterr(all="ignore")

plt.rcParams.update({
    "font.family": "serif",
    "font.serif": ["Times New Roman", "Times", "DejaVu Serif"],
    "font.size": 8,
    "axes.labelsize": 8,
    "axes.titlesize": 8.5,
    "xtick.labelsize": 7,
    "ytick.labelsize": 7,
    "legend.fontsize": 7,
    "axes.linewidth": 0.6,
    "xtick.major.width": 0.6,
    "ytick.major.width": 0.6,
})

CASE = "#B2182B"
CTRL = "#2166AC"
GREY = "#666666"


def wilson(k, n, z=1.959963985):
    if n == 0:
        return (0.0, 0.0)
    p = k / n
    d = 1 + z * z / n
    c = (p + z * z / (2 * n)) / d
    h = z * math.sqrt(p * (1 - p) / n + z * z / (4 * n * n)) / d
    return (max(0.0, c - h), min(1.0, c + h))


def isx(c):
    return c.replace("chr", "") in ("X", "23")


def main() -> int:
    ap = argparse.ArgumentParser()
    ap.add_argument("--scan", default="hubisz_tmrca_5kb_8afr.tsv")
    ap.add_argument("--posterior", default="catalog_annotation/site_posterior.tsv")
    ap.add_argument("-o", "--out", default="figures/calibration.png")
    args = ap.parse_args()

    recs = load(args.scan)
    coord, _ = default_coord(recs)
    for d in recs:
        for tag in ("nea", "den"):
            mu, sg = fit_site(coord, d[f"{tag}_p025"], d[f"{tag}_p975"])
            for p in PCUTS:
                d[f"{tag}_hi{p}"] = coord.to_t(mu + z_of(p) * sg)
                d[f"{tag}_lo{p}"] = coord.to_t(mu + z_of(1 - p) * sg)
    auto = [d for d in recs if not isx(d["chrom"])]
    print(f"{len(auto):,} autosomal grid points")

    fig, axes = plt.subplots(2, 2, figsize=(6.9, 5.2))
    (axA, axB), (axC, axD) = axes

    # ---- (a) case and control counts against the crossing threshold ----
    xs = list(range(500, 826, 25))
    ncase, nctrl = [], []
    for x in xs:
        a = sum(1 for d in auto if d["nea_hi0.8"] <= x and d["den_lo0.8"] > x)
        b = sum(1 for d in auto if d["den_hi0.8"] <= x and d["nea_lo0.8"] > x)
        ncase.append(a)
        nctrl.append(b)
    axA.plot(xs, ncase, "-o", color=CASE, ms=2.6, lw=1.2,
             label="Neandertal direction (case)")
    axA.plot(xs, nctrl, "-s", color=CTRL, ms=2.6, lw=1.2,
             label="Denisova direction (control)")
    axA.set_yscale("log")
    axA.set_xlabel("crossing threshold (ka)")
    axA.set_ylabel("grid points")
    axA.set_title("(a) the mirrored control, 80/20 cut", loc="left")
    axA.legend(frameon=False, loc="upper left")
    axA.grid(alpha=0.25, lw=0.4)

    # ---- cells, per cut, for (b) and (c) ----
    rows = []
    theta0 = {}
    for p in PCUTS:
        cells = defaultdict(lambda: [0, 0])
        for d in auto:
            iu, iv = binof(d[f"nea_hi{p}"], U_EDGES), binof(d[f"den_lo{p}"], V_EDGES)
            if iu is not None and iv is not None:
                cells[(iu, iv)][0] += 1
            ju, jv = binof(d[f"den_hi{p}"], U_EDGES), binof(d[f"nea_lo{p}"], V_EDGES)
            if ju is not None and jv is not None:
                cells[(ju, jv)][1] += 1
        a = sum(v[0] for v in cells.values())
        b = sum(v[1] for v in cells.values())
        theta0[p] = (a, b)
        for (iu, iv), (kn, kd) in cells.items():
            rows.append({"p": p,
                         "u": (U_EDGES[iu] + U_EDGES[iu + 1]) / 2.0,
                         "v": (V_EDGES[iv] + V_EDGES[iv + 1]) / 2.0,
                         "nea": kn, "n": kn + kd})

    # ---- (b) the null skew theta0 per cut ----
    ps = list(PCUTS)
    th = [theta0[p][0] / (theta0[p][0] + theta0[p][1]) for p in ps]
    lo, hi = zip(*[wilson(theta0[p][0], theta0[p][0] + theta0[p][1]) for p in ps])
    xpos = np.arange(len(ps))
    axB.errorbar(xpos, th,
                 yerr=[np.array(th) - np.array(lo), np.array(hi) - np.array(th)],
                 fmt="o", color=CASE, ms=3.5, lw=1.0, capsize=2.2)
    axB.axhline(0.5, color=GREY, ls="--", lw=0.9)
    axB.set_ylim(0.488, 0.638)
    axB.text(0.97, 0.502, "symmetric null assumed by the mirror",
             transform=axB.get_yaxis_transform(), va="bottom", ha="right",
             color=GREY, fontsize=6.5)
    axB.set_xticks(xpos)
    axB.set_xticklabels([f"{int(p*100) if p != 0.975 else 97.5}" for p in ps])
    axB.set_xlabel("percentile cut (%)")
    axB.set_ylabel(r"background skew $\theta_0$")
    axB.set_title(r"(b) the null is not 0.5", loc="left")
    axB.grid(alpha=0.25, lw=0.4)

    # ---- (c) raw versus fitted cell skew ----
    u = np.array([r["u"] for r in rows]) / 100.0
    v = np.array([r["v"] for r in rows]) / 100.0
    pp = np.array([r["p"] for r in rows])
    k = np.array([r["nea"] for r in rows], float)
    n = np.array([r["n"] for r in rows], float)
    beta, _ = irls(design(u, v, pp), k, n)
    fitted = 1 / (1 + np.exp(-np.clip(design(u, v, pp) @ beta, -30, 30)))
    raw = k / n
    sc = axC.scatter(fitted, raw, s=np.clip(n / 4, 1.5, 60),
                     c=np.log10(n), cmap="viridis", alpha=0.65, linewidths=0)
    axC.plot([0, 1], [0, 1], color=GREY, ls="--", lw=0.9)
    cb = fig.colorbar(sc, ax=axC, pad=0.02, fraction=0.046)
    cb.set_label(r"$\log_{10}$ sites in cell", fontsize=6.5)
    cb.ax.tick_params(labelsize=6)
    axC.set_xlabel("fitted skew (joint surface)")
    axC.set_ylabel("raw skew in cell")
    axC.set_title("(c) pooling stabilizes small cells", loc="left")
    axC.grid(alpha=0.25, lw=0.4)

    # ---- (d) loci retained against the P(real) cutoff ----
    pr = {"autosome": [], "X": []}
    with open(args.posterior, encoding="utf-8") as f:
        for r in csv.DictReader(f, delimiter="\t"):
            pr["autosome" if r["chrom_set"] == "autosome" else "X"].append(
                float(r["p_real"]))
    cuts = np.linspace(0.0, 0.98, 60)
    for key, col, lab in (("autosome", CASE, "autosomes"), ("X", CTRL, "X chromosome")):
        arr = np.array(pr[key])
        cnt = np.array([(arr >= c).sum() for c in cuts], float)
        # break the line where the count hits zero rather than flooring it,
        # so the X curve ends where the X runs out of sites
        axD.plot(cuts, np.where(cnt > 0, cnt, np.nan), color=col, lw=1.3, label=lab)
        last = cuts[cnt > 0][-1] if (cnt > 0).any() else None
        if key == "X" and last is not None:
            axD.plot([last], [cnt[cnt > 0][-1]], "o", color=col, ms=3.5)
            axD.annotate(f"max {arr.max():.2f}", xy=(last, cnt[cnt > 0][-1]),
                         xytext=(last + 0.04, cnt[cnt > 0][-1] * 3.2),
                         fontsize=6.5, color=col,
                         arrowprops=dict(arrowstyle="-", color=col, lw=0.6))
    axD.set_yscale("log")
    axD.axvline(0.5, color=GREY, ls=":", lw=0.9)
    axD.text(0.51, 0.94, "catalog cutoff", transform=axD.get_xaxis_transform(),
             fontsize=6.5, color=GREY, va="top")
    axD.set_xlabel(r"$P(\mathrm{real})$ cutoff")
    axD.set_ylabel("grid points retained")
    axD.set_title(r"(d) the X never reaches 0.5", loc="left")
    axD.legend(frameon=False, loc="upper right")
    axD.grid(alpha=0.25, lw=0.4)

    fig.tight_layout(pad=0.7, w_pad=1.4, h_pad=1.2)
    os.makedirs(os.path.dirname(args.out) or ".", exist_ok=True)
    fig.savefig(args.out, dpi=400)
    fig.savefig(os.path.splitext(args.out)[0] + ".pdf")
    print("wrote", args.out)

    print("\ntheta0 by cut:")
    for p in ps:
        a, b = theta0[p]
        t = a / (a + b)
        l, h = wilson(a, a + b)
        print(f"  p={p:<6} Nea {a:>6,}  Den {b:>6,}  theta0 {t:.4f} "
              f"[{l:.4f}, {h:.4f}]  odds {t/(1-t):.3f}")
    return 0


if __name__ == "__main__":
    raise SystemExit(main())
```
