## Supplementary Information section 4 code base for "Hypothesis: A modern human range expansion ∼300,000 years ago had a critical role in Neandertal origins": fig_locus_zoom.py

```
#!/usr/bin/env python3
"""Per-locus scan figures: P(real), TMRCA traces, grid points, sites, genes.

Each figure answers one question: does this locus look like a sweep shared by
modern humans and Neandertals but not Denisovans, and how far does the signal
extend? So the two traces that matter are TAfr+Nea and TAfr+Den, each against
its own genome-wide average, and the thing that separates signal from noise is
the gap between them. Where the modern-Neandertal coalescence drops toward
600 ka while the modern-Denisovan coalescence stays near 2 Ma, the locus is a
candidate; where the two traces sit together on their dotted baselines, it is
not.

P(real) is on the left because it is the quantity the reader is being asked to
judge; the TMRCAs that produce it are on the right.

WINDOW CHOICE
Three conditions, unioned, so the window can only grow:
  * outward from the catalog locus while P(real) stays at or above 0.2;
  * further left until TAfr+Nea recovers above 1500 ka, so the figure always
    shows the Neandertal trace climbing back to something near baseline
    rather than being cropped mid-signal;
  * a 20 kb margin beyond whichever bound wins.
The recovery condition is applied on the left only, as asked, which is why
some figures are visibly lopsided: at FADS the left edge runs out to where the
Neandertal trace clears 1500 ka while the right edge stops at the P(real)
bound. Extending both sides is a one-line change if the asymmetry grates.

P(real) has a floor near 0.057 rather than 0 because almost every grid point
in the genome lies outside the fitted surface's support and is scored at its
boundary. The floor is the surface's estimate for an ordinary grid point, so
it is the right visual zero, but it should not be read as "this position has a
6% chance of being a real sweep".
"""

import argparse
import csv
import gzip
import os
import re
import sys
from collections import defaultdict

import matplotlib
matplotlib.use("Agg")
import matplotlib.pyplot as plt  # noqa: E402
from matplotlib.patches import Rectangle  # noqa: E402
from matplotlib.lines import Line2D  # noqa: E402
from matplotlib.offsetbox import (  # noqa: E402
    AnchoredOffsetbox, HPacker, TextArea)
import bisect  # noqa: E402

# The five inputs are overridable from the environment so the figure can be
# regenerated against a rebuilt catalog without editing this file; unset, they
# are the published paths.
GRID = os.environ.get("ZOOM_GRID",
                      "catalog_annotation/preal_gridpoints.tsv")
REGIONS = os.environ.get(
    "ZOOM_REGIONS",
    "catalog_annotation/catalog_final_median_x625_y700_auto_annot.regions.tsv")
CAT_SITES = os.environ.get(
    "ZOOM_CAT_SITES",
    "catalog_annotation/catalog_final_median_x625_y700_auto.sites.tsv")
# The screen rerun over five Neandertals and two Denisovans, functionally
# annotated. The earlier files came from a two-Neandertal panel over a narrower
# region set, so both the tick marks and which sites are starred change.
SITES = os.environ.get("ZOOM_SITES",
                       "catalog_annotation/fd_af99.functional.tsv")
# make_functional_table.py's own output, so the stars cannot drift from the
# table in the note. This used to point at a second copy that was left behind
# by an earlier run and went stale, still starring the synonymous sites after
# they were dropped from Table S4.2.
TABLE2 = os.environ.get("ZOOM_TABLE2",
                        "catalog_annotation/table_functional_sites.tsv")
GTF = "refdata/gencode.v19.annotation.gtf.gz"

C_NEA = "#1f4e9c"     # modern + Neandertal
C_DEN = "#c0392b"     # modern + Denisovan
C_PR = "#2e7d32"      # P(real)
STEP = 5000
PR_CUT = 0.20
NEA_RECOVER = 1500.0  # ka; left edge runs out until the trace clears this
MARGIN = 20000

# --- comparator -------------------------------------------------------------
# Default traces are the subset TMRCAs T_Afr+Nea and T_Afr+Den, the quantities
# the catalog rule is written on. A subset TMRCA is a maximum over the set, so
# it cannot fall below T_Afr and is dragged down wherever the African tree is
# shallow: genome-wide corr(T_Afr, T_Afr+Den) is 0.82, and at 17.7% of grid
# points T_Afr+Den equals T_Afr exactly, the Denisovan sitting inside the
# African clade so the comparator adds nothing to the African tree.
#
# --pairwise switches both traces to mean pairwise split times from
# hubisz_pairwise_split_scan.py: T_mean(Afr,Den) over 16 x 2 = 32 haplotype
# pairs and T_mean(Afr,Nea) over 16 x 4 = 64. Both traces are switched, not
# just the Denisovan one, because a mean against a maximum is not a like
# comparison. Window bounds and P(real) still come from the subset grid, so a
# panel covers exactly the same span in either mode and the two versions of the
# figure can be read side by side.
#
# --pairwise-den-only switches the Denisovan trace alone, keeping T_Afr+Nea on
# the blue line. The two traces are then not the same kind of statistic, one a
# maximum over a set and the other a mean over pairs, so their difference at a
# point carries no interpretation and only the shape of each against its own
# genome average can be read.
LAB_NEA = r"$T_{\mathrm{Afr+Nea}}$"
LAB_DEN = r"$T_{\mathrm{Afr+Den}}$"
PW_COLS = ("AfrNea", "AfrDen")
PW_DEN_ONLY = False


def load_grid():
    g = defaultdict(dict)
    with open(GRID, encoding="utf-8") as f:
        for r in csv.DictReader(f, delimiter="\t"):
            g[r["chrom"]][int(r["pos"])] = (
                float(r["nea_mean_ka"]), float(r["den_mean_ka"]),
                float(r["preal"]))
    return g


def load_pairwise(path):
    """chrom -> pos -> (T_mean(Afr,Nea), T_mean(Afr,Den)), in ka.

    Rows whose requested pair is NA, which happens when a tree is missing a
    sample, are skipped rather than filled, so a gap shows as a gap.
    """
    g = defaultdict(dict)
    a, b = PW_COLS
    with open(path, encoding="utf-8") as f:
        for r in csv.DictReader(f, delimiter="\t"):
            try:
                v = (float(r[f"{a}_mean_kya"]), float(r[f"{b}_mean_kya"]))
            except (KeyError, ValueError):
                continue
            g[str(int(r["chrom"]))].setdefault(int(r["pos"]), v)
    return g


def nea_distribution(pw=None):
    """Sorted genome-wide Neandertal trace, so a tick can be given a percentile."""
    if pw is not None and not PW_DEN_ONLY:
        vals = [v[0] for c in pw for v in pw[c].values()]
    else:
        vals = []
        with open(GRID, encoding="utf-8") as f:
            for r in csv.DictReader(f, delimiter="\t"):
                vals.append(float(r["nea_mean_ka"]))
    vals.sort()
    return vals


def genome_means(pw=None):
    if pw is not None and not PW_DEN_ONLY:
        vs = [v for c in pw for v in pw[c].values()]
        return (sum(v[0] for v in vs) / len(vs),
                sum(v[1] for v in vs) / len(vs))
    n = d = k = 0
    with open(GRID, encoding="utf-8") as f:
        for r in csv.DictReader(f, delimiter="\t"):
            n += float(r["nea_mean_ka"])
            d += float(r["den_mean_ka"])
            k += 1
    if pw is None:
        return n / k, d / k
    # Den-only: each average must come from the scan its trace is drawn from.
    vs = [v for c in pw for v in pw[c].values()]
    return n / k, sum(v[1] for v in vs) / len(vs)


def load_regions():
    out = defaultdict(list)
    with open(REGIONS, encoding="utf-8") as f:
        for r in csv.DictReader(f, delimiter="\t"):
            out[r["chrom"].replace("chr", "")].append(
                (int(r["start"]), int(r["end"])))
    for c in out:
        out[c].sort()
    return out


def load_catalog_points():
    """The individual 5 kb grid points that passed the rule."""
    out = defaultdict(set)
    with open(CAT_SITES, encoding="utf-8") as f:
        for r in csv.DictReader(f, delimiter="\t"):
            out[r["chrom"].replace("chr", "")].add(int(r["pos"]))
    return out


def load_sites():
    """Fixed-derived sites, flagged against the main-text table.

    "High functional significance" means the site made Table S4.2, not merely
    that it fell in some qualifying annotation class. The two differ a lot:
    the annotation classes are broad enough to flag 10 of the 13 sites around
    FADS, none of which is in the table, so using the class flag would put
    stars on the figure that the text does not support.
    """
    hi = set()
    if os.path.exists(TABLE2):
        with open(TABLE2, encoding="utf-8") as f:
            for r in csv.DictReader(f, delimiter="\t"):
                c, mb = r["locus"].split(":")
                hi.add((c, round(float(mb), 3)))
    out = defaultdict(list)
    with open(SITES, encoding="utf-8") as f:
        for r in csv.DictReader(f, delimiter="\t"):
            pos = int(r["pos"])
            out[r["chrom"]].append({
                "pos": pos, "gene": r.get("gene", ""),
                "classes": r.get("classes", ""),
                "tabled": (r["chrom"], round(pos / 1e6, 3)) in hi,
            })
    return out, hi


def window(grid, chrom, start, end):
    pos = sorted(grid[chrom])
    lo = hi = None
    for p in pos:
        if start <= p <= end:
            lo = p if lo is None else lo
            hi = p
    if lo is None:
        sys.exit(f"no grid points in chr{chrom}:{start}-{end}")
    while grid[chrom].get(lo - STEP, (0, 0, 0))[2] >= PR_CUT:
        lo -= STEP
    while grid[chrom].get(hi + STEP, (0, 0, 0))[2] >= PR_CUT:
        hi += STEP
    # Then run both edges outward while TAfr+Nea is still depressed, stopping
    # on the first grid point that clears 1500 ka, which is included so the
    # reader can see the trace recover. Both sides are chosen the same way.
    for sign in (-1, 1):
        edge = lo if sign < 0 else hi
        guard = 0
        while guard < 400:
            nxt = grid[chrom].get(edge + sign * STEP)
            edge += sign * STEP
            if nxt is None or nxt[0] > NEA_RECOVER:
                break
            guard += 1
        if sign < 0:
            lo = edge
        else:
            hi = edge
    return lo, hi


def apply_adjust(grid, chrom, lo, hi, adj):
    """Hand tweaks on top of the automatic window.

    left:N pulls the left edge in by N grid points (negative pushes it out),
    right:N pulls the right edge in by N. The automatic rule sets both edges
    from where TAfr+Nea recovers, so these are only for overriding it.
    """
    if not adj:
        return lo, hi
    lo += int(adj.get("left", 0)) * STEP
    hi -= int(adj.get("right", 0)) * STEP
    return lo, hi


def load_genes(chrom, lo, hi):
    want = f"chr{chrom}"
    genes, exons, cds = {}, defaultdict(list), defaultdict(list)
    with gzip.open(GTF, "rt", encoding="utf-8") as f:
        for line in f:
            if line[0] == "#":
                continue
            p = line.split("\t", 9)
            if p[0] != want or p[2] not in ("gene", "exon", "CDS"):
                continue
            s, e = int(p[3]), int(p[4])
            if e < lo or s > hi:
                continue
            at = p[8]
            name = re.search(r'gene_name "([^"]+)"', at)
            if not name:
                continue
            name = name.group(1)
            if p[2] == "gene":
                gt = re.search(r'gene_type "([^"]+)"', at)
                genes[name] = {"start": s, "end": e, "strand": p[6],
                               "type": gt.group(1) if gt else ""}
            elif '"basic"' in at:
                (cds if p[2] == "CDS" else exons)[name].append((s, e))

    def merge(iv):
        out, cur = [], None
        for s, e in sorted(iv):
            if cur and s <= cur[1] + 1:
                cur = (cur[0], max(cur[1], e))
            else:
                if cur:
                    out.append(cur)
                cur = (s, e)
        if cur:
            out.append(cur)
        return out

    for n in genes:
        genes[n]["exons"] = merge(exons.get(n, []))
        genes[n]["cds"] = merge(cds.get(n, []))
    return genes


def stack(genes, lo, hi):
    span = hi - lo
    order = sorted(genes.items(), key=lambda kv: kv[1]["start"])
    rows = []
    for name, g in order:
        pad = 0.10 * span + len(name) * 0.010 * span
        a, b = g["start"] - pad, g["end"] + pad
        for i, occ in enumerate(rows):
            if all(b < s or a > e for s, e in occ):
                occ.append((a, b))
                g["row"] = i
                break
        else:
            rows.append([(a, b)])
            g["row"] = len(rows) - 1
    return order, max(1, len(rows))


def panel_label(ax, letter, display, span):
    """(a) in bold, gene symbols italic, span in brackets, panel top left.

    Assembled from separate text boxes rather than mathtext because mathtext
    leaves digits upright, which left symbols like FADS1/2 only half italic.

    A label carrying no * markers is a bare gene symbol and is italicized
    whole. Only multi-gene titles need explicit markers, to separate the
    symbols from the connecting words, so before this the panels named for a
    single gene -- MED12L, SPATA31D1, ARHGAP32 -- fell through as segment 0
    and printed upright while their multi-gene neighbours came out italic.
    """
    if "*" not in display:
        display = f"*{display}*"
    kids = [TextArea(f"({letter})",
                     textprops=dict(fontsize=13.5, fontweight="bold",
                                    color="black"))]
    for i, seg in enumerate(display.split("*")):
        seg = seg.strip()
        if not seg:
            continue
        kids.append(TextArea(seg, textprops=dict(
            fontsize=12, color="black",
            style="italic" if i % 2 else "normal")))
    kids.append(TextArea(f"({span})",
                         textprops=dict(fontsize=12, color="black")))
    ax.add_artist(AnchoredOffsetbox(
        loc="lower left", pad=0, frameon=False, borderpad=0,
        child=HPacker(children=kids, align="baseline", pad=0, sep=5),
        bbox_to_anchor=(0.0, 1.01), bbox_transform=ax.transAxes))


def panel(fig, gs_row, name, chrom, lo, hi, data, gmean, label=None,
          display=None, legend=True, neadist=None):
    """Draw one locus into a 4-row sub-gridspec. Returns a stats tuple."""
    grid, regions, cat_pts, sites, pw = data
    pts = [(p, *grid[chrom][p]) for p in sorted(grid[chrom]) if lo <= p <= hi]
    xs = [p / 1e6 for p, _, _, _ in pts]
    pr = [v[3] for v in pts]
    if pw is None:
        txs = xs
        nea = [v[1] for v in pts]
        den = [v[2] for v in pts]
    else:
        # The pairwise scan is its own run, so a position present in the subset
        # grid can be absent here. Those are left out of the traces instead of
        # interpolated across, and P(real) keeps its own full x array.
        keep = [p for p, *_ in pts if p in pw.get(chrom, {})]
        txs = [p / 1e6 for p in keep]
        nea = ([grid[chrom][p][0] for p in keep] if PW_DEN_ONLY
               else [pw[chrom][p][0] for p in keep])
        den = [pw[chrom][p][1] for p in keep]
        if not keep:
            sys.exit(f"{name}: no pairwise values in chr{chrom}:{lo}-{hi}")
        if len(keep) < len(pts):
            sys.stderr.write(
                f"  {name}: {len(pts) - len(keep)} of {len(pts)} grid points "
                f"missing from the pairwise scan\n")

    genes = load_genes(chrom, lo, hi)
    order, nrow = stack(genes, lo, hi)
    site_list = [s for s in sites.get(chrom, []) if lo <= s["pos"] <= hi]

    sub = gs_row.subgridspec(4, 1, hspace=0.10,
                             height_ratios=[4.1, 0.40, 0.52, 0.34 * nrow + 0.3])
    A = fig.add_subplot(sub[0])
    W = fig.add_subplot(sub[1], sharex=A)
    S = fig.add_subplot(sub[2], sharex=A)
    G = fig.add_subplot(sub[3], sharex=A)

    # ---- left axis: P(real) -------------------------------------------------
    prpc = [100.0 * v for v in pr]
    A.plot(xs, prpc, color=C_PR, lw=1.7, zorder=4,
           label=r"$P(\mathrm{real})$")
    A.fill_between(xs, 0, prpc, color=C_PR, alpha=0.13, lw=0, zorder=1)
    A.set_ylabel(r"$P(\mathrm{real})$ (%)", fontsize=9, color=C_PR)
    A.tick_params(axis="y", labelsize=8, colors=C_PR)
    A.tick_params(axis="x", labelbottom=False)
    A.set_ylim(0, 102)
    A.grid(alpha=0.18, lw=0.5)

    # ---- right axis: the two TMRCAs ----------------------------------------
    A2 = A.twinx()
    A2.plot(txs, nea, "-o", color=C_NEA, ms=3.0, lw=1.6, zorder=5,
            label=LAB_NEA)
    A2.plot(txs, den, "-o", color=C_DEN, ms=3.0, lw=1.6, zorder=5,
            label=LAB_DEN)
    # In subset mode these are 1885 and 1889 ka and would overprint, so the
    # dash phases differ. In pairwise mode they are well separated.
    A2.axhline(gmean[0], color=C_NEA, ls=(0, (1, 3)), lw=1.4, zorder=2,
               label="genome average")
    A2.axhline(gmean[1], color=C_DEN, ls=(2, (1, 3)), lw=1.4, zorder=2)
    A2.tick_params(axis="y", labelsize=8)
    if neadist:
        def _pct(v, _):
            q = 100.0 * bisect.bisect_right(neadist, v) / len(neadist)
            d = 2 if q < 1 else 0
            return f"{v:.0f} ({q:.{d}f}%)"
        A2.yaxis.set_major_formatter(matplotlib.ticker.FuncFormatter(_pct))
        if pw is None or PW_DEN_ONLY:
            A2.yaxis.set_major_locator(matplotlib.ticker.FixedLocator(
                [500, 1000, 1500, 2000, 2500, 3000]))
        else:
            # Pairwise means live on a lower, narrower scale than subset
            # maxima, so the hand-picked subset ticks would fall off the axis.
            A2.yaxis.set_major_locator(matplotlib.ticker.MaxNLocator(
                nbins=6, steps=[1, 2, 2.5, 5, 10]))
        A2.set_ylabel("TMRCA in ka (genome percentile of "
                      f"{LAB_NEA})", fontsize=9, labelpad=9)
    else:
        A2.set_ylabel("TMRCA (ka)", fontsize=9)
    # Both genome averages must be inside the axis. With subset TMRCAs they are
    # 1885 and 1889 ka so either would do, but the pairwise means are well
    # separated and using only the Neandertal one clipped the Denisovan line.
    A2.set_ylim(min(min(nea), min(den)) * 0.55,
                max(max(nea), max(den), gmean[0], gmean[1]) * 1.08)

    if legend:
        hA, lA = A.get_legend_handles_labels()
        hB, lB = A2.get_legend_handles_labels()
        A.legend(hB + hA, lB + lA, fontsize=7.2, ncol=4, handlelength=1.8,
                 frameon=False, loc="lower left", bbox_to_anchor=(0.045, 1.13))
    span = f"chr{chrom}:{lo/1e6:.3f}-{hi/1e6:.3f} Mb"
    if label:
        panel_label(A, label, display or name, span)
    else:
        A.set_title(f"{(display or name).replace('*', '')}    {span}",
                    fontsize=9.2, loc="right", pad=15)

    # ---- grid-point track, replacing the shaded spans ----------------------
    incat = cat_pts.get(chrom, set())
    for p, _, _, _ in pts:
        member = p in incat
        W.add_patch(Rectangle((p / 1e6, 0.18), STEP / 1e6, 0.64,
                              facecolor="#c98f00" if member else "#f5f5f5",
                              edgecolor="#6f6f6f", lw=0.5,
                              zorder=3 if member else 2))
    W.set_ylim(0, 1)
    W.set_xlim(lo / 1e6, hi / 1e6)
    W.set_yticks([])
    for sp in ("top", "right", "left"):
        W.spines[sp].set_visible(False)
    W.tick_params(labelbottom=False)
    n_in = sum(1 for p, _, _, _ in pts if p in incat)
    W.set_ylabel(f"grid points\n{n_in}/{len(pts)} in catalog", fontsize=6.3,
                 rotation=0, ha="right", va="center", labelpad=14)

    # ---- fixed-derived sites ----------------------------------------------
    S.set_ylim(0, 1)
    S.set_yticks([])
    for sp in ("top", "right", "left"):
        S.spines[sp].set_visible(False)
    for s in site_list:
        if s["tabled"]:
            S.plot([s["pos"] / 1e6], [0.5], marker="*", ms=10,
                   color="#d4a017", mec="black", mew=0.5, zorder=4)
        else:
            S.plot([s["pos"] / 1e6], [0.5], marker="|", ms=8,
                   color="#5a5a5a", mew=1.3, zorder=3)
    S.tick_params(labelbottom=False)
    n_big = sum(1 for s in site_list if s["tabled"])
    tail = (f"\n{len(site_list)}"
            + (f", {n_big} in Table S4.2" if n_big else "")) if site_list \
        else "\nnone"
    S.set_ylabel("fixed derived" + tail, fontsize=6.3, rotation=0,
                 ha="right", va="center", labelpad=14)

    # ---- gene track --------------------------------------------------------
    for gname, g in order:
        y = nrow - 1 - g["row"] + 0.5
        pc = g["type"] == "protein_coding"
        col = "#123f7a" if pc else "#7a7a7a"
        a, b = max(g["start"], lo), min(g["end"], hi)
        G.plot([a / 1e6, b / 1e6], [y, y], color=col, lw=0.9, zorder=2)
        n_arrow = max(2, int((b - a) / (hi - lo) * 14))
        for k in range(n_arrow):
            xa = a + (b - a) * (k + 0.5) / n_arrow
            G.plot([xa / 1e6], [y], marker=">" if g["strand"] == "+" else "<",
                   ms=2.6, color=col, zorder=3)
        for iv, half in ((g["exons"], 0.15), (g["cds"], 0.26)):
            for s, e in iv:
                if e < lo or s > hi:
                    continue
                G.add_patch(Rectangle(
                    (max(s, lo) / 1e6, y - half),
                    (min(e, hi) - max(s, lo)) / 1e6, 2 * half,
                    facecolor=col, edgecolor="none",
                    zorder=4 if half > 0.2 else 3))
        G.text((a + b) / 2e6, y + 0.30, gname, fontsize=6.2, ha="center",
               va="bottom", style="italic", color=col, zorder=5)
    G.set_ylim(0, nrow)
    G.set_yticks([])
    for sp in ("top", "right", "left"):
        G.spines[sp].set_visible(False)
    G.set_xlabel(f"chromosome {chrom} position (Mb, hg19)", fontsize=8.5)
    G.tick_params(labelsize=8)
    G.set_xlim(lo / 1e6, hi / 1e6)
    G.ticklabel_format(axis="x", useOffset=False, style="plain")
    G.xaxis.set_major_formatter(
        matplotlib.ticker.FuncFormatter(lambda v, _: f"{v:.3f}"))
    G.xaxis.set_major_locator(matplotlib.ticker.MaxNLocator(6))
    return len(site_list), len(order), (hi - lo) / 1000.0, nrow


def main():
    ap = argparse.ArgumentParser()
    ap.add_argument("--locus", action="append", required=True,
                    help="name=chrom:start-end of the catalog locus, in bp")
    ap.add_argument("--outdir", default="figures")
    ap.add_argument("--combined",
                    help="also tile every locus into this single figure")
    ap.add_argument("--ncol", type=int, default=2)
    ap.add_argument("--panel-w", type=float, default=10.2,
                    help="width of one panel in the composite, inches")
    ap.add_argument("--panel-h", type=float, default=5.75,
                    help="height of one panel in the composite, inches; "
                         "lower values flatten the panels so more rows fit")
    ap.add_argument("--hspace", type=float, default=0.42,
                    help="vertical gap between rows of the composite")
    ap.add_argument("--adjust", action="append", default=[],
                    help="NAME=left:3,right:2  pull an edge in by N grid pts")
    ap.add_argument("--display", action="append", default=[],
                    help="NAME=*GENE* text  label, *...* marking italics")
    ap.add_argument("--pairwise", default=None,
                    help="collated hubisz_pairwise_split_scan.py output; plot "
                         "mean pairwise split times instead of subset TMRCAs")
    ap.add_argument("--pairwise-den-only", action="store_true",
                    help="with --pairwise, switch only the Denisovan trace and "
                         "keep T_Afr+Nea in blue")
    a = ap.parse_args()

    adjust = {}
    for spec in a.adjust:
        nm, rest = spec.split("=", 1)
        adjust[nm] = {k: int(v) for k, v in
                      (part.split(":") for part in rest.split(","))}
    display = dict(spec.split("=", 1) for spec in a.display)

    os.makedirs(a.outdir, exist_ok=True)
    global LAB_NEA, LAB_DEN, PW_DEN_ONLY
    pw = None
    if a.pairwise_den_only and not a.pairwise:
        sys.exit("--pairwise-den-only requires --pairwise")
    if a.pairwise:
        pw = load_pairwise(a.pairwise)
        PW_DEN_ONLY = a.pairwise_den_only
        if not PW_DEN_ONLY:
            LAB_NEA = r"$T_{\mathrm{mean}}(\mathrm{Afr},\mathrm{Nea})$"
        LAB_DEN = r"$T_{\mathrm{mean}}(\mathrm{Afr},\mathrm{Den})$"
        sys.stderr.write(
            f"pairwise comparator from {a.pairwise}: "
            f"{sum(len(v) for v in pw.values()):,} grid points\n")
    # Window bounds and P(real) always come from the subset grid, so a panel
    # spans the same interval whichever comparator is plotted.
    data = (load_grid(), load_regions(), load_catalog_points(),
            load_sites()[0], pw)
    gmean = genome_means(pw)
    neadist = nea_distribution(pw)
    kind = ("subset TMRCA" if pw is None else
            "subset Nea + mean pairwise Den" if PW_DEN_ONLY else "mean pairwise")
    sys.stderr.write(f"genome averages ({kind}): Nea {gmean[0]:.0f} ka, "
                     f"Den {gmean[1]:.0f} ka\n")

    specs = []
    for spec in a.locus:
        name, coord = spec.split("=", 1)
        chrom, rng = coord.split(":")
        s, e = (int(x) for x in rng.split("-"))
        lo, hi = window(data[0], chrom, s, e)
        lo, hi = apply_adjust(data[0], chrom, lo, hi, adjust.get(name))
        specs.append((name, chrom, lo, hi))

    for name, chrom, lo, hi in specs:
        fig = plt.figure(figsize=(8.4, 6.0))
        gs = fig.add_gridspec(1, 1)
        ns, ng, kb, _ = panel(fig, gs[0], name, chrom, lo, hi, data, gmean,
                              display=display.get(name), neadist=neadist)
        out = os.path.join(a.outdir, f"locus_{name}.png")
        fig.savefig(out, dpi=200, bbox_inches="tight")
        plt.close(fig)
        sys.stderr.write(f"{name}: chr{chrom}:{lo:,}-{hi:,} ({kb:.0f} kb), "
                         f"{ng} genes, {ns} sites -> {out}\n")

    if a.combined:
        ncol = a.ncol
        nrow = -(-len(specs) // ncol)
        # Wider and shorter than square panels would be, so the composite
        # fits the text width of a portrait page without overrunning it.
        # --panel-h flattens the panels further, which is what lets ten of
        # them onto one Letter page: at the default 5.75 a five-row figure is
        # taller than it is wide and has to be shrunk until the axis labels
        # are unreadable, where at 4.6 the whole thing is width-limited.
        pw, ph = a.panel_w, a.panel_h
        fig = plt.figure(figsize=(pw * ncol, ph * nrow))
        top = 1.0 - 1.7 / (ph * nrow)
        gs = fig.add_gridspec(nrow, ncol, hspace=a.hspace, wspace=0.38,
                              top=top)
        for i, (name, chrom, lo, hi) in enumerate(specs):
            panel(fig, gs[i // ncol, i % ncol], name, chrom, lo, hi, data,
                  gmean, label=chr(ord("a") + i),
                  display=display.get(name), legend=False, neadist=neadist)
        # one legend for all eight panels, spread across the page
        shared = [
            Line2D([], [], color=C_NEA, marker="o", ms=7, lw=2.6,
                   label=LAB_NEA),
            Line2D([], [], color=C_DEN, marker="o", ms=7, lw=2.6,
                   label=LAB_DEN),
            Line2D([], [], color="#555555", ls=(0, (1, 2.4)), lw=2.2,
                   label="genome average"),
            Line2D([], [], color=C_PR, lw=2.6,
                   label=r"$P(\mathrm{real})$")]
        fig.legend(handles=shared, loc="upper center", ncol=4, frameon=False,
                   bbox_to_anchor=(0.5, 1.0), fontsize=21, handlelength=2.4,
                   columnspacing=2.2, handletextpad=0.6, borderaxespad=0.4)
        fig.savefig(a.combined, dpi=170, bbox_inches="tight")
        plt.close(fig)
        sys.stderr.write(f"combined {len(specs)} panels -> {a.combined}\n")
    return 0


if __name__ == "__main__":
    sys.exit(main())
```
