## Supplementary Information section 4 code base for "Hypothesis: A modern human range expansion ∼300,000 years ago had a critical role in Neandertal origins": fig_s51_composite.py

```
#!/usr/bin/env python3
"""Figure S4.1, two panels.

(A) Case and control counts against the Neandertal threshold, with P(real) on
    the right axis. The counts and the ratio are the same information read
    two ways -- the ratio is literally the lower curve divided by the upper --
    so they belong on one pair of axes rather than two panels. What the panel
    shows is where the two directions separate: they track each other above
    about 700 ka, where the rule has no power, and diverge below it.

(B) Everything in the (size, purity) plane, which is the only plane in which
    rules of different shapes are comparable: any tightening buys purity by
    shrinking the catalog, so a rule is only better if it sits HIGHER at the
    SAME size. Purity is P(real) on a single left axis; the panel used to
    carry the control/case ratio on the left and P(real) on the right, which
    labelled one quantity twice on scales running opposite ways. Each parameter of the final rule is varied in turn from the
    operating point, and the fitted logistic surface is drawn on the same
    full-genome, cross-validated footing.

WHY EVERY SWEEP STARTS FROM THE SAME PLACE
Earlier drafts varied the threshold with the percentile cut at 80/20 and the
Denisovan threshold tied to the Neandertal one, which is not the rule the
catalog uses. Here all three sweeps pass through the operating point actually
adopted -- 625 ka on the Neandertal median, median cut, no Denisovan condition
-- so each curve answers "what would changing this one thing have bought?"

Usage:
  python3 scripts/fig_s51_composite.py -o catalog_annotation/figure_s5_1.png
"""

from __future__ import annotations

import argparse
import os
import sys
from collections import defaultdict

import numpy as np

sys.path.insert(0, os.path.dirname(os.path.abspath(__file__)))
from approx_percentiles import default_coord, load  # noqa: E402
from fig_s51_sensitivity import counts, site_percentiles, wilson  # noqa: E402
from fit_fdr_surface import PCUTS  # noqa: E402
from optimize_rule_by_size import NULL_RATIO, p_real  # noqa: E402

# The operating point the catalog actually uses. The Denisovan threshold is
# set above the Neandertal one, which forces the Denisovan lineage to be an
# outgroup to the African-plus-Neandertal clade. It is adopted for
# interpretability, not for purity: panel B shows it costs a little of both
# size and ratio, and the sweeps show no setting of it improves on leaving it
# off. 700 ka is the lower bound of the published modern-archaic split
# interval, so it is fixed by an external estimate rather than tuned here.
BASE_X, BASE_Y, BASE_CUT = 625.0, 700.0, 50.0

X_PANEL_A = [500, 525, 550, 575, 600, 625, 650, 675, 700, 725, 750, 800, 900]
X_SWEEP = [525, 550, 575, 600, 625, 650, 675, 700, 750, 800]
Y_SWEEP = [400, 550, 600, 650, 700, 800, 900, 1000, 1200, 1300]
CUT_SWEEP = [50, 55, 60, 65, 70, 75, 80, 85, 90]
SURFACE_TARGETS = [50, 75, 100, 150, 200, 300, 400, 550, 750, 1000, 1400]


def surface_cv(recs, coord, targets):
    """Held-out purity of the fitted surface, on a full-genome footing.

    The surface is fitted on one half of the chromosomes and used to rank the
    other half, both ways round, and the two halves are then pooled. Pooling
    matters: a curve scored on half the genome sits at half the catalog sizes
    and cannot be laid over the full-genome parameter sweeps, which is why the
    earlier draft needed a separate panel for it.

    The percentile cut is chosen per size AFTER seeing the held-out result,
    which favours the surface. That is deliberate -- the claim being tested is
    that the surface does not beat the rule, so the comparison is made on terms
    generous to the surface.
    """
    from fig_s51_model_curve import add_tails, fit_on, in_support
    from fit_fdr_surface import design

    add_tails(recs, coord)
    odd = [d for d in recs if int(d["chrom"].replace("chr", "")) % 2 == 1]
    even = [d for d in recs if int(d["chrom"].replace("chr", "")) % 2 == 0]

    # (fold, p) -> (sorted case scores desc, sorted control scores desc)
    scored = {}
    for name, train, test in (("odd->even", odd, even), ("even->odd", even, odd)):
        beta, ncell = fit_on(train)
        print(f"  surface {name}: fitted on {ncell:,} cells")
        for p in PCUTS:
            ec, et = [], []
            for d in test:
                u, v = d[f"nea_hi{p}"], d[f"den_lo{p}"]
                if in_support(u, v):
                    ec.append(design(np.array([u / 100.0]),
                                     np.array([v / 100.0]),
                                     np.array([p]))[0] @ beta)
                us, vs = d[f"den_hi{p}"], d[f"nea_lo{p}"]
                if in_support(us, vs):
                    et.append(design(np.array([us / 100.0]),
                                     np.array([vs / 100.0]),
                                     np.array([p]))[0] @ beta)
            scored[(name, p)] = (np.sort(np.array(ec))[::-1],
                                 np.sort(np.array(et))[::-1])

    pts = []
    for N in targets:
        half = N // 2
        best = None
        for p in PCUTS:
            tot_case = tot_ctrl = 0
            ok = True
            for name in ("odd->even", "even->odd"):
                ec, et = scored[(name, p)]
                if half > len(ec) or half == 0:
                    ok = False
                    break
                thresh = ec[half - 1]
                tot_case += half
                tot_ctrl += int(np.searchsorted(-et, -thresh, side="right"))
            if ok and (best is None or tot_ctrl / tot_case < best[1] / best[0]):
                best = (tot_case, tot_ctrl, p)
        if best:
            pts.append(best)
    return pts


def main():
    ap = argparse.ArgumentParser()
    ap.add_argument("--scan", default="hubisz_tmrca_5kb_8afr.tsv")
    ap.add_argument("-o", "--out", default="catalog_annotation/figure_s5_1.png")
    ap.add_argument("--no-surface", action="store_true",
                    help="skip the fitted surface (slow) while iterating")
    a = ap.parse_args()

    import matplotlib
    matplotlib.use("Agg")
    import matplotlib.pyplot as plt
    import matplotlib.ticker as mticker

    recs = load(a.scan)
    coord, desc = default_coord(recs)
    recs = [d for d in recs if d["chrom"].replace("chr", "") not in ("X", "23")]
    print(f"time coordinate: {desc}; {len(recs):,} autosomal windows")
    sites = site_percentiles(recs, coord)

    bc, bk = counts(sites, coord, BASE_X, BASE_Y, BASE_CUT)
    print(f"operating point: n={bc} ctrl={bk} ratio={bk/bc:.4f} "
          f"P(real)={p_real(bk/bc):.3f}\n")

    fig, (A, B) = plt.subplots(1, 2, figsize=(13.2, 5.4))

    # ================= Panel A: counts, with the ratio on the right =========
    xs, ca, ct = [], [], []
    for x in X_PANEL_A:
        c, k = counts(sites, coord, x, BASE_Y, BASE_CUT)
        xs.append(x); ca.append(c); ct.append(k)
    A.plot(xs, ca, "o-", color="#1f4e79", lw=2.0, ms=5,
           label="case: Neandertal direction")
    A.plot(xs, ct, "s-", color="#c00000", lw=1.8, ms=5,
           label="control: Denisovan direction")
    A.set_yscale("log")
    A.set_xlabel("Neandertal threshold x (ka)")
    A.set_ylabel("grid points retained (log scale)")
    A.axvline(BASE_X, color="#555", ls=":", lw=1.3)
    A.text(BASE_X + 14, 4600, f"catalog\n{BASE_X:.0f} ka", fontsize=8.5,
           color="#333", ha="left", va="center")
    A.grid(alpha=0.25, lw=0.5)

    A2 = A.twinx()
    ratios = [k / c if c else np.nan for c, k in zip(ca, ct)]
    # P(real) rather than the raw ratio, so the figure carries one purity
    # scale instead of two. P(real) falls as the ratio rises, so the Wilson
    # bounds swap sides on the way through.
    prs = [p_real(r) for r in ratios]
    lo = [p_real(wilson(k, c)[1]) for c, k in zip(ca, ct)]
    hi = [p_real(wilson(k, c)[0]) for c, k in zip(ca, ct)]
    A2.plot(xs, prs, "^--", color="#2e7d32", lw=1.8, ms=6,
            label="P(real) (right axis)")
    A2.fill_between(xs, lo, hi, color="#2e7d32", alpha=0.13, lw=0)
    A2.set_ylabel("P(real)", color="#2e7d32")
    # Headroom below the P(real) trace. It descends to about 0.1 at the right
    # edge, which on a 0-1 axis lands in the bottom corner where the legend now
    # sits; dropping the floor below zero lifts the whole trace clear of it.
    # Ticks stay on 0-1 so the axis still reads as a probability.
    A2.set_ylim(-0.34, 1.04)
    A2.set_yticks([0, 0.2, 0.4, 0.6, 0.8, 1.0])
    A2.tick_params(axis="y", colors="#2e7d32")

    # The counts climb the diagonal from lower left to upper right, so the
    # lower right is clear once P(real) has been lifted off the floor above.
    # Keeping the legend inside means neither panel overhangs the figure box,
    # so the whole thing can be set to full text width.
    h1, l1 = A.get_legend_handles_labels()
    h2, l2 = A2.get_legend_handles_labels()
    A.set_ylim(2.5, 2.5e4)
    A.legend(h1 + h2, l1 + l2, loc="lower right", ncol=1, frameon=True,
             framealpha=0.92, edgecolor="#cccccc", fontsize=8.6,
             handlelength=1.9, borderpad=0.6, labelspacing=0.45)
    A.set_title("A", loc="left", fontweight="bold")
    print("A: threshold   case  control    ratio   P(real)")
    for x, c, k, r in zip(xs, ca, ct, ratios):
        print(f"   {x:>8} {c:>6} {k:>7} {r:>8.3f} {p_real(r):>8.3f}")

    # ================= Panel B: the size-purity plane =======================
    sweeps = (
        ("vary Neandertal threshold (525-800 ka)",
         [(x, BASE_Y, BASE_CUT) for x in X_SWEEP], "#1f4e79", "o"),
        ("vary percentile cut (50-90)",
         [(BASE_X, BASE_Y, c) for c in CUT_SWEEP], "#c00000", "s"),
        ("vary Denisovan threshold (off-1300 ka)",
         [(BASE_X, y, BASE_CUT) for y in Y_SWEEP], "#ef8a00", "D"),
    )
    print(f"  (purity plotted as P(real) = 1 - ratio/{NULL_RATIO:.4f})")
    for label, specs, col, mk in sweeps:
        pts = []
        for sp in specs:
            c, k = counts(sites, coord, *sp)
            if c >= 5:
                pts.append((c, k))
        pts.sort()
        B.plot([c for c, _ in pts], [p_real(k / c) for c, k in pts], mk + "-",
               color=col, lw=1.8, ms=5, label=label, zorder=3)
        B.fill_between([c for c, _ in pts],
                       [p_real(wilson(k, c)[1]) for c, k in pts],
                       [p_real(wilson(k, c)[0]) for c, k in pts],
                       color=col, alpha=0.11, lw=0, zorder=2)
        print(f"\nB: {label}")
        for c, k in pts:
            print(f"   n={c:>5} ctrl={k:>5} ratio={k/c:.3f} "
                  f"P(real)={p_real(k/c):.3f}")

    if not a.no_surface:
        print("\nB: fitted logistic surface, two-fold cross-validated")
        recs2 = load(a.scan)
        recs2 = [d for d in recs2
                 if d["chrom"].replace("chr", "") not in ("X", "23")]
        spts = surface_cv(recs2, coord, SURFACE_TARGETS)
        if spts:
            B.plot([c for c, _, _ in spts], [p_real(k / c) for c, k, _ in spts],
                   "^--", color="#2e7d32", lw=1.7, ms=6, zorder=4,
                   label="fitted surface (cross-validated, best cut)")
            for c, k, p in spts:
                print(f"   n={c:>5} ctrl={k:>5} ratio={k/c:.3f} "
                      f"P(real)={p_real(k/c):.3f}  (cut {p})")

    # A line at the operating point makes the claim readable directly: no
    # sweep drops materially below it, at any size, in any direction.
    B.axhline(p_real(bk / bc), color="#888", ls="-", lw=0.9, zorder=1)
    # Marked on the axes rather than in the legend: a star sized to be visible
    # against the curves is far too large as a legend key.
    B.plot([bc], [p_real(bk / bc)], "*", color="black", ms=18, zorder=6)
    # Upper right. Every series falls away past the operating point, so above
    # 800 grid points nothing rises over 0.7 and the top of that side is
    # empty; the strip low and right, where this used to sit, is now crossed
    # by the descending traces.
    B.annotate(f"catalog\n{bc} grid points\nP(real) {p_real(bk/bc):.2f}",
               xy=(bc, p_real(bk / bc)), xytext=(bc * 2.4, 0.945),
               fontsize=8.6, color="#111", ha="left", va="top",
               arrowprops=dict(arrowstyle="->", color="#444", lw=0.9,
                               connectionstyle="arc3,rad=0.2"))
    # A log axis is kept because the informative part of this plane is the
    # 100-400 window range, which a linear axis running to 1,400 compresses
    # into its left quarter. What was actually wrong was the labelling: the
    # default log formatter writes 10^2 and 10^3. These are written out
    # instead, at values a reader would choose, with the minor ticks silenced
    # so nothing is labelled in scientific notation.
    B.set_xscale("log")
    xticks = [25, 50, 100, 200, 400, 800, 1600]
    B.set_xticks(xticks)
    B.set_xticklabels([f"{t:,}" for t in xticks])
    B.set_xticks([], minor=True)
    B.xaxis.set_major_formatter(mticker.FuncFormatter(lambda v, _: f"{v:,.0f}"))
    B.set_xlabel("catalog size (Neandertal-direction grid points)")
    B.set_ylabel("P(real) = 1 - control/case")
    B.set_ylim(0.10, 0.95)
    B.grid(alpha=0.25, lw=0.5)
    # Every series enters above 0.4 at the left edge and the catalog callout
    # holds the right, so the lower left is the one corner that stays clear.
    B.legend(loc="lower left", ncol=1, frameon=True, framealpha=0.92,
             edgecolor="#cccccc", fontsize=8.6, handlelength=1.9,
             borderpad=0.6, labelspacing=0.45)
    B.set_title("B", loc="left", fontweight="bold")

    # No twin axis here. The old right axis re-expressed the left one through
    # P(real) = 1 - ratio/null, so the panel labelled a single quantity twice
    # on two scales that ran in opposite directions. P(real) is the quantity
    # worth reading, so it is now the left axis and the ratio is gone.

    fig.tight_layout()
    os.makedirs(os.path.dirname(a.out) or ".", exist_ok=True)
    fig.savefig(a.out, dpi=200, bbox_inches="tight")
    print(f"\nwrote {a.out}")
    return 0


if __name__ == "__main__":
    raise SystemExit(main())
```
