## Supplementary Information section 4 code base for "Hypothesis: A modern human range expansion ∼300,000 years ago had a critical role in Neandertal origins": fig_s51_model_curve.py

```
#!/usr/bin/env python3
"""Does the fitted three-way score beat the raw box rule?

THE COMPARISON
Every curve is drawn in the same plane -- Neandertal-direction windows retained
on x, Denisovan/Neandertal ratio on y -- so a curve lying lower is strictly
better: a cleaner catalog at equal size. Four rules are compared:

  box, vary threshold   nea_hi(p) <= x and den_lo(p) > x, cut held at 80/20
  box, vary cut         the same, threshold held at 625 ka
  score, cut 80/20      rank windows by the fitted eta at p = 0.80
  score, best cut       rank by eta, cut chosen freely

CROSS-VALIDATION
The score is fitted on odd chromosomes and every curve is evaluated on even
chromosomes. This matters more than it usually would: the response the model
fits (Neandertal versus Denisovan direction) is the very quantity plotted on
the y-axis, so an in-sample curve would be guaranteed to look good and would
demonstrate nothing. The box curves are evaluated on the same even-chromosome
half so that all four are strictly comparable.

The control is the identical rule with the two archaics swapped -- for the
score, eta evaluated at the swapped coordinates -- so it inherits the ARG's
idiosyncrasies and its count estimates how many case windows arise with no
real signal behind them.

MATCHED CATALOG SIZES
The score curves use top-N selection: taking the N highest-scoring windows
gives exactly N cases, so the score is read at the same catalog sizes the box
rule produces, and no interpolation is needed to compare them.

Scoring is confined to the region where the model was fitted (u in 350-900 ka,
v in 350-1500 ka); the basis carries quadratic terms, which extrapolate wildly
outside their support.

Usage:
  python3 scripts/fig_s51_model_curve.py -o catalog_annotation/figure_s5_1_model.png
"""

from __future__ import annotations

import argparse
import os
import sys
from collections import defaultdict

import numpy as np

sys.path.insert(0, os.path.dirname(os.path.abspath(__file__)))
from approx_percentiles import default_coord, fit_site, load, z_of  # noqa: E402
from fig_s51_sensitivity import wilson  # noqa: E402
from fit_fdr_surface import (PCUTS, U_EDGES, V_EDGES, binof, design,  # noqa: E402
                             irls)

BASE_X = 625.0
BASE_P = 0.80
TARGETS = [50, 75, 100, 150, 200, 300, 400, 550, 750, 1000, 1400, 2000]


def add_tails(recs, coord):
    for d in recs:
        for tag in ("nea", "den"):
            mu, sg = fit_site(coord, d[f"{tag}_p025"], d[f"{tag}_p975"])
            for p in PCUTS:
                d[f"{tag}_hi{p}"] = coord.to_t(mu + z_of(p) * sg)
                d[f"{tag}_lo{p}"] = coord.to_t(mu + z_of(1 - p) * sg)


def fit_on(recs):
    """Cell-level binomial fit, identical in form to fit_fdr_surface."""
    cells = defaultdict(lambda: [0, 0])
    for p in PCUTS:
        for d in recs:
            iu, iv = binof(d[f"nea_hi{p}"], U_EDGES), binof(d[f"den_lo{p}"], V_EDGES)
            if iu is not None and iv is not None:
                cells[(p, iu, iv)][0] += 1
            ju, jv = binof(d[f"den_hi{p}"], U_EDGES), binof(d[f"nea_lo{p}"], V_EDGES)
            if ju is not None and jv is not None:
                cells[(p, ju, jv)][1] += 1
    u, v, pp, k, n = [], [], [], [], []
    for (p, iu, iv), (kn, kd) in sorted(cells.items()):
        u.append((U_EDGES[iu] + U_EDGES[iu + 1]) / 200.0)
        v.append((V_EDGES[iv] + V_EDGES[iv + 1]) / 200.0)
        pp.append(p); k.append(kn); n.append(kn + kd)
    X = design(np.array(u), np.array(v), np.array(pp))
    beta, _ = irls(X, np.array(k, dtype=float), np.array(n, dtype=float))
    return beta, len(u)


def in_support(uu, vv):
    return (U_EDGES[0] < uu <= U_EDGES[-1]) and (V_EDGES[0] < vv <= V_EDGES[-1])


def box_curve(recs, specs):
    """(count, ratio) for box rules given (x_thresh, cut) specs."""
    out = []
    for x, p in specs:
        case = sum(1 for d in recs
                   if d[f"nea_hi{p}"] <= x and d[f"den_lo{p}"] > x)
        ctrl = sum(1 for d in recs
                   if d[f"den_hi{p}"] <= x and d[f"nea_lo{p}"] > x)
        if case:
            out.append((case, ctrl, x, p))
    return out


def score_curve(recs, beta, pcuts):
    """(count, ratio) by top-N selection on the fitted score, best p per N."""
    per_p = {}
    for p in pcuts:
        ec, et = [], []
        for d in recs:
            u, v = d[f"nea_hi{p}"], d[f"den_lo{p}"]
            if in_support(u, v):
                ec.append(design(np.array([u / 100.0]), np.array([v / 100.0]),
                                 np.array([p]))[0] @ beta)
            us, vs = d[f"den_hi{p}"], d[f"nea_lo{p}"]
            if in_support(us, vs):
                et.append(design(np.array([us / 100.0]), np.array([vs / 100.0]),
                                 np.array([p]))[0] @ beta)
        per_p[p] = (np.sort(np.array(ec))[::-1], np.sort(np.array(et))[::-1])

    curves = {}
    for p, (ec, et) in per_p.items():
        pts = []
        for N in TARGETS:
            if N > len(ec):
                continue
            c = ec[N - 1]
            ctrl = int(np.searchsorted(-et, -c, side="right"))
            pts.append((N, ctrl))
        curves[p] = pts
    return curves


def main():
    ap = argparse.ArgumentParser()
    ap.add_argument("--scan", default="hubisz_tmrca_5kb_8afr.tsv")
    ap.add_argument("-o", "--out",
                    default="catalog_annotation/figure_s5_1_model.png")
    args = ap.parse_args()

    import matplotlib
    matplotlib.use("Agg")
    import matplotlib.pyplot as plt

    recs = load(args.scan)
    coord, grid_desc = default_coord(recs)
    print(f"time coordinate: {grid_desc}")
    recs = [d for d in recs if d["chrom"].replace("chr", "") not in ("X", "23")]
    add_tails(recs, coord)

    odd = [d for d in recs if int(d["chrom"].replace("chr", "")) % 2 == 1]
    even = [d for d in recs if int(d["chrom"].replace("chr", "")) % 2 == 0]
    print(f"fit on {len(odd):,} odd-chromosome windows; "
          f"evaluate on {len(even):,} even-chromosome windows")

    beta, ncells = fit_on(odd)
    print(f"fitted on {ncells:,} cells; beta = "
          + ", ".join(f"{b:+.3f}" for b in beta))

    fig, ax = plt.subplots(figsize=(8.2, 5.6))

    # --- box rules, held-out half ---
    thr = box_curve(even, [(x, BASE_P) for x in
                           (550, 575, 600, 625, 650, 675, 700, 750, 800, 900)])
    cut = box_curve(even, [(BASE_X, p) for p in PCUTS])
    for pts, color, lab in (
            (thr, "#c00000", "box rule, vary threshold (cut 80/20)"),
            (cut, "#1f4e79", "box rule, vary cut (threshold 625 ka)")):
        xs = [c for c, k, _, _ in pts]; ys = [k / c for c, k, _, _ in pts]
        lo = [wilson(k, c)[0] for c, k, _, _ in pts]
        hi = [wilson(k, c)[1] for c, k, _, _ in pts]
        ax.plot(xs, ys, "o-", color=color, lw=1.7, ms=5, label=lab, zorder=3)
        ax.fill_between(xs, lo, hi, color=color, alpha=0.12, lw=0, zorder=2)
        print(f"\n{lab}")
        for c, k, x, p in pts:
            print(f"  x={x:>5.0f} p={p:<5}  case={c:>6} ctrl={k:>6} "
                  f"ratio={k/c:.3f}")

    # --- fitted score, held-out half ---
    curves = score_curve(even, beta, PCUTS)
    base = curves.get(BASE_P, [])
    if base:
        xs = [N for N, _ in base]; ys = [k / N for N, k in base]
        ax.plot(xs, ys, "^-", color="#2e7d32", lw=1.9, ms=6,
                label="fitted score, cut 80/20 (held out)", zorder=4)
        print("\nfitted score, cut 80/20")
        for N, k in base:
            print(f"  N={N:>6} ctrl={k:>6} ratio={k/N:.3f}")

    best = {}
    for p, pts in curves.items():
        for N, k in pts:
            if N not in best or k < best[N][0]:
                best[N] = (k, p)
    if best:
        xs = sorted(best); ys = [best[N][0] / N for N in xs]
        ax.plot(xs, ys, "^--", color="#66bb6a", lw=1.5, ms=5,
                label="fitted score, best cut (held out)", zorder=4)
        print("\nfitted score, best cut per size")
        for N in xs:
            k, p = best[N]
            print(f"  N={N:>6} ctrl={k:>6} ratio={k/N:.3f}  (p={p})")

    ax.set_xscale("log")
    ax.set_xlabel("Neandertal-direction windows retained (catalog size)")
    ax.set_ylabel("Denisovan / Neandertal windows  (false-discovery proxy)")
    ax.set_ylim(0, 0.62)
    ax.grid(alpha=0.25, lw=0.5)
    ax.legend(loc="upper left", frameon=False, fontsize=9)
    ax.set_title("Does the fitted score earn its parameters? "
                 "Lower is better; all curves held out",
                 loc="left", fontweight="bold", fontsize=11)
    fig.tight_layout()
    os.makedirs(os.path.dirname(args.out) or ".", exist_ok=True)
    fig.savefig(args.out, dpi=200, bbox_inches="tight")
    print(f"\nwrote {args.out}")
    return 0


if __name__ == "__main__":
    raise SystemExit(main())
```
