## Supplementary Information section 4 code base for "Hypothesis: A modern human range expansion ∼300,000 years ago had a critical role in Neandertal origins": fig_s51_sensitivity.py

```
#!/usr/bin/env python3
"""One-at-a-time sensitivity of the catalog rule, and the counting helpers
that the Figure S4.1 scripts import.

site_percentiles(), counts() and wilson() are imported by fig_s51_composite.py,
fig_s51_model_curve.py, build_preal_grid.py, site_exact_preal.py,
make_loci_table.py and den_threshold_test.py. Those callers pass the rule in,
so the module is required even when its own main() is not run.

Running this file directly reproduces an earlier single-panel draft whose
baseline was

    case     p80[ TMRCA(Africans + both Neandertals) ] <= 625 ka
        and  p20[ TMRCA(Africans + Denisova)         ] >  625 ka
    control  the same with the archaics swapped

which is NOT the rule the catalog was built with. The published rule takes the
median (percentile cut 50) and separates the two thresholds, 625 ka on
TMRCA(Africans + Neandertals) and 700 ka on TMRCA(Africans + Denisovans); it is
defined in fig_s51_composite.py as BASE_X, BASE_Y, BASE_CUT = 625, 700, 50 and
that is the script that draws Figure S4.1. The BASE_X and BASE_CUT constants
below belong to the superseded draft and are not used by the callers above.

Three panels vary one parameter each over a wide range while the other two are
held at baseline: the Neandertal threshold, the Denisovan threshold, and the
percentile cut.

Each panel plots BOTH the Denisovan/Neandertal ratio and the Neandertal-
direction count, on twin axes. The ratio alone has no interior optimum -- it
falls monotonically as the rule tightens, reaching 0.02 at 500 ka, so a
ratio-only figure argues for the tightest threshold available and cannot
justify any particular choice. It is the pairing with the count that shows why
the baseline sits where it does: the ratio is still low while the catalog is
still large enough to be worth having.

The control is the same rule with Neandertal and Denisovan swapped, so it
inherits every idiosyncrasy of the ARG inference. Its count is an estimate of
how many case windows arise without a real signal, making the ratio an FDR
proxy rather than an arbitrary index.

Usage:
  python3 scripts/fig_s51_sensitivity.py --scan 2_hubisz_tmrca_5kb.tsv \
      -o catalog_annotation/figure_s5_1_sensitivity.png
"""

from __future__ import annotations

import argparse
import math
import os
import sys

sys.path.insert(0, os.path.dirname(os.path.abspath(__file__)))
from approx_percentiles import default_coord, fit_site, load, z_of  # noqa: E402

# load() renames the scan's verbose column prefixes to these short tags.
NEA = "nea"
DEN = "den"
BASE_X = 625.0
BASE_CUT = 80.0


def site_percentiles(recs, coord):
    """Per-window (mu, sigma) in the log-spaced time coordinate, for both queries.

    Done once: the percentile reconstruction is the expensive part, and every
    panel reuses it.
    """
    out = []
    for d in recs:
        try:
            mn, sn = fit_site(coord, d[f"{NEA}_p025"], d[f"{NEA}_p975"])
            md, sd = fit_site(coord, d[f"{DEN}_p025"], d[f"{DEN}_p975"])
        except (KeyError, TypeError, ValueError):
            continue
        out.append((mn, sn, md, sd))
    return out


def counts(sites, coord, x_nea, x_den, cut):
    """(n_case, n_control) under a given rule.

    cut is the upper percentile; its mirror (100-cut) is the lower one, so the
    80/20 baseline is cut=80.
    """
    z_hi = z_of(cut / 100.0)
    z_lo = z_of(1.0 - cut / 100.0)
    u_nea, u_den = coord.to_u(x_nea), coord.to_u(x_den)
    case = ctrl = 0
    for mn, sn, md, sd in sites:
        # case: Neandertal query shallow, Denisovan query deep
        if (mn + z_hi * sn) <= u_nea and (md + z_lo * sd) > u_den:
            case += 1
        # control: identical rule, archaics swapped
        if (md + z_hi * sd) <= u_nea and (mn + z_lo * sn) > u_den:
            ctrl += 1
    return case, ctrl


def wilson(k, n, z=1.96):
    """Confidence interval for the Denisovan/Neandertal ratio k/n.

    That ratio is an odds, not a proportion -- it is unbounded above -- so the
    interval is built on the binomial proportion q = k/(k+n), which is what the
    counts actually are, and then mapped through q/(1-q). Applying a Wilson
    interval to k/n directly breaks once k exceeds n.
    """
    if n == 0:
        return (float("nan"), float("nan"))
    tot = k + n
    q = k / tot
    d = 1 + z * z / tot
    c = (q + z * z / (2 * tot)) / d
    h = z * math.sqrt(q * (1 - q) / tot + z * z / (4 * tot * tot)) / d
    qlo, qhi = max(0.0, c - h), min(1.0 - 1e-12, c + h)
    return (qlo / (1 - qlo), qhi / (1 - qhi))


def main():
    ap = argparse.ArgumentParser()
    ap.add_argument("--scan", default="hubisz_tmrca_5kb_8afr.tsv")
    ap.add_argument("--x", type=float, default=BASE_X)
    ap.add_argument("--cut", type=float, default=BASE_CUT)
    ap.add_argument("-o", "--out", default="catalog_annotation/figure_s5_1_sensitivity.png")
    args = ap.parse_args()

    import matplotlib
    matplotlib.use("Agg")
    import matplotlib.pyplot as plt

    recs = load(args.scan)
    coord, grid_desc = default_coord(recs)
    print(f"time coordinate: {grid_desc}")
    # X-linked windows are excluded throughout this project: their Denisovan/
    # Neandertal ratio is near 1, so they are noise and including them inflates
    # every ratio reported here.
    n0 = len(recs)
    recs = [d for d in recs if d["chrom"].replace("chr", "") not in ("X", "23")]
    print(f"excluded {n0 - len(recs):,} X-linked windows")
    sites = site_percentiles(recs, coord)
    print(f"windows usable: {len(sites)}")

    b_case, b_ctrl = counts(sites, coord, args.x, args.x, args.cut)
    print(f"baseline x={args.x:.0f} cut={args.cut:.0f}/{100-args.cut:.0f}: "
          f"case={b_case} control={b_ctrl} ratio={b_ctrl/b_case:.3f}")

    panels = [
        ("A", "Neandertal threshold (ka)",
         [(x, args.x, args.cut) for x in
          (450, 500, 550, 575, 600, 625, 650, 675, 700, 750, 800, 900)],
         lambda t: t[0], args.x),
        ("B", "Denisovan threshold (ka)",
         [(args.x, x, args.cut) for x in
          (450, 500, 550, 575, 600, 625, 650, 675, 700, 750, 800, 900)],
         lambda t: t[1], args.x),
        ("C", "percentile cut (upper/lower)",
         [(args.x, args.x, c) for c in
          (50, 55, 60, 65, 70, 75, 80, 85, 90, 95)],
         lambda t: t[2], args.cut),
    ]

    fig, axes = plt.subplots(1, 3, figsize=(15, 4.6))
    twin_handle = None
    for ax, (tag, xlabel, grid, getx, basev) in zip(axes, panels):
        xs, ratios, los, his, ncase = [], [], [], [], []
        for spec in grid:
            c, k = counts(sites, coord, *spec)
            if c == 0:
                continue
            lo, hi = wilson(k, c)
            xs.append(getx(spec)); ratios.append(k / c)
            los.append(lo); his.append(hi); ncase.append(c)
            print(f"  {tag} x={getx(spec):>5}  case={c:>6} ctrl={k:>6} "
                  f"ratio={k/c:.3f}")

        ax.plot(xs, ratios, "o-", color="#1f4e79", lw=1.8, ms=5,
                label="Denisovan / Neandertal", zorder=3)
        ax.fill_between(xs, los, his, color="#1f4e79", alpha=0.15, lw=0,
                        zorder=2)
        ax.axvline(basev, color="#c00000", ls="--", lw=1.2, zorder=1)
        ax.set_xlabel(xlabel)
        ax.set_ylim(0, 1.0)
        ax.grid(alpha=0.25, lw=0.5)
        ax.set_title(f"{tag}", loc="left", fontweight="bold")

        ax2 = ax.twinx()
        ln2, = ax2.plot(xs, ncase, "s--", color="#7f7f7f", lw=1.2, ms=4,
                        label="Neandertal-direction windows (right axis)",
                        zorder=3)
        twin_handle = twin_handle or ln2
        ax2.set_yscale("log")
        # All three panels share a count range, so a common axis makes the
        # panels comparable and stops the log ticks from colliding with the
        # ratio axis of the panel to the left.
        ax2.set_ylim(5, 1e4)
        ax2.tick_params(axis="y", colors="#7f7f7f", labelsize=8)
        if tag == "C":
            ax2.set_ylabel("Neandertal-direction windows (log)",
                           color="#7f7f7f")
        else:
            ax2.set_yticklabels([])
        if tag == "A":
            ax.set_ylabel("Denisovan / Neandertal windows")

    # One legend for the figure: both series mean the same thing in every panel.
    h1, l1 = axes[0].get_legend_handles_labels()
    fig.legend(h1 + ([twin_handle] if twin_handle else []),
               l1 + ([twin_handle.get_label()] if twin_handle else []),
               loc="lower center", ncol=2, frameon=False,
               bbox_to_anchor=(0.5, -0.04))
    fig.suptitle(
        f"Sensitivity of the catalog rule around the baseline "
        f"(dashed red: {args.x:.0f} ka, {args.cut:.0f}/{100-args.cut:.0f} cut)",
        y=1.02)
    fig.tight_layout()
    os.makedirs(os.path.dirname(args.out) or ".", exist_ok=True)
    fig.savefig(args.out, dpi=200, bbox_inches="tight")
    print(f"wrote {args.out}")
    return 0


if __name__ == "__main__":
    raise SystemExit(main())
```
