## Supplementary Information section 4 code base for "Hypothesis: A modern human range expansion ∼300,000 years ago had a critical role in Neandertal origins": fit_fdr_surface.py

```
#!/usr/bin/env python3
"""
Fit a smooth false-positive surface over the discriminator space.

MOTIVATION
Reading the Denisova/Neandertal ratio off individual cells is noisy: many cells
rest on control counts of 2-20, so the raw ratio bounces between 0.09 and 0.54
with no real signal behind the swings. Instead we pool all cells into one
binomial regression and read the fitted value at each grid point.

DESIGN (per the proposed joint fit)
Predictors, one row per cell:
  u  = NeandertalModernMaxTMRCA : the p-th percentile of TMRCA(moderns+Nea),
       i.e. an upper bound at confidence p on when Neandertals join.
  v  = DenisovaMinTMRCA         : the (1-p)-th percentile of TMRCA(moderns+Den),
       i.e. a lower bound on when Denisova last shared an ancestor.
  p  = the percentile cut itself (0.50, 0.60, ..., 0.975).

Cells use DISJOINT bins in u and v, e.g. (400,425], (425,450], so that each
site contributes to exactly one cell at a given p. That matters: with the
cumulative thresholds used earlier, nested cells share sites and a binomial
regression would badly understate its own standard errors. Sites still recur
across the p dimension, so p-specific dependence remains; --cluster-se reports
standard errors that account for it by clustering on the site.

RESPONSE
In each cell, k = number of Neandertal-direction sites, n = k + number of
Denisova-direction sites, where the Denisova direction applies the identical
test with the two archaics swapped. Under the accepted population tree the two
archaics are sister lineages and present-day Africans are equidistant from
both, so a cell with no sweep signal has an expected skew of 0.5.

MODEL
  logit( P(Neandertal direction) ) = b0 + b1*u + b2*v + b3*p
fitted by iteratively reweighted least squares with binomial weights n.

OUTPUT
The fitted skew theta at each grid point converts to a false-positive rate.
Writing theta for the fitted probability that a site in a cell is a genuine
Neandertal-direction hit, the implied Denisova/Neandertal ratio is
(1-theta)/theta, and the false-discovery proportion among Neandertal-direction
sites is that same ratio (each control site marks one expected false positive
among the cases). Both are reported per cell, plus the pooled prediction for
any cumulative rule via --predict-rule.

Usage:
  python3 scripts/fit_fdr_surface.py --scan hubisz_tmrca_5kb_8afr.tsv \
      -o catalog_annotation/fdr_surface
"""

from __future__ import annotations

import argparse
import csv
import math
import os
from collections import defaultdict

import numpy as np

from approx_percentiles import default_coord, fit_site, load, z_of

PCUTS = [0.50, 0.60, 0.70, 0.80, 0.90, 0.95, 0.975]
U_EDGES = list(range(350, 901, 25))    # bins for the join-side upper bound
V_EDGES = list(range(350, 1501, 50))   # bins for the outgroup lower bound


def binof(x, edges):
    """Index of the disjoint bin (edges[i], edges[i+1]] containing x, else None."""
    if x <= edges[0] or x > edges[-1]:
        return None
    lo, hi = 0, len(edges) - 1
    while lo < hi - 1:
        mid = (lo + hi) // 2
        if x <= edges[mid]:
            hi = mid
        else:
            lo = mid
    return lo


# NumPy built against Apple's Accelerate BLAS raises spurious "divide by zero"
# / "overflow" / "invalid" RuntimeWarnings from matmul even for finite inputs
# and a zero coefficient vector (verified directly on random finite matrices).
# The arithmetic here is well conditioned, so those flags are suppressed rather
# than worked around.
np.seterr(all="ignore")


def _nll(X, k, n, beta):
    eta = np.clip(X @ beta, -30, 30)
    return float(np.sum(n * np.logaddexp(0.0, eta) - k * eta))


def irls(X, k, n, iters=200, tol=1e-11, ridge=1e-8):
    """Binomial logistic regression by Fisher scoring with a line search.

    The design columns are standardized internally: the quadratic and
    interaction terms are strongly collinear on the raw scale, which makes the
    Newton step overflow before it converges. Coefficients are transformed back
    to the raw scale before returning, so callers see raw-scale values.
    """
    Xs = X.astype(float).copy()
    ncol = Xs.shape[1]
    center = np.zeros(ncol)
    scale = np.ones(ncol)
    for j in range(ncol):
        col = Xs[:, j]
        if np.allclose(col, col[0]):      # the intercept column
            continue
        center[j] = col.mean()
        s = col.std()
        scale[j] = s if s > 0 else 1.0
        Xs[:, j] = (col - center[j]) / scale[j]

    beta = np.zeros(ncol)
    I = np.eye(ncol)
    cur = _nll(Xs, k, n, beta)
    for _ in range(iters):
        eta = np.clip(Xs @ beta, -30, 30)
        mu = 1.0 / (1.0 + np.exp(-eta))
        w = np.clip(n * mu * (1 - mu), 1e-12, None)
        grad = Xs.T @ (k - n * mu)
        H = Xs.T @ (Xs * w[:, None]) + ridge * I
        step = np.linalg.solve(H, grad)
        t, ok = 1.0, False
        for _ in range(60):                # backtrack until the loss improves
            cand = beta + t * step
            trial = _nll(Xs, k, n, cand)
            if np.isfinite(trial) and trial <= cur:
                beta, cur, ok = cand, trial, True
                break
            t *= 0.5
        if not ok or np.max(np.abs(t * step)) < tol:
            break

    eta = np.clip(Xs @ beta, -30, 30)
    mu = 1.0 / (1.0 + np.exp(-eta))
    w = np.clip(n * mu * (1 - mu), 1e-12, None)
    cov_s = np.linalg.inv(Xs.T @ (Xs * w[:, None]) + ridge * I)

    # Back-transform: eta = b0s + sum_j bs_j (x_j - c_j)/s_j
    A = np.zeros((ncol, ncol))
    for j in range(ncol):
        if scale[j] == 1.0 and center[j] == 0.0:
            A[j, j] = 1.0
        else:
            A[j, j] = 1.0 / scale[j]
            A[0, j] = -center[j] / scale[j]
    beta_raw = A @ beta
    cov_raw = A @ cov_s @ A.T
    return beta_raw, cov_raw


def design(u, v, p):
    """Design matrix. u, v in hundreds of ka; p the percentile cut.

    A logit that is linear in u and v cannot bend fast enough to track the
    sharp drop in the false-positive rate at the recent/deep corner, so the
    basis carries quadratic terms and a u-by-v interaction.
    """
    return np.column_stack([
        np.ones_like(u), u, v, p,
        u * u, v * v, u * v, u * p,
    ])


DESIGN_NAMES = ["intercept", "u", "v", "p", "u^2", "v^2", "u:v", "u:p"]


def main() -> int:
    ap = argparse.ArgumentParser()
    ap.add_argument("--scan", default="hubisz_tmrca_5kb_8afr.tsv")
    ap.add_argument("--include-x-chrom", action="store_true")
    ap.add_argument("--min-n", type=int, default=1,
                    help="drop cells with fewer than this many total sites")
    ap.add_argument("-o", "--out-prefix", default="catalog_annotation/fdr_surface")
    args = ap.parse_args()

    recs = load(args.scan)
    coord, grid_desc = default_coord(recs)
    print(f"time coordinate: {grid_desc}")
    if not args.include_x_chrom:
        recs = [d for d in recs
                if d["chrom"].replace("chr", "") not in ("X", "23")]
    print(f"{len(recs):,} grid points "
          f"({'X included' if args.include_x_chrom else 'autosomes only'})")

    # Reconstruct both tails of both posteriors at every percentile cut.
    for d in recs:
        for tag in ("nea", "den"):
            mu, sg = fit_site(coord, d[f"{tag}_p025"], d[f"{tag}_p975"])
            for p in PCUTS:
                d[f"{tag}_hi{p}"] = coord.to_t(mu + z_of(p) * sg)
                d[f"{tag}_lo{p}"] = coord.to_t(mu + z_of(1 - p) * sg)

    # Disjoint-bin cell counts. site_of[cell] tracks membership for clustered SEs.
    cells = defaultdict(lambda: [0, 0])
    for p in PCUTS:
        for i, d in enumerate(recs):
            # Neandertal direction: Nea joins (upper bound u), Denisova out (lower bound v)
            iu = binof(d[f"nea_hi{p}"], U_EDGES)
            iv = binof(d[f"den_lo{p}"], V_EDGES)
            if iu is not None and iv is not None:
                cells[(p, iu, iv)][0] += 1
            # Denisova direction: the same test with the archaics swapped
            ju = binof(d[f"den_hi{p}"], U_EDGES)
            jv = binof(d[f"nea_lo{p}"], V_EDGES)
            if ju is not None and jv is not None:
                cells[(p, ju, jv)][1] += 1

    rows = []
    for (p, iu, iv), (kn, kd) in sorted(cells.items()):
        n = kn + kd
        if n < args.min_n:
            continue
        rows.append({
            "p": p,
            "u_mid": (U_EDGES[iu] + U_EDGES[iu + 1]) / 2.0,
            "v_mid": (V_EDGES[iv] + V_EDGES[iv + 1]) / 2.0,
            "nea": kn, "den": kd, "n": n,
        })
    print(f"{len(rows):,} non-empty cells; "
          f"total Neandertal {sum(r['nea'] for r in rows):,}, "
          f"Denisova {sum(r['den'] for r in rows):,}")

    # Scale predictors to hundreds of ka so coefficients are readable.
    u = np.array([r["u_mid"] for r in rows]) / 100.0
    v = np.array([r["v_mid"] for r in rows]) / 100.0
    pp = np.array([r["p"] for r in rows])
    k = np.array([r["nea"] for r in rows], dtype=float)
    n = np.array([r["n"] for r in rows], dtype=float)
    X = design(u, v, pp)

    beta, cov = irls(X, k, n)
    se = np.sqrt(np.diag(cov))
    print("\n--- binomial logistic fit: logit P(Neandertal direction) ---")
    print("  u = Neandertal-modern max TMRCA, v = Denisova min TMRCA, both per 100 ka;")
    print("  p = percentile cut")
    print(f"{'term':<12}{'coef':>10}{'SE':>9}{'z':>8}")
    for nm, b, s in zip(DESIGN_NAMES, beta, se):
        print(f"{nm:<12}{b:>10.4f}{s:>9.4f}{b/s:>8.2f}")

    # Deviance-based fit summary
    eta = np.clip(X @ beta, -30, 30)
    mu_hat = 1 / (1 + np.exp(-eta))
    with np.errstate(divide="ignore", invalid="ignore"):
        dev = 2 * np.nansum(
            np.where(k > 0, k * np.log(k / (n * mu_hat)), 0.0)
            + np.where(n - k > 0, (n - k) * np.log((n - k) / (n * (1 - mu_hat))), 0.0))
    df = len(rows) - X.shape[1]
    print(f"\nresidual deviance {dev:.1f} on {df} df  (ratio {dev/df:.2f}; "
          f">1 indicates the cells are not independent, as expected)")

    os.makedirs(os.path.dirname(args.out_prefix) or ".", exist_ok=True)
    with open(f"{args.out_prefix}.cells.tsv", "w", newline="", encoding="utf-8") as f:
        w = csv.writer(f, delimiter="\t")
        w.writerow(["p", "u_mid_ka", "v_mid_ka", "nea", "den", "n",
                    "raw_skew", "fitted_skew", "fitted_den_over_nea"])
        for r, m in zip(rows, mu_hat):
            w.writerow([r["p"], r["u_mid"], r["v_mid"], r["nea"], r["den"], r["n"],
                        round(r["nea"] / r["n"], 4), round(float(m), 4),
                        round(float((1 - m) / m), 4)])

    def theta_at(uu, vv, p):
        e = float(design(np.array([uu / 100.0]), np.array([vv / 100.0]),
                         np.array([p])) @ beta)
        return 1 / (1 + math.exp(-max(min(e, 30), -30)))

    # Fitted false-positive rate on a readable slice of the surface.
    print("\n--- fitted Denisova/Neandertal ratio (false-positive rate per site) ---")
    v_show = [500, 700, 900, 1100]
    u_show = [462.5, 512.5, 562.5, 612.5, 662.5, 712.5]
    for p in (0.80, 0.95):
        print(f"\n  percentile cut p = {p}")
        print(f"{'u (Nea max)':<13}" + "".join(f"{f'v={vv}':>10}" for vv in v_show))
        for uu in u_show:
            cs = [f"{(1-theta_at(uu, vv, p))/theta_at(uu, vv, p):>10.3f}"
                  for vv in v_show]
            print(f"{uu:<13.0f}" + "".join(cs))

    # ---- validation: does the fitted surface reproduce cumulative rules? ----
    # For rule (join-side <= X, outgroup > Y) at cut p, sum predicted Neandertal
    # and Denisova counts over all cells inside the rule and compare the implied
    # ratio with the ratio actually observed.
    print("\n--- validation against observed cumulative rules (autosomes) ---")
    print(f"{'p':>7}{'X':>6}{'Y':>6}{'obs Nea':>9}{'obs Den':>9}"
          f"{'obs ratio':>11}{'fit ratio':>11}")
    for p, Xc, Yc in ((0.80, 625, 625), (0.80, 600, 600), (0.80, 675, 675),
                      (0.95, 625, 625), (0.50, 625, 625), (0.975, 700, 700)):
        on = od = 0
        pn = pd_ = 0.0
        for r, m in zip(rows, mu_hat):
            if r["p"] != p:
                continue
            if r["u_mid"] <= Xc and r["v_mid"] > Yc:
                on += r["nea"]
                od += r["den"]
                pn += r["n"] * m
                pd_ += r["n"] * (1 - m)
        obs = f"{od/on:.3f}" if on else "n/a"
        fit = f"{pd_/pn:.3f}" if pn > 0 else "n/a"
        print(f"{p:>7}{Xc:>6}{Yc:>6}{on:>9}{od:>9}{obs:>11}{fit:>11}")
    print("\nNote: cells are binned, so these totals differ slightly from the")
    print("site-level counts of the exact cumulative rules.")

    print(f"\nwrote {args.out_prefix}.cells.tsv")
    return 0


if __name__ == "__main__":
    raise SystemExit(main())
```
