## Supplementary Information section 4 code base for "Hypothesis: A modern human range expansion ∼300,000 years ago had a critical role in Neandertal origins": great_annotation_bias.py

```
#!/usr/bin/env python3
"""Is the proteasome enrichment annotation depth rather than biology?

The B-matched permutation in great_test.py controls where the catalog looks. It
cannot control what GO knows. The proteasome is one of the best characterised
complexes in the cell, and well studied genes accumulate GO terms, so a gene set
that happens to contain famous genes will look enriched for whatever those genes
are famous for. Three tests separate the two explanations.

  1. degree of the hit genes. If the catalog's genes carry no more GO terms than
     genes drawn from matched genomic locations, annotation depth cannot be
     inflating the result, because the null already has the same depth.

  2. degree-matched gene null. Gene sets of the same size are drawn from the
     universe matched on number of GO terms, ignoring genomic position. This
     asks whether any set of equally well annotated genes would show the same
     excess. It is deliberately harsh: it discards the spatial evidence.

  3. IEA annotations included. Electronic annotations are transferred by
     homology rather than earned by publication, so they cover obscure genes far
     more evenly than curated ones. If the signal is study bias it should shrink
     when they are added.
"""

import argparse
import bisect
import csv
import collections
import os
import random
import sys

from scipy.stats import hypergeom

sys.path.insert(0, os.path.dirname(os.path.abspath(__file__)))
from great_domains import (ANN, HG19, domains, genes_for_regions,  # noqa: E402
                           load_grid_preal, load_tss, regions_at_threshold)
from great_test import (ancestors, load_bstat, load_gaf, load_obo,  # noqa: E402
                        matched_block_draws, merge_regions)

TERM = "GO:0043161"  # proteasome-mediated ubiquitin-dependent protein catabolic


def degree_matched_sets(observed, universe, degree, nrep, rng, nbin=10):
    """Gene sets of len(observed) drawn from the universe, matched on GO degree."""
    vals = sorted(degree[g] for g in universe)
    edges = [vals[int(len(vals) * i / nbin)] for i in range(1, nbin)]
    pool = collections.defaultdict(list)
    for g in universe:
        pool[bisect.bisect_right(edges, degree[g])].append(g)
    want = collections.Counter(
        bisect.bisect_right(edges, degree[g]) for g in observed)
    out = []
    for _ in range(nrep):
        pick = []
        for b, k in want.items():
            src = pool[b]
            pick.extend(rng.sample(src, min(k, len(src))))
        out.append(set(pick))
    return out


def main():
    ap = argparse.ArgumentParser()
    ap.add_argument("--threshold", type=float, default=0.5)
    ap.add_argument("--regions", default=None,
                    help="region TSV to test instead of thresholding P(real)")
    ap.add_argument("--merge-gap", type=int, default=100000)
    ap.add_argument("--spatial-reps", type=int, default=2000)
    ap.add_argument("--degree-reps", type=int, default=10000)
    ap.add_argument("--keep-iea", action="store_true")
    ap.add_argument("--seed", type=int, default=11)
    a = ap.parse_args()

    terms, alt = load_obo(os.path.join(
        os.path.dirname(os.path.dirname(os.path.abspath(__file__))),
        "refdata", "go", "go-basic.obo"))
    anc = ancestors(terms)
    sym2go = load_gaf(terms, alt, anc, {"P"}, not a.keep_iea)

    tss = load_tss(("protein_coding",))
    doms = domains(tss)
    gene_domains = collections.defaultdict(list)
    for c, ds in doms.items():
        for s, e, sym in ds:
            gene_domains[sym].append((c, s, e))
    universe = sorted(set(gene_domains) & set(sym2go))
    degree = {g: len(sym2go[g]) for g in universe}

    preal = load_grid_preal()
    if a.regions:
        regs = []
        with open(a.regions, encoding="utf-8") as f:
            for r in csv.DictReader(f, delimiter="\t"):
                c = r["chrom"].replace("chr", "")
                if c in HG19:
                    regs.append((c, int(r["start"]), int(r["end"])))
    else:
        regs, _, _ = regions_at_threshold(preal, a.threshold)
    regs = merge_regions(regs, a.merge_gap)
    by_gene, _ = genes_for_regions(regs, doms)
    hits = sorted(set(by_gene) & set(universe))

    term_genes = {g for g in universe if TERM in sym2go[g]}
    k_obs = len(term_genes & set(hits))
    p_obs = hypergeom.sf(k_obs - 1, len(universe), len(term_genes), len(hits))
    label = "with IEA" if a.keep_iea else "curated only"
    print(f"\n=== {label} ===")
    print(f"universe {len(universe):,} genes; term has {len(term_genes)} genes; "
          f"catalog hits {len(hits)} genes, {k_obs} in term "
          f"(expected {len(term_genes)*len(hits)/len(universe):.1f}), "
          f"p_hyper {p_obs:.2e}")

    # ---- test 1 and 2 need the degree of the observed hits --------------
    med_u = sorted(degree[g] for g in universe)[len(universe) // 2]
    med_h = sorted(degree[g] for g in hits)[len(hits) // 2]
    mean_h = sum(degree[g] for g in hits) / len(hits)
    print(f"\nGO terms per gene: universe median {med_u}, "
          f"catalog-hit median {med_h}, hit mean {mean_h:.1f}")
    in_term = sorted(term_genes & set(hits))
    if in_term:
        print(f"  the {len(in_term)} term genes hit carry a median of "
              f"{sorted(degree[g] for g in in_term)[len(in_term)//2]} terms")

    rng = random.Random(a.seed)
    bstat = load_bstat()
    if bstat and a.spatial_reps:
        draws = matched_block_draws(regs, preal, bstat, a.spatial_reps, rng)
        means, meds = [], []
        for nregs in draws:
            nby, _ = genes_for_regions(nregs, doms)
            ng = [g for g in nby if g in degree]
            if ng:
                means.append(sum(degree[g] for g in ng) / len(ng))
                meds.append(sorted(degree[g] for g in ng)[len(ng) // 2])
        means.sort()
        q = bisect.bisect_left(means, mean_h) / len(means)
        print(f"\ntest 1  B-matched null genes carry a mean of "
              f"{sum(means)/len(means):.1f} GO terms "
              f"(2.5-97.5%: {means[int(.025*len(means))]:.1f}-"
              f"{means[int(.975*len(means))]:.1f}); "
              f"catalog hits {mean_h:.1f}, percentile {100*q:.0f}")
        print("        -> annotation depth of the catalog's genes is "
              + ("TYPICAL of matched locations; depth cannot explain the excess"
                 if 0.05 < q < 0.95 else
                 "ATYPICAL; depth may contribute"))

    sets = degree_matched_sets(hits, universe, degree, a.degree_reps, rng)
    beat = sum(1 for s in sets if len(term_genes & s) >= k_obs)
    p_deg = (beat + 1) / (a.degree_reps + 1)
    exp_deg = sum(len(term_genes & s) for s in sets) / len(sets)
    print(f"\ntest 2  degree-matched gene sets: mean {exp_deg:.1f} term genes, "
          f"{beat}/{a.degree_reps} reached {k_obs}; p = {p_deg:.1e}")
    print("        -> " + ("survives a null of equally well annotated genes"
                           if p_deg < 0.05 else
                           "does NOT survive; consistent with annotation depth"))


if __name__ == "__main__":
    main()
```
