## Supplementary Information section 4 code base for "Hypothesis: A modern human range expansion ∼300,000 years ago had a critical role in Neandertal origins": great_domains.py

```
#!/usr/bin/env python3
"""GREAT-style regulatory domains from GENCODE v19, and the genes a region set hits.

GREAT (McLean et al. 2010, Nat Biotechnol 28:495) does not assign a region to the
gene it overlaps. It gives every gene a regulatory domain and assigns the region
to each gene whose domain it falls in, which is what lets a small set of mostly
non-coding intervals say anything about function at all. The domain is

  basal      TSS - 5 kb to TSS + 1 kb, strand aware, always kept;
  extension  outward from the basal domain to the nearest neighbouring basal
             domain, capped at 1 Mb either side of the TSS.

The cap and the basal sizes are GREAT's defaults. Using GENCODE v19 protein-coding
genes rather than GREAT's own hg19 gene set keeps this consistent with the gene
annotation used everywhere else in this note, at the cost of not reproducing
GREAT's numbers exactly.

Written to be imported by great_test.py as well as run on its own for the
region-count and gene-count ladder that decides which threshold is worth testing.
"""

import argparse
import csv
import gzip
import os
import re
import sys
from collections import defaultdict

REF = os.path.join(os.path.dirname(os.path.dirname(os.path.abspath(__file__))),
                   "refdata")
GTF = os.path.join(REF, "gencode.v19.annotation.gtf.gz")
ANN = os.path.join(os.path.dirname(os.path.dirname(os.path.abspath(__file__))),
                   "catalog_annotation")
GRID = os.path.join(ANN, os.environ.get("GREAT_PREAL_GRID",
                                        "preal_gridpoints.tsv"))

BASAL_UP = 5000
BASAL_DOWN = 1000
MAX_EXT = 1_000_000
# bp a scan grid point stands for. The scan moved from a 5 kb to a 1 kb lattice,
# and the matched-null machinery needs the true spacing: it tests contiguity as
# ps[i+k-1] == ps[i] + GRID_STEP*(k-1) and converts a locus span into a block
# length as span // GRID_STEP, both of which silently produce empty candidate
# pools if the constant disagrees with the file it is reading.
GRID_STEP = int(os.environ.get("GREAT_GRID_STEP", "5000"))

# hg19 autosome lengths, for the genome fraction the binomial test needs.
HG19 = {
    "1": 249250621, "2": 243199373, "3": 198022430, "4": 191154276,
    "5": 180915260, "6": 171115067, "7": 159138663, "8": 146364022,
    "9": 141213431, "10": 135534747, "11": 135006516, "12": 133851895,
    "13": 115169878, "14": 107349540, "15": 102531392, "16": 90354753,
    "17": 81195210, "18": 78077248, "19": 59128983, "20": 63025520,
    "21": 48129895, "22": 51304566,
}
GENOME = sum(HG19.values())


def load_tss(biotypes=("protein_coding",)):
    """chrom -> list of (tss, symbol, strand), one entry per gene."""
    out = defaultdict(list)
    with gzip.open(GTF, "rt") as fh:
        for line in fh:
            if line.startswith("#"):
                continue
            f = line.split("\t", 9)
            if f[2] != "gene":
                continue
            c = f[0].replace("chr", "")
            if c not in HG19:
                continue
            bt = re.search(r'gene_type "([^"]+)"', f[8]).group(1)
            if biotypes and bt not in biotypes:
                continue
            sym = re.search(r'gene_name "([^"]+)"', f[8]).group(1)
            strand = f[6]
            tss = int(f[3]) if strand == "+" else int(f[4])
            out[c].append((tss, sym, strand))
    for c in out:
        out[c].sort()
    return out


def domains(tss_by_chrom):
    """chrom -> sorted list of (start, end, symbol), GREAT basal plus extension.

    A gene's domain runs outward from its basal domain until it meets the basal
    domain of the nearest gene on that side, or 1 Mb from the TSS, whichever is
    nearer. Genes are processed in TSS order, so the neighbour on each side is
    the adjacent entry. Overlapping basal domains are left overlapping, as
    GREAT does: a region in two basal domains belongs to both genes.
    """
    out = {}
    for c, genes in tss_by_chrom.items():
        basal = []
        for tss, sym, strand in genes:
            if strand == "+":
                basal.append((tss - BASAL_UP, tss + BASAL_DOWN))
            else:
                basal.append((tss - BASAL_DOWN, tss + BASAL_UP))
        doms = []
        for i, (tss, sym, _strand) in enumerate(genes):
            bs, be = basal[i]
            lo = max(0, tss - MAX_EXT)
            if i > 0:
                lo = max(lo, basal[i - 1][1])
            hi = min(HG19[c], tss + MAX_EXT)
            if i + 1 < len(genes):
                hi = min(hi, basal[i + 1][0])
            # The basal domain is never given up, even to a closer neighbour.
            doms.append((min(lo, bs), max(hi, be), sym))
        out[c] = sorted(doms)
    return out


def total_domain_bp(doms):
    """Genome bp covered by at least one regulatory domain, and per gene."""
    per_gene = defaultdict(int)
    covered = 0
    for c, ds in doms.items():
        for s, e, sym in ds:
            per_gene[sym] += e - s
        merged = []
        for s, e, _ in sorted(ds):
            if merged and s <= merged[-1][1]:
                merged[-1][1] = max(merged[-1][1], e)
            else:
                merged.append([s, e])
        covered += sum(e - s for s, e in merged)
    return covered, per_gene


def genes_for_regions(regions, doms):
    """Symbols whose regulatory domain overlaps any region, and the hit regions.

    Returns (symbol -> set of region indices, region index -> set of symbols).
    """
    by_gene = defaultdict(set)
    by_region = defaultdict(set)
    for c, ds in doms.items():
        starts = [d[0] for d in ds]
        for ri, (rc, rs, re_) in enumerate(regions):
            if rc != c:
                continue
            # Domains are sorted by start; walk back far enough to catch any
            # domain that starts before the region but extends into it. A
            # domain is at most 2 Mb wide, so a linear scan from the first
            # candidate is bounded.
            import bisect
            j = bisect.bisect_left(starts, rs - 2 * MAX_EXT - 1)
            while j < len(ds) and ds[j][0] < re_:
                s, e, sym = ds[j]
                if s < re_ and e > rs:
                    by_gene[sym].add(ri)
                    by_region[ri].add(sym)
                j += 1
    return by_gene, by_region


def load_grid_preal():
    """(chrom, pos) -> P(real), autosomes only."""
    out = {}
    with open(GRID, encoding="utf-8") as f:
        for r in csv.DictReader(f, delimiter="\t"):
            c = r["chrom"].replace("chr", "")
            if c in HG19:
                out[(c, int(r["pos"]))] = float(r["preal"])
    return out


def regions_at_threshold(preal, t):
    """Merge gridpoints with P(real) >= t into regions; return list and sum P."""
    pts = sorted((c, p) for (c, p), v in preal.items() if v >= t)
    regions, psum = [], 0.0
    for c, p in pts:
        psum += preal[(c, p)]
        if regions and regions[-1][0] == c and p == regions[-1][2]:
            regions[-1][2] = p + GRID_STEP
        else:
            regions.append([c, p, p + GRID_STEP])
    return [tuple(r) for r in regions], psum, len(pts)


def main():
    ap = argparse.ArgumentParser()
    ap.add_argument("--thresholds", default="0.8,0.7,0.6,0.5,0.4,0.3,0.2")
    ap.add_argument("--biotypes", default="protein_coding",
                    help="comma separated GENCODE gene_type values, or 'all'")
    a = ap.parse_args()

    bts = () if a.biotypes == "all" else tuple(a.biotypes.split(","))
    tss = load_tss(bts)
    ngene = sum(len(v) for v in tss.values())
    doms = domains(tss)
    covered, per_gene = total_domain_bp(doms)
    sys.stderr.write(
        f"{ngene:,} genes ({a.biotypes}) on {len(tss)} autosomes\n"
        f"regulatory domains cover {covered/1e6:,.0f} Mb "
        f"({100*covered/GENOME:.1f}% of the autosomes); "
        f"median domain {sorted(per_gene.values())[len(per_gene)//2]/1000:,.0f} kb\n\n")

    preal = load_grid_preal()
    sys.stderr.write(f"{len(preal):,} autosomal grid points\n\n")

    print(f"{'P(real)>=':>10}{'points':>9}{'regions':>9}{'span Mb':>9}"
          f"{'E[real]':>9}{'purity':>8}{'genes':>7}{'genes/reg':>10}")
    for t in [float(x) for x in a.thresholds.split(",")]:
        regs, psum, npts = regions_at_threshold(preal, t)
        by_gene, by_region = genes_for_regions(regs, doms)
        span = sum(e - s for _, s, e in regs)
        print(f"{t:>10.2f}{npts:>9,}{len(regs):>9,}{span/1e6:>9.2f}"
              f"{psum:>9.0f}{psum/max(npts,1):>8.2f}{len(by_gene):>7,}"
              f"{sum(len(v) for v in by_region.values())/max(len(regs),1):>10.1f}")


if __name__ == "__main__":
    main()
```
