## Supplementary Information section 4 code base for "Hypothesis: A modern human range expansion ∼300,000 years ago had a critical role in Neandertal origins": great_matched_100k.py

```
#!/usr/bin/env python3
"""Push the B-matched permutation null from 10,000 to 100,000 replicates.

great_test.py --matched N reports (beat + 1) / (N + 1), so 10,000 replicates
put a floor of 1e-4 under the permutation p value. Eight of the nine categories
in Table S4.3 sit on that floor, which is only a factor of 2.5 below the
Bonferroni threshold for the 200 categories permuted. 100,000 replicates move
the floor to 1e-5 and let those rows be quoted as clearing Bonferroni by a
factor of 25 rather than 2.5.

Nothing about the null changes. This is the same matched_block_draws from
great_test.py, drawing the same blocks from the same B bins; only the number of
draws and the wall clock differ.

Speed. genes_for_regions rebuilds a per chromosome list of domain starts on
every call, about 20,000 elements over 22 chromosomes, which at 100,000 calls
dominates everything else. Here the starts are built once and the regions are
bucketed by chromosome so only chromosomes carrying a region are visited.
--verify asserts that this returns exactly what great_domains.genes_for_regions
returns, on the observed catalog and on null draws, before any of it is used.

Run as ten independent chunks with different seeds and aggregate; permutation
counts are additive over independent draws.

  for i in $(seq 0 9); do
    python3 scripts/great_matched_100k.py --regions REGIONS.tsv \
        --nrep 10000 --seed $((100 + i)) -o /tmp/counts.$i.tsv &
  done; wait
  python3 scripts/great_matched_100k.py --aggregate /tmp/counts.*.tsv \
      --results catalog_annotation/great_go_catalog132_matched10k.tsv \
      -o catalog_annotation/great_go_catalog132_matched100k.tsv
"""

import argparse
import bisect
import collections
import csv
import glob
import os
import random
import sys

from scipy.stats import hypergeom

sys.path.insert(0, os.path.dirname(os.path.abspath(__file__)))
from great_domains import (ANN, GRID_STEP, HG19, domains,  # noqa: E402
                           genes_for_regions, load_grid_preal, load_tss)
from great_test import (ancestors, load_bstat, load_gaf,  # noqa: E402
                        load_obo, matched_block_draws, merge_regions)

REF = os.path.join(os.path.dirname(os.path.dirname(os.path.abspath(__file__))),
                   "refdata")


def build_index(doms):
    """chrom -> (starts, ends, syms) as parallel lists, sorted by start."""
    idx = {}
    for c, ds in doms.items():
        ds = sorted(ds)
        idx[c] = ([d[0] for d in ds], [d[1] for d in ds], [d[2] for d in ds])
    return idx


def hit_genes(regions, idx, span=2_000_001):
    """Symbols whose regulatory domain overlaps any region.

    Same answer as great_domains.genes_for_regions, first return value keyed to
    a set of symbols rather than symbol -> region indices, which is all the
    permutation needs.
    """
    out = set()
    by_chrom = collections.defaultdict(list)
    for c, s, e in regions:
        by_chrom[c].append((s, e))
    for c, rs_list in by_chrom.items():
        got = idx.get(c)
        if got is None:
            continue
        starts, ends, syms = got
        n = len(starts)
        for rs, re_ in rs_list:
            j = bisect.bisect_left(starts, rs - span)
            while j < n and starts[j] < re_:
                if ends[j] > rs:
                    out.add(syms[j])
                j += 1
    return out


def setup(regions_path, merge_gap, min_genes, max_genes, keep_iea, aspects):
    terms, alt = load_obo(os.path.join(REF, "go", "go-basic.obo"))
    anc = ancestors(terms)
    # load_gaf's fifth argument is drop_iea, so it is the negation of the flag.
    sym2go = load_gaf(terms, alt, anc, set(aspects), not keep_iea)
    tss = load_tss(("protein_coding",))
    doms = domains(tss)
    universe = sorted({s for c in doms for _, _, s in doms[c]} & set(sym2go))
    go2genes = collections.defaultdict(set)
    for g in universe:
        for t in sym2go[g]:
            go2genes[t].add(g)
    tested = {t: g for t, g in go2genes.items()
              if min_genes <= len(g) <= max_genes}

    regs = []
    with open(regions_path, encoding="utf-8") as f:
        for r in csv.DictReader(f, delimiter="\t"):
            c = r["chrom"].replace("chr", "")
            if c in HG19:
                regs.append((c, int(r["start"]), int(r["end"])))
    regs = merge_regions(regs, merge_gap)
    return doms, universe, tested, regs


def main():
    ap = argparse.ArgumentParser()
    ap.add_argument("--regions")
    ap.add_argument("--nrep", type=int, default=10000)
    ap.add_argument("--seed", type=int, default=100)
    ap.add_argument("--merge-gap", type=int, default=100000)
    ap.add_argument("--min-genes", type=int, default=10)
    ap.add_argument("--max-genes", type=int, default=1000)
    ap.add_argument("--aspects", default="P")
    ap.add_argument("--keep-iea", action="store_true")
    ap.add_argument("--top", type=int, default=200)
    ap.add_argument("--verify", action="store_true")
    ap.add_argument("--aggregate", nargs="*", default=None)
    ap.add_argument("--results", default=None)
    ap.add_argument("-o", "--out", required=True)
    a = ap.parse_args()

    # ------------------------------------------------------------- aggregate
    if a.aggregate is not None:
        files = []
        for pat in a.aggregate:
            files.extend(sorted(glob.glob(pat)) or [pat])
        beat, nrep = collections.Counter(), 0
        nloci, ngenes, seeds = [], [], []
        for fn in files:
            with open(fn, encoding="utf-8") as f:
                rd = csv.DictReader(f, delimiter="\t")
                seen = None
                for r in rd:
                    beat[r["go_id"]] += int(r["beat"])
                    seen = r
                if seen is None:
                    sys.exit(f"{fn} is empty")
                nrep += int(seen["nrep"])
                nloci.append(float(seen["mean_loci"]))
                ngenes.append(float(seen["mean_genes"]))
                seeds.append(seen["seed"])
        sys.stderr.write(
            f"{len(files)} chunks, seeds {','.join(seeds)}, {nrep:,} "
            f"replicates; null averages {sum(nloci)/len(nloci):.1f} loci and "
            f"{sum(ngenes)/len(ngenes):.1f} annotated genes\n")
        if not a.results:
            sys.exit("--aggregate needs --results")
        with open(a.results, encoding="utf-8") as f:
            rows = list(csv.DictReader(f, delimiter="\t"))
        cols = list(rows[0].keys())
        # --results only has to carry the observed statistics. p_matched is
        # recomputed here from the chunk counts whatever it held before, so
        # requiring it to already exist meant the aggregation could only run
        # against a great_test.py --matched run, and died with a bare KeyError
        # on the cheap --matched 0 output that is otherwise identical.
        if "p_matched" not in cols:
            cols.append("p_matched")
        for r in rows:
            if r["go_id"] in beat or r.get("p_matched") not in (None, ""):
                k = beat.get(r["go_id"], 0)
                r["p_matched"] = f"{(k + 1) / (nrep + 1):.3g}"
                r["n_beat"] = k
            else:
                r["p_matched"], r["n_beat"] = "", ""
        if "n_beat" not in cols:
            cols.insert(cols.index("p_matched") + 1, "n_beat")
        with open(a.out, "w", newline="", encoding="utf-8") as f:
            w = csv.DictWriter(f, fieldnames=cols, delimiter="\t",
                               extrasaction="ignore")
            w.writeheader()
            w.writerows(rows)
        floor = 1 / (nrep + 1)
        print(f"wrote {a.out}; floor is {floor:.3g} "
              f"(Bonferroni for {a.top} categories is {0.05/a.top:.2g})")
        for r in rows[:12]:
            if r["p_matched"]:
                print(f"  {r['name'][:58]:<60} beat {str(r['n_beat']):>6} "
                      f"p {r['p_matched']}")
        return 0

    # ----------------------------------------------------------------- setup
    if not a.regions:
        sys.exit("--regions is required unless --aggregate is used")
    doms, universe, tested, regs = setup(
        a.regions, a.merge_gap, a.min_genes, a.max_genes, a.keep_iea,
        a.aspects)
    idx = build_index(doms)
    preal = load_grid_preal()
    bstat = load_bstat()
    if not bstat:
        sys.exit("no B map found; the matched null needs one")
    rng = random.Random(a.seed)

    hits = hit_genes(regs, idx) & set(universe)
    sys.stderr.write(f"{len(regs)} loci, {len(hits)} annotated genes, "
                     f"universe {len(universe)}, {len(tested)} terms\n")

    # -------------------------------------------------- verify the fast path
    if a.verify:
        ref_by_gene, _ = genes_for_regions(regs, doms)
        assert set(ref_by_gene) == hit_genes(regs, idx), "observed set differs"
        vrng = random.Random(12345)
        for i, nregs in enumerate(matched_block_draws(
                regs, preal, bstat, 50, vrng), 1):
            ref, _ = genes_for_regions(nregs, doms)
            fast = hit_genes(nregs, idx)
            assert set(ref) == fast, f"replicate {i} differs"
        sys.stderr.write("verify: fast path matches genes_for_regions on the "
                         "observed catalog and 50 null draws\n")

    # ------------------------------------------------------- the top N terms
    if not a.results:
        sys.exit("--results (the great_test.py output) is required")
    with open(a.results, encoding="utf-8") as f:
        rows = list(csv.DictReader(f, delimiter="\t"))[:a.top]
    top = [(r["go_id"], float(r["p_hyper"])) for r in rows]
    N = len(universe)

    # The permutation compares null p values against the observed ones in the
    # results file, so this setup has to be the same setup that produced them.
    # Recomputing p_hyper is a strict check on every input at once: universe
    # size, annotated gene count, IEA handling, aspect and term membership.
    missing = [gid for gid, _ in top if gid not in tested]
    if missing:
        sys.exit(f"{len(missing)} of the top {a.top} terms are absent from "
                 f"this setup, e.g. {missing[:3]}; configuration differs from "
                 f"the run that produced {a.results}")
    sizes = {gid: len(tested[gid]) for gid, _ in top}
    for r in rows:
        gid = r["go_id"]
        mine = hypergeom.sf(int(r["k_genes"]) - 1, N, sizes[gid], len(hits))
        want = float(r["p_hyper"])
        if abs(mine - want) > 1e-9 * max(want, 1e-12) + 1e-15:
            sys.exit(f"p_hyper mismatch for {gid}: recomputed {mine:.6e} "
                     f"against {want:.6e} in {a.results}; setup differs")
    sys.stderr.write(f"observed statistics reproduced for all {len(rows)} "
                     f"top terms\n")

    # ----------------------------------------------------------- permutation
    beat = collections.Counter()
    nloci, ngenes = 0, 0
    draws = matched_block_draws(regs, preal, bstat, a.nrep, rng)
    for i, nregs in enumerate(draws, 1):
        nh = hit_genes(nregs, idx) & set(universe)
        nloci += len(nregs)
        ngenes += len(nh)
        for gid, p_obs in top:
            kn = len(tested[gid] & nh)
            if kn and hypergeom.sf(kn - 1, N, sizes[gid], len(nh)) <= p_obs:
                beat[gid] += 1
        if i % 1000 == 0:
            sys.stderr.write(f"  seed {a.seed}: {i}/{a.nrep}\n")

    with open(a.out, "w", newline="", encoding="utf-8") as f:
        w = csv.writer(f, delimiter="\t")
        w.writerow(["go_id", "beat", "nrep", "seed", "mean_loci",
                    "mean_genes"])
        for gid, _ in top:
            w.writerow([gid, beat.get(gid, 0), a.nrep, a.seed,
                        f"{nloci/a.nrep:.2f}", f"{ngenes/a.nrep:.2f}"])
    sys.stderr.write(f"seed {a.seed}: wrote {a.out}\n")
    return 0


if __name__ == "__main__":
    sys.exit(main())
```
