## Supplementary Information section 4 code base for "Hypothesis: A modern human range expansion ∼300,000 years ago had a critical role in Neandertal origins": great_negative_control.py

```
#!/usr/bin/env python3
"""Does the GO pipeline manufacture the proteasome signal from any swept-looking set?

Three region sets go through identical code:

  case            P(real) >= 0.5, the set the enrichment was found in.
  mirror-exact    the published control regions, which are the catalog rule with
                  Denisovan and Neandertal exchanged. Only 26 regions, because
                  the asymmetry between the two is the catalog's central claim,
                  so this is the right comparison but an underpowered one.
  mirror-matched  the same exchange, loosened on the posterior mean until it
                  yields as many loci as the case set. This is powered, and is
                  the set that matters: it is depressed TMRCA of the same depth
                  and extent, on the Denisovan side instead.

If the enrichment reflects Neandertal-shared sweeps it should be absent from
both mirrors. If it reflects a generic property of regions where the ARG gives
shallow coalescence, it should be present in mirror-matched too, and the result
is an artefact of the scan rather than a finding about modern humans.
"""

import argparse
import collections
import csv
import os
import sys

from scipy.stats import hypergeom

sys.path.insert(0, os.path.dirname(os.path.abspath(__file__)))
from great_domains import (ANN, GRID_STEP, HG19, domains,  # noqa: E402
                           genes_for_regions, load_tss)
from great_test import (ancestors, bh, load_gaf, load_obo,  # noqa: E402
                        merge_regions)

# Overridable so the same script runs on either grid; the defaults reproduce the
# published 5 kb run. GREAT_PREAL_GRID and GREAT_GRID_STEP must be set together,
# since the grid file and the step it implies have to agree, and GREAT_CONTROL
# has to name the control regions from the same catalog build as that grid.
GRID = os.path.join(ANN, os.environ.get("GREAT_PREAL_GRID",
                                        "preal_gridpoints.tsv"))
CTRL = os.environ.get(
    "GREAT_CONTROL",
    os.path.join(ANN, "catalog_final_median_x625_y700_auto.control_regions.tsv"))
TERM = "GO:0043161"
Y_MIN = 700.0


def load_full_grid():
    rows = []
    with open(GRID, encoding="utf-8") as f:
        for r in csv.DictReader(f, delimiter="\t"):
            c = r["chrom"].replace("chr", "")
            if c not in HG19:
                continue
            rows.append((c, int(r["pos"]), float(r["nea_mean_ka"]),
                         float(r["den_mean_ka"]), float(r["preal"])))
    return rows


def to_regions(pts, gap):
    regs = []
    for c, p in sorted(pts):
        if regs and regs[-1][0] == c and p == regs[-1][2]:
            regs[-1][2] = p + GRID_STEP
        else:
            regs.append([c, p, p + GRID_STEP])
    return merge_regions([tuple(r) for r in regs], gap)


def read_regions(path, gap):
    out = []
    with open(path, encoding="utf-8") as f:
        for r in csv.DictReader(f, delimiter="\t"):
            c = r["chrom"].replace("chr", "")
            if c in HG19:
                out.append((c, int(r["start"]), int(r["end"])))
    return merge_regions(out, gap)


def evaluate(name, regs, doms, universe, sym2go, go2genes, terms, tested):
    by_gene, by_region = genes_for_regions(regs, doms)
    hits = sorted(set(by_gene) & set(universe))
    if not hits:
        print(f"\n### {name}: {len(regs)} loci, no annotated genes")
        return
    tg = {g for g in universe if TERM in sym2go[g]}
    k = len(tg & set(hits))
    exp = len(tg) * len(hits) / len(universe)
    p = hypergeom.sf(k - 1, len(universe), len(tg), len(hits)) if k else 1.0
    print(f"\n### {name}: {len(regs)} loci, {len(hits)} annotated genes")
    print(f"    proteasome-mediated ubiquitin-dependent catabolism: "
          f"{k} genes vs {exp:.1f} expected, p = {p:.2e}"
          + (f", genes {', '.join(sorted(tg & set(hits)))}" if k else ""))
    rows = []
    for t, g in tested.items():
        kk = len(g & set(hits))
        if kk < 2:
            continue
        rows.append((hypergeom.sf(kk - 1, len(universe), len(g), len(hits)),
                     t, kk, len(g), len(g) * len(hits) / len(universe)))
    if not rows:
        return
    qs = bh([r[0] for r in rows])
    ranked = sorted(zip(rows, qs), key=lambda x: x[0][0])
    print(f"    strongest terms (of {len(rows)} tested, best q = {min(qs):.3f}):")
    for (pv, t, kk, n, e), q in ranked[:5]:
        print(f"      {terms[t]['name'][:52]:<54}{kk}/{n:<6} exp {e:>4.1f}  "
              f"p {pv:.1e}  q {q:.3f}")


def main():
    ap = argparse.ArgumentParser()
    ap.add_argument("--threshold", type=float, default=0.5)
    ap.add_argument("--merge-gap", type=int, default=100000)
    ap.add_argument("--min-genes", type=int, default=10)
    ap.add_argument("--max-genes", type=int, default=1000)
    a = ap.parse_args()

    terms, alt = load_obo(os.path.join(
        os.path.dirname(os.path.dirname(os.path.abspath(__file__))),
        "refdata", "go", "go-basic.obo"))
    anc = ancestors(terms)
    sym2go = load_gaf(terms, alt, anc, {"P"}, True)
    tss = load_tss(("protein_coding",))
    doms = domains(tss)
    universe = sorted({s for c in doms for _, _, s in doms[c]} & set(sym2go))
    go2genes = collections.defaultdict(set)
    for g in universe:
        for t in sym2go[g]:
            go2genes[t].add(g)
    tested = {t: g for t, g in go2genes.items()
              if a.min_genes <= len(g) <= a.max_genes}

    grid = load_full_grid()
    case = to_regions([(c, p) for c, p, _n, _d, pr in grid
                       if pr >= a.threshold], a.merge_gap)
    target = len(case)

    # Loosen the mirrored rule on the posterior mean until it gives as many
    # loci as the case set, so the two are compared at equal catalog size.
    best = None
    for x in range(400, 1201, 5):
        pts = [(c, p) for c, p, n, d, _pr in grid if d <= x and n >= Y_MIN]
        regs = to_regions(pts, a.merge_gap)
        if best is None or abs(len(regs) - target) < abs(best[1] - target):
            best = (x, len(regs), regs)
        if len(regs) >= target:
            break
    x, nloci, mirror_matched = best
    sys.stderr.write(f"mirror matched at T_Afr+Den <= {x} ka and "
                     f"T_Afr+Nea >= {Y_MIN:.0f} ka: {nloci} loci "
                     f"(target {target})\n")

    evaluate(f"case  P(real) >= {a.threshold}", case, doms, universe,
             sym2go, go2genes, terms, tested)
    evaluate("mirror-exact  published control regions",
             read_regions(CTRL, a.merge_gap), doms, universe,
             sym2go, go2genes, terms, tested)
    evaluate(f"mirror-matched  T_Afr+Den <= {x} ka, T_Afr+Nea >= 700 ka",
             mirror_matched, doms, universe, sym2go, go2genes, terms, tested)


if __name__ == "__main__":
    main()
```
