## Supplementary Information section 4 code base for "Hypothesis: A modern human range expansion ∼300,000 years ago had a critical role in Neandertal origins": great_test.py

```
#!/usr/bin/env python3
"""GO enrichment on the sweep catalog, GREAT style, with a matched background.

Two tests are run for every GO term, as GREAT does, because each is blind to a
different artefact:

  binomial      over regions. The null probability is the fraction of the
                autosomes covered by the regulatory domains of the term's genes,
                and the count is the number of catalog regions falling in them.
                This is insensitive to how many genes a region happens to sit
                near, but it assumes regions are placed uniformly at random.

  hypergeometric  over genes. The universe is every annotated gene with a
                domain, and the draw is the genes the catalog hits. This is
                insensitive to region size but double counts gene-dense loci.

GREAT reports a term only when both agree, and that convention is kept here.

The uniform-placement assumption behind the binomial is wrong for this catalog
in a specific and known direction: a TMRCA-depression scan finds regions where
diversity is low, and diversity is low where background selection is strong,
which is near genes and in low-recombination regions. Against a whole-genome
null, any gene-linked category is therefore expected to come out enriched
whether or not it was swept. --matched replaces the null with grid points drawn
to match the catalog's own distribution of the B statistic, which removes most
of that confound at the cost of being conservative: if sweeps really do target
conserved, gene-rich sequence, matching on B removes some real signal too.
Both are reported, and the honest reading is the matched one.
"""

import argparse
import bisect
import csv
import gzip
import os
import random
import sys
from collections import defaultdict

import numpy as np
from scipy.stats import binom, hypergeom

sys.path.insert(0, os.path.dirname(os.path.abspath(__file__)))
from great_domains import (ANN, GENOME, GRID_STEP, HG19, domains,  # noqa: E402
                           genes_for_regions, load_grid_preal, load_tss,
                           regions_at_threshold)

REF = os.path.join(os.path.dirname(os.path.dirname(os.path.abspath(__file__))),
                   "refdata", "go")
OBO = os.path.join(REF, "go-basic.obo")
GAF = os.path.join(REF, "goa_human.gaf.gz")
ASPECT = {"P": "biological_process", "F": "molecular_function",
          "C": "cellular_component"}


def load_obo(path):
    """id -> (name, namespace, parents). go-basic is acyclic over is_a/part_of."""
    terms, cur = {}, None
    with open(path, encoding="utf-8") as f:
        for line in f:
            line = line.rstrip("\n")
            if line == "[Term]":
                cur = {"parents": set(), "alt": []}
            elif line.startswith("[") and line != "[Term]":
                cur = None
            elif cur is not None and line:
                k, _, v = line.partition(": ")
                if k == "id":
                    cur["id"] = v
                elif k == "name":
                    cur["name"] = v
                elif k == "namespace":
                    cur["ns"] = v
                elif k == "alt_id":
                    cur["alt"].append(v)
                elif k == "is_a":
                    cur["parents"].add(v.split(" ! ")[0])
                elif k == "relationship" and v.startswith("part_of "):
                    cur["parents"].add(v.split()[1])
                elif k == "is_obsolete" and v == "true":
                    cur["obsolete"] = True
            elif cur is not None and not line and "id" in cur:
                if not cur.get("obsolete"):
                    terms[cur["id"]] = cur
                cur = None
    alt = {a: t["id"] for t in terms.values() for a in t["alt"]}
    return terms, alt


def ancestors(terms):
    """id -> frozenset of itself and all is_a/part_of ancestors, memoised."""
    cache = {}

    def walk(t, stack):
        if t in cache:
            return cache[t]
        if t in stack or t not in terms:
            return frozenset([t]) if t in terms else frozenset()
        stack.add(t)
        acc = {t}
        for p in terms[t]["parents"]:
            acc |= walk(p, stack)
        stack.discard(t)
        cache[t] = frozenset(acc)
        return cache[t]

    return {t: walk(t, set()) for t in terms}


def load_gaf(terms, alt, anc, aspects, drop_iea):
    """symbol -> frozenset of GO ids, propagated to ancestors."""
    g = defaultdict(set)
    kept = skipped = 0
    with gzip.open(GAF, "rt") as f:
        for line in f:
            if line.startswith("!"):
                continue
            c = line.split("\t")
            if len(c) < 15:
                continue
            sym, qual, goid, ev, asp = c[2], c[3], c[4], c[6], c[8]
            if "NOT" in qual or asp not in aspects:
                skipped += 1
                continue
            if drop_iea and ev == "IEA":
                skipped += 1
                continue
            goid = alt.get(goid, goid)
            if goid not in terms:
                skipped += 1
                continue
            g[sym] |= anc[goid]
            kept += 1
    sys.stderr.write(f"GAF: {kept:,} annotations kept, {skipped:,} skipped, "
                     f"{len(g):,} symbols\n")
    return {k: frozenset(v) for k, v in g.items()}


def term_genome_fraction(term_genes, gene_domains):
    """Fraction of the autosomes covered by the union of a term's domains."""
    per_chrom = defaultdict(list)
    for sym in term_genes:
        for c, s, e in gene_domains.get(sym, ()):
            per_chrom[c].append((s, e))
    total = 0
    for c, iv in per_chrom.items():
        iv.sort()
        ce = -1
        for s, e in iv:
            if s > ce:
                total += e - s
                ce = e
            elif e > ce:
                total += e - ce
                ce = e
    return total / GENOME


def bh(pvals):
    """Benjamini-Hochberg q values, order preserved."""
    n = len(pvals)
    order = sorted(range(n), key=lambda i: pvals[i])
    q = [0.0] * n
    prev = 1.0
    for rank, i in enumerate(reversed(order), 1):
        v = pvals[i] * n / (n - rank + 1)
        prev = min(prev, v)
        q[i] = min(1.0, prev)
    return q


def load_bstat():
    """(chrom, pos) -> B, from whichever B map is present.

    The map has to be on the same lattice as the scan, because the matched null
    reads B at grid points by exact (chrom, pos) key: a 5 kb map queried on a
    1 kb grid would miss four of every five points and quietly shrink the
    candidate pool. GREAT_BSTAT names the file when it is not the default.
    """
    override = os.environ.get("GREAT_BSTAT")
    names = ((override,) if override else
             ("bstat.phastCons_bestfit.5kb.tsv", "bstat.CADD_bestfit.5kb.tsv"))
    for name in names:
        p = os.path.join(ANN, name)
        if not os.path.exists(p):
            continue
        out = {}
        with open(p, encoding="utf-8") as f:
            rd = csv.DictReader(f, delimiter="\t")
            bcol = next((c for c in rd.fieldnames
                         if c.lower() in ("b", "bstat", "b_stat", "bvalue")), None)
            if bcol is None:
                continue
            for r in rd:
                c = r["chrom"].replace("chr", "")
                try:
                    out[(c, int(r["pos"]))] = float(r[bcol])
                except (ValueError, KeyError):
                    continue
        if out:
            sys.stderr.write(f"B statistic from {name}: {len(out):,} points\n")
            return out
    sys.stderr.write("no B map found; --matched unavailable\n")
    return None


def block_candidates(preal, bstat, ks):
    """For each block length k, every contiguous k-point window and its mean B.

    Returns k -> (list of (chrom, start_pos), list of mean B), index aligned.
    Only windows whose k grid points are contiguous in the scan are kept, so a
    null locus lies on the same footprint the real scan covered.

    Done with prefix sums rather than by re-reading each window. The obvious
    nested loop costs O(n * sum(k)), which was tolerable on the 5 kb lattice
    (~510k points, blocks of a few) but is not on the 1 kb one: ~2.6M points
    with blocks up to ~100 long is of order 10^10 dict lookups. Cumulative sums
    over each maximal contiguous run make every k a single vectorised slice, so
    the cost is O(n) per k and the result is identical.
    """
    by_chrom = defaultdict(list)
    for (c, p) in preal:
        by_chrom[c].append(p)

    # One maximal contiguous run at a time: positions GRID_STEP apart with a B
    # value at every one of them. Windows may not straddle a break, and a point
    # with no B value is a break, which is what the old `any(v is None)` test
    # enforced window by window.
    # Insertion order, not sorted: the pools built from this feed rng.choice by
    # index, so reordering them would change which null blocks a given seed
    # draws and make the permutation p values irreproducible against earlier
    # runs, for no gain.
    runs = []  # (chrom, positions array, B array)
    for c in by_chrom:
        ps = sorted(by_chrom[c])
        cur_p, cur_b = [], []
        for p in ps:
            b = bstat.get((c, p))
            if b is None:
                if cur_p:
                    runs.append((c, np.asarray(cur_p), np.asarray(cur_b)))
                cur_p, cur_b = [], []
                continue
            if cur_p and p != cur_p[-1] + GRID_STEP:
                runs.append((c, np.asarray(cur_p), np.asarray(cur_b)))
                cur_p, cur_b = [], []
            cur_p.append(p)
            cur_b.append(b)
        if cur_p:
            runs.append((c, np.asarray(cur_p), np.asarray(cur_b)))

    out = {}
    for k in ks:
        spots, means = [], []
        for c, ps, bs in runs:
            if len(ps) < k:
                continue
            csum = np.concatenate(([0.0], np.cumsum(bs)))
            mk = (csum[k:] - csum[:-k]) / k
            spots.extend((c, int(p)) for p in ps[:len(ps) - k + 1])
            means.extend(mk.tolist())
        out[k] = (spots, means)
    return out


def matched_block_draws(loci, preal, bstat, nrep, rng, nbin=20):
    """Relocate each locus to a random block of equal length and similar mean B.

    Matching point by point is not enough. A scattered set of points merges into
    far more loci than the real catalog does, so the gene draw is much larger and
    the p values are not comparable. Here the number of loci, and each locus's
    length, are preserved by construction, and only position changes.
    """
    want = []
    for c, s, e in loci:
        k = max(1, (e - s) // GRID_STEP)
        pts = [bstat.get((c, s + GRID_STEP * j)) for j in range(k)]
        pts = [v for v in pts if v is not None]
        if not pts:
            continue
        want.append((k, sum(pts) / len(pts)))
    cand = block_candidates(preal, bstat, sorted({k for k, _ in want}))
    binned = {}
    for k, (spots, means) in cand.items():
        srt = sorted(means)
        if not srt:
            continue
        edges = [srt[int(len(srt) * i / nbin)] for i in range(1, nbin)]
        pool = defaultdict(list)
        for idx, m in enumerate(means):
            pool[bisect.bisect_right(edges, m)].append(idx)
        binned[k] = (spots, edges, pool)
    out = []
    for _ in range(nrep):
        regs = []
        for k, m in want:
            if k not in binned:
                continue
            spots, edges, pool = binned[k]
            b = bisect.bisect_right(edges, m)
            src = pool.get(b) or [i for v in pool.values() for i in v]
            c, s = spots[rng.choice(src)]
            regs.append((c, s, s + GRID_STEP * k))
        out.append(regs)
    return out


def points_to_regions(pts, gap=0):
    regs = []
    for c, p in sorted(pts):
        if regs and regs[-1][0] == c and p <= regs[-1][2] + gap:
            regs[-1][2] = p + GRID_STEP
        else:
            regs.append([c, p, p + GRID_STEP])
    return [tuple(r) for r in regs]


def merge_regions(regs, gap):
    """Join regions within gap bp so one sweep contributes one draw.

    The binomial test treats regions as independent placements. Contiguous and
    near-contiguous grid points come from the same depression in the same ARG
    window, so without this the same sweep is counted many times and the
    binomial p value is inflated by orders of magnitude.
    """
    out = []
    for c, s, e in sorted(regs):
        if out and out[-1][0] == c and s <= out[-1][2] + gap:
            out[-1][2] = max(out[-1][2], e)
        else:
            out.append([c, s, e])
    return [tuple(r) for r in out]


def main():
    ap = argparse.ArgumentParser()
    ap.add_argument("--threshold", type=float, default=0.3)
    ap.add_argument("--regions", default=None,
                    help="region TSV (chrom/start/end) to test instead of "
                         "thresholding P(real); the published catalog file")
    ap.add_argument("--aspects", default="P")
    ap.add_argument("--keep-iea", action="store_true")
    ap.add_argument("--min-genes", type=int, default=10)
    ap.add_argument("--max-genes", type=int, default=1000)
    ap.add_argument("--merge-gap", type=int, default=100000,
                    help="join regions within this many bp into one locus")
    ap.add_argument("--matched", type=int, default=0,
                    help="replicates for the B-matched null (0 = skip)")
    ap.add_argument("--seed", type=int, default=1)
    ap.add_argument("--top", type=int, default=25)
    ap.add_argument("-o", "--out", default=None)
    a = ap.parse_args()

    terms, alt = load_obo(OBO)
    anc = ancestors(terms)
    sym2go = load_gaf(terms, alt, anc, set(a.aspects), not a.keep_iea)

    tss = load_tss(("protein_coding",))
    doms = domains(tss)
    gene_domains = defaultdict(list)
    for c, ds in doms.items():
        for s, e, sym in ds:
            gene_domains[sym].append((c, s, e))

    universe = sorted(set(gene_domains) & set(sym2go))
    sys.stderr.write(f"universe: {len(universe):,} genes with a domain and "
                     f"{a.aspects} annotation\n")

    go2genes = defaultdict(set)
    for sym in universe:
        for t in sym2go[sym]:
            go2genes[t].add(sym)
    tested = {t: g for t, g in go2genes.items()
              if a.min_genes <= len(g) <= a.max_genes}
    sys.stderr.write(f"{len(tested):,} terms with {a.min_genes}-{a.max_genes} genes\n")

    preal = load_grid_preal()
    if a.regions:
        regs, psum, npts = [], 0.0, 0
        with open(a.regions, encoding="utf-8") as f:
            for r in csv.DictReader(f, delimiter="\t"):
                c = r["chrom"].replace("chr", "")
                if c not in HG19:
                    continue
                s0, e0 = int(r["start"]), int(r["end"])
                regs.append((c, s0, e0))
                for q in range(s0, e0, GRID_STEP):
                    if (c, q) in preal:
                        psum += preal[(c, q)]
                        npts += 1
    else:
        regs, psum, npts = regions_at_threshold(preal, a.threshold)
    raw = len(regs)
    regs = merge_regions(regs, a.merge_gap)
    by_gene, by_region = genes_for_regions(regs, doms)
    hits = sorted(set(by_gene) & set(universe))
    tag = a.regions if a.regions else f"P(real) >= {a.threshold}"
    sys.stderr.write(f"\n{tag}: {npts:,} points -> {raw:,} "
                     f"contiguous regions -> {len(regs):,} loci at "
                     f"{a.merge_gap/1000:.0f} kb merge, E[real] {psum:.0f}, "
                     f"{len(hits):,} annotated genes hit\n\n")

    rows = []
    for t, tg in tested.items():
        k_gene = len(tg & set(hits))
        if k_gene < 2:
            continue
        p_hyper = hypergeom.sf(k_gene - 1, len(universe), len(tg), len(hits))
        frac = term_genome_fraction(tg, gene_domains)
        k_reg = sum(1 for ri, syms in by_region.items() if syms & tg)
        p_binom = binom.sf(k_reg - 1, len(regs), frac) if frac > 0 else 1.0
        rows.append({
            "go_id": t, "name": terms[t]["name"], "aspect": terms[t]["ns"],
            "n_genes_term": len(tg), "k_genes": k_gene,
            "exp_genes": len(tg) * len(hits) / len(universe),
            "p_hyper": p_hyper, "k_regions": k_reg,
            "exp_regions": frac * len(regs), "genome_frac": frac,
            "p_binom": p_binom,
            "genes": ",".join(sorted(tg & set(hits))),
        })
    if not rows:
        sys.exit("no term had two or more genes hit")

    for key, qk in (("p_hyper", "q_hyper"), ("p_binom", "q_binom")):
        qs = bh([r[key] for r in rows])
        for r, q in zip(rows, qs):
            r[qk] = q
    for r in rows:
        r["q_both"] = max(r["q_hyper"], r["q_binom"])
    rows.sort(key=lambda r: (r["q_both"], r["p_hyper"]))

    if a.matched:
        bstat = load_bstat()
        if bstat:
            rng = random.Random(a.seed)
            draws = matched_block_draws(regs, preal, bstat, a.matched, rng)
            top = rows[:200]
            beat = defaultdict(int)
            nloci, ngenes = [], []
            for i, nregs in enumerate(draws, 1):
                nby_gene, _ = genes_for_regions(nregs, doms)
                nhits = set(nby_gene) & set(universe)
                nloci.append(len(nregs))
                ngenes.append(len(nhits))
                for r in top:
                    tg = tested[r["go_id"]]
                    kn = len(tg & nhits)
                    pn = hypergeom.sf(kn - 1, len(universe), len(tg), len(nhits))
                    if pn <= r["p_hyper"]:
                        beat[r["go_id"]] += 1
                if i % 10 == 0:
                    sys.stderr.write(f"  matched replicate {i}/{a.matched}\n")
            sys.stderr.write(
                f"matched null: {sum(nloci)/len(nloci):.0f} loci and "
                f"{sum(ngenes)/len(ngenes):.0f} annotated genes per replicate, "
                f"against {len(regs)} loci and {len(hits)} genes observed\n")
            ids = {r["go_id"] for r in top}
            for r in rows:
                r["p_matched"] = (
                    round((beat.get(r["go_id"], 0) + 1) / (a.matched + 1), 5)
                    if r["go_id"] in ids else "")

    cols = ["go_id", "name", "aspect", "n_genes_term", "k_genes", "exp_genes",
            "p_hyper", "q_hyper", "k_regions", "exp_regions", "genome_frac",
            "p_binom", "q_binom", "q_both"]
    if a.matched:
        cols.append("p_matched")
    cols.append("genes")
    out = a.out or os.path.join(ANN, f"great_go_preal{int(a.threshold*100)}.tsv")
    with open(out, "w", newline="", encoding="utf-8") as f:
        w = csv.DictWriter(f, fieldnames=cols, delimiter="\t", extrasaction="ignore")
        w.writeheader()
        for r in rows:
            w.writerow(r)
    sys.stderr.write(f"wrote {out} ({len(rows):,} terms)\n\n")

    hdr = (f"{'GO term':<52}{'k/term':>10}{'exp':>7}{'p_hyper':>10}"
           f"{'q_hyp':>8}{'reg':>5}{'exp':>7}{'p_binom':>10}{'q_both':>8}")
    print(hdr)
    print("-" * len(hdr))
    for r in rows[:a.top]:
        nm = r["name"][:50]
        print(f"{nm:<52}{r['k_genes']}/{r['n_genes_term']:<8}"
              f"{r['exp_genes']:>6.1f}{r['p_hyper']:>10.2e}{r['q_hyper']:>8.3f}"
              f"{r['k_regions']:>5}{r['exp_regions']:>7.1f}"
              f"{r['p_binom']:>10.2e}{r['q_both']:>8.3f}")


if __name__ == "__main__":
    main()
```
