## Supplementary Information section 4 code base for "Hypothesis: A modern human range expansion ∼300,000 years ago had a critical role in Neandertal origins": great_threshold_ladder.sh

```
#!/bin/bash
# Robustness of the gene ontology enrichment to the catalog threshold
# (Table S4.4).
#
# The catalog rule has three parameters: the modern-plus-Neandertal ceiling x,
# the Denisovan floor y and the percentile cut pcut. This driver holds
# y = 700 ka and pcut = 50 fixed and rebuilds the catalog across a ladder of x,
# then runs the identical GREAT-style test on each rebuild so that the only
# thing differing between rungs is the threshold. x = 625 ka reproduces the
# published catalog of 141 loci and the Table S4.3 numbers exactly.
#
# The permutation is the same 100,000 B-matched replicates used for
# Table S4.3, run through the same chunked driver, so the two tables are
# directly comparable and share a p value floor of 1/100,001.

set -euo pipefail
cd "$(dirname "$0")/.."

SCAN="${SCAN:-hubisz_tmrca_1kb_8afr.tight.tsv}"
LADDER="${LADDER:-catalog_annotation/threshold_ladder}"
XS="${XS:-575 600 625 650 675 700}"
CHUNKS="${CHUNKS:-5}"
NREP="${NREP:-20000}"

mkdir -p "$LADDER"

for X in $XS; do
    echo "=== x = $X ka ==="

    python3 scripts/make_catalog_2080.py \
        --scan "$SCAN" --grid-step 1000 --cluster-gap 5000 \
        --x "$X" --y 700 --cut 0.50 --exclude-x \
        -o "$LADDER/x$X" > "$LADDER/x$X.build.log" 2>&1
    grep -E '^(CASE|P\(real\))' "$LADDER/x$X.build.log"

    # hypergeometric and binomial tests; the permutation is left to the
    # chunked driver below, so no replicates are wasted here
    python3 scripts/great_test.py \
        --regions "$LADDER/x$X.regions.tsv" --matched 0 \
        -o "$LADDER/go_x$X.base.tsv" > "$LADDER/go_x$X.log" 2>&1

    REGIONS="$LADDER/x$X.regions.tsv" \
    RESULTS="$LADDER/go_x$X.base.tsv" \
    OUT="$LADDER/perm_x$X" \
    FINAL="$LADDER/go_x$X.tsv" \
    CHUNKS="$CHUNKS" NREP="$NREP" \
        bash scripts/run_great_100k.sh > "$LADDER/perm_x$X.log" 2>&1

    echo "  permutation done -> $LADDER/go_x$X.tsv"
done

echo
echo "wrote $LADDER/go_x*.tsv"
echo "Table S4.4 is the fold enrichment and matched-permutation p from these files."
```
