## Supplementary Information section 4 code base for "Hypothesis: A modern human range expansion ∼300,000 years ago had a critical role in Neandertal origins": hubisz_out_bed_tmrca_scan.py

```
#!/usr/bin/env python3
"""
Genome-wide subset-TMRCA scan from Hubisz 2020 out.bed.gz (local trees).

Each line of out.bed.gz is: chrom, start, end, MCMC_rep, newick
(see README in the dataset). NHX comments [&&NHX:...] are stripped before
parsing.

For each genomic position on a grid (default 5 kb), for each query tip-set,
TMRCA is computed from Newick branch lengths (generations), then written as
**thousands of years (kya)**: round(generations * years_per_gen / 1000) with
default years_per_gen=29 (Hubisz et al. 2020). Summary columns are integers;
_n is still the MCMC sample count.

Requires: pip install biopython

Example (one window, for testing):
  python3 hubisz_out_bed_tmrca_scan.py \\
    --dataset-root /path/to/Hubisz.Seipel2020__PlosGenetics__args.Nean.Deni.amh \\
    --windows-bed filtered_windows.bed \\
    --queries scripts/tmrca_queries.hubisz_2afr.json \\
    --only-dir chr4.71 \\
    --out chr4.71.tmrca.tsv

Full genome (all rows in filtered_windows.bed):
  python3 hubisz_out_bed_tmrca_scan.py \\
    --dataset-root ... \\
    --windows-bed filtered_windows.bed \\
    --queries scripts/tmrca_queries.hubisz_2afr.json \\
    --out hubisz_tmrca_5kb.tsv
"""

from __future__ import annotations

import argparse
import bisect
import gzip
import json
import math
import re
import sys
from collections import defaultdict
from io import StringIO
from typing import Dict, List, Optional, Sequence, Tuple

try:
    from Bio import Phylo
except ImportError:
    sys.stderr.write("Install biopython: pip install biopython\n")
    raise


def strip_nhx(newick: str) -> str:
    """Remove bracketed NHX annotations from a Newick string."""
    return re.sub(r"\[[^\]]*\]", "", newick)


def tmrca_subset_generations(newick: str, tip_names: Sequence[str]) -> Optional[float]:
    """
    Return TMRCA in generations for tip_names: height of MRCA above the tips
    (max patristic distance from MRCA to any selected tip, in branch-length units).
    """
    tips_set = set(tip_names)
    if len(tips_set) < 2:
        return None
    clean = strip_nhx(newick.strip())
    if not clean.endswith(";"):
        clean += ";"
    try:
        tree = Phylo.read(StringIO(clean), "newick")
    except Exception:
        return None

    tips = [t for t in tree.get_terminals() if t.name in tips_set]
    if len(tips) < len(tips_set):
        return None
    if len(tips) < 2:
        return None

    mrca = tree.common_ancestor(*tips)
    # Bio.Phylo does not set Clade.parent when parsing Newick; use path length API.
    try:
        depths = [tree.distance(mrca, t) for t in tips]
    except (TypeError, ValueError):
        return None
    return max(depths)


def generations_to_kya_int(gen: float, years_per_gen: float) -> int:
    """Convert generations to kya and round to nearest integer."""
    return int(round(gen * years_per_gen / 1000.0))


def percentile_nearest_rank(vals: List[float], p: float) -> float:
    """p in (0,1); linear interpolation on sorted data (simple)."""
    if not vals:
        return float("nan")
    x = sorted(vals)
    n = len(x)
    if n == 1:
        return x[0]
    idx = p * (n - 1)
    lo = int(math.floor(idx))
    hi = int(math.ceil(idx))
    if lo == hi:
        return x[lo]
    w = idx - lo
    return x[lo] * (1 - w) + x[hi] * w


def parse_positions_tsv(path: str) -> Dict[int, List[int]]:
    """chrom -> sorted exact positions, from a TSV with chrom and pos columns.

    Used instead of a regular lattice when the question is about particular
    bases rather than a scan. The ARG carries a local tree per interval, so a
    named base gets the tree that spans it: this is an exact read of the
    posterior at that base, not an interpolation between lattice points.
    """
    out: Dict[int, List[int]] = defaultdict(list)
    with open(path, encoding="utf-8") as fh:
        head = fh.readline().rstrip("\n").split("\t")
        try:
            ic, ip = head.index("chrom"), head.index("pos")
        except ValueError:
            sys.exit("%s: need chrom and pos columns" % path)
        for line in fh:
            f = line.rstrip("\n").split("\t")
            if len(f) <= max(ic, ip):
                continue
            c = parse_chrom_field(f[ic])
            if c is None:
                continue
            out[c].append(int(float(f[ip])))
    for c in out:
        out[c] = sorted(set(out[c]))
    return out


def grid_positions(region_start: int, region_end: int, step: int) -> List[int]:
    """Positions i with region_start <= i < region_end and (i - region_start) % step == 0."""
    if region_end <= region_start or step <= 0:
        return []
    out = []
    i = region_start
    while i < region_end:
        out.append(i)
        i += step
    return out


def grid_hits_interval(
    interval_start: int, interval_end: int, grid: Sequence[int]
) -> List[int]:
    """Grid points in [interval_start, interval_end) — assumes sorted grid."""
    import bisect

    lo = bisect.bisect_left(grid, interval_start)
    hi = bisect.bisect_left(grid, interval_end)
    return list(grid[lo:hi])


def load_queries(path: str) -> List[dict]:
    with open(path) as fh:
        data = json.load(fh)
    return data["queries"]


def parse_chrom_field(token: str) -> Optional[int]:
    """
    Chromosome from BED / out.bed.gz first column.
    Accepts 1-22, optional 'chr' prefix, X/Y/MT (mapped to 23/24/25 to match
    integer coding in tree files).
    """
    s = token.strip()
    if s.lower().startswith("chr"):
        s = s[3:]
    if not s:
        return None
    su = s.upper()
    if su == "X":
        return 23
    if su == "Y":
        return 24
    if su in ("MT", "M"):
        return 25
    try:
        return int(s)
    except ValueError:
        return None


def parse_windows_bed(path: str) -> List[Tuple[int, int, int, str]]:
    rows = []
    with open(path) as fh:
        for line in fh:
            line = line.strip()
            if not line or line.startswith("#"):
                continue
            p = line.split("\t")
            if len(p) < 4:
                continue
            chrom = parse_chrom_field(p[0])
            if chrom is None:
                continue
            a, b = int(p[1]), int(p[2])
            dirname = p[3].strip()
            rows.append((chrom, a, b, dirname))
    return rows


def process_out_bed_gz(
    bed_gz_path: str,
    chrom: int,
    region_start: int,
    region_end: int,
    grid_step: int,
    queries: List[dict],
    burnin_rep: int,
    max_rep: Optional[int],
    positions: Optional[List[int]] = None,
) -> Dict[Tuple[int, str], List[float]]:
    """
    Accumulate TMRCA samples per (position, query_id) across MCMC rows.
    """
    if positions is None:
        grid = grid_positions(region_start, region_end, grid_step)
    else:
        lo = bisect.bisect_left(positions, region_start)
        hi = bisect.bisect_left(positions, region_end)
        grid = list(positions[lo:hi])
    accum: Dict[Tuple[int, str], List[float]] = defaultdict(list)

    try:
        fh = gzip.open(bed_gz_path, "rt", encoding="utf-8", errors="replace")
    except OSError as e:
        sys.stderr.write("Could not open %s: %s\n" % (bed_gz_path, e))
        return accum

    with fh:
        for line in fh:
            line = line.rstrip("\n")
            if not line:
                continue
            parts = line.split("\t", 4)
            if len(parts) < 5:
                continue
            try:
                c = parse_chrom_field(parts[0])
                s, e = int(parts[1]), int(parts[2])
                rep = int(parts[3])
                nw = parts[4]
            except ValueError:
                continue
            if c is None or c != chrom:
                continue
            if rep < burnin_rep:
                continue
            if max_rep is not None and rep > max_rep:
                continue

            hits = grid_hits_interval(s, e, grid)
            if not hits:
                continue

            vals_by_q = {}
            for q in queries:
                qid = q["id"]
                v = tmrca_subset_generations(nw, q["tips"])
                if v is not None and not (math.isnan(v) or math.isinf(v)):
                    vals_by_q[qid] = v

            for pos in hits:
                for qid, v in vals_by_q.items():
                    accum[(pos, qid)].append(v)

    return accum


def write_chunk_stats(
    accum: Dict[Tuple[int, str], List[float]],
    chrom: int,
    query_order: List[str],
    out_fh,
    years_per_gen: float,
) -> None:
    """Write one TSV line per position; TMRCA summaries as integer kya."""
    by_pos: Dict[int, Dict[str, List[float]]] = defaultdict(lambda: defaultdict(list))
    for (pos, qid), vals in accum.items():
        by_pos[pos][qid].extend(vals)

    for pos in sorted(by_pos.keys()):
        row = [str(chrom), str(pos)]
        for qid in query_order:
            vals = by_pos[pos].get(qid, [])
            n = len(vals)
            if n == 0:
                row.extend(["NA", "NA", "NA", "0"])
            else:
                m = sum(vals) / n
                lo = percentile_nearest_rank(vals, 0.025)
                hi = percentile_nearest_rank(vals, 0.975)
                row.extend(
                    [
                        str(generations_to_kya_int(m, years_per_gen)),
                        str(generations_to_kya_int(lo, years_per_gen)),
                        str(generations_to_kya_int(hi, years_per_gen)),
                        str(n),
                    ]
                )
        out_fh.write("\t".join(row) + "\n")


def main() -> None:
    ap = argparse.ArgumentParser(description=__doc__)
    ap.add_argument(
        "--dataset-root",
        required=True,
        help="Path to Hubisz.Seipel2020__PlosGenetics__args.Nean.Deni.amh",
    )
    ap.add_argument(
        "--windows-bed",
        required=True,
        help="filtered_windows.bed (chrom start end dirname)",
    )
    ap.add_argument(
        "--queries",
        required=True,
        help="JSON file with { 'queries': [ {id, tips}, ... ] }",
    )
    ap.add_argument("--grid-step", type=int, default=5000, help="BP between output sites")
    ap.add_argument(
        "--positions-tsv",
        default=None,
        help="TSV with chrom and pos columns; report at exactly these bases "
             "instead of a --grid-step lattice",
    )
    ap.add_argument(
        "--burnin-rep",
        type=int,
        default=500,
        help="Ignore MCMC rep index < this (see dataset README)",
    )
    ap.add_argument(
        "--max-rep",
        type=int,
        default=None,
        help="Optional upper bound on MCMC rep index (inclusive)",
    )
    ap.add_argument(
        "--only-dir",
        default=None,
        help="Process only this subdirectory name (e.g. chr4.71)",
    )
    ap.add_argument(
        "--years-per-gen",
        type=float,
        default=29.0,
        help="Years per generation for kya output (default 29, Hubisz et al. 2020)",
    )
    ap.add_argument(
        "-o",
        "--out",
        required=True,
        help="Output TSV (genome-wide); sorted by chrom, pos within each run",
    )
    args = ap.parse_args()

    queries = load_queries(args.queries)
    positions = (parse_positions_tsv(args.positions_tsv)
                 if args.positions_tsv else None)
    if positions is not None:
        sys.stderr.write("exact positions: %d on %d chromosomes\n"
                         % (sum(len(v) for v in positions.values()),
                            len(positions)))
    query_order = [q["id"] for q in queries]
    windows = parse_windows_bed(args.windows_bed)

    header = ["chrom", "pos"]
    for qid in query_order:
        header.extend(
            [
                f"{qid}_mean_kya",
                f"{qid}_p025_kya",
                f"{qid}_p975_kya",
                f"{qid}_n",
            ]
        )

    root = args.dataset_root.rstrip("/")
    sys.stderr.write(
        "TMRCA columns: integer kya = round(generations * years_per_gen / 1000); years_per_gen=%s\n"
        % args.years_per_gen
    )
    n_done = 0
    with open(args.out, "w") as out_fh:
        out_fh.write("\t".join(header) + "\n")
        for chrom, r0, r1, dirname in windows:
            if args.only_dir is not None and dirname != args.only_dir:
                continue
            if positions is not None:
                have = positions.get(chrom, [])
                if bisect.bisect_left(have, r0) >= bisect.bisect_left(have, r1):
                    continue
            bed_path = "%s/%s/out.bed.gz" % (root, dirname)
            accum = process_out_bed_gz(
                bed_path,
                chrom,
                r0,
                r1,
                args.grid_step,
                queries,
                args.burnin_rep,
                args.max_rep,
                None if positions is None else positions.get(chrom, []),
            )
            write_chunk_stats(accum, chrom, query_order, out_fh, args.years_per_gen)
            n_done += 1
            sys.stderr.write("Wrote chunk %s (%d positions with data)\n" % (dirname, len({p for (p, _) in accum})))

    sys.stderr.write("Done. Chunks processed: %d. Output: %s\n" % (n_done, args.out))


if __name__ == "__main__":
    main()
```
