## Supplementary Information section 4 code base for "Hypothesis: A modern human range expansion ∼300,000 years ago had a critical role in Neandertal origins": hubisz_tmrca_collate_parts.sh

```
#!/usr/bin/env bash
# Merge per-chunk TSVs from parallel Slurm runs into one genome file, sorted by chrom, pos.
#
# Usage:
#   bash hubisz_tmrca_collate_parts.sh PARTS_DIR FINAL_TSV
#
# Example:
#   bash scripts/hubisz_tmrca_collate_parts.sh "$OUTDIR/parts" "$OUTDIR/hubisz_tmrca_5kb.tsv"
#
# Optional:
#   EXPECT_CHUNKS=1217   # if set, warn when the number of part files differs

set -euo pipefail

PARTS_DIR="${1:?Usage: $0 PARTS_DIR FINAL_TSV}"
FINAL_TSV="${2:?Usage: $0 PARTS_DIR FINAL_TSV}"

shopt -s nullglob
parts=( "$PARTS_DIR"/*.tsv )
n="${#parts[@]}"

if [[ "$n" -eq 0 ]]; then
  echo "ERROR: no *.tsv files under $PARTS_DIR"
  exit 1
fi

if [[ -n "${EXPECT_CHUNKS:-}" ]] && [[ "$n" -ne "$EXPECT_CHUNKS" ]]; then
  echo "WARNING: found $n part files, expected EXPECT_CHUNKS=$EXPECT_CHUNKS (missing or extra chunks?)" >&2
fi

header="$(head -1 "${parts[0]}")"
for f in "${parts[@]}"; do
  h="$(head -1 "$f")"
  if [[ "$h" != "$header" ]]; then
    echo "ERROR: header mismatch in $f" >&2
    exit 1
  fi
done

TMPDIR="${TMPDIR:-/tmp}"
TMP_SORT="$(mktemp "$TMPDIR/hubisz_tmrca_sort.XXXXXX")"
trap 'rm -f "$TMP_SORT"' EXIT

for f in "${parts[@]}"; do
  tail -n +2 "$f"
done | sort -k1,1n -k2,2n >"$TMP_SORT"

{
  echo "$header"
  cat "$TMP_SORT"
} >"${FINAL_TSV}.tmp"

mv "${FINAL_TSV}.tmp" "$FINAL_TSV"
echo "Wrote $FINAL_TSV ($n chunks merged; body sorted by chrom, pos)"
```
