## Supplementary Information section 4 code base for "Hypothesis: A modern human range expansion ∼300,000 years ago had a critical role in Neandertal origins": make_catalog_2080.py

```
#!/usr/bin/env python3
"""
Build the sweep catalog under the selected discriminator: the symmetric
reconstructed 20/80 rule at a single crossing threshold x (default 675 ka).

CASE (catalog) rule, applied independently to every 5-kb grid point:
    p80[ TMRCA(Africans + both Neandertals) ]  <=  x
    p20[ TMRCA(Africans + Denisova)         ]  >   x

CONTROL rule (identical, archaics swapped) is used only to estimate the FDR:
    p80[ TMRCA(Africans + Denisova)         ]  <=  x
    p20[ TMRCA(Africans + both Neandertals) ]  >   x

The 20th and 80th percentiles are reconstructed by approx_percentiles.py from
the stored 2.5 / 97.5 percentiles, assuming the posterior is Normal in
ARGweaver's own log-spaced time coordinate.

Usage:
  python3 scripts/make_catalog_2080.py --x 675 \
      -o catalog_annotation/catalog_2080_x675
"""

from __future__ import annotations

import argparse
import csv
import os
from collections import defaultdict

from approx_percentiles import default_coord, fit_site, load, z_of
from optimize_rule_by_size import NULL_RATIO


def chrom_key(c):
    return (int(c), "") if c.isdigit() else (99, c)


def cluster(sites, gap=5000, step=5000):
    """Merge grid points no more than `gap` apart into contiguous regions.

    `gap` and `step` are different things and were both 5000 while the scan was
    on a 5 kb lattice, which hid the distinction. `step` is the width a grid
    point stands for, so it is what a region's end has to be extended by to
    cover the last point's territory. `gap` is the choice of how far apart two
    points can be and still be called one locus. On a 1 kb lattice `step` must
    become 1000, but `gap` should stay at 5000 if a locus is to keep meaning
    the same physical thing: dropping it to 1000 would split one sweep into
    several wherever the signal dips below threshold for a couple of kilobases.
    """
    by = defaultdict(list)
    for c, p in sites:
        by[c].append(p)
    regions = []
    for c in sorted(by, key=chrom_key):
        ps = sorted(by[c])
        start = prev = ps[0]
        n = 1
        for p in ps[1:]:
            if p - prev <= gap:
                prev = p
                n += 1
            else:
                regions.append((c, start, prev + step, n))
                start = prev = p
                n = 1
        regions.append((c, start, prev + step, n))
    return regions


def main() -> int:
    ap = argparse.ArgumentParser()
    ap.add_argument("--scan", default="hubisz_tmrca_5kb_8afr.tsv")
    ap.add_argument("--grid-step", type=int, default=5000,
                    help="bp a grid point stands for; 1000 for the 1 kb scan")
    ap.add_argument("--cluster-gap", type=int, default=5000,
                    help="merge grid points this far apart into one locus; "
                         "kept at 5000 independent of --grid-step so a locus "
                         "means the same physical thing on either lattice")
    ap.add_argument("--x", type=float, default=675.0)
    ap.add_argument("--cut", type=float, default=0.80,
                    help="upper percentile of the rule; its mirror (1-cut) is "
                         "the lower one. 0.80 is the 80/20 rule, 0.50 the "
                         "median rule.")
    ap.add_argument("--y", type=float, default=None,
                    help="Denisovan threshold, in ka, applied to the lower "
                         "percentile of TMRCA(Afr+Den). Defaults to x, i.e. "
                         "tied to the Neandertal threshold. Setting y above x "
                         "additionally forces the Denisovan lineage to be an "
                         "outgroup to the African-plus-Neandertal clade, since "
                         "that clade must close below x while the Denisovan "
                         "join must lie above y.")
    ap.add_argument("--no-den", action="store_true",
                    help="drop the Denisovan half of the rule. A free grid "
                         "search placed that threshold where it never binds at "
                         "every catalog size up to 1,000 windows, so it "
                         "contributes nothing to purity over the useful range. "
                         "It may still be wanted for interpretability.")
    ap.add_argument("--exclude-x", action="store_true",
                    help="drop X-linked grid points (their FDR is near 1, i.e. noise)")
    ap.add_argument("-o", "--out-prefix", default="catalog_annotation/catalog_2080_x675")
    args = ap.parse_args()

    recs = load(args.scan)
    # Recover the ARGweaver time grid from the whole scan before any filtering,
    # so the percentile reconstruction is identical across chromosome subsets.
    coord, grid_desc = default_coord(recs)
    print(f"time coordinate: {grid_desc}")
    if args.exclude_x:
        n0 = len(recs)
        recs = [d for d in recs
                if d["chrom"].replace("chr", "") not in ("X", "23")]
        print(f"excluded {n0 - len(recs):,} X-linked grid points")
    # p20/p80 are the rule's lower and upper percentiles; with --cut they need
    # not be the 20th and 80th, and the output headers carry the real values.
    z20, z80 = z_of(1.0 - args.cut), z_of(args.cut)
    for d in recs:
        for tag in ("nea", "den"):
            mu, sg = fit_site(coord, d[f"{tag}_p025"], d[f"{tag}_p975"])
            d[f"{tag}_p20"] = coord.to_t(mu + z20 * sg)
            d[f"{tag}_p80"] = coord.to_t(mu + z80 * sg)

    x = args.x
    y = x if args.y is None else args.y
    if args.no_den:
        case = [d for d in recs if d["nea_p80"] <= x]
        ctrl = [d for d in recs if d["den_p80"] <= x]
    else:
        case = [d for d in recs if d["nea_p80"] <= x and d["den_p20"] > y]
        ctrl = [d for d in recs if d["den_p80"] <= x and d["nea_p20"] > y]

    cs = [(d["chrom"].replace("chr", ""), d["pos"]) for d in case]
    cs = [("X" if c == "23" else c, p) for c, p in cs]
    ks = [(d["chrom"].replace("chr", ""), d["pos"]) for d in ctrl]
    ks = [("X" if c == "23" else c, p) for c, p in ks]

    creg = cluster(cs, args.cluster_gap, args.grid_step)
    kreg = cluster(ks, args.cluster_gap, args.grid_step)
    # Null skew, imported rather than restated so the two cannot drift apart.
    #
    # Two things it is NOT. It is not the limit of the curve as x grows without
    # bound: that limit is 1.0, because a rule loose enough to admit every
    # window makes case and control the same set (the observed ratio climbs
    # 0.871 at 800 ka, 0.964 at 1000 ka, 0.990 at 1500 ka). And it is not the
    # odds inside the fitted support, which was the previous basis for 1.150 --
    # that region contains the catalog itself, so it measures signal, not a
    # null. It is now measured in shells above the published modern-archaic
    # split, where a shallow join-side coalescence carries no sweep evidence.
    null_ratio = NULL_RATIO
    print(f"usable grid points : {len(recs):,}")
    print(f"threshold x        : {x:.0f} ka")
    print(f"percentile cut     : {args.cut:.2f}/{1-args.cut:.2f}"
          + ("   Denisovan condition: OFF" if args.no_den else ""))
    # Territory covered, which is the merged extent of the clusters rather than
    # points x step: with a cluster gap wider than the step, a locus can span
    # bases that no grid point sits on, and those bases are part of the locus.
    print(f"CASE    sites {len(case):>5}   clusters {len(creg):>4}   "
          f"{sum(e - s for _, s, e, _ in creg)/1e6:.2f} Mb")
    print(f"CONTROL sites {len(ctrl):>5}   clusters {len(kreg):>4}")
    ratio = len(ctrl) / len(case)
    print(f"Den/Nea ratio (sites)   = {ratio:.3f}")
    print(f"Den/Nea ratio (clusters)= {len(kreg)/len(creg):.3f}")
    print(f"P(real) = 1 - ratio/{null_ratio:.4f} = "
          f"{max(0.0, 1 - ratio/null_ratio):.3f}")

    bych = defaultdict(int)
    for c, _ in cs:
        bych[c] += 1
    print("case sites by chromosome: " +
          ", ".join(f"{c}:{bych[c]}" for c in sorted(bych, key=chrom_key)))

    os.makedirs(os.path.dirname(args.out_prefix) or ".", exist_ok=True)

    with open(f"{args.out_prefix}.sites.tsv", "w", newline="", encoding="utf-8") as f:
        w = csv.writer(f, delimiter="\t")
        w.writerow(["chrom", "pos", "nea_mean_kya", "nea_p025_kya", "nea_p20_kya",
                    "nea_p80_kya", "nea_p975_kya", "den_mean_kya", "den_p025_kya",
                    "den_p20_kya", "den_p80_kya", "den_p975_kya", "afr_mean_kya"])
        for d in sorted(case, key=lambda r: (chrom_key(r["chrom"].replace("chr", "")), r["pos"])):
            w.writerow([d["chrom"], d["pos"],
                        round(d["nea_mean"]), round(d["nea_p025"]), round(d["nea_p20"]),
                        round(d["nea_p80"]), round(d["nea_p975"]),
                        round(d["den_mean"]), round(d["den_p025"]), round(d["den_p20"]),
                        round(d["den_p80"]), round(d["den_p975"]), round(d["afr_mean"])])

    with open(f"{args.out_prefix}.regions.tsv", "w", newline="", encoding="utf-8") as f:
        w = csv.writer(f, delimiter="\t")
        w.writerow(["chrom", "start", "end", "n_sites", "length_kb"])
        for c, s, e, n in creg:
            w.writerow([c, s, e, n, (e - s) // 1000])

    with open(f"{args.out_prefix}.control_regions.tsv", "w", newline="", encoding="utf-8") as f:
        w = csv.writer(f, delimiter="\t")
        w.writerow(["chrom", "start", "end", "n_sites", "length_kb"])
        for c, s, e, n in kreg:
            w.writerow([c, s, e, n, (e - s) // 1000])

    print(f"\nwrote {args.out_prefix}.sites.tsv, .regions.tsv, .control_regions.tsv")
    return 0


if __name__ == "__main__":
    raise SystemExit(main())
```
