## Supplementary Information section 4 code base for "Hypothesis: A modern human range expansion ∼300,000 years ago had a critical role in Neandertal origins": make_functional_table.py

```
#!/usr/bin/env python3
"""
The main-text table of fixed-derived sites worth following up.

WHAT THE ORDERING COLUMN IS, AND IS NOT
Preal is a property of a set: it is one minus the control-to-case ratio over a
collection of grid points, and a single point has no such ratio. There is
nonetheless an honest per-locus quantity of the same kind. The thresholds are
nested, so every locus has a tightest rule that still admits it, and that rule's
catalog has a measured purity. A locus in the catalog proper sits at 0.800; one
that only appears when the Neandertal threshold is relaxed to 650 ka sits at
0.701, and to 675 ka at 0.663. The column therefore reads "the purity of the
smallest catalog containing this locus", which is ordinal, measured, and not a
claim about the individual site. Sorting on it descending puts the best-supported
loci first.

WHY THE TABLE IS NOT THE QUALIFYING SET
Scored against random positions drawn from the same regions, the full qualifying
definition sits at 28.8% background against 32.8% observed (p = 0.24), and no
narrower version reaches significance either. The annotation does not show these
sites to be functional, and a table of every qualifying site would be a table of
mostly background. What it can do is say which sites have something specific
about them worth reading, and that is what is tabulated: a protein-altering
change, an essential splice position, a CADD score in the top 1% of possible
substitutions, or conservation coinciding with a transcribed regulatory feature.
Permissive annotation alone -- an ENCODE cluster and nothing else, which covers
about a third of these regions -- does not earn a row.

Example:
  python3 make_functional_table.py \
      -i catalog_annotation/fixed_derived_functional.tsv \
      -r catalog_annotation/screen_preal30.regions.tsv \
      -o catalog_annotation/table_functional_sites.tsv
"""

from __future__ import annotations

import argparse
import csv
import os

ALTERING = {"protein_altering"}
STRONG_SINGLE = {"protein_altering", "splice_essential", "cadd"}
# Conservation on its own is weak; conservation plus a transcribed or
# regulatory feature at the same base is the combination worth reading.
PAIR_A = {"phastcons", "phylop"}
PAIR_B = {"utr5", "utr3", "promoter", "splice_region"}
# A synonymous change is not evidence of function and cannot stand in for a
# regulatory feature. CADD is scaled across all substitution types and being
# inside a coding exon lifts it on its own -- median 11.0 at the synonymous
# fixed-derived sites against 1.9 at the non-coding ones -- so a synonymous
# site in the CADD top 1% is mostly telling us about the exon around it. The
# base-level evidence does not agree: every synonymous site in the catalog has
# phyloP within noise of zero, where two of the five missense sites exceed 1. A
# synonymous change acting through splicing or codon usage should have left
# conservation behind, so these are excluded even when CADD is high.
NEVER_ALONE = {"synonymous"}


def qualifies(classes):
    """The Table S4.2 rule. Single source of truth; build_supp_table2 imports it.

    Either one strong line of evidence on its own -- a protein-altering change,
    an essential splice position, or CADD in the top 1% -- or conservation
    coinciding with a transcribed or regulatory feature at the same base. A
    synonymous change never qualifies on its own account.
    """
    cls = set(classes.split(";")) - {""} if isinstance(classes, str) else set(classes)
    if cls & NEVER_ALONE and not (cls & {"protein_altering",
                                         "splice_essential"}):
        return False
    strong = bool(cls & STRONG_SINGLE) or "coding_unresolved" in cls
    return strong or (bool(cls & PAIR_A) and bool(cls & PAIR_B))

PRETTY = {"protein_altering": "protein-altering", "synonymous": "synonymous",
          "splice_essential": "essential splice site",
          "splice_region": "splice region", "utr5": "5' UTR",
          "utr3": "3' UTR", "promoter": "promoter",
          "phastcons": "conserved element", "phylop": "phyloP",
          "cadd": "CADD top 1%", "encode_dnase": "DNase cluster",
          "encode_tfbs": "TFBS cluster",
          "coding_unresolved": "coding, consequence not resolved"}


def load_tiers(path):
    """chrom -> list of (start, end, min_x, set_preal, in_catalog)."""
    out = {}
    if not path or not os.path.exists(path):
        return out
    with open(path, newline="") as f:
        for r in csv.DictReader(f, delimiter="\t"):
            out.setdefault(r["chrom"].replace("chr", ""), []).append(
                (int(r["start"]), int(r["end"]), float(r["min_x_ka"]),
                 float(r["set_preal_at_min_x"]), r["in_catalog"]))
    for c in out:
        out[c].sort()
    return out


def tier_of(tiers, c, p):
    for s, e, x, pr, inc in tiers.get(c, []):
        if s <= p <= e:
            return x, pr, inc
    return None


def main():
    ap = argparse.ArgumentParser()
    ap.add_argument("-i", "--infile",
                    default="catalog_annotation/"
                            "fd_af99.functional.canonical.tsv")
    ap.add_argument("-r", "--regions",
                    default="catalog_annotation/screen_preal30.padded.regions.tsv")
    ap.add_argument("-o", "--outfile",
                    default="catalog_annotation/table_functional_sites.tsv")
    ap.add_argument("--cadd-min", type=float, default=20.0)
    a = ap.parse_args()

    tiers = load_tiers(a.regions)
    with open(a.infile, newline="") as f:
        rows = list(csv.DictReader(f, delimiter="\t"))

    keep = []
    for r in rows:
        cls = set(r["classes"].split(";")) - {""}
        # "coding_unresolved" means the screen recorded a CDS hit without a
        # consequence, which happens when the input predates the protein
        # annotation. Those are kept rather than dropped: silently losing a
        # nonsynonymous change is the one failure this table cannot afford.
        if not qualifies(cls):
            continue
        c, p = r["chrom"].replace("chr", ""), int(r["pos"])
        t = tier_of(tiers, c, p)
        keep.append({
            "chr": c, "pos": p,
            "locus": f"{c}:{p/1e6:.3f}",
            "gene": r.get("gene") or "-",
            "classes": ", ".join(PRETTY.get(k, k) for k in sorted(cls)),
            "cadd_max": r.get("cadd_max") or "",
            "phylop": r.get("phylop") or "",
            "tier_x_ka": f"{t[0]:.0f}" if t else "",
            "set_preal": f"{t[1]:.3f}" if t else "",
            "in_catalog": t[2] if t else "outside current region set",
        })

    def key(row):
        pr = float(row["set_preal"]) if row["set_preal"] else -1.0
        cd = float(row["cadd_max"]) if row["cadd_max"] else -1.0
        return (-pr, -cd)

    keep.sort(key=key)
    fields = ["locus", "gene", "classes", "cadd_max", "phylop", "tier_x_ka",
              "set_preal", "in_catalog"]
    with open(a.outfile, "w", newline="") as f:
        w = csv.DictWriter(f, fieldnames=fields, delimiter="\t",
                           extrasaction="ignore")
        w.writeheader()
        w.writerows(keep)

    wid = [max(len(str(r[k])) for r in keep + [dict(zip(fields, fields))])
           for k in fields]
    print(f"{len(keep)} sites of {len(rows)} screened\n")
    print("  ".join(h.ljust(w) for h, w in zip(fields, wid)))
    print("  ".join("-" * w for w in wid))
    for r in keep:
        print("  ".join(str(r[k]).ljust(w) for k, w in zip(fields, wid)))
    print(f"\nwrote {a.outfile}")
    return 0


if __name__ == "__main__":
    raise SystemExit(main())
```
