## Supplementary Information section 4 code base for "Hypothesis: A modern human range expansion ∼300,000 years ago had a critical role in Neandertal origins": make_loci_table.py

```
#!/usr/bin/env python3
"""Build the full 132-locus catalog table for Table S4.1.

Rows are ordered by chromosome and position, and the added column is the best
P(real) attained anywhere in the locus.

WHY THE FITTED SURFACE AND NOT A LADDER OF NESTED CATALOGS
The tempting definition of a per-locus purity is the purity of the tightest
catalog that still contains the locus, since P(real) is a property of a set
rather than of a site. That definition fails in practice because the ladder is
not monotone: at y = 700 ka and pcut = 50 the rungs run 0.558 at x <= 525 ka,
0.727 at 550, 0.798 at 575, 0.767 at 600 and 0.800 at 625. The tightest rung
rests on 32 cases against 14 controls, so its purity is mostly noise, and a
column built this way would rank the shallowest loci worst -- the exact
opposite of what it is meant to convey.

The cross-validated FDR surface gives a smooth alternative. Each grid point is
scored by the model fitted on the chromosomes of the opposite parity, so no
grid point contributes to the model that ranks it, and P(real) follows from
theta as one minus the implied control-to-case odds over the signal-free odds.

CLAMPING TO THE FITTED SUPPORT
The surface is fitted over u in (350, 900] and v in (350, 1500], and 116 of the
329 catalog grid points fall outside that box. Discarding them is not an
option: 108 do so because the Denisovan median is DEEPER than 1500 ka, which
makes them the strongest evidence in the catalog, and the remaining 8 because
the Neandertal median is shallower than 350 ka, likewise a mark of strength.
Both are therefore scored at the boundary of the support rather than dropped.
This is not extrapolation -- the surface is evaluated inside its own support --
and it understates rather than flatters those loci, because a Denisovan
coalescence deeper than the boundary, or a Neandertal one shallower than it,
can only be stronger evidence for a sweep than the boundary itself.

Usage:
  python3 scripts/make_loci_table.py -o catalog_annotation/table_s51_loci.tsv
"""

from __future__ import annotations

import argparse
import csv
import math
import os
import sys
from collections import Counter

import numpy as np

sys.path.insert(0, os.path.dirname(os.path.abspath(__file__)))
from approx_percentiles import default_coord, fit_site, load, z_of  # noqa: E402
from fig_s51_model_curve import U_EDGES, V_EDGES, fit_on  # noqa: E402
from fit_fdr_surface import PCUTS, design  # noqa: E402
from optimize_rule_by_size import NULL_RATIO, THETA0  # noqa: E402

Y_FLOOR = 700.0
RULE_CUT = 0.50
NULL_ODDS = THETA0 / (1 - THETA0)
EPS = 1e-6


def preal_from_theta(th):
    return max(0.0, 1.0 - ((1 - th) / th) / NULL_ODDS)


def fmt_t(mean, lo, hi):
    """Point estimate with its 95% central interval, e.g. '706 (600-1300)'.

    Both parts are averaged over the same catalog grid points, so the interval
    is the typical posterior spread across the locus rather than the spread of
    the locus mean. A wide interval therefore flags a locus where ARGweaver-D
    localized the coalescence poorly at every grid point, which is the case the
    reader needs to discount.
    """
    return f"{mean:.0f} ({lo:.0f}\u2013{hi:.0f})"


def fmt_central90(median, lo, hi):
    """Median with its 90% central interval, e.g. '706 (450-1300)'.

    The bounds of a 90% central interval are the 5th and the 95th percentiles,
    which are exactly the two numbers the catalog argument turns on: the 95th
    bounds how old the modern and modern-Neandertal coalescences can be, and
    the 5th bounds how young the Denisovan one can be. Quoting the 90% interval
    therefore carries both relevant bounds at once, and each is the same number
    a one-sided 95% statement would give, rather than the 97.5% tails a 95%
    central interval spends its width on.

    Unlike the central style this reports the median rather than the posterior
    mean, so all three printed numbers are quantiles of the same reconstructed
    posterior. TMRCA posteriors are right skewed, so the mean sits above the
    median and mixing the two invites the reader to misread the interval as
    symmetric about the point estimate.
    """
    return f"{median:.0f} ({lo:.0f}\u2013{hi:.0f})"


# Nodes for the two-dimensional quadrature behind diff_quantiles. A product of
# two 41-point standard-normal grids over +/- 4 sigma gives 1,681 atoms, which
# resolves a 5th and a 95th percentile far more finely than the one-kiloyear
# precision the column is printed to, and unlike Monte Carlo it returns the
# same answer every run.
_DZ = [-4.0 + 8.0 * i / 40.0 for i in range(41)]
_DW = [math.exp(-0.5 * z * z) for z in _DZ]
_DWSUM = sum(_DW)
_DW = [w / _DWSUM for w in _DW]


def diff_quantiles(coord, d, ps=(0.05, 0.5, 0.95)):
    """Percentiles of TAfr+Den - TAfr+Nea at one grid point.

    The scan stores only a marginal summary per haplotype set, so the joint
    posterior of the two coalescences is not available and the difference has
    to be built from the two marginals. Treating them as independent is the
    conservative choice rather than a neutral one: the Denisovan and Neandertal
    sets both contain the same 16 African haplotypes, so their TMRCAs are
    positively correlated, and positive correlation subtracts from the variance
    of a difference. The interval this returns is therefore at least as wide as
    the true one, which is the right direction for a column whose purpose is to
    show that the separation is real.

    Both marginals are normal in grid-index coordinates, not in time, so the
    difference is taken after mapping each draw back to time and is itself
    skewed; that is why it is read off a quadrature rather than from a
    closed-form sum of variances.
    """
    tn = [coord.to_t(d["nea_mu"] + z * d["nea_sg"]) for z in _DZ]
    td = [coord.to_t(d["den_mu"] + z * d["den_sg"]) for z in _DZ]
    atoms = sorted((td[i] - tn[j], _DW[i] * _DW[j])
                   for i in range(len(_DZ)) for j in range(len(_DZ)))
    out, acc, k = [], 0.0, 0
    for p in ps:
        while k < len(atoms) and acc + atoms[k][1] < p:
            acc += atoms[k][1]
            k += 1
        out.append(atoms[min(k, len(atoms) - 1)][0])
    return out


def fmt_one_sided(median, bound, direction):
    """Median with the one bound that carries the argument, e.g. '300 (<550)'.

    A central interval spends half its width on the side the reader does not
    need. The catalog claims that moderns and Neandertals coalesce recently and
    that Denisova does not, so for the two modern-Neandertal columns only the
    upper bound can embarrass the claim, and for the Denisovan column only the
    lower bound can. Printing the informative tail at 95% instead of both tails
    at 97.5% also tightens the number the reader actually weighs: the bound
    moves inward because it is now a one-sided statement.

    direction "<" prints an upper bound, ">" prints a lower bound.
    """
    return f"{median:.0f} ({direction}{bound:.0f})"


def gene_label(field, nearest):
    """Compact gene list. Every entry in the genes field is already inside the
    5 kb window, so the distance is used only to format the fallback."""
    coding, noncoding = [], []
    for g in (field or "").split(";"):
        if not g:
            continue
        parts = g.split("|")
        name = parts[0]
        biotype = parts[1] if len(parts) > 1 else ""
        (coding if biotype == "protein_coding" else noncoding).append(name)
    if coding:
        out = ", ".join(dict.fromkeys(coding))
        n_nc = len(set(noncoding))
        return out + (f" (+{n_nc} non-coding)" if n_nc else "")
    if noncoding:
        names = list(dict.fromkeys(noncoding))
        if len(names) > 2:
            return ", ".join(names[:2]) + f" & {len(names) - 2} other non-coding"
        return ", ".join(names) + " (non-coding)"
    if nearest:
        parts = nearest.split("|")
        name = parts[0]
        try:
            kb = int(parts[2]) / 1000.0
            return f"none within 5 kb; {name} at {kb:.0f} kb"
        except (IndexError, ValueError):
            return f"none within 5 kb; {name} nearest"
    return "none within 5 kb"


def main():
    ap = argparse.ArgumentParser()
    ap.add_argument("--scan", default="hubisz_tmrca_5kb_8afr.tsv")
    ap.add_argument("--regions",
                    default="catalog_annotation/"
                            "catalog_final_median_x625_y700_auto_annot.regions.tsv")
    ap.add_argument("--sites",
                    default="catalog_annotation/"
                            "catalog_final_median_x625_y700_auto.sites.tsv")
    ap.add_argument("-o", "--out", required=True)
    ap.add_argument("--interval-style",
                    choices=("central", "one-sided", "central90"),
                    default="central",
                    help="central prints the posterior mean with its 95%% "
                         "central interval, '706 (600-1300)', and is what the "
                         "published tables used. one-sided prints the median "
                         "with only the bound that bears on the claim: an "
                         "upper 95th percentile for the two modern-Neandertal "
                         "columns, '300 (<550)', and a lower 5th percentile "
                         "for the Denisovan column, '300 (>220)'. central90 "
                         "prints the median with its 90%% central interval, "
                         "'706 (450-1300)', whose bounds are the 5th and 95th "
                         "percentiles, so one interval carries both of the "
                         "bounds the argument rests on.")
    a = ap.parse_args()

    recs = load(a.scan)
    coord, desc = default_coord(recs)
    recs = [d for d in recs
            if d["chrom"].replace("chr", "") not in ("X", "Y", "23", "24")]
    # "afr" is fitted alongside the other two because the one-sided columns need
    # a median and a 95th percentile for it as well. The catalog rule itself
    # only ever reads nea and den, so before this the African posterior was
    # never reconstructed and only its recorded mean was available.
    for d in recs:
        for tag in ("afr", "nea", "den"):
            mu, sg = fit_site(coord, d[f"{tag}_p025"], d[f"{tag}_p975"])
            # Kept so the Denisovan-minus-Neandertal column can be given as a
            # distribution rather than a single subtraction; see diff_quantiles.
            d[f"{tag}_mu"], d[f"{tag}_sg"] = mu, sg
            for p in PCUTS:
                d[f"{tag}_hi{p}"] = coord.to_t(mu + z_of(p) * sg)
                d[f"{tag}_lo{p}"] = coord.to_t(mu + z_of(1 - p) * sg)
    print(f"time coordinate: {desc}; {len(recs):,} autosomal grid points")

    odd = [d for d in recs if int(d["chrom"].replace("chr", "")) % 2 == 1]
    even = [d for d in recs if int(d["chrom"].replace("chr", "")) % 2 == 0]
    beta_for_even, n1 = fit_on(odd)
    beta_for_odd, n2 = fit_on(even)
    print(f"surface fitted on {n1:,} odd and {n2:,} even cells; "
          f"every locus scored by the model it did not train")

    u_lo, u_hi = U_EDGES[0] + EPS, U_EDGES[-1]
    v_lo, v_hi = V_EDGES[0] + EPS, V_EDGES[-1]
    pr_of, clamped = {}, {}
    for d in recs:
        c = d["chrom"].replace("chr", "")
        u, v = d[f"nea_hi{RULE_CUT}"], d[f"den_lo{RULE_CUT}"]
        cu, cv = min(max(u, u_lo), u_hi), min(max(v, v_lo), v_hi)
        if cu != u or cv != v:
            clamped[(int(c), int(d["pos"]))] = ("u" if cu != u else "") + \
                                               ("v" if cv != v else "")
        beta = beta_for_odd if int(c) % 2 == 1 else beta_for_even
        eta = design(np.array([cu / 100.0]), np.array([cv / 100.0]),
                     np.array([RULE_CUT]))[0] @ beta
        pr_of[(int(c), int(d["pos"]))] = preal_from_theta(
            1.0 / (1.0 + np.exp(-eta)))

    with open(a.sites, encoding="utf-8") as fh:
        sites = list(csv.DictReader(fh, delimiter="\t"))
    with open(a.regions, encoding="utf-8") as fh:
        regions = list(csv.DictReader(fh, delimiter="\t"))
    ncl = sum(1 for r in sites
              if (int(r["chrom"]), int(r["pos"])) in pr_of)
    print(f"{len(sites)} catalog grid points in {len(regions)} loci; "
          f"{ncl} scored")

    by_chr = {}
    for r in sites:
        by_chr.setdefault(int(r["chrom"]), []).append(int(r["pos"]))
    post = {(int(d["chrom"].replace("chr", "")), d["pos"]): d for d in recs}

    out, widths = [], []
    for r in regions:
        c, s, e = int(r["chrom"]), int(r["start"]), int(r["end"])
        pts = [p for p in by_chr.get(c, []) if s <= p <= e]
        vals = [pr_of[(c, p)] for p in pts if (c, p) in pr_of]
        if not vals:
            print(f"WARNING: no scored grid point in {c}:{s}-{e}")
            continue
        span = int(r["span_kb"])
        interval = (f"{s / 1e6:.3f}" if e == s
                    else f"{s / 1e6:.3f}-{e / 1e6:.3f}")
        ds = [post[(c, p)] for p in pts if (c, p) in post]
        # Every printed quantity is averaged over the locus's grid points, so
        # the interval is the typical posterior spread within the locus rather
        # than the spread of the locus-level estimate.
        def avg(k):
            return sum(d[k] for d in ds) / len(ds)
        ci, centre = {}, {}
        for tag in ("afr", "nea", "den"):
            if a.interval_style == "central":
                centre[tag] = avg(f"{tag}_mean")
                lo, hi = avg(f"{tag}_p025"), avg(f"{tag}_p975")
                ci[tag] = fmt_t(centre[tag], lo, hi)
                spread = hi - lo
            elif a.interval_style == "central90":
                centre[tag] = avg(f"{tag}_hi0.5")
                lo, hi = avg(f"{tag}_lo0.95"), avg(f"{tag}_hi0.95")
                ci[tag] = fmt_central90(centre[tag], lo, hi)
                spread = hi - lo
            else:
                centre[tag] = avg(f"{tag}_hi0.5")
                if tag == "den":
                    bound = avg(f"{tag}_lo0.95")
                    ci[tag] = fmt_one_sided(centre[tag], bound, "\u003e")
                else:
                    bound = avg(f"{tag}_hi0.95")
                    ci[tag] = fmt_one_sided(centre[tag], bound, "\u003c")
                spread = abs(bound - centre[tag])
            if tag == "nea":
                widths.append((spread, c, interval))
        lab = {"central": "95% CI", "central90": "90% CI",
               "one-sided": "95th pct"}[a.interval_style]
        dlab = {"central": "95% CI", "central90": "90% CI",
                "one-sided": "5th pct"}[a.interval_style]
        # How much deeper the Denisova-inclusive coalescence is than the
        # Neandertal-inclusive one. This is the catalog's claim reduced to a
        # single number, and it is positive whenever the locus behaves as the
        # rule intends. In the central90 style it is given as a distribution:
        # the difference is drawn at each grid point by diff_quantiles and the
        # resulting percentiles are averaged over the locus, exactly as the
        # three TMRCA columns are. The point estimate stays the difference of
        # the two printed medians so that the column reconciles by eye with the
        # columns on either side of it; the median of the difference is within
        # a few kiloyears of it but would not subtract cleanly on the page.
        if a.interval_style == "central90":
            dq = [diff_quantiles(coord, d) for d in ds]
            dlo = sum(q[0] for q in dq) / len(dq)
            dhi = sum(q[2] for q in dq) / len(dq)
            diff_cell = fmt_central90(centre["den"] - centre["nea"], dlo, dhi)
        else:
            diff_cell = f"{centre['den'] - centre['nea']:.0f}"
        out.append({
            "chr": c, "start": s,
            "interval (Mb)": interval, "span (kb)": span,
            "P (real)": f"{max(vals):.2f}",
            f"TAfr, ka ({lab})": ci["afr"],
            f"TAfr+Nea, ka ({lab})": ci["nea"],
            f"TAfr+Den, ka ({dlab})": ci["den"],
            "TAfr+Den \u2212 TAfr+Nea, ka": diff_cell,
            "genes within 5 kb": gene_label(r["genes"],
                                            r.get("nearest_if_none", "")),
        })

    out.sort(key=lambda r: (r["chr"], r["start"]))
    cols = [k for k in out[0] if k != "start"]
    with open(a.out, "w", newline="", encoding="utf-8") as fh:
        w = csv.DictWriter(fh, fieldnames=cols, delimiter="\t",
                           extrasaction="ignore")
        w.writeheader(); w.writerows(out)
    keys = {(int(r["chrom"]), int(r["pos"])) for r in sites}
    cu = sum(1 for k, v in clamped.items() if k in keys and "u" in v)
    cv = sum(1 for k, v in clamped.items() if k in keys and "v" in v)
    print(f"wrote {a.out} ({len(out)} rows); of the {len(sites)} catalog "
          f"grid points, {cv} were scored at the deep-Denisovan boundary "
          f"and {cu} at the shallow-Neandertal boundary")

    pv = sorted(float(r["P (real)"]) for r in out)
    print(f"P(real) over the {len(out)} loci: min {pv[0]:.2f}, "
          f"median {pv[len(pv) // 2]:.2f}, max {pv[-1]:.2f}")

    widths.sort()
    ws = [w for w, _, _ in widths]
    wlab = {"central": "95% interval", "central90": "90% interval",
            "one-sided": "median-to-95th gap"}[a.interval_style]
    print(f"\nwidth of the TAfr+Nea {wlab}, over {len(ws)} loci: "
          f"min {ws[0]:.0f}, median {ws[len(ws) // 2]:.0f}, "
          f"max {ws[-1]:.0f} ka")
    dl = sorted(float(r["TAfr+Den \u2212 TAfr+Nea, ka"].split()[0])
                for r in out)
    neg = sum(1 for d in dl if d <= 0)
    print(f"TAfr+Den minus TAfr+Nea over {len(dl)} loci: "
          f"min {dl[0]:.0f}, median {dl[len(dl) // 2]:.0f}, "
          f"max {dl[-1]:.0f} ka; {neg} loci at or below zero")
    if a.interval_style == "central90":
        # A locus whose interval covers zero has not, on its own, established
        # that the two archaic coalescences differ, however large the gap
        # between the two point estimates looks.
        cov = sum(1 for r in out if float(
            r["TAfr+Den \u2212 TAfr+Nea, ka"].split("(")[1]
            .split("\u2013")[0]) <= 0)
        print(f"  {cov} of {len(out)} loci have a 90% interval on the "
              f"difference that reaches zero or below")
    print("  widest 8 loci (the ones to discount):")
    for w, c, iv in widths[-8:][::-1]:
        print(f"    chr{c}:{iv:<16} interval spans {w:.0f} ka")
    print("top 8 loci by P(real):")
    for r in sorted(out, key=lambda r: -float(r["P (real)"]))[:8]:
        print(f"  {r['chr']:>2}:{r['interval (Mb)']:<16} "
              f"P={r['P (real)']}  {r['genes within 5 kb'][:44]}")
    return 0


if __name__ == "__main__":
    sys.exit(main())
```
