## Supplementary Information section 4 code base for "Hypothesis: A modern human range expansion ∼300,000 years ago had a critical role in Neandertal origins": null_ratio_audit.py

```
#!/usr/bin/env python3
"""
Is omega0 = 1.150 a null, or does it contain real signal?

omega0 normalises the case/control ratio into P(real) = 1 - omega0/omega, so it
has to be the ratio the mirrored design would produce with no sweeps anywhere.
The value in use, 1.150, came from 2,708 Neandertal-direction against 2,355
Denisovan-direction windows "inside the fitted support". But that support is
the whole binned domain of the surface fit -- nea_hi anywhere in [350, 900] and
den_lo anywhere in [350, 1500] -- which contains every catalog window and every
other strong-signal window in the genome. Whatever real signal exists is inside
that count, so it cannot serve as a null: normalising by it subtracts the signal
from itself.

This script does three things.

1. Confirms the contamination directly, by asking how many catalog windows sit
   inside the region where omega0 was measured.

2. Estimates the null in regions where signal is not possible. The published
   modern-archaic split is 825-694 ka, and the catalog's premise is that a
   modern-archaic coalescence MORE RECENT than the split is the sweep
   signature. So a window whose join-side bound sits above 825 ka carries no
   such evidence. The cumulative test u <= x cannot isolate those windows
   because it always includes everything below x, so this uses shells,
   x_lo < u <= x_hi, entirely above the split. The mirrored rule gets the
   identical shell, so the comparison stays fair.

3. Checks the SIGN of the structural bias that omega0 is supposed to capture.
   The stated mechanism is that TMRCA(Afr+Nea) is the MRCA of 20 haplotypes and
   TMRCA(Afr+Den) of 18, and subset-MRCA depth is non-linear in lineage count.
   But that mechanism predicts a DEEPER Neandertal statistic, which makes the
   case rule (shallow Nea, deep Den) harder and the control rule easier, so it
   predicts omega0 < 1. The value in use is above 1. If the genome-wide
   distributions confirm the Neandertal side is the deeper one, then 1.150 has
   the wrong sign for its own explanation, which is independent evidence that
   it is signal and not structure.

Example:
  python3 null_ratio_audit.py --scan hubisz_tmrca_5kb_8afr.tsv
"""

from __future__ import annotations

import argparse
import csv
import math
import os
import sys

import numpy as np

sys.path.insert(0, os.path.dirname(os.path.abspath(__file__)))

from approx_percentiles import default_coord, load  # noqa: E402
from optimize_rule_by_size import tails  # noqa: E402

CUT = 0.50          # the catalog's percentile cut (posterior median)
CAT_X = 625.0       # catalog Neandertal threshold
CAT_Y = 700.0       # catalog Denisovan threshold
SPLIT_HI = 825.0    # upper end of the published modern-archaic split interval


def wilson_ratio(k, n, z=1.96):
    """95% CI for the odds k/n, via the Wilson interval on k/(k+n)."""
    if k + n == 0:
        return float("nan"), float("nan")
    m = k + n
    q = k / m
    d = 1 + z * z / m
    c = (q + z * z / (2 * m)) / d
    h = z * math.sqrt(q * (1 - q) / m + z * z / (4 * m * m)) / d
    out = []
    for b in (max(0.0, c - h), min(1.0, c + h)):
        out.append(float("inf") if b >= 1 else b / (1 - b))
    return out[0], out[1]


def main():
    ap = argparse.ArgumentParser()
    ap.add_argument("--scan", default="hubisz_tmrca_5kb_8afr.tsv")
    ap.add_argument("--catalog", default="catalog_annotation/"
                    "catalog_final_median_x625_y700_auto.sites.tsv")
    args = ap.parse_args()

    recs = load(args.scan)
    coord, grid_desc = default_coord(recs)
    recs = [d for d in recs if d["chrom"].replace("chr", "") not in ("X", "23")]
    T = tails(recs, coord)
    print(f"time coordinate: {grid_desc}")
    print(f"{len(recs):,} autosomal windows\n")

    nh = T[("nea", "hi", CUT)]
    nl = T[("nea", "lo", CUT)]
    dh = T[("den", "hi", CUT)]
    dl = T[("den", "lo", CUT)]

    # ---------------------------------------------------------------- part 1
    # Reproduce the support-wide count and show what is inside it.
    print("=" * 72)
    print("1. WHAT REGION WAS omega0 MEASURED OVER?")
    print("=" * 72)
    # The surface's binned support, as fit_fdr_surface defines it.
    in_case = (nh >= 350) & (nh <= 900) & (dl >= 350) & (dl <= 1500)
    in_ctrl = (dh >= 350) & (dh <= 900) & (nl >= 350) & (nl <= 1500)
    kn, kd = int(in_case.sum()), int(in_ctrl.sum())
    print(f"  Neandertal-direction windows in support: {kn:,}")
    print(f"  Denisovan-direction  windows in support: {kd:,}")
    print(f"  odds = {kn / kd:.4f}   <- the 1.150 in the note")
    # How much of that is the catalog and its immediate neighbourhood?
    cat = (nh <= CAT_X) & (dl >= CAT_Y)
    print(f"\n  catalog windows (nea_hi<={CAT_X:.0f}, den_lo>={CAT_Y:.0f}): "
          f"{int(cat.sum()):,}")
    print(f"  of those, inside the omega0 region: {int((cat & in_case).sum()):,}")
    strong = (nh <= 700) & in_case
    print(f"  windows with nea_hi <= 700 ka inside it: {int(strong.sum()):,}")
    print("\n  -> the region omega0 was measured over contains the catalog and")
    print("     every other shallow-Neandertal window, so it is not a null.")

    # ---------------------------------------------------------------- part 2
    print("\n" + "=" * 72)
    print("2. THE NULL, MEASURED ABOVE THE SPLIT (SHELLS, NOT CUMULATIVE)")
    print("=" * 72)
    print(f"  A window whose join-side bound exceeds {SPLIT_HI:.0f} ka (the top of")
    print("  the published 825-694 ka split) carries no sweep evidence.")
    print("  Each shell applies the identical test in both directions.\n")
    print(f"{'shell (ka)':>14} {'y (ka)':>7} {'nea':>7} {'den':>7} "
          f"{'odds':>7} {'95% CI':>17}")
    print("-" * 66)
    rows = []
    for lo, hi in [(825, 875), (875, 925), (925, 1000), (1000, 1100),
                   (1100, 1300), (825, 925), (825, 1000), (825, 1100)]:
        for y in (CAT_Y, float(hi) + 75.0):
            c = int(np.count_nonzero((nh > lo) & (nh <= hi) & (dl > y)))
            k = int(np.count_nonzero((dh > lo) & (dh <= hi) & (nl > y)))
            if c == 0 or k == 0:
                continue
            r = c / k
            a, b = wilson_ratio(c, k)
            print(f"{f'{lo}-{hi}':>14} {y:>7.0f} {c:>7,} {k:>7,} "
                  f"{r:>7.3f} {f'{a:.3f}-{b:.3f}':>17}")
            rows.append((lo, hi, y, c, k, r))
    if rows:
        wide = [r for r in rows if (r[0], r[1]) == (825, 1100)]
        print()
        for lo, hi, y, c, k, r in wide:
            a, b = wilson_ratio(c, k)
            print(f"  widest signal-free shell {lo}-{hi} ka at y={y:.0f}: "
                  f"odds {r:.3f} (95% CI {a:.3f}-{b:.3f})")

    # ---------------------------------------------------------------- part 3
    print("\n" + "=" * 72)
    print("3. DOES 1.150 EVEN HAVE THE SIGN ITS EXPLANATION PREDICTS?")
    print("=" * 72)
    print("  Mechanism claimed: T(Afr+Nea) spans 20 haplotypes, T(Afr+Den) 18;")
    print("  more lineages means a deeper MRCA. A deeper Neandertal statistic")
    print("  makes the case rule harder and the control rule easier, i.e. it")
    print("  predicts omega0 < 1.\n")
    nmed = T[("nea", "hi", CUT)]
    dmed = T[("den", "hi", CUT)]
    print(f"  genome-wide median of the Neandertal median bound: {np.median(nmed):>8.1f} ka")
    print(f"  genome-wide median of the Denisovan  median bound: {np.median(dmed):>8.1f} ka")
    frac = float(np.count_nonzero(nmed > dmed)) / len(nmed)
    print(f"  windows where the Neandertal side is DEEPER: {frac * 100:.1f}%")
    verdict = ("deeper -> mechanism predicts omega0 < 1, so 1.150 has the "
               "WRONG SIGN" if np.median(nmed) > np.median(dmed) else
               "shallower -> mechanism is consistent with omega0 > 1")
    print(f"  Neandertal side is {verdict}")

    # ---------------------------------------------------------------- part 4
    print("\n" + "=" * 72)
    print("4. CONSEQUENCE FOR THE CATALOG'S PURITY")
    print("=" * 72)
    ncase = int(np.count_nonzero((nh <= CAT_X) & (dl > CAT_Y)))
    nctrl = int(np.count_nonzero((dh <= CAT_X) & (nl > CAT_Y)))
    omega = ncase / nctrl
    print(f"  catalog: {ncase} case, {nctrl} control -> ratio "
          f"{nctrl / ncase:.4f}, omega = {omega:.4f}\n")
    print(f"{'omega0':>8} {'P(real)':>9}  basis")
    print("-" * 62)
    for w0, basis in [(1.150, "support-wide count (signal-contaminated)"),
                      (1.000, "exact mirror symmetry"),
                      (0.900, "if the structural bias runs the other way")]:
        print(f"{w0:>8.3f} {1 - w0 / omega:>9.3f}  {basis}")
    if rows:
        for lo, hi, y, c, k, r in rows:
            if (lo, hi, y) == (825, 1100, CAT_Y):
                print(f"{r:>8.3f} {1 - r / omega:>9.3f}  "
                      f"measured in the {lo}-{hi} ka signal-free shell")


if __name__ == "__main__":
    main()
```
