## Supplementary Information section 4 code base for "Hypothesis: A modern human range expansion ∼300,000 years ago had a critical role in Neandertal origins": null_ratio_shell_bstat.py

```
#!/usr/bin/env python3
"""
Where did omega0 = 1.150 come from, and is the shell null flat in B?

Two questions, both follow-ups to null_ratio_audit.py.

(a) PROVENANCE. The note attributes 1.150 to 2,708 Neandertal-direction against
    2,355 Denisovan-direction windows. Recomputing the case/control counts over
    the surface's binned support gives 7,844 / 7,448, which is 1.053, not 1.150,
    and 15,292 windows rather than 5,063. The suspected reason is that those
    counts came out of the B-statistic diagnostic, which silently drops every
    window the B map does not cover. If so, 1.150 is the odds within an
    arbitrary third of the support -- arbitrary for this purpose, since B
    coverage has nothing to do with whether a window is a null.

(b) FLATNESS. omega0 is now measured in shells above the published split, where
    sweep signal is not possible. The reason for binning by B in the first place
    was that the structural bias might itself depend on B, which would force a
    local rather than a global normalisation. That check has to be redone where
    omega0 is now measured, rather than over the contaminated support.

Example:
  python3 null_ratio_shell_bstat.py --scan hubisz_tmrca_5kb_8afr.tsv
"""

from __future__ import annotations

import argparse
import csv
import math
import os
import sys

import numpy as np

sys.path.insert(0, os.path.dirname(os.path.abspath(__file__)))

from approx_percentiles import default_coord, load  # noqa: E402
from optimize_rule_by_size import tails  # noqa: E402

CUT = 0.50
CAT_Y = 700.0
SHELL = (825.0, 1100.0)


def load_bstat(path):
    """(chrom, pos) -> B, applying the same coverage filter as the diagnostic."""
    b = {}
    with open(path) as fh:
        for r in csv.DictReader(fh, delimiter="\t"):
            v = r.get("bstat", "NA")
            if v in ("", "NA"):
                continue
            try:
                frac = float(r.get("bstat_covered_frac", 1.0))
            except ValueError:
                frac = 1.0
            if frac < 0.5:
                continue
            try:
                b[(r["chrom"].replace("chr", ""), int(r["pos"]))] = float(v)
            except ValueError:
                continue
    return b


def wilson(k, n, z=1.96):
    m = k + n
    if m == 0:
        return float("nan"), float("nan")
    q = k / m
    d = 1 + z * z / m
    c = (q + z * z / (2 * m)) / d
    h = z * math.sqrt(q * (1 - q) / m + z * z / (4 * m * m)) / d
    out = []
    for x in (max(0.0, c - h), min(1.0, c + h)):
        out.append(float("inf") if x >= 1 else x / (1 - x))
    return out[0], out[1]


def main():
    ap = argparse.ArgumentParser()
    ap.add_argument("--scan", default="hubisz_tmrca_5kb_8afr.tsv")
    ap.add_argument("--bstat",
                    default="catalog_annotation/bstat.phastCons_bestfit.5kb.tsv")
    ap.add_argument("--bstat2",
                    default="catalog_annotation/bstat.CADD_bestfit.5kb.tsv")
    args = ap.parse_args()

    recs = load(args.scan)
    coord, _ = default_coord(recs)
    recs = [d for d in recs if d["chrom"].replace("chr", "") not in ("X", "23")]
    T = tails(recs, coord)
    nh, nl = T[("nea", "hi", CUT)], T[("nea", "lo", CUT)]
    dh, dl = T[("den", "hi", CUT)], T[("den", "lo", CUT)]

    lo, hi = SHELL
    case = (nh > lo) & (nh <= hi) & (dl > CAT_Y)
    ctrl = (dh > lo) & (dh <= hi) & (nl > CAT_Y)

    # ------------------------------------------------------------- provenance
    print("=" * 70)
    print("(a) PROVENANCE OF 1.150")
    print("=" * 70)
    in_case = (nh >= 350) & (nh <= 900) & (dl >= 350) & (dl <= 1500)
    in_ctrl = (dh >= 350) & (dh <= 900) & (nl >= 350) & (nl <= 1500)
    print(f"  support-wide, all windows:  {int(in_case.sum()):>6,} / "
          f"{int(in_ctrl.sum()):>6,} = {in_case.sum() / in_ctrl.sum():.4f}")

    for label, path in (("phastCons", args.bstat), ("CADD", args.bstat2)):
        if not os.path.exists(path):
            continue
        B = load_bstat(path)
        keys = [(d["chrom"].replace("chr", ""), int(d[coord.poskey]))
                if hasattr(coord, "poskey") else
                (d["chrom"].replace("chr", ""), int(d["pos"])) for d in recs]
        hasB = np.array([k in B for k in keys])
        c = int((in_case & hasB).sum())
        k = int((in_ctrl & hasB).sum())
        print(f"  support-wide, B-covered ({label:>9}): {c:>6,} / {k:>6,} "
              f"= {c / k:.4f}")
        if abs(c / k - 1.150) < 0.02:
            print(f"       ^ reproduces the note's 1.150 "
                  f"(cited counts were 2,708 / 2,355)")

    # ---------------------------------------------------------------- flatness
    print("\n" + "=" * 70)
    print(f"(b) IS THE SHELL NULL FLAT IN B?  shell {lo:.0f}-{hi:.0f} ka, "
          f"y={CAT_Y:.0f}")
    print("=" * 70)
    for label, path in (("phastCons", args.bstat), ("CADD", args.bstat2)):
        if not os.path.exists(path):
            continue
        B = load_bstat(path)
        keys = [(d["chrom"].replace("chr", ""), int(d["pos"])) for d in recs]
        bv = np.array([B.get(k, np.nan) for k in keys])
        ok = ~np.isnan(bv)
        tagged = ok & (case | ctrl)
        if tagged.sum() == 0:
            continue
        qs = np.quantile(bv[tagged], np.linspace(0, 1, 11))
        print(f"\n  {label} map, {int(tagged.sum()):,} shell windows with B")
        print(f"  {'dec':>4} {'B range':>15} {'nea':>7} {'den':>7} "
              f"{'odds':>7} {'95% CI':>15}")
        print("  " + "-" * 60)
        ks, ns, xs = [], [], []
        for i in range(10):
            a, b = qs[i], qs[i + 1]
            sel = tagged & (bv >= a) & (bv <= b if i == 9 else bv < b)
            c = int((sel & case).sum())
            k = int((sel & ctrl).sum())
            if c + k == 0:
                continue
            w, v = wilson(c, k)
            print(f"  {i + 1:>4} {f'{a:.3f}-{b:.3f}':>15} {c:>7,} {k:>7,} "
                  f"{c / k if k else float('nan'):>7.3f} "
                  f"{f'{w:.3f}-{v:.3f}':>15}")
            ks.append(c)
            ns.append(c + k)
            xs.append(i + 1)
        # Cochran-Armitage trend test on the Neandertal-direction fraction.
        tot_k, tot_n = sum(ks), sum(ns)
        pbar = tot_k / tot_n
        xbar = sum(n * x for n, x in zip(ns, xs)) / tot_n
        num = sum((k - n * pbar) * (x - xbar) for k, n, x in zip(ks, ns, xs))
        den = pbar * (1 - pbar) * sum(n * (x - xbar) ** 2
                                      for n, x in zip(ns, xs))
        z = num / math.sqrt(den) if den > 0 else float("nan")
        p = 2 * (1 - 0.5 * (1 + math.erf(abs(z) / math.sqrt(2))))
        odds = [k / (n - k) for k, n in zip(ks, ns) if n - k > 0]
        print(f"  pooled odds {tot_k / (tot_n - tot_k):.4f}; "
              f"spread {min(odds):.3f}-{max(odds):.3f}")
        print(f"  Cochran-Armitage trend: z = {z:+.2f}, p = {p:.2f} -> "
              f"{'no trend, global omega0 suffices' if p > 0.05 else 'TREND'}")


if __name__ == "__main__":
    main()
```
