## Supplementary Information section 4 code base for "Hypothesis: A modern human range expansion ∼300,000 years ago had a critical role in Neandertal origins": o2_build_venv.sh

```
#!/bin/bash
# Build the python env the fixed-derived screen needs, then verify it.
#
#   ssh 'bash -l -s' < scripts/o2_build_venv.sh
#
# Note 'bash -l -s' rather than 'bash -s': `module` is a shell function defined
# by the login profile, so a non-login shell reports "module: command not found".
#
# Idempotent -- reuses an existing venv if pysam already imports, so it is safe
# to re-run.

set -u

ENRICH=/n/groups/reich/DAVID/CURSOR_ENRICH
VENV="$ENRICH/work/venv"

echo "=== loading modules ==="
# gcc/9.2.0 and python/3.9.14 no longer exist on O2 (Lmod reports them unknown);
# the default python3 is /n/app/python/3.13.1-gcc-14.2.0, which is fine -- pysam
# ships a cp313 manylinux wheel. Try the module anyway in case a site reinstates
# it, but do not let Lmod's error text look like a failure of this script.
module load gcc/9.2.0 python/3.9.14 >/dev/null 2>&1 \
  || echo "  (no gcc/python module; using default python3, which is expected)"
echo "  python3: $(command -v python3)"
echo "  version: $(python3 --version 2>&1)"

if [ -f "$VENV/bin/activate" ]; then
  # shellcheck disable=SC1091
  source "$VENV/bin/activate"
  if python3 -c 'import pysam' 2>/dev/null; then
    echo "=== venv already good: pysam $(python3 -c 'import pysam;print(pysam.__version__)') ==="
    exit 0
  fi
  echo "=== venv exists but pysam missing; installing into it ==="
else
  echo "=== creating $VENV ==="
  mkdir -p "$ENRICH/work" || { echo "ERROR: cannot mkdir $ENRICH/work" >&2; exit 1; }
  python3 -m venv "$VENV" || { echo "ERROR: venv creation failed" >&2; exit 1; }
  # shellcheck disable=SC1091
  source "$VENV/bin/activate"
fi

echo "=== installing pysam ==="
python3 -m pip install --upgrade pip 2>&1 | tail -2 | sed 's/^/  /'
# pysam publishes manylinux wheels for cp39, so this should not compile htslib.
python3 -m pip install pysam 2>&1 | tail -5 | sed 's/^/  /'

echo "=== verifying ==="
python3 - <<'PY'
import sys
try:
    import pysam
except Exception as e:
    print(f"FAIL: pysam did not import: {e}")
    sys.exit(1)
print(f"pysam {pysam.__version__} at {pysam.__file__}")
# The screen uses exactly these three; check them rather than just the import,
# because a wheel built without libcurl silently lacks remote/tabix support.
for attr in ("FastaFile", "TabixFile", "VariantFile"):
    print(f"  {attr}: {'OK' if hasattr(pysam, attr) else 'MISSING'}")
PY
rc=$?

echo
if [ "$rc" = "0" ]; then
  echo "=== venv ready: $VENV ==="
  echo "Activate with:  source $VENV/bin/activate"
else
  echo "=== venv build FAILED ===" >&2
fi
exit "$rc"
```
