## Supplementary Information section 4 code base for "Hypothesis: A modern human range expansion ∼300,000 years ago had a critical role in Neandertal origins": o2_sbatch_fixed_derived.sh

```
#!/bin/bash
# Slurm array: screen catalog clusters for fixed derived sites, one task per
# chromosome. Each task is small (a few hundred kb of sequence and a handful of
# tabix lookups), so this is short and low-memory; the whole array should
# finish in minutes now that nothing is fetched over the network.
#
# SETUP (once, on O2)
#   WORK=/n/groups/reich/DAVID/CURSOR_ENRICH/fixed_derived
#   mkdir -p $WORK/out $WORK/logs && cd $WORK
#   # copy from the laptop:
#   #   scripts/o2_screen_fixed_derived.py
#   #   scripts/o2_sbatch_fixed_derived.sh
#   #   catalog_annotation/preal50.regions.tsv
#
# STEP 1 - verify inputs and the codon logic before burning any allocation.
#   module load gcc/9.2.0 python/3.9.14
#   source /n/groups/reich/DAVID/CURSOR_ENRICH/work/run1/venv/bin/activate 2>/dev/null || true
#   python3 o2_screen_fixed_derived.py --check-paths --selftest
#   # expect every path 'ok' and the selftest to print SELFTEST PASS
#   #   (Leu111Arg at chr9:84,605,717; the old Ile->Met @205 case was
#   #    built on a rounded coordinate and could never pass)
#
# STEP 2 - launch.
#   sbatch o2_sbatch_fixed_derived.sh
#
# STEP 3 - collate when the array finishes.
#   python3 o2_screen_fixed_derived.py --collate out/fixed_derived
#   # writes out/fixed_derived.all.tsv and prints the missense list
#
#SBATCH --job-name=fixdrv
#SBATCH --array=1-22
#SBATCH --partition=short
#SBATCH --time=0-02:00:00
#SBATCH --mem=8G
#SBATCH --cpus-per-task=1
#SBATCH --output=logs/fixdrv_%A_%a.out
#SBATCH --error=logs/fixdrv_%A_%a.err

set -euo pipefail

CHROM="${SLURM_ARRAY_TASK_ID}"
REGIONS="${REGIONS:-preal50.regions.tsv}"

# The screen adds PAD to each region's end only, never its start. Region files
# that already carry their padding must therefore set PAD=0, or every interval
# is silently shifted downstream by one pad width.
PAD="${PAD:-5000}"
OUTPREFIX="${OUTPREFIX:-out/fixed_derived}"

# Require Neandertals to carry the derived allele as well: the catalog premise
# is a variant shared by modern humans and Neandertals but not Denisova.
# Set NEANDERTAL=none to reproduce the looser screen that only conditions on
# modern humans and Denisova.
NEANDERTAL="${NEANDERTAL:-both}"

# Add Vindija here if a local copy exists; the CURSOR_ENRICH tree has Altai
# only, so by default the Neandertal condition rests on one genome. This is the
# legacy path and is not how the panel is populated: PANEL inside the screen
# script carries five Neandertals and two Denisovans off the Reich filesystem.
EXTRA_NEA="${EXTRA_NEA:-}"

# Restrict the archaic panel by genome id, empty meaning all of PANEL. Set this
# to run the four classic high-coverage Neandertals and drop GoyetQ56-1, which
# comes from the late-Neandertal release rather than the high-coverage set:
#   PANEL_ONLY=Altai,Vindija33.19,Chagyrskaya8,Denisova17,Denisova25
# The driver genome is added back automatically.
PANEL_ONLY="${PANEL_ONLY:-}"

# Maximum ancestral allele frequency in 1000 Genomes, i.e. one minus the floor
# on modern derived frequency. The screen applies this while scanning rather
# than at reporting, so a site above the cutoff is never emitted and the count
# at a looser threshold cannot be recovered from a stricter run's output; it
# needs its own pass. 0.01 is the >99% screen in the note, 0.05 the >95% one.
MAXAF="${MAXAF:-0.01}"

# No module load: gcc/9.2.0 and python/3.9.14 are gone from O2, and the venv
# carries its own interpreter (python 3.13.1, pysam 0.24.0) which activate puts
# first on PATH, so nothing else is needed.
# work/venv is the current env; work/run1/venv was an earlier one that no longer
# exists. Try both, then fail fast rather than running 22 tasks that each die on
# the pysam import.
for V in /n/groups/reich/DAVID/CURSOR_ENRICH/work/venv \
         /n/groups/reich/DAVID/CURSOR_ENRICH/work/run1/venv; do
  if [ -f "$V/bin/activate" ]; then
    # shellcheck disable=SC1091
    source "$V/bin/activate"
    echo "venv: $V"
    break
  fi
done
python3 -c 'import pysam' 2>/dev/null || {
  echo "ERROR: pysam not importable; build the venv first (see o2_build_venv.sh)" >&2
  exit 2
}

mkdir -p "$(dirname "$OUTPREFIX")" logs

echo "chr${CHROM}  regions=${REGIONS}  pad=${PAD}  neandertal=${NEANDERTAL}"
echo "max ancestral AF=${MAXAF}"
echo "panel_only=${PANEL_ONLY:-<all of PANEL>}"
python3 o2_screen_fixed_derived.py \
  --chrom "${CHROM}" \
  --regions "${REGIONS}" \
  --pad "${PAD}" \
  --neandertal "${NEANDERTAL}" \
  --max-af "${MAXAF}" \
  ${EXTRA_NEA:+--extra-nea "${EXTRA_NEA}"} \
  ${PANEL_ONLY:+--panel-only "${PANEL_ONLY}"} \
  -o "${OUTPREFIX}"
```
