## Supplementary Information section 4 code base for "Hypothesis: A modern human range expansion ∼300,000 years ago had a critical role in Neandertal origins": o2_screen_fixed_derived.py

```
#!/usr/bin/env python3
"""
O2 version of the fixed-derived screen: all inputs local, one chromosome per
task, and amino acid consequences computed from GRCh37.fa rather than by
querying Ensembl.

WHAT IT LOOKS FOR
At every position inside a catalog cluster:

  ancestral   Ensembl 6-primate EPO ancestral allele (GRCh37 e71). Uppercase
              only, which are the high-confidence calls.
  derived     the hg19 reference base, required to differ from the ancestral
              base, so the modern human reference carries a derived allele.
  modern      derived and fixed or near-fixed: absent from 1000 Genomes
              Phase 3, or present with ancestral-allele frequency <= --max-af.
  Denisova    homozygous ANCESTRAL, at depth >= --min-dp and GQ >= --min-gq.
  Neandertal  homozygous DERIVED (--neandertal both|any|none controls whether
              this is required or merely recorded).

The Neandertal requirement is the part the earlier streaming screen recorded
but did not enforce. The catalog premise is that the derived allele is shared
by modern humans AND Neandertals to the exclusion of Denisova, so `both` is
the default here.

AMINO ACID CONSEQUENCES
For sites in a CDS, the script rebuilds each affected transcript's coding
sequence from the GENCODE v19 CDS exons and GRCh37.fa, maps the genomic
position to a coding index, and translates the codon with the derived base
against the codon with the ancestral base. Handling the full transcript rather
than a single exon means codons that straddle an exon boundary are correct.

Sanity check to run first (validated against Ensembl GRCh37 VEP):
  chr9:84,606,000 A(ancestral) vs G(derived) in SPATA31D1, transcript
  ENST00000344803, should come out missense, Ile->Met, protein position 205.
  Run with --selftest to check exactly this before launching the array.

Usage
  # 0. verify every input path exists and the codon logic is right
  python3 o2_screen_fixed_derived.py --check-paths --selftest

  # 1. one chromosome (what the Slurm array runs)
  python3 o2_screen_fixed_derived.py --chrom 9 \
      --regions preal50.regions.tsv -o out/fixed_derived

  # 2. collate
  python3 o2_screen_fixed_derived.py --collate out/fixed_derived
"""

from __future__ import annotations

import argparse
import csv
import glob
import gzip
import os
import sys
from collections import defaultdict

# pysam is imported lazily so that --check-paths and --collate, which only
# stat files and concatenate TSVs, stay usable on a login node where pysam is
# not installed. A top-level import made them fail before argparse ever ran.
# --selftest and the main per-chromosome run do need it, since both read the
# reference FASTA.
pysam = None


def need_pysam():
    global pysam
    if pysam is None:
        try:
            import pysam as _p
        except ImportError:
            sys.stderr.write(
                "ERROR: pysam is required for this mode but is not installed.\n"
                "On O2, try:\n"
                "  source /n/groups/reich/DAVID/CURSOR_ENRICH/work/run1/venv/bin/activate\n"
                "or build one:\n"
                "  module load gcc/9.2.0 python/3.9.14\n"
                "  python3 -m venv ~/pysam-venv && source ~/pysam-venv/bin/activate\n"
                "  pip install --upgrade pip && pip install pysam\n")
            raise SystemExit(2)
        pysam = _p
    return pysam


# ---------------------------------------------------------------- data paths
ENRICH = "/n/groups/reich/DAVID/CURSOR_ENRICH"
DEF_REF = f"{ENRICH}/data/ref/GRCh37.fa"
DEF_ANC = f"{ENRICH}/data/ref/homo_sapiens_ancestor_GRCh37_merged.fa"
DEF_GTF = f"{ENRICH}/data/gencode/gencode.v19.annotation.gtf.gz"
DEF_DEN = f"{ENRICH}/data/archaic_den/den_meyer_autosomes.vcf.gz"
DEF_NEA = f"{ENRICH}/data/archaic_nea/nea_altai_autosomes.vcf.gz"
# Phase 3 lives in two places in this tree; both patterns are tried.
DEF_G1K_GLOBS = [f"{ENRICH}/data/phase3/ALL.chr{{c}}.*.vcf.gz",
                 f"{ENRICH}/refs/phase3_20130502/ALL.chr{{c}}.*.vcf.gz"]

CODONS = {}
_B = "TCAG"
_AA = ("FFLLSSSSYY**CC*W" "LLLLPPPPHHQQRRRR"
       "IIIMTTTTNNKKSSRR" "VVVVAAAADDEEGGGG")
for _i, _a in enumerate(_AA):
    CODONS[_B[_i // 16] + _B[(_i // 4) % 4] + _B[_i % 4]] = _a

COMP = str.maketrans("ACGTNacgtn", "TGCANtgcan")

THREE = {"A": "Ala", "R": "Arg", "N": "Asn", "D": "Asp", "C": "Cys",
         "Q": "Gln", "E": "Glu", "G": "Gly", "H": "His", "I": "Ile",
         "L": "Leu", "K": "Lys", "M": "Met", "F": "Phe", "P": "Pro",
         "S": "Ser", "T": "Thr", "W": "Trp", "Y": "Tyr", "V": "Val",
         "*": "Ter"}


def rc(s):
    return s.translate(COMP)[::-1]


def translate(codon):
    return CODONS.get(codon.upper(), "?")


# ------------------------------------------------------------------ helpers
def contig_name(tbx_or_fasta, chrom, is_fasta=False):
    """Match '9' vs 'chr9' vs ancestral-style naming against the file.

    The Ensembl ancestral-sequence FASTAs name each record

        ANCESTOR_for_chromosome:GRCh37:10:1:135534747:1

    i.e. the chromosome is the *third* colon-separated field, not the first.
    Matching any field would be wrong rather than merely incomplete: looking for
    chromosome '1' would hit the start coordinate '1' in chromosome 10's name
    and silently return the wrong sequence. So the assembly-style name is
    matched at its known field position, and the loose any-field fallback is
    only accepted when exactly one record matches.
    """
    c = chrom.replace("chr", "")
    names = (list(tbx_or_fasta.references) if is_fasta
             else list(tbx_or_fasta.contigs))

    for cand in (c, f"chr{c}"):
        if cand in names:
            return cand

    # ANCESTOR_for_chromosome:<assembly>:<chrom>:<start>:<end>:<strand>
    for n in names:
        f = n.split(":")
        if len(f) >= 3 and f[0].startswith("ANCESTOR") and f[2] == c:
            return n

    # Some builds drop the ANCESTOR prefix but keep the field layout.
    for n in names:
        f = n.split(":")
        if len(f) >= 3 and f[2] == c and f[1].upper().startswith("GRC"):
            return n

    for n in names:                       # short prefix form, e.g. '10:1-135e6'
        if n.split(":")[0].replace("chr", "") == c:
            return n

    hits = [n for n in names
            if c in n.replace("chr", ":").replace("_", ":").split(":")]
    if len(hits) == 1:
        return hits[0]
    return None


def g1k_index_ok(path):
    """Is there a usable tabix/csi index next to this VCF?

    Existence is not enough. Phase 3 under data/phase3 had .tbi files that
    htslib rejected with EINVAL ("Invalid argument"), which is what a truncated
    or misnamed index looks like -- so the size is sanity-checked and, when
    pysam is already loaded, the index is actually opened.
    """
    for ext in (".tbi", ".csi"):
        idx = path + ext
        if not os.path.exists(idx):
            continue
        # A real index for a whole-chromosome VCF is never this small; a
        # zero-length or stub file is the common breakage.
        try:
            if os.path.getsize(idx) < 1000:
                continue
        except OSError:
            continue
        if pysam is None:          # cannot verify without pysam; size is enough
            return True
        try:
            pysam.TabixFile(path).close()
            return True
        except Exception:
            continue
    return False


def find_g1k(chrom, globs=None):
    """First phase 3 VCF for this chromosome that has a *loadable* index.

    Both candidate trees are scanned for a working index before falling back to
    merely existing, because the screen preferred data/phase3 and failed there
    while a second intact copy sat unused in refs/phase3_20130502.
    """
    pats = globs or DEF_G1K_GLOBS
    first = None
    for pat in pats:
        for hit in sorted(glob.glob(pat.format(c=chrom))):
            if first is None:
                first = hit
            if g1k_index_ok(hit):
                return hit
    # Nothing indexed. Return the first hit so the caller's error names a real
    # file rather than reporting "not found" for a VCF that is plainly there.
    return first


def sample_index(vcf_path, prefer=None):
    """Column index of the sample to use in a (possibly multi-sample) VCF."""
    op = gzip.open if vcf_path.endswith(".gz") else open
    with op(vcf_path, "rt", errors="replace") as f:
        for line in f:
            if line.startswith("#CHROM"):
                cols = line.rstrip("\n").split("\t")
                samples = cols[9:]
                if not samples:
                    return None, []
                if prefer:
                    for i, s in enumerate(samples):
                        if prefer.lower() in s.lower():
                            return 9 + i, samples
                return 9, samples
            if not line.startswith("#"):
                break
    return None, []


# --------------------------------------------------------- the archaic panel
# Five Neandertals and two Denisovans, all present on the Reich filesystem, all
# ALL-SITES hg19 VCFs with plain (non-chr) contig names -- verified 2026-09-05.
# Callers are mixed: Denisova 3 and Altai are GATK-era (FORMAT carries AD/PL/IR),
# the other four are snpAD (FORMAT carries PP). The mix is accepted deliberately.
# It matters mainly through GQ, which is not on a common scale between the two
# callers, so per-genome DP and GQ are written out for every site and the
# caller is recorded, letting any threshold effect be audited after the fact.
PANEL = [
    # id, lineage, directory, filename pattern ({c} = chromosome), caller
    ("Denisova3", "Denisovan",
     "/n/data1/hms/genetics/reich/1000Genomes/REORG/FOR_DATA_SHARING_DIRECTORY"
     "/sample_sets/denisova_dec11/vcfs",
     "DenisovaPinky.hg19_1000g.{c}.mod.vcf.gz", "GATK-era"),
    ("Altai", "Neandertal",
     "/n/data1/hms/genetics/reich/1000Genomes/nean_altai_jul12/hg19/A-raw/VCF_mod",
     "AltaiNea.hg19_1000g.{c}.mod.vcf.gz", "GATK-era"),
    ("Vindija33.19", "Neandertal",
     "/n/data1/hms/genetics/reich/1000Genomes/nean_mez_2016jul/hg19/A-raw"
     "/genotypes/Vindija33.19",
     "chr{c}_mq25_mapab100.vcf.gz", "snpAD"),
    ("Chagyrskaya8", "Neandertal",
     "/n/data1/hms/genetics/reich/1000Genomes/nean_chagyrskaya_2019mar"
     "/D-vcf_hicov/AA.raw",
     "chr{c}.noRB.vcf.gz", "snpAD"),
    ("Denisova25", "Denisovan",
     "/n/data1/hms/genetics/reich/1000Genomes/amh_samples"
     "/peyregneKelso2025__200kDenisova_sample__2025.11.05/AA.raw/VCF",
     "chr{c}.Den25.L35MQ25.B30.map35_100.vcf.gz", "snpAD"),
    ("GoyetQ56-1", "Neandertal",
     "/n/data1/hms/genetics/reich/1000Genomes/amh_samples"
     "/Mesa.Hajdinjak2026__Nature__PRJEB98484__edid586__27.late.neandertals"
     "__2026.09.05/A.round1/A.raw",
     "GoyetQ56_1_final.uniq.L35MQ25.sorted.chr_{c}.indel_realn_mq25"
     ".map35_100.vcf.gz", "snpAD"),
    # The fifth high-coverage Neandertal, released 2026-09-07. Its filenames
    # were not known when this was wired in, so the pattern is "*" and the
    # per-chromosome file is discovered by resolve_vcf below.
    ("Denisova17", "Neandertal",
     "/n/data1/hms/genetics/reich/1000Genomes/amh_samples"
     "/Massilani.Paabo2026__PNAS__prjeb108287__edid540__highCovNeanderthal"
     "/A.round1/A.raw/vcfs",
     "*", "snpAD"),
]

# Filename shapes seen across the archaic releases, each anchored by the
# delimiter that follows the chromosome number. The anchor is the point: a bare
# "*{c}*" glob would let chromosome 1 match chr11, chr12 and chr21, and picking
# the wrong chromosome's VCF would corrupt every call on it silently.
CHROM_GLOBS = ("*chr{c}.*", "*chr_{c}.*", "*chr{c}_*", "*.{c}.*", "*_{c}.*")


def resolve_vcf(d, pat, chrom):
    """Locate one genome's VCF for one chromosome.

    Returns (path, note). A concrete pattern is used verbatim. The pattern "*"
    means the release's filenames are not known in advance, which is normal for
    a directory belonging to another group; the chromosome is then matched
    through the anchored globs above. An ambiguous match returns no path and
    says so, because guessing between two candidate files is exactly the kind
    of error that would not show up in the output.
    """
    if pat != "*":
        return os.path.join(d, pat.replace("{c}", str(chrom))), ""
    hits = []
    for g in CHROM_GLOBS:
        hits += glob.glob(os.path.join(d, g.format(c=chrom) + "vcf.gz"))
    hits = sorted(set(h for h in hits if not h.endswith(".tbi")))
    if len(hits) == 1:
        return hits[0], f"discovered {os.path.basename(hits[0])}"
    if not hits:
        return "", f"nothing in {d} matched chromosome {chrom}"
    return "", ("ambiguous, refusing to guess between "
                + ", ".join(os.path.basename(h) for h in hits))

# Indexes live in our own tree, not beside the sources, which belong to other
# users (see sbatch_index_archaics.sh).
DEF_ARCH_IDX = "/n/groups/reich/DAVID/CURSOR_ENRICH/refs/archaic_index"

# Region-extracted copies we produced ourselves, which take precedence over the
# originals when present (see sbatch_extract_chagyrskaya.sh).
DEF_ARCH_LOCAL = "/n/groups/reich/DAVID/CURSOR_ENRICH/refs/archaic_local"

# Denisova 3 drives the scan: it is ALL-SITES, 52x, and fully indexed, so
# iterating its records enumerates every position we could possibly call. Any
# all-sites genome would do; the highest-coverage one wastes the fewest queries.
DRIVER = "Denisova3"


def panel_only(args):
    """The --panel-only set, with the driver added back.

    The driver is what enumerates candidate positions, so excluding it would
    not shrink the panel, it would empty the scan.
    """
    if not getattr(args, "panel_only", ""):
        return None
    ids = {t.strip() for t in args.panel_only.split(",") if t.strip()}
    unknown = ids - {e[0] for e in PANEL}
    if unknown:
        sys.exit(f"--panel-only: unknown genome id(s) {sorted(unknown)}\n"
                 f"  known: {', '.join(e[0] for e in PANEL)}")
    return ids | {DRIVER}


def open_panel(chrom, idx_dir=DEF_ARCH_IDX, only=None):
    """Open the panel for one chromosome.

    Returns (driver, others, missing) where driver and others are dicts with
    keys id/lineage/caller/tb/si/contig, and missing lists genomes that could
    not be opened. A genome is skipped rather than fatal, because losing one
    genome should degrade the panel, not abort the chromosome -- but the strict
    rule counts a skipped genome as no_data, so the loss stays visible.
    """
    pysam = need_pysam()
    opened, missing = [], []
    for gid, lineage, d, pat, caller in PANEL:
        if only and gid not in only:
            continue
        # A region-extracted copy in our own tree wins over the original. This
        # is how Chagyrskaya 8 becomes usable at all: its source VCFs are plain
        # gzip and cannot be tabix-indexed, so sbatch_extract_chagyrskaya.sh
        # writes a BGZF subset covering the catalog regions.
        local = os.path.join(DEF_ARCH_LOCAL, f"{gid}.chr{chrom}.vcf.gz")
        if os.path.exists(local):
            path, note = local, ""
        else:
            path, note = resolve_vcf(d, pat, chrom)
        if not path:
            missing.append((gid, note))
            continue
        if not os.path.exists(path):
            missing.append((gid, f"no VCF at {path}"))
            continue
        idx = os.path.join(idx_dir, f"{gid}.chr{chrom}.tbi")
        try:
            tb = (pysam.TabixFile(path, index=idx) if os.path.exists(idx)
                  else pysam.TabixFile(path))
        except (OSError, ValueError) as e:
            missing.append((gid, f"index unusable ({e}); "
                                f"expected {idx} -- run sbatch_index_archaics.sh"))
            continue
        si, _ = sample_index(path)
        c = contig_name(tb, chrom)
        if si is None or c is None:
            missing.append((gid, f"sample_index={si} contig={c}"))
            continue
        opened.append({"id": gid, "lineage": lineage, "caller": caller,
                       "tb": tb, "si": si, "contig": c, "path": path,
                       "note": note})
    driver = next((g for g in opened if g["id"] == DRIVER), None)
    others = [g for g in opened if g["id"] != DRIVER]
    return driver, others, missing


def archaic_state(g, chrom, pos, anc, der, min_dp, min_gq):
    """Classify one archaic genome at one position.

    Distinguishes no_data (position absent from an all-sites VCF, so genuinely
    unobserved) from low_qual (observed but below DP/GQ threshold). Collapsing
    those two would be the single easiest way to bias a Neandertal-versus-
    Denisovan contrast, since coverage differs systematically between genomes.
    """
    try:
        for nl in g["tb"].fetch(g["contig"], pos - 1, pos):
            nf = nl.rstrip("\n").split("\t")
            if int(nf[1]) != pos:
                continue
            nb, ndp, ngq = call_of(nf, g["si"])
            if not nb or None in nb:
                return "no_call", ndp, ngq
            if ndp < min_dp or ngq < min_gq:
                return "low_qual", ndp, ngq
            if set(nb) == {der}:
                return "derived_hom", ndp, ngq
            if set(nb) == {anc}:
                return "ancestral_hom", ndp, ngq
            return "het", ndp, ngq
    except (ValueError, OSError):
        return "no_data", 0, 0.0
    return "no_data", 0, 0.0


def call_of(row, idx):
    """(genotype bases, dp, gq) for one sample column of a VCF row."""
    if idx is None or idx >= len(row):
        return None, 0, 0.0
    fmt = row[8].split(":")
    d = dict(zip(fmt, row[idx].split(":")))
    gt = (d.get("GT") or "./.").replace("|", "/")
    dp = int(d["DP"]) if str(d.get("DP", "")).isdigit() else 0
    try:
        gq = float(d.get("GQ", "") or 0)
    except ValueError:
        gq = 0.0
    alts = [] if row[4] in (".", "") else row[4].split(",")
    alleles = [row[3]] + alts
    bases = []
    for h in gt.split("/"):
        bases.append(alleles[int(h)].upper()
                     if h.isdigit() and int(h) < len(alleles) else None)
    return bases, dp, gq


# ------------------------------------------------------- transcript machinery
def load_cds(gtf, chroms):
    """chrom -> list of (start, end, strand, tx, gene); plus tx -> exon list."""
    by_chrom = defaultdict(list)
    by_tx = defaultdict(list)
    tx_meta = {}
    op = gzip.open if gtf.endswith(".gz") else open
    with op(gtf, "rt", errors="replace") as f:
        for line in f:
            if line.startswith("#"):
                continue
            fl = line.rstrip("\n").split("\t")
            if len(fl) < 9 or fl[2] != "CDS":
                continue
            c = fl[0].replace("chr", "")
            if c not in chroms:
                continue
            s, e, strand = int(fl[3]), int(fl[4]), fl[6]
            tx = gene = ""
            for item in fl[8].split(";"):
                item = item.strip()
                if item.startswith("transcript_id "):
                    tx = item[len("transcript_id "):].strip('"')
                elif item.startswith("gene_name "):
                    gene = item[len("gene_name "):].strip('"')
            by_chrom[c].append((s, e, strand, tx, gene))
            by_tx[tx].append((s, e))
            tx_meta[tx] = (c, strand, gene)
    for c in by_chrom:
        by_chrom[c].sort()
    return by_chrom, by_tx, tx_meta


class TxSeq:
    """Coding sequence of one transcript, with a genomic->CDS index map."""

    def __init__(self, fasta, fa_contig, exons, strand):
        exons = sorted(exons)
        self.strand = strand
        self.pos2idx = {}
        pieces = []
        order = exons if strand == "+" else list(reversed(exons))
        idx = 0
        for s, e in order:
            seq = fasta.fetch(fa_contig, s - 1, e).upper()
            if strand == "-":
                seq = rc(seq)
            pieces.append(seq)
            span = range(s, e + 1) if strand == "+" else range(e, s - 1, -1)
            for p in span:
                self.pos2idx[p] = idx
                idx += 1
        self.seq = "".join(pieces)

    def consequence(self, pos, derived_base, ancestral_base):
        i = self.pos2idx.get(pos)
        if i is None:
            return None
        d = derived_base.upper()
        a = ancestral_base.upper()
        if self.strand == "-":
            d, a = rc(d), rc(a)
        start = (i // 3) * 3
        codon = self.seq[start:start + 3]
        if len(codon) < 3:
            return None
        off = i - start
        if codon[off] != d:                 # reference disagrees; report it
            return {"mismatch": f"{codon[off]}!={d}"}
        anc_codon = codon[:off] + a + codon[off + 1:]
        aa_d, aa_a = translate(codon), translate(anc_codon)
        return {
            "codon_derived": codon, "codon_ancestral": anc_codon,
            "aa_derived": aa_d, "aa_ancestral": aa_a,
            "protein_position": i // 3 + 1,
            "consequence": ("synonymous" if aa_d == aa_a else
                            "stop_gained" if aa_d == "*" else
                            "stop_lost" if aa_a == "*" else "missense"),
        }


# ------------------------------------------------------------------- selftest
def selftest(ref_fa, gtf):
    """chr9:84,605,717 in SPATA31D1 should be missense Leu->Arg at codon 111.

    The previous case asserted Ile->Met at codon 205 for chr9:84,606,000 with a
    derived allele of G. That coordinate is a rounded region boundary, not a
    site: hg19 carries C there, so the assertion could never hold, and the test
    duly failed with 'mismatch: C!=G' -- which was the consequence machinery
    working correctly, reporting that the reference contradicted the allele it
    had been handed. A test that cannot pass is worse than no test, because it
    trains you to launch through a red light.

    The real site is 84,605,717, the SPATA31D1 entry in the catalog: hg19 has G,
    the ancestral allele is T, and the derived allele is the reference, as it is
    at every site the screen returns. In ENST00000344803.2 that is CDS index
    331, the middle base of codon 111, turning the ancestral CTG into CGG and so
    leucine into arginine -- the Leu111Arg the note discusses. Verified against
    GENCODE v19 and the UCSC hg19 sequence independently of this code.
    """
    need_pysam()
    print("--- selftest: chr9:84,605,717 SPATA31D1 "
          "(expect missense L->R @111) ---")
    fasta = pysam.FastaFile(ref_fa)
    fc = contig_name(fasta, "9", is_fasta=True)
    by_chrom, by_tx, tx_meta = load_cds(gtf, {"9"})
    pos, anc, der = 84605717, "T", "G"
    got = False
    for s, e, strand, tx, gene in by_chrom["9"]:
        if s <= pos <= e and gene == "SPATA31D1":
            t = TxSeq(fasta, fc, by_tx[tx], strand)
            r = t.consequence(pos, der, anc)
            if r:
                print(f"  {tx} ({gene}, {strand}): {r}")
                got = got or (r.get("consequence") == "missense"
                              and r.get("aa_ancestral") == "L"
                              and r.get("aa_derived") == "R"
                              and r.get("protein_position") == 111)
    print("  SELFTEST PASS" if got else
          "  SELFTEST FAIL - do not launch; the codon logic or the reference "
          "does not agree with the expected Leu111Arg")
    fasta.close()
    return 0 if got else 1


def check_paths(args):
    ok = True
    print("--- input paths ---")
    for label, p in (("GRCh37.fa", args.ref), ("ancestral", args.anc),
                     ("gencode", args.gtf), ("Denisova", args.den),
                     ("Neandertal", args.nea)):
        e = os.path.exists(p)
        ok &= e
        print(f"  [{'ok' if e else 'MISSING'}] {label:<12} {p}")
        # Only the VCFs are queried by region and so actually need an index.
        # The GTF is streamed once per chromosome by load_cds(), so a missing
        # .tbi there is harmless and should not read as a problem.
        if (e and p.endswith(".gz") and not os.path.exists(p + ".tbi")
                and label in ("Denisova", "Neandertal")):
            print(f"         WARNING: no .tbi index; region queries will fail")
            ok = False
        if e and p.endswith(".fa") and not os.path.exists(p + ".fai"):
            print(f"         (no .fai index alongside)")
    # Check the index, not just existence. The first version of this reported
    # all 22 phase 3 VCFs as 'ok' while every one of their .tbi files was
    # unloadable, so the array failed 17/17 on something --check-paths had
    # already blessed.
    for c in ("1", "9", "11"):
        g = find_g1k(c)
        if not g:
            print(f"  [MISSING] 1000G chr{c:<3}  no VCF found")
            ok = False
            continue
        if g1k_index_ok(g):
            print(f"  [ok] 1000G chr{c:<3}  {g}")
        else:
            print(f"  [BAD INDEX] 1000G chr{c:<3}  {g}")
            for ext in (".tbi", ".csi"):
                if os.path.exists(g + ext):
                    print(f"         {ext} present, "
                          f"{os.path.getsize(g + ext)} bytes, but unloadable")
                    break
            else:
                print(f"         no .tbi or .csi alongside")
            print(f"         rebuild with: tabix -f -p vcf {g}")
            ok = False
    if args.extra_nea:
        for p in args.extra_nea.split(","):
            print(f"  [{'ok' if os.path.exists(p) else 'MISSING'}] extra Nea   {p}")
    print("\nsample columns:")
    for label, p in (("Denisova", args.den), ("Neandertal", args.nea)):
        if os.path.exists(p):
            i, s = sample_index(p)
            print(f"  {label:<12} column {i}, samples {s}")
    return 0 if ok else 1


# ---------------------------------------------------------------------- main
def run_chrom(args):
    chrom = args.chrom.replace("chr", "")
    regs = []
    with open(args.regions, encoding="utf-8") as f:
        for r in csv.DictReader(f, delimiter="\t"):
            if r["chrom"].replace("chr", "") == chrom:
                regs.append((int(r["start"]), int(r["end"]) + args.pad, r))
    if not regs:
        print(f"no clusters on chromosome {chrom}", file=sys.stderr)
        return 0
    print(f"chr{chrom}: {len(regs)} clusters, "
          f"{sum(e-s+1 for s, e, _ in regs)/1e6:.2f} Mb", file=sys.stderr)

    need_pysam()
    fasta = pysam.FastaFile(args.ref)
    anc_fa = pysam.FastaFile(args.anc)
    fc = contig_name(fasta, chrom, is_fasta=True)
    ac = contig_name(anc_fa, chrom, is_fasta=True)
    if fc is None or ac is None:
        # Print the candidate names: the first version of this message said only
        # "anc=None", which told us a lookup failed but not why, and cost a
        # whole 22-task array to find out.
        sys.stderr.write(f"could not resolve contig for {chrom} "
                         f"(ref={fc}, anc={ac})\n")
        for label, fh in (("ref", fasta), ("anc", anc_fa)):
            names = list(fh.references)
            sys.stderr.write(f"  {label}: {len(names)} contigs, first 5: "
                             f"{names[:5]}\n")
        return 1

    if args.legacy_pair:
        # The original 1 Denisovan + 1 Neandertal screen, kept because both
        # staged genomes are GATK-era and so give a caller-consistent result
        # to compare the mixed-caller panel against.
        den_tb = pysam.TabixFile(args.den)
        den_i, _ = sample_index(args.den, prefer="deni")
        driver = {"id": "Denisova(staged)", "lineage": "Denisovan",
                  "caller": "GATK-era", "tb": den_tb, "si": den_i,
                  "contig": contig_name(den_tb, chrom), "path": args.den}
        others = []
        for name, p in [("Altai", args.nea)] + [
                (os.path.basename(p).split(".")[0], p)
                for p in filter(None, (args.extra_nea or "").split(","))]:
            if not os.path.exists(p):
                continue
            tb = pysam.TabixFile(p)
            others.append({"id": name, "lineage": "Neandertal",
                           "caller": "GATK-era", "tb": tb,
                           "si": sample_index(p)[0],
                           "contig": contig_name(tb, chrom), "path": p})
        missing = []
    else:
        driver, others, missing = open_panel(chrom, args.archaic_index,
                                            only=panel_only(args))

    for gid, why in missing:
        sys.stderr.write(f"  PANEL WARNING {gid}: {why}\n")
    if driver is None:
        sys.stderr.write(f"chr{chrom}: cannot open driver genome {DRIVER}; "
                         f"nothing to iterate over\n")
        return 1

    panel_all = [driver] + others
    nea_ids = [g["id"] for g in panel_all if g["lineage"] == "Neandertal"]
    den_ids = [g["id"] for g in panel_all if g["lineage"] == "Denisovan"]
    sys.stderr.write(f"  panel: {len(nea_ids)} Neandertal {nea_ids} + "
                     f"{len(den_ids)} Denisovan {den_ids}\n")

    g1k_path = find_g1k(chrom)
    g1k_tb = pysam.TabixFile(g1k_path) if g1k_path else None

    dc = driver["contig"]
    den_tb, den_i = driver["tb"], driver["si"]
    gc = contig_name(g1k_tb, chrom) if g1k_tb else None

    cds_by_chrom, by_tx, tx_meta = load_cds(args.gtf, {chrom})
    txcache = {}

    rows = []
    for start, end, reg in regs:
        anc_seq = anc_fa.fetch(ac, start - 1, end)
        ref_seq = fasta.fetch(fc, start - 1, end).upper()

        af = {}
        if g1k_tb and gc:
            for ln in g1k_tb.fetch(gc, start - 1, end):
                fl = ln.split("\t")
                info = dict(kv.split("=", 1) for kv in fl[7].split(";") if "=" in kv)
                try:
                    af[int(fl[1])] = max(float(x) for x in info.get("AF", "0").split(","))
                except ValueError:
                    af[int(fl[1])] = 0.0

        # Candidate positions come from the sequences, not from any one genome's
        # records. Driving the loop off a single archaic VCF would give that
        # genome veto power over which sites are even considered, which breaks
        # an "at least one of the two has data" rule: a site where Denisova 3
        # is low-quality but Denisova 25 is cleanly ancestral must survive.
        cand_pos = []
        for k in range(end - start + 1):
            a = anc_seq[k] if k < len(anc_seq) else "N"
            if a not in "ACGT":             # uppercase == high-confidence call
                continue
            ref = ref_seq[k] if k < len(ref_seq) else "N"
            if ref not in "ACGT" or ref == a:
                continue                    # need the modern reference derived
            pos = start + k
            if af.get(pos, 0.0) > args.max_af:
                continue                    # not fixed/near-fixed in moderns
            cand_pos.append((pos, a, ref))

        n_cand = 0
        for pos, a, ref in cand_pos:
            state, dp_of, gq_of = {}, {}, {}
            for g in panel_all:
                st, dp, gq = archaic_state(g, chrom, pos, a, ref,
                                           args.min_dp, args.min_gq)
                state[g["id"]], dp_of[g["id"]], gq_of[g["id"]] = st, dp, gq

            # "Has data" means a usable genotype call. no_data (absent from an
            # all-sites VCF), no_call and low_qual are all treated as absence of
            # evidence rather than as evidence of the ancestral state -- the
            # distinction that keeps differing coverage from masquerading as a
            # lineage difference.
            def verdict(ids, want):
                called = [i for i in ids if state[i] not in
                          ("no_data", "no_call", "low_qual")]
                agree = [i for i in called if state[i] == want]
                return called, agree, bool(called) and len(agree) == len(called)

            nea_called, nea_agree, nea_ok = verdict(nea_ids, "derived_hom")
            den_called, den_agree, den_ok = verdict(den_ids, "ancestral_hom")

            # The snpAD genomes (Vindija, Chagyrskaya, Goyet, Denisova 17,
            # Denisova 25)
            # were all filtered through mappability masks -- mapab100, map35_100
            # -- so positions failing those masks are simply absent from their
            # VCFs. The GATK-era pair (Altai, Denisova 3) carries no such mask
            # and reports everywhere. A position therefore goes missing from
            # every snpAD genome at once while both GATK genomes call it, which
            # looks like independent corroboration failing but is one shared
            # filter. Flagged rather than dropped: such sites are real calls in
            # sequence the modern pipelines treat as unmappable, and that is a
            # confidence caveat the reader should see, not a deletion.
            snp_called = [g["id"] for g in panel_all if g["caller"] == "snpAD"
                          and state[g["id"]] not in
                          ("no_data", "no_call", "low_qual")]
            gatk_called = [g["id"] for g in panel_all if g["caller"] != "snpAD"
                           and state[g["id"]] not in
                           ("no_data", "no_call", "low_qual")]
            snpad_masked = "yes" if (not snp_called and gatk_called) else "no"

            # The catalog rule: at least one genome with data on each side, and
            # unanimity among those with data.
            #
            # The archaic panel and 1000 Genomes are CONFIRMATORY: they describe
            # how well each candidate is corroborated, they do not decide which
            # candidates exist. The catalog's site list is set by the ARGweaver
            # regions plus the modern fixed-derived criterion, so the unanimity
            # rule is recorded as a column and every candidate reaches the
            # output. Filtering here would have hidden the three sites where all
            # archaics are ancestral (modern-specific derived alleles, and
            # interesting in their own right) and would have made the ARGweaver
            # self-consistency check impossible to run, since the rule discards
            # precisely the sites that check inspects.
            why = []
            if not nea_called:
                why.append("no_nea_data")
            elif len(nea_agree) != len(nea_called):
                why.append("nea_disagree:" + ",".join(
                    f"{i}={state[i]}" for i in nea_called
                    if state[i] != "derived_hom"))
            if not den_called:
                why.append("no_den_data")
            elif len(den_agree) != len(den_called):
                why.append("den_disagree:" + ",".join(
                    f"{i}={state[i]}" for i in den_called
                    if state[i] != "ancestral_hom"))
            passes = "yes" if (nea_ok and den_ok) else "no"
            if args.require_pass and passes == "no":
                continue

            cons = {}
            gene = ""
            for s, e, strand, tx, gname in cds_by_chrom.get(chrom, []):
                if s <= pos <= e:
                    if tx not in txcache:
                        txcache[tx] = TxSeq(fasta, fc, by_tx[tx], strand)
                    r = txcache[tx].consequence(pos, ref, a)
                    if r and "mismatch" not in r:
                        gene = gname
                        if not cons or r["consequence"] == "missense":
                            cons = dict(r, transcript=tx, gene=gname)
            n_cand += 1
            rows.append({
                "chrom": chrom, "pos": pos, "ancestral": a, "derived": ref,
                "g1k_anc_af": af.get(pos, 0.0),
                "g1k_polymorphic": "yes" if pos in af else "no",
                # Per-genome state, depth and quality for all six genomes. This
                # is what makes the calling rule revisable: any alternative rule
                # ("every Neandertal called", "Denisova 3 only", a different
                # GQ cut) is a filter over these columns, so it never costs
                # another pass over ~280 GB of archaic VCFs.
                **{f"{g['id']}_state": state[g["id"]] for g in panel_all},
                **{f"{g['id']}_dp": dp_of[g["id"]] for g in panel_all},
                **{f"{g['id']}_gq": round(gq_of[g["id"]], 1) for g in panel_all},
                "nea_called": len(nea_called), "nea_derived": len(nea_agree),
                "den_called": len(den_called), "den_ancestral": len(den_agree),
                # Sites in sequence the snpAD mappability masks exclude: every
                # snpAD genome silent, GATK-era genomes calling. Roughly a
                # quarter of the catalog, and they rest on Altai alone.
                "snpad_masked": snpad_masked,
                "snpad_called": len(snp_called), "gatk_called": len(gatk_called),
                # Summary of the corroboration, as a column rather than a filter.
                "archaic_pattern": passes, "pattern_detail": ";".join(why) or "pass",
                # Caller mix is recorded per site so any GQ-threshold artefact
                # can be tested for, rather than assumed absent: GQ is not on a
                # common scale between GATK-era and snpAD genomes.
                "callers_nea": ",".join(sorted({g["caller"] for g in panel_all
                                                if g["lineage"] == "Neandertal"
                                                and g["id"] in nea_called})),
                "callers_den": ",".join(sorted({g["caller"] for g in panel_all
                                                if g["lineage"] == "Denisovan"
                                                and g["id"] in den_called})),
                "context": "CDS" if cons else "non_coding",
                "gene": cons.get("gene", gene),
                "transcript": cons.get("transcript", ""),
                "consequence": cons.get("consequence", ""),
                "aa_derived": THREE.get(cons.get("aa_derived", ""), ""),
                "aa_ancestral": THREE.get(cons.get("aa_ancestral", ""), ""),
                "protein_position": cons.get("protein_position", ""),
                "codon_derived": cons.get("codon_derived", ""),
                "codon_ancestral": cons.get("codon_ancestral", ""),
                "cluster_start": start, "cluster_end": end,
            })
        print(f"  {chrom}:{start}-{end} cand={n_cand}", file=sys.stderr)

    os.makedirs(os.path.dirname(args.out_prefix) or ".", exist_ok=True)
    out = f"{args.out_prefix}.chr{chrom}.tsv"
    if rows:
        with open(out, "w", newline="", encoding="utf-8") as f:
            w = csv.DictWriter(f, fieldnames=list(rows[0].keys()), delimiter="\t")
            w.writeheader()
            w.writerows(rows)
    else:
        open(out, "w").close()
    print(f"chr{chrom}: {len(rows)} sites -> {out}", file=sys.stderr)

    return 0


def collate(prefix):
    files = sorted(glob.glob(f"{prefix}.chr*.tsv"))
    allrows, hdr = [], None
    for p in files:
        with open(p, encoding="utf-8") as f:
            rd = csv.DictReader(f, delimiter="\t")
            if rd.fieldnames:
                hdr = hdr or rd.fieldnames
                allrows.extend(rd)
    out = f"{prefix}.all.tsv"
    if hdr:
        with open(out, "w", newline="", encoding="utf-8") as f:
            w = csv.DictWriter(f, fieldnames=hdr, delimiter="\t")
            w.writeheader()
            w.writerows(allrows)
    print(f"{len(files)} chromosome files -> {len(allrows)} sites -> {out}")
    cons = defaultdict(int)
    for r in allrows:
        cons[r.get("consequence") or r.get("context") or "?"] += 1
    print("  by consequence:", dict(cons))
    mis = [r for r in allrows if r.get("consequence") == "missense"]
    if mis:
        print(f"\n  {len(mis)} missense:")
        for r in mis:
            print(f"    {r['gene']:<12} chr{r['chrom']}:{r['pos']:>11} "
                  f"{r['aa_ancestral']}->{r['aa_derived']} @{r['protein_position']} "
                  f"({r['transcript']})")
    return 0


def main() -> int:
    ap = argparse.ArgumentParser()
    ap.add_argument("--chrom", default="")
    ap.add_argument("--regions", default="preal50.regions.tsv")
    ap.add_argument("--ref", default=DEF_REF)
    ap.add_argument("--anc", default=DEF_ANC)
    ap.add_argument("--gtf", default=DEF_GTF)
    ap.add_argument("--den", default=DEF_DEN)
    ap.add_argument("--nea", default=DEF_NEA)
    ap.add_argument("--extra-nea", default="",
                    help="comma-separated extra Neandertal VCFs, e.g. Vindija")
    ap.add_argument("--neandertal", choices=("both", "any", "none"), default="both",
                    help="only used with --legacy-pair")
    ap.add_argument("--legacy-pair", action="store_true",
                    help="run the original 1 Denisovan + 1 Neandertal screen off "
                         "the staged merged VCFs instead of the 6-genome panel. "
                         "Both staged genomes are GATK-era, so this gives a "
                         "caller-consistent comparison for the mixed panel.")
    ap.add_argument("--audit", action="store_true",
                    help="accepted for backward compatibility and ignored: the "
                         "main output now always contains every candidate.")
    ap.add_argument("--require-pass", action="store_true",
                    help="emit only sites where every called Neandertal is "
                         "derived and every called Denisovan ancestral. Off by "
                         "default: the archaic panel is confirmatory, so the "
                         "pattern is reported per site in archaic_pattern "
                         "rather than deciding membership.")
    ap.add_argument("--archaic-index", default=DEF_ARCH_IDX,
                    help="directory of tabix indexes for the panel, built by "
                         "sbatch_index_archaics.sh (indexes are kept out of the "
                         "source directories, which belong to other users)")
    ap.add_argument("--max-af", type=float, default=0.01)
    ap.add_argument("--min-dp", type=int, default=10)
    ap.add_argument("--min-gq", type=float, default=30.0)
    ap.add_argument("--pad", type=int, default=5000)
    ap.add_argument("--check-paths", action="store_true")
    ap.add_argument("--panel-only", default="",
                    help="comma-separated genome ids to restrict the archaic "
                         "panel to; the driver is always kept. Use to drop a "
                         "genome without editing PANEL, e.g. to run the four "
                         "classic high-coverage Neandertals only")
    ap.add_argument("--panel-check", default="",
                    help="comma-separated chromosomes; open the archaic panel "
                         "on each and report which genomes resolved, with the "
                         "filename discovered for globbed entries, then exit")
    ap.add_argument("--selftest", action="store_true")
    ap.add_argument("--collate", default="")
    ap.add_argument("-o", "--out-prefix", default="out/fixed_derived")
    args = ap.parse_args()

    rc_ = 0
    if args.panel_check:
        bad = 0
        for c in args.panel_check.split(","):
            c = c.strip()
            driver, others, missing = open_panel(c, args.archaic_index,
                                                 only=panel_only(args))
            got = ([driver] if driver else []) + others
            nea = [g["id"] for g in got if g["lineage"] == "Neandertal"]
            den = [g["id"] for g in got if g["lineage"] == "Denisovan"]
            print(f"--- chr{c}: {len(nea)} Neandertal + {len(den)} Denisovan")
            for g in got:
                extra = f"  [{g['note']}]" if g.get("note") else ""
                print(f"  [ok]   {g['id']:<13} {g['lineage']:<10} "
                      f"{g['caller']:<9} {os.path.basename(g['path'])}{extra}")
            for gid, why in missing:
                print(f"  [MISS] {gid:<13} {why}")
                bad += 1
        print(f"\nNeandertal genomes expected: "
              f"{sum(1 for e in PANEL if e[1] == 'Neandertal')}")
        return 1 if bad else 0
    if args.check_paths:
        rc_ |= check_paths(args)
    if args.selftest:
        rc_ |= selftest(args.ref, args.gtf)
    if args.collate:
        return collate(args.collate)
    if args.check_paths or args.selftest:
        return rc_
    if not args.chrom:
        ap.error("give --chrom, or --check-paths/--selftest/--collate")
    return run_chrom(args)


if __name__ == "__main__":
    raise SystemExit(main())
```
