## Supplementary Information section 4 code base for "Hypothesis: A modern human range expansion ∼300,000 years ago had a critical role in Neandertal origins": optimize_relate_rule.py

```
#!/usr/bin/env python3
"""Case/control optimisation of a sweep rule on the Relate archaic scan.

The ARGweaver rule is three-parameter: a Neandertal threshold x, a Denisovan
floor y, and a percentile cut through the posterior. The released Relate trees
carry point-estimate branch lengths and no posterior, so the percentile
dimension does not exist here and the rule collapses to (x, y) applied to point
estimates. That is a smaller search, not a weaker one -- the percentile cut had
already collapsed to the median in the ARGweaver rule, and the Denisovan floor
was retained for interpretability rather than because tuning it helped.

Case and control are mirror images, exactly as in the ARGweaver scan:

    case = (root(moderns + Neandertals) <= x) and (root(moderns + Den)  > y)
    ctrl = (root(moderns + Den)         <= x) and (root(moderns + Nea)  > y)

P(real) = 1 - omega0 * ctrl/case, where omega0 is the ratio expected when no
sweep signal is present. omega0 is measured here rather than assumed to be 1,
for two reasons. The Denisovan is a slightly more diverged lineage, which puts
the control root marginally deeper than the case root even under the null (in
the ARGweaver scan the two medians differ by 0.3%, giving omega0 = 1.011). And
this scan pools three Neandertals against one Denisovan, so the case side must
clear six archaic haplotypes where the control clears two; root times saturate
with sample size, so that asymmetry is expected to be small, but measuring
omega0 absorbs whatever it is as long as it does not vary with the threshold.
Whether it varies is checked, not assumed: --shell-report prints the marginal
ratio across the high-threshold shell, which should be flat.

Selection uses odd chromosomes and reports on even ones. Optimising the ratio
and then quoting the ratio at the chosen operating point is winner's curse; the
held-out half is the only honest number, and the Wilson upper bound is used as
the objective so that a rule surviving on six grid points cannot win on noise.
"""

from __future__ import annotations

import argparse
import collections
import csv
import glob
import os
import sys

sys.path.insert(0, os.path.dirname(os.path.abspath(__file__)))
from optimize_rule_by_size import wilson_hi, wilson_lo  # noqa: E402


NEA_COL = {"nea3": "02_mod_nea", "nea2": "02b_mod_nea2",
           "nea1": "02c_mod_nea1"}


Rec = collections.namedtuple("Rec", "chrom pos case ctrl nea_root den_root")


def load_grids(paths, panel, years_per_gen, nea="nea2"):
    """One Rec per grid point, autosomes only, both union roots called.

    Grid points where either root is missing are dropped rather than imputed. A
    missing root means the subset did not coalesce inside that tree's interval,
    which is not the same as a deep one, and treating it as deep would push it
    into the control tail and quietly inflate the denominator.

    The two archaic-internal roots are carried alongside because they set the
    structural floors on the case and control statistics; see eligible().
    """
    nea_col = f"{panel}_{NEA_COL[nea]}_tmrca_gen"
    den_col = f"{panel}_03_mod_den_tmrca_gen"
    out, skipped, missing = [], 0, 0
    for p in sorted(paths):
        with open(p, encoding="utf-8") as f:
            rd = csv.DictReader(f, delimiter="\t")
            if nea_col not in rd.fieldnames or den_col not in rd.fieldnames:
                sys.exit(f"{p}: no columns {nea_col} / {den_col}\n"
                         f"  available: {', '.join(rd.fieldnames)}")
            for r in rd:
                c = r["chrom"].replace("chr", "")
                if c in ("X", "Y", "23", "24"):
                    skipped += 1
                    continue
                a, b = r[nea_col], r[den_col]
                if a in ("NA", "") or b in ("NA", ""):
                    missing += 1
                    continue
                k = years_per_gen / 1000.0

                def root(col):
                    v = r.get(col, "NA")
                    return float(v) * k if v not in ("NA", "") else float("nan")
                out.append(Rec(int(c), int(r["pos"]),
                               float(a) * k, float(b) * k,
                               root("arch_04_nea_only_tmrca_gen"),
                               root("arch_05_den_only_tmrca_gen")))
    sys.stderr.write(f"loaded {len(out):,} autosomal grid points from "
                     f"{len(paths)} file(s); {missing:,} uncalled, "
                     f"{skipped:,} non-autosomal\n")
    return out


def coincidence_report(recs):
    """How often the case and control statistics are the same number.

    This is the ceiling on anything a Relate-only rule can do, and it is worth
    printing before any threshold is scanned. The case statistic is the root of
    moderns plus Neandertals and the control that of moderns plus the
    Denisovan, so both are bounded below by the modern-only root. With a large
    modern panel that root is itself older than the modern-archaic split, the
    archaic tips attach beneath it, and neither statistic moves: the two become
    equal and the contrast the catalog is built on carries no information. The
    ARGweaver-D scan, whose modern panel is 16 haplotypes rather than 216, has
    them equal at 49% of grid points for comparison.
    """
    n = len(recs)
    same = sum(1 for r in recs if r.case == r.ctrl)
    d = sorted(r.ctrl - r.case for r in recs)
    sys.stderr.write(
        f"case and control statistics are exactly equal at {same:,} of "
        f"{n:,} grid points ({100.0 * same / n:.0f}%); "
        f"control minus case spans {d[n // 20]:.0f} ka at the 5th percentile "
        f"to {d[min(n - 1, 19 * n // 20)]:.0f} ka at the 95th\n")
    return same / n


def eligible(recs, cap):
    """Drop grid points where either archaic panel's own root exceeds cap.

    WHY THIS IS NEEDED, AND WHY IT IS NOT AD HOC
    The case statistic is the root of moderns plus the Neandertals, so it can
    never fall below the Neandertals' own root; the control statistic is the
    root of moderns plus Denisova, floored likewise by the two Denisovan
    haplotypes' root. Those two floors are not equal. With three Neandertals
    pooled the Neandertal-internal root has a median of 313 ka against 188 ka
    for Denisova, and at a 700 ka threshold that difference blocks 14.0% of
    grid points from ever being counted as cases against 2.8% for controls.
    The ratio then measures which side is structurally free to fire rather
    than which side coalesces recently, which is what drove P(real) negative
    in the shallow tail on the first pass.

    Requiring both internal roots at or below the lowest threshold in the scan
    removes the asymmetry using columns already in the grid, so it is an
    alternative to regenerating the grid with fewer Neandertals. Within the
    surviving set both conditions can fire at every threshold considered, so
    the comparison is symmetric by construction.

    It does not discard sweep candidates. A sweep shared by moderns and
    Neandertals requires the Neandertal lineages to sit in a recent clade with
    moderns, which forces their internal root to be recent too. What is
    removed is the regions where the archaic panel could not have shown the
    signal either way.
    """
    keep = [r for r in recs
            if not (r.nea_root != r.nea_root or r.den_root != r.den_root)
            and r.nea_root <= cap and r.den_root <= cap]
    nb = sum(1 for r in recs if r.nea_root == r.nea_root and r.nea_root > cap)
    db = sum(1 for r in recs if r.den_root == r.den_root and r.den_root > cap)
    sys.stderr.write(
        f"symmetric-eligibility cap {cap:.0f} ka: kept {len(keep):,} of "
        f"{len(recs):,} ({100 * len(keep) / max(1, len(recs)):.1f}%); "
        f"case side was blocked at {100 * nb / max(1, len(recs)):.1f}% and "
        f"control side at {100 * db / max(1, len(recs)):.1f}% before it\n")
    return keep


def counts(recs, x, y):
    case = ctrl = 0
    for r in recs:
        if r.case <= x and r.ctrl > y:
            case += 1
        if r.ctrl <= x and r.case > y:
            ctrl += 1
    return case, ctrl


def floor_for(x, y, tie, gap):
    """The Denisovan floor to pair with threshold x.

    Two regimes. With an absolute floor the rule is (nea <= x, den > y), which
    is what the ARGweaver catalog uses: x = 625 ka, y = 700 ka. With --tie-y the
    floor tracks the threshold, (nea <= x, den > x + gap), which keeps the case
    and control conditions mutually exclusive and mirror-symmetric at every x.

    The distinction is not cosmetic. If the floor does not exclude the control
    statistic, both conditions can fire at the same grid point, and the counts
    stop measuring the contrast between the two roots and start measuring their
    marginal distributions instead -- which are nearly identical here, since
    both are dominated by the same modern-archaic split, so the ratio collapses
    toward 1 and no discrimination is visible whether or not any exists.
    """
    return (x + gap) if tie else y


def shell_report(recs, xs, y, tie=False, gap=0.0):
    """Marginal ctrl/case between consecutive thresholds.

    The cumulative ratio is contaminated by real signal at every threshold that
    admits any. The marginal ratio over a band that admits only noise estimates
    omega0 directly, and its flatness across the high shell is the assumption
    the whole normalisation rests on.
    """
    rows, prev = [], None
    for x in xs:
        case, ctrl = counts(recs, x, floor_for(x, y, tie, gap))
        marg = float("nan")
        if prev is not None:
            dc, dk = case - prev[0], ctrl - prev[1]
            marg = dk / dc if dc > 0 else float("nan")
        rows.append((x, case, ctrl, marg))
        prev = (case, ctrl)
    return rows


def estimate_omega0(recs, xs, y, shell_lo, tie=False, gap=0.0):
    """Pooled marginal ratio over thresholds at or above shell_lo."""
    prev, dc_tot, dk_tot = None, 0, 0
    for x in xs:
        case, ctrl = counts(recs, x, floor_for(x, y, tie, gap))
        if prev is not None and x > shell_lo:
            dc_tot += case - prev[0]
            dk_tot += ctrl - prev[1]
        prev = (case, ctrl)
    if dc_tot <= 0:
        sys.exit(f"no grid points admitted above the shell floor {shell_lo} ka")
    return dk_tot / dc_tot, dc_tot, dk_tot


def main():
    ap = argparse.ArgumentParser()
    ap.add_argument("--grids", default="relate_archaic/chr*.tsv",
                    help="glob of per-chromosome grid TSVs")
    ap.add_argument("--panel", default="yri", choices=("yri", "ygc"),
                    help="yri = YRI only; ygc = YRI+GBR+CHB")
    ap.add_argument("--nea", default="nea2", choices=tuple(NEA_COL),
                    help="which Neandertal set forms the case side. nea2 "
                         "(Altai+Vindija) matches ARGweaver; nea1 (Altai) is "
                         "the exactly symmetric 2-vs-2 contrast; nea3 pools "
                         "all three and breaks the shallow tail")
    ap.add_argument("--years-per-gen", type=float, default=28.0)
    ap.add_argument("--x-grid", default="500:2600:50",
                    help="lo:hi:step in ka for the Neandertal threshold")
    ap.add_argument("--y-grid", default="",
                    help="lo:hi:step in ka for the Denisovan floor; "
                         "default ties y to a single value via --y")
    ap.add_argument("--y", type=float, default=0.0,
                    help="fixed Denisovan floor in ka (0 = does not bind)")
    ap.add_argument("--shell-lo", type=float, default=0.0,
                    help="threshold above which the scan is treated as "
                         "signal-free for omega0; 0 = pick the top third")
    ap.add_argument("--tie-y", action="store_true",
                    help="derive the Denisovan floor as x + --gap instead of "
                         "using a fixed --y, keeping case and control mutually "
                         "exclusive at every threshold")
    ap.add_argument("--gap", type=float, default=0.0,
                    help="with --tie-y, the ka by which the floor exceeds the "
                         "threshold (ARGweaver's catalog uses 625 -> 700, "
                         "i.e. a gap of 75)")
    ap.add_argument("--sym-cap", type=float, default=-1.0,
                    help="restrict to grid points where BOTH archaic-internal "
                         "roots are at or below this ka value, so neither the "
                         "case nor the control condition is structurally "
                         "blocked at any threshold in the scan. -1 uses the "
                         "low end of --x-grid; 0 disables the restriction")
    ap.add_argument("--catalog", default="",
                    help="write the selected rule's case grid points here")
    ap.add_argument("--shell-report", action="store_true")
    ap.add_argument("--omega0", type=float, default=0.0,
                    help="use this null ratio instead of measuring one; set "
                         "when omega0 comes from a separate estimate, or to "
                         "reproduce a published operating point exactly")
    ap.add_argument("--target", type=int, default=0,
                    help="select the rule whose held-out case count is nearest "
                         "this size, for comparison at matched catalog size")
    ap.add_argument("-o", "--out", default="")
    args = ap.parse_args()

    paths = sorted(glob.glob(args.grids))
    if not paths:
        sys.exit(f"no grid files matched {args.grids}\n"
                 "  (the archaic run writes relate_archaic/tsv/chrN.tsv on O2; "
                 "copy them down first)")
    recs = load_grids(paths, args.panel, args.years_per_gen, args.nea)
    if not recs:
        sys.exit("no usable grid points")
    coincidence_report(recs)

    lo, hi, step = (float(v) for v in args.x_grid.split(":"))
    cap = lo if args.sym_cap < 0 else args.sym_cap
    if cap:
        recs = eligible(recs, cap)
        if not recs:
            sys.exit("the eligibility cap removed every grid point")
    xs = [lo + i * step for i in range(int((hi - lo) / step) + 1)]
    ys = [args.y]
    if args.y_grid:
        ylo, yhi, ystep = (float(v) for v in args.y_grid.split(":"))
        ys = [ylo + i * ystep for i in range(int((yhi - ylo) / ystep) + 1)]

    if args.omega0:
        omega0 = args.omega0
        sys.stderr.write(f"omega0 = {omega0:.4f} (supplied, not measured)\n")
    else:
        shell_lo = args.shell_lo or (lo + 2.0 * (hi - lo) / 3.0)
        omega0, dc, dk = estimate_omega0(recs, xs, ys[0], shell_lo,
                                         args.tie_y, args.gap)
        sys.stderr.write(f"omega0 = {omega0:.4f} from the shell above "
                         f"{shell_lo:.0f} ka ({dk:,} control / {dc:,} case "
                         f"marginal admissions)\n")

    if args.shell_report:
        print(f"{'x (ka)':>8} {'case':>8} {'ctrl':>8} "
              f"{'cum ratio':>10} {'marginal':>10}")
        for x, case, ctrl, marg in shell_report(recs, xs, ys[0],
                                                args.tie_y, args.gap):
            cr = ctrl / case if case else float("nan")
            print(f"{x:>8.0f} {case:>8,} {ctrl:>8,} {cr:>10.3f} {marg:>10.3f}")
        return 0

    odd = [r for r in recs if r[0] % 2 == 1]
    even = [r for r in recs if r[0] % 2 == 0]
    sys.stderr.write(f"odd (select): {len(odd):,};  even (report): "
                     f"{len(even):,} grid points\n")

    rows = []
    for y in ys:
        for x in xs:
            yy = floor_for(x, y, args.tie_y, args.gap)
            co, ko = counts(odd, x, yy)
            if co < 10:
                continue
            ce, ke = counts(even, x, yy)
            rows.append({
                "x_ka": x, "y_ka": yy,
                "case_odd": co, "ctrl_odd": ko,
                "preal_odd": max(0.0, 1.0 - omega0 * ko / co),
                "case_even": ce, "ctrl_even": ke,
                "preal_even": (max(0.0, 1.0 - omega0 * ke / ce)
                               if ce else float("nan")),
                "preal_even_lo": (max(0.0, 1.0 - omega0 * wilson_hi(ke, ce))
                                  if ce else float("nan")),
                "preal_even_hi": (max(0.0, 1.0 - omega0 * wilson_lo(ke, ce))
                                  if ce else float("nan")),
                "sel_objective": wilson_hi(ko, co),
            })
    if not rows:
        sys.exit("no rule admitted at least 10 case grid points on odd "
                 "chromosomes; widen --x-grid")

    best = min(rows, key=lambda r: (r["sel_objective"], -r["case_odd"]))
    print(f"\nselected on odd chromosomes (Wilson upper bound on ctrl/case):")
    print(f"  x = {best['x_ka']:.0f} ka, y = {best['y_ka']:.0f} ka")
    print(f"  odd : {best['case_odd']:,} case, {best['ctrl_odd']:,} ctrl, "
          f"P(real) = {best['preal_odd']:.3f}")
    print(f"  even: {best['case_even']:,} case, {best['ctrl_even']:,} ctrl, "
          f"P(real) = {best['preal_even']:.3f} "
          f"[{best['preal_even_lo']:.3f}, {best['preal_even_hi']:.3f}]")
    print("  the even-chromosome value is the one to quote; the odd one is "
          "the value that was optimised and is biased upward")

    if args.target:
        near = min(rows, key=lambda r: abs(r["case_odd"] + r["case_even"]
                                           - args.target))
        print(f"\nat matched catalog size ~{args.target:,} grid points:")
        print(f"  x = {near['x_ka']:.0f} ka, y = {near['y_ka']:.0f} ka, "
              f"total case = {near['case_odd'] + near['case_even']:,}, "
              f"held-out P(real) = {near['preal_even']:.3f}")

    print(f"\n{'x':>7} {'y':>6} {'case':>8} {'ctrl':>8} {'P(real) odd':>12} "
          f"{'P(real) even':>13}")
    for r in rows:
        tot = r["case_odd"] + r["case_even"]
        if tot < 20:
            continue
        print(f"{r['x_ka']:>7.0f} {r['y_ka']:>6.0f} {tot:>8,} "
              f"{r['ctrl_odd'] + r['ctrl_even']:>8,} "
              f"{r['preal_odd']:>12.3f} {r['preal_even']:>13.3f}")

    if args.catalog:
        yy = best["y_ka"]
        hits = [r for r in recs if r.case <= best["x_ka"] and r.ctrl > yy]
        with open(args.catalog, "w", newline="", encoding="utf-8") as f:
            w = csv.writer(f, delimiter="\t")
            w.writerow(["chrom", "pos", "case_ka", "ctrl_ka",
                        "nea_root_ka", "den_root_ka"])
            for r in sorted(hits, key=lambda r: (r.chrom, r.pos)):
                w.writerow([r.chrom, r.pos, f"{r.case:.1f}", f"{r.ctrl:.1f}",
                            f"{r.nea_root:.1f}", f"{r.den_root:.1f}"])
        sys.stderr.write(f"wrote {args.catalog} ({len(hits):,} grid points "
                         f"at x={best['x_ka']:.0f}, y={yy:.0f})\n")

    if args.out:
        with open(args.out, "w", newline="", encoding="utf-8") as f:
            w = csv.DictWriter(f, delimiter="\t", fieldnames=list(rows[0]))
            w.writeheader()
            w.writerows(rows)
        sys.stderr.write(f"wrote {args.out}\n")
    return 0


if __name__ == "__main__":
    sys.exit(main())
```
