## Supplementary Information section 4 code base for "Hypothesis: A modern human range expansion ∼300,000 years ago had a critical role in Neandertal origins": optimize_rule_by_size.py

```
#!/usr/bin/env python3
"""For each target catalog size, find the best (Nea cut, Den cut, percentile cut).

THE SEARCH
The rule family is the same box rule as the catalog, but with the two archaic
thresholds decoupled:

    case     nea_hi(p) <= x   and   den_lo(p) >  y
    control  den_hi(p) <= x   and   nea_lo(p) >  y     (archaics swapped)

A full grid over (x, y, p) is scanned. For each target catalog size, the rule
minimising the Denisovan/Neandertal ratio is reported.

WHY SELECTION AND REPORTING ARE SPLIT
The grid holds a few thousand rules while control counts at the small-catalog
end are single digits, so the minimum ratio over the grid is biased downward --
some rule will reach zero controls by luck alone. Rules are therefore SELECTED
on odd chromosomes and REPORTED on even ones. The gap between the two columns
is the size of the winner's curse, and it is the honest measure of whether the
optimisation found signal or noise.

WHY THE OBJECTIVE IS A WILSON UPPER BOUND
At a target of 50 windows many rules tie at zero controls, and the raw ratio
cannot separate them. Selecting on the 95% Wilson upper bound instead penalises
a low ratio resting on a tiny catalog, which is precisely the pathology that
naive minimisation would chase.

BASELINE FOR COMPARISON
The same procedure is run over the constrained two-parameter family (y tied to
x, cut fixed at 80/20). If the three-way search does not beat it out of sample,
the extra freedom is not doing any work -- consistent with the Denisovan
threshold being non-binding.

Usage:
  python3 scripts/optimize_rule_by_size.py
"""

from __future__ import annotations

import argparse
import math
import os
import sys

import numpy as np

sys.path.insert(0, os.path.dirname(os.path.abspath(__file__)))
from approx_percentiles import default_coord, fit_site, load, z_of  # noqa: E402
from fit_fdr_surface import PCUTS  # noqa: E402

# Genome-wide targets; each half-genome carries about half of them.
TARGETS = [50, 75, 100, 150, 200, 300, 500, 750, 1000, 1500, 2000, 3000]

# Null skew. The two archaics are sister lineages and present-day Africans
# should be equidistant from both, so a signal-free genome would sit at an odds
# of 1; it does not sit exactly there, and the excess must be divided out
# before a ratio can be read as a false-discovery proportion:
# P(real) = 1 - ratio/NULL_RATIO, not 1 - ratio.
#
# This was previously 1.150 (theta0 = 0.5349), from 2708 Neandertal-direction
# against 2355 Denisovan-direction windows "inside the fitted support". That
# figure was wrong twice over, and both errors inflated it:
#
#   1. The support is the surface's whole binned domain, nea_hi in [350, 900]
#      and den_lo in [350, 1500]. It contains all 329 catalog windows and 1,126
#      windows with nea_hi <= 700 ka -- that is, it contains the signal. A
#      quantity measured there cannot be a null, and normalising by it
#      subtracts the signal from itself.
#   2. It was measured at percentile cut p = 0.80 while the catalog rule uses
#      p = 0.50. The same construction at p = 0.50 gives 1.053.
#
# The null is now measured where sweep signal is not possible: in shells
# strictly above the published 825-694 ka modern-archaic split, so that a
# join-side coalescence carries no sweep evidence, using the catalog's own cut
# (p = 0.50) and its own Denisovan threshold (y = 700 ka). A cumulative u <= x
# test cannot isolate those windows because it always admits everything below
# x, hence shells. Over 825-1100 ka: 25,260 Neandertal-direction against
# 24,977 Denisovan-direction, odds 1.0113 (95% CI 0.994-1.029), from 50,237
# windows. Stable across neighbouring shells (1.011-1.020 at p = 0.50) and flat
# across B-statistic deciles in both Murphy maps (Cochran-Armitage z = +1.20,
# p = 0.23 for phastCons; z = +0.84, p = 0.40 for CADD), so a single global
# value suffices. See null_ratio_audit.py and null_ratio_shell_bstat.py.
#
# The structural asymmetry is therefore real but worth about 1%, not 15%. Note
# that it is indistinguishable from the exact-symmetry value of 1: using 1
# instead moves the catalog's P(real) from 0.800 to 0.802.
#
# Re-measured on the 1 kb scan, which is the grid the catalog now uses: the
# same 825-1100 ka shell at p = 0.50 and y = 700 ka gives 128,198 against
# 126,611, odds 1.0125 (95% CI 1.005-1.020), from 254,809 grid points. The 5 kb
# figure of 1.0113 lies inside that interval, so the denser grid confirms the
# value rather than changing it; the extra points mainly tighten the CI.
OMEGA0 = 1.0125
THETA0 = OMEGA0 / (1 + OMEGA0)
NULL_RATIO = 1.0 / OMEGA0


def p_real(ratio):
    """P(real) implied by an observed Denisovan/Neandertal ratio."""
    return max(0.0, 1.0 - ratio / NULL_RATIO)
X_GRID = list(range(500, 926, 25))
Y_GRID = list(range(400, 1501, 50))
BAND_LO, BAND_HI = 0.75, 1.34


def wilson_hi(k, n, z=1.96):
    """Upper confidence limit on the ratio k/n, via the proportion k/(k+n).

    The ratio is an odds and can exceed 1, so it is not itself a binomial
    proportion; the interval is built on the proportion the counts really form
    and then mapped through q/(1-q).
    """
    if n == 0:
        return float("inf")
    tot = k + n
    q = k / tot
    d = 1 + z * z / tot
    c = (q + z * z / (2 * tot)) / d
    h = z * math.sqrt(q * (1 - q) / tot + z * z / (4 * tot * tot)) / d
    qhi = min(1.0 - 1e-12, c + h)
    return qhi / (1 - qhi)


def wilson_lo(k, n, z=1.96):
    """Lower confidence limit on the ratio k/n, matching wilson_hi."""
    if n == 0:
        return float("nan")
    tot = k + n
    q = k / tot
    d = 1 + z * z / tot
    c = (q + z * z / (2 * tot)) / d
    h = z * math.sqrt(q * (1 - q) / tot + z * z / (4 * tot * tot)) / d
    qlo = max(0.0, c - h)
    return qlo / (1 - qlo)


def tails(recs, coord):
    """Per-percentile arrays of the four bounds the rule needs."""
    out = {}
    mu_s, sg_s = {}, {}
    for tag in ("nea", "den"):
        ms, ss = [], []
        for d in recs:
            m, s = fit_site(coord, d[f"{tag}_p025"], d[f"{tag}_p975"])
            ms.append(m); ss.append(s)
        mu_s[tag] = np.array(ms); sg_s[tag] = np.array(ss)
    for p in PCUTS:
        zhi, zlo = z_of(p), z_of(1 - p)
        for tag in ("nea", "den"):
            out[(tag, "hi", p)] = np.array(
                [coord.to_t(v) for v in mu_s[tag] + zhi * sg_s[tag]])
            out[(tag, "lo", p)] = np.array(
                [coord.to_t(v) for v in mu_s[tag] + zlo * sg_s[tag]])
    return out


def scan(T, x_grid, y_grid, pcuts, tie_y_to_x=False):
    """Yield (x, y, p, case, ctrl) over the grid, vectorised per rule."""
    for p in pcuts:
        nh, dl = T[("nea", "hi", p)], T[("den", "lo", p)]
        dh, nl = T[("den", "hi", p)], T[("nea", "lo", p)]
        for x in x_grid:
            nh_ok, dh_ok = nh <= x, dh <= x
            ys = [x] if tie_y_to_x else y_grid
            for y in ys:
                case = int(np.count_nonzero(nh_ok & (dl > y)))
                ctrl = int(np.count_nonzero(dh_ok & (nl > y)))
                yield x, y, p, case, ctrl


def select(rows, target):
    """Best rule whose case count lands in a band around the target."""
    best = None
    lo, hi = BAND_LO * target, BAND_HI * target
    for x, y, p, case, ctrl in rows:
        if not (lo <= case <= hi):
            continue
        key = (wilson_hi(ctrl, case), ctrl / case, -case)
        if best is None or key < best[0]:
            best = (key, (x, y, p, case, ctrl))
    return best[1] if best else None


def main():
    ap = argparse.ArgumentParser()
    ap.add_argument("--scan", default="hubisz_tmrca_5kb_8afr.tsv")
    ap.add_argument("-o", "--out", default="catalog_annotation/rule_by_size.tsv")
    args = ap.parse_args()

    recs = load(args.scan)
    coord, grid_desc = default_coord(recs)
    print(f"time coordinate: {grid_desc}")
    recs = [d for d in recs if d["chrom"].replace("chr", "") not in ("X", "23")]
    odd = [d for d in recs if int(d["chrom"].replace("chr", "")) % 2 == 1]
    even = [d for d in recs if int(d["chrom"].replace("chr", "")) % 2 == 0]
    print(f"odd (select): {len(odd):,} windows;  even (report): {len(even):,}")

    To, Te = tails(odd, coord), tails(even, coord)
    rows_o = list(scan(To, X_GRID, Y_GRID, PCUTS))
    # Two-way: the Neandertal cut and the percentile cut are free, the Denisovan
    # cut is pinned where it cannot bind. Comparing this against the three-way
    # search isolates what the Denisovan threshold contributes, which the
    # one-parameter family below cannot do.
    rows_o2way = list(scan(To, X_GRID, [Y_GRID[0]], PCUTS))
    rows_o1 = list(scan(To, X_GRID, Y_GRID, [0.80], tie_y_to_x=True))
    print(f"grid: {len(rows_o):,} three-way rules, "
          f"{len(rows_o2way):,} two-way rules, "
          f"{len(rows_o1):,} one-parameter rules")

    # Even-chromosome counts for a specific rule, looked up on demand.
    def eval_even(x, y, p):
        nh, dl = Te[("nea", "hi", p)], Te[("den", "lo", p)]
        dh, nl = Te[("den", "hi", p)], Te[("nea", "lo", p)]
        return (int(np.count_nonzero((nh <= x) & (dl > y))),
                int(np.count_nonzero((dh <= x) & (nl > y))))

    hdr = (f"{'target':>7} {'x':>5} {'y':>6} {'p':>6} "
           f"{'odd n':>7} {'odd r':>7} {'even n':>7} {'even r':>7} "
           f"{'P(real)':>8} {'family':>9}")
    print("\n" + hdr)
    print("-" * len(hdr))
    out = []
    for target in TARGETS:
        half = target / 2.0
        for label, rows in (("three-way", rows_o), ("two-way", rows_o2way),
                            ("one-param", rows_o1)):
            sel = select(rows, half)
            if sel is None:
                print(f"{target:>7} {'':>5} {'':>6} {'':>6} "
                      f"{'no rule in band':>33} {label:>9}")
                continue
            x, y, p, co, ko = sel
            ce, ke = eval_even(x, y, p)
            ro = ko / co if co else float("nan")
            re_ = ke / ce if ce else float("nan")
            print(f"{target:>7} {x:>5} {y:>6} {p:>6} {co:>7} {ro:>7.3f} "
                  f"{ce:>7} {re_:>7.3f} {p_real(re_):>8.3f} {label:>9}")
            out.append({"target": target, "family": label, "x": x, "y": y,
                        "p": p, "odd_n": co, "odd_ctrl": ko, "odd_ratio": ro,
                        "even_n": ce, "even_ctrl": ke, "even_ratio": re_,
                        "p_real": round(p_real(re_), 4)})

    os.makedirs(os.path.dirname(args.out) or ".", exist_ok=True)
    with open(args.out, "w", encoding="utf-8") as f:
        keys = ["target", "family", "x", "y", "p", "odd_n", "odd_ctrl",
                "odd_ratio", "even_n", "even_ctrl", "even_ratio", "p_real"]
        f.write("\t".join(keys) + "\n")
        for r in out:
            f.write("\t".join(str(r[k]) for k in keys) + "\n")
    print(f"\nwrote {args.out}")

    # --- the curve: catalog size against ratio, with P(real) on the right ---
    import matplotlib
    matplotlib.use("Agg")
    import matplotlib.pyplot as plt

    fig, ax = plt.subplots(figsize=(8.4, 5.6))
    names = {
        "three-way": "all three cuts free (Nea, Den, percentile)",
        "two-way": "Nea + percentile cut free, Den cut non-binding",
        "one-param": "Nea cut only (Den tied, percentile fixed 80/20)",
    }
    for label, color, mk in (("three-way", "#1f4e79", "o"),
                             ("two-way", "#2e7d32", "^"),
                             ("one-param", "#c00000", "s")):
        pts = [r for r in out if r["family"] == label and r["even_n"] > 0]
        pts.sort(key=lambda r: r["even_n"])
        xs = [r["even_n"] for r in pts]
        ys = [r["even_ratio"] for r in pts]
        lo = [wilson_lo(r["even_ctrl"], r["even_n"]) for r in pts]
        hi = [wilson_hi(r["even_ctrl"], r["even_n"]) for r in pts]
        ax.plot(xs, ys, mk + "-", color=color, lw=1.8, ms=5,
                label=names[label], zorder=3)
        ax.fill_between(xs, lo, hi, color=color, alpha=0.13, lw=0, zorder=2)

    ax.axhline(NULL_RATIO, color="#555555", ls=":", lw=1.4, zorder=1)
    ax.text(0.99, NULL_RATIO, f" null skew {NULL_RATIO:.3f} (P(real)=0) ",
            transform=ax.get_yaxis_transform(), ha="right", va="bottom",
            fontsize=8, color="#555555")

    ax.set_xscale("log")
    ax.set_xlabel("catalog size (Neandertal-direction windows, held-out half)")
    ax.set_ylabel("Denisovan / Neandertal ratio")
    ax.set_ylim(0, 0.95)
    ax.grid(alpha=0.25, lw=0.5)
    ax.legend(loc="upper left", frameon=False, fontsize=9)
    ax.set_title("Best achievable purity at each catalog size "
                 "(rules selected on odd chromosomes, scored on even)",
                 loc="left", fontweight="bold", fontsize=10.5)

    # Right axis: the same quantity read as P(real), normalised by the null skew.
    ax2 = ax.twinx()
    ax2.set_ylim(ax.get_ylim())
    ticks = [t for t in (0, 0.2, 0.4, 0.6, 0.8) if t <= 0.95]
    ax2.set_yticks(ticks)
    ax2.set_yticklabels([f"{p_real(t):.2f}" for t in ticks])
    ax2.set_ylabel("P(real) = 1 - ratio / null skew")

    out_png = os.path.splitext(args.out)[0] + ".png"
    fig.tight_layout()
    fig.savefig(out_png, dpi=200, bbox_inches="tight")
    print(f"wrote {out_png}")
    return 0


if __name__ == "__main__":
    raise SystemExit(main())
```
