## Supplementary Information section 4 code base for "Hypothesis: A modern human range expansion ∼300,000 years ago had a critical role in Neandertal origins": p_nea_lt700.py

```
#!/usr/bin/env python3
"""
Posterior probability that the modern-plus-Neandertal TMRCA is below the
catalog threshold, P(T_Afr+Nea < 700 ka), at every catalog grid point.

P(real) measures how much a grid point is enriched for the catalog pattern
relative to the genome-wide null. It does not say how confident the underlying
TMRCA read itself is. This script reports the complementary quantity: given the
reconstructed posterior at a grid point, how much of its mass falls below the
700 ka threshold that defines the modern-Neandertal side of the catalog rule.

The posterior is reconstructed exactly as in approx_percentiles.py: Normal in
the ARGweaver time coordinate, with mu and sigma fixed by the stored 2.5 and
97.5 percentiles. 700 ka is itself a node of the run's time grid (times2.txt),
so no interpolation error enters at the threshold.

The Denisovan side, P(T_Afr+Den > 700 ka), is reported alongside it, and the
product of the two is given as a joint score. The product treats the two reads
as independent, which they are not (they share the same local tree), so it is
a convenient ranking score and not a calibrated probability.

Usage:
  python3 scripts/p_nea_lt700.py --scan hubisz_tmrca_1kb_8afr.tight.tsv \
      --regions k1/catX_k1v2_annot.regions.tsv -o k1/p_nea_lt700_1kb.tsv
"""

from __future__ import annotations

import argparse
import bisect
import csv
import math
import os
import statistics
import sys

sys.path.insert(0, os.path.dirname(os.path.abspath(__file__)))

from approx_percentiles import default_coord, fit_site  # noqa: E402

THRESH = 700.0
NEA = "02_modern_plus_both_Nea"
DEN = "03_modern_plus_Den"


def norm_cdf(z: float) -> float:
    return 0.5 * (1.0 + math.erf(z / math.sqrt(2.0)))


def p_below(coord, p025: float, p975: float, thresh: float) -> float:
    """Posterior mass below thresh under the reconstructed Normal-in-u model."""
    mu, sg = fit_site(coord, p025, p975)
    if sg <= 0:
        return 1.0 if coord.to_u(thresh) > mu else 0.0
    return norm_cdf((coord.to_u(thresh) - mu) / sg)


def load_regions(path):
    """Catalog loci as chrom -> sorted list of (start, end, index)."""
    by_chrom, rows = {}, []
    with open(path, encoding="utf-8") as f:
        for i, r in enumerate(csv.DictReader(f, delimiter="\t")):
            c, s, e = r["chrom"], int(r["start"]), int(r["end"])
            by_chrom.setdefault(c, []).append((s, e, i))
            rows.append(r)
    for c in by_chrom:
        by_chrom[c].sort()
    return by_chrom, rows


def cluster_ids(rows, join=100_000):
    """Group loci into clusters, pooling any pair within `join` bp."""
    order = sorted(range(len(rows)),
                   key=lambda i: (int(rows[i]["chrom"]), int(rows[i]["start"])))
    cid, out, prev = 0, {}, None
    for i in order:
        r = rows[i]
        if prev is not None:
            pc, pe = prev
            if r["chrom"] != pc or int(r["start"]) - pe > join:
                cid += 1
        out[i] = cid
        prev = (r["chrom"], int(r["end"]))
    return out


def main():
    ap = argparse.ArgumentParser()
    ap.add_argument("--scan", default="hubisz_tmrca_1kb_8afr.tight.tsv")
    ap.add_argument("--regions", default="k1/catX_k1v2_annot.regions.tsv")
    ap.add_argument("--threshold", type=float, default=THRESH)
    ap.add_argument("-o", "--out", default="k1/p_nea_lt700_1kb.tsv")
    ap.add_argument("--per-point", help="optional per-grid-point dump")
    a = ap.parse_args()

    by_chrom, rows = load_regions(a.regions)
    coord, desc = default_coord([])
    sys.stderr.write(f"time coordinate: {desc}\n")
    sys.stderr.write(f"threshold {a.threshold:.0f} ka -> grid index "
                     f"{coord.to_u(a.threshold):.4f}\n")

    per_locus = {i: [] for i in range(len(rows))}
    pts = []
    n_read = 0
    with open(a.scan, encoding="utf-8") as f:
        for r in csv.DictReader(f, delimiter="\t"):
            c = r["chrom"]
            iv = by_chrom.get(c)
            if not iv:
                continue
            pos = int(float(r["pos"]))
            j = bisect.bisect_right(iv, (pos, float("inf"), 0)) - 1
            # `end` is the last grid point of the locus, not one past it: a
            # 1 kb locus has start == end, and a 3 kb locus spans three points
            # start, start+1000, end. The test has to be inclusive at both
            # ends or every locus loses its last point and the 24 loci that
            # are a single grid point wide vanish altogether.
            if j < 0 or not (iv[j][0] <= pos <= iv[j][1]):
                continue
            try:
                pn = p_below(coord, float(r[f"{NEA}_p025_kya"]),
                             float(r[f"{NEA}_p975_kya"]), a.threshold)
                pd = 1.0 - p_below(coord, float(r[f"{DEN}_p025_kya"]),
                                   float(r[f"{DEN}_p975_kya"]), a.threshold)
            except (KeyError, ValueError):
                continue
            per_locus[iv[j][2]].append((pn, pd))
            n_read += 1
            if a.per_point:
                pts.append((c, pos, pn, pd))
    sys.stderr.write(f"{n_read:,} catalog grid points scored\n")

    cl = cluster_ids(rows)
    with open(a.out, "w", newline="", encoding="utf-8") as f:
        w = csv.writer(f, delimiter="\t")
        w.writerow(["chrom", "start", "end", "span_kb", "cluster",
                    "n_points", "p_nea_lt700_max", "p_nea_lt700_median",
                    "p_den_gt700_max", "p_joint_max", "genes"])
        for i, r in enumerate(rows):
            v = per_locus[i]
            if not v:
                continue
            pn = [x[0] for x in v]
            pd = [x[1] for x in v]
            jt = [x[0] * x[1] for x in v]
            w.writerow([r["chrom"], r["start"], r["end"], r["span_kb"], cl[i],
                        len(v), f"{max(pn):.3f}",
                        f"{statistics.median(pn):.3f}",
                        f"{max(pd):.3f}", f"{max(jt):.3f}",
                        r.get("genes", "")])
    sys.stderr.write(f"wrote {a.out}\n")

    if a.per_point:
        with open(a.per_point, "w", newline="", encoding="utf-8") as f:
            w = csv.writer(f, delimiter="\t")
            w.writerow(["chrom", "pos", "p_nea_lt700", "p_den_gt700"])
            for c, pos, pn, pd in pts:
                w.writerow([c, pos, f"{pn:.4f}", f"{pd:.4f}"])
        sys.stderr.write(f"wrote {a.per_point}\n")
    return 0


if __name__ == "__main__":
    raise SystemExit(main())
```
