## Supplementary Information section 4 code base for "Hypothesis: A modern human range expansion ∼300,000 years ago had a critical role in Neandertal origins": preal30_extension.py

```
#!/usr/bin/env python3
"""
How far can the fixed-derived screen be extended before P(real) reaches 0.3?

The catalog proper is the operating point: cut = 50, x = 625 ka, y = 700 ka,
which yields 329 grid points at P(real) = 0.80. For the fixed-derived screen
there is a reason to cast a wider net than that, since a coding change is
interesting even at moderate confidence and the screen's own evidence (fixation
in moderns, both Neandertals derived, Denisova ancestral) is independent of the
ARG. This finds the loosest rule of the catalog's own shape whose purity is
still 0.3.

TWO SENSES OF "DOWN TO P(real) = 0.3", REPORTED SEPARATELY
  cumulative  -- the whole extended set has P(real) = 0.3. Weaker demand,
                 larger set, and the number quoted describes the set a reader
                 is handed.
  marginal    -- the grid points added last have P(real) = 0.3, measured in the
                 shell between successive thresholds. Stricter, smaller, and it
                 is the honest reading of "going to lower probabilities of being
                 real", since it says what the least reliable member is worth.

Only x is relaxed; the percentile cut and the Denisovan threshold stay at the
catalog's values, so the extension is nested inside the catalog's rule shape
rather than being a different rule.

Example:
  python3 preal30_extension.py --scan hubisz_tmrca_5kb_8afr.tsv
"""

from __future__ import annotations

import argparse
import csv
import os
import sys
from collections import defaultdict

import numpy as np

sys.path.insert(0, os.path.dirname(os.path.abspath(__file__)))

from approx_percentiles import default_coord, load  # noqa: E402
from optimize_rule_by_size import NULL_RATIO, p_real, tails  # noqa: E402

CUT = 0.50
BASE_X = 625.0
Y = 700.0
PAD = 5000
TARGET = 0.30


def cluster(points, gap=5000, step=5000):
    """Merge grid points into loci, joining any within `gap` of each other.

    `step` is the width a grid point stands for and only sets how far a
    region's end reaches past its last point; `gap` is the separate choice of
    which points count as one locus. Both were 5000 while the scan was a 5 kb
    lattice. On the 1 kb scan `step` becomes 1000 and `gap` stays at 5000.
    """
    by = defaultdict(list)
    for c, p in points:
        by[c].append(p)
    out = []
    for c in sorted(by, key=lambda s: (len(s), s)):
        ps = sorted(by[c])
        s = e = ps[0]
        n = 1
        for p in ps[1:]:
            if p - e <= gap:
                e = p
                n += 1
            else:
                out.append((c, s, e, n))
                s = e = p
                n = 1
        out.append((c, s, e, n))
    return out


def main():
    ap = argparse.ArgumentParser()
    ap.add_argument("--scan", default="hubisz_tmrca_5kb_8afr.tsv")
    ap.add_argument("--grid-step", type=int, default=5000,
                    help="bp a grid point stands for; 1000 for the 1 kb scan")
    ap.add_argument("--cluster-gap", type=int, default=5000,
                    help="merge grid points this far apart into one locus")
    ap.add_argument("-o", "--out-prefix",
                    default="catalog_annotation/screen_preal30")
    a = ap.parse_args()

    recs = load(a.scan)
    coord, _ = default_coord(recs)
    recs = [d for d in recs if d["chrom"].replace("chr", "") not in ("X", "23")]
    T = tails(recs, coord)
    nh, nl = T[("nea", "hi", CUT)], T[("nea", "lo", CUT)]
    dh, dl = T[("den", "hi", CUT)], T[("den", "lo", CUT)]
    print(f"{len(recs):,} autosomal grid points; null omega0 = "
          f"{1/NULL_RATIO:.4f}\n")

    grid = list(range(int(BASE_X), 1001, 25))
    print(f"{'x (ka)':>7} {'case':>7} {'ctrl':>6} {'ratio':>7} {'P(real)':>8} "
          f"{'added':>7} {'shell P(real)':>14}")
    print("-" * 62)
    rows = []
    prev = None
    for x in grid:
        case = (nh <= x) & (dl > Y)
        ctrl = (dh <= x) & (nl > Y)
        c, k = int(case.sum()), int(ctrl.sum())
        pr = p_real(k / c)
        if prev is None:
            sh = ""
            shv = None
        else:
            dc, dk = c - prev[0], k - prev[1]
            shv = p_real(dk / dc) if dc > 0 else None
            sh = f"{shv:.3f}" if shv is not None else "-"
        print(f"{x:>7} {c:>7,} {k:>6,} {k/c:>7.3f} {pr:>8.3f} "
              f"{c - (prev[0] if prev else c):>7,} {sh:>14}")
        rows.append((x, c, k, pr, shv))
        prev = (c, k)

    cum = [r for r in rows if r[3] >= TARGET]
    x_cum = cum[-1][0] if cum else BASE_X
    mar = [r for r in rows if r[4] is None or r[4] >= TARGET]
    # walk from the base until the first shell that falls below target
    x_mar = BASE_X
    for x, c, k, pr, shv in rows:
        if shv is None or shv >= TARGET:
            x_mar = x
        else:
            break

    print(f"\ncumulative P(real) >= {TARGET}: x <= {x_cum:.0f} ka")
    print(f"marginal  P(real) >= {TARGET}: x <= {x_mar:.0f} ka")

    for label, x in (("cumulative", x_cum), ("marginal", x_mar)):
        case = (nh <= x) & (dl > Y)
        ctrl = (dh <= x) & (nl > Y)
        c, k = int(case.sum()), int(ctrl.sum())
        pts = [(d["chrom"].replace("chr", ""), int(d["pos"]))
               for d, m in zip(recs, case) if m]
        loci = cluster(pts, a.cluster_gap, a.grid_step)
        span = sum(e - s + a.grid_step for _, s, e, _ in loci)
        print(f"\n{label} set at x = {x:.0f} ka: {c} grid points, "
              f"{k} control, ratio {k/c:.3f}, P(real) {p_real(k/c):.3f}")
        print(f"  {len(loci)} loci, {span/1e6:.2f} Mb before padding, "
              f"{(span + 2*PAD*len(loci))/1e6:.2f} Mb with \u00b15 kb padding")

    # Write the marginal set: it is the defensible one to hand to the screen.
    x = x_mar
    case = (nh <= x) & (dl > Y)
    pts = [(d["chrom"].replace("chr", ""), int(d["pos"]))
           for d, m in zip(recs, case) if m]
    loci = cluster(pts, a.cluster_gap, a.grid_step)
    # Per-locus tier: the strictest x at which the locus already appears, which
    # is an ordinal confidence label, not a per-locus probability.
    tier_x = {}
    for xx in grid:
        m = (nh <= xx) & (dl > Y)
        for c2, p2 in [(d["chrom"].replace("chr", ""), int(d["pos"]))
                       for d, mm in zip(recs, m) if mm]:
            tier_x.setdefault((c2, p2), xx)
    prx = {r[0]: r[3] for r in rows}
    path = f"{a.out_prefix}.regions.tsv"
    os.makedirs(os.path.dirname(path) or ".", exist_ok=True)
    with open(path, "w", newline="") as fh:
        w = csv.writer(fh, delimiter="\t")
        w.writerow(["chrom", "start", "end", "n_sites", "span_kb",
                    "min_x_ka", "set_preal_at_min_x", "in_catalog"])
        for c2, s, e, n in loci:
            xs = [tier_x[(c2, p)] for p in range(s, e + 1, a.grid_step)
                  if (c2, p) in tier_x]
            mx = min(xs) if xs else x
            w.writerow([c2, s, e + a.grid_step, n,
                        (e - s + a.grid_step) // 1000,
                        f"{mx:.0f}", f"{prx.get(mx, float('nan')):.3f}",
                        "yes" if mx <= BASE_X else "no"])
    print(f"\nwrote {path} ({len(loci)} loci)")
    print("  in_catalog=yes marks loci already in the 329-point catalog;")
    print("  min_x_ka is the strictest threshold at which the locus appears.")


if __name__ == "__main__":
    raise SystemExit(main())
```
