## Supplementary Information section 4 code base for "Hypothesis: A modern human range expansion ∼300,000 years ago had a critical role in Neandertal origins": README.md

# Code for Supplementary Information section 4:
# "Catalog of variants that may be important in uniquely modern human biology"

Python 3.9+; dependencies in requirements.txt (numpy, scipy, matplotlib, pandas,
pysam, pyBigWig, tskit). Every script takes
its inputs and outputs as arguments and defaults to the paths used for the note, so the
commands below can be run as written from one directory. The sbatch_* and slurm_* scripts
write SLURM jobs (Harvard Medical School O2 conventions); each has a plain-Python or
single-chromosome equivalent that runs on one machine, shown in its docstring. All genomic
coordinates are GRCh37 / hg19, all dates are thousands of years assuming 29 years per
generation, and the grid is every 1 kb on the autosomes throughout. The
published run was originally at 5 kb; every script takes the grid step as an
argument, so both are reproducible from the same code, and the 5 kb numbers
quoted in earlier versions of the note are recovered by passing
--grid-step 5000 and the 5 kb scan.

Three external inputs are not redistributed here: the ARGweaver-D round-2 ARG (Hubisz et al.
2020, shared by Melissa Hubisz), the Relate genealogy inferred over 1000 Genomes phase 3
together with three Neandertals and a Denisovan (Speidel et al., shared by Leo Speidel), and
the high-coverage archaic VCFs (five Neandertals, two Denisovans) held under the laboratory's
data-sharing directory. Paths to all three are arguments, collected in archaic_panel.json for
the archaic genomes.

## Stage 1. The TMRCA scan
    python hubisz_out_bed_tmrca_scan.py --queries tmrca_queries.hubisz_8afr.json ...
    QUERIES=$REPO/scripts/tmrca_queries.hubisz_8afr.json GRID_STEP=1000 \
      bash submit_hubisz_tmrca_slurm.sh                # one array task per ARG chunk
    bash slurm_hubisz_tmrca_array_task.sh              # the array-task wrapper
    bash hubisz_tmrca_collate_parts.sh                 # -> hubisz_tmrca_1kb_8afr.tsv
    python resolve_by_interval.py                      # overlapping ARG windows
Parses the local Newick trees and writes, at every grid point, the posterior mean and the
2.5 and 97.5 percentiles of the TMRCA of each haplotype set: Africans alone, Africans plus
Neandertals, Africans plus Denisovans, and four further sets carried for reference.
The collated scan overlaps at chunk boundaries; resolve_by_interval.py keeps the read whose
95% posterior interval is narrower in absolute time, which is what leaves 2,440,090 autosomal
grid points. compare_hubisz_tmrca_runs.py and audit_argweaver_consistency.py are the checks
that the eight-genome run agrees with the earlier two-genome run and that the sets nest
correctly. The query file is tmrca_queries.hubisz_8afr.json (eight African diploids);
tmrca_queries.hubisz_8modern.json is the earlier four-modern-genome panel and is not the
scan behind this note.

QUERIES and GRID_STEP must both be set on the submit line, as shown above.
submit_hubisz_tmrca_slurm.sh submits with --export=ALL, so they reach the array task, whose
own defaults (a two-African query file, not redistributed here, at a 5 kb grid) are not the
scan behind this note. submit_hubisz_tmrca_slurm_8modern.sh and its array task
slurm_hubisz_tmrca_array_task_8modern.sh are the fixed-parameter pair that ran the earlier
four-modern-genome panel at 5 kb; they are kept only for the comparison reported below and
take no QUERIES or GRID_STEP argument.

    python approx_percentiles.py                       # intermediate percentiles
Reconstructs percentiles between the two reported ones by mapping the mean and the 2.5/97.5
bounds onto ARGweaver-D's own log-spaced time grid (times2.txt), fitting a normal in
grid-index coordinates, and reading off the percentile wanted. Imported by everything below
that needs a percentile cut other than 2.5 or 97.5.

## Stage 2. Building the catalog
    python discriminator_grid.py                       # case and control counts over (x, y, pcut)
    python threshold_landscape.py                      # the same counts as a surface
    python den_threshold_test.py                       # does decoupling the two thresholds help?
    python optimize_rule_by_size.py                    # cross-validated search over all three
    python make_catalog_2080.py --cut 0.50 --x 625 --y 700 --grid-step 1000
    python make_loci_table.py --interval-style central90   # -> Table S4.1
The catalog rule is x = 625 ka on the African-plus-Neandertal side, y = 700 ka on the
African-plus-Denisovan side, at the median (pcut = 50). Controls are the same rule with the
two archaic labels interchanged, and P(real) is one minus the control-to-case ratio.
den_threshold_test.py and optimize_rule_by_size.py are the two answers to "could this rule be
better", the second by chromosome-parity cross-validation so no grid point is scored by a
model fitted on its own chromosome.

    python fit_fdr_surface.py                          # binomial logistic on the case/control counts
    python build_preal_grid.py                         # -> preal_gridpoints_1kb.tsv (all 2,440,090)
    python p_nea_lt700.py                              # P(TAfr+Nea < 700 ka) for Table S4.1
The surface is fitted by Fisher scoring with internal standardization and a deviance-based
line search, and evaluated at the boundary of its support for grid points whose TMRCAs fall
outside the fitted region; build_preal_grid.py flags those as extrapolated. The locus figures
need P(real) away from catalog members, which is why the surface is cached genome-wide rather
than recomputed per figure.

## Stage 3. The null, and background selection
    python null_ratio_audit.py                         # is the control catalog an honest null?
    python bstat_to_grid.py                            # B statistic onto the 1-kb grid
    python null_ratio_shell_bstat.py                   # control/case ratio within B shells
    python bstat_incremental_value.py                  # B-conditioned rules vs the flat rule
Tests both B-statistic maps (CADD- and phastCons-calibrated), comparing the single-threshold
rule against a threshold that shifts with B decile and against a free threshold per decile.
At matched held-out catalog size none of them improves on the flat rule, which is the result
reported in the note.

## Stage 4. Gene and functional annotation
    python annotate_catalog_regions.py                 # GENCODE v19 + MyGene.info -> Table S4.1 genes
    python annotate_gene_list_function.py              # the prose summaries of each gene
Merges catalog grid points within 5 kb, extends each by 5 kb, and intersects with GENCODE v19
gene bodies (all biotypes, introns included), recording the distance to the nearest gene
where nothing overlaps.

## Stage 5. The fixed-derived screen
    python preal30_extension.py                        # -> screen_preal30.padded.regions.tsv
    bash sbatch_extract_chagyrskaya.sh                 # Chagyrskaya 8 VCFs are plain gzip, not BGZF
    bash o2_build_venv.sh                              # the pysam venv the two sbatch runs need
    sbatch sbatch_index_archaics.sh                    # tabix indices the screen requires
    MAXAF=0.01 sbatch o2_sbatch_fixed_derived.sh       # the >99% modern-frequency screen
    MAXAF=0.05 sbatch o2_sbatch_fixed_derived.sh       # the >95% sensitivity run
    python o2_screen_fixed_derived.py --help           # the same screen for one region, locally
The screen runs on a deliberately looser region set than the catalog: x relaxed from 625 to
675 ka with the other two parameters held, giving 2,621 grid points in 287 regions covering
2.98 Mb before padding (5.56 Mb after), against 1.82 Mb for the catalog itself. Within it, every
1000 Genomes phase 3
biallelic SNV is polarized against the Ensembl six-primate ancestral sequence and kept if the
derived allele is above the modern-frequency threshold, derived in every Neandertal with a
call, and ancestral in every Denisovan with a call. Because the rule demands unanimity among
genomes that have data, the per-genome state columns are carried through to the output rather
than collapsed. archaic_gt_check_derived_pattern.py and compare_screen_processings.py verify
the genotype logic and that the two callers (GATK-era and snpAD) agree where they overlap.

    python fetch_cadd_regions.py                       # CADD scores by region, not site-by-site
    python assemble_fd_annotation.py                   # join annotation; resolve overlap duplicates
    python annotate_functional_classes.py              # GENCODE v19, phastCons, phyloP, CADD, ENCODE
    python annotate_fixed_derived_consequences.py      # codon and amino-acid consequences
    python canonicalize_protein_positions.py           # residue numbers onto canonical transcripts
    python make_functional_table.py                    # -> Table S4.2
Table S4.2 is the strongest subset: a change to coding sequence, an essential splice position,
a CADD v1.6 score in the top 1% of possible substitutions, or a conserved element coinciding
with a UTR or promoter. Synonymous changes never qualify: CADD is scaled across all
substitution types and being inside a coding exon raises it on its own, while none of the
synonymous fixed-derived sites is at a conserved base. The screen records whichever transcript
its annotation source returned first, which is often not the canonical one, so
canonicalize_protein_positions.py re-reads the coding rows through Ensembl VEP on GRCh37 and
overwrites the residue numbers; without it the reported positions cannot be checked against
the domain literature.

P(real) for these sites is read at the substitution itself rather than at the nearest point of
the 1-kb lattice. ARGweaver-D carries a local tree per recombination interval, so the scan
script will report at named bases instead of a grid:

    bash submit_hubisz_tmrca_sites.sh                  # scan at --positions-tsv, not a lattice
    bash slurm_hubisz_tmrca_sites_task.sh              # the array-task wrapper
    python resolve_by_interval.py                      # pick between overlapping ARG windows
    python site_exact_preal.py                         # evaluate the fitted surface at those bases
The surface is still fitted on the genome-wide scan under the same odd/even chromosome
cross-validation; only the TMRCAs it is evaluated at change. 450 of the 1,223 sites move by
more than 0.05 against their nearest lattice point, because TMRCA changes in steps at
recombination boundaries. The ARG windows overlap by 100 kb, so a site in an overlap is read
twice from separately inferred trees; resolve_by_interval.py keeps the read whose 95% posterior
interval is narrower in absolute time, which is the same rule applied to the genome-wide grid.
Applying it in both places is the point: when the halves disagreed, a base could be assigned a
different TMRCA by its own read than by the grid it sits on, which is how TOMM20L came to carry
a locus P(real) of 0.83 beside a site value of 0.00.

## Stage 6. The Relate replication
    bash sbatch_relate_archaic.sh                      # or cluster_relate_one_chr.sh per chromosome
    sbatch sbatch_relate_tmrca_chr_array.sh            # TMRCA at the same 1-kb grid
    python relate_ts_tmrca_grid.py                     # tree sequence -> per-grid-point TMRCA
    python relate_marginal_tmrca_grid.py               # called by cluster_relate_one_chr.sh
    python relate_locus_percentiles.py                 # per-catalog-locus percentile
    python relate_modern_vs_argweaver.py               # concordance between the two methods
    python optimize_relate_rule.py                     # how well any Relate rule could do
Takes the TMRCA of the modern haplotypes only from a genealogy that also contains the
archaics, and places each catalog locus in the genome-wide distribution of the same quantity
averaged over blocks of equal size. Chromosome lists and hg19 lengths are
relate_chrom_list_autosomes_22.txt and relate_hg19_chrom_sizes.tsv. Percentiles rather than
dates, because ARGweaver-D and Relate do not share a timescale and a percentile survives any
monotone rescaling.
optimize_relate_rule.py is the ceiling calculation: the best a Relate-only catalog could do,
which is what makes the replication rate interpretable.

## Figures
    python fig_s51_model_curve.py                      # the cross-validated surface for panel B
    python fig_s51_composite.py                        # -> Figure S4.1
    python fig_s51_sensitivity.py                      # counting helpers imported by both
    python fig_locus_zoom.py                           # -> Figure S4.2, the ten locus panels
fig_locus_zoom.py reads preal_gridpoints.tsv, the fixed-derived sites, and GENCODE v19, and
runs each window outward until the African-plus-Neandertal TMRCA recovers above 1500 ka.

## Supplementary tables
    python build_supp_table1.py --bounds 5-95          # -> supplementary_table2_scan_1kb.tsv.gz
Both this and relate_locus_percentiles.py read the Relate scan from relate_archaic_1kb/,
the 1 kb grid. The 5 kb scan in relate_archaic/ is the earlier run; it is reproduced
exactly by the 1 kb one at every shared position, but it can only score 111 of the 141
catalog loci and leaves four rows in five of Supplementary Table 2 blank, so it is not
the grid either table is built from.
    python build_supp_table2.py                        # -> supplementary_table3_fixed_derived.tsv
    python screen_coverage.py                          # fraction of the autosomes with ARG trees
Supplementary Table 2 is one row per autosomal grid point (2,440,090), carrying the ARGweaver-D
dates with 5th and 95th percentile bounds (the 90% central interval used by Table S4.1), P(real),
the catalog flag, and where Relate's modern-only TMRCA at that point falls in the genome-wide
Relate distribution for two panels (216 Yoruba and 604 Yoruba/British/Han haplotypes).
Supplementary Table 3 is one row per fixed-derived site (1,223: 991 at >99% derived in 1000
Genomes plus 232 added at >95%), with derived-over-observed haplotype counts for moderns,
Neandertals and Denisovans, the per-genome states behind them, 90% TMRCA intervals, and the
functional annotation.

## Stage 7. Gene ontology
    python great_domains.py                            # GENCODE v19 regulatory domains
    python great_test.py                               # hypergeometric and binomial tests
    bash run_great_100k.sh                             # 100,000 B-matched block permutations
    python great_matched_100k.py                       # the permutation worker
    python great_annotation_bias.py                    # annotation-depth controls
    python great_negative_control.py                   # Denisovan-side mirror catalog
These read go-basic.obo and goa_human.gaf from the Gene Ontology Consortium (release 2026-07-26).
The permutation relocates all 66 clusters to blocks of equal length and similar mean B.

## Calibration onto a published timescale
    python recalibrate_peyregne.py                     # ARGweaver-D dates -> the published scale
    python fig_calibration.py
Reported for orientation only. The catalog rule is applied on the ARGweaver-D scale, and no
result in the note depends on the recalibration.
