## Supplementary Information section 4 code base for "Hypothesis: A modern human range expansion ∼300,000 years ago had a critical role in Neandertal origins": recalibrate_peyregne.py

```
#!/usr/bin/env python3
"""
Put the Hubisz 2020 ARGweaver-D TMRCA dates onto the Peyregne 2025 calibration.

The common quantity
-------------------
Both papers report a modern-human vs archaic (Neandertal+Denisovan) POPULATION
SPLIT TIME, and both use the same clock (mu = 1.45e-8 /bp/gen, 29 yr/gen,
0.5e-9 /bp/yr; see catalog_annotation/recalibration_notes.md sections 1-2):

    Hubisz 2020:    S_H = 575 ka   (fixed model input, from Prufer 2017)
    Peyregne 2025:  S_P = 694-825 ka  (estimated, neutral regions, divergence)

Because the clocks agree, this is NOT a clock rescaling and a multiplicative
factor S_P/S_H is wrong: it would stretch the coalescent waiting time W as well
as the split S, and under Hubisz's own model W dominates (E[W] = 2*18,500 gen =
1.07 Myr, versus S_H = 0.575 Myr). See recalibration_notes.md section 3c.

Why truncation, not an additive shift
-------------------------------------
recalibration_notes.md prescribes T_P = T_H + (S_P - S_H), flagged there as an
upper bound because ARGweaver posteriors are part prior, part data. That bound
can be replaced by something exact under the same model.

For two lineages sampled either side of a split at S, the prior is

    pi(T | S) = (1/2N) exp(-(T-S)/2N) * 1{T >= S},   N = Ne_anc, constant.

Revising S while holding the clock and the likelihood L(data|T) fixed:

    pi_P(T) / pi_H(T) = exp((S_P - S_H)/2N) * 1{T >= S_P}   (a constant, times
                                                             an indicator)

because the exponential ancestral coalescent is shift-invariant in S. So

    p(T | data, S_P)  =  p(T | data, S_H) truncated below at S_P, renormalised.

No free parameters, and it self-adjusts across the two regimes the note had to
bracket by hand:

  * a window pinned at the prior floor (posterior mass just above S_H) loses
    almost all its mass to the truncation and re-forms just above S_P, giving
    close to the full additive shift;
  * a deep, mutation-anchored window (all mass already above S_P) is untouched,
    giving no shift.

The posterior per window is reconstructed from the two stored percentiles by
approx_percentiles.py (Normal in ARGweaver's own log-spaced grid coordinate,
validated against the held-out posterior mean).

Does the floor actually bind?
-----------------------------
The truncation argument needs the prior to be hard-floored at S, which is not
automatic: ARGweaver-D models gene flow, and an introgressed lineage could in
principle coalesce across populations more recently than the split. Measured on
all 509,601 autosomal windows of the 8afr scan, using the sampler's own stored
posterior means:

    posterior mean below 575 ka    0.02% of windows
    p025 below 575 ka              0.22%
    density either side of 575 ka  26 windows in 575-600, 8,992 in 700-800

The distribution stops at the split instead of passing through it. The floor
binds, and truncation is the right operation.

Why the BASELINE is also truncated
----------------------------------
The reconstructed posterior is Normal in the grid coordinate, fitted to p025
and p975. For a floored window that shape is wrong in a specific way: the true
posterior piles against the floor with a long left tail (p025 as low as 300 ka
for catalog windows), so a symmetric fit misplaces a large share of the mass
below 575 ka -- a median of 60% for catalog windows, which cannot be real given
the genome-wide result above.

That sub-floor mass is a reconstruction artefact, not gene flow, and the fix is
to impose the floor the model already implies. Checked against the stored
posterior mean, which is never used in the fit:

    catalog T(Afr+Nea), median stored mean          634 ka
      untruncated fit                               585 ka   (-8%)
      fit truncated at S_H = 575 ka                 678 ka   (+7%)

So the Hubisz-scale baseline is the fit truncated at S_H, and the Peyregne
scale is the same fit truncated at S_P. Both scales pass through identical
machinery, so the residual bias largely cancels in the difference, which is the
quantity being reported. The `*_subfloor` column records the discarded mass so
the size of the correction is visible per window.

Scope and caveats
-----------------
  * Applies to TMRCAs that must cross the modern-archaic split: T(Afr+Nea) and
    T(Afr+Den). Both are floored at S up to the gene-flow leak above.
  * T(Afr) (modern-only) is NOT floored at S, so the truncation does not apply
    and it is reported unchanged, as recalibration_notes.md section 3c
    prescribes. Second-order effect not modelled: with a later split, modern
    lineages spend longer at modern Ne before entering the ancestral
    population, which would make deep T(Afr) values slightly younger.
  * Shift-invariance of the prior assumes the number of lineages entering the
    ancestral population is unchanged. A later split gives modern lineages more
    time to coalesce below it, so slightly fewer enter; that makes the
    truncation result a mild overestimate of the true shift.
  * Exact recalibration would require re-running ARGweaver-D with the revised
    demography. This is the closest thing available without the sampler.

Usage
-----
  python3 scripts/recalibrate_peyregne.py \
      --catalog catalog_annotation/catalog_final_median_x625_auto.sites.tsv \
      --scan hubisz_tmrca_5kb_8afr.tsv \
      --emit catalog_annotation/catalog_recalibrated_peyregne.tsv \
      --figure catalog_annotation/fig_recalibration.png
"""

from __future__ import annotations

import argparse
import csv
import math
import os
import random
import sys

import numpy as np

sys.path.insert(0, os.path.dirname(os.path.abspath(__file__)))
from approx_percentiles import default_coord, fit_site, load  # noqa: E402

# ---------------------------------------------------------------------------
# Calibration constants. Sources in catalog_annotation/recalibration_notes.md.
# ---------------------------------------------------------------------------
S_HUBISZ = 575.0                     # ka, fixed modern-archaic split (S 1a/1c)
S_PEYREGNE = {"low": 694.0, "mid": 759.5, "high": 825.0}   # ka (S 2a)
NE_ANC_GEN = 18500.0                 # diploid Ne above the split (pop_sizes2)
YEARS_PER_GEN = 29.0
# Nea-Den split, for reference only: 415 ka fixed vs 504-585 ka estimated.
S_NEADEN_HUBISZ, S_NEADEN_PEYREGNE = 415.0, (504.0, 585.0)

PCTL = (0.025, 0.5, 0.975)


# ---------------------------------------------------------------------------
# Vectorised version of approx_percentiles.GridCoord.to_t, needed because the
# quadrature below evaluates it a few thousand times per window.
# ---------------------------------------------------------------------------
def make_to_t(coord):
    g = np.asarray(coord.g, dtype=float)
    idx = np.arange(len(g), dtype=float)
    lo_step = max(g[1] - g[0], 1e-9)
    hi_step = max(g[-1] - g[-2], 1e-9)

    def to_t(u):
        u = np.asarray(u, dtype=float)
        t = np.interp(u, idx, g)
        hi = u > idx[-1]
        if hi.any():
            t[hi] = g[-1] + (u[hi] - idx[-1]) * hi_step
        lo = u < 0
        if lo.any():
            t[lo] = g[0] + u[lo] * lo_step
        return t

    return to_t


def trunc_stats(to_t, mu, sigma, u_lo, ps=PCTL, n=2001):
    """Mean and quantiles of Normal(mu, sigma) in the grid coordinate u,
    truncated to u >= u_lo and mapped through t(u).

    Weights are held in log space and re-centred before exponentiating, so a
    window whose posterior sits far below the truncation point (alpha of 30+,
    where the raw density underflows) is still handled correctly.
    """
    if sigma <= 1e-12:
        t = float(to_t(np.array([max(mu, u_lo)]))[0])
        return t, {p: t for p in ps}

    alpha = (u_lo - mu) / sigma
    if alpha <= 0:
        hi = mu + 8.0 * sigma
    else:
        # Above the truncation point the density decays with length
        # sigma/alpha, so the integration window must shrink with alpha or the
        # quadrature grid misses the mass entirely.
        hi = u_lo + 10.0 * sigma / max(alpha, 1.0)
    hi = max(hi, u_lo + 1e-9)

    u = np.linspace(u_lo, hi, n)
    z = (u - mu) / sigma
    logw = -0.5 * z * z
    logw -= logw.max()
    w = np.exp(logw)
    t = to_t(u)

    mass = np.trapezoid(w, u) if hasattr(np, "trapezoid") else np.trapz(w, u)
    mean = (np.trapezoid(w * t, u) if hasattr(np, "trapezoid")
            else np.trapz(w * t, u)) / mass

    cdf = np.concatenate([[0.0], np.cumsum(0.5 * (w[1:] + w[:-1]) * np.diff(u))])
    cdf /= cdf[-1]
    q = {p: float(np.interp(p, cdf, t)) for p in ps}
    return float(mean), q


def recalibrate(coord, to_t, p025, p975, s_p, ps=PCTL):
    mu, sg = fit_site(coord, p025, p975)
    return trunc_stats(to_t, mu, sg, coord.to_u(s_p), ps)


def hubisz_stats(coord, to_t, p025, p975, ps=PCTL):
    """Same machinery with the truncation at S_H, so the before/after
    comparison is like-for-like rather than mixing stored and modelled values.
    """
    mu, sg = fit_site(coord, p025, p975)
    return trunc_stats(to_t, mu, sg, coord.to_u(S_HUBISZ), ps)


def fitted_mean(to_t, coord, p025, p975, n=2001):
    """Untruncated mean of the fitted posterior. This is the quantity that
    reproduces the stored posterior mean (approx_percentiles validates it to
    ~1.5% genome-wide), so it is the honest baseline to quote shifts against.
    """
    mu, sg = fit_site(coord, p025, p975)
    if sg <= 1e-12:
        return float(to_t(np.array([mu]))[0])
    u = np.linspace(mu - 8 * sg, mu + 8 * sg, n)
    w = np.exp(-0.5 * ((u - mu) / sg) ** 2)
    t = to_t(u)
    tz = np.trapezoid if hasattr(np, "trapezoid") else np.trapz
    return float(tz(w * t, u) / tz(w, u))


def subfloor_mass(coord, p025, p975, t):
    """Share of the reconstructed posterior lying below t. Evaluated at t = S_H
    it measures the reconstruction artefact described in the header, i.e. how
    much mass the symmetric fit misplaces below the model's own floor.
    """
    mu, sg = fit_site(coord, p025, p975)
    if sg <= 1e-12:
        return 1.0 if mu < coord.to_u(t) else 0.0
    z = (coord.to_u(t) - mu) / sg
    return 0.5 * math.erfc(-z / math.sqrt(2.0))


def med(x):
    return float(np.median(np.asarray(x, dtype=float))) if len(x) else float("nan")


def read_catalog(path):
    with open(path, encoding="utf-8") as f:
        return list(csv.DictReader(f, delimiter="\t"))


# ---------------------------------------------------------------------------
def main() -> int:
    ap = argparse.ArgumentParser()
    ap.add_argument("--catalog",
                    default="catalog_annotation/catalog_final_median_x625_auto.sites.tsv")
    ap.add_argument("--scan", default="hubisz_tmrca_5kb_8afr.tsv")
    ap.add_argument("--emit", default="")
    ap.add_argument("--figure", default="")
    ap.add_argument("--threshold", type=float, default=625.0,
                    help="the catalog rule's Hubisz-scale Neandertal cut")
    ap.add_argument("--genome-sample", type=int, default=20000)
    ap.add_argument("--seed", type=int, default=0)
    args = ap.parse_args()

    coord, which = default_coord([])
    to_t = make_to_t(coord)
    grid = coord.g

    print("=" * 78)
    print("Recalibrating Hubisz 2020 TMRCAs onto the Peyregne 2025 split")
    print("=" * 78)
    print(f"time coordinate: {which}")
    print(f"grid (ka): {[int(g) for g in grid[:14]]} ...")
    print(f"\nanchor  : modern-human vs archaic population split time")
    print(f"  Hubisz   S_H = {S_HUBISZ:.0f} ka  (fixed input)")
    print(f"  Peyregne S_P = {S_PEYREGNE['low']:.0f}-{S_PEYREGNE['high']:.0f} ka"
          f"  (mid {S_PEYREGNE['mid']:.1f})")
    print(f"  additive shift S_P - S_H = "
          f"+{S_PEYREGNE['low']-S_HUBISZ:.0f} to "
          f"+{S_PEYREGNE['high']-S_HUBISZ:.0f} ka"
          f"  (mid +{S_PEYREGNE['mid']-S_HUBISZ:.1f})")
    ew = 2 * NE_ANC_GEN * YEARS_PER_GEN / 1000.0
    print(f"  clock identical in both papers, so this is a demographic "
          f"correction, not a rescale")
    print(f"  E[W] in the ancestral population = 2Ne = "
          f"{2*NE_ANC_GEN:.0f} gen = {ew:.0f} ka, i.e. {ew/S_HUBISZ:.2f}x the "
          f"split itself -> multiplicative rescaling is wrong")

    # -- how the three candidate maps differ, at the grid points ------------
    print("\n" + "-" * 78)
    print("Map comparison at the ARGweaver grid points (Peyregne mid, S_P = "
          f"{S_PEYREGNE['mid']:.1f} ka)")
    print("-" * 78)
    print(f"{'T_Hubisz':>9} | {'multiplic.':>10} {'additive':>9} | "
          f"{'truncation, by posterior width':>34}")
    print(f"{'(ka)':>9} | {'(wrong)':>10} {'(note 3c)':>9} | "
          f"{'narrow':>10} {'typical':>10} {'wide':>10}")
    f_mid = S_PEYREGNE["mid"] / S_HUBISZ
    u_p_mid = coord.to_u(S_PEYREGNE["mid"])
    # Posterior widths in grid units, spanning what the scan actually shows.
    widths = (0.5, 1.5, 3.0)
    for g in [x for x in grid if 400 <= x <= 3000]:
        cells = []
        for wgu in widths:
            sg = wgu / (2 * 1.959964)
            mu = coord.to_u(g)
            m, _ = trunc_stats(to_t, mu, sg, u_p_mid, ps=(0.5,))
            cells.append(f"{m:10.0f}")
        add = g + (S_PEYREGNE["mid"] - S_HUBISZ) if g > S_HUBISZ else g
        print(f"{g:9.0f} | {g*f_mid:10.0f} {add:9.0f} | " + " ".join(cells))
    print("  'narrow/typical/wide' = 95% posterior interval spanning 0.5 / 1.5 /")
    print("  3.0 grid steps. Note the truncation map converges to the additive")
    print("  shift near the floor and to the identity when the window is deep.")

    # -- the catalog --------------------------------------------------------
    rows = read_catalog(args.catalog)
    print("\n" + "-" * 78)
    print(f"Catalog: {os.path.basename(args.catalog)}  ({len(rows)} sites)")
    print("-" * 78)

    out = []
    for r in rows:
        rec = {"chrom": r["chrom"], "pos": int(float(r["pos"]))}
        for tag in ("nea", "den"):
            lo = float(r[f"{tag}_p025_kya"])
            hi = float(r[f"{tag}_p975_kya"])
            hm, hq = hubisz_stats(coord, to_t, lo, hi)
            rec[f"{tag}_mean_hubisz"] = hm
            rec[f"{tag}_p50_hubisz"] = hq[0.5]
            rec[f"{tag}_mean_stored"] = float(r[f"{tag}_mean_kya"])
            rec[f"{tag}_mean_fitted"] = fitted_mean(to_t, coord, lo, hi)
            rec[f"{tag}_subfloor"] = subfloor_mass(coord, lo, hi, S_HUBISZ)
            for k, s_p in S_PEYREGNE.items():
                m, q = recalibrate(coord, to_t, lo, hi, s_p)
                rec[f"{tag}_mean_peyregne_{k}"] = m
                rec[f"{tag}_p50_peyregne_{k}"] = q[0.5]
                rec[f"{tag}_shift_{k}"] = m - hm
                # The value to report. The modelled absolute level carries a
                # reconstruction bias (+7% for Nea, +20% for Den against the
                # stored mean), but the shift is a difference through identical
                # machinery on both scales, so that bias cancels. Take the
                # level from the sampler and the movement from the model.
                rec[f"{tag}_recal_{k}"] = rec[f"{tag}_mean_stored"] + (m - hm)
                if k == "mid":
                    rec[f"{tag}_p025_peyregne_mid"] = q[0.025]
                    rec[f"{tag}_p975_peyregne_mid"] = q[0.975]
            # Like-for-like: both scales are the same fitted posterior, one
            # truncated at S_H and one at S_P, so the reconstruction bias
            # largely cancels. See the header for why the baseline is
            # truncated rather than raw.
            rec[f"{tag}_shift_mid"] = (rec[f"{tag}_mean_peyregne_mid"]
                                       - rec[f"{tag}_mean_hubisz"])
            rec[f"{tag}_shift_vs_stored"] = (rec[f"{tag}_mean_peyregne_mid"]
                                             - rec[f"{tag}_mean_stored"])
        rec["afr_mean_kya"] = float(r["afr_mean_kya"])   # unchanged, see header
        out.append(rec)

    # -- how far into the fitted posterior the truncation reaches -----------
    # The posterior is reconstructed from p025 and p975 only, so mass beyond
    # p975 is Normal-tail extrapolation. If S_P lands above p975 the answer for
    # that window rests on the extrapolation rather than on the fit, and should
    # be read as the additive shift rather than as a measurement.
    print("\nwhere the truncation point falls inside the stored posterior:")
    for tag, label in (("nea", "T(Afr+Nea)"), ("den", "T(Afr+Den)")):
        lo = np.array([float(r[f"{tag}_p025_kya"]) for r in rows])
        hi = np.array([float(r[f"{tag}_p975_kya"]) for r in rows])
        s = S_PEYREGNE["high"]     # worst case of the three
        print(f"  {label}: S_P(high)={s:.0f} ka is below p025 for "
              f"{100*(s < lo).mean():4.0f}% of windows (no truncation at all), "
              f"inside [p025,p975] for {100*((s >= lo) & (s <= hi)).mean():4.0f}%, "
              f"above p975 for {100*(s > hi).mean():4.0f}% (extrapolated tail)")

    # -- validation of the reconstruction, before any recalibration ---------
    # The stored posterior mean is never used in the fit, so it is a genuine
    # held-out check on which baseline to use.
    print("\nchoosing the baseline: both candidates against the stored mean")
    print(f"{'':<12} {'stored':>8} {'untrunc fit':>12} {'fit trunc@S_H':>14} "
          f"{'subfloor mass':>14}")
    for tag, label in (("nea", "T(Afr+Nea)"), ("den", "T(Afr+Den)")):
        s = med([d[f"{tag}_mean_stored"] for d in out])
        u = med([d[f"{tag}_mean_fitted"] for d in out])
        h = med([d[f"{tag}_mean_hubisz"] for d in out])
        sf = med([d[f"{tag}_subfloor"] for d in out])
        print(f"  {label:<10} {s:8.0f} {u:8.0f}{f'({(u-s)/s:+.0%})':>4} "
              f"{h:10.0f}{f'({(h-s)/s:+.0%})':>4} {sf:13.0%}")
    print("  The untruncated fit puts a median 60% of a catalog window's")
    print("  Neandertal posterior below 575 ka, which the genome-wide floor")
    print("  test rules out. That mass is a reconstruction artefact of forcing")
    print("  a symmetric shape onto a floor-piled posterior, so the baseline")
    print("  is the fit truncated at S_H, and shifts below are like-for-like.")

    print()
    print("VALUES TO REPORT: the sampler's own date, moved by the modelled")
    print("shift. Absolute modelled levels are not quoted because they carry")
    print("the reconstruction bias shown above; the shift does not.")
    hdr = (f"{'quantity':<40} {'Hubisz':>8} {'Peyr low':>9} {'Peyr mid':>9} "
           f"{'Peyr high':>10}")
    print(hdr)
    print("-" * len(hdr))
    for tag, label in (("nea", "T(Afr+Nea)"), ("den", "T(Afr+Den)")):
        base = med([d[f"{tag}_mean_stored"] for d in out])
        vals = [med([d[f"{tag}_recal_{k}"] for d in out])
                for k in ("low", "mid", "high")]
        print(f"{label + ', median over catalog (ka)':<40} {base:8.0f} "
              f"{vals[0]:9.0f} {vals[1]:9.0f} {vals[2]:10.0f}")
    print("\nfor reference, the modelled absolutes (biased level, see above):")
    for tag, label in (("nea", "T(Afr+Nea)"), ("den", "T(Afr+Den)")):
        base = med([d[f"{tag}_mean_hubisz"] for d in out])
        vals = [med([d[f"{tag}_mean_peyregne_{k}"] for d in out])
                for k in ("low", "mid", "high")]
        print(f"  {label + ', modelled mean':<38} {base:8.0f} "
              f"{vals[0]:9.0f} {vals[1]:9.0f} {vals[2]:10.0f}")

    # realised shift, which is the whole point
    print()
    full = S_PEYREGNE["mid"] - S_HUBISZ
    for tag, label in (("nea", "T(Afr+Nea)"), ("den", "T(Afr+Den)")):
        sh = [d[f"{tag}_shift_mid"] for d in out]
        st = [d[f"{tag}_shift_vs_stored"] for d in out]
        print(f"{label} shift at Peyregne mid: median {med(sh):+.0f} ka "
              f"({100*med(sh)/full:.0f}% of the flat +{full:.0f} ka shift); "
              f"against the raw stored mean, {med(st):+.0f} ka")
    print("  The Neandertal figure can exceed 100% because truncation is not")
    print("  translation: for a floored window the posterior is right-skewed,")
    print("  so conditioning on T >= S_P keeps only the upper tail, whose mean")
    print("  sits deeper than a translated mean would.")

    gap_h = [d["den_mean_stored"] - d["nea_mean_stored"] for d in out]
    gaps = {k: [d[f"den_recal_{k}"] - d[f"nea_recal_{k}"] for d in out]
            for k in S_PEYREGNE}
    print(f"\nDen-minus-Nea gap (the contrast the catalog selects on):")
    print(f"  Hubisz scale       median {med(gap_h):+.0f} ka")
    for k in ("low", "mid", "high"):
        print(f"  Peyregne {k:<10} median {med(gaps[k]):+.0f} ka   "
              f"({100*(med(gaps[k])-med(gap_h))/med(gap_h):+.0f}% vs Hubisz)")

    # -- what the rule's threshold becomes ---------------------------------
    print("\n" + "-" * 78)
    print(f"Translating the catalog rule (Hubisz-scale Neandertal cut at "
          f"{args.threshold:.0f} ka)")
    print("-" * 78)
    near = [d for d in out if abs(d["nea_p50_hubisz"] - args.threshold) <= 40]
    print(f"  {len(near)} catalog sites sit within 40 ka of the cut; their "
          f"recalibrated posterior medians:")
    for k in ("low", "mid", "high"):
        v = [d[f"nea_p50_peyregne_{k}"] for d in near]
        if v:
            print(f"    Peyregne {k:<5} median {med(v):.0f} ka "
                  f"(range {min(v):.0f}-{max(v):.0f})")
    print("  The map is monotone, so catalog MEMBERSHIP is unchanged by")
    print("  recalibration: it re-labels the axis, it does not re-rank windows.")

    # -- genome-wide context -----------------------------------------------
    if args.genome_sample and os.path.exists(args.scan):
        recs = load(args.scan)
        recs = [d for d in recs if d["chrom"] != "X"]
        random.seed(args.seed)
        sub = random.sample(recs, min(args.genome_sample, len(recs)))
        print("\n" + "-" * 78)
        print(f"Does the prior floor bind? (all {len(recs):,} autosomal windows)")
        print("-" * 78)
        print("  Truncation needs a hard floor at S_H. ARGweaver-D models gene")
        print("  flow, so an introgressed lineage could in principle coalesce")
        print("  across populations more recently. Measured:")
        for tag, label in (("nea", "T(Afr+Nea)"), ("den", "T(Afr+Den)")):
            p025 = np.array([d[f"{tag}_p025"] for d in recs])
            mean = np.array([d[f"{tag}_mean"] for d in recs])
            print(f"    {label}: posterior mean below S_H for "
                  f"{100*(mean < S_HUBISZ).mean():.2f}% of windows, p025 below "
                  f"S_H for {100*(p025 < S_HUBISZ).mean():.2f}%, "
                  f"genome minimum mean {mean.min():.0f} ka")
        nm = np.array([d["nea_mean"] for d in recs])
        near = ((nm >= 575) & (nm < 600)).sum()
        far = ((nm >= 700) & (nm < 800)).sum()
        print(f"  Density either side of the floor: {near:,} windows in "
              f"575-600 ka against {far:,} in 700-800 ka, i.e. the "
              f"distribution stops at the split rather than passing through "
              f"it. The floor binds; truncation is the right operation.")

        print("\n" + "-" * 78)
        print(f"Genome-wide context (random {len(sub):,} autosomal windows)")
        print("-" * 78)
        for tag, label in (("nea", "T(Afr+Nea)"), ("den", "T(Afr+Den)")):
            h, p, sh = [], [], []
            for d in sub:
                hm, _ = hubisz_stats(coord, to_t, d[f"{tag}_p025"],
                                     d[f"{tag}_p975"], ps=(0.5,))
                pm, _ = recalibrate(coord, to_t, d[f"{tag}_p025"],
                                    d[f"{tag}_p975"], S_PEYREGNE["mid"],
                                    ps=(0.5,))
                h.append(hm); p.append(pm); sh.append(pm - hm)
            full = S_PEYREGNE["mid"] - S_HUBISZ
            print(f"  {label}: median {med(h):.0f} -> {med(p):.0f} ka, "
                  f"realised shift {med(sh):+.0f} ka "
                  f"({100*med(sh)/full:.0f}% of full)")
        print("  Catalog windows shift more than genome-wide ones because the")
        print("  rule selects windows sitting on the prior floor.")

    # -- outputs ------------------------------------------------------------
    if args.emit:
        cols = ["chrom", "pos"]
        for tag in ("nea", "den"):
            cols += [f"{tag}_mean_stored", f"{tag}_mean_fitted",
                     f"{tag}_mean_hubisz", f"{tag}_p50_hubisz",
                     f"{tag}_subfloor"]
            for k in ("low", "mid", "high"):
                cols += [f"{tag}_recal_{k}", f"{tag}_shift_{k}",
                         f"{tag}_mean_peyregne_{k}", f"{tag}_p50_peyregne_{k}"]
            cols += [f"{tag}_p025_peyregne_mid", f"{tag}_p975_peyregne_mid",
                     f"{tag}_shift_mid", f"{tag}_shift_vs_stored"]
        cols.append("afr_mean_kya")
        with open(args.emit, "w", encoding="utf-8", newline="") as f:
            w = csv.writer(f, delimiter="\t")
            w.writerow(cols)
            for d in out:
                w.writerow([d["chrom"], d["pos"]] +
                           [(round(d[c], 1) if isinstance(d[c], float) else d[c])
                            for c in cols[2:]])
        print(f"\nwrote {args.emit}  ({len(out)} sites, {len(cols)} columns)")

    if args.figure:
        make_figure(args.figure, coord, to_t, out, grid, args.threshold)
        print(f"wrote {args.figure}")
    return 0


def make_figure(path, coord, to_t, out, grid, threshold):
    import matplotlib
    matplotlib.use("Agg")
    import matplotlib.pyplot as plt

    fig, axes = plt.subplots(1, 3, figsize=(15.0, 4.6))
    full = S_PEYREGNE["mid"] - S_HUBISZ
    u_p = coord.to_u(S_PEYREGNE["mid"])

    # (a) the maps
    ax = axes[0]
    t = np.linspace(400, 2600, 400)
    ax.plot(t, t, color="0.6", lw=1.0, ls=":", label="identity (no change)")
    ax.plot(t, t * S_PEYREGNE["mid"] / S_HUBISZ, color="#c0392b", lw=1.4,
            ls="--", label="multiplicative $S_P/S_H$ (wrong)")
    ax.plot(t, np.where(t > S_HUBISZ, t + full, t), color="#2980b9", lw=1.4,
            ls="-.", label=f"additive +{full:.0f} ka (note 3c)")
    for wgu, sty in ((0.5, "-"), (1.5, "-"), (3.0, "-")):
        sg = wgu / (2 * 1.959964)
        y = [trunc_stats(to_t, coord.to_u(x), sg, u_p, ps=(0.5,))[0] for x in t]
        ax.plot(t, y, sty, lw=2.0, alpha=0.85,
                label=f"truncation, 95% CI = {wgu} grid steps")
    ax.axvline(S_HUBISZ, color="k", lw=0.7, alpha=0.4)
    ax.axhline(S_PEYREGNE["mid"], color="k", lw=0.7, alpha=0.4)
    ax.set_xlabel("TMRCA on the Hubisz 2020 scale (ka)")
    ax.set_ylabel("TMRCA on the Peyregne 2025 scale (ka)")
    ax.set_title("a. Three candidate recalibration maps", loc="left",
                 fontsize=10, fontweight="bold")
    ax.legend(fontsize=6.5, loc="upper left")

    # (b) realised shift, per catalog window
    ax = axes[1]
    for tag, col, label in (("nea", "#1f77b4", "T(Afr+Nea)"),
                            ("den", "#d62728", "T(Afr+Den)")):
        x = [d[f"{tag}_mean_stored"] for d in out]
        y = [d[f"{tag}_shift_mid"] for d in out]
        ax.scatter(x, y, s=14, alpha=0.55, color=col, edgecolor="none",
                   label=label)
    ax.axhline(full, color="#2980b9", lw=1.2, ls="-.",
               label=f"full additive shift (+{full:.0f} ka)")
    ax.axhline(0, color="0.6", lw=1.0, ls=":", label="no shift")
    ax.set_xscale("log")
    ax.set_xlim(400, 5000)
    ax.set_xlabel("TMRCA on the Hubisz scale (ka, log)")
    ax.set_ylabel("realised shift (ka)")
    ax.set_title("b. The correction is self-limiting with depth", loc="left",
                 fontsize=10, fontweight="bold")
    ax.legend(fontsize=7, loc="center right")

    # (c) the catalog's defining contrast, both scales
    ax = axes[2]
    pairs = [("Hubisz", "nea_mean_stored", "den_mean_stored", "0.45"),
             ("Peyregne mid", "nea_recal_mid", "den_recal_mid", "#8e44ad")]
    for i, (label, nk, dk, col) in enumerate(pairs):
        n = [d[nk] for d in out]
        dd = [d[dk] for d in out]
        ax.boxplot([n, dd], positions=[i * 3 + 0.6, i * 3 + 1.4], widths=0.6,
                   patch_artist=True, showfliers=False,
                   boxprops=dict(facecolor=col, alpha=0.5),
                   medianprops=dict(color="k", lw=1.4))
    ax.set_yscale("log")
    ax.set_xticks([0.6, 1.4, 3.6, 4.4])
    ax.set_xticklabels(["Nea\nHubisz", "Den\nHubisz", "Nea\nPeyregne",
                        "Den\nPeyregne"], fontsize=8)
    ax.set_ylabel("TMRCA (ka, log)")
    ax.set_title("c. Nea/Den contrast compresses under recalibration",
                 loc="left", fontsize=10, fontweight="bold")

    fig.tight_layout()
    fig.savefig(path, dpi=170)


if __name__ == "__main__":
    raise SystemExit(main())
```
