## Supplementary Information section 4 code base for "Hypothesis: A modern human range expansion ∼300,000 years ago had a critical role in Neandertal origins": relate_locus_percentiles.py

```
#!/usr/bin/env python3
"""Per-locus Relate modern-TMRCA percentiles, for Table S4.1.

For each catalog locus this reports where its Relate modern TMRCA sits in the
genome-wide distribution of the same quantity. A low percentile means Relate
independently finds the locus recent, which is what a real sweep in the modern
lineage should look like.

Two choices worth stating because they change the number:

  * The locus value is the MEAN over its 5-kb grid points, to parallel the
    "mean TAfr" columns already in the table. The alternative, the minimum,
    rewards long loci purely for having more chances to contain one recent
    point, which would make the column a function of span.

  * The reference distribution is the genome-wide set of locus-sized BLOCK
    MEANS, not of individual grid points. Averaging over a locus shrinks
    variance, so scoring a locus mean against single-point spread would push
    every percentile toward the middle and understate the signal. Blocks are
    drawn at the same span as the locus being scored, which removes that
    mismatch. Adjacent grid points are strongly autocorrelated, so contiguous
    blocks also carry the right correlation structure.
"""

import argparse
import csv
import glob
import sys

import numpy as np


PANELS = {
    "yri216": ("yri_01_modern_tmrca_years", "YRI, 216 haplotypes"),
    "ygc604": ("ygc_01_modern_tmrca_years", "YRI+GBR+CHB, 604 haplotypes"),
}


def load(paths, cols):
    """chrom -> (sorted positions, {col: values in ka})."""
    per = {}
    for p in sorted(paths):
        with open(p, encoding="utf-8") as f:
            rd = csv.DictReader(f, delimiter="\t")
            miss = [c for c in cols if c not in rd.fieldnames]
            if miss:
                sys.exit(f"{p}: missing {miss}\n  has: {rd.fieldnames}")
            for r in rd:
                c = r["chrom"].replace("chr", "")
                if c in ("X", "Y", "23", "24"):
                    continue
                vals = [r[c2] for c2 in cols]
                if any(v in ("", "NA") for v in vals):
                    continue
                per.setdefault(int(c), []).append(
                    (int(r["pos"]), [float(v) / 1000.0 for v in vals]))
    out = {}
    for c, rows in per.items():
        rows.sort()
        pos = np.array([r[0] for r in rows])
        arr = np.array([r[1] for r in rows])
        out[c] = (pos, {cols[i]: arr[:, i] for i in range(len(cols))})
    n = sum(len(v[0]) for v in out.values())
    sys.stderr.write(f"loaded {n:,} grid points on {len(out)} autosomes\n")
    return out


def main():
    ap = argparse.ArgumentParser()
    ap.add_argument("--grids", default="relate_archaic_1kb/chr*.tsv")
    ap.add_argument("--loci", default="catalog_annotation/table_s51_loci.tsv")
    ap.add_argument("--perms", type=int, default=4000)
    ap.add_argument("--seed", type=int, default=11)
    ap.add_argument("-o", "--out",
                    default="catalog_annotation/table_s51_relate_pct.tsv")
    args = ap.parse_args()

    cols = [PANELS[k][0] for k in ("yri216", "ygc604")]
    data = load(sorted(glob.glob(args.grids)), cols)

    # one flat array per panel for block sampling
    flat = {c: np.concatenate([data[k][1][c] for k in sorted(data)])
            for c in cols}

    loci = []
    with open(args.loci, encoding="utf-8") as f:
        for r in csv.DictReader(f, delimiter="\t"):
            c = int(r["chr"])
            # single-grid-point loci read "16.815"; longer ones read
            # "35.765-35.775", the first and last grid point starts. Span is
            # carried separately, so only the left edge is needed here.
            start = int(round(float(r["interval (Mb)"].split("-")[0]) * 1e6))
            span = int(r["span (kb)"]) * 1000
            loci.append((c, start, span, r))
    sys.stderr.write(f"{len(loci)} loci\n")

    rng = np.random.default_rng(args.seed)
    # cache the null per (panel, n_points) since span takes few distinct values
    cache = {}

    rows = []
    for c, start, span, src in loci:
        pos, vals = data[c]
        lo = np.searchsorted(pos, start, "left")
        hi = np.searchsorted(pos, start + span, "left")
        rec = {"chr": c, "interval (Mb)": src["interval (Mb)"],
               "span (kb)": src["span (kb)"], "n_grid": int(hi - lo)}
        for key in ("yri216", "ygc604"):
            col = PANELS[key][0]
            if hi <= lo:
                rec[f"{key}_ka"] = ""
                rec[f"{key}_pct"] = ""
                continue
            m = float(vals[col][lo:hi].mean())
            k = (col, hi - lo)
            if k not in cache:
                v = flat[col]
                n = hi - lo
                st = rng.integers(0, len(v) - n, size=args.perms)
                cache[k] = np.sort(np.array(
                    [v[s:s + n].mean() for s in st]))
            null = cache[k]
            pct = float(np.searchsorted(null, m, "right") / len(null))
            rec[f"{key}_ka"] = f"{m:.0f}"
            rec[f"{key}_pct"] = f"{100 * pct:.0f}"
        rows.append(rec)

    with open(args.out, "w", newline="", encoding="utf-8") as f:
        w = csv.DictWriter(f, delimiter="\t", fieldnames=list(rows[0]))
        w.writeheader()
        w.writerows(rows)
    sys.stderr.write(f"wrote {args.out}\n")

    for key in ("yri216", "ygc604"):
        p = np.array([float(r[f"{key}_pct"]) for r in rows
                      if r[f"{key}_pct"] != ""])
        print(f"\n{PANELS[key][1]}")
        print(f"  loci scored           {len(p)}")
        print(f"  median percentile     {np.median(p):.0f}  (50 = no signal)")
        print(f"  loci below 10th pct   {int((p < 10).sum())} "
              f"({100 * (p < 10).mean():.0f}%)")
        print(f"  loci below 25th pct   {int((p < 25).sum())} "
              f"({100 * (p < 25).mean():.0f}%)")
        print(f"  loci above 50th pct   {int((p > 50).sum())} "
              f"({100 * (p > 50).mean():.0f}%)")
    return 0


if __name__ == "__main__":
    sys.exit(main())
```
