## Supplementary Information section 4 code base for "Hypothesis: A modern human range expansion ∼300,000 years ago had a critical role in Neandertal origins": relate_marginal_tmrca_grid.py

```
#!/usr/bin/env python3
"""
Marginal TMRCA on a regular grid from Relate's Newick export (RelateExtract
--mode AncToNewick), for the full sample or an arbitrary subset of tips.

No third-party dependencies (the previous version needed biopython; that was
both a cluster-environment failure mode and far too slow for 5,008-tip trees).

Inputs:
  --pos     .pos file (starting bp of each tree, one per line)
  --newick  .newick file (one tree per line, aligned with .pos)
  Trees are piecewise constant between consecutive positions. Branch lengths
  are in generations (Relate convention).

Full-sample TMRCA (default): height of the tree = max root-to-tip depth.

Subset TMRCA (optional): give --poplabels (Relate .poplabels: sample ID,
population, group, sex; one line per diploid individual, tips 2i and 2i+1
belong to individual i) and --subset-pops (comma-separated population and/or
group labels, e.g. AFR or YRI,LWK,GWD,MSL,ESN or ALL). TMRCA of the subset =
tree height minus depth of the most recent common ancestor of the subset tips.

Only trees hit by a grid point are parsed (lazy), which typically cuts work
by a large factor relative to parsing every tree.

Output TSV columns follow the ARGweaver scan convention:
  chrom, pos, <prefix>_mean_kya, <prefix>_p025_kya, <prefix>_p975_kya, <prefix>_n
Relate gives a single point estimate per tree, so mean = p025 = p975 and n=1.

Example (full sample, all modern humans):
  python3 relate_marginal_tmrca_grid.py \
    --pos chr1_extract.pos --newick chr1_extract.newick \
    --chrom 1 --chrom-len 249250621 --grid-step 5000 \
    --years-per-gen 28 -o chr1_tmrca_all_samples_5kb.tsv

Example (African haplotypes only):
  ... --poplabels 1000GP_Phase3.poplabels --subset-pops AFR \
      --column-prefix relate_afr_tmrca
"""

from __future__ import annotations

import argparse
import bisect
import gzip
import sys
from typing import List, Optional, TextIO, Tuple


def _open_text(path: str) -> TextIO:
    if path.endswith(".gz"):
        return gzip.open(path, "rt", encoding="utf-8", errors="replace")
    return open(path, encoding="utf-8", errors="replace")


def parse_newick(s: str) -> Tuple[List[int], List[float], List[str]]:
    """One-pass Newick parse.

    Returns (parent, blen, tip_label) over nodes indexed in creation order:
      parent[i]  index of parent (-1 for root)
      blen[i]    branch length above node i (0.0 if absent)
      tip_label[i]  label for tips, "" for internal nodes
    """
    parent: List[int] = []
    blen: List[float] = []
    label: List[str] = []

    def new_node(par: int) -> int:
        parent.append(par)
        blen.append(0.0)
        label.append("")
        return len(parent) - 1

    s = s.strip()
    if s.endswith(";"):
        s = s[:-1]
    root = new_node(-1)
    cur = root
    i, n = 0, len(s)
    while i < n:
        c = s[i]
        if c == "(":
            cur = new_node(cur)
            i += 1
        elif c == ",":
            cur = new_node(parent[cur])
            i += 1
        elif c == ")":
            cur = parent[cur]
            i += 1
        elif c == ":":
            j = i + 1
            while j < n and s[j] not in ",():;":
                j += 1
            blen[cur] = float(s[i + 1 : j])
            i = j
        else:  # label (tip name, or internal node label — ignored for internals)
            j = i
            while j < n and s[j] not in ",():;":
                j += 1
            if not label[cur]:
                label[cur] = s[i:j].strip()
            i = j
    return parent, blen, label


def node_depths(parent: List[int], blen: List[float]) -> List[float]:
    """Depth from root for every node. Nodes are created parent-before-child,
    so a single forward pass works."""
    depth = [0.0] * len(parent)
    for i in range(1, len(parent)):
        depth[i] = depth[parent[i]] + blen[i]
    return depth


def tree_height(parent: List[int], blen: List[float], label: List[str],
                depth: List[float]) -> Optional[float]:
    tips = [i for i in range(len(parent)) if label[i]]
    if len(tips) < 2:
        # fall back: leaves = nodes with no children
        has_child = [False] * len(parent)
        for i in range(1, len(parent)):
            has_child[parent[i]] = True
        tips = [i for i in range(len(parent)) if not has_child[i]]
        if len(tips) < 2:
            return None
    return max(depth[t] for t in tips)


def subset_mrca_depth(parent: List[int], depth: List[float],
                      tips: List[int]) -> Optional[float]:
    """Depth of the MRCA of the given tip node indices (LCA by parent chains)."""
    if not tips:
        return None

    def ancestors(v: int) -> set:
        out = set()
        while v != -1:
            out.add(v)
            v = parent[v]
        return out

    anc = ancestors(tips[0])  # ancestors of current running MRCA
    mrca = tips[0]
    for t in tips[1:]:
        v = t
        while v not in anc:
            v = parent[v]
        if v != mrca:
            mrca = v
            anc = ancestors(mrca)
    return depth[mrca]


def load_subset_haplotypes(poplabels: str, pops: List[str]) -> Optional[set]:
    """Relate .poplabels: header then one line per diploid sample:
    ID POP GROUP SEX. Tip labels in AncToNewick trees are haplotype indices
    2i and 2i+1 for the i-th sample. Returns set of haplotype-label strings,
    or None if pops == ['ALL']."""
    if pops == ["ALL"]:
        return None
    want = set(p.upper() for p in pops)
    haps: set = set()
    with _open_text(poplabels) as f:
        header = f.readline()
        for i, line in enumerate(f):
            fields = line.split()
            if len(fields) < 3:
                continue
            pop, group = fields[1].upper(), fields[2].upper()
            if pop in want or group in want:
                haps.add(str(2 * i))
                haps.add(str(2 * i + 1))
    if not haps:
        sys.stderr.write("ERROR: no haplotypes matched --subset-pops\n")
        sys.exit(1)
    return haps


def generations_to_kya_int(gen: float, years_per_gen: float) -> int:
    return int(round(gen * years_per_gen / 1000.0))


def main() -> None:
    ap = argparse.ArgumentParser(
        description=__doc__, formatter_class=argparse.RawDescriptionHelpFormatter)
    ap.add_argument("--pos", required=True, help="RelateExtract .pos file")
    ap.add_argument("--newick", required=True, help="RelateExtract .newick file")
    ap.add_argument("--chrom", required=True)
    ap.add_argument("--chrom-len", type=int, required=True)
    ap.add_argument("--grid-step", type=int, default=5000)
    ap.add_argument("--years-per-gen", type=float, default=28.0)
    ap.add_argument("--poplabels", help="Relate .poplabels file (for subsets)")
    ap.add_argument("--subset-pops", default="ALL",
                    help="comma-separated POP/GROUP labels (e.g. AFR or YRI,LWK); "
                         "default ALL = full sample")
    ap.add_argument("--column-prefix", default="relate_full_sample_tmrca",
                    help="output column prefix")
    ap.add_argument("--subset", action="append", default=[], metavar="PREFIX=POPS",
                    help="repeatable: compute an extra subset TMRCA in the SAME "
                         "pass, e.g. --subset relate_afr=AFR --subset relate_yri=YRI. "
                         "Parsing 5,008-tip trees dominates runtime, so folding "
                         "several subsets into one pass is much cheaper than "
                         "rerunning. Overrides --subset-pops/--column-prefix.")
    ap.add_argument("-o", "--out", required=True)
    args = ap.parse_args()

    # Each entry: (column_prefix, set-of-haplotype-labels or None for full sample)
    specs: List[Tuple[str, Optional[set]]] = []
    if args.subset:
        if not args.poplabels:
            sys.stderr.write("ERROR: --subset requires --poplabels\n")
            sys.exit(1)
        for spec in args.subset:
            if "=" not in spec:
                sys.stderr.write(f"ERROR: --subset needs PREFIX=POPS, got {spec!r}\n")
                sys.exit(1)
            pre, popstr = spec.split("=", 1)
            pre = pre.strip()
            pops = [p.strip() for p in popstr.split(",") if p.strip()]
            if not pre or not pops:
                sys.stderr.write(f"ERROR: malformed --subset {spec!r}\n")
                sys.exit(1)
            haps = load_subset_haplotypes(args.poplabels, pops)
            n = "all" if haps is None else len(haps)
            sys.stderr.write(f"subset {pre}: {n} haplotypes from {pops}\n")
            specs.append((pre, haps))
    else:
        pops = [p.strip() for p in args.subset_pops.split(",") if p.strip()]
        subset_haps: Optional[set] = None
        if pops != ["ALL"]:
            if not args.poplabels:
                sys.stderr.write("ERROR: --subset-pops requires --poplabels\n")
                sys.exit(1)
            subset_haps = load_subset_haplotypes(args.poplabels, pops)
            sys.stderr.write(f"subset: {len(subset_haps)} haplotypes from {pops}\n")
        specs.append((args.column_prefix, subset_haps))

    positions: List[int] = []
    with _open_text(args.pos) as fh:
        for line in fh:
            line = line.strip()
            if line and not line.startswith("#"):
                positions.append(int(line.split()[0]))
    if not positions:
        sys.stderr.write("ERROR: empty pos file\n")
        sys.exit(1)

    # Which tree index does each grid point use?
    grid = list(range(0, args.chrom_len, args.grid_step))
    tree_idx: List[Optional[int]] = []
    needed = set()
    for p in grid:
        if p < positions[0]:
            tree_idx.append(None)
            continue
        j = bisect.bisect_right(positions, p) - 1
        tree_idx.append(j)
        needed.add(j)
    sys.stderr.write(
        f"{len(positions)} trees on file; {len(needed)} needed for "
        f"{len(grid)} grid points\n")

    # Single pass over the newick file; parse only needed lines. Every subset
    # is evaluated on each parsed tree, so cost is one parse regardless of
    # how many subsets were requested.
    tmrca_gen: List[dict] = [{} for _ in specs]
    with _open_text(args.newick) as fh:
        for j, line in enumerate(fh):
            if j not in needed:
                continue
            line = line.strip()
            if not line:
                continue
            try:
                parent, blen, label = parse_newick(line)
                depth = node_depths(parent, blen)
                height = tree_height(parent, blen, label, depth)
                if height is None:
                    continue
                for k_spec, (_, haps) in enumerate(specs):
                    if haps is None:
                        tmrca_gen[k_spec][j] = height
                        continue
                    tips = [i for i in range(len(parent)) if label[i] in haps]
                    if len(tips) < 2:
                        continue
                    d = subset_mrca_depth(parent, depth, tips)
                    if d is not None:
                        tmrca_gen[k_spec][j] = height - d
            except (ValueError, IndexError) as e:
                sys.stderr.write(f"WARN: bad tree at index {j}: {e}\n")

    header = ["chrom", "pos"]
    for pre, _ in specs:
        header += [f"{pre}_mean_kya", f"{pre}_p025_kya",
                   f"{pre}_p975_kya", f"{pre}_n"]
    with open(args.out, "w") as out:
        out.write("\t".join(header) + "\n")
        for p, j in zip(grid, tree_idx):
            row = [str(args.chrom), str(p)]
            for k_spec in range(len(specs)):
                g = tmrca_gen[k_spec].get(j) if j is not None else None
                if g is None:
                    row += ["NA", "NA", "NA", "0"]
                else:
                    k = generations_to_kya_int(g, args.years_per_gen)
                    row += [str(k), str(k), str(k), "1"]
            out.write("\t".join(row) + "\n")

    what = ", ".join(pre for pre, _ in specs)
    sys.stderr.write(
        f"Wrote {args.out} (years_per_gen={args.years_per_gen}, "
        f"grid_step={args.grid_step}, subsets=[{what}])\n")


if __name__ == "__main__":
    main()
```
