## Supplementary Information section 4 code base for "Hypothesis: A modern human range expansion ∼300,000 years ago had a critical role in Neandertal origins": relate_modern_vs_argweaver.py

```
#!/usr/bin/env python3
"""Do Relate's modern-only TMRCAs recognise the ARGweaver sweep catalog?

The catalog was built from ARGweaver-D's archaic contrast: the modern-plus-
Neandertal root shallow, the modern-plus-Denisova root deep. Nothing in that
rule looks at the modern-only root. But a sweep that carried a haplotype
through both moderns and Neandertals has to have carried it through moderns,
so the modern-only TMRCA at a catalog locus should be recent. Relate infers
that quantity from the same 1000 Genomes haplotypes by a completely different
algorithm, which makes it a check on the catalog rather than a restatement of
it.

Two things are measured, and they are not the same question:

  1. Concordance. Across the whole genome, do the two methods agree about
     where the modern root is recent? This is the calibration question, and it
     is inflated by covariates both methods respond to -- accessibility, local
     recombination rate, mutation density -- so a high value is necessary but
     not sufficient.

  2. Recognition. At the catalog's grid points specifically, is Relate's
     modern TMRCA unusually recent relative to the rest of the genome? This is
     scored as a percentile rank against the genome-wide Relate distribution,
     so it does not depend on the two methods sharing a timescale. Under the
     null the mean percentile is 0.5.

The per-locus version takes the minimum over each locus's grid points, which
is biased downward by construction, so the null is built by permutation over
random position-matched blocks of the same grid-point count rather than
assumed.
"""

import argparse
import csv
import glob
import os
import sys

import numpy as np


def load_argweaver(path):
    """chrom, pos -> (modern-only mean ka, modern+Nea mean ka)."""
    out = {}
    with open(path, encoding="utf-8") as f:
        rd = csv.DictReader(f, delimiter="\t")
        for r in rd:
            c = r["chrom"].replace("chr", "")
            if c in ("X", "Y", "23", "24"):
                continue
            a = r["01_modern_only_mean_kya"]
            b = r["02_modern_plus_both_Nea_mean_kya"]
            if a in ("", "NA") or b in ("", "NA"):
                continue
            out[(int(c), int(r["pos"]))] = (float(a), float(b))
    sys.stderr.write(f"ARGweaver: {len(out):,} autosomal grid points\n")
    return out


def load_relate(paths, cols):
    """chrom, pos -> tuple of ka values for the requested columns."""
    out, have = {}, None
    for p in sorted(paths):
        with open(p, encoding="utf-8") as f:
            rd = csv.DictReader(f, delimiter="\t")
            if have is None:
                have = [c for c in cols if c in rd.fieldnames]
                missing = [c for c in cols if c not in rd.fieldnames]
                if missing:
                    sys.stderr.write(f"  note: absent here: "
                                     f"{', '.join(missing)}\n")
                if not have:
                    sys.exit(f"{p}: none of the requested columns exist\n"
                             f"  available: {', '.join(rd.fieldnames)}")
            for r in rd:
                c = r["chrom"].replace("chr", "")
                if c in ("X", "Y", "23", "24"):
                    continue
                vals = [r[c2] for c2 in have]
                if any(v in ("", "NA") for v in vals):
                    continue
                out[(int(c), int(r["pos"]))] = tuple(
                    float(v) / 1000.0 for v in vals)
    sys.stderr.write(f"Relate: {len(out):,} autosomal grid points "
                     f"[{', '.join(have)}]\n")
    return out, have


def spearman(x, y):
    """Rank correlation, ties averaged, without pulling in scipy."""
    def rank(v):
        order = np.argsort(v, kind="mergesort")
        r = np.empty(len(v), float)
        r[order] = np.arange(len(v), dtype=float)
        # average ties
        sv = v[order]
        i = 0
        while i < len(sv):
            j = i
            while j + 1 < len(sv) and sv[j + 1] == sv[i]:
                j += 1
            if j > i:
                r[order[i:j + 1]] = (i + j) / 2.0
            i = j + 1
        return r
    rx, ry = rank(x), rank(y)
    rx -= rx.mean()
    ry -= ry.mean()
    d = np.sqrt((rx * rx).sum() * (ry * ry).sum())
    return float((rx * ry).sum() / d) if d else float("nan")


def pearson(x, y):
    x = x - x.mean()
    y = y - y.mean()
    d = np.sqrt((x * x).sum() * (y * y).sum())
    return float((x * y).sum() / d) if d else float("nan")


def mannwhitney_z(a, b):
    """Normal approximation to the rank-sum statistic, tie-corrected."""
    n1, n2 = len(a), len(b)
    v = np.concatenate([a, b])
    order = np.argsort(v, kind="mergesort")
    r = np.empty(len(v), float)
    r[order] = np.arange(1, len(v) + 1, dtype=float)
    sv = v[order]
    i, tie = 0, 0.0
    while i < len(sv):
        j = i
        while j + 1 < len(sv) and sv[j + 1] == sv[i]:
            j += 1
        if j > i:
            r[order[i:j + 1]] = (i + j + 2) / 2.0
            t = j - i + 1
            tie += t ** 3 - t
        i = j + 1
    u = r[:n1].sum() - n1 * (n1 + 1) / 2.0
    mu = n1 * n2 / 2.0
    n = n1 + n2
    sd = np.sqrt(n1 * n2 / 12.0 * ((n + 1) - tie / (n * (n - 1))))
    return float((u - mu) / sd) if sd else float("nan")


def two_sided_p(z):
    """Survival function of |z| under the normal, via erfc."""
    import math
    return math.erfc(abs(z) / math.sqrt(2.0))


def percentile_of(values, ref_sorted):
    """Fraction of ref at or below each value, vectorised."""
    return np.searchsorted(ref_sorted, values, side="right") / len(ref_sorted)


def main():
    ap = argparse.ArgumentParser()
    ap.add_argument("--argweaver", default="hubisz_tmrca_5kb_8afr.tsv")
    ap.add_argument("--relate-glob", default="relate_archaic/chr*.tsv")
    ap.add_argument("--relate-cols",
                    default="yri_01_modern_tmrca_years,"
                            "ygc_01_modern_tmrca_years")
    ap.add_argument("--catalog-sites",
                    default="catalog_annotation/"
                            "catalog_final_median_x625_y700_auto.sites.tsv")
    ap.add_argument("--catalog-regions",
                    default="catalog_annotation/"
                            "catalog_final_median_x625_y700_auto.regions.tsv")
    ap.add_argument("--perms", type=int, default=5000)
    ap.add_argument("--seed", type=int, default=17)
    args = ap.parse_args()

    aw = load_argweaver(args.argweaver)
    paths = sorted(glob.glob(args.relate_glob))
    if not paths:
        sys.exit(f"no Relate grids matched {args.relate_glob}")
    cols = args.relate_cols.split(",")
    rel, have = load_relate(paths, cols)

    keys = sorted(set(aw) & set(rel))
    if not keys:
        sys.exit("the two grids do not share any positions")
    sys.stderr.write(f"joined on {len(keys):,} shared grid points "
                     f"({100 * len(keys) / len(aw):.1f}% of ARGweaver)\n\n")

    aw_mod = np.array([aw[k][0] for k in keys])
    aw_nea = np.array([aw[k][1] for k in keys])
    R = {c: np.array([rel[k][i] for k in keys]) for i, c in enumerate(have)}
    index = {k: i for i, k in enumerate(keys)}

    print("=" * 74)
    print("1. CONCORDANCE  genome-wide, on the shared 5-kb grid")
    print("=" * 74)
    print(f"{'Relate panel':<34} {'vs AW modern':>14} {'vs AW mod+Nea':>15}")
    for c in have:
        s1 = spearman(R[c], aw_mod)
        s2 = spearman(R[c], aw_nea)
        print(f"{c:<34} {s1:>14.3f} {s2:>15.3f}")
    print("\nSpearman. For scale, the two ARGweaver columns correlate with")
    print(f"each other at {spearman(aw_mod, aw_nea):.3f}, which is the "
          f"ceiling a perfect")
    print("replication of the same quantity could reach here.")
    print("\nmedian TMRCA, as a reminder that the timescales differ:")
    print(f"  ARGweaver modern-only   {np.median(aw_mod):>7.0f} ka")
    print(f"  ARGweaver modern+Nea    {np.median(aw_nea):>7.0f} ka")
    for c in have:
        print(f"  {c:<22}  {np.median(R[c]):>7.0f} ka")

    # ---- catalog grid points
    cat = []
    with open(args.catalog_sites, encoding="utf-8") as f:
        for r in csv.DictReader(f, delimiter="\t"):
            k = (int(r["chrom"].replace("chr", "")), int(r["pos"]))
            if k in index:
                cat.append(index[k])
    cat = np.array(sorted(set(cat)))
    print()
    print("=" * 74)
    print("2. RECOGNITION  are catalog grid points recent in Relate?")
    print("=" * 74)
    print(f"{len(cat):,} of the catalog's grid points survive the join\n")
    print(f"{'panel':<34} {'cat median':>11} {'bg median':>10} "
          f"{'mean pct':>9} {'z':>7} {'p':>10}")
    bg_mask = np.ones(len(keys), bool)
    bg_mask[cat] = False
    for c in list(have) + ["ARGWEAVER_modern_only"]:
        v = aw_mod if c.startswith("ARGWEAVER") else R[c]
        ref = np.sort(v[bg_mask])
        pct = percentile_of(v[cat], ref)
        z = mannwhitney_z(v[cat], v[bg_mask])
        print(f"{c:<34} {np.median(v[cat]):>9.0f} ka "
              f"{np.median(v[bg_mask]):>7.0f} ka "
              f"{pct.mean():>9.3f} {z:>7.1f} {two_sided_p(z):>10.2e}")
    print("\nmean pct is the average percentile of catalog grid points in the")
    print("genome-wide distribution of that column; 0.5 is the null. The")
    print("ARGweaver row is the same test on the data the catalog came from,")
    print("so it is the internal benchmark, not an independent result.")

    # ---- per locus, permutation null on the minimum
    regions = []
    with open(args.catalog_regions, encoding="utf-8") as f:
        for r in csv.DictReader(f, delimiter="\t"):
            c = int(r["chrom"].replace("chr", ""))
            idx = [index[(c, p)]
                   for p in range(int(r["start"]), int(r["end"]) + 1, 5000)
                   if (c, p) in index]
            if idx:
                regions.append(idx)
    print()
    print("=" * 74)
    print("3. PER LOCUS  minimum over each locus, permutation null")
    print("=" * 74)
    print(f"{len(regions)} of 132 loci have at least one joined grid point")
    sizes = [len(r) for r in regions]
    rng = np.random.default_rng(args.seed)
    n = len(keys)
    print(f"\n{'panel':<34} {'obs median of minima':>21} "
          f"{'null median':>12} {'p':>9}")
    for c in list(have) + ["ARGWEAVER_modern_only"]:
        v = aw_mod if c.startswith("ARGWEAVER") else R[c]
        obs = np.median([v[np.array(r)].min() for r in regions])
        null = np.empty(args.perms)
        for b in range(args.perms):
            mins = np.empty(len(sizes))
            for i, s in enumerate(sizes):
                st = rng.integers(0, n - s)
                mins[i] = v[st:st + s].min()
            null[b] = np.median(mins)
        p = (np.sum(null <= obs) + 1) / (args.perms + 1)
        print(f"{c:<34} {obs:>19.0f} ka {np.median(null):>10.0f} ka "
              f"{p:>9.4f}")
    print("\nBlocks are drawn contiguously so each draw carries the same")
    print("within-locus autocorrelation as the real loci. p is one-sided:")
    print("the chance a random block set is at least this recent.")
    return 0


if __name__ == "__main__":
    sys.exit(main())
```
