## Supplementary Information section 4 code base for "Hypothesis: A modern human range expansion ∼300,000 years ago had a critical role in Neandertal origins": relate_ts_tmrca_grid.py

```
#!/usr/bin/env python3
"""Marginal TMRCA on a fixed bp grid from a Relate tree sequence.

Replaces the AncToNewick text route, which did not scale: exporting every
marginal tree as newick spent 11 h on chr22 -- the smallest autosome -- and was
killed before finishing, having written 1.3 GB for ~8,600 trees. The tree
sequence holds the same genealogies in a form built for positional queries.

Two ideas make this fast:

  1. simplify() down to the haplotypes of interest first. After simplification
     the TMRCA of that subset at a position is just the root time of the local
     tree, so we never search for an MRCA across 1,322 samples per position.

  2. Trees carry an interval, so one linear pass assigns every grid point in
     [left, right) at once, instead of seeking per grid point.

Usage:
  python3 relate_ts_tmrca_grid.py --trees chr22.trees --chrom 22 \\
      --chrom-len 51304566 --poplabels x.poplabels \\
      --subset relate_afr1322=AFR --subset relate_yri216=YRI \\
      --grid-step 5000 --years-per-gen 28 -o chr22_tmrca_5kb.tsv
"""

from __future__ import annotations

import argparse
import csv
import sys


def load_poplabels(path):
    """Return (pop_of_sample_row, groups) for the poplabels file.

    Relate poplabels are one row per *individual*: sample, population, group,
    sex. A diploid individual contributes two haplotypes, and the tree sequence
    orders them consecutively, so individual i owns nodes 2i and 2i+1.
    """
    rows = []
    with open(path, encoding="utf-8") as f:
        header = f.readline().split()
        if header and header[0].upper() not in ("SAMPLE", "ID"):
            f.seek(0)                      # no header after all
        for line in f:
            p = line.split()
            if len(p) >= 3:
                rows.append((p[0], p[1], p[2]))
    return rows


def subset_nodes(rows, want, ts_num_samples):
    """Sample node ids for a comma-separated union of labels or sample ids.

    Each token matches a population label ('YRI'), a group label ('AFR'), an
    individual's sample id ('AltaiNeandertal'), or 'ALL'. Unions are the point:
    the ARGweaver scan's statistics are roots of *combined* sets -- moderns plus
    both Neandertals, moderns plus the Denisovan -- and no single label names
    such a set, so replicating those columns needs 'YRI,GBR,CHB,Altai,Vindija'
    to be expressible as one subset.

    Returns (nodes, unmatched). Unmatched tokens are handed back rather than
    ignored because the failure they cause is silent and severe: misspell an
    archaic's sample id and the subset quietly becomes moderns-only, which
    still produces a full column of plausible root times that no longer mean
    what the column name claims.
    """
    toks = [t.strip().upper() for t in want.split(",") if t.strip()]
    nodes, seen, matched = [], set(), set()
    for i, (sample, pop, grp) in enumerate(rows):
        keys = {sample.upper(), pop.upper(), grp.upper()}
        hit = [t for t in toks if t == "ALL" or t in keys]
        if not hit:
            continue
        matched.update(hit)
        for n in (2 * i, 2 * i + 1):
            if n < ts_num_samples and n not in seen:
                seen.add(n)
                nodes.append(n)
    return nodes, [t for t in toks if t not in matched and t != "ALL"]


def main():
    ap = argparse.ArgumentParser()
    ap.add_argument("--trees", required=True)
    ap.add_argument("--chrom", required=True)
    ap.add_argument("--chrom-len", type=int, required=True)
    ap.add_argument("--poplabels", required=True)
    ap.add_argument("--subset", action="append", default=[],
                    help="PREFIX=POP, repeatable, e.g. relate_yri216=YRI")
    ap.add_argument("--grid-step", type=int, default=5000)
    ap.add_argument("--years-per-gen", type=float, default=28.0)
    ap.add_argument("-o", "--out", required=True)
    args = ap.parse_args()

    try:
        import tskit
    except ImportError:
        sys.exit("tskit not installed: pip install tskit")

    ts = tskit.load(args.trees)
    rows = load_poplabels(args.poplabels)
    sys.stderr.write(f"tree sequence: {ts.num_samples} samples, "
                     f"{ts.num_trees} trees, {ts.sequence_length:.0f} bp\n")
    sys.stderr.write(f"poplabels: {len(rows)} individuals "
                     f"({2 * len(rows)} haplotypes)\n")
    if 2 * len(rows) != ts.num_samples:
        sys.stderr.write("WARNING: poplabels haplotype count does not match the "
                         "tree sequence sample count; subset node ids may be "
                         "misaligned. Check that this poplabels file belongs to "
                         "this .trees file.\n")

    ngrid = args.chrom_len // args.grid_step + 1
    specs = [s.split("=", 1) for s in args.subset] or [["relate_all", "ALL"]]

    results = {}
    bad = []
    for prefix, pop in specs:
        nodes, unmatched = subset_nodes(rows, pop, ts.num_samples)
        if unmatched:
            # Fatal, not a warning. An unmatched token means the subset is not
            # the set the column name claims -- typically an archaic silently
            # dropped, leaving a full column of modern-only root times that
            # look entirely reasonable and are wrong.
            bad.append(f"{prefix}: no sample, population or group named "
                       f"{', '.join(unmatched)}")
            continue
        if not nodes:
            bad.append(f"{prefix}: '{pop}' matched nothing")
            continue
        sys.stderr.write(f"  {prefix}: {len(nodes)} haplotypes for '{pop}' ... ")
        sys.stderr.flush()

        # Do NOT simplify. Relate's ConvertToTreeSequence stores every tree
        # independently with no edge sharing between adjacent trees, so chr22
        # alone is 31,332 trees x ~10k edges = ~313M edges in a 13.8 GB file.
        # Rewriting that edge table is what simplify() does, and it was killed
        # at the 48 GB cap.
        #
        # tracked_samples gets the same answer without touching the tables:
        # tskit maintains a per-node count of tracked samples in C as it moves
        # along the sequence, so the subset MRCA is found by walking up from any
        # one member until the count covers the whole subset. Memory is just
        # the loaded tree sequence; the walk is O(depth) per tree.
        want = len(nodes)
        first = nodes[0]
        gen = [""] * ngrid
        n_multi = 0
        for tree in ts.trees(tracked_samples=nodes):
            u = first
            while u != tskit.NULL and tree.num_tracked_samples(u) < want:
                u = tree.parent(u)
            if u == tskit.NULL:
                # Walked off the top without gathering the subset, i.e. it has
                # not coalesced in this interval. Recorded as missing rather
                # than as the deepest root, which would masquerade as a very
                # old TMRCA.
                n_multi += 1
                continue
            t = tree.time(u)
            left, right = tree.interval
            i0 = int(left) // args.grid_step
            i1 = min(int(right - 1) // args.grid_step, ngrid - 1)
            if int(left) % args.grid_step:
                i0 += 1
            for i in range(max(i0, 0), i1 + 1):
                gen[i] = t
        filled = sum(1 for g in gen if g != "")
        sys.stderr.write(f"{filled}/{ngrid} grid points"
                         + (f", {n_multi} uncoalesced trees" if n_multi else "")
                         + "\n")
        results[prefix] = gen

    if bad:
        sys.exit("subset specification errors:\n  " + "\n  ".join(bad)
                 + "\n(check the poplabels: this dataset spells the sample id "
                   "AltaiNeandertal and its population AltaiNeanderthal)")
    if not results:
        sys.exit("no subsets produced output")

    cols = list(results)
    with open(args.out, "w", newline="", encoding="utf-8") as f:
        w = csv.writer(f, delimiter="\t")
        head = ["chrom", "pos"]
        for c in cols:
            head += [f"{c}_tmrca_gen", f"{c}_tmrca_years"]
        w.writerow(head)
        for i in range(ngrid):
            row = [args.chrom, i * args.grid_step]
            for c in cols:
                g = results[c][i]
                row += ([f"{g:.2f}", f"{g * args.years_per_gen:.1f}"]
                        if g != "" else ["NA", "NA"])
            w.writerow(row)
    sys.stderr.write(f"wrote {args.out}\n")
    return 0


if __name__ == "__main__":
    raise SystemExit(main())
```
