## Supplementary Information section 4 code base for "Hypothesis: A modern human range expansion ∼300,000 years ago had a critical role in Neandertal origins": resolve_by_interval.py

```
#!/usr/bin/env python3
"""Resolve overlapping-ARG-window duplicates by keeping the tighter posterior.

ARGweaver-D was inferred in 2.2 Mb windows overlapping by 100 kb, so positions
in an overlap carry two independent reads. resolve_window_overlaps.py breaks the
tie by which window the position sits further inside. This is the alternative:
keep the read whose posterior is tighter at that position, on the grounds that
the window with more information about the local tree reports less spread.

Two practical advantages over the margin rule. It needs no window annotation,
so it runs on the collated scan as it stands rather than requiring the
per-window parts to be re-collated on the cluster. And it is a property of the
inference at that base rather than of the tiling geometry.

One design point matters. A position is resolved as a whole -- one window wins
and supplies all of its columns -- rather than per statistic. Picking the
tighter interval separately for each haplotype set would take TMRCAs from
different inferred genealogies and combine them, which can manufacture or
destroy the case/control contrast that P(real) is built from.

Width is the absolute range of the 95% central interval, hi-lo in ka, summed
over the three haplotype sets. An earlier version measured width in
ARGweaver-D's grid-index coordinate on the theory that the grid is log-spaced.
It is not: times2.txt is hand-tuned with 50 ka steps between 400 and 600 ka and
200-1000 ka steps elsewhere, so 300-600 ka crosses five grid cells while the
far wider 600-1300 ka crosses only four. That coordinate therefore measures how
much of the dense part of the grid an interval straddles rather than how
concentrated the posterior is, and it inverted the comparison at real loci.
Absolute range is the intended inference: the window that localizes the
coalescence to a narrower span of time is the better-informed one.

Usage:
  python3 scripts/resolve_by_interval.py -i hubisz_tmrca_5kb_8afr.tsv \
      -o hubisz_tmrca_5kb_8afr.tight.tsv
  python3 scripts/resolve_by_interval.py \
      -i catalog_annotation/hubisz_tmrca_sites_8afr.tsv \
      -o catalog_annotation/hubisz_tmrca_sites_8afr.tight.tsv --report 14:58862763
"""

import argparse
import csv
import math
import os
import sys

sys.path.insert(0, os.path.dirname(os.path.abspath(__file__)))
from approx_percentiles import COLS, GridCoord, true_grid  # noqa: E402

SETS = ("afr", "nea", "den")


def width(row, coord, tags, units="linear"):
    """Total width of the 95% central intervals at this position.

    units="linear" (default, and the rule used for the note) is the absolute
    range hi-lo in ka, summed over the sets.

    units="log" measures log10(hi/lo) and units="index" measures the span in
    ARGweaver-D's grid-index coordinate. Both are kept only for the sensitivity
    check that shows the choice is not free; see the module docstring for why
    the grid-index coordinate is misleading.
    """
    tot = 0.0
    for t in tags:
        pre = COLS[t]
        try:
            lo = float(row[f"{pre}_p025_kya"])
            hi = float(row[f"{pre}_p975_kya"])
        except (KeyError, ValueError):
            return None
        if units == "index":
            tot += abs(coord.to_u(hi) - coord.to_u(lo))
        elif units == "log":
            lo = max(lo, 1.0)
            hi = max(hi, lo + 1e-9)
            tot += math.log10(hi / lo)
        else:
            tot += abs(hi - lo)
    return tot


def main():
    ap = argparse.ArgumentParser()
    ap.add_argument("-i", "--infile", required=True)
    ap.add_argument("-o", "--out", required=True)
    ap.add_argument("--tags", default="afr,nea,den",
                    help="haplotype sets whose widths are summed")
    ap.add_argument("--times", default="times2.txt")
    ap.add_argument("--units", default="linear",
                    choices=("linear", "log", "index"),
                    help="how interval width is measured; linear = absolute ka")
    ap.add_argument("--report", default=None,
                    help="chrom:pos to describe in detail")
    a = ap.parse_args()

    tags = [t for t in a.tags.split(",") if t]
    for t in tags:
        if t not in COLS:
            raise SystemExit(f"unknown set {t!r}; choose from {list(COLS)}")
    coord = GridCoord(true_grid(a.times))

    with open(a.infile, encoding="utf-8") as fh:
        rdr = csv.DictReader(fh, delimiter="\t")
        cols = rdr.fieldnames
        best, order, n_in, skipped = {}, [], 0, 0
        first = {}
        for row in rdr:
            n_in += 1
            key = (row["chrom"], row["pos"])
            w = width(row, coord, tags, a.units)
            if w is None:
                skipped += 1
                continue
            if key not in best:
                order.append(key)
                best[key] = (w, row)
                first[key] = row
            elif w < best[key][0]:
                best[key] = (w, row)

    n_dup = n_in - len(best) - skipped
    changed = sum(1 for k in best if best[k][1] is not first[k])
    with open(a.out, "w", newline="", encoding="utf-8") as fo:
        wr = csv.DictWriter(fo, fieldnames=cols, delimiter="\t",
                            lineterminator="\n")
        wr.writeheader()
        for k in order:
            wr.writerow(best[k][1])

    sys.stderr.write(
        f"{a.infile}\n"
        f"  rows in                  {n_in:,}\n"
        f"  unique positions out     {len(best):,}\n"
        f"  duplicated positions     {n_dup:,}\n"
        f"  where tighter != first   {changed:,}"
        f"{'' if not n_dup else f'  ({100*changed/n_dup:.1f}% of duplicates)'}\n"
        + (f"  rows dropped (bad cols)  {skipped:,}\n" if skipped else "")
        + f"  width measured over      {'+'.join(tags)}"
          f" ({a.units}{' ka' if a.units == 'linear' else ''})\n")

    if a.report:
        c, p = a.report.split(":")
        print(f"\n=== chr{c}:{int(p):,} ===")
        rows = []
        with open(a.infile, encoding="utf-8") as fh:
            for row in csv.DictReader(fh, delimiter="\t"):
                if row["chrom"] == c and row["pos"] == p:
                    rows.append(row)
        for i, row in enumerate(rows, 1):
            parts = []
            for t in SETS:
                pre = COLS[t]
                lo = float(row[f"{pre}_p025_kya"])
                hi = float(row[f"{pre}_p975_kya"])
                parts.append(f"{t} {lo:.0f}-{hi:.0f} ka (w={hi - lo:.0f})")
            tot = width(row, coord, tags, a.units)
            print(f"  read {i}: " + "  ".join(parts))
            print(f"          summed {a.units} width over "
                  f"{'+'.join(tags)} = {tot:.2f}")
        w_all = [width(r, coord, tags, a.units) for r in rows]
        if len(rows) > 1:
            k = min(range(len(rows)), key=lambda i: w_all[i])
            print(f"  -> keeps read {k+1} (tighter)")
            for t in SETS:
                pre = COLS[t]
                print(f"       {t} mean {float(rows[k][f'{pre}_mean_kya']):.0f} ka")


if __name__ == "__main__":
    main()
```
