## Supplementary Information section 4 code base for "Hypothesis: A modern human range expansion ∼300,000 years ago had a critical role in Neandertal origins": run_great_100k.sh

```
#!/bin/bash
# 100,000 B-matched permutation replicates for the GO enrichment, as five
# independent 20,000-replicate chunks run in parallel. Permutation counts are
# additive over independent draws, so the chunks aggregate to a single null of
# 100,000 with a p value floor of 1/100001.
#
# Five workers rather than ten: each holds the GO annotations, the GENCODE
# domains, the P(real) grid and the B map, about 2 GB resident, and five is the
# most that comfortably fits alongside everything else.

set -euo pipefail
cd "$(dirname "$0")/.."

# Overridable so the same driver can be re-run on a rebuilt catalog without
# editing the file; the defaults reproduce the published run.
REGIONS="${REGIONS:-catalog_annotation/catalog_final_median_x625_y700_auto.regions.tsv}"
RESULTS="${RESULTS:-catalog_annotation/great_go_catalog132_matched10k.tsv}"
OUT="${OUT:-catalog_annotation/perm100k}"
FINAL="${FINAL:-catalog_annotation/great_go_catalog132_matched100k.tsv}"
CHUNKS="${CHUNKS:-5}"
NREP="${NREP:-20000}"

rm -rf "$OUT"
mkdir -p "$OUT"

echo "launching $CHUNKS chunks of $NREP replicates each"
pids=()
for i in $(seq 0 $((CHUNKS - 1))); do
    python3 scripts/great_matched_100k.py \
        --regions "$REGIONS" --results "$RESULTS" \
        --nrep "$NREP" --seed $((100 + i)) \
        -o "$OUT/counts.$i.tsv" > "$OUT/log.$i.txt" 2>&1 &
    pid=$!
    pids+=("$pid")
    echo "  chunk $i pid $pid seed $((100 + i))"
done

fail=0
for p in "${pids[@]}"; do
    if ! wait "$p"; then
        echo "chunk with pid $p FAILED"
        fail=1
    fi
done
if [ "$fail" -ne 0 ]; then
    tail -5 "$OUT"/log.*.txt
    exit 1
fi

echo "all chunks finished; aggregating"
python3 scripts/great_matched_100k.py \
    --aggregate "$OUT"/counts.*.tsv \
    --results "$RESULTS" \
    -o "$FINAL"
```
