## Supplementary Information section 4 code base for "Hypothesis: A modern human range expansion ∼300,000 years ago had a critical role in Neandertal origins": sbatch_extract_chagyrskaya.sh

```
#!/bin/bash
#SBATCH --job-name=chagex
#SBATCH --partition=short
#SBATCH --time=4:00:00
#SBATCH --mem=4G
#SBATCH --cpus-per-task=2
#SBATCH --array=1-22%8
#SBATCH --output=/n/groups/reich/DAVID/CURSOR_ENRICH/fixed_derived/logs/chagex_%A_%a.out
#SBATCH --error=/n/groups/reich/DAVID/CURSOR_ENRICH/fixed_derived/logs/chagex_%A_%a.err
#
# Make Chagyrskaya 8 queryable.
#
# Its VCFs are plain gzip (magic 1f8b0800), not BGZF (1f8b0804), which is why
# the directory has no .tbi and why the indexing job refused all 22 files. Tabix
# fundamentally cannot index plain gzip: random access needs BGZF's independently
# inflatable blocks.
#
# The obvious fix -- recompress all 45 GB as BGZF -- is the wrong one. The screen
# only ever looks at the ~1.5 Mb of catalog regions, so instead we stream each
# chromosome once and keep only records inside those regions (generously padded),
# producing a small BGZF VCF that indexes in seconds. One sequential read of the
# source either way, but a few MB of output instead of 45 GB, and the source is
# never modified.
#
# Submit:  FORCE=1 sbatch --export=ALL,FORCE=1 sbatch_extract_chagyrskaya.sh
# Verify:  bash sbatch_extract_chagyrskaya.sh --report

set -uo pipefail

E=/n/groups/reich/DAVID/CURSOR_ENRICH
SRC=/n/data1/hms/genetics/reich/1000Genomes/nean_chagyrskaya_2019mar/D-vcf_hicov/AA.raw
OUTDIR=$E/refs/archaic_local
IDXDIR=$E/refs/archaic_index
# The region set must match the one the screen will use. The first extraction
# was built for preal50.regions.tsv; the current screen covers the wider
# P(real)>=0.3 set, so re-extracting is mandatory or Chagyrskaya 8 silently
# has no records over the added regions and its haplotypes drop out of the
# denominator at exactly the sites this rerun is meant to strengthen.
REGIONS=${REGIONS:-$E/fixed_derived/screen_preal30.padded.regions.tsv}
VENV=$E/work/venv
PAD=20000          # generous: the screen pads its own regions, and extra
                   # records only cost bytes, whereas missing ones are silent

mkdir -p "$OUTDIR" "$IDXDIR"

if [ "${1:-}" = "--report" ]; then
  echo "=== Chagyrskaya extraction ==="
  n=0
  for c in $(seq 1 22); do
    f=$OUTDIR/Chagyrskaya8.chr$c.vcf.gz
    i=$IDXDIR/Chagyrskaya8.chr$c.tbi
    if [ -s "$f" ] && [ -s "$i" ]; then
      n=$((n+1))
      printf "  chr%-3s %8s  %s recs\n" "$c" "$(du -h "$f" | cut -f1)" \
        "$(zcat "$f" 2>/dev/null | grep -vc '^#')"
    else
      printf "  chr%-3s MISSING\n" "$c"
    fi
  done
  echo "  ---- $n/22 ready"
  exit 0
fi

C="${SLURM_ARRAY_TASK_ID:?}"
IN=$SRC/chr${C}.noRB.vcf.gz
OUT=$OUTDIR/Chagyrskaya8.chr${C}.vcf.gz
TMP=$OUTDIR/.chr${C}.$$.vcf

echo "=== Chagyrskaya8 chr$C ==="
echo "  in:  $IN"
echo "  out: $OUT"

[ -f "$IN" ] || { echo "  ERROR: source missing"; exit 1; }
# FORCE=1 overrides the resume check, which is what a change of region set
# requires: the old output looks complete but covers the wrong intervals.
if [ "${FORCE:-0}" != "1" ] \
   && [ -s "$OUT" ] && [ -s "$IDXDIR/Chagyrskaya8.chr${C}.tbi" ]; then
  echo "  already done (FORCE=1 to redo)"; exit 0
fi

# Intervals for this chromosome, padded. Written to a file so awk can read them
# without any quoting games.
IV=$OUTDIR/.iv.chr${C}.$$
awk -F'\t' -v c="$C" -v pad="$PAD" '
  NR==1 { for (i=1;i<=NF;i++) h[$i]=i; next }
  { ch=$h["chrom"]; sub(/^chr/,"",ch)
    if (ch==c) { s=$h["start"]-pad; if (s<1) s=1; print s"\t"$h["end"]+pad } }
' "$REGIONS" > "$IV"

nreg=$(wc -l < "$IV")
echo "  regions on this chromosome: $nreg"
if [ "$nreg" -eq 0 ]; then
  echo "  no catalog regions here; writing header-only VCF for uniformity"
fi

# Single streaming pass. zcat handles plain gzip fine -- it is only random
# access that BGZF is needed for, and we are reading start to finish.
zcat "$IN" | awk -v ivf="$IV" '
  BEGIN { n=0
          while ((getline line < ivf) > 0) { split(line,a,"\t"); lo[n]=a[1]; hi[n]=a[2]; n++ } }
  /^#/  { print; next }
  n==0  { next }
  { p=$2+0
    for (i=0;i<n;i++) if (p>=lo[i] && p<=hi[i]) { print; next } }
' > "$TMP"

nrec=$(grep -vc '^#' "$TMP" || true)
echo "  records kept: $nrec"
if [ "$nrec" -eq 0 ] && [ "$nreg" -gt 0 ]; then
  echo "  ERROR: regions exist but no records matched -- contig/position mismatch?"
  head -3 "$IV"; grep -v '^#' "$TMP" | head -2
  rm -f "$TMP" "$IV"; exit 1
fi

# shellcheck disable=SC1091
source "$VENV/bin/activate" 2>/dev/null || { echo "  ERROR: no venv"; rm -f "$TMP" "$IV"; exit 1; }

python3 - "$TMP" "$OUT" "$IDXDIR/Chagyrskaya8.chr${C}.tbi" <<'PYEOF'
import os, shutil, sys
import pysam

tmp, out, idx = sys.argv[1], sys.argv[2], sys.argv[3]

# tabix_compress writes real BGZF; gzip.open would reproduce the original problem.
pysam.tabix_compress(tmp, out, force=True)
pysam.tabix_index(out, preset="vcf", force=True)

made = out + ".tbi"
if not os.path.exists(made):
    print("  ERROR: no index produced", file=sys.stderr); sys.exit(1)
shutil.copyfile(made, idx)

tb = pysam.TabixFile(out, index=idx)
contigs = list(tb.contigs)
got = sum(1 for _ in tb.fetch(contigs[0])) if contigs else 0
print(f"  OK  {os.path.getsize(out)} bytes  contigs={contigs}  fetchable={got}")
if got == 0 and contigs:
    print("  ERROR: index loads but returns nothing", file=sys.stderr); sys.exit(1)
PYEOF
rc=$?
rm -f "$TMP" "$IV"
exit $rc
```
