## Supplementary Information section 4 code base for "Hypothesis: A modern human range expansion ∼300,000 years ago had a critical role in Neandertal origins": sbatch_index_archaics.sh

```
#!/bin/bash
#SBATCH --job-name=archidx
#SBATCH --partition=short
#SBATCH --time=6:00:00
#SBATCH --mem=4G
#SBATCH --cpus-per-task=1
#SBATCH --array=0-137
#SBATCH --output=/n/groups/reich/DAVID/CURSOR_ENRICH/fixed_derived/logs/archidx_%A_%a.out
#SBATCH --error=/n/groups/reich/DAVID/CURSOR_ENRICH/fixed_derived/logs/archidx_%A_%a.err
#
# Build tabix indexes for the 6-genome archaic panel (4 Neandertal + 2 Denisovan).
#
# The source VCFs sit under other users' directories in /n/data1, so indexes are
# written into OUR tree, never beside the source. Two reasons: we may not hold
# write permission there, and even where we do, dropping files into a colleague's
# data directory is the kind of thing that gets noticed for the wrong reasons.
# pysam.TabixFile(path, index=...) reads an index from an arbitrary location, so
# the screen can point at these explicitly.
#
# Indexing does NOT touch the source file -- it only reads it. But there is one
# real hazard: if a file is plain gzip rather than BGZF, tabix cannot index it,
# and some code paths respond by recompressing IN PLACE. Every file is therefore
# checked for the BGZF magic first and skipped if it is not BGZF, so we can never
# rewrite someone else's data.
#
# Submit:  sbatch sbatch_index_archaics.sh
# Collate: bash sbatch_index_archaics.sh --report

set -uo pipefail

ROOT=/n/groups/reich/DAVID/CURSOR_ENRICH
IDXDIR=$ROOT/refs/archaic_index
VENV=$ROOT/work/venv

# id : lineage : dir : filename pattern with {c} for the chromosome token
PANEL=(
"Denisova3:Denisovan:/n/data1/hms/genetics/reich/1000Genomes/REORG/FOR_DATA_SHARING_DIRECTORY/sample_sets/denisova_dec11/vcfs:DenisovaPinky.hg19_1000g.{c}.mod.vcf.gz"
"Altai:Neandertal:/n/data1/hms/genetics/reich/1000Genomes/nean_altai_jul12/hg19/A-raw/VCF_mod:AltaiNea.hg19_1000g.{c}.mod.vcf.gz"
"Vindija33.19:Neandertal:/n/data1/hms/genetics/reich/1000Genomes/nean_mez_2016jul/hg19/A-raw/genotypes/Vindija33.19:chr{c}_mq25_mapab100.vcf.gz"
"Chagyrskaya8:Neandertal:/n/data1/hms/genetics/reich/1000Genomes/nean_chagyrskaya_2019mar/D-vcf_hicov/AA.raw:chr{c}.noRB.vcf.gz"
"Denisova25:Denisovan:/n/data1/hms/genetics/reich/1000Genomes/amh_samples/peyregneKelso2025__200kDenisova_sample__2025.11.05/AA.raw/VCF:chr{c}.Den25.L35MQ25.B30.map35_100.vcf.gz"
"GoyetQ56-1:Neandertal:/n/data1/hms/genetics/reich/1000Genomes/amh_samples/Mesa.Hajdinjak2026__Nature__PRJEB98484__edid586__27.late.neandertals__2026.09.05/A.round1/A.raw:GoyetQ56_1_final.uniq.L35MQ25.sorted.chr_{c}.indel_realn_mq25.map35_100.vcf.gz"
)

mkdir -p "$IDXDIR" "$ROOT/fixed_derived/logs"

# ---------------------------------------------------------------- report mode
if [ "${1:-}" = "--report" ]; then
  echo "=== archaic index inventory: $IDXDIR ==="
  tot=0; have=0
  for spec in "${PANEL[@]}"; do
    IFS=: read -r id lin dir pat <<< "$spec"
    n=0
    for c in $(seq 1 22); do
      f="$dir/${pat//\{c\}/$c}"
      [ -e "$f" ] || continue
      tot=$((tot+1))
      if [ -s "$IDXDIR/$id.chr$c.tbi" ]; then n=$((n+1)); have=$((have+1)); fi
    done
    printf "  %-14s %-11s %2d/22 indexed\n" "$id" "$lin" "$n"
  done
  echo "  ---- $have/$tot total"
  echo
  echo "failures, if any:"
  grep -l "SKIP\|ERROR" "$ROOT"/fixed_derived/logs/archidx_*.err 2>/dev/null \
    | head -20 | while read -r f; do echo "  == $f"; grep "SKIP\|ERROR" "$f" | sed 's/^/     /'; done
  exit 0
fi

# ------------------------------------------------------- build the task table
# 6 genomes x 22 autosomes = 132 tasks (array is 0-137 for slack).
TASKS=()
for spec in "${PANEL[@]}"; do
  IFS=: read -r id lin dir pat <<< "$spec"
  for c in $(seq 1 22); do
    TASKS+=("$id|$dir/${pat//\{c\}/$c}|$c")
  done
done

T="${SLURM_ARRAY_TASK_ID:-0}"
if [ "$T" -ge "${#TASKS[@]}" ]; then
  echo "task $T beyond panel (${#TASKS[@]} real tasks); nothing to do"; exit 0
fi

IFS='|' read -r ID VCF CHR <<< "${TASKS[$T]}"
OUT=$IDXDIR/$ID.chr$CHR.tbi

echo "=== $ID chr$CHR ==="
echo "  src: $VCF"
echo "  idx: $OUT"

if [ ! -e "$VCF" ]; then echo "  SKIP: source missing"; exit 0; fi
if [ -s "$OUT" ]; then echo "  already indexed ($(wc -c < "$OUT") bytes)"; exit 0; fi

# BGZF check before anything else. A BGZF member is a gzip member carrying an
# extra subfield tagged 'BC'; plain gzip lacks it and cannot be tabix-indexed.
if ! od -An -tx1 -N4 "$VCF" | tr -d ' \n' | grep -qi '^1f8b0804'; then
  echo "  SKIP: not BGZF (plain gzip?) -- refusing to touch it, since"
  echo "        recompressing would rewrite another user's file"
  od -An -tx1 -N4 "$VCF"
  exit 0
fi

# shellcheck disable=SC1091
source "$VENV/bin/activate" 2>/dev/null || { echo "  ERROR: no venv at $VENV"; exit 1; }

python3 - "$VCF" "$OUT" <<'PYEOF'
import os, shutil, sys, tempfile
import pysam

vcf, out = sys.argv[1], sys.argv[2]

# pysam.tabix_index writes <vcf>.tbi beside the source, which is exactly what we
# must not do. So index a symlink inside a scratch dir we own, then move the
# resulting .tbi to its final name. The symlink target is only ever read.
tmp = tempfile.mkdtemp(prefix="tbi_", dir=os.path.dirname(out))
try:
    link = os.path.join(tmp, os.path.basename(vcf))
    os.symlink(os.path.abspath(vcf), link)
    pysam.tabix_index(link, preset="vcf", force=True, keep_original=True)
    made = link + ".tbi"
    if not os.path.exists(made):
        print("  ERROR: tabix produced no index", file=sys.stderr)
        sys.exit(1)
    shutil.move(made, out)

    # An index that exists is not an index that works: confirm it loads and that
    # a region query on it actually returns records.
    tb = pysam.TabixFile(vcf, index=out)
    contigs = list(tb.contigs)
    got = 0
    for c in contigs[:1]:
        for i, _ in enumerate(tb.fetch(c)):
            got = i + 1
            if got >= 5:
                break
    print(f"  OK  {os.path.getsize(out)} bytes  contigs={contigs[:3]}  fetched={got}")
    if got == 0:
        print("  ERROR: index loads but returns no records", file=sys.stderr)
        sys.exit(1)
finally:
    shutil.rmtree(tmp, ignore_errors=True)
PYEOF
rc=$?
[ $rc -ne 0 ] && echo "  ERROR: indexing failed rc=$rc"
exit $rc
```
