## Supplementary Information section 4 code base for "Hypothesis: A modern human range expansion ∼300,000 years ago had a critical role in Neandertal origins": sbatch_relate_archaic.sh

```
#!/bin/bash
#SBATCH --job-name=relarch
#SBATCH --partition=medium
#SBATCH --time=1-00:00:00
# Far cheaper than the 1000GP_Phase3 run, and for one reason: 612 haplotypes
# instead of 5,008. Relate's converter writes every tree with its own full edge
# set, so the .trees file scales with (trees x 2n). Measured on chr22 of the
# 5,008-haplotype set: 31,332 trees, ~313M edges, 13.8 GB, 32 GB peak RSS.
# Scaling both factors by 612/5008 puts this set's chr22 near 0.2 GB and its
# whole genome in the tens of GB, so no per-chromosome memory tiering is needed
# and 48G carries chr1. If that estimate proves badly wrong, the fix is
# relate_lib's `Convert --compress`, which shares node ages across adjacent
# trees and reads .anc.gz directly; it is not needed at this size.
#SBATCH --mem=48G
#SBATCH --cpus-per-task=1
#SBATCH --array=1-22%6
#SBATCH --output=/n/groups/reich/DAVID/CURSOR_ARGWEAVER/logs/relarch_%A_%a.out
#SBATCH --error=/n/groups/reich/DAVID/CURSOR_ARGWEAVER/logs/relarch_%A_%a.err
#
# TMRCA on a 5-kb grid from the Relate ARG that contains archaic genomes.
#
# Dataset: relate_YRICHBGBRNEA.zip, from the Relate homepage's "Inferred trees"
# section, labelled "1000G CHB, GBR, YRI, Neanderthals (hg37)". The label
# undersells it -- the Denisovan is in there too, which is what makes this the
# only released Relate ARG able to reproduce the catalog's case/control design.
# Composition confirmed from its poplabels: YRI 108, CHB 103, GBR 91, plus
# AltaiNeandertal, Vindija, Chagyrskaya and Denisova; 306 individuals, 612
# haplotypes, matching sample_ages_hap.txt (604 contemporary + 8 archaic).
#
# Sample ages are in generations and, at 28 years each, come out to exactly 55,
# 68, 73 and 110 ka -- so 28 is this dataset's own generation time and is used
# here rather than the scan's.
#
# Unlike 1000GP_Phase3_mask_prene, this release carries no "prene" marker and
# ships a companion .coal, so its branch lengths appear to be Ne-fitted already.
# That should be confirmed before its TMRCAs are read as absolute dates.
#
# Submit:  sbatch sbatch_relate_archaic.sh
# Report:  bash sbatch_relate_archaic.sh --report

set -uo pipefail

A=/n/groups/reich/DAVID/CURSOR_ARGWEAVER
ARCH=$A/relate_archaic
ZIP=$ARCH/relate_YRICHBGBRNEA.zip
BIN=$A/relate-build/bin
BASE=1000G_sub_Nea
POPLABELS=$ARCH/data/1000GP_Phase3_sub_Nea.poplabels
VENV="${VENV:-$A/relate_work/venv}"
[ -x "$VENV/bin/python3" ] || VENV=/n/groups/reich/DAVID/CURSOR_ENRICH/work/venv
GRID=5000
YPG=28

TSDIR=$ARCH/ts
TSVDIR=$ARCH/tsv
mkdir -p "$TSDIR" "$TSVDIR" "$A/logs"

# Three Neandertal variants are emitted on the case side, because pooling all
# three turned out to be the wrong default and the first run proved it.
#
# The case statistic is the ROOT of moderns plus Neandertals, so it is bounded
# below by the Neandertal-internal root. Pooling three structured Neandertals
# puts that floor at a median of 313 ka against the single Denisovan's 188 ka,
# and at a threshold of 700 ka it bars 14.0% of grid points from ever being a
# case while barring only 2.8% from being a control -- a 5.1x asymmetry that
# shrinks to 4.6x by 1500 ka. Being threshold-dependent, it does not cancel
# into omega0, and it drove P(real) to -2.0 in the shallow tail where the
# signal actually lives. The medians hid this completely: adding archaic
# genomes moves the median root by under 1%, which is why checking medians was
# not a valid test of the decision.
#
#   nea3  all three; better evidence per locus, unusable in a mirror contrast
#   nea2  Altai + Vindija; matches ARGweaver's 02_modern_plus_both_Nea, and
#         reproduces its 4-versus-2 haplotype asymmetry, which is mild enough
#         there that the shallow tail behaves
#   nea1  Altai alone; 2 versus 2, the only exactly symmetric contrast, and so
#         the control for whether any residual asymmetry still matters
#
# The archaic-internal roots are emitted for each variant too, since they are
# what quantifies the floor.
#
# Both modern panels are emitted in one pass. YRI-only mirrors the original
# African-only design; YRI+GBR+CHB asks the stronger question of whether every
# sampled population coalesces recently, at some cost in sensitivity. Panel
# size matters less than it appears: under a neutral coalescent, 16 -> 216
# haplotypes deepens the expected root by only 6%, and the root's variance is
# dominated by the final coalescent interval and does not shrink at all.
#
# Note the spelling: the sample id is AltaiNeandertal while its population
# label is AltaiNeanderthal. The subset matcher accepts either.
NEA3=AltaiNeandertal,Vindija,Chagyrskaya
NEA2=AltaiNeandertal,Vindija
NEA1=AltaiNeandertal
SUBSETS=(
  "yri_01_modern=YRI"
  "yri_02_mod_nea=YRI,$NEA3"
  "yri_02b_mod_nea2=YRI,$NEA2"
  "yri_02c_mod_nea1=YRI,$NEA1"
  "yri_03_mod_den=YRI,Denisova"
  "ygc_01_modern=YRI,GBR,CHB"
  "ygc_02_mod_nea=YRI,GBR,CHB,$NEA3"
  "ygc_02b_mod_nea2=YRI,GBR,CHB,$NEA2"
  "ygc_02c_mod_nea1=YRI,GBR,CHB,$NEA1"
  "ygc_03_mod_den=YRI,GBR,CHB,Denisova"
  "arch_04_nea_only=$NEA3"
  "arch_04b_nea2_only=$NEA2"
  "arch_05_den_only=Denisova"
  "arch_06_all=ALL"
)

if [ "${1:-}" = "--report" ]; then
  n=$(ls -1 "$TSVDIR"/chr*.tsv 2>/dev/null | wc -l)
  echo "grids: $n/22"
  for f in "$TSVDIR"/chr*.tsv; do
    [ -e "$f" ] || continue
    printf "  %-14s %7d rows  %s\n" "$(basename "$f")" \
      "$(( $(wc -l < "$f") - 1 ))" "$(du -h "$f" | cut -f1)"
  done
  echo "leftover .trees (should be none once grids are complete):"
  ls -la "$TSDIR"/*.trees 2>/dev/null || echo "  none"
  exit 0
fi

C="${SLURM_ARRAY_TASK_ID:?}"
declare -A LEN=( [1]=249250621 [2]=243199373 [3]=198022430 [4]=191154276
  [5]=180915260 [6]=171115067 [7]=159138663 [8]=146364022 [9]=141213431
  [10]=135534747 [11]=135006516 [12]=133851895 [13]=115169878 [14]=107349540
  [15]=102531392 [16]=90354753 [17]=81195210 [18]=78077248 [19]=59128983
  [20]=63025520 [21]=48129895 [22]=51304566 )
CLEN=${LEN[$C]}

ts() { date +%H:%M:%S; }
echo "=== chr$C  len=$CLEN  ==="
[ -f "$ZIP" ] || { echo "ERROR: missing $ZIP"; exit 1; }
[ -f "$POPLABELS" ] || { echo "ERROR: missing $POPLABELS"; exit 1; }

STAGE=$TSDIR/stage_chr$C
mkdir -p "$STAGE"
IN=$STAGE/chr$C
TS=$TSDIR/chr$C

if [ ! -s "$TS.trees" ]; then
  # ConvertToTreeSequence wants an extensionless prefix with <prefix>.anc and
  # <prefix>.mut beside each other, and it must be given UNCOMPRESSED input:
  # handed a .gz it exits zero having written nothing, which is how the first
  # attempt on the other dataset failed silently on every chromosome.
  echo "[$(ts)] extracting chr$C from the archive ..."
  unzip -p "$ZIP" "${BASE}_chr${C}.anc.gz" | gunzip -c > "$IN.anc" || exit 1
  unzip -p "$ZIP" "${BASE}_chr${C}.mut.gz" | gunzip -c > "$IN.mut" || exit 1
  echo "[$(ts)] staged: $(du -sh "$STAGE" | cut -f1)"

  echo "[$(ts)] converting to tree sequence ..."
  "$BIN/RelateFileFormats" --mode ConvertToTreeSequence -i "$IN" -o "$TS" || true
  echo "[$(ts)] conversion returned"
fi
[ -s "$TS.trees" ] || { echo "ERROR: no tree sequence at $TS.trees"; exit 1; }
echo "  $TS.trees  $(du -h "$TS.trees" | cut -f1)"
rm -f "$IN.anc" "$IN.mut"

# shellcheck disable=SC1091
source "$VENV/bin/activate" 2>/dev/null || { echo "ERROR: no venv"; exit 1; }
python3 -c "import tskit" 2>/dev/null || pip install --quiet tskit || {
  echo "ERROR: cannot import or install tskit"; exit 1; }

SUB=()
for s in "${SUBSETS[@]}"; do SUB+=(--subset "$s"); done

echo "[$(ts)] computing TMRCA grid over ${#SUBSETS[@]} subsets ..."
python3 "$A/scripts/relate_ts_tmrca_grid.py" \
  --trees "$TS.trees" --chrom "$C" --chrom-len "$CLEN" \
  --poplabels "$POPLABELS" "${SUB[@]}" \
  --grid-step "$GRID" --years-per-gen "$YPG" \
  -o "$TSVDIR/chr${C}.tsv" || exit 1

# Delete the intermediate only against a verified-complete grid. Deleting on
# mere existence throws away the conversion whenever a job dies mid-write,
# which is precisely when rebuilding costs most.
EXP=$(( CLEN / GRID ))
GOT=$(( $(wc -l < "$TSVDIR/chr${C}.tsv" 2>/dev/null || echo 1) - 1 ))
if [ "$GOT" -ge "$(( EXP - 1 ))" ]; then
  SZ=$(du -h "$TS.trees" | cut -f1)
  rm -f "$TS.trees"; rmdir "$STAGE" 2>/dev/null
  echo "[$(ts)] grid complete ($GOT rows, expected ~$EXP); reclaimed $SZ"
else
  echo "[$(ts)] WARNING grid has $GOT rows, expected ~$EXP; keeping $TS.trees"
fi
echo "[$(ts)] Done chr$C"
```
