## Supplementary Information section 4 code base for "Hypothesis: A modern human range expansion ∼300,000 years ago had a critical role in Neandertal origins": sbatch_relate_tmrca_chr_array.sh

```
#!/usr/bin/env bash
# Slurm array: one task per chromosome listed in CHROM_LIST (one label per line, e.g. 1..22).
# Use relate_chrom_list_autosomes_22.txt for autosomes only; use relate_chrom_list_autosomes_x.txt to include X.
#
# Submit example:
# Paths below were verified on O2 in Sep 2026.
#   export REPO_ROOT=/n/groups/reich/DAVID/CURSOR_ARGWEAVER
#   export RELATE_BIN=$REPO_ROOT/relate-build/bin/RelateExtract
#   export RELATE_DATA=/n/data1/hms/genetics/reich/1000Genomes/datasets/relate_1000g_v37/raw
#   export ANC_BASENAME='results/anc/1000GP_Phase3_mask_prene_chr{CHR}.anc.gz'
#   export MUT_BASENAME='results/mut/1000GP_Phase3_mask_prene_chr{CHR}.mut.gz'
#   export OUTNW=$REPO_ROOT/relate_work/newick
#   export OUTTSV=$REPO_ROOT/relate_work/tsv
#   export CHROM_LIST=$REPO_ROOT/scripts/relate_chrom_list_autosomes_x.txt
#   sbatch --array=1-23%5 $REPO_ROOT/scripts/sbatch_relate_tmrca_chr_array.sh
#
# VENV is no longer needed; relate_marginal_tmrca_grid.py has no third-party deps.
# The partition/walltime below (long, 7 days) matter: AncToNewick on chr1 exceeded
# the 12 h short-partition limit in the May 2026 attempt.
#
# Edit #SBATCH resources and partition for your site.

#SBATCH -J relate_tmrca
#SBATCH --cpus-per-task=2
#SBATCH --mem=64G
#SBATCH --time=7-00:00:00
#SBATCH --partition=long

set -euo pipefail

: "${REPO_ROOT:?}"
: "${RELATE_BIN:?}"
: "${RELATE_DATA:?}"
: "${ANC_BASENAME:?}"
: "${MUT_BASENAME:?}"
: "${OUTNW:?}"
: "${OUTTSV:?}"
: "${CHROM_LIST:?}"
: "${POPLABELS:?}"

TASK="${SLURM_ARRAY_TASK_ID:?}"
CHROM="$(sed -n "${TASK}p" "$CHROM_LIST")"
if [[ -z "$CHROM" ]]; then
  echo "ERROR: empty CHROM for task $TASK"
  exit 1
fi

export CHROM RELATE_BIN RELATE_DATA ANC_BASENAME MUT_BASENAME REPO_ROOT OUTNW OUTTSV
export POPLABELS SUBSETS
export YEARS_PER_GEN="${YEARS_PER_GEN:-28}"
export GRID_STEP="${GRID_STEP:-5000}"
export VENV="${VENV:-}"
# COAL enables branch-length re-inference in cluster_relate_one_chr.sh. Without
# it the pre-Ne branch lengths are used and the TMRCAs are not coalescent-
# calibrated, so it should normally be set.
export COAL="${COAL:-}"
export MU="${MU:-1.25e-8}"

bash "$REPO_ROOT/scripts/cluster_relate_one_chr.sh"
```
