## Supplementary Information section 4 code base for "Hypothesis: A modern human range expansion ∼300,000 years ago had a critical role in Neandertal origins": screen_coverage.py

```
#!/usr/bin/env python3
"""How much of the autosomes the scan actually looked at.

The catalog can only contain sweeps at positions ARGweaver-D produced a local
tree for, and that is not the whole genome. This prints the shortfall three
ways: bases, protein-coding bases, and whole genes, because they answer
different objections. Bases are the headline number but include assembly gaps,
which nothing could have screened; coding bases are immune to that complaint,
since a coding base is by construction real sequence; and the gene count is
what a reader actually wants when asking "could my favourite gene have shown
up at all?".

The screened territory is read off the scan output rather than off the
ARGweaver-D window file, because what matters is where a TMRCA was recorded,
not where a window was nominally defined. A grid point stands for --grid-step
bases. As a check that this is the right reading, the 5 kb and 1 kb runs must
agree: 488,021 points x 5 kb and 2,440,090 points x 1 kb are both 2.440 Gb, so
the denser grid resampled the same territory rather than reaching new territory.

Coverage is very uneven and the per-chromosome table is worth reading, not just
the total. Chromosome 19 is under a third screened, which is the reason no
catalog locus lies there; it is a fact about the input data, not a biological
result.

Usage:
  python3 scripts/screen_coverage.py \
      --scan k1/supplementary_table2_scan_1kb.tsv.gz --grid-step 1000
"""

import argparse
import csv
import gzip
import os
import re
import sys
from collections import defaultdict

# hg19 / GRCh37 autosome lengths. Hard-coded rather than read from a .fai so the
# script runs against the release on its own; these are assembly constants.
HG19 = {"1": 249250621, "2": 243199373, "3": 198022430, "4": 191154276,
        "5": 180915260, "6": 171115067, "7": 159138663, "8": 146364022,
        "9": 141213431, "10": 135534747, "11": 135006516, "12": 133851895,
        "13": 115169878, "14": 107349540, "15": 102531392, "16": 90354753,
        "17": 81195210, "18": 78077248, "19": 59128983, "20": 63025520,
        "21": 48129895, "22": 51304566}
AUTO = set(HG19)


def opener(path):
    return gzip.open(path, "rt") if path.endswith(".gz") else open(path)


def union(iv):
    """Merge overlapping/abutting [start, end) intervals."""
    iv = sorted(iv)
    out = []
    for a, b in iv:
        if out and a <= out[-1][1]:
            out[-1][1] = max(out[-1][1], b)
        else:
            out.append([a, b])
    return out


def overlap(a_iv, b_iv):
    """Total bases in both interval lists; both must be sorted and disjoint."""
    i = j = tot = 0
    while i < len(a_iv) and j < len(b_iv):
        a0, a1 = a_iv[i]
        b0, b1 = b_iv[j]
        tot += max(0, min(a1, b1) - max(a0, b0))
        if a1 < b1:
            i += 1
        else:
            j += 1
    return tot


def screened_intervals(scan, step):
    """Union of [pos, pos + step) over every grid point in the scan."""
    pts = defaultdict(list)
    with opener(scan) as f:
        rd = csv.reader(f, delimiter="\t")
        head = next(rd)
        ci, pi = head.index("chrom"), head.index("pos")
        for row in rd:
            if row[ci] in AUTO:
                pts[row[ci]].append(int(row[pi]))
    out = {}
    for c, ps in pts.items():
        ps.sort()
        iv, s, e = [], ps[0], ps[0]
        for p in ps[1:]:
            if p <= e + step:
                e = p
            else:
                iv.append([s, e + step])
                s = e = p
        iv.append([s, e + step])
        out[c] = iv
    return out


def coding(gtf):
    """Per-gene autosomal CDS intervals, protein-coding genes only."""
    per_gene = defaultdict(lambda: defaultdict(list))
    with gzip.open(gtf, "rt") as f:
        for line in f:
            if line.startswith("#"):
                continue
            p = line.split("\t")
            if p[2] != "CDS":
                continue
            c = p[0][3:] if p[0].startswith("chr") else p[0]
            if c not in AUTO or 'gene_type "protein_coding"' not in p[8]:
                continue
            m = re.search(r'gene_name "([^"]+)"', p[8])
            if m:
                per_gene[m.group(1)][c].append([int(p[3]) - 1, int(p[4])])
    for g in per_gene:
        for c in per_gene[g]:
            per_gene[g][c] = union(per_gene[g][c])
    return per_gene


def main():
    ap = argparse.ArgumentParser()
    ap.add_argument("--scan", default="k1/supplementary_table2_scan_1kb.tsv.gz",
                    help="genome-wide scan table; one row per grid point")
    ap.add_argument("--grid-step", type=int, default=1000,
                    help="bases a grid point stands for (1000 for the 1 kb "
                         "scan, 5000 for the 5 kb one)")
    ap.add_argument("--gtf", default="refdata/gencode.v19.annotation.gtf.gz")
    ap.add_argument("-o", "--out", default="k1/screen_coverage.tsv")
    args = ap.parse_args()

    scr = screened_intervals(args.scan, args.grid_step)

    rows = []
    tot_len = tot_scr = 0
    for c in sorted(HG19, key=int):
        iv = scr.get(c, [])
        s = sum(b - a for a, b in iv)
        rows.append({"chrom": c, "length_bp": HG19[c], "screened_bp": s,
                     "screened_pct": f"{100 * s / HG19[c]:.1f}"})
        tot_len += HG19[c]
        tot_scr += s

    # Coding bases and whole genes.
    genes = coding(args.gtf)
    cds_all = defaultdict(list)
    for g in genes:
        for c, iv in genes[g].items():
            cds_all[c].extend(iv)
    for c in cds_all:
        cds_all[c] = union(cds_all[c])
    cds_tot = sum(b - a for c in cds_all for a, b in cds_all[c])
    cds_scr = sum(overlap(cds_all[c], scr.get(c, [])) for c in cds_all)

    full = partial = missing = 0
    missing_names = []
    for g, by_c in genes.items():
        tot = sum(b - a for c in by_c for a, b in by_c[c])
        cov = sum(overlap(by_c[c], scr.get(c, [])) for c in by_c)
        if cov == 0:
            missing += 1
            missing_names.append(g)
        elif cov >= tot:
            full += 1
        else:
            partial += 1
    n_genes = len(genes)

    with open(args.out, "w", newline="") as f:
        w = csv.DictWriter(f, fieldnames=list(rows[0]), delimiter="\t")
        w.writeheader()
        w.writerows(rows)
        w.writerow({"chrom": "all", "length_bp": tot_len, "screened_bp": tot_scr,
                    "screened_pct": f"{100 * tot_scr / tot_len:.1f}"})

    p = print
    p(f"{'chr':>4}{'length Mb':>11}{'screened Mb':>13}{'screened %':>12}")
    for r in rows:
        p(f"{r['chrom']:>4}{r['length_bp'] / 1e6:>11.1f}"
          f"{r['screened_bp'] / 1e6:>13.1f}{r['screened_pct']:>12}")
    p(f"{'all':>4}{tot_len / 1e6:>11.1f}{tot_scr / 1e6:>13.1f}"
      f"{100 * tot_scr / tot_len:>12.1f}")
    p("")
    p(f"autosomal bases        {tot_len / 1e9:.3f} Gb")
    p(f"  screened             {tot_scr / 1e9:.3f} Gb "
      f"({100 * tot_scr / tot_len:.1f}%)")
    p(f"  NOT screened         {(tot_len - tot_scr) / 1e6:.0f} Mb "
      f"({100 * (tot_len - tot_scr) / tot_len:.1f}%)")
    p("")
    p(f"autosomal coding bases {cds_tot / 1e6:.1f} Mb")
    p(f"  screened             {cds_scr / 1e6:.1f} Mb "
      f"({100 * cds_scr / cds_tot:.1f}%)")
    p(f"  NOT screened         {(cds_tot - cds_scr) / 1e6:.1f} Mb "
      f"({100 * (cds_tot - cds_scr) / cds_tot:.1f}%)")
    p("")
    p(f"autosomal protein-coding genes {n_genes:,}")
    p(f"  fully screened       {full:>6,} ({100 * full / n_genes:.1f}%)")
    p(f"  partly screened      {partial:>6,} ({100 * partial / n_genes:.1f}%)")
    p(f"  not screened at all  {missing:>6,} ({100 * missing / n_genes:.1f}%)")
    p(f"  e.g. {', '.join(sorted(missing_names)[:10])}")
    p("")
    p(f"wrote {args.out}")


if __name__ == "__main__":
    sys.exit(main())
```
