## Supplementary Information section 4 code base for "Hypothesis: A modern human range expansion ∼300,000 years ago had a critical role in Neandertal origins": site_exact_preal.py

```
#!/usr/bin/env python3
"""P(real) at exact substitution positions, from the site-exact TMRCA read.

The surface is fitted on the genome-wide 5 kb scan, exactly as for Table S4.1,
and then evaluated at the substitution positions. Fitting on the sites
themselves would be meaningless -- 936 points cannot define the case/control
surface -- so the two inputs are kept separate: --scan supplies the fit,
--sites supplies the positions to score.

Chromosome-parity cross-validation is preserved, so a site on chromosome 3 is
scored by a surface fitted on the even chromosomes and never on its own.

The output is directly comparable to the preal column of Supplementary Table 1,
with one difference that is the whole point: the TMRCAs here are read from the
local tree spanning the substitution itself, not from the nearest lattice
point up to 2.5 kb away.
"""

import argparse
import os
import sys

import numpy as np

sys.path.insert(0, os.path.dirname(os.path.abspath(__file__)))
from approx_percentiles import default_coord, fit_site, load, z_of  # noqa: E402
from fig_s51_model_curve import U_EDGES, V_EDGES, fit_on  # noqa: E402
from fit_fdr_surface import PCUTS, design  # noqa: E402
from make_loci_table import EPS, RULE_CUT, preal_from_theta  # noqa: E402


def percentiles(recs, coord):
    for d in recs:
        for tag in ("nea", "den"):
            mu, sg = fit_site(coord, d[f"{tag}_p025"], d[f"{tag}_p975"])
            for p in PCUTS:
                d[f"{tag}_hi{p}"] = coord.to_t(mu + z_of(p) * sg)
                d[f"{tag}_lo{p}"] = coord.to_t(mu + z_of(1 - p) * sg)
    return recs


def autosomal(recs):
    return [d for d in recs
            if d["chrom"].replace("chr", "") not in ("X", "Y", "23", "24")]


def dedup(recs):
    """First-wins on (chrom, pos), matching Supplementary Table 1.

    The ARG windows overlap at their boundaries, so a position sitting in an
    overlap is read once per neighbouring window. The reads agree to within
    MCMC noise; taking the first keeps this table consistent with the scan
    tables rather than silently averaging two posteriors.
    """
    seen, out = set(), []
    for d in recs:
        k = (d["chrom"].replace("chr", ""), d["pos"])
        if k in seen:
            continue
        seen.add(k)
        out.append(d)
    return out


def main():
    ap = argparse.ArgumentParser()
    ap.add_argument("--scan", default="hubisz_tmrca_5kb_8afr.tsv",
                    help="genome-wide 5 kb scan; supplies the fitted surface")
    ap.add_argument("--sites",
                    default="catalog_annotation/hubisz_tmrca_sites_8afr.tsv",
                    help="site-exact read from hubisz_out_bed_tmrca_scan.py "
                         "--positions-tsv")
    ap.add_argument("-o", "--out",
                    default="catalog_annotation/site_exact_preal.tsv")
    a = ap.parse_args()

    scan = autosomal(load(a.scan))
    coord, desc = default_coord(scan)
    scan = percentiles(scan, coord)
    sys.stderr.write(f"time coordinate: {desc}\n"
                     f"surface fit set: {len(scan):,} grid points\n")

    odd = [d for d in scan if int(d["chrom"].replace("chr", "")) % 2 == 1]
    even = [d for d in scan if int(d["chrom"].replace("chr", "")) % 2 == 0]
    beta_for_even, n1 = fit_on(odd)
    beta_for_odd, n2 = fit_on(even)
    sys.stderr.write(f"fitted on {n1:,} odd and {n2:,} even cells\n")

    raw = autosomal(load(a.sites))
    sites = percentiles(dedup(raw), coord)
    if len(raw) != len(sites):
        sys.stderr.write(f"window overlaps: {len(raw):,} reads -> "
                         f"{len(sites):,} unique positions\n")
    sys.stderr.write(f"sites to score:  {len(sites):,}\n")

    # the same lattice value, for the side-by-side comparison
    grid = {(int(d["chrom"].replace("chr", "")), d["pos"]): d for d in scan}

    u_lo, u_hi = U_EDGES[0] + EPS, U_EDGES[-1]
    v_lo, v_hi = V_EDGES[0] + EPS, V_EDGES[-1]

    def score(d, c):
        u, v = d[f"nea_hi{RULE_CUT}"], d[f"den_lo{RULE_CUT}"]
        cu, cv = min(max(u, u_lo), u_hi), min(max(v, v_lo), v_hi)
        tag = ("u" if cu != u else "") + ("v" if cv != v else "")
        beta = beta_for_odd if c % 2 == 1 else beta_for_even
        eta = design(np.array([cu / 100.0]), np.array([cv / 100.0]),
                     np.array([RULE_CUT]))[0] @ beta
        return preal_from_theta(1.0 / (1.0 + np.exp(-eta))), tag or "-"

    n_moved = 0
    with open(a.out, "w", encoding="utf-8") as f:
        f.write("chrom\tpos\tafr_mean_ka\tnea_mean_ka\tden_mean_ka\t"
                "preal_site\tclamped\tnearest_grid_pos\tpreal_nearest_grid\t"
                "delta\n")
        for d in sorted(sites, key=lambda d: (int(d["chrom"].replace("chr", "")),
                                              d["pos"])):
            c = int(d["chrom"].replace("chr", ""))
            pr, tag = score(d, c)
            lo = (d["pos"] // 5000) * 5000
            near = min((g for g in (lo, lo + 5000) if (c, g) in grid),
                       key=lambda g: abs(g - d["pos"]), default=None)
            if near is None:
                f.write(f"{c}\t{d['pos']}\t{d['afr_mean']:.0f}\t"
                        f"{d['nea_mean']:.0f}\t{d['den_mean']:.0f}\t"
                        f"{pr:.4f}\t{tag}\tNA\tNA\tNA\n")
                continue
            gpr, _ = score(grid[(c, near)], c)
            n_moved += abs(pr - gpr) > 0.05
            f.write(f"{c}\t{d['pos']}\t{d['afr_mean']:.0f}\t"
                    f"{d['nea_mean']:.0f}\t{d['den_mean']:.0f}\t"
                    f"{pr:.4f}\t{tag}\t{near}\t{gpr:.4f}\t{pr - gpr:+.4f}\n")

    sys.stderr.write(f"wrote {a.out}\n"
                     f"  {n_moved:,} of {len(sites):,} sites move by more "
                     f"than 0.05 against the nearest lattice point\n")
    return 0


if __name__ == "__main__":
    sys.exit(main())
```
