## Supplementary Information section 4 code base for "Hypothesis: A modern human range expansion ∼300,000 years ago had a critical role in Neandertal origins": slurm_hubisz_tmrca_array_task.sh

```
#!/usr/bin/env bash
# Slurm array: one task per line of WINDOWS_LIST (dirname = column 4 of filtered_windows.bed).
# Submit via submit_hubisz_tmrca_slurm.sh (sets --array and exports env).
#
# Required env (export before sbatch or via --export):
#   WINDOWS_LIST  absolute path to newline-separated dirnames (e.g. chr4.71)
#   HUBISZ        dataset root (read-only)
#   REPO          repo with scripts/hubisz_out_bed_tmrca_scan.py and queries JSON
#   OUTDIR        writable; receives parts/ and logs/
#
# Optional env:
#   QUERIES       default: $REPO/scripts/tmrca_queries.hubisz_2afr.json
#   WINDOWS_BED   default: $HUBISZ/filtered_windows.bed
#   VENV          venv dir to activate; default: $OUTDIR/venv
#   GRID_STEP     default: 5000
#   BURNIN_REP    default: 500
#   MAX_REP       if set, passed as --max-rep
#   YEARS_PER_GEN default: 29

#SBATCH -J hubisz_tmrca
#SBATCH --cpus-per-task=1
#SBATCH --mem=8G
#SBATCH --time=04:00:00
# Partition: set by submit_hubisz_tmrca_slurm.sh via sbatch --partition=$PARTITION (do not hardcode here).
# If sbatch requires an account, add a line here, e.g.  #SBATCH --account=reich_lab

set -euo pipefail

: "${WINDOWS_LIST:?Set WINDOWS_LIST to absolute path of dirname list}"
: "${HUBISZ:?Set HUBISZ}"
: "${REPO:?Set REPO}"
: "${OUTDIR:?Set OUTDIR}"

QUERIES="${QUERIES:-$REPO/scripts/tmrca_queries.hubisz_2afr.json}"
WINDOWS_BED="${WINDOWS_BED:-$HUBISZ/filtered_windows.bed}"
VENV="${VENV:-$OUTDIR/venv}"
GRID_STEP="${GRID_STEP:-5000}"
BURNIN_REP="${BURNIN_REP:-500}"
YEARS_PER_GEN="${YEARS_PER_GEN:-29}"

LOG_DIR="$OUTDIR/logs"
PARTS_DIR="$OUTDIR/parts"
mkdir -p "$LOG_DIR" "$PARTS_DIR"

TASK="${SLURM_ARRAY_TASK_ID:?Not running under Slurm array?}"
LOG_OUT="$LOG_DIR/chunk_${SLURM_ARRAY_JOB_ID}_${TASK}.out"
LOG_ERR="$LOG_DIR/chunk_${SLURM_ARRAY_JOB_ID}_${TASK}.err"
exec >"$LOG_OUT" 2>"$LOG_ERR"

DIRNAME="$(sed -n "${TASK}p" "$WINDOWS_LIST")"
if [[ -z "$DIRNAME" ]]; then
  echo "ERROR: empty dirname for SLURM_ARRAY_TASK_ID=$TASK"
  exit 1
fi

PART_TSV="$PARTS_DIR/${DIRNAME}.tsv"
if [[ ! -f "$HUBISZ/$DIRNAME/out.bed.gz" ]]; then
  echo "ERROR: missing $HUBISZ/$DIRNAME/out.bed.gz"
  exit 1
fi

if [[ ! -f "$VENV/bin/activate" ]]; then
  echo "ERROR: venv not found at $VENV (set VENV or create $OUTDIR/venv)"
  exit 1
fi

# shellcheck source=/dev/null
source "$VENV/bin/activate"

MAX_REP_ARGS=()
if [[ -n "${MAX_REP:-}" ]]; then
  MAX_REP_ARGS=(--max-rep "$MAX_REP")
fi

echo "Task $TASK dirname=$DIRNAME -> $PART_TSV"
date -Is

python3 "$REPO/scripts/hubisz_out_bed_tmrca_scan.py" \
  --dataset-root "$HUBISZ" \
  --windows-bed "$WINDOWS_BED" \
  --queries "$QUERIES" \
  --grid-step "$GRID_STEP" \
  --burnin-rep "$BURNIN_REP" \
  "${MAX_REP_ARGS[@]}" \
  --years-per-gen "$YEARS_PER_GEN" \
  --only-dir "$DIRNAME" \
  -o "$PART_TSV"

echo -n "Exit code: "
echo $?
date -Is
```
