## Supplementary Information section 4 code base for "Hypothesis: A modern human range expansion ∼300,000 years ago had a critical role in Neandertal origins": submit_hubisz_tmrca_sites.sh

```
#!/usr/bin/env bash
# Submit a Slurm array that reads subset TMRCAs at exact base positions rather
# than on the 5 kb lattice, for the fixed-derived sites.
#
# The ARG carries one local tree per interval, so a named base gets the tree
# spanning it: this is an exact read of the posterior at that base, not an
# interpolation between lattice points. Only windows that actually contain a
# requested position are submitted, which is why this array is far smaller
# than the genome-wide scan.
#
# Usage:
#   export HUBISZ=...          # dataset root (holds filtered_windows.bed)
#   export REPO=...            # path with scripts/
#   export OUTDIR=...          # writable; use a NEW directory
#   export POSITIONS=...       # TSV with chrom and pos columns
#   export PARTITION=short
#   bash scripts/submit_hubisz_tmrca_sites.sh
#
# After completion:
#   bash scripts/hubisz_tmrca_collate_parts.sh "$OUTDIR/parts" \
#        "$OUTDIR/hubisz_tmrca_sites_8afr.tsv"

set -euo pipefail

: "${HUBISZ:?export HUBISZ (dataset root)}"
: "${REPO:?export REPO (path with scripts/hubisz_out_bed_tmrca_scan.py)}"
: "${OUTDIR:?export OUTDIR (writable; use a new directory)}"
: "${POSITIONS:?export POSITIONS (TSV with chrom and pos columns)}"
: "${PARTITION:?export PARTITION (e.g. short, medium, priority)}"

WINDOWS_BED="${WINDOWS_BED:-$HUBISZ/filtered_windows.bed}"
MAX_PARALLEL="${MAX_PARALLEL:-100}"

SCRIPT_DIR="$(cd "$(dirname "${BASH_SOURCE[0]}")" && pwd)"
SLURM_JOB_SCRIPT="${SLURM_JOB_SCRIPT:-$SCRIPT_DIR/slurm_hubisz_tmrca_sites_task.sh}"

for f in "$WINDOWS_BED" "$POSITIONS" "$SLURM_JOB_SCRIPT"; do
  [[ -f "$f" ]] || { echo "ERROR: not found: $f"; exit 1; }
done

META="$OUTDIR/metadata"
mkdir -p "$META" "$OUTDIR/parts" "$OUTDIR/logs"

# Keep only the windows holding at least one requested position. The windows
# bed is (chrom start end dirname); the positions file has a header row.
WINDOWS_LIST="$META/windows_dirs.txt"
awk -v POS="$POSITIONS" '
  BEGIN {
    while ((getline line < POS) > 0) {
      n = split(line, f, "\t")
      if (n < 2) continue
      c = f[1]; sub(/^chr/, "", c)
      if (c !~ /^[0-9]+$/) continue          # skips the header row
      np[c]++; p[c, np[c]] = f[2] + 0
    }
  }
  !/^#/ && NF >= 4 {
    c = $1; sub(/^chr/, "", c)
    for (i = 1; i <= np[c]; i++)
      if (p[c, i] >= $2 && p[c, i] < $3) { print $4; next }
  }
' "$WINDOWS_BED" | sort -u >"$WINDOWS_LIST"

N="$(wc -l <"$WINDOWS_LIST" | tr -d ' ')"
NPOS="$(($(wc -l <"$POSITIONS" | tr -d ' ') - 1))"
if [[ "$N" -lt 1 ]]; then
  echo "ERROR: no window in $WINDOWS_BED contains any position in $POSITIONS"
  exit 1
fi

echo "Positions requested: $NPOS"
echo "Windows containing at least one: $N (listed in $WINDOWS_LIST)"
echo "Submitting Slurm array 1-${N}%${MAX_PARALLEL} on partition ${PARTITION}"

SBATCH_OPTS=( --array="1-${N}%${MAX_PARALLEL}" --export=ALL --partition="$PARTITION" )
if [[ "$PARTITION" == "medium" || "$PARTITION" == "long" ]]; then
  SBATCH_OPTS+=( --time=12:00:01 )
fi
if [[ -n "${SLURM_ACCOUNT:-}" ]]; then
  SBATCH_OPTS+=( -A "$SLURM_ACCOUNT" )
fi

export WINDOWS_LIST HUBISZ REPO OUTDIR POSITIONS WINDOWS_BED
export QUERIES="${QUERIES:-$REPO/scripts/tmrca_queries.hubisz_8afr.json}"
echo "QUERIES=$QUERIES"

sbatch "${SBATCH_OPTS[@]}" "$SLURM_JOB_SCRIPT"

echo "Submitted. Logs: $OUTDIR/logs/site_JOBID_TASK.out"
echo "When complete:"
echo "  bash $SCRIPT_DIR/hubisz_tmrca_collate_parts.sh \"$OUTDIR/parts\" \"$OUTDIR/hubisz_tmrca_sites_8afr.tsv\""
```
