## Supplementary Information section 4 code base for "Hypothesis: A modern human range expansion ∼300,000 years ago had a critical role in Neandertal origins": submit_hubisz_tmrca_slurm_8modern.sh

```
#!/usr/bin/env bash
# Submit Slurm array for Hubisz TMRCA scan using tmrca_queries.hubisz_8modern.json
# (8 modern haplotypes: Mandenka, Khomani San, Papuan, French Basque / Basque_2F).
#
# Use a fresh OUTDIR so you do not overwrite the prior 4-haplotype run, e.g.:
#   export OUTDIR=/n/groups/reich/DAVID/CURSOR_ARGWEAVER/tmrca_scan_out_8modern
#
# Usage:
#   export HUBISZ=...
#   export REPO=...
#   export OUTDIR=...   # e.g. .../tmrca_scan_out_8modern
#   export PARTITION=short
#   bash scripts/submit_hubisz_tmrca_slurm_8modern.sh
#
# After completion:
#   bash scripts/hubisz_tmrca_collate_parts.sh "$OUTDIR/parts" "$OUTDIR/hubisz_tmrca_5kb_8modern.tsv"

set -euo pipefail

: "${HUBISZ:?export HUBISZ (dataset root)}"
: "${REPO:?export REPO (path with scripts/hubisz_out_bed_tmrca_scan.py)}"
: "${OUTDIR:?export OUTDIR (writable; use a new directory)}"
: "${PARTITION:?export PARTITION (e.g. short, medium, priority)}"

WINDOWS_BED="${WINDOWS_BED:-$HUBISZ/filtered_windows.bed}"
MAX_PARALLEL="${MAX_PARALLEL:-150}"

SCRIPT_DIR="$(cd "$(dirname "${BASH_SOURCE[0]}")" && pwd)"
SLURM_JOB_SCRIPT="${SLURM_JOB_SCRIPT:-$SCRIPT_DIR/slurm_hubisz_tmrca_array_task_8modern.sh}"

if [[ ! -f "$WINDOWS_BED" ]]; then
  echo "ERROR: WINDOWS_BED not found: $WINDOWS_BED"
  exit 1
fi
if [[ ! -f "$SLURM_JOB_SCRIPT" ]]; then
  echo "ERROR: Slurm task script not found: $SLURM_JOB_SCRIPT"
  exit 1
fi

META="$OUTDIR/metadata"
mkdir -p "$META" "$OUTDIR/parts" "$OUTDIR/logs"

WINDOWS_LIST="$META/windows_dirs.txt"
awk '!/^#/ && NF >= 4 && !seen[$4]++ { print $4 }' "$WINDOWS_BED" >"$WINDOWS_LIST"

N="$(wc -l <"$WINDOWS_LIST" | tr -d ' ')"
if [[ "$N" -lt 1 ]]; then
  echo "ERROR: no rows in $WINDOWS_LIST"
  exit 1
fi

echo "Using QUERIES=$REPO/scripts/tmrca_queries.hubisz_8modern.json (override with export QUERIES=... before sbatch if needed)"
echo "Windows list: $WINDOWS_LIST ($N unique dirnames)"
echo "Submitting Slurm array 1-${N}%${MAX_PARALLEL} on partition ${PARTITION}"

SBATCH_OPTS=( --array="1-${N}%${MAX_PARALLEL}" --export=ALL --partition="$PARTITION" )
if [[ "$PARTITION" == "medium" || "$PARTITION" == "long" ]]; then
  SBATCH_OPTS+=( --time=12:00:01 )
fi
if [[ -n "${SLURM_ACCOUNT:-}" ]]; then
  SBATCH_OPTS+=( -A "$SLURM_ACCOUNT" )
fi

export WINDOWS_LIST HUBISZ REPO OUTDIR
export QUERIES="${QUERIES:-$REPO/scripts/tmrca_queries.hubisz_8modern.json}"

sbatch "${SBATCH_OPTS[@]}" "$SLURM_JOB_SCRIPT"

echo "Submitted. Logs: $OUTDIR/logs/chunk_JOBID_TASK.out"
echo "When complete, collate with:"
echo "  bash $SCRIPT_DIR/hubisz_tmrca_collate_parts.sh \"$OUTDIR/parts\" \"$OUTDIR/hubisz_tmrca_5kb_8modern.tsv\""
```
