## Supplementary Information section 4 code base for "Hypothesis: A modern human range expansion ∼300,000 years ago had a critical role in Neandertal origins": threshold_landscape.py

```
#!/usr/bin/env python3
"""
Explore the (x, y, stringency) threshold landscape for the sweep catalog and
score each combination by the Neandertal-to-Denisovan count ratio.

Case  : we are confident that TMRCA(moderns + Neandertals) <= x
        AND confident that TMRCA(moderns + Denisova)      >  y
Control: the same statement with Neandertal and Denisova swapped.

"Confident" has three available levels per side, because the scan stored the
posterior mean and the 2.5 / 97.5 percentiles over MCMC replicates:

  level   "<= x" uses      "> y" uses     interpretation
  strict  p975 <= x        p025 >  y      whole 95% interval on the right side
  mean    mean <= x        mean >  y      point estimate only (current catalog)
  loose   p025 <= x        p975 >  y      interval merely touches the region

Under the standard population tree the two archaics are sister lineages, so
moderns are equidistant from both and the case/control counts should be equal.
The ratio is therefore an enrichment statistic; the binomial p-value tests
case vs control against 50/50.

Usage:
  python3 scripts/threshold_landscape.py --scan hubisz_tmrca_5kb_8afr.tsv \
      -o catalog_annotation/threshold_landscape.tsv
"""

from __future__ import annotations

import argparse
import csv
import math
from collections import defaultdict

LEVELS = ("strict", "mean", "loose")
IN_KEY = {"strict": "p975", "mean": "mean", "loose": "p025"}
OUT_KEY = {"strict": "p025", "mean": "mean", "loose": "p975"}


def binom_two_sided(k: int, n: int, p: float = 0.5) -> float:
    """Two-sided binomial p-value against p=0.5.

    Exact (log-space) summation for n <= 2000, normal approximation above,
    where the exact tail is numerically irrelevant anyway.
    """
    if n == 0:
        return 1.0
    if n > 2000:
        z = abs(k - n * p) / math.sqrt(n * p * (1 - p))
        return math.erfc(z / math.sqrt(2))

    def log_pmf(i):
        return (math.lgamma(n + 1) - math.lgamma(i + 1) - math.lgamma(n - i + 1)
                + i * math.log(p) + (n - i) * math.log(1 - p))

    obs = log_pmf(k)
    tot = 0.0
    for i in range(n + 1):
        lp = log_pmf(i)
        if lp <= obs + 1e-12:
            tot += math.exp(lp)
    return min(1.0, tot)


def load(scan_path):
    seen = set()
    recs = []
    with open(scan_path, encoding="utf-8") as f:
        for r in csv.DictReader(f, delimiter="\t"):
            key = (r["chrom"], r["pos"])
            if key in seen:
                continue
            seen.add(key)
            d = {}
            ok = True
            for tag, pre in (("nea", "02_modern_plus_both_Nea"),
                             ("den", "03_modern_plus_Den")):
                for suf in ("mean", "p025", "p975"):
                    try:
                        d[f"{tag}_{suf}"] = float(r[f"{pre}_{suf}_kya"])
                    except (KeyError, ValueError):
                        ok = False
            if not ok:
                continue
            c = r["chrom"].replace("chr", "")
            d["chrom"] = "X" if c == "23" else c
            d["pos"] = int(float(r["pos"]))
            recs.append(d)
    return recs


def n_clusters(hits, gap=5000):
    by = defaultdict(list)
    for c, p in hits:
        by[c].append(p)
    tot = 0
    for ps in by.values():
        ps.sort()
        tot += 1
        for i in range(1, len(ps)):
            if ps[i] - ps[i - 1] > gap:
                tot += 1
    return tot


def count(recs, level, x, y, flip=False):
    ik, ok = IN_KEY[level], OUT_KEY[level]
    a, b = ("den", "nea") if flip else ("nea", "den")
    return [(d["chrom"], d["pos"]) for d in recs
            if d[f"{a}_{ik}"] <= x and d[f"{b}_{ok}"] > y]


def main() -> int:
    ap = argparse.ArgumentParser()
    ap.add_argument("--scan", default="hubisz_tmrca_5kb_8afr.tsv")
    ap.add_argument("--min-case", type=int, default=25,
                    help="minimum case sites for a combination to be 'usable'")
    ap.add_argument("-o", "--out", default="catalog_annotation/threshold_landscape.tsv")
    args = ap.parse_args()

    recs = load(args.scan)
    print(f"usable grid sites: {len(recs):,}\n")

    xs = [450, 500, 550, 600, 650, 694, 700, 750, 800]
    ys = [550, 600, 650, 694, 700, 750, 800, 900, 1000]

    rows = []
    for level in LEVELS:
        for x in xs:
            for y in ys:
                case = count(recs, level, x, y, flip=False)
                ctrl = count(recs, level, x, y, flip=True)
                nc, nk = len(case), len(ctrl)
                rows.append({
                    "level": level, "x_ka": x, "y_ka": y,
                    "case_sites": nc, "control_sites": nk,
                    "case_clusters": n_clusters(case),
                    "control_clusters": n_clusters(ctrl),
                    "case_X": sum(1 for c, _ in case if c == "X"),
                    "ratio": round(nc / nk, 3) if nk else None,
                    "binom_p": round(binom_two_sided(nc, nc + nk), 6) if nc + nk else None,
                })

    import os
    os.makedirs(os.path.dirname(args.out) or ".", exist_ok=True)
    with open(args.out, "w", encoding="utf-8", newline="") as f:
        w = csv.DictWriter(f, fieldnames=list(rows[0].keys()), delimiter="\t")
        w.writeheader()
        w.writerows(rows)

    for level in LEVELS:
        print(f"===== level = {level}  ('<=x' via {IN_KEY[level]},  '>y' via {OUT_KEY[level]}) =====")
        print("ratio (case/control); '.' = no control sites; blank = no case sites")
        print("      y> " + "".join(f"{y:>8}" for y in ys))
        for x in xs:
            cells = []
            for y in ys:
                r = next(q for q in rows if q["level"] == level and q["x_ka"] == x and q["y_ka"] == y)
                if r["case_sites"] == 0:
                    cells.append(f"{'-':>8}")
                elif r["ratio"] is None:
                    cells.append(f"{'inf':>8}")
                else:
                    cells.append(f"{r['ratio']:>8.2f}")
            print(f"x<={x:<5}" + "".join(cells))
        print("      case sites")
        print("      y> " + "".join(f"{y:>8}" for y in ys))
        for x in xs:
            cells = []
            for y in ys:
                r = next(q for q in rows if q["level"] == level and q["x_ka"] == x and q["y_ka"] == y)
                cells.append(f"{r['case_sites']:>8}")
            print(f"x<={x:<5}" + "".join(cells))
        print()

    usable = [r for r in rows if r["case_sites"] >= args.min_case and r["control_sites"] > 0]
    usable.sort(key=lambda r: (-r["ratio"], -r["case_sites"]))
    print(f"--- best combinations with >= {args.min_case} case sites, ranked by ratio ---")
    print(f"{'level':<8}{'x':>6}{'y':>6}{'case':>7}{'ctrl':>6}{'ratio':>8}{'clust':>7}{'X':>5}{'binom p':>12}")
    for r in usable[:18]:
        print(f"{r['level']:<8}{r['x_ka']:>6}{r['y_ka']:>6}{r['case_sites']:>7}"
              f"{r['control_sites']:>6}{r['ratio']:>8.2f}{r['case_clusters']:>7}"
              f"{r['case_X']:>5}{r['binom_p']:>12.2e}")
    print(f"\nwrote {args.out}")
    return 0


if __name__ == "__main__":
    raise SystemExit(main())
```
